## Supplementary Figures and files for "Hominoid-specific transposable elements rewired the expression of neural crest migration genes"

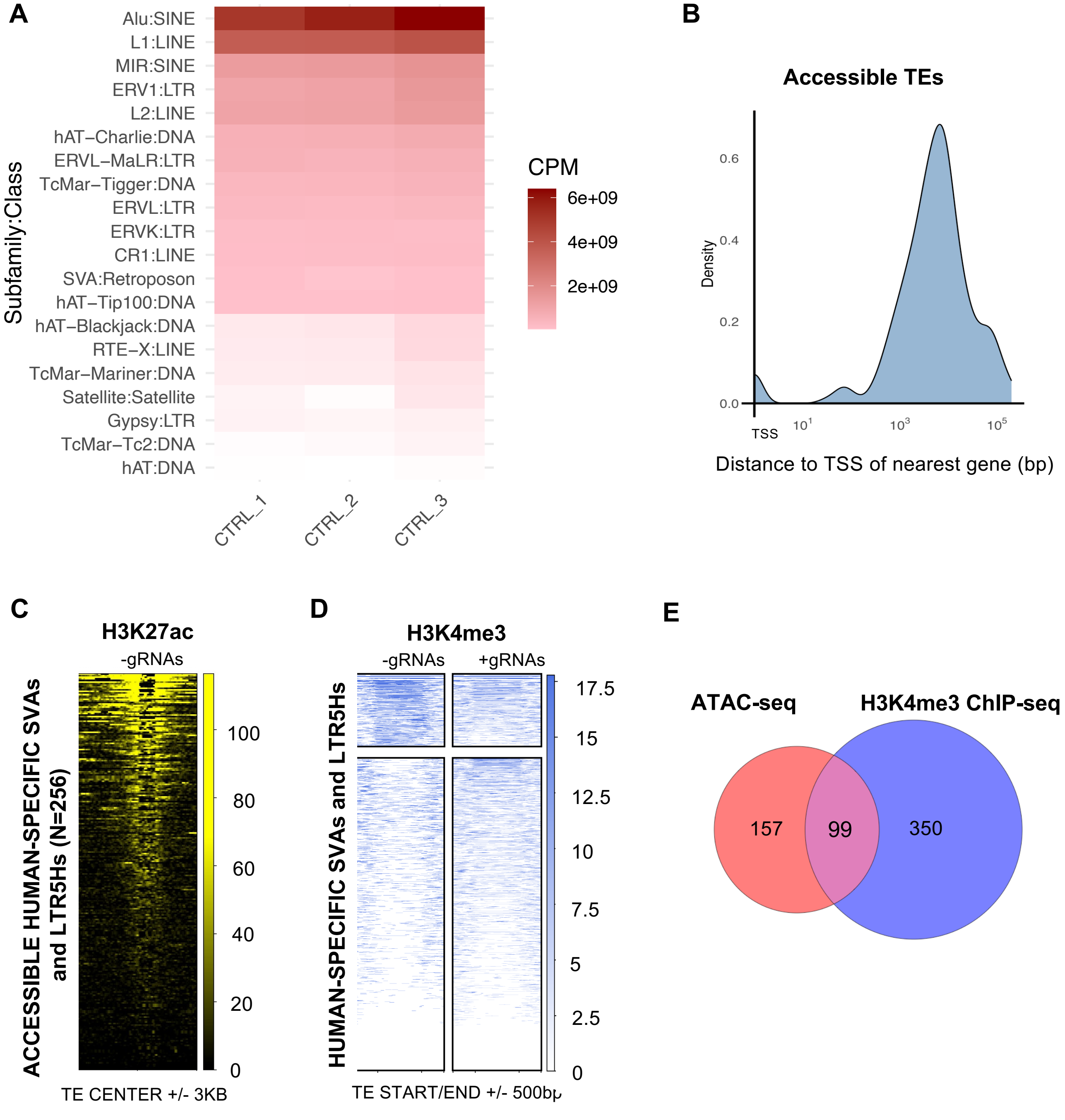

**Appendix Figure S1 – (a)** Top 20 transposable element (TE) subfamilies ranked by counts per million (CPM) expression across three CNCC replicates. **(b)** Density plot of accessible human-specific SVAs and LTR5Hs in relation to the transcription start site of their nearest genes. **(c)** ChIP-seq heatmap displaying H3K27ac signal at SVAs and LTR5Hs accessible in CNCCs. Bigwig used for heatmap was normalized by sequencing depth. **(d)** ChIP-seq heatmap displaying H3K4me3 signal at human-specific SVAs and LTR5Hs. Bigwig used for heatmap was normalized by sequencing depth. **(e)** Venn diagram illustrating the overlap between human-specific SVAs and LTR5Hs that are both accessible and marked by H3K4me3.  Bigwigs used for heatmap are normalised by sequencing depth.

**A**

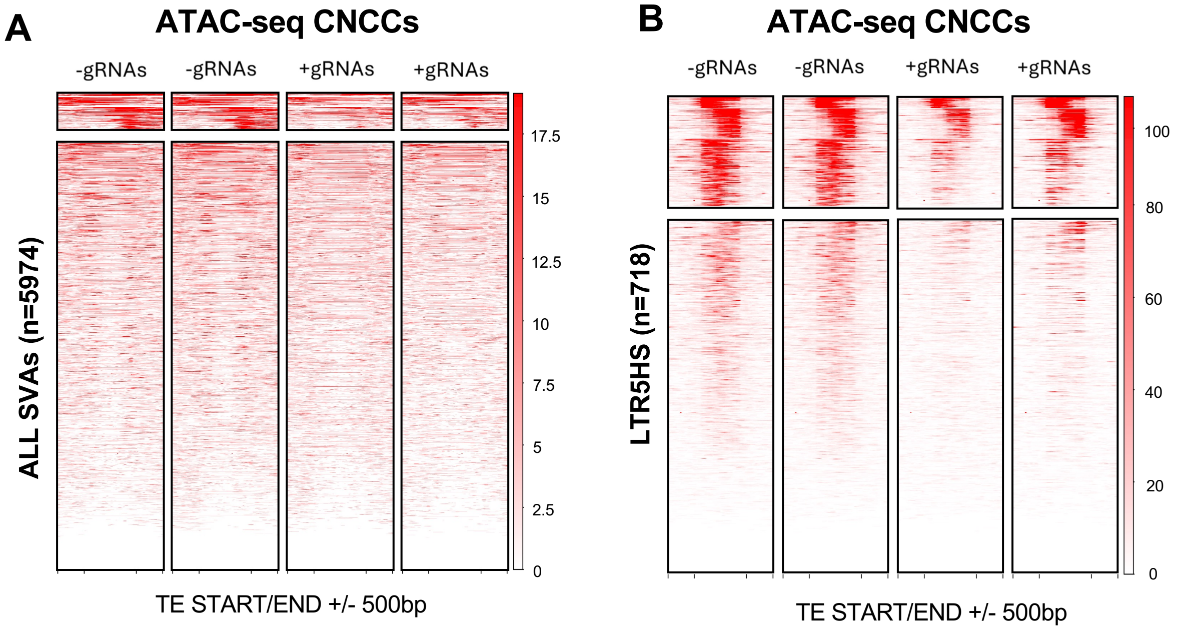

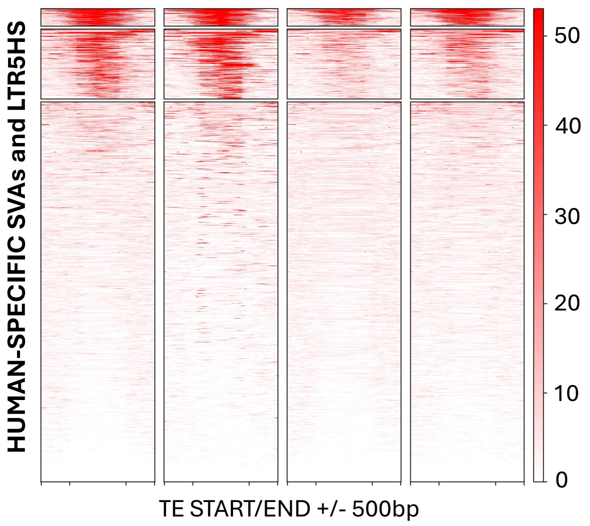

**D**

**Motifs enriched in accessible SVAs**

| | | |
|---|---|---|
| TWIST1 | 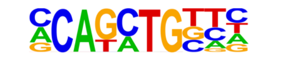 | 46% |
| HOMEOBOX | 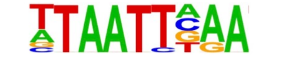 | 36% |
| SOX9 | 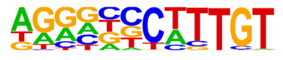 | 43% |
| SOX10 | 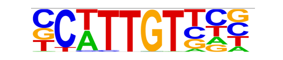 | 50% |

**E**

**Motifs enriched in accessible LTR5HS**

| | | |
|---|---|---|
| TWIST1 | 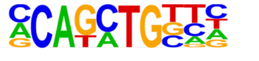 | 89% |
| HOMEOBOX | 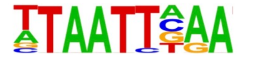 | 66% |
| TFAP2A | 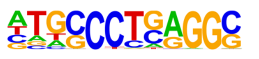 | 75% |
| SLUG | 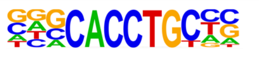 | 67% |
| SNAIL1 | 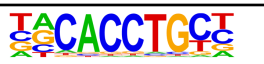 | 76% |
| ZEB1 | 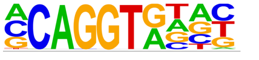 | 93% |
| ZEB2 | 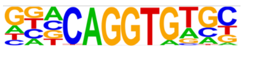 | 75% |
| SOX9 | 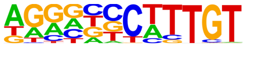 | 66% |
| SOX10 | 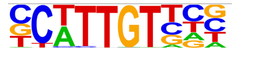 | 95% |
| FOXD3 | 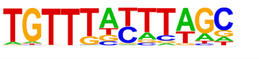 | 37% |

**Appendix Figure S2 – (a,b)** ATAC-seq heatmap showing chromatin accessibility at all SVAs **(a)** and LTR5Hs **(b)** in CNCCs. **(c)** ATAC-seq heatmap showing chromatin accessibility at human-specific SVAs and LTR5Hs in CNCCs generated with alternative protocol (Bajpai et al. 2010). Bigwigs used for heatmaps were normalized by sequencing depth. **(d,e)** Motif analysis for all accessilbe SVAs **(d)** and LTR5Hs **(e)**.

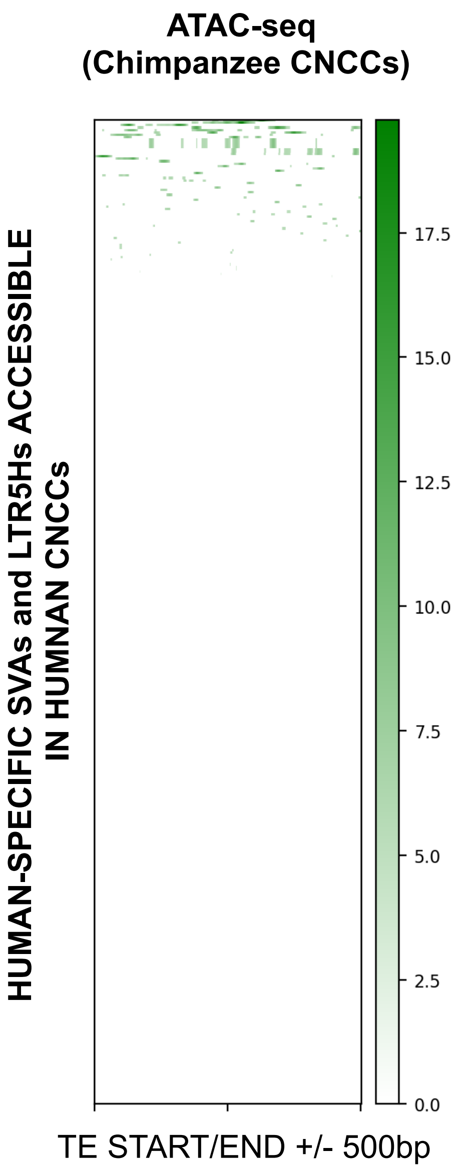

**Appendix Figure S3.** Chimpanzee CNCC ATAC-seq heatmap centered on regions up and downstream (+/- 250 bp) to the 256 accessible LTR5Hs and human-specific SVAs. The bigwig used for the heatmap is normalized by sequencing depth. The chimpanzee ATAC-seq data were publicly available (Prescott et al. 2015).

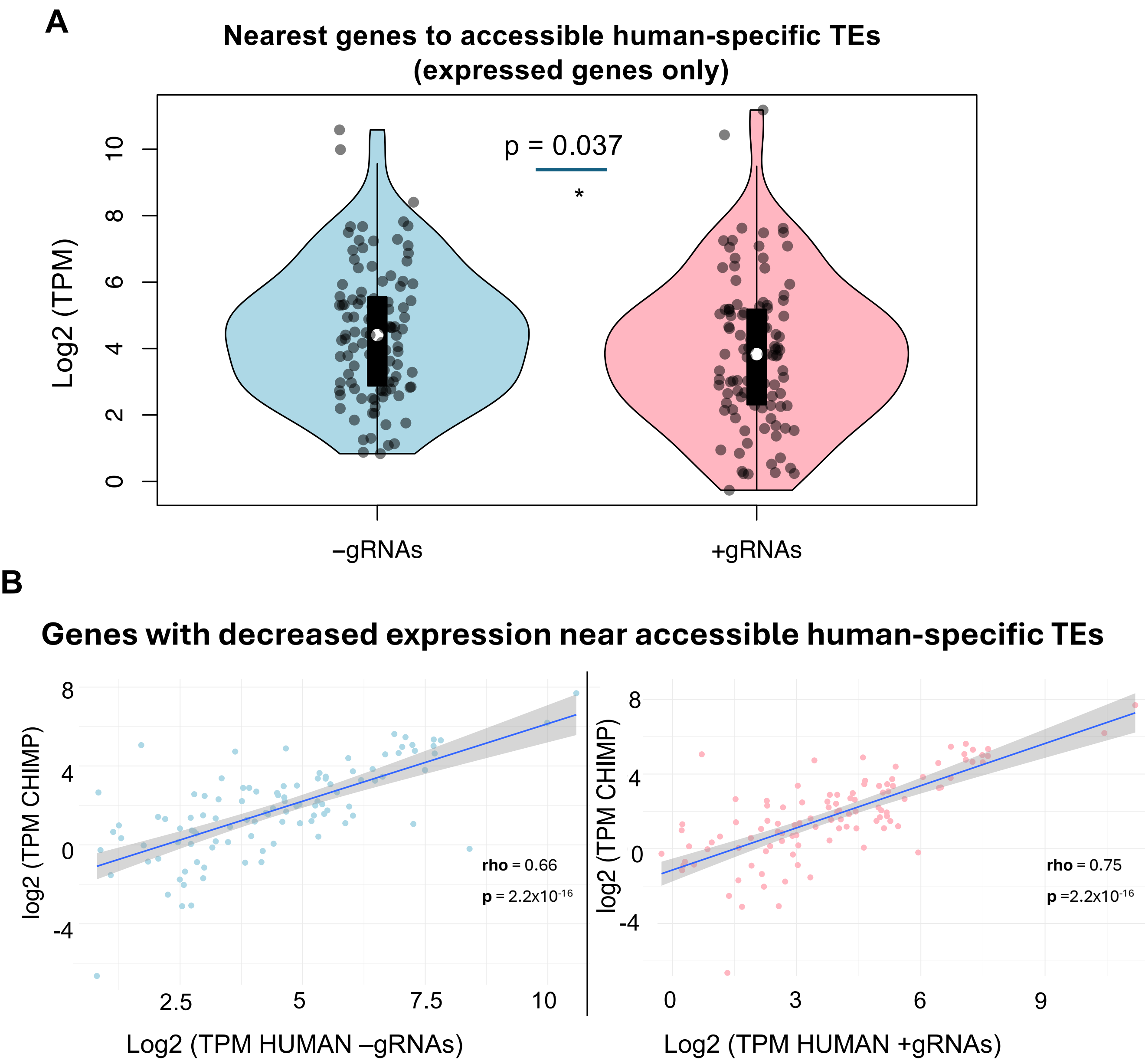

**Appendix Figure S4 – (a)** Violin plots showing the expression of the genes representing the nearest gene to an accessible human-specific LTR5Hs or SVA (expressed genes only, N = 107; p-value calculated with Wilcoxon's Rank Sum test). **(b)** Correlation plots showing correlation of expression for the same set of genes between human (+ and – gRNAs) and chimpanzee CNCCs.

Appendix File 1 - CPMs of TEs in CNCCs

| | CTRL_1 | CTRL_2 | CTRL_3 |
|---|---|---|---|
| (CATTC)n:Satellite:Satellite | 10964431384.5884 | 833333333.333333 | 1.25e+09 |
| (GAATG)n:Satellite:Satellite | 9867988246.12957 | 1111111111.11111 | 0 |
| 6kbHsap:acro:Satellite | 0 | 0 | 0 |
| 7SK:RNA:RNA | 15350203938.4237 | 9444444444.44445 | 9.375e+09 |
| ACRO1:acro:Satellite | 2192886276.91768 | 277777777.777778 | 4.375e+09 |
| ALR/Alpha:centr:Satellite | 149116266830.402 | 16666666666.6667 | 1.75e+10 |
| Alu:Alu:SINE | 254374808122.451 | 131666666666.667 | 2.21875e+11 |
| AluJb:Alu:SINE | 18341300820139.5 | 10653888888888.9 | 1.8189375e+13 |
| AluJo:Alu:SINE | 11159598263234.1 | 6328888888888.88 | 1.084875e+13 |
| AluJr4:Alu:SINE | 2507565457655.36 | 1500555555555.55 | 2.505e+12 |
| AluJr:Alu:SINE | 11797728169817.1 | 6803888888888.88 | 1.158e+13 |
| AluSc5:Alu:SINE | 867286522520.944 | 485277777777.778 | 8.5625e+11 |
| AluSc8:Alu:SINE | 3131441603438.45 | 1703055555555.55 | 2.851875e+12 |
| AluSc:Alu:SINE | 4282706898820.23 | 2426388888888.88 | 3.998125e+12 |
| AluSg4:Alu:SINE | 938555326520.768 | 592777777777.778 | 9.18125e+11 |
| AluSg7:Alu:SINE | 1263102495504.58 | 789722222222.222 | 1.2425e+12 |
| AluSg:Alu:SINE | 6034823034077.45 | 3.6e+12 | 5.854375e+12 |
| AluSp:Alu:SINE | 8838428139116.72 | 4846388888888.88 | 8.28875e+12 |
| AluSq10:Alu:SINE | 217095741414.85 | 118888888888.889 | 1.8625e+11 |
| AluSq2:Alu:SINE | 9592781018376.39 | 5448333333333.33 | 9.2625e+12 |
| AluSq4:Alu:SINE | 285075215999.298 | 174444444444.445 | 3.1625e+11 |
| AluSq:Alu:SINE | 2465900618393.93 | 1396944444444.45 | 2.32375e+12 |
| AluSx1:Alu:SINE | 17343537564141.9 | 9783333333333.33 | 1.6393125e+13 |
| AluSx3:Alu:SINE | 4650015350203.94 | 2673611111111.12 | 4.503125e+12 |
| AluSx4:Alu:SINE | 1517477303627.03 | 889166666666.667 | 1.505625e+12 |
| AluSx:Alu:SINE | 16764615587035.7 | 9589444444444.45 | 1.5944375e+13 |
| AluSz6:Alu:SINE | 6247533002938.48 | 3773055555555.55 | 6.468125e+12 |
| AluSz:Alu:SINE | 15437919389500.5 | 8926388888888.88 | 1.480875e+13 |
| AluY:Alu:SINE | 14226349721503.4 | 8072222222222.22 | 1.2328125e+13 |
| AluYa5:Alu:SINE | 267532125783.957 | 148888888888.889 | 1.99375e+11 |
| AluYa8:Alu:SINE | 20832419630.718 | 1.5e+10 | 2.125e+10 |
| AluYb10:Alu:SINE | 0 | 0 | 0 |
| AluYb11:Alu:SINE | 0 | 0 | 0 |
| AluYb8:Alu:SINE | 233542388491.733 | 114722222222.222 | 1.525e+11 |
| AluYb8a1:Alu:SINE | 0 | 0 | 0 |
| AluYb9:Alu:SINE | 29603964738.3887 | 11388888888.8889 | 1.3125e+10 |
| AluYc3:Alu:SINE | 133766062891.978 | 70277777777.7778 | 1.0125e+11 |
| AluYc:Alu:SINE | 414455506337.44 | 214444444444.445 | 3.6375e+11 |
| AluYd8:Alu:SINE | 5482215692.2942 | 6111111111.11112 | 1.1875e+10 |
| AluYe5:Alu:SINE | 138151835445.814 | 85833333333.3333 | 1.25e+11 |
| AluYe6:Alu:SINE | 32893294153.7652 | 25833333333.3333 | 1.5625e+10 |
| AluYf1:Alu:SINE | 201745537476.427 | 148611111111.111 | 2.41875e+11 |
| AluYg6:Alu:SINE | 53725713784.4831 | 25833333333.3333 | 4.375e+10 |
| AluYh3:Alu:SINE | 281785886583.922 | 158333333333.333 | 2.2625e+11 |
| AluYh3a3:Alu:SINE | 49339941230.6477 | 26388888888.8888 | 3.9375e+10 |
| AluYh7:Alu:SINE | 3289329415.37652 | 4722222222.22222 | 3.75e+09 |
| AluYh9:Alu:SINE | 4385772553.83536 | 1666666666.66667 | 4.375e+09 |
| AluYi6:Alu:SINE | 57015043199.8596 | 31666666666.6667 | 8.4375e+10 |
| AluYi6_4d:Alu:SINE | 17543090215.3414 | 13055555555.5556 | 3.0625e+10 |
| AluYj4:Alu:SINE | 360729792552.957 | 194166666666.667 | 3.59375e+11 |
| AluYk11:Alu:SINE | 71268803999.8245 | 33611111111.1112 | 7.0625e+10 |
| AluYk12:Alu:SINE | 21928862769.1768 | 8611111111.11112 | 2.0625e+10 |
| AluYk2:Alu:SINE | 73461690276.7424 | 48888888888.8888 | 6.5e+10 |
| AluYk3:Alu:SINE | 143634051138.108 | 80833333333.3333 | 1.6125e+11 |
| AluYk4:Alu:SINE | 142537607999.649 | 75277777777.7778 | 1.08125e+11 |
| AluYm1:Alu:SINE | 719266698829 | 400277777777.778 | 6.6625e+11 |
| AmnHarb1:PIF-Harbinger:DNA | 0 | 0 | 0 |
| AmnL2-1:L2:LINE | 0 | 0 | 0 |
| AmnSINE1:5S-Deu-L2:SINE | 114030086399.719 | 60833333333.3333 | 1.21875e+11 |
| Arthur1:hAT-Tip100:DNA | 91004780492.0838 | 45833333333.3333 | 1.05e+11 |
| Arthur1A:hAT-Tip100:DNA | 58111486338.3185 | 28333333333.3333 | 7.25e+10 |

| | | | |
|---|---|---|---|
| Arthur1B:hAT-Tip100:DNA | 95390553045.9192 | 39722222222.2222 | 7.9375e+10 |
| Arthur1C:hAT-Tip100:DNA | 37279066707.6007 | 18888888888.8888 | 3.8125e+10 |
| Arthur2:hAT-Tip100:DNA | 0 | 0 | 0 |
| BLACKJACK:hAT-Blackjack:DNA | 169948686461.12 | 85833333333.3333 | 1.725e+11 |
| BSR/Beta:Satellite:Satellite | 75654576553.6599 | 21666666666.6667 | 4.0625e+10 |
| CER:Satellite:Satellite | 4385772553.83536 | 1111111111.11111 | 1.25e+09 |
| CR1-11_Crp:CR1:LINE | 1096443138.45884 | 555555555.555555 | 0 |
| CR1-12_AMi:CR1:LINE | 21928862769.1768 | 1388888888.88889 | 7.5e+09 |
| CR1-13_AMi:CR1:LINE | 1096443138.45884 | 833333333.333333 | 1.875e+09 |
| CR1-16_AMi:CR1:LINE | 8771545107.67073 | 5e+09 | 8.125e+09 |
| CR1-1_Amn:CR1:LINE | 0 | 0 | 0 |
| CR1-3_Croc:CR1:LINE | 46050611815.2712 | 24166666666.6667 | 3.75e+10 |
| CR1-8_Crp:CR1:LINE | 0 | 1111111111.11111 | 1.25e+09 |
| CR1-L3A_Croc:CR1:LINE | 5482215692.2942 | 7222222222.22222 | 6.25e+09 |
| CR1-L3B_Croc:CR1:LINE | 1096443138.45884 | 0 | 0 |
| CR1_Mam:CR1:LINE | 139248278584.273 | 82222222222.2222 | 1.4375e+11 |
| Chap1_Mam:hAT-Charlie:DNA | 54822156922.9421 | 39166666666.6667 | 5.8125e+10 |
| Chap1a_Mam:hAT-Charlie:DNA | 0 | 0 | 0 |
| Charlie10:hAT-Charlie:DNA | 401298188675.934 | 162222222222.222 | 3.36875e+11 |
| Charlie10a:hAT-Charlie:DNA | 21928862769.1768 | 14166666666.6667 | 1.9375e+10 |
| Charlie10b:hAT-Charlie:DNA | 26314635323.0122 | 12222222222.2222 | 2e+10 |
| Charlie11:hAT-Charlie:DNA | 8771545107.67073 | 2.5e+09 | 3.125e+09 |
| Charlie12:hAT-Charlie:DNA | 35086180430.6828 | 14444444444.4444 | 2e+10 |
| Charlie13a:hAT-Charlie:DNA | 32893294153.7652 | 15277777777.7778 | 2.0625e+10 |
| Charlie13b:hAT-Charlie:DNA | 104162098153.59 | 4e+10 | 6.3125e+10 |
| Charlie14a:hAT-Charlie:DNA | 21928862769.1768 | 3888888888.88888 | 1.625e+10 |
| Charlie15a:hAT-Charlie:DNA | 243410376737.862 | 154166666666.667 | 2.25625e+11 |
| Charlie15b:hAT-Charlie:DNA | 145826937415.026 | 7.25e+10 | 1.325e+11 |
| Charlie16:hAT-Charlie:DNA | 0 | 0 | 0 |
| Charlie16a:hAT-Charlie:DNA | 123898074645.849 | 75555555555.5555 | 1.35625e+11 |
| Charlie17:hAT-Charlie:DNA | 118415858953.555 | 52222222222.2222 | 1.175e+11 |
| Charlie17a:hAT-Charlie:DNA | 61400815753.695 | 33888888888.8888 | 7.5625e+10 |
| Charlie17b:hAT-Charlie:DNA | 10964431384.5884 | 5833333333.33333 | 1.4375e+10 |
| Charlie18a:hAT-Charlie:DNA | 249989035568.616 | 141111111111.111 | 2.10625e+11 |
| Charlie19a:hAT-Charlie:DNA | 100872768738.213 | 63055555555.5555 | 8.5625e+10 |
| Charlie1:hAT-Charlie:DNA | 201745537476.427 | 104444444444.445 | 1.85e+11 |
| Charlie1a:hAT-Charlie:DNA | 577825533967.808 | 398055555555.555 | 7.15e+11 |
| Charlie1b:hAT-Charlie:DNA | 143634051138.108 | 83888888888.8888 | 1.35e+11 |
| Charlie20a:hAT-Charlie:DNA | 40568396122.9771 | 25555555555.5555 | 4.9375e+10 |
| Charlie21a:hAT-Charlie:DNA | 61400815753.695 | 3.75e+10 | 5.8125e+10 |
| Charlie22a:hAT-Charlie:DNA | 88811894215.1659 | 53333333333.3333 | 9.9375e+10 |
| Charlie23a:hAT-Charlie:DNA | 61400815753.695 | 34166666666.6667 | 7.625e+10 |
| Charlie24:hAT-Charlie:DNA | 80040349107.4954 | 38333333333.3333 | 8.375e+10 |
| Charlie25:hAT-Charlie:DNA | 40568396122.9771 | 22222222222.2222 | 3.9375e+10 |
| Charlie26a:hAT-Charlie:DNA | 13157317661.5061 | 1e+10 | 1.8125e+10 |
| Charlie29a:hAT-Charlie:DNA | 55918600061.4007 | 51111111111.1112 | 9.8125e+10 |
| Charlie29b:hAT-Charlie:DNA | 0 | 0 | 0 |
| Charlie2a:hAT-Charlie:DNA | 298232533660.804 | 118888888888.889 | 2.7625e+11 |
| Charlie2b:hAT-Charlie:DNA | 225867286522.521 | 128333333333.333 | 3.50625e+11 |
| Charlie30a:hAT-Charlie:DNA | 14253760799.9649 | 1.25e+10 | 2e+10 |
| Charlie30b:hAT-Charlie:DNA | 0 | 0 | 0 |
| Charlie31a:hAT-Charlie:DNA | 0 | 0 | 0 |
| Charlie32a:hAT-Charlie:DNA | 0 | 0 | 0 |
| Charlie3:hAT-Charlie:DNA | 44954168676.8126 | 26944444444.4445 | 4.3125e+10 |
| Charlie4:hAT-Charlie:DNA | 10964431384.5884 | 4166666666.66667 | 9.375e+09 |
| Charlie4a:hAT-Charlie:DNA | 286171659137.757 | 142222222222.222 | 2.51875e+11 |
| Charlie4z:hAT-Charlie:DNA | 509846059383.361 | 273611111111.112 | 5.5875e+11 |
| Charlie5:hAT-Charlie:DNA | 126090960922.767 | 65555555555.5555 | 1.16875e+11 |
| Charlie6:hAT-Charlie:DNA | 1096443138.45884 | 555555555.555555 | 1.25e+09 |
| Charlie7:hAT-Charlie:DNA | 461602561291.172 | 176944444444.445 | 4.5625e+11 |
| Charlie7a:hAT-Charlie:DNA | 55918600061.4007 | 5.25e+10 | 7.5625e+10 |
| Charlie8:hAT-Charlie:DNA | 275207227753.169 | 138611111111.111 | 2.58125e+11 |
| Charlie9:hAT-Charlie:DNA | 62497258892.154 | 51111111111.1112 | 7.75e+10 |
| Cheshire:hAT-Charlie:DNA | 89908337353.6248 | 64444444444.4445 | 1.4375e+11 |

| | | | |
|---|---|---|---|
| Chompy-6_Croc:PIF-Harbinger:DNA | 0 | 0 | 0 |
| Chompy-7_Croc:PIF-Harbinger:DNA | 0 | 0 | 0 |
| D20S16:Satellite:Satellite | 9867988246.12957 | 6666666666.66667 | 1.5625e+10 |
| DNA1_Mam:TcMar-Tc1:DNA | 0 | 0 | 0 |
| DNA1_Mam:TcMar:DNA | 7675101969.21189 | 6666666666.66667 | 1.25e+10 |
| ERV24B_Prim-int:ERV1:LTR | 36182623569.1417 | 19444444444.4445 | 4.125e+10 |
| ERV24_Prim-int:ERV1:LTR | 38375509846.0593 | 15833333333.3333 | 2.625e+10 |
| ERV3-16A3_I-int:ERVL:LTR | 278496557168.545 | 112777777777.778 | 2.275e+11 |
| ERV3-16A3_I:ERVL:LTR | 0 | 0 | 0 |
| ERV3-16A3_LTR:ERVL:LTR | 32893294153.7652 | 18333333333.3333 | 2.875e+10 |
| ERVL-B4-int:ERVL:LTR | 362922678829.877 | 149722222222.222 | 3.39375e+11 |
| ERVL-E-int:ERVL:LTR | 1149072409104.86 | 348611111111.112 | 1.33375e+12 |
| ERVL-int:ERVL:LTR | 18639533353.8003 | 10833333333.3333 | 1.4375e+10 |
| ERVL47-int:ERVL:LTR | 3289329415.37652 | 1944444444.44445 | 1.25e+09 |
| EUTREP14:EUTREP14:Unknown | 0 | 0 | 0 |
| EUTREP15:ERV1:LTR | 0 | 0 | 0 |
| EUTREP16:LTR:LTR | 0 | 0 | 0 |
| EUTREP2:hAT:DNA | 0 | 0 | 0 |
| EUTREP2:hAT:DNA? | 0 | 0 | 0 |
| EUTREP4:PiggyBac:DNA | 0 | 0 | 0 |
| EUTREP6:EUTREP6:Unknown | 0 | 0 | 0 |
| EUTREP7:ERV1:LTR | 0 | 0 | 0 |
| EUTREP8:LTR:LTR | 0 | 0 | 0 |
| Eulor10:DNA:DNA? | 1096443138.45884 | 1666666666.66667 | 6.25e+08 |
| Eulor11:DNA:DNA | 1096443138.45884 | 277777777.777778 | 6.25e+08 |
| Eulor12:DNA:DNA? | 0 | 0 | 0 |
| Eulor1:DNA:DNA | 3289329415.37652 | 277777777.777778 | 1.25e+09 |
| Eulor2A:DNA:DNA? | 2192886276.91768 | 277777777.777778 | 5.625e+09 |
| Eulor2B:DNA:DNA? | 0 | 0 | 0 |
| Eulor2C:DNA:DNA? | 0 | 277777777.777778 | 1.25e+09 |
| Eulor3:DNA:DNA? | 0 | 0 | 0 |
| Eulor4:Eulor4:Unknown | 0 | 555555555.555555 | 0 |
| Eulor5A:Crypton-A:DNA | 0 | 0 | 0 |
| Eulor5A:DNA:DNA? | 10964431384.5884 | 8611111111.11112 | 1.375e+10 |
| Eulor5B:DNA:DNA? | 0 | 277777777.777778 | 6.25e+08 |
| Eulor6A:Crypton:DNA | 0 | 0 | 0 |
| Eulor6A:DNA:DNA? | 0 | 0 | 0 |
| Eulor6B:Crypton:DNA | 0 | 0 | 0 |
| Eulor6B:DNA:DNA? | 1096443138.45884 | 1666666666.66667 | 2.5e+09 |
| Eulor6C:DNA:DNA? | 0 | 0 | 0 |
| Eulor6D:DNA:DNA? | 1096443138.45884 | 277777777.777778 | 0 |
| Eulor6E:DNA:DNA? | 0 | 0 | 0 |
| Eulor7:DNA:DNA? | 0 | 0 | 0 |
| Eulor8:TcMar:DNA | 1096443138.45884 | 1388888888.88889 | 0 |
| Eulor9A:DNA:DNA | 4385772553.83536 | 833333333.333333 | 1.25e+09 |
| Eulor9B:DNA:DNA | 0 | 277777777.777778 | 0 |
| Eulor9C:DNA:DNA | 4385772553.83536 | 277777777.777778 | 2.5e+09 |
| EutTc1-N2:TcMar-Tc1:DNA | 0 | 0 | 0 |
| EuthAT-2:hAT-Ac:DNA | 0 | 0 | 0 |
| EuthAT-N1:hAT-Tip100:DNA | 0 | 0 | 0 |
| Eutr10:ERVL:LTR | 0 | 0 | 0 |
| Eutr11:DNA:DNA | 0 | 0 | 0 |
| Eutr15:hAT-Tip100:DNA | 0 | 0 | 0 |
| Eutr16:hAT-Tip100:DNA | 0 | 0 | 0 |
| Eutr16:hAT-Tip100:DNA? | 0 | 0 | 0 |
| Eutr18:LTR:LTR | 0 | 0 | 0 |
| Eutr18:LTR:LTR? | 0 | 0 | 0 |
| Eutr2:DNA:DNA | 0 | 0 | 0 |
| Eutr2:MULE-MuDR:DNA | 0 | 0 | 0 |
| Eutr2B:MULE-MuDR:DNA | 0 | 0 | 0 |
| Eutr5:LTR:LTR | 0 | 0 | 0 |
| Eutr5:LTR:LTR? | 0 | 0 | 0 |
| FAM:Alu:SINE | 502170957414.149 | 318333333333.333 | 5.69375e+11 |
| FLAM_A:Alu:SINE | 1451690715319.5 | 833055555555.555 | 1.545e+12 |

| | | | |
|---|---|---|---|
| FLAM_C:Alu:SINE | 2595280908732.07 | 1472222222222.22 | 2.385625e+12 |
| FRAM:Alu:SINE | 834393228367.176 | 501111111111.112 | 7.88125e+11 |
| FordPrefect:hAT-Tip100:DNA | 28507521599.9298 | 10833333333.3333 | 1.25e+10 |
| FordPrefect_a:hAT-Tip100:DNA | 24121749046.0945 | 5833333333.33333 | 2.125e+10 |
| GSAT:centr:Satellite | 1096443138.45884 | 555555555.555555 | 1.875e+09 |
| GSATII:centr:Satellite | 53725713784.4831 | 23888888888.8888 | 2.75e+10 |
| GSATX:centr:Satellite | 8771545107.67073 | 3333333333.33333 | 5e+09 |
| HAL1:L1:LINE | 2076663304241.04 | 1083888888888.89 | 1.85625e+12 |
| HAL1M8:L1:LINE | 220385070830.227 | 1.4e+11 | 2.71875e+11 |
| HAL1ME:L1:LINE | 366212008245.252 | 228888888888.888 | 3.54375e+11 |
| HAL1b:L1:LINE | 324547168983.817 | 1.675e+11 | 2.8625e+11 |
| HERV-Fc1-int:ERV1:LTR | 3289329415.37652 | 0 | 0 |
| HERV-Fc1_LTR1:ERV1:LTR | 4385772553.83536 | 277777777.777778 | 0 |
| HERV-Fc1_LTR2:ERV1:LTR | 0 | 0 | 0 |
| HERV-Fc1_LTR3:ERV1:LTR | 0 | 0 | 0 |
| HERV-Fc2-int:ERV1:LTR | 0 | 0 | 0 |
| HERV-Fc2_LTR:ERV1:LTR | 0 | 0 | 0 |
| HERV15-int:ERV1:LTR | 3289329415.37652 | 8055555555.55555 | 5.625e+09 |
| HERV16-int:ERVL:LTR | 41664839261.4358 | 19444444444.4445 | 2e+10 |
| HERV17-int:ERV1:LTR | 155694925661.155 | 74722222222.2222 | 1.35625e+11 |
| HERV1_I-int:ERV1:LTR | 59207929476.7775 | 31111111111.1112 | 2.75e+10 |
| HERV1_LTRa:ERV1:LTR | 2192886276.91768 | 555555555.555555 | 1.875e+09 |
| HERV1_LTRb:ERV1:LTR | 6578658830.75305 | 2.5e+09 | 0 |
| HERV1_LTRc:ERV1:LTR | 3289329415.37652 | 1111111111.11111 | 6.25e+08 |
| HERV1_LTRd:ERV1:LTR | 3289329415.37652 | 277777777.777778 | 1.875e+09 |
| HERV1_LTRe:ERV1:LTR | 2192886276.91768 | 2222222222.22222 | 6.25e+08 |
| HERV3-int:ERV1:LTR | 63593702030.6126 | 46666666666.6667 | 9.625e+10 |
| HERV30-int:ERV1:LTR | 12060874523.0473 | 17777777777.7778 | 5e+09 |
| HERV35I-int:ERV1:LTR | 36182623569.1417 | 11944444444.4444 | 1.8125e+10 |
| HERV4_I-int:ERV1:LTR | 36182623569.1417 | 18888888888.8888 | 4.125e+10 |
| HERV4_I:ERV1:LTR | 0 | 0 | 0 |
| HERV9-int:ERV1:LTR | 47147054953.7301 | 28333333333.3333 | 6.125e+10 |
| HERV9N-int:ERV1:LTR | 6578658830.75305 | 6111111111.11112 | 4.375e+09 |
| HERV9NC-int:ERV1:LTR | 646901451690.715 | 333888888888.888 | 5.31875e+11 |
| HERVE-int:ERV1:LTR | 331125827814.57 | 1.525e+11 | 2.58125e+11 |
| HERVE_a-int:ERV1:LTR | 282882329722.38 | 173888888888.888 | 1.525e+11 |
| HERVFH19-int:ERV1:LTR | 13157317661.5061 | 3888888888.88888 | 1.4375e+10 |
| HERVFH21-int:ERV1:LTR | 104162098153.59 | 36388888888.8888 | 6.8125e+10 |
| HERVH-int:ERV1:LTR | 14755931757379.1 | 8701111111111.12 | 2.185875e+13 |
| HERVH48-int:ERV1:LTR | 49339941230.6477 | 32777777777.7778 | 3.1875e+10 |
| HERVI-int:ERV1:LTR | 150212709968.861 | 51944444444.4445 | 1.15e+11 |
| HERVIP10B3-int:ERV1:LTR | 145826937415.026 | 105555555555.555 | 1.675e+11 |
| HERVIP10F-int:ERV1:LTR | 239024604184.027 | 88611111111.1112 | 2.95625e+11 |
| HERVIP10FH-int:ERV1:LTR | 249989035568.616 | 140833333333.333 | 2.2375e+11 |
| HERVK-int:ERVK:LTR | 1719222841103.46 | 1240555555555.55 | 1.4025e+12 |
| HERVK11-int:ERVK:LTR | 451734573045.043 | 3e+11 | 2.8625e+11 |
| HERVK11D-int:ERVK:LTR | 69075917722.907 | 11111111111.1111 | 3.375e+10 |
| HERVK13-int:ERVK:LTR | 23025305907.6356 | 10277777777.7778 | 3e+10 |
| HERVK14-int:ERVK:LTR | 23025305907.6356 | 8888888888.88888 | 1.1875e+10 |
| HERVK14C-int:ERVK:LTR | 9867988246.12957 | 5277777777.77778 | 6.25e+09 |
| HERVK22-int:ERVK:LTR | 226963729660.98 | 114722222222.222 | 2.14375e+11 |
| HERVK3-int:ERVK:LTR | 853032761720.977 | 736666666666.667 | 5.05e+11 |
| HERVK9-int:ERVK:LTR | 132669619753.52 | 93333333333.3333 | 9.75e+10 |
| HERVKC4-int:ERVK:LTR | 9867988246.12957 | 31944444444.4445 | 2.6875e+10 |
| HERVL-int:ERVL:LTR | 493399412306.477 | 254722222222.222 | 3.8125e+11 |
| HERVL18-int:ERVL:LTR | 58111486338.3185 | 24444444444.4445 | 4.0625e+10 |
| HERVL32-int:ERVL:LTR | 3289329415.37652 | 2222222222.22222 | 5e+09 |
| HERVL40-int:ERVL:LTR | 218192184553.309 | 58888888888.8888 | 1.68125e+11 |
| HERVL66-int:ERVL:LTR | 30700407876.8475 | 10277777777.7778 | 2.5e+10 |
| HERVL74-int:ERVL:LTR | 28507521599.9298 | 11388888888.8889 | 1.4375e+10 |
| HERVP71A-int:ERV1:LTR | 18639533353.8003 | 8055555555.55555 | 1.8125e+10 |
| HERVS71-int:ERV1:LTR | 255471251260.91 | 1.675e+11 | 1.9e+11 |
| HSAT4:centr:Satellite | 9867988246.12957 | 833333333.333333 | 0 |
| HSAT5:Satellite:Satellite | 0 | 0 | 6.25e+08 |

| | | | |
|---|---|---|---|
| HSAT6:Satellite:Satellite | 0 | 0 | 0 |
| HSATI:Satellite:Satellite | 0 | 833333333.333333 | 6.25e+08 |
| HSATII:Satellite:Satellite | 0 | 0 | 0 |
| HSMAR1:TcMar-Mariner:DNA | 220385070830.227 | 96388888888.8888 | 1.3625e+11 |
| HSMAR2:TcMar-Mariner:DNA | 184202447261.085 | 73055555555.5555 | 1.5625e+11 |
| HUERS-P1-int:ERV1:LTR | 100872768738.213 | 5e+10 | 8.8125e+10 |
| HUERS-P2-int:ERV1:LTR | 40568396122.9771 | 18888888888.8888 | 3.375e+10 |
| HUERS-P3-int:ERV1:LTR | 154598482522.696 | 61111111111.1112 | 1.18125e+11 |
| HUERS-P3b-int:ERV1:LTR | 149116266830.402 | 62777777777.7778 | 1.3375e+11 |
| Harlequin-int:ERV1:LTR | 766413753782.728 | 1.0475e+12 | 3.58125e+11 |
| Helitron1Na_Mam:Helitron:RC | 10964431384.5884 | 1e+10 | 1.75e+10 |
| Helitron1Nb_Mam:Helitron:RC | 6578658830.75305 | 5833333333.33333 | 6.875e+09 |
| Helitron2Na_Mam:Helitron:RC | 0 | 0 | 0 |
| Helitron2Na_Mam:Helitron:RC? | 27411078461.471 | 15277777777.7778 | 2e+10 |
| Helitron3Na_Mam:Helitron:RC | 41664839261.4358 | 23055555555.5555 | 5.125e+10 |
| Kanga11a:TcMar-Tc2:DNA | 73461690276.7424 | 39166666666.6667 | 6.5625e+10 |
| Kanga1:TcMar-Tc2:DNA | 27411078461.471 | 16388888888.8889 | 3.4375e+10 |
| Kanga1a:TcMar-Tc2:DNA | 65786588307.5305 | 53611111111.1112 | 7.6875e+10 |
| Kanga1b:TcMar-Tc2:DNA | 24121749046.0945 | 11944444444.4444 | 2.0625e+10 |
| Kanga1c:TcMar-Tc2:DNA | 69075917722.907 | 39166666666.6667 | 7.875e+10 |
| Kanga1d:TcMar-Tc2:DNA | 41664839261.4358 | 24444444444.4445 | 4.75e+10 |
| Kanga2_a:TcMar-Tc2:DNA | 52629270646.0242 | 40833333333.3333 | 7.75e+10 |
| L1HS:L1:LINE | 1630410946888.29 | 735277777777.778 | 9.08125e+11 |
| L1M1:L1:LINE | 1188544362089.38 | 696944444444.445 | 9.3e+11 |
| L1M2:L1:LINE | 722556028244.374 | 473333333333.333 | 6.675e+11 |
| L1M2a1:L1:LINE | 16446647076.8826 | 3888888888.88888 | 8.125e+09 |
| L1M2a:L1:LINE | 65786588307.5305 | 28055555555.5555 | 5.1875e+10 |
| L1M2b:L1:LINE | 13157317661.5061 | 10277777777.7778 | 1.1875e+10 |
| L1M2c:L1:LINE | 88811894215.1659 | 76388888888.8888 | 1.18125e+11 |
| L1M3:L1:LINE | 430902153414.325 | 245833333333.333 | 3.50625e+11 |
| L1M3a:L1:LINE | 122801631507.39 | 5.5e+10 | 1.0625e+11 |
| L1M3b:L1:LINE | 26314635323.0122 | 17777777777.7778 | 2.4375e+10 |
| L1M3c:L1:LINE | 29603964738.3887 | 2e+10 | 3.3125e+10 |
| L1M3d:L1:LINE | 18639533353.8003 | 8333333333.33333 | 2.5e+10 |
| L1M3de:L1:LINE | 26314635323.0122 | 2.75e+10 | 3.0625e+10 |
| L1M3e:L1:LINE | 53725713784.4831 | 50277777777.7778 | 5.75e+10 |
| L1M3f:L1:LINE | 245603263014.78 | 105833333333.333 | 1.6e+11 |
| L1M4:L1:LINE | 2770711810885.49 | 1431944444444.45 | 2.355e+12 |
| L1M4a1:L1:LINE | 304811192491.557 | 142222222222.222 | 2.2e+11 |
| L1M4a2:L1:LINE | 121705188368.931 | 80833333333.3333 | 1.4125e+11 |
| L1M4b:L1:LINE | 430902153414.325 | 250555555555.555 | 3.675e+11 |
| L1M4c:L1:LINE | 372790667076.007 | 205277777777.778 | 3.05625e+11 |
| L1M5:L1:LINE | 4205955879128.11 | 2306666666666.67 | 4.208125e+12 |
| L1M6:L1:LINE | 292750317968.51 | 191666666666.667 | 2.81875e+11 |
| L1M6B:L1:LINE | 6578658830.75305 | 12222222222.2222 | 1.875e+10 |
| L1M7:L1:LINE | 175430902153.414 | 111666666666.667 | 1.55e+11 |
| L1M8:L1:LINE | 152405596245.779 | 78333333333.3333 | 1.25e+11 |
| L1M:L1:LINE | 16446647076.8826 | 6666666666.66667 | 2.1875e+10 |
| L1MA10:L1:LINE | 265339239507.039 | 150277777777.778 | 2.0375e+11 |
| L1MA1:L1:LINE | 605236612429.278 | 350833333333.333 | 4.2125e+11 |
| L1MA2:L1:LINE | 1126047103197.23 | 683055555555.555 | 8.95625e+11 |
| L1MA3:L1:LINE | 1702776194026.58 | 1018888888888.89 | 1.424375e+12 |
| L1MA4:L1:LINE | 1635893162580.59 | 948888888888.888 | 1.2625e+12 |
| L1MA4A:L1:LINE | 916626463751.589 | 558611111111.112 | 7.375e+11 |
| L1MA5:L1:LINE | 735713345905.881 | 392777777777.778 | 7.8125e+11 |
| L1MA5A:L1:LINE | 403491074952.854 | 210555555555.555 | 3.09375e+11 |
| L1MA6:L1:LINE | 392526643568.265 | 240555555555.555 | 3.6e+11 |
| L1MA7:L1:LINE | 482434980921.891 | 2.875e+11 | 4.59375e+11 |
| L1MA8:L1:LINE | 622779702644.623 | 391111111111.112 | 5.33125e+11 |
| L1MA9:L1:LINE | 1138107977720.27 | 589444444444.445 | 8.74375e+11 |
| L1MB1:L1:LINE | 412262620060.523 | 221388888888.888 | 3.31875e+11 |
| L1MB2:L1:LINE | 1349721503442.83 | 621666666666.667 | 9.6125e+11 |
| L1MB3:L1:LINE | 2123810359194.77 | 1114444444444.45 | 1.968125e+12 |
| L1MB4:L1:LINE | 1015306346212.88 | 568611111111.112 | 9.79375e+11 |

| | | | |
|---|---|---|---|
| L1MB5:L1:LINE | 1166615499320.21 | 522222222222.222 | 9.775e+11 |
| L1MB7:L1:LINE | 2930792509100.48 | 1504444444444.45 | 2.72375e+12 |
| L1MB8:L1:LINE | 1906714617779.92 | 1128333333333.33 | 1.874375e+12 |
| L1MC1:L1:LINE | 1133722205166.44 | 644444444444.445 | 1.003125e+12 |
| L1MC2:L1:LINE | 459409675014.255 | 201666666666.667 | 4.23125e+11 |
| L1MC3:L1:LINE | 1401254330950.4 | 6.075e+11 | 1.111875e+12 |
| L1MC4:L1:LINE | 2283891057409.76 | 1099722222222.22 | 2.096875e+12 |
| L1MC4a:L1:LINE | 1431954738827.25 | 6.05e+11 | 1.2525e+12 |
| L1MC5:L1:LINE | 1246655848427.7 | 728055555555.555 | 1.27625e+12 |
| L1MC5a:L1:LINE | 1799263190210.96 | 809444444444.445 | 1.645e+12 |
| L1MC:L1:LINE | 436384369106.619 | 226944444444.445 | 4.20625e+11 |
| L1MCa:L1:LINE | 634840577167.669 | 337222222222.222 | 5.03125e+11 |
| L1MCb:L1:LINE | 111837200122.802 | 64722222222.2222 | 9.0625e+10 |
| L1MCc:L1:LINE | 140344721722.731 | 80277777777.7778 | 1.11875e+11 |
| L1MD1:L1:LINE | 1604096311565.28 | 641388888888.888 | 1.10625e+12 |
| L1MD2:L1:LINE | 1117275558089.56 | 605833333333.333 | 9.45e+11 |
| L1MD3:L1:LINE | 647997894829.176 | 297222222222.222 | 5.1375e+11 |
| L1MD:L1:LINE | 827814569536.424 | 413055555555.555 | 7.40625e+11 |
| L1MDa:L1:LINE | 809175036182.623 | 459166666666.667 | 6.66875e+11 |
| L1MDb:L1:LINE | 55918600061.4007 | 25833333333.3333 | 4.625e+10 |
| L1ME1:L1:LINE | 3151177579930.71 | 1603888888888.89 | 2.856875e+12 |
| L1ME2:L1:LINE | 1410025876058.07 | 696388888888.888 | 1.176875e+12 |
| L1ME2z:L1:LINE | 550414455506.338 | 231666666666.667 | 5.18125e+11 |
| L1ME3:L1:LINE | 758738651813.517 | 368888888888.888 | 7.025e+11 |
| L1ME3A:L1:LINE | 1020788561905.18 | 555277777777.778 | 9.20625e+11 |
| L1ME3B:L1:LINE | 675408973290.646 | 365277777777.778 | 6.44375e+11 |
| L1ME3C:L1:LINE | 274110784614.71 | 158055555555.556 | 3.01875e+11 |
| L1ME3Cz:L1:LINE | 927590895136.179 | 513888888888.888 | 8.93125e+11 |
| L1ME3D:L1:LINE | 271917898337.792 | 143055555555.556 | 2.83125e+11 |
| L1ME3E:L1:LINE | 450638129906.583 | 206388888888.888 | 3.85e+11 |
| L1ME3F:L1:LINE | 426516380860.49 | 260555555555.555 | 4.43125e+11 |
| L1ME3G:L1:LINE | 1160036840489.45 | 713333333333.333 | 1.123125e+12 |
| L1ME4a:L1:LINE | 2037191351256.52 | 1162777777777.78 | 2.041875e+12 |
| L1ME4b:L1:LINE | 1903425288364.55 | 1.0075e+12 | 1.76375e+12 |
| L1ME4c:L1:LINE | 656769439936.844 | 386944444444.445 | 7.2875e+11 |
| L1ME5:L1:LINE | 100872768738.213 | 86388888888.8888 | 1.44375e+11 |
| L1MEa:L1:LINE | 10964431384.5884 | 4444444444.44445 | 1.0625e+10 |
| L1MEb:L1:LINE | 175430902153.414 | 84166666666.6667 | 1.30625e+11 |
| L1MEc:L1:LINE | 1765273452918.73 | 960277777777.778 | 1.4e+12 |
| L1MEd:L1:LINE | 1587649664488.4 | 840277777777.778 | 1.435e+12 |
| L1MEf:L1:LINE | 7047936494013.41 | 1744166666666.67 | 4.85125e+12 |
| L1MEg1:L1:LINE | 26314635323.0122 | 14444444444.4444 | 2.25e+10 |
| L1MEg2:L1:LINE | 8771545107.67073 | 14444444444.4444 | 2.5625e+10 |
| L1MEg:L1:LINE | 1166615499320.21 | 686388888888.888 | 1.013125e+12 |
| L1MEh:L1:LINE | 180913117845.709 | 86388888888.8888 | 1.39375e+11 |
| L1MEi:L1:LINE | 359633349414.5 | 224166666666.667 | 3.5875e+11 |
| L1MEj:L1:LINE | 400201745537.477 | 156944444444.444 | 3.325e+11 |
| L1P1:L1:LINE | 490110082891.103 | 224444444444.445 | 4.6125e+11 |
| L1P2:L1:LINE | 665540985044.516 | 340833333333.333 | 5.0625e+11 |
| L1P3:L1:LINE | 354151133722.205 | 217222222222.222 | 2.93125e+11 |
| L1P3b:L1:LINE | 6578658830.75305 | 5555555555.55555 | 7.5e+09 |
| L1P4:L1:LINE | 245603263014.78 | 108888888888.889 | 2.14375e+11 |
| L1P4a:L1:LINE | 41664839261.4358 | 21944444444.4445 | 3.3125e+10 |
| L1P4b:L1:LINE | 15350203938.4237 | 7.5e+09 | 6.25e+09 |
| L1P4c:L1:LINE | 3289329415.37652 | 555555555.555555 | 0 |
| L1P4d:L1:LINE | 10964431384.5884 | 5555555555.55555 | 4.375e+09 |
| L1P4e:L1:LINE | 10964431384.5884 | 6388888888.88888 | 5e+09 |
| L1P5:L1:LINE | 25218192184.5533 | 9166666666.66667 | 2.125e+10 |
| L1P:L1:LINE | 1096443138.45884 | 555555555.555555 | 0 |
| L1PA10:L1:LINE | 1381518354458.14 | 689722222222.222 | 9.75e+11 |
| L1PA11:L1:LINE | 987895267751.414 | 534166666666.667 | 7.59375e+11 |
| L1PA12:L1:LINE | 171045129599.579 | 78611111111.1112 | 1.26875e+11 |
| L1PA13:L1:LINE | 958291303013.026 | 481944444444.445 | 7.40625e+11 |
| L1PA14:L1:LINE | 588789965352.397 | 376944444444.445 | 4.0625e+11 |

| | | | |
|---|---|---|---|
| L1PA15-16:L1:LINE | 143634051138.108 | 78333333333.3333 | 1.25625e+11 |
| L1PA15:L1:LINE | 779571071444.235 | 395555555555.555 | 6.16875e+11 |
| L1PA16:L1:LINE | 1354107275996.67 | 746388888888.888 | 1.100625e+12 |
| L1PA17:L1:LINE | 392526643568.265 | 181666666666.667 | 2.84375e+11 |
| L1PA2:L1:LINE | 6168589096969.44 | 2891666666666.67 | 3.7475e+12 |
| L1PA3:L1:LINE | 5950396912416.13 | 2.8225e+12 | 4.223125e+12 |
| L1PA4:L1:LINE | 2898995658085.17 | 1593888888888.89 | 2.39875e+12 |
| L1PA5:L1:LINE | 3434059909653.08 | 1738333333333.33 | 2.729375e+12 |
| L1PA6:L1:LINE | 3245471689838.17 | 1.5775e+12 | 2.42e+12 |
| L1PA7:L1:LINE | 3227928599622.82 | 1579722222222.22 | 2.71625e+12 |
| L1PA8:L1:LINE | 749967106705.848 | 426944444444.445 | 6.01875e+11 |
| L1PA8A:L1:LINE | 281785886583.922 | 141111111111.111 | 2.33125e+11 |
| L1PB1:L1:LINE | 2596377351870.53 | 1368055555555.55 | 1.88375e+12 |
| L1PB2:L1:LINE | 277400114030.086 | 191666666666.667 | 2.3625e+11 |
| L1PB3:L1:LINE | 469277663260.384 | 2.675e+11 | 3.41875e+11 |
| L1PB4:L1:LINE | 516424718214.113 | 440277777777.778 | 4.61875e+11 |
| L1PB:L1:LINE | 120608745230.472 | 8.25e+10 | 9.375e+10 |
| L1PBa1:L1:LINE | 121705188368.931 | 59444444444.4445 | 9.25e+10 |
| L1PBa:L1:LINE | 518617604491.03 | 2.55e+11 | 5.6375e+11 |
| L1PBb:L1:LINE | 49339941230.6477 | 18333333333.3333 | 3.625e+10 |
| L1PREC2:L1:LINE | 779571071444.235 | 367222222222.222 | 6.2875e+11 |
| L2-1_AMi:L2:LINE | 48243498092.1891 | 26111111111.1112 | 6e+10 |
| L2-1_Crp:L2:LINE | 2192886276.91768 | 0 | 0 |
| L2-2_Mam:L2:LINE | 0 | 0 | 0 |
| L2-3_AMi:L2:LINE | 0 | 0 | 0 |
| L2-3_Crp:L2:LINE | 141441164861.19 | 63055555555.5555 | 1.49375e+11 |
| L2:L2:LINE | 5160957852725.76 | 2703333333333.33 | 4.979375e+12 |
| L2a:L2:LINE | 14535546686548.8 | 7585833333333.33 | 1.3856875e+13 |
| L2b:L2:LINE | 7035875619490.36 | 3588055555555.55 | 6.67e+12 |
| L2c:L2:LINE | 9679400026314.64 | 5217777777777.78 | 9.483125e+12 |
| L2d2:L2:LINE | 0 | 0 | 0 |
| L2d:L2:LINE | 0 | 0 | 0 |
| L3:CR1:LINE | 3584272619621.95 | 2116388888888.88 | 3.49625e+12 |
| L3b:CR1:LINE | 752159992982.765 | 415833333333.333 | 6.79375e+11 |
| L4_A_Mam:RTE-X:LINE | 387044427875.97 | 204722222222.222 | 4.45e+11 |
| L4_B_Mam:RTE-X:LINE | 273014341476.251 | 119444444444.445 | 2.63125e+11 |
| L4_C_Mam:RTE-X:LINE | 232445945353.274 | 139722222222.222 | 2.65625e+11 |
| L5:RTE-X:LINE | 39471952984.5182 | 19166666666.6667 | 4.875e+10 |
| LOR1-int:ERV1:LTR | 137055392307.355 | 51944444444.4445 | 1.075e+11 |
| LOR1a:ERV1:LTR | 23025305907.6356 | 7.5e+09 | 2.9375e+10 |
| LOR1b:ERV1:LTR | 52629270646.0242 | 24444444444.4445 | 4.125e+10 |
| LSAU:Satellite:Satellite | 4385772553.83536 | 555555555.555555 | 0 |
| LTR06:ERV1:LTR | 2192886276.91768 | 2777777777.77778 | 2.5e+09 |
| LTR101_Mam:ERVL:LTR | 43857725538.3536 | 2.25e+10 | 3.5625e+10 |
| LTR102_Mam:ERVL:LTR | 55918600061.4007 | 14722222222.2222 | 4.75e+10 |
| LTR103_Mam:ERV1:LTR | 5482215692.2942 | 6388888888.88888 | 7.5e+09 |
| LTR103b_Mam:ERV1:LTR | 10964431384.5884 | 4166666666.66667 | 8.75e+09 |
| LTR104_Mam:Gypsy:LTR | 89908337353.6248 | 38333333333.3333 | 9.5625e+10 |
| LTR105_Mam:ERVL:LTR | 36182623569.1417 | 1.25e+10 | 2.4375e+10 |
| LTR106_Mam:LTR:LTR | 27411078461.471 | 29444444444.4445 | 4.4375e+10 |
| LTR107_Mam:LTR:LTR | 21928862769.1768 | 1.25e+10 | 1.9375e+10 |
| LTR108a_Mam:ERVL:LTR | 7675101969.21189 | 1944444444.44445 | 1.25e+09 |
| LTR108b_Mam:ERVL:LTR | 0 | 1111111111.11111 | 0 |
| LTR108c_Mam:ERVL:LTR | 0 | 277777777.777778 | 6.25e+08 |
| LTR108d_Mam:ERVL:LTR | 4385772553.83536 | 2222222222.22222 | 3.75e+09 |
| LTR108e_Mam:ERVL:LTR | 0 | 2.5e+09 | 3.125e+09 |
| LTR109A2:ERV1:LTR | 0 | 277777777.777778 | 0 |
| LTR10A:ERV1:LTR | 32893294153.7652 | 17222222222.2222 | 2.875e+10 |
| LTR10B1:ERV1:LTR | 9867988246.12957 | 833333333.333333 | 1.375e+10 |
| LTR10B2:ERV1:LTR | 28507521599.9298 | 19722222222.2222 | 2.5625e+10 |
| LTR10B:ERV1:LTR | 2192886276.91768 | 833333333.333333 | 1.875e+09 |
| LTR10C:ERV1:LTR | 69075917722.907 | 50277777777.7778 | 6.25e+10 |
| LTR10D:ERV1:LTR | 20832419630.718 | 8333333333.33333 | 1.5e+10 |
| LTR10E:ERV1:LTR | 63593702030.6126 | 25833333333.3333 | 6.25e+10 |

| | | | |
|---|---|---|---|
| LTR10F:ERV1:LTR | 41664839261.4358 | 15833333333.3333 | 3e+10 |
| LTR10G:ERV1:LTR | 117319415815.096 | 39166666666.6667 | 8.875e+10 |
| LTR12:ERV1:LTR | 67979474584.448 | 41944444444.4445 | 5.875e+10 |
| LTR12B:ERV1:LTR | 19735976492.2591 | 10833333333.3333 | 1.8125e+10 |
| LTR12C:ERV1:LTR | 777378185167.318 | 286111111111.112 | 5.40625e+11 |
| LTR12D:ERV1:LTR | 42761282399.8947 | 14444444444.4444 | 2.5e+10 |
| LTR12E:ERV1:LTR | 24121749046.0945 | 5277777777.77778 | 6.25e+09 |
| LTR12F:ERV1:LTR | 41664839261.4358 | 18888888888.8888 | 4.3125e+10 |
| LTR12_:ERV1:LTR | 32893294153.7652 | 1.25e+10 | 3.25e+10 |
| LTR12_v:ERV1:LTR | 0 | 0 | 0 |
| LTR13:ERVK:LTR | 324547168983.817 | 1.3e+11 | 2.6125e+11 |
| LTR13A:ERVK:LTR | 203938423753.344 | 64166666666.6667 | 1.425e+11 |
| LTR13_:ERVK:LTR | 14253760799.9649 | 6666666666.66667 | 1.375e+10 |
| LTR13_v:ERVK:LTR | 0 | 0 | 0 |
| LTR14:ERVK:LTR | 16446647076.8826 | 17222222222.2222 | 2.4375e+10 |
| LTR14A:ERVK:LTR | 5482215692.2942 | 2.5e+09 | 3.125e+09 |
| LTR14B:ERVK:LTR | 26314635323.0122 | 13611111111.1111 | 2.1875e+10 |
| LTR14C:ERVK:LTR | 1096443138.45884 | 1111111111.11111 | 6.25e+08 |
| LTR15:ERV1:LTR | 27411078461.471 | 1e+10 | 1.75e+10 |
| LTR16:ERVL:LTR | 82233235384.4129 | 41388888888.8888 | 6.375e+10 |
| LTR16A1:ERVL:LTR | 92101223630.5424 | 43888888888.8888 | 7.5625e+10 |
| LTR16A2:ERVL:LTR | 53725713784.4831 | 3.25e+10 | 4.875e+10 |
| LTR16A:ERVL:LTR | 200649094337.968 | 95555555555.5555 | 2.40625e+11 |
| LTR16B1:ERVL:LTR | 13157317661.5061 | 1.5e+10 | 1.6875e+10 |
| LTR16B2:ERVL:LTR | 78943905969.0364 | 35833333333.3333 | 6.125e+10 |
| LTR16B:ERVL:LTR | 71268803999.8245 | 21666666666.6667 | 3.4375e+10 |
| LTR16C:ERVL:LTR | 340993816060.699 | 154722222222.222 | 2.91875e+11 |
| LTR16D1:ERVL:LTR | 5482215692.2942 | 1388888888.88889 | 2.5e+09 |
| LTR16D2:ERVL:LTR | 6578658830.75305 | 10277777777.7778 | 1.875e+10 |
| LTR16D:ERVL:LTR | 36182623569.1417 | 26666666666.6667 | 3.375e+10 |
| LTR16E1:ERVL:LTR | 49339941230.6477 | 2.5e+10 | 4.5625e+10 |
| LTR16E2:ERVL:LTR | 72365247138.2834 | 30555555555.5555 | 4.875e+10 |
| LTR17:ERV1:LTR | 160080698214.991 | 61944444444.4445 | 1.06875e+11 |
| LTR18A:ERVL:LTR | 9867988246.12957 | 9166666666.66667 | 1.0625e+10 |
| LTR18B:ERVL:LTR | 29603964738.3887 | 14166666666.6667 | 4.875e+10 |
| LTR18C:ERVL:LTR | 0 | 0 | 0 |
| LTR19-int:ERV1:LTR | 18639533353.8003 | 11388888888.8889 | 1.375e+10 |
| LTR19A:ERV1:LTR | 16446647076.8826 | 1.5e+10 | 2.875e+10 |
| LTR19B:ERV1:LTR | 14253760799.9649 | 3888888888.88888 | 1.25e+10 |
| LTR19C:ERV1:LTR | 13157317661.5061 | 7.5e+09 | 5.625e+09 |
| LTR1:ERV1:LTR | 12060874523.0473 | 2222222222.22222 | 5.625e+09 |
| LTR1A1:ERV1:LTR | 52629270646.0242 | 18888888888.8888 | 2.5e+10 |
| LTR1A2:ERV1:LTR | 70172360861.3659 | 36388888888.8888 | 3.9375e+10 |
| LTR1B0:ERV1:LTR | 3289329415.37652 | 555555555.555555 | 1.25e+09 |
| LTR1B1:ERV1:LTR | 15350203938.4237 | 13055555555.5556 | 1.0625e+10 |
| LTR1B:ERV1:LTR | 9867988246.12957 | 4166666666.66667 | 5e+09 |
| LTR1C1:ERV1:LTR | 4385772553.83536 | 2.5e+09 | 0 |
| LTR1C3:ERV1:LTR | 3289329415.37652 | 1944444444.44445 | 1.875e+09 |
| LTR1C:ERV1:LTR | 3289329415.37652 | 1388888888.88889 | 3.75e+09 |
| LTR1D1:ERV1:LTR | 44954168676.8126 | 21944444444.4445 | 2.875e+10 |
| LTR1D:ERV1:LTR | 15350203938.4237 | 8888888888.88888 | 1e+10 |
| LTR1E:ERV1:LTR | 6578658830.75305 | 3611111111.11112 | 6.875e+09 |
| LTR1F1:ERV1:LTR | 18639533353.8003 | 8333333333.33333 | 6.875e+09 |
| LTR1F2:ERV1:LTR | 3289329415.37652 | 1111111111.11111 | 5.625e+09 |
| LTR1F:ERV1:LTR | 3289329415.37652 | 4444444444.44445 | 6.875e+09 |
| LTR21A:ERV1:LTR | 4385772553.83536 | 5e+09 | 9.375e+09 |
| LTR21B:ERV1:LTR | 13157317661.5061 | 3888888888.88888 | 1.0625e+10 |
| LTR21C:ERVL:LTR | 0 | 555555555.555555 | 0 |
| LTR22:ERVK:LTR | 3289329415.37652 | 555555555.555555 | 2.5e+09 |
| LTR22A:ERVK:LTR | 3289329415.37652 | 1944444444.44445 | 1.1875e+10 |
| LTR22B1:ERVK:LTR | 18639533353.8003 | 5e+09 | 8.125e+09 |
| LTR22B2:ERVK:LTR | 1096443138.45884 | 2222222222.22222 | 5.625e+09 |
| LTR22B:ERVK:LTR | 6578658830.75305 | 9722222222.22222 | 1.6875e+10 |
| LTR22C0:ERVK:LTR | 1096443138.45884 | 3333333333.33333 | 9.375e+09 |

| | | | |
|---|---|---|---|
| LTR22C2:ERVK:LTR | 3289329415.37652 | 1111111111.11111 | 1.875e+09 |
| LTR22C:ERVK:LTR | 1096443138.45884 | 0 | 2.5e+09 |
| LTR22E:ERVK:LTR | 87715451076.7073 | 25277777777.7778 | 6.4375e+10 |
| LTR23-int:ERV1:LTR | 5482215692.2942 | 1666666666.66667 | 4.375e+09 |
| LTR23:ERV1:LTR | 55918600061.4007 | 23055555555.5555 | 4.125e+10 |
| LTR24:ERV1:LTR | 116222972676.637 | 51388888888.8888 | 1.15e+11 |
| LTR24B:ERV1:LTR | 30700407876.8475 | 10833333333.3333 | 2.9375e+10 |
| LTR24C:ERV1:LTR | 9867988246.12957 | 6388888888.88888 | 1.25e+10 |
| LTR25-int:ERV1:LTR | 40568396122.9771 | 25277777777.7778 | 6.5625e+10 |
| LTR25:ERV1:LTR | 5482215692.2942 | 1388888888.88889 | 6.25e+09 |
| LTR26:ERV1:LTR | 40568396122.9771 | 27777777777.7778 | 2.9375e+10 |
| LTR26B:ERV1:LTR | 3289329415.37652 | 833333333.333333 | 1.25e+09 |
| LTR26C:ERV1:LTR | 14253760799.9649 | 8333333333.33333 | 3.125e+09 |
| LTR26D:ERV1:LTR | 4385772553.83536 | 1388888888.88889 | 3.75e+09 |
| LTR26E:ERV1:LTR | 7675101969.21189 | 2222222222.22222 | 2.5e+09 |
| LTR2752:ERV1:LTR | 3289329415.37652 | 6944444444.44445 | 8.125e+09 |
| LTR27:ERV1:LTR | 4385772553.83536 | 2222222222.22222 | 6.875e+09 |
| LTR27B:ERV1:LTR | 6578658830.75305 | 3888888888.88888 | 1.875e+09 |
| LTR27C:ERV1:LTR | 17543090215.3414 | 6944444444.44445 | 1.4375e+10 |
| LTR27D:ERV1:LTR | 5482215692.2942 | 2222222222.22222 | 2.5e+09 |
| LTR27E:ERV1:LTR | 5482215692.2942 | 5e+09 | 8.125e+09 |
| LTR28:ERV1:LTR | 25218192184.5533 | 8611111111.11112 | 2.25e+10 |
| LTR28B:ERV1:LTR | 15350203938.4237 | 9722222222.22222 | 1.0625e+10 |
| LTR28C:ERV1:LTR | 60304372615.2361 | 30277777777.7778 | 6.5e+10 |
| LTR29:ERV1:LTR | 81136792245.954 | 34444444444.4445 | 5.9375e+10 |
| LTR2:ERV1:LTR | 183106004122.626 | 79444444444.4445 | 1.43125e+11 |
| LTR2B:ERV1:LTR | 77847462830.5775 | 128611111111.111 | 5.75e+10 |
| LTR2C:ERV1:LTR | 48243498092.1891 | 11666666666.6667 | 2.625e+10 |
| LTR30:ERV1:LTR | 7675101969.21189 | 4722222222.22222 | 3.125e+09 |
| LTR31:ERV1:LTR | 3289329415.37652 | 2222222222.22222 | 4.375e+09 |
| LTR32:ERVL:LTR | 30700407876.8475 | 19722222222.2222 | 3.8125e+10 |
| LTR33:ERVL:LTR | 354151133722.205 | 148055555555.556 | 2.675e+11 |
| LTR33A:ERVL:LTR | 89908337353.6248 | 40833333333.3333 | 7.75e+10 |
| LTR33A_:ERVL:LTR | 215999298276.391 | 74722222222.2222 | 1.65625e+11 |
| LTR33A_v:ERVL:LTR | 0 | 0 | 0 |
| LTR33B:ERVL:LTR | 32893294153.7652 | 13333333333.3333 | 2.75e+10 |
| LTR33C:ERVL:LTR | 117319415815.096 | 23888888888.8888 | 1.3625e+11 |
| LTR34:ERV1:LTR | 37279066707.6007 | 26388888888.8888 | 2.3125e+10 |
| LTR35:ERV1:LTR | 33989737292.2242 | 15555555555.5556 | 2.375e+10 |
| LTR35A:ERV1:LTR | 3289329415.37652 | 1111111111.11111 | 2.5e+09 |
| LTR35B:ERV1:LTR | 10964431384.5884 | 4166666666.66667 | 4.375e+09 |
| LTR36:ERV1:LTR | 9867988246.12957 | 9166666666.66667 | 1.8125e+10 |
| LTR37-int:ERV1:LTR | 48243498092.1891 | 20833333333.3333 | 3.75e+10 |
| LTR37A:ERV1:LTR | 70172360861.3659 | 35555555555.5555 | 5.125e+10 |
| LTR37B:ERV1:LTR | 47147054953.7301 | 16666666666.6667 | 2.6875e+10 |
| LTR38-int:ERV1:LTR | 0 | 555555555.555555 | 1.875e+09 |
| LTR38:ERV1:LTR | 26314635323.0122 | 8055555555.55555 | 2e+10 |
| LTR38A1:ERV1:LTR | 2192886276.91768 | 833333333.333333 | 2.5e+09 |
| LTR38B:ERV1:LTR | 21928862769.1768 | 8611111111.11112 | 1.75e+10 |
| LTR38C:ERV1:LTR | 15350203938.4237 | 5277777777.77778 | 2.125e+10 |
| LTR39-int:ERV1:LTR | 4385772553.83536 | 5277777777.77778 | 6.875e+09 |
| LTR39:ERV1:LTR | 52629270646.0242 | 26111111111.1112 | 4.125e+10 |
| LTR3:ERVK:LTR | 3289329415.37652 | 5555555555.55555 | 3.125e+09 |
| LTR3A:ERVK:LTR | 18639533353.8003 | 11666666666.6667 | 1.625e+10 |
| LTR3B:ERVK:LTR | 2192886276.91768 | 1111111111.11111 | 5e+09 |
| LTR3B_:ERVK:LTR | 7675101969.21189 | 4444444444.44445 | 5e+09 |
| LTR3B_v:ERVK:LTR | 0 | 0 | 0 |
| LTR40A1:ERVL:LTR | 12060874523.0473 | 4722222222.22222 | 4.375e+09 |
| LTR40a:ERVL:LTR | 107451427568.966 | 51944444444.4445 | 7.0625e+10 |
| LTR40b:ERVL:LTR | 59207929476.7775 | 4.5e+10 | 7.6875e+10 |
| LTR40c:ERVL:LTR | 75654576553.6599 | 32777777777.7778 | 6.0625e+10 |
| LTR41:ERVL:LTR | 110740756984.343 | 65277777777.7778 | 8.4375e+10 |
| LTR41B:ERVL:LTR | 198456208061.05 | 68888888888.8888 | 9.5e+10 |
| LTR41C:ERVL:LTR | 53725713784.4831 | 23888888888.8888 | 4.0625e+10 |

| | | | |
|---|---|---|---|
| LTR42:ERVL:LTR | 80040349107.4954 | 4e+10 | 7.6875e+10 |
| LTR43-int:ERV1:LTR | 0 | 0 | 1.875e+09 |
| LTR43:ERV1:LTR | 92101223630.5424 | 38611111111.1112 | 9.75e+10 |
| LTR43B:ERV1:LTR | 6578658830.75305 | 2.5e+09 | 4.375e+09 |
| LTR44:ERV1:LTR | 17543090215.3414 | 8333333333.33333 | 5.625e+09 |
| LTR45:ERV1:LTR | 3289329415.37652 | 2777777777.77778 | 5.625e+09 |
| LTR45B:ERV1:LTR | 33989737292.2242 | 18888888888.8888 | 2.125e+10 |
| LTR45C:ERV1:LTR | 28507521599.9298 | 17222222222.2222 | 2.3125e+10 |
| LTR46-int:ERV1:LTR | 46050611815.2712 | 29722222222.2222 | 5.3125e+10 |
| LTR46:ERV1:LTR | 35086180430.6828 | 10555555555.5556 | 2e+10 |
| LTR47A2:ERVL:LTR | 3289329415.37652 | 2222222222.22222 | 6.25e+09 |
| LTR47A:ERVL:LTR | 62497258892.154 | 31666666666.6667 | 4.875e+10 |
| LTR47B2:ERVL:LTR | 15350203938.4237 | 3333333333.33333 | 4.375e+09 |
| LTR47B3:ERVL:LTR | 12060874523.0473 | 3888888888.88888 | 2.3125e+10 |
| LTR47B4:ERVL:LTR | 15350203938.4237 | 6388888888.88888 | 1.9375e+10 |
| LTR47B:ERVL:LTR | 1096443138.45884 | 277777777.777778 | 1.875e+09 |
| LTR48:ERV1:LTR | 50436384369.1066 | 28611111111.1112 | 5.1875e+10 |
| LTR48B:ERV1:LTR | 64690145169.0715 | 32222222222.2222 | 4.625e+10 |
| LTR49-int:ERV1:LTR | 93197666769.0013 | 6e+10 | 8.6875e+10 |
| LTR49:ERV1:LTR | 50436384369.1066 | 24166666666.6667 | 3.6875e+10 |
| LTR4:ERV1:LTR | 7675101969.21189 | 6944444444.44445 | 9.375e+09 |
| LTR50:ERVL:LTR | 96486996184.3778 | 63055555555.5555 | 1.13125e+11 |
| LTR51:ERV1:LTR | 14253760799.9649 | 5555555555.55555 | 1.8125e+10 |
| LTR52-int:ERVL:LTR | 10964431384.5884 | 10555555555.5556 | 1.3125e+10 |
| LTR52:ERVL:LTR | 19735976492.2591 | 16388888888.8889 | 1.875e+10 |
| LTR53-int:ERVL:LTR | 23025305907.6356 | 18055555555.5555 | 1.625e+10 |
| LTR53:ERVL:LTR | 66883031445.9891 | 40555555555.5555 | 8.25e+10 |
| LTR53B:ERVL:LTR | 7675101969.21189 | 3333333333.33333 | 1.875e+09 |
| LTR54:ERV1:LTR | 27411078461.471 | 19166666666.6667 | 3.1875e+10 |
| LTR54B:ERV1:LTR | 143634051138.108 | 40833333333.3333 | 1.05e+11 |
| LTR55:ERVL:LTR | 3289329415.37652 | 6388888888.88888 | 1.0625e+10 |
| LTR56:ERV1:LTR | 50436384369.1066 | 35277777777.7778 | 5.1875e+10 |
| LTR57-int:ERVL:LTR | 16446647076.8826 | 4444444444.44445 | 8.125e+09 |
| LTR57:ERVL:LTR | 10964431384.5884 | 4444444444.44445 | 2.5e+09 |
| LTR58:ERV1:LTR | 1096443138.45884 | 0 | 0 |
| LTR59:ERV1:LTR | 6578658830.75305 | 833333333.333333 | 3.75e+09 |
| LTR5:ERVK:LTR | 4385772553.83536 | 277777777.777778 | 1.875e+09 |
| LTR5A:ERVK:LTR | 150212709968.861 | 56666666666.6667 | 1.08125e+11 |
| LTR5B:ERVK:LTR | 194070435507.215 | 92222222222.2222 | 1.74375e+11 |
| LTR5_Hs:ERVK:LTR | 448445243629.666 | 178888888888.888 | 3.36875e+11 |
| LTR60:ERV1:LTR | 29603964738.3887 | 11388888888.8889 | 3.5e+10 |
| LTR60B:ERV1:LTR | 2192886276.91768 | 4166666666.66667 | 7.5e+09 |
| LTR61:ERV1:LTR | 65786588307.5305 | 21944444444.4445 | 4.75e+10 |
| LTR62:ERVL:LTR | 32893294153.7652 | 18333333333.3333 | 1.5e+10 |
| LTR64:ERV1:LTR | 41664839261.4358 | 17777777777.7778 | 3.1875e+10 |
| LTR65:ERV1:LTR | 41664839261.4358 | 14444444444.4444 | 2.75e+10 |
| LTR66:ERVL:LTR | 28507521599.9298 | 12222222222.2222 | 3e+10 |
| LTR67B:ERVL:LTR | 101969211876.672 | 46944444444.4445 | 8.5625e+10 |
| LTR68:ERV1:LTR | 8771545107.67073 | 1666666666.66667 | 4.375e+09 |
| LTR69:ERVL:LTR | 13157317661.5061 | 4722222222.22222 | 1.0625e+10 |
| LTR6A:ERV1:LTR | 127187404061.225 | 46944444444.4445 | 6.9375e+10 |
| LTR6B:ERV1:LTR | 122801631507.39 | 136666666666.667 | 5.75e+10 |
| LTR70:ERV1:LTR | 268628568922.416 | 1.65e+11 | 2.7e+11 |
| LTR71A:ERV1:LTR | 32893294153.7652 | 9444444444.44445 | 2e+10 |
| LTR71B:ERV1:LTR | 14253760799.9649 | 10555555555.5556 | 9.375e+09 |
| LTR72:ERV1:LTR | 19735976492.2591 | 8333333333.33333 | 1.3125e+10 |
| LTR72B:ERV1:LTR | 15350203938.4237 | 2777777777.77778 | 8.125e+09 |
| LTR73:ERV1:LTR | 5482215692.2942 | 2222222222.22222 | 5e+09 |
| LTR75:ERVL:LTR | 35086180430.6828 | 21944444444.4445 | 3.25e+10 |
| LTR75B:ERVL:LTR | 8771545107.67073 | 5277777777.77778 | 2.5e+09 |
| LTR75_1:ERV1:LTR | 15350203938.4237 | 5833333333.33333 | 1.125e+10 |
| LTR76:ERV1:LTR | 27411078461.471 | 19166666666.6667 | 2.4375e+10 |
| LTR77:ERV1:LTR | 3289329415.37652 | 2777777777.77778 | 6.25e+09 |
| LTR78:ERV1:LTR | 225867286522.521 | 1.05e+11 | 1.5875e+11 |

| | | | |
|---|---|---|---|
| LTR78B:ERV1:LTR | 150212709968.861 | 48611111111.1112 | 8.25e+10 |
| LTR79:ERVL:LTR | 120608745230.472 | 58611111111.1112 | 9.9375e+10 |
| LTR7:ERV1:LTR | 946230428489.98 | 558888888888.888 | 1.1525e+12 |
| LTR7A:ERV1:LTR | 1096443138.45884 | 555555555.555555 | 6.25e+08 |
| LTR7B:ERV1:LTR | 120608745230.472 | 46388888888.8888 | 8.3125e+10 |
| LTR7C:ERV1:LTR | 50436384369.1066 | 14444444444.4444 | 3.4375e+10 |
| LTR7Y:ERV1:LTR | 87715451076.7073 | 24166666666.6667 | 1.425e+11 |
| LTR80A:ERVL:LTR | 4385772553.83536 | 2222222222.22222 | 4.375e+09 |
| LTR80B:ERVL:LTR | 18639533353.8003 | 10833333333.3333 | 1.25e+10 |
| LTR81:Gypsy:LTR | 2192886276.91768 | 1111111111.11111 | 1.25e+09 |
| LTR81A:Gypsy:LTR | 21928862769.1768 | 14722222222.2222 | 1.5625e+10 |
| LTR81AB:Gypsy:LTR | 14253760799.9649 | 3611111111.11112 | 8.125e+09 |
| LTR81B:Gypsy:LTR | 13157317661.5061 | 7777777777.77778 | 2.1875e+10 |
| LTR81C:Gypsy:LTR | 10964431384.5884 | 5833333333.33333 | 9.375e+09 |
| LTR82A:ERVL:LTR | 18639533353.8003 | 11944444444.4444 | 1.8125e+10 |
| LTR82B:ERVL:LTR | 29603964738.3887 | 1.75e+10 | 3.8125e+10 |
| LTR83:ERVL:LTR | 14253760799.9649 | 15555555555.5556 | 1.5625e+10 |
| LTR84a:ERVL:LTR | 88811894215.1659 | 4.25e+10 | 4.6875e+10 |
| LTR84b:ERVL:LTR | 116222972676.637 | 76944444444.4445 | 5.5625e+10 |
| LTR85a:Gypsy:LTR | 59207929476.7775 | 33055555555.5555 | 4.1875e+10 |
| LTR85b:Gypsy:LTR | 44954168676.8126 | 18611111111.1112 | 4.125e+10 |
| LTR85c:Gypsy:LTR | 30700407876.8475 | 13333333333.3333 | 3.1875e+10 |
| LTR86A1:ERVL:LTR | 111837200122.802 | 6e+10 | 1.16875e+11 |
| LTR86A2:ERVL:LTR | 7675101969.21189 | 4444444444.44445 | 8.125e+09 |
| LTR86B1:ERVL:LTR | 5482215692.2942 | 3333333333.33333 | 7.5e+09 |
| LTR86B2:ERVL:LTR | 10964431384.5884 | 4722222222.22222 | 1.125e+10 |
| LTR86C:ERVL:LTR | 7675101969.21189 | 5555555555.55555 | 5e+09 |
| LTR87:ERVL:LTR | 89908337353.6248 | 66388888888.8888 | 1.28125e+11 |
| LTR88a:Gypsy:LTR | 18639533353.8003 | 5833333333.33333 | 8.75e+09 |
| LTR88b:Gypsy:LTR | 57015043199.8596 | 2.5e+10 | 5.875e+10 |
| LTR88c:Gypsy:LTR | 28507521599.9298 | 10555555555.5556 | 2.4375e+10 |
| LTR89:ERVL:LTR | 15350203938.4237 | 11666666666.6667 | 2.8125e+10 |
| LTR89B:ERVL:LTR | 0 | 0 | 0 |
| LTR8:ERV1:LTR | 372790667076.007 | 178888888888.888 | 3.4875e+11 |
| LTR8A:ERV1:LTR | 119512302092.014 | 65277777777.7778 | 1.09375e+11 |
| LTR8B:ERV1:LTR | 74558133415.201 | 27777777777.7778 | 4.3125e+10 |
| LTR90A:LTR:LTR | 33989737292.2242 | 13888888888.8889 | 2.4375e+10 |
| LTR90B:LTR:LTR | 15350203938.4237 | 6111111111.11112 | 6.875e+09 |
| LTR91:ERVL:LTR | 9867988246.12957 | 3055555555.55555 | 6.875e+09 |
| LTR91:LTR:LTR | 0 | 0 | 0 |
| LTR91A:LTR:LTR | 0 | 0 | 0 |
| LTR9:ERV1:LTR | 37279066707.6007 | 10833333333.3333 | 3.9375e+10 |
| LTR9A1:ERV1:LTR | 41664839261.4358 | 11944444444.4444 | 2.8125e+10 |
| LTR9B:ERV1:LTR | 58111486338.3185 | 20277777777.7778 | 4.4375e+10 |
| LTR9C:ERV1:LTR | 13157317661.5061 | 7777777777.77778 | 1.1875e+10 |
| LTR9D:ERV1:LTR | 57015043199.8596 | 31666666666.6667 | 5.5625e+10 |
| Looper:PiggyBac:DNA | 93197666769.0013 | 66388888888.8888 | 1.0875e+11 |
| MADE1:TcMar-Mariner:DNA | 164466470768.826 | 110833333333.333 | 2.04375e+11 |
| MADE2:TcMar-Mariner:DNA | 87715451076.7073 | 54722222222.2222 | 1.13125e+11 |
| MARE4:hAT-Tip100:DNA | 0 | 0 | 0 |
| MARE6:L1-Tx1:LINE | 0 | 0 | 0 |
| MARE8:DNA:DNA | 0 | 0 | 0 |
| MARE9:MARE9:Unknown | 0 | 0 | 0 |
| MARNA:TcMar-Mariner:DNA | 228060172799.439 | 127777777777.778 | 2.34375e+11 |
| MER101-int:ERV1:LTR | 33989737292.2242 | 18055555555.5555 | 3.625e+10 |
| MER101:ERV1:LTR | 13157317661.5061 | 3611111111.11112 | 1.875e+10 |
| MER101B:ERV1:LTR | 6578658830.75305 | 2222222222.22222 | 5.625e+09 |
| MER102a:hAT-Charlie:DNA | 317968510153.064 | 159166666666.667 | 2.8625e+11 |
| MER102b:hAT-Charlie:DNA | 340993816060.699 | 190277777777.778 | 3.1e+11 |
| MER102c:hAT-Charlie:DNA | 256567694399.369 | 148333333333.333 | 2.50625e+11 |
| MER103C:hAT-Charlie:DNA | 685276961536.775 | 407222222222.222 | 6.45625e+11 |
| MER104:TcMar-Tc2:DNA | 150212709968.861 | 68333333333.3333 | 1.3625e+11 |
| MER105:DNA:DNA | 0 | 0 | 0 |
| MER105:hAT-Charlie:DNA | 81136792245.954 | 73888888888.8888 | 1.1625e+11 |

| | | | |
|---|---|---|---|
| MER106A:hAT-Charlie:DNA | 31796851015.3064 | 23611111111.1112 | 5.6875e+10 |
| MER106B:hAT-Charlie:DNA | 39471952984.5182 | 27777777777.7778 | 3.6875e+10 |
| MER107:hAT-Charlie:DNA | 77847462830.5775 | 62777777777.7778 | 1.25625e+11 |
| MER110-int:ERV1:LTR | 23025305907.6356 | 9444444444.44445 | 1.75e+10 |
| MER110:ERV1:LTR | 21928862769.1768 | 8611111111.11112 | 9.375e+09 |
| MER110A:ERV1:LTR | 21928862769.1768 | 13888888888.8889 | 1.9375e+10 |
| MER112:hAT-Charlie:DNA | 309196965045.393 | 175277777777.778 | 3.08125e+11 |
| MER113:hAT-Charlie:DNA | 183106004122.626 | 109722222222.222 | 1.90625e+11 |
| MER113A:hAT-Charlie:DNA | 96486996184.3778 | 63055555555.5555 | 9.625e+10 |
| MER113B:hAT-Charlie:DNA | 43857725538.3536 | 2.25e+10 | 4.1875e+10 |
| MER115:hAT-Tip100:DNA | 146923380553.484 | 85277777777.7778 | 1.225e+11 |
| MER117:hAT-Charlie:DNA | 303714749353.099 | 1.575e+11 | 3.14375e+11 |
| MER119:hAT-Charlie:DNA | 77847462830.5775 | 4.5e+10 | 9.9375e+10 |
| MER11A:ERVK:LTR | 145826937415.026 | 60555555555.5555 | 1.0625e+11 |
| MER11B:ERVK:LTR | 156791368799.614 | 58611111111.1112 | 1.30625e+11 |
| MER11C:ERVK:LTR | 330029384676.111 | 136944444444.445 | 2.08125e+11 |
| MER11D:ERVK:LTR | 47147054953.7301 | 16944444444.4445 | 5.0625e+10 |
| MER121:TcMar:DNA | 0 | 0 | 0 |
| MER121:hAT:DNA | 27411078461.471 | 20555555555.5555 | 2.8125e+10 |
| MER121B:hAT:DNA | 1096443138.45884 | 0 | 1.25e+09 |
| MER123:DNA:DNA? | 0 | 0 | 1.875e+09 |
| MER124:DNA:DNA | 0 | 0 | 0 |
| MER124:DNA:DNA? | 15350203938.4237 | 11666666666.6667 | 1.1875e+10 |
| MER125:DNA:DNA | 5482215692.2942 | 833333333.333333 | 4.375e+09 |
| MER125:Merlin:DNA | 0 | 0 | 0 |
| MER126:DNA:DNA | 2192886276.91768 | 2777777777.77778 | 7.5e+09 |
| MER127:TcMar-Tigger:DNA | 6578658830.75305 | 3055555555.55555 | 3.75e+09 |
| MER129:LTR:LTR | 0 | 0 | 0 |
| MER129:LTR:LTR? | 5482215692.2942 | 6388888888.88888 | 5e+09 |
| MER130:DNA:DNA | 4385772553.83536 | 3333333333.33333 | 5.625e+09 |
| MER131:DNA:DNA | 0 | 0 | 0 |
| MER131:DNA:DNA? | 36182623569.1417 | 14444444444.4444 | 2.3125e+10 |
| MER132:TcMar-Pogo:DNA | 1096443138.45884 | 1944444444.44445 | 0 |
| MER133A:DNA:DNA? | 21928862769.1768 | 7777777777.77778 | 5e+09 |
| MER133A:PIF-Harbinger:DNA | 0 | 0 | 0 |
| MER133B:DNA:DNA? | 6578658830.75305 | 1666666666.66667 | 6.25e+08 |
| MER134:Crypton:DNA | 0 | 0 | 0 |
| MER134:DNA:DNA? | 2192886276.91768 | 1944444444.44445 | 5e+09 |
| MER135:DNA:DNA | 21928862769.1768 | 12222222222.2222 | 3.25e+10 |
| MER136:DNA:DNA | 0 | 0 | 0 |
| MER136:Merlin:DNA | 0 | 0 | 0 |
| MER1A:hAT-Charlie:DNA | 400201745537.477 | 216944444444.445 | 4.18125e+11 |
| MER1B:hAT-Charlie:DNA | 652383667383.01 | 364722222222.222 | 6.65625e+11 |
| MER20:hAT-Charlie:DNA | 1484584009473.27 | 816388888888.888 | 1.44125e+12 |
| MER20B:hAT-Charlie:DNA | 177623788430.332 | 126944444444.445 | 2.2e+11 |
| MER21-int:ERVL:LTR | 33989737292.2242 | 18333333333.3333 | 3.125e+10 |
| MER21A:ERVL:LTR | 245603263014.78 | 116111111111.111 | 1.975e+11 |
| MER21B:ERVL:LTR | 437480812245.076 | 207777777777.778 | 4.29375e+11 |
| MER21C:ERVL:LTR | 478049208368.053 | 250277777777.778 | 5.14375e+11 |
| MER2:TcMar-Tigger:DNA | 978027279505.285 | 495555555555.555 | 1.00875e+12 |
| MER2B:TcMar-Tigger:DNA | 142537607999.649 | 1.025e+11 | 1.8125e+11 |
| MER30:hAT-Charlie:DNA | 436384369106.619 | 258888888888.888 | 5.5125e+11 |
| MER30B:hAT-Charlie:DNA | 17543090215.3414 | 3611111111.11112 | 1.8125e+10 |
| MER31-int:ERV1:LTR | 48243498092.1891 | 23611111111.1112 | 6.125e+10 |
| MER31A:ERV1:LTR | 74558133415.201 | 28333333333.3333 | 5.6875e+10 |
| MER31B:ERV1:LTR | 44954168676.8126 | 27777777777.7778 | 4.125e+10 |
| MER33:hAT-Charlie:DNA | 905662032367 | 424166666666.667 | 8.49375e+11 |
| MER34-int:ERV1:LTR | 64690145169.0715 | 47777777777.7778 | 4.3125e+10 |
| MER34:ERV1:LTR | 50436384369.1066 | 3e+10 | 4.625e+10 |
| MER34A1:ERV1:LTR | 54822156922.9421 | 2.75e+10 | 3.3125e+10 |
| MER34A:ERV1:LTR | 144730494276.567 | 55277777777.7778 | 9.9375e+10 |
| MER34B-int:ERV1:LTR | 70172360861.3659 | 23333333333.3333 | 3.6875e+10 |
| MER34B:ERV1:LTR | 26314635323.0122 | 10277777777.7778 | 3.25e+10 |
| MER34C2:ERV1:LTR | 57015043199.8596 | 25555555555.5555 | 5.625e+10 |

| | | | |
|---|---|---|---|
| MER34C:ERV1:LTR | 7675101969.21189 | 1944444444.44445 | 5e+09 |
| MER34C_:ERV1:LTR | 23025305907.6356 | 17777777777.7778 | 3.25e+10 |
| MER34C_v:ERV1:LTR | 0 | 0 | 0 |
| MER34D:ERV1:LTR | 6578658830.75305 | 7777777777.77778 | 8.125e+09 |
| MER39:ERV1:LTR | 179816674707.25 | 9.5e+10 | 1.31875e+11 |
| MER39B:ERV1:LTR | 75654576553.6599 | 47777777777.7778 | 7.125e+10 |
| MER3:hAT-Charlie:DNA | 1149072409104.86 | 614166666666.667 | 1.15e+12 |
| MER4-int:ERV1:LTR | 570150431998.596 | 288055555555.555 | 4.48125e+11 |
| MER41-int:ERV1:LTR | 218192184553.309 | 127777777777.778 | 3.2375e+11 |
| MER41A:ERV1:LTR | 224770843384.062 | 83888888888.8888 | 1.46875e+11 |
| MER41B:ERV1:LTR | 364019121968.334 | 159444444444.444 | 2.54375e+11 |
| MER41C:ERV1:LTR | 30700407876.8475 | 1e+10 | 2.5e+10 |
| MER41D:ERV1:LTR | 50436384369.1066 | 16111111111.1111 | 2.5625e+10 |
| MER41E:ERV1:LTR | 23025305907.6356 | 8333333333.33333 | 1.875e+10 |
| MER41G:ERV1:LTR | 30700407876.8475 | 20833333333.3333 | 1.625e+10 |
| MER44A:TcMar-Tigger:DNA | 201745537476.427 | 123333333333.333 | 2.35e+11 |
| MER44B:TcMar-Tigger:DNA | 160080698214.991 | 84722222222.2222 | 1.6375e+11 |
| MER44C:TcMar-Tigger:DNA | 60304372615.2361 | 46111111111.1112 | 6.25e+10 |
| MER44D:TcMar-Tigger:DNA | 35086180430.6828 | 19444444444.4445 | 4.125e+10 |
| MER45A:hAT-Tip100:DNA | 142537607999.649 | 75833333333.3333 | 1.15625e+11 |
| MER45B:hAT-Tip100:DNA | 35086180430.6828 | 15555555555.5556 | 3.8125e+10 |
| MER45C:hAT-Tip100:DNA | 46050611815.2712 | 2e+10 | 6e+10 |
| MER45R:hAT-Tip100:DNA | 9867988246.12957 | 8611111111.11112 | 6.25e+09 |
| MER46C:TcMar-Tigger:DNA | 334415157229.947 | 199722222222.222 | 3.5e+11 |
| MER47A:TcMar-Tigger:DNA | 249989035568.616 | 150277777777.778 | 3.19375e+11 |
| MER47B:TcMar-Tigger:DNA | 47147054953.7301 | 28888888888.8888 | 6.3125e+10 |
| MER47C:TcMar-Tigger:DNA | 18639533353.8003 | 17777777777.7778 | 2.375e+10 |
| MER48:ERV1:LTR | 23025305907.6356 | 2777777777.77778 | 8.125e+09 |
| MER49:ERV1:LTR | 174334459014.956 | 85555555555.5555 | 1.7625e+11 |
| MER4A1:ERV1:LTR | 168852243322.661 | 91666666666.6667 | 1.2125e+11 |
| MER4A1_:ERV1:LTR | 63593702030.6126 | 39166666666.6667 | 4.9375e+10 |
| MER4A1_v:ERV1:LTR | 0 | 0 | 0 |
| MER4A:ERV1:LTR | 148019823691.943 | 85833333333.3333 | 9.4375e+10 |
| MER4B-int:ERV1:LTR | 93197666769.0013 | 65277777777.7778 | 1.1e+11 |
| MER4B:ERV1:LTR | 141441164861.19 | 67777777777.7778 | 1.30625e+11 |
| MER4C:ERV1:LTR | 81136792245.954 | 4e+10 | 6.5625e+10 |
| MER4CL34:ERV1:LTR | 73461690276.7424 | 33888888888.8888 | 5.6875e+10 |
| MER4D0:ERV1:LTR | 50436384369.1066 | 26666666666.6667 | 1.9375e+10 |
| MER4D1:ERV1:LTR | 184202447261.085 | 73888888888.8888 | 1.38125e+11 |
| MER4D:ERV1:LTR | 96486996184.3778 | 46944444444.4445 | 8.875e+10 |
| MER4E1:ERV1:LTR | 112933643261.261 | 48055555555.5555 | 7.6875e+10 |
| MER4E:ERV1:LTR | 62497258892.154 | 38611111111.1112 | 6.4375e+10 |
| MER50-int:ERV1:LTR | 33989737292.2242 | 14444444444.4444 | 2.9375e+10 |
| MER50:ERV1:LTR | 269725012060.874 | 85833333333.3333 | 1.55625e+11 |
| MER50B:ERV1:LTR | 25218192184.5533 | 8333333333.33333 | 2.25e+10 |
| MER50C:ERV1:LTR | 18639533353.8003 | 6388888888.88888 | 1.6875e+10 |
| MER51-int:ERV1:LTR | 124994517784.308 | 56944444444.4445 | 8.1875e+10 |
| MER51A:ERV1:LTR | 89908337353.6248 | 48333333333.3333 | 7.25e+10 |
| MER51B:ERV1:LTR | 77847462830.5775 | 3.25e+10 | 5.625e+10 |
| MER51C:ERV1:LTR | 25218192184.5533 | 6666666666.66667 | 1.8125e+10 |
| MER51D:ERV1:LTR | 12060874523.0473 | 1388888888.88889 | 5e+09 |
| MER51E:ERV1:LTR | 42761282399.8947 | 10277777777.7778 | 1.875e+10 |
| MER52-int:ERV1:LTR | 10964431384.5884 | 2777777777.77778 | 2.5e+09 |
| MER52A:ERV1:LTR | 258760580676.286 | 77777777777.7778 | 1.09375e+11 |
| MER52C:ERV1:LTR | 36182623569.1417 | 13333333333.3333 | 2.125e+10 |
| MER52D:ERV1:LTR | 89908337353.6248 | 41111111111.1112 | 4.75e+10 |
| MER53:hAT-Blackjack:DNA | 0 | 0 | 0 |
| MER53:hAT:DNA | 174334459014.956 | 90833333333.3333 | 1.70625e+11 |
| MER54A:ERVL:LTR | 58111486338.3185 | 15833333333.3333 | 3.1875e+10 |
| MER54B:ERVL:LTR | 8771545107.67073 | 2.5e+09 | 5e+09 |
| MER57-int:ERV1:LTR | 95390553045.9192 | 35833333333.3333 | 4.75e+10 |
| MER57A-int:ERV1:LTR | 235735274768.65 | 1.25e+11 | 2.90625e+11 |
| MER57A1:ERV1:LTR | 65786588307.5305 | 28888888888.8888 | 5.0625e+10 |
| MER57B1:ERV1:LTR | 52629270646.0242 | 18611111111.1112 | 4.25e+10 |

| | | | |
|---|---|---|---|
| MER57B2:ERV1:LTR | 37279066707.6007 | 19166666666.6667 | 3.9375e+10 |
| MER57C1:ERV1:LTR | 3289329415.37652 | 3611111111.11112 | 3.75e+09 |
| MER57C2:ERV1:LTR | 9867988246.12957 | 2.5e+09 | 6.875e+09 |
| MER57D:ERV1:LTR | 20832419630.718 | 7.5e+09 | 1.9375e+10 |
| MER57E1:ERV1:LTR | 71268803999.8245 | 23888888888.8888 | 5e+10 |
| MER57E2:ERV1:LTR | 0 | 277777777.777778 | 1.25e+09 |
| MER57E3:ERV1:LTR | 3289329415.37652 | 3888888888.88888 | 7.5e+09 |
| MER57F:ERV1:LTR | 32893294153.7652 | 14166666666.6667 | 2.3125e+10 |
| MER58A:hAT-Charlie:DNA | 997763255997.543 | 628611111111.112 | 1e+12 |
| MER58B:hAT-Charlie:DNA | 729134687075.129 | 399722222222.222 | 7.25e+11 |
| MER58C:hAT-Charlie:DNA | 267532125783.957 | 185277777777.778 | 2.9125e+11 |
| MER58D:hAT-Charlie:DNA | 87715451076.7073 | 4.5e+10 | 1.0875e+11 |
| MER5A1:hAT-Charlie:DNA | 1060260514889.7 | 6.45e+11 | 8.86875e+11 |
| MER5A:hAT-Charlie:DNA | 2237840445594.49 | 1145555555555.55 | 2.093125e+12 |
| MER5B:hAT-Charlie:DNA | 1429761852550.33 | 805555555555.555 | 1.469375e+12 |
| MER5C1:hAT-Charlie:DNA | 23025305907.6356 | 18611111111.1112 | 2.375e+10 |
| MER5C:hAT-Charlie:DNA | 35086180430.6828 | 30833333333.3333 | 3.9375e+10 |
| MER61-int:ERV1:LTR | 305907635630.016 | 136944444444.445 | 2.98125e+11 |
| MER61A:ERV1:LTR | 61400815753.695 | 18611111111.1112 | 4.375e+10 |
| MER61B:ERV1:LTR | 23025305907.6356 | 16666666666.6667 | 2e+10 |
| MER61C:ERV1:LTR | 10964431384.5884 | 3888888888.88888 | 1e+10 |
| MER61D:ERV1:LTR | 26314635323.0122 | 17222222222.2222 | 5.75e+10 |
| MER61E:ERV1:LTR | 20832419630.718 | 9166666666.66667 | 2.3125e+10 |
| MER61F:ERV1:LTR | 3289329415.37652 | 4166666666.66667 | 6.875e+09 |
| MER63A:hAT-Blackjack:DNA | 109644313845.884 | 73333333333.3333 | 1.725e+11 |
| MER63A:hAT-Tip100:DNA | 0 | 0 | 0 |
| MER63B:hAT-Blackjack:DNA | 129380290338.143 | 84166666666.6667 | 1.4e+11 |
| MER63B:hAT-Tip100:DNA | 0 | 0 | 0 |
| MER63C:hAT-Blackjack:DNA | 36182623569.1417 | 17222222222.2222 | 4.0625e+10 |
| MER63C:hAT-Tip100:DNA | 0 | 0 | 0 |
| MER63D:hAT-Blackjack:DNA | 87715451076.7073 | 41944444444.4445 | 7.75e+10 |
| MER63D:hAT-Tip100:DNA | 0 | 0 | 0 |
| MER65-int:ERV1:LTR | 25218192184.5533 | 23055555555.5555 | 2.5625e+10 |
| MER65A:ERV1:LTR | 58111486338.3185 | 30555555555.5555 | 5.3125e+10 |
| MER65B:ERV1:LTR | 14253760799.9649 | 5e+09 | 1.5e+10 |
| MER65C:ERV1:LTR | 27411078461.471 | 16666666666.6667 | 2.375e+10 |
| MER65D:ERV1:LTR | 57015043199.8596 | 19722222222.2222 | 1.9375e+10 |
| MER66-int:ERV1:LTR | 29603964738.3887 | 11111111111.1111 | 1.9375e+10 |
| MER66A:ERV1:LTR | 9867988246.12957 | 6111111111.11112 | 9.375e+09 |
| MER66B:ERV1:LTR | 117319415815.096 | 64722222222.2222 | 1.175e+11 |
| MER66C:ERV1:LTR | 59207929476.7775 | 23333333333.3333 | 4.125e+10 |
| MER66D:ERV1:LTR | 1096443138.45884 | 1111111111.11111 | 5e+09 |
| MER67A:ERV1:LTR | 24121749046.0945 | 11388888888.8889 | 2.4375e+10 |
| MER67B:ERV1:LTR | 58111486338.3185 | 35277777777.7778 | 4.4375e+10 |
| MER67C:ERV1:LTR | 129380290338.143 | 70555555555.5555 | 1.31875e+11 |
| MER67D:ERV1:LTR | 28507521599.9298 | 21944444444.4445 | 2.75e+10 |
| MER68-int:ERVL:LTR | 81136792245.954 | 3e+10 | 6.9375e+10 |
| MER68:ERVL:LTR | 119512302092.014 | 61944444444.4445 | 1.14375e+11 |
| MER68B:ERVL:LTR | 48243498092.1891 | 23888888888.8888 | 4.125e+10 |
| MER68C:ERVL:LTR | 2192886276.91768 | 1388888888.88889 | 0 |
| MER6:TcMar-Tigger:DNA | 71268803999.8245 | 39444444444.4445 | 7.75e+10 |
| MER6A:TcMar-Tigger:DNA | 99776325599.7543 | 31944444444.4445 | 6.5625e+10 |
| MER6B:TcMar-Tigger:DNA | 60304372615.2361 | 46666666666.6667 | 6.9375e+10 |
| MER6C:TcMar-Tigger:DNA | 5482215692.2942 | 3888888888.88888 | 7.5e+09 |
| MER70-int:ERVL:LTR | 53725713784.4831 | 2.75e+10 | 3.625e+10 |
| MER70A:ERVL:LTR | 25218192184.5533 | 10277777777.7778 | 2.25e+10 |
| MER70B:ERVL:LTR | 14253760799.9649 | 6666666666.66667 | 8.125e+09 |
| MER70C:ERVL:LTR | 12060874523.0473 | 4166666666.66667 | 9.375e+09 |
| MER72:ERV1:LTR | 23025305907.6356 | 7777777777.77778 | 2.0625e+10 |
| MER72B:ERV1:LTR | 3289329415.37652 | 1666666666.66667 | 8.125e+09 |
| MER73:ERVL:LTR | 9867988246.12957 | 5e+09 | 4.375e+09 |
| MER74A:ERVL:LTR | 101969211876.672 | 36944444444.4445 | 7.25e+10 |
| MER74B:ERVL:LTR | 39471952984.5182 | 20555555555.5555 | 4.3125e+10 |
| MER74C:ERVL:LTR | 3289329415.37652 | 3055555555.55555 | 7.5e+09 |

| | | | |
|---|---|---|---|
| MER75:PiggyBac:DNA | 52629270646.0242 | 28055555555.5555 | 3.625e+10 |
| MER75A:PiggyBac:DNA | 1096443138.45884 | 0 | 4.375e+09 |
| MER75B:PiggyBac:DNA | 0 | 1666666666.66667 | 1.875e+09 |
| MER76-int:ERVL:LTR | 6578658830.75305 | 1111111111.11111 | 2.5e+09 |
| MER76:ERVL:LTR | 30700407876.8475 | 13611111111.1111 | 1.5625e+10 |
| MER77:ERVL:LTR | 94294109907.4603 | 38333333333.3333 | 9.125e+10 |
| MER77B:ERVL:LTR | 71268803999.8245 | 29444444444.4445 | 4.25e+10 |
| MER81:hAT-Blackjack:DNA | 210517082584.097 | 96111111111.1112 | 2.0625e+11 |
| MER83:ERV1:LTR | 44954168676.8126 | 28055555555.5555 | 6.3125e+10 |
| MER83A-int:ERV1:LTR | 8771545107.67073 | 2.5e+09 | 5e+09 |
| MER83B-int:ERV1:LTR | 8771545107.67073 | 2222222222.22222 | 8.125e+09 |
| MER83B:ERV1:LTR | 3289329415.37652 | 1666666666.66667 | 4.375e+09 |
| MER83C:ERV1:LTR | 4385772553.83536 | 555555555.555555 | 4.375e+09 |
| MER84-int:ERV1:LTR | 2192886276.91768 | 2222222222.22222 | 0 |
| MER84:ERV1:LTR | 26314635323.0122 | 16666666666.6667 | 1.25e+10 |
| MER85:PiggyBac:DNA | 157887811938.073 | 79722222222.2222 | 1.625e+11 |
| MER87:ERV1:LTR | 1096443138.45884 | 2222222222.22222 | 0 |
| MER87B:ERV1:LTR | 8771545107.67073 | 1111111111.11111 | 1.875e+09 |
| MER88:ERVL:LTR | 0 | 555555555.555555 | 1.875e+09 |
| MER89-int:ERV1:LTR | 20832419630.718 | 13333333333.3333 | 1.4375e+10 |
| MER89:ERV1:LTR | 64690145169.0715 | 2.5e+10 | 4.5e+10 |
| MER8:TcMar-Tigger:DNA | 199552651199.509 | 130833333333.333 | 2.35e+11 |
| MER90:ERV1:LTR | 25218192184.5533 | 11666666666.6667 | 2.4375e+10 |
| MER90a:ERV1:LTR | 98679882461.2957 | 48611111111.1112 | 7.6875e+10 |
| MER91A:hAT-Tip100:DNA | 232445945353.274 | 98055555555.5555 | 1.675e+11 |
| MER91B:hAT-Tip100:DNA | 74558133415.201 | 56111111111.1112 | 9.3125e+10 |
| MER91C:hAT-Tip100:DNA | 103065655015.131 | 39444444444.4445 | 7.75e+10 |
| MER92-int:ERV1:LTR | 18639533353.8003 | 4166666666.66667 | 9.375e+09 |
| MER92A:ERV1:LTR | 8771545107.67073 | 1111111111.11111 | 3.125e+09 |
| MER92B:ERV1:LTR | 109644313845.884 | 31388888888.8888 | 7.5e+10 |
| MER92C:ERV1:LTR | 12060874523.0473 | 2222222222.22222 | 3.125e+09 |
| MER92D:ERV1:LTR | 3289329415.37652 | 555555555.555555 | 3.125e+09 |
| MER94:hAT-Blackjack:DNA | 215999298276.391 | 118888888888.889 | 2.1375e+11 |
| MER94B:hAT-Blackjack:DNA | 12060874523.0473 | 5833333333.33333 | 1.25e+10 |
| MER95:ERV1:LTR | 5482215692.2942 | 1944444444.44445 | 5.625e+09 |
| MER96:hAT-Tip100:DNA | 110740756984.343 | 58611111111.1112 | 1.2875e+11 |
| MER96:hAT:DNA | 0 | 0 | 0 |
| MER96B:hAT-Tip100:DNA | 82233235384.4129 | 30833333333.3333 | 8.25e+10 |
| MER96B:hAT:DNA | 1096443138.45884 | 4444444444.44445 | 5e+09 |
| MER97a:hAT-Tip100:DNA | 5482215692.2942 | 5e+09 | 1.3125e+10 |
| MER97b:hAT-Tip100:DNA | 3289329415.37652 | 277777777.777778 | 3.125e+09 |
| MER97c:hAT-Tip100:DNA | 9867988246.12957 | 2.5e+09 | 7.5e+09 |
| MER97d:hAT-Tip100:DNA | 8771545107.67073 | 1388888888.88889 | 3.125e+09 |
| MER99:hAT:DNA | 21928862769.1768 | 8611111111.11112 | 1.375e+10 |
| MER9B:ERVK:LTR | 2192886276.91768 | 1111111111.11111 | 1.25e+09 |
| MER9a1:ERVK:LTR | 39471952984.5182 | 21666666666.6667 | 3.1875e+10 |
| MER9a2:ERVK:LTR | 27411078461.471 | 16666666666.6667 | 3.625e+10 |
| MER9a3:ERVK:LTR | 99776325599.7543 | 43888888888.8888 | 1.10625e+11 |
| MERX:TcMar-Tigger:DNA | 12060874523.0473 | 14166666666.6667 | 3.625e+10 |
| MIR1_Amn:MIR:SINE | 1048199640366.65 | 534722222222.222 | 8.20625e+11 |
| MIR3:MIR:SINE | 6363755975615.1 | 3185555555555.55 | 5.629375e+12 |
| MIR:MIR:SINE | 13594798473751.2 | 7287777777777.78 | 1.295875e+13 |
| MIRb:MIR:SINE | 16959782465681.3 | 9018611111111.12 | 1.629e+13 |
| MIRc:MIR:SINE | 7451427568966.26 | 3776388888888.88 | 6.5225e+12 |
| MLT-int:ERVL-MaLR:LTR | 23025305907.6356 | 13333333333.3333 | 2.25e+10 |
| MLT1-int:ERVL-MaLR:LTR | 8771545107.67073 | 8333333333.33333 | 1e+10 |
| MLT1A-int:ERVL-MaLR:LTR | 42761282399.8947 | 19444444444.4445 | 3.625e+10 |
| MLT1A0-int:ERVL-MaLR:LTR | 40568396122.9771 | 26388888888.8888 | 4.375e+10 |
| MLT1A0:ERVL-MaLR:LTR | 854129204859.437 | 508333333333.333 | 7.3625e+11 |
| MLT1A1-int:ERVL-MaLR:LTR | 69075917722.907 | 32222222222.2222 | 6.125e+10 |
| MLT1A1:ERVL-MaLR:LTR | 260953466953.204 | 119722222222.222 | 2.3e+11 |
| MLT1A:ERVL-MaLR:LTR | 414455506337.44 | 178055555555.555 | 3.475e+11 |
| MLT1B-int:ERVL-MaLR:LTR | 28507521599.9298 | 10277777777.7778 | 2.125e+10 |
| MLT1B:ERVL-MaLR:LTR | 843164773474.848 | 393055555555.555 | 6.9e+11 |

| | | | |
|---|---|---|---|
| MLT1C-int:ERVL-MaLR:LTR | 2192886276.91768 | 4444444444.44445 | 8.125e+09 |
| MLT1C2-int:ERVL-MaLR:LTR | 0 | 0 | 0 |
| MLT1C2:ERVL-MaLR:LTR | 0 | 0 | 0 |
| MLT1C:ERVL-MaLR:LTR | 1332178413227.49 | 535555555555.555 | 1.03625e+12 |
| MLT1D-int:ERVL-MaLR:LTR | 23025305907.6356 | 18888888888.8888 | 3.0625e+10 |
| MLT1D:ERVL-MaLR:LTR | 1348625060304.37 | 683888888888.888 | 1.115625e+12 |
| MLT1E-int:ERVL-MaLR:LTR | 0 | 0 | 0 |
| MLT1E1-int:ERVL-MaLR:LTR | 0 | 0 | 0 |
| MLT1E1:ERVL-MaLR:LTR | 50436384369.1066 | 25277777777.7778 | 3.6875e+10 |
| MLT1E1A-int:ERVL-MaLR:LTR | 12060874523.0473 | 8888888888.88888 | 4.375e+09 |
| MLT1E1A:ERVL-MaLR:LTR | 205034866891.803 | 90277777777.7778 | 1.5125e+11 |
| MLT1E2-int:ERVL-MaLR:LTR | 78943905969.0364 | 3.25e+10 | 5.375e+10 |
| MLT1E2:ERVL-MaLR:LTR | 314679180737.687 | 138888888888.889 | 2.4375e+11 |
| MLT1E3-int:ERVL-MaLR:LTR | 1096443138.45884 | 277777777.777778 | 6.25e+08 |
| MLT1E3:ERVL-MaLR:LTR | 131573176615.061 | 61111111111.1112 | 1.125e+11 |
| MLT1E:ERVL-MaLR:LTR | 52629270646.0242 | 12777777777.7778 | 2.25e+10 |
| MLT1F-int:ERVL-MaLR:LTR | 61400815753.695 | 24166666666.6667 | 3.4375e+10 |
| MLT1F1-int:ERVL-MaLR:LTR | 39471952984.5182 | 15555555555.5556 | 2.8125e+10 |
| MLT1F1:ERVL-MaLR:LTR | 162273584491.908 | 81666666666.6667 | 1.30625e+11 |
| MLT1F2-int:ERVL-MaLR:LTR | 18639533353.8003 | 7.5e+09 | 2.4375e+10 |
| MLT1F2:ERVL-MaLR:LTR | 268628568922.416 | 131388888888.889 | 2.39375e+11 |
| MLT1F:ERVL-MaLR:LTR | 253278364983.992 | 116666666666.667 | 2.40625e+11 |
| MLT1G-int:ERVL-MaLR:LTR | 13157317661.5061 | 4166666666.66667 | 6.875e+09 |
| MLT1G1-int:ERVL-MaLR:LTR | 8771545107.67073 | 4722222222.22222 | 5e+09 |
| MLT1G1:ERVL-MaLR:LTR | 179816674707.25 | 59722222222.2222 | 1.18125e+11 |
| MLT1G3-int:ERVL-MaLR:LTR | 14253760799.9649 | 833333333.333333 | 1e+10 |
| MLT1G3:ERVL-MaLR:LTR | 104162098153.59 | 45277777777.7778 | 9.5625e+10 |
| MLT1G:ERVL-MaLR:LTR | 154598482522.696 | 55277777777.7778 | 1.81875e+11 |
| MLT1H-int:ERVL-MaLR:LTR | 21928862769.1768 | 11944444444.4444 | 2.125e+10 |
| MLT1H1-int:ERVL-MaLR:LTR | 4385772553.83536 | 2.5e+09 | 2.5e+09 |
| MLT1H1:ERVL-MaLR:LTR | 185298890399.544 | 84722222222.2222 | 1.25625e+11 |
| MLT1H2-int:ERVL-MaLR:LTR | 27411078461.471 | 11944444444.4444 | 1.6875e+10 |
| MLT1H2:ERVL-MaLR:LTR | 162273584491.908 | 6.75e+10 | 1.175e+11 |
| MLT1H:ERVL-MaLR:LTR | 400201745537.477 | 156111111111.111 | 2.71875e+11 |
| MLT1I-int:ERVL-MaLR:LTR | 50436384369.1066 | 17777777777.7778 | 2.9375e+10 |
| MLT1I:ERVL-MaLR:LTR | 654576553659.927 | 2.425e+11 | 5.48125e+11 |
| MLT1J-int:ERVL-MaLR:LTR | 49339941230.6477 | 24444444444.4445 | 4.5625e+10 |
| MLT1J1-int:ERVL-MaLR:LTR | 35086180430.6828 | 1.25e+10 | 2.1875e+10 |
| MLT1J1:ERVL-MaLR:LTR | 122801631507.39 | 55555555555.5555 | 1.09375e+11 |
| MLT1J2-int:ERVL-MaLR:LTR | 15350203938.4237 | 6666666666.66667 | 1.0625e+10 |
| MLT1J2:ERVL-MaLR:LTR | 230253059076.356 | 1.025e+11 | 2.06875e+11 |
| MLT1J:ERVL-MaLR:LTR | 525196263321.785 | 243055555555.555 | 4.3875e+11 |
| MLT1K-int:ERVL-MaLR:LTR | 4385772553.83536 | 1111111111.11111 | 6.25e+08 |
| MLT1K:ERVL-MaLR:LTR | 623876145783.08 | 256111111111.112 | 5.3375e+11 |
| MLT1L-int:ERVL-MaLR:LTR | 0 | 277777777.777778 | 0 |
| MLT1L:ERVL-MaLR:LTR | 444059471075.831 | 198888888888.888 | 3.83125e+11 |
| MLT1M-int:ERVL-MaLR:LTR | 0 | 0 | 0 |
| MLT1M:ERVL-MaLR:LTR | 70172360861.3659 | 39444444444.4445 | 6.625e+10 |
| MLT1N2-int:ERVL-MaLR:LTR | 0 | 555555555.555555 | 0 |
| MLT1N2:ERVL-MaLR:LTR | 139248278584.273 | 70833333333.3333 | 1.5e+11 |
| MLT1O-int:ERVL-MaLR:LTR | 0 | 0 | 0 |
| MLT1O:ERVL-MaLR:LTR | 251085478707.074 | 106388888888.889 | 1.91875e+11 |
| MLT2A1:ERVL:LTR | 137055392307.355 | 64722222222.2222 | 1.23125e+11 |
| MLT2A2:ERVL:LTR | 223674400245.603 | 101388888888.889 | 1.375e+11 |
| MLT2B1:ERVL:LTR | 222577957107.144 | 64722222222.2222 | 1.475e+11 |
| MLT2B2:ERVL:LTR | 70172360861.3659 | 26666666666.6667 | 5.0625e+10 |
| MLT2B3:ERVL:LTR | 210517082584.097 | 71388888888.8888 | 1.6e+11 |
| MLT2B4:ERVL:LTR | 248892592430.157 | 112222222222.222 | 1.6625e+11 |
| MLT2B5:ERVL:LTR | 88811894215.1659 | 27222222222.2222 | 8.625e+10 |
| MLT2C1:ERVL:LTR | 65786588307.5305 | 30833333333.3333 | 4.9375e+10 |
| MLT2C2:ERVL:LTR | 37279066707.6007 | 11944444444.4444 | 3.375e+10 |
| MLT2D:ERVL:LTR | 134862506030.437 | 7.5e+10 | 1.2e+11 |
| MLT2E:ERVL:LTR | 5482215692.2942 | 3055555555.55555 | 1.125e+10 |
| MLT2F:ERVL:LTR | 115126529538.178 | 49722222222.2222 | 6.625e+10 |

| | | | |
|---|---|---|---|
| MSR1:Satellite:Satellite | 233542388491.733 | 76666666666.6667 | 1.60625e+11 |
| MST-int:ERVL-MaLR:LTR | 15350203938.4237 | 8888888888.88888 | 2.5e+10 |
| MSTA-int:ERVL-MaLR:LTR | 610718828121.573 | 250277777777.778 | 4.89375e+11 |
| MSTA1-int:ERVL-MaLR:LTR | 0 | 277777777.777778 | 0 |
| MSTA1:ERVL-MaLR:LTR | 23025305907.6356 | 8333333333.33333 | 2e+10 |
| MSTA:ERVL-MaLR:LTR | 1061356958028.16 | 497777777777.778 | 8.79375e+11 |
| MSTB-int:ERVL-MaLR:LTR | 51532827507.5656 | 34444444444.4445 | 5.8125e+10 |
| MSTB1-int:ERVL-MaLR:LTR | 23025305907.6356 | 9444444444.44445 | 1.3125e+10 |
| MSTB1:ERVL-MaLR:LTR | 190781106091.838 | 118888888888.889 | 1.8e+11 |
| MSTB2-int:ERVL-MaLR:LTR | 2192886276.91768 | 1111111111.11111 | 3.75e+09 |
| MSTB2:ERVL-MaLR:LTR | 86619007938.2484 | 38888888888.8888 | 7.9375e+10 |
| MSTB:ERVL-MaLR:LTR | 359633349414.5 | 203888888888.888 | 3.65625e+11 |
| MSTC-int:ERVL-MaLR:LTR | 35086180430.6828 | 9166666666.66667 | 1.5e+10 |
| MSTC:ERVL-MaLR:LTR | 255471251260.91 | 86388888888.8888 | 2.1125e+11 |
| MSTD-int:ERVL-MaLR:LTR | 91004780492.0838 | 40833333333.3333 | 7.3125e+10 |
| MSTD:ERVL-MaLR:LTR | 354151133722.205 | 138888888888.889 | 3.0125e+11 |
| MamGyp-int:Gypsy:LTR | 18639533353.8003 | 4722222222.22222 | 1e+10 |
| MamGypLTR1a:Gypsy:LTR | 25218192184.5533 | 14722222222.2222 | 2.5e+10 |
| MamGypLTR1b:Gypsy:LTR | 28507521599.9298 | 12222222222.2222 | 1.8125e+10 |
| MamGypLTR1c:Gypsy:LTR | 103065655015.131 | 28333333333.3333 | 6.75e+10 |
| MamGypLTR1d:Gypsy:LTR | 21928862769.1768 | 6666666666.66667 | 6.25e+09 |
| MamGypLTR2b:Gypsy:LTR | 15350203938.4237 | 8333333333.33333 | 8.75e+09 |
| MamGypLTR2c:Gypsy:LTR | 8771545107.67073 | 14722222222.2222 | 7.5e+09 |
| MamGypLTR3:Gypsy:LTR | 8771545107.67073 | 3333333333.33333 | 6.875e+09 |
| MamGypLTR3a:Gypsy:LTR | 33989737292.2242 | 19722222222.2222 | 2.8125e+10 |
| MamGypLTR4:Gypsy:LTR | 0 | 0 | 0 |
| MamGypsy2-I:Gypsy:LTR | 92101223630.5424 | 4.5e+10 | 6.4375e+10 |
| MamGypsy2-LTR:Gypsy:LTR | 9867988246.12957 | 5555555555.55555 | 7.5e+09 |
| MamRTE1:RTE-Bo.B:LINE | 392526643568.265 | 214444444444.445 | 3.88125e+11 |
| MamRTE1:RTE-BovB:LINE | 0 | 0 | 0 |
| MamRTE2:RTE-BovB:LINE | 0 | 0 | 0 |
| MamRep1151:LTR:LTR | 0 | 0 | 0 |
| MamRep1151:LTR:LTR? | 20832419630.718 | 16666666666.6667 | 2.8125e+10 |
| MamRep1161:TcMar-Tigger:DNA | 0 | 0 | 1.25e+09 |
| MamRep137:TcMar-Tigger:DNA | 263146353230.122 | 124722222222.222 | 2.76875e+11 |
| MamRep137:TcMar:DNA | 13157317661.5061 | 6388888888.88888 | 1.75e+10 |
| MamRep1527:LTR:LTR | 51532827507.5656 | 12777777777.7778 | 4.5e+10 |
| MamRep1879:hAT-Tip100:DNA | 64690145169.0715 | 41666666666.6667 | 7.0625e+10 |
| MamRep1894:hAT-Tip100:DNA | 0 | 0 | 0 |
| MamRep1894:hAT:DNA | 28507521599.9298 | 9722222222.22222 | 3e+10 |
| MamRep38:hAT-Tip100:DNA | 0 | 0 | 0 |
| MamRep38:hAT:DNA | 184202447261.085 | 81111111111.1112 | 1.2625e+11 |
| MamRep4096:hAT-Tip100:DNA | 38375509846.0593 | 13333333333.3333 | 2.125e+10 |
| MamRep4096:hAT:DNA | 9867988246.12957 | 2.5e+09 | 5.625e+09 |
| MamRep434:TcMar-Tigger:DNA | 76751019692.1189 | 50277777777.7778 | 7.875e+10 |
| MamRep488:hAT-Tip100:DNA | 8771545107.67073 | 4166666666.66667 | 1e+10 |
| MamRep564:MamRep564:Unknown | 25218192184.5533 | 9166666666.66667 | 2e+10 |
| MamRep605:LTR:LTR | 0 | 0 | 0 |
| MamRep605:LTR:LTR? | 241217490460.945 | 115555555555.555 | 1.9e+11 |
| MamRep605b:LTR:LTR | 0 | 0 | 0 |
| MamRep605b:LTR:LTR? | 0 | 0 | 0 |
| MamTip1:hAT-Tip100:DNA | 50436384369.1066 | 35555555555.5555 | 5.875e+10 |
| MamTip1b:hAT-Tip100:DNA | 0 | 0 | 0 |
| MamTip2:hAT-Tip100:DNA | 322354282706.899 | 134166666666.667 | 2.35625e+11 |
| MamTip2b:hAT-Tip100:DNA | 0 | 0 | 0 |
| MamTip3:hAT-Tip100:DNA | 40568396122.9771 | 18055555555.5555 | 5.375e+10 |
| MamTip3B:hAT-Tip100:DNA | 0 | 0 | 0 |
| Mam_R4:Dong-R4:LINE | 25218192184.5533 | 9722222222.22222 | 2.125e+10 |
| Merlin1_HS:Merlin:DNA | 81136792245.954 | 38333333333.3333 | 6.5e+10 |
| ORSL-2a:hAT-Tip100:DNA | 19735976492.2591 | 6388888888.88888 | 1e+10 |
| ORSL-2b:hAT-Tip100:DNA | 25218192184.5533 | 19166666666.6667 | 2.5e+10 |
| ORSL:hAT-Tip100:DNA | 30700407876.8475 | 23333333333.3333 | 4e+10 |
| OldhAT1:hAT-Ac:DNA | 305907635630.016 | 125833333333.333 | 2.375e+11 |
| PABL_A-int:ERV1:LTR | 52629270646.0242 | 20555555555.5555 | 3.5625e+10 |

| | | | |
|---|---|---|---|
| PABL_A:ERV1:LTR | 25218192184.5533 | 18611111111.1112 | 4.375e+10 |
| PABL_B-int:ERV1:LTR | 78943905969.0364 | 48333333333.3333 | 1.3625e+11 |
| PABL_B:ERV1:LTR | 117319415815.096 | 39444444444.4445 | 7.9375e+10 |
| PRIMA4-int:ERV1:LTR | 106354984430.507 | 48611111111.1112 | 9.9375e+10 |
| PRIMA41-int:ERV1:LTR | 67979474584.448 | 29166666666.6667 | 7.125e+10 |
| PRIMA4_LTR:ERV1:LTR | 29603964738.3887 | 12222222222.2222 | 2.0625e+10 |
| PRIMAX-int:ERV1:LTR | 5482215692.2942 | 1388888888.88889 | 4.375e+09 |
| Penelope1_Vert:Penelope:LINE | 70172360861.3659 | 23333333333.3333 | 4.6875e+10 |
| Plat_L3:CR1:LINE | 160080698214.991 | 86111111111.1112 | 1.425e+11 |
| PrimLTR79:ERV1:LTR | 2192886276.91768 | 1944444444.44445 | 1.875e+09 |
| REP522:REP522:Unknown | 0 | 0 | 0 |
| REP522:telo:Satellite | 31796851015.3064 | 7222222222.22222 | 8.75e+09 |
| Ricksha:MULE-MuDR:DNA | 8771545107.67073 | 4166666666.66667 | 8.125e+09 |
| Ricksha_0:MULE-MuDR:DNA | 25218192184.5533 | 18333333333.3333 | 3.0625e+10 |
| Ricksha_a:MULE-MuDR:DNA | 3289329415.37652 | 2222222222.22222 | 4.375e+09 |
| Ricksha_b:MULE-MuDR:DNA | 31796851015.3064 | 11944444444.4444 | 2.125e+10 |
| Ricksha_c:MULE-MuDR:DNA | 54822156922.9421 | 28333333333.3333 | 5.3125e+10 |
| SAR:Satellite:Satellite | 24121749046.0945 | 3611111111.11112 | 6.25e+08 |
| SATR1:Satellite:Satellite | 30700407876.8475 | 15833333333.3333 | 2.625e+10 |
| SATR2:Satellite:Satellite | 317968510153.064 | 132222222222.222 | 5.38125e+11 |
| SST1:centr:Satellite | 19735976492.2591 | 7.5e+09 | 1.9375e+10 |
| SVA_A:SVA:Retroposon | 395815972983.642 | 130277777777.778 | 2.15e+11 |
| SVA_B:SVA:Retroposon | 595368624183.149 | 191388888888.888 | 3.58125e+11 |
| SVA_C:SVA:Retroposon | 212709968861.015 | 79444444444.4445 | 1.05625e+11 |
| SVA_D:SVA:Retroposon | 1395772115258.1 | 397777777777.778 | 5.86875e+11 |
| SVA_E:SVA:Retroposon | 171045129599.579 | 65833333333.3333 | 9.0625e+10 |
| SVA_F:SVA:Retroposon | 215999298276.391 | 63888888888.8888 | 1.09375e+11 |
| TAR1:subtelo:Satellite | 0 | 0 | 0 |
| TAR1:telo:Satellite | 9867988246.12957 | 4166666666.66667 | 3.75e+09 |
| THE1-int:ERVL-MaLR:LTR | 29603964738.3887 | 8333333333.33333 | 2.125e+10 |
| THE1A-int:ERVL-MaLR:LTR | 491206526029.56 | 211944444444.445 | 3.625e+11 |
| THE1A:ERVL-MaLR:LTR | 230253059076.356 | 94166666666.6667 | 2.01875e+11 |
| THE1B-int:ERVL-MaLR:LTR | 1187447918950.92 | 461111111111.112 | 8.09375e+11 |
| THE1B:ERVL-MaLR:LTR | 1270777597473.79 | 505277777777.778 | 8.71875e+11 |
| THE1C-int:ERVL-MaLR:LTR | 223674400245.603 | 107222222222.222 | 1.40625e+11 |
| THE1C:ERVL-MaLR:LTR | 508749616244.901 | 181666666666.667 | 4.0375e+11 |
| THE1D-int:ERVL-MaLR:LTR | 513135388798.738 | 230555555555.555 | 3.9125e+11 |
| THE1D:ERVL-MaLR:LTR | 653480110521.467 | 265833333333.333 | 5.10625e+11 |
| Tigger10:TcMar-Tigger:DNA | 57015043199.8596 | 37222222222.2222 | 8.125e+10 |
| Tigger11a:TcMar-Tigger:DNA | 27411078461.471 | 19722222222.2222 | 3.3125e+10 |
| Tigger12:TcMar-Tigger:DNA | 66883031445.9891 | 34722222222.2222 | 6.4375e+10 |
| Tigger12A:TcMar-Tigger:DNA | 29603964738.3887 | 2e+10 | 3.1875e+10 |
| Tigger12c:TcMar-Tigger:DNA | 50436384369.1066 | 22777777777.7778 | 6.625e+10 |
| Tigger13a:TcMar-Tigger:DNA | 230253059076.356 | 102777777777.778 | 1.74375e+11 |
| Tigger14a:TcMar-Tigger:DNA | 93197666769.0013 | 8e+10 | 6.375e+10 |
| Tigger15a:TcMar-Tigger:DNA | 419937722029.735 | 272777777777.778 | 4.95625e+11 |
| Tigger16a:TcMar-Tigger:DNA | 126090960922.767 | 75555555555.5555 | 1.575e+11 |
| Tigger16b:TcMar-Tigger:DNA | 89908337353.6248 | 61666666666.6667 | 1.11875e+11 |
| Tigger17:TcMar-Tigger:DNA | 8771545107.67073 | 4166666666.66667 | 1.5625e+10 |
| Tigger17a:TcMar-Tigger:DNA | 146923380553.484 | 114166666666.667 | 1.73125e+11 |
| Tigger17b:TcMar-Tigger:DNA | 19735976492.2591 | 7.5e+09 | 1.5e+10 |
| Tigger17c:TcMar-Tigger:DNA | 73461690276.7424 | 48055555555.5555 | 5.75e+10 |
| Tigger17d:TcMar-Tigger:DNA | 0 | 0 | 0 |
| Tigger18a:TcMar-Tigger:DNA | 142537607999.649 | 71388888888.8888 | 1.21875e+11 |
| Tigger19a:TcMar-Tigger:DNA | 260953466953.204 | 153888888888.889 | 3.41875e+11 |
| Tigger19b:TcMar-Tigger:DNA | 404587518091.311 | 230555555555.555 | 3.86875e+11 |
| Tigger1:TcMar-Tigger:DNA | 3492171395991.39 | 1.555e+12 | 3.08875e+12 |
| Tigger1a_Art:TcMar-Tigger:DNA | 14253760799.9649 | 6944444444.44445 | 1.375e+10 |
| Tigger1a_Mars:TcMar-Tigger:DNA | 0 | 1944444444.44445 | 6.25e+08 |
| Tigger20a:TcMar-Tigger:DNA | 129380290338.143 | 76111111111.1112 | 1.23125e+11 |
| Tigger21a:TcMar-Tigger:DNA | 0 | 0 | 0 |
| Tigger22N1:TcMar-Tigger:DNA | 0 | 0 | 0 |
| Tigger23a:TcMar-Tigger:DNA | 0 | 0 | 0 |
| Tigger2:TcMar-Tigger:DNA | 882636726459.364 | 455277777777.778 | 7.16875e+11 |

| | | | |
|---|---|---|---|
| Tigger2a:TcMar-Tigger:DNA | 152405596245.779 | 67777777777.7778 | 1.25625e+11 |
| Tigger2b_Pri:TcMar-Tigger:DNA | 154598482522.696 | 86944444444.4445 | 1.5e+11 |
| Tigger3:TcMar-Tigger:DNA | 116222972676.637 | 88611111111.1112 | 1.69375e+11 |
| Tigger3a:TcMar-Tigger:DNA | 544932239814.043 | 343055555555.555 | 6.09375e+11 |
| Tigger3b:TcMar-Tigger:DNA | 563571773167.844 | 296944444444.445 | 5.625e+11 |
| Tigger3c:TcMar-Tigger:DNA | 126090960922.767 | 73333333333.3333 | 1.45625e+11 |
| Tigger3d:TcMar-Tigger:DNA | 23025305907.6356 | 7222222222.22222 | 2.125e+10 |
| Tigger4:TcMar-Tigger:DNA | 185298890399.544 | 104722222222.222 | 1.79375e+11 |
| Tigger4a:TcMar-Tigger:DNA | 444059471075.831 | 276111111111.112 | 5.025e+11 |
| Tigger4b:TcMar-Tigger:DNA | 392526643568.265 | 238055555555.555 | 3.925e+11 |
| Tigger5:TcMar-Tigger:DNA | 67979474584.448 | 2.75e+10 | 6.1875e+10 |
| Tigger5b:TcMar-Tigger:DNA | 178720231568.791 | 102222222222.222 | 1.86875e+11 |
| Tigger6:TcMar-Tigger:DNA | 0 | 0 | 0 |
| Tigger6a:TcMar-Tigger:DNA | 35086180430.6828 | 34166666666.6667 | 4.5625e+10 |
| Tigger6b:TcMar-Tigger:DNA | 33989737292.2242 | 16944444444.4445 | 2.5625e+10 |
| Tigger7:TcMar-Tigger:DNA | 201745537476.427 | 105833333333.333 | 2.09375e+11 |
| Tigger8:TcMar-Tigger:DNA | 83329678522.8719 | 32222222222.2222 | 7.0625e+10 |
| Tigger9a:TcMar-Tigger:DNA | 38375509846.0593 | 24722222222.2222 | 4.3125e+10 |
| Tigger9b:TcMar-Tigger:DNA | 41664839261.4358 | 16666666666.6667 | 3.875e+10 |
| UCON100:DNA:DNA? | 0 | 0 | 0 |
| UCON101:UCON101:Unknown | 0 | 0 | 0 |
| UCON103:UCON103:Unknown | 0 | 0 | 0 |
| UCON105:UCON105:Unknown | 31796851015.3064 | 19166666666.6667 | 4e+10 |
| UCON105:hAT:DNA | 0 | 0 | 0 |
| UCON107:hAT-Tag1:DNA | 5482215692.2942 | 3888888888.88888 | 5e+09 |
| UCON11:TcMar-Tigger:DNA | 0 | 555555555.555555 | 0 |
| UCON12:UCON12:Unknown | 0 | 277777777.777778 | 0 |
| UCON12A:UCON12A:Unknown | 0 | 0 | 0 |
| UCON132b:hAT-Tip100:DNA | 29603964738.3887 | 15555555555.5556 | 2e+10 |
| UCON14:DNA:DNA? | 2192886276.91768 | 555555555.555555 | 0 |
| UCON15:UCON15:Unknown | 0 | 0 | 0 |
| UCON15:hAT-Ac:DNA | 0 | 0 | 0 |
| UCON16:UCON16:Unknown | 0 | 277777777.777778 | 2.5e+09 |
| UCON17:UCON17:Unknown | 0 | 0 | 0 |
| UCON18:UCON18:Unknown | 1096443138.45884 | 555555555.555555 | 1.25e+09 |
| UCON19:UCON19:Unknown | 1096443138.45884 | 0 | 0 |
| UCON1:UCON1:Unknown | 1096443138.45884 | 1666666666.66667 | 1.25e+09 |
| UCON20:UCON20:Unknown | 0 | 0 | 0 |
| UCON21:DNA:DNA? | 0 | 0 | 0 |
| UCON22:UCON22:Unknown | 0 | 0 | 0 |
| UCON23:hAT-Tip100:DNA | 5482215692.2942 | 1666666666.66667 | 6.25e+09 |
| UCON24:UCON24:Unknown | 0 | 0 | 0 |
| UCON25:UCON25:Unknown | 0 | 555555555.555555 | 6.25e+08 |
| UCON26:UCON26:Unknown | 2192886276.91768 | 2.5e+09 | 6.25e+08 |
| UCON27:UCON27:Unknown | 0 | 555555555.555555 | 1.25e+09 |
| UCON28a:UCON28a:Unknown | 3289329415.37652 | 555555555.555555 | 6.25e+08 |
| UCON28b:UCON28b:Unknown | 16446647076.8826 | 5e+09 | 1.25e+10 |
| UCON28c:UCON28c:Unknown | 1096443138.45884 | 277777777.777778 | 6.25e+08 |
| UCON29:Kolobok:DNA | 0 | 0 | 0 |
| UCON29:PiggyBac:DNA | 2192886276.91768 | 1388888888.88889 | 6.25e+09 |
| UCON2:DNA:DNA? | 8771545107.67073 | 2777777777.77778 | 4.375e+09 |
| UCON31:UCON31:Unknown | 3289329415.37652 | 0 | 0 |
| UCON33:TcMar-Tigger:DNA | 9867988246.12957 | 5833333333.33333 | 4.375e+09 |
| UCON34:UCON34:Unknown | 10964431384.5884 | 2.5e+09 | 1.875e+09 |
| UCON34:hAT-Charlie:DNA | 0 | 0 | 0 |
| UCON35:DNA:DNA | 0 | 0 | 0 |
| UCON35:UCON35:Unknown | 9867988246.12957 | 3055555555.55555 | 1.875e+09 |
| UCON37:UCON37:Unknown | 5482215692.2942 | 555555555.555555 | 3.75e+09 |
| UCON37:hAT-Ac:DNA | 0 | 0 | 0 |
| UCON38:UCON38:Unknown | 3289329415.37652 | 0 | 0 |
| UCON38:hAT-Charlie:DNA | 0 | 0 | 0 |
| UCON39:TcMar-Tigger:DNA | 0 | 1666666666.66667 | 6.25e+08 |
| UCON40:TcMar-Mariner:DNA | 0 | 0 | 0 |
| UCON40:UCON40:Unknown | 1096443138.45884 | 1388888888.88889 | 1.875e+09 |

| | | | |
|---|---|---|---|
| UCON41:UCON41:Unknown | 1096443138.45884 | 0 | 0 |
| UCON43:UCON43:Unknown | 1096443138.45884 | 555555555.555555 | 1.25e+09 |
| UCON44:TcMar-Tigger:DNA | 0 | 0 | 0 |
| UCON44:UCON44:Unknown | 4385772553.83536 | 3611111111.11112 | 1.875e+09 |
| UCON46:UCON46:Unknown | 2192886276.91768 | 555555555.555555 | 0 |
| UCON47:UCON47:Unknown | 1096443138.45884 | 833333333.333333 | 3.75e+09 |
| UCON48:UCON48:Unknown | 3289329415.37652 | 1388888888.88889 | 4.375e+09 |
| UCON49:L2:LINE | 4385772553.83536 | 1666666666.66667 | 5e+09 |
| UCON4:DNA:DNA | 0 | 0 | 0 |
| UCON4:UCON4:Unknown | 6578658830.75305 | 4166666666.66667 | 1e+10 |
| UCON50:UCON50:Unknown | 1096443138.45884 | 1111111111.11111 | 3.125e+09 |
| UCON51:LTR:LTR? | 1096443138.45884 | 277777777.777778 | 6.25e+08 |
| UCON53:UCON53:Unknown | 0 | 833333333.333333 | 0 |
| UCON54:UCON54:Unknown | 2192886276.91768 | 2222222222.22222 | 3.125e+09 |
| UCON55:TcMar-Tigger:DNA | 25218192184.5533 | 10277777777.7778 | 2.125e+10 |
| UCON56:UCON56:Unknown | 0 | 0 | 0 |
| UCON57:UCON57:Unknown | 0 | 0 | 0 |
| UCON58:DNA:DNA | 0 | 0 | 0 |
| UCON58:UCON58:Unknown | 0 | 277777777.777778 | 6.25e+08 |
| UCON59:DNA:DNA | 0 | 0 | 0 |
| UCON59:UCON59:Unknown | 0 | 0 | 0 |
| UCON5:UCON5:Unknown | 0 | 4722222222.22222 | 5e+09 |
| UCON60:UCON60:Unknown | 0 | 0 | 0 |
| UCON61:UCON61:Unknown | 0 | 0 | 0 |
| UCON62:DNA:DNA? | 0 | 555555555.555555 | 1.25e+09 |
| UCON63:UCON63:Unknown | 0 | 0 | 0 |
| UCON64:UCON64:Unknown | 0 | 0 | 0 |
| UCON65:UCON65:Unknown | 5482215692.2942 | 277777777.777778 | 6.25e+08 |
| UCON66:UCON66:Unknown | 0 | 277777777.777778 | 1.25e+09 |
| UCON67:UCON67:Unknown | 0 | 0 | 0 |
| UCON68:UCON68:Unknown | 0 | 0 | 0 |
| UCON69:hAT:DNA | 0 | 0 | 0 |
| UCON6:UCON6:Unknown | 2192886276.91768 | 555555555.555555 | 2.5e+09 |
| UCON70:UCON70:Unknown | 9867988246.12957 | 1111111111.11111 | 6.875e+09 |
| UCON71:UCON71:Unknown | 2192886276.91768 | 0 | 1.875e+09 |
| UCON71_Crp:UCON71_Crp:Unknown | 16446647076.8826 | 6388888888.88888 | 7.5e+09 |
| UCON72:UCON72:Unknown | 1096443138.45884 | 833333333.333333 | 1.25e+09 |
| UCON73:UCON73:Unknown | 8771545107.67073 | 2.5e+09 | 3.125e+09 |
| UCON73:hAT:DNA | 0 | 0 | 0 |
| UCON74:DNA:DNA? | 7675101969.21189 | 2777777777.77778 | 3.75e+09 |
| UCON75:UCON75:Unknown | 0 | 0 | 0 |
| UCON76:UCON76:Unknown | 1096443138.45884 | 0 | 0 |
| UCON77:UCON77:Unknown | 0 | 277777777.777778 | 1.25e+09 |
| UCON78:DNA:DNA | 0 | 555555555.555555 | 5e+09 |
| UCON79:DNA:DNA? | 7675101969.21189 | 1111111111.11111 | 1.25e+09 |
| UCON7:DNA:DNA? | 3289329415.37652 | 833333333.333333 | 6.25e+08 |
| UCON80:hAT:DNA | 1096443138.45884 | 1111111111.11111 | 1.25e+09 |
| UCON80_AMi:UCON80_AMi:Unknown | 18639533353.8003 | 5555555555.55555 | 7.5e+09 |
| UCON81:hAT-Charlie:DNA | 0 | 833333333.333333 | 0 |
| UCON82:UCON82:Unknown | 2192886276.91768 | 555555555.555555 | 6.25e+08 |
| UCON83:DNA:DNA | 0 | 0 | 0 |
| UCON83:SINE:SINE? | 0 | 0 | 0 |
| UCON84:UCON84:Unknown | 3289329415.37652 | 555555555.555555 | 1.25e+09 |
| UCON85:UCON85:Unknown | 0 | 0 | 0 |
| UCON86:L2:LINE | 0 | 2222222222.22222 | 1.875e+09 |
| UCON87:UCON87:Unknown | 0 | 0 | 0 |
| UCON87B:UCON87B:Unknown | 0 | 0 | 0 |
| UCON88:UCON88:Unknown | 1096443138.45884 | 0 | 6.25e+08 |
| UCON89:hAT:DNA | 1096443138.45884 | 0 | 1.25e+09 |
| UCON8:DNA:DNA | 33989737292.2242 | 8333333333.33333 | 3.25e+10 |
| UCON92:UCON92:Unknown | 0 | 833333333.333333 | 0 |
| UCON93:UCON93:Unknown | 0 | 277777777.777778 | 6.25e+08 |
| UCON94:UCON94:Unknown | 0 | 555555555.555555 | 3.125e+09 |
| UCON96:UCON96:Unknown | 0 | 0 | 0 |

| | | | |
|---|---|---|---|
| UCON97:DNA:DNA | 0 | 0 | 0 |
| UCON99:DNA:DNA | 0 | 277777777.777778 | 0 |
| UCON99:Helitron:RC | 0 | 0 | 0 |
| UCON9:UCON9:Unknown | 0 | 0 | 0 |
| UCON9B:UCON9B:Unknown | 0 | 0 | 0 |
| X10b_DNA:TcMar-Tigger:DNA | 0 | 0 | 0 |
| X11_DNA:TcMar-Tigger:DNA | 0 | 0 | 0 |
| X12_DNA:TcMar-Tigger:DNA | 0 | 0 | 0 |
| X12_LINE:I-Jockey:LINE | 0 | 0 | 0 |
| X13_LINE:L2:LINE | 0 | 0 | 0 |
| X15_LINE:L2:LINE | 0 | 0 | 0 |
| X17_LINE:CR1:LINE | 0 | 0 | 0 |
| X1_DNA:TcMar-Tigger:DNA | 0 | 0 | 0 |
| X1_LINE:CR1:LINE | 1096443138.45884 | 277777777.777778 | 6.25e+08 |
| X20_LINE:CR1:LINE | 0 | 0 | 0 |
| X21_LINE:CR1:LINE | 0 | 0 | 0 |
| X23_DNA:DNA:DNA | 0 | 0 | 0 |
| X24_DNA:DNA:DNA | 0 | 0 | 0 |
| X24_LINE:L2:LINE | 0 | 0 | 0 |
| X26_DNA:TcMar-Tigger:DNA | 0 | 0 | 0 |
| X2_LINE:CR1:LINE | 5482215692.2942 | 1.25e+10 | 1.375e+10 |
| X2a_DNA:DNA:DNA | 0 | 0 | 0 |
| X2b_DNA:DNA:DNA | 0 | 0 | 0 |
| X30_DNA:DNA:DNA | 0 | 0 | 0 |
| X32_DNA:TcMar-Tc1:DNA | 0 | 0 | 0 |
| X34_DNA:DNA:DNA | 0 | 0 | 0 |
| X4a_DNA:TcMar-Tigger:DNA | 0 | 0 | 0 |
| X4b_DNA:TcMar-Tigger:DNA | 0 | 0 | 0 |
| X5B_LINE:CR1:LINE | 1096443138.45884 | 1388888888.88889 | 2.5e+09 |
| X5a_DNA:TcMar-Tigger:DNA | 0 | 0 | 0 |
| X6A_LINE:CR1:LINE | 94294109907.4603 | 37222222222.2222 | 7.125e+10 |
| X6B_LINE:CR1:LINE | 39471952984.5182 | 17222222222.2222 | 4.1875e+10 |
| X6a_DNA:TcMar-Tigger:DNA | 0 | 0 | 0 |
| X6b_DNA:TcMar-Tigger:DNA | 0 | 0 | 0 |
| X7A_LINE:CR1:LINE | 61400815753.695 | 46111111111.1112 | 6.4375e+10 |
| X7B_LINE:CR1:LINE | 29603964738.3887 | 16944444444.4445 | 2.625e+10 |
| X7C_LINE:CR1:LINE | 43857725538.3536 | 25555555555.5555 | 5.25e+10 |
| X7D_LINE:CR1:LINE | 0 | 277777777.777778 | 0 |
| X8_LINE:CR1:LINE | 12060874523.0473 | 5833333333.33333 | 1.125e+10 |
| X9_LINE:L1:LINE | 39471952984.5182 | 15555555555.5556 | 3.1875e+10 |
| X9a_DNA:DNA:DNA | 0 | 0 | 0 |
| X9b_DNA:TcMar-Tigger:DNA | 0 | 0 | 0 |
| Zaphod2:hAT-Tip100:DNA | 29603964738.3887 | 15277777777.7778 | 2.9375e+10 |
| Zaphod3:hAT-Tip100:DNA | 103065655015.131 | 39722222222.2222 | 8.4375e+10 |
| Zaphod4a:hAT-Tip100:DNA | 0 | 0 | 0 |
| Zaphod5a:hAT-Tip100:DNA | 0 | 0 | 0 |
| Zaphod5b:hAT-Tip100:DNA | 0 | 0 | 0 |
| Zaphod:hAT-Tip100:DNA | 57015043199.8596 | 32222222222.2222 | 5.25e+10 |
| hAT-16_Crp:hAT-Charlie:DNA | 29603964738.3887 | 3055555555.55555 | 1.1875e+10 |
| hAT-1_Mam:hAT-Tag1:DNA | 0 | 0 | 0 |
| hAT-5_Mam:hAT-Tag1:DNA | 0 | 0 | 0 |
| hAT-N1_Mam:hAT-Tip100:DNA | 24121749046.0945 | 16388888888.8889 | 3.25e+10 |
| hAT-N1a_Mam:hAT-Tip100:DNA | 0 | 0 | 0 |

Appendix File S2 - Accessible Human-Specific SVAs and LTF5HS

| | | | |
|---|---|---|---|
| chr1 | 1409806 | 1410773 | LTR5_Hs |
| chr1 | 10425394 | 10425655 | LTR5_Hs |
| chr1 | 15135337 | 15136296 | LTR5_Hs |
| chr1 | 25587849 | 25588817 | LTR5_Hs |
| chr1 | 46320295 | 46321263 | LTR5_Hs |
| chr1 | 52006244 | 52007211 | LTR5_Hs |
| chr1 | 65139795 | 65140762 | LTR5_Hs |
| chr1 | 70460424 | 70461399 | LTR5_Hs |
| chr1 | 77687252 | 77687779 | SVA_F |
| chr1 | 77982895 | 77983837 | LTR5_Hs |
| chr1 | 79702340 | 79703294 | LTR5_Hs |
| chr1 | 99835833 | 99836768 | LTR5_Hs |
| chr1 | 120424750 | 120425723 | LTR5_Hs |
| chr1 | 144450893 | 144451866 | LTR5_Hs |
| chr1 | 145414872 | 145415845 | LTR5_Hs |
| chr1 | 145932413 | 145933382 | LTR5_Hs |
| chr1 | 146948312 | 146949300 | LTR5_Hs |
| chr1 | 147240099 | 147241067 | LTR5_Hs |
| chr1 | 148137832 | 148138805 | LTR5_Hs |
| chr1 | 149092876 | 149093849 | LTR5_Hs |
| chr1 | 150634829 | 150634922 | SVA_E |
| chr1 | 150634923 | 150635885 | LTR5_Hs |
| chr1 | 155599536 | 155600505 | LTR5_Hs |
| chr1 | 156179222 | 156180190 | LTR5_Hs |
| chr1 | 156181460 | 156182421 | LTR5_Hs |
| chr1 | 160652138 | 160653095 | LTR5_Hs |
| chr1 | 161315707 | 161316604 | LTR5_Hs |
| chr1 | 207635111 | 207636074 | LTR5_Hs |
| chr1 | 207636074 | 207636164 | SVA_F |
| chr10 | 3061468 | 3062440 | LTR5_Hs |
| chr10 | 6824178 | 6825146 | LTR5_Hs |
| chr10 | 6832673 | 6833641 | LTR5_Hs |
| chr10 | 19509806 | 19510764 | LTR5_Hs |
| chr10 | 43337202 | 43338170 | LTR5_Hs |
| chr10 | 90301652 | 90302607 | LTR5_Hs |
| chr10 | 97417834 | 97418118 | LTR5_Hs |
| chr10 | 102454374 | 102455335 | LTR5_Hs |
| chr10 | 102753234 | 102753255 | SVA_F |
| chr10 | 103129261 | 103130573 | SVA_F |
| chr10 | 103129261 | 103130573 | SVA_F |
| chr10 | 110305283 | 110305386 | SVA_F |
| chr11 | 207579 | 207645 | SVA_F |
| chr11 | 9318775 | 9319704 | LTR5_Hs |
| chr11 | 33049341 | 33050285 | LTR5_Hs |
| chr11 | 37858937 | 37859901 | LTR5_Hs |
| chr11 | 56057660 | 56058622 | LTR5_Hs |
| chr11 | 62326599 | 62327566 | LTR5_Hs |
| chr11 | 62382199 | 62383091 | LTR5_Hs |
| chr11 | 62858517 | 62859476 | LTR5_Hs |
| chr11 | 63527970 | 63528938 | LTR5_Hs |
| chr11 | 63530313 | 63531281 | LTR5_Hs |
| chr11 | 67757888 | 67758849 | LTR5_Hs |
| chr11 | 71767204 | 71767231 | LTR5_Hs |
| chr11 | 71767234 | 71767904 | LTR5_Hs |
| chr11 | 90218722 | 90218887 | SVA_F |
| chr11 | 96586132 | 96587100 | LTR5_Hs |
| chr11 | 118721014 | 118721981 | LTR5_Hs |
| chr11 | 119039271 | 119040160 | LTR5_Hs |
| chr12 | 4721230 | 4722199 | LTR5_Hs |
| chr12 | 6885859 | 6886821 | LTR5_Hs |
| chr12 | 8461931 | 8462899 | LTR5_Hs |

| | | | |
|---|---|---|---|
| chr12 | 8635660 | 8636609 | LTR5_Hs |
| chr12 | 9600464 | 9601432 | LTR5_Hs |
| chr12 | 15912016 | 15912381 | SVA_E |
| chr12 | 21644941 | 21645931 | LTR5_Hs |
| chr12 | 55333430 | 55334399 | LTR5_Hs |
| chr12 | 73008722 | 73009695 | LTR5_Hs |
| chr12 | 73853208 | 73853666 | SVA_F |
| chr12 | 85468939 | 85469903 | LTR5_Hs |
| chr12 | 122750859 | 122751831 | LTR5_Hs |
| chr13 | 22813649 | 22814620 | LTR5_Hs |
| chr13 | 28805330 | 28805453 | SVA_F |
| chr13 | 43040543 | 43041509 | LTR5_Hs |
| chr13 | 54266806 | 54267766 | LTR5_Hs |
| chr13 | 69614877 | 69615842 | LTR5_Hs |
| chr13 | 95404162 | 95404252 | SVA_E |
| chr14 | 55024549 | 55025503 | LTR5_Hs |
| chr14 | 64894101 | 64894126 | LTR5_Hs |
| chr14 | 77661397 | 77662365 | LTR5_Hs |
| chr14 | 87020141 | 87020979 | LTR5_Hs |
| chr14 | 100311790 | 100312758 | LTR5_Hs |
| chr15 | 58833341 | 58834296 | LTR5_Hs |
| chr15 | 65226541 | 65227509 | LTR5_Hs |
| chr16 | 10681410 | 10681489 | SVA_E |
| chr16 | 13919492 | 13919574 | SVA_E |
| chr16 | 21221786 | 21222748 | LTR5_Hs |
| chr16 | 23599483 | 23600443 | LTR5_Hs |
| chr16 | 47864458 | 47865426 | LTR5_Hs |
| chr17 | 5074679 | 5075646 | LTR5_Hs |
| chr17 | 8367835 | 8367948 | SVA_E |
| chr17 | 16834879 | 16835857 | LTR5_Hs |
| chr17 | 17610927 | 17611884 | LTR5_Hs |
| chr17 | 30700040 | 30701008 | LTR5_Hs |
| chr17 | 43033948 | 43034010 | SVA_E |
| chr17 | 59290106 | 59290917 | LTR5_Hs |
| chr18 | 4917278 | 4918247 | LTR5_Hs |
| chr18 | 27476539 | 27476584 | LTR5_Hs |
| chr18 | 67076782 | 67077752 | LTR5_Hs |
| chr18 | 68942268 | 68943233 | LTR5_Hs |
| chr18 | 79960165 | 79961063 | LTR5_Hs |
| chr19 | 9584765 | 9584849 | SVA_F |
| chr19 | 20082341 | 20083300 | LTR5_Hs |
| chr19 | 20248489 | 20248737 | LTR5_Hs |
| chr19 | 20285751 | 20286703 | LTR5_Hs |
| chr19 | 20466330 | 20467299 | LTR5_Hs |
| chr19 | 23723657 | 23724624 | LTR5_Hs |
| chr19 | 28130629 | 28131598 | LTR5_Hs |
| chr19 | 36233353 | 36234332 | LTR5_Hs |
| chr19 | 36247143 | 36247924 | LTR5_Hs |
| chr19 | 36247143 | 36247924 | LTR5_Hs |
| chr19 | 36332168 | 36333136 | LTR5_Hs |
| chr19 | 37320787 | 37321678 | LTR5_Hs |
| chr19 | 38631138 | 38632089 | LTR5_Hs |
| chr19 | 38638753 | 38638818 | SVA_F |
| chr19 | 39479461 | 39480441 | LTR5_Hs |
| chr19 | 39959230 | 39960162 | LTR5_Hs |
| chr19 | 51905691 | 51906661 | LTR5_Hs |
| chr19 | 51908191 | 51909157 | LTR5_Hs |
| chr19 | 52485537 | 52486432 | LTR5_Hs |
| chr19 | 52558696 | 52559661 | LTR5_Hs |
| chr19 | 53838625 | 53839592 | LTR5_Hs |
| chr2 | 3104480 | 3105423 | LTR5_Hs |
| chr2 | 15219321 | 15220292 | SVA_E |
| chr2 | 26749864 | 26750871 | LTR5_Hs |
| chr2 | 38277783 | 38278760 | LTR5_Hs |

| | | | |
|---|---|---|---|
| chr2 | 55279270 | 55280234 | LTR5_Hs |
| chr2 | 98129886 | 98130831 | LTR5_Hs |
| chr2 | 108726632 | 108727229 | SVA_F |
| chr2 | 111962842 | 111963809 | LTR5_Hs |
| chr2 | 186521528 | 186522372 | LTR5_Hs |
| chr2 | 192541941 | 192542927 | LTR5_Hs |
| chr2 | 193142415 | 193143387 | LTR5_Hs |
| chr2 | 201139001 | 201139967 | LTR5_Hs |
| chr2 | 202577999 | 202578506 | SVA_F |
| chr2 | 207036832 | 207037793 | LTR5_Hs |
| chr2 | 208199554 | 208200169 | LTR5_Hs |
| chr2 | 214802891 | 214803852 | LTR5_Hs |
| chr2 | 223190520 | 223191443 | LTR5_Hs |
| chr2 | 231576783 | 231577743 | LTR5_Hs |
| chr2 | 232422992 | 232423965 | LTR5_Hs |
| chr2 | 234168064 | 234169033 | LTR5_Hs |
| chr20 | 4032945 | 4034099 | LTR5_Hs |
| chr20 | 4032945 | 4034099 | SVA_E |
| chr20 | 4034076 | 4034982 | SVA_E |
| chr20 | 4034076 | 4034982 | LTR5_Hs |
| chr20 | 23756279 | 23757247 | LTR5_Hs |
| chr20 | 23982640 | 23983609 | LTR5_Hs |
| chr20 | 25233837 | 25234802 | LTR5_Hs |
| chr20 | 25241245 | 25242215 | LTR5_Hs |
| chr20 | 35262123 | 35263091 | LTR5_Hs |
| chr20 | 60041178 | 60042140 | LTR5_Hs |
| chr21 | 14280939 | 14281912 | LTR5_Hs |
| chr21 | 18568678 | 18569644 | LTR5_Hs |
| chr21 | 44282859 | 44283777 | LTR5_Hs |
| chr22 | 18938673 | 18939641 | LTR5_Hs |
| chr22 | 18946884 | 18947848 | LTR5_Hs |
| chr22 | 23536061 | 23537026 | LTR5_Hs |
| chr22 | 24630445 | 24631411 | LTR5_Hs |
| chr22 | 38981115 | 38981255 | SVA_F |
| chr22 | 40500391 | 40501374 | SVA_F |
| chr22_KI270878v1_alt | 160041 | 161006 | LTR5_Hs |
| chr3 | 9847661 | 9848644 | LTR5_Hs |
| chr3 | 9853634 | 9854552 | LTR5_Hs |
| chr3 | 14272844 | 14273811 | LTR5_Hs |
| chr3 | 39418298 | 39419272 | LTR5_Hs |
| chr3 | 47259533 | 47260501 | LTR5_Hs |
| chr3 | 50519891 | 50520848 | LTR5_Hs |
| chr3 | 75393192 | 75394152 | LTR5_Hs |
| chr3 | 100272493 | 100273445 | LTR5_Hs |
| chr3 | 101691892 | 101692861 | LTR5_Hs |
| chr3 | 112564079 | 112565009 | LTR5_Hs |
| chr3 | 125898792 | 125899573 | LTR5_Hs |
| chr3 | 125898792 | 125899573 | LTR5_Hs |
| chr3 | 127411652 | 127412622 | LTR5_Hs |
| chr3 | 129641274 | 129642062 | LTR5_Hs |
| chr3 | 129641274 | 129642062 | LTR5_Hs |
| chr3 | 130057288 | 130058265 | LTR5_Hs |
| chr3 | 146522264 | 146523231 | LTR5_Hs |
| chr3 | 186886873 | 186887834 | LTR5_Hs |
| chr3 | 186893219 | 186894186 | LTR5_Hs |
| chr3 | 196944079 | 196945028 | LTR5_Hs |
| chr4 | 3925642 | 3926609 | LTR5_Hs |
| chr4 | 9731773 | 9732691 | LTR5_Hs |
| chr4 | 23372101 | 23372551 | SVA_E |
| chr4 | 66358756 | 66359723 | LTR5_Hs |
| chr4 | 159976487 | 159977451 | LTR5_Hs |
| chr4 | 164905513 | 164906483 | LTR5_Hs |
| chr4 | 164990382 | 164991350 | LTR5_Hs |
| chr4 | 164995687 | 164996655 | LTR5_Hs |

| | | | |
|---|---|---|---|
| chr4 | 165001952 | 165002916 | LTR5_Hs |
| chr4 | 174383028 | 174384002 | LTR5_Hs |
| chr4 | 174400759 | 174401732 | LTR5_Hs |
| chr4 | 188068119 | 188068743 | SVA_F |
| chr5 | 1595976 | 1596944 | LTR5_Hs |
| chr5 | 55571079 | 55572039 | LTR5_Hs |
| chr5 | 56156996 | 56157967 | LTR5_Hs |
| chr5 | 116821138 | 116822098 | LTR5_Hs |
| chr5 | 120194838 | 120195651 | LTR5_Hs |
| chr5 | 122809986 | 122810418 | SVA_E |
| chr5 | 181266861 | 181267826 | LTR5_Hs |
| chr6 | 1263272 | 1263316 | SVA_F |
| chr6 | 2908662 | 2909630 | LTR5_Hs |
| chr6 | 3054799 | 3055162 | LTR5_Hs |
| chr6 | 24400046 | 24401019 | LTR5_Hs |
| chr6 | 25999230 | 26000000 | LTR5_Hs |
| chr6 | 27825845 | 27826625 | LTR5_Hs |
| chr6 | 33809959 | 33810919 | LTR5_Hs |
| chr6 | 34449846 | 34449919 | SVA_E |
| chr6 | 34719459 | 34720428 | LTR5_Hs |
| chr6 | 44328047 | 44329009 | LTR5_Hs |
| chr6 | 52922679 | 52923640 | LTR5_Hs |
| chr6 | 89890820 | 89891765 | LTR5_Hs |
| chr6 | 93173365 | 93174330 | LTR5_Hs |
| chr6 | 95560622 | 95561586 | LTR5_Hs |
| chr6 | 111255096 | 111256055 | LTR5_Hs |
| chr6 | 115414996 | 115415149 | LTR5_Hs |
| chr6 | 120931437 | 120932413 | LTR5_Hs |
| chr6 | 151455209 | 151456185 | LTR5_Hs |
| chr7 | 2267299 | 2269533 | SVA_F |
| chr7 | 2389406 | 2390372 | LTR5_Hs |
| chr7 | 6901996 | 6902963 | LTR5_Hs |
| chr7 | 7001776 | 7002700 | LTR5_Hs |
| chr7 | 16197721 | 16198689 | LTR5_Hs |
| chr7 | 47989678 | 47990639 | LTR5_Hs |
| chr7 | 54675200 | 54676131 | LTR5_Hs |
| chr7 | 65260236 | 65261141 | SVA_E |
| chr7 | 65260236 | 65261141 | LTR5_Hs |
| chr7 | 66618957 | 66619926 | LTR5_Hs |
| chr7 | 119100246 | 119101210 | LTR5_Hs |
| chr7 | 123780636 | 123781604 | LTR5_Hs |
| chr7 | 139467404 | 139468373 | LTR5_Hs |
| chr7 | 140552983 | 140553942 | LTR5_Hs |
| chr7 | 151027013 | 151027969 | LTR5_Hs |
| chr8 | 58671738 | 58672704 | LTR5_Hs |
| chr8 | 112772173 | 112773142 | LTR5_Hs |
| chr8 | 119917331 | 119918257 | LTR5_Hs |
| chr8 | 143831988 | 143832957 | LTR5_Hs |
| chr8 | 144795893 | 144796861 | LTR5_Hs |
| chr9 | 17445360 | 17446328 | LTR5_Hs |
| chr9 | 26715149 | 26716178 | LTR5_Hs |
| chr9 | 31632878 | 31633846 | LTR5_Hs |
| chr9 | 69763136 | 69764104 | LTR5_Hs |
| chr9 | 83966145 | 83966210 | SVA_E |
| chr9 | 97232135 | 97233093 | LTR5_Hs |
| chr9 | 121429105 | 121430073 | LTR5_Hs |
| chr9 | 133957779 | 133958741 | LTR5_Hs |
| chrUn_KI270442v1 | 390369 | 391354 | LTR5_Hs |
| chrX | 23659882 | 23660832 | LTR5_Hs |
| chrX | 57335163 | 57336040 | LTR5_Hs |
| chrX | 72733184 | 72734156 | LTR5_Hs |
| chrX | 113990609 | 113990894 | LTR5_Hs |
| chrX | 113990913 | 113991079 | LTR5_Hs |
| chrX | 135302738 | 135303701 | LTR5_Hs |

| | | | |
|---|---|---|---|
| chrX | 135406098 | 135407066 | LTR5_Hs |
| chrX | 138333486 | 138334454 | LTR5_Hs |
| chrX | 139341914 | 139342888 | LTR5_Hs |

### Homer Known Motif Enrichment Results (/rds/general/user/ldeelen/home)

Homer *de novo* Motif Results
Gene Ontology Enrichment Results
Known Motif Enrichment Results (txt file)
Total Target Sequences = 187, Total Background Sequences = 1123

| Rank | Motif | Name | P-value | log P-pvalue | q-value (Benjamini) | # Target Sequences with Motif | % of Targets Sequences with Motif | # Background Sequences with Motif | % of Background Sequences with Motif | Motif File | SVG |
|---|---|---|---|---|---|---|---|---|---|---|---|
| 1 | | FoxD3(forkhead)/ZebrafishEmbryo-Foxd3.biotin-ChIP-seq(GSE106676)/Homer | 1e-12 | -2.866e+01 | 0.0000 | 61.0 | 32.62% | 120.1 | 10.69% | motif file (matrix) | svg |
| 2 | | KANADI1(Myb)/Seedling-KAN1-ChIP-Seq(GSE48081)/Homer | 1e-10 | -2.356e+01 | 0.0000 | 81.0 | 43.32% | 227.0 | 20.21% | motif file (matrix) | svg |
| 3 | | At5g29000(G2like)/col-At5g29000-DAP-Seq(GSE60143)/Homer | 1e-10 | -2.334e+01 | 0.0000 | 67.0 | 35.83% | 166.0 | 14.77% | motif file (matrix) | svg |
| 4 | | At1g49010(MYBrelated)/col-At1g49010-DAP-Seq(GSE60143)/Homer | 1e-10 | -2.308e+01 | 0.0000 | 117.0 | 62.57% | 420.0 | 37.38% | motif file (matrix) | svg |
| 5 | | At2g03500(G2like)/col-At2g03500-DAP-Seq(GSE60143)/Homer | 1e-9 | -2.296e+01 | 0.0000 | 68.0 | 36.36% | 172.0 | 15.30% | motif file (matrix) | svg |
| 6 | | AT2G20400(G2like)/colamp-AT2G20400-DAP-Seq(GSE60143)/Homer | 1e-9 | -2.238e+01 | 0.0000 | 66.0 | 35.29% | 165.3 | 14.72% | motif file (matrix) | svg |
| 7 | | KAN2(G2like)/colamp-KAN2-DAP-Seq(GSE60143)/Homer | 1e-9 | -2.126e+01 | 0.0000 | 97.0 | 51.87% | 320.8 | 28.55% | motif file (matrix) | svg |
| 8 | | At5g08520(MYBrelated)/colamp-At5g08520-DAP-Seq(GSE60143)/Homer | 1e-9 | -2.083e+01 | 0.0000 | 135.0 | 72.19% | 545.6 | 48.56% | motif file (matrix) | svg |
| 9 | | Hnf1(Homeobox)/Liver-Foxa2-Chip-Seq(GSE25694)/Homer | 1e-9 | -2.081e+01 | 0.0000 | 122.0 | 65.24% | 464.3 | 41.32% | motif file (matrix) | svg |
| 10 | | HNF1b(Homeobox)/PDAC-HNF1B-ChIP-Seq(GSE64557)/Homer | 1e-8 | -2.053e+01 | 0.0000 | 120.0 | 64.17% | 454.0 | 40.41% | motif file (matrix) | svg |
| 11 | | At3g04030(G2like)/col-At3g04030-DAP-Seq(GSE60143)/Homer | 1e-8 | -2.041e+01 | 0.0000 | 73.0 | 39.04% | 206.8 | 18.40% | motif file (matrix) | svg |
| 12 | | At5g58900(MYBrelated)/colamp-At5g58900-DAP-Seq(GSE60143)/Homer | 1e-8 | -1.997e+01 | 0.0000 | 120.0 | 64.17% | 458.6 | 40.81% | motif file (matrix) | svg |
| 13 | | Six2(Homeobox)/NephronProgenitor-Six2-ChIP-Seq(GSE39837)/Homer | 1e-8 | -1.965e+01 | 0.0000 | 96.0 | 51.34% | 325.0 | 28.93% | motif file (matrix) | svg |
| 14 | | ATHB53(HB)/col-ATHB53-DAP-Seq(GSE60143)/Homer | 1e-8 | -1.943e+01 | 0.0000 | 33.0 | 17.65% | 51.4 | 4.57% | motif file (matrix) | svg |
| 15 | | Znf263(Zf)/K562-Znf263-ChIP-Seq(GSE31477)/Homer | 1e-8 | -1.923e+01 | 0.0000 | 88.0 | 47.06% | 286.4 | 25.49% | motif file (matrix) | svg |
| 16 | | BPC1(BBRBPC)/colamp-BPC1-DAP-Seq(GSE60143)/Homer | 1e-8 | -1.912e+01 | 0.0000 | 109.0 | 58.29% | 400.3 | 35.62% | motif file (matrix) | svg |
| 17 | | ATHB21(HB)/colamp-ATHB21-DAP-Seq(GSE60143)/Homer | 1e-8 | -1.855e+01 | 0.0000 | 29.0 | 15.51% | 41.6 | 3.71% | motif file (matrix) | svg |
| 18 | | At1g74840(MYBrelated)/col100-At1g74840-DAP-Seq(GSE60143)/Homer | 1e-7 | -1.810e+01 | 0.0000 | 128.0 | 68.45% | 521.8 | 46.44% | motif file (matrix) | svg |
| 19 | | AT2G33550(Trihelix)/colamp-AT2G33550-DAP-Seq(GSE60143)/Homer | 1e-7 | -1.790e+01 | 0.0000 | 101.0 | 54.01% | 364.6 | 32.45% | motif file (matrix) | svg |
| 20 | | COG1(C2C2dof)/col-COG1-DAP-Seq(GSE60143)/Homer | 1e-7 | -1.760e+01 | 0.0000 | 72.0 | 38.50% | 217.4 | 19.35% | motif file (matrix) | svg |
| 21 | | At1g19000(MYBrelated)/colamp-At1g19000-DAP-Seq(GSE60143)/Homer | 1e-7 | -1.748e+01 | 0.0000 | 63.0 | 33.69% | 176.9 | 15.74% | motif file (matrix) | svg |
| 22 | | ATHB5(HB)/colamp-ATHB5-DAP-Seq(GSE60143)/Homer | 1e-7 | -1.744e+01 | 0.0000 | 36.0 | 19.25% | 67.9 | 6.04% | motif file (matrix) | svg |
| 23 | | ATHB6(Homeobox)/Arabidopsis-HB6-ChIP-Seq(GSE80564)/Homer | 1e-7 | -1.744e+01 | 0.0000 | 36.0 | 19.25% | 67.4 | 6.00% | motif file (matrix) | svg |
| 24 | | LBD19(LOBAS2)/colamp-LBD19-DAP-Seq(GSE60143)/Homer | 1e-7 | -1.730e+01 | 0.0000 | 96.0 | 51.34% | 341.1 | 30.36% | motif file (matrix) | svg |
| 25 | | AT4G00250(GeBP)/col-AT4G00250-DAP-Seq(GSE60143)/Homer | 1e-7 | -1.730e+01 | 0.0000 | 61.0 | 32.62% | 168.3 | 14.98% | motif file (matrix) | svg |
| 26 | | MafB(bZIP)/BMM-Mafb-ChIP-Seq(GSE75722)/Homer | 1e-7 | -1.674e+01 | 0.0000 | 96.0 | 51.34% | 345.3 | 30.73% | motif file (matrix) | svg |
| 27 | | ZNF652/HepG2-ZNF652.Flag-ChIP-Seq(Encode)/Homer | 1e-7 | -1.661e+01 | 0.0000 | 96.0 | 51.34% | 346.5 | 30.84% | motif file (matrix) | svg |
| 28 | | Zelda(Zf)/Embryo-zld-ChIP-Seq(GSE65441)/Homer | 1e-7 | -1.659e+01 | 0.0000 | 27.0 | 14.44% | 40.3 | 3.59% | motif file (matrix) | svg |
| 29 | | Unknown2/Drosophila-Promoters/Homer | 1e-7 | -1.657e+01 | 0.0000 | 165.0 | 88.24% | 794.0 | 70.67% | motif file (matrix) | svg |
| 30 | | ATHB40(HB)/col-ATHB40-DAP-Seq(GSE60143)/Homer | 1e-7 | -1.612e+01 | 0.0000 | 35.0 | 18.72% | 68.5 | 6.09% | motif file (matrix) | svg |
| 31 | | LMI1(HB)/colamp-LMI1-DAP-Seq(GSE60143)/Homer | 1e-7 | -1.612e+01 | 0.0000 | 35.0 | 18.72% | 68.3 | 6.08% | motif file (matrix) | svg |
| 32 | | ATHB6(Homeobox)/col-ATHB6-DAP-Seq(GSE60143)/Homer | 1e-6 | -1.592e+01 | 0.0000 | 37.0 | 19.79% | 76.5 | 6.81% | motif file (matrix) | svg |
| 33 | | AT5G56840(MYBrelated)/colamp-AT5G56840-DAP-Seq(GSE60143)/Homer | 1e-6 | -1.526e+01 | 0.0000 | 69.0 | 36.90% | 217.2 | 19.33% | motif file (matrix) | svg |
| 34 | | Phox2b(Homeobox)/CLBGA-PHOX2B-ChIP-Seq(GSE90683)/Homer | 1e-6 | -1.521e+01 | 0.0000 | 116.0 | 62.03% | 472.0 | 42.01% | motif file (matrix) | svg |
| 35 | | GTL1(Trihelix)/colamp-GTL1-DAP-Seq(GSE60143)/Homer | 1e-6 | -1.492e+01 | 0.0000 | 110.0 | 58.82% | 438.5 | 39.03% | motif file (matrix) | svg |
| 36 | | Phox2a(Homeobox)/Neuron-Phox2a-ChIP-Seq(GSE31456)/Homer | 1e-6 | -1.486e+01 | 0.0000 | 116.0 | 62.03% | 474.1 | 42.20% | motif file (matrix) | svg |
| 37 | | ATHB33(ZFHD)/col-ATHB33-DAP-Seq(GSE60143)/Homer | 1e-6 | -1.478e+01 | 0.0000 | 44.0 | 23.53% | 108.8 | 9.69% | motif file (matrix) | svg |
| 38 | | At5g47390(MYBrelated)/col-At5g47390-DAP-Seq(GSE60143)/Homer | 1e-6 | -1.469e+01 | 0.0000 | 67.0 | 35.83% | 211.9 | 18.86% | motif file (matrix) | svg |
| 39 | | CRC(C2C2YABBY)/col-CRC-DAP-Seq(GSE60143)/Homer | 1e-6 | -1.445e+01 | 0.0000 | 65.0 | 34.76% | 203.0 | 18.07% | motif file (matrix) | svg |
| 40 | | AT5G61620(MYBrelated)/colamp-AT5G61620-DAP-Seq(GSE60143)/Homer | 1e-6 | -1.436e+01 | 0.0000 | 68.0 | 36.36% | 218.2 | 19.42% | motif file (matrix) | svg |
| 41 | | KLF5(Zf)/LoVo-KLF5-ChIP-Seq(GSE49402)/Homer | 1e-6 | -1.408e+01 | 0.0000 | 154.0 | 82.35% | 730.4 | 65.01% | motif file (matrix) | svg |
| 42 | | ATHB13(Homeobox)/col-ATHB13-DAP-Seq(GSE60143)/Homer | 1e-6 | -1.396e+01 | 0.0000 | 36.0 | 19.25% | 80.4 | 7.15% | motif file (matrix) | svg |
| 43 | | MYB41(MYB)/col-MYB41-DAP-Seq(GSE60143)/Homer | 1e-5 | -1.339e+01 | 0.0000 | 65.0 | 34.76% | 210.3 | 18.72% | motif file (matrix) | svg |
| 44 | | ATHB7(Homeobox)/col-ATHB7-DAP-Seq(GSE60143)/Homer | 1e-5 | -1.323e+01 | 0.0000 | 29.0 | 15.51% | 57.4 | 5.11% | motif file (matrix) | svg |
| 45 | | Otx2(Homeobox)/EpiLC-Otx2-ChIP-Seq(GSE56098)/Homer | 1e-5 | -1.295e+01 | 0.0001 | 167.0 | 89.30% | 839.3 | 74.70% | motif file (matrix) | svg |
| 46 | | Eomes(T-box)/H9-Eomes-ChIP-Seq(GSE26097)/Homer | 1e-5 | -1.230e+01 | 0.0001 | 161.0 | 86.10% | 798.5 | 71.07% | motif file (matrix) | svg |
| 47 | | PR(NR)/T47D-PR-ChIP-Seq(GSE31130)/Homer | 1e-5 | -1.216e+01 | 0.0001 | 166.0 | 88.77% | 838.5 | 74.63% | motif file (matrix) | svg |
| 48 | | FOXP1(Forkhead)/H9-FOXP1-ChIP-Seq(GSE31006)/Homer | 1e-5 | -1.209e+01 | 0.0001 | 68.0 | 36.36% | 235.0 | 20.91% | motif file (matrix) | svg |
| 49 | | AT2G38300(G2like)/col-AT2G38300-DAP-Seq(GSE60143)/Homer | 1e-5 | -1.199e+01 | 0.0001 | 157.0 | 83.96% | 771.8 | 68.69% | motif file (matrix) | svg |
| 50 | | ANAC083(NAC)/col-ANAC083-DAP-Seq(GSE60143)/Homer | 1e-5 | -1.195e+01 | 0.0001 | 34.0 | 18.18% | 81.4 | 7.25% | motif file (matrix) | svg |
| 51 | | AR-halfsite(NR)/LNCaP-AR-ChIP-Seq(GSE27824)/Homer | 1e-5 | -1.194e+01 | 0.0001 | 172.0 | 91.98% | 889.1 | 79.13% | motif file (matrix) | svg |
| 52 | | SUT1?/SacCer-Promoters/Homer | 1e-5 | -1.172e+01 | 0.0002 | 177.0 | 94.65% | 935.4 | 83.25% | motif file (matrix) | svg |
| 53 | | ASHR1(ND)/col-ASHR1-DAP-Seq(GSE60143)/Homer | 1e-5 | -1.165e+01 | 0.0002 | 159.0 | 85.03% | 789.6 | 70.28% | motif file (matrix) | svg |
| 54 | | At3g24120(G2like)/col-At3g24120-DAP-Seq(GSE60143)/Homer | 1e-4 | -1.149e+01 | 0.0002 | 169.0 | 90.37% | 868.9 | 77.34% | motif file (matrix) | svg |
| 55 | | EBF2(EBF)/BrownAdipose-EBF2-ChIP-Seq(GSE97114)/Homer | 1e-4 | -1.138e+01 | 0.0002 | 99.0 | 52.94% | 406.8 | 36.20% | motif file (matrix) | svg |
| 56 | | Foxa2(Forkhead)/Liver-Foxa2-ChIP-Seq(GSE25694)/Homer | 1e-4 | -1.137e+01 | 0.0002 | 55.0 | 29.41% | 176.8 | 15.74% | motif file (matrix) | svg |
| 57 | | FOXK2(Forkhead)/U2OS-FOXK2-ChIP-Seq(E-MTAB-2204)/Homer | 1e-4 | -1.134e+01 | 0.0002 | 105.0 | 56.15% | 441.4 | 39.28% | motif file (matrix) | svg |
| 58 | | Nkx6.1(Homeobox)/Islet-Nkx6.1-ChIP-Seq(GSE40975)/Homer | 1e-4 | -1.122e+01 | 0.0002 | 101.0 | 54.01% | 419.9 | 37.38% | motif file (matrix) | svg |
| 59 | | Isl1(Homeobox)/Neuron-Isl1-ChIP-Seq(GSE31456)/Homer | 1e-4 | -1.116e+01 | 0.0002 | 166.0 | 88.77% | 847.4 | 75.42% | motif file (matrix) | svg |
| 60 | | Cdx2(Homeobox)/mES-Cdx2-ChIP-Seq(GSE14586)/Homer | 1e-4 | -1.112e+01 | 0.0002 | 73.0 | 39.04% | 267.8 | 23.83% | motif file (matrix) | svg |
| 61 | | BBX31(Orphan)/col-BBX31-DAP-Seq(GSE60143)/Homer | 1e-4 | -1.110e+01 | 0.0002 | 83.0 | 44.39% | 320.9 | 28.56% | motif file (matrix) | svg |
| 62 | | Sox10(HMG)/SciaticNerve-Sox3-ChIP-Seq(GSE35132)/Homer | 1e-4 | -1.108e+01 | 0.0003 | 165.0 | 88.24% | 840.7 | 74.82% | motif file (matrix) | svg |
| 63 | | LRF(Zf)/Erythroblasts-ZBTB7A-ChIP-Seq(GSE74977)/Homer | 1e-4 | -1.103e+01 | 0.0003 | 74.0 | 39.57% | 273.6 | 24.35% | motif file (matrix) | svg |
| 64 | | dof43(C2C2dof)/colamp-dof43-DAP-Seq(GSE60143)/Homer | 1e-4 | -1.102e+01 | 0.0003 | 81.0 | 43.32% | 310.9 | 27.67% | motif file (matrix) | svg |
| 65 | | Bach2(bZIP)/OCILy7-Bach2-ChIP-Seq(GSE44420)/Homer | 1e-4 | -1.100e+01 | 0.0003 | 107.0 | 57.22% | 456.7 | 40.65% | motif file (matrix) | svg |
| 66 | | NF1-halfsite(CTF)/LNCaP-NF1-ChIP-Seq(Unpublished)/Homer | 1e-4 | -1.087e+01 | 0.0003 | 72.0 | 38.50% | 264.5 | 23.54% | motif file (matrix) | svg |
| 67 | | ATY19(MYB)/col-ATY19-DAP-Seq(GSE60143)/Homer | 1e-4 | -1.079e+01 | 0.0003 | 64.0 | 34.22% | 224.2 | 19.96% | motif file (matrix) | svg |
| 68 | | MNT(bHLH)/HepG2-MNT-ChIP-Seq(Encode)/Homer | 1e-4 | -1.073e+01 | 0.0003 | 166.0 | 88.77% | 851.6 | 75.79% | motif file (matrix) | svg |
| 69 | | PHA-4(Forkhead)/cElegans-Embryos-PHA4-ChIP-Seq(modEncode)/Homer | 1e-4 | -1.073e+01 | 0.0003 | 170.0 | 90.91% | 883.4 | 78.63% | motif file (matrix) | svg |
| 70 | | MYB107(MYB)/col-MYB107-DAP-Seq(GSE60143)/Homer | 1e-4 | -1.073e+01 | 0.0003 | 71.0 | 37.97% | 260.8 | 23.22% | motif file (matrix) | svg |
| 71 | | At2g01060(G2like)/colamp-At2g01060-DAP-Seq(GSE60143)/Homer | 1e-4 | -1.072e+01 | 0.0003 | 168.0 | 89.84% | 867.3 | 77.20% | motif file (matrix) | svg |
| 72 | | Tgif2(Homeobox)/mES-Tgif2-ChIP-Seq(GSE55404)/Homer | 1e-4 | -1.069e+01 | 0.0003 | 145.0 | 77.54% | 699.6 | 62.26% | motif file (matrix) | svg |
| 73 | | GSC(Homeobox)/FrogEmbryos-GSC-ChIP-Seq(DRA000576)/Homer | 1e-4 | -1.068e+01 | 0.0003 | 174.0 | 93.05% | 917.1 | 81.62% | motif file (matrix) | svg |
| 74 | | CDX4(Homeobox)/ZebrafishEmbryos-Cdx4.Myc-ChIP-Seq(GSE48254)/Homer | 1e-4 | -1.056e+01 | 0.0004 | 74.0 | 39.57% | 277.4 | 24.69% | motif file (matrix) | svg |
| 75 | | Elf4(ETS)/BMDM-Elf4-ChIP-Seq(GSE88699)/Homer | 1e-4 | -1.042e+01 | 0.0004 | 156.0 | 83.42% | 779.3 | 69.36% | motif file (matrix) | svg |
| 76 | | MYB17(MYB)/colamp-MYB17-DAP-Seq(GSE60143)/Homer | 1e-4 | -1.041e+01 | 0.0004 | 62.0 | 33.16% | 217.5 | 19.36% | motif file (matrix) | svg |
| 77 | | FOXK1(Forkhead)/HEK293-FOXK1-ChIP-Seq(GSE51673)/Homer | 1e-4 | -1.035e+01 | 0.0004 | 162.0 | 86.63% | 824.8 | 73.41% | motif file (matrix) | svg |
| 78 | | AT1G72740(MYBrelated)/colamp-AT1G72740-DAP-Seq(GSE60143)/Homer | 1e-4 | -1.033e+01 | 0.0004 | 91.0 | 48.66% | 371.6 | 33.07% | motif file (matrix) | svg |
| 79 | | BMAL1(bHLH)/Liver-Bmal1-ChIP-Seq(GSE39860)/Homer | 1e-4 | -1.029e+01 | 0.0004 | 167.0 | 89.30% | 863.0 | 76.81% | motif file (matrix) | svg |
| 80 | | bcd(Homeobox)/Embryo-Bcd-ChIP-Seq(GSE86966)/Homer | 1e-4 | -1.029e+01 | 0.0004 | 170.0 | 90.91% | 887.0 | 78.95% | motif file (matrix) | svg |
| 81 | | FoxL2(Forkhead)/Ovary-FoxL2-ChIP-Seq(GSE60858)/Homer | 1e-4 | -1.025e+01 | 0.0004 | 75.0 | 40.11% | 285.1 | 25.37% | motif file (matrix) | svg |
| 82 | | ANAC045(NAC)/col-ANAC045-DAP-Seq(GSE60143)/Homer | 1e-4 | -1.024e+01 | 0.0004 | 48.0 | 25.67% | 151.5 | 13.49% | motif file (matrix) | svg |
| 83 | | AT3G25990(Trihelix)/colamp-AT3G25990-DAP-Seq(GSE60143)/Homer | 1e-4 | -1.020e+01 | 0.0004 | 123.0 | 65.78% | 561.4 | 49.97% | motif file (matrix) | svg |
| 84 | | CRX(Homeobox)/Retina-Crx-ChIP-Seq(GSE20012)/Homer | 1e-4 | -1.016e+01 | 0.0005 | 173.0 | 92.51% | 913.0 | 81.26% | motif file (matrix) | svg |
| 85 | | At1g64620(C2C2dof)/colamp-At1g64620-DAP-Seq(GSE60143)/Homer | 1e-4 | -1.016e+01 | 0.0005 | 79.0 | 42.25% | 307.8 | 27.40% | motif file (matrix) | svg |
| 86 | | Sp2(Zf)/HEK293-Sp2.eGFP-ChIP-Seq(Encode)/Homer | 1e-4 | -1.016e+01 | 0.0005 | 137.0 | 73.26% | 651.4 | 57.98% | motif file (matrix) | svg |
| 87 | | caudal(Homeobox)/Drosophila-Embryos-ChIP-Chip(modEncode)/Homer | 1e-4 | -1.016e+01 | 0.0005 | 86.0 | 45.99% | 345.7 | 30.77% | motif file (matrix) | svg |
| 88 | | SeqBias: CG bias | 1e-4 | -1.015e+01 | 0.0005 | 182.0 | 97.33% | 996.0 | 88.65% | motif file (matrix) | svg |
| 89 | | ZNF341(Zf)/EBV-ZNF341-ChIP-Seq(GSE113194)/Homer | 1e-4 | -1.011e+01 | 0.0005 | 14.0 | 7.49% | 18.2 | 1.62% | motif file (matrix) | svg |
| 90 | | JGL(C2H2)/col-JGL-DAP-Seq(GSE60143)/Homer | 1e-4 | -1.003e+01 | 0.0005 | 118.0 | 63.10% | 532.2 | 47.36% | motif file (matrix) | svg |
| 91 | | MYB(HTH)/ERMYB-Myb-ChIPSeq(GSE22095)/Homer | 1e-4 | -1.001e+01 | 0.0005 | 159.0 | 85.03% | 805.6 | 71.70% | motif file (matrix) | svg |
| 92 | | AT5G47660(Trihelix)/colamp-AT5G47660-DAP-Seq(GSE60143)/Homer | 1e-4 | -1.000e+01 | 0.0005 | 109.0 | 58.29% | 478.1 | 42.55% | motif file (matrix) | svg |
| 93 | | Sox3(HMG)/NPC-Sox3-ChIP-Seq(GSE33059)/Homer | 1e-4 | -9.967e+00 | 0.0005 | 168.0 | 89.84% | 874.5 | 77.83% | motif file (matrix) | svg |
| 94 | | Trl(Zf)/S2-GAGAfactor-ChIP-Seq(GSE40646)/Homer | 1e-4 | -9.967e+00 | 0.0005 | 168.0 | 89.84% | 874.3 | 77.82% | motif file (matrix) | svg |
| 95 | | SEP3(MADS)/Arabidopsis-Flower-Sep3-ChIP-Seq/Homer | 1e-4 | -9.934e+00 | 0.0005 | 58.0 | 31.02% | 201.9 | 17.97% | motif file (matrix) | svg |
| 96 | | TBP3(MYBrelated)/col-TBP3-DAP-Seq(GSE60143)/Homer | 1e-4 | -9.925e+00 | 0.0005 | 145.0 | 77.54% | 707.7 | 62.99% | motif file (matrix) | svg |
| 97 | | AT4G26030(C2H2)/col-AT4G26030-DAP-Seq(GSE60143)/Homer | 1e-4 | -9.919e+00 | 0.0005 | 63.0 | 33.69% | 226.3 | 20.14% | motif file (matrix) | svg |
| 98 | | Tgif1(Homeobox)/mES-Tgif1-ChIP-Seq(GSE55404)/Homer | 1e-4 | -9.874e+00 | 0.0005 | 143.0 | 76.47% | 694.3 | 61.80% | motif file (matrix) | svg |
| 99 | | MYB40(MYB)/col-MYB40-DAP-Seq(GSE60143)/Homer | 1e-4 | -9.795e+00 | 0.0006 | 60.0 | 32.09% | 213.0 | 18.96% | motif file (matrix) | svg |
| 100 | | MYB51(MYB)/col-MYB51-DAP-Seq(GSE60143)/Homer | 1e-4 | -9.795e+00 | 0.0006 | 62.0 | 33.16% | 222.1 | 19.77% | motif file (matrix) | svg |
| 101 | | Foxf1(Forkhead)/Lung-Foxf1-ChIP-Seq(GSE77951)/Homer | 1e-4 | -9.740e+00 | 0.0006 | 162.0 | 86.63% | 830.7 | 73.93% | motif file (matrix) | svg |
| 102 | | ETV1(ETS)/GIST48-ETV1-ChIP-Seq(GSE22441)/Homer | 1e-4 | -9.713e+00 | 0.0006 | 155.0 | 82.89% | 779.3 | 69.36% | motif file (matrix) | svg |
| 103 | | Foxo3(Forkhead)/U2OS-Foxo3-ChIP-Seq(E-MTAB-2701)/Homer | 1e-4 | -9.693e+00 | 0.0006 | 161.0 | 86.10% | 823.4 | 73.29% | motif file (matrix) | svg |
| 104 | | At3g11280(MYBrelated)/col-At3g11280-DAP-Seq(GSE60143)/Homer | 1e-4 | -9.515e+00 | 0.0007 | 76.0 | 40.64% | 297.7 | 26.49% | motif file (matrix) | svg |
| 105 | | ANAC094(NAC)/col-ANAC094-DAP-Seq(GSE60143)/Homer | 1e-4 | -9.447e+00 | 0.0008 | 38.0 | 20.32% | 111.9 | 9.96% | motif file (matrix) | svg |
| 106 | | At3g60580(C2H2)/col-At3g60580-DAP-Seq(GSE60143)/Homer | 1e-4 | -9.441e+00 | 0.0008 | 177.0 | 94.65% | 954.8 | 84.98% | motif file (matrix) | svg |
| 107 | | GATA3(Zf)/iTreg-Gata3-ChIP-Seq(GSE20898)/Homer | 1e-4 | -9.438e+00 | 0.0008 | 169.0 | 90.37% | 887.3 | 78.97% | motif file (matrix) | svg |
| 108 | | At5g05790(MYBrelated)/col-At5g05790-DAP-Seq(GSE60143)/Homer | 1e-4 | -9.409e+00 | 0.0008 | 76.0 | 40.64% | 298.4 | 26.56% | motif file (matrix) | svg |
| 109 | | MYB10(MYB)/col-MYB10-DAP-Seq(GSE60143)/Homer | 1e-4 | -9.311e+00 | 0.0008 | 60.0 | 32.09% | 216.3 | 19.25% | motif file (matrix) | svg |
| 110 | | TCFL2(HMG)/K562-TCF7L2-ChIP-Seq(GSE29196)/Homer | 1e-4 | -9.288e+00 | 0.0008 | 51.0 | 27.27% | 172.1 | 15.32% | motif file (matrix) | svg |
| 111 | | MYB61(MYB)/colamp-MYB61-DAP-Seq(GSE60143)/Homer | 1e-4 | -9.270e+00 | 0.0009 | 67.0 | 35.83% | 252.8 | 22.50% | motif file (matrix) | svg |
| 112 | | AT5G25475(ABI3VP1)/col-AT5G25475-DAP-Seq(GSE60143)/Homer | 1e-4 | -9.263e+00 | 0.0009 | 160.0 | 85.56% | 820.2 | 73.00% | motif file (matrix) | svg |
| 113 | | Usf2(bHLH)/C2C12-Usf2-ChIP-Seq(GSE36030)/Homer | 1e-4 | -9.263e+00 | 0.0009 | 139.0 | 74.33% | 674.8 | 60.06% | motif file (matrix) | svg |
| 114 | | AT2G40260(G2like)/colamp-AT2G40260-DAP-Seq(GSE60143)/Homer | 1e-3 | -9.184e+00 | 0.0009 | 166.0 | 88.77% | 866.0 | 77.08% | motif file (matrix) | svg |
| 115 | | MYB99(MYB)/colamp-MYB99-DAP-Seq(GSE60143)/Homer | 1e-3 | -9.089e+00 | 0.0010 | 62.0 | 33.16% | 228.8 | 20.37% | motif file (matrix) | svg |
| 116 | | ZNF189(Zf)/HEK293-ZNF189.GFP-ChIP-Seq(GSE58341)/Homer | 1e-3 | -8.957e+00 | 0.0011 | 162.0 | 86.63% | 838.8 | 74.65% | motif file (matrix) | svg |
| 117 | | FOXA1(Forkhead)/LNCAP-FOXA1-ChIP-Seq(GSE27824)/Homer | 1e-3 | -8.941e+00 | 0.0011 | 67.0 | 35.83% | 255.4 | 22.73% | motif file (matrix) | svg |
| 118 | | PU.1-IRF(ETS:IRF)/Bcell-PU.1-ChIP-Seq(GSE21512)/Homer | 1e-3 | -8.937e+00 | 0.0011 | 168.0 | 89.84% | 884.1 | 78.68% | motif file (matrix) | svg |
| 119 | | TRPS1(Zf)/MCF7-TRPS1-ChIP-Seq(GSE107013)/Homer | 1e-3 | -8.937e+00 | 0.0011 | 168.0 | 89.84% | 884.6 | 78.73% | motif file (matrix) | svg |
| 120 | | BIM2(bHLH)/col-BIM2-DAP-Seq(GSE60143)/Homer | 1e-3 | -8.882e+00 | 0.0012 | 160.0 | 85.56% | 824.2 | 73.36% | motif file (matrix) | svg |
| 121 | | Gata1(Zf)/K562-GATA1-ChIP-Seq(GSE18829)/Homer | 1e-3 | -8.852e+00 | 0.0012 | 61.0 | 32.62% | 225.4 | 20.07% | motif file (matrix) | svg |
| 122 | | MYB55(MYB)/colamp-MYB55-DAP-Seq(GSE60143)/Homer | 1e-3 | -8.852e+00 | 0.0012 | 61.0 | 32.62% | 225.0 | 20.03% | motif file (matrix) | svg |
| 123 | | Prop1(Homeobox)/GHFT1-PROP1.biotin-ChIP-Seq(GSE77302)/Homer | 1e-3 | -8.847e+00 | 0.0012 | 136.0 | 72.73% | 659.8 | 58.73% | motif file (matrix) | svg |
| 124 | | MYB73(MYB)/col-MYB73-DAP-Seq(GSE60143)/Homer | 1e-3 | -8.821e+00 | 0.0012 | 165.0 | 88.24% | 862.8 | 76.79% | motif file (matrix) | svg |
| 125 | | Hoxa10(Homeobox)/ChickenMSG-Hoxa10.Flag-ChIP-Seq(GSE86088)/Homer | 1e-3 | -8.683e+00 | 0.0014 | 138.0 | 73.80% | 674.5 | 60.03% | motif file (matrix) | svg |
| 126 | | EWS:FLI1-fusion(ETS)/SK_N_MC-EWS:FLI1-ChIP-Seq(SRA014231)/Homer | 1e-3 | -8.661e+00 | 0.0014 | 11.0 | 5.88% | 13.9 | 1.24% | motif file (matrix) | svg |
| 127 | | MYB58(MYB)/colamp-MYB58-DAP-Seq(GSE60143)/Homer | 1e-3 | -8.627e+00 | 0.0014 | 61.0 | 32.62% | 227.2 | 20.23% | motif file (matrix) | svg |
| 128 | | LEF1(HMG)/H1-LEF1-ChIP-Seq(GSE64758)/Homer | 1e-3 | -8.608e+00 | 0.0014 | 55.0 | 29.41% | 197.3 | 17.56% | motif file (matrix) | svg |
| 129 | | PIF5ox(bHLH)/Arabidopsis-PIF5ox-ChIP-Seq(GSE35062)/Homer | 1e-3 | -8.587e+00 | 0.0015 | 154.0 | 82.35% | 784.2 | 69.80% | motif file (matrix) | svg |
| 130 | | AT3G10030(Trihelix)/colamp-AT3G10030-DAP-Seq(GSE60143)/Homer | 1e-3 | -8.578e+00 | 0.0015 | 129.0 | 68.98% | 617.0 | 54.92% | motif file (matrix) | svg |
| 131 | | Ptf1a(bHLH)/Panc1-Ptf1a-ChIP-Seq(GSE47459)/Homer | 1e-3 | -8.520e+00 | 0.0015 | 169.0 | 90.37% | 896.8 | 79.82% | motif file (matrix) | svg |
| 132 | | MYB92(MYB)/colamp-MYB92-DAP-Seq(GSE60143)/Homer | 1e-3 | -8.517e+00 | 0.0015 | 61.0 | 32.62% | 228.9 | 20.38% | motif file (matrix) | svg |
| 133 | | ANAC042(NAC)/col-ANAC042-DAP-Seq(GSE60143)/Homer | 1e-3 | -8.503e+00 | 0.0015 | 53.0 | 28.34% | 188.5 | 16.77% | motif file (matrix) | svg |
| 134 | | MYB83(MYB)/colamp-MYB83-DAP-Seq(GSE60143)/Homer | 1e-3 | -8.471e+00 | 0.0016 | 65.0 | 34.76% | 249.3 | 22.19% | motif file (matrix) | svg |
| 135 | | Pitx1(Homeobox)/Chicken-Pitx1-ChIP-Seq(GSE38910)/Homer | 1e-3 | -8.455e+00 | 0.0016 | 176.0 | 94.12% | 954.5 | 84.95% | motif file (matrix) | svg |
| 136 | | Tcf3(HMG)/mES-Tcf3-ChIP-Seq(GSE11724)/Homer | 1e-3 | -8.393e+00 | 0.0017 | 52.0 | 27.81% | 184.3 | 16.41% | motif file (matrix) | svg |
| 137 | | NPAS2(bHLH)/Liver-NPAS2-ChIP-Seq(GSE39860)/Homer | 1e-3 | -8.389e+00 | 0.0017 | 162.0 | 86.63% | 844.2 | 75.14% | motif file (matrix) | svg |
| 138 | | Tbr1(T-box)/Cortex-Tbr1-ChIP-Seq(GSE71384)/Homer | 1e-3 | -8.295e+00 | 0.0018 | 128.0 | 68.45% | 614.2 | 54.67% | motif file (matrix) | svg |
| 139 | | OBP4(C2C2dof)/col-OBP4-DAP-Seq(GSE60143)/Homer | 1e-3 | -8.286e+00 | 0.0018 | 78.0 | 41.71% | 320.6 | 28.54% | motif file (matrix) | svg |
| 140 | | Gata4(Zf)/Heart-Gata4-ChIP-Seq(GSE35151)/Homer | 1e-3 | -8.242e+00 | 0.0019 | 163.0 | 87.17% | 853.6 | 75.98% | motif file (matrix) | svg |
| 141 | | Gata2(Zf)/K562-GATA2-ChIP-Seq(GSE18829)/Homer | 1e-3 | -8.205e+00 | 0.0019 | 95.0 | 50.80% | 416.1 | 37.04% | motif file (matrix) | svg |
| 142 | | NPAS(bHLH)/Liver-NPAS-ChIP-Seq(GSE39860)/Homer | 1e-3 | -8.169e+00 | 0.0020 | 161.0 | 86.10% | 839.7 | 74.74% | motif file (matrix) | svg |
| 143 | | MYB93(MYB)/colamp-MYB93-DAP-Seq(GSE60143)/Homer | 1e-3 | -8.136e+00 | 0.0021 | 64.0 | 34.22% | 247.9 | 22.06% | motif file (matrix) | svg |
| 144 | | Bcl6(Zf)/Liver-Bcl6-ChIP-Seq(GSE31578)/Homer | 1e-3 | -7.995e+00 | 0.0024 | 37.0 | 19.79% | 116.2 | 10.34% | motif file (matrix) | svg |
| 145 | | Tcf4(HMG)/Hct116-Tcf4-ChIP-Seq(SRA012054)/Homer | 1e-3 | -7.923e+00 | 0.0025 | 52.0 | 27.81% | 188.2 | 16.75% | motif file (matrix) | svg |
| 146 | | GATA15(C2C2gata)/col-GATA15-DAP-Seq(GSE60143)/Homer | 1e-3 | -7.781e+00 | 0.0029 | 163.0 | 87.17% | 858.3 | 76.39% | motif file (matrix) | svg |
| 147 | | GAGA-repeat/SacCer-Promoters/Homer | 1e-3 | -7.717e+00 | 0.0030 | 170.0 | 90.91% | 912.0 | 81.18% | motif file (matrix) | svg |
| 148 | | AT5G05550(Trihelix)/col-AT5G05550-DAP-Seq(GSE60143)/Homer | 1e-3 | -7.679e+00 | 0.0031 | 30.0 | 16.04% | 87.4 | 7.78% | motif file (matrix) | svg |
| 149 | | Gata6(Zf)/HUG1N-GATA6-ChIP-Seq(GSE51936)/Homer | 1e-3 | -7.606e+00 | 0.0034 | 95.0 | 50.80% | 423.8 | 37.72% | motif file (matrix) | svg |
| 150 | | Initiator/Drosophila-Promoters/Homer | 1e-3 | -7.454e+00 | 0.0039 | 146.0 | 78.07% | 742.4 | 66.08% | motif file (matrix) | svg |
| 151 | | SeqBias: CA-repeat | 1e-3 | -7.450e+00 | 0.0039 | 175.0 | 93.58% | 955.4 | 85.03% | motif file (matrix) | svg |
| 152 | | At3g12730(G2like)/colamp-At3g12730-DAP-Seq(GSE60143)/Homer | 1e-3 | -7.436e+00 | 0.0039 | 117.0 | 62.57% | 556.0 | 49.49% | motif file (matrix) | svg |
| 153 | | TATA-Box(TBP)/Promoter/Homer | 1e-3 | -7.413e+00 | 0.0040 | 79.0 | 42.25% | 335.5 | 29.86% | motif file (matrix) | svg |
| 154 | | Max(bHLH)/K562-Max-ChIP-Seq(GSE31477)/Homer | 1e-3 | -7.310e+00 | 0.0044 | 150.0 | 80.21% | 771.6 | 68.67% | motif file (matrix) | svg |
| 155 | | MYB62(MYB)/colamp-MYB62-DAP-Seq(GSE60143)/Homer | 1e-3 | -7.309e+00 | 0.0044 | 63.0 | 33.69% | 250.1 | 22.26% | motif file (matrix) | svg |
| 156 | | LBD18(LOBAS2)/colamp-LBD18-DAP-Seq(GSE60143)/Homer | 1e-3 | -7.288e+00 | 0.0044 | 153.0 | 81.82% | 792.4 | 70.53% | motif file (matrix) | svg |
| 157 | | Sox6(HMG)/Myotubes-Sox6-ChIP-Seq(GSE32627)/Homer | 1e-3 | -7.188e+00 | 0.0048 | 161.0 | 86.10% | 850.3 | 75.68% | motif file (matrix) | svg |
| 158 | | PCF/Arabidopsis-Promoters/Homer | 1e-3 | -7.178e+00 | 0.0049 | 17.0 | 9.09% | 37.0 | 3.30% | motif file (matrix) | svg |
| 159 | | LHY(Myb)/Seedling-LHY-ChIP-Seq(GSE52175)/Homer | 1e-3 | -7.134e+00 | 0.0050 | 35.0 | 18.72% | 113.9 | 10.14% | motif file (matrix) | svg |
| 160 | | TCF4(bHLH)/SHSY5Y-TCF4-ChIP-Seq(GSE96915)/Homer | 1e-3 | -7.038e+00 | 0.0055 | 155.0 | 82.89% | 809.8 | 72.08% | motif file (matrix) | svg |
| 161 | | PIF4(bHLH)/Seedling-PIF4-ChIP-Seq(GSE35315)/Homer | 1e-3 | -7.015e+00 | 0.0056 | 148.0 | 79.14% | 761.9 | 67.81% | motif file (matrix) | svg |
| 162 | | HIC1(Zf)/Treg-ZBTB29-ChIP-Seq(GSE99889)/Homer | 1e-3 | -6.997e+00 | 0.0057 | 71.0 | 37.97% | 296.6 | 26.40% | motif file (matrix) | svg |
| 163 | | TEAD(TEA)/Fibroblast-PU.1-ChIP-Seq(Unpublished)/Homer | 1e-3 | -6.977e+00 | 0.0058 | 140.0 | 74.87% | 708.7 | 63.08% | motif file (matrix) | svg |
| 164 | | PRDM1(Zf)/Hela-PRDM1-ChIP-Seq(GSE31477)/Homer | 1e-2 | -6.863e+00 | 0.0064 | 22.0 | 11.76% | 58.9 | 5.24% | motif file (matrix) | svg |
| 165 | | MYB67(MYB)/col-MYB67-DAP-Seq(GSE60143)/Homer | 1e-2 | -6.861e+00 | 0.0064 | 64.0 | 34.22% | 260.6 | 23.19% | motif file (matrix) | svg |
| 166 | | Atoh1(bHLH)/Cerebellum-Atoh1-ChIP-Seq(GSE22111)/Homer | 1e-2 | -6.841e+00 | 0.0065 | 149.0 | 79.68% | 770.5 | 68.57% | motif file (matrix) | svg |
| 167 | | RBFox2(?)/Heart-RBFox2-CLIP-Seq(GSE57926)/Homer | 1e-2 | -6.828e+00 | 0.0065 | 160.0 | 85.56% | 847.4 | 75.43% | motif file (matrix) | svg |
| 168 | | ZEB1(Zf)/PDAC-ZEB1-ChIP-Seq(GSE64557)/Homer | 1e-2 | -6.792e+00 | 0.0067 | 155.0 | 82.89% | 812.0 | 72.27% | motif file (matrix) | svg |
| 169 | | Nkx2.1(Homeobox)/LungAC-Nkx2.1-ChIP-Seq(GSE43252)/Homer | 1e-2 | -6.779e+00 | 0.0068 | 167.0 | 89.30% | 899.1 | 80.02% | motif file (matrix) | svg |
| 170 | | OBP3(C2C2dof)/col-OBP3-DAP-Seq(GSE60143)/Homer | 1e-2 | -6.716e+00 | 0.0072 | 141.0 | 75.40% | 718.9 | 63.99% | motif file (matrix) | svg |
| 171 | | NLP7(RWPRK)/col-NLP7-DAP-Seq(GSE60143)/Homer | 1e-2 | -6.715e+00 | 0.0072 | 157.0 | 83.96% | 827.3 | 73.63% | motif file (matrix) | svg |
| 172 | | ARF2(ARF)/col-ARF2-DAP-Seq(GSE60143)/Homer | 1e-2 | -6.650e+00 | 0.0076 | 168.0 | 89.84% | 908.9 | 80.90% | motif file (matrix) | svg |
| 173 | | EBF1(EBF)/Near-E2A-ChIP-Seq(GSE21512)/Homer | 1e-2 | -6.622e+00 | 0.0077 | 148.0 | 79.14% | 766.0 | 68.18% | motif file (matrix) | svg |
| 174 | | MYB105(MYB)/colamp-MYB105-DAP-Seq(GSE60143)/Homer | 1e-2 | -6.609e+00 | 0.0078 | 123.0 | 65.78% | 604.1 | 53.77% | motif file (matrix) | svg |
| 175 | | Tcf7(HMG)/GM12878-TCF7-ChIP-Seq(Encode)/Homer | 1e-2 | -6.605e+00 | 0.0078 | 57.0 | 30.48% | 226.9 | 20.19% | motif file (matrix) | svg |
| 176 | | TCP1(TCP)/col-TCP1-DAP-Seq(GSE60143)/Homer | 1e-2 | -6.574e+00 | 0.0080 | 60.0 | 32.09% | 242.1 | 21.54% | motif file (matrix) | svg |
| 177 | | Olig2(bHLH)/Neuron-Olig2-ChIP-Seq(GSE30882)/Homer | 1e-2 | -6.551e+00 | 0.0081 | 155.0 | 82.89% | 815.9 | 72.62% | motif file (matrix) | svg |
| 178 | | MYB33(MYB)/col-MYB33-DAP-Seq(GSE60143)/Homer | 1e-2 | -6.521e+00 | 0.0083 | 140.0 | 74.87% | 714.7 | 63.61% | motif file (matrix) | svg |
| 179 | | Foxo1(Forkhead)/RAW-Foxo1-ChIP-Seq(Fan_et_al.)/Homer | 1e-2 | -6.427e+00 | 0.0091 | 161.0 | 86.10% | 859.2 | 76.47% | motif file (matrix) | svg |
| 180 | | TEAD4(TEA)/Tropoblast-Tead4-ChIP-Seq(GSE37350)/Homer | 1e-2 | -6.405e+00 | 0.0092 | 139.0 | 74.33% | 709.7 | 63.17% | motif file (matrix) | svg |
| 181 | | NeuroD1(bHLH)/Islet-NeuroD1-ChIP-Seq(GSE30298)/Homer | 1e-2 | -6.298e+00 | 0.0102 | 149.0 | 79.68% | 777.5 | 69.20% | motif file (matrix) | svg |
| 182 | | HSF3(HSF)/colamp-HSF3-DAP-Seq(GSE60143)/Homer | 1e-2 | -6.295e+00 | 0.0102 | 10.0 | 5.35% | 16.0 | 1.43% | motif file (matrix) | svg |
| 183 | | NGA4(ABI3VP1)/col-NGA4-DAP-Seq(GSE60143)/Homer | 1e-2 | -6.283e+00 | 0.0103 | 150.0 | 80.21% | 784.8 | 69.85% | motif file (matrix) | svg |
| 184 | | bZIP18(bZIP)/colamp-bZIP18-DAP-Seq(GSE60143)/Homer | 1e-2 | -6.268e+00 | 0.0104 | 165.0 | 88.24% | 890.8 | 79.28% | motif file (matrix) | svg |
| 185 | | TEAD3(TEA)/HepG2-TEAD3-ChIP-Seq(Encode)/Homer | 1e-2 | -6.267e+00 | 0.0104 | 141.0 | 75.40% | 724.9 | 64.52% | motif file (matrix) | svg |
| 186 | | Ascl1(bHLH)/NeuralTubes-Ascl1-ChIP-Seq(GSE55840)/Homer | 1e-2 | -6.257e+00 | 0.0104 | 152.0 | 81.28% | 798.6 | 71.08% | motif file (matrix) | svg |
| 187 | | MITF(bHLH)/MastCells-MITF-ChIP-Seq(GSE48085)/Homer | 1e-2 | -6.248e+00 | 0.0104 | 153.0 | 81.82% | 805.4 | 71.68% | motif file (matrix) | svg |
| 188 | | At4g38000(C2C2dof)/col-At4g38000-DAP-Seq(GSE60143)/Homer | 1e-2 | -6.192e+00 | 0.0110 | 46.0 | 24.60% | 174.7 | 15.55% | motif file (matrix) | svg |
| 189 | | Zic(Zf)/Cerebellum-ZIC1.2-ChIP-Seq(GSE60731)/Homer | 1e-2 | -6.063e+00 | 0.0124 | 30.0 | 16.04% | 98.4 | 8.76% | motif file (matrix) | svg |
| 190 | | At1g25550(G2like)/colamp-At1g25550-DAP-Seq(GSE60143)/Homer | 1e-2 | -6.045e+00 | 0.0125 | 109.0 | 58.29% | 526.6 | 46.87% | motif file (matrix) | svg |
| 191 | | PRDM9(Zf)/Testis-DMC1-ChIP-Seq(GSE35498)/Homer | 1e-2 | -5.979e+00 | 0.0133 | 7.0 | 3.74% | 8.6 | 0.76% | motif file (matrix) | svg |
| 192 | | KLF3(Zf)/MEF-Klf3-ChIP-Seq(GSE44748)/Homer | 1e-2 | -5.963e+00 | 0.0135 | 10.0 | 5.35% | 17.1 | 1.52% | motif file (matrix) | svg |
| 193 | | CEBP(bZIP)/ThioMac-CEBPb-ChIP-Seq(GSE21512)/Homer | 1e-2 | -5.937e+00 | 0.0138 | 145.0 | 77.54% | 755.6 | 67.25% | motif file (matrix) | svg |
| 194 | | TEAD1(TEAD)/HepG2-TEAD1-ChIP-Seq(Encode)/Homer | 1e-2 | -5.936e+00 | 0.0138 | 140.0 | 74.87% | 722.5 | 64.31% | motif file (matrix) | svg |
| 195 | | Twist2(bHLH)/Myoblast-Twist2.Ty1-ChIP-Seq(GSE127998)/Homer | 1e-2 | -5.859e+00 | 0.0147 | 154.0 | 82.35% | 817.2 | 72.73% | motif file (matrix) | svg |
| 196 | | At5g04390(C2H2)/col200-At5g04390-DAP-Seq(GSE60143)/Homer | 1e-2 | -5.846e+00 | 0.0148 | 143.0 | 76.47% | 743.9 | 66.21% | motif file (matrix) | svg |
| 197 | | ATY13(MYB)/col-ATY13-DAP-Seq(GSE60143)/Homer | 1e-2 | -5.810e+00 | 0.0153 | 163.0 | 87.17% | 881.1 | 78.42% | motif file (matrix) | svg |
| 198 | | ELT-3(Gata)/cElegans-L1-ELT3-ChIP-Seq(modEncode)/Homer | 1e-2 | -5.806e+00 | 0.0153 | 54.0 | 28.88% | 219.5 | 19.54% | motif file (matrix) | svg |
| 199 | | GLIS3(Zf)/Thyroid-Glis3.GFP-ChIP-Seq(GSE103297)/Homer | 1e-2 | -5.793e+00 | 0.0154 | 153.0 | 81.82% | 811.3 | 72.21% | motif file (matrix) | svg |
| 200 | | At5g62940(C2C2dof)/col-At5g62940-DAP-Seq(GSE60143)/Homer | 1e-2 | -5.770e+00 | 0.0157 | 165.0 | 88.24% | 896.4 | 79.78% | motif file (matrix) | svg |
| 201 | | AP-2gamma(AP2)/MCF7-TFAP2C-ChIP-Seq(GSE21234)/Homer | 1e-2 | -5.744e+00 | 0.0160 | 126.0 | 67.38% | 635.2 | 56.53% | motif file (matrix) | svg |
| 202 | | bZIP48(bZIP)/colamp-bZIP48-DAP-Seq(GSE60143)/Homer | 1e-2 | -5.743e+00 | 0.0160 | 91.0 | 48.66% | 424.3 | 37.76% | motif file (matrix) | svg |
| 203 | | GBF6(bZIP)/colamp-GBF6-DAP-Seq(GSE60143)/Homer | 1e-2 | -5.743e+00 | 0.0160 | 91.0 | 48.66% | 424.5 | 37.78% | motif file (matrix) | svg |
| 204 | | NeuroG2(bHLH)/Fibroblast-NeuroG2-ChIP-Seq(GSE75910)/Homer | 1e-2 | -5.709e+00 | 0.0164 | 154.0 | 82.35% | 819.2 | 72.91% | motif file (matrix) | svg |
| 205 | | STAT6(Stat)/Macrophage-Stat6-ChIP-Seq(GSE38377)/Homer | 1e-2 | -5.676e+00 | 0.0168 | 6.0 | 3.21% | 6.3 | 0.56% | motif file (matrix) | svg |
| 206 | | AT1G69570(C2C2dof)/col-AT1G69570-DAP-Seq(GSE60143)/Homer | 1e-2 | -5.651e+00 | 0.0172 | 55.0 | 29.41% | 227.0 | 20.20% | motif file (matrix) | svg |
| 207 | | GATA14(C2C2gata)/col-GATA14-DAP-Seq(GSE60143)/Homer | 1e-2 | -5.634e+00 | 0.0174 | 103.0 | 55.08% | 496.9 | 44.23% | motif file (matrix) | svg |
| 208 | | Cbf1(bHLH)/Yeast-Cbf1-ChIP-Seq(GSE29506)/Homer | 1e-2 | -5.632e+00 | 0.0174 | 94.0 | 50.27% | 443.2 | 39.45% | motif file (matrix) | svg |
| 209 | | MYB77(MYB)/col-MYB77-DAP-Seq(GSE60143)/Homer | 1e-2 | -5.537e+00 | 0.0190 | 144.0 | 77.01% | 754.2 | 67.13% | motif file (matrix) | svg |
| 210 | | Slug(Zf)/Mesoderm-Snai2-ChIP-Seq(GSE61475)/Homer | 1e-2 | -5.513e+00 | 0.0193 | 113.0 | 60.43% | 558.4 | 49.70% | motif file (matrix) | svg |
| 211 | | AT2G28810(C2C2dof)/colamp-AT2G28810-DAP-Seq(GSE60143)/Homer | 1e-2 | -5.483e+00 | 0.0198 | 137.0 | 73.26% | 709.2 | 63.12% | motif file (matrix) | svg |
| 212 | | CLOCK(bHLH)/Liver-Clock-ChIP-Seq(GSE39860)/Homer | 1e-2 | -5.431e+00 | 0.0208 | 103.0 | 55.08% | 499.7 | 44.47% | motif file (matrix) | svg |
| 213 | | HY5(bZIP)/colamp-HY5-DAP-Seq(GSE60143)/Homer | 1e-2 | -5.397e+00 | 0.0214 | 97.0 | 51.87% | 464.5 | 41.34% | motif file (matrix) | svg |
| 214 | | THRb(NR)/Liver-NR1A2-ChIP-Seq(GSE52613)/Homer | 1e-2 | -5.396e+00 | 0.0214 | 161.0 | 86.10% | 872.1 | 77.62% | motif file (matrix) | svg |
| 215 | | ZBTB18(Zf)/HEK293-ZBTB18.GFP-ChIP-Seq(GSE58341)/Homer | 1e-2 | -5.392e+00 | 0.0214 | 12.0 | 6.42% | 26.3 | 2.34% | motif file (matrix) | svg |
| 216 | | Gfi1b(Zf)/HPC7-Gfi1b-ChIP-Seq(GSE22178)/Homer | 1e-2 | -5.355e+00 | 0.0220 | 135.0 | 72.19% | 698.9 | 62.21% | motif file (matrix) | svg |
| 217 | | Tal1 | 1e-2 | -5.314e+00 | 0.0228 | 151.0 | 80.75% | 804.4 | 71.59% | motif file (matrix) | svg |
| 218 | | Fli1(ETS)/CD8-FLI-ChIP-Seq(GSE20898)/Homer | 1e-2 | -5.301e+00 | 0.0230 | 63.0 | 33.69% | 273.4 | 24.34% | motif file (matrix) | svg |
| 219 | | Hand2(bHLH)/Mesoderm-Hand2-ChIP-Seq(GSE61475)/Homer | 1e-2 | -5.292e+00 | 0.0231 | 16.0 | 8.56% | 42.1 | 3.75% | motif file (matrix) | svg |
| 220 | | Unknown-ESC-element(?)/mES-Nanog-ChIP-Seq(GSE11724)/Homer | 1e-2 | -5.285e+00 | 0.0232 | 25.0 | 13.37% | 81.1 | 7.22% | motif file (matrix) | svg |
| 221 | | AP-2alpha(AP2)/Hela-AP2alpha-ChIP-Seq(GSE31477)/Homer | 1e-2 | -5.277e+00 | 0.0232 | 122.0 | 65.24% | 617.5 | 54.96% | motif file (matrix) | svg |
| 222 | | MYB13(MYB)/col-MYB13-DAP-Seq(GSE60143)/Homer | 1e-2 | -5.254e+00 | 0.0237 | 48.0 | 25.67% | 194.0 | 17.27% | motif file (matrix) | svg |
| 223 | | Nanog(Homeobox)/mES-Nanog-ChIP-Seq(GSE11724)/Homer | 1e-2 | -5.245e+00 | 0.0238 | 161.0 | 86.10% | 874.6 | 77.85% | motif file (matrix) | svg |
| 224 | | SeqBias: GA-repeat | 1e-2 | -5.218e+00 | 0.0243 | 175.0 | 93.58% | 978.6 | 87.10% | motif file (matrix) | svg |
| 225 | | Barx1(Homeobox)/Stomach-Barx1.3xFlag-ChIP-Seq(GSE69483)/Homer | 1e-2 | -5.217e+00 | 0.0243 | 71.0 | 37.97% | 318.9 | 28.39% | motif file (matrix) | svg |
| 226 | | IDD5(C2H2)/colamp-IDD5-DAP-Seq(GSE60143)/Homer | 1e-2 | -5.195e+00 | 0.0247 | 134.0 | 71.66% | 694.6 | 61.82% | motif file (matrix) | svg |
| 227 | | MYB96(MYB)/colamp-MYB96-DAP-Seq(GSE60143)/Homer | 1e-2 | -5.183e+00 | 0.0249 | 49.0 | 26.20% | 200.1 | 17.81% | motif file (matrix) | svg |
| 228 | | bHLH34(bHLH)/colamp-bHLH34-DAP-Seq(GSE60143)/Homer | 1e-2 | -5.179e+00 | 0.0249 | 92.0 | 49.20% | 438.4 | 39.02% | motif file (matrix) | svg |
| 229 | | HOXB13(Homeobox)/ProstateTumor-HOXB13-ChIP-Seq(GSE56288)/Homer | 1e-2 | -5.148e+00 | 0.0255 | 141.0 | 75.40% | 740.5 | 65.90% | motif file (matrix) | svg |
| 230 | | AT1G76870(Trihelix)/col-AT1G76870-DAP-Seq(GSE60143)/Homer | 1e-2 | -5.144e+00 | 0.0255 | 116.0 | 62.03% | 582.4 | 51.84% | motif file (matrix) | svg |
| 231 | | PRDM15(Zf)/ESC-Prdm15-ChIP-Seq(GSE73694)/Homer | 1e-2 | -5.143e+00 | 0.0255 | 23.0 | 12.30% | 73.4 | 6.53% | motif file (matrix) | svg |
| 232 | | bHLHE40(bHLH)/HepG2-BHLHE40-ChIP-Seq(GSE31477)/Homer | 1e-2 | -5.132e+00 | 0.0256 | 97.0 | 51.87% | 468.8 | 41.72% | motif file (matrix) | svg |
| 233 | | AIL7(AP2EREBP)/colamp-AIL7-DAP-Seq(GSE60143)/Homer | 1e-2 | -5.127e+00 | 0.0256 | 7.0 | 3.74% | 11.0 | 0.98% | motif file (matrix) | svg |
| 234 | | At4g36780(BZR)/col-At4g36780-DAP-Seq(GSE60143)/Homer | 1e-2 | -5.075e+00 | 0.0269 | 96.0 | 51.34% | 463.9 | 41.29% | motif file (matrix) | svg |
| 235 | | SCL(bHLH)/HPC7-Scl-ChIP-Seq(GSE13511)/Homer | 1e-2 | -5.043e+00 | 0.0276 | 164.0 | 87.70% | 898.1 | 79.94% | motif file (matrix) | svg |
| 236 | | OCT:OCT(POU,Homeobox)/NPC-OCT6-ChIP-Seq(GSE43916)/Homer | 1e-2 | -5.003e+00 | 0.0286 | 125.0 | 66.84% | 640.1 | 56.97% | motif file (matrix) | svg |
| 237 | | SeqBias: GCW-triplet | 1e-2 | -4.993e+00 | 0.0288 | 187.0 | 100.00% | 1091.9 | 97.18% | motif file (matrix) | svg |
| 238 | | SPL9(SBP)/colamp-SPL9-DAP-Seq(GSE60143)/Homer | 1e-2 | -4.920e+00 | 0.0308 | 142.0 | 75.94% | 750.3 | 66.78% | motif file (matrix) | svg |
| 239 | | NFAT(RHD)/Jurkat-NFATC1-ChIP-Seq(Jolma_et_al.)/Homer | 1e-2 | -4.920e+00 | 0.0308 | 43.0 | 22.99% | 172.4 | 15.34% | motif file (matrix) | svg |
| 240 | | AREB3(bZIP)/col-AREB3-DAP-Seq(GSE60143)/Homer | 1e-2 | -4.864e+00 | 0.0322 | 91.0 | 48.66% | 437.0 | 38.90% | motif file (matrix) | svg |
| 241 | | bZIP53(bZIP)/colamp-bZIP53-DAP-Seq(GSE60143)/Homer | 1e-2 | -4.864e+00 | 0.0322 | 91.0 | 48.66% | 437.3 | 38.92% | motif file (matrix) | svg |
| 242 | | AMYB(HTH)/Testes-AMYB-ChIP-Seq(GSE44588)/Homer | 1e-2 | -4.851e+00 | 0.0325 | 150.0 | 80.21% | 804.1 | 71.57% | motif file (matrix) | svg |
| 243 | | HDG1(Homeobox)/col100-HDG1-DAP-Seq(GSE60143)/Homer | 1e-2 | -4.850e+00 | 0.0325 | 127.0 | 67.91% | 655.5 | 58.34% | motif file (matrix) | svg |
| 244 | | Zfp281(Zf)/ES-Zfp281-ChIP-Seq(GSE81042)/Homer | 1e-2 | -4.824e+00 | 0.0331 | 10.0 | 5.35% | 21.3 | 1.89% | motif file (matrix) | svg |
| 245 | | SPL15(SBP)/colamp-SPL15-DAP-Seq(GSE60143)/Homer | 1e-2 | -4.822e+00 | 0.0331 | 141.0 | 75.40% | 745.8 | 66.38% | motif file (matrix) | svg |
| 246 | | At1g78700(BZR)/col-At1g78700-DAP-Seq(GSE60143)/Homer | 1e-2 | -4.812e+00 | 0.0332 | 97.0 | 51.87% | 473.2 | 42.12% | motif file (matrix) | svg |
| 247 | | LBD23(LOBAS2)/colamp-LBD23-DAP-Seq(GSE60143)/Homer | 1e-2 | -4.771e+00 | 0.0345 | 45.0 | 24.06% | 184.2 | 16.39% | motif file (matrix) | svg |
| 248 | | AT1G47655(C2C2dof)/colamp-AT1G47655-DAP-Seq(GSE60143)/Homer | 1e-2 | -4.744e+00 | 0.0353 | 146.0 | 78.07% | 779.3 | 69.36% | motif file (matrix) | svg |
| 249 | | ZNF165(Zf)/WHIM12-ZNF165-ChIP-Seq(GSE65937)/Homer | 1e-2 | -4.740e+00 | 0.0353 | 48.0 | 25.67% | 200.1 | 17.81% | motif file (matrix) | svg |
| 250 | | SRS7(SRS)/colamp-SRS7-DAP-Seq(GSE60143)/Homer | 1e-2 | -4.715e+00 | 0.0361 | 154.0 | 82.35% | 833.5 | 74.18% | motif file (matrix) | svg |
| 251 | | c-Myc(bHLH)/LNCAP-cMyc-ChIP-Seq(Unpublished)/Homer | 1e-2 | -4.688e+00 | 0.0369 | 97.0 | 51.87% | 475.2 | 42.29% | motif file (matrix) | svg |
| 252 | | bZIP3(bZIP)/col-bZIP3-DAP-Seq(GSE60143)/Homer | 1e-2 | -4.647e+00 | 0.0383 | 94.0 | 50.27% | 458.8 | 40.83% | motif file (matrix) | svg |
| 253 | | ETV4(ETS)/HepG2-ETV4-ChIP-Seq(ENCODE)/Homer | 1e-2 | -4.635e+00 | 0.0386 | 61.0 | 32.62% | 271.6 | 24.18% | motif file (matrix) | svg |
| 254 | | KLF6(Zf)/PDAC-KLF6-ChIP-Seq(GSE64557)/Homer | 1e-2 | -4.614e+00 | 0.0393 | 152.0 | 81.28% | 821.6 | 73.12% | motif file (matrix) | svg |
| 255 | | TF3A(C2H2)/col-TF3A-DAP-Seq(GSE60143)/Homer | 1e-2 | -4.606e+00 | 0.0394 | 144.0 | 77.01% | 768.3 | 68.38% | motif file (matrix) | svg |

Appendix File S4 - Normalized expression values (TPM) of expressed genes near accessible human-specific SVAs and LTR5HS

| GENE | NOG1 | NOG2 | NOG3 | HSTE1 | HSTE3 | CHIMP |
|---|---|---|---|---|---|---|
| STOM | 338.6927 | 368.0246 | 277.11236 | 26.81615 | 95.6402 | 4.306915 |
| TRIML2 | 153.02222 | 461.54504 | 65.643892 | 1.289704 | 7.505061 | 10.3745705 |
| SNX2 | 204.567731 | 179.112639 | 180.325473 | 142.002262 | 68.754442 | 69.185645 |
| DERA | 96.489615 | 100.435735 | 99.103987 | 25.48269911 | 24.13401039 | 54.857585 |
| SLC3A2 | 218.087728 | 343.90514 | 225.674258 | 150.546958 | 155.438003 | 197.1565031 |
| PRODH | 39.957813 | 30.426749 | 51.1103 | 11.527249 | 0.959688 | 6.638564 |
| TIMM50 | 144.7896887 | 115.610233 | 136.865572 | 117.0986777 | 94.561327 | 220.3145651 |
| ARMT1 | 65.431614 | 65.169659 | 62.905177 | 47.447522 | 29.275061 | 13.910845 |
| ZNF90 | 28.459687 | 52.096173 | 43.362305 | 16.319451 | 17.142462 | 14.8092265 |
| SDHC | 199.666242 | 206.866744 | 214.964646 | 186.831027 | 174.13002 | 123.9574185 |
| PFKP | 61.768106 | 56.026419 | 70.97457 | 41.518914 | 30.6643281 | 19.173705 |
| RIC8A | 65.103424 | 65.276123 | 59.552289 | 53.634338 | 25.5079183 | 66.23072 |
| EIF3K | 176.162851 | 206.634786 | 203.6151731 | 152.838598 | 205.116368 | 161.60682 |
| DNAJC15 | 58.5008 | 62.8475 | 41.7985 | 32.86735 | 36.1469 | 10.66295 |
| ENSA | 174.0938671 | 156.355251 | 118.194654 | 163.9781 | 107.516553 | 136.083875 |
| DENR | 74.75354 | 86.06632 | 91.56546 | 65.953996 | 66.549556 | 71.646755 |
| RPUSD3 | 63.6247731 | 60.982965 | 55.485307 | 44.97777402 | 38.882625 | 105.248655 |
| EIF3B | 150.939238 | 127.601903 | 172.761717 | 173.296855 | 91.6097411 | 167.60323 |
| NCBP2 | 89.839008 | 102.8171503 | 118.656183 | 95.373694 | 75.861608 | 120.246667 |
| MRS2 | 38.234056 | 48.860257 | 37.364138 | 22.354897 | 20.578914 | 19.881795 |
| MSTO1 | 35.20645668 | 40.597738 | 53.088307 | 40.3735072 | 9.9291152 | 62.4511714 |
| ZNF649 | 30.876404 | 30.49426 | 35.018718 | 22.756772 | 9.752741 | 10.628018 |
| NSMAF | 23.242473 | 30.574494 | 31.360044 | 23.4800983 | 9.1598092 | 20.68682 |
| RANBP2 | 20.359826 | 29.267606 | 33.545678 | 22.726388 | 8.78899 | 19.49315 |
| WBP1L | 24.0072491 | 26.341471 | 20.1150606 | 13.6780462 | 8.2647749 | 11.20178 |
| ABCB8 | 24.815917 | 31.5282159 | 22.006623 | 22.4052658 | 4.9117027 | 23.15368 |
| AGL | 14.326482 | 18.18789356 | 19.62175401 | 8.933412007 | 6.06659862 | 3.98568825 |
| KLRG2 | 17.811217 | 14.0096003 | 14.185828 | 5.615728 | 1.87773 | 2.62703 |
| SNX8 | 47.368743 | 50.37898222 | 46.789568 | 35.68902777 | 38.442986 | 42.028195 |
| APELA | 17.856143 | 21.356786 | 17.829138 | 10.91944 | 5.363467 | 2.67813 |
| SMNDC1 | 48.188759 | 42.227491 | 39.42996 | 48.655113 | 17.144826 | 20.808525 |
| VAV2 | 25.701086 | 21.773997 | 22.604485 | 18.93218 | 7.80209 | 28.72027 |
| PLSCR1 | 19.512552 | 31.5374371 | 24.667185 | 14.831594 | 16.0877152 | 26.535545 |
| MMP24OS | 56.1805157 | 45.387626 | 40.2331894 | 28.811673 | 43.801148 | 16.720215 |
| ASRGL1 | 30.743948 | 27.087235 | 35.327964 | 29.84702 | 13.633117 | 10.614892 |
| TMCC1 | 19.757471 | 18.980254 | 16.536868 | 13.925197 | 6.229255 | 15.366399 |
| ENTPD6 | 47.396488 | 35.3858473 | 39.4894985 | 42.637939 | 20.27688 | 23.07206383 |
| USP33 | 44.346545 | 43.1813537 | 46.751208 | 41.710167 | 31.10541473 | 54.889545 |
| PSMC4 | 38.7784701 | 45.414263 | 39.628651 | 35.6186211 | 27.946043 | 51.476805 |
| MRTFA | 14.09309881 | 13.54868719 | 12.5384693 | 8.790056006 | 3.9175425 | 23.20533977 |
| SLC35A5 | 37.511289 | 35.6301913 | 41.150228 | 45.253766 | 15.581974 | 14.00149 |
| RBM8A | 204.011987 | 214.437312 | 198.613366 | 171.932179 | 222.410714 | 202.55648 |
| HSBP1L1 | 12.977331 | 15.752373 | 14.53834 | 3.09209 | 12.343234 | 12.02412 |
| CR1L | 6.765421 | 7.954092 | 14.267864 | 0.986449 | 1.3441903 | 2.20289 |
| ZNF701 | 6.669049 | 13.309874 | 11.13296 | 4.799059 | 5.444288 | 5.46629 |
| SERPINB9 | 5.74565 | 18.0717 | 7.17799 | 0.832976 | 1.55089 | 2.677875 |
| SLC35F6 | 9.916161 | 13.70873 | 11.451181 | 5.52163 | 6.1206 | 9.3657 |
| NBPF15 | 19.58212 | 16.32336 | 22.146217 | 22.292302 | 6.1811554 | 14.54613 |
| SDHAP3 | 8.719765 | 9.735468 | 11.1349 | 5.573897 | 4.17137 | 24.72334 |
| TRIM38 | 7.83166 | 7.40432 | 8.29185 | 0.952764 | 5.05575 | 1.499745 |
| CHD1L | 44.26924321 | 40.99676575 | 48.43409008 | 56.00145026 | 24.1465319 | 51.30866495 |
| SMIM10L2A | 5.47856 | 4.92294 | 8.28796 | 1.83006 | 0.789673 | 5.4254 |
| CACNA2D2 | 8.755006 | 8.914848 | 11.395641 | 7.0766019 | 2.463349 | 4.513045175 |

| | | | | | | |
|---|---|---|---|---|---|---|
| PYGB | 28.06726 | 20.18493 | 24.45504 | 28.95902 | 11.9501 | 39.32822 |
| MFSD4B | 3.64906 | 7.77279 | 6.58817 | 4.08572 | 1.95594 | 5.11405 |
| TCP11L1 | 9.347057 | 10.60045 | 9.260063 | 8.160111 | 3.487937 | 12.6392739 |
| PACSIN1 | 4.812044 | 4.571797 | 3.490020048 | 1.693120139 | 0.645959 | 12.351625 |
| NBPF11 | 5.5423 | 7.724746 | 7.098247 | 5.069955 | 2.337206601 | 7.41195 |
| HYOU1 | 52.0240003 | 45.88562341 | 55.3934474 | 59.380064 | 37.9811986 | 31.6459175 |
| LDLRAP1 | 7.824388 | 10.77293 | 7.485557 | 6.534545 | 3.247159 | 14.079405 |
| ZNF578 | 2.4910734 | 4.1501445 | 4.4140246 | 1.1706529 | 1.275296 | 3.029755 |
| ECEL1P1 | 5.80991 | 3.94788 | 10.9445 | 0.872093 | 5.51563 | 0.53082 |
| ZNF146 | 83.591126 | 88.977897 | 109.50771 | 122.372425 | 50.883066 | 47.674345 |
| APOBEC3B | 3.887361 | 7.764471 | 4.749639365 | 2.138344 | 2.99294 | 0.82037 |
| KRBA2 | 2.55691 | 3.587704045 | 4.588893504 | 2.157393 | 0.689476 | 2.72993 |
| BARD1 | 6.9039187 | 6.534843 | 8.703234 | 7.280767 | 2.3993502 | 29.58318001 |
| NBPF9 | 8.342214 | 10.17013806 | 13.460654 | 10.53907633 | 5.7425225 | 51.819505 |
| TMEM41B | 18.626564 | 21.025895 | 22.072816 | 15.91073426 | 22.480863 | 18.044994 |
| ALX1 | 3.2578 | 6.5462 | 1.11285 | 1.90434 | 1.34143 | 166.285 |
| TBC1D23 | 15.219766 | 15.4557 | 16.710506 | 16.042746 | 11.666935 | 6.7603305 |
| DSTNP2 | 6.05597 | 5.68299 | 9.3592 | 5.31323 | 3.599638776 | 1.90609 |
| GALNT8 | 3.126064 | 3.372623 | 3.374464 | 1.808728 | 1.7671174 | 4.8583 |
| PUF60 | 42.80258141 | 12.32280264 | 6.150515 | 21.298382 | 0.2107722 | 132.6338496 |
| NBPF8 | 5.95377 | 5.11975 | 7.37674 | 7.25078 | 1.978989 | 1.175195 |
| ZNF517 | 4.1418453 | 4.1336127 | 3.851378 | 4.2235279 | 1.546685 | 13.309685 |
| SEMA4A | 2.278135303 | 2.371563087 | 3.2152212 | 1.801899 | 0.510677 | 9.869059625 |
| PTAR1 | 7.54406 | 9.358883 | 15.60004 | 12.248593 | 4.170151 | 5.620755 |
| NBAS | 23.6191821 | 21.658773 | 22.28398 | 17.694426 | 24.588219 | 30.837455 |
| SPTB | 2.314385386 | 1.142477547 | 1.826061092 | 1.277710214 | 0.366655044 | 4.110609 |
| PEMT | 26.852139 | 20.523603 | 25.22508 | 28.078504 | 20.466607 | 148.01576 |
| FAAH2 | 6.620793 | 8.10587 | 5.23218 | 2.862782 | 8.97191 | 0.54461 |
| ZNF527 | 2.60131 | 2.450193 | 2.069585 | 2.682205 | 1.155209 | 6.277085 |
| ZNF565 | 1.746892 | 3.104125 | 2.187917 | 3.224394 | 1.1894096 | 7.826857 |
| RRP12 | 13.39812192 | 13.7803582 | 15.16177208 | 16.36307831 | 11.495413 | 36.9634586 |
| ERCC4 | 6.87213 | 6.347114 | 9.200773 | 9.229252 | 5.650641 | 8.01779 |
| KCTD7 | 5.638524886 | 4.773298984 | 5.851686 | 9.7521408 | 2.758422 | 8.8501518 |
| CTH | 13.889928 | 15.013404 | 22.874579 | 14.64033 | 16.88297 | 37.00286 |
| DENND2A | 7.993525 | 5.257436 | 10.793876 | 14.346552 | 3.17685039 | 27.87986652 |
| ZNF681 | 5.62617 | 5.403581 | 6.43122 | 9.95632 | 3.220215 | 1.430225 |
| CENPS-CORT | 3.455793 | 1.7741602 | 1.740973 | 4.272103 | 1.455883 | 31.45888524 |
| KIF9 | 6.78734094 | 4.8201299 | 6.674405601 | 7.120935502 | 8.79235 | 8.251171 |
| AK4 | 6.56396 | 6.158176893 | 10.8304685 | 11.40399 | 5.341270603 | 12.91487 |
| DNAJB4 | 16.248492 | 15.548723 | 16.324557 | 28.743235 | 8.134622 | 11.31638 |
| CAAP1 | 13.777195 | 16.825776 | 16.618635 | 23.773887 | 15.508416 | 32.500301 |
| CHORDC1 | 21.2069027 | 25.0355312 | 19.55656404 | 34.489742 | 14.7434365 | 5.1663645 |
| WDHD1 | 26.49974 | 29.515062 | 36.71122 | 38.641325 | 29.11726 | 40.141915 |
| ZNF121 | 25.357659 | 20.268579 | 47.6118 | 40.9347 | 23.477414 | 16.059128 |
| RND2 | 38.535337 | 22.037637 | 36.24485 | 55.34807 | 32.0431 | 11.4875 |
| EPHA7 | 4.401424 | 1.1749565 | 2.113967 | 19.32684 | 0.77025 | 1.67942 |
| ZC3H15 | 89.842969 | 71.962311 | 73.091954 | 105.0517143 | 73.924519 | 48.5834 |
| PRDX4 | 119.634272 | 116.925925 | 112.399851 | 131.1674 | 140.529179 | 245.4997 |
| MRPL20 | 124.290014 | 125.49607 | 118.024882 | 123.72759 | 180.236725 | 125.82333 |
| NDUFAB1 | 138.221958 | 112.230689 | 130.711432 | 153.425705 | 239.689892 | 156.3325 |
| UQCRH | 1034.47424 | 939.18506 | 1015.44567 | 1093.65982 | 1668.51567 | 365.430395 |
| LDHB | 1531.518188 | 1366.470509 | 1569.917343 | 2190.457461 | 2443.493007 | 1030.105225 |

Appendix File S5 - Deseq2 output for differential gene expression analysis

| GENE | baseMean | log2FoldChange | lfcSE | stat | pvalue | padj |
|---|---|---|---|---|---|---|
| A1BG | 56.703831 | 1.340071865 | 0.551527357 | 2.429746862 | 0.015109372 | 0.139261407 |
| A2ML1 | 124.0999122 | -2.533497387 | 0.681340148 | -3.718403201 | 0.000200486 | 0.009416639 |
| A4GALT | 219.2342317 | 0.068043372 | 0.419502976 | 0.162199974 | 0.87114838 | 0.948553225 |
| AAAS | 1887.681556 | 0.001220163 | 0.24547076 | 0.004970707 | 0.996033966 | 0.998272751 |
| AACS | 853.016537 | -0.022829751 | 0.307467685 | -0.074250896 | 0.940810748 | 0.97807441 |
| AACSP1 | 16.54639206 | -0.398461008 | 0.842940704 | -0.472703484 | 0.63642474 | 0.841268221 |
| AADAT | 434.9579816 | -0.195274792 | 0.308947518 | -0.632064607 | 0.527344661 | 0.778906646 |
| AAGAB | 1750.078323 | 0.149081741 | 0.268295003 | 0.555663503 | 0.578440911 | 0.810209453 |
| AAK1 | 757.4568219 | -0.273951727 | 0.363471857 | -0.753708222 | 0.451024444 | 0.732208864 |
| AAMDC | 301.709175 | 1.018542827 | 0.493674527 | 2.063186922 | 0.039094872 | 0.227855669 |
| AAMP | 2970.154122 | 0.100008791 | 0.375561728 | 0.266291221 | 0.790014934 | 0.912949942 |
| AAR2 | 915.9785468 | -0.334739261 | 0.250457293 | -1.336512333 | 0.181381872 | 0.485749663 |
| AARD | 43.2057477 | 1.380112213 | 0.608553557 | 2.267856619 | 0.023337947 | 0.174385097 |
| AARS | 3663.821983 | -0.200432009 | 0.296754124 | -0.675414401 | 0.499412519 | 0.76281557 |
| AARS2 | 1016.738427 | -0.383838867 | 0.364512722 | -1.053019124 | 0.292332227 | 0.605539626 |
| AARSD1 | 1085.828932 | 0.418712962 | 0.271112565 | 1.544424774 | 0.122485462 | 0.403241325 |
| AASDH | 594.0589809 | 0.104423025 | 0.265061346 | 0.393957951 | 0.693612084 | 0.868790368 |
| AASDHPPT | 1368.56101 | 0.561016587 | 0.297170362 | 1.887861839 | 0.059044501 | 0.281686831 |
| AASS | 3326.794021 | 0.023511111 | 0.308845666 | 0.076125761 | 0.939319045 | 0.977796709 |
| AATF | 1281.42299 | -0.391477257 | 0.305456302 | -1.2816146 | 0.199977876 | 0.50598138 |
| AATK | 24.77075888 | 2.303026081 | 0.996574787 | 2.310941549 | 0.020836084 | 0.163640591 |
| ABAT | 413.3583367 | -0.143993856 | 0.441635262 | -0.326047009 | 0.744388789 | 0.892725315 |
| ABCA1 | 573.1988106 | -0.454515671 | 0.652207192 | -0.696888468 | 0.485872595 | 0.75593662 |
| ABCA10 | 14.95028154 | 0.080391725 | 0.935721742 | 0.085914136 | 0.931534674 | 0.974341807 |
| ABCA11P | 134.1052779 | -0.261555678 | 0.386474506 | -0.676773433 | 0.498549717 | 0.762775559 |
| ABCA12 | 17.35524499 | -2.396719834 | 0.974719231 | -2.458882269 | 0.013937031 | 0.134161613 |
| ABCA2 | 294.1305269 | 0.796550735 | 0.697124365 | 1.142623576 | 0.253194913 | 0.565800563 |
| ABCA3 | 293.9905685 | 0.714296816 | 0.617110114 | 1.157486809 | 0.247073526 | 0.559706205 |
| ABCA4 | 261.4934349 | -2.798131044 | 0.700457015 | -3.994721995 | 6.48E-05 | 0.004569989 |
| ABCA5 | 137.2377178 | -0.81182921 | 0.427989634 | -1.896843158 | 0.057848641 | 0.278761201 |
| ABCA6 | 14.60756706 | -1.221176568 | 0.937653633 | -1.302374911 | 0.192788255 | 0.498908723 |
| ABCA7 | 433.2263977 | -0.831018981 | 0.3703636 | -2.243792263 | 0.02484577 | 0.180179342 |
| ABCA9 | 12.35261758 | -0.047885108 | 0.938179534 | -0.051040453 | 0.959293286 | 0.985871727 |
| ABCB1 | 16.48166201 | -3.798310196 | 1.788226657 | -2.124065302 | 0.033664683 | 0.209086405 |
| ABCB10 | 526.8999994 | 0.116130114 | 0.256766349 | 0.452279338 | 0.651067759 | 0.847876685 |

| | | | | | | |
|---|---|---|---|---|---|---|
| ABCB4 | 521.4953889 | -4.692752508 | 0.827486489 | -5.671092605 | 1.42E-08 | 1.03E-05 |
| ABCB6 | 401.1543277 | 0.657503548 | 0.373777342 | 1.75907813 | 0.078564239 | 0.321658491 |
| ABCB7 | 1406.209856 | -0.022472715 | 0.245831941 | -0.091414951 | 0.927162882 | 0.972482583 |
| ABCB8 | 796.9402467 | -0.791279674 | 0.436410398 | -1.813154952 | 0.069807926 | 0.303603614 |
| ABCB9 | 102.7209094 | 0.503732108 | 0.413870427 | 1.217125156 | 0.223556601 | 0.53268495 |
| ABCC1 | 826.25249 | -0.122107795 | 0.519972721 | -0.234835001 | 0.8143368 | 0.92237594 |
| ABCC10 | 624.3248976 | -0.540953701 | 0.454014386 | -1.191490221 | 0.233461193 | 0.543310288 |
| ABCC2 | 26.80315627 | -0.10474041 | 0.597331914 | -0.175347084 | 0.860806917 | 0.943659349 |
| ABCC3 | 46.10754794 | 0.63545637 | 0.549836887 | 1.155717969 | 0.247796536 | 0.560507118 |
| ABCC4 | 977.9086366 | -0.844539593 | 0.309330366 | -2.730218835 | 0.00632923 | 0.086470416 |
| ABCC5 | 1491.289011 | -0.555377007 | 0.42800864 | -1.297583636 | 0.194430448 | 0.500298624 |
| ABCC6 | 19.849261 | -1.169353438 | 0.862300854 | -1.356085214 | 0.175072057 | 0.479137654 |
| ABCC6P1 | 44.3134736 | -0.739920601 | 0.717256791 | -1.031597902 | 0.302260523 | 0.613115536 |
| ABCC8 | 43.83051879 | 0.17232489 | 0.883877786 | 0.194964612 | 0.845420651 | 0.937778401 |
| ABCC9 | 69.62340282 | -1.088577967 | 0.447555491 | -2.432274857 | 0.015004316 | 0.139113039 |
| ABCD1 | 96.43291796 | -0.205530031 | 0.534425077 | -0.384581562 | 0.700547457 | 0.872001128 |
| ABCD3 | 1327.431567 | -0.061739016 | 0.323455603 | -0.190873231 | 0.848624913 | 0.938569494 |
| ABCD4 | 1844.729708 | 0.205329319 | 0.230117474 | 0.892280434 | 0.372242639 | 0.677458382 |
| ABCE1 | 3177.34446 | 0.498143783 | 0.253009331 | 1.968875146 | 0.04896743 | 0.254849017 |
| ABCF1 | 1431.124168 | -0.113985267 | 0.327323295 | -0.348234508 | 0.727664074 | 0.88517603 |
| ABCF2 | 1739.703527 | -0.258551592 | 0.250154718 | -1.033566723 | 0.301338763 | 0.612159932 |
| ABCF3 | 1089.395594 | -0.365906153 | 0.246305927 | -1.485575917 | 0.137391329 | 0.425712285 |
| ABCG2 | 1206.57842 | -3.552063204 | 0.83952491 | -4.231039678 | 2.33E-05 | 0.002063732 |
| ABCG4 | 12.25901548 | 0.77239989 | 0.983717308 | 0.785184812 | 0.432345219 | 0.718639059 |
| ABCG8 | 14.47436878 | -0.146890813 | 0.898410337 | -0.163500805 | 0.870124139 | 0.947914212 |
| ABHD10 | 782.4102858 | 1.067984409 | 0.268074752 | 3.983905244 | 6.78E-05 | 0.004653715 |
| ABHD11 | 517.5532652 | -0.256086366 | 0.474365225 | -0.539850631 | 0.589300047 | 0.818256545 |
| ABHD12 | 1715.932794 | -0.144073096 | 0.329681477 | -0.437006949 | 0.662106309 | 0.853787914 |
| ABHD12B | 80.23615803 | -0.484706138 | 0.664856333 | -0.729038913 | 0.46597786 | 0.742755984 |
| ABHD13 | 696.7672853 | 0.219267341 | 0.269908973 | 0.812375145 | 0.416576404 | 0.708268589 |
| ABHD14A | 62.41557166 | 1.270446446 | 0.558345599 | 2.275376485 | 0.022883355 | 0.172718097 |
| ABHD14A-ACY1 | 22.46484871 | -0.820922213 | 0.814014336 | -1.008486186 | 0.313221116 | 0.623018103 |
| ABHD14B | 226.8605005 | 0.54509026 | 0.402811955 | 1.353212717 | 0.175987684 | 0.480071547 |
| ABHD15 | 75.52855219 | 0.057492854 | 0.463460382 | 0.124051282 | 0.901274672 | 0.961878265 |
| ABHD16A | 31.22399939 | 0.966010628 | 0.908673445 | 1.063099877 | 0.28773666 | 0.601673803 |
| ABHD17A | 653.0067432 | -0.127766798 | 0.52724155 | -0.242330671 | | |

0.80852395 0.91963689
ABHD17B 1013.260577 0.246098917 0.229870506 1.070598055 0.284350199
0.598246396
ABHD17C 583.6385403 0.261961538 0.461277513 0.56790442 0.570099871
0.806205765
ABHD18 419.9714163 -0.240006994 0.303249458 -0.791450693
0.428681038 0.715644829
ABHD2 685.6797402 -0.439199874 0.493408763 -0.890133914 0.373393985
0.678294888
ABHD3 398.2355036 0.459289755 0.328728395 1.397170922 0.162362179 0.463093929
ABHD4 533.3781712 -0.944001484 0.396128709 -2.38306758 0.017169042
0.147741524
ABHD5 354.8132785 -0.101568325 0.358920208 -0.282983021 0.777189852
0.907898743
ABHD6 175.7146114 0.476850171 0.424764093 1.122623542 0.261597412 0.575089747
ABHD8 222.0385218 -0.311366318 0.619464864 -0.502637577 0.615219105
0.830958873
ABI1 1653.065186 -0.307603858 0.240551045 -1.278746714 0.200986264
0.506881795
ABI2 4920.654742 0.305907356 0.287313954 1.064714582 0.287005109 0.601236469
ABI3 13.78692898 0.639125026 0.848514877 0.753227838 0.451313013 0.732520566
ABL1 3166.825726 -0.3029751 0.429798628 -0.704923378 0.480857921
0.752563699
ABL2 770.0595333 -0.109613173 0.35053315 -0.312704156 0.754505437
0.897438517
ABLIM1 1343.906761 -0.03736596 0.361384853 -0.103396586 0.917648221
0.969677217
ABLIM2 11.45220551 0.250486673 0.865511768 0.289408743 0.772268605
0.905685717
ABLIM3 62.41553194 -2.313908912 0.704269765 -3.285543448
0.001017858 0.027467362
ABO 12.70810572 -1.102730138 1.038799561 -1.061542746 0.288443318
0.601673803
ABR 990.8790901 -0.544970848 0.432986082 -1.25863364 0.208162691
0.515093162
ABRACL 3289.223793 0.850754341 0.560514475 1.517809761 0.129062364
0.413453061
ABRAXAS1 1574.157749 0.679609754 0.371615544 1.828797976 0.067429878
0.298244514
ABRAXAS2 604.3485853 -0.042817951 0.280474263 -0.152662675
0.878664296 0.951592678
ABT1 749.1786696 0.166955107 0.369821028 0.451448387 0.651666416 0.8478779
ABTB1 189.609768 -0.871854209 0.486926412 -1.790525608 0.073369454
0.311344591
ABTB2 111.6186016 0.422493263 0.367917458 1.148337091 0.250829433 0.563811922
AC000093.1 11.70092756 2.226801624 1.049675351 2.121419371 0.033886527
0.209833671
AC000120.3 77.35520975 -0.610945364 2.025085685 -0.301688649
0.762889422 0.901526361
AC001226.2 92.01520861 -2.061331928 0.660474757 -3.120985179
0.001802471 0.039360945
AC002472.2 21.51816281 -0.111641858 0.677831577 -0.164704422
0.869176636 0.947227711
AC003005.1 13.07377217 0.162887162 1.603937689 0.101554545 0.919110261
0.969677217
AC003072.1 15.64087067 0.568531117 0.815958236 0.696764972 0.485949891
0.75593662
AC003665.1 303.3432403 -0.490390544 0.374834761 -1.308284594
0.190776798 0.496058205
AC004057.1 5215.963896 0.790110612 0.503848844 1.568150094 0.116846108
0.394102335
AC004233.2 19.5809369 -0.663831557 0.872532048 -0.760810514
0.446770253 0.728709125
AC004475.1 1031.292988 -0.327833871 0.289273491 -1.133300776

| | | | | | | |
|---|---|---|---|---|---|---|
| | | | | | 0.257087966 | 0.56963364 |
| AC004687.2 | 22.33164069 | -0.548189855 | 0.774600339 | -0.707706707 | 0.479127411 | 0.751534528 |
| AC004890.2 | 98.29550681 | 0.116802265 | 0.461095354 | 0.253314773 | 0.800024981 | 0.916311991 |
| AC004922.1 | 18.71744539 | -1.209773579 | 0.812193759 | -1.489513512 | 0.136352198 | 0.424384947 |
| AC004951.2 | 22.13544498 | -1.107965322 | 0.710467341 | -1.559488042 | 0.118880913 | 0.397626945 |
| AC004951.3 | 23.39754714 | -0.277288465 | 0.774500185 | -0.358022464 | 0.720326502 | 0.88214152 |
| AC004967.1 | 13.05046294 | -0.305175501 | 0.833604824 | -0.366091333 | 0.714296925 | 0.878457604 |
| AC004980.1 | 22.57125678 | -0.126337711 | 0.632484479 | -0.199748318 | 0.841677423 | 0.935190287 |
| AC004997.1 | 21.55766245 | -3.25843485 | 1.480639748 | -2.200693891 | 0.027757702 | 0.190634717 |
| AC005041.1 | 10.39610063 | -1.53417306 | 1.084537108 | -1.414587891 | 0.157189361 | 0.456574487 |
| AC005077.4 | 14.30663781 | -0.30016039 | 1.084243367 | -0.276838576 | 0.781904059 | 0.909909835 |
| AC005154.2 | 18.19611846 | 0.665583234 | 0.765458903 | 0.86952184 | 0.384561768 | 0.68524625 |
| AC005154.5 | 31.4317852 | 0.40405607 | 0.566764178 | 0.712917444 | 0.475896844 | 0.749091526 |
| AC005324.4 | 81.13976676 | 0.8828197 | 0.998715798 | 0.883954877 | 0.376720572 | 0.68029735 |
| AC005520.1 | 21.74575093 | -0.918059401 | 0.705805717 | -1.300725368 | 0.193352476 | 0.499293687 |
| AC005670.2 | 21.41045896 | 0.186431642 | 1.008376303 | 0.184883006 | 0.8533208 | 0.940799171 |
| AC005726.1 | 140.826447 | 0.564365653 | 0.416193116 | 1.356018712 | 0.175093214 | 0.479137654 |
| AC005832.4 | 24.26283269 | -0.591053395 | 1.416367556 | -0.417302269 | 0.676457329 | 0.860199564 |
| AC005833.1 | 366.0270249 | -0.215291516 | 0.265215529 | -0.811760597 | 0.416929016 | 0.708661116 |
| AC005943.1 | 108.3396673 | 0.359919374 | 3.981320689 | 0.090402005 | 0.927967763 | 0.972633911 |
| AC006001.3 | 528.8238359 | -0.218588502 | 0.301512292 | -0.724973769 | 0.468468123 | 0.744337782 |
| AC006001.4 | 11.96499845 | 3.251684494 | 1.76975416 | 1.837365079 | 0.066156018 | 0.296938744 |
| AC006011.1 | 44.2859064 | 2.339314847 | 1.108853328 | 2.109670223 | 0.034886771 | 0.213176891 |
| AC006030.1 | 14.21495345 | 0.244078936 | 1.153452039 | 0.211607356 | 0.83241337 | 0.931375477 |
| AC006064.6 | 159.0760796 | -0.340857471 | 2.272789621 | -0.149973173 | 0.880785781 | 0.952620024 |
| AC006213.1 | 56.37947436 | -0.947162643 | 0.714467005 | -1.325691231 | 0.184941993 | 0.489406194 |
| AC006254.1 | 98.16227472 | -0.348819861 | 0.834801887 | -0.417847475 | 0.676058637 | 0.860110127 |
| AC006453.2 | 22.64889952 | 0.76080144 | 0.800589547 | 0.950301491 | 0.341959081 | 0.648704346 |
| AC006978.1 | 611.4875955 | -0.559908001 | 0.534327518 | -1.047874164 | 0.294696587 | 0.6068228 |
| AC006978.2 | 10.27069758 | -3.305462269 | 1.410941442 | -2.342735263 | 0.019142961 | 0.157152521 |
| AC007068.1 | 51.44012205 | 0.19046791 | 0.699594072 | 0.272254894 | 0.785426036 | 0.910931322 |
| AC007244.1 | 19.50449106 | 0.367114982 | 0.858056378 | 0.427844826 | 0.668764098 | 0.85734495 |
| AC007318.1 | 13.55615196 | -0.040062835 | 0.916493345 | -0.043713176 | | |

0.965133036 0.988700292
AC007325.2 1368.143409 -1.521836334 0.430927706 -3.531535134
0.000413155 0.015709208
AC007325.4 135.135393 -1.658174736 0.978670808 -1.694313065
0.090205809 0.344106696
AC007326.4 396.4030874 -1.002251094 0.589718606 -1.699541245
0.08921725 0.342726775
AC007546.3 29.32760561 -0.496833093 0.671995737 -0.739339649
0.459700779 0.737689921
AC007842.1 10.69880732 -0.991549527 0.952025702 -1.041515502
0.297636362 0.60928029
AC007906.2 98.39600029 0.969153948 0.624384941 1.552173803 0.120620643
0.400973879
AC007950.1 1120.517894 0.32981881 0.26697274 1.235402572 0.216680716
0.524312093
AC007969.1 10.28620449 0.686505288 0.945562681 0.726028324 0.467821407
0.744166872
AC008012.1 367.6312205 0.428901206 0.286581585 1.496611186 0.134494458
0.420896533
AC008013.1 14.76253176 0.510894014 0.831363203 0.614525652 0.538868027
0.786598616
AC008013.2 48.80414547 -0.082926275 0.710703835 -0.116681902
0.907112132 0.965296417
AC008038.1 31.77186294 -0.079995015 0.603179181 -0.132622307
0.89449209 0.958237013
AC008073.3 31.97090294 -0.649637585 0.560514456 -1.159002373
0.246455218 0.558555738
AC008132.1 17.24737137 -1.736101079 1.235693365 -1.404961076
0.160032853 0.460344789
AC008267.2 13.48390596 -0.546066193 0.899836975 -0.606850139
0.54395037 0.790296022
AC008397.2 19.93810455 1.088503328 0.801656467 1.357817684 0.174521542
0.478578744
AC008403.1 191.2285981 -0.588949588 0.451853097 -1.30340943 0.192435019
0.49845171
AC008554.1 945.2713318 -0.165423099 0.323001958 -0.512142712
0.608551133 0.827863096
AC008581.2 86.48819424 0.035630958 0.957146893 0.037226217 0.970304635
0.98957539
AC008736.1 87.74321394 -1.185317817 0.570086859 -2.079188108
0.037600064 0.22230342
AC008752.3 13.72222564 -0.596166035 0.82282126 -0.724538929
0.468734936 0.744337782
AC008755.1 1092.889216 -0.156990626 0.247441427 -0.63445571 0.525783463
0.778438656
AC008758.1 15.53181906 2.949460516 4.185588108 0.704670512 0.481015306
0.75260258
AC008758.5 20.53316051 -0.798712104 1.249979242 -0.638980294
0.522835751 0.777662198
AC008764.4 37.15843753 -0.534535563 1.357802445 -0.393676978
0.693819541 0.868802318
AC008770.4 83.41510691 0.513532196 0.403106267 1.273937517 0.202685561
0.509290789
AC008878.1 63.23103087 -0.522384215 0.453879898 -1.150930493
0.249760832 0.562685117
AC008878.3 85.28607607 -2.953535532 3.680514818 -0.802478913
0.422275981 0.712133049
AC008982.1 41.05055933 7.237316678 4.260436651 1.698726509 0.089370726
0.342726775
AC009053.1 100.0215816 -0.74391358 0.762413716 -0.975734781 0.329195912
0.638424836
AC009086.2 24.412772 1.151063812 1.223574318 0.940738781 0.34683874
0.65343598
AC009093.3 10.34998753 -0.799376314 1.016463976 -0.786428573

0.43161645 0.718224543
AC009093.8 10.42656636 -1.315948984 1.221594507 -1.077238786
0.281373602 0.595211573
AC009119.2 16.91266707 -2.033179012 0.90895126 -2.236840525
0.025296765 0.181849597
AC009133.6 470.5860475 1.250901832 0.618116161 2.023732608 0.042997672
0.238392703
AC009244.1 43.11552133 1.698983154 0.771396404 2.202477409 0.027631603
0.190244919
AC009412.1 317.5656933 -0.681720826 0.580280716 -1.174812133
0.240069885 0.551666195
AC009533.1 204.4315073 -0.302144419 0.685222572 -0.440943471
0.659253922 0.851702397
AC009690.1 14.45823484 0.945394845 0.871979423 1.084193984 0.278278798
0.592005173
AC009779.3 85.99540334 0.38099997 0.439834966 0.866233928 0.386361891
0.686679504
AC009879.3 21.4043749 -1.093281053 0.867549264 -1.260194779
0.207599106 0.514453779
AC010132.3 522.8865697 0.23852403 0.249644572 0.955454503 0.339347903
0.646260212
AC010186.2 173.3707278 -0.138203698 0.438347677 -0.31528329 0.752546571
0.896586461
AC010197.2 11.57465973 0.227843562 0.853453361 0.266966623 0.78949486
0.91284403
AC010203.1 38.96519903 -0.978507417 0.74083033 -1.320825265
0.186559633 0.49118221
AC010323.1 29.43283736 -0.612264652 1.245824259 -0.491453468
0.623105756 0.833907812
AC010326.4 86.63233169 0.291207967 0.618187567 0.47106733 0.637592654
0.841782104
AC010422.3 20.15901768 1.18452213 1.048461148 1.129772078 0.258572276
0.571503497
AC010422.6 28.63175767 -2.658932955 1.018765824 -2.609954999
0.009055413 0.107495835
AC010422.8 209.4558536 -0.004225164 0.346013725 -0.012210973
0.990257295 0.996629523
AC010487.1 45.53041753 -0.198273727 0.532161359 -0.372581968
0.709459595 0.875983684
AC010487.3 52.84060812 -0.781676838 1.170645201 -0.667731638
0.504304911 0.765923496
AC010507.2 66.78941721 -1.086980694 0.442378662 -2.457127316
0.014005304 0.134477513
AC010522.1 15.36982146 -0.189981653 1.269842191 -0.149610443
0.881071969 0.952848873
AC010547.4 12.06053472 -2.835737266 1.256388512 -2.257054436
0.024004673 0.177207433
AC010615.1 111.7552615 -0.585406392 0.482267836 -1.213861567
0.224800591 0.534134654
AC010618.1 969.1643406 0.28201497 0.424437069 0.66444472 0.506405719
0.767347299
AC010655.3 24.04795495 0.551961186 0.665873031 0.82892858 0.407144822
0.701739367
AC010761.1 415.9179207 -0.135941961 0.313593622 -0.433497213
0.664653591 0.855037526
AC010970.1 17.06081756 0.631773203 0.949415197 0.665434053 0.50577291
0.767164369
AC010980.1 84.15984018 0.664014752 0.916900537 0.724194965 0.468946049
0.744337782
AC011043.1 934.6783958 0.139369521 0.309984986 0.449600874 0.652998258
0.848824999
AC011447.1 25.04214245 -1.775685458 1.180519715 -1.5041557 0.132541281
0.418641974
AC011448.1 45.1842708 0.111070133 0.495285198 0.224254901 0.822558966

| | | | | | | |
|---|---|---|---|---|---|---|
| | | | | | | 0.925840841 |
| AC011455.2 | 33.95914579 | -1.162288013 | 1.223923158 | -0.949641328 | 0.342294532 | 0.649037179 |
| AC011462.1 | 65.58169194 | -0.719136109 | 0.591121498 | -1.216562266 | 0.223770807 | 0.53293986 |
| AC011477.1 | 25.35249503 | 0.101922397 | 0.761001897 | 0.133931856 | 0.893456459 | 0.95780189 |
| AC011481.1 | 1707.661504 | 0.364074816 | 0.2539164 | 1.43383734 | 0.151618686 | 0.447172604 |
| AC011497.2 | 14.57422594 | -0.546981266 | 1.001172873 | -0.546340478 | 0.584831917 | 0.815111163 |
| AC011498.4 | 13.74239456 | 0.076383045 | 0.773489833 | 0.098751195 | 0.92133582 | 0.970205128 |
| AC011503.3 | 25.35384359 | 0.10425628 | 0.760087709 | 0.137163486 | 0.890901572 | 0.957017706 |
| AC011511.1 | 37.90800843 | 0.600472537 | 0.565639533 | 1.061581629 | 0.288425657 | 0.601673803 |
| AC011511.4 | 551.3441227 | 0.304754061 | 0.471886195 | 0.6458211 | 0.518395222 | 0.774169079 |
| AC011529.1 | 119.4675219 | 0.789428141 | 0.558421415 | 1.413678128 | 0.15745643 | 0.456731315 |
| AC012184.2 | 42.18629029 | -2.230848914 | 1.20627024 | -1.849377395 | 0.064403337 | 0.293255043 |
| AC012213.5 | 63.95546365 | -0.284482688 | 0.568859366 | -0.500093177 | 0.61700947 | 0.831835425 |
| AC012254.2 | 70.95713756 | 1.910913715 | 0.557947458 | 3.424899041 | 0.000615028 | 0.020152459 |
| AC012435.1 | 223.4095031 | -0.128705445 | 0.453702854 | -0.283677838 | 0.77665728 | 0.907755593 |
| AC012501.2 | 10.63201524 | -0.290930244 | 1.101969225 | -0.264009409 | 0.791772668 | 0.913528265 |
| AC012618.3 | 77.62754325 | -0.184968818 | 0.606114102 | -0.305171613 | 0.760235451 | 0.899760802 |
| AC012651.1 | 20.58423272 | -1.021587366 | 0.856099657 | -1.193304258 | 0.232750236 | 0.542441738 |
| AC013394.1 | 712.7056817 | 1.081163552 | 0.425227558 | 2.542552877 | 0.011004595 | 0.119239998 |
| AC013489.1 | 68.44590831 | 0.793744222 | 0.452283389 | 1.75497098 | 0.079264272 | 0.322931692 |
| AC015802.6 | 38.44438449 | 0.109779884 | 0.809044353 | 0.13569081 | 0.892065713 | 0.957468421 |
| AC015813.2 | 583.5030827 | 0.688879615 | 0.37057426 | 1.858951603 | 0.063033999 | 0.289778852 |
| AC015818.6 | 31.18535456 | 0.312660924 | 0.6574937 | 0.475534479 | 0.634406064 | 0.84020728 |
| AC016582.1 | 54.82078039 | -1.3057812 | 0.731162074 | -1.785898429 | 0.074115705 | 0.313031561 |
| AC016582.2 | 10.84137544 | 0.276696006 | 1.050025771 | 0.263513538 | 0.79215479 | 0.913570955 |
| AC016582.3 | 191.1110225 | -1.243498389 | 0.378430896 | -3.285932525 | 0.001016454 | 0.027467362 |
| AC017083.3 | 15.02167678 | -6.528191902 | 1.701813292 | -3.83602122 | 0.000125044 | 0.007078495 |
| AC017104.4 | 27.02819599 | 1.390843323 | 0.674876156 | 2.060886745 | 0.039313846 | 0.228518481 |
| AC017104.5 | 14.58404433 | -0.827606094 | 1.032093921 | -0.801870913 | 0.422627633 | 0.712269066 |
| AC018362.3 | 29.30618244 | -0.575566637 | 0.664893781 | -0.865652009 | 0.386681025 | 0.687008387 |
| AC018523.2 | 10.53958675 | 1.31912885 | 1.575219232 | 0.837425561 | 0.402353403 | 0.699450381 |
| AC018630.2 | 33.56496249 | -0.277328939 | 1.480602896 | -0.187308116 | 0.85141906 | 0.939477675 |
| AC018638.2 | 36.09628381 | -1.460374352 | 0.710899819 | -2.054261814 | | |

0.039950359 0.230276174
AC018638.4 152.8820666 -1.83084891 0.89236412 -2.051683691 0.040200413 0.231327132
AC018638.5 51.49339616 -0.562162276 0.573146158 -0.98083581 0.326673716 0.636466385
AC018641.1 28.66744822 -1.114996413 0.736778762 -1.513339513 0.130193441 0.415299556
AC018804.1 17.0567303 0.63625564 0.805187754 0.790195375 0.429413676 0.71613003
AC020907.6 12.07141297 1.300920999 1.634128296 0.796094776 0.42597695 0.71395509
AC020915.1 36.05419919 0.119483981 0.703247335 0.169903212 0.865086255 0.945248223
AC020915.5 165.0161511 -0.856387854 2.143444266 -0.399538195 0.689496686 0.867647165
AC021087.5 83.31264872 -0.106624512 0.607456475 -0.175526176 0.860666205 0.943645503
AC022137.3 133.7661048 0.441168751 0.490186241 0.900002313 0.36811902 0.67424045
AC022182.3 10.26246551 0.136288426 0.949595729 0.143522577 0.885877482 0.954877821
AC022384.1 83.97501472 0.042256143 2.095729899 0.020162972 0.983913366 0.994061643
AC022400.7 167.5938223 -0.432438308 0.335121058 -1.290394313 0.196913786 0.50258876
AC022414.1 17.87435243 0.199437887 1.325740734 0.15043506 0.880421382 0.952483146
AC022415.1 202.8781256 0.643658504 0.585283455 1.099738081 0.271446258 0.583632536
AC022415.2 277.1080719 -0.11763407 0.685353265 -0.171640052 0.863720518 0.94507782
AC022826.2 29.42601132 0.855391611 1.327330284 0.644445185 0.519286792 0.77471727
AC023055.1 28.37413045 0.066867417 0.934002007 0.071592369 0.942926313 0.978574897
AC023794.4 10.62497549 -1.721514428 1.155442577 -1.48991777 0.136245859 0.424171131
AC024592.3 13.80699435 0.547386995 1.16682489 0.46912523 0.638980126 0.842953695
AC024940.2 72.61718602 -1.335644977 0.877638542 -1.521862263 0.128043597 0.412255816
AC025423.2 89.49943516 -0.671727647 0.514151881 -1.30647708 0.191390369 0.496933644
AC025449.1 12.97519872 0.519068393 1.040965989 0.498641068 0.618032268 0.832176095
AC026316.4 11.08909306 0.068787059 1.00469771 0.068465429 0.945415139 0.979754792
AC026412.1 108.5769984 -0.605888718 0.480703266 -1.26042147 0.207517361 0.514335219
AC026464.4 837.947813 -0.297509179 0.58939878 -0.504767211 0.613722348 0.830316752
AC026464.6 280.5517134 1.301531383 0.518030051 2.512463091 0.011989165 0.124327283
AC026470.1 1000.522675 -0.254028119 0.334716565 -0.758935007 0.447891437 0.729386108
AC026877.1 16.38784402 -0.904932358 1.182400715 -0.765334752 0.444072235 0.727127105
AC026954.2 25.78551177 4.567777198 2.879887649 1.586095624 0.112717619 0.38618767
AC027559.1 31.79450072 0.987298671 0.723050492 1.365462968 0.172107585 0.475426833
AC027644.4 13.45353214 -0.033575106 0.842794818 -0.039837818 0.968222426 0.989351548
AC034102.2 47.71628666 1.432797085 3.678055847 0.389552836 0.696867234

0.870026319
AC040162.1 33.30136992 0.783710072 0.561054546 1.396851834 0.162458131 0.463216519
AC044860.1 39.44299544 1.049650082 0.813776207 1.289851035 0.197102384 0.50258876
AC055811.2 10.49007559 -1.740403134 1.187941715 -1.465057681 0.142905181 0.433233718
AC055839.2 619.592136 0.986076086 0.546678952 1.803757181 0.071269381 0.306992555
AC058822.1 19.42260706 -0.14744248 1.250542464 -0.117902818 0.906144661 0.964913361
AC060766.1 81.30017217 -0.504973343 0.461375018 -1.094496502 0.273737281 0.586847211
AC060766.5 60.07641777 -0.041575022 0.479490465 -0.086706671 0.930904674 0.974050266
AC064799.1 19.31603807 1.084935945 0.736050987 1.473995638 0.140482765 0.430123922
AC067968.1 73.67341133 -1.556383152 0.751588591 -2.07079135 0.038378299 0.225585886
AC068134.3 33.00655055 -1.220272928 1.125716779 -1.083996393 0.278366398 0.592005173
AC068533.4 59.43282012 0.669064442 0.736858722 0.907995551 0.363880572 0.670443546
AC068547.1 177.7840885 -0.683864602 0.382462351 -1.788057308 0.073766762 0.311905115
AC068580.4 52.32714946 -0.01592161 0.610867355 -0.02606394 0.979206339 0.992858175
AC068620.3 20.33804443 -0.079364792 0.8236087 -0.096362255 0.923232869 0.971120333
AC068631.2 208.3247558 -0.006261671 0.340083698 -0.018412148 0.985310062 0.99460071
AC068831.7 378.2239879 0.222702861 0.830328862 0.26821043 0.788537347 0.912309638
AC068896.1 12.42947737 -0.38264424 1.065648734 -0.359071641 0.719541496 0.881544378
AC068946.1 12.76949417 -0.64282994 1.253275737 -0.5129198 0.608007423 0.827607804
AC068946.2 62.85572226 1.832534653 0.732263289 2.502562508 0.012329785 0.126347892
AC069288.1 151.1452248 -0.065334121 0.513353861 -0.127269172 0.898727359 0.960250212
AC069499.1 20.2530654 -1.203797006 1.165842437 -1.032555488 0.301811968 0.612629029
AC073111.5 248.2502896 -0.921995761 0.470659382 -1.958944827 0.050119249 0.257487003
AC073508.2 20.48971404 -1.269041351 0.802741271 -1.580884647 0.113904415 0.388324469
AC073611.1 63.1383564 0.117338797 0.689098201 0.170278775 0.864790903 0.945248223
AC073869.1 395.6529969 -0.240292081 0.390181495 -0.615846944 0.537995541 0.786237391
AC073896.1 46.98806248 -0.266272221 2.260460888 -0.117795544 0.90622966 0.964936143
AC074141.1 40.91883309 -0.845868757 0.583228717 -1.450320829 0.146969073 0.440037066
AC074143.1 114.4541492 0.311071574 0.42055727 0.739665194 0.459503173 0.737476588
AC079140.3 47.68817369 1.286819351 0.608397907 2.115094966 0.034421854 0.211851485
AC079140.5 10.30786205 0.269374989 1.02016578 0.26405021 0.791741229 0.913528265
AC079447.1 12.97997461 -0.398147509 0.914719904 -0.435267132 0.66336854 0.854409382
AC079594.2 15.59609106 0.217977785 1.443505198 0.151005889 0.879971071

| | | | | | | |
|---|---|---|---|---|---|---|
| | | | | | | 0.952214059 |
| AC080038.1 | 1087.970742 | 0.401004102 | 0.523328586 | 0.766256828 | 0.443523501 | 0.726848446 |
| AC083899.1 | 97.10136339 | -0.017966166 | 0.513463103 | -0.034990179 | 0.972087572 | 0.989900695 |
| AC087289.4 | 86.7272761 | 0.972738201 | 0.643981231 | 1.510507068 | 0.130914087 | 0.416533681 |
| AC087632.1 | 121.8914847 | 0.32522383 | 0.52119408 | 0.623997552 | 0.532629193 | 0.782684541 |
| AC087651.1 | 24.4008659 | -0.760498514 | 0.848273621 | -0.896525007 | 0.369972431 | 0.67567735 |
| AC087721.2 | 1242.467847 | 0.231225477 | 0.477838661 | 0.483898638 | 0.628457807 | 0.836791621 |
| AC087783.2 | 77.55727053 | 0.302599639 | 0.427225251 | 0.708290622 | 0.4787648 | 0.751165197 |
| AC087894.1 | 11.29722086 | 0.042466258 | 1.03277684 | 0.041118523 | 0.967201408 | 0.988774431 |
| AC090099.1 | 118.6607722 | 0.818142877 | 0.549925393 | 1.487734314 | 0.136820975 | 0.424728769 |
| AC090114.3 | 77.00738331 | -1.528189129 | 1.914886296 | -0.798057374 | 0.424837196 | 0.71350603 |
| AC090360.1 | 17.73662599 | 1.25942134 | 0.733234173 | 1.717624991 | 0.085865039 | 0.335996519 |
| AC090517.4 | 117.0963954 | -0.112881012 | 0.392995387 | -0.287232411 | 0.773934371 | 0.906248714 |
| AC090607.2 | 18.8590382 | -1.196394832 | 0.940367034 | -1.27226369 | 0.203279445 | 0.509659967 |
| AC090616.2 | 39.94133779 | 1.728838928 | 0.699958948 | 2.469914748 | 0.013514526 | 0.132362083 |
| AC090971.3 | 35.24617344 | -0.778529557 | 0.616884944 | -1.262033648 | 0.206936679 | 0.513642439 |
| AC091057.6 | 90.84004661 | 1.126436834 | 0.3722297 | 3.026187417 | 0.002476587 | 0.048014245 |
| AC091133.4 | 25.22383384 | 0.245888501 | 0.828243613 | 0.296879441 | 0.766558553 | 0.903470079 |
| AC091390.4 | 51.23259578 | 0.318224578 | 0.522491905 | 0.609051691 | 0.542490177 | 0.789223006 |
| AC091551.1 | 11.3004661 | -0.263894509 | 1.393065483 | -0.18943439 | 0.849752373 | 0.938580023 |
| AC091959.3 | 106.6232756 | 0.529987133 | 2.211774556 | 0.239620775 | 0.810624258 | 0.920714278 |
| AC092070.2 | 138.1147118 | -1.618697043 | 0.408813153 | -3.959503341 | 7.51E-05 | 0.004975677 |
| AC092072.1 | 197.9017391 | -0.096899579 | 0.376441822 | -0.257409176 | 0.79686292 | 0.915710299 |
| AC092299.1 | 29.62507113 | -0.198543111 | 1.437079061 | -0.138157403 | 0.89011602 | 0.956709184 |
| AC092329.3 | 153.7886451 | -0.222283437 | 0.549310763 | -0.404658805 | 0.685728327 | 0.86493781 |
| AC092338.1 | 548.8230108 | 0.805881707 | 0.331352762 | 2.432095944 | 0.01501173 | 0.139113039 |
| AC092587.1 | 20.83970691 | 1.059529405 | 0.908334227 | 1.166453243 | 0.243431243 | 0.554768722 |
| AC092647.5 | 47.12691645 | -0.784840181 | 0.746924999 | -1.0507617 | 0.29336805 | 0.606314063 |
| AC092651.1 | 33.50047001 | 9.216302938 | 4.330163768 | 2.12839593 | 0.03330427 | 0.208274771 |
| AC092651.2 | 10.42049146 | -0.551860594 | 1.13899926 | -0.484513567 | 0.628021438 | 0.836571537 |
| AC092685.1 | 14.06068184 | -2.85412212 | 1.162407338 | -2.455354528 | 0.014074571 | 0.134821147 |
| AC092718.3 | 129.1676032 | 0.570445099 | 0.328643214 | 1.735758034 | 0.082606627 | 0.328745819 |
| AC092821.1 | 248.3648177 | 0.029357189 | 0.35196069 | 0.083410419 | 0.933525204 | |

0.974775337
AC092835.1 30.06660315 -0.760727058 1.172634419 -0.648733353
0.516510742 0.773950426
AC093155.3 15.97787737 0.173571122 0.803099325 0.216126595 0.82888907
0.929284369
AC093249.4 13.23319879 1.248756817 0.931410871 1.340715313 0.180012901
0.484754911
AC093323.1 503.4317599 0.539988246 0.396515579 1.36183362 0.173250399 0.4771935
AC093512.2 136.4699621 -0.11990716 0.687276847 -0.174467044 0.861498429
0.94403207
AC093616.1 10.88841087 -0.868394614 1.049678892 -0.827295491
0.408069599 0.702586667
AC093668.2 46.18423808 -0.477093597 0.607690052 -0.785093644
0.432398666 0.718639059
AC093668.3 318.7599238 -1.315138265 0.34915918 -3.766586538
0.000165495 0.008482164
AC093752.1 285.7695538 0.414871163 0.27346227 1.517105682 0.129240004
0.413472911
AC093827.5 29.14457343 0.075636232 0.66630659 0.11351566 0.909621748
0.96630013
AC097636.2 14.47242202 -1.014547086 0.867639716 -1.169318401
0.242275372 0.553231259
AC098484.3 119.0501131 0.789823599 0.514059676 1.536443403 0.124429669
0.405999855
AC098850.3 16.30005217 2.016006479 1.619561821 1.244785134 0.213210742
0.52037004
AC098934.1 806.0722589 -0.368286406 0.452201733 -0.814429444
0.415398977 0.707513171
AC098934.2 95.53182421 -0.633600278 0.401846648 -1.57672157 0.114859605
0.389985713
AC099489.1 188.9091399 0.158324165 0.56397615 0.280728476 0.778918666
0.908299392
AC099850.2 39.00600829 5.007065185 2.39589212 2.089854189 0.0366309
0.219135375
AC104389.4 20.54400743 -0.258147022 0.75205096 -0.343257353
0.731404861 0.887123693
AC104452.1 152.8088028 -1.419905833 2.05282518 -0.691683757
0.48913595 0.75752159
AC104662.2 73.30519627 -0.757611328 0.659967854 -1.147951863
0.250988436 0.564026895
AC105052.2 21.64191532 0.706091667 0.679323471 1.039404198 0.298616793
0.609979179
AC105052.3 264.8791872 -0.72014347 0.601057901 -1.198126618 0.230867728
0.539623726
AC106795.1 211.9219974 0.68644491 0.45342105 1.513923778 0.130045173
0.415175781
AC106886.5 778.8415985 -0.771099276 0.316311834 -2.437781941
0.014777685 0.138219645
AC107075.1 13.17434214 7.868937892 4.333535044 1.81582422 0.069397341
0.30336211
AC107871.1 123.0711113 0.028320957 0.941905747 0.030067719 0.976013046
0.991758053
AC107918.4 15.33765529 0.262816007 0.941422507 0.279169029 0.78011511
0.909065699
AC108488.2 15.42141122 0.272826277 0.755143518 0.36129063 0.717882198
0.880671367
AC109583.3 45.84084178 -0.530564375 0.830302367 -0.639001399
0.522822021 0.777662198
AC110079.1 409.60062 -1.938388669 1.507004913 -1.286252389
0.198354993 0.50414535
AC112128.1 56.67178682 -1.21074619 0.762485945 -1.587893125 0.112310508
0.385402109
AC112184.1 21.38725035 1.600712009 0.784324912 2.040878703 0.041262885
0.233407618

| | | | | | | |
|---|---|---|---|---|---|---|
| AC112206.3 | 33.27720934 | -0.867564958 | 0.665142294 | -1.304329864 | 0.192121137 | 0.498063871 |
| AC112206.4 | 18.74768051 | -0.495532058 | 0.786523003 | -0.630028691 | 0.528675813 | 0.779677336 |
| AC114267.1 | 26.55253271 | -0.198390255 | 0.664368455 | -0.298614803 | 0.765233969 | 0.903026935 |
| AC114490.2 | 73.74264295 | -0.266997269 | 2.037987907 | -0.131010232 | 0.895767213 | 0.95884318 |
| AC114490.3 | 12.09953649 | 0.71467342 | 0.892841575 | 0.800448186 | 0.423451172 | 0.712688899 |
| AC114801.3 | 20.23086039 | 0.766739767 | 1.459501394 | 0.525343634 | 0.599344329 | 0.823816451 |
| AC115223.1 | 102.6223867 | 0.857743396 | 0.420814706 | 2.038292351 | 0.041520704 | 0.234080493 |
| AC116366.3 | 396.9652752 | 0.053214288 | 0.634199517 | 0.083907803 | 0.933129735 | 0.97462713 |
| AC116533.1 | 326.4424531 | 4.785003254 | 3.815625819 | 1.254054638 | 0.209822147 | 0.517783588 |
| AC116562.4 | 218.4144591 | -0.66188745 | 0.65973592 | -1.0032612 | 0.315734851 | 0.625491438 |
| AC117378.1 | 70.89705681 | 1.74726011 | 0.571040601 | 3.059782625 | 0.002214977 | 0.044579397 |
| AC117529.1 | 29.57753739 | -1.091382148 | 0.976845334 | -1.117251739 | 0.263886704 | 0.577444856 |
| AC118549.1 | 1614.211925 | -0.076161615 | 0.345463759 | -0.220461953 | 0.825511401 | 0.92714195 |
| AC118553.2 | 79.44997257 | -0.765259728 | 0.875026607 | -0.874555953 | 0.381815563 | 0.683241792 |
| AC118754.1 | 246.6884991 | -2.472551172 | 0.817180556 | -3.025709746 | 0.002480503 | 0.048014245 |
| AC120114.4 | 155.7729785 | 1.384352971 | 0.539646759 | 2.565294702 | 0.010308821 | 0.115220384 |
| AC124312.1 | 182.4850927 | -0.726250441 | 0.483637511 | -1.501642085 | 0.133189568 | 0.419488075 |
| AC124319.1 | 1344.214695 | -1.037872085 | 0.4442172 | -2.336406796 | 0.01947005 | 0.158211103 |
| AC124944.3 | 249.005454 | -0.593894443 | 0.467257202 | -1.271022555 | 0.203720625 | 0.509986087 |
| AC125232.1 | 464.7240086 | -0.065754527 | 0.317403613 | -0.207163764 | 0.835881965 | 0.932644355 |
| AC126177.6 | 17.05687277 | -0.202144794 | 0.831383844 | -0.243142557 | 0.807894965 | 0.919552343 |
| AC126755.1 | 125.6418072 | 0.679626119 | 1.106218404 | 0.614368841 | 0.538971621 | 0.786603027 |
| AC129492.1 | 473.6594463 | 0.127464479 | 0.346562941 | 0.367796044 | 0.713025319 | 0.877743193 |
| AC131160.1 | 16.8543504 | -6.688677117 | 1.626779216 | -4.111607189 | 3.93E-05 | 0.003088551 |
| AC131206.1 | 276.1211947 | 0.711941864 | 0.450763147 | 1.579414529 | 0.114241007 | 0.389050381 |
| AC131392.1 | 43.25393955 | -0.443834283 | 1.031199347 | -0.4304059 | 0.666900404 | 0.856258128 |
| AC132008.2 | 381.1482543 | -0.178719059 | 0.364736151 | -0.489995463 | 0.624137109 | 0.834707114 |
| AC133552.2 | 23.31521378 | 0.023218948 | 0.659308664 | 0.035217113 | 0.971906617 | 0.989782861 |
| AC135178.2 | 26.22042635 | 0.672311068 | 1.35023802 | 0.49792041 | 0.618540141 | 0.83229541 |
| AC135506.1 | 30.88774941 | 0.782130765 | 0.873699848 | 0.895193889 | 0.370683451 | 0.676128642 |
| AC135721.1 | 15.02867379 | -1.222725588 | 0.823618185 | -1.484578182 | 0.137655599 | 0.425849298 |
| AC135977.1 | 32.09004988 | -0.042253996 | 1.620525391 | -0.026074257 | 0.97919811 | 0.992858175 |

| | | | | | | |
|---|---|---|---|---|---|---|
| AC137695.2 | 15.12595816 | -0.411806908 | 0.937280955 | -0.439363359 | 0.660398272 | 0.852355524 |
| AC137834.1 | 50.30634739 | -1.128117578 | 1.137962647 | -0.991348513 | 0.321515432 | 0.631253641 |
| AC138207.8 | 42.90566703 | -0.394174721 | 0.673577248 | -0.585196015 | 0.558415924 | 0.798855459 |
| AC138393.1 | 181.397968 | -0.052531945 | 0.470703866 | -0.11160296 | 0.911138225 | 0.966866396 |
| AC138409.2 | 439.1749383 | 0.022783504 | 0.359406209 | 0.06339207 | 0.949454302 | 0.981593436 |
| AC138811.2 | 20.16344188 | -0.688998894 | 0.871927403 | -0.790202133 | 0.42940973 | 0.71613003 |
| AC138866.1 | 136.5653186 | -0.785109379 | 0.47028973 | -1.669416379 | 0.09503489 | 0.353833719 |
| AC138866.2 | 11.07858925 | -1.433295805 | 1.272955169 | -1.125959373 | 0.260182716 | 0.57337005 |
| AC138932.1 | 117.3905456 | 0.635541547 | 2.479934125 | 0.25627356 | 0.797739612 | 0.915916373 |
| AC138969.1 | 868.2044807 | -0.442097006 | 0.428987101 | -1.030560138 | 0.302747137 | 0.613538084 |
| AC139491.6 | 17.0262149 | -2.038689811 | 0.929074454 | -2.194323397 | 0.028212169 | 0.192229399 |
| AC139530.3 | 12.1414149 | -2.297267138 | 1.992459439 | -1.152980629 | 0.248918335 | 0.56168903 |
| AC139768.1 | 83.42414424 | -0.403941847 | 0.444211257 | -0.909346264 | 0.363167377 | 0.670041454 |
| AC139769.1 | 527.697873 | 0.26578989 | 0.25342207 | 1.048803247 | 0.294268683 | 0.606538766 |
| AC139795.1 | 21.86419509 | 0.008524195 | 0.967514345 | 0.008810407 | 0.992970403 | 0.997243081 |
| AC140134.1 | 12.8385076 | -1.361671385 | 1.36514509 | -0.997455432 | 0.318543496 | 0.6285113 |
| AC141557.1 | 19.93223048 | 0.267118875 | 0.834945262 | 0.319923816 | 0.749026083 | 0.895096304 |
| AC141586.1 | 172.7357303 | -0.149924203 | 0.454215437 | -0.330072892 | 0.741344886 | 0.892005215 |
| AC146944.1 | 28.35432392 | -0.21959351 | 0.814582877 | -0.269577861 | 0.787485034 | 0.911979806 |
| AC211476.3 | 18.40381855 | -1.633323531 | 1.238553388 | -1.318734862 | 0.187257768 | 0.49211797 |
| AC211486.3 | 10.69197457 | -0.013712303 | 1.455180419 | -0.009423095 | 0.992481569 | 0.997035603 |
| AC233699.1 | 15.89770083 | -0.962337264 | 1.250672236 | -0.769456006 | 0.441622652 | 0.725092776 |
| AC233968.1 | 186.9553985 | -0.682199783 | 0.419827221 | -1.624953668 | 0.104172432 | 0.369857233 |
| AC234031.1 | 461.8918858 | 0.210005205 | 0.372112852 | 0.564358916 | 0.572509893 | 0.807582296 |
| AC234783.1 | 34.1328031 | 0.411976747 | 0.666774683 | 0.617865012 | 0.536664324 | 0.785348759 |
| AC239811.1 | 48.21517835 | 1.882908344 | 0.732091443 | 2.571957865 | 0.01011252 | 0.114259456 |
| AC239868.1 | 77.48654245 | -0.409029077 | 0.510030682 | -0.80196955 | 0.422570572 | 0.712269066 |
| AC240274.1 | 458.1329125 | -0.249951125 | 0.302940803 | -0.825082399 | 0.409324811 | 0.703508181 |
| AC241952.1 | 152.6672831 | -0.708393374 | 0.48878226 | -1.449302548 | 0.14725311 | 0.440626613 |
| AC242376.2 | 131.6911529 | 0.208344063 | 0.345057331 | 0.603795499 | 0.545979612 | 0.791677099 |
| AC242843.1 | 254.8862249 | 0.966450226 | 0.479366721 | 2.016097873 | 0.043789742 | 0.240941849 |
| AC244154.1 | 185.4715517 | -0.554628542 | 0.45640205 | -1.215219218 | 0.224282493 | 0.53365585 |

| | | | | | | |
|---|---|---|---|---|---|---|
| AC244197.3 | 274.1516222 | -0.195201747 | 0.811385494 | -0.240578305 | 0.809881967 | 0.920215647 |
| AC244669.1 | 77.66189008 | 0.704777309 | 0.585657121 | 1.203395782 | 0.228823199 | 0.537506211 |
| AC245060.4 | 393.7475465 | 0.225389953 | 0.270988938 | 0.831731193 | 0.40556069 | 0.701249284 |
| AC245100.4 | 10.26909703 | -0.771689665 | 0.962934891 | -0.801393399 | 0.422903935 | 0.712273914 |
| ACAA1 | 664.3888588 | -0.656869131 | 0.386607806 | -1.699058115 | 0.089308234 | 0.342726775 |
| ACAA2 | 2902.333771 | -0.219977302 | 0.309280832 | -0.711254237 | 0.476926705 | 0.749746108 |
| ACACA | 629.2964054 | -1.052210409 | 0.390732819 | -2.692915356 | 0.007083025 | 0.092634974 |
| ACACB | 321.6090788 | -0.223378026 | 0.466143903 | -0.479204006 | 0.631793505 | 0.838768903 |
| ACAD10 | 964.3648363 | -0.720263837 | 0.383478797 | -1.878236404 | 0.060348832 | 0.283945763 |
| ACAD11 | 1505.955345 | 0.973469142 | 0.436297083 | 2.231207083 | 0.025667415 | 0.182645569 |
| ACAD8 | 1136.244114 | 0.097551677 | 0.270385035 | 0.360788003 | 0.718257932 | 0.88075421 |
| ACAD9 | 916.142123 | -0.515338254 | 0.288069201 | -1.78893909 | 0.073624625 | 0.311651561 |
| ACADL | 17.70680109 | -1.167436659 | 0.821566349 | -1.420988896 | 0.155319988 | 0.453268877 |
| ACADM | 1621.815928 | 0.156915707 | 0.266446282 | 0.588920611 | 0.555914531 | 0.796719574 |
| ACADS | 115.438544 | -0.190836857 | 0.517214075 | -0.368970735 | 0.712149538 | 0.877474733 |
| ACADSB | 641.9428877 | -0.327035866 | 0.28895865 | -1.131773925 | 0.257729493 | 0.570388641 |
| ACADVL | 4075.881154 | 0.011494633 | 0.290387698 | 0.039583745 | 0.968424987 | 0.989435867 |
| ACAP1 | 111.1901385 | 0.514259643 | 0.423394972 | 1.214609707 | 0.224514985 | 0.533874112 |
| ACAP2 | 1509.700847 | -0.364005451 | 0.252876407 | -1.439459917 | 0.150020259 | 0.444740017 |
| ACAP3 | 488.9999075 | -0.036262983 | 0.53334345 | -0.067991804 | 0.945792159 | 0.979962631 |
| ACAT1 | 703.4210666 | 0.930899273 | 0.280197012 | 3.322302639 | 0.000892778 | 0.025375039 |
| ACAT2 | 2747.744061 | 0.333024255 | 0.223734122 | 1.488482187 | 0.136623776 | 0.424728769 |
| ACBD3 | 1070.323945 | -0.322864319 | 0.232322645 | -1.38972384 | 0.164612753 | 0.466967104 |
| ACBD4 | 231.6491547 | 0.145743556 | 0.378651408 | 0.384901662 | 0.700310275 | 0.871925086 |
| ACBD5 | 629.6063225 | -0.170129716 | 0.288417112 | -0.589873862 | 0.555275219 | 0.796360966 |
| ACBD6 | 1147.15571 | 0.403390332 | 0.305950513 | 1.318482284 | 0.187342252 | 0.492150528 |
| ACBD7 | 715.6840117 | 0.93812291 | 0.366666361 | 2.558519157 | 0.010511903 | 0.116405896 |
| ACCS | 332.8213956 | -0.477822061 | 0.369211962 | -1.294167335 | 0.195607627 | 0.501359174 |
| ACD | 437.8047966 | 0.171992861 | 0.301998661 | 0.569515308 | 0.569006485 | 0.805450586 |
| ACE | 26.21719924 | -3.258396074 | 0.818983051 | -3.978587923 | 6.93E-05 | 0.004731696 |
| ACER2 | 167.8175349 | -1.256117829 | 0.715850843 | -1.754720052 | 0.079307204 | 0.322931692 |
| ACER3 | 977.5605019 | -0.563538102 | 0.44002459 | -1.280696839 | 0.20030017 | 0.506121566 |
| ACHE | 32.3823172 | -0.977291828 | 0.887688721 | -1.100939783 | 0.270922866 | 0.583394591 |
| ACIN1 | 8720.503989 | -0.252211549 | 0.249758657 | -1.009821048 | 0.312581034 | 0.622907489 |
| ACKR2 | 17.76587374 | -1.050056389 | 0.844234126 | -1.243797612 | 0.21357406 | 0.520678417 |
| ACKR3 | 640.7223661 | -4.387516316 | 1.354568551 | -3.239050776 | 0.001199282 | 0.030476232 |
| ACLY | 10482.89383 | -0.427506441 | 0.305921746 | -1.397437243 | 0.162282126 | |

| | | | | | | |
|---|---|---|---|---|---|---|
| | | | | | | 0.46295264 |
| ACO1 | 541.8114599 | 0.227441468 | 0.438364429 | 0.518841068 | 0.603871577 | 0.825641353 |
| ACO2 | 1937.733669 | -0.088893583 | 0.29450214 | -0.301843589 | 0.762771301 | 0.901485479 |
| ACOT1 | 46.61093171 | -0.131414823 | 0.642259663 | -0.204613227 | 0.83787432 | 0.933631385 |
| ACOT11 | 150.0862127 | -0.644755287 | 0.428442515 | -1.50488167 | 0.132354502 | 0.418524946 |
| ACOT13 | 1273.037225 | 0.685799161 | 0.352671478 | 1.944583566 | 0.051825119 | 0.262457735 |
| ACOT2 | 221.4056199 | 0.224138997 | 0.450271682 | 0.497786128 | 0.618634795 | 0.83229541 |
| ACOT4 | 80.18374078 | -0.547585897 | 0.547883864 | -0.99945615 | 0.317573771 | 0.627634461 |
| ACOT7 | 1517.07917 | 0.760036676 | 0.269451644 | 2.820679303 | 0.004792208 | 0.073585621 |
| ACOT8 | 359.201647 | 0.565279582 | 0.40742087 | 1.387458582 | 0.165301972 | 0.467523531 |
| ACOT9 | 790.7424997 | 0.24480618 | 0.261335319 | 0.936751225 | 0.34888654 | 0.655555821 |
| ACOX1 | 572.2664151 | -0.705057673 | 0.374446538 | -1.882932814 | 0.059709474 | 0.282669434 |
| ACOX2 | 56.13978855 | 0.026957804 | 0.550068134 | 0.049008119 | 0.960912826 | 0.986559996 |
| ACOX3 | 311.3737716 | -0.130611759 | 0.338018433 | -0.38640425 | 0.699197303 | 0.871039767 |
| ACOXL | 373.4150148 | -0.59823595 | 0.403232997 | -1.483598698 | 0.137915415 | 0.426133352 |
| ACP1 | 3930.977229 | 0.453689003 | 0.274108716 | 1.655142565 | 0.097895582 | 0.359375073 |
| ACP2 | 761.8773064 | -0.586319528 | 0.298807455 | -1.962198458 | 0.049739383 | 0.25701505 |
| ACP5 | 115.8137816 | -0.742431247 | 0.531847676 | -1.39594715 | 0.162730411 | 0.463882574 |
| ACP6 | 403.5681213 | -0.085411488 | 0.27872024 | -0.306441642 | 0.759268407 | 0.899449351 |
| ACP7 | 10.98863864 | -0.67965944 | 0.994445365 | -0.683455787 | 0.494318879 | 0.760354598 |
| ACPP | 924.7112667 | -1.914924954 | 0.668054479 | -2.866420349 | 0.004151426 | 0.066435955 |
| ACRBP | 44.96701626 | -0.013611418 | 0.622864326 | -0.021852942 | 0.982565263 | 0.993518428 |
| ACSBG1 | 26.96222499 | 0.509066085 | 0.745979743 | 0.682412747 | 0.494977998 | 0.760514211 |
| ACSF2 | 710.9893018 | -0.314815061 | 0.379699029 | -0.829117372 | 0.407037994 | 0.701644519 |
| ACSF3 | 482.2912007 | -0.271789783 | 0.321352709 | -0.84576783 | 0.397682275 | 0.696357086 |
| ACSL1 | 64.26394221 | -0.821206383 | 0.921953912 | -0.890723898 | 0.373077311 | 0.678163896 |
| ACSL3 | 2468.868549 | 0.049888049 | 0.35419123 | 0.140850605 | 0.887987965 | 0.955943827 |
| ACSL4 | 1322.082773 | 0.332391426 | 0.333826965 | 0.99569975 | 0.319396052 | 0.629115732 |
| ACSL6 | 10.5089345 | 0.346452095 | 0.912521648 | 0.379664522 | 0.704194459 | 0.873421952 |
| ACSM2A | 74.13742882 | -0.487963447 | 0.792038704 | -0.616085356 | 0.537838187 | 0.786158891 |
| ACSM3 | 144.6323878 | 1.099175672 | 0.381305555 | 2.882663678 | 0.003943282 | 0.064934636 |
| ACSS1 | 54.32546525 | -1.3729339 | 0.896667452 | -1.531151708 | 0.125731903 | 0.407929576 |
| ACSS2 | 726.4407244 | -0.771983922 | 0.259211237 | -2.978203926 | 0.00289943 | 0.052996691 |
| ACSS3 | 1121.367118 | -0.323408395 | 0.259571656 | -1.245931084 | 0.212789697 | 0.519927926 |
| ACTA1 | 255.5247755 | 0.472479998 | 0.572483474 | 0.825316398 | 0.409191984 | 0.703508181 |
| ACTA2 | 513.9610225 | 1.249480041 | 0.391366065 | 3.192612117 | 0.001409922 | 0.034006242 |
| ACTB | 63582.07408 | 0.002405565 | 0.518431342 | 0.004640083 | 0.996297763 | 0.998431718 |
| ACTC1 | 263.138712 | -1.132465904 | 0.644431392 | -1.75731027 | 0.078864938 | 0.322507823 |
| ACTG1 | 76343.75564 | -0.064274509 | 0.50282707 | -0.12782627 | 0.89828646 | 0.960048173 |
| ACTG1P17 | 20.23604376 | 0.612135132 | 0.7910597 | 0.773816606 | 0.439039251 | |

0.722630418

| | | | | | | |
|---|---|---|---|---|---|---|
| ACTG2 | 12.55368637 | -0.348333387 | 1.05260583 | -0.330924812 | 0.740701279 | 0.891917214 |
| ACTL6A | 1893.489461 | 0.477762528 | 0.238596738 | 2.002384992 | 0.045243341 | 0.244986785 |
| ACTN1 | 5200.705168 | -0.531285432 | 0.372962479 | -1.424501022 | 0.154301489 | 0.451476933 |
| ACTN2 | 45.42714829 | -0.620585244 | 0.560969435 | -1.106272828 | 0.268608449 | 0.581542504 |
| ACTN3 | 449.6505896 | 0.65322741 | 0.624415924 | 1.046141498 | 0.295495708 | 0.60699582 |
| ACTN4 | 2458.719894 | -0.548372053 | 0.314955264 | -1.74111093 | 0.081664136 | 0.326743321 |
| ACTR10 | 1667.539334 | 0.476122936 | 0.30448113 | 1.563719024 | 0.117883568 | 0.395839103 |
| ACTR1A | 3421.758043 | -0.081811861 | 0.244679878 | -0.334362849 | 0.73810577 | 0.891242475 |
| ACTR1B | 1257.16852 | 0.213580949 | 0.321193608 | 0.66496015 | 0.506075982 | 0.767209743 |
| ACTR2 | 8342.786224 | -0.232681327 | 0.229039836 | -1.01589894 | 0.309677514 | 0.61915097 |
| ACTR3 | 4251.473078 | 0.058220063 | 0.222916656 | 0.261174126 | 0.793958231 | 0.914519802 |
| ACTR3B | 635.5308653 | 0.606916718 | 0.32229607 | 1.883103069 | 0.059686401 | 0.282669434 |
| ACTR5 | 287.1413217 | 0.301418765 | 0.335549908 | 0.89828296 | 0.369034715 | 0.674858445 |
| ACTR6 | 1248.486475 | 0.575062896 | 0.322574141 | 1.782730921 | 0.074630115 | 0.313779586 |
| ACTR8 | 752.6838559 | -0.415362012 | 0.280139215 | -1.482698568 | 0.138154515 | 0.426306273 |
| ACTRT3 | 32.90686348 | -0.155361559 | 0.604506088 | -0.25700578 | 0.797174311 | 0.915732089 |
| ACVR1 | 687.2473712 | -0.628212562 | 0.43648334 | -1.43925897 | 0.150077163 | 0.444740017 |
| ACVR1B | 1070.62508 | -0.483136481 | 0.278910915 | -1.732225075 | 0.083233496 | 0.3302996 |
| ACVR1C | 66.2151188 | -0.084245997 | 0.566909544 | -0.148605714 | 0.881864764 | 0.953217539 |
| ACVR2A | 399.5262148 | 0.206037888 | 0.349403726 | 0.589684289 | 0.55540233 | 0.796360966 |
| ACVR2B | 3792.831821 | -0.063370648 | 0.29958143 | -0.211530627 | 0.832473236 | 0.931375477 |
| ACVRL1 | 11.48482372 | 0.334427437 | 1.03195099 | 0.324072984 | 0.74588278 | 0.893892697 |
| ACY1 | 493.9078357 | 0.573792691 | 0.278356617 | 2.061358185 | 0.039268881 | 0.228431056 |
| ACYP1 | 312.962863 | 1.268368533 | 0.383136471 | 3.310487592 | 0.000931336 | 0.02597297 |
| ACYP2 | 155.746551 | 1.140551448 | 0.468268624 | 2.435677706 | 0.014863922 | 0.138598988 |
| AD000671.1 | 45.00397453 | 0.990741234 | 0.822884328 | 1.203986029 | 0.228594979 | 0.53738497 |
| AD000864.1 | 96.12778647 | -0.219543427 | 0.495552287 | -0.443027775 | 0.657745645 | 0.850760404 |
| ADA | 521.3526376 | 0.374805818 | 0.407407962 | 0.919976663 | 0.357584954 | 0.665144843 |
| ADA2 | 549.0380947 | -0.653886797 | 0.354054996 | -1.846850924 | 0.064768749 | 0.293666076 |
| ADAL | 369.26727 | 0.020061185 | 0.265590625 | 0.075534237 | 0.939789658 | 0.978003717 |
| ADAM10 | 4324.103554 | -0.136411036 | 0.230397454 | -0.592068329 | 0.553804834 | 0.796301122 |
| ADAM11 | 185.5319093 | 0.448777477 | 0.439741076 | 1.020549367 | 0.307467989 | 0.617319594 |
| ADAM12 | 26.33929798 | 3.19426042 | 1.359305428 | 2.349921036 | 0.018777394 | 0.1554129 |
| ADAM15 | 691.1820148 | 0.270466205 | 0.357353879 | 0.756858176 | 0.449134835 | 0.730485072 |
| ADAM17 | 2294.214247 | -0.235693112 | 0.233400541 | -1.009822477 | 0.312580349 | 0.622907489 |
| ADAM19 | 469.4003649 | -0.09223997 | 0.36776369 | -0.25081315 | 0.801958574 | 0.917465766 |
| ADAM22 | 326.3044821 | -0.244824603 | 0.434206583 | -0.5638436 | 0.572860576 | |

| | | | | | | |
|---|---|---|---|---|---|---|
| | | | | | | 0.807701469 |
| ADAM23 | 964.396424 | 0.334825132 | 0.42846775 | 0.781447685 | 0.434539234 | 0.719554351 |
| ADAM28 | 73.04360143 | -1.480291259 | 0.732556525 | -2.020719504 | 0.04330881 | 0.239441079 |
| ADAM32 | 36.7778452 | -1.111901303 | 1.029165057 | -1.08039162 | 0.279967826 | 0.593790959 |
| ADAM33 | 307.650316 | -0.432154773 | 0.389247058 | -1.110232601 | 0.266898808 | 0.580103412 |
| ADAM8 | 65.48393672 | -0.286015283 | 0.513712421 | -0.55676147 | 0.577690412 | 0.809936484 |
| ADAM9 | 1308.498968 | -0.748366158 | 0.744229902 | -1.005557766 | 0.314628348 | 0.624045314 |
| ADAMTS1 | 569.1928482 | -1.47891054 | 0.842988582 | -1.75436604 | 0.079367806 | 0.322931692 |
| ADAMTS10 | 94.14946422 | -0.198498819 | 0.524073034 | -0.378761748 | 0.704864795 | 0.873723655 |
| ADAMTS12 | 720.0286544 | -0.442354708 | 0.402793838 | -1.098216174 | 0.272110105 | 0.584314565 |
| ADAMTS13 | 138.1758127 | -0.465152205 | 0.435541613 | -1.067985677 | 0.285526973 | 0.599411008 |
| ADAMTS14 | 16.35732845 | -0.330031815 | 0.730306791 | -0.451908457 | 0.651334932 | 0.847876685 |
| ADAMTS16 | 18.96544829 | 0.739057459 | 0.736645398 | 1.003274385 | 0.315728491 | 0.625491438 |
| ADAMTS17 | 56.28670678 | -2.410699621 | 0.648014699 | -3.720131077 | 0.000199119 | 0.00940977 |
| ADAMTS18 | 338.3216798 | -2.77154246 | 1.53446379 | -1.806196065 | 0.070887719 | 0.306392476 |
| ADAMTS19 | 4576.353324 | -0.831444406 | 0.261247328 | -3.182594869 | 0.001459617 | 0.034762713 |
| ADAMTS2 | 169.6732481 | -1.497111434 | 0.424567482 | -3.526203724 | 0.000421563 | 0.015778519 |
| ADAMTS20 | 123.088635 | -1.657311204 | 0.799606831 | -2.072657636 | 0.038204153 | 0.224998139 |
| ADAMTS3 | 147.7189436 | 0.036474819 | 0.373211333 | 0.097732344 | 0.922144831 | 0.970640219 |
| ADAMTS4 | 163.1171059 | 1.774919797 | 0.6122202 | 2.89915262 | 0.003741727 | 0.063073047 |
| ADAMTS5 | 38.39708082 | 0.696221027 | 0.99680939 | 0.698449507 | 0.484896125 | 0.755350561 |
| ADAMTS6 | 177.8232145 | -1.0063619 | 0.434169131 | -2.317902931 | 0.020454597 | 0.161737081 |
| ADAMTS7 | 785.0517776 | 0.358679401 | 0.593845551 | 0.603994422 | 0.54584735 | 0.791658199 |
| ADAMTS9 | 121.9861216 | -0.295373237 | 0.775163723 | -0.381046259 | 0.703168923 | 0.872833885 |
| ADAMTSL2 | 36.70736363 | -0.953092969 | 0.695379993 | -1.370607406 | 0.170497373 | 0.473631094 |
| ADAMTSL3 | 88.78636006 | 0.690924558 | 0.518017106 | 1.333787149 | 0.182273624 | 0.486415681 |
| ADAMTSL4 | 478.8581656 | -3.475873914 | 0.885946021 | -3.923347283 | 8.73E-05 | 0.005590042 |
| ADAMTSL5 | 21.43184515 | -0.174810025 | 0.684860512 | -0.255249094 | 0.798530715 | 0.916056735 |
| ADAP1 | 16.85126249 | 0.314919073 | 0.837504793 | 0.376020621 | 0.706901566 | 0.87508797 |
| ADAP2 | 21.93251889 | -0.979231164 | 0.773539176 | -1.26591024 | 0.20554522 | 0.511932925 |
| ADAR | 6521.242896 | -0.41248817 | 0.349884269 | -1.178927453 | 0.238427065 | 0.549388973 |
| ADARB1 | 330.6323098 | -0.351345887 | 0.362738361 | -0.968593136 | 0.33274823 | 0.640543509 |
| ADARB2 | 12.42600362 | -0.186499249 | 0.964656484 | -0.193332292 | 0.846698738 | 0.938145276 |

| | | | | | | |
|---|---|---|---|---|---|---|
| ADAT1 | 445.4257446 | -0.290737211 | 0.414540986 | -0.701347325 | 0.483086286 | 0.753955635 |
| ADAT2 | 346.8770328 | 0.43254171 | 0.388707184 | 1.112770044 | 0.265807204 | 0.579572403 |
| ADAT3 | 27.02934904 | -0.348084958 | 1.139772572 | -0.305398609 | 0.760062581 | 0.899727639 |
| ADCK1 | 222.8546492 | -0.015420922 | 0.328735081 | -0.046909876 | 0.962585057 | 0.987448637 |
| ADCK2 | 256.0314691 | -0.829079782 | 0.411306878 | -2.015720686 | 0.043829191 | 0.240941849 |
| ADCK5 | 88.89513708 | -0.416022909 | 0.546104028 | -0.761801577 | 0.446178439 | 0.728589711 |
| ADCY1 | 182.1231792 | 0.377195448 | 0.578735999 | 0.651757362 | 0.514557712 | 0.77285333 |
| ADCY10 | 44.09488474 | -4.226942644 | 1.060304751 | -3.986535606 | 6.70E-05 | 0.004643142 |
| ADCY10P1 | 110.3326998 | -0.798658289 | 0.427668735 | -1.86746943 | 0.061836062 | 0.286447296 |
| ADCY2 | 78.86773724 | 0.787484393 | 0.660076234 | 1.19302037 | 0.232861395 | 0.542441738 |
| ADCY3 | 658.5390919 | 0.266767003 | 0.440514293 | 0.605580811 | 0.544793148 | 0.790763189 |
| ADCY5 | 71.13864997 | -0.852665695 | 0.487335826 | -1.749647059 | 0.080179234 | 0.324202401 |
| ADCY6 | 1324.091564 | -0.298346083 | 0.325061672 | -0.917813785 | 0.358716364 | 0.666432275 |
| ADCY7 | 201.9436439 | -0.573656663 | 0.439553052 | -1.305090844 | 0.191861916 | 0.497612799 |
| ADCY9 | 115.8342035 | 0.823143738 | 0.796652035 | 1.033253794 | 0.301485144 | 0.612375301 |
| ADCYAP1R1 | 180.1599358 | 0.682444641 | 0.699454417 | 0.975681366 | 0.329222389 | 0.638424836 |
| ADD1 | 4396.319697 | -0.691111007 | 0.374843228 | -1.843733475 | 0.065221992 | 0.294576612 |
| ADD2 | 1630.402623 | 0.338292569 | 0.444981355 | 0.760239874 | 0.447111214 | 0.728897941 |
| ADD3 | 4271.136553 | 0.012187965 | 0.274258867 | 0.04443964 | 0.964553965 | 0.988532608 |
| ADGRA2 | 20.85027841 | 0.653998418 | 0.962761005 | 0.679294669 | 0.496951174 | 0.761325016 |
| ADGRA3 | 3312.48986 | -0.15478455 | 0.278606348 | -0.555567203 | 0.578506757 | 0.810209453 |
| ADGRB1 | 33.99033657 | 0.116840626 | 0.818546768 | 0.14274154 | 0.886494308 | 0.95525287 |
| ADGRB2 | 249.1828173 | -0.26835346 | 0.519204671 | -0.516854865 | 0.605257477 | 0.826077544 |
| ADGRD1 | 21.49188997 | -0.891412879 | 0.763540581 | -1.167472825 | 0.243019477 | 0.554163307 |
| ADGRD2 | 11.84221998 | 0.645264163 | 0.845787277 | 0.762915429 | 0.445513834 | 0.728328705 |
| ADGRE2 | 85.55617136 | -0.343659598 | 0.643619781 | -0.533948161 | 0.5933774 | 0.819865987 |
| ADGRE5 | 285.548213 | -0.63882193 | 0.341801182 | -1.868986898 | 0.061624637 | 0.286166933 |
| ADGRG1 | 1128.593232 | -1.645970479 | 0.29554679 | -5.569238227 | 2.56E-08 | 1.55E-05 |
| ADGRG2 | 17.90215917 | -1.218598302 | 0.771859482 | -1.578782577 | 0.114385936 | 0.38927175 |
| ADGRG3 | 29.54844733 | 0.17053082 | 0.874299451 | 0.195048527 | 0.845354958 | 0.937778401 |
| ADGRG5 | 70.53275252 | -0.197055943 | 0.399986814 | -0.492656097 | 0.622255603 | 0.833640266 |
| ADGRG6 | 285.9330494 | -3.875493448 | 0.788109682 | -4.917454429 | 8.77E-07 | 0.000170531 |
| ADGRL1 | 2803.829568 | 0.51547817 | 0.354543386 | 1.453921272 | 0.145968135 | 0.438251053 |
| ADGRL2 | 5250.302325 | 0.095720392 | 0.412857556 | 0.231848469 | 0.81665571 | 0.923554512 |
| ADGRL3 | 618.3860473 | 0.625882768 | 0.466940765 | 1.340390076 | 0.18011856 | 0.484754911 |
| ADGRV1 | 1397.671601 | 0.25267471 | 0.335208079 | 0.753784667 | 0.450978533 | |

0.732208864

| | | | | | | |
|---|---|---|---|---|---|---|
| ADH5 | 9285.529319 | 0.235016694 | 0.228158501 | 1.030058897 | 0.302982359 | 0.613538084 |
| ADHFE1 | 332.2901325 | -0.603667015 | 0.289719886 | -2.083622988 | 0.037194474 | 0.220938673 |
| ADI1 | 1504.745917 | -0.184923698 | 0.265226424 | -0.697229543 | 0.485659153 | 0.755840687 |
| ADIPOR1 | 1750.624824 | 0.303388897 | 0.238897045 | 1.269956674 | 0.204100064 | 0.510452039 |
| ADIPOR2 | 767.2201693 | -0.14762902 | 0.368883626 | -0.40020486 | 0.689005635 | 0.867388971 |
| ADK | 1013.015912 | 0.850526318 | 0.330579273 | 2.572836191 | 0.010086894 | 0.114259456 |
| ADM | 106.212575 | -0.732863151 | 0.625388075 | -1.171853414 | 0.241255913 | 0.552673201 |
| ADM2 | 152.8457044 | 0.440484501 | 0.39367476 | 1.118904601 | 0.26318084 | 0.576857184 |
| ADM5 | 41.28277226 | -0.495991664 | 0.690028245 | -0.718799074 | 0.472264728 | 0.746559153 |
| ADNP | 5171.312996 | -0.489956038 | 0.315688905 | -1.552021722 | 0.120657026 | 0.400973879 |
| ADNP2 | 1698.255544 | 0.01262364 | 0.289945923 | 0.043537912 | 0.965272744 | 0.988700292 |
| ADO | 1009.428328 | 0.607356536 | 0.267693029 | 2.268854507 | 0.023277175 | 0.174130017 |
| ADORA1 | 32.97854216 | 1.664170364 | 0.578217598 | 2.87810397 | 0.004000733 | 0.065575642 |
| ADORA2A | 54.26311666 | 0.502946749 | 0.610112309 | 0.824351094 | 0.409740095 | 0.703966274 |
| ADORA2B | 148.4031389 | -1.073519363 | 0.506990395 | -2.117435308 | 0.03422292 | 0.211055253 |
| ADPGK | 990.611161 | -0.271864674 | 0.311849811 | -0.871780786 | 0.383327978 | 0.684494911 |
| ADPRH | 76.14381129 | -0.462949399 | 0.427595701 | -1.082680199 | 0.278950396 | 0.592545009 |
| ADPRHL1 | 40.10560417 | -0.6376242 | 0.625586059 | -1.019242981 | 0.308087626 | 0.617841234 |
| ADPRHL2 | 604.5998213 | 0.29192864 | 0.283849222 | 1.028463768 | 0.303731726 | 0.614002998 |
| ADPRM | 230.6650284 | 0.467868222 | 0.329028248 | 1.421969766 | 0.155035028 | 0.452837199 |
| ADRA2A | 17.46872638 | 0.741759057 | 1.35045029 | 0.549267946 | 0.582821581 | 0.813654754 |
| ADRA2B | 221.7261612 | 0.557299364 | 0.543252694 | 1.025856605 | 0.30495918 | 0.615135352 |
| ADRA2C | 149.7369248 | 0.866428175 | 0.520414528 | 1.664880837 | 0.095936532 | 0.355595682 |
| ADRB2 | 30.63912608 | 0.408084557 | 0.623604865 | 0.654396045 | 0.512856682 | 0.771819984 |
| ADRM1 | 1345.543283 | 0.121250582 | 0.386198902 | 0.313958899 | 0.753552256 | 0.896919918 |
| ADSL | 3308.915466 | 0.289353885 | 0.273729113 | 1.057081149 | 0.290474554 | 0.603504445 |
| ADSS | 2420.271002 | 0.501727239 | 0.257613556 | 1.947596417 | 0.051463273 | 0.261207532 |
| ADSSL1 | 325.2516683 | 0.552775013 | 0.318491771 | 1.735602181 | 0.0826342 | 0.328745819 |
| AEBP1 | 452.059806 | 0.154091667 | 0.419776509 | 0.367080253 | 0.713559157 | 0.878044121 |
| AEBP2 | 936.496579 | 0.241924811 | 0.250638664 | 0.965233406 | 0.334427918 | 0.642472578 |
| AEN | 1694.176364 | 0.675940217 | 0.387921846 | 1.74246494 | 0.081427121 | 0.326289185 |
| AES | 787.8642754 | 0.853908999 | 0.284084087 | 3.005831862 | 0.002648554 | 0.050167035 |
| AF228730.5 | 84.72839191 | 0.255156308 | 0.400517491 | 0.637066579 | 0.524081477 | 0.777662198 |
| AF274858.1 | 14.19592215 | 2.420771345 | 0.886355656 | 2.731151235 | 0.00631135 | 0.086338588 |
| AFAP1 | 1620.836153 | -1.328154456 | 0.246263866 | -5.393216952 | 6.92E-08 | 3.12E-05 |
| AFAP1L1 | 268.3010508 | -3.427893338 | 0.714608236 | -4.796884736 | 1.61E-06 | 0.000260089 |
| AFAP1L2 | 596.867205 | -1.069689368 | 0.56861594 | -1.881215937 | 0.059942551 | 0.283136022 |
| AFDN | 7415.607821 | -0.555679515 | 0.303089331 | -1.83338527 | 0.066745297 | 0.297817842 |
| AFF1 | 2013.160959 | -0.170084097 | 0.458228092 | -0.371177804 | 0.710505104 | |

0.876838602

| | | | | | | |
|---|---|---|---|---|---|---|
| AFF2 | 377.569708 | 0.170298864 | 0.581055778 | 0.293085227 | 0.769457009 | 0.904597192 |
| AFF3 | 154.5879889 | 2.387612243 | 1.026233217 | 2.326578602 | 0.019987704 | 0.159764734 |
| AFF4 | 2988.975682 | -0.622518251 | 0.378814743 | -1.643331632 | 0.100314337 | 0.36243601 |
| AFG1L | 170.2513994 | -0.30611462 | 0.320661347 | -0.954635232 | 0.339762194 | 0.646629378 |
| AFG3L1P | 620.4813528 | -0.443168995 | 0.276501958 | -1.602769825 | 0.108985481 | 0.379241044 |
| AFG3L2 | 1940.039805 | 0.307127513 | 0.254127615 | 1.208556236 | 0.226833378 | 0.535481989 |
| AFMID | 510.8662149 | 0.703676867 | 0.276918453 | 2.541097785 | 0.011050501 | 0.11934855 |
| AFTPH | 718.8773386 | 0.012857116 | 0.239430022 | 0.053698846 | 0.957175102 | 0.984903668 |
| AGA | 346.3942094 | 0.142606853 | 0.370418411 | 0.384988566 | 0.700245887 | 0.871916477 |
| AGAP1 | 1287.604599 | -0.071553611 | 0.300590338 | -0.238043615 | 0.811847267 | 0.921333624 |
| AGAP10P | 59.87857092 | -0.597806617 | 0.530123476 | -1.127674294 | 0.259457498 | 0.572792447 |
| AGAP12P | 94.95051487 | -0.1129622 | 0.56154154 | -0.201164459 | 0.840569981 | 0.934712833 |
| AGAP14P | 20.63789944 | -0.341366805 | 0.801505599 | -0.42590695 | 0.670175651 | 0.857922099 |
| AGAP2 | 61.60970528 | 0.376848578 | 0.594866681 | 0.633500899 | 0.526406594 | 0.778525486 |
| AGAP3 | 1690.594142 | 0.26025594 | 0.369546546 | 0.704257536 | 0.481272406 | 0.752722364 |
| AGAP4 | 349.4069427 | -0.191183267 | 0.297585495 | -0.642448204 | 0.52058221 | 0.775597169 |
| AGAP5 | 179.4284189 | -0.642965854 | 0.345252681 | -1.862305176 | 0.062560086 | 0.288484612 |
| AGAP6 | 408.0551991 | -0.065427046 | 0.279551266 | -0.234043103 | 0.814951516 | 0.922747905 |
| AGAP7P | 26.17491345 | 0.40935512 | 0.756375031 | 0.541206549 | 0.588365224 | 0.817482262 |
| AGAP9 | 203.9839919 | -0.679518509 | 0.361909654 | -1.877591551 | 0.060437063 | 0.28404296 |
| AGBL2 | 59.13723584 | 0.125436088 | 0.491062224 | 0.255438276 | 0.798384611 | 0.916056735 |
| AGBL3 | 58.78124845 | 0.549483588 | 0.510740614 | 1.075856458 | 0.281991458 | 0.595751624 |
| AGBL4 | 26.51822449 | -0.112805996 | 0.646967114 | -0.174361253 | 0.861581564 | 0.94403207 |
| AGBL5 | 2457.404705 | -0.092391764 | 0.334269075 | -0.276399376 | 0.782241336 | 0.909909835 |
| AGER | 57.58458022 | 2.934790502 | 1.070090577 | 2.7425627 | 0.006096181 | 0.084270328 |
| AGFG1 | 2093.823064 | -0.066741901 | 0.271581904 | -0.245752388 | 0.805873916 | 0.919034291 |
| AGFG2 | 86.76850879 | -0.061202256 | 0.548183912 | -0.11164548 | 0.911104509 | 0.966866396 |
| AGGF1 | 1240.87623 | -0.142363277 | 0.245117579 | -0.58079587 | 0.561378039 | 0.801493152 |
| AGK | 711.7445882 | 0.098125465 | 0.319908762 | 0.306729533 | 0.759049249 | 0.899440455 |
| AGL | 1739.146447 | -0.840850109 | 0.26923718 | -3.123083183 | 0.001789672 | 0.039332648 |
| AGMAT | 23.1559548 | 0.541671683 | 0.690943235 | 0.783959746 | 0.43306373 | 0.718657207 |
| AGMO | 46.32821736 | -0.849792141 | 0.50456084 | -1.684221353 | 0.092138898 | 0.348501426 |
| AGO1 | 1462.354334 | -0.37343727 | 0.335736666 | -1.112292186 | 0.266012543 | 0.579643865 |
| AGO2 | 1720.343686 | -0.094871587 | 0.288294823 | -0.329078359 | 0.742096461 | 0.892150449 |
| AGO3 | 538.5461152 | -0.317408987 | 0.320715905 | -0.989688947 | 0.322326179 | 0.632029271 |
| AGO4 | 904.7954486 | -0.43269347 | 0.277463301 | -1.559461984 | 0.118887076 | 0.397626945 |
| AGPAT2 | 144.1690436 | -0.138113242 | 0.567248208 | -0.24347938 | 0.807634057 | 0.919453224 |
| AGPAT3 | 229.5565771 | -1.494235724 | 0.458665912 | -3.257786735 | | |

0.001122848 0.029019967
AGPAT4 2390.840662 0.096716603 0.243259886 0.397585497 0.690935753
0.868004248
AGPAT5 1022.059418 0.819165653 0.303653608 2.697697744 0.00698208
0.091869159
AGPS 2110.27832 0.042174409 0.267108766 0.157892268 0.874541691 0.949740014
AGRN 4703.008323 -0.173406913 0.389143607 -0.445611619 0.655877816
0.849801208
AGTPBP1 1853.629083 0.46637336 0.36162844 1.289647905 0.197172935
0.50258876
AGTRAP 1077.688916 -0.012162211 0.349353573 -0.034813471
0.972228478 0.989954357
AHCTF1 1654.319125 0.143076078 0.434302328 0.329438893 0.741823975
0.892150449
AHCY 9357.209741 0.233903404 0.241335418 0.969204628 0.332443103 0.640222942
AHCYL1 4411.334399 -0.199312991 0.260860202 -0.764060557
0.444831156 0.727526247
AHCYL2 1120.795415 -0.333824958 0.314443477 -1.061637408
0.288400325 0.601673803
AHDC1 1729.389286 -0.16682368 0.398986961 -0.418118124 0.675860754
0.860110127
AHI1 441.3095302 0.244744582 0.332871257 0.735252973 0.462185443 0.739799045
AHNAK 12425.89212 -2.107573782 0.439311867 -4.797443319 1.61E-06
0.000260089
AHNAK2 97.94757902 -0.890180247 0.631907214 -1.408719868
0.158918018 0.458846746
AHR 182.9277469 -1.495167509 0.551326761 -2.711944374 0.006688982
0.089444885
AHRR 29.35143601 -0.716138759 0.611697993 -1.170739103 0.241703663
0.552977821
AHSA1 1897.293044 0.198214492 0.290526901 0.682258654 0.495075412 0.760514211
AHSA2P 1417.513545 0.422029533 0.233905981 1.804269952 0.071188998
0.306992555
AIDA 3698.796925 0.066169769 0.235700056 0.280737179 0.77891199 0.908299392
AIF1L 7541.675808 -0.560859049 0.339974172 -1.649710758 0.099002109
0.36122198
AIFM1 1143.197891 0.20262251 0.250557842 0.808685564 0.418696034 0.709782331
AIFM2 70.61909961 -0.083248044 0.494279323 -0.168423076 0.866250455
0.9456635
AIFM3 49.46552071 0.785494298 0.548764003 1.431388162 0.152318999 0.448182998
AIG1 704.4907657 0.223953042 0.32577296 0.687451292 0.491798385 0.759010508
AIM2 47.45468169 0.120328373 0.492374637 0.244383777 0.806933605 0.919137032
AIMP1 1704.792448 0.55744528 0.31137274 1.790282859 0.07340845 0.311344591
AIMP2 1132.969564 0.758657655 0.350563522 2.164108948 0.030455989 0.199054492
AIP 798.1639712 0.775547312 0.345464464 2.244940921 0.024771925 0.17998797
AIPL1 84.56225948 -0.85925833 0.841444122 -1.021170994 0.307173432
0.617205881
AJAP1 33.33260031 -1.493724969 0.915323618 -1.63190913 0.10269863
0.367636293
AJM1 16.49021812 0.784074614 0.940060819 0.834067964 0.404242686 0.700367583
AJUBA 2684.336162 0.143980857 0.314840088 0.457314245 0.647445193 0.846597949
AK1 384.0594944 0.91454277 0.330441706 2.767637232 0.005646427 0.080660961
AK2 1937.393758 -0.015512381 0.242375869 -0.064001343 0.948969157
0.98134156
AK3 1628.927173 0.297818558 0.345886399 0.861029977 0.389221533 0.689097897
AK4 662.142946 0.675195267 0.435186006 1.551509601 0.120779608 0.401039054
AK6 328.0092745 1.196603668 0.308305783 3.881223562 0.000103932 0.006332354
AK7 167.5518855 0.460959796 0.418906109 1.100389292 0.271162542 0.583394591
AK8 29.47639743 0.161935382 0.623908771 0.259549775 0.795211082 0.915124686
AK9 152.6463411 -0.089001169 0.378569112 -0.235098866 0.814131999
0.922281702
AKAP1 711.3986196 0.040795149 0.370608355 0.110076172 0.912348965 0.967315188
AKAP10 746.8498671 -0.046766284 0.260768543 -0.179340204
0.85767058 0.942343063

| | | | | | | |
|---|---|---|---|---|---|---|
| AKAP11 | 1025.683102 | 0.127330862 | 0.403490554 | 0.31557334 | 0.752326376 | 0.896586461 |
| AKAP12 | 2376.111187 | 0.221881975 | 0.456804314 | 0.485726531 | 0.627161071 | 0.836041947 |
| AKAP13 | 1732.491474 | -0.393670664 | 0.411134807 | -0.957522099 | 0.338303797 | 0.645504579 |
| AKAP17A | 1070.264007 | -0.027898818 | 0.29538299 | -0.094449644 | 0.924751982 | 0.971780852 |
| AKAP2 | 1163.489643 | -0.06865063 | 0.300170169 | -0.228705705 | 0.819097661 | 0.924316427 |
| AKAP3 | 68.89811701 | -1.123069993 | 0.433070866 | -2.593270714 | 0.009506791 | 0.110434548 |
| AKAP5 | 111.730942 | -0.252939318 | 0.386594637 | -0.65427529 | 0.512934463 | 0.771819984 |
| AKAP6 | 62.76937948 | 0.175883637 | 0.433753578 | 0.405492073 | 0.685115844 | 0.864428848 |
| AKAP7 | 181.9813768 | 0.110495602 | 0.32517912 | 0.339799191 | 0.734007757 | 0.88891398 |
| AKAP8 | 1069.20611 | -0.256779279 | 0.254386728 | -1.00940517 | 0.31278036 | 0.622907489 |
| AKAP8L | 2635.171766 | -0.164564035 | 0.394705692 | -0.416928456 | 0.676730739 | 0.860288423 |
| AKAP9 | 1472.616993 | -0.335382618 | 0.282462418 | -1.187353067 | 0.235088384 | 0.545674093 |
| AKIP1 | 380.6706484 | 0.90157445 | 0.276015074 | 3.266395694 | 0.001089259 | 0.028511711 |
| AKIRIN1 | 4358.593261 | 0.068998811 | 0.257090753 | 0.268383091 | 0.788404453 | 0.912309638 |
| AKIRIN2 | 1265.72821 | 0.390561756 | 0.318233026 | 1.227282288 | 0.21971651 | 0.528259775 |
| AKNA | 591.580823 | -0.609440674 | 0.466019146 | -1.307758877 | 0.190955106 | 0.496058205 |
| AKR1A1 | 4755.201302 | 0.832989229 | 0.327514347 | 2.543367146 | 0.01097898 | 0.119221957 |
| AKR1B1 | 3969.977599 | 0.668451389 | 0.271248406 | 2.464351401 | 0.013726147 | 0.133230564 |
| AKR1B10 | 10.09738529 | 0.505171797 | 1.165965939 | 0.433264627 | 0.664822533 | 0.855037526 |
| AKR1B15 | 60.6407023 | 0.2098829 | 0.634245739 | 0.330917319 | 0.74070694 | 0.891917214 |
| AKR1C1 | 97.51228105 | 0.089728484 | 0.396174082 | 0.226487516 | 0.820822268 | 0.925132406 |
| AKR1C2 | 113.3480976 | 0.116886557 | 0.519280132 | 0.225093451 | 0.821906575 | 0.92563193 |
| AKR1C3 | 71.10097989 | 0.480466087 | 0.511235924 | 0.939812843 | 0.34731357 | 0.653977184 |
| AKR1D1 | 22.7891697 | -2.801515909 | 1.169832836 | -2.394800199 | 0.016629424 | 0.146056278 |
| AKR1E2 | 71.10655694 | 0.222676226 | 0.448381661 | 0.496622065 | 0.61945559 | 0.832691898 |
| AKR7A2 | 975.4407304 | 0.688358801 | 0.376356952 | 1.829005141 | 0.067398838 | 0.298244514 |
| AKR7L | 26.16398008 | -0.213134425 | 0.661189701 | -0.322349887 | 0.747187644 | 0.89418504 |
| AKT1 | 2051.008181 | -0.259870849 | 0.284712558 | -0.91274811 | 0.361375035 | 0.668292036 |
| AKT1S1 | 947.313156 | -0.194544205 | 0.28251487 | -0.688615805 | 0.491065072 | 0.758593654 |
| AKT2 | 2495.534544 | -0.317583922 | 0.439882416 | -0.721974578 | 0.470310107 | 0.7454951 |
| AKT3 | 389.3208648 | 1.129920044 | 0.462359335 | 2.44381363 | 0.014532932 | 0.137370161 |
| AKTIP | 898.1968728 | -0.54499098 | 0.346760928 | -1.571662017 | 0.116028955 | 0.392480553 |
| AL021920.2 | 16.092974 | -1.644393456 | 0.961965091 | -1.709410737 | 0.087374893 | 0.339540089 |
| AL022238.4 | 128.6058274 | -1.327402478 | 2.06229581 | -0.643652803 | 0.519800602 | 0.775026528 |

| | | | | | | |
|---|---|---|---|---|---|---|
| AL022318.4 | 11.36951877 | 3.223107589 | 1.755075058 | 1.836449999 | 0.066291131 | 0.297369232 |
| AL031281.3 | 182.9045838 | 0.447550586 | 0.410084937 | 1.091360705 | 0.275114196 | 0.588252711 |
| AL031315.1 | 24.30841734 | -0.184485862 | 0.767255053 | -0.240449198 | 0.809982042 | 0.920260433 |
| AL031432.2 | 10.88482601 | 0.564089125 | 1.386776342 | 0.40676287 | 0.68418216 | 0.863826466 |
| AL031708.1 | 60.2940734 | -1.133499831 | 0.660293856 | -1.716659667 | 0.086041374 | 0.33625288 |
| AL033529.1 | 60.63948427 | -0.777604504 | 0.537262523 | -1.447345518 | 0.147800177 | 0.44174616 |
| AL034430.1 | 175.3443388 | 2.163082129 | 0.770937492 | 2.805781467 | 0.005019472 | 0.075621065 |
| AL034430.2 | 55.89480715 | 0.82161307 | 0.440983566 | 1.863137616 | 0.062442907 | 0.288202417 |
| AL035078.4 | 499.0262102 | 0.038632274 | 2.47177912 | 0.015629339 | 0.987530099 | 0.995541774 |
| AL035425.2 | 44.80790598 | -1.605030721 | 0.864090409 | -1.857480078 | 0.063242883 | 0.289778852 |
| AL035461.3 | 88.76261813 | -1.981042642 | 2.107630834 | -0.939938157 | 0.347249284 | 0.653937299 |
| AL049697.1 | 33.29694181 | -1.311429563 | 0.960673732 | -1.365114419 | 0.172217091 | 0.475642725 |
| AL096711.2 | 175.4009754 | 1.043038142 | 0.925855668 | 1.126566675 | 0.259925735 | 0.573325579 |
| AL109811.3 | 107.4902206 | 0.871290775 | 0.688177534 | 1.266084305 | 0.205482901 | 0.511932925 |
| AL109918.1 | 365.8434369 | 0.524116955 | 0.45064539 | 1.16303632 | 0.244814767 | 0.556250536 |
| AL117339.5 | 45.44402567 | -0.00901295 | 0.851881674 | -0.010580049 | 0.9915585 | 0.996814923 |
| AL121594.1 | 65.7074218 | 0.314086514 | 0.838791288 | 0.37445133 | 0.70806856 | 0.875585221 |
| AL121768.1 | 159.578994 | -2.984601786 | 0.590118674 | -5.057629793 | 4.24E-07 | 0.000111036 |
| AL121845.3 | 80.35997958 | 0.341372015 | 0.759857673 | 0.449257838 | 0.653245669 | 0.848827707 |
| AL132780.3 | 354.4112543 | 1.887988446 | 0.651724229 | 2.896913081 | 0.003768541 | 0.063384194 |
| AL133216.2 | 42.76185379 | -1.751956727 | 0.674261922 | -2.598332592 | 0.009367771 | 0.109379514 |
| AL133352.1 | 29.34505006 | 1.080278184 | 0.981681445 | 1.100436591 | 0.271141943 | 0.583394591 |
| AL133353.2 | 47.78807234 | -0.652799911 | 0.48459257 | -1.347110855 | 0.177944532 | 0.482243034 |
| AL133461.1 | 36.90468539 | -0.518045346 | 1.249629403 | -0.414559184 | 0.678464633 | 0.861138002 |
| AL133517.1 | 17.76668271 | 0.198531348 | 0.766708815 | 0.258939697 | 0.795681769 | 0.915261082 |
| AL135905.2 | 1250.174033 | -0.726080729 | 0.367304374 | -1.976782148 | 0.048066261 | 0.252967897 |
| AL136038.4 | 11.23390913 | -1.65963358 | 1.306097327 | -1.270681399 | 0.203842016 | 0.509986087 |
| AL136116.3 | 13.66902672 | -0.073234157 | 0.87951102 | -0.083266901 | 0.933639318 | 0.974827398 |
| AL136295.1 | 210.5934922 | -0.677456082 | 0.662151748 | -1.023113032 | 0.306254406 | 0.616286722 |
| AL136295.4 | 14.73170529 | -0.63329758 | 1.369705726 | -0.462360322 | 0.643822948 | 0.845466381 |
| AL136295.5 | 84.9519109 | -0.534500453 | 0.895940943 | -0.596580006 | 0.550787823 | 0.794461475 |
| AL136981.2 | 13.52032861 | -1.39377571 | 0.906257752 | -1.537946249 | 0.124061758 | 0.405284438 |

| | | | | | | |
|---|---|---|---|---|---|---|
| AL136982.4 | 18.78084913 | 0.138619767 | 0.714340424 | 0.194052811 | 0.846134531 | 0.938145276 |
| AL137058.1 | 25.03141108 | 0.129372519 | 0.721924306 | 0.179205102 | 0.857776658 | 0.942343063 |
| AL138686.2 | 53.89432645 | -0.584746534 | 0.864247957 | -0.676595795 | 0.498662448 | 0.762775559 |
| AL138752.2 | 12.59360225 | 1.028937054 | 1.390836776 | 0.739797129 | 0.459423101 | 0.737476588 |
| AL139011.2 | 27.33793047 | -0.566003907 | 0.89815573 | -0.630184597 | 0.528573816 | 0.779660133 |
| AL139082.2 | 28.46040379 | 0.481773457 | 0.784985968 | 0.613735119 | 0.539390377 | 0.786987056 |
| AL139260.3 | 84.18471089 | 2.222864982 | 0.73621412 | 3.019318596 | 0.00253344 | 0.048651661 |
| AL139300.1 | 701.5541425 | -0.295668245 | 0.421221724 | -0.70193019 | 0.482722701 | 0.753667396 |
| AL139353.1 | 10.81368573 | 0.482614444 | 1.019450025 | 0.473406672 | 0.635923071 | 0.841016203 |
| AL157392.5 | 49.19679016 | 0.212481569 | 0.529948055 | 0.400947917 | 0.68845847 | 0.867059891 |
| AL157935.2 | 102.2235127 | 1.041643554 | 0.619395912 | 1.681708796 | 0.092625321 | 0.3491405 |
| AL162171.1 | 714.8666189 | -0.412515904 | 0.372446974 | -1.10758291 | 0.268041987 | 0.581152025 |
| AL162231.1 | 118.5421499 | 1.290723497 | 0.493594579 | 2.614946665 | 0.008924139 | 0.10668882 |
| AL162417.1 | 13.64185019 | 0.356631208 | 4.181878417 | 0.085280147 | 0.932038674 | 0.974430344 |
| AL353625.1 | 39.02986807 | -0.96805484 | 0.559473223 | -1.730297 | 0.083577225 | 0.330798352 |
| AL353743.1 | 94.70333508 | -0.252730129 | 0.444676485 | -0.56834606 | 0.569800009 | 0.806129205 |
| AL353813.1 | 31.15245369 | 1.320165875 | 0.785650256 | 1.68034805 | 0.092889617 | 0.349511008 |
| AL354751.1 | 34.90998432 | 0.020031965 | 0.601937342 | 0.033279153 | 0.973451978 | 0.99055872 |
| AL354798.1 | 21.09212447 | 2.947147893 | 1.609750765 | 1.83081006 | 0.067128898 | 0.297817842 |
| AL355877.1 | 21.09197317 | 0.653875465 | 0.886734992 | 0.737396709 | 0.460881136 | 0.738646494 |
| AL355916.3 | 33.3549901 | -0.81020738 | 0.600811039 | -1.348522792 | 0.177490294 | 0.481837595 |
| AL356235.1 | 10.9165995 | -0.285521284 | 0.97498991 | -0.292845374 | 0.769640345 | 0.904694332 |
| AL356432.3 | 104.5716383 | 8.019437558 | 1.507772439 | 5.318732027 | 1.04E-07 | 3.96E-05 |
| AL356534.1 | 42.32806957 | -0.038905738 | 0.546624488 | -0.071174524 | 0.943258857 | 0.978607681 |
| AL358113.1 | 18.48103335 | -0.446059599 | 0.939382143 | -0.474843601 | 0.634898453 | 0.840313679 |
| AL358472.6 | 20.46232724 | -1.102499731 | 0.890481435 | -1.23809401 | 0.215681205 | 0.523284752 |
| AL360181.3 | 16.16232152 | 3.522595072 | 4.190453102 | 0.84062391 | 0.40055866 | 0.698117882 |
| AL365205.1 | 291.8713863 | -1.229846938 | 0.608950238 | -2.019618127 | 0.043423013 | 0.239726164 |
| AL390334.1 | 34.28862775 | 0.006080674 | 0.57235397 | 0.010623974 | 0.991523455 | 0.996814923 |
| AL390719.1 | 79.35302884 | 0.260960329 | 0.475505777 | 0.548805801 | 0.583138729 | 0.81387283 |
| AL390728.4 | 660.668653 | -0.950226329 | 0.389179816 | -2.441612566 | 0.014621829 | 0.137627695 |
| AL390879.1 | 119.6534485 | 0.600538009 | 0.546260752 | 1.099361442 | 0.271610442 | 0.583820065 |
| AL391987.2 | 14.92009936 | 0.632572709 | 0.843690078 | 0.749769051 | 0.453393812 | 0.733497409 |

| | | | | | | |
|---|---|---|---|---|---|---|
| AL441992.2 | 28.29219211 | -0.02397112 | 0.613703654 | -0.039059764 | 0.96884274 | 0.989496332 |
| AL445238.1 | 42.61510401 | -1.960927573 | 0.788977713 | -2.485402997 | 0.012940485 | 0.128476822 |
| AL445423.2 | 34.60525441 | -1.046563449 | 0.744070651 | -1.406537736 | 0.159564507 | 0.459433126 |
| AL445685.3 | 1054.77262 | 1.051297889 | 0.495850383 | 2.120191746 | 0.033989878 | 0.210275984 |
| AL445989.1 | 25.49585753 | 0.149638942 | 1.467836269 | 0.101945254 | 0.918800129 | 0.969677217 |
| AL450124.1 | 27.42301576 | -1.517892902 | 0.649025736 | -2.338725289 | 0.019349655 | 0.157622736 |
| AL512590.2 | 51.94059272 | -0.619728824 | 0.734329507 | -0.843938339 | 0.398703863 | 0.697260669 |
| AL513477.1 | 52.72216156 | -0.824512938 | 0.60936805 | -1.353062304 | 0.176035727 | 0.480071547 |
| AL513523.2 | 86.4434846 | 2.661938107 | 1.897733881 | 1.40269304 | 0.160708394 | 0.461238562 |
| AL589666.1 | 24.34459608 | 0.396956683 | 0.700103021 | 0.56699753 | 0.57071586 | 0.80670179 |
| AL589743.1 | 16.30852156 | 0.605687561 | 0.763452475 | 0.793353327 | 0.427571995 | 0.714914829 |
| AL589843.2 | 24.42065937 | -0.562442655 | 0.679728398 | -0.827452048 | 0.40798089 | 0.702586667 |
| AL590065.1 | 12.4760389 | 0.392733603 | 0.991284761 | 0.396186462 | 0.691967478 | 0.86812349 |
| AL590327.1 | 91.00539268 | -1.145176926 | 0.446408514 | -2.56531157 | 0.010308319 | 0.115220384 |
| AL591806.3 | 517.9665578 | -2.377150348 | 1.633662634 | -1.455104804 | 0.145640249 | 0.437613037 |
| AL591846.1 | 19.58166879 | 2.489840042 | 1.014003316 | 2.455455522 | 0.014070616 | 0.134821147 |
| AL592293.1 | 15.88704918 | 1.703976306 | 0.741710097 | 2.297361616 | 0.02159815 | 0.166611729 |
| AL603832.3 | 24.86286784 | -1.42824965 | 1.089368277 | -1.311080633 | 0.189830523 | 0.495280824 |
| AL627309.6 | 24.88323901 | 1.893968225 | 1.461614668 | 1.295805431 | 0.195042527 | 0.500590454 |
| AL627309.7 | 13.01453728 | -0.710744682 | 0.897170774 | -0.792206682 | 0.428240173 | 0.715316008 |
| AL669831.1 | 1028.508036 | -1.13382835 | 0.443608786 | -2.555919506 | 0.010590761 | 0.116977919 |
| AL669831.3 | 74.82247911 | -1.469395078 | 0.532860192 | -2.757562114 | 0.005823415 | 0.082336465 |
| AL713999.1 | 13.52793415 | 1.983014626 | 1.037627758 | 1.911104064 | 0.055991209 | 0.274013752 |
| AL732372.2 | 638.3673104 | -1.407860536 | 0.77736548 | -1.811066444 | 0.070130567 | 0.304159758 |
| AL732372.3 | 40.20992699 | -0.214617259 | 0.910323592 | -0.235759307 | 0.813619447 | 0.921959809 |
| ALAD | 1292.008643 | -1.128103745 | 0.349755139 | -3.225410066 | 0.001257924 | 0.031407111 |
| ALAS1 | 842.3356362 | 0.08521852 | 0.333473604 | 0.255548024 | 0.798299857 | 0.916056735 |
| ALCAM | 2990.346139 | -0.351518892 | 0.48336555 | -0.727231992 | 0.467083851 | 0.743482227 |
| ALDH16A1 | 697.669654 | 0.408805195 | 0.497781155 | 0.821254865 | 0.411501111 | 0.704686727 |
| ALDH18A1 | 2688.883557 | -0.008976077 | 0.237060719 | -0.037864043 | 0.969796082 | 0.989496332 |
| ALDH1A2 | 30.8579339 | -0.600936492 | 0.71988887 | -0.834762859 | 0.403851242 | 0.700367583 |
| ALDH1A3 | 72.88924176 | 0.580940539 | 0.653563438 | 0.888881637 | 0.374066695 | 0.678520659 |
| ALDH1B1 | 609.6478288 | 1.022624883 | 0.427080959 | 2.394452065 | 0.016645219 | |

0.146056278
ALDH1L1 51.74715993 0.060542559 0.447206635 0.135379384 0.892311923 0.957550733
ALDH1L2 55.5837937 -0.521287739 0.597921647 -0.87183286 0.383299566 0.684494911
ALDH2 1797.258202 0.658171607 0.294479832 2.235031177 0.025415302 0.18194777
ALDH3A2 2571.0469 -0.28716324 0.3081895 -0.931774898 0.35145287 0.658827565
ALDH3B1 19.98046246 -0.110976371 0.792057937 -0.14011143 0.888571949 0.955997805
ALDH3B2 116.3006274 -1.809257114 0.725960544 -2.492225132 0.012694554 0.127363842
ALDH4A1 2315.574399 -1.002522727 0.396474503 -2.528593184 0.011452068 0.121279761
ALDH5A1 216.0178784 0.00607959 0.350243017 0.017358205 0.986150852 0.994937085
ALDH6A1 1379.109948 -0.245802366 0.280042945 -0.877730971 0.380089733 0.68184242
ALDH7A1 6066.252239 -0.162132449 0.249638913 -0.649467854 0.516036018 0.773896629
ALDH9A1 2482.597265 0.168283454 0.264171776 0.637022839 0.524109967 0.777662198
ALDOA 13962.43927 0.233395084 0.248105434 0.940709279 0.346853863 0.65343598
ALDOC 400.691606 0.264128314 0.315608817 0.836885091 0.402657162 0.699577622
ALG1 721.918649 0.201065274 0.309916328 0.648772768 0.516485262 0.773950426
ALG10 178.5576463 -0.14226416 0.494100941 -0.287925296 0.773403923 0.906158747
ALG10B 660.1431108 0.172706014 0.369236746 0.467737885 0.639972045 0.843538624
ALG11 1321.974857 -0.403483128 0.485442216 -0.83116613 0.405879786 0.701371854
ALG12 487.4355772 -0.251246666 0.404279514 -0.621467716 0.53429193 0.783689739
ALG13 1030.4882 -0.05043422 0.296459332 -0.170121884 0.864914283 0.945248223
ALG14 212.9173129 0.554363845 0.34226625 1.619685972 0.105299751 0.372117988
ALG1L 72.33024979 1.006668531 0.554752395 1.814626742 0.06958129 0.303425952
ALG1L2 18.36555968 -0.938127659 0.814647367 -1.151575144 0.2494957 0.562413758
ALG2 1050.042274 0.059877524 0.464882332 0.128801462 0.897514748 0.959562808
ALG3 647.9589165 -0.09374974 0.326961792 -0.286729956 0.774319099 0.906279513
ALG5 504.1630118 0.733623691 0.480943784 1.525383454 0.127163483 0.410771662
ALG6 685.8657652 0.545191406 0.254771259 2.139925075 0.032360822 0.205073532
ALG8 1769.980321 0.247566843 0.240710017 1.028485837 0.30372135 0.614002998
ALG9 800.7320117 -0.378182883 0.291349443 -1.298038806 0.194274 0.500187881
ALKBH1 307.125968 0.003684034 0.285850581 0.012887972 0.989717171 0.996285567
ALKBH2 385.552086 0.245542953 0.343889495 0.714017021 0.475216655 0.748884582
ALKBH3 286.1072088 0.207593086 0.381481478 0.544176056 0.586320327 0.815986211
ALKBH4 432.1589946 0.43167263 0.359660828 1.20022142 0.230053358 0.53893445
ALKBH5 1204.432299 0.378804596 0.292968344 1.292988148 0.196015158 0.501797297
ALKBH6 426.4975901 -0.363400205 0.282506205 -1.286344155 0.198322979 0.50414535
ALKBH7 622.256763 1.046243404 0.60898871 1.718001315 0.085796375 0.335901113
ALKBH8 348.9095706 0.08685766 0.305400302 0.284405939 0.776099308 0.907383061
ALMS1 1234.669785 -0.302552727 0.526814437 -0.574306065 0.565760706

| | | | | | | |
|---|---|---|---|---|---|---|
| | | | | | | 0.804118013 |
| ALMS1P1 | 18.51048303 | -0.253141758 | 0.769588767 | -0.328931201 | 0.74220769 | 0.892150449 |
| ALOX12B | 13.30023679 | 0.518514098 | 0.809502653 | 0.640534155 | 0.521825391 | 0.776900197 |
| ALOX12P2 | 74.93199932 | -1.233399613 | 0.457555165 | -2.6956304 | 0.007025557 | 0.0921438 |
| ALOX15P1 | 13.42095516 | -0.866965806 | 0.841403789 | -1.030380202 | 0.302831563 | 0.613538084 |
| ALOX5 | 16.61772817 | -4.022340035 | 1.465479871 | -2.744725543 | 0.006056152 | 0.084137259 |
| ALPG | 47.36519185 | -1.809080014 | 1.007287337 | -1.795992015 | 0.072495787 | 0.309899573 |
| ALPK1 | 16.38342079 | -0.627938634 | 0.797695645 | -0.787190751 | 0.43117021 | 0.717875687 |
| ALPK2 | 1556.122034 | -4.459799302 | 0.615567235 | -7.245023853 | 4.32E-13 | 1.64E-09 |
| ALPK3 | 461.7992311 | -0.627319412 | 0.458601362 | -1.367896966 | 0.171344329 | 0.474701392 |
| ALPL | 5383.846059 | 0.168248023 | 0.363542324 | 0.462801748 | 0.643506478 | 0.845466381 |
| ALPP | 13.36260362 | -1.43537985 | 0.932002513 | -1.540102982 | 0.123535253 | 0.40487218 |
| ALS2 | 2716.518358 | -0.76363586 | 0.330452034 | -2.310882613 | 0.02083934 | 0.163640591 |
| ALS2CL | 131.5838272 | -0.660790304 | 0.419845601 | -1.573888834 | 0.115513152 | 0.391434001 |
| ALX1 | 63.07546778 | -0.823562648 | 0.779426913 | -1.05662588 | 0.290682364 | 0.60368818 |
| ALYREF | 2446.298969 | 0.69828944 | 0.376526537 | 1.85455571 | 0.063659698 | 0.290852073 |
| AMACR | 344.4455652 | 0.436964724 | 0.297069155 | 1.470919204 | 0.141312969 | 0.431039476 |
| AMBN | 21.74132092 | -4.438961215 | 1.439334865 | -3.084036469 | 0.002042126 | 0.042686766 |
| AMBP | 23.93753722 | 1.65222213 | 1.039658963 | 1.589196255 | 0.112016092 | 0.384920166 |
| AMBRA1 | 909.2518191 | -0.545176245 | 0.382496999 | -1.425308556 | 0.154068028 | 0.451087275 |
| AMD1 | 3030.054856 | 0.522785996 | 0.27454143 | 1.904215317 | 0.056882148 | 0.2761211 |
| AMDHD1 | 36.63561706 | -2.747270875 | 0.762674761 | -3.602152603 | 0.000315593 | 0.013085543 |
| AMDHD2 | 624.8739922 | 0.105470254 | 0.477148622 | 0.221042773 | 0.825059129 | 0.926977119 |
| AMER1 | 488.7492341 | -0.345469165 | 0.534525772 | -0.646309651 | 0.518078839 | 0.773958483 |
| AMER2 | 376.1812836 | -2.417022882 | 0.636226081 | -3.799000002 | 0.000145281 | 0.007733542 |
| AMER3 | 56.43396483 | -1.941476263 | 0.844701622 | -2.298416638 | 0.021538087 | 0.166469811 |
| AMFR | 930.1413888 | -0.11636613 | 0.29354225 | -0.396420379 | 0.691794935 | 0.86812349 |
| AMIGO1 | 139.112069 | 0.44456636 | 0.524974042 | 0.846834938 | 0.397087131 | 0.695636128 |
| AMIGO2 | 38.91329752 | -0.204646734 | 0.655151253 | -0.31236563 | 0.754762668 | 0.897503925 |
| AMIGO3 | 25.76170434 | -0.821666042 | 1.483691202 | -0.553798554 | 0.579716718 | 0.811340629 |
| AMMECR1 | 593.6104572 | 0.225816361 | 0.271703078 | 0.831114475 | 0.405908963 | 0.701371854 |
| AMMECR1L | 1224.646527 | -0.265495861 | 0.326080058 | -0.814204531 | 0.41552779 | 0.707542731 |
| AMN1 | 305.4305338 | 1.106667724 | 0.333165913 | 3.321671524 | 0.0008948 | 0.025375039 |
| AMOT | 1475.966206 | -1.627683999 | 0.43743291 | -3.720991175 | 0.000198442 | 0.009408029 |
| AMOTL1 | 2076.609634 | -0.303296019 | 0.34512771 | -0.87879359 | 0.379513202 | 0.681253679 |

| | | | | | | |
|---|---|---|---|---|---|---|
| AMOTL2 | 2060.410073 | -0.871704874 | 0.464128778 | -1.878153038 | 0.060360232 | 0.283945763 |
| AMPD2 | 712.44094 | 0.581037571 | 0.416705308 | 1.394360859 | 0.163208661 | 0.464573649 |
| AMPD3 | 39.65488509 | -0.80299058 | 0.537105793 | -1.495032432 | 0.134905978 | 0.421556868 |
| AMPH | 306.962392 | 1.373078117 | 0.703155578 | 1.952737287 | 0.050850733 | 0.259366654 |
| AMT | 1090.580641 | 0.424280725 | 0.284094921 | 1.493447059 | 0.135320203 | 0.422590118 |
| AMY2B | 123.2522354 | -0.00818462 | 0.561304181 | -0.014581434 | 0.988366111 | 0.995803391 |
| AMZ2 | 1877.861897 | 0.44504773 | 0.276268423 | 1.610925077 | 0.107196053 | 0.375111686 |
| AMZ2P1 | 326.2274163 | 0.983650002 | 0.364522697 | 2.698460235 | 0.006966106 | 0.091738536 |
| ANAPC1 | 2246.02236 | -0.245158465 | 0.360301197 | -0.680426452 | 0.496234476 | 0.761056737 |
| ANAPC10 | 409.4251126 | 0.772692677 | 0.293765701 | 2.630302561 | 0.008530891 | 0.10430915 |
| ANAPC11 | 1665.786409 | 1.237352808 | 0.47013828 | 2.631891216 | 0.008491105 | 0.104053754 |
| ANAPC13 | 1306.487491 | 0.856151961 | 0.312469011 | 2.739957986 | 0.006144704 | 0.084669666 |
| ANAPC15 | 903.4429187 | 0.555705551 | 0.378603788 | 1.467775994 | 0.14216508 | 0.432308365 |
| ANAPC16 | 2240.442045 | 0.760480897 | 0.37901648 | 2.006458653 | 0.044807336 | 0.243645911 |
| ANAPC2 | 683.0568582 | -0.410073506 | 0.448518619 | -0.914284243 | 0.360567508 | 0.667663818 |
| ANAPC4 | 1089.738411 | -0.226745904 | 0.238301628 | -0.951507992 | 0.341346564 | 0.648152255 |
| ANAPC5 | 2520.588896 | 0.037668917 | 0.238623633 | 0.157859122 | 0.87456781 | 0.949740014 |
| ANAPC7 | 1770.195212 | 0.044722911 | 0.22373689 | 0.199890642 | 0.841566109 | 0.935153081 |
| ANG | 35.82966247 | -0.902955213 | 0.721720658 | -1.251114545 | 0.21089269 | 0.518294063 |
| ANGEL1 | 790.0926618 | -0.197617675 | 0.400952247 | -0.492870851 | 0.622103845 | 0.833640266 |
| ANGEL2 | 1057.116089 | -0.177658115 | 0.299589576 | -0.593004993 | 0.553177811 | 0.79574357 |
| ANGPT1 | 73.73096202 | 2.484076929 | 1.057494648 | 2.349020805 | 0.018822855 | 0.155449937 |
| ANGPT2 | 18.03951408 | -0.777695072 | 0.934121243 | -0.832541898 | 0.405103139 | 0.700857535 |
| ANGPTL2 | 375.7822035 | -0.040583962 | 0.552523503 | -0.073452009 | 0.941446432 | 0.97846707 |
| ANGPTL4 | 69.01653784 | 2.171613908 | 0.597162052 | 3.636557111 | 0.000276306 | 0.011709082 |
| ANGPTL7 | 21.71762336 | 0.241341611 | 1.126010406 | 0.214333375 | 0.830287093 | 0.930253375 |
| ANK1 | 142.8157555 | -0.329153123 | 0.502176081 | -0.655453606 | 0.512175747 | 0.771687791 |
| ANK2 | 622.6048135 | -0.324481338 | 0.492508393 | -0.658834129 | 0.510002289 | 0.769949736 |
| ANK3 | 3001.547717 | -1.133311001 | 0.262435876 | -4.318430152 | 1.57E-05 | 0.001528215 |
| ANKAR | 44.85953237 | 0.464573846 | 0.477532499 | 0.972863307 | 0.330621244 | 0.638962406 |
| ANKDD1A | 116.1932438 | 1.003201136 | 0.409538276 | 2.449590661 | 0.01430187 | 0.135778267 |
| ANKDD1B | 21.8531831 | -7.06780856 | 1.624432095 | -4.35094122 | 1.36E-05 | 0.001389524 |
| ANKEF1 | 326.8042336 | -0.683048871 | 0.312667519 | -2.184585315 | 0.028919266 | 0.19447437 |
| ANKFY1 | 2116.127169 | -0.533405729 | 0.392157264 | -1.360183216 | 0.173771952 | 0.477935875 |
| ANKH | 57.90004495 | -0.758848132 | 0.653649022 | -1.16094128 | 0.24566578 | |

0.557432777
ANKHD1 2239.925319 0.512605916 0.23972032 2.138349876 0.032488357
0.205554752
ANKHD1-EIF4EBP3 1039.440881 -0.734856134 0.623118671 -1.179319714
0.238270891 0.549112515
ANKIB1 1724.045825 -0.405595751 0.275749051 -1.470887204
0.141321624 0.431039476
ANKLE1 132.6862351 -0.179117884 0.34475102 -0.519557227
0.603372217 0.825211561
ANKLE2 1864.348021 -0.285450487 0.320955057 -0.889378373
0.373799764 0.678520659
ANKMY1 157.0396159 -0.333205381 0.372446473 -0.894639647
0.370979751 0.676265004
ANKMY2 914.321125 -0.458922505 0.363237973 -1.26342106 0.206437899
0.512832712
ANKRA2 520.2252589 0.564524558 0.292854593 1.927661612 0.05389723
0.268354078
ANKRD1 118.4629662 -1.39556597 0.584900456 -2.385988856 0.017033267
0.14773715
ANKRD10 2766.431244 0.530722524 0.4018641 1.320651745 0.186617511
0.49118221
ANKRD11 3997.730673 -0.119111491 0.301426988 -0.395158681
0.692725784 0.86854217
ANKRD12 2011.906056 0.684387622 0.453307501 1.50976461 0.131103498
0.416813082
ANKRD13A 752.2153615 -0.735622746 0.359085032 -2.0486032 0.040500932
0.231601821
ANKRD13B 255.9277454 0.803231878 0.523946237 1.533042554 0.12526537
0.407112453
ANKRD13C 874.8838225 0.141495006 0.259769611 0.544694223 0.585963839
0.815639728
ANKRD13D 538.4245124 0.028150606 0.332774898 0.084593539 0.932584536
0.97453841
ANKRD16 153.6134174 0.154935016 0.346293025 0.447410154 0.654578945
0.849149183
ANKRD17 3361.64888 -0.192239697 0.269397908 -0.713590164
0.475480642 0.74892861
ANKRD18A 92.93716274 0.417968388 0.421394822 0.991868828 0.321261517
0.63106605
ANKRD18B 183.5329662 1.277725568 0.473792957 2.696801526 0.007000898
0.091957254
ANKRD18CP 40.14681034 -0.692875921 0.685263069 -1.011109387
0.311964078 0.622030306
ANKRD20A19P 15.26292309 0.268393629 0.858052358 0.312794 0.754437174
0.897438517
ANKRD20A5P 22.96921083 -1.210745705 0.700502963 -1.728394838
0.08391746 0.331434265
ANKRD22 11.80399525 0.724518125 1.324506004 0.547010072 0.584371812
0.814844188
ANKRD23 91.42610216 0.458410583 0.500247213 0.916368091 0.359473866
0.66703422
ANKRD24 72.68872413 0.547652636 0.452262143 1.210918588 0.225926608
0.534727976
ANKRD26 1052.176093 0.102809366 0.259094104 0.396803185 0.691512602
0.86812349
ANKRD27 1286.99491 -0.405396212 0.286793891 -1.413545492
0.157495395 0.456731315
ANKRD28 1000.071289 -0.372612453 0.259311079 -1.436932251
0.150737245 0.445603028
ANKRD33 12.09311553 0.205223428 1.007776037 0.203639916 0.838634898
0.933879959
ANKRD33B 661.8867193 -1.999598402 0.667561405 -2.995377482
0.002741054 0.051338925
ANKRD34B 59.60184604 -0.149155345 0.705890338 -0.211301015

| | | | | | | |
|---|---|---|---|---|---|---|
| | | | | | 0.832652391 | 0.931432899 |
| ANKRD35 | 49.25865416 | 0.69097801 | 0.648044455 | 1.066250941 | 0.28631022 | 0.60033547 |
| ANKRD36 | 478.1061093 | -0.151848872 | 0.250441332 | -0.606325126 | 0.544298877 | 0.790726637 |
| ANKRD36B | 250.6439745 | 0.102399249 | 0.288453995 | 0.354993345 | 0.722594577 | 0.882935039 |
| ANKRD36BP2 | 11.62355334 | -0.917289107 | 0.927936763 | -0.98852545 | 0.322895377 | 0.632818209 |
| ANKRD36C | 74.03204663 | 0.312013515 | 0.401753048 | 0.776630112 | 0.437377031 | 0.721468765 |
| ANKRD37 | 26.93630932 | 0.335545603 | 0.8143109 | 0.412060803 | 0.680294858 | 0.862002279 |
| ANKRD39 | 204.5813986 | 0.409389115 | 0.318650775 | 1.284757944 | 0.198876883 | 0.504782074 |
| ANKRD40 | 1490.677467 | 0.009528354 | 0.303157346 | 0.031430392 | 0.974926304 | 0.991261106 |
| ANKRD42 | 150.7399969 | -0.577940595 | 0.327834539 | -1.762903313 | 0.077916798 | 0.320085498 |
| ANKRD44 | 18.36680758 | -1.314585196 | 0.78172736 | -1.681641533 | 0.092638372 | 0.3491405 |
| ANKRD45 | 32.82446371 | 0.787829283 | 0.708149043 | 1.112519025 | 0.265915055 | 0.579614326 |
| ANKRD46 | 788.471915 | 0.601603445 | 0.327005768 | 1.839733433 | 0.065807383 | 0.29619856 |
| ANKRD49 | 461.8160586 | -0.30526746 | 0.278017723 | -1.098014389 | 0.272198206 | 0.584421028 |
| ANKRD50 | 958.149575 | -0.425388042 | 0.320823366 | -1.325925996 | 0.184864211 | 0.489406194 |
| ANKRD52 | 1354.931454 | -0.688022282 | 0.421028699 | -1.634145806 | 0.102228239 | 0.366730813 |
| ANKRD53 | 21.75130805 | -0.143490554 | 0.7255733 | -0.197761624 | 0.843231573 | 0.936162912 |
| ANKRD54 | 392.1467236 | 0.378120912 | 0.298494538 | 1.266759901 | 0.205241154 | 0.511787237 |
| ANKRD6 | 671.2247115 | -0.78351457 | 0.341737878 | -2.292735515 | 0.021863237 | 0.167857876 |
| ANKRD65 | 149.4810502 | -0.013447721 | 0.524691543 | -0.025629766 | 0.979552644 | 0.992903933 |
| ANKRD9 | 436.0832325 | 0.738449617 | 0.3741323 | 1.973766012 | 0.048408355 | 0.253679847 |
| ANKS1A | 1859.201096 | -0.880409612 | 0.428333907 | -2.055428247 | 0.039837659 | 0.229888601 |
| ANKS1B | 29.26694498 | -0.631792804 | 0.639558749 | -0.98785734 | 0.323222521 | 0.63322621 |
| ANKS3 | 391.3795168 | -0.28169717 | 0.33440135 | -0.842392442 | 0.399568322 | 0.697589493 |
| ANKS6 | 1245.790156 | 0.147980772 | 0.297663344 | 0.497141403 | 0.61908934 | 0.832495306 |
| ANKZF1 | 1127.891143 | -0.659548252 | 0.327609097 | -2.013217147 | 0.04409179 | 0.241443505 |
| ANLN | 2702.080153 | 0.386795563 | 0.405207063 | 0.954562736 | 0.339798869 | 0.646629378 |
| ANO1 | 16.9805762 | -1.54966895 | 1.174190276 | -1.319776685 | 0.186909588 | 0.491529111 |
| ANO10 | 1312.716362 | -0.710043362 | 0.357386968 | -1.986763438 | 0.046948618 | 0.249564641 |
| ANO2 | 132.6004418 | -2.071456585 | 0.453720726 | -4.565488124 | 4.98E-06 | 0.00065741 |
| ANO3 | 20.63343462 | 2.76576508 | 1.319150618 | 2.096625694 | 0.036026723 | 0.217061721 |
| ANO4 | 17.12747555 | -1.345849911 | 0.885919685 | -1.519155668 | 0.128723317 | 0.413329156 |
| ANO5 | 171.6588716 | 0.206476512 | 0.365984455 | 0.564167438 | 0.572640186 | 0.807670159 |
| ANO6 | 2291.127944 | -1.060428092 | 0.391805763 | -2.706514786 | 0.006799355 | 0.090392202 |
| ANO7 | 54.63016593 | -0.217853119 | 0.437210227 | -0.498280016 | 0.618286691 | |

| | | | | | | |
|---|---|---|---|---|---|---|
| | | | | | | 0.832248827 |
| ANO8 | 453.9124245 | -0.158566819 | 0.571382804 | -0.277514161 | 0.781385334 | 0.909755868 |
| ANO9 | 711.8884709 | -1.44157686 | 0.526720686 | -2.736890536 | 0.006202293 | 0.085385647 |
| ANOS1 | 361.0955173 | 0.12106172 | 0.325658139 | 0.371744802 | 0.710082865 | 0.876599663 |
| ANOS2P | 135.6488039 | 0.129794184 | 0.519122132 | 0.250026297 | 0.802567012 | 0.917815781 |
| ANP32A | 8915.628049 | 0.42874368 | 0.330528282 | 1.297146731 | 0.194580705 | 0.500370345 |
| ANP32B | 7473.650426 | 0.574425963 | 0.445036198 | 1.290739865 | 0.196793896 | 0.50258876 |
| ANP32E | 5635.555785 | 0.448740205 | 0.228766815 | 1.96156162 | 0.049813544 | 0.257223037 |
| ANPEP | 41.34166783 | -0.738726328 | 0.736874368 | -1.002513263 | 0.316095764 | 0.625961211 |
| ANTXR1 | 943.034564 | 0.169298893 | 0.341710029 | 0.495446075 | 0.620285278 | 0.833216571 |
| ANTXR2 | 367.2699193 | 0.23753793 | 0.430948081 | 0.551198487 | 0.581497617 | 0.812928525 |
| ANXA1 | 596.111893 | -1.497201951 | 0.773609751 | -1.935345241 | 0.052947944 | 0.266072627 |
| ANXA11 | 1513.587944 | -1.273476167 | 0.497320484 | -2.560675075 | 0.010446901 | 0.116176818 |
| ANXA2 | 18097.31845 | -0.524138326 | 0.380888896 | -1.376092431 | 0.168793019 | 0.471334233 |
| ANXA3 | 2358.917474 | -1.740314997 | 0.686374638 | -2.535517634 | 0.011228128 | 0.120297968 |
| ANXA4 | 669.0487803 | -0.536454731 | 0.409023171 | -1.311550954 | 0.189671691 | 0.495097939 |
| ANXA5 | 6586.358875 | 0.29915975 | 0.254004057 | 1.177775477 | 0.238886128 | 0.550029055 |
| ANXA6 | 3713.979317 | 0.117047044 | 0.399339263 | 0.293101768 | 0.769444366 | 0.904597192 |
| ANXA7 | 2265.573518 | -0.188688022 | 0.238368144 | -0.791582376 | 0.428604226 | 0.715644829 |
| ANXA8 | 13.85094385 | -0.651955166 | 1.220222115 | -0.534292206 | 0.593139383 | 0.819859199 |
| ANXA9 | 510.3859725 | -1.830672945 | 0.697734128 | -2.623740005 | 0.008697015 | 0.105298369 |
| AOC2 | 92.64838236 | -0.01539809 | 0.456585568 | -0.033724433 | 0.973096895 | 0.990423175 |
| AOC3 | 83.43017047 | -0.283064003 | 0.540742342 | -0.523472977 | 0.600645152 | 0.824475293 |
| AOX1 | 58.50839302 | 0.686680065 | 0.761106539 | 0.902212804 | 0.366943834 | 0.673470229 |
| AP000295.1 | 76.91301421 | -0.084966461 | 1.041280758 | -0.081598033 | 0.934966366 | 0.975224148 |
| AP000311.1 | 25.10731885 | 0.681074439 | 0.673611584 | 1.01107887 | 0.311978683 | 0.622030306 |
| AP000346.2 | 70.11914721 | -0.407279704 | 0.505387466 | -0.805876146 | 0.420314267 | 0.710879347 |
| AP000347.1 | 539.2497479 | -0.473642998 | 0.432789487 | -1.094395804 | 0.273781423 | 0.586847211 |
| AP000553.5 | 35.40717527 | -0.567166043 | 0.702271708 | -0.80761625 | 0.419311529 | 0.709924862 |
| AP000580.1 | 15.85667334 | 0.847897112 | 0.906596004 | 0.935253529 | 0.349657658 | 0.656273207 |
| AP000769.1 | 29.74950258 | 1.113104115 | 0.61872021 | 1.799042762 | 0.072011917 | 0.308875541 |
| AP000812.5 | 19.47885752 | 0.483422001 | 0.664322453 | 0.727691799 | 0.466802272 | 0.743189974 |
| AP000944.2 | 145.2467896 | 0.213971187 | 1.068473549 | 0.200258759 | 0.841278214 | 0.935153081 |
| AP001267.5 | 104.5757342 | 0.6231101 | 0.446900863 | 1.394291556 | 0.16322958 | 0.464573649 |
| AP001273.2 | 63.83043505 | -0.826852075 | 3.647167747 | -0.226710734 | | |

0.82064868 0.925132406
AP001324.1 22.26166605 1.147410923 0.734167388 1.562873728 0.118082298
0.395958766
AP001453.5 26.78809908 0.675957952 0.787661174 0.858183663 0.390791056
0.690469214
AP001781.2 11.11787984 -1.116509545 1.750849108 -0.637696041
0.523671563 0.777662198
AP001931.1 10.7696413 -2.062156222 1.404776357 -1.467960514
0.142114949 0.432308365
AP002495.2 44.60033331 -0.461636844 0.581917241 -0.793303261
0.427601158 0.714914829
AP002748.4 72.77348314 1.168321977 2.256225766 0.517821396 0.604582888
0.825894731
AP002956.1 231.0987876 0.254484099 0.382936802 0.664559002 0.506332599
0.767347299
AP002990.1 137.6763209 -1.380686203 0.638712151 -2.161672046
0.030643464 0.199470916
AP003108.2 291.9335716 -0.522839371 0.29005066 -1.80257949 0.071454281
0.307527061
AP003170.1 10.69109296 -3.67281579 1.521448501 -2.414025705 0.015777349
0.142221132
AP003419.1 207.2015866 0.263349244 0.720485882 0.365516176 0.714726134
0.87855355
AP005061.1 24.06055724 0.508299502 0.607977491 0.83604987 0.403126848
0.699766232
AP006222.1 179.3463419 -0.798160788 0.512488999 -1.557420333
0.119370722 0.398357612
AP1AR 693.8439421 0.524268073 0.303557366 1.727080717 0.084153168 0.332133444
AP1B1 2118.708899 -0.087891885 0.384429292 -0.228629522 0.819156877
0.924316427
AP1G1 2257.021414 -0.43739382 0.327153109 -1.336969777 0.181232502
0.485749663
AP1G2 1791.868267 -1.123684732 0.289750447 -3.878112161 0.00010527
0.006358893
AP1M1 826.7914326 0.045743025 0.367128999 0.124596601 0.90084292 0.961704658
AP1M2 2083.080621 -1.094158469 0.821210604 -1.332372553 0.182737797
0.486683209
AP1S1 2501.782761 0.015821925 0.413217559 0.038289575 0.969456802 0.989496332
AP1S2 1690.222391 0.287598906 0.297277675 0.967441991 0.333323127 0.641324687
AP1S3 138.0154837 0.277907769 0.343393204 0.809298979 0.418343193 0.709602481
AP2A1 3421.924631 -0.342936768 0.359765915 -0.953221954 0.340477625
0.647332681
AP2A2 131.7551861 -0.949839199 0.644231225 -1.47437622 0.140380324
0.430070656
AP2B1 8995.802054 -0.406692969 0.252149835 -1.612901987 0.106765803
0.374599763
AP2M1 10002.92182 -0.066790587 0.216330095 -0.308743853 0.757516383
0.898914948
AP2S1 2076.032218 0.791732243 0.452735998 1.748772456 0.08033036 0.324292679
AP3B1 2998.052839 -0.569443967 0.272294563 -2.091279239 0.03650304
0.218801903
AP3B2 123.8635386 -0.726226093 0.367415407 -1.976580405 0.048089079
0.252967897
AP3D1 4326.804914 -0.24039466 0.325757148 -0.737956669 0.460540781
0.738395128
AP3M1 2080.492579 -0.070772886 0.304201206 -0.232651562 0.816031985
0.923222505
AP3M2 509.2956855 0.608112981 0.442266425 1.374992418 0.169133795 0.471613806
AP3S1 1983.147865 0.70451708 0.386541086 1.82261888 0.068361134 0.301221831
AP3S2 796.0737157 -0.0826317 0.29716703 -0.27806483 0.780962593 0.909494396
AP4B1 451.9160557 -0.077969672 0.253823755 -0.307180359 0.758706095
0.899440455
AP4E1 485.0379685 -0.372309627 0.345136188 -1.078732511 0.280706988
0.594538239

| | | | | | | |
|---|---|---|---|---|---|---|
| AP4M1 | 530.3541977 | 0.076928229 | 0.321999576 | 0.238907857 | 0.811177033 | 0.920897407 |
| AP4S1 | 177.5720384 | -0.211622888 | 0.376628614 | -0.56188744 | 0.574192712 | 0.808359708 |
| AP5B1 | 295.5831409 | -0.344608268 | 0.443085298 | -0.777747016 | 0.436718169 | 0.720784523 |
| AP5M1 | 1712.233668 | -0.51377206 | 0.26161211 | -1.963869564 | 0.04954522 | 0.25654413 |
| AP5S1 | 147.8664988 | -0.429869575 | 0.364724524 | -1.178614399 | 0.238551755 | 0.549478686 |
| AP5Z1 | 465.6677516 | -0.445509077 | 0.505681679 | -0.881006957 | 0.378314055 | 0.681253679 |
| APAF1 | 465.1665401 | -0.552011584 | 0.372159897 | -1.483264554 | 0.138004136 | 0.426233866 |
| APBA2 | 345.5013552 | 0.728144499 | 0.451318456 | 1.613371864 | 0.106663742 | 0.374599763 |
| APBA3 | 256.3456632 | -0.193371337 | 0.457901959 | -0.422298558 | 0.67280711 | 0.859151631 |
| APBB1 | 1340.373365 | 0.043758617 | 0.362059973 | 0.120860134 | 0.903801819 | 0.963500625 |
| APBB1IP | 15.85927286 | -3.451440823 | 1.188624644 | -2.903726455 | 0.003687502 | 0.062506247 |
| APBB2 | 1536.573985 | -0.350481032 | 0.231686405 | -1.512738874 | 0.130346001 | 0.415496314 |
| APBB3 | 519.0991544 | -0.557690671 | 0.366063861 | -1.523479182 | 0.127638865 | 0.411564128 |
| APC | 732.7289611 | -0.404889792 | 0.371444043 | -1.090042497 | 0.275694425 | 0.588927079 |
| APC2 | 30.01466258 | 1.463053087 | 1.040913032 | 1.405547862 | 0.159858427 | 0.460017489 |
| APCDD1 | 151.0928205 | 0.224755223 | 0.434671476 | 0.517069179 | 0.605107868 | 0.8260678 |
| APCDD1L | 24.78960425 | -0.277956101 | 0.679328928 | -0.409162763 | 0.682420219 | 0.862821664 |
| APEH | 3779.527431 | -0.179375558 | 0.259991398 | -0.68992882 | 0.490238951 | 0.758593654 |
| APELA | 443.0722817 | -1.236042879 | 0.311893612 | -3.963027241 | 7.40E-05 | 0.004945969 |
| APEX1 | 10662.58988 | 0.197851761 | 0.305572637 | 0.647478658 | 0.517322203 | 0.773950969 |
| APEX2 | 798.5741414 | -0.161480168 | 0.258543391 | -0.624576661 | 0.532248941 | 0.782369637 |
| APH1A | 3440.556512 | -0.154925161 | 0.285393042 | -0.542848418 | 0.587234174 | 0.816733558 |
| APH1B | 377.257041 | -0.850646846 | 0.291765596 | -2.915514573 | 0.003551025 | 0.061218868 |
| API5 | 5255.706962 | -0.055495222 | 0.248344738 | -0.223460432 | 0.823177175 | 0.926014081 |
| APIP | 501.5280579 | 1.271068095 | 0.368565462 | 3.448690199 | 0.000563313 | 0.018823822 |
| APLF | 171.0880938 | 0.230757346 | 0.363856399 | 0.634198949 | 0.525950993 | 0.778458782 |
| APLNR | 153.7296258 | 2.634870586 | 0.822388071 | 3.203926078 | 0.001355673 | 0.033172439 |
| APLP1 | 919.0863582 | 1.336733874 | 0.477325354 | 2.800466942 | 0.005102873 | 0.075897737 |
| APLP2 | 18386.98955 | -0.498189077 | 0.2312092 | -2.154711307 | 0.031184432 | 0.201233103 |
| APMAP | 3067.216742 | -0.256967855 | 0.312276469 | -0.822885746 | 0.410572969 | 0.704296983 |
| APOBEC2 | 14.61626901 | 1.014583877 | 0.810307741 | 1.252096982 | 0.210534527 | 0.518294063 |
| APOBEC3B | 119.5034176 | -0.653771156 | 0.570158139 | -1.146648817 | 0.251526792 | 0.564383659 |
| APOBEC3C | 1407.062599 | 0.469267452 | 0.527741494 | 0.889199461 | 0.373895893 | 0.678520659 |
| APOBEC3D | 43.69173222 | -1.913075535 | 0.615537256 | -3.107976839 | 0.001883728 | 0.040513318 |
| APOBEC3F | 93.45344291 | -0.741397115 | 0.472515946 | -1.569041469 | 0.116638277 | 0.393839149 |
| APOBEC3G | 90.10278171 | -1.715060395 | 0.627831716 | -2.731719903 | 0.006300468 | 0.086267508 |
| APOBEC3H | 47.37665647 | -0.109260934 | 0.482946895 | -0.226237989 | 0.821016326 | 0.925158936 |

| | | | | | | |
|---|---|---|---|---|---|---|
| APOC1 | 520.7455923 | 0.567460256 | 0.378975289 | 1.497354242 | 0.134301108 | 0.420619758 |
| APOE | 5904.304897 | -0.164907981 | 0.523070025 | -0.315269416 | 0.752557104 | 0.896586461 |
| APOL2 | 185.5600419 | 0.0893451 | 0.36293864 | 0.246171362 | 0.805549583 | 0.918930547 |
| APOL4 | 34.36292101 | 0.341829129 | 0.570863729 | 0.598792868 | 0.549311019 | 0.79348329 |
| APOL6 | 73.26484678 | -0.800608341 | 0.641380463 | -1.248258073 | 0.211936563 | 0.519175172 |
| APOLD1 | 67.4912624 | 1.4206639 | 0.455669188 | 3.117752828 | 0.001822356 | 0.03966565 |
| APOO | 579.0352686 | 1.257002315 | 0.398843293 | 3.151619536 | 0.001623677 | 0.03731384 |
| APOOL | 633.049555 | -0.212134154 | 0.269979944 | -0.785740417 | 0.432019579 | 0.71850132 |
| APOPT1 | 551.2699327 | 0.738902404 | 0.466587393 | 1.583631307 | 0.113277645 | 0.387319169 |
| APP | 21171.86421 | -0.706184286 | 0.249931894 | -2.825506883 | 0.004720587 | 0.072780519 |
| APPBP2 | 786.1801859 | -0.001394859 | 0.35276416 | -0.003954085 | 0.996845105 | 0.998556427 |
| APPL1 | 786.2284646 | -0.732563504 | 0.349325191 | -2.097081809 | 0.035986334 | 0.217048632 |
| APPL2 | 903.6427415 | -0.224979023 | 0.260279792 | -0.864373762 | 0.387382601 | 0.687545052 |
| APRT | 1063.197717 | 0.666470417 | 0.420290817 | 1.585736327 | 0.112799135 | 0.386379696 |
| APTX | 996.3005456 | 0.468850953 | 0.311388147 | 1.505680153 | 0.132149301 | 0.418458996 |
| AQP11 | 48.84353314 | -0.286852103 | 0.667221481 | -0.429920366 | 0.66725357 | 0.856421651 |
| AQP3 | 97.38497501 | -0.086750336 | 0.518497648 | -0.167310953 | 0.867125386 | 0.946278179 |
| AQR | 1891.995214 | -0.525454272 | 0.365031678 | -1.439475816 | 0.150015757 | 0.444740017 |
| AR | 212.3521418 | 0.581463275 | 0.374930961 | 1.550854251 | 0.120936616 | 0.401219509 |
| ARAF | 1168.535397 | -0.119350646 | 0.245810034 | -0.485540174 | 0.627293223 | 0.836041947 |
| ARAP1 | 811.9451625 | 0.268146384 | 0.375854065 | 0.713432178 | 0.475578368 | 0.74892861 |
| ARAP2 | 81.89473452 | -1.612954635 | 0.509785414 | -3.163987417 | 0.001556235 | 0.0365054 |
| ARAP3 | 413.2002051 | 0.287505894 | 0.454774895 | 0.632193855 | 0.527260212 | 0.778906646 |
| ARC | 31.77866357 | 0.46906696 | 0.718756877 | 0.652608656 | 0.514008604 | 0.772630754 |
| ARCN1 | 3885.127757 | -0.271932319 | 0.288124512 | -0.943801404 | 0.345271143 | 0.652154944 |
| AREL1 | 1351.188305 | -0.260736357 | 0.329180213 | -0.792077854 | 0.428315282 | 0.715316008 |
| ARF1 | 6235.318576 | -0.007699844 | 0.239942089 | -0.032090425 | 0.974399939 | 0.991190926 |
| ARF3 | 7001.756996 | -0.101882146 | 0.269198781 | -0.378464365 | 0.705085661 | 0.873783251 |
| ARF4 | 3057.57353 | 0.404011697 | 0.354467565 | 1.139770566 | 0.254381899 | 0.566949875 |
| ARF5 | 3810.911922 | 0.506511799 | 0.445728968 | 1.136367243 | 0.255802899 | 0.567999769 |
| ARF6 | 3426.495774 | -0.227388062 | 0.381205209 | -0.596497783 | 0.550842735 | 0.794461475 |
| ARFGAP1 | 2402.927453 | -0.022902526 | 0.358965094 | -0.06380154 | 0.949128252 | 0.98134156 |
| ARFGAP2 | 2234.768736 | -0.28338385 | 0.251674275 | -1.125994502 | 0.260167846 | 0.57337005 |
| ARFGAP3 | 817.1862223 | 0.279795733 | 0.26668192 | 1.049173985 | 0.29409805 | 0.606454309 |
| ARFGEF1 | 2418.382105 | -0.972807735 | 0.285530577 | -3.407017723 | 0.000656769 | 0.02093243 |
| ARFGEF2 | 1121.508524 | -0.635781486 | 0.347127723 | -1.831549153 | 0.067018618 | 0.297817842 |
| ARFGEF3 | 308.3918839 | -0.255984394 | 0.342410567 | -0.747594902 | 0.454704541 | 0.734645643 |
| ARFIP1 | 1251.904836 | 0.082730421 | 0.264972317 | 0.312222885 | 0.754871141 | 0.897503925 |

| | | | | | | |
|---|---|---|---|---|---|---|
| ARFIP2 | 1297.866101 | -0.169529506 | 0.23155441 | -0.732136806 | 0.464085071 | 0.741119432 |
| ARFRP1 | 444.2382952 | 0.07473495 | 0.350988608 | 0.212926997 | 0.831383905 | 0.930997148 |
| ARG2 | 1398.874989 | 0.492857057 | 0.2618427 | 1.882263884 | 0.059800196 | 0.282832055 |
| ARGFX | 16.37514987 | -0.481871945 | 0.76965259 | -0.626090201 | 0.531255779 | 0.781658561 |
| ARGLU1 | 6135.284167 | 0.220429111 | 0.226844256 | 0.971720049 | 0.33118984 | 0.639084337 |
| ARHGAP1 | 1008.079982 | -1.282423812 | 0.356167729 | -3.600617648 | 0.000317462 | 0.01312321 |
| ARHGAP10 | 1124.962671 | -1.226883 | 0.335363622 | -3.658366386 | 0.000253828 | 0.011033882 |
| ARHGAP11A | 1582.790802 | 0.394832224 | 0.497448381 | 0.793714965 | 0.427361386 | 0.714829062 |
| ARHGAP11B | 31.67557344 | -0.98726677 | 1.008250275 | -0.979188199 | 0.327486999 | 0.637370783 |
| ARHGAP12 | 1210.469807 | -0.200108256 | 0.23817149 | -0.840185597 | 0.400804333 | 0.698357934 |
| ARHGAP15 | 52.32812612 | 1.675273759 | 0.452568025 | 3.701705969 | 0.000214155 | 0.009905308 |
| ARHGAP17 | 1850.526835 | -0.298100149 | 0.265752529 | -1.121720838 | 0.261981154 | 0.575516375 |
| ARHGAP18 | 384.2069027 | -1.218920203 | 0.453824277 | -2.685885848 | 0.00723378 | 0.094039138 |
| ARHGAP19 | 1512.190292 | 0.674382239 | 0.463924857 | 1.45364541 | 0.14604464 | 0.438393992 |
| ARHGAP19-SLIT1 | 27.98586566 | -0.261299865 | 1.093855237 | -0.23887975 | 0.811198828 | 0.920897407 |
| ARHGAP21 | 2042.835543 | -0.710953114 | 0.454956872 | -1.562682439 | 0.118127307 | 0.395958766 |
| ARHGAP22 | 38.27035719 | -0.196937088 | 0.636442938 | -0.309434005 | 0.756991406 | 0.898563152 |
| ARHGAP23 | 277.2376977 | 0.19928742 | 0.371134688 | 0.536967916 | 0.591289776 | 0.819070234 |
| ARHGAP24 | 303.0156306 | -1.556282848 | 0.749701119 | -2.075871049 | 0.037905879 | 0.223664427 |
| ARHGAP25 | 137.4471796 | -0.629788013 | 0.348766407 | -1.80575881 | 0.070956022 | 0.306451904 |
| ARHGAP26 | 367.5809278 | -0.332220886 | 0.279196342 | -1.189918477 | 0.234078435 | 0.544210545 |
| ARHGAP27 | 90.64917702 | -1.08336032 | 0.442064125 | -2.450685902 | 0.014258431 | 0.135535502 |
| ARHGAP28 | 1391.014229 | -1.614695698 | 0.561171478 | -2.877365942 | 0.004010103 | 0.065575642 |
| ARHGAP29 | 1784.423571 | -1.942395264 | 0.584089551 | -3.325509352 | 0.000882571 | 0.02526319 |
| ARHGAP31 | 30.6428091 | -1.43516111 | 0.763885463 | -1.878764787 | 0.060276617 | 0.283904551 |
| ARHGAP32 | 1173.134402 | -0.919605309 | 0.346671575 | -2.652670064 | 0.007985789 | 0.100486153 |
| ARHGAP33 | 376.3741268 | 0.502584105 | 0.499435578 | 1.006304171 | 0.314269275 | 0.623891543 |
| ARHGAP35 | 2887.055339 | -0.328039288 | 0.513589809 | -0.638718452 | 0.523006106 | 0.777662198 |
| ARHGAP39 | 328.8342012 | 0.039747397 | 0.514591373 | 0.077240698 | 0.938432066 | 0.977342821 |
| ARHGAP4 | 164.7313977 | -0.125200763 | 0.502328415 | -0.249240853 | 0.80317448 | 0.917969611 |
| ARHGAP40 | 34.1045496 | -0.186712957 | 0.696154971 | -0.268206024 | 0.788540738 | 0.912309638 |
| ARHGAP42 | 943.6548943 | -1.852253172 | 0.369875193 | -5.007778857 | 5.51E-07 | 0.000128176 |
| ARHGAP44 | 425.3375451 | -0.425317911 | 0.310846066 | -1.368258949 | | |

0.171231035 0.474474162
ARHGAP45 187.6098176 0.448393025 0.359409191 1.247583634 0.212183575
0.519367057
ARHGAP5 2210.818338 0.001951432 0.348656578 0.005597004 0.995534261
0.998116797
ARHGAP6 23.49973815 0.206852609 0.877592949 0.235704502 0.813661976
0.921959809
ARHGAP8 754.0214455 -1.41584432 0.601956938 -2.35206911 0.018669306
0.155025747
ARHGDIA 1813.100338 -0.253416608 0.422353264 -0.600011008 0.5484989
0.793409306
ARHGDIB 31.85231091 -1.441182795 1.317205246 -1.094121664
0.273901622 0.586847211
ARHGEF1 719.1122589 -0.620483402 0.394961416 -1.57099751 0.116183227
0.392827221
ARHGEF10 742.7758909 -0.510860423 0.344472361 -1.483022965
0.138068309 0.426306273
ARHGEF10L 422.8197875 -0.032805047 0.521001926 -0.062965309
0.949794129 0.981628635
ARHGEF11 1406.454916 -0.702105315 0.444746589 -1.578663744
0.114413205 0.38927175
ARHGEF12 1589.323324 -0.705152172 0.370897967 -1.901202581
0.057275483 0.277081108
ARHGEF16 358.8504112 -1.237391845 0.464829078 -2.662036227
0.007766952 0.098687125
ARHGEF17 1244.913074 -0.177824172 0.543296701 -0.32730582 0.743436591
0.892534058
ARHGEF18 710.2618154 -0.350131994 0.51391351 -0.681305292
0.495678336 0.760859008
ARHGEF19 1341.304369 -0.122166532 0.423414236 -0.288527219
0.772943199 0.905717252
ARHGEF2 2022.079531 0.055799473 0.306778183 0.181888663 0.855670102
0.942139318
ARHGEF25 403.9984555 0.227677266 0.385527338 0.590560626 0.554814853
0.796360966
ARHGEF26 298.2350657 -0.04764488 0.418006301 -0.113981249 0.909252658
0.966018007
ARHGEF3 99.65018569 -1.955788014 0.430613508 -4.541864051 5.58E-06
0.000693377
ARHGEF33 13.93139752 -0.569045456 0.904210759 -0.62932834 0.529134122
0.779974131
ARHGEF34P 44.30045124 -1.284109458 0.681280601 -1.884846649
0.059450545 0.282486927
ARHGEF35 289.6643357 -1.294242327 0.669780916 -1.932336824
0.053317946 0.267056023
ARHGEF37 40.1667171 -0.890722972 0.514687362 -1.730609761
0.08352139 0.330745685
ARHGEF38 284.0058222 -1.750314757 0.679678237 -2.575210829 0.0100179
0.114014677
ARHGEF39 434.3845485 0.750889948 0.328834029 2.283492224 0.02240139
0.170396445
ARHGEF4 839.7998509 0.985888773 0.808217361 1.219831224 0.222528862
0.531400184
ARHGEF40 382.1529499 0.567259673 0.479262942 1.18360846 0.23656809
0.547336452
ARHGEF5 289.3436152 -1.745280544 0.464002942 -3.761356636
0.000168994 0.008574622
ARHGEF6 105.9647546 0.272803495 0.531395694 0.513371669 0.607691359
0.827550096
ARHGEF7 1865.385867 -0.321668489 0.260200085 -1.236235143
0.216371171 0.524036883
ARHGEF9 864.2567361 0.112725712 0.444949033 0.253345223 0.800001453
0.916311991
ARID1A 4978.824817 -0.287318173 0.284645667 -1.009388889

0.312788165 0.622907489
ARID1B 2057.973323 -0.437253196 0.365858783 -1.195141996 0.232031556 0.541311816
ARID2 1887.39985 -0.2341068 0.266225763 -0.879354415 0.37920914 0.681253679
ARID3A 4917.887126 -0.451180681 0.313586078 -1.438777781 0.150213494 0.444771508
ARID3B 5550.663153 -0.55799775 0.375042677 -1.487824679 0.136797136 0.424728769
ARID4A 690.1450483 -0.274459485 0.316714913 -0.866582132 0.386171007 0.686679504
ARID4B 1905.488087 0.067866275 0.244437996 0.277642087 0.781287121 0.909732666
ARID5A 18.72072287 0.897569056 0.707293652 1.269018963 0.204434301 0.510835602
ARID5B 462.8403184 -1.788176941 0.749080844 -2.387161486 0.016979031 0.147615406
ARIH1 2615.82951 -0.236578289 0.227851002 -1.038302604 0.299129197 0.610246278
ARIH2 2165.602458 -0.146809451 0.277095915 -0.529814562 0.596240508 0.821812117
ARIH2OS 27.14593877 0.017994362 0.658082992 0.027343606 0.978185677 0.992395125
ARL1 1904.245923 0.346684156 0.292705339 1.1844135 0.236249419 0.547141583
ARL10 351.650709 0.027713609 0.469683591 0.059004849 0.952948246 0.983108339
ARL11 74.14992064 -0.075643338 0.6039054 -0.125256932 0.900320147 0.961410357
ARL13B 314.0117981 0.18368823 0.306999255 0.59833445 0.549616794 0.793815345
ARL14EP 708.6110312 0.516220649 0.281680058 1.832648902 0.066854801 0.297817842
ARL15 658.4039195 -0.146843056 0.282124116 -0.520490975 0.602721417 0.825021776
ARL16 437.6685613 0.386767709 0.338398092 1.142937025 0.253064739 0.565800563
ARL17A 430.3565682 -0.347912039 0.391943954 -0.887657623 0.374724947 0.679043499
ARL17B 119.8233316 0.505952616 0.469409498 1.077849124 0.281101093 0.595171527
ARL2 2649.126403 0.501398553 0.307290927 1.631673796 0.102748223 0.367639927
ARL2BP 744.2577824 0.246021336 0.256458598 0.959302351 0.337406452 0.645009233
ARL3 1862.01432 1.127962027 0.343146439 3.287115642 0.001012193 0.027467362
ARL4A 860.2149167 1.670704112 0.376082714 4.442384748 8.90E-06 0.000999795
ARL4C 8461.546072 -0.173948003 0.384491838 -0.452410131 0.65097355 0.847876685
ARL4D 676.3549354 0.568643798 0.542880439 1.047456782 0.294888955 0.606966925
ARL5A 1749.624669 0.456102852 0.227818601 2.002043955 0.045280004 0.244986785
ARL5B 3627.857013 -0.194390331 0.259357037 -0.749508605 0.453550714 0.733497409
ARL6 261.6557788 0.469249516 0.31985034 1.467090879 0.142351335 0.432590018
ARL6IP1 10616.96413 0.628112086 0.225390737 2.786769738 0.005323629 0.077733184
ARL6IP4 1896.616832 0.483741457 0.40389548 1.197689701 0.23103784 0.53990682
ARL6IP5 1703.24501 0.498831914 0.239608537 2.081862026 0.037355074 0.221199778
ARL6IP6 1187.297223 0.660304131 0.268449297 2.45969775 0.013905407 0.134161613
ARL8A 833.754199 0.03928163 0.254200572 0.154530063 0.877191809 0.950622089
ARL8B 1352.816041 0.183490293 0.294951739 0.622102766 0.533874296 0.783156734
ARL9 24.86501386 1.172568683 0.696955004 1.682416621 0.092488081 0.348952169
ARMC1 1857.339969 0.41379288 0.229800544 1.800661011 0.071756326 0.308302245
ARMC10 815.408868 -0.086524798 0.374593768 -0.230983015 0.817327997 0.923820833

| | | | | | | |
|---|---|---|---|---|---|---|
| ARMC12 | 32.76448624 | 1.226453119 | 0.646898485 | 1.895897344 | 0.05797362 | 0.278988926 |
| ARMC2 | 96.99423642 | 0.166645829 | 0.455793046 | 0.365617313 | 0.714650654 | 0.87855355 |
| ARMC3 | 22.21639654 | 1.465698057 | 0.849665256 | 1.72503 | 0.084522066 | 0.332801523 |
| ARMC4 | 59.69514029 | 1.297849264 | 0.755291984 | 1.718341106 | 0.085734416 | 0.335745178 |
| ARMC5 | 234.2632575 | 0.198971214 | 0.50963815 | 0.390416641 | 0.696228485 | 0.870023227 |
| ARMC6 | 727.065517 | 0.400388502 | 0.390415083 | 1.025545681 | 0.305105782 | 0.615135352 |
| ARMC7 | 273.2283516 | 0.093966244 | 0.340706734 | 0.275798025 | 0.782703201 | 0.909909835 |
| ARMC8 | 1126.838197 | 0.001341962 | 0.233062254 | 0.005757954 | 0.995405843 | 0.998116797 |
| ARMC9 | 955.0885475 | 0.057613045 | 0.278640746 | 0.20676461 | 0.836193696 | 0.93270414 |
| ARMCX1 | 1103.404708 | 0.135825636 | 0.28286741 | 0.480174214 | 0.631103521 | 0.838320891 |
| ARMCX2 | 2967.714217 | 0.191713489 | 0.240557591 | 0.796954644 | 0.425477373 | 0.713673173 |
| ARMCX3 | 1474.694179 | 0.064242907 | 0.269953352 | 0.237977807 | 0.811898307 | 0.921333624 |
| ARMCX4 | 681.8910308 | -0.045305499 | 0.444396905 | -0.101948278 | 0.918797729 | 0.969677217 |
| ARMCX5 | 661.4053436 | -0.031663382 | 0.272010425 | -0.116405033 | 0.907331547 | 0.965296417 |
| ARMCX5-GPRASP2 | 17.67392526 | -2.389730126 | 1.138442155 | -2.099123012 | 0.035806058 | 0.216454473 |
| ARMCX6 | 1152.158304 | -0.320564989 | 0.279415604 | -1.14726946 | 0.251270272 | 0.564075361 |
| ARMCX7P | 93.72603409 | 24.63024489 | 4.328759976 | 5.689907739 | 1.27E-08 | 1.01E-05 |
| ARMH1 | 77.22670015 | -0.039338787 | 0.476700968 | -0.082522984 | 0.934230842 | 0.97507153 |
| ARMH3 | 606.7118716 | -0.498916971 | 0.301467891 | -1.654958905 | 0.097932834 | 0.359375073 |
| ARMH4 | 598.8896939 | 0.34894512 | 0.328982091 | 1.060681205 | 0.288834805 | 0.601911102 |
| ARMT1 | 2273.487449 | -0.45918807 | 0.232801384 | -1.972445621 | 0.048558758 | 0.25374113 |
| ARNT | 1353.93025 | -0.489521649 | 0.367579442 | -1.331743819 | 0.182944385 | 0.486835514 |
| ARNT2 | 455.0721543 | 1.112701786 | 0.691496056 | 1.60912239 | 0.107589582 | 0.376005887 |
| ARNTL | 97.87126622 | -0.358653875 | 0.637516511 | -0.562579744 | 0.573721088 | 0.808163661 |
| ARNTL2 | 259.8979356 | -0.110775839 | 0.284462676 | -0.389421347 | 0.696964483 | 0.870026319 |
| ARPC1A | 4176.517804 | 0.103623561 | 0.326324323 | 0.31754777 | 0.750827999 | 0.896207755 |
| ARPC1B | 1121.725386 | 0.343966802 | 0.410430916 | 0.838062603 | 0.401995545 | 0.699116854 |
| ARPC2 | 4941.225758 | 0.100778272 | 0.348958758 | 0.288797084 | 0.772736664 | 0.905717252 |
| ARPC3 | 4406.632555 | 0.337181741 | 0.38095212 | 0.885102676 | 0.376101256 | 0.680079109 |
| ARPC4 | 3377.591505 | 0.249853916 | 0.445298274 | 0.561093387 | 0.574733876 | 0.808359708 |
| ARPC4-TTLL3 | 166.1693417 | 0.111613121 | 0.837180641 | 0.133320236 | 0.893940124 | 0.958104248 |
| ARPC5 | 3563.750825 | 0.23723674 | 0.370127978 | 0.640958679 | 0.52154953 | 0.776633705 |
| ARPC5L | 861.3532734 | 0.369074753 | 0.415718096 | 0.887800547 | 0.374648049 | 0.678987738 |
| ARPIN | 608.414282 | -0.370809205 | 0.2894898 | -1.280905944 | 0.200226705 | 0.506121566 |
| ARPIN-AP3S2 | 579.3357717 | -0.172575664 | 0.317000458 | -0.544401939 | 0.586164912 | 0.815844759 |
| ARPP19 | 4997.189019 | 0.421390996 | 0.255697324 | 1.648007067 | 0.099351221 | 0.361473741 |
| ARRB1 | 817.7370527 | -1.051057451 | 0.525908771 | -1.998554709 | 0.045656555 | 0.245971447 |
| ARRB2 | 1441.162295 | 0.611667562 | 0.322654376 | 1.895736141 | 0.057994944 | 0.278988926 |
| ARRDC1 | 417.7303817 | -0.46874174 | 0.410513671 | -1.141841975 | 0.253519711 | 0.565942797 |
| ARRDC2 | 61.85440773 | -0.251648716 | 0.447297938 | -0.562597532 | 0.573708973 | 0.808163661 |

| | | | | | | |
|---|---|---|---|---|---|---|
| ARRDC3 | 2131.807757 | -0.035216521 | 0.33003877 | -0.106704193 | 0.915023656 | 0.968859847 |
| ARRDC4 | 739.4546753 | 1.612461421 | 0.62073333 | 2.597671727 | 0.009385818 | 0.1094483 |
| ARSA | 321.8706388 | 0.000805433 | 0.45533113 | 0.001768896 | 0.998588626 | 0.999115481 |
| ARSB | 483.2598787 | -0.083114101 | 0.276459351 | -0.300637693 | 0.763690785 | 0.902262511 |
| ARSD | 276.3305546 | -0.995486064 | 0.488078459 | -2.03960254 | 0.041389929 | 0.233690589 |
| ARSE | 45.22738071 | -0.704914161 | 0.529001976 | -1.33253597 | 0.18268413 | 0.486683209 |
| ARSG | 245.6034364 | -0.320786046 | 0.312029693 | -1.028062561 | 0.3039204 | 0.61411513 |
| ARSI | 235.2386984 | 0.511548817 | 0.288970477 | 1.770245948 | 0.076686178 | 0.317958513 |
| ARSK | 463.7702695 | 0.175072324 | 0.33569858 | 0.521516428 | 0.602007065 | 0.825021776 |
| ART5 | 265.7358965 | 0.86413139 | 0.407184528 | 2.122210769 | 0.033820043 | 0.209507499 |
| ARTN | 34.8959878 | 1.397074288 | 0.568054686 | 2.459401044 | 0.013916906 | 0.134161613 |
| ARV1 | 438.074324 | 0.574579362 | 0.290062575 | 1.980880717 | 0.047604652 | 0.251378411 |
| ARVCF | 764.8254578 | 0.32022372 | 0.562391882 | 0.569396057 | 0.569087392 | 0.805450586 |
| AS3MT | 789.7803593 | 0.66621629 | 0.329866314 | 2.019655423 | 0.043419142 | 0.239726164 |
| ASAH1 | 1887.323599 | 0.009691667 | 0.300343218 | 0.032268639 | 0.974257819 | 0.991112812 |
| ASAH2 | 47.31175445 | -0.528600341 | 0.540370502 | -0.97821835 | 0.327966343 | 0.637787615 |
| ASAH2B | 398.7738484 | -0.037142283 | 0.305364886 | -0.121632462 | 0.903190104 | 0.963390077 |
| ASAP1 | 1001.298661 | -0.434747622 | 0.393591961 | -1.104564284 | 0.269348434 | 0.582188661 |
| ASAP2 | 1077.456912 | -0.144393418 | 0.284621832 | -0.507316735 | 0.611932599 | 0.829098987 |
| ASAP3 | 421.1214797 | -0.654420853 | 0.329104342 | -1.988490487 | 0.04675747 | 0.249119645 |
| ASB1 | 1000.803786 | 0.13702244 | 0.283733183 | 0.482927089 | 0.629147507 | 0.837367367 |
| ASB13 | 392.4562196 | 0.382683595 | 0.363553225 | 1.05262055 | 0.292514935 | 0.605587347 |
| ASB3 | 996.2044289 | 0.271231078 | 0.231434395 | 1.171956647 | 0.241214462 | 0.552673201 |
| ASB6 | 588.7948564 | -0.2654878 | 0.345105784 | -0.769293974 | 0.441718814 | 0.72517218 |
| ASB7 | 570.6388979 | -0.413565051 | 0.336449051 | -1.229205579 | 0.218994738 | 0.527527654 |
| ASB8 | 242.3914855 | 0.00705783 | 0.33509327 | 0.021062285 | 0.983195971 | 0.993766984 |
| ASCC1 | 855.2872494 | -0.111200614 | 0.232965124 | -0.477327302 | 0.633129073 | 0.839434984 |
| ASCC2 | 1142.829967 | -0.764260512 | 0.347360769 | -2.200192367 | 0.02779325 | 0.190782329 |
| ASCC3 | 2146.895767 | 0.350047722 | 0.426887372 | 0.82000018 | 0.412216005 | 0.70515068 |
| ASCL4 | 26.22122455 | 1.442425306 | 1.447215608 | 0.996689988 | 0.318915011 | 0.62891715 |
| ASDURF | 21.16895016 | 0.553202144 | 1.107282411 | 0.499603479 | 0.617354308 | 0.831835425 |
| ASF1A | 932.4735594 | 0.574503135 | 0.263108765 | 2.183519559 | 0.02899757 | 0.194741982 |
| ASF1B | 642.5164104 | 0.670177972 | 0.280231101 | 2.391518892 | 0.01677882 | 0.146402531 |
| ASGR1 | 138.0646796 | 0.663184771 | 0.533344493 | 1.243445428 | 0.213703739 | 0.520735532 |
| ASH1L | 968.6274971 | -0.591601143 | 0.534455231 | -1.106923665 | 0.268326933 | 0.581480913 |
| ASH2L | 1865.301705 | -0.087562563 | 0.249868711 | -0.350434286 | 0.726012799 | 0.884472832 |
| ASIC1 | 140.9847638 | 1.004942359 | 0.556532736 | 1.805720119 | 0.070962068 | 0.306451904 |
| ASIC3 | 114.9764403 | -0.586009036 | 0.403042569 | -1.453963133 | 0.145956528 | 0.438251053 |
| ASIC4 | 14.64040873 | 1.393788158 | 0.837013074 | 1.665192817 | 0.095874294 | 0.355538721 |
| ASL | 754.0256346 | -0.238805245 | 0.286322743 | -0.834042182 | 0.404257214 | 0.700367583 |
| ASLP1 | 27.150317 | 0.170154054 | 0.663544603 | 0.256431977 | 0.797617299 | 0.915916373 |
| ASMTL | 2577.762298 | 0.41535346 | 0.392252733 | 1.058892457 | 0.28964876 | 0.602448771 |
| ASNA1 | 2556.563073 | 0.313854224 | 0.299584445 | 1.047631909 | 0.29480823 | 0.606939295 |
| ASNS | 1574.659383 | 0.352727975 | 0.329924545 | 1.069117106 | 0.285016905 | 0.598807847 |
| ASNSD1 | 1783.989758 | 0.324957007 | 0.227060556 | 1.431146883 | 0.152388123 | |

0.448299441
ASPA 36.75798256 -1.57994104 0.653544561 -2.417495506 0.015627727
0.141376414
ASPH 2560.070228 0.12003868 0.271135429 0.442725911 0.657963999 0.850760404
ASPHD1 71.93870193 0.957641246 0.510801179 1.874782764 0.060822615
0.284357441
ASPHD2 149.9894307 0.896883473 0.439034924 2.042852227 0.041067069
0.232934829
ASPM 1857.846875 0.736971616 0.485402161 1.51827016 0.128946307 0.413453061
ASPRV1 22.09400946 0.409269115 0.622018974 0.65796886 0.510558141
0.770330936
ASPSCR1 675.0346346 0.371942268 0.381782886 0.974224569 0.329945048
0.638681113
ASRGL1 689.9230074 -0.199216537 0.242669993 -0.820936016
0.411682715 0.704686727
ASS1 3718.019881 0.199007263 0.256480193 0.775916693 0.437798176 0.7217818
ASTE1 212.7158148 0.065012037 0.290831054 0.223538843 0.823116156 0.926014081
ASTN1 167.1119484 1.000854006 0.655678785 1.526439515 0.126900443 0.41022941
ASTN2 1021.966638 -0.810078507 0.320861156 -2.524701081 0.011579674
0.121704879
ASXL1 2373.457308 -0.436549724 0.48886168 -0.892992315 0.371861288
0.677193964
ASXL2 1073.498251 -0.394531654 0.361594898 -1.091087449 0.275234405
0.588275734
ASXL3 65.26460278 0.158905908 0.892974607 0.177951206 0.858761296 0.942495112
ATAD1 839.0345582 0.370015337 0.285323558 1.29682715 0.194690667 0.500449357
ATAD2 958.9541705 0.243432175 0.443074108 0.549416386 0.582719731 0.813587433
ATAD2B 589.5474709 -0.214656827 0.382419923 -0.56131183 0.574584978
0.808359708
ATAD3A 561.4671718 0.039484832 0.465999757 0.084731444 0.932474897
0.97453841
ATAD3B 596.5082768 -0.004970992 0.378353833 -0.013138475
0.989517315 0.996285567
ATAD3C 40.82756934 0.36245962 0.743231231 0.487680826 0.625775935
0.835487698
ATAD5 855.2747276 0.248615866 0.392754577 0.633005649 0.526729954 0.778755887
ATAT1 42.62044272 -0.506236367 0.588140632 -0.860740339 0.389381071
0.689219487
ATCAY 262.2519269 1.183523085 0.655555444 1.805374503 0.071016098 0.306597959
ATE1 734.4589863 -0.103025557 0.271382501 -0.379632277 0.704218398
0.873421952
ATF1 564.3969526 0.413111551 0.25055945 1.648756617 0.099197506 0.361452832
ATF2 1066.731486 0.2487587 0.427024464 0.582539692 0.560203214 0.800493827
ATF3 485.0410742 -0.917638297 0.505797492 -1.814240504 0.069640707
0.303425952
ATF4 9279.07669 0.034039456 0.274456464 0.124024973 0.901295502 0.961878265
ATF5 74.91661344 0.325746166 0.398419208 0.81759654 0.413587604 0.705778726
ATF6 1448.984291 -0.19002372 0.237747729 -0.799266183 0.424136083
0.71328772
ATF6B 320.6850493 -0.935995524 0.47117717 -1.986504406 0.046977344
0.249626249
ATF7 706.7761379 -0.510407563 0.333797223 -1.529094693 0.126240971
0.409038191
ATF7IP 5725.955321 -0.309112421 0.311262094 -0.993093688 0.3206643
0.630327731
ATF7IP2 268.2807741 -0.412232018 0.628273827 -0.656134315
0.511737707 0.771343542
ATG10 703.975809 -0.127755812 0.340268082 -0.375456349 0.707321104
0.87520286
ATG101 772.743578 0.660112938 0.368067155 1.793457877 0.07289974
0.310624251
ATG12 1769.982556 0.446362527 0.270792685 1.648355183 0.099279807 0.361452832
ATG13 1982.224587 -0.234035937 0.222941028 -1.049766118 0.293825657
0.606397638

| | | | | | | |
|---|---|---|---|---|---|---|
| ATG14 | 750.7072112 | -0.57261244 | 0.266367582 | -2.149707694 | 0.031578343 | 0.202592072 |
| ATG16L1 | 348.5050603 | -0.068597428 | 0.275993451 | -0.248547303 | 0.803710975 | 0.917969611 |
| ATG16L2 | 508.2888361 | -0.434719053 | 0.343254109 | -1.266464236 | 0.205346926 | 0.511820585 |
| ATG2A | 466.6184377 | -0.815272041 | 0.496161272 | -1.643159367 | 0.100349964 | 0.362478407 |
| ATG2B | 584.5746729 | -0.65435484 | 0.384378342 | -1.702371772 | 0.088685698 | 0.342301733 |
| ATG3 | 1154.433284 | 0.431780065 | 0.30966069 | 1.394365122 | 0.163207375 | 0.464573649 |
| ATG4A | 156.1553116 | -0.377568365 | 0.324168068 | -1.164730282 | 0.244128187 | 0.555457814 |
| ATG4B | 1691.072445 | 0.126010241 | 0.277179404 | 0.454616178 | 0.649385394 | 0.847281162 |
| ATG4C | 732.3479063 | 0.57346516 | 0.279707459 | 2.050231916 | 0.040341805 | 0.231564707 |
| ATG4D | 420.6149731 | 0.444818554 | 0.351420787 | 1.26577189 | 0.205594762 | 0.511932925 |
| ATG5 | 1145.966712 | 0.326228819 | 0.252753741 | 1.29069828 | 0.196808321 | 0.50258876 |
| ATG7 | 611.8022875 | -0.311859055 | 0.27388014 | -1.138669841 | 0.254840886 | 0.567256913 |
| ATG9A | 1478.433266 | -0.259024724 | 0.271393395 | -0.954425306 | 0.339868402 | 0.646629378 |
| ATG9B | 489.6543522 | -2.830198718 | 0.579476153 | -4.88406417 | 1.04E-06 | 0.000185481 |
| ATIC | 4938.968926 | 0.041192117 | 0.31376072 | 0.131285131 | 0.895549754 | 0.95881336 |
| ATL1 | 116.7005143 | -0.351559882 | 0.343384098 | -1.023809443 | 0.305925289 | 0.615951236 |
| ATL2 | 1716.976342 | 0.48411162 | 0.284954718 | 1.698907192 | 0.089336672 | 0.342726775 |
| ATL3 | 436.8932573 | -0.028311849 | 0.509754628 | -0.055540151 | 0.955708143 | 0.984323736 |
| ATM | 1502.836427 | -0.234752594 | 0.369705233 | -0.634972332 | 0.52544646 | 0.778371208 |
| ATMIN | 2417.853258 | -0.141216067 | 0.242295746 | -0.582825201 | 0.560010978 | 0.800294513 |
| ATN1 | 3193.161534 | -0.452140767 | 0.319041821 | -1.417183382 | 0.156429319 | 0.455090998 |
| ATOX1 | 909.9242622 | 0.811299894 | 0.549122531 | 1.477447833 | 0.139555632 | 0.428322576 |
| ATP10A | 11.5125421 | 0.492234462 | 1.208337126 | 0.407365173 | 0.683739802 | 0.863715015 |
| ATP10B | 73.21414509 | 0.078781665 | 0.957885534 | 0.082245385 | 0.934451584 | 0.97507153 |
| ATP10D | 219.9385614 | -0.771439732 | 0.35479741 | -2.174310495 | 0.029681825 | 0.196982924 |
| ATP11A | 1987.664562 | -1.328983197 | 0.410453846 | -3.23783833 | 0.00120439 | 0.030537472 |
| ATP11B | 859.1093961 | -0.410111019 | 0.265361495 | -1.545480514 | 0.122230079 | 0.402857381 |
| ATP11C | 1387.650846 | -0.459415078 | 0.320249091 | -1.434555446 | 0.151413818 | 0.446906428 |
| ATP12A | 101.5747457 | -2.477715442 | 0.716278529 | -3.459150793 | 0.000541881 | 0.018350186 |
| ATP13A1 | 1098.504058 | -0.633371091 | 0.44724403 | -1.416164439 | 0.156727365 | 0.455611392 |
| ATP13A2 | 874.8540347 | -0.453801246 | 0.430239463 | -1.054764346 | 0.291533115 | 0.604602035 |
| ATP13A3 | 1776.605109 | 0.097702475 | 0.399898833 | 0.244317979 | 0.80698456 | 0.919137032 |
| ATP13A4 | 62.70328779 | -0.84117122 | 0.867099393 | -0.970097808 | 0.331997742 | 0.639828376 |
| ATP1A1 | 13189.09796 | -0.503481496 | 0.267493858 | -1.882217031 | 0.059806554 | 0.282832055 |
| ATP1A2 | 5266.261517 | 1.350597541 | 0.677081848 | 1.994733051 | 0.046072003 | 0.246950947 |
| ATP1A3 | 91.82482612 | 0.12158423 | 0.61033982 | 0.199207435 | 0.842100484 | 0.935422069 |

| | | | | | | |
|---|---|---|---|---|---|---|
| ATP1A4 | 21.8653518 | -1.333122305 | 0.984708338 | -1.353824532 | 0.175792365 | 0.480011874 |
| ATP1B1 | 1105.727127 | 0.018687931 | 0.253124494 | 0.073829012 | 0.941146442 | 0.978289323 |
| ATP1B2 | 375.6474713 | 0.311509367 | 0.343922589 | 0.905754312 | 0.365065909 | 0.671700076 |
| ATP1B3 | 4182.4952 | 0.721375103 | 0.340161623 | 2.120683387 | 0.033948456 | 0.210131383 |
| ATP23 | 89.78368383 | 0.294991477 | 0.468147498 | 0.630125073 | 0.528612756 | 0.779660133 |
| ATP2A1 | 83.94973745 | 0.28812193 | 0.492427197 | 0.585105641 | 0.558476686 | 0.798855459 |
| ATP2A2 | 6660.701349 | -0.231786595 | 0.358276356 | -0.64694918 | 0.517664835 | 0.773950969 |
| ATP2A3 | 336.4325454 | -0.094549955 | 0.446333476 | -0.211837024 | 0.832234183 | 0.931245375 |
| ATP2B1 | 4775.96915 | -1.755350547 | 0.564093163 | -3.111809645 | 0.001859444 | 0.04012748 |
| ATP2B4 | 3698.693323 | -0.96693886 | 0.345228637 | -2.800865153 | 0.005096581 | 0.075897737 |
| ATP2C1 | 2934.712188 | -0.443645764 | 0.246861318 | -1.797145731 | 0.072312487 | 0.309552129 |
| ATP2C2 | 249.6319702 | -2.293032507 | 0.59140042 | -3.877292662 | 0.000105625 | 0.006358893 |
| ATP5F1A | 20848.78807 | 0.185717846 | 0.2225749 | 0.834406061 | 0.404052203 | 0.700367583 |
| ATP5F1B | 18095.91353 | 0.291568669 | 0.22550685 | 1.29294817 | 0.196028986 | 0.501797297 |
| ATP5F1C | 4691.602898 | 0.605545822 | 0.263254429 | 2.300230328 | 0.021435174 | 0.166254106 |
| ATP5F1D | 562.2581717 | 0.869495712 | 0.521203474 | 1.668246193 | 0.095266864 | 0.354499289 |
| ATP5F1E | 3473.526046 | 1.246117659 | 0.417795003 | 2.982605465 | 0.002858062 | 0.052557154 |
| ATP5IF1 | 755.2529251 | 0.893371738 | 0.428872473 | 2.083070827 | 0.037244768 | 0.220977855 |
| ATP5MC1 | 1113.852776 | 1.061127473 | 0.402384381 | 2.637099061 | 0.008361841 | 0.103556802 |
| ATP5MC2 | 9186.099867 | 0.891630849 | 0.387202654 | 2.30274984 | 0.021292922 | 0.165489202 |
| ATP5MC3 | 6026.817986 | 1.020561121 | 0.428064672 | 2.384128352 | 0.01711963 | 0.14773715 |
| ATP5MD | 2652.778646 | 1.148768157 | 0.450002088 | 2.552806279 | 0.010685891 | 0.117606289 |
| ATP5ME | 2922.029478 | 1.269646401 | 0.471172599 | 2.694652456 | 0.007046208 | 0.092312631 |
| ATP5MF | 3093.034273 | 0.827453371 | 0.404792782 | 2.044140624 | 0.040939657 | 0.232707809 |
| ATP5MF-PTCD1 | 239.5206713 | -1.018856498 | 0.990684535 | -1.028436866 | 0.303744374 | 0.614002998 |
| ATP5MG | 4234.43022 | 1.041779769 | 0.42241346 | 2.46625609 | 0.013653368 | 0.132885181 |
| ATP5MPL | 2629.28229 | 1.071228448 | 0.394363869 | 2.716345316 | 0.006600703 | 0.089064189 |
| ATP5PB | 6248.774503 | 0.648935955 | 0.327330597 | 1.982509309 | 0.047422267 | 0.25076445 |
| ATP5PD | 3254.344516 | 1.024846009 | 0.42978705 | 2.384543714 | 0.017100316 | 0.14773715 |
| ATP5PF | 2082.580803 | 0.841963064 | 0.412857809 | 2.039353612 | 0.041414749 | 0.233743731 |
| ATP5PO | 7303.121489 | 0.939895016 | 0.359008851 | 2.618027421 | 0.00884397 | 0.106517932 |
| ATP6AP1 | 2378.544882 | -0.52693576 | 0.287075441 | -1.835530612 | 0.066427108 | 0.297462837 |
| ATP6AP1L | 272.0262932 | -0.267289233 | 0.321522151 | -0.831324472 | | |

| | | | | | | |
|---|---|---|---|---|---|---|
| | | | | | 0.405790353 | 0.701371854 |
| ATP6AP2 | 3215.647951 | -0.044895978 | 0.384501045 | -0.116764254 | 0.907046871 | 0.965296417 |
| ATP6V0A1 | 1440.748807 | -0.770080449 | 0.353678171 | -2.177347975 | 0.029454614 | 0.196075451 |
| ATP6V0A2 | 819.6974123 | 0.098454617 | 0.344712047 | 0.285614087 | 0.775173713 | 0.906557532 |
| ATP6V0A4 | 30.25016284 | -2.278476168 | 0.898284013 | -2.536476364 | 0.01119743 | 0.120297968 |
| ATP6V0B | 1909.523995 | 0.682466438 | 0.336020314 | 2.031027319 | 0.042252223 | 0.236528876 |
| ATP6V0C | 2001.904437 | 0.230706831 | 0.439741169 | 0.52464233 | 0.599831855 | 0.824047262 |
| ATP6V0D1 | 1300.500093 | -0.072360169 | 0.376490399 | -0.192196585 | 0.847588221 | 0.938207222 |
| ATP6V0D2 | 88.49181236 | -0.480470892 | 0.934126502 | -0.514353132 | 0.60700512 | 0.82679787 |
| ATP6V0E1 | 1607.294388 | 0.432444018 | 0.45436002 | 0.951765119 | 0.341216116 | 0.648152255 |
| ATP6V0E2 | 810.2307405 | -0.092060198 | 0.356290155 | -0.258385466 | 0.796109433 | 0.915329761 |
| ATP6V1A | 2162.783527 | 0.403534998 | 0.326170971 | 1.23718857 | 0.216017084 | 0.523764612 |
| ATP6V1B1 | 510.5879341 | -1.908622788 | 0.499165501 | -3.823627203 | 0.000131503 | 0.007254645 |
| ATP6V1B2 | 2270.48649 | -0.327135782 | 0.27891102 | -1.172903751 | 0.240834405 | 0.552437022 |
| ATP6V1C1 | 1314.126029 | -0.47350023 | 0.290269686 | -1.631242437 | 0.102839174 | 0.367817927 |
| ATP6V1C2 | 95.15077003 | -1.463760669 | 0.647424548 | -2.260897695 | 0.023765593 | 0.176393253 |
| ATP6V1D | 1908.085628 | 0.531695372 | 0.28956257 | 1.836202008 | 0.066327786 | 0.297445711 |
| ATP6V1E1 | 2490.269725 | 0.218074074 | 0.316576642 | 0.688850803 | 0.490917162 | 0.758593654 |
| ATP6V1E2 | 80.74129184 | 0.329300115 | 0.466860314 | 0.705350413 | 0.480592195 | 0.752509463 |
| ATP6V1F | 1540.466261 | 0.538598731 | 0.311987375 | 1.72634784 | 0.084284853 | 0.332430848 |
| ATP6V1G1 | 3823.631427 | 0.456927662 | 0.369807383 | 1.235582852 | 0.216613662 | 0.524289385 |
| ATP6V1H | 1339.004172 | -0.004700796 | 0.400031089 | -0.011751076 | 0.990624214 | 0.996726674 |
| ATP7A | 159.4383913 | -0.549131624 | 0.451658986 | -1.215810248 | 0.224057214 | 0.533370782 |
| ATP7B | 498.7892821 | -1.384347761 | 0.388942556 | -3.559260202 | 0.000371901 | 0.014489772 |
| ATP8A1 | 232.5685134 | 0.134255048 | 0.424488159 | 0.316275132 | 0.751793685 | 0.896586461 |
| ATP8A2 | 183.2159534 | -0.977129012 | 0.574284693 | -1.701471456 | 0.088854493 | 0.34239561 |
| ATP8B1 | 580.8907485 | -1.831982663 | 0.660894727 | -2.771973492 | 0.005571757 | 0.079919047 |
| ATP8B2 | 1283.377902 | 0.101268815 | 0.454702459 | 0.222714465 | 0.823757742 | 0.926468056 |
| ATP8B3 | 87.38681607 | 1.217479322 | 0.551333052 | 2.20824657 | 0.02722709 | 0.188612866 |
| ATP8B4 | 10.926022 | -0.951861177 | 1.052186698 | -0.904650456 | 0.365650596 | 0.672153785 |
| ATP9A | 500.3233474 | -1.526292509 | 0.316838165 | -4.817262181 | 1.46E-06 | 0.000240001 |
| ATP9B | 622.4466558 | -0.791165771 | 0.482290673 | -1.640433489 | 0.100915067 | 0.363826635 |
| ATPAF1 | 1705.483396 | 0.449071605 | 0.247933261 | 1.81126003 | 0.07010061 | |

0.304116772

| | | | | | | |
|---|---|---|---|---|---|---|
| ATPAF2 | 326.1375993 | 0.003944367 | 0.306356348 | 0.012875093 | 0.989727446 | 0.996285567 |
| ATPSCKMT | 295.8291974 | 0.596980411 | 0.270459061 | 2.207285671 | 0.027294108 | 0.188904612 |
| ATR | 678.4191575 | -0.13751187 | 0.399885097 | -0.343878456 | 0.730937695 | 0.886911603 |
| ATRAID | 2104.83377 | 0.618323194 | 0.366032399 | 1.689258096 | 0.09116998 | 0.346217714 |
| ATRIP | 437.7090587 | -0.146982532 | 0.28666832 | -0.512726807 | 0.608142435 | 0.827607804 |
| ATRN | 1544.173326 | -0.370819683 | 0.399599384 | -0.927978615 | 0.353418663 | 0.660553718 |
| ATRNL1 | 81.83830967 | 0.607677665 | 0.581140134 | 1.0456646 | 0.295715912 | 0.60699582 |
| ATRX | 2964.117601 | -0.12472125 | 0.29296368 | -0.425722566 | 0.670310017 | 0.857949318 |
| ATXN1 | 137.7549635 | -1.517037828 | 0.406669972 | -3.73039057 | 0.000191183 | 0.009160128 |
| ATXN10 | 3565.960605 | 0.590444133 | 0.269383009 | 2.191838809 | 0.028391148 | 0.192544525 |
| ATXN1L | 1536.269 | -0.572503648 | 0.33251852 | -1.721719582 | 0.085120328 | 0.334289542 |
| ATXN2 | 1117.937704 | -0.135459534 | 0.35054351 | -0.386427163 | 0.699180337 | 0.871039767 |
| ATXN2L | 4033.244506 | -0.391002594 | 0.369748235 | -1.057483328 | 0.29029106 | 0.60328736 |
| ATXN3 | 677.5890413 | -0.025690837 | 0.255586288 | -0.100517274 | 0.91993367 | 0.969770994 |
| ATXN7 | 704.1838666 | -0.730036559 | 0.297880398 | -2.450770726 | 0.014255072 | 0.135535502 |
| ATXN7L1 | 359.0071265 | -1.142553308 | 0.451383263 | -2.531226569 | 0.01136644 | 0.120838052 |
| ATXN7L2 | 176.9538279 | 0.28117271 | 0.441944378 | 0.636217416 | 0.524634723 | 0.777662198 |
| ATXN7L3 | 3486.316867 | -0.513681606 | 0.309957371 | -1.657265335 | 0.097465842 | 0.358846393 |
| ATXN7L3B | 3813.916894 | -0.256971468 | 0.239507734 | -1.072915115 | 0.283309204 | 0.597121968 |
| AUH | 531.5913499 | -0.190056516 | 0.3737866 | -0.508462625 | 0.611128944 | 0.829058038 |
| AUNIP | 283.1490124 | 0.256517731 | 0.287301321 | 0.8928526 | 0.371936114 | 0.677193964 |
| AUP1 | 1852.581363 | -0.145543106 | 0.249568358 | -0.583179324 | 0.559772588 | 0.800179962 |
| AURKA | 1723.889588 | 0.957688448 | 0.321488723 | 2.978917704 | 0.002892685 | 0.052996691 |
| AURKAIP1 | 1505.245116 | 0.861193652 | 0.493117683 | 1.746426223 | 0.080736917 | 0.325104967 |
| AURKB | 1534.53962 | 1.15371417 | 0.278611723 | 4.140939075 | 3.46E-05 | 0.002806123 |
| AURKC | 97.2471834 | 0.617857953 | 0.614301258 | 1.005789823 | 0.314516684 | 0.624045314 |
| AUTS2 | 2072.726777 | 0.167069626 | 0.321606916 | 0.519483934 | 0.603423314 | 0.825211561 |
| AVEN | 395.6835773 | 0.01028309 | 0.371122631 | 0.027708064 | 0.977894992 | 0.99226352 |
| AVIL | 15.65089253 | -0.590083413 | 0.873687424 | -0.675394194 | 0.499425354 | 0.76281557 |
| AVL9 | 1306.326125 | -0.117474178 | 0.274648298 | -0.427725854 | 0.668850725 | 0.857383552 |
| AVPI1 | 104.7751907 | 0.168446196 | 0.390153701 | 0.431743172 | 0.665928086 | 0.855444115 |
| AXIN1 | 969.7626283 | -0.029250617 | 0.467066667 | -0.062626214 | 0.950064155 | 0.981636196 |
| AXIN2 | 1851.565785 | -0.141539014 | 0.360368247 | -0.392762167 | 0.69449515 | 0.86892666 |
| AXL | 1290.149186 | 0.150527581 | 0.237821901 | 0.63294247 | 0.526771212 | 0.778755887 |
| AZI2 | 1108.733338 | 0.325551332 | 0.286279299 | 1.137180834 | 0.255462698 | 0.567840162 |
| AZIN1 | 7949.211064 | -0.56007314 | 0.376948882 | -1.485806609 | 0.137330281 | 0.425712285 |

| | | | | | | |
|---|---|---|---|---|---|---|
| AZIN2 | 88.18086761 | -0.386767024 | 0.381863755 | -1.012840362 | 0.311136418 | 0.621167338 |
| AZU1 | 10.95174594 | -0.426653107 | 1.128779491 | -0.377977373 | 0.705447403 | 0.873945986 |
| B2M | 500.0254652 | 0.805208273 | 0.396742205 | 2.02955033 | 0.042402268 | 0.236937315 |
| B3GALNT1 | 269.3820919 | -0.199365629 | 0.292542861 | -0.681492033 | 0.495560206 | 0.760859008 |
| B3GALNT2 | 589.3126676 | -0.164613701 | 0.278991145 | -0.590031992 | 0.555169202 | 0.796360966 |
| B3GALT1 | 149.7830709 | -0.34571768 | 0.507363461 | -0.68140043 | 0.495618151 | 0.760859008 |
| B3GALT4 | 15.55972709 | -0.016891981 | 0.910279176 | -0.018556923 | 0.985194567 | 0.99460071 |
| B3GALT5 | 86.37666635 | -0.368793043 | 0.532249622 | -0.69289489 | 0.488375517 | 0.757013457 |
| B3GALT6 | 679.7275611 | 0.614190599 | 0.342529359 | 1.79310352 | 0.072956373 | 0.310730246 |
| B3GAT1 | 620.5914996 | 0.274009033 | 0.400447015 | 0.684257898 | 0.493812328 | 0.759876947 |
| B3GAT2 | 16.76239083 | 0.304949794 | 0.927669097 | 0.328726908 | 0.742362114 | 0.892150449 |
| B3GAT3 | 921.8524534 | 1.090347153 | 0.417953792 | 2.608774398 | 0.009086713 | 0.107783048 |
| B3GLCT | 294.5358468 | 0.187905344 | 0.406318834 | 0.462457872 | 0.643753006 | 0.845466381 |
| B3GNT10 | 77.74320369 | -0.16578064 | 0.41212674 | -0.402256452 | 0.6874953 | 0.866494242 |
| B3GNT2 | 5134.89136 | -0.349415947 | 0.404515489 | -0.863788795 | 0.387703924 | 0.687775519 |
| B3GNT4 | 16.11218743 | -0.152843649 | 0.817007946 | -0.18707731 | 0.851600018 | 0.939609009 |
| B3GNT5 | 520.5715261 | 0.26542149 | 0.354018439 | 0.749739168 | 0.453411812 | 0.733497409 |
| B3GNT7 | 717.1714909 | -1.023043793 | 0.746477987 | -1.370494255 | 0.170532668 | 0.473631094 |
| B3GNT8 | 52.25561528 | 0.040731868 | 0.716020185 | 0.05688648 | 0.954635624 | 0.984285514 |
| B3GNT9 | 56.35675645 | -0.342924516 | 0.451289398 | -0.759877182 | 0.447328002 | 0.728949036 |
| B3GNTL1 | 97.81249095 | 0.509595608 | 0.531600446 | 0.958606436 | 0.337757049 | 0.645135506 |
| B4GALNT1 | 18.78192678 | 0.979984458 | 0.758166452 | 1.292571645 | 0.196159252 | 0.501797297 |
| B4GALNT2 | 24.15109027 | -2.079711828 | 0.874529769 | -2.378091519 | 0.017402506 | 0.149075901 |
| B4GALNT3 | 1910.459159 | 0.031741591 | 0.3443066 | 0.092189899 | 0.926547163 | 0.972261514 |
| B4GALNT4 | 1096.4075 | 0.124908575 | 0.418974628 | 0.298129211 | 0.765604548 | 0.903026935 |
| B4GALT1 | 1196.378458 | -0.657157088 | 0.342956465 | -1.916153084 | 0.055345613 | 0.272191885 |
| B4GALT2 | 1365.369518 | 0.553581497 | 0.27744991 | 1.995248424 | 0.046015792 | 0.246950947 |
| B4GALT3 | 1185.950064 | 0.189140442 | 0.247513708 | 0.764161482 | 0.444771017 | 0.727526247 |
| B4GALT4 | 1154.400721 | -0.243851476 | 0.307336999 | -0.793433519 | 0.427525288 | 0.714914829 |
| B4GALT5 | 2186.343467 | -0.253888337 | 0.341228319 | -0.744042401 | 0.456850824 | 0.735735031 |
| B4GALT6 | 541.6614836 | 0.346596202 | 0.307802385 | 1.126034816 | 0.260150783 | 0.57337005 |
| B4GALT7 | 354.828603 | 0.120682586 | 0.370300716 | 0.325904274 | 0.744496782 | 0.892725315 |
| B4GAT1 | 788.0482121 | 0.927822619 | 0.288765077 | 3.213070738 | 0.001313239 | |

| | | | | | | |
|---|---|---|---|---|---|---|
| | 0.032290365 | | | | | |
| B9D1 | 312.8479208 | 0.783725698 | 0.361782449 | 2.166289989 | 0.030289034 | 0.198494574 |
| B9D2 | 63.94529127 | 0.699355179 | 0.562754144 | 1.242736614 | 0.213964907 | 0.520920862 |
| BAALC | 39.56137459 | 1.201263842 | 0.876558643 | 1.370431804 | 0.17055215 | 0.473631094 |
| BABAM1 | 2675.107054 | 0.189236559 | 0.361645184 | 0.523265806 | 0.600789293 | 0.824475293 |
| BABAM2 | 752.1835879 | -0.266263351 | 0.336436931 | -0.791421295 | 0.428698188 | 0.715644829 |
| BACE1 | 742.7269369 | -0.582577985 | 0.321535964 | -1.81185948 | 0.070007912 | 0.303975395 |
| BACE2 | 125.4262415 | 0.367266376 | 0.404928143 | 0.90699148 | 0.364411302 | 0.671176869 |
| BACH1 | 1714.147452 | -0.154054774 | 0.266640364 | -0.57776239 | 0.563424551 | 0.801924557 |
| BACH2 | 237.0972215 | -0.545350767 | 0.301401818 | -1.80938115 | 0.07039181 | 0.304770018 |
| BAD | 243.0010033 | 1.08173434 | 0.53696293 | 2.014541936 | 0.043952667 | 0.241112355 |
| BAG1 | 969.4546767 | 0.696350434 | 0.339340655 | 2.052068986 | 0.040162958 | 0.231238043 |
| BAG2 | 1000.999101 | 0.363054742 | 0.243546597 | 1.490699298 | 0.136040461 | 0.423792572 |
| BAG3 | 464.5638528 | -0.213835938 | 0.330398461 | -0.647206218 | 0.517498487 | 0.773950969 |
| BAG4 | 1095.898382 | 0.489146409 | 0.29498287 | 1.658219709 | 0.097273128 | 0.358274005 |
| BAG5 | 903.9635741 | -0.178636069 | 0.239006109 | -0.747412146 | 0.454814816 | 0.734745562 |
| BAG6 | 5015.797833 | -0.532232558 | 0.432053542 | -1.231867134 | 0.21799872 | 0.526298311 |
| BAGE2 | 30.89752575 | 0.008387123 | 0.758898284 | 0.01105171 | 0.991182191 | 0.996766871 |
| BAHCC1 | 805.7896072 | 0.139115509 | 0.395130619 | 0.352074738 | 0.724782213 | 0.883935898 |
| BAHD1 | 977.4216557 | 0.484449353 | 0.300883895 | 1.610087351 | 0.107378788 | 0.375442174 |
| BAIAP2 | 549.9456302 | -0.438879537 | 0.392751928 | -1.117447187 | 0.263803169 | 0.577444856 |
| BAIAP2L1 | 1296.769371 | -0.310137606 | 0.326396606 | -0.95018637 | 0.342017563 | 0.648704346 |
| BAIAP2L2 | 79.06539643 | 0.137869299 | 0.451199551 | 0.305561693 | 0.759938391 | 0.899725004 |
| BAIAP3 | 61.51510053 | 0.776028785 | 0.553104374 | 1.403042214 | 0.160604251 | 0.461026887 |
| BAK1 | 961.8951209 | 0.313997247 | 0.25756349 | 1.219106194 | 0.222803889 | 0.531805822 |
| BAMBI | 1340.51774 | -2.030470091 | 0.705712072 | -2.877193367 | 0.004012296 | 0.065575642 |
| BANF1 | 4903.020996 | 0.899138545 | 0.364584681 | 2.466199465 | 0.013655527 | 0.132885181 |
| BANK1 | 39.86143517 | -0.947273747 | 0.982059217 | -0.964579051 | 0.334755697 | 0.642776696 |
| BANP | 467.4087222 | -0.339150652 | 0.428121912 | -0.792182418 | 0.428254319 | 0.715316008 |
| BAP1 | 2623.722005 | -0.206000185 | 0.385367681 | -0.534554906 | 0.592957673 | 0.819859199 |
| BARD1 | 438.9349836 | -0.129540451 | 0.310283242 | -0.417490967 | 0.67631933 | 0.860199564 |
| BARX2 | 24.03156961 | -2.092510553 | 1.260508372 | -1.66005288 | 0.096903815 | 0.357434422 |
| BASP1 | 5690.00869 | 0.292343523 | 0.482460381 | 0.605943068 | 0.544552559 | 0.790739196 |
| BATF2 | 14.39141101 | 0.068184998 | 0.774822008 | 0.088000853 | 0.929875998 | 0.97351106 |
| BAX | 3745.304925 | 0.75443166 | 0.364880314 | 2.067614038 | 0.038676331 | 0.226576733 |
| BAZ1A | 1050.873797 | 0.298670116 | 0.243592408 | 1.226106013 | 0.220158783 | 0.528727493 |
| BAZ1B | 3214.637429 | 0.091853928 | 0.369316867 | 0.24871306 | 0.803582746 | 0.917969611 |
| BAZ2A | 5361.961374 | -0.671740473 | 0.340117127 | -1.975026896 | 0.048265096 | 0.253279062 |
| BAZ2B | 1339.293383 | -0.127274567 | 0.286080905 | -0.444890117 | 0.656399165 | 0.849955395 |
| BBC3 | 155.6364368 | 1.070785741 | 0.459354436 | 2.331066508 | 0.019749853 | 0.159121094 |
| BBIP1 | 547.2223965 | 0.9442422 | 0.296996253 | 3.179306777 | 0.001476278 | 0.0351044 |
| BBOF1 | 74.43445316 | -0.291323976 | 0.445743049 | -0.653569308 | 0.513389323 | 0.772068737 |

| | | | | | | |
|---|---|---|---|---|---|---|
| BBS1 | 799.2601391 | -0.626350267 | 0.423526742 | -1.478891897 | 0.139169209 | 0.427760708 |
| BBS10 | 550.3729803 | 0.26965346 | 0.356405569 | 0.756591602 | 0.449294574 | 0.730519016 |
| BBS12 | 177.4905126 | -0.160394376 | 0.381436323 | -0.420501055 | 0.674119459 | 0.859672768 |
| BBS2 | 1013.938279 | -0.213388879 | 0.234612263 | -0.909538471 | 0.36306596 | 0.670000448 |
| BBS4 | 521.6005031 | 0.547577597 | 0.3515274 | 1.557709578 | 0.119302109 | 0.398224928 |
| BBS5 | 139.5435157 | 0.120455235 | 0.383195093 | 0.314344408 | 0.753259473 | 0.896641779 |
| BBS7 | 709.6168663 | 0.53040658 | 0.247600233 | 2.142189342 | 0.032178249 | 0.204582474 |
| BBS9 | 2375.876555 | 0.232364199 | 0.392927391 | 0.591366763 | 0.554274705 | 0.796360966 |
| BBX | 2089.776267 | -0.173124207 | 0.236012167 | -0.73353933 | 0.463229549 | 0.740745005 |
| BCAM | 3856.737184 | -1.283991455 | 0.498663259 | -2.57486677 | 0.010027871 | 0.114042596 |
| BCAN | 16.57800939 | -0.601974533 | 0.926226265 | -0.649921682 | 0.515742812 | 0.773832307 |
| BCAP29 | 509.8084781 | 0.181098628 | 0.251436552 | 0.720255774 | 0.47136753 | 0.746147696 |
| BCAP31 | 2526.33581 | -0.172453325 | 0.327215229 | -0.527033308 | 0.598170458 | 0.822891188 |
| BCAR1 | 343.9022277 | -1.13643038 | 0.495196346 | -2.294908654 | 0.02173836 | 0.16740744 |
| BCAR3 | 502.9238125 | -0.039335599 | 0.283174086 | -0.138909599 | 0.889521586 | 0.956341292 |
| BCAS2 | 1524.491067 | 0.242898301 | 0.306119777 | 0.793474708 | 0.427501299 | 0.714914829 |
| BCAS3 | 720.1849384 | -0.76330466 | 0.301481458 | -2.53184612 | 0.011346377 | 0.120838052 |
| BCAS4 | 503.6745848 | 0.804383683 | 0.38427675 | 2.093240569 | 0.036327685 | 0.218181832 |
| BCAT1 | 5124.446011 | 0.588144208 | 0.350606003 | 1.677507524 | 0.093443279 | 0.350724392 |
| BCAT2 | 1060.589625 | 0.37099932 | 0.354287952 | 1.047168886 | 0.295021693 | 0.606966925 |
| BCCIP | 2745.498605 | 0.493291149 | 0.252166752 | 1.95621011 | 0.050440408 | 0.258046297 |
| BCDIN3D | 144.7765101 | 0.077639388 | 0.333447817 | 0.232838194 | 0.815887053 | 0.923166951 |
| BCHE | 42.7768164 | 0.951482362 | 0.667921056 | 1.424543144 | 0.154289305 | 0.451476933 |
| BCKDHA | 681.7481024 | -0.080887022 | 0.265363883 | -0.304815489 | 0.760506684 | 0.899902262 |
| BCKDHB | 955.1141423 | 0.147773138 | 0.270838424 | 0.545613639 | 0.585331547 | 0.815432957 |
| BCKDK | 987.9246294 | -0.352989812 | 0.276890347 | -1.27483611 | 0.202367257 | 0.509139909 |
| BCL10 | 753.7014051 | 0.061977015 | 0.371957613 | 0.166623866 | 0.867666014 | 0.946527727 |
| BCL11A | 69.52403763 | 0.810329087 | 0.567845222 | 1.427024576 | 0.153572811 | 0.450300177 |
| BCL11B | 69.32710202 | 0.545830856 | 0.469938319 | 1.161494678 | 0.245440786 | 0.557172253 |
| BCL2 | 127.0361582 | -1.386216889 | 0.881842295 | -1.571955549 | 0.115960861 | 0.39242521 |
| BCL2L1 | 2289.916926 | 0.095278319 | 0.218901858 | 0.435255871 | 0.663376713 | 0.854409382 |
| BCL2L11 | 1025.361966 | 0.027320451 | 0.436834046 | 0.062541946 | 0.950131259 | 0.981636196 |
| BCL2L12 | 1455.830811 | 0.664380289 | 0.384120706 | 1.729613318 | 0.083699385 | 0.330936505 |
| BCL2L13 | 1278.38134 | -0.202776267 | 0.253282489 | -0.800593315 | 0.423367122 | 0.712688899 |
| BCL2L15 | 15.6390055 | 0.543694229 | 0.773991135 | 0.70245537 | 0.482395225 | 0.753310328 |
| BCL2L2 | 850.7884458 | -0.271052791 | 0.282703766 | -0.958787337 | 0.33766589 | 0.645098755 |
| BCL2L2-PABPN1 | 54.56475753 | 0.635253075 | 0.916880029 | 0.692842089 | 0.488408656 | 0.757013457 |
| BCL3 | 24.30364507 | -0.03320352 | 0.783320751 | -0.042388153 | 0.966189273 | 0.988700292 |
| BCL6 | 594.5033413 | -0.288108438 | 0.362262694 | -0.795302532 | 0.426437541 | |

0.714361085

| | | | | | | |
|---|---|---|---|---|---|---|
| BCL6B | 82.60097583 | 1.59495715 | 0.584471332 | 2.728888592 | 0.006354817 | 0.086487511 |
| BCL7A | 359.5476693 | 0.261754512 | 0.485962884 | 0.538630666 | 0.590141724 | 0.818321084 |
| BCL7B | 692.0660014 | -0.246317491 | 0.294236298 | -0.837141755 | 0.402512892 | 0.699487181 |
| BCL7C | 613.5414643 | 0.295769159 | 0.452967747 | 0.652958539 | 0.513783008 | 0.772584202 |
| BCL9 | 2759.780988 | -0.488749418 | 0.397364167 | -1.229978592 | 0.218705122 | 0.527143548 |
| BCL9L | 1840.215049 | -1.109029064 | 0.42226595 | -2.626375778 | 0.008629949 | 0.104839578 |
| BCLAF1 | 5684.99676 | -0.343306759 | 0.309160051 | -1.110449934 | 0.266805191 | 0.579983028 |
| BCLAF3 | 351.8151302 | -0.365539217 | 0.274258401 | -1.332827786 | 0.182588325 | 0.486683209 |
| BCO2 | 17.14784072 | 0.406860178 | 0.774748688 | 0.525151168 | 0.599478108 | 0.823816451 |
| BCOR | 590.0952018 | -0.361724082 | 0.444439832 | -0.813887631 | 0.415709327 | 0.707609161 |
| BCORL1 | 1161.584453 | -0.266654564 | 0.407623694 | -0.65416846 | 0.513003279 | 0.771819984 |
| BCORP1 | 77.71214999 | -0.341403973 | 0.47369236 | -0.720729321 | 0.471076069 | 0.74600157 |
| BCR | 3466.408399 | -0.209952662 | 0.397276899 | -0.528479413 | 0.597166632 | 0.821867951 |
| BCRP2 | 79.89040448 | 1.026968554 | 0.412011398 | 2.492573162 | 0.01268212 | 0.127358401 |
| BCS1L | 545.6919331 | 0.082255784 | 0.271377831 | 0.303104286 | 0.761810388 | 0.900742374 |
| BDH1 | 213.4784254 | -0.010948276 | 0.290226111 | -0.037723262 | 0.969908329 | 0.989496332 |
| BDH2 | 1532.201352 | 0.093514576 | 0.304729864 | 0.306876966 | 0.758937022 | 0.899440455 |
| BDKRB2 | 29.93733803 | -0.826067159 | 0.642150935 | -1.286406536 | 0.198301219 | 0.50414535 |
| BDNF | 57.97263154 | -0.155289336 | 0.447616292 | -0.346925121 | 0.728647574 | 0.885833187 |
| BDP1 | 728.408158 | -0.115262205 | 0.565065934 | -0.203980099 | 0.838369051 | 0.933879959 |
| BEAN1 | 11.04206348 | -0.83706387 | 1.037289192 | -0.806972516 | 0.419682318 | 0.710229302 |
| BECN1 | 1463.74807 | -0.002831269 | 0.328767739 | -0.008611759 | 0.993128895 | 0.997283174 |
| BEND3 | 494.3879377 | -0.100127702 | 0.264902391 | -0.377979609 | 0.705445742 | 0.873945986 |
| BEND4 | 346.0161491 | -0.201688297 | 0.36431326 | -0.553612287 | 0.579844215 | 0.811340629 |
| BEND5 | 193.580265 | 0.61400958 | 0.347280664 | 1.768050003 | 0.077052545 | 0.31844659 |
| BEND6 | 115.9745353 | -0.399885701 | 0.471630796 | -0.847878689 | 0.396505534 | 0.69525953 |
| BEND7 | 88.28501526 | 0.099313401 | 0.44455712 | 0.223398517 | 0.823225358 | 0.926014081 |
| BEST1 | 88.18875652 | 0.938245861 | 0.680252973 | 1.379260214 | 0.167814539 | 0.470246466 |
| BEST4 | 53.86609541 | 0.696272053 | 0.623198901 | 1.117254945 | 0.263885334 | 0.577444856 |
| BET1 | 804.8095074 | 0.739500216 | 0.32857202 | 2.250648783 | 0.024407791 | 0.178492903 |
| BET1L | 1417.362924 | -0.290629363 | 0.228487712 | -1.271969337 | 0.203384014 | 0.509735688 |
| BEX1 | 412.4644174 | 1.066763732 | 0.3835017 | 2.781640165 | 0.005408497 | 0.07827792 |
| BEX2 | 153.9212589 | 0.552044732 | 0.522457583 | 1.056630719 | 0.290680155 | 0.60368818 |
| BEX3 | 5220.612604 | 0.702197435 | 0.474036892 | 1.481313893 | 0.138522948 | 0.426880285 |
| BEX4 | 1477.461762 | 0.139573954 | 0.297381969 | 0.469342354 | 0.638824945 | 0.842822267 |
| BEX5 | 64.02476214 | 1.115602532 | 0.532006068 | 2.096973324 | 0.035995937 | 0.217048632 |
| BFAR | 1845.206864 | 0.108873001 | 0.224107951 | 0.485806062 | 0.627104676 | 0.836041947 |
| BFSP1 | 120.2969427 | -0.050177345 | 0.55821151 | -0.089889484 | 0.928375038 | 0.972633911 |
| BGLAP | 24.29079594 | 2.170298319 | 1.157300303 | 1.875311286 | 0.060749912 | 0.284105089 |
| BGN | 82.78458109 | 0.309498992 | 0.535953566 | 0.57747352 | 0.563619622 | 0.801976485 |
| BHLHB9 | 395.9332873 | 0.170691745 | 0.33858974 | 0.504125569 | 0.61417314 | 0.830316752 |
| BHLHE22 | 22.03172005 | -0.38710606 | 0.940736588 | -0.41149251 | 0.680711432 | |

| | | | | | | |
|---|---|---|---|---|---|---|
| | | | | | | 0.862305386 |
| BHLHE40 | 79.32049005 | -4.067200572 | 1.012733641 | -4.016061489 | 5.92E-05 | 0.004234915 |
| BHLHE41 | 20.49172268 | 0.178571946 | 1.267808533 | 0.140850879 | 0.887987748 | 0.955943827 |
| BICC1 | 30.86866338 | -2.153848987 | 0.828537181 | -2.599580364 | 0.009333782 | 0.109346547 |
| BICD1 | 663.1752959 | -0.33860359 | 0.291249824 | -1.162588136 | 0.244996647 | 0.556497099 |
| BICD2 | 932.131504 | -0.173026172 | 0.376160016 | -0.459980234 | 0.645530408 | 0.845794147 |
| BICDL1 | 194.5824329 | -1.364380706 | 0.373067054 | -3.657199666 | 0.000254986 | 0.01105254 |
| BICDL2 | 165.758608 | -2.261328162 | 0.815922051 | -2.771500244 | 0.005579863 | 0.079935885 |
| BICRA | 386.6607542 | -0.182848236 | 0.336672969 | -0.543103405 | 0.587058608 | 0.81660769 |
| BICRAL | 356.8444556 | -0.49293699 | 0.329106446 | -1.497804119 | 0.13418415 | 0.420619758 |
| BID | 444.3475305 | 1.015596352 | 0.42590203 | 2.384577393 | 0.017098751 | 0.14773715 |
| BIK | 128.8324371 | 0.304627519 | 0.774664693 | 0.393237902 | 0.694143778 | 0.86892666 |
| BIN1 | 1822.673743 | -0.51039281 | 0.284833347 | -1.791899777 | 0.073149022 | 0.311055732 |
| BIN2 | 29.10856211 | -0.364999967 | 0.701496131 | -0.520316437 | 0.602843041 | 0.825021776 |
| BIN3 | 638.1011728 | -0.012354366 | 0.319129739 | -0.038712674 | 0.969119469 | 0.989496332 |
| BIRC2 | 1876.331237 | 0.006827686 | 0.262621428 | 0.025998204 | 0.979258771 | 0.992858175 |
| BIRC3 | 13.38848691 | -0.417046904 | 0.834075406 | -0.500011031 | 0.61706731 | 0.831835425 |
| BIRC5 | 2858.391095 | 1.296222887 | 0.272568365 | 4.75558815 | 1.98E-06 | 0.000306095 |
| BIRC6 | 1856.184155 | -0.507381861 | 0.420840624 | -1.205638981 | 0.227956722 | 0.536924612 |
| BIVM | 396.055657 | -0.010097853 | 0.280025998 | -0.036060412 | 0.971234189 | 0.989629517 |
| BLACAT1 | 315.5921914 | -0.305893058 | 0.349120676 | -0.876181443 | 0.380931403 | 0.682545212 |
| BLCAP | 1405.815088 | 0.461386312 | 0.370540038 | 1.245172626 | 0.213068302 | 0.520189768 |
| BLM | 938.6071092 | 0.180888882 | 0.406109933 | 0.445418512 | 0.656017337 | 0.849909395 |
| BLMH | 2982.238284 | -0.202577608 | 0.280179805 | -0.723027158 | 0.4696632 | 0.744930517 |
| BLOC1S1 | 966.1206878 | 0.918811684 | 0.423750432 | 2.168284948 | 0.030137013 | 0.198185614 |
| BLOC1S2 | 1718.413215 | 0.851723644 | 0.379353531 | 2.245197617 | 0.024755449 | 0.17998149 |
| BLOC1S3 | 139.3248712 | -0.209860957 | 0.446730007 | -0.469771347 | 0.638518388 | 0.842491082 |
| BLOC1S4 | 432.3303366 | 1.067190828 | 0.512921213 | 2.080613554 | 0.037469292 | 0.221788225 |
| BLOC1S5 | 707.4369482 | 0.158912321 | 0.255427037 | 0.622143695 | 0.533847386 | 0.783156734 |
| BLOC1S5-TXNDC5 | 289.1839288 | -0.487693184 | 2.329666934 | -0.209340304 | 0.834182591 | 0.932107564 |
| BLOC1S6 | 2159.988412 | -0.168784797 | 0.232340282 | -0.726455161 | 0.467559786 | 0.744005613 |
| BLVRA | 410.0379679 | 1.035844971 | 0.465258473 | 2.226386042 | 0.025988335 | 0.184130533 |
| BLVRB | 438.5008474 | 0.96380293 | 0.390526297 | 2.467959101 | 0.013588584 | 0.132501135 |
| BLZF1 | 478.0385984 | -0.148753563 | 0.362175099 | -0.41072278 | 0.681275821 | 0.862490397 |
| BMF | 2437.480114 | -0.908962046 | 0.304750002 | -2.982648203 | 0.002857663 | 0.052557154 |
| BMI1 | 853.2145203 | 0.498800378 | 0.356674739 | 1.398474081 | 0.161970751 | 0.462586269 |
| BMP1 | 200.1966823 | 0.384051332 | 0.543700661 | 0.706365394 | 0.479960936 | 0.752036291 |
| BMP2 | 152.3095047 | 0.792055822 | 0.563908132 | 1.404583084 | 0.16014529 | 0.460493592 |

| | | | | | | |
|---|---|---|---|---|---|---|
| BMP2K | 692.0957667 | -0.010366233 | 0.331510657 | -0.031269683 | 0.975054468 | 0.99132451 |
| BMP3 | 24.66218539 | 0.341160622 | 0.875099343 | 0.389853592 | 0.696644812 | 0.870023227 |
| BMP4 | 797.946916 | -0.199069338 | 0.404704574 | -0.491888035 | 0.622798497 | 0.833787151 |
| BMP5 | 805.6199149 | 0.066273774 | 0.55401768 | 0.119623932 | 0.904781061 | 0.964070338 |
| BMP6 | 15.75103496 | 1.34626272 | 0.918911549 | 1.465062357 | 0.142903906 | 0.433233718 |
| BMP7 | 882.2385327 | -0.279325716 | 0.490361541 | -0.569632184 | 0.568927195 | 0.805450586 |
| BMP8A | 18.9513594 | -0.468844176 | 0.744703754 | -0.629571387 | 0.528975049 | 0.779891202 |
| BMPER | 198.103098 | -1.381916808 | 0.431156867 | -3.205136952 | 0.001349982 | 0.033086562 |
| BMPR1A | 1689.013066 | -0.181164537 | 0.234998706 | -0.770917169 | 0.44075603 | 0.724218535 |
| BMPR1B | 34.86073562 | 0.984819592 | 0.738360682 | 1.333792029 | 0.182272024 | 0.486415681 |
| BMPR2 | 656.7369567 | -0.678168027 | 0.35101183 | -1.932037523 | 0.053354875 | 0.267056023 |
| BMS1 | 2198.105971 | -0.290620269 | 0.302590184 | -0.960441824 | 0.336832897 | 0.644459616 |
| BMS1P1 | 344.9350271 | -0.501718115 | 0.50824996 | -0.987148361 | 0.323569913 | 0.633361998 |
| BMS1P10 | 35.13579253 | 0.007964799 | 0.559902726 | 0.014225326 | 0.988650215 | 0.995803391 |
| BMS1P2 | 102.277879 | -0.467393683 | 0.823126756 | -0.567827105 | 0.570152373 | 0.806205765 |
| BMS1P4 | 100.8421879 | -0.409287602 | 0.407328099 | -1.004810626 | 0.314988046 | 0.624501261 |
| BMT2 | 391.1693736 | -0.514835714 | 0.36735592 | -1.401462955 | 0.161075678 | 0.46167973 |
| BNC1 | 319.0478708 | -5.912353234 | 1.066064237 | -5.545963393 | 2.92E-08 | 1.71E-05 |
| BNC2 | 178.5309926 | -2.653089448 | 0.838076698 | -3.165688123 | 0.001547166 | 0.036390779 |
| BNIP1 | 258.3327666 | 0.586738471 | 0.312577607 | 1.877096946 | 0.060504808 | 0.284105089 |
| BNIP2 | 1562.837644 | -0.035051688 | 0.287006443 | -0.122128574 | 0.902797192 | 0.963038687 |
| BNIP3 | 1533.124576 | 0.66119544 | 0.319441573 | 2.069847808 | 0.038466599 | 0.225742345 |
| BNIP3L | 2128.173343 | 0.347399062 | 0.244484202 | 1.42094687 | 0.155332206 | 0.453268877 |
| BNIPL | 1550.855096 | -0.880459328 | 0.710139154 | -1.239840562 | 0.215034372 | 0.522216497 |
| BOC | 2125.806108 | 0.880571727 | 0.566535338 | 1.554310326 | 0.120110412 | 0.400130666 |
| BOD1 | 2870.586165 | 0.628176801 | 0.278300991 | 2.257184921 | 0.023996522 | 0.177207433 |
| BOD1L1 | 2301.366751 | -0.365512599 | 0.269944263 | -1.354029883 | 0.175726844 | 0.479919343 |
| BOK | 480.9906737 | 0.085169556 | 0.336882238 | 0.252816999 | 0.800409631 | 0.916453926 |
| BOLA1 | 159.1463711 | 0.704551434 | 0.515474395 | 1.366801999 | 0.171687376 | 0.474879827 |
| BOLA2 | 69.27175882 | -0.285617329 | 0.5344786 | -0.534384968 | 0.593075217 | 0.819859199 |
| BOLA2-SMG1P6 | 1146.366433 | 0.960561416 | 0.376759472 | 2.549534881 | 0.010786672 | 0.117984572 |
| BOLA2B | 623.5509055 | 1.069043242 | 0.436584217 | 2.448652977 | 0.014339152 | 0.135962051 |
| BOLA3 | 465.5697541 | 1.286495385 | 0.416834599 | 3.086345013 | 0.002026336 | 0.042460685 |
| BOP1 | 463.2147765 | 0.795500318 | 0.436784309 | 1.821265789 | 0.068566464 | 0.301671663 |
| BORA | 742.3460054 | 0.27473832 | 0.324905759 | 0.845593877 | 0.397779343 | 0.696366306 |
| BORCS5 | 105.1816 | 0.141704859 | 0.39398923 | 0.359666835 | 0.719096296 | 0.881260702 |
| BORCS6 | 102.1332545 | 0.076170432 | 0.493138652 | 0.154460478 | 0.877246671 | 0.950622089 |
| BORCS7 | 332.0746304 | 0.454170552 | 0.390577126 | 1.162819126 | 0.244902895 | 0.556367449 |

| | | | | | | |
|---|---|---|---|---|---|---|
| BORCS8 | 330.6602349 | -0.287148075 | 0.39410132 | -0.728614852 | 0.466237291 | 0.742914184 |
| BORCS8-MEF2B | 50.01251817 | -0.285914561 | 0.955485476 | -0.299234858 | 0.764760854 | 0.902893924 |
| BPGM | 540.6897051 | 0.298615371 | 0.263714738 | 1.132342369 | 0.257490524 | 0.570028755 |
| BPHL | 922.0121869 | 0.092888503 | 0.234619039 | 0.395912043 | 0.692169916 | 0.868131192 |
| BPNT1 | 497.457491 | 0.474726956 | 0.335089688 | 1.416716102 | 0.156565947 | 0.455379983 |
| BPTF | 4717.214671 | -0.279245244 | 0.296414412 | -0.942077149 | 0.346153143 | 0.652845515 |
| BPTFP1 | 18.75748873 | -1.757193115 | 0.895701027 | -1.961807638 | 0.049784884 | 0.257162573 |
| BRAF | 995.1260353 | -0.329587206 | 0.28049486 | -1.175020481 | 0.239986522 | 0.551618176 |
| BRAP | 775.8073122 | -0.487145989 | 0.252652748 | -1.928124637 | 0.053839625 | 0.268155271 |
| BRAT1 | 782.0727365 | 0.194850442 | 0.471065666 | 0.413637538 | 0.679139578 | 0.861617874 |
| BRCA1 | 1037.964982 | -0.009710826 | 0.410753397 | -0.023641498 | 0.981138571 | 0.993241851 |
| BRCA2 | 490.160532 | 0.016788609 | 0.562299069 | 0.029857081 | 0.976181035 | 0.991761418 |
| BRCC3 | 732.5721225 | -0.457322322 | 0.311736469 | -1.467015792 | 0.142371759 | 0.432590018 |
| BRD1 | 1045.338995 | -0.211265911 | 0.366903064 | -0.57580852 | 0.564744614 | 0.803200575 |
| BRD2 | 702.573929 | -0.334169981 | 1.406329117 | -0.237618617 | 0.812176909 | 0.921390083 |
| BRD3 | 2134.669569 | -0.083095944 | 0.263524026 | -0.315325873 | 0.752514243 | 0.896586461 |
| BRD3OS | 465.4178609 | 0.028190291 | 0.355596167 | 0.079276139 | 0.936812985 | 0.976155095 |
| BRD4 | 2193.15677 | -0.273708021 | 0.311438177 | -0.878851859 | 0.379481604 | 0.681253679 |
| BRD7 | 4199.627395 | 0.179849422 | 0.218363958 | 0.823622284 | 0.41015421 | 0.704109399 |
| BRD8 | 2446.792271 | -0.120715125 | 0.339995629 | -0.355049049 | 0.722552846 | 0.882935039 |
| BRD9 | 1424.85233 | -0.034930504 | 0.250504492 | -0.139440627 | 0.88910197 | 0.956228979 |
| BRF1 | 446.1604224 | -0.204658397 | 0.445337078 | -0.459558404 | 0.645833221 | 0.845889304 |
| BRF2 | 426.1548754 | -0.153031766 | 0.260976093 | -0.586382317 | 0.557618624 | 0.798455135 |
| BRI3 | 1816.785954 | 0.784293558 | 0.475013795 | 1.65109638 | 0.098718896 | 0.360796043 |
| BRI3BP | 318.4700817 | 0.427545142 | 0.370964048 | 1.152524468 | 0.249105621 | 0.561876506 |
| BRICD5 | 75.14474143 | 0.331758007 | 0.571332285 | 0.580674358 | 0.561459947 | 0.801534662 |
| BRINP1 | 28.92281175 | -1.668164981 | 0.805695978 | -2.070464575 | 0.03840886 | 0.225590715 |
| BRIP1 | 737.8488772 | 0.328581609 | 0.450153171 | 0.729932898 | 0.465431202 | 0.742335301 |
| BRIX1 | 1513.30799 | 0.607262676 | 0.257603631 | 2.357352938 | 0.018405744 | 0.153593808 |
| BRK1 | 3313.768466 | 0.316558367 | 0.376059313 | 0.841777762 | 0.399912362 | 0.697652919 |
| BRMS1 | 809.4827813 | -0.074063028 | 0.292494263 | -0.253211899 | 0.800104472 | 0.916311991 |
| BRMS1L | 548.1061843 | 0.33314014 | 0.263311896 | 1.265192135 | 0.205802461 | 0.512178669 |
| BROX | 1182.007818 | -0.238209991 | 0.306023279 | -0.778404807 | 0.436330406 | 0.720784523 |
| BRPF1 | 784.223935 | -0.244928144 | 0.276460133 | -0.88594381 | 0.375647807 | 0.67966045 |
| BRPF3 | 1729.98953 | -0.832604104 | 0.32292603 | -2.57831214 | 0.009928427 | 0.11342181 |
| BRSK1 | 814.6215632 | -0.147928168 | 0.469885172 | -0.314817697 | 0.752900074 | 0.896586461 |
| BRSK2 | 73.76699642 | 1.491046018 | 0.820921059 | 1.816308648 | 0.069323039 | 0.30336211 |
| BRWD1 | 1813.833256 | -0.040525122 | 0.26998626 | -0.150100682 | 0.880685182 | |

| | | | | | | |
|---|---|---|---|---|---|---|
| | | | | | | 0.952579132 |
| BRWD3 | 2035.885206 | -0.536521066 | 0.27591194 | -1.94453733 | 0.051830688 | 0.262457735 |
| BSCL2 | 1048.581851 | 0.885366519 | 0.494294964 | 1.791170419 | 0.073265952 | 0.311262322 |
| BSDC1 | 1320.751429 | -0.627993887 | 0.285848678 | -2.196945222 | 0.028024359 | 0.191340029 |
| BSG | 12861.31822 | 0.361900088 | 0.376674084 | 0.960777773 | 0.336663917 | 0.644459616 |
| BSN | 30.14720317 | -0.209693455 | 1.229353888 | -0.170572084 | 0.86456025 | 0.945178609 |
| BSPRY | 295.6162172 | -1.49986893 | 0.830476172 | -1.806034876 | 0.070912892 | 0.30641398 |
| BST1 | 94.66505263 | -0.271268144 | 0.524349368 | -0.517342368 | 0.604917183 | 0.82602102 |
| BST2 | 1822.364578 | 1.169174717 | 0.704595231 | 1.659356558 | 0.097043965 | 0.357864363 |
| BTAF1 | 2679.29586 | -0.00636939 | 0.569501103 | -0.011184158 | 0.991076519 | 0.996726674 |
| BTBD1 | 1448.066939 | 0.653923645 | 0.2742199 | 2.384668824 | 0.017094502 | 0.14773715 |
| BTBD10 | 888.3011791 | 0.090738668 | 0.333155202 | 0.272361552 | 0.785344032 | 0.910931322 |
| BTBD11 | 41.8407529 | 0.808954207 | 0.784230402 | 1.031526201 | 0.302294128 | 0.613115536 |
| BTBD17 | 21.69302277 | 2.471985959 | 1.101295783 | 2.244615839 | 0.024792805 | 0.180028929 |
| BTBD19 | 44.31780817 | -0.046668551 | 0.504180186 | -0.092563239 | 0.92625055 | 0.972261514 |
| BTBD2 | 2212.2483 | -0.226178065 | 0.567714129 | -0.398401332 | 0.690334376 | 0.867938158 |
| BTBD3 | 4793.905823 | -0.079658385 | 0.291604857 | -0.273172351 | 0.784720732 | 0.910931322 |
| BTBD6 | 375.5052498 | -0.519427211 | 0.368263764 | -1.410476027 | 0.15839917 | 0.457902784 |
| BTBD7 | 362.4066844 | -0.300325005 | 0.36349085 | -0.826224388 | 0.408676816 | 0.703031634 |
| BTBD8 | 119.9589746 | -0.12955491 | 0.391961882 | -0.33052936 | 0.741000012 | 0.891943909 |
| BTBD9 | 209.5870247 | -0.513607832 | 0.314244926 | -1.634418853 | 0.102170932 | 0.366611924 |
| BTC | 14.27285617 | 0.155420213 | 0.960067847 | 0.161884614 | 0.871396719 | 0.948581876 |
| BTD | 456.1074932 | -0.146617675 | 0.330116608 | -0.444139043 | 0.65694206 | 0.85024732 |
| BTF3 | 18582.39453 | 0.75574046 | 0.42344689 | 1.78473494 | 0.074304321 | 0.313236304 |
| BTF3L4 | 3232.921855 | 0.527791105 | 0.301656298 | 1.749643911 | 0.080179778 | 0.324202401 |
| BTF3L4P2 | 329.7447507 | 0.299243457 | 0.348104837 | 0.859636022 | 0.389989712 | 0.689813912 |
| BTG1 | 1455.268498 | 0.429629562 | 0.411538142 | 1.043960494 | 0.296503668 | 0.607790454 |
| BTG2 | 2026.110206 | 0.296810399 | 0.410460156 | 0.723116226 | 0.469608482 | 0.744930517 |
| BTG3 | 1058.31714 | 1.355463776 | 0.269661768 | 5.026533004 | 4.99E-07 | 0.000120267 |
| BTK | 37.65063583 | 0.795617398 | 0.545576571 | 1.458305653 | 0.144756313 | 0.435928729 |
| BTN2A1 | 485.0816299 | -0.147858302 | 0.393499404 | -0.37575229 | 0.70710106 | 0.875135606 |
| BTN2A2 | 397.3793407 | -0.327404579 | 0.414811126 | -0.789285914 | 0.429944918 | 0.716554273 |
| BTN2A3P | 85.93168229 | -0.303418075 | 0.486093326 | -0.624197163 | 0.53249811 | 0.782597698 |
| BTN3A1 | 110.3072051 | 0.22009293 | 0.690989345 | 0.318518558 | 0.75009162 | 0.895752516 |
| BTN3A2 | 153.7030532 | 0.578914105 | 0.444816828 | 1.301466286 | 0.193098897 | 0.498908723 |
| BTN3A3 | 18.00360834 | 0.9578451 | 0.879435363 | 1.089159182 | 0.276083696 | 0.589426646 |
| BTRC | 988.6440477 | -0.456921947 | 0.359574075 | -1.270731064 | 0.20382434 | 0.509986087 |
| BUB1 | 3578.409451 | 0.789059018 | 0.395581811 | 1.994679725 | 0.046077822 | 0.246950947 |
| BUB1B | 2722.348465 | 0.754943713 | 0.429494388 | 1.757749889 | 0.078790075 | 0.3223636 |

| | | | | | | |
|---|---|---|---|---|---|---|
| BUB3 | 3207.966373 | 0.769180789 | 0.261223149 | 2.944535324 | 0.0032344 | 0.057190077 |
| BUD13 | 885.559604 | 0.104667312 | 0.269740934 | 0.388029028 | 0.697994552 | 0.870467 |
| BUD23 | 1337.817687 | 0.460607362 | 0.367910162 | 1.251956073 | 0.210585871 | 0.518294063 |
| BUD31 | 1370.582339 | 0.611022196 | 0.39056614 | 1.564452555 | 0.117711327 | 0.395700983 |
| BVES | 103.8889825 | 0.303089128 | 0.401988572 | 0.753974488 | 0.450864541 | 0.732184323 |
| BX255925.3 | 274.9017888 | -0.044581606 | 0.521256411 | -0.085527209 | 0.931842265 | 0.974359294 |
| BX322639.1 | 392.7395192 | 0.460867275 | 0.376323951 | 1.224655711 | 0.220704968 | 0.529377876 |
| BX571818.1 | 18.74173399 | -0.997698851 | 0.990556365 | -1.007210581 | 0.31383359 | 0.623434514 |
| BX664615.1 | 12.19932556 | 0.269135467 | 0.818749118 | 0.328715429 | 0.742370791 | 0.892150449 |
| BX679664.3 | 17.16331905 | -0.034596558 | 0.760860106 | -0.045470328 | 0.963732426 | 0.988130579 |
| BYSL | 855.0882875 | 0.381877624 | 0.260157739 | 1.467869552 | 0.14213966 | 0.432308365 |
| BZW1 | 8005.757725 | 0.229910272 | 0.21365291 | 1.076092398 | 0.281885935 | 0.595704168 |
| BZW1P2 | 180.1315119 | 0.038173739 | 2.265527117 | 0.016849827 | 0.98655642 | 0.995107011 |
| BZW2 | 3466.448814 | -0.302279705 | 0.407346253 | -0.742070665 | 0.458044522 | 0.736518606 |
| C10orf105 | 38.20404761 | 0.853079453 | 0.738799542 | 1.154683246 | 0.248220163 | 0.56079644 |
| C10orf143 | 61.49902865 | -0.383384698 | 0.509942308 | -0.75181975 | 0.452159463 | 0.732888936 |
| C10orf82 | 451.2039279 | -0.922877566 | 0.523784933 | -1.761939889 | 0.078079454 | 0.320406654 |
| C10orf88 | 362.7344506 | 0.324751608 | 0.302286132 | 1.074318582 | 0.28267992 | 0.596532022 |
| C11orf1 | 593.3065502 | 0.342181339 | 0.457947109 | 0.747207118 | 0.454938548 | 0.734867197 |
| C11orf24 | 526.0210388 | 0.17151674 | 0.263170171 | 0.651733206 | 0.514573297 | 0.77285333 |
| C11orf45 | 10.3463397 | 0.056252417 | 1.07227636 | 0.052460745 | 0.958161573 | 0.985242593 |
| C11orf49 | 953.6666199 | 0.261465817 | 0.288524779 | 0.906216162 | 0.36482145 | 0.671614642 |
| C11orf52 | 189.0940374 | -1.212858128 | 0.809554335 | -1.498180018 | 0.134086486 | 0.420619758 |
| C11orf54 | 809.765958 | -0.073127176 | 0.264602474 | -0.276366186 | 0.782266826 | 0.909909835 |
| C11orf58 | 6287.67881 | 0.382704536 | 0.258131895 | 1.482592984 | 0.138182582 | 0.426306273 |
| C11orf65 | 27.64977597 | 0.112347795 | 0.589242642 | 0.19066474 | 0.848788266 | 0.938569494 |
| C11orf68 | 494.9846941 | 0.137688253 | 0.327349263 | 0.420615742 | 0.674035697 | 0.859672768 |
| C11orf71 | 214.4741951 | 0.287024591 | 0.419102273 | 0.684855726 | 0.493434967 | 0.759455148 |
| C11orf74 | 750.3626882 | 0.845733498 | 0.387380631 | 2.183210592 | 0.029020305 | 0.194746536 |
| C11orf80 | 336.1942636 | -0.682907987 | 0.26832946 | -2.545035448 | 0.010926665 | 0.119001025 |
| C11orf87 | 23.29121465 | -0.03091315 | 1.108428707 | -0.027889164 | 0.977750551 | 0.99226352 |
| C11orf95 | 1487.749774 | -0.164809237 | 0.314923256 | -0.523331427 | 0.600743635 | 0.824475293 |
| C11orf96 | 334.15519 | 0.999735062 | 0.677004845 | 1.476702966 | 0.139755277 | 0.428675153 |
| C11orf98 | 1499.564866 | 0.969600392 | 0.469896565 | 2.063433668 | 0.039071444 | 0.22780664 |
| C12orf10 | 1122.63352 | 0.051391082 | 0.291935424 | 0.176035787 | 0.860265827 | 0.943475231 |
| C12orf29 | 525.3361555 | 0.22709532 | 0.311328914 | 0.729438579 | 0.465733426 | |

0.742657327

| | | | | | | |
|---|---|---|---|---|---|---|
| C12orf4 | 533.787664 | -0.07853261 | 0.324960886 | -0.241667884 | 0.809037517 | 0.919669426 |
| C12orf43 | 558.9648729 | 0.294788127 | 0.281507693 | 1.047176098 | 0.295018367 | 0.606966925 |
| C12orf45 | 739.1311124 | 0.468596052 | 0.389882964 | 1.201889016 | 0.229406531 | 0.538167075 |
| C12orf49 | 1955.706745 | -0.279362407 | 0.264540655 | -1.056028261 | 0.290955304 | 0.604006967 |
| C12orf50 | 154.3953915 | -0.21811277 | 0.563651182 | -0.386964096 | 0.698782788 | 0.870873025 |
| C12orf56 | 42.94083159 | 0.505496539 | 0.492713499 | 1.025944164 | 0.304917904 | 0.615135352 |
| C12orf57 | 1989.952565 | 1.164719629 | 0.426824171 | 2.728804291 | 0.006356442 | 0.086487511 |
| C12orf60 | 119.3537132 | 0.393777682 | 0.391955356 | 1.004649318 | 0.315065741 | 0.624573677 |
| C12orf65 | 395.7315285 | 0.503430609 | 0.317819845 | 1.584012508 | 0.113190873 | 0.387109723 |
| C12orf66 | 175.7779637 | 0.294981986 | 0.391770269 | 0.75294633 | 0.451482166 | 0.732638351 |
| C12orf73 | 300.010589 | 1.246726144 | 0.4094162 | 3.045131444 | 0.002325786 | 0.046063323 |
| C12orf75 | 346.9371034 | 1.065701126 | 0.303471897 | 3.511696255 | 0.000445257 | 0.016477615 |
| C12orf76 | 462.4190861 | 0.629718777 | 0.365843615 | 1.721278576 | 0.085200285 | 0.334358075 |
| C13orf42 | 62.35813837 | 0.270232164 | 0.470175627 | 0.574747283 | 0.565462224 | 0.803844397 |
| C14orf119 | 1622.121879 | 0.207204762 | 0.364282701 | 0.56880209 | 0.569490456 | 0.805870694 |
| C14orf132 | 537.0402591 | 0.534908432 | 0.488925281 | 1.094049444 | 0.273933293 | 0.586847211 |
| C14orf28 | 162.2554453 | 0.220788223 | 0.358892658 | 0.615192922 | 0.538427321 | 0.786560766 |
| C14orf39 | 44.43841243 | 0.144501376 | 0.611195047 | 0.236424324 | 0.813103423 | 0.921802034 |
| C14orf93 | 378.4488245 | -0.327408239 | 0.262717137 | -1.246238609 | 0.212676809 | 0.519735803 |
| C15orf39 | 449.5408866 | -1.865463258 | 0.591487278 | -3.153851869 | 0.001611309 | 0.037175747 |
| C15orf40 | 522.9414681 | 0.257826118 | 0.285970527 | 0.901582832 | 0.367278514 | 0.673921424 |
| C15orf41 | 636.7009927 | -0.487850573 | 0.35701662 | -1.3664646 | 0.171793185 | 0.474879827 |
| C15orf53 | 21.38002353 | -1.107502931 | 0.923966514 | -1.198639685 | 0.23066808 | 0.539536045 |
| C15orf61 | 69.19684508 | 1.002744944 | 0.68992853 | 1.453404084 | 0.146111593 | 0.438508206 |
| C15orf65 | 54.77690864 | 0.893225973 | 0.522245765 | 1.710355608 | 0.087200135 | 0.339208524 |
| C16orf45 | 415.523643 | 1.17630852 | 0.59121853 | 1.989634054 | 0.046631262 | 0.248574447 |
| C16orf46 | 35.5274645 | 1.066877585 | 0.604395694 | 1.765197196 | 0.07753063 | 0.319277194 |
| C16orf58 | 1356.382448 | -0.362717623 | 0.285510968 | -1.270415722 | 0.203936586 | 0.510127278 |
| C16orf70 | 675.2193469 | -0.188375823 | 0.294695704 | -0.639221477 | 0.522678863 | 0.777662198 |
| C16orf72 | 3213.148668 | -0.116995417 | 0.268419165 | -0.435868344 | 0.662932254 | 0.854171175 |
| C16orf74 | 71.53501453 | -0.071745133 | 0.775310695 | -0.092537267 | 0.926271184 | 0.972261514 |
| C16orf86 | 47.98527035 | 0.903831145 | 0.629857114 | 1.434978069 | 0.151293347 | |

0.446637745

| | | | | | | |
|---|---|---|---|---|---|---|
| C16orf87 | 1138.43308 | 0.46712377 | 0.350729685 | 1.331862656 | 0.182905324 | 0.486816961 |
| C16orf89 | 17.199239 | 2.187037722 | 0.89294934 | 2.44922934 | 0.014316226 | 0.13582956 |
| C16orf91 | 203.6420941 | 1.171680821 | 0.473541086 | 2.474296012 | 0.013349903 | 0.131319811 |
| C16orf95 | 53.0490508 | 0.84465946 | 0.522655781 | 1.616091298 | 0.106074575 | 0.373724427 |
| C16orf96 | 28.01985079 | 0.468846178 | 0.848204987 | 0.552751028 | 0.580433907 | 0.811740671 |
| C17orf100 | 49.15398248 | 1.152922119 | 0.508637856 | 2.266685628 | 0.023409437 | 0.174689901 |
| C17orf107 | 59.009849 | -0.606170033 | 0.513037569 | -1.181531472 | 0.237391659 | 0.547879744 |
| C17orf113 | 120.5137113 | -0.688290935 | 0.484487719 | -1.420657136 | 0.155416459 | 0.453320705 |
| C17orf49 | 1783.276543 | 1.06814647 | 0.421838097 | 2.532124237 | 0.011337381 | 0.120838052 |
| C17orf51 | 218.8242662 | 0.31360142 | 0.516565295 | 0.607089604 | 0.543791449 | 0.790216482 |
| C17orf53 | 293.5199838 | -0.247525107 | 0.33542919 | -0.737935499 | 0.460553646 | 0.738395128 |
| C17orf58 | 402.7778234 | 0.650199372 | 0.286265528 | 2.271315642 | 0.023127877 | 0.173699516 |
| C17orf67 | 19.32754403 | -0.685910401 | 0.904779456 | -0.75809679 | 0.448393041 | 0.729796208 |
| C17orf75 | 1123.439427 | 0.728950802 | 0.284768766 | 2.559799 | 0.010473272 | 0.116232631 |
| C17orf80 | 1047.979315 | -0.173532746 | 0.280208067 | -0.619299609 | 0.535719002 | 0.784724629 |
| C17orf97 | 52.39667651 | 0.344011659 | 0.520092767 | 0.661442882 | 0.508328333 | 0.768801629 |
| C18orf21 | 620.7706635 | 0.685950359 | 0.40307074 | 1.701811345 | 0.088790739 | 0.342301733 |
| C18orf25 | 721.2909307 | -0.183121487 | 0.30807931 | -0.594397226 | 0.552246463 | 0.795643979 |
| C18orf32 | 774.9710877 | 1.214638642 | 0.493245543 | 2.462543572 | 0.013795541 | 0.133647607 |
| C18orf54 | 1197.835419 | 0.216812707 | 0.329151598 | 0.658701669 | 0.510087361 | 0.769949736 |
| C19orf12 | 648.3998193 | 0.050684311 | 0.31638599 | 0.160197709 | 0.872725334 | 0.949330774 |
| C19orf18 | 19.40585387 | 0.203592575 | 0.888208519 | 0.229217093 | 0.818700188 | 0.924316427 |
| C19orf24 | 222.3997226 | 0.763964568 | 0.572223469 | 1.335080802 | 0.181849902 | 0.48591777 |
| C19orf25 | 360.8747321 | 0.318294142 | 0.442580481 | 0.719177993 | 0.472031257 | 0.746399031 |
| C19orf33 | 26.53125559 | -0.617696751 | 0.924933868 | -0.667828016 | 0.504243381 | 0.765917338 |
| C19orf44 | 229.3684483 | -0.180120203 | 0.383306871 | -0.469911231 | 0.63841844 | 0.842491082 |
| C19orf47 | 239.6856877 | -0.32448256 | 0.369996052 | -0.876989248 | 0.380492478 | 0.682161591 |
| C19orf48 | 2049.991136 | 0.23919297 | 0.293897498 | 0.813865283 | 0.415722131 | 0.707609161 |
| C19orf53 | 1651.442487 | 0.596961191 | 0.424711704 | 1.405568027 | 0.159852436 | 0.460017489 |
| C19orf54 | 887.5686058 | -0.410796866 | 0.256391751 | -1.602223412 | 0.109106215 | 0.379241044 |
| C19orf57 | 24.67946032 | 0.297172626 | 0.744630675 | 0.399087274 | 0.689828899 | 0.867683896 |
| C19orf66 | 19.79374165 | 1.206860147 | 0.82391831 | 1.464781317 | 0.142980591 | |

0.433233718
C19orf84 68.46141464 0.178326316 0.664433113 0.268388664 0.788400164 0.912309638
C1D 900.1452185 1.022216205 0.437105831 2.338601165 0.019356084 0.157622736
C1DP1 36.70777509 0.411063989 0.507733712 0.809605467 0.418166965 0.709589334
C1GALT1 1795.165052 -0.402792399 0.408963633 -0.984910065 0.324668249 0.634653009
C1GALT1C1 705.3961804 0.494537411 0.356058844 1.388920453 0.16485694 0.467115865
C1orf105 33.53265361 -3.272764424 1.568751501 -2.086222338 0.036958488 0.220226715
C1orf109 583.0770324 0.253310597 0.263479765 0.961404366 0.33634889 0.64425371
C1orf112 646.8150396 -0.16823529 0.320276955 -0.525280659 0.599388101 0.823816451
C1orf115 695.3368825 -1.025437964 0.300630719 -3.410955368 0.000647357 0.020763322
C1orf116 45.60667166 -1.076173335 0.571552496 -1.882894998 0.059714599 0.282669434
C1orf122 376.1439645 0.999836025 0.510790073 1.957430415 0.050296885 0.257701469
C1orf131 543.3642234 0.307780145 0.349405898 0.880867057 0.37838978 0.681253679
C1orf159 203.8191763 -0.227495122 0.607116041 -0.374714398 0.707872883 0.875585221
C1orf162 11.64940876 0.814708629 0.933131774 0.873090652 0.382613669 0.683824871
C1orf167 22.24992597 -0.502293292 0.743821362 -0.675287532 0.499493104 0.76281557
C1orf174 604.8398125 0.080520214 0.250092822 0.321961317 0.747482 0.894202292
C1orf198 1837.493638 -0.171869783 0.259386427 -0.662601297 0.507585939 0.768444894
C1orf21 936.2779894 0.367553499 0.299183362 1.228522522 0.21925088 0.52778634
C1orf210 182.3956107 -1.075550233 0.793439122 -1.35555483 0.175240852 0.479195921
C1orf216 196.3935331 0.326111504 0.335446362 0.972171829 0.330965073 0.639084337
C1orf226 256.7563819 -0.604151422 0.448253062 -1.347790953 0.177725628 0.482042363
C1orf35 455.3751218 0.829873166 0.549073867 1.511405324 0.130685214 0.416052367
C1orf43 4680.490361 0.496759753 0.25220228 1.969687796 0.048874163 0.254713129
C1orf50 209.1329258 -0.235452644 0.304808002 -0.77246215 0.439840756 0.723341367
C1orf52 1131.950717 0.399605759 0.3839099 1.040884225 0.297929284 0.609375237
C1orf54 485.2944679 1.324011098 0.418478061 3.163872189 0.001556851 0.0365054
C1orf56 54.01874175 0.316860697 0.507848783 0.623927255 0.53267536 0.782684541
C1orf61 66.20331337 -2.118395877 0.861426565 -2.459171752 0.013925798 0.134161613
C1orf74 278.8146574 0.018742491 0.273393663 0.068554958 0.945343872 0.979754792
C1QBP 4432.411904 0.858911644 0.306246863 2.804638177 0.005037309 0.075664373
C1QTNF1 148.8816189 -0.25636854 0.533180525 -0.480828778 0.630638195 0.838192723
C1QTNF12 13.40123923 0.950151867 0.929206164 1.022541502 0.30652468 0.61666701
C1QTNF2 11.14593063 0.954155869 0.986925885 0.966795869 0.33364609 0.641713975
C1QTNF3 71.13356628 -0.376293091 0.528519862 -0.711975307

0.476480067 0.74962969
C1QTNF3-AMACR 54.42733506 -1.346465961 1.118399917 -1.203921728
0.228619833 0.53738497
C1QTNF5 13.69833955 2.261177201 1.009706575 2.23943991 0.025127307
0.181440445
C1QTNF6 517.4120168 0.14369874 0.318710857 0.450874945 0.652079682
0.848139649
C1R 171.2421718 0.803342063 0.482995356 1.663250075 0.096262388 0.356020645
C1RL 900.5696275 -0.858908456 0.27928883 -3.075341235 0.002102618
0.043389195
C1S 48.21886972 0.255166985 0.598167829 0.426580924 0.669684595 0.857762448
C2 15.07863898 -1.135860132 1.284553375 -0.884245181 0.376563873
0.68026322
C20orf194 526.0827743 -0.268202107 0.435404892 -0.615983218
0.537905596 0.786181676
C20orf27 1087.056615 0.957807826 0.375664045 2.549639335 0.010783441
0.117984572
C20orf96 685.3599698 0.154689097 0.272957943 0.566714036 0.570908483
0.806748565
C21orf58 309.4312054 0.481464759 0.470099081 1.024177197 0.305751587
0.615683213
C21orf59-TCP10L 261.437987 -0.805832804 0.815959711 -0.987588962
0.323353994 0.63322621
C21orf91 1121.472604 0.497725098 0.401319735 1.240220837 0.214893723
0.522125669
C22orf15 12.20791431 1.107777618 0.951266437 1.164529278 0.244209585
0.555457814
C22orf23 10.91436074 0.129395317 0.941085131 0.137495868 0.890638859
0.957017706
C22orf39 720.2681428 0.429754184 0.283912453 1.513685572 0.130105606
0.415181944
C22orf46 680.9146235 -0.88604672 0.392565508 -2.257067171 0.024003878
0.177207433
C2CD2 181.7063869 0.386555803 0.468993128 0.824224877 0.409811795 0.703966274
C2CD2L 102.3803458 0.491126687 0.391982691 1.252929524 0.210231356
0.51816316
C2CD3 895.4439336 -0.088036731 0.284479125 -0.309466401 0.756966767
0.898563152
C2CD4C 77.46305481 1.144535582 0.479797174 2.385457111 0.017057911
0.14773715
C2CD5 1430.515209 0.075039227 0.292635102 0.256425926 0.797621971 0.915916373
C2orf15 136.0117065 -0.911297999 0.596852271 -1.526840131
0.12680077 0.410063292
C2orf27A 11.76782895 0.754112047 0.945039994 0.797968394 0.424888831
0.71350603
C2orf42 198.2840701 -0.116990135 0.319569623 -0.366086531
0.714300508 0.878457604
C2orf49 821.6909681 0.042331629 0.252534594 0.167627048 0.866876691
0.946142897
C2orf68 1363.44452 0.491786257 0.348498837 1.411156095 0.158198593
0.457584529
C2orf69 730.0597942 0.318258655 0.275610304 1.154741497 0.248196301
0.56079644
C2orf72 62.07019548 -2.469907624 0.99121826 -2.491789876
0.012710121 0.127363842
C2orf74 470.3745705 0.508830259 0.370804439 1.372233463 0.169990771
0.472591166
C2orf76 192.7708262 1.121074678 0.504052823 2.224121414 0.026140276
0.184538918
C2orf80 16.12609604 2.446807125 1.289624588 1.897301856 0.05778811
0.278583864
C2orf81 72.36284435 0.829535374 0.603852033 1.373739474 0.169522577
0.472289317
C2orf91 20.31508144 -1.717751207 0.802235751 -2.141205008

| | | | | | | |
|---|---|---|---|---|---|---|
| | | | | | 0.032257509 | 0.204675314 |
| C2orf92 | 81.33968728 | -1.281992748 | 0.474137871 | -2.703839593 | 0.006854337 | 0.090803834 |
| C3 | 40.92354735 | -0.459520487 | 0.674982975 | -0.680788262 | 0.496005478 | 0.761056737 |
| C3orf14 | 326.9220969 | 0.432936808 | 0.357727371 | 1.210242332 | 0.226185919 | 0.534746233 |
| C3orf18 | 169.9130659 | 0.539743846 | 0.491393669 | 1.098393977 | 0.272032491 | 0.584230595 |
| C3orf33 | 164.1568483 | 0.581177258 | 0.438886075 | 1.324209837 | 0.185433365 | 0.490031772 |
| C3orf38 | 932.8946021 | 0.414788807 | 0.272551881 | 1.521871015 | 0.128041403 | 0.412255816 |
| C3orf52 | 72.44664543 | 0.136818766 | 0.459010669 | 0.298073171 | 0.765647318 | 0.903026935 |
| C3orf62 | 103.7330101 | -0.143148429 | 0.425670604 | -0.336289205 | 0.73665279 | 0.890498763 |
| C3orf67 | 396.2735984 | 0.041711743 | 0.320006313 | 0.130346626 | 0.896292192 | 0.959013488 |
| C3orf70 | 45.08359335 | -0.783433615 | 1.147862116 | -0.682515438 | 0.494913085 | 0.760514211 |
| C4A | 21.50490892 | 1.549201393 | 0.793454885 | 1.952475713 | 0.050881752 | 0.259366654 |
| C4B | 38.38372085 | 0.97111494 | 0.632443835 | 1.535495939 | 0.124662054 | 0.406173955 |
| C4orf19 | 69.1024529 | -0.202820753 | 0.746233931 | -0.271792456 | 0.785781606 | 0.910984215 |
| C4orf3 | 2278.266921 | 0.633551895 | 0.522144955 | 1.213364006 | 0.224990681 | 0.534418918 |
| C4orf33 | 428.169269 | -0.23804322 | 0.356015404 | -0.668631797 | 0.503730384 | 0.765510733 |
| C4orf36 | 29.58914474 | 0.997566068 | 0.559161036 | 1.784040738 | 0.074417046 | 0.313367219 |
| C4orf46 | 1137.835193 | 0.655417356 | 0.299757207 | 2.186494073 | 0.028779478 | 0.194136709 |
| C4orf47 | 51.82093729 | 0.854430939 | 0.475660936 | 1.796302523 | 0.072446417 | 0.309775814 |
| C4orf48 | 28.84454813 | 1.530525642 | 1.011636464 | 1.512920597 | 0.130299829 | 0.41546421 |
| C5 | 89.54486006 | 0.109732493 | 0.559149229 | 0.196249029 | 0.844415255 | 0.936997062 |
| C5AR2 | 10.57208135 | 0.05864404 | 0.919009527 | 0.063812222 | 0.949119747 | 0.98134156 |
| C5orf15 | 1623.9967 | 0.34612686 | 0.285589007 | 1.211975431 | 0.225521786 | 0.53459368 |
| C5orf22 | 1262.728201 | 0.1875687 | 0.251723857 | 0.745136765 | 0.456189046 | 0.735399427 |
| C5orf24 | 1327.753481 | 0.344997356 | 0.29112923 | 1.185031667 | 0.236004927 | 0.546881128 |
| C5orf30 | 1157.623716 | -0.419268068 | 0.512193878 | -0.818572977 | 0.41303009 | 0.705607651 |
| C5orf34 | 206.0442476 | 0.491780728 | 0.33162633 | 1.482936315 | 0.138091332 | 0.426306273 |
| C5orf47 | 12.35248298 | 1.066701775 | 0.95267226 | 1.119694379 | 0.262844021 | 0.576453244 |
| C5orf51 | 1434.662902 | -0.394516487 | 0.280086554 | -1.408552041 | 0.158967668 | 0.458846746 |
| C5orf63 | 322.7214072 | 1.218759156 | 0.438032121 | 2.782351103 | 0.005396662 | 0.078197481 |
| C6orf106 | 2442.524422 | -0.689922891 | 0.276354854 | -2.496510849 | 0.012542183 | 0.12669427 |
| C6orf118 | 39.53911284 | 2.484348458 | 1.049951843 | 2.366154672 | 0.017973934 | 0.151238239 |
| C6orf120 | 1042.646306 | 0.120634855 | 0.250426841 | 0.481716954 | 0.63000703 | 0.838039163 |
| C6orf132 | 356.194939 | -1.266192226 | 0.454899398 | -2.783455485 | 0.005378324 | 0.078006269 |
| C6orf136 | 65.65467469 | 0.665393146 | 0.684533108 | 0.972039392 | 0.331030952 | |

| | | | | | | |
|---|---|---|---|---|---|---|
| | | | | | | 0.639084337 |
| C6orf141 | 285.295646 | -1.180619589 | 0.631147013 | -1.870593641 | 0.061401427 | 0.285640507 |
| C6orf163 | 88.96491112 | -0.456136848 | 0.397880793 | -1.14641585 | 0.251623127 | 0.564383659 |
| C6orf203 | 298.4629029 | 0.821819405 | 0.39165171 | 2.098342439 | 0.035874906 | 0.216490931 |
| C6orf226 | 32.9351227 | 1.228506642 | 0.704906792 | 1.742793028 | 0.081369775 | 0.326231726 |
| C6orf48 | 604.2629599 | 0.748707921 | 0.539313904 | 1.38826 | 0.165057888 | 0.46718157 |
| C6orf62 | 2985.486462 | 0.124487452 | 0.404446792 | 0.307796857 | 0.758236913 | 0.89938619 |
| C6orf89 | 993.226045 | -0.333954544 | 0.364027805 | -0.917387461 | 0.358939641 | 0.666738847 |
| C7orf25 | 38.44242179 | -0.08873589 | 0.531137878 | -0.167067524 | 0.867316919 | 0.946404214 |
| C7orf26 | 591.7594003 | -0.082862949 | 0.281823679 | -0.294024083 | 0.768739501 | 0.90442396 |
| C7orf31 | 146.1194722 | -0.735807801 | 0.462700389 | -1.590246775 | 0.111779191 | 0.384888109 |
| C7orf43 | 319.0607136 | 0.098395035 | 0.338452021 | 0.290720778 | 0.771264879 | 0.905067213 |
| C7orf50 | 632.6296379 | 0.861883561 | 0.462131417 | 1.865018326 | 0.062178835 | 0.287461403 |
| C7orf61 | 14.97102708 | -0.018665658 | 0.913215873 | -0.02043948 | 0.98369279 | 0.994045382 |
| C8orf33 | 1655.296943 | 0.169124549 | 0.250519815 | 0.675094498 | 0.499615729 | 0.762849343 |
| C8orf34 | 25.72219292 | 0.355266581 | 0.972448286 | 0.365332107 | 0.714863514 | 0.87855355 |
| C8orf37 | 115.4853699 | 0.050352341 | 0.39435759 | 0.127681938 | 0.898400684 | 0.960048173 |
| C8orf44 | 98.90270529 | 0.565520384 | 0.529531382 | 1.067963869 | 0.28553681 | 0.599411008 |
| C8orf48 | 76.32788362 | -0.086261466 | 0.488196402 | -0.176694187 | 0.859748607 | 0.943248924 |
| C8orf58 | 486.1828982 | -0.187536314 | 0.36851717 | -0.508894373 | 0.610826264 | 0.829058038 |
| C8orf59 | 1713.107047 | 0.819081598 | 0.423566774 | 1.93377207 | 0.053141158 | 0.266955137 |
| C8orf76 | 486.607726 | 0.552152251 | 0.322385661 | 1.712707229 | 0.086766414 | 0.33809963 |
| C8orf82 | 169.3788344 | 0.382894354 | 0.45406964 | 0.843250287 | 0.399088479 | 0.697352242 |
| C8orf88 | 218.0968809 | 0.815709859 | 0.380792251 | 2.142138809 | 0.032182314 | 0.204582474 |
| C9orf116 | 58.05066584 | 0.604607879 | 0.436651249 | 1.384647084 | 0.166160407 | 0.468031848 |
| C9orf135 | 38.85985061 | -2.893931316 | 0.862902503 | -3.353717607 | 0.000797337 | 0.023672012 |
| C9orf16 | 435.723374 | 0.479455536 | 0.510427839 | 0.939320898 | 0.347566012 | 0.654127772 |
| C9orf3 | 337.1637082 | -0.19194569 | 0.315074233 | -0.609207831 | 0.54238669 | 0.789157809 |
| C9orf40 | 412.5574519 | 1.400769903 | 0.380031041 | 3.685935497 | 0.000227864 | 0.010319181 |
| C9orf66 | 17.92927549 | -0.813419989 | 0.760827844 | -1.069124896 | 0.285013395 | 0.598807847 |
| C9orf72 | 911.8698317 | 0.778026934 | 0.360367697 | 2.158980789 | 0.030851657 | 0.200535773 |
| C9orf78 | 1385.526802 | 0.159941321 | 0.318772455 | 0.501741346 | 0.615849477 | 0.831232421 |
| C9orf85 | 388.5884431 | 0.258723708 | 0.401349276 | 0.644634794 | 0.519163881 | |

0.774631448
CA11 1324.327584 0.672597932 0.35657862 1.886254235 0.059260703 0.282340574
CA12 80.84886182 -1.957088703 0.810198501 -2.415566928 0.015710735
0.141958046
CA13 91.50920801 -1.11087214 0.468275289 -2.372262994 0.017679502
0.150008791
CA14 1178.838365 1.005325272 0.360346036 2.78988853 0.005272619 0.077250742
CA2 408.4441218 0.032985852 0.342051296 0.096435395 0.923174782 0.971120333
CA3 23.08392303 -0.181195968 0.675620545 -0.268191916 0.788551598
0.912309638
CA4 125.1984029 0.836077606 0.650786473 1.284718782 0.198890572 0.504782074
CA5A 12.57910416 0.254610519 1.052712371 0.24186143 0.808887537 0.919641737
CA5B 445.323813 -1.675924739 0.62530201 -2.680184474 0.00735816
0.095085725
CA5BP1 248.9384383 -0.435971457 0.350896309 -1.242450965
0.214070222 0.520920862
CA8 151.0474688 0.612725571 0.574222489 1.067052549 0.285948108 0.599871951
CA9 13.86362736 1.700730985 1.025225837 1.658884242 0.097139121 0.357954238
CAAP1 757.7267173 0.833093417 0.296810235 2.806821726 0.005003292 0.07561376
CAB39 1322.310972 0.008513369 0.241063115 0.035315934 0.971827818 0.989769054
CAB39L 638.8000253 -0.45944626 0.282911385 -1.623993535 0.104377188
0.370237626
CABIN1 1677.999118 -0.481489346 0.551023351 -0.873809332
0.382222097 0.683628312
CABLES1 184.124633 0.219893254 0.385527886 0.570369256 0.568427277
0.805450586
CABLES2 413.9018784 -0.001406731 0.32842954 -0.004283203
0.996582509 0.998431718
CABP1 15.84054707 0.298153741 0.838030642 0.355779044 0.722006044 0.882710428
CABYR 238.1784819 0.128897098 0.359192505 0.358852416 0.719705498 0.881674105
CACFD1 385.8203161 0.353716831 0.426455134 0.829435038 0.406858282
0.701626421
CACHD1 2544.377382 0.436573035 0.47198728 0.924967798 0.354982658
0.662415978
CACNA1A 247.3040425 0.785608115 0.427037614 1.839669596 0.06581676
0.29619856
CACNA1B 49.65992435 0.281034908 0.801220249 0.350758618 0.725769445
0.884318388
CACNA1C 33.10644336 -1.12870613 0.607075048 -1.859253042 0.06299128
0.289778852
CACNA1D 26.73216257 -0.947825838 0.832040568 -1.139158202
0.254637175 0.567256913
CACNA1E 17.38808074 0.773398184 0.751056291 1.029747295 0.303128648
0.61357768
CACNA1F 48.77582009 -0.578383553 0.588989216 -0.981993452
0.326103074 0.635960913
CACNA1G 473.9654705 -0.83092072 0.561715157 -1.479256363 0.139071811
0.427760708
CACNA1H 679.6941625 0.64043116 0.654564147 0.978408553 0.3278723
0.637775693
CACNA1I 53.52008291 -0.544707973 0.462618769 -1.177444604
0.239018096 0.550165913
CACNA1S 15.72766149 -0.796082636 0.930007743 -0.855995707
0.392000154 0.691496659
CACNA2D1 368.3764361 -0.359058173 0.459771635 -0.780948945
0.434832524 0.719786603
CACNA2D2 638.6603817 -1.032731735 0.50897942 -2.029024542
0.042455791 0.237061759
CACNA2D3 167.773665 -0.261798989 0.384905275 -0.680164721
0.496400168 0.761167958
CACNA2D4 14.68564022 -0.616384111 0.892546121 -0.690590768
0.489822751 0.75819824
CACNB1 166.7281303 0.296086044 0.508589007 0.582171537 0.560451143
0.800546493

| | | | | | | |
|---|---|---|---|---|---|---|
| CACNB2 | 57.14089612 | -0.393665099 | 0.696331451 | -0.565341546 | 0.571841475 | 0.807351345 |
| CACNB3 | 1270.610874 | -0.543947852 | 0.260004279 | -2.092072693 | 0.036432013 | 0.218635315 |
| CACNB4 | 102.291104 | 0.58803241 | 0.488759181 | 1.203112766 | 0.228932686 | 0.537637427 |
| CACNG4 | 307.7783028 | -1.30673231 | 0.46425675 | -2.814675954 | 0.004882644 | 0.074671968 |
| CACNG6 | 143.9665323 | -1.4237438 | 0.815159312 | -1.746583495 | 0.080709612 | 0.325104967 |
| CACNG7 | 250.826697 | 0.789950575 | 0.554092315 | 1.425666001 | 0.153964775 | 0.450926563 |
| CACNG8 | 156.703691 | -1.262492142 | 0.523747252 | -2.41049884 | 0.015930722 | 0.142746012 |
| CACTIN | 607.6490152 | -0.09918052 | 0.438349873 | -0.226258808 | 0.821000134 | 0.925158936 |
| CACUL1 | 1556.035064 | 0.022203295 | 0.297332606 | 0.074674941 | 0.940473346 | 0.97807441 |
| CACYBP | 3037.207935 | 0.85358666 | 0.301097989 | 2.834913185 | 0.004583814 | 0.071324146 |
| CAD | 3039.969995 | 0.039520398 | 0.481663794 | 0.082049758 | 0.934607145 | 0.975099718 |
| CADM1 | 2849.745789 | 0.075622639 | 0.2854197 | 0.264952417 | 0.791046118 | 0.913361935 |
| CADM2 | 46.03697247 | 1.284020512 | 0.599621089 | 2.14138651 | 0.032242882 | 0.204668101 |
| CADM3 | 242.54097 | 0.473496388 | 0.352084692 | 1.344836623 | 0.178678001 | 0.483022799 |
| CADM4 | 559.4177981 | 0.549661985 | 0.243303516 | 2.259161701 | 0.023873328 | 0.176933201 |
| CADPS | 153.6278582 | -0.907004421 | 0.495071809 | -1.83206639 | 0.06694153 | 0.297817842 |
| CADPS2 | 129.8169378 | -0.159107351 | 0.450996294 | -0.352790817 | 0.724245269 | 0.883667763 |
| CAGE1 | 11.5606156 | -0.860248361 | 0.983927159 | -0.87430086 | 0.381954431 | 0.683329089 |
| CALB1 | 212.8532936 | -0.819812583 | 0.680433006 | -1.204839529 | 0.228265257 | 0.537150955 |
| CALB2 | 49.85549725 | 1.892158369 | 0.514641663 | 3.676652133 | 0.000236315 | 0.010544512 |
| CALCB | 42.76552517 | -0.307527356 | 0.751159914 | -0.409403311 | 0.68224371 | 0.862735748 |
| CALCOCO1 | 1182.576473 | -1.231120353 | 0.410107351 | -3.001946564 | 0.002682593 | 0.050458917 |
| CALCOCO2 | 977.5072971 | 0.237576698 | 0.259052675 | 0.917098032 | 0.359091272 | 0.666802166 |
| CALCRL | 42.93814851 | 0.151521567 | 0.654933444 | 0.231354145 | 0.817039686 | 0.923735281 |
| CALD1 | 4457.246405 | -0.228686769 | 0.293280562 | -0.779754265 | 0.435535531 | 0.720542613 |
| CALHM2 | 44.23629975 | -0.952399007 | 0.60889251 | -1.56414965 | 0.117782428 | 0.395839103 |
| CALHM5 | 40.87807305 | 1.653020231 | 0.651774098 | 2.536185831 | 0.011206725 | 0.120297968 |
| CALM1 | 7348.199278 | 0.304612254 | 0.376568494 | 0.808915931 | 0.418563505 | 0.709738117 |
| CALM2 | 12635.98844 | 1.188844916 | 0.217999539 | 5.453428578 | 4.94E-08 | 2.53E-05 |
| CALM3 | 11101.87103 | 0.096466107 | 0.319623198 | 0.301811971 | 0.762795405 | 0.901485479 |
| CALML4 | 129.1488275 | 1.99856043 | 0.472098446 | 4.233355243 | 2.30E-05 | 0.002054603 |
| CALML6 | 13.21799665 | -0.152695476 | 0.849326827 | -0.179784119 | 0.857322052 | 0.942343063 |
| CALN1 | 15.04167912 | -0.082757027 | 0.895610322 | -0.09240294 | 0.926377904 | 0.972261514 |
| CALR | 15258.0517 | 0.037384893 | 0.246052013 | 0.151938984 | 0.879235061 | 0.951892893 |
| CALU | 7733.689555 | 0.067680105 | 0.265834814 | 0.25459459 | 0.799036238 | 0.916056735 |
| CAMK1 | 164.8633063 | 1.225592476 | 0.40486978 | 3.027127578 | 0.002468897 | 0.047998079 |
| CAMK1D | 164.9494451 | 0.442979289 | 0.535725859 | 0.82687681 | 0.408306889 | 0.702713988 |
| CAMK1G | 267.5346978 | -2.49939288 | 0.758662485 | -3.294472744 | 0.000986065 | 0.026954222 |

| | | | | | | |
|---|---|---|---|---|---|---|
| CAMK2A | 23.5959619 | 0.122006535 | 0.682064887 | 0.178878194 | 0.858033345 | 0.942343063 |
| CAMK2D | 1119.778422 | -0.508735513 | 0.353839088 | -1.437759508 | 0.150502302 | 0.445342388 |
| CAMK2G | 1880.006046 | -0.30542923 | 0.24703395 | -1.236385646 | 0.216315249 | 0.523985094 |
| CAMK2N1 | 461.850743 | 0.750253251 | 0.320577648 | 2.340316786 | 0.019267389 | 0.157577123 |
| CAMK2N2 | 28.49234913 | 1.314864446 | 0.657745829 | 1.999046422 | 0.045603331 | 0.245771986 |
| CAMK4 | 38.1905476 | -0.203738058 | 0.531201947 | -0.383541626 | 0.701318213 | 0.872388416 |
| CAMKK1 | 171.8940245 | 0.910029558 | 0.489215427 | 1.860181645 | 0.062859831 | 0.289510169 |
| CAMKK2 | 1090.938645 | -0.037134135 | 0.398358478 | -0.093217885 | 0.925730466 | 0.972124891 |
| CAMKMT | 464.8884579 | 0.464055908 | 0.47260401 | 0.98191276 | 0.326142829 | 0.635960913 |
| CAMKV | 261.4631603 | 0.742964192 | 0.536572308 | 1.384648781 | 0.166159888 | 0.468031848 |
| CAMLG | 1849.315022 | 0.952926167 | 0.368332829 | 2.587133411 | 0.009677811 | 0.111651769 |
| CAMSAP1 | 1629.348336 | -0.034998786 | 0.35089709 | -0.099740883 | 0.920550042 | 0.969843271 |
| CAMSAP2 | 1178.885371 | -0.00906631 | 0.413649667 | -0.021917847 | 0.982513488 | 0.993518428 |
| CAMSAP3 | 482.3822489 | -0.782205266 | 0.423884159 | -1.845327905 | 0.064989854 | 0.293878113 |
| CAMTA1 | 1496.425975 | 1.030409225 | 0.438465455 | 2.350035135 | 0.018771639 | 0.1554129 |
| CAMTA2 | 500.8334102 | -1.971407265 | 0.421065699 | -4.681946944 | 2.84E-06 | 0.000420026 |
| CAND1 | 6109.350194 | 0.283071816 | 0.339537191 | 0.833698999 | 0.404450623 | 0.700367583 |
| CAND2 | 226.4080448 | 0.309572646 | 0.522864321 | 0.592070703 | 0.553803244 | 0.796301122 |
| CANT1 | 1761.507237 | -0.21264263 | 0.288070433 | -0.738161941 | 0.460416047 | 0.738395128 |
| CANX | 12825.16864 | -0.395179329 | 0.244134275 | -1.618696631 | 0.105512552 | 0.372436233 |
| CAP1 | 6791.887112 | -0.277709691 | 0.219128855 | -1.267335104 | 0.205035493 | 0.51158559 |
| CAP2 | 1001.01001 | 0.142921242 | 0.267244403 | 0.534796017 | 0.592790918 | 0.819859199 |
| CAPG | 1011.509081 | -0.500797028 | 0.55746752 | -0.898342972 | 0.36900273 | 0.674858445 |
| CAPN1 | 3574.05486 | -1.293443573 | 0.30941327 | -4.180310605 | 2.91E-05 | 0.002413359 |
| CAPN10 | 612.8340566 | 0.005497289 | 0.466456638 | 0.01178521 | 0.990596981 | 0.996726674 |
| CAPN11 | 10.29355639 | 0.662940951 | 1.020054535 | 0.649907362 | 0.515752063 | 0.773832307 |
| CAPN12 | 129.1428511 | -0.649094288 | 0.52373583 | -1.239354368 | 0.215214293 | 0.522486163 |
| CAPN13 | 72.78869049 | -1.350692148 | 0.558820071 | -2.417043013 | 0.015647168 | 0.141467929 |
| CAPN15 | 331.4842955 | 0.171762729 | 0.655091415 | 0.26219658 | 0.79316989 | 0.914167013 |
| CAPN2 | 3419.575574 | -0.845272718 | 0.376656259 | -2.244148868 | 0.024822825 | 0.18009903 |
| CAPN3 | 43.81377984 | -1.145830047 | 0.702571928 | -1.630907813 | 0.102909773 | 0.367870915 |
| CAPN5 | 717.8870756 | -0.697510556 | 0.279294836 | -2.497398685 | 0.012510821 | 0.12669427 |
| CAPN6 | 267.9391171 | -1.877958087 | 0.309019891 | -6.077143065 | 1.22E-09 | 1.55E-06 |
| CAPN7 | 994.4677173 | 0.161149307 | 0.247551909 | 0.65097178 | 0.515064704 | 0.773208651 |
| CAPN9 | 24.24939516 | -1.271387063 | 0.723748519 | -1.756669657 | 0.078974132 | 0.322507823 |
| CAPNS1 | 5262.953283 | -0.27345719 | 0.318839333 | -0.85766454 | 0.391077726 | |

| | | | | | | |
|---|---|---|---|---|---|---|
| | | | | | | 0.690530747 |
| CAPRIN1 | 10499.14223 | -0.064290388 | 0.258728251 | -0.248486156 | 0.80375828 | 0.917969611 |
| CAPRIN2 | 1041.888456 | -0.165324811 | 0.343690351 | -0.481028376 | 0.630496332 | 0.838177265 |
| CAPS | 218.4471217 | -0.912189303 | 0.423919856 | -2.151796595 | 0.031413378 | 0.202023044 |
| CAPS2 | 209.4694394 | 0.037934996 | 0.411103677 | 0.092275984 | 0.926478769 | 0.972261514 |
| CAPZA1 | 4901.5141 | 0.150777721 | 0.270534742 | 0.557332193 | 0.577300484 | 0.809901794 |
| CAPZA2 | 2343.374709 | 0.582790077 | 0.242726836 | 2.401012127 | 0.016349795 | 0.144378309 |
| CAPZB | 5787.873248 | -0.034759982 | 0.312909723 | -0.111086294 | 0.911547917 | 0.966866396 |
| CARD10 | 123.2871838 | -0.94630543 | 0.430209186 | -2.199640221 | 0.027832431 | 0.190853383 |
| CARD11 | 103.6451428 | -1.536206 | 0.638354472 | -2.406509342 | 0.016105792 | 0.143654707 |
| CARD14 | 110.5538927 | -0.691210666 | 0.42559588 | -1.624100934 | 0.104354269 | 0.370237626 |
| CARD19 | 145.8839239 | 1.064114276 | 0.349889843 | 3.041283701 | 0.002355718 | 0.046595294 |
| CARD6 | 11.2146086 | -1.97467869 | 1.116724891 | -1.768276777 | 0.077014644 | 0.31844659 |
| CARD8 | 839.6051158 | 0.116686032 | 0.248052259 | 0.470409069 | 0.638062786 | 0.842182924 |
| CARD9 | 11.16088295 | 0.312288347 | 0.872994507 | 0.357720862 | 0.720552219 | 0.88214152 |
| CARF | 394.2997599 | -0.288757941 | 0.296780385 | -0.972968416 | 0.330568999 | 0.638942832 |
| CARHSP1 | 4811.990442 | 0.453584256 | 0.374420824 | 1.211429032 | 0.225731018 | 0.53459368 |
| CARM1 | 3239.235648 | -0.487638366 | 0.335282507 | -1.454410404 | 0.145832557 | 0.438104104 |
| CARMIL1 | 675.3811798 | -1.570857162 | 0.308006737 | -5.100074041 | 3.40E-07 | 9.51E-05 |
| CARMIL2 | 133.9903305 | -0.581613668 | 0.39509365 | -1.472090648 | 0.140996401 | 0.430713136 |
| CARMIL3 | 36.82426593 | -0.012235231 | 0.753034449 | -0.016247904 | 0.987036619 | 0.995287969 |
| CARNMT1 | 710.9986405 | 0.433182863 | 0.272924448 | 1.587189661 | 0.112469696 | 0.385686647 |
| CARNS1 | 19.11704209 | 0.29006681 | 0.95381391 | 0.304112581 | 0.761042122 | 0.9003659 |
| CARS | 643.5117414 | -0.363355425 | 0.27492571 | -1.321649492 | 0.186284895 | 0.490817669 |
| CARS2 | 771.0773692 | -0.190938486 | 0.391343866 | -0.487904634 | 0.625617392 | 0.835423066 |
| CASC1 | 15.1095328 | 0.078490642 | 0.912772636 | 0.085991449 | 0.931473214 | 0.974341807 |
| CASC10 | 670.5217368 | 0.080074163 | 0.302568471 | 0.264648074 | 0.791280582 | 0.913361935 |
| CASC3 | 4500.925141 | -0.236658476 | 0.286791072 | -0.82519471 | 0.409261056 | 0.703508181 |
| CASC4 | 1158.76161 | 0.339852691 | 0.259560243 | 1.309340317 | 0.190419099 | 0.496058205 |
| CASD1 | 999.1728054 | -0.09548227 | 0.367254889 | -0.259989105 | 0.794872178 | 0.914877916 |
| CASK | 2392.629759 | 0.02106205 | 0.224014487 | 0.094020926 | 0.925092534 | 0.971856438 |
| CASKIN1 | 61.82917779 | 0.513123196 | 0.629454086 | 0.815187648 | 0.414964907 | 0.707272509 |
| CASKIN2 | 361.2853987 | -0.472906779 | 0.432154546 | -1.094300136 | 0.273823366 | 0.586847211 |
| CASP10 | 95.3667881 | -0.836780592 | 0.398754012 | -2.09848821 | 0.03586204 | 0.216490931 |
| CASP2 | 1074.435938 | -0.501113034 | 0.28694958 | -1.74634524 | 0.080750979 | 0.325104967 |
| CASP3 | 1524.798997 | 0.210340169 | 0.26868378 | 0.78285399 | 0.433712859 | 0.71918874 |
| CASP6 | 1006.415357 | -0.164627764 | 0.450108277 | -0.365751469 | 0.714550536 | |

| | | | | | | |
|---|---|---|---|---|---|---|
| | | | | | | 0.878551437 |
| CASP7 | 250.8466386 | 0.238429996 | 0.278876792 | 0.854965357 | 0.392570326 | 0.691717122 |
| CASP8 | 86.89083245 | -0.214192255 | 0.553618368 | -0.386895138 | 0.698833841 | 0.870873025 |
| CASP8AP2 | 1173.770432 | -0.134456913 | 0.385359476 | -0.348912953 | 0.727154661 | 0.8850111 |
| CASP9 | 452.6187297 | -0.120898954 | 0.291686016 | -0.414483205 | 0.678520265 | 0.861138002 |
| CASQ1 | 29.5109321 | -1.954774256 | 1.075421593 | -1.817681799 | 0.069112781 | 0.303010569 |
| CASQ2 | 31.92886592 | -2.191798704 | 0.959331525 | -2.28471456 | 0.02232957 | 0.170232116 |
| CASR | 26.56034658 | -0.938010131 | 0.676537254 | -1.386487033 | 0.165598237 | 0.467560858 |
| CAST | 5240.507407 | -0.530654561 | 0.490080363 | -1.082790908 | 0.278901242 | 0.59252356 |
| CASTOR1 | 247.0464682 | -0.113333527 | 0.449851417 | -0.251935468 | 0.801090944 | 0.916680548 |
| CASTOR2 | 63.90329354 | -1.093615109 | 0.792286038 | -1.38032864 | 0.167485481 | 0.469758223 |
| CASTOR3 | 756.5294059 | -0.110667696 | 0.338610657 | -0.326828746 | 0.743797416 | 0.892535157 |
| CASZ1 | 577.2853668 | -1.029409055 | 0.398306758 | -2.584462939 | 0.009753079 | 0.112404636 |
| CAT | 223.7893479 | 0.2242076 | 0.395574698 | 0.566789538 | 0.570857179 | 0.806748565 |
| CATIP | 16.67418541 | 0.921591869 | 0.779967471 | 1.181577313 | 0.23737346 | 0.547879744 |
| CATSPER2 | 93.5698374 | -0.555507221 | 0.385457403 | -1.44116371 | 0.149538438 | 0.444386222 |
| CATSPER2P1 | 26.68359951 | 0.498297931 | 0.635068272 | 0.784636792 | 0.432666552 | 0.718639059 |
| CATSPER4 | 14.92959997 | 0.410342363 | 0.768342233 | 0.534061965 | 0.593298664 | 0.819859199 |
| CATSPERG | 125.2875852 | -1.77163575 | 0.419169194 | -4.226540921 | 2.37E-05 | 0.002093167 |
| CAV1 | 55.21959637 | -0.84316439 | 0.713649257 | -1.181482896 | 0.237410944 | 0.547879744 |
| CAV2 | 31.56153556 | 0.555381868 | 0.946366107 | 0.586857311 | 0.557299542 | 0.79822426 |
| CAVIN1 | 868.4070209 | 0.642190924 | 0.378979591 | 1.694526406 | 0.090165298 | 0.344106696 |
| CAVIN3 | 65.67004086 | 0.18879079 | 0.656433966 | 0.287600582 | 0.7736525 | 0.906198431 |
| CAVIN4 | 317.4952122 | -0.247600441 | 0.420072217 | -0.58942351 | 0.555577209 | 0.796434078 |
| CBARP | 68.65360443 | 0.629699574 | 0.797404636 | 0.789686373 | 0.429710952 | 0.716389544 |
| CBFA2T2 | 706.8587748 | -0.578783406 | 0.263771226 | -2.194262863 | 0.028216518 | 0.192229399 |
| CBFA2T3 | 24.70208331 | -0.111770105 | 0.69965674 | -0.159749915 | 0.873078079 | 0.94953238 |
| CBFB | 1162.535823 | 0.424886513 | 0.24636951 | 1.724590488 | 0.084601299 | 0.332854332 |
| CBL | 1226.172184 | -0.440053548 | 0.339553392 | -1.295977478 | 0.194983244 | 0.500590454 |
| CBLB | 1418.417241 | -1.159364053 | 0.368490397 | -3.146253097 | 0.001653768 | 0.037682927 |
| CBLC | 301.9872357 | -0.560372278 | 0.803365282 | -0.697531112 | 0.485470476 | 0.755840687 |
| CBLL1 | 1102.776438 | -0.22746133 | 0.249246343 | -0.912596459 | 0.361454818 | 0.668327975 |
| CBLN1 | 52.28985096 | -0.101339206 | 0.483333988 | -0.209667039 | 0.833927553 | 0.932107564 |
| CBLN3 | 11.89873148 | -0.212067859 | 0.897562453 | -0.236270867 | 0.813222492 | 0.921802034 |
| CBR1 | 349.4658909 | 0.686014628 | 0.342025154 | 2.005743203 | 0.044883653 | 0.243886068 |
| CBR4 | 1690.887655 | 0.25278695 | 0.340634566 | 0.742105984 | 0.458023124 | 0.736518606 |
| CBS | 2310.393087 | 0.133087961 | 0.31775025 | 0.418844551 | 0.675329743 | 0.860095396 |

| | | | | | | |
|---|---|---|---|---|---|---|
| CBSL | 2931.601129 | -0.147803796 | 0.311296943 | -0.474800022 | 0.634929517 | 0.840313679 |
| CBWD1 | 951.4015841 | -0.068519496 | 0.239604794 | -0.285968802 | 0.774902017 | 0.906450643 |
| CBWD2 | 936.4752656 | 0.122563083 | 0.270931399 | 0.452376813 | 0.650997548 | 0.847876685 |
| CBWD3 | 463.2970559 | -0.068378966 | 0.281552891 | -0.242863662 | 0.808111016 | 0.919580875 |
| CBWD4P | 45.2995073 | 1.259425462 | 0.741359242 | 1.69880591 | 0.08935576 | 0.342726775 |
| CBWD5 | 718.277398 | 0.232285661 | 0.303701825 | 0.764847759 | 0.444362205 | 0.727151225 |
| CBWD6 | 219.4656138 | 0.032896221 | 0.323661551 | 0.101637716 | 0.919044241 | 0.969677217 |
| CBX1 | 6110.920889 | 0.124702533 | 0.231594602 | 0.538451812 | 0.590265165 | 0.818321084 |
| CBX2 | 3174.37206 | 0.203896511 | 0.333398544 | 0.611569891 | 0.540822364 | 0.787864037 |
| CBX3 | 6963.895668 | 0.570109846 | 0.25177734 | 2.264341364 | 0.023553127 | 0.175417032 |
| CBX4 | 154.9861909 | -0.080377093 | 0.60687227 | -0.132444827 | 0.89463246 | 0.958237013 |
| CBX5 | 7146.79141 | 0.196096486 | 0.357090457 | 0.549150734 | 0.582902011 | 0.813692161 |
| CBX6 | 674.7636906 | 0.900105587 | 0.375981071 | 2.394018363 | 0.016664914 | 0.146056278 |
| CBX7 | 169.8756625 | 0.046814876 | 0.337553087 | 0.13868893 | 0.889695966 | 0.956447219 |
| CBX8 | 140.4337976 | 0.732929839 | 0.546043134 | 1.342256303 | 0.179512907 | 0.484443207 |
| CBY1 | 695.2154029 | 0.418382899 | 0.3545289 | 1.180109432 | 0.237956694 | 0.54863845 |
| CC2D1A | 402.1588217 | -0.503736086 | 0.448806652 | -1.122389973 | 0.261696666 | 0.57522459 |
| CC2D1B | 877.1953718 | -0.408823188 | 0.253716474 | -1.611338758 | 0.107105908 | 0.375093196 |
| CC2D2A | 581.3478555 | -0.167007348 | 0.285282769 | -0.585409866 | 0.558272157 | 0.798855459 |
| CC2D2B | 12.65329275 | -0.292062835 | 0.860139947 | -0.339552693 | 0.734193409 | 0.889013346 |
| CCAR1 | 3751.327072 | 0.129399081 | 0.32495064 | 0.398211497 | 0.690474292 | 0.867938158 |
| CCAR2 | 4008.575302 | -0.223431705 | 0.325810017 | -0.685772977 | 0.49285628 | 0.759389046 |
| CCBE1 | 906.7381915 | -3.588261537 | 0.750459973 | -4.781416286 | 1.74E-06 | 0.00027224 |
| CCDC102A | 165.7216261 | -0.316579681 | 0.490477834 | -0.645451556 | 0.518634603 | 0.774353465 |
| CCDC102B | 155.959646 | 0.269609992 | 0.336440282 | 0.801360619 | 0.422922907 | 0.712273914 |
| CCDC103 | 74.66875156 | -1.077150904 | 2.119016073 | -0.508325972 | 0.611224759 | 0.829058038 |
| CCDC106 | 302.8317876 | -0.14564471 | 0.469072073 | -0.310495376 | 0.756184275 | 0.898288428 |
| CCDC107 | 190.1887433 | 0.81484885 | 0.426923641 | 1.90865244 | 0.056306942 | 0.274938082 |
| CCDC110 | 16.01284127 | 1.111589348 | 0.878272766 | 1.265653897 | 0.205637021 | 0.511932925 |
| CCDC112 | 524.9492256 | 1.039539258 | 0.316320421 | 3.286348871 | 0.001014952 | 0.027467362 |
| CCDC113 | 379.3849332 | -0.264667486 | 0.329308988 | -0.803705627 | 0.421567004 | 0.711966271 |
| CCDC114 | 32.356031 | -0.008441286 | 0.593738195 | -0.014217186 | 0.988656709 | 0.995803391 |
| CCDC115 | 241.3372198 | 0.350425657 | 0.337425541 | 1.038527361 | 0.299024604 | 0.610246278 |
| CCDC117 | 1328.41883 | 0.263001211 | 0.231426569 | 1.136434818 | 0.255774631 | 0.567999769 |
| CCDC12 | 1080.259829 | 0.762724806 | 0.460767182 | 1.655336653 | 0.097856228 | 0.359375073 |
| CCDC120 | 701.2188031 | -1.602380879 | 0.439151386 | -3.648812071 | 0.000263456 | 0.011290641 |
| CCDC121 | 131.8769499 | 0.248927776 | 0.33726719 | 0.73807291 | 0.460470145 | 0.738395128 |
| CCDC122 | 183.7450474 | -0.280853749 | 0.422448427 | -0.664823755 | 0.506163227 | 0.767209743 |

| | | | | | | |
|---|---|---|---|---|---|---|
| CCDC124 | 812.0115511 | 0.214263901 | 0.420284865 | 0.509806369 | 0.610187123 | 0.828605673 |
| CCDC125 | 194.5101151 | -0.343828517 | 0.330416445 | -1.040591417 | 0.298065216 | 0.609375237 |
| CCDC126 | 302.7098107 | -0.411382984 | 0.474332728 | -0.867287791 | 0.385784344 | 0.686679504 |
| CCDC127 | 734.0900488 | 0.1132364 | 0.277564813 | 0.407963815 | 0.683300241 | 0.863415274 |
| CCDC13 | 33.81657775 | 0.660315692 | 0.53394561 | 1.236672199 | 0.216208804 | 0.523980406 |
| CCDC130 | 518.0917873 | -0.25563626 | 0.397431924 | -0.643220246 | 0.520081198 | 0.775216335 |
| CCDC134 | 75.41953935 | 0.230550425 | 0.407494921 | 0.565774966 | 0.571546766 | 0.807274555 |
| CCDC136 | 587.4420536 | 0.882916102 | 0.542769815 | 1.626686079 | 0.103803787 | 0.369190637 |
| CCDC137 | 898.3372191 | 0.070580201 | 0.328730361 | 0.214705451 | 0.829996972 | 0.93010439 |
| CCDC138 | 646.3080928 | 0.533019552 | 0.324318913 | 1.643504375 | 0.100278621 | 0.36243601 |
| CCDC14 | 3093.85283 | -0.0925872 | 0.323122389 | -0.286539104 | 0.774465247 | 0.90635568 |
| CCDC140 | 33.04597785 | 0.088574698 | 0.787258949 | 0.112510246 | 0.910418846 | 0.966517087 |
| CCDC141 | 12.10927149 | -0.131154171 | 0.883419362 | -0.148461961 | 0.881978204 | 0.953217539 |
| CCDC142 | 182.9698085 | -0.452506105 | 0.529928702 | -0.853899974 | 0.393160414 | 0.692183315 |
| CCDC144A | 97.47195609 | 0.268235759 | 0.431125632 | 0.622175392 | 0.533826545 | 0.783156734 |
| CCDC144B | 33.92249668 | -0.199002703 | 0.703632081 | -0.2828221 | 0.777313212 | 0.907898743 |
| CCDC144NL | 14.29415213 | 0.725422423 | 0.992383506 | 0.730990003 | 0.464785259 | 0.74184715 |
| CCDC146 | 55.91576571 | -0.804390243 | 0.570015929 | -1.411171517 | 0.158194047 | 0.457584529 |
| CCDC148 | 87.10107973 | -0.055504483 | 0.446174114 | -0.124400949 | 0.900997822 | 0.961727852 |
| CCDC149 | 165.250622 | -0.762603272 | 0.403241326 | -1.891183324 | 0.058599877 | 0.280288923 |
| CCDC15 | 106.3090266 | 0.577979198 | 0.379575088 | 1.522700556 | 0.127833639 | 0.412017025 |
| CCDC150 | 606.1956624 | -0.539170354 | 0.340803843 | -1.582054797 | 0.113637062 | 0.388022186 |
| CCDC151 | 22.0599193 | 0.752368603 | 0.912846381 | 0.824200674 | 0.409825544 | 0.703966274 |
| CCDC152 | 422.2319533 | 0.566866882 | 0.295951279 | 1.915406086 | 0.055440736 | 0.272416332 |
| CCDC153 | 117.0127717 | 0.171473243 | 0.531681324 | 0.322511314 | 0.747065368 | 0.89418504 |
| CCDC154 | 22.32539617 | 0.222205314 | 0.809478051 | 0.274504434 | 0.783696997 | 0.910512111 |
| CCDC157 | 101.4066925 | -0.020549532 | 0.42966172 | -0.047827234 | 0.961853932 | 0.9870991 |
| CCDC159 | 96.03667502 | 0.317613192 | 0.407838018 | 0.778772891 | 0.436113509 | 0.720784523 |
| CCDC160 | 327.0223298 | 1.275123154 | 0.520200028 | 2.451217004 | 0.014237409 | 0.135535502 |
| CCDC162P | 134.3350523 | -0.299819032 | 0.436834377 | -0.686344866 | 0.492495663 | 0.759174768 |
| CCDC163 | 82.9632734 | -0.838192098 | 0.456972789 | -1.834227591 | 0.066620217 | 0.297701125 |
| CCDC167 | 758.0801453 | 1.18203175 | 0.407628744 | 2.899775269 | 0.003734303 | 0.063017919 |

CCDC169 638.7718873 0.105830547 0.317586837 0.333233418 0.738958095 0.891443823
CCDC169-SOHLH2 23.12254265 0.638078036 1.014119771 0.629193961 0.529222082 0.780028
CCDC17 35.5757057 -1.216120158 0.583940134 -2.082611019 0.037286694 0.221140123
CCDC171 135.1915778 -0.480434314 0.447510025 -1.073572182 0.283014471 0.596961028
CCDC173 22.71537244 0.928718185 0.689264357 1.347404917 0.177849858 0.482075465
CCDC174 644.7432587 -0.31809238 0.282157046 -1.127359337 0.259590584 0.572948272
CCDC177 35.14214409 0.871834705 0.683207406 1.27609083 0.201923413 0.508529233
CCDC18 475.0880151 0.460778192 0.321830418 1.431742204 0.152217613 0.448119169
CCDC180 264.4839368 -0.223782722 0.495308254 -0.451804952 0.651409502 0.847876685
CCDC181 304.0107032 0.735965637 0.360930793 2.039076885 0.041442355 0.233812556
CCDC183 98.80037973 -0.219459837 0.68069979 -0.322403269 0.747147208 0.89418504
CCDC184 18.2744016 1.518765433 0.904252286 1.679581525 0.093038765 0.349658165
CCDC186 778.934173 -0.725963748 0.313345134 -2.316818322 0.020513631 0.161921066
CCDC189 49.46669287 0.071312094 0.481620937 0.148066847 0.882290013 0.953230092
CCDC191 192.1054793 -0.384244756 0.338181342 -1.13620921 0.255869017 0.568011246
CCDC192 11.21038674 -0.275352409 1.009341732 -0.272803948 0.785003925 0.910931322
CCDC200 13.59638894 0.054812709 0.988411262 0.055455367 0.955775687 0.984323736
CCDC22 429.3623974 0.137830911 0.413308176 0.333482179 0.738770339 0.89135397
CCDC24 293.1311977 -0.758044281 0.416852017 -1.818497335 0.068988154 0.302794854
CCDC25 1998.378552 0.356056617 0.252391703 1.410730275 0.158324161 0.45780246
CCDC28A 291.9646803 -0.030386202 0.538838345 -0.056392056 0.955029485 0.984323736
CCDC28B 200.4598971 0.877999308 0.376997216 2.328927828 0.019862889 0.159523502
CCDC3 191.7963951 -2.558451887 0.871698215 -2.935020219 0.00333526 0.058361272
CCDC30 100.9050888 -0.414615646 0.397703323 -1.042524973 0.297168353 0.608609612
CCDC32 335.7409438 0.339111011 0.263206137 1.288385652 0.197611751 0.503183597
CCDC33 236.0732053 -0.369186733 0.423046331 -0.87268629 0.382834092 0.683935
CCDC34 627.3451122 1.116280538 0.469513396 2.377526494 0.017429191 0.149104584
CCDC39 27.47588 -0.161149583 0.611790378 -0.263406535 0.792237254 0.913570955
CCDC40 151.5000594 -0.080293901 0.50743757 -0.158234048 0.874272375 0.949740014
CCDC43 1048.036192 0.203572296 0.307437785 0.662157696 0.507870163 0.768645076
CCDC47 2637.115163 -0.321184325 0.257051912 -1.249492068 0.211485153 0.518745554
CCDC50 1382.553334 0.015060379 0.226269373 0.066559512 0.946932379 0.980608267

| | | | | | | |
|---|---|---|---|---|---|---|
| CCDC51 | 430.110162 | 0.41373592 | 0.355237778 | 1.164673202 | 0.2441513 | 0.555457814 |
| CCDC57 | 604.3071472 | -0.843495416 | 0.404109773 | -2.087292794 | 0.036861676 | 0.219736143 |
| CCDC58 | 668.4867038 | 0.764322424 | 0.282355778 | 2.706948056 | 0.006790488 | 0.090367101 |
| CCDC59 | 998.3476201 | 0.739367236 | 0.423816919 | 1.744543936 | 0.081064287 | 0.325604941 |
| CCDC6 | 1798.66142 | -0.652747641 | 0.366188353 | -1.782546155 | 0.074660212 | 0.313779586 |
| CCDC60 | 15.97431885 | 0.572402599 | 0.773894829 | 0.739638743 | 0.459519227 | 0.737476588 |
| CCDC61 | 140.6687289 | 0.363128472 | 0.38332295 | 0.947317326 | 0.343477105 | 0.650427919 |
| CCDC62 | 11.68139996 | -0.229147971 | 0.872903344 | -0.262512422 | 0.79292641 | 0.913966604 |
| CCDC65 | 15.78337353 | -0.107021396 | 0.765549056 | -0.139796915 | 0.888820451 | 0.956197352 |
| CCDC66 | 229.9687016 | 0.204131074 | 0.347631048 | 0.587206105 | 0.557065293 | 0.798114795 |
| CCDC68 | 15.25970778 | -2.264937944 | 0.961914493 | -2.354614635 | 0.018541924 | 0.154302741 |
| CCDC69 | 352.3616647 | -1.598020098 | 0.423724611 | -3.771364835 | 0.000162357 | 0.008377957 |
| CCDC7 | 34.39646302 | -0.171076566 | 0.529707266 | -0.322964357 | 0.746722236 | 0.894109554 |
| CCDC71 | 613.2428465 | 0.11540828 | 0.309393164 | 0.373014963 | 0.709137307 | 0.875799584 |
| CCDC71L | 968.2693321 | 0.127858827 | 0.570005383 | 0.224311613 | 0.822514841 | 0.925840841 |
| CCDC74A | 126.0500737 | 1.735420194 | 0.563401135 | 3.080256831 | 0.002068222 | 0.043016015 |
| CCDC74B | 115.0423112 | 1.251385608 | 0.729776106 | 1.714752782 | 0.086390562 | 0.337171176 |
| CCDC77 | 830.4024269 | 0.536870155 | 0.386672198 | 1.388437435 | 0.165003884 | 0.467115865 |
| CCDC78 | 28.12282092 | 1.183154357 | 0.684007856 | 1.729737965 | 0.083677102 | 0.330936505 |
| CCDC8 | 423.9709829 | 0.273701958 | 0.346279498 | 0.79040763 | 0.429289747 | 0.71613003 |
| CCDC80 | 297.2493321 | 0.475389235 | 0.485371156 | 0.979434458 | 0.327365358 | 0.637370783 |
| CCDC81 | 24.39775983 | -0.493626171 | 0.688739286 | -0.716709764 | 0.4735532 | 0.747811725 |
| CCDC82 | 1016.806803 | -0.071154557 | 0.374398148 | -0.190050504 | 0.849269555 | 0.938580023 |
| CCDC84 | 238.6051258 | -0.773324208 | 0.43923891 | -1.760600415 | 0.078306058 | 0.320989249 |
| CCDC85A | 17.89255752 | 0.77895295 | 0.708245322 | 1.099834938 | 0.271404047 | 0.583632536 |
| CCDC85B | 100.3584341 | 1.421381591 | 0.809133381 | 1.756671552 | 0.078973809 | 0.322507823 |
| CCDC85C | 2695.785328 | -0.273232028 | 0.321249086 | -0.850530135 | 0.395030414 | 0.693813886 |
| CCDC86 | 569.0791868 | 0.319439781 | 0.273768735 | 1.166823454 | 0.243281673 | 0.554511158 |
| CCDC87 | 18.41222614 | -0.415789286 | 0.718663673 | -0.578558931 | 0.562886823 | 0.801767863 |
| CCDC88A | 2820.843464 | 0.786349454 | 0.382032996 | 2.058328629 | 0.039558598 | 0.229237914 |
| CCDC88B | 238.6707657 | -0.366463781 | 0.404730804 | -0.905450679 | 0.365226678 | 0.671700076 |
| CCDC88C | 1336.538669 | -0.47564531 | 0.295181901 | -1.611363396 | 0.107100541 | 0.375093196 |
| CCDC9 | 427.5746522 | 0.190932911 | 0.334352983 | 0.571051915 | 0.567964452 | 0.805288664 |

| | | | | | | |
|---|---|---|---|---|---|---|
| CCDC90B | 2021.657595 | 0.570277237 | 0.361731489 | 1.576520856 | 0.114905816 | 0.389985713 |
| CCDC91 | 395.3942286 | 0.251581952 | 0.296382316 | 0.848842654 | 0.395968852 | 0.694559308 |
| CCDC92 | 78.80463197 | -2.214015478 | 0.892986011 | -2.479339486 | 0.013162596 | 0.130006343 |
| CCDC93 | 859.1016136 | -0.596043914 | 0.409468685 | -1.455652008 | 0.145488843 | 0.437244698 |
| CCDC96 | 11.40709432 | -0.601465977 | 1.000315876 | -0.601276048 | 0.54765614 | 0.792794971 |
| CCDC97 | 909.9100417 | -0.468181263 | 0.317343457 | -1.475314056 | 0.140128129 | 0.429639018 |
| CCDC9B | 14.10340103 | -0.145336359 | 0.836387702 | -0.173766734 | 0.862048789 | 0.944228049 |
| CCER2 | 12.7658825 | 0.94250846 | 0.856070904 | 1.100970089 | 0.270909675 | 0.583394591 |
| CCHCR1 | 647.4610443 | 0.173031712 | 0.338574681 | 0.511059218 | 0.609309591 | 0.8283007 |
| CCKBR | 200.4201634 | -1.560030428 | 0.786848374 | -1.982631571 | 0.047408598 | 0.25076445 |
| CCL2 | 65.52644712 | 1.168344447 | 0.638267906 | 1.830492238 | 0.067176366 | 0.297817842 |
| CCL27 | 21.01752531 | 1.092516535 | 0.69797927 | 1.565256423 | 0.117522796 | 0.395641036 |
| CCM2 | 658.2564611 | 0.061597812 | 0.372629332 | 0.16530586 | 0.868703246 | 0.947046732 |
| CCM2L | 89.306556 | 0.182933047 | 0.540819755 | 0.338251415 | 0.735173733 | 0.889632345 |
| CCN1 | 2877.978582 | -1.627040638 | 0.634968379 | -2.56239632 | 0.010395262 | 0.115896273 |
| CCN2 | 1668.243733 | -1.477848575 | 0.680338672 | -2.172224859 | 0.029838707 | 0.197226575 |
| CCN3 | 12.16131903 | 1.705343688 | 0.827971229 | 2.059665394 | 0.03943054 | 0.228786879 |
| CCNA1 | 16.45037776 | 0.15569053 | 0.749536818 | 0.207715653 | 0.835450995 | 0.932531417 |
| CCNA2 | 2131.472454 | 0.991197045 | 0.307325371 | 3.225236631 | 0.001258686 | 0.031407111 |
| CCNB1 | 4382.867167 | 0.942485887 | 0.323077326 | 2.917214585 | 0.003531727 | 0.061024865 |
| CCNB1IP1 | 4414.815855 | 0.391676726 | 0.287420521 | 1.362730555 | 0.172967445 | 0.476760374 |
| CCNB2 | 2905.482881 | 0.941265682 | 0.268065536 | 3.511326724 | 0.000445876 | 0.016477615 |
| CCNC | 3190.734889 | 0.222527974 | 0.234201628 | 0.950155541 | 0.342033225 | 0.648704346 |
| CCND1 | 3775.342764 | 0.616420201 | 0.502776635 | 1.226031916 | 0.220186665 | 0.528727493 |
| CCND2 | 826.5777697 | 0.319293929 | 0.381977348 | 0.835897551 | 0.40321254 | 0.699821239 |
| CCND3 | 283.851649 | -0.224246552 | 0.564586644 | -0.397187135 | 0.691229468 | 0.868004248 |
| CCNDBP1 | 1074.702169 | -0.030529944 | 0.404272195 | -0.075518287 | 0.939802348 | 0.978003717 |
| CCNE1 | 562.3073943 | -0.384903733 | 0.345502279 | -1.114041081 | 0.265261563 | 0.579200226 |
| CCNE2 | 84.39651886 | 0.8095993 | 0.426504516 | 1.898219757 | 0.057667139 | 0.278177478 |
| CCNF | 1105.765765 | 0.60762243 | 0.308924203 | 1.966898109 | 0.049194959 | 0.255507264 |
| CCNG1 | 17274.40931 | 0.674932414 | 0.28366748 | 2.379308391 | 0.017345158 | 0.14883676 |
| CCNG2 | 1753.679469 | -0.117322096 | 0.256062612 | -0.458177379 | 0.64682501 | 0.846556814 |
| CCNH | 739.5288685 | 0.844855836 | 0.293587947 | 2.877692506 | 0.004005954 | 0.065575642 |
| CCNI | 11067.3157 | 0.399713303 | 0.467820216 | 0.85441648 | 0.392874268 | 0.692011554 |
| CCNJ | 1069.43362 | 0.074551449 | 0.306066487 | 0.243579262 | 0.807556691 | 0.919453224 |
| CCNJL | 581.2706202 | 0.112937897 | 0.338648502 | 0.333495928 | 0.738759963 | 0.89135397 |
| CCNK | 1807.718481 | -0.398434448 | 0.226762361 | -1.757057238 | 0.078908053 | 0.322507823 |
| CCNL1 | 2665.684393 | -0.425816061 | 0.235999795 | -1.804306911 | 0.071183207 | 0.306992555 |
| CCNL2 | 4259.847843 | -0.511121392 | 0.279054398 | -1.831619199 | 0.067008174 | 0.297817842 |
| CCNQ | 243.5883916 | 0.657528757 | 0.362931612 | 1.811715308 | 0.070030197 | 0.303985156 |
| CCNT1 | 1336.702019 | -0.172560068 | 0.318495531 | -0.541797454 | 0.587958045 | 0.817215896 |
| CCNT2 | 1973.374239 | -0.081024748 | 0.313894846 | -0.25812704 | 0.796308863 | 0.915420305 |
| CCNY | 1611.898245 | 0.128901428 | 0.238000217 | 0.541602144 | 0.588092614 | 0.817253188 |
| CCNYL1 | 849.5307595 | 0.071651606 | 0.29633535 | 0.2417923 | 0.808941105 | |

0.919641737

| | | | | | | |
|---|---|---|---|---|---|---|
| CCNYL2 | 32.81542175 | -1.415850716 | 0.684601673 | -2.068137974 | 0.038627051 | 0.226521449 |
| CCP110 | 581.1815971 | 0.194518384 | 0.390681034 | 0.497895642 | 0.618557599 | 0.83229541 |
| CCPG1 | 625.9296275 | 0.06306972 | 0.259406587 | 0.243130757 | 0.807904106 | 0.919552343 |
| CCR1 | 41.60464715 | -1.65373311 | 0.654149594 | -2.528065637 | 0.011469291 | 0.121279761 |
| CCR10 | 23.45034588 | 0.851592049 | 0.964694592 | 0.882758186 | 0.377366937 | 0.680741236 |
| CCR7 | 103.0270871 | -0.661860602 | 0.529400529 | -1.250207671 | 0.211223695 | 0.518607329 |
| CCS | 847.0396698 | 0.349554633 | 0.422896263 | 0.826573 | 0.408479126 | 0.702771243 |
| CCSAP | 466.1677476 | 0.187049883 | 0.311991687 | 0.599534829 | 0.548816291 | 0.793468239 |
| CCSER2 | 817.8475316 | -0.334520845 | 0.25187892 | -1.32810179 | 0.184144482 | 0.488572217 |
| CCT2 | 9040.568754 | 0.496628414 | 0.233007265 | 2.131385961 | 0.033057358 | 0.207677316 |
| CCT3 | 13495.84408 | 0.170674309 | 0.23293222 | 0.732720914 | 0.463728666 | 0.740940241 |
| CCT4 | 12723.00932 | 0.667622566 | 0.309435071 | 2.157552999 | 0.030962602 | 0.20078452 |
| CCT5 | 10222.30821 | 0.464880241 | 0.262916898 | 1.768164179 | 0.077033461 | 0.31844659 |
| CCT6A | 10306.00855 | 0.389276635 | 0.228876591 | 1.700814543 | 0.088977818 | 0.342496199 |
| CCT6B | 95.07002885 | 0.590382142 | 0.433068116 | 1.363254696 | 0.172802255 | 0.476478191 |
| CCT6P1 | 157.2558522 | -0.498358461 | 0.357443746 | -1.394229068 | 0.163248443 | 0.464573649 |
| CCT6P3 | 245.5277519 | 0.397861439 | 0.491490566 | 0.809499647 | 0.418227806 | 0.709589334 |
| CCT7 | 8588.457419 | 0.073713443 | 0.320870676 | 0.229729446 | 0.818302012 | 0.924313888 |
| CCT8 | 15644.73091 | 0.556132474 | 0.256906128 | 2.16473028 | 0.030408347 | 0.198932742 |
| CCZ1 | 2029.74689 | 0.059289658 | 0.24453365 | 0.242460121 | 0.808423654 | 0.91963689 |
| CCZ1B | 1431.574127 | 0.693277375 | 0.409157763 | 1.69440113 | 0.090189085 | 0.344106696 |
| CD101 | 20.76483961 | 0.764019921 | 0.698474064 | 1.093841505 | 0.274024497 | 0.586847211 |
| CD109 | 110.6943757 | -1.19584087 | 0.379482087 | -3.151244585 | 0.001625763 | 0.03731384 |
| CD151 | 2176.79982 | 0.163923029 | 0.31655049 | 0.517841654 | 0.604568754 | 0.825894731 |
| CD163L1 | 100.8850759 | -0.278533152 | 0.550686021 | -0.50579303 | 0.613001952 | 0.829603266 |
| CD164 | 4133.850001 | -0.267626472 | 0.373532247 | -0.716474881 | 0.473698172 | 0.747889996 |
| CD164L2 | 29.63474172 | 1.014043929 | 0.838955011 | 1.208698816 | 0.226778577 | 0.535481989 |
| CD200 | 532.5910648 | 0.876407804 | 0.340306922 | 2.575345217 | 0.010014008 | 0.114014677 |
| CD24 | 24687.21423 | -0.333383433 | 0.428107145 | -0.778738307 | 0.436133885 | 0.720784523 |
| CD247 | 44.64586975 | 0.32383786 | 0.632217291 | 0.512225566 | 0.608493152 | 0.827858453 |
| CD248 | 46.68139899 | 0.410557181 | 0.759684769 | 0.540430976 | 0.58889985 | 0.818050255 |
| CD274 | 27.83893895 | -3.145188512 | 1.007372518 | -3.122170257 | 0.001795231 | 0.039357583 |
| CD276 | 2856.641439 | 0.227579315 | 0.249440322 | 0.912359771 | 0.361579359 | 0.66847678 |
| CD2AP | 2223.309766 | -0.484171094 | 0.275348328 | -1.758394895 | 0.078680341 | 0.322001471 |
| CD2BP2 | 2486.875508 | 0.175884863 | 0.264901097 | 0.66396427 | 0.50671318 | 0.767639613 |
| CD302 | 35.68739246 | 0.865470103 | 0.619048431 | 1.398065255 | 0.162093472 | 0.462717357 |
| CD320 | 671.4691184 | 1.295246016 | 0.410991093 | 3.151518459 | 0.001624239 | 0.03731384 |
| CD34 | 14.61826692 | -0.53604874 | 1.005480499 | -0.533126938 | 0.593945713 | 0.820352368 |
| CD38 | 13.91302511 | 0.630528918 | 1.050506242 | 0.600214347 | 0.548363393 | 0.793364585 |
| CD3EAP | 402.5750265 | 0.511967226 | 0.40925176 | 1.250983565 | 0.210940474 | 0.518294063 |
| CD4 | 189.3579314 | 0.79230248 | 0.576771939 | 1.373684166 | 0.169539755 | 0.472289317 |
| CD40 | 200.6435871 | 0.738360486 | 0.376813254 | 1.959486506 | 0.050055839 | 0.257487003 |
| CD44 | 66.03418521 | 0.736717241 | 0.580694237 | 1.268683575 | 0.204553943 | 0.510835602 |
| CD46 | 2624.306304 | 0.042119369 | 0.294611413 | 0.142965843 | 0.886317157 | 0.955129819 |
| CD47 | 1324.880376 | -0.175100896 | 0.304089489 | -0.575820285 | 0.564736661 | 0.803200575 |

| | | | | | | |
|---|---|---|---|---|---|---|
| CD52 | 31.36833797 | 0.012216957 | 1.052383156 | 0.011608849 | 0.990737687 | 0.996726674 |
| CD55 | 843.5341576 | -0.135201916 | 0.429597429 | -0.314717703 | 0.752976002 | 0.896586461 |
| CD58 | 117.5391705 | 0.709158097 | 0.3671416 | 1.931565634 | 0.053413141 | 0.26722241 |
| CD59 | 2105.680135 | 0.193871081 | 0.244786233 | 0.792001569 | 0.428359761 | 0.715316008 |
| CD63 | 7774.754701 | 0.758431897 | 0.47344319 | 1.601949111 | 0.109166863 | 0.379246733 |
| CD68 | 36.7010069 | 0.128413535 | 0.541536483 | 0.237128132 | 0.812557388 | 0.921654347 |
| CD7 | 16.09711351 | 0.614203608 | 0.819510589 | 0.74947611 | 0.453570293 | 0.733497409 |
| CD70 | 246.0433169 | 2.187364734 | 0.537378001 | 4.07043967 | 4.69E-05 | 0.003559457 |
| CD72 | 53.16741706 | 0.033388019 | 0.59866923 | 0.055770394 | 0.95552472 | 0.984323736 |
| CD74 | 400.8674824 | -0.027292928 | 0.323143089 | -0.084460814 | 0.932690058 | 0.974543059 |
| CD79A | 10.42304141 | 1.010402668 | 1.014037707 | 0.996415282 | 0.319048411 | 0.629098445 |
| CD79B | 31.93176358 | 0.988944907 | 0.771494304 | 1.281856394 | 0.199893027 | 0.50598138 |
| CD81 | 4850.731127 | 0.026076689 | 0.390488501 | 0.06677966 | 0.946757117 | 0.980560637 |
| CD82 | 316.7006874 | 0.425164763 | 0.506652342 | 0.839164705 | 0.401376891 | 0.698793758 |
| CD83 | 94.66498499 | -0.058238441 | 0.369419752 | -0.157648422 | 0.874733846 | 0.949740014 |
| CD8A | 10.17599046 | 0.202053023 | 1.283623539 | 0.157408319 | 0.874923058 | 0.949740014 |
| CD8B | 26.13357765 | 0.394949884 | 0.948369865 | 0.416451322 | 0.67707978 | 0.86029961 |
| CD9 | 1099.091881 | 0.440933804 | 0.379307343 | 1.162471045 | 0.24504418 | 0.556521744 |
| CD99 | 3741.790377 | -0.080689312 | 0.550506999 | -0.146572727 | 0.883469282 | 0.953822408 |
| CD99L2 | 563.6527046 | 0.747923144 | 0.589530236 | 1.268676479 | 0.204556475 | 0.510835602 |
| CD99P1 | 425.9264068 | -0.336831622 | 0.438215304 | -0.768644131 | 0.442104599 | 0.725491496 |
| CDA | 187.9868186 | 0.368174475 | 0.766330698 | 0.480438114 | 0.630915899 | 0.838320891 |
| CDADC1 | 144.7447275 | 0.592034029 | 0.378332391 | 1.56485155 | 0.117617721 | 0.395641036 |
| CDAN1 | 579.0872895 | -0.246478186 | 0.296045695 | -0.832568048 | 0.405088385 | 0.700857535 |
| CDC123 | 2701.168886 | 0.173224578 | 0.263051316 | 0.658520097 | 0.510203988 | 0.769949736 |
| CDC14A | 367.6369636 | -0.177922177 | 0.290324558 | -0.612838882 | 0.539982878 | 0.787321504 |
| CDC14B | 1055.959379 | -1.061411553 | 0.321280037 | -3.303695939 | 0.000954193 | 0.026416161 |
| CDC16 | 1858.474822 | -0.129042627 | 0.276443957 | -0.466794893 | 0.64064663 | 0.843724522 |
| CDC20 | 3711.399556 | 0.809337848 | 0.299677001 | 2.700700574 | 0.00691936 | 0.091360846 |
| CDC20B | 30.35505334 | -4.0239253 | 1.574906092 | -2.555025548 | 0.010618 | 0.117068085 |
| CDC23 | 1722.459449 | -0.197310266 | 0.246088265 | -0.801786571 | 0.422676428 | 0.712269066 |
| CDC25A | 952.6463024 | 0.251475392 | 0.343554112 | 0.731981902 | 0.464179614 | 0.741159393 |
| CDC25B | 1788.273602 | 0.262711286 | 0.29011311 | 0.905547789 | 0.365175254 | 0.671700076 |
| CDC25C | 315.0852224 | 1.031487077 | 0.423456158 | 2.435876908 | 0.014855739 | 0.138598988 |
| CDC26 | 770.4939641 | 0.877148378 | 0.49469483 | 1.773110058 | 0.076210472 | 0.316642999 |
| CDC27 | 1640.534955 | 0.443882699 | 0.334625677 | 1.326505195 | 0.184672415 | 0.489202936 |
| CDC34 | 1515.559163 | 0.37700064 | 0.301773653 | 1.249282818 | 0.21156165 | 0.51879501 |
| CDC37 | 3401.474004 | -0.254405429 | 0.328495641 | -0.774456025 | 0.438661163 | 0.722366178 |
| CDC37L1 | 493.177704 | 0.448597538 | 0.298900633 | 1.500824982 | 0.133400835 | 0.419880513 |
| CDC40 | 1140.264676 | -0.019489336 | 0.25526478 | -0.076349489 | 0.939141054 | 0.977796709 |
| CDC42 | 8504.937049 | -0.233738745 | 0.302083225 | -0.773756123 | 0.439075024 | 0.722630418 |
| CDC42BPA | 937.66163 | -0.318798777 | 0.406831557 | -0.783613689 | 0.433266821 | 0.718841967 |

| | | | | | | |
|---|---|---|---|---|---|---|
| CDC42BPB | 3040.41056 | -0.672667795 | 0.51321809 | -1.310686057 | 0.189963851 | 0.49534919 |
| CDC42BPG | 883.8377393 | -1.279317326 | 0.364692404 | -3.507935212 | 0.000451599 | 0.016498831 |
| CDC42EP1 | 128.0711859 | -0.770195788 | 0.588360631 | -1.309053916 | 0.190516088 | 0.496058205 |
| CDC42EP2 | 60.27713219 | 0.577102812 | 0.758908657 | 0.760437777 | 0.446992949 | 0.728783454 |
| CDC42EP3 | 668.2785952 | -0.178564823 | 0.378131113 | -0.472229914 | 0.63676269 | 0.841268221 |
| CDC42EP4 | 374.9421741 | 0.210107787 | 0.465110374 | 0.451737478 | 0.651458116 | 0.847876685 |
| CDC42SE1 | 4314.478804 | -0.885997639 | 0.320177119 | -2.767210979 | 0.005653815 | 0.080690528 |
| CDC42SE2 | 1363.808557 | 0.075864886 | 0.232245425 | 0.326658256 | 0.743926377 | 0.892535157 |
| CDC45 | 479.1255643 | 0.222054037 | 0.317592049 | 0.699180089 | 0.484439492 | 0.75494931 |
| CDC5L | 2270.712715 | 0.213844403 | 0.277744563 | 0.769931913 | 0.441340282 | 0.724864504 |
| CDC6 | 1258.550555 | 0.003359595 | 0.313903456 | 0.010702639 | 0.991460693 | 0.996814923 |
| CDC7 | 1666.409074 | 0.582628229 | 0.320920867 | 1.815488766 | 0.069448831 | 0.30336211 |
| CDC73 | 1351.106561 | -0.187000172 | 0.242052135 | -0.772561548 | 0.439781909 | 0.723322998 |
| CDCA2 | 966.8831573 | 0.673319588 | 0.398948098 | 1.687737306 | 0.091461667 | 0.347000104 |
| CDCA3 | 576.2729456 | 1.08240691 | 0.289140421 | 3.743533702 | 0.00018145 | 0.008851388 |
| CDCA4 | 1123.361203 | 0.795853052 | 0.26713548 | 2.979211339 | 0.002889914 | 0.052996691 |
| CDCA5 | 1588.257616 | 0.320595037 | 0.273560231 | 1.171935829 | 0.241222821 | 0.552673201 |
| CDCA7 | 1456.36977 | 0.107180141 | 0.265782808 | 0.403262129 | 0.686755399 | 0.865835142 |
| CDCA7L | 1735.640989 | 0.368005548 | 0.473950356 | 0.776464336 | 0.437474871 | 0.721468765 |
| CDCA8 | 1374.613561 | 0.742494675 | 0.352079135 | 2.108885765 | 0.034954443 | 0.213397931 |
| CDCP1 | 248.6891352 | -2.042601919 | 0.602254663 | -3.391591707 | 0.000694879 | 0.021675701 |
| CDH1 | 11224.1046 | -2.231642813 | 0.736094227 | -3.031735246 | 0.002431524 | 0.047414712 |
| CDH10 | 75.10467554 | -1.875954005 | 1.089490788 | -1.721863117 | 0.085094318 | 0.334273924 |
| CDH11 | 242.0990696 | -0.698801887 | 0.354729414 | -1.96995755 | 0.048843236 | 0.254713129 |
| CDH13 | 72.1166529 | -0.970078125 | 0.558884574 | -1.735739667 | 0.082609876 | 0.328745819 |
| CDH17 | 10.24150372 | -0.763343132 | 0.936254032 | -0.815316256 | 0.414891306 | 0.707226517 |
| CDH18 | 39.93978213 | -1.611388916 | 0.635923294 | -2.533935982 | 0.011278934 | 0.120501902 |
| CDH2 | 2735.822367 | 1.038025769 | 0.523801659 | 1.981715314 | 0.047511112 | 0.251059242 |
| CDH23 | 218.6611351 | 1.550755777 | 0.733298521 | 2.11476736 | 0.03444978 | 0.211851485 |
| CDH24 | 879.7121089 | -0.09550689 | 0.444967536 | -0.214637883 | 0.830049656 | 0.93010439 |
| CDH26 | 10.41673945 | -1.076451233 | 1.063478598 | -1.012198304 | 0.311443247 | 0.621452782 |
| CDH3 | 3978.993809 | -1.935483791 | 0.63823943 | -3.032535594 | 0.002425085 | 0.047411036 |
| CDH4 | 82.27955492 | 0.553302439 | 0.927970661 | 0.596249927 | 0.551008277 | 0.794554689 |
| CDH6 | 835.6753344 | -0.330699195 | 0.484571622 | -0.682456793 | 0.494950154 | 0.760514211 |
| CDH7 | 57.69824713 | 0.168883791 | 0.545263556 | 0.309728734 | 0.75676725 | 0.898563152 |
| CDH8 | 341.3379529 | 0.102546949 | 0.406776664 | 0.252096439 | 0.800966522 | 0.916680548 |
| CDH9 | 20.89828968 | -1.149992337 | 1.01495195 | -1.133051015 | 0.257192831 | 0.569767157 |
| CDHR1 | 67.68278075 | -0.778582051 | 0.646115945 | -1.205019094 | 0.228195931 | 0.537150955 |
| CDHR3 | 64.98483352 | -0.640617222 | 0.62185953 | -1.030163872 | 0.302933086 | 0.613538084 |
| CDIP1 | 314.2664385 | 0.00882076 | 2.373110974 | 0.003716961 | 0.997034301 | 0.998614074 |

| | | | | | | |
|---|---|---|---|---|---|---|
| CDIPT | 1432.907141 | 0.253273983 | 0.400702613 | 0.632074698 | 0.527338068 | 0.778906646 |
| CDIPTOSP | 11.72642443 | 0.311898115 | 0.834764679 | 0.373635975 | 0.708675165 | 0.875585221 |
| CDK1 | 2610.839529 | 1.62495091 | 0.309766549 | 5.245727524 | 1.56E-07 | 5.13E-05 |
| CDK10 | 1455.631355 | 0.1899767 | 0.288942662 | 0.657489272 | 0.510866367 | 0.770599585 |
| CDK11A | 973.4795626 | 0.000914069 | 0.353711294 | 0.002584224 | 0.99793809 | 0.998859851 |
| CDK11B | 1660.633913 | 0.113464116 | 0.324444808 | 0.349717773 | 0.726550514 | 0.884630646 |
| CDK12 | 1677.510015 | -0.618701794 | 0.345461818 | -1.790941173 | 0.073302736 | 0.311331411 |
| CDK13 | 1410.911768 | -0.342317157 | 0.240392464 | -1.423992878 | 0.154448534 | 0.451820036 |
| CDK14 | 351.6509708 | 0.385087709 | 0.375515069 | 1.025492026 | 0.305131086 | 0.615135352 |
| CDK16 | 3351.062553 | -0.06308189 | 0.289046006 | -0.218241693 | 0.827240802 | 0.928082319 |
| CDK17 | 856.1355411 | 0.367838587 | 0.289709499 | 1.269680795 | 0.204198357 | 0.510613692 |
| CDK18 | 928.7975827 | -0.237124396 | 0.253428913 | -0.935664336 | 0.349446038 | 0.656038342 |
| CDK19 | 510.6438357 | 0.12854036 | 0.263633675 | 0.487571856 | 0.625853134 | 0.835517239 |
| CDK2 | 1345.366396 | -0.221194618 | 0.315503486 | -0.701084546 | 0.483250254 | 0.754025466 |
| CDK20 | 187.3182288 | 0.228408287 | 0.391740839 | 0.583059675 | 0.559853128 | 0.800219692 |
| CDK2AP1 | 3524.012803 | 0.60852622 | 0.346993663 | 1.753709896 | 0.079480229 | 0.323002882 |
| CDK2AP2 | 469.2505352 | 0.503678362 | 0.382280614 | 1.317561873 | 0.187650356 | 0.492618714 |
| CDK3 | 350.4137197 | -0.30908015 | 0.310403692 | -0.995736061 | 0.319378404 | 0.629115732 |
| CDK4 | 3987.400704 | 0.556189862 | 0.280689713 | 1.981511384 | 0.047533954 | 0.251092485 |
| CDK5 | 410.4600558 | 0.243125347 | 0.373387825 | 0.651133569 | 0.514960269 | 0.773145336 |
| CDK5R1 | 290.5453971 | 1.085275539 | 0.411459803 | 2.637622268 | 0.008348952 | 0.103542973 |
| CDK5RAP1 | 888.9432168 | 0.19185175 | 0.28013658 | 0.684850763 | 0.493438099 | 0.759455148 |
| CDK5RAP2 | 2228.88061 | -0.135983333 | 0.482245562 | -0.281979438 | 0.777959277 | 0.908018171 |
| CDK5RAP3 | 4004.427592 | 0.022524046 | 0.231298986 | 0.097380652 | 0.922424109 | 0.970733641 |
| CDK6 | 725.9772859 | -0.863585771 | 0.458023832 | -1.885460343 | 0.059367714 | 0.282486927 |
| CDK7 | 546.1988219 | 0.676788453 | 0.652100308 | 1.037859429 | 0.299335504 | 0.610246278 |
| CDK8 | 896.6027638 | 0.269576588 | 0.246703391 | 1.092715374 | 0.274518788 | 0.587352209 |
| CDK9 | 1150.85305 | 0.098896042 | 0.333909846 | 0.296175879 | 0.767095769 | 0.903470997 |
| CDKAL1 | 496.9622045 | -0.160019609 | 0.266201164 | -0.601122876 | 0.547758148 | 0.792866984 |
| CDKL1 | 15.79816851 | 0.351109061 | 0.758467939 | 0.46291879 | 0.643422578 | 0.845432525 |
| CDKL2 | 25.80636158 | 0.621884916 | 0.828783876 | 0.750358367 | 0.453038898 | 0.733436323 |
| CDKL3 | 120.9708425 | 0.776366522 | 0.427529409 | 1.815937117 | 0.069380019 | 0.30336211 |
| CDKL5 | 484.6291863 | -0.359932487 | 0.403852747 | -0.891246846 | 0.372796757 | 0.67811865 |
| CDKN1A | 1507.619528 | 0.722927145 | 0.292814918 | 2.468887688 | 0.013553375 | 0.132400677 |
| CDKN1B | 1408.451911 | 0.732323368 | 0.309237238 | 2.368160358 | 0.017876786 | 0.150854036 |
| CDKN1C | 80.14872634 | -0.92098456 | 0.963989832 | -0.955388252 | 0.339381393 | 0.646260212 |
| CDKN2AIP | 869.026301 | 0.291444475 | 0.302074416 | 0.96481019 | 0.334639891 | 0.64263567 |
| CDKN2AIPNL | 1234.983358 | 0.846673764 | 0.346774759 | 2.441566872 | 0.01462368 | 0.137627695 |
| CDKN2C | 17.52504899 | 1.142886312 | 0.774991491 | 1.474708205 | 0.140291009 | 0.429883841 |
| CDKN2D | 33.35295528 | 1.225608437 | 0.558770167 | 2.193403493 | 0.028278321 | |

| | | | | | | |
|---|---|---|---|---|---|---|
| | | | | | | 0.192356209 |
| CDKN3 | 571.1029541 | 1.641899358 | 0.379220731 | 4.329666666 | 1.49E-05 | 0.001490503 |
| CDNF | 32.26074164 | -1.33013445 | 0.633304373 | -2.100308329 | 0.035701727 | 0.216129905 |
| CDO1 | 450.0141286 | 0.436923377 | 0.266765752 | 1.637854089 | 0.101452135 | 0.365328826 |
| CDON | 2189.12023 | -0.057973839 | 0.322852513 | -0.179567563 | 0.857492071 | 0.942343063 |
| CDPF1 | 227.9109385 | 0.854179912 | 0.413070782 | 2.067877831 | 0.038651513 | 0.226576733 |
| CDR2L | 509.1328403 | 0.095705952 | 0.274286507 | 0.348926941 | 0.727144159 | 0.8850111 |
| CDRT4 | 209.9091245 | 1.096917917 | 0.906198025 | 1.210461607 | 0.226101814 | 0.534727976 |
| CDS1 | 560.6984727 | -0.509394396 | 0.469503023 | -1.084965105 | 0.277937111 | 0.591980875 |
| CDS2 | 2144.157619 | -0.41344312 | 0.259291091 | -1.594513405 | 0.110821091 | 0.383119107 |
| CDT1 | 970.1912627 | 0.344771392 | 0.378369393 | 0.911203175 | 0.362188331 | 0.669276392 |
| CDV3 | 6140.220448 | -0.455935099 | 0.226838347 | -2.009956012 | 0.044435845 | 0.242233633 |
| CDX1 | 13.14269779 | -1.488233537 | 1.406098506 | -1.058413426 | 0.289867001 | 0.60258519 |
| CDX2 | 504.2757663 | -0.563413256 | 0.401750635 | -1.402395433 | 0.160797196 | 0.461318885 |
| CDYL | 1855.144933 | 0.305094801 | 0.277892606 | 1.097887435 | 0.272253645 | 0.584457344 |
| CDYL2 | 28.20435421 | 0.070121096 | 0.724919196 | 0.096729535 | 0.922941185 | 0.971075713 |
| CEACAM19 | 34.4030718 | -0.414629729 | 0.56257411 | -0.737022415 | 0.461108718 | 0.73885513 |
| CEACAM6 | 12.52209418 | -1.671152998 | 1.458523347 | -1.145784194 | 0.251884457 | 0.564702097 |
| CEBPA | 294.6704695 | -0.682845276 | 0.859748956 | -0.794237982 | 0.427056901 | 0.714556109 |
| CEBPB | 206.5002459 | 0.669034065 | 0.695089552 | 0.96251492 | 0.335791012 | 0.644086928 |
| CEBPD | 27.85717408 | -0.840769265 | 1.154856742 | -0.728029057 | 0.466595799 | 0.743017201 |
| CEBPG | 1333.605088 | 0.312332959 | 0.237408618 | 1.315592341 | 0.188310906 | 0.493498834 |
| CEBPZ | 2556.604937 | 0.334959128 | 0.241539555 | 1.386767179 | 0.165512768 | 0.467560858 |
| CEBPZOS | 1090.254502 | 0.406020267 | 0.336242962 | 1.207520493 | 0.227231758 | 0.536089352 |
| CECR2 | 3127.437789 | -0.479161844 | 0.414491329 | -1.156023806 | 0.24767142 | 0.560334446 |
| CECR7 | 38.80459629 | 0.120133771 | 0.707593029 | 0.169778059 | 0.865184683 | 0.945248223 |
| CEL | 98.23107547 | -0.127106154 | 0.473935424 | -0.268192982 | 0.788550777 | 0.912309638 |
| CELF1 | 4081.744052 | 0.247541473 | 0.302246933 | 0.819004085 | 0.412784083 | 0.705474445 |
| CELF2 | 754.1535204 | 0.312634704 | 0.375263952 | 0.833106143 | 0.40478487 | 0.70065889 |
| CELF3 | 22.01028288 | -0.592049711 | 0.78069277 | -0.758364538 | 0.448232781 | 0.729628705 |
| CELF4 | 20.2919961 | 0.401741844 | 0.880095878 | 0.456475088 | 0.648048381 | 0.846597949 |
| CELF5 | 128.7645484 | 1.143862923 | 0.407629655 | 2.806132744 | 0.005014003 | 0.07561376 |
| CELP | 31.92223086 | 0.193063553 | 0.645987106 | 0.298865955 | 0.765042324 | 0.902991439 |
| CELSR1 | 1536.565016 | 0.0880972 | 0.428160251 | 0.20575754 | 0.83698031 | 0.933183529 |
| CELSR2 | 1899.425488 | 0.481057385 | 0.542134095 | 0.887340215 | 0.374895761 | 0.67920525 |
| CELSR3 | 947.9993534 | 0.260012494 | 0.313226872 | 0.830109154 | 0.406477072 | 0.701474651 |
| CEMIP2 | 5414.886199 | -0.338758761 | 0.353023466 | -0.959592758 | 0.337260215 | 0.64481093 |
| CEND1 | 24.9382539 | 1.93787135 | 0.775484875 | 2.498915728 | 0.012457393 | 0.126670974 |
| CENPA | 517.8612046 | 1.265089206 | 0.323350888 | 3.912434616 | 9.14E-05 | 0.005775744 |
| CENPB | 1123.677902 | 0.405283977 | 0.362645356 | 1.117576635 | 0.263747853 | 0.577444856 |
| CENPBD1 | 519.3800076 | 0.154485289 | 0.29454095 | 0.524495114 | 0.599934217 | 0.824047262 |
| CENPBD1P1 | 825.7449347 | 0.439347766 | 0.295189591 | 1.488357925 | 0.136656526 | 0.424728769 |
| CENPC | 876.5020286 | 0.107739015 | 0.311631018 | 0.345726223 | 0.729548475 | 0.886372257 |

| | | | | | | |
|---|---|---|---|---|---|---|
| CENPE | 2461.411535 | 1.21870342 | 0.501977753 | 2.427803647 | 0.015190565 | 0.13975504 |
| CENPF | 6370.350609 | 0.640331078 | 0.531903301 | 1.203848663 | 0.228648077 | 0.53738497 |
| CENPH | 2110.266384 | 0.98689308 | 0.369212178 | 2.672970016 | 0.007518297 | 0.096389041 |
| CENPI | 403.8378599 | 0.144725739 | 0.540320771 | 0.267851518 | 0.788813613 | 0.912533261 |
| CENPJ | 985.6694568 | -0.059718859 | 0.435144833 | -0.137239039 | 0.890841854 | 0.957017706 |
| CENPK | 902.189843 | 0.866223753 | 0.263791182 | 3.283747955 | 0.001024365 | 0.027467362 |
| CENPL | 279.9427436 | 0.376814152 | 0.397833339 | 0.947165848 | 0.343554275 | 0.650450756 |
| CENPM | 226.7896791 | 1.458687097 | 0.311393886 | 4.684379368 | 2.81E-06 | 0.000420026 |
| CENPN | 863.4129963 | 0.968200469 | 0.240326876 | 4.028681626 | 5.61E-05 | 0.004071528 |
| CENPO | 1106.816803 | 0.007452225 | 0.467730951 | 0.015932718 | 0.987288068 | 0.995424157 |
| CENPP | 230.5389023 | 0.464017705 | 0.346527288 | 1.33905098 | 0.180554077 | 0.485155137 |
| CENPQ | 600.2226251 | 0.98702036 | 0.261079459 | 3.780536261 | 0.000156491 | 0.008176843 |
| CENPS | 393.1818912 | 0.790643958 | 0.38927112 | 2.031088149 | 0.042246053 | 0.236528876 |
| CENPS-CORT | 27.19176174 | 0.66945467 | 0.641559159 | 1.043480808 | 0.296725665 | 0.607999063 |
| CENPT | 2770.988659 | -1.034183322 | 0.409195485 | -2.52735761 | 0.011492442 | 0.121279761 |
| CENPU | 1344.909022 | 0.580292874 | 0.265441192 | 2.186144768 | 0.028805016 | 0.194174362 |
| CENPV | 3132.255448 | 1.496842601 | 0.398751759 | 3.753820684 | 0.000174159 | 0.008728094 |
| CENPVL3 | 79.63223935 | 1.238686531 | 0.428345371 | 2.891793896 | 0.003830491 | 0.063812059 |
| CENPW | 1181.006654 | 1.142620992 | 0.510698553 | 2.237368767 | 0.025262248 | 0.181849597 |
| CENPX | 732.8180619 | 0.682369618 | 0.3334533 | 2.046372363 | 0.040719749 | 0.23195346 |
| CEP104 | 571.0277076 | -0.478281592 | 0.342778955 | -1.395306173 | 0.162923531 | 0.464171436 |
| CEP112 | 155.1571655 | 1.012695152 | 0.531233924 | 1.906307384 | 0.056610337 | 0.275490291 |
| CEP120 | 728.2673002 | 0.046470938 | 0.357739595 | 0.129901578 | 0.896644295 | 0.959240576 |
| CEP126 | 182.629064 | -0.539439746 | 0.396920528 | -1.359062352 | 0.174126831 | 0.478455902 |
| CEP128 | 307.9951851 | -0.291552259 | 0.42028923 | -0.693694336 | 0.487873919 | 0.75669844 |
| CEP131 | 469.4916169 | -0.009004894 | 0.582430724 | -0.015460884 | 0.987664491 | 0.995541774 |
| CEP135 | 564.5846969 | 0.059189803 | 0.364381403 | 0.162439145 | 0.870960047 | 0.948416227 |
| CEP152 | 519.5420716 | 0.687323603 | 0.448622151 | 1.532076829 | 0.125503477 | 0.407449871 |
| CEP162 | 713.1396773 | -0.341166888 | 0.463156273 | -0.736612905 | 0.461357785 | 0.738942035 |
| CEP164 | 881.2129785 | -0.338303223 | 0.38247222 | -0.884517111 | 0.37641713 | 0.680079109 |
| CEP170 | 1194.954495 | 0.625761609 | 0.313778156 | 1.994280346 | 0.046121426 | 0.247053322 |
| CEP170B | 498.1247703 | -0.959501077 | 0.466785797 | -2.055548995 | 0.039826008 | 0.229888601 |
| CEP19 | 114.0373881 | 0.680435924 | 0.388476186 | 1.751551183 | 0.079851013 | 0.323453279 |
| CEP192 | 1163.966499 | -0.048956974 | 0.465542992 | -0.105161017 | 0.916248043 | 0.969184139 |
| CEP250 | 1034.498419 | -0.156894232 | 0.513600023 | -0.30547941 | 0.760001049 | 0.899725004 |
| CEP290 | 1072.744927 | 0.078688028 | 0.354705608 | 0.221840384 | 0.824438141 | 0.92681808 |
| CEP295 | 591.2535005 | -0.028251931 | 0.397878524 | -0.071006425 | 0.943392642 | 0.978679552 |
| CEP350 | 1023.87692 | -0.300283468 | 0.377422436 | -0.795616368 | 0.42625505 | 0.714238498 |
| CEP41 | 448.6242495 | 0.497350231 | 0.293431912 | 1.69494254 | 0.090086322 | 0.343910313 |
| CEP44 | 536.5065669 | 0.136862642 | 0.260913175 | 0.524552437 | 0.599894358 | 0.824047262 |
| CEP55 | 616.7055002 | 0.829272302 | 0.431292097 | 1.922762573 | 0.054509874 | 0.269872826 |
| CEP57 | 2892.577505 | 0.085542669 | 0.266085025 | 0.321486219 | 0.747841954 | 0.894202292 |
| CEP57L1 | 414.7321143 | 0.648125759 | 0.340003156 | 1.906234538 | 0.056619783 | |

| | | | | | | |
|---|---|---|---|---|---|---|
| | | | | | | 0.275490291 |
| CEP63 | 760.9113052 | -0.004652788 | 0.238805893 | -0.019483558 | 0.984455353 | 0.994286144 |
| CEP68 | 1111.848015 | 0.323557452 | 0.409137937 | 0.790827304 | 0.429044775 | 0.715829569 |
| CEP70 | 1305.904278 | 0.564956696 | 0.261908234 | 2.15707879 | 0.030999526 | 0.200870165 |
| CEP72 | 322.8545041 | 0.280017637 | 0.313604696 | 0.892900013 | 0.37191072 | 0.677193964 |
| CEP76 | 458.022171 | 0.109103617 | 0.279834845 | 0.389885744 | 0.696621036 | 0.870023227 |
| CEP78 | 2265.613283 | -0.076315978 | 0.252035099 | -0.302799006 | 0.762043041 | 0.900877034 |
| CEP83 | 662.8391829 | 0.135934471 | 0.361413118 | 0.376119363 | 0.70682816 | 0.875084872 |
| CEP85 | 1070.072292 | 0.64666104 | 0.546371716 | 1.183555116 | 0.236589217 | 0.547336452 |
| CEP85L | 340.2347609 | -0.350961843 | 0.432008569 | -0.81239556 | 0.416564693 | 0.708268589 |
| CEP89 | 738.1402245 | -0.002443381 | 0.238057618 | -0.010263822 | 0.991810798 | 0.99688215 |
| CEP95 | 1438.493317 | -0.325840826 | 0.239257096 | -1.361885736 | 0.173233948 | 0.4771935 |
| CEP97 | 566.1428892 | 0.312328408 | 0.438913635 | 0.711594226 | 0.476716084 | 0.749746108 |
| CEPT1 | 1059.941685 | 0.369891227 | 0.248112101 | 1.490822998 | 0.136007972 | 0.423778383 |
| CERCAM | 890.4583255 | -0.22404739 | 0.361303821 | -0.620108002 | 0.535186685 | 0.784193183 |
| CERK | 2499.239021 | 0.34853761 | 0.335252777 | 1.039626318 | 0.298513545 | 0.609920825 |
| CERS1 | 300.05047 | 0.822423376 | 0.297877313 | 2.760946668 | 0.005763409 | 0.081869544 |
| CERS2 | 4202.449668 | 0.076763384 | 0.221328293 | 0.346830415 | 0.728718726 | 0.885848701 |
| CERS3 | 27.10111911 | 1.061500447 | 0.851685943 | 1.246351962 | 0.212635209 | 0.519735803 |
| CERS4 | 1220.209753 | 0.037962391 | 0.360290984 | 0.10536592 | 0.916085457 | 0.969184139 |
| CERS5 | 1319.747234 | 0.186751613 | 0.250866509 | 0.744426244 | 0.456618647 | 0.735622968 |
| CERS6 | 1750.56069 | -0.173461769 | 0.353659371 | -0.490476947 | 0.623796438 | 0.834534018 |
| CES2 | 794.3923026 | 0.001489464 | 0.28097079 | 0.005301134 | 0.995770327 | 0.998171604 |
| CES3 | 148.768961 | -0.93999761 | 0.408170859 | -2.302951299 | 0.021281583 | 0.165485854 |
| CES4A | 22.97515187 | -0.760744542 | 0.879894327 | -0.864586255 | 0.387265919 | 0.68748084 |
| CETN2 | 1159.777235 | 0.493132404 | 0.375754564 | 1.312379011 | 0.189392285 | 0.494879496 |
| CETN3 | 1361.769244 | 1.239966906 | 0.353984114 | 3.50288857 | 0.000460242 | 0.016624597 |
| CETN4P | 21.0530044 | 1.014378518 | 0.659133654 | 1.538957254 | 0.123814732 | 0.405045568 |
| CFAP20 | 1048.693046 | 0.829508343 | 0.331490931 | 2.502356071 | 0.012336978 | 0.126347892 |
| CFAP206 | 29.54173069 | -1.317373956 | 0.739043569 | -1.782538962 | 0.074661384 | 0.313779586 |
| CFAP221 | 28.87321847 | 0.49482914 | 0.580192362 | 0.852870828 | 0.393730941 | 0.692653237 |
| CFAP298 | 1236.509124 | 0.534285526 | 0.415002171 | 1.287428266 | 0.197945058 | 0.503609671 |
| CFAP300 | 153.050953 | 0.598369096 | 0.365442346 | 1.637383031 | 0.101550461 | 0.36559612 |
| CFAP36 | 774.2792677 | 0.606719219 | 0.33124907 | 1.831610333 | 0.067009496 | 0.297817842 |
| CFAP410 | 335.0439484 | 0.184603175 | 0.449829126 | 0.410385109 | 0.681523468 | 0.862590493 |
| CFAP43 | 30.6654237 | -0.334963377 | 0.569916289 | -0.587741364 | 0.556705907 | 0.797750573 |
| CFAP44 | 54.60285584 | -1.857265514 | 0.600422075 | -3.093266537 | 0.001979662 | 0.041771153 |
| CFAP45 | 35.60974374 | -1.155818573 | 0.988224503 | -1.169591089 | 0.242165565 | 0.553231259 |
| CFAP46 | 45.59876688 | 0.078519495 | 0.559395637 | 0.140364868 | 0.888371714 | 0.955943827 |
| CFAP47 | 14.73745795 | -1.671120372 | 0.917613126 | -1.821160057 | 0.06858253 | 0.301671663 |
| CFAP53 | 28.56292118 | -0.67081916 | 0.710499764 | -0.944151136 | 0.345092421 | 0.651979716 |

| | | | | | | |
|---|---|---|---|---|---|---|
| CFAP54 | 50.75318951 | -0.18997356 | 0.521028927 | -0.364612308 | 0.715400826 | 0.879026964 |
| CFAP57 | 21.57859904 | -0.166454491 | 0.851041348 | -0.195589193 | 0.84493172 | 0.937501581 |
| CFAP58 | 21.7891521 | 0.087839547 | 0.688030945 | 0.127668018 | 0.898411701 | 0.960048173 |
| CFAP65 | 30.71662083 | -0.819215898 | 0.607346961 | -1.348843332 | 0.177387293 | 0.481794969 |
| CFAP69 | 76.68913186 | -0.292930217 | 0.394672783 | -0.742210331 | 0.45795991 | 0.736518606 |
| CFAP70 | 33.35294635 | -0.741537273 | 0.671309741 | -1.104612712 | 0.26932744 | 0.582188661 |
| CFAP97 | 1407.373078 | 0.219032163 | 0.271188189 | 0.807675895 | 0.419277183 | 0.709924862 |
| CFC1 | 133.4127035 | -2.236659692 | 0.687440191 | -3.253606235 | 0.001139501 | 0.029400291 |
| CFC1B | 10.87002907 | -0.76485671 | 1.953996441 | -0.391431987 | 0.695477952 | 0.869563155 |
| CFDP1 | 3242.415828 | 0.676363056 | 0.310073589 | 2.181298507 | 0.029161345 | 0.195495697 |
| CFL1 | 17559.05288 | 0.32357008 | 0.379427682 | 0.85278459 | 0.393778771 | 0.692653237 |
| CFL2 | 732.3391778 | 0.499427721 | 0.241641054 | 2.066816513 | 0.038751447 | 0.226725106 |
| CFLAR | 703.491413 | -0.165941138 | 0.262264007 | -0.632725549 | 0.526912882 | 0.778755887 |
| CFP | 16.5318282 | -0.216103314 | 0.743513875 | -0.290651353 | 0.771317981 | 0.905067213 |
| CFTR | 34.26604325 | -2.62035898 | 1.043523296 | -2.511068981 | 0.012036617 | 0.124561748 |
| CGAS | 15.97041282 | 0.97125708 | 0.737322131 | 1.317276451 | 0.187745976 | 0.492689771 |
| CGB7 | 50.05557725 | -1.946849155 | 0.593693637 | -3.279215126 | 0.001040962 | 0.027754731 |
| CGGBP1 | 3238.111567 | 0.023407815 | 0.222987318 | 0.10497375 | 0.916396638 | 0.969211928 |
| CGN | 1867.045579 | -1.458109471 | 0.37851469 | -3.852187267 | 0.000117067 | 0.006837179 |
| CGNL1 | 853.6240235 | -1.540596932 | 0.285762777 | -5.391174276 | 7.00E-08 | 3.12E-05 |
| CGREF1 | 40.01079523 | 0.679202786 | 0.558785505 | 1.215498219 | 0.224176128 | 0.533486437 |
| CGRRF1 | 454.4214611 | 0.742031811 | 0.324290237 | 2.288171908 | 0.022127512 | 0.169031462 |
| CHAC1 | 104.613747 | 0.077034975 | 0.679815678 | 0.113317444 | 0.909778888 | 0.96630013 |
| CHAC2 | 83.98468348 | 1.443555807 | 0.400756838 | 3.602074052 | 0.000315688 | 0.013085543 |
| CHAF1A | 1569.170102 | 0.100962083 | 0.285204681 | 0.353998689 | 0.723339866 | 0.883273431 |
| CHAF1B | 674.4943315 | 0.117821232 | 0.257072355 | 0.45831934 | 0.646723032 | 0.84654315 |
| CHAMP1 | 1253.403527 | -0.082222083 | 0.30150196 | -0.272708286 | 0.785077465 | 0.910931322 |
| CHCHD1 | 1178.179577 | 1.267151305 | 0.440678467 | 2.875455462 | 0.004034451 | 0.065655774 |
| CHCHD10 | 124.8160698 | 1.274504947 | 0.521763744 | 2.44268591 | 0.014578419 | 0.13754303 |
| CHCHD3 | 2371.342677 | 0.438025955 | 0.246149805 | 1.779509657 | 0.075156243 | 0.314600339 |
| CHCHD4 | 235.9856505 | 0.333316153 | 0.370262647 | 0.900215443 | 0.36800561 | 0.674179083 |
| CHCHD5 | 432.6893505 | 1.005752841 | 0.509911085 | 1.972408269 | 0.048563019 | 0.25374113 |
| CHCHD6 | 352.9300725 | 0.879473403 | 0.366076765 | 2.402428911 | 0.0162866 | 0.144292038 |
| CHCHD7 | 846.0112155 | 0.486596281 | 0.338348099 | 1.438152844 | 0.150390692 | 0.445185792 |
| CHD1 | 2287.128529 | 0.2546167 | 0.371559283 | 0.685265346 | 0.493176495 | 0.759389046 |
| CHD1L | 1932.891672 | -0.029662276 | 0.36147609 | -0.08205875 | 0.934599995 | |

| | | | | | | |
|---|---|---|---|---|---|---|
| | | | | | | 0.975099718 |
| CHD2 | 3191.634714 | -0.461777221 | 0.26522469 | -1.741079312 | 0.081669677 | 0.326743321 |
| CHD3 | 2088.844882 | -0.793191103 | 0.316973386 | -2.502390227 | 0.012335787 | 0.126347892 |
| CHD4 | 15140.21801 | -0.200382048 | 0.261174542 | -0.767234228 | 0.442942266 | 0.726237665 |
| CHD5 | 24.00939262 | -0.456305844 | 0.69645071 | -0.655187563 | 0.512347001 | 0.771687791 |
| CHD6 | 1922.133842 | -0.323582907 | 0.386989028 | -0.836155251 | 0.403067569 | 0.699766232 |
| CHD7 | 5801.881483 | -0.072304463 | 0.315978144 | -0.228827417 | 0.819003056 | 0.924316427 |
| CHD8 | 3533.376488 | -0.529768207 | 0.40501332 | -1.308026627 | 0.190864278 | 0.496058205 |
| CHD9 | 1396.93181 | 0.101828553 | 0.312041627 | 0.326330029 | 0.744174671 | 0.892691858 |
| CHDH | 160.1596474 | -0.097612071 | 0.363005046 | -0.268900039 | 0.788006607 | 0.912309638 |
| CHEK1 | 1722.464735 | 0.126463351 | 0.256634916 | 0.492775312 | 0.622171357 | 0.833640266 |
| CHEK2 | 828.8750291 | 0.827231011 | 0.318282274 | 2.599048324 | 0.009348261 | 0.109346547 |
| CHERP | 1646.35895 | -0.160527954 | 0.324915324 | -0.49406089 | 0.621263177 | 0.833349572 |
| CHFR | 1199.96092 | -0.39545528 | 0.305458969 | -1.294626513 | 0.195449102 | 0.50126155 |
| CHGB | 44.5754039 | -1.008822145 | 0.525980306 | -1.917984634 | 0.05511296 | 0.271793713 |
| CHIC1 | 457.4617541 | 0.303079251 | 0.327392613 | 0.925736376 | 0.354582999 | 0.661914443 |
| CHIC2 | 114.7758251 | -0.107372461 | 0.530171644 | -0.20252396 | 0.839507129 | 0.934215701 |
| CHID1 | 1742.161543 | 0.598873857 | 0.347089084 | 1.725418296 | 0.084452116 | 0.332698795 |
| CHKA | 1199.702949 | -0.462222465 | 0.312952664 | -1.476972457 | 0.139683021 | 0.428626842 |
| CHKB | 1042.948599 | -0.189746392 | 0.244600427 | -0.775740231 | 0.437902381 | 0.721875151 |
| CHKB-CPT1B | 339.8549779 | -0.408813501 | 0.568203274 | -0.719484592 | 0.471842393 | 0.746399031 |
| CHM | 590.4303716 | -0.361759877 | 0.300762681 | -1.202808394 | 0.229050476 | 0.537758004 |
| CHML | 1376.978859 | 0.06750431 | 0.373532613 | 0.180718653 | 0.856588417 | 0.942343063 |
| CHMP1A | 1612.71993 | -0.130843893 | 0.323491674 | -0.404473758 | 0.685864371 | 0.865013582 |
| CHMP1B | 2157.268959 | 0.139308796 | 0.404163946 | 0.344683877 | 0.730332041 | 0.886744089 |
| CHMP1B2P | 154.2199289 | -1.231003597 | 0.390208245 | -3.154734973 | 0.00160644 | 0.037175747 |
| CHMP2A | 2094.189532 | 0.771031777 | 0.462541438 | 1.666946385 | 0.095525066 | 0.354819129 |
| CHMP2B | 1389.723855 | 0.353005122 | 0.31753933 | 1.111689446 | 0.266271699 | 0.57970088 |
| CHMP3 | 1249.513181 | -0.193591985 | 0.314722749 | -0.615119133 | 0.538476047 | 0.786560766 |
| CHMP4A | 494.3242567 | 0.570263023 | 0.530013555 | 1.075940449 | 0.28195389 | 0.595751624 |
| CHMP4B | 2142.292609 | -0.139793257 | 0.377119861 | -0.37068654 | 0.710871017 | 0.876976263 |
| CHMP4C | 224.2441175 | -0.4048281 | 0.616123018 | -0.657057257 | 0.511144101 | 0.770636467 |
| CHMP5 | 2414.725018 | 0.627976487 | 0.32212872 | 1.949458238 | 0.051240725 | 0.260426481 |
| CHMP6 | 370.5875737 | 0.218919474 | 0.423720251 | 0.516660399 | 0.605393245 | 0.826077544 |
| CHMP7 | 1225.70917 | 0.267956276 | 0.228455846 | 1.172901816 | 0.240835181 | 0.552437022 |
| CHN1 | 874.8768143 | -0.15610462 | 0.289893683 | -0.538489208 | 0.590239354 | 0.818321084 |
| CHN2 | 176.7202574 | -0.491708473 | 0.525709824 | -0.935322968 | 0.349621882 | 0.656273207 |

| | | | | | | |
|---|---|---|---|---|---|---|
| CHODL | 131.7355484 | 0.355204477 | 0.607733298 | 0.584474272 | 0.55890127 | 0.799176463 |
| CHORDC1 | 771.3961803 | 0.499638612 | 0.278805383 | 1.792069459 | 0.07312184 | 0.311055732 |
| CHP1 | 4182.980589 | -0.179510027 | 0.348234028 | -0.51548675 | 0.606212926 | 0.826611695 |
| CHP2 | 14.34890112 | -0.083726119 | 0.840848817 | -0.099573333 | 0.920683066 | 0.969843271 |
| CHPF | 344.287124 | -0.264799693 | 0.493765121 | -0.536286753 | 0.591760385 | 0.819494655 |
| CHPF2 | 569.0187981 | -0.631653273 | 0.314365344 | -2.009296779 | 0.044505669 | 0.242353018 |
| CHPT1 | 333.1532726 | 0.512641054 | 0.29340157 | 1.747233505 | 0.080596842 | 0.325022513 |
| CHRAC1 | 504.2616709 | 0.065002318 | 0.277370246 | 0.234352166 | 0.81471159 | 0.922623448 |
| CHRD | 184.3116976 | 1.09303281 | 0.463973271 | 2.355809868 | 0.018482375 | 0.154148494 |
| CHRDL1 | 136.4415255 | 0.732311017 | 0.584445716 | 1.253000915 | 0.210205374 | 0.51816316 |
| CHRM3 | 366.7547963 | -0.176287795 | 0.327349398 | -0.538530987 | 0.590210518 | 0.818321084 |
| CHRNA10 | 10.6217128 | -0.762442842 | 1.085518594 | -0.702376584 | 0.482444344 | 0.753310328 |
| CHRNA3 | 20.38610177 | 0.982695948 | 0.738781556 | 1.330157663 | 0.18346633 | 0.487796651 |
| CHRNA5 | 404.6778677 | 1.432705582 | 0.315348571 | 4.543244251 | 5.54E-06 | 0.000693377 |
| CHRNA7 | 90.59635828 | -0.084434834 | 0.538151168 | -0.156897984 | 0.875325248 | 0.949829007 |
| CHRNA9 | 27.52665631 | 1.22575495 | 0.708199174 | 1.730805394 | 0.08348648 | 0.330745685 |
| CHRNB1 | 267.8908973 | -0.315227361 | 0.294799281 | -1.06929488 | 0.284936817 | 0.598807847 |
| CHRNB2 | 16.38098217 | -0.989757651 | 0.924500282 | -1.07058664 | 0.284355333 | 0.598246396 |
| CHRNB4 | 13.21580079 | 0.347649684 | 1.094588123 | 0.317607762 | 0.750782486 | 0.896207755 |
| CHRNE | 16.76749633 | -1.67885787 | 0.821641569 | -2.043297142 | 0.041023032 | 0.232934829 |
| CHRNG | 82.01486503 | -0.871520442 | 0.804118821 | -1.083820475 | 0.278444405 | 0.592005173 |
| CHST1 | 11.2441338 | -0.556913388 | 1.067186555 | -0.521851953 | 0.601773414 | 0.825021776 |
| CHST10 | 600.1939062 | 0.307918447 | 0.516219527 | 0.596487406 | 0.550849664 | 0.794461475 |
| CHST11 | 607.9575769 | 0.568162413 | 0.324851256 | 1.748992507 | 0.080292315 | 0.324292679 |
| CHST12 | 281.1616838 | 0.264447894 | 0.302336651 | 0.874680237 | 0.381747916 | 0.683201326 |
| CHST14 | 598.9776352 | 0.214399944 | 0.379422625 | 0.565068949 | 0.572026868 | 0.807351345 |
| CHST15 | 487.3026783 | -1.167155329 | 0.446633581 | -2.613227889 | 0.008969147 | 0.106919917 |
| CHST2 | 109.9379314 | 0.984271027 | 0.516949941 | 1.903996789 | 0.056910602 | 0.2761211 |
| CHST3 | 411.2075406 | -0.685772496 | 0.334699373 | -2.04892077 | 0.040469863 | 0.231601821 |
| CHST4 | 16.04680176 | -0.519346274 | 1.01491223 | -0.511715455 | 0.608850168 | 0.828009963 |
| CHST6 | 572.6572363 | 0.27667998 | 0.427749473 | 0.646827166 | 0.517743808 | 0.773950969 |
| CHST7 | 109.6820254 | 0.640648948 | 0.602746455 | 1.06288298 | 0.287835022 | 0.601673803 |
| CHST8 | 38.79197004 | -0.023407157 | 0.565230411 | -0.041411708 | 0.966967679 | 0.988749915 |
| CHST9 | 41.86560902 | 0.2320491 | 0.542215434 | 0.427964764 | 0.668676774 | 0.857305446 |
| CHSY1 | 1379.764866 | -0.24634922 | 0.244953932 | -1.005696126 | 0.314561767 | 0.624045314 |
| CHSY3 | 11.44483873 | 0.252494233 | 0.871478236 | 0.289730968 | 0.772022063 | 0.905547141 |

| | | | | | | |
|---|---|---|---|---|---|---|
| CHTF18 | 1051.257858 | 0.156927754 | 0.595430202 | 0.263553568 | 0.792123941 | 0.913570955 |
| CHTF8 | 1782.088668 | -0.221149561 | 0.375068111 | -0.589625069 | 0.555442041 | 0.796360966 |
| CHTOP | 2945.27242 | 0.337251872 | 0.292522987 | 1.152907249 | 0.248948457 | 0.56168903 |
| CHUK | 807.7474891 | 0.293754343 | 0.29918576 | 0.981846002 | 0.326175721 | 0.635960913 |
| CHURC1 | 1336.349488 | 0.374134407 | 0.322330615 | 1.160716324 | 0.245757281 | 0.557557008 |
| CHURC1-FNTB | 11.73455433 | 0.247958062 | 1.401054652 | 0.176979579 | 0.85952443 | 0.943071175 |
| CIAO1 | 1857.453489 | -0.01803562 | 0.236706813 | -0.076193921 | 0.939264819 | 0.977796709 |
| CIAO2A | 1510.774364 | 1.012133673 | 0.358683301 | 2.821803163 | 0.004775448 | 0.073402552 |
| CIAO2B | 1043.658772 | 0.864534616 | 0.406857556 | 2.124907361 | 0.033594344 | 0.208960645 |
| CIAO3 | 507.9749411 | -0.084170264 | 0.461455261 | -0.182401786 | 0.855267425 | 0.941874426 |
| CIAPIN1 | 2119.159802 | 0.160693777 | 0.224183824 | 0.7167947 | 0.473500782 | 0.747811725 |
| CIART | 239.7442481 | 0.832965506 | 0.489733855 | 1.700853426 | 0.088970514 | 0.342496199 |
| CIB1 | 258.5513698 | 0.494845972 | 0.37483235 | 1.320179467 | 0.186775106 | 0.491426488 |
| CIB2 | 854.5786486 | 0.256137288 | 0.341964678 | 0.749016797 | 0.453847082 | 0.73365094 |
| CIC | 988.5038646 | 0.123659611 | 0.502787328 | 0.245948145 | 0.805722374 | 0.918930547 |
| CICP18 | 10.9086356 | -0.442760747 | 1.184391408 | -0.373829753 | 0.708530982 | 0.875585221 |
| CIDECP1 | 143.743547 | 0.007320676 | 0.380162066 | 0.019256725 | 0.984636306 | 0.994402703 |
| CIITA | 17.15920977 | -0.821990614 | 0.783488888 | -1.049141382 | 0.294113053 | 0.606454309 |
| CILP2 | 32.06801712 | 0.734549192 | 0.666777896 | 1.101639987 | 0.270618217 | 0.583257419 |
| CINP | 632.0883572 | 0.645414248 | 0.332796573 | 1.939365671 | 0.052456828 | 0.264480735 |
| CIP2A | 713.3247083 | 0.360438302 | 0.412217421 | 0.874388816 | 0.381906546 | 0.683324001 |
| CIPC | 1146.544269 | -0.037630754 | 0.435299188 | -0.086448022 | 0.931110274 | 0.974131023 |
| CIR1 | 1074.793398 | 0.196506239 | 0.500499387 | 0.392620339 | 0.694599915 | 0.86892666 |
| CIRBP | 7746.228521 | 0.128401252 | 0.360327368 | 0.3563461 | 0.72158139 | 0.882498649 |
| CISD1 | 808.1284607 | 0.824287626 | 0.402994864 | 2.045404792 | 0.040814967 | 0.232172427 |
| CISD2 | 1123.413265 | 0.823254319 | 0.286380878 | 2.874683274 | 0.00404433 | 0.065655774 |
| CISD3 | 53.12987604 | 0.410257146 | 0.499886436 | 0.820700695 | 0.411816775 | 0.704836675 |
| CISH | 42.8546564 | 0.151773148 | 0.532035356 | 0.285268915 | 0.775438126 | 0.906749793 |
| CIT | 1477.574443 | 0.097704479 | 0.46755958 | 0.208966906 | 0.834474074 | 0.932107564 |
| CITED1 | 17.15212637 | 1.778201889 | 0.962246859 | 1.847968504 | 0.064606899 | 0.293546352 |
| CITED2 | 5649.809059 | 0.466625986 | 0.277332829 | 1.68254868 | 0.092462494 | 0.34894241 |
| CITED4 | 137.7253103 | 0.523389381 | 0.636536662 | 0.822245461 | 0.41093721 | 0.704426187 |
| CIZ1 | 2353.381873 | -0.558327123 | 0.338280618 | -1.650485108 | 0.098843757 | 0.361016186 |
| CKAP2 | 3075.725001 | 0.644537632 | 0.340777312 | 1.891374834 | 0.058574325 | 0.280288923 |
| CKAP2L | 532.4906212 | 0.495083156 | 0.482367201 | 1.026361566 | 0.304721187 | 0.614996026 |
| CKAP4 | 3922.217273 | 0.639731044 | 0.235378668 | 2.717880299 | 0.00657016 | 0.088793705 |
| CKAP5 | 6520.126299 | 0.227640637 | 0.500656912 | 0.4546839 | 0.649336666 | 0.847281162 |
| CKB | 5277.04661 | 1.626500243 | 0.423400362 | 3.841518305 | 0.000122276 | 0.007000164 |
| CKLF | 217.7782205 | 0.78344088 | 0.501848286 | 1.561110999 | 0.118497563 | 0.396849124 |
| CKMT1A | 349.7048845 | -0.08836856 | 0.302455928 | -0.292170037 | 0.77015662 | 0.90489824 |
| CKMT1B | 292.5897518 | 0.077047318 | 0.396126112 | 0.194501991 | 0.845782837 | 0.937897187 |
| CKS1B | 3710.803323 | 1.50339215 | 0.409735418 | 3.669177928 | 0.000243332 | 0.010669321 |
| CKS2 | 3552.679503 | 1.770171563 | 0.340756858 | 5.194823004 | 2.05E-07 | 6.61E-05 |
| CLASP1 | 2032.649392 | -0.667822825 | 0.47708511 | -1.399798089 | | |

0.16157379 0.461974363
CLASP2 1617.217107 0.117643848 0.340833643 0.345165011 0.729970322 0.886517751
CLASRP 1148.752785 -0.127539416 0.340941358 -0.374080214 0.708344639 0.875585221
CLBA1 182.102056 -0.156404208 0.430673824 -0.36316163 0.716484149 0.879930464
CLCA2 39.94304619 -1.19028811 0.709896881 -1.676705648 0.093600056 0.351044935
CLCC1 674.3903933 -0.325848476 0.270658905 -1.203908203 0.228625061 0.53738497
CLCF1 59.66226409 -0.928129665 0.606846613 -1.529430411 0.126157778 0.408874098
CLCN1 11.20168486 -2.755826554 1.310167051 -2.103416166 0.035429407 0.215430672
CLCN2 489.2489956 1.025222078 0.477500876 2.147058005 0.031788664 0.203069232
CLCN3 2054.625795 -0.504083121 0.267902559 -1.881591288 0.05989153 0.283057446
CLCN3P1 81.67596273 -0.274175655 0.462445095 -0.592882609 0.553259718 0.79574357
CLCN4 212.9680227 1.040318576 0.617055966 1.685938768 0.091807594 0.347768541
CLCN5 1283.0203 -1.435511502 0.503015412 -2.853812163 0.004319806 0.068337626
CLCN6 958.3687824 -0.378077308 0.393038442 -0.961934679 0.336082417 0.64425371
CLCN7 1026.261958 -0.300573232 0.431601277 -0.69641414 0.48616951 0.756169535
CLCNKA 16.78093669 -0.34291557 0.725850866 -0.472432542 0.636618081 0.841268221
CLCNKB 169.857515 -0.486202046 0.369203258 -1.31689533 0.187873712 0.492689771
CLDN1 12.62891115 -2.686983114 1.279131028 -2.100631644 0.035673314 0.216083122
CLDN10 394.4119165 0.052456754 0.460378854 0.113942578 0.909283313 0.966018007
CLDN11 47.24877758 1.985590747 0.553702696 3.58602326 0.000335759 0.013547304
CLDN12 1749.062365 0.173033085 0.279046882 0.620086072 0.535201122 0.784193183
CLDN15 171.6854085 -0.697423527 0.537321098 -1.297964158 0.194299651 0.500187881
CLDN19 185.1850239 -0.341519254 0.50279665 -0.679239319 0.496986238 0.761325016
CLDN23 54.64026641 -0.680573753 0.686522592 -0.991334824 0.321522114 0.631253641
CLDN3 139.1920038 -0.519318607 0.799989711 -0.649156608 0.516237155 0.773896629
CLDN4 6508.462823 -0.013165611 0.783721877 -0.016798831 0.986597102 0.995107011
CLDN6 16396.05519 -0.475379201 0.596651671 -0.796744942 0.425599177 0.713770188
CLDN7 2013.639448 -0.720899619 0.782248715 -0.921573413 0.356751132 0.664245388
CLDN9 53.27257228 -0.947544629 0.783430811 -1.209480934 0.226478131 0.535186874
CLDND1 1899.518902 0.183089038 0.228608787 0.800883644 0.42319901 0.712580708
CLDND2 14.95144777 2.043327575 1.179470352 1.732411139 0.083200386 0.330254594
CLEC10A 32.60936001 1.999759947 0.939081479 2.129485024 0.033214153 0.207932851
CLEC16A 612.8114078 -0.555100279 0.344100046 -1.613194434 0.106702272 0.374599763
CLEC18A 46.41301609 1.231257929 0.545782828 2.255948457 0.024073859

0.177207433
CLEC18B 92.05877325 2.228353704 0.816415038 2.729437357 0.00634425
0.086476746
CLEC18C 25.16610543 1.485192077 1.269340441 1.170050231 0.241980755
0.553231259
CLEC19A 14.99049853 1.498631362 1.059492897 1.414479857 0.157221058
0.456574487
CLEC2D 259.4643391 -0.593146548 0.538087205 -1.102324202
0.270320751 0.583170438
CLEC4A 10.17404571 1.053052086 0.877938576 1.199459865 0.230349182
0.539112606
CLEC4E 11.44973382 0.107753584 1.016830565 0.105970048 0.915606116
0.969158739
CLEC4GP1 29.76277579 0.113838533 1.012203368 0.112466069 0.910453872
0.966517087
CLGN 149.7377872 2.497930739 0.760852464 3.283068476 0.001026837 0.027474686
CLHC1 180.8861153 -0.025857827 0.329812525 -0.078401592 0.937508607
0.976716322
CLIC1 1051.029943 0.280201248 0.316763576 0.884575339 0.376385712 0.680079109
CLIC2 14.60810115 0.49403013 0.842144352 0.586633548 0.557449846 0.798288807
CLIC4 10383.22724 -0.369835161 0.231524013 -1.597394399 0.11017782
0.381884329
CLIC5 38.88528797 -3.734477148 0.980861779 -3.807342918 0.000140468
0.007590614
CLIC6 100.7043975 0.767294312 0.705811159 1.08710992 0.276988234 0.590738803
CLINT1 2924.283864 -0.269138266 0.224826744 -1.197091864
0.231270751 0.540284656
CLIP1 2277.665201 -1.308245705 0.506523611 -2.582793136 0.009800407
0.112404636
CLIP2 649.4281971 0.200633664 0.477341969 0.420314317 0.674255852 0.859672768
CLIP3 1288.865621 -0.209684142 0.4242163 -0.494285915 0.621104271
0.833349572
CLIP4 25.92161409 -0.375152735 0.644192738 -0.582361011 0.560323538
0.800514963
CLK1 1365.455178 0.191871119 0.278243812 0.689579103 0.490458912 0.758593654
CLK2 1026.407802 -0.229264364 0.306007812 -0.749210819 0.453730149
0.733540078
CLK3 1094.121688 0.272639749 0.387132105 0.704255074 0.481273939 0.752722364
CLK4 369.1488292 -0.176553101 0.403807282 -0.437221191 0.661950943
0.853756491
CLMN 247.7134951 -0.357795685 0.373042713 -0.95912793 0.337494302
0.645014619
CLMP 173.5404588 0.378647494 0.486122921 0.778913065 0.436030927 0.720784523
CLN3 997.3603811 -0.237965163 0.330794893 -0.719373749 0.471910667
0.746399031
CLN5 500.8006742 0.115238713 0.276635817 0.416571917 0.676991554 0.86029961
CLN6 1158.682058 -0.152995486 0.34804856 -0.439580862 0.660240704
0.852284938
CLN8 154.6852781 0.478988183 0.49432456 0.968975086 0.33255762 0.640257824
CLNS1A 4038.712537 0.671983009 0.415368927 1.617797974 0.105706143
0.372686009
CLOCK 593.1710566 0.205538349 0.306072914 0.671533938 0.501880447 0.764473972
CLP1 568.2113871 0.119825043 0.332209419 0.360691286 0.71833024 0.880771685
CLPB 703.9024469 -0.265063694 0.30793387 -0.860781225 0.389358547
0.689219487
CLPP 852.6554248 0.728167493 0.416323704 1.749041638 0.080283822 0.324292679
CLPTM1 1408.864256 -0.811518235 0.408194675 -1.988066687
0.046804315 0.249225774
CLPTM1L 1684.543468 0.06722403 0.312292567 0.215259783 0.829564783
0.929835784
CLPX 1640.807146 -0.105446971 0.248763889 -0.423883751 0.671650595
0.858578629
CLSPN 1027.459284 0.357534696 0.307048561 1.164423942 0.24425225 0.555471576
CLSTN1 5001.590185 -0.485329332 0.357451366 -1.357749271

| | | | | | | |
|---|---|---|---|---|---|---|
| | | | | | 0.174543257 | 0.478578744 |
| CLSTN2 | 61.34910834 | -0.260587472 | 0.469759865 | -0.554724853 | 0.579082874 | 0.810713649 |
| CLSTN3 | 1048.752581 | -0.548503109 | 0.399654348 | -1.372443742 | 0.169925341 | 0.472495848 |
| CLTA | 4359.274308 | 0.257971149 | 0.360570496 | 0.715452738 | 0.474329335 | 0.748263527 |
| CLTB | 385.5441816 | 0.272592084 | 0.405351435 | 0.672483331 | 0.501276048 | 0.763991735 |
| CLTC | 11231.03353 | -0.437476094 | 0.272122441 | -1.607644313 | 0.107913101 | 0.376615979 |
| CLTCL1 | 417.5958256 | -0.320384624 | 0.435897583 | -0.734999772 | 0.462339633 | 0.73986733 |
| CLTRN | 73.7238497 | 0.777805383 | 0.587081364 | 1.32486812 | 0.185214896 | 0.489784764 |
| CLU | 5958.056526 | 0.48104195 | 0.376808651 | 1.276621302 | 0.201735977 | 0.508310333 |
| CLUAP1 | 574.942706 | 0.117667635 | 0.307223922 | 0.38300284 | 0.701717658 | 0.872479299 |
| CLUH | 1142.841155 | -0.396322708 | 0.466461666 | -0.84963618 | 0.395527391 | 0.69426198 |
| CLUHP3 | 404.9425344 | -0.732365469 | 0.343669299 | -2.131018022 | 0.033087658 | 0.20777022 |
| CLVS1 | 55.71545209 | -0.08521188 | 0.751220278 | -0.113431282 | 0.90968864 | 0.96630013 |
| CLVS2 | 24.94471684 | -0.44214396 | 0.728230768 | -0.607148145 | 0.543752601 | 0.790216482 |
| CLYBL | 399.9060497 | 0.864390422 | 0.380975964 | 2.2688844 | 0.023275357 | 0.174130017 |
| CMAS | 1225.33848 | 0.532502046 | 0.264598962 | 2.012487287 | 0.044168595 | 0.241589846 |
| CMBL | 618.8746433 | 0.981999211 | 0.372459081 | 2.636529116 | 0.008375901 | 0.103556802 |
| CMC1 | 305.0782613 | 0.847223026 | 0.447548944 | 1.893028769 | 0.058354044 | 0.279977608 |
| CMC2 | 622.796462 | 1.307696547 | 0.368011195 | 3.553415123 | 0.000380264 | 0.014675547 |
| CMC4 | 153.674894 | 0.408919523 | 0.356407708 | 1.147336363 | 0.251242631 | 0.564075361 |
| CMIP | 1458.165108 | -0.213441237 | 0.26304792 | -0.811415791 | 0.417126934 | 0.708661116 |
| CMPK1 | 1355.410154 | 0.264215725 | 0.300014261 | 0.880677218 | 0.378492551 | 0.681253679 |
| CMSS1 | 941.7812008 | 0.808788655 | 0.3111812 | 2.599092279 | 0.009347064 | 0.109346547 |
| CMTM3 | 1232.585218 | 0.183180631 | 0.334200276 | 0.548116335 | 0.583612025 | 0.8142179 |
| CMTM4 | 1134.313029 | -0.626832096 | 0.376867749 | -1.663268078 | 0.096258786 | 0.356020645 |
| CMTM5 | 43.28313834 | 1.151835247 | 0.633402557 | 1.81848847 | 0.068989507 | 0.302794854 |
| CMTM6 | 2219.058794 | -0.073713791 | 0.290918487 | -0.253382973 | 0.799972284 | 0.916311991 |
| CMTM7 | 751.4881258 | 0.772272583 | 0.484960161 | 1.592445412 | 0.111284659 | 0.383708973 |
| CMTM8 | 176.30452 | 0.851812249 | 0.517944689 | 1.644600799 | 0.100052162 | 0.36243601 |
| CMTR1 | 1471.97703 | -0.346119628 | 0.43154199 | -0.802053187 | 0.422522193 | 0.712269066 |
| CMTR2 | 793.4872135 | -0.229762947 | 0.35850696 | -0.640888386 | 0.521595202 | 0.776633705 |
| CMYA5 | 55.85981961 | 0.378794694 | 0.560113444 | 0.676282097 | 0.498861558 | 0.76281557 |
| CNBD2 | 58.45504008 | 1.333883836 | 0.569934546 | 2.340415833 | 0.01926228 | 0.157577123 |
| CNBP | 8970.247822 | 0.403231968 | 0.278155998 | 1.44966124 | 0.147153009 | 0.440413949 |
| CNDP2 | 3707.130716 | -0.353896598 | 0.24028742 | -1.472805353 | 0.140803529 | 0.430693807 |
| CNEP1R1 | 484.389411 | 0.408489644 | 0.312022699 | 1.309166433 | 0.19047798 | 0.496058205 |
| CNFN | 34.90959201 | 0.921816455 | 0.56332353 | 1.636389049 | 0.10175819 | 0.365921972 |
| CNGA1 | 17.95971638 | -0.019559658 | 1.080393349 | -0.018104201 | 0.985555727 | 0.994669101 |
| CNGB1 | 21.46485695 | -0.767787124 | 0.872706578 | -0.879776942 | 0.378980158 | 0.681253679 |
| CNIH1 | 2632.959512 | 0.873077421 | 0.314024226 | 2.780286833 | 0.00543109 | 0.078471497 |
| CNIH2 | 120.0304744 | 0.997108818 | 0.383508352 | 2.599966368 | 0.00932329 | 0.109346547 |
| CNIH3 | 57.00414086 | 0.193659404 | 0.476331255 | 0.406564554 | 0.684327835 | 0.863857185 |
| CNIH4 | 1427.447215 | 0.514494041 | 0.351179805 | 1.465044496 | 0.142908778 | 0.433233718 |
| CNKSR1 | 684.6275687 | -0.706076083 | 0.330596502 | -2.135763927 | 0.032698661 | 0.206178012 |
| CNKSR2 | 450.5116621 | -0.733286124 | 0.317870946 | -2.3068674 | 0.021062213 | |

| | | | | | | |
|---|---|---|---|---|---|---|
| | | | | | | 0.16470868 |
| CNKSR3 | 1121.421014 | 1.886018669 | 0.718900285 | 2.623477428 | 0.008703721 | 0.105298369 |
| CNMD | 154.6718814 | 0.016658538 | 0.460595658 | 0.036167379 | 0.971148897 | 0.989629517 |
| CNN1 | 61.59813017 | -0.12553241 | 0.45029714 | -0.278776831 | 0.780416096 | 0.909206926 |
| CNN2 | 7065.297403 | -0.353061006 | 0.308986372 | -1.142642646 | 0.253186992 | 0.565800563 |
| CNN3 | 7297.466514 | 0.451594676 | 0.232311995 | 1.943914591 | 0.051905751 | 0.262699338 |
| CNNM1 | 10.40686011 | 0.583604179 | 0.902215297 | 0.646856889 | 0.51772457 | 0.773950969 |
| CNNM2 | 647.4524101 | -0.491964521 | 0.334892139 | -1.469023796 | 0.141826335 | 0.43212631 |
| CNNM3 | 845.5348214 | -0.640652607 | 0.314368304 | -2.037904579 | 0.041559477 | 0.234125072 |
| CNNM4 | 1334.440904 | -1.146671572 | 0.283317646 | -4.047300224 | 5.18E-05 | 0.0038721 |
| CNOT1 | 8394.251889 | -0.266192772 | 0.367509553 | -0.724315245 | 0.46887222 | 0.744337782 |
| CNOT10 | 1210.763499 | -0.224910652 | 0.255208774 | -0.881281035 | 0.378165728 | 0.681253679 |
| CNOT11 | 1424.962193 | -0.045130156 | 0.346685061 | -0.130176234 | 0.896426997 | 0.959075738 |
| CNOT2 | 2203.50038 | 0.10662322 | 0.258292816 | 0.412799791 | 0.679753305 | 0.861682789 |
| CNOT3 | 1315.044662 | -1.001255946 | 0.635091771 | -1.576553173 | 0.114898375 | 0.389985713 |
| CNOT4 | 502.1605653 | -0.494960965 | 0.365269367 | -1.355057418 | 0.175399264 | 0.479452242 |
| CNOT6 | 1691.931506 | -0.227693425 | 0.249291787 | -0.913361117 | 0.361052649 | 0.667991431 |
| CNOT6L | 347.3414519 | -0.599663215 | 0.419867454 | -1.428220286 | 0.153228463 | 0.44961176 |
| CNOT7 | 3905.369489 | 0.110994539 | 0.224574475 | 0.494243785 | 0.621134021 | 0.833349572 |
| CNOT8 | 1825.476462 | 0.473577526 | 0.271700929 | 1.743010329 | 0.081331811 | 0.326165715 |
| CNOT9 | 1947.726545 | 0.097070017 | 0.229407122 | 0.423134279 | 0.672197294 | 0.858915619 |
| CNP | 2313.170897 | 0.074081014 | 0.297771038 | 0.248785156 | 0.803526974 | 0.917969611 |
| CNPPD1 | 1107.21137 | 0.00160819 | 0.304042828 | 0.005289353 | 0.995779727 | 0.998171604 |
| CNPY2 | 2742.401797 | 0.714991069 | 0.398618185 | 1.793673987 | 0.07286522 | 0.310624251 |
| CNPY3 | 1392.43915 | 0.199384444 | 0.276916787 | 0.720015737 | 0.471515307 | 0.746229785 |
| CNPY4 | 387.74964 | 0.580721978 | 0.26204303 | 2.216132129 | 0.026682454 | 0.186715644 |
| CNR1 | 16.03865469 | -1.823829106 | 0.883219628 | -2.064978006 | 0.038925081 | 0.227316695 |
| CNRIP1 | 26.46264859 | 1.745393316 | 0.98768786 | 1.767150723 | 0.07720299 | 0.31844659 |
| CNST | 543.0026015 | 0.213274925 | 0.272612567 | 0.7823371 | 0.434016489 | 0.719296316 |
| CNTD2 | 12.11879412 | 0.591831821 | 0.897825051 | 0.659183903 | 0.509777682 | 0.769834282 |
| CNTF | 10.60777539 | -4.594962553 | 1.668596796 | -2.753788431 | 0.005890984 | 0.082905485 |
| CNTFR | 466.1544668 | 1.533327202 | 0.61502107 | 2.493129546 | 0.012662264 | 0.127358401 |
| CNTLN | 513.4121884 | 0.175190075 | 0.314403884 | 0.557213456 | 0.577381597 | 0.809901794 |
| CNTN1 | 91.94532873 | -2.227441888 | 0.685569745 | -3.249037614 | 0.001157962 | 0.029724934 |
| CNTN2 | 39.79342409 | 0.805939322 | 0.9348197 | 0.862133437 | 0.388614093 | 0.688604437 |
| CNTNAP1 | 679.4053167 | 0.401142931 | 0.620720387 | 0.646253836 | 0.518114979 | 0.773958483 |
| CNTNAP2 | 1865.922079 | 0.288650454 | 0.51435595 | 0.561188131 | 0.574669293 | 0.808359708 |
| CNTNAP3 | 184.5042814 | 0.433569526 | 0.590428507 | 0.734330272 | 0.462747472 | 0.74030812 |
| CNTNAP3B | 132.2959244 | 0.87544107 | 0.427746538 | 2.046635083 | 0.040693928 | 0.23195346 |
| CNTNAP3C | 10.78596441 | -0.158774907 | 1.351348371 | -0.11749369 | 0.906468844 | 0.965055357 |
| CNTNAP3P2 | 12.2681992 | 1.347244907 | 1.474675883 | 0.91358713 | 0.360933832 | |

0.667991431
CNTNAP5 12.01576001 -1.705934067 0.964048762 -1.769551638
0.076801861 0.31831258
CNTRL 665.471396 0.226120573 0.454429002 0.497592741 0.618771121 0.832361827
CNTROB 1218.223574 0.153227739 0.329512614 0.465013274 0.641921951
0.844118741
COA1 2476.8239 0.885194677 0.291700355 3.034602668 0.002408528 0.047209009
COA3 796.1777506 0.566092671 0.440615634 1.284776631 0.198870351 0.504782074
COA4 1256.543169 0.845255244 0.378607981 2.232534144 0.025579681 0.182449151
COA5 688.0625902 0.625494746 0.281697573 2.220447761 0.026388388 0.185660023
COA6 579.2947082 1.34254338 0.439685989 3.053414056 0.002262535 0.045223875
COA7 1005.065798 0.141300984 0.34189409 0.413288758 0.679395065 0.861682789
COASY 1626.977874 -0.276462289 0.224303636 -1.232535923 0.217748956
0.526030161
COBL 1420.908944 -1.250093546 0.26704415 -4.681224242 2.85E-06
0.000420026
COBLL1 108.2539437 -1.640767269 0.56435302 -2.907342057
0.003645144 0.062135365
COCH 341.523084 0.214708019 0.508566617 0.422182683 0.67289168 0.859151631
COG1 473.8175817 -0.600494972 0.337117732 -1.781261901 0.074869675
0.314030369
COG2 849.6912047 -0.044750827 0.238868045 -0.187345391 0.851389836
0.939477675
COG3 841.3660305 -0.171755 0.251746596 -0.682253517 0.49507866
0.760514211
COG4 1838.614946 -0.441114372 0.304056213 -1.450765854 0.146845071
0.439926258
COG5 448.9964631 -0.004210125 0.373563945 -0.01127016 0.991007904
0.996726674
COG6 954.8208981 0.121967174 0.280492209 0.434832663 0.663683894 0.854667747
COG7 486.8007698 -0.081711943 0.305949684 -0.26707641 0.78941033
0.912815863
COG8 859.1379716 -0.186260996 0.27978939 -0.66571858 0.505590995
0.76710881
COIL 777.6882631 0.286015235 0.237419633 1.204682322 0.228325963 0.537209827
COL11A1 452.5588252 1.269304571 0.700135508 1.812941289 0.069840877
0.303603614
COL12A1 95.811596 0.580151364 0.841808693 0.689172456 0.490714748
0.758593654
COL13A1 320.4881691 0.682288391 0.428012286 1.594085995 0.110916775
0.383132604
COL14A1 286.9068748 0.824460413 0.630173034 1.308307986 0.190768868
0.496058205
COL15A1 14.59633447 -0.021087753 0.759208001 -0.027775989
0.977840816 0.99226352
COL16A1 276.3970114 -0.754156173 0.521262368 -1.446788065
0.147956291 0.441755032
COL17A1 229.9842725 -1.369424103 0.546427202 -2.506141892
0.012205663 0.125625583
COL18A1 5706.788283 -0.050274668 0.497833947 -0.100986821
0.919560922 0.969677217
COL1A1 1254.608484 0.882251952 0.502375772 1.756159435 0.079061189
0.322602824
COL1A2 6932.376635 0.486661581 0.491025574 0.991112493 0.321630653
0.631318236
COL21A1 158.9827146 -1.654355972 0.629848888 -2.626591877
0.008624471 0.104839578
COL23A1 34.39198809 -0.03876031 0.597519703 -0.064868672 0.948278564
0.98134156
COL25A1 28.95774669 -0.876374295 0.876129874 -1.000278978
0.317175517 0.627198876
COL26A1 892.0128091 -0.22131085 0.253404157 -0.873351299 0.382471628
0.683824871
COL27A1 849.0946689 -0.048035656 0.272134902 -0.176514133

| | | | | | | |
|---|---|---|---|---|---|---|
| | | | | | 0.859890047 | 0.943267672 |
| COL28A1 | 28.29565979 | -0.228805787 | 0.692822847 | -0.330251504 | 0.741209933 | 0.891956524 |
| COL2A1 | 1662.931741 | -0.473452441 | 0.540055215 | -0.87667414 | 0.380663657 | 0.682307223 |
| COL3A1 | 69.89834003 | -5.383059331 | 1.516329841 | -3.550058297 | 0.000385146 | 0.014792526 |
| COL4A1 | 3536.858815 | 0.28834944 | 0.487397554 | 0.591610355 | 0.554111538 | 0.796360966 |
| COL4A2 | 3518.62943 | 0.551150831 | 0.554799787 | 0.993422933 | 0.32050389 | 0.63024811 |
| COL4A3BP | 1597.072715 | -0.471582505 | 0.264900527 | -1.780224864 | 0.075039168 | 0.314480446 |
| COL4A4 | 35.32765882 | 0.447251753 | 0.570169065 | 0.784419534 | 0.43279398 | 0.718657207 |
| COL4A5 | 2435.468969 | -0.511969129 | 0.289663723 | -1.767460291 | 0.077151174 | 0.31844659 |
| COL4A6 | 1567.059038 | -0.582759154 | 0.291326482 | -2.000364509 | 0.045460918 | 0.245452751 |
| COL5A1 | 1159.979423 | -0.47938426 | 0.531329838 | -0.902234782 | 0.366932162 | 0.673470229 |
| COL5A2 | 2365.779548 | -1.856619003 | 0.603776897 | -3.075008355 | 0.002104967 | 0.043389195 |
| COL5A3 | 13.29475021 | 1.377220477 | 0.818333861 | 1.682956728 | 0.092383469 | 0.348900321 |
| COL6A1 | 705.7463209 | -0.351134122 | 0.523757574 | -0.67041345 | 0.502594263 | 0.764968568 |
| COL6A2 | 518.8105794 | -0.835721604 | 0.500318086 | -1.670380558 | 0.094844094 | 0.353533108 |
| COL6A3 | 166.5102313 | -1.679134139 | 0.930427164 | -1.804691655 | 0.071122946 | 0.30688459 |
| COL6A4P2 | 30.9929398 | -0.304377417 | 0.597606349 | -0.509327617 | 0.610522603 | 0.828909828 |
| COL6A6 | 119.077917 | -5.925459234 | 1.163229126 | -5.093974269 | 3.51E-07 | 9.51E-05 |
| COL7A1 | 527.949623 | 0.020041602 | 0.30291687 | 0.066162053 | 0.947248808 | 0.980790989 |
| COL9A1 | 222.6852782 | 1.550294848 | 0.453657853 | 3.417321747 | 0.000632405 | 0.020419026 |
| COL9A2 | 602.8127791 | 0.196937253 | 0.482662855 | 0.408022394 | 0.683257234 | 0.863415274 |
| COL9A3 | 784.783301 | -0.332116219 | 0.477492037 | -0.695542947 | 0.486715107 | 0.756340124 |
| COLCA2 | 25.49767573 | -1.191921349 | 0.83429974 | -1.428648831 | 0.153105192 | 0.449450244 |
| COLEC11 | 26.4617737 | 1.035630399 | 0.686151654 | 1.509331637 | 0.131214054 | 0.416813082 |
| COLEC12 | 1256.850434 | -1.567269652 | 0.51368918 | -3.051007715 | 0.002280747 | 0.045527914 |
| COLGALT1 | 1543.028365 | 0.221621815 | 0.422034421 | 0.525127344 | 0.599494669 | 0.823816451 |
| COLGALT2 | 127.2584283 | 1.418023314 | 0.726339012 | 1.952288519 | 0.05090396 | 0.259366654 |
| COLQ | 90.12322089 | -0.976520641 | 0.572827623 | -1.704737344 | 0.088243422 | 0.34168988 |
| COMMD1 | 443.149035 | 0.442984107 | 0.368126114 | 1.203348772 | 0.228841383 | 0.537506211 |
| COMMD10 | 761.2603613 | 0.505293039 | 0.248804538 | 2.030883534 | 0.04226681 | 0.236528876 |
| COMMD2 | 1459.313481 | 0.180603365 | 0.230106348 | 0.784869111 | 0.432530314 | 0.718639059 |
| COMMD3 | 379.431269 | 0.903648136 | 0.357495651 | 2.527717841 | 0.011480658 | 0.121279761 |
| COMMD3-BMI1 | 17.48324434 | 1.664218443 | 1.637827056 | 1.016113659 | 0.309575265 | |

0.61915097
COMMD4 1715.69763 0.503610214 0.419890264 1.199385309 0.230378157
0.539112606
COMMD5 350.7217649 0.45505459 0.404171842 1.125893848 0.260210454
0.57337005
COMMD6 1788.313391 1.027072048 0.455075779 2.256925318 0.024012742
0.177207433
COMMD7 692.8357496 -0.441114195 0.378753941 -1.164645823
0.244162387 0.555457814
COMMD8 566.7393635 0.911473636 0.319152771 2.855916409 0.004291281
0.067957227
COMMD9 540.7084732 -0.778844267 0.299181735 -2.603248046
0.009234512 0.108795292
COMT 223.2815323 0.604222472 0.370144734 1.63239516 0.102596268 0.367636293
COMTD1 80.23995341 1.104474969 0.738065396 1.496445945 0.134537484
0.420896533
COP1 1569.766605 -0.024049885 0.229959187 -0.104583276 0.916706486
0.969211928
COPA 5245.695307 -0.685620679 0.318061023 -2.155626217 0.031112864
0.200942211
COPB1 4648.726766 -0.02548672 0.226574741 -0.11248703 0.910437253 0.966517087
COPB2 4014.808064 -0.263435433 0.227886338 -1.155994849 0.247683264
0.560334446
COPE 1896.777335 0.002392723 0.429103774 0.005576094 0.995550944 0.998116797
COPG1 3140.057801 -0.441482045 0.230404161 -1.916120105 0.05534981
0.272191885
COPG2 3219.427619 0.119807247 0.275906239 0.434231742 0.664120165 0.854858903
COPRS 705.1066383 1.211561072 0.427823455 2.831918303 0.004626967 0.071864367
COPS2 3338.934937 0.37413007 0.292408918 1.279475579 0.200729634 0.506869054
COPS3 1743.73108 0.421047072 0.228838663 1.839929786 0.065778546 0.29619856
COPS4 1398.679656 0.464574 0.263136083 1.765527535 0.077475147 0.319212925
COPS5 2103.430867 0.390035775 0.294652222 1.323715711 0.185597479 0.490113028
COPS6 1990.782714 0.123951048 0.223040075 0.555734427 0.578392418 0.810209453
COPS7A 907.5203516 0.316013679 0.244940671 1.290164176 0.196993661
0.50258876
COPS7B 1659.847288 0.160208244 0.227567117 0.704004366 0.481430054
0.752733727
COPS8 2181.523489 0.594082701 0.224459123 2.646730033 0.008127421 0.101315612
COPS9 929.0855251 1.320352037 0.513561093 2.570973649 0.010141304 0.114304404
COPZ1 3591.015201 0.427262937 0.308560382 1.384697976 0.166144838 0.468031848
COQ10A 700.4579005 0.736088788 0.392013671 1.877712035 0.060420569
0.28404296
COQ10B 478.6242933 0.265188188 0.359949412 0.736737384 0.461282068
0.738942035
COQ2 496.3796015 0.622793266 0.282344536 2.205791814 0.02739858 0.189349252
COQ3 271.4367005 0.801973607 0.376187729 2.131844142 0.033019661 0.207677316
COQ4 749.3380436 -0.074906359 0.311185109 -0.240713185 0.809777419
0.920165773
COQ5 1403.329144 0.335269401 0.265762794 1.26153626 0.207115704 0.513927437
COQ6 283.3524531 -0.01970136 0.301918474 -0.065253908 0.94797184
0.981274993
COQ7 454.0228926 0.392624004 0.283533839 1.384751833 0.166128364 0.468031848
COQ8A 546.9825107 0.538398377 0.307371109 1.751623239 0.079838614 0.323453279
COQ8B 1154.593183 0.122752478 0.365949859 0.335435237 0.737296791 0.89093937
COQ9 523.3048899 0.147179904 0.318177873 0.462571148 0.643671793 0.845466381
CORIN 12.91994684 -1.560605209 0.916488821 -1.702808778 0.08860386
0.342280113
CORO1A 106.0161729 0.583378303 0.556537396 1.048228398 0.29453339
0.606702791
CORO1B 1703.698482 -0.523033542 0.447765501 -1.168097008
0.242767637 0.553922067
CORO1C 4811.414674 0.048304867 0.292926109 0.164904614 0.869019061
0.947201167
CORO2A 901.7550364 -1.76291553 0.524516163 -3.361031849 0.000776519

0.023327852
CORO2B	56.63481518 1.365976569 0.76109103 1.794761094 0.072691772
0.310433977
CORO6 44.45010583 1.377688029 0.680687055 2.023966841 0.042973564 0.238392703
CORO7 343.7964965 0.237493713 0.394238536 0.60241121 0.54690045 0.792249935
COTL1 5030.351526 0.825933604 0.325824807 2.534900921 0.011247913 0.120339982
COX10 482.1145328 -0.083453605 0.263217218 -0.317052225 0.751203975
0.896349431
COX11 1204.495812 0.514808067 0.292181783 1.761944436 0.078078686 0.320406654
COX14 307.0837397 0.960411165 0.510099836 1.882790578 0.059728754 0.282669434
COX15 1271.024549 -0.083171862 0.233442029 -0.356284864 0.721627243
0.882498649
COX16 1350.800434 1.352656507 0.457948039 2.953733596 0.003139549 0.055790951
COX17 753.1945442 1.604344697 0.393381586 4.078342135 4.54E-05 0.00345792
COX18 550.6551873 0.710213436 0.39929898 1.778650766 0.075297036 0.314600339
COX19 455.451284 0.309025391 0.310655599 0.994752362 0.319856721 0.629465081
COX20 652.0325634 0.325997414 0.324842803 1.003554367 0.31559346 0.625406234
COX4I1	8741.035946 0.825441901 0.450216744 1.833432259 0.066738314
0.297817842
COX5A 3165.480063 1.080778013 0.480736457 2.248171525 0.024565255 0.179037045
COX5B 2667.463537 1.113802508 0.442382811 2.517734598 0.011811228 0.12332288
COX6A1	3884.829036 1.185541775 0.449744894 2.63603165 0.008388191
0.103556802
COX6B1	4432.878399 1.020214208 0.479786232 2.126393256 0.033470529
0.208789229
COX6B2	22.92587615 -0.022504768 0.728432462 -0.03089479 0.975353445
0.991429622
COX6C 3185.586433 1.340326464 0.471009074 2.84564892 0.004432103 0.069675779
COX7A2	4023.648959 1.342265496 0.413417676 3.246754007 0.001167292
0.029913838
COX7A2L	4295.163351 0.782609222 0.410734225 1.905390821 0.056729288
0.27582404
COX7B 2355.983374 1.262849339 0.410552635 3.075974262 0.00209816 0.043389195
COX7C 9425.575667 0.978527105 0.405136234 2.415303849 0.015722088 0.141976071
COX8A 1690.036485 1.126955334 0.416423242 2.706273862 0.006804291 0.090392202
CP	13.99939142 -0.519846127 0.825640915 -0.62962738 0.528938405
0.779891202
CPAMD8	296.0767 -0.675499491 0.438476881 -1.540558968
0.123424161 0.404779768
CPD	2865.690001 -0.163085184 0.296752815 -0.549565752 0.582617254
0.813577525
CPE	2845.668734 0.432090193 0.411427138 1.050222877 0.293615654 0.606359511
CPEB1 41.97002247 0.956707853 0.922263667 1.037347438 0.299573965 0.610246278
CPEB2 118.4176295 -0.163815579 0.429582405 -0.381336799 0.702953351
0.872703476
CPEB3 52.61487626 -0.575187176 0.504494499 -1.140125764 0.25423391
0.566703298
CPEB4 665.0343978 -1.41038395 0.381657997 -3.695413071 0.00021953
0.010092374
CPLANE1	741.351466 -0.694862701 0.382708137 -1.815646529
0.069424611 0.30336211
CPLANE2	74.69861446 0.260041014 0.513214555 0.506690646 0.612371895
0.829194397
CPM	22.88949624 -0.029391037 0.621296105 -0.047306006 0.962269341
0.987343615
CPNE1 2712.617958 -0.190075155 0.288380096 -0.659113294 0.509823019
0.769834282
CPNE2 429.0032246 -0.347663207 0.313555106 -1.108778648 0.267525684
0.580717148
CPNE3 1986.47911 -0.332732129 0.237368278 -1.401754826 0.160988472
0.46167973
CPNE4 20.14262218 -0.81028554 0.813500751 -0.996047685 0.319226977
0.629115732
CPNE7 22.39468643 0.088791653 0.644004579 0.137874257 0.890339796 0.956814128

| | | | | | | |
|---|---|---|---|---|---|---|
| CPNE8 | 352.1879998 | -0.443242543 | 0.331642567 | -1.336506792 | 0.181383682 | 0.485749663 |
| CPNE9 | 37.35992651 | -0.088321455 | 0.609159173 | -0.144989124 | 0.884719459 | 0.95440284 |
| CPOX | 951.0371158 | 0.302942826 | 0.23590856 | 1.2841536 | 0.199088217 | 0.504867077 |
| CPPED1 | 370.4000118 | 0.015665091 | 0.284798098 | 0.055004198 | 0.956135119 | 0.984426596 |
| CPQ | 48.08584287 | -0.089110648 | 0.627502395 | -0.142008459 | 0.887073324 | 0.955673157 |
| CPS1 | 1073.37487 | -0.307990246 | 0.244512621 | -1.259608791 | 0.207810523 | 0.514566354 |
| CPSF1 | 1709.568838 | -0.297853464 | 0.479830194 | -0.620747648 | 0.534765674 | 0.784009474 |
| CPSF2 | 2543.658218 | -0.108251678 | 0.316832665 | -0.341668301 | 0.73260053 | 0.888122198 |
| CPSF3 | 3389.172687 | 0.153672843 | 0.244010764 | 0.62977895 | 0.52883922 | 0.779842516 |
| CPSF4 | 2207.450134 | 0.265758019 | 0.270776631 | 0.98146586 | 0.326363062 | 0.636162664 |
| CPSF6 | 5301.239133 | 0.170069127 | 0.278692382 | 0.610239598 | 0.541703102 | 0.788617001 |
| CPSF7 | 3058.884528 | -0.458356411 | 0.357389333 | -1.282512849 | 0.199662801 | 0.505774645 |
| CPT1A | 111.18515 | 0.447074608 | 0.382921474 | 1.167536006 | 0.242993976 | 0.554163307 |
| CPT1B | 1171.580108 | -0.487289071 | 0.333262932 | -1.462176031 | 0.143692978 | 0.434862592 |
| CPT1C | 1477.863008 | 0.417873159 | 0.439064377 | 0.95173551 | 0.341231135 | 0.648152255 |
| CPT2 | 727.9540154 | 0.033871738 | 0.24966847 | 0.135666864 | 0.892084644 | 0.957468421 |
| CPTP | 105.5931364 | -0.113508694 | 0.58966582 | -0.192496649 | 0.847353193 | 0.938171381 |
| CPVL | 2014.249384 | -0.155826932 | 0.266402013 | -0.58493151 | 0.55859377 | 0.798871238 |
| CPXM1 | 2869.841318 | 0.627707333 | 0.437760036 | 1.433907349 | 0.151598704 | 0.447172604 |
| CPXM2 | 545.4851455 | -0.5729412 | 0.709223053 | -0.807843452 | 0.419180707 | 0.709924862 |
| CPZ | 847.7523423 | -0.64298988 | 0.430825719 | -1.492459366 | 0.135578762 | 0.423136268 |
| CR1 | 11.11124434 | -0.055038899 | 0.924015981 | -0.059564877 | 0.952502193 | 0.982820565 |
| CR1L | 60.99199978 | -2.767815206 | 0.710000318 | -3.898329531 | 9.69E-05 | 0.005997719 |
| CR383656.13 | 73.58975036 | 1.221752461 | 0.702776951 | 1.738464045 | 0.08212908 | 0.328062211 |
| CRABP1 | 23353.09755 | 2.270652391 | 1.284844486 | 1.76725854 | 0.07718494 | 0.31844659 |
| CRABP2 | 16858.93881 | 0.826711845 | 0.359881238 | 2.29717962 | 0.021608525 | 0.166611729 |
| CRACR2A | 23.07698284 | 0.997997233 | 0.666928886 | 1.496407269 | 0.134547556 | 0.420896533 |
| CRACR2B | 106.668062 | 0.478356854 | 0.462473917 | 1.034343423 | 0.300975644 | 0.611749932 |
| CRADD | 198.0168915 | 0.391508308 | 0.362169729 | 1.081007816 | 0.279693637 | 0.593293088 |
| CRAMP1 | 1196.53858 | -0.824739983 | 0.430092136 | -1.917589077 | 0.055163136 | 0.271802514 |
| CRAT | 368.6043992 | 0.679136716 | 0.556382059 | 1.22063015 | 0.222226085 | 0.531069497 |
| CRB2 | 805.9261983 | 2.122856041 | 0.794248127 | 2.672786964 | 0.007522401 | 0.096389041 |
| CRB3 | 107.833533 | -1.489691891 | 0.74063883 | -2.011360775 | 0.044287363 | 0.2418127 |
| CRBN | 1139.422638 | 0.231604547 | 0.269890969 | 0.858141151 | 0.390814527 | 0.690469214 |
| CRCP | 863.0048633 | -0.035827737 | 0.250298912 | -0.143139805 | 0.886179769 | 0.955049604 |
| CREB1 | 2368.106818 | -0.366398053 | 0.263885421 | -1.388474029 | 0.164992748 | 0.467115865 |
| CREB3 | 667.9372116 | -0.252433851 | 0.339848597 | -0.742783267 | 0.457612908 | 0.736329582 |
| CREB3L1 | 274.3243577 | -0.49187387 | 0.316941372 | -1.551939611 | 0.120676674 | 0.400973879 |

| | | | | | | |
|---|---|---|---|---|---|---|
| CREB3L2 | 1967.321797 | -1.009241631 | 0.384142354 | -2.627259451 | 0.008607568 | 0.104830807 |
| CREB3L4 | 661.9843495 | 0.199786288 | 0.257672153 | 0.775350717 | 0.438132448 | 0.721949744 |
| CREB5 | 52.16616381 | -2.259182703 | 0.87215869 | -2.590334453 | 0.009588273 | 0.111098245 |
| CREBBP | 1921.511882 | -0.579203056 | 0.277728975 | -2.085497406 | 0.037024173 | 0.220531501 |
| CREBL2 | 877.3245048 | -0.047006634 | 0.26163598 | -0.179664258 | 0.857416155 | 0.942343063 |
| CREBRF | 297.834827 | -0.515131502 | 0.310259027 | -1.660327202 | 0.096848646 | 0.357404722 |
| CREBZF | 1601.054925 | 0.099947178 | 0.251036872 | 0.398137443 | 0.690528876 | 0.867938158 |
| CREG1 | 1581.084835 | 0.228153497 | 0.33837796 | 0.674256377 | 0.500148333 | 0.763360896 |
| CREG2 | 37.92417077 | -0.729692874 | 0.846105029 | -0.862414061 | 0.388459706 | 0.688552657 |
| CRELD1 | 581.4671828 | -0.610472668 | 0.307250614 | -1.986888356 | 0.04693477 | 0.249564641 |
| CRELD2 | 494.6228823 | 0.412961403 | 0.400854294 | 1.030203265 | 0.302914597 | 0.613538084 |
| CREM | 544.5437202 | 0.503275645 | 0.292024137 | 1.72340427 | 0.084815443 | 0.33352387 |
| CRHR1 | 28.9099521 | 0.106433375 | 1.00753524 | 0.105637372 | 0.915870072 | 0.969169304 |
| CRIM1 | 572.7030719 | -0.837660651 | 0.376266947 | -2.226240327 | 0.025998088 | 0.184130533 |
| CRIP1 | 282.3820229 | -0.089763956 | 0.72094664 | -0.124508461 | 0.900912702 | 0.961704658 |
| CRIP2 | 34.41713546 | 0.782517834 | 0.775647297 | 1.008857812 | 0.313042831 | 0.622994042 |
| CRIP3 | 219.2586645 | 1.660562933 | 0.520112438 | 3.192699907 | 0.001409493 | 0.034006242 |
| CRIPT | 545.5226457 | 0.736279123 | 0.471817433 | 1.560516997 | 0.118637756 | 0.396986757 |
| CRISP3 | 17.41346374 | -0.949355049 | 1.06524468 | -0.89120844 | 0.372817357 | 0.67811865 |
| CRISPLD1 | 1549.960095 | -0.270373543 | 0.261571682 | -1.033649902 | 0.301299862 | 0.612159932 |
| CRISPLD2 | 96.97438188 | -1.36744323 | 0.597075477 | -2.290235127 | 0.02200769 | 0.168540468 |
| CRK | 2336.077191 | 0.02627742 | 0.245874526 | 0.106873292 | 0.914889501 | 0.968785414 |
| CRKL | 3466.878944 | -0.153813348 | 0.227298791 | -0.676701125 | 0.498595603 | 0.762775559 |
| CRLF1 | 10.28176971 | 0.678363376 | 0.895090604 | 0.757871184 | 0.448528103 | 0.729796208 |
| CRLF3 | 567.2969824 | 0.047230284 | 0.257403737 | 0.183487173 | 0.854415782 | 0.941485939 |
| CRLS1 | 986.7762901 | 0.78137661 | 0.335064463 | 2.332018748 | 0.019699705 | 0.158893482 |
| CRMP1 | 2343.870102 | -0.525965417 | 0.230732064 | -2.279550606 | 0.022634355 | 0.171427007 |
| CRNKL1 | 1244.05568 | -0.051770704 | 0.247484608 | -0.209187572 | 0.834301814 | 0.932107564 |
| CROCC | 923.5196289 | 0.228774558 | 0.418026327 | 0.547273086 | 0.584191131 | 0.814692336 |
| CROCCP2 | 926.1525865 | 0.483166346 | 0.290443642 | 1.663545956 | 0.0962032 | 0.355975303 |
| CROCCP3 | 146.4411248 | -0.203561356 | 0.53798198 | -0.378379506 | 0.70514869 | 0.873789984 |
| CROT | 841.2674203 | -0.413900109 | 0.404531465 | -1.023159246 | 0.306232559 | 0.616286722 |
| CRTAC1 | 25.91906519 | 1.008276682 | 0.584889775 | 1.723874694 | 0.084730467 | 0.333276098 |
| CRTAP | 4526.774355 | -0.657865455 | 0.318208655 | -2.067402769 | 0.038696218 | 0.226576733 |
| CRTC1 | 322.5229378 | -0.426116555 | 0.465997971 | -0.91441719 | 0.360497673 | 0.667663818 |
| CRTC2 | 717.601369 | -0.510525407 | 0.308725113 | -1.653656882 | 0.098197248 | 0.359930044 |
| CRTC3 | 338.5999158 | -0.836419834 | 0.299499679 | -2.792723639 | 0.005226632 | 0.076983729 |
| CRX | 16.0061241 | 1.263370708 | 0.979686841 | 1.289565865 | 0.197201434 | 0.50258876 |

| | | | | | | |
|---|---|---|---|---|---|---|
| CRY1 | 982.8856639 | 0.33912714 | 0.254513049 | 1.332454825 | 0.182710777 | 0.486683209 |
| CRY2 | 758.7101711 | -0.29540606 | 0.278696293 | -1.059956903 | 0.289164207 | 0.602227073 |
| CRYAB | 41.25334445 | 0.171114144 | 0.548369336 | 0.312041781 | 0.755008771 | 0.897581331 |
| CRYBA1 | 29.37391285 | -0.755878971 | 0.718074424 | -1.052647114 | 0.292502755 | 0.605587347 |
| CRYBA4 | 10.23879179 | 1.87295993 | 0.948151531 | 1.975380378 | 0.048224998 | 0.253243836 |
| CRYBB2P1 | 809.1638861 | 0.765992759 | 0.338624655 | 2.26207025 | 0.023693063 | 0.176113407 |
| CRYBG1 | 73.32490524 | -1.716630195 | 0.873490778 | -1.965252798 | 0.049384986 | 0.25614346 |
| CRYBG2 | 233.3441546 | -0.409862511 | 0.480300786 | -0.853345494 | 0.393467738 | 0.69249409 |
| CRYBG3 | 256.9165089 | -0.143035251 | 0.515682577 | -0.277370727 | 0.781495457 | 0.909755868 |
| CRYGN | 36.27858166 | -2.130509322 | 0.93974485 | -2.267114656 | 0.023383223 | 0.174580152 |
| CRYGS | 39.91479499 | 0.589642227 | 0.651724808 | 0.904741112 | 0.365602555 | 0.672153785 |
| CRYL1 | 61.6932388 | -0.458503696 | 0.700179255 | -0.654837591 | 0.512572325 | 0.771819984 |
| CRYM | 12.20479213 | -0.033186753 | 0.883275682 | -0.037572361 | 0.970028645 | 0.989496332 |
| CRYZ | 1767.552068 | 0.175075625 | 0.241620517 | 0.724589233 | 0.468704066 | 0.744337782 |
| CRYZL1 | 834.1305839 | 0.627158061 | 0.266936144 | 2.349468497 | 0.018800235 | 0.155449937 |
| CRYZL2P | 50.73319743 | -1.40251643 | 0.518813895 | -2.703313161 | 0.006865203 | 0.090803834 |
| CS | 5131.681759 | -0.305713445 | 0.279763804 | -1.092755531 | 0.274501151 | 0.587352209 |
| CSAD | 483.6858091 | 0.262199613 | 0.308979021 | 0.84860005 | 0.396103879 | 0.694715833 |
| CSDC2 | 19.70044446 | 0.601049292 | 0.80192117 | 0.749511691 | 0.453548855 | 0.733497409 |
| CSDE1 | 23081.85438 | -0.289388921 | 0.249218596 | -1.161185103 | 0.245566632 | 0.557291155 |
| CSE1L | 9532.078693 | 0.396999639 | 0.352770485 | 1.12537657 | 0.260429495 | 0.573686057 |
| CSF1 | 172.5428732 | 0.029422013 | 0.647747096 | 0.045422069 | 0.96377089 | 0.988130579 |
| CSF2RA | 622.5679931 | -2.012055136 | 0.712249011 | -2.824932159 | 0.004729063 | 0.072837169 |
| CSF2RB | 31.64655718 | -0.938971471 | 0.907975427 | -1.034137536 | 0.301071871 | 0.61178159 |
| CSF3 | 138.425068 | -0.779282428 | 0.514856135 | -1.513592584 | 0.130129203 | 0.415181944 |
| CSF3R | 23.50085464 | -1.467058717 | 0.97952244 | -1.497728543 | 0.134203793 | 0.420619758 |
| CSGALNACT1 | 133.085508 | -2.618544084 | 0.916400917 | -2.857421938 | 0.004270977 | 0.067777184 |
| CSGALNACT2 | 369.8877155 | -0.027124937 | 0.313690444 | -0.086470396 | 0.931092488 | 0.974131023 |
| CSH1 | 20.99345583 | 1.10913281 | 1.485637629 | 0.746570219 | 0.455323031 | 0.735096914 |
| CSK | 1097.297676 | -0.07643664 | 0.245003751 | -0.311981511 | 0.755054575 | 0.897581331 |
| CSKMT | 303.4070215 | 0.667457812 | 0.296512829 | 2.251025073 | 0.024383949 | 0.178492903 |
| CSMD2 | 84.89197778 | 0.514791072 | 0.782765179 | 0.657657092 | 0.5107585 | 0.770556603 |
| CSMD3 | 44.11737696 | -1.829486273 | 0.775767992 | -2.358290484 | 0.01835932 | 0.153593808 |
| CSNK1A1 | 6971.987354 | 0.096509379 | 0.286152003 | 0.337266131 | 0.735916289 | 0.890104921 |
| CSNK1A1P1 | 22.35282457 | 1.881617658 | 1.041493051 | 1.806654068 | 0.070816234 | 0.306170726 |
| CSNK1D | 3170.848033 | -0.361414127 | 0.230336961 | -1.569067009 | 0.116632326 | 0.393839149 |
| CSNK1E | 3650.535252 | 0.182168144 | 0.264322284 | 0.689189505 | 0.49070402 | 0.758593654 |
| CSNK1G1 | 725.2769021 | -0.21946725 | 0.343596547 | -0.63873532 | 0.522995131 | |

0.777662198

| | | | | | | |
|---|---|---|---|---|---|---|
| CSNK1G2 | 1122.145507 | -0.236745074 | 0.514836383 | -0.459845267 | 0.645627289 | 0.845794147 |
| CSNK1G3 | 1390.644444 | 0.220767123 | 0.310352457 | 0.711343242 | 0.476871562 | 0.749746108 |
| CSNK2A1 | 5835.030913 | 0.16550818 | 0.260708436 | 0.634840142 | 0.52553268 | 0.778371208 |
| CSNK2A2 | 3560.287826 | -0.251757467 | 0.264158889 | -0.953053169 | 0.340563132 | 0.647332681 |
| CSPG4 | 50.85365036 | 0.437371828 | 0.58869355 | 0.742953322 | 0.457509941 | 0.736277004 |
| CSPG4P12 | 17.99659299 | -0.584427479 | 0.705750429 | -0.828093693 | 0.407617441 | 0.702164665 |
| CSPG5 | 12.63788026 | 1.291360493 | 0.839292598 | 1.53862967 | 0.123894731 | 0.405045568 |
| CSPP1 | 648.9466508 | 0.040232899 | 0.254779005 | 0.15791293 | 0.874525409 | 0.949740014 |
| CSRNP1 | 512.2423292 | -0.687187763 | 0.409224653 | -1.679243316 | 0.093104633 | 0.349799503 |
| CSRNP2 | 1189.148824 | -0.040797374 | 0.299662967 | -0.136144196 | 0.891707289 | 0.957210151 |
| CSRNP3 | 222.9696124 | -0.106872384 | 0.352819674 | -0.302909366 | 0.761958934 | 0.900847801 |
| CSRP1 | 748.2947101 | -2.274580316 | 0.750198353 | -3.031971888 | 0.002429618 | 0.047414712 |
| CSRP2 | 2198.204206 | 1.400225939 | 0.364450972 | 3.842014559 | 0.000122029 | 0.007000164 |
| CST3 | 1134.30923 | 0.685914053 | 0.410921439 | 1.669209704 | 0.095075828 | 0.353875216 |
| CSTB | 2082.058634 | 0.567204264 | 0.542109652 | 1.046290658 | 0.295426857 | 0.60699582 |
| CSTF1 | 1402.702151 | 0.103708937 | 0.299519661 | 0.346250851 | 0.729154202 | 0.886086417 |
| CSTF2 | 759.0907876 | -0.073668354 | 0.362211142 | -0.203385113 | 0.838834035 | 0.933915137 |
| CSTF2T | 1307.00343 | 0.089408539 | 0.254040502 | 0.351946001 | 0.724878759 | 0.883935898 |
| CSTF3 | 1640.277063 | 0.360337158 | 0.2663494 | 1.352873925 | 0.176095911 | 0.480085722 |
| CTAGE4 | 55.41764707 | -0.843195074 | 2.117662319 | -0.398172582 | 0.690502975 | 0.867938158 |
| CTAGE8 | 14.95449503 | -3.580685656 | 1.310521415 | -2.732260317 | 0.006290142 | 0.086267508 |
| CTBP1 | 2365.10297 | 0.247191387 | 0.435216385 | 0.56797353 | 0.570052942 | 0.806205765 |
| CTBP2 | 3923.065894 | -0.067341913 | 0.248685506 | -0.270791466 | 0.786551424 | 0.911315996 |
| CTBS | 218.6204788 | 0.287462199 | 0.369457028 | 0.778066667 | 0.436529712 | 0.720784523 |
| CTC1 | 747.6577524 | -0.89476968 | 0.335232697 | -2.66910026 | 0.007605474 | 0.097123444 |
| CTCF | 2676.599593 | -0.195263086 | 0.306226075 | -0.637643563 | 0.523705731 | 0.777662198 |
| CTCFL | 20.37107899 | -0.03225542 | 0.85927025 | -0.037538155 | 0.970055918 | 0.989496332 |
| CTDNEP1 | 2393.238236 | 0.031000125 | 0.309408832 | 0.100191467 | 0.920192321 | 0.969770994 |
| CTDP1 | 638.9828233 | -0.355639869 | 0.38642137 | -0.920342138 | 0.357393995 | 0.664952698 |
| CTDSP1 | 1638.337989 | -0.073443529 | 0.287828817 | -0.255163918 | 0.798596498 | 0.916056735 |
| CTDSP2 | 4793.530865 | -0.408737772 | 0.384462821 | -1.063139918 | 0.287718504 | 0.601673803 |
| CTDSPL | 2962.423287 | -0.676829717 | 0.548621494 | -1.233691577 | 0.217317854 | 0.525468356 |
| CTDSPL2 | 1882.232893 | 0.376120031 | 0.352864465 | 1.065905094 | 0.286466546 | 0.600439896 |
| CTF1 | 46.1544395 | -1.016466587 | 0.639587848 | -1.589252502 | 0.112003398 | 0.384920166 |
| CTH | 534.1121402 | 0.252098625 | 0.39028064 | 0.645941919 | 0.518316971 | 0.774169079 |
| CTHRC1 | 398.7990173 | -0.914338084 | 0.697509969 | -1.310860237 | 0.189904987 | 0.495280824 |
| CTIF | 206.8250212 | 0.194617224 | 0.587670812 | 0.331167075 | 0.740518288 | 0.891917214 |
| CTNNA1 | 13736.76992 | -0.638639715 | 0.224197044 | -2.848564388 | 0.004391697 | 0.069186323 |

| | | | | | | |
|---|---|---|---|---|---|---|
| CTNNA2 | 499.1855696 | 0.182612004 | 0.433494055 | 0.421256074 | 0.673568102 | 0.85950893 |
| CTNNA3 | 152.9527858 | 0.223477257 | 0.453480752 | 0.492804284 | 0.622150884 | 0.833640266 |
| CTNNAL1 | 1846.745584 | 0.572795587 | 0.229449144 | 2.496394521 | 0.012546298 | 0.12669427 |
| CTNNB1 | 17410.12683 | -0.278960958 | 0.291390544 | -0.957343895 | 0.338393707 | 0.645594935 |
| CTNNBIP1 | 1773.607749 | 0.340905599 | 0.282418557 | 1.207093479 | 0.227396146 | 0.536186965 |
| CTNNBL1 | 1486.337136 | -0.187529714 | 0.239900669 | -0.781697335 | 0.434392468 | 0.719529221 |
| CTNND1 | 13759.13342 | -1.035804569 | 0.239489966 | -4.325043696 | 1.53E-05 | 0.001503507 |
| CTNND2 | 212.5718179 | 1.262030595 | 0.606178663 | 2.081944932 | 0.0373475 | 0.221199778 |
| CTNS | 397.5524078 | -0.440030401 | 0.285870511 | -1.53926475 | 0.123739676 | 0.405045568 |
| CTPS1 | 2253.547742 | 0.6016204 | 0.323881371 | 1.857533198 | 0.063235333 | 0.289778852 |
| CTPS2 | 1383.154893 | -0.267569062 | 0.254338432 | -1.052019781 | 0.292790474 | 0.605827098 |
| CTR9 | 1741.347018 | -0.094843339 | 0.246725208 | -0.384408793 | 0.700675484 | 0.872001128 |
| CTSA | 981.45795 | -0.501667579 | 0.292373736 | -1.715843516 | 0.086190687 | 0.336662954 |
| CTSB | 3367.833386 | -0.163749469 | 0.285181314 | -0.574194244 | 0.565836364 | 0.804126907 |
| CTSC | 5104.993861 | 0.110473054 | 0.331963405 | 0.332786845 | 0.739295189 | 0.891550525 |
| CTSD | 3332.395363 | -0.413410432 | 0.384181064 | -1.076082272 | 0.281890464 | 0.595704168 |
| CTSH | 220.8809298 | -0.029966593 | 0.403568422 | -0.074254058 | 0.940808232 | 0.97807441 |
| CTSK | 119.7562407 | -0.302389199 | 0.394562664 | -0.766390808 | 0.443443801 | 0.726848446 |
| CTSL | 1062.827268 | -0.105869345 | 0.374887434 | -0.282403024 | 0.777634497 | 0.908018171 |
| CTSO | 266.0507564 | -1.227343305 | 0.394174335 | -3.113706799 | 0.00184753 | 0.039984131 |
| CTSV | 15602.47681 | -2.23926628 | 0.748496189 | -2.991686951 | 0.002774406 | 0.051552064 |
| CTSZ | 1622.445697 | -0.852129299 | 0.349005949 | -2.44158961 | 0.014622759 | 0.137627695 |
| CTTN | 4120.773827 | -0.593610414 | 0.26700482 | -2.223219842 | 0.026200979 | 0.184791771 |
| CTTNBP2 | 1735.283776 | -2.170729583 | 0.543695413 | -3.99254717 | 6.54E-05 | 0.004569989 |
| CTTNBP2NL | 1108.310538 | -0.229890358 | 0.299451834 | -0.767703958 | 0.442663084 | 0.726083385 |
| CTU1 | 48.97805366 | 1.347761656 | 0.470779434 | 2.862830356 | 0.004198753 | 0.066981369 |
| CTU2 | 258.1211088 | 0.627555983 | 0.428317977 | 1.465163775 | 0.14287624 | 0.433233718 |
| CTXN1 | 240.047557 | 1.002693641 | 0.612088112 | 1.638152453 | 0.101389895 | 0.365191382 |
| CU634019.7 | 10.49504699 | -0.962148663 | 0.983974079 | -0.977819115 | 0.328163795 | 0.637787615 |
| CUBN | 21.49946708 | -0.016104663 | 0.995081568 | -0.016184264 | 0.987087389 | 0.995287969 |
| CUEDC1 | 189.2332788 | 0.661452306 | 0.513229797 | 1.288803397 | 0.197466445 | 0.503066907 |
| CUEDC2 | 1550.366615 | 0.966605946 | 0.385334784 | 2.508483499 | 0.012125063 | 0.125055076 |
| CUL1 | 2317.797736 | -0.126039016 | 0.366669711 | -0.343739917 | 0.731041889 | 0.886936313 |
| CUL2 | 1296.932231 | 0.025368449 | 0.295988851 | 0.085707447 | 0.931698982 | 0.974341807 |
| CUL3 | 3056.274143 | -0.028146704 | 0.237686843 | -0.118419277 | 0.905735452 | 0.964545314 |

| | | | | | | |
|---|---|---|---|---|---|---|
| CUL4A | 2445.668654 | 0.063186942 | 0.233119503 | 0.271049573 | 0.786352905 | 0.911294777 |
| CUL4B | 2658.978687 | -0.084035632 | 0.245830748 | -0.341843452 | 0.732468707 | 0.888105842 |
| CUL5 | 1529.333899 | 0.211455867 | 0.232857912 | 0.908089682 | 0.363830842 | 0.670433341 |
| CUL7 | 1797.475459 | -0.367913706 | 0.284284383 | -1.294174875 | 0.195605023 | 0.501359174 |
| CUL9 | 920.9628995 | -0.763284611 | 0.398011659 | -1.917744354 | 0.055143435 | 0.271793713 |
| CUTA | 2133.20556 | 0.710915294 | 0.397574694 | 1.78813014 | 0.073755014 | 0.311905115 |
| CUTALP | 858.4259697 | -0.640342704 | 0.330278338 | -1.938797161 | 0.052526042 | 0.264550665 |
| CUTC | 572.6675189 | 0.565727436 | 0.344170991 | 1.643739451 | 0.100230033 | 0.36243601 |
| CUX1 | 2169.786433 | -0.565593234 | 0.241041252 | -2.346458253 | 0.018952789 | 0.156140235 |
| CUX2 | 145.8362981 | -0.283095756 | 0.363088246 | -0.779688572 | 0.435574207 | 0.720542613 |
| CUZD1 | 31.67269173 | 2.722245144 | 0.870051734 | 3.128831355 | 0.00175503 | 0.039027635 |
| CWC15 | 1837.096549 | 0.775253943 | 0.355922008 | 2.1781568 | 0.029394365 | 0.195932299 |
| CWC22 | 1539.627175 | 0.209188312 | 0.356452408 | 0.586861831 | 0.557296507 | 0.79822426 |
| CWC25 | 572.7194217 | 0.158245983 | 0.298756147 | 0.529682769 | 0.596331897 | 0.821812117 |
| CWC27 | 1887.979895 | 0.552952248 | 0.301729127 | 1.832611433 | 0.066860377 | 0.297817842 |
| CWF19L1 | 636.4849422 | -0.551440901 | 0.263239397 | -2.094826639 | 0.036186406 | 0.217764366 |
| CWF19L2 | 1049.044492 | 0.576577991 | 0.281731229 | 2.046553351 | 0.040701959 | 0.23195346 |
| CX3CL1 | 58.84509404 | 1.207550768 | 0.611912249 | 1.973405124 | 0.048449424 | 0.253705068 |
| CXADR | 5062.665935 | 0.439618958 | 0.317737899 | 1.383589933 | 0.166484053 | 0.46837989 |
| CXCL12 | 386.0116466 | -3.248208068 | 0.77529786 | -4.189625997 | 2.79E-05 | 0.002341988 |
| CXCL14 | 608.9478198 | 1.094862984 | 1.323731462 | 0.827103544 | 0.408178376 | 0.702652234 |
| CXCL5 | 11.85436725 | 0.972741247 | 0.863829033 | 1.126080751 | 0.260131341 | 0.57337005 |
| CXCL6 | 13.38562683 | 0.772860974 | 0.872167919 | 0.886137815 | 0.375543268 | 0.67966045 |
| CXCR4 | 85.76575036 | 2.087799821 | 0.59519246 | 3.507772631 | 0.000451875 | 0.016498831 |
| CXorf38 | 499.6899504 | 0.070308687 | 0.306292536 | 0.229547505 | 0.818443403 | 0.924316427 |
| CXorf40A | 268.1935819 | 0.143230979 | 0.294260421 | 0.486749044 | 0.626436185 | 0.835767952 |
| CXorf40B | 703.2373299 | 0.458535716 | 0.349126388 | 1.313380287 | 0.189054837 | 0.494253132 |
| CXorf56 | 820.5046307 | 0.125433891 | 0.315113774 | 0.398059054 | 0.690586656 | 0.867938875 |
| CXXC1 | 1306.884766 | -0.195741682 | 0.392669959 | -0.498489069 | 0.618139372 | 0.832176095 |
| CXXC4 | 709.8980487 | 0.571614901 | 0.385209774 | 1.483905498 | 0.137833993 | 0.425968529 |
| CXXC5 | 1906.612942 | 0.256152632 | 0.429110464 | 0.596938675 | 0.550548325 | 0.794328924 |
| CYB561 | 658.5047102 | -0.616929038 | 0.400669801 | -1.53974429 | 0.123622696 | 0.405045568 |
| CYB561A3 | 432.5724828 | -0.128951519 | 0.297820125 | -0.43298457 | 0.66502598 | 0.855040241 |
| CYB561D1 | 279.0349843 | -0.460736056 | 0.312902581 | -1.472458472 | 0.140897113 | 0.430713136 |
| CYB561D2 | 277.1358175 | 0.427456837 | 0.32903031 | 1.299141217 | 0.19389547 | 0.499845061 |
| CYB5A | 1054.704273 | 0.631916448 | 0.523336491 | 1.207476372 | 0.227248739 | 0.536089352 |
| CYB5B | 1969.242479 | 0.193877836 | 0.2189225 | 0.885600316 | 0.375832941 | 0.679780547 |
| CYB5D1 | 427.209454 | -0.185475933 | 0.454497269 | -0.408090313 | 0.683207371 | 0.863415274 |
| CYB5D2 | 383.9466821 | -0.166446734 | 0.408054862 | -0.407902833 | 0.683345013 | 0.863415274 |
| CYB5R1 | 434.7257487 | -0.03100016 | 0.415408489 | -0.074625726 | 0.940512505 | 0.97807441 |
| CYB5R2 | 557.0887636 | 0.13942165 | 0.277665439 | 0.502121007 | 0.615582404 | |

| | | | | | | |
|---|---|---|---|---|---|---|
| | | | | | | 0.830968556 |
| CYB5R3 | 1898.064454 | -0.208522844 | 0.25975209 | -0.802776386 | 0.422103993 | 0.712080472 |
| CYB5R4 | 627.7155747 | 0.240716161 | 0.297554348 | 0.808982166 | 0.418525404 | 0.709738117 |
| CYB5RL | 416.7603349 | 0.571336367 | 0.480645633 | 1.188685235 | 0.234563553 | 0.544872709 |
| CYBA | 691.7797252 | 0.182469545 | 0.534894341 | 0.341131942 | 0.733004256 | 0.888353376 |
| CYBC1 | 1092.679933 | -0.26952914 | 0.296043042 | -0.91043903 | 0.362591022 | 0.669723897 |
| CYBRD1 | 80.79379311 | 0.975009082 | 0.564842333 | 1.72616149 | 0.084318364 | 0.332430848 |
| CYC1 | 1969.27805 | 0.451672506 | 0.238977705 | 1.890019431 | 0.058755361 | 0.28066045 |
| CYCS | 4648.151431 | 1.097294056 | 0.369354723 | 2.970840734 | 0.002969858 | 0.053894388 |
| CYFIP1 | 2990.96118 | -0.406451614 | 0.266486735 | -1.525222687 | 0.127203564 | 0.410771662 |
| CYFIP2 | 2128.612136 | 0.226371842 | 0.522741267 | 0.433047582 | 0.664980202 | 0.855040241 |
| CYGB | 10.47907074 | 0.087629531 | 0.884467672 | 0.099076013 | 0.921077917 | 0.970124485 |
| CYHR1 | 996.7179741 | -0.037772533 | 0.395449792 | -0.095517897 | 0.923903476 | 0.971414487 |
| CYLD | 356.0214808 | -0.287469199 | 0.318519176 | -0.902517715 | 0.366781916 | 0.673335968 |
| CYP11A1 | 143.0455028 | -2.468864701 | 0.746269218 | -3.308276211 | 0.000938722 | 0.026083055 |
| CYP1A1 | 50.96582389 | -0.140656842 | 0.691537035 | -0.20339741 | 0.838824424 | 0.933915137 |
| CYP1A2 | 10.09532588 | 0.679130853 | 1.028271166 | 0.660458909 | 0.508959379 | 0.769344882 |
| CYP1B1 | 895.5351507 | -0.075481067 | 0.653558766 | -0.115492394 | 0.908054849 | 0.965583536 |
| CYP20A1 | 355.7141433 | -0.123934978 | 0.383688513 | -0.323009352 | 0.74668816 | 0.894109554 |
| CYP26A1 | 198.4609462 | 2.952895214 | 0.836030181 | 3.532043795 | 0.000412361 | 0.015709208 |
| CYP26B1 | 14.03023637 | 0.124921377 | 0.771067837 | 0.162010877 | 0.871297288 | 0.948581876 |
| CYP26C1 | 134.3843848 | 1.971968768 | 1.009896161 | 1.952645077 | 0.050861666 | 0.259366654 |
| CYP27A1 | 85.65661449 | -0.01484537 | 0.512004804 | -0.028994592 | 0.976868904 | 0.992055261 |
| CYP27B1 | 13.46811088 | -0.251290028 | 0.816607431 | -0.307724396 | 0.758292055 | 0.89938619 |
| CYP27C1 | 452.1681403 | -0.468662301 | 0.401856123 | -1.166244021 | 0.243515799 | 0.554794742 |
| CYP2C18 | 16.39550205 | -2.612268879 | 1.103035331 | -2.368254947 | 0.017872216 | 0.150854036 |
| CYP2C8 | 39.30920791 | -0.783557155 | 0.554956352 | -1.411925735 | 0.157971829 | 0.457277356 |
| CYP2J2 | 11.27797169 | -0.928872436 | 0.971011677 | -0.956602745 | 0.338767803 | 0.645902518 |
| CYP2R1 | 294.9354328 | 0.003404467 | 0.291827377 | 0.011666032 | 0.990692064 | 0.996726674 |
| CYP2S1 | 1575.259255 | -1.522730891 | 0.275301172 | -5.531145698 | 3.18E-08 | 1.79E-05 |
| CYP2T1P | 44.00221086 | -0.000698864 | 0.569140069 | -0.001227931 | 0.999020253 | 0.999324484 |
| CYP2U1 | 169.4545872 | 0.273809632 | 0.409017922 | 0.66943187 | 0.503220027 | 0.765246655 |
| CYP39A1 | 93.59002067 | -0.510602038 | 0.449758119 | -1.135281425 | 0.256257422 | 0.568457575 |
| CYP3A5 | 54.61790622 | 0.167296403 | 0.703903397 | 0.237669549 | 0.812137404 | 0.921390083 |
| CYP46A1 | 128.0207857 | 0.247455403 | 0.399164649 | 0.619933162 | 0.535301792 | 0.784264943 |

| | | | | | | |
|---|---|---|---|---|---|---|
| CYP4V2 | 23.78720118 | 0.226994529 | 0.739271333 | 0.307051713 | 0.758804011 | 0.899440455 |
| CYP51A1 | 1357.802725 | 0.131717692 | 0.237859484 | 0.553762621 | 0.579741313 | 0.811340629 |
| CYP7B1 | 84.5356907 | -0.697667818 | 0.387986836 | -1.798173938 | 0.072149448 | 0.309203186 |
| CYREN | 729.750602 | -0.134029733 | 0.275289518 | -0.486868274 | 0.626351683 | 0.835767952 |
| CYSLTR1 | 38.27791632 | -0.515596382 | 0.576775562 | -0.893928966 | 0.371359899 | 0.67682226 |
| CYSRT1 | 18.65444085 | 0.771944592 | 0.775184914 | 0.995819937 | 0.319337642 | 0.629115732 |
| CYSTM1 | 553.4541732 | 0.690085543 | 0.51425306 | 1.341918205 | 0.179622518 | 0.484443366 |
| CYTH1 | 548.6233361 | -0.012994855 | 0.348546509 | -0.037282986 | 0.970259371 | 0.98957539 |
| CYTH2 | 3579.740534 | -0.255330961 | 0.27570728 | -0.926094375 | 0.354396937 | 0.66172996 |
| CYTH3 | 326.4002686 | -0.459273903 | 0.437961907 | -1.048661759 | 0.294333821 | 0.606538766 |
| CYTIP | 21.61549027 | 0.318350009 | 0.6922812 | 0.459856499 | 0.645619226 | 0.845794147 |
| CYTL1 | 29.03874122 | 1.160733807 | 0.636326358 | 1.824117124 | 0.068134367 | 0.300309844 |
| CYYR1 | 901.5807125 | 0.614134653 | 0.327642418 | 1.874405204 | 0.060874597 | 0.284375094 |
| CZIB | 1429.034601 | 0.472890663 | 0.307848673 | 1.536114021 | 0.124510418 | 0.405999855 |
| D2HGDH | 206.2463868 | 0.077673196 | 0.450837584 | 0.172286427 | 0.863212355 | 0.94479249 |
| DAAM1 | 1880.853179 | -0.138047177 | 0.390143372 | -0.353837042 | 0.723461012 | 0.883279173 |
| DAAM2 | 173.5105194 | -0.508633032 | 0.488140808 | -1.04198015 | 0.297420882 | 0.608930122 |
| DAB1 | 34.47197077 | -0.116921988 | 0.604211613 | -0.193511653 | 0.846558281 | 0.938145276 |
| DAB2 | 369.3086839 | -0.456504366 | 0.434763863 | -1.050005313 | 0.29371567 | 0.606359511 |
| DAB2IP | 1872.085824 | -0.228315066 | 0.418394239 | -0.545693617 | 0.585276561 | 0.815432095 |
| DACH1 | 1321.42876 | 0.256745854 | 0.351034805 | 0.731397143 | 0.46453661 | 0.741528294 |
| DACH2 | 23.84702576 | 0.447746692 | 0.773385693 | 0.578943593 | 0.562627235 | 0.801706045 |
| DACT1 | 2659.950111 | -2.278036003 | 0.687746122 | -3.312321116 | 0.000925253 | 0.02597297 |
| DACT2 | 67.03558805 | -0.78211094 | 0.626064471 | -1.249249839 | 0.211573708 | 0.51879501 |
| DACT3 | 62.06511325 | 1.500703767 | 0.499938533 | 3.001776556 | 0.002684091 | 0.050458917 |
| DAD1 | 2337.51867 | 0.762685556 | 0.342313281 | 2.228033788 | 0.025878262 | 0.183710607 |
| DAG1 | 6935.216546 | -0.667224183 | 0.245506144 | -2.717749434 | 0.006572759 | 0.088793705 |
| DAGLA | 109.8210108 | -0.192405461 | 0.657482818 | -0.292639527 | 0.769797698 | 0.904694332 |
| DAGLB | 694.5567875 | -0.265003612 | 0.253871881 | -1.043847832 | 0.296555797 | 0.607815185 |
| DALRD3 | 707.1149951 | -0.060778218 | 0.265059153 | -0.229300583 | 0.8186353 | 0.924316427 |
| DAP | 1540.420403 | -0.351130858 | 0.269939103 | -1.300778045 | 0.193334439 | 0.499293687 |
| DAP3 | 2948.280599 | 0.183501956 | 0.335052045 | 0.547681946 | 0.583910311 | 0.81442524 |
| DAPK1 | 3217.417439 | -1.188386382 | 0.264870755 | -4.486665131 | 7.23E-06 | 0.000857477 |
| DAPK2 | 42.37593621 | 0.972127333 | 0.596862179 | 1.628729992 | 0.103370191 | 0.368734814 |
| DAPK3 | 723.9427184 | -0.529268505 | 0.33963387 | -1.558350188 | 0.119150258 | 0.398078527 |
| DARS | 3515.098912 | 0.317955465 | 0.232114833 | 1.369819676 | 0.170743198 | 0.473986265 |
| DARS2 | 1085.55973 | -0.04887544 | 0.339587007 | -0.143926119 | 0.88555881 | 0.954787344 |
| DAXX | 859.5226892 | -0.018323765 | 0.436409092 | -0.041987587 | 0.966508594 | |

| | | | | | | |
|---|---|---|---|---|---|---|
| | | | | | | 0.988700292 |
| DAZAP1 | 4327.447746 | 0.127105881 | 0.3025256 | 0.420149173 | 0.674376482 | 0.859672768 |
| DAZAP2 | 1304.898733 | 0.028520698 | 0.31798794 | 0.089691132 | 0.928532663 | 0.972633911 |
| DAZL | 84.24902677 | 2.240333206 | 0.645794765 | 3.469110202 | 0.000522185 | 0.017923239 |
| DBF4 | 1165.311927 | 0.741929653 | 0.276993812 | 2.678506231 | 0.007395136 | 0.095319968 |
| DBF4B | 693.6959981 | -0.177461542 | 0.520943931 | -0.340653824 | 0.733364204 | 0.888457729 |
| DBI | 5356.13084 | 0.750363211 | 0.522489362 | 1.436131078 | 0.150965046 | 0.446102593 |
| DBIL5P | 48.54782874 | 0.123913369 | 0.499314763 | 0.248166844 | 0.804005318 | 0.918076523 |
| DBN1 | 7417.785573 | -0.09922874 | 0.40587665 | -0.244480042 | 0.806859057 | 0.919137032 |
| DBNDD1 | 1319.302186 | -0.462701959 | 0.314263942 | -1.472335502 | 0.140930301 | 0.430713136 |
| DBNDD2 | 595.2574615 | 0.194005444 | 0.596558581 | 0.3252077 | 0.745023882 | 0.893216162 |
| DBNL | 1226.974705 | -0.315386614 | 0.34534014 | -0.913263699 | 0.361103869 | 0.668004731 |
| DBP | 296.3608229 | 0.686962554 | 0.443446589 | 1.549143845 | 0.121347145 | 0.401692676 |
| DBR1 | 564.7308218 | 0.129587035 | 0.253091449 | 0.512016649 | 0.608639358 | 0.827908876 |
| DBT | 1176.315536 | -0.154485504 | 0.245476794 | -0.629328344 | 0.529134119 | 0.779974131 |
| DBX1 | 12.43915683 | 2.112395127 | 1.42922899 | 1.477996278 | 0.139408775 | 0.428072582 |
| DCAF1 | 958.2461729 | -0.225467417 | 0.449489831 | -0.501607382 | 0.615943726 | 0.831265761 |
| DCAF10 | 839.3761429 | -0.04430731 | 0.274896993 | -0.161177863 | 0.871953316 | 0.948884854 |
| DCAF11 | 1124.262989 | -0.403041151 | 0.294250382 | -1.369721759 | 0.170773774 | 0.473986265 |
| DCAF12 | 1429.61636 | 0.000559261 | 0.253452213 | 0.002206574 | 0.99823941 | 0.998939187 |
| DCAF12L1 | 20.59741082 | 0.639565005 | 0.657490105 | 0.972737082 | 0.33068399 | 0.63900227 |
| DCAF12L2 | 13.65980705 | 0.228680479 | 0.896894912 | 0.254969089 | 0.798746974 | 0.916056735 |
| DCAF13 | 2777.41814 | 0.671826279 | 0.244023266 | 2.753123874 | 0.005902956 | 0.082996978 |
| DCAF15 | 1025.500756 | 0.075549898 | 0.377559255 | 0.200100771 | 0.841401771 | 0.935153081 |
| DCAF16 | 1584.190245 | 0.114418475 | 0.35532183 | 0.322013638 | 0.747442363 | 0.894202292 |
| DCAF17 | 823.0457919 | 0.062132653 | 0.363648635 | 0.170859031 | 0.864334611 | 0.945136254 |
| DCAF4 | 371.5235143 | -0.554754262 | 0.274136082 | -2.023645547 | 0.043006635 | 0.238392703 |
| DCAF5 | 1675.931418 | -0.646530376 | 0.271179982 | -2.38413754 | 0.017119203 | 0.14773715 |
| DCAF6 | 686.2212692 | -0.877388448 | 0.392810596 | -2.233617059 | 0.025508281 | 0.182111117 |
| DCAF7 | 3282.096857 | 0.013123096 | 0.24963695 | 0.052568723 | 0.958075538 | 0.985220903 |
| DCAF8 | 2877.826237 | -0.343404933 | 0.263311725 | -1.304176383 | 0.19217345 | 0.498075763 |
| DCAKD | 863.1219641 | 0.072741036 | 0.231374639 | 0.314386382 | 0.753227598 | 0.896641779 |
| DCBLD1 | 105.2530911 | -0.745012338 | 0.489327851 | -1.522521837 | 0.127878378 | 0.412073677 |
| DCBLD2 | 2160.216579 | 0.040329908 | 0.25401416 | 0.158770316 | 0.873849837 | 0.949740014 |
| DCDC1 | 39.55900834 | 0.231339595 | 0.850012748 | 0.272160147 | 0.785498883 | 0.910931322 |
| DCDC2 | 40.08079413 | -3.198100508 | 0.84879527 | -3.767811417 | 0.000164685 | 0.008469277 |
| DCHS1 | 539.788633 | 0.418804856 | 0.605296942 | 0.691899838 | 0.489000232 | 0.757388721 |
| DCK | 648.2579223 | 0.640566311 | 0.278366343 | 2.301162933 | 0.021382423 | 0.16601471 |

| | | | | | | |
|---|---|---|---|---|---|---|
| DCLK1 | 206.9654204 | -0.454311978 | 0.544503829 | -0.834359564 | 0.404078396 | 0.700367583 |
| DCLK2 | 44.63906497 | 1.440543707 | 0.94130028 | 1.530376372 | 0.125923594 | 0.408202317 |
| DCLRE1A | 786.6275027 | 0.291105374 | 0.409507681 | 0.710866699 | 0.477166844 | 0.749854795 |
| DCLRE1B | 417.9543998 | -0.278919788 | 0.302234954 | -0.922857482 | 0.356081481 | 0.663487122 |
| DCLRE1C | 339.3691624 | -0.309345786 | 0.440099615 | -0.702899469 | 0.482118402 | 0.753266558 |
| DCN | 15.24213907 | -5.138643607 | 1.871360171 | -2.745940459 | 0.006033771 | 0.083980132 |
| DCP1A | 1106.536671 | -0.179779062 | 0.250273944 | -0.718329118 | 0.47255438 | 0.746939206 |
| DCP1B | 234.3651039 | -0.008516408 | 0.372909126 | -0.022837758 | 0.981779689 | 0.993518428 |
| DCP2 | 3405.962583 | 0.492939389 | 0.390910774 | 1.261002308 | 0.207308016 | 0.514236414 |
| DCPS | 762.832402 | 0.320447998 | 0.299462476 | 1.070077301 | 0.284584516 | 0.598479579 |
| DCST2 | 29.79688411 | -0.314817957 | 0.688724508 | -0.457102882 | 0.647597099 | 0.846597949 |
| DCTD | 1585.849675 | 0.222742838 | 0.277133587 | 0.803738156 | 0.421548213 | 0.711966271 |
| DCTN1 | 5668.362612 | -0.721813762 | 0.343618098 | -2.100627896 | 0.035673643 | 0.216083122 |
| DCTN2 | 3751.12576 | -0.121162914 | 0.270509311 | -0.447906629 | 0.654220584 | 0.849068645 |
| DCTN3 | 740.3897848 | 0.557972301 | 0.296754093 | 1.880251407 | 0.060073822 | 0.283477437 |
| DCTN4 | 2239.450466 | -0.125563704 | 0.233258064 | -0.538303808 | 0.590367322 | 0.818321084 |
| DCTN5 | 2610.336413 | -0.262280956 | 0.302055955 | -0.868319102 | 0.385219668 | 0.685935162 |
| DCTN6 | 964.6730046 | 0.808974904 | 0.343170859 | 2.357353146 | 0.018405734 | 0.153593808 |
| DCTPP1 | 1490.384425 | 1.112049789 | 0.362242515 | 3.069904121 | 0.002141275 | 0.043839788 |
| DCUN1D1 | 1281.171698 | 0.148702393 | 0.245498685 | 0.605715639 | 0.544703597 | 0.790739196 |
| DCUN1D2 | 608.9691177 | 0.015954155 | 0.331947989 | 0.048062212 | 0.961666662 | 0.9870991 |
| DCUN1D3 | 325.2684587 | -0.069220676 | 0.315629873 | -0.219309647 | 0.826408848 | 0.927806455 |
| DCUN1D4 | 1265.230355 | 0.689392179 | 0.423918653 | 1.626236956 | 0.103899257 | 0.369319502 |
| DCUN1D5 | 1376.002853 | 0.723813252 | 0.354418692 | 2.042254738 | 0.041126269 | 0.232946328 |
| DCX | 86.86113805 | -0.722888008 | 0.568568796 | -1.271416954 | 0.203580354 | 0.509986087 |
| DCXR | 1122.115904 | 1.113377112 | 0.39129982 | 2.845329987 | 0.004436544 | 0.069675779 |
| DDA1 | 793.3466978 | 0.261836864 | 0.35098272 | 0.746010699 | 0.455660952 | 0.735306344 |
| DDAH1 | 793.722568 | 0.015847631 | 0.243128397 | 0.065182148 | 0.948028974 | 0.981274993 |
| DDB1 | 11705.34219 | -0.544642999 | 0.258480184 | -2.107097696 | 0.035109112 | 0.213997725 |
| DDB2 | 2085.20959 | 1.171054313 | 0.396355221 | 2.954557561 | 0.003131177 | 0.055756493 |
| DDC | 19.45375612 | -1.967773648 | 0.935593693 | -2.103235263 | 0.035445209 | 0.215440412 |
| DDHD1 | 1470.558363 | -1.128915059 | 0.2594425 | -4.35131121 | 1.35E-05 | 0.001389524 |
| DDHD2 | 1733.254512 | -0.226299193 | 0.271663884 | -0.833011695 | 0.404838134 | 0.70065889 |
| DDI2 | 2303.811386 | -0.820996984 | 0.317725327 | -2.58398344 | 0.009766648 | 0.112404636 |
| DDIAS | 596.2748062 | 0.190603596 | 0.360507122 | 0.528709656 | 0.597006878 | 0.821812117 |
| DDIT3 | 141.1101989 | 0.041334602 | 0.670289446 | 0.061666795 | 0.950828183 | 0.981826318 |
| DDIT4 | 2634.191834 | 0.682040233 | 0.423588913 | 1.610146566 | 0.107365863 | 0.375442174 |
| DDIT4L | 127.8741754 | -1.012397656 | 0.540130754 | -1.874356622 | 0.060881288 | 0.284375094 |
| DDN | 23.42590603 | 0.825840265 | 0.818648877 | 1.00878446 | 0.313078015 | 0.622994042 |
| DDOST | 7379.184286 | -0.345361433 | 0.232499242 | -1.485430361 | 0.137429858 | |

| | | | | | | |
|---|---|---|---|---|---|---|
| | | | | | | 0.425712285 |
| DDR1 | 422.330218 | 0.430275272 | 1.661153919 | 0.259021917 | 0.79561833 | 0.915261082 |
| DDRGK1 | 788.928069 | 0.027616458 | 0.384549626 | 0.071815068 | 0.942749081 | 0.978574897 |
| DDT | 279.1573908 | 0.609899882 | 0.48289535 | 1.263006326 | 0.206586906 | 0.513118853 |
| DDX1 | 4348.705317 | 0.35410672 | 0.228509895 | 1.549634079 | 0.121229369 | 0.401420514 |
| DDX10 | 1239.755428 | 0.524788627 | 0.297370911 | 1.764761138 | 0.077603919 | 0.319405602 |
| DDX11 | 2166.83965 | -0.037739499 | 0.360487589 | -0.104690148 | 0.91662168 | 0.969211928 |
| DDX11L2 | 21.74916227 | 0.208146468 | 0.725529908 | 0.286888887 | 0.7741974 | 0.906279513 |
| DDX12P | 140.3719579 | -0.103095772 | 0.928738806 | -0.111006206 | 0.911611425 | 0.966866396 |
| DDX17 | 15965.14289 | -0.260114098 | 0.315834191 | -0.823578022 | 0.410179368 | 0.704109399 |
| DDX18 | 3201.25077 | 0.21445661 | 0.219767968 | 0.975831972 | 0.329147738 | 0.638424836 |
| DDX19A | 1767.683132 | -0.088703282 | 0.249000371 | -0.35623755 | 0.721662673 | 0.882498649 |
| DDX19B | 1164.879371 | -0.374697889 | 0.252370282 | -1.484714784 | 0.137619394 | 0.425842052 |
| DDX20 | 887.4125016 | 0.052158527 | 0.281898512 | 0.185025905 | 0.853208717 | 0.940799171 |
| DDX21 | 5523.759909 | 0.356829419 | 0.286787436 | 1.244229606 | 0.21341507 | 0.520617468 |
| DDX23 | 2559.156833 | -0.515588725 | 0.29755024 | -1.732778724 | 0.083135006 | 0.330129196 |
| DDX24 | 401.9059757 | 0.815473247 | 0.717606906 | 1.13637876 | 0.255798081 | 0.567999769 |
| DDX25 | 129.2881069 | 0.495184665 | 0.458608456 | 1.079754764 | 0.2802514 | 0.594060918 |
| DDX27 | 2028.920961 | -0.482126978 | 0.239574305 | -2.012431915 | 0.044174427 | 0.241589846 |
| DDX28 | 355.9232219 | 0.247829833 | 0.388671905 | 0.637632485 | 0.523712944 | 0.777662198 |
| DDX31 | 568.2155957 | 0.111569041 | 0.28325787 | 0.39387799 | 0.693671121 | 0.868792585 |
| DDX39A | 1631.6525 | 0.450163758 | 0.225315208 | 1.99792887 | 0.045724372 | 0.246249361 |
| DDX39B | 148.8467597 | 0.247769518 | 0.55053353 | 0.450053457 | 0.652671896 | 0.8486189 |
| DDX3X | 7812.634506 | -0.382192576 | 0.296788423 | -1.287761066 | 0.197829149 | 0.503399198 |
| DDX3Y | 2718.346665 | 0.062954937 | 0.355531834 | 0.177072573 | 0.859451385 | 0.943059234 |
| DDX41 | 1798.712951 | -0.312390037 | 0.36678873 | -0.851689299 | 0.394386561 | 0.693260965 |
| DDX42 | 4522.149613 | -0.457801869 | 0.250242263 | -1.829434654 | 0.06733452 | 0.298083457 |
| DDX46 | 4251.7159 | -0.046868196 | 0.243120309 | -0.192777792 | 0.847132997 | 0.938145276 |
| DDX47 | 1777.795788 | 0.407422296 | 0.270615444 | 1.505539706 | 0.132185377 | 0.418485882 |
| DDX49 | 575.1221347 | -0.223870625 | 0.456156712 | -0.490775691 | 0.623585105 | 0.834324864 |
| DDX5 | 16737.9094 | -0.092716292 | 0.252575286 | -0.367083785 | 0.713556522 | 0.878044121 |
| DDX50 | 2395.515067 | 0.388208071 | 0.290211715 | 1.33767195 | 0.181003399 | 0.485674312 |
| DDX50P1 | 12.23843427 | -0.213683504 | 0.875114146 | -0.244177865 | 0.807093069 | 0.919137032 |
| DDX51 | 804.626974 | -0.067875365 | 0.249205771 | -0.272366748 | 0.785340038 | 0.910931322 |
| DDX52 | 977.9682432 | -0.483428099 | 0.3540089 | -1.365581766 | 0.172070273 | 0.475410327 |
| DDX54 | 1636.278291 | 0.183849225 | 0.410999502 | 0.447322258 | 0.654642398 | 0.849149183 |
| DDX55 | 825.4889981 | 0.124807967 | 0.268526936 | 0.464787516 | 0.642083629 | 0.844148268 |
| DDX56 | 1495.233992 | -0.207537825 | 0.236410934 | -0.877868979 | 0.380014826 | 0.68184242 |
| DDX58 | 145.9671346 | -0.332087872 | 0.362771017 | -0.915420076 | 0.359971143 | 0.66745566 |
| DDX59 | 413.471074 | 0.073306814 | 0.27605223 | 0.265554146 | 0.790582604 | 0.91311918 |
| DDX6 | 2887.420425 | -0.413546679 | 0.319414324 | -1.294702983 | 0.195422711 | 0.50126155 |
| DDX60 | 52.92891584 | -0.022140393 | 0.667998283 | -0.033144385 | 0.973559448 | |

0.990601635
DDX60L 30.74800622 0.229178671 0.596101192 0.384462695 0.70063554
0.872001128
DEAF1 756.4826379 0.208514653 0.53063108 0.392955975 0.694351998 0.86892666
DEC1 48.136141 -3.016984871 0.786230357 -3.837278534 0.000124405
0.007078495
DECR1 657.1748898 -0.161395866 0.359686665 -0.448712397 0.65363914
0.84883826
DECR2 256.6383452 0.271912845 0.379950052 0.715654186 0.474204906 0.748145033
DEDD 1069.576543 -0.379533159 0.258009089 -1.471006933 0.141289242
0.431039476
DEDD2 465.0880983 0.175935022 0.371706263 0.473317347 0.635986788 0.841016203
DEF6 129.570702 -0.71812229 0.62783864 -1.143800722 0.252706291
0.565423035
DEF8 1399.956191 -0.101627764 0.326510667 -0.311254038 0.755607505
0.897896779
DEGS1 1082.562807 0.637621856 0.297509794 2.143196183 0.032097349 0.204342793
DEK 8111.108419 0.742883869 0.284820774 2.608250298 0.009100638 0.107863893
DELE1 1717.367257 -0.156118242 0.292280227 -0.534138909 0.593245432
0.819859199
DENND1A 432.3098408 -0.203799821 0.328235478 -0.620895163
0.534668604 0.784009474
DENND1B 111.8722041 -1.053561186 0.447440231 -2.354641161 0.0185406
0.154302741
DENND1C 36.3864204 -0.383932645 0.527228618 -0.728209039
0.466485633 0.743017201
DENND2A 469.4798373 0.43389499 0.485430319 0.893835785 0.37140976
0.67682226
DENND2C 41.19113108 -0.336280752 0.520115741 -0.646549846
0.517923327 0.773950969
DENND2D 44.23312299 -1.064878138 0.575927117 -1.848980725
0.064460596 0.293286926
DENND3 83.99040285 -0.826668571 0.413162573 -2.000831208
0.045410583 0.245452751
DENND4A 237.6119349 -0.329627198 0.348484283 -0.945888276
0.344205574 0.65118919
DENND4B 923.2819974 -0.339397112 0.240039028 -1.413924704
0.157384012 0.456731315
DENND4C 909.9547266 -0.710471283 0.581010275 -1.222820514
0.221397496 0.530451896
DENND5A 2222.324868 -0.164106725 0.453319482 -0.362011191
0.717343668 0.880516207
DENND5B 760.3226669 0.211163203 0.409689959 0.515421962 0.606258189
0.826611695
DENND6A 581.2892979 0.021337781 0.289788688 0.073632207 0.941303043
0.978385069
DENND6B 103.6089286 0.39025816 0.49820891 0.783322321 0.433437859
0.71890019
DENR 3762.71415 0.103125306 0.348734665 0.295712806 0.767449416 0.903747194
DEPDC1 1654.124198 0.895220275 0.364273137 2.457552269 0.013988745
0.134477513
DEPDC1B 1486.949785 1.300287465 0.332609991 3.909345775 9.25E-05
0.005825816
DEPDC4 104.0812637 0.15727896 0.375423268 0.418937699 0.675261664
0.860095396
DEPDC5 430.172928 -0.325305591 0.433673461 -0.750116436
0.453184581 0.733497409
DEPDC7 48.16892069 0.906748168 0.512199941 1.770301193 0.076676979
0.317958513
DEPP1 103.4342889 -1.498722811 0.757657499 -1.978100675 0.04791735
0.252590036
DEPTOR 16.59938599 -0.964797478 0.772866423 -1.248336645 0.2119078
0.519175172
DERA 1707.799557 -1.483127629 0.349542501 -4.243053782 2.20E-05

0.002015171
DERL1 1334.835254 -0.079525457 0.240250114 -0.331011111 0.740636093
0.891917214
DERL2 1040.972004 0.278781915 0.337698773 0.825534283 0.409068327 0.703484892
DERL3 42.33246035 1.014327878 0.48117485 2.108023473 0.035028959 0.213680876
DERPC 1355.968111 -0.521719111 0.284951026 -1.830907991 0.067114277
0.297817842
DES 62.69170009 2.089705227 0.584784928 3.573459453 0.000352296 0.013991308
DESI1 948.0905337 0.250040321 0.234999897 1.064001831 0.28732787 0.601618729
DESI2 2097.904778 0.367418106 0.345046799 1.064835571 0.286950345 0.601236469
DET1 306.2042771 -0.215872739 0.327116611 -0.659925948 0.509301351
0.769562053
DEXI 438.2227192 0.40610749 0.299541344 1.355764397 0.175174143 0.479186245
DFFA 3272.651494 0.229275815 0.391747686 0.585263994 0.558370221 0.798855459
DFFB 123.4488266 -0.139857685 0.501334967 -0.278970538 0.780267435
0.909103544
DGAT1 264.5081314 0.242693973 0.339782255 0.714263237 0.47506442 0.748791879
DGAT2 90.24172464 0.289755819 0.675339494 0.429052086 0.66788532 0.856696248
DGCR2 1696.488457 -0.106406443 0.375670515 -0.283244063 0.776989753
0.907794497
DGCR5 48.65163765 -0.254028233 0.609032116 -0.417101539 0.676604139
0.860199564
DGCR6 182.4863967 -1.001792126 0.569399572 -1.75938335 0.078512418
0.321574485
DGCR6L 490.5515974 0.528025879 0.461114765 1.14510729 0.252164716
0.564998953
DGCR8 1207.938801 -0.37700325 0.312971904 -1.204591357 0.228361095
0.537209827
DGKA 450.4464594 -0.513090935 0.3446693 -1.488647045 0.136580336
0.424728769
DGKD 502.3178287 -0.90073404 0.52787439 -1.706341617 0.087944494
0.340967522
DGKE 210.7751582 -1.599008473 0.582043879 -2.747230115 0.006010095
0.083881464
DGKH 222.9200547 0.603558442 0.583009034 1.035247151 0.300553504 0.611214323
DGKI 114.1168852 1.118067546 0.749543061 1.491665528 0.13578685 0.423669306
DGKK 16.66404349 -1.261894095 0.847432761 -1.489078725 0.136466639
0.424510024
DGKQ 185.3628072 -0.103929025 0.493081024 -0.21077474 0.83306305
0.931554473
DGKZ 1510.748901 -0.484489323 0.51652086 -0.937985975 0.348251617
0.654732583
DGLUCY 488.0032613 0.30015282 0.471426814 0.636690174 0.524326675
0.777662198
DGUOK 1422.928965 0.847022733 0.425143997 1.992319632 0.046335999 0.247615641
DHCR24 5095.841346 -0.310460548 0.429377878 -0.723047377
0.469650778 0.744930517
DHCR7 1710.041417 0.001622362 0.285793503 0.005676695 0.995470677 0.998116797
DHDDS 999.7081422 -0.318417197 0.243033377 -1.310178878 0.190135329
0.495711133
DHDH 15.43769078 1.003541871 0.993836817 1.009765239 0.312607778 0.622907489
DHFR 3202.590279 0.827846529 0.217046593 3.814142021 0.000136657 0.007484554
DHFR2 268.6660535 0.438776702 0.319454686 1.373517812 0.169591428 0.47234653
DHFRP1 158.2763343 -0.311091777 2.209490422 -0.140797975
0.888029543 0.955943827
DHODH 426.4839561 0.254362789 0.26646273 0.954590494 0.339784826 0.646629378
DHPS 1536.233891 0.567371592 0.336634595 1.685422712 0.091907045 0.347971494
DHRS1 162.3862726 -0.182575301 0.396531058 -0.460431275 0.64520669
0.845794147
DHRS11 441.7273899 0.216697482 0.275924344 0.785351083 0.432247753
0.718639059
DHRS12 184.926946 0.06449849 0.35559937 0.181379652 0.85606959
0.942245037
DHRS13 380.2449767 0.655398638 0.406349406 1.612894291 0.106767476

0.374599763

| | | | | | | |
|---|---|---|---|---|---|---|
| DHRS2 | 12.60538618 | -0.529302881 | 0.832188516 | -0.636037233 | 0.524752154 | 0.777746671 |
| DHRS3 | 362.0122474 | -0.437605113 | 0.588249765 | -0.743910392 | 0.456930688 | 0.735735031 |
| DHRS4 | 74.67756095 | 0.019496698 | 0.452226219 | 0.04311271 | 0.965611688 | 0.988700292 |
| DHRS4L2 | 134.1272094 | 0.811634733 | 0.60208627 | 1.348037272 | 0.177646395 | 0.482042363 |
| DHRS7 | 877.8107011 | -0.059138605 | 0.537278779 | -0.110070614 | 0.912353373 | 0.967315188 |
| DHRS7B | 354.8956823 | -0.029631981 | 0.388953411 | -0.076183883 | 0.939272805 | 0.977796709 |
| DHRSX | 420.3738062 | 0.116765661 | 0.332520384 | 0.351153392 | 0.725473274 | 0.88422336 |
| DHTKD1 | 2812.760405 | -0.051694791 | 0.302113691 | -0.171110389 | 0.864136967 | 0.9451245 |
| DHX15 | 5917.298916 | 0.093200461 | 0.32291526 | 0.288622041 | 0.772870627 | 0.905717252 |
| DHX16 | 761.283123 | -0.671879831 | 0.653495743 | -1.028131917 | 0.303887779 | 0.61411513 |
| DHX29 | 1379.072554 | -0.083216637 | 0.285522786 | -0.291453575 | 0.770704445 | 0.905067213 |
| DHX30 | 2459.203177 | 0.001080727 | 0.379131355 | 0.002850534 | 0.997725606 | 0.998851994 |
| DHX32 | 1177.624814 | 0.162646464 | 0.280218475 | 0.580427341 | 0.561626472 | 0.801570466 |
| DHX33 | 1590.843373 | 0.082026239 | 0.291812641 | 0.281092137 | 0.778639732 | 0.908183546 |
| DHX34 | 526.2817011 | -0.258440424 | 0.475183827 | -0.543874622 | 0.586527754 | 0.816200014 |
| DHX35 | 626.0608728 | 0.124386246 | 0.309540416 | 0.401841697 | 0.687800533 | 0.866689498 |
| DHX36 | 1839.290975 | -0.254998408 | 0.366376639 | -0.696000729 | 0.486428373 | 0.756340124 |
| DHX37 | 628.0299884 | -0.403084916 | 0.362717024 | -1.111293071 | 0.266442221 | 0.579775521 |
| DHX38 | 1871.729598 | -0.466302686 | 0.34339512 | -1.357918789 | 0.174489455 | 0.478578744 |
| DHX40 | 2069.377282 | -0.174998049 | 0.312791583 | -0.559471732 | 0.575839818 | 0.808969969 |
| DHX40P1 | 30.14839402 | -0.927767508 | 0.789619456 | -1.174955229 | 0.240012628 | 0.551618176 |
| DHX57 | 1135.841375 | -0.10920052 | 0.258422786 | -0.42256537 | 0.672612397 | 0.859011927 |
| DHX58 | 29.06605019 | 0.837603209 | 0.693771723 | 1.207318172 | 0.227309635 | 0.536149638 |
| DHX8 | 1446.225738 | -0.480060868 | 0.354421554 | -1.35449118 | 0.175579725 | 0.47969026 |
| DHX9 | 7739.577736 | -0.217508969 | 0.321065482 | -0.677459838 | 0.498114244 | 0.762537616 |
| DIABLO | 1618.549266 | 0.167936983 | 0.289565937 | 0.579961112 | 0.561940842 | 0.801706045 |
| DIAPH1 | 2963.98876 | -0.742437829 | 0.276417193 | -2.685932162 | 0.007232777 | 0.094039138 |
| DIAPH2 | 683.1804224 | -0.804102263 | 0.335741244 | -2.395005911 | 0.016620097 | 0.146056278 |
| DIAPH3 | 602.1031363 | -0.165940218 | 0.296873705 | -0.558958964 | 0.576189727 | 0.808993188 |
| DICER1 | 1704.266029 | -0.277945099 | 0.395669366 | -0.702468078 | 0.482387303 | 0.753310328 |
| DIDO1 | 1582.536123 | -0.286362179 | 0.326818685 | -0.876211161 | 0.38091525 | 0.682545212 |
| DIMT1 | 822.5517994 | 0.626222226 | 0.243984142 | 2.566651347 | 0.01026858 | 0.11507205 |
| DIO2 | 99.86089018 | -2.255321226 | 0.568837624 | -3.964789126 | 7.35E-05 | 0.004931304 |
| DIO3 | 61.07528543 | -0.751786499 | 0.915385475 | -0.821278597 | 0.411487596 | 0.704686727 |
| DIP2A | 1469.270461 | -0.132113681 | 0.344238979 | -0.383784779 | 0.70113797 | 0.872239782 |
| DIP2B | 1489.524075 | -0.338054064 | 0.399251633 | -0.8467193 | 0.397151598 | 0.695668733 |
| DIP2C | 588.8123671 | 0.021776891 | 0.359694265 | 0.06054278 | 0.951723345 | 0.982350991 |

| | | | | | | |
|---|---|---|---|---|---|---|
| DIPK1A | 139.1574851 | -0.436103594 | 0.390499419 | -1.116784234 | 0.264086591 | 0.577448862 |
| DIPK1B | 685.5032108 | 0.592264585 | 0.388389942 | 1.524922561 | 0.127278414 | 0.410838472 |
| DIPK2A | 392.5076545 | 0.309111849 | 0.255274802 | 1.210898399 | 0.225934346 | 0.534727976 |
| DIPK2B | 11.08796935 | -0.315874132 | 0.953320312 | -0.331341028 | 0.740386904 | 0.891917214 |
| DIRAS1 | 187.7274446 | 0.665579498 | 0.454654989 | 1.463922126 | 0.143215227 | 0.433589745 |
| DIRAS2 | 94.03874941 | -3.669822608 | 0.977732897 | -3.753399952 | 0.000174452 | 0.008728094 |
| DIRAS3 | 41.69022748 | 0.429656396 | 0.669786694 | 0.641482429 | 0.521209293 | 0.776211464 |
| DIS3 | 2303.763184 | -0.016372578 | 0.236813699 | -0.069136953 | 0.944880607 | 0.979553348 |
| DIS3L | 454.0304309 | -0.506552973 | 0.381043347 | -1.329384118 | 0.183721275 | 0.488132305 |
| DIS3L2 | 627.9544465 | -0.307944568 | 0.404076732 | -0.762094283 | 0.446003734 | 0.728501578 |
| DISC1 | 94.73929981 | -0.496750539 | 0.394488481 | -1.259226979 | 0.207948359 | 0.514730716 |
| DISP1 | 290.5734244 | -0.946837781 | 0.46955589 | -2.016453849 | 0.04375254 | 0.240932768 |
| DISP3 | 103.8454341 | -0.035787772 | 0.481645588 | -0.074303124 | 0.940769191 | 0.97807441 |
| DIXDC1 | 609.096162 | 0.281740038 | 0.404306745 | 0.696847237 | 0.485898401 | 0.75593662 |
| DKC1 | 2411.965575 | 0.373410624 | 0.293288977 | 1.273183287 | 0.20295301 | 0.509506436 |
| DKK1 | 284.6981476 | 1.177452883 | 0.882328603 | 1.33448341 | 0.18204548 | 0.486131134 |
| DKK3 | 1063.288851 | -0.988089136 | 0.315441148 | -3.132404076 | 0.001733811 | 0.038738796 |
| DKKL1 | 37.76008521 | 0.933229251 | 0.507262426 | 1.8397366 | 0.065806917 | 0.29619856 |
| DLAT | 2084.577263 | 0.184660651 | 0.264746158 | 0.697500777 | 0.485489453 | 0.755840687 |
| DLC1 | 248.2008029 | 0.171451787 | 0.808112885 | 0.212163164 | 0.831979743 | 0.931142584 |
| DLD | 3008.311197 | 0.159989899 | 0.219689564 | 0.728254435 | 0.466457849 | 0.743017201 |
| DLEC1 | 80.59295602 | 1.616727396 | 0.953591874 | 1.695408109 | 0.089998029 | 0.343659727 |
| DLEU7 | 29.67312385 | 0.937853179 | 0.620044587 | 1.512557642 | 0.130392061 | 0.415496314 |
| DLG1 | 2356.241407 | -0.244928751 | 0.306805522 | -0.79831924 | 0.424685256 | 0.713499448 |
| DLG2 | 134.150379 | -1.857246315 | 0.536153946 | -3.464016872 | 0.000532173 | 0.018147267 |
| DLG3 | 1777.109236 | -0.057209265 | 0.294100972 | -0.19452253 | 0.845766757 | 0.937897187 |
| DLG4 | 1282.138655 | 0.235604695 | 0.486360731 | 0.484423762 | 0.628085158 | 0.836582962 |
| DLG5 | 431.5874927 | -0.805929911 | 0.479219213 | -1.68175626 | 0.092616114 | 0.3491405 |
| DLGAP1 | 416.3015237 | 0.344003763 | 0.434101419 | 0.792450215 | 0.42809821 | 0.715316008 |
| DLGAP3 | 12.17951902 | -0.45954663 | 0.995927478 | -0.461425797 | 0.644493147 | 0.845794147 |
| DLGAP4 | 1597.828937 | -0.686548421 | 0.363785013 | -1.887236682 | 0.059128499 | 0.281976232 |
| DLGAP5 | 3478.10285 | 0.963584383 | 0.430815905 | 2.236649976 | 0.025309226 | 0.181849597 |
| DLK1 | 2990.318007 | 2.088375662 | 0.867497678 | 2.40735591 | 0.016068502 | 0.143566101 |
| DLL1 | 306.3674814 | -2.131129747 | 2.188476851 | -0.973795883 | 0.330157898 | 0.638717861 |
| DLL3 | 22.42527554 | 1.724031164 | 0.758988305 | 2.27148581 | 0.023117585 | 0.173699516 |
| DLL4 | 10.09614663 | -0.188141357 | 1.011056963 | -0.186083835 | 0.852379018 | 0.940058308 |
| DLST | 3095.253171 | -0.333379806 | 0.290889028 | -1.146072124 | 0.251765311 | 0.564518405 |
| DLX2 | 43.54632508 | -3.460120014 | 1.049076535 | -3.298253177 | 0.000972884 | 0.026690088 |

| | | | | | | |
|---|---|---|---|---|---|---|
| DLX3 | 1500.336455 | -0.462900194 | 0.715529077 | -0.646934148 | 0.517674564 | 0.773950969 |
| DLX4 | 307.1872245 | -0.48472595 | 0.725618465 | -0.668017661 | 0.504122319 | 0.765890246 |
| DLX5 | 2741.378854 | -1.057309271 | 0.757620175 | -1.395566414 | 0.162845102 | 0.464122308 |
| DLX6 | 735.1777166 | -1.251963092 | 0.757930184 | -1.651818491 | 0.098571558 | 0.360518107 |
| DMAC1 | 916.1463207 | 1.271648792 | 0.37261058 | 3.412809135 | 0.00064297 | 0.020666295 |
| DMAC2 | 1834.632545 | -0.111036571 | 0.257941515 | -0.430471889 | 0.666852411 | 0.856258128 |
| DMAC2L | 592.6053321 | 0.300083627 | 0.299324247 | 1.002536983 | 0.316084314 | 0.625961211 |
| DMAP1 | 1125.497176 | -0.288185811 | 0.272947367 | -1.055829237 | 0.291046238 | 0.604113077 |
| DMBX1 | 22.61451081 | -0.464671629 | 1.123216962 | -0.413697126 | 0.679095932 | 0.861617874 |
| DMC1 | 71.17361537 | 0.634154819 | 0.418817678 | 1.514154851 | 0.12998657 | 0.415175781 |
| DMD | 955.6143149 | -0.29750324 | 0.337625264 | -0.881164033 | 0.378229043 | 0.681253679 |
| DMKN | 1995.343279 | -0.246895475 | 0.459754804 | -0.537015542 | 0.591256878 | 0.819070234 |
| DMPK | 723.7854571 | -0.161835376 | 0.36027232 | -0.449202915 | 0.653285285 | 0.848827707 |
| DMRT3 | 21.11905025 | -7.016199314 | 1.834908311 | -3.823732919 | 0.000131446 | 0.007254645 |
| DMRTA1 | 58.99006821 | 0.282314123 | 0.477911795 | 0.590724327 | 0.554705145 | 0.796360966 |
| DMRTC1B | 23.82197915 | -0.566331184 | 0.851227258 | -0.665311382 | 0.505851352 | 0.767164369 |
| DMTF1 | 2227.407231 | -0.460013608 | 0.305906988 | -1.503769533 | 0.132640719 | 0.418704191 |
| DMWD | 631.6220038 | -0.128235365 | 0.299308017 | -0.428439459 | 0.668331201 | 0.856934809 |
| DMXL1 | 703.4652936 | -0.332756855 | 0.376159709 | -0.884615888 | 0.376363835 | 0.680079109 |
| DMXL2 | 858.196044 | 0.022347939 | 0.406141321 | 0.055025031 | 0.956118522 | 0.984426596 |
| DNA2 | 1268.857591 | 0.53435489 | 0.354891506 | 1.505685207 | 0.132148003 | 0.418458996 |
| DNAAF1 | 16.43106641 | -1.079717042 | 0.905051511 | -1.192989602 | 0.232873445 | 0.542441738 |
| DNAAF2 | 954.9380865 | 0.720243651 | 0.293607068 | 2.453086897 | 0.014163612 | 0.135106002 |
| DNAAF3 | 77.49642224 | 0.729765743 | 0.432720852 | 1.686458463 | 0.091707529 | 0.347649906 |
| DNAAF4 | 157.4674966 | 0.72817572 | 0.518654037 | 1.403971949 | 0.160327203 | 0.460580192 |
| DNAAF5 | 1244.757522 | 0.221779409 | 0.290866017 | 0.762479617 | 0.445773804 | 0.728439722 |
| DNAH1 | 117.3928616 | -0.370676721 | 0.71607035 | -0.517654057 | 0.604699659 | 0.825894731 |
| DNAH10 | 19.40682561 | -1.065314328 | 0.840848874 | -1.266951008 | 0.205172808 | 0.511700916 |
| DNAH11 | 54.58653203 | -0.50217239 | 0.705557482 | -0.711738453 | 0.476626751 | 0.749735677 |
| DNAH14 | 520.7341471 | 0.575883059 | 0.323948728 | 1.777698163 | 0.075453442 | 0.314738565 |
| DNAH2 | 19.94790968 | -1.535770392 | 0.788182554 | -1.948495794 | 0.051355668 | 0.260792387 |
| DNAH3 | 15.26740922 | -0.334026313 | 0.951593272 | -0.351017944 | 0.725574887 | 0.88422336 |
| DNAH5 | 13.88861124 | 0.427584339 | 0.970447188 | 0.440605469 | 0.659498643 | 0.851946007 |
| DNAH6 | 80.82757684 | -0.444855302 | 0.485505487 | -0.91627245 | 0.359524014 | 0.66703422 |
| DNAH7 | 10.71247678 | -0.25125947 | 0.911026344 | -0.275798248 | 0.782703029 | |

0.909909835
DNAI1 176.0936505 -1.671009886 0.508849802 -3.283896111 0.001023827 0.027467362
DNAJA1 4555.886323 0.534057083 0.263775458 2.024665551 0.042901718 0.238392703
DNAJA2 2727.881601 -0.108711341 0.242074736 -0.449081728 0.653372701 0.848827707
DNAJA3 765.6110163 0.253012263 0.266798577 0.948326883 0.342963069 0.649840387
DNAJA4 16.23347796 -3.682341622 1.211542538 -3.039382857 0.002370634 0.046768381
DNAJB1 1588.74309 -0.240823803 0.373507475 -0.644763008 0.519080777 0.774631448
DNAJB11 1770.943903 -0.162581966 0.267308267 -0.608218997 0.543042237 0.78943022
DNAJB12 734.423132 -0.220926526 0.246257756 -0.897135299 0.369646725 0.675245088
DNAJB13 17.44098643 0.131160517 0.760076675 0.172562219 0.862995551 0.944730333
DNAJB14 1009.847063 0.104706066 0.3029312 0.345643055 0.729610985 0.886372257
DNAJB2 1220.847425 0.072050508 0.303242294 0.237600458 0.812190995 0.921390083
DNAJB4 540.6142332 0.738383255 0.278828621 2.648161629 0.008093082 0.100970518
DNAJB5 864.1479563 1.164403119 0.477540083 2.438335881 0.014755057 0.138131624
DNAJB6 3226.891844 -0.000904133 0.250460589 -0.003609881 0.997119738 0.998633717
DNAJB9 349.2897407 0.31333751 0.378035828 0.82885665 0.407185528 0.701739367
DNAJC1 1515.995859 -0.365894917 0.509949403 -0.717512198 0.473058112 0.747346103
DNAJC10 3919.683906 -0.2765355 0.264635284 -1.044968367 0.29603759 0.607327421
DNAJC11 1641.015239 -0.561331528 0.436520052 -1.285923809 0.198469653 0.504246161
DNAJC12 26.23218892 1.084759651 0.693113324 1.565053814 0.117570292 0.395641036
DNAJC13 1323.801949 -0.684846549 0.391125556 -1.750963442 0.079952208 0.323627252
DNAJC14 1242.566902 -0.121440972 0.237781381 -0.510725318 0.60954341 0.828395867
DNAJC15 2130.852193 0.101434067 0.434905158 0.233232614 0.815580781 0.922975941
DNAJC16 688.87465 -0.341197115 0.272237971 -1.253304647 0.210094858 0.51816316
DNAJC17 471.8580217 0.444765927 0.318206911 1.397725541 0.162195501 0.462792543
DNAJC18 617.3760886 0.484614986 0.347626803 1.394066805 0.163297432 0.464625909
DNAJC19 1194.900101 0.889670912 0.400620675 2.220731396 0.02636916 0.185636437
DNAJC2 1486.527912 0.365550247 0.296868942 1.231352272 0.21819114 0.526595256
DNAJC21 1920.552127 0.199991477 0.225424036 0.887179027 0.374982522 0.67920525
DNAJC22 24.90113759 -0.333747488 1.071456845 -0.311489435 0.755428573 0.897831634
DNAJC24 766.8048258 0.888571202 0.305763178 2.906076555 0.003659919 0.062311776
DNAJC25 305.1204127 0.474091487 0.357223686 1.327155802 0.18445715 0.488888787
DNAJC25-GNG10 288.3196944 -0.187578408 0.407240652 -0.460608261

0.645079684 0.845794147
DNAJC27 231.3562394 0.187960061 0.448565645 0.419024647 0.67519812 0.860095396
DNAJC28 66.76809204 0.238022091 0.429093583 0.554709043 0.57909369 0.810713649
DNAJC3 963.443971 -0.545267698 0.252205933 -2.161993939 0.030618643 0.199448449
DNAJC30 249.5661225 0.048575134 0.485695541 0.100011489 0.920335204 0.969813529
DNAJC4 336.7237386 -0.033397881 0.370448368 -0.090155293 0.928163811 0.972633911
DNAJC5 1351.683796 -0.402802504 0.41329991 -0.974600995 0.32975822 0.638548751
DNAJC6 475.0802842 -1.255996834 0.450924313 -2.785382816 0.005346456 0.077809098
DNAJC7 2827.400229 -0.029971322 0.231425436 -0.129507467 0.896956117 0.95928311
DNAJC8 2353.805002 0.480262463 0.267978786 1.792165978 0.073106383 0.311055732
DNAJC9 1354.191506 0.185270015 0.271424976 0.682582779 0.494870519 0.760514211
DNAL1 381.1466824 0.509107515 0.270870772 1.879521779 0.060173283 0.283770243
DNAL4 501.9764903 -0.086946959 0.367622636 -0.236511441 0.813035831 0.921802034
DNALI1 703.1086685 0.496945822 0.270221994 1.839028032 0.065911063 0.296365364
DNASE1 417.4568446 -0.433450619 0.277499204 -1.561988692 0.118290653 0.39633116
DNASE1L1 493.2647694 -0.158523094 0.290857201 -0.545020352 0.58573952 0.815639728
DNASE2 981.481152 0.006866943 0.361946699 0.01897225 0.984863242 0.994433294
DND1 22.15432459 1.480929373 0.681811252 2.172051821 0.029851754 0.197226575
DNER 114.648417 0.687880952 0.555487567 1.238337261 0.215591033 0.523214349
DNHD1 329.1467636 -0.40352817 0.529391211 -0.76224947 0.445911126 0.728501578
DNLZ 113.545226 0.983616913 0.518581615 1.896744668 0.057861645 0.278761201
DNM1 1033.174077 -0.182323263 0.465553497 -0.391626879 0.695333925 0.86951129
DNM1L 2848.910156 0.010208857 0.286453875 0.035638745 0.971570414 0.989665121
DNM2 2726.392551 -0.364462338 0.506071264 -0.720179872 0.471414255 0.746147696
DNM3 12.29494447 0.871412521 0.828306472 1.052041183 0.292780655 0.605827098
DNMBP 2371.292148 -0.873833243 0.243354872 -3.590777674 0.000329693 0.013445616
DNMT1 3511.13614 -0.145704104 0.439488966 -0.331530744 0.740243622 0.891917214
DNMT3A 3057.51914 -0.397277816 0.298095695 -1.332719063 0.182624015 0.486683209
DNMT3B 3781.54642 -0.368762907 0.373120523 -0.988321158 0.322995387 0.632932448
DNMT3L 62.05825182 -0.999795913 1.098920042 -0.909798598 0.362928734 0.669987492
DNPEP 802.9599495 0.2053046 0.242887625 0.845265787 0.397962459 0.696517252
DNPH1 438.0978231 1.685460233 0.480714413 3.506157056 0.000454627 0.016498831
DNTTIP1 813.5779788 0.493026548 0.351518714 1.40256131 0.160747696 0.4612641
DNTTIP2 2279.883568 0.160490603 0.281758316 0.569603781 0.568946463 0.805450586
DOC2A 105.5246159 0.26196981 0.441838473 0.592908553 0.553242353 0.79574357
DOC2B 32.35645462 -5.150200541 1.955615449 -2.633544618 0.008449873 0.103848246
DOC2GP 57.5043324 0.402840444 0.605632541 0.665156537 0.505950376 0.767193719
DOCK1 2480.655183 -0.542017611 0.428466128 -1.265018574 0.20586467 0.512218582

| | | | | | | |
|---|---|---|---|---|---|---|
| DOCK11 | 537.0717419 | -0.357172687 | 0.383714143 | -0.930830133 | 0.35194144 | 0.65929447 |
| DOCK2 | 12.87319749 | -1.044685096 | 0.926761181 | -1.127243045 | 0.259639736 | 0.572948272 |
| DOCK3 | 902.7688001 | -1.070884619 | 0.362176949 | -2.956799492 | 0.003108502 | 0.055612125 |
| DOCK4 | 543.4841131 | -0.326453137 | 0.373113893 | -0.874942325 | 0.381605288 | 0.683026643 |
| DOCK5 | 179.2813805 | -1.071082436 | 0.416874949 | -2.569313506 | 0.010190022 | 0.114598094 |
| DOCK6 | 1035.783194 | -0.53421481 | 0.316020657 | -1.690442694 | 0.090943293 | 0.34565427 |
| DOCK7 | 1346.107019 | -0.229853423 | 0.375348724 | -0.612372999 | 0.540291002 | 0.787481586 |
| DOCK8 | 177.9368391 | -1.343131602 | 0.396183492 | -3.390175589 | 0.000698479 | 0.021675701 |
| DOCK9 | 1451.460613 | -3.24054051 | 0.702767837 | -4.611111012 | 4.01E-06 | 0.000552393 |
| DOHH | 137.6676726 | 0.638395498 | 0.499888002 | 1.277077055 | 0.201575045 | 0.5080736 |
| DOK1 | 163.0803574 | 1.453711344 | 0.393259359 | 3.696571511 | 0.000218531 | 0.01007699 |
| DOK3 | 55.92459103 | -1.300037262 | 0.652587773 | -1.992126294 | 0.046357203 | 0.247615641 |
| DOK4 | 1265.327964 | -1.362393291 | 0.563318273 | -2.418514287 | 0.015584034 | 0.141315649 |
| DOK5 | 152.9260578 | 0.869788592 | 0.566068968 | 1.536541729 | 0.124405571 | 0.405999855 |
| DOK6 | 22.22327479 | -2.023844417 | 0.956605768 | -2.115651487 | 0.034374459 | 0.211724136 |
| DOLK | 481.5923953 | 0.035676966 | 0.265003543 | 0.134628259 | 0.892905798 | 0.957604542 |
| DOLPP1 | 846.943897 | 0.399831944 | 0.237593031 | 1.682843741 | 0.092405345 | 0.348900321 |
| DONSON | 1144.077297 | 0.360978319 | 0.261872357 | 1.378451408 | 0.168063961 | 0.470384431 |
| DOP1A | 605.056767 | -0.957347802 | 0.345378967 | -2.771876381 | 0.00557342 | 0.079919047 |
| DOP1B | 446.398087 | -1.045306825 | 0.580400148 | -1.801010609 | 0.071701208 | 0.308152698 |
| DOT1L | 1554.636499 | -0.128183009 | 0.478816428 | -0.267708043 | 0.788924058 | 0.912533261 |
| DPAGT1 | 698.2855047 | -0.361577689 | 0.34611114 | -1.044686654 | 0.296167815 | 0.607412916 |
| DPCD | 414.2033013 | 0.703759453 | 0.422263303 | 1.666636548 | 0.095586697 | 0.354819129 |
| DPF1 | 95.80304082 | 0.691745802 | 0.422551921 | 1.63706699 | 0.101616473 | 0.365596699 |
| DPF2 | 1997.665611 | -0.232982918 | 0.308776011 | -0.754536978 | 0.45052685 | 0.731870954 |
| DPF3 | 33.32975777 | 1.073282528 | 0.97526026 | 1.10050883 | 0.271110485 | 0.583394591 |
| DPH1 | 513.2132044 | -0.131120056 | 0.432137021 | -0.303422409 | 0.76156797 | 0.90066633 |
| DPH2 | 639.377375 | -0.144198566 | 0.322816331 | -0.446689194 | 0.655099485 | 0.849331122 |
| DPH3 | 911.4133118 | 0.621392321 | 0.312132827 | 1.99079452 | 0.046503482 | 0.248067625 |
| DPH5 | 1767.84397 | 0.543365081 | 0.263674927 | 2.060738528 | 0.039327992 | 0.228518481 |
| DPH6 | 723.810287 | 0.562444604 | 0.313889157 | 1.791857384 | 0.073155814 | 0.311055732 |
| DPH7 | 762.3243011 | 0.260559622 | 0.240984001 | 1.081232038 | 0.27959391 | 0.593247442 |
| DPM1 | 1600.886279 | 0.894323173 | 0.348481372 | 2.566344273 | 0.010277676 | 0.11507205 |
| DPM2 | 803.5149023 | 0.289113925 | 0.351242521 | 0.823117669 | 0.410441082 | 0.704150362 |
| DPM3 | 242.7442407 | 1.633412503 | 0.592534357 | 2.756654502 | 0.005839602 | 0.082488453 |
| DPP10 | 267.2690849 | -1.647840673 | 0.538696693 | -3.058939647 | 0.002221219 | 0.044607685 |
| DPP3 | 1658.124885 | -0.212628863 | 0.305626999 | -0.69571361 | 0.486608201 | 0.756340124 |
| DPP7 | 75.31950983 | 0.151408125 | 0.517540756 | 0.292553047 | 0.769863807 | 0.904694332 |
| DPP8 | 796.7087562 | -0.138877165 | 0.304090189 | -0.456697289 | 0.64788864 | 0.846597949 |
| DPP9 | 593.250837 | -0.609447272 | 0.339380595 | -1.795763462 | 0.072532144 | |

0.30996765
DPPA2 40.74559457 0.45459757 0.583473804 0.779122503 0.435907556 0.720784523
DPPA4 3721.893002 0.224332204 0.31424608 0.71387431 0.475304905 0.748902679
DPY19L1 1148.678178 0.4820639 0.310359152 1.553245319 0.120364538
0.400562008
DPY19L1P1 68.23836537 -0.04323056 0.432763344 -0.099894228 0.920428299
0.969843271
DPY19L2 34.6084666 0.689002706 0.555056325 1.24132034 0.214487433
0.521483923
DPY19L2P1 24.39666161 0.173220131 0.676095649 0.256206546 0.797791354
0.915916373
DPY19L2P2 41.01824176 1.130770389 0.542449407 2.084563786 0.037108915
0.220693972
DPY19L2P3 12.1818944 -0.265922118 0.828562019 -0.320944132
0.748252729 0.894401367
DPY19L3 1885.859903 -0.092525813 0.312185194 -0.296381171
0.766939003 0.903470997
DPY19L4 1028.486578 -0.186126659 0.277938189 -0.669669253
0.503068656 0.765194965
DPY30 2124.325422 0.748910028 0.487757628 1.535414281 0.124682098 0.406173955
DPYSL2 3007.458506 0.458177248 0.327126268 1.400612832 0.161329882
0.46167973
DPYSL3 11739.47934 0.331732533 0.444232599 0.746754141 0.455211982
0.73507408
DPYSL4 124.0204622 0.82373767 0.646349065 1.274447067 0.202505021
0.509290789
DPYSL5 1494.529343 0.614882524 0.566938426 1.084566675 0.278113622
0.592005173
DQX1 60.55833022 -1.150066668 0.461334595 -2.492912263 0.012670015
0.127358401
DR1 2156.750785 0.31723365 0.253061798 1.253581744 0.20999407 0.518103763
DRAM1 298.0458321 0.107508704 0.466267249 0.230573142 0.817646435 0.92398615
DRAM2 1957.982805 0.493761161 0.243770824 2.025513769 0.042814636 0.238139903
DRAP1 1395.065157 1.074444193 0.43653042 2.461327192 0.013842406 0.133991065
DRAXIN 1872.128265 0.835618504 0.476576471 1.753377589 0.079537215
0.323002882
DRC3 35.1432222 -0.14827301 0.635999251 -0.233133939 0.815657402
0.922975941
DRC7 15.91521856 -1.099132782 0.812580836 -1.352644234 0.176169314
0.480176907
DRG1 2136.072213 -0.057689086 0.21793416 -0.264708782 0.791233812
0.913361935
DRG2 1078.503159 -0.017893876 0.25449063 -0.070312512 0.943944924
0.978984717
DROSHA 2637.262851 -0.517313287 0.345079648 -1.499112715
0.133844394 0.420559542
DRP2 36.62372756 0.552632326 0.825780435 0.669224291 0.503352413 0.765246655
DSC1 19.62253739 -1.635105716 0.929558731 -1.759012812 0.078575332
0.321658491
DSC2 5356.575351 -1.536299662 0.501951307 -3.060654772 0.002208536
0.044579397
DSC3 134.7114794 -0.047526543 0.529761132 -0.089713156 0.928515161
0.972633911
DSCC1 1007.241868 0.844774287 0.292054442 2.892523331 0.003821608 0.063781754
DSE 1015.068407 0.326762512 0.321606757 1.016031238 0.309614511 0.61915097
DSEL 174.2940365 -0.46850095 0.450173455 -1.040712073 0.298009198
0.609375237
DSG2 6224.227028 -1.29344266 0.326117555 -3.966185324 7.30E-05
0.00492431
DSG3 10.46496566 0.164185445 1.057227272 0.155298155 0.87658627 0.950324661
DSN1 790.2893981 0.324461687 0.254384857 1.275475637 0.202140942 0.508823666
DSP 44281.44041 -1.867964531 0.350042775 -5.336389329 9.48E-08
3.89E-05
DST 3678.879763 -1.203652448 0.449993504 -2.674821829 0.007476897

| | | | | | | |
|---|---|---|---|---|---|---|
| | | | | | | 0.096128823 |
| DSTN | 6597.232489 | -0.006890747 | 0.379545838 | -0.018155243 | 0.985515007 | 0.994669101 |
| DSTNP2 | 82.81143307 | 0.023142915 | 0.456346318 | 0.05071349 | 0.959553827 | 0.985942023 |
| DSTYK | 794.7141619 | 0.033099753 | 0.317427224 | 0.104275091 | 0.916951044 | 0.969344596 |
| DTD1 | 1231.360862 | 0.624730697 | 0.269239522 | 2.320352867 | 0.020321796 | 0.161005801 |
| DTD2 | 225.0165233 | 0.843251065 | 0.346745449 | 2.43190233 | 0.015019757 | 0.139113039 |
| DTL | 1225.44588 | 0.217144498 | 0.316840977 | 0.68534222 | 0.493127996 | 0.759389046 |
| DTNA | 84.00944329 | 0.872466982 | 0.854188105 | 1.021399124 | 0.30706538 | 0.617205881 |
| DTNB | 718.8951637 | -0.306931603 | 0.342460371 | -0.896254367 | 0.370116925 | 0.675859878 |
| DTNBP1 | 645.2333187 | 0.374086662 | 0.279758159 | 1.337178738 | 0.181164301 | 0.485749663 |
| DTWD1 | 838.7817033 | 0.323877055 | 0.242392512 | 1.336167742 | 0.181494452 | 0.485768051 |
| DTWD2 | 202.6671788 | -0.054748191 | 0.30309199 | -0.180632259 | 0.856656233 | 0.942343063 |
| DTX1 | 35.07078997 | 0.860690686 | 0.572285546 | 1.503953213 | 0.132593414 | 0.418641974 |
| DTX2 | 212.1106897 | -0.976625089 | 0.534775982 | -1.826232144 | 0.067815301 | 0.299425477 |
| DTX2P1 | 33.58492059 | -0.698895375 | 0.574394806 | -1.216750861 | 0.223699021 | 0.532852544 |
| DTX3 | 1335.067205 | 0.031694236 | 0.241729971 | 0.131114218 | 0.895684954 | 0.958837464 |
| DTX3L | 126.4409767 | -0.943554515 | 0.40301839 | -2.341219502 | 0.019220863 | 0.157389967 |
| DTX4 | 1688.078304 | 0.264463298 | 0.391339477 | 0.675789982 | 0.499173996 | 0.76281557 |
| DTYMK | 1053.722448 | 1.011907074 | 0.361265505 | 2.801006629 | 0.005094347 | 0.075897737 |
| DUOX1 | 13.32332341 | -0.71662746 | 0.933323283 | -0.767823403 | 0.442592109 | 0.726055891 |
| DUS1L | 1363.698643 | 0.33884108 | 0.346526965 | 0.97782024 | 0.328163239 | 0.637787615 |
| DUS2 | 409.2949087 | -0.096629971 | 0.315667351 | -0.30611329 | 0.759518389 | 0.899504565 |
| DUS3L | 267.3609062 | 0.179517179 | 0.425336272 | 0.422059418 | 0.672981648 | 0.859194192 |
| DUS4L | 205.7440878 | 0.423410995 | 0.338077795 | 1.252406995 | 0.210421599 | 0.51816316 |
| DUSP1 | 302.2086678 | 0.181929948 | 0.4865892 | 0.373888175 | 0.708487515 | 0.875585221 |
| DUSP10 | 37.661997 | -0.028728867 | 0.718058439 | -0.040009093 | 0.968085877 | 0.98927865 |
| DUSP11 | 553.9056341 | 0.222146871 | 0.366679134 | 0.605834503 | 0.544624656 | 0.790739196 |
| DUSP12 | 735.2317982 | 0.813543355 | 0.366923388 | 2.217202235 | 0.026609275 | 0.18654774 |
| DUSP13 | 15.65886358 | 0.344577949 | 0.90837694 | 0.37933366 | 0.704440107 | 0.873435301 |
| DUSP14 | 255.7096575 | 1.521004035 | 0.945903859 | 1.607990094 | 0.107837348 | 0.37651706 |
| DUSP15 | 35.0326162 | 0.882789642 | 0.780851207 | 1.130547835 | 0.258245454 | 0.571113962 |
| DUSP16 | 446.1475951 | -0.303620141 | 0.328637711 | -0.923874926 | 0.355551439 | 0.663069562 |
| DUSP18 | 150.4638506 | -0.126388322 | 0.356313361 | -0.354711149 | 0.722805998 | 0.883047978 |
| DUSP19 | 50.97251434 | 1.27646804 | 0.511440754 | 2.495827776 | 0.01256636 | 0.126734979 |
| DUSP2 | 57.82230065 | -0.65007139 | 0.817516801 | -0.795178019 | 0.426509956 | 0.714361085 |
| DUSP22 | 1063.025164 | 0.19749348 | 0.245907221 | 0.803121921 | 0.421904269 | 0.712063713 |
| DUSP23 | 321.5941215 | 0.941696464 | 0.506472097 | 1.859325459 | 0.062981021 | 0.289778852 |
| DUSP26 | 16.4019516 | 1.530960975 | 0.818604184 | 1.870209077 | 0.06145479 | 0.285640507 |
| DUSP27 | 32.61470236 | -1.550865638 | 0.758036783 | -2.045897605 | 0.040766446 | 0.231983403 |
| DUSP28 | 38.76876581 | -0.259654704 | 0.525722554 | -0.493900636 | | |

0.621376355 0.833354021
DUSP3 899.2084519 0.074139779 0.246833782 0.300363177 0.763900148 0.902439585
DUSP4 32.84076478 0.517453707 0.595214764 0.869356303 0.384652276 0.685277276
DUSP5 369.1506104 -1.556094855 0.774338619 -2.009579292 0.044475735
0.242276974
DUSP6 139.2641879 -0.293985521 0.957100142 -0.30716276 0.758719489
0.899440455
DUSP7 215.1265852 0.838763688 0.434549729 1.930190339 0.053583258 0.267581175
DUSP8P5 26.75085312 -0.680130249 0.67867711 -1.002141135
0.316275433 0.626189654
DUT 1408.311496 0.983937534 0.393663896 2.499435542 0.012439133 0.126653745
DUXAP9 15.96639546 0.706922512 0.798819956 0.884958502 0.376179013
0.680079109
DVL1 963.0675246 0.393138703 0.516709454 0.760850609 0.446746302 0.728709125
DVL2 1802.637093 -0.064045085 0.333861351 -0.191831383 0.847874287
0.938207222
DVL3 2207.527876 -0.973794354 0.372312505 -2.615529536 0.008908922
0.10668882
DXO 263.6174545 0.224554022 0.291164009 0.77122864 0.440571421 0.7241465
DYDC2 55.25173021 0.959234981 0.518814306 1.848898477 0.064472474 0.293286926
DYM 1537.122272 -0.023505345 0.231556999 -0.101509975 0.91914564
0.969677217
DYNAP 40.40756027 0.572100267 0.653595273 0.875312736 0.381403766 0.682984572
DYNC1H1 9732.736675 -0.626952577 0.460683466 -1.360918338
0.173539498 0.477540818
DYNC1I1 511.1057482 -0.726605183 0.695013845 -1.045454258
0.295813071 0.607113109
DYNC1I2 4778.199968 -0.049676228 0.256735282 -0.19349202 0.846573655
0.938145276
DYNC1LI1 1888.855182 0.496543383 0.236264192 2.101644686 0.035584412
0.215965226
DYNC1LI2 2893.752021 -0.544879237 0.28617959 -1.903976579
0.056913235 0.2761211
DYNC2H1 926.2874289 -0.67757442 0.429723105 -1.576769811 0.1148485
0.389985713
DYNC2LI1 851.2243219 0.600962343 0.310962592 1.932587256 0.053287064
0.267056023
DYNLL1 5804.842583 0.676141864 0.409003873 1.653142937 0.098301777
0.359965304
DYNLL2 5214.740279 -0.170260008 0.244729204 -0.695707766
0.486611861 0.756340124
DYNLRB1 1994.661508 0.32427539 0.398014655 0.814732285 0.415225569
0.707398889
DYNLT1 938.7087277 1.142356609 0.403634431 2.83017632 0.004652236
0.072024426
DYNLT3 272.6530611 0.570647079 0.287878684 1.982248463 0.047451439
0.250831282
DYRK1A 1407.13936 0.05946131 0.288897991 0.205821126 0.836930638
0.933183529
DYRK1B 228.0393117 -0.309951775 0.439682723 -0.704944175
0.480844978 0.752563699
DYRK2 1414.344469 0.377772695 0.403441921 0.93637442 0.349080444 0.655726918
DYRK3 172.9936654 1.148902645 0.533776763 2.152402886 0.031365637 0.202023044
DYRK4 233.8300224 -0.026837084 0.292752129 -0.091671696 0.926958886
0.972403074
DYSF 165.5295109 -2.286272917 0.510447811 -4.47895528 7.50E-06
0.000875358
DZANK1 205.1180974 -0.195606303 0.342184313 -0.571640182
0.567565767 0.805151447
DZIP1 1284.671873 0.338870491 0.376019809 0.901203827 0.367479957 0.673938238
DZIP1L 77.29886687 0.443015175 0.484698614 0.914001324 0.36071615
0.667857544
DZIP3 1912.551728 -0.009770597 0.264245223 -0.03697549 0.970504548
0.98957539

| | | | | | | |
|---|---|---|---|---|---|---|
| E2F1 | 583.5930863 | 0.75892413 | 0.279429671 | 2.715975462 | 0.006608082 | 0.089064189 |
| E2F2 | 70.4856576 | -0.031267538 | 0.442431183 | -0.070672094 | 0.943658731 | 0.97883756 |
| E2F3 | 973.277876 | 0.096178465 | 0.306841114 | 0.313447125 | 0.753940986 | 0.897171441 |
| E2F4 | 1637.600748 | 0.002440901 | 0.235858419 | 0.010349009 | 0.991742833 | 0.99688215 |
| E2F5 | 1493.963215 | 0.921463059 | 0.29054621 | 3.171485384 | 0.001516615 | 0.035838892 |
| E2F6 | 697.5933427 | 0.339436213 | 0.266462306 | 1.273862025 | 0.202712319 | 0.509290789 |
| E2F7 | 460.9744788 | 0.021993551 | 0.466268007 | 0.047169333 | 0.962378269 | 0.987343615 |
| E2F8 | 121.1739649 | -0.468244602 | 0.727144549 | -0.643949821 | 0.519607974 | 0.774983751 |
| E4F1 | 376.7056584 | -0.245481371 | 0.516531166 | -0.475249873 | 0.634608883 | 0.840313679 |
| EAF1 | 1232.206225 | -0.206547166 | 0.23660559 | -0.872959787 | 0.382684997 | 0.683829692 |
| EAF2 | 64.12280325 | 0.187854779 | 0.430291445 | 0.436575678 | 0.662419105 | 0.853900946 |
| EAPP | 955.0688574 | 0.337326716 | 0.35026782 | 0.963053688 | 0.335520581 | 0.643919384 |
| EARS2 | 1206.137524 | -0.102518323 | 0.414344975 | -0.247422629 | 0.804581163 | 0.918595231 |
| EBAG9 | 497.9150522 | 0.105984528 | 0.331340672 | 0.319865737 | 0.749070112 | 0.895096304 |
| EBF1 | 10.48888586 | 1.354414689 | 1.003551215 | 1.349621891 | 0.177137299 | 0.481566699 |
| EBF4 | 21.54954438 | 0.530641598 | 0.68149472 | 0.77864374 | 0.436189605 | 0.720784523 |
| EBLN2 | 59.65654526 | -1.074487366 | 0.579285117 | -1.854850633 | 0.063617559 | 0.290852073 |
| EBNA1BP2 | 1318.918287 | 0.395262037 | 0.334702509 | 1.180935386 | 0.237628393 | 0.548298151 |
| EBP | 1428.990733 | 0.571878618 | 0.361580436 | 1.581608295 | 0.113739019 | 0.388022186 |
| EBPL | 1374.936889 | 1.49601258 | 0.487038612 | 3.071650876 | 0.002128786 | 0.043702036 |
| ECD | 1166.382934 | -0.222657253 | 0.245715661 | -0.906158166 | 0.364852142 | 0.671614642 |
| ECE1 | 1750.636927 | -0.23061287 | 0.311620641 | -0.740043629 | 0.459273522 | 0.737472333 |
| ECEL1 | 156.0198052 | -0.789805935 | 0.489515798 | -1.613443201 | 0.106648254 | 0.374599763 |
| ECEL1P1 | 172.1260742 | -0.428295183 | 0.855636583 | -0.500557353 | 0.616682683 | 0.831694047 |
| ECEL1P2 | 89.50955132 | -0.044542355 | 0.801431125 | -0.05557852 | 0.955677577 | 0.984323736 |
| ECH1 | 816.562496 | 0.275868381 | 0.291285966 | 0.947070623 | 0.343602793 | 0.650461439 |
| ECHDC1 | 1402.482783 | 0.320431301 | 0.247662436 | 1.293822778 | 0.195726643 | 0.501413188 |
| ECHDC2 | 555.2829275 | -1.390502872 | 0.454119857 | -3.061973288 | 0.002198831 | 0.044537331 |
| ECHDC3 | 356.4178173 | 0.768055714 | 0.352491945 | 2.17893125 | 0.029336776 | 0.195861365 |
| ECHS1 | 2433.147429 | 0.542810822 | 0.385757492 | 1.407129693 | 0.159388935 | 0.459189048 |
| ECI1 | 935.1192574 | 0.796542575 | 0.444058612 | 1.793778012 | 0.072848608 | 0.310624251 |
| ECI2 | 1400.753602 | 0.734692305 | 0.261908404 | 2.805149794 | 0.00502932 | 0.075658979 |
| ECM1 | 34.31288661 | 0.080783876 | 0.598173714 | 0.135050863 | 0.89257166 | 0.957550733 |
| ECPAS | 3100.214458 | -0.35671306 | 0.34864964 | -1.023127573 | 0.306247531 | 0.616286722 |
| ECSIT | 686.908904 | 0.270657462 | 0.318229081 | 0.850511401 | 0.395040824 | 0.693813886 |
| ECT2 | 2801.662475 | 0.730725543 | 0.37198117 | 1.96441541 | 0.049481937 | 0.256470949 |
| EDA | 394.1917556 | -0.123784023 | 0.380822818 | -0.32504361 | 0.745148066 | 0.893294454 |
| EDA2R | 431.1231457 | -0.036419371 | 0.488031351 | -0.074625064 | 0.940513032 | 0.97807441 |
| EDAR | 50.20930467 | -3.019689498 | 0.698690139 | -4.321929462 | 1.55E-05 | 0.001513876 |
| EDARADD | 135.1707803 | -1.016123845 | 0.483253687 | -2.102671684 | 0.035494479 | 0.21565348 |
| EDC3 | 1464.951213 | -0.24236101 | 0.33063536 | -0.733016002 | 0.463548669 | 0.740758218 |
| EDC4 | 1479.054252 | -0.447865836 | 0.376664438 | -1.189031379 | 0.234427319 | 0.544723061 |

| | | | | | | |
|---|---|---|---|---|---|---|
| EDEM1 | 425.3598478 | -0.184028626 | 0.303835951 | -0.605684171 | 0.544724497 | 0.790739196 |
| EDEM2 | 562.2638011 | -0.72071408 | 0.362457508 | -1.988409854 | 0.04676638 | 0.249119645 |
| EDEM3 | 573.1549486 | 0.107475178 | 0.341580871 | 0.314640505 | 0.753034622 | 0.896586461 |
| EDF1 | 4220.417471 | 0.616686891 | 0.434965428 | 1.417783693 | 0.156253927 | 0.454843067 |
| EDIL3 | 412.4476482 | 0.00936836 | 0.416707906 | 0.022481839 | 0.982063599 | 0.993518428 |
| EDN1 | 312.6865202 | -0.038695849 | 0.536578711 | -0.072115886 | 0.942509684 | 0.978574897 |
| EDN3 | 15.12533864 | 0.088388807 | 0.839736959 | 0.105257731 | 0.916171302 | 0.969184139 |
| EDNRA | 359.820366 | -0.911642918 | 0.349746814 | -2.606579623 | 0.009145156 | 0.108141373 |
| EDNRB | 2916.336255 | -3.198535066 | 0.746778922 | -4.283108389 | 1.84E-05 | 0.001758503 |
| EDRF1 | 1056.501523 | -0.250020973 | 0.419656212 | -0.595775699 | 0.55132508 | 0.794844891 |
| EEA1 | 1013.496176 | 0.10390129 | 0.290856572 | 0.357225178 | 0.720923239 | 0.882227889 |
| EED | 926.1835544 | 0.818841616 | 0.244677944 | 3.346609845 | 0.000818063 | 0.024051992 |
| EEF1A1 | 509211.0526 | 0.316509577 | 0.263566261 | 1.200872889 | 0.229800512 | 0.538758086 |
| EEF1A1P13 | 22.77617057 | 0.041828352 | 0.743610841 | 0.056250326 | 0.955142391 | 0.984323736 |
| EEF1A1P14 | 34.47422826 | -1.178275151 | 0.767107045 | -1.535998344 | 0.124538786 | 0.405999855 |
| EEF1A1P19 | 21.95910061 | 0.038839136 | 0.641122575 | 0.060579891 | 0.951693789 | 0.982350991 |
| EEF1A1P5 | 385.2450857 | -0.362537593 | 0.294213721 | -1.232225309 | 0.217864932 | 0.526142769 |
| EEF1A1P6 | 150.4913968 | -0.340346421 | 0.459865515 | -0.740099899 | 0.45923938 | 0.737472333 |
| EEF1A1P9 | 24.03174764 | 1.108082692 | 0.855227666 | 1.295658146 | 0.195093287 | 0.500636038 |
| EEF1A2 | 93.29291875 | -0.863699329 | 0.620756564 | -1.3913656 | 0.164114591 | 0.466164099 |
| EEF1AKMT1 | 202.9703076 | 0.795251886 | 0.420331841 | 1.891962036 | 0.058496039 | 0.280248566 |
| EEF1AKMT2 | 748.0088641 | 0.794319976 | 0.352471004 | 2.253575376 | 0.024222894 | 0.178105241 |
| EEF1AKMT3 | 94.50034678 | 0.027473841 | 0.44531911 | 0.061694728 | 0.950805939 | 0.981826318 |
| EEF1AKMT4 | 133.3376559 | 0.05406801 | 0.387408761 | 0.139563209 | 0.889005111 | 0.956217735 |
| EEF1AKNMT | 1361.399942 | -0.070311748 | 0.230019137 | -0.305677816 | 0.759849966 | 0.899725004 |
| EEF1B2 | 7395.487085 | 0.70667922 | 0.372213448 | 1.898585942 | 0.057618938 | 0.27812183 |
| EEF1D | 5514.286338 | 0.814588277 | 0.41563091 | 1.959883778 | 0.050009376 | 0.257487003 |
| EEF1E1 | 1402.785397 | 1.300493632 | 0.307077136 | 4.235071518 | 2.28E-05 | 0.002051038 |
| EEF1G | 78185.82136 | -0.019014632 | 0.294030739 | -0.064668858 | 0.948437658 | 0.98134156 |
| EEF2 | 81852.47627 | -0.433719278 | 0.450716811 | -0.962287777 | 0.335905067 | 0.644086928 |
| EEF2K | 107.3108027 | 2.544451674 | 1.07617855 | 2.364339705 | 0.018062242 | 0.151560991 |
| EEF2KMT | 218.9118187 | 0.425127618 | 0.369614229 | 1.150192782 | 0.250064479 | 0.562794702 |
| EEFSEC | 13.39425631 | -0.595275088 | 0.838255202 | -0.710135871 | 0.477619884 | 0.750061298 |
| EEPD1 | 49.75811854 | 0.11198559 | 0.622208824 | 0.179980717 | 0.857167707 | 0.942343063 |
| EFCAB1 | 13.61809753 | 1.612110947 | 0.998823203 | 1.614010311 | 0.10652519 | 0.374529236 |
| EFCAB10 | 31.22044314 | 1.150219207 | 0.583628515 | 1.970807077 | 0.048745947 | 0.254481951 |
| EFCAB11 | 320.6722232 | 0.259039242 | 0.392673008 | 0.659681813 | 0.50945804 | |

| | | | | | | |
|---|---|---|---|---|---|---|
| | | | | | | 0.769564758 |
| EFCAB12 | 16.69009055 | 0.16504472 | 0.97879322 | 0.16862062 | 0.86609506 | 0.945612136 |
| EFCAB13 | 67.82329223 | 0.022882681 | 0.421405601 | 0.054300847 | 0.956695475 | 0.984539826 |
| EFCAB14 | 1606.439401 | -0.678651373 | 0.277924387 | -2.441856152 | 0.014611968 | 0.137627695 |
| EFCAB2 | 190.0297263 | 0.037542207 | 0.328654168 | 0.114230126 | 0.909055371 | 0.966016507 |
| EFCAB5 | 10.5446045 | 0.34553463 | 0.904836618 | 0.381875162 | 0.702553964 | 0.872703476 |
| EFCAB6 | 36.05749156 | -0.441549003 | 0.682349021 | -0.647101395 | 0.517566323 | 0.773950969 |
| EFCAB7 | 253.4312971 | 0.177826881 | 0.372006884 | 0.478020404 | 0.632635683 | 0.838887855 |
| EFCC1 | 141.0263394 | 1.939483373 | 0.700384475 | 2.769169567 | 0.005619938 | 0.080434034 |
| EFEMP1 | 764.1652895 | -0.260382676 | 0.360616116 | -0.722049472 | 0.470264062 | 0.7454951 |
| EFEMP2 | 287.1031472 | -0.030101365 | 0.319666088 | -0.094165026 | 0.924978066 | 0.97180348 |
| EFHB | 38.71801331 | 0.410299212 | 0.668718765 | 0.613560189 | 0.539505998 | 0.78700361 |
| EFHC1 | 611.7659005 | 0.032571632 | 0.316986436 | 0.102754024 | 0.918158196 | 0.969677217 |
| EFHC2 | 33.92478907 | 0.304143366 | 0.603959686 | 0.503582231 | 0.614554982 | 0.830593641 |
| EFHD1 | 147.5857948 | -0.583429901 | 0.498345394 | -1.170734009 | 0.24170571 | 0.552977821 |
| EFHD2 | 263.2017485 | 0.375233169 | 0.414700237 | 0.904829888 | 0.365555515 | 0.672153785 |
| EFL1 | 1012.98934 | -0.171637141 | 0.366391231 | -0.468453191 | 0.639460537 | 0.843436176 |
| EFNA1 | 1153.906078 | -1.670712329 | 0.786584322 | -2.124009192 | 0.033669375 | 0.209086405 |
| EFNA2 | 214.3074343 | 1.343122441 | 0.351069308 | 3.825804222 | 0.000130346 | 0.007243505 |
| EFNA3 | 82.16963029 | 0.95480744 | 0.382489589 | 2.496296547 | 0.012549764 | 0.12669427 |
| EFNA4 | 348.2896786 | 0.185138623 | 0.323544708 | 0.5722196 | 0.567173211 | 0.804843774 |
| EFNA5 | 1636.537436 | 0.376583035 | 0.413866373 | 0.90991455 | 0.362867576 | 0.669987492 |
| EFNB1 | 1290.111422 | 0.761452717 | 0.44219159 | 1.721997284 | 0.085070011 | 0.334264991 |
| EFNB2 | 2824.401688 | -0.601937105 | 0.543910888 | -1.106683315 | 0.268430871 | 0.581517171 |
| EFNB3 | 806.7905171 | 0.132867592 | 0.410291148 | 0.323837336 | 0.746061188 | 0.89396527 |
| EFR3A | 1128.873251 | -0.553794446 | 0.301731758 | -1.835386669 | 0.066448418 | 0.297462837 |
| EFR3B | 543.4817246 | 0.63916532 | 0.517762643 | 1.234475544 | 0.217025754 | 0.524951804 |
| EFS | 2278.463229 | 0.348159736 | 0.327146793 | 1.064230929 | 0.287224099 | 0.601529101 |
| EFTUD2 | 3617.269275 | -0.394508042 | 0.325170749 | -1.213233489 | 0.225040564 | 0.534448858 |
| EGF | 107.1625726 | 0.792738422 | 0.781880002 | 1.013887578 | 0.310636399 | 0.620659135 |
| EGFEM1P | 35.22112627 | -0.338326905 | 0.667414981 | -0.506921354 | 0.612210002 | 0.829098987 |
| EGFL7 | 196.2594047 | 0.773282237 | 0.588373192 | 1.314271704 | 0.188754786 | 0.493962743 |
| EGFL8 | 56.83823318 | 1.862537465 | 0.740275878 | 2.516004534 | 0.011869366 | 0.123589668 |
| EGFLAM | 63.56426266 | -1.213732022 | 0.61975064 | -1.958419959 | 0.050180755 | 0.257598027 |
| EGFR | 450.5251373 | -1.575455298 | 0.390756513 | -4.031808161 | 5.53E-05 | 0.004042687 |
| EGLN1 | 261.155476 | 0.522004688 | 0.378668657 | 1.37852626 | 0.168040866 | 0.470384431 |
| EGLN2 | 1634.342414 | 0.382261679 | 0.317867226 | 1.202582864 | 0.229137782 | 0.537806762 |
| EGLN3 | 420.5051494 | 1.12356629 | 0.618906462 | 1.815405653 | 0.069461593 | 0.30336211 |
| EGR1 | 6540.71468 | -1.776824262 | 0.683233019 | -2.6006124 | 0.009305753 | 0.10932398 |
| EGR2 | 518.6107352 | -1.505139407 | 1.511421085 | -0.99584386 | 0.319326016 | 0.629115732 |
| EGR3 | 150.1083571 | -3.554484874 | 1.137465143 | -3.124917624 | 0.001778549 | 0.039161626 |
| EGR4 | 25.12020065 | -4.478879661 | 3.172507707 | -1.411778969 | 0.158015053 | 0.457315216 |

| | | | | | | |
|---|---|---|---|---|---|---|
| EHBP1 | 1151.076982 | -0.241802478 | 0.347457307 | -0.695919969 | 0.486478951 | 0.756340124 |
| EHBP1L1 | 476.7364416 | -0.785387075 | 0.29962995 | -2.62119016 | 0.008762337 | 0.105800902 |
| EHD1 | 799.8527508 | -0.41713794 | 0.386299104 | -1.079831497 | 0.280217223 | 0.594060918 |
| EHD2 | 518.3753729 | -0.303237126 | 0.439957885 | -0.689241258 | 0.490671458 | 0.758593654 |
| EHD3 | 260.3004723 | 0.415898 | 0.479714293 | 0.866970207 | 0.385958334 | 0.686679504 |
| EHD4 | 790.1437601 | -0.037151496 | 0.247063386 | -0.150372325 | 0.880470875 | 0.952483146 |
| EHHADH | 243.5789093 | -0.932695254 | 0.368500593 | -2.531054962 | 0.011372003 | 0.120838052 |
| EHMT1 | 2332.845102 | -0.297617301 | 0.388999481 | -0.765084057 | 0.444221493 | 0.727151225 |
| EHMT2 | 166.3276983 | 0.309088676 | 0.440814489 | 0.701176308 | 0.483192993 | 0.75401367 |
| EI24 | 5675.933054 | 0.271712451 | 0.341120437 | 0.79652938 | 0.425724406 | 0.713822387 |
| EID1 | 3930.982412 | 0.986678137 | 0.303456174 | 3.251468318 | 0.001148106 | 0.029521884 |
| EID2 | 452.9683648 | 0.699763211 | 0.299156293 | 2.339122488 | 0.019329095 | 0.157622736 |
| EID2B | 71.09853603 | 1.77185858 | 0.450739599 | 3.931002702 | 8.46E-05 | 0.005437924 |
| EID3 | 43.29210154 | 0.65959371 | 0.537118375 | 1.228022985 | 0.21943834 | 0.527925634 |
| EIF1 | 9764.64394 | 0.511491801 | 0.398929174 | 1.282161934 | 0.199785846 | 0.505887637 |
| EIF1AD | 545.1832701 | 0.004319028 | 0.301245539 | 0.014337234 | 0.988560934 | 0.995803391 |
| EIF1AX | 4017.321257 | 0.323203096 | 0.26254419 | 1.231042653 | 0.218306912 | 0.526623336 |
| EIF1AXP1 | 50.00802397 | -0.048278126 | 1.301389058 | -0.037097381 | 0.97040736 | 0.98957539 |
| EIF1AY | 898.0578984 | 1.170818094 | 0.315313099 | 3.713192058 | 0.000204661 | 0.009553597 |
| EIF1B | 2056.99015 | 0.535409336 | 0.419331264 | 1.276817118 | 0.201666821 | 0.508220488 |
| EIF2A | 3476.018954 | 0.30588313 | 0.265061587 | 1.154007767 | 0.248496984 | 0.561181618 |
| EIF2AK1 | 3620.750027 | -0.449130559 | 0.280332168 | -1.602137074 | 0.109125301 | 0.379241044 |
| EIF2AK2 | 1014.123531 | -0.401585596 | 0.288131939 | -1.393755922 | 0.163391323 | 0.464631632 |
| EIF2AK3 | 533.8239612 | -0.411613482 | 0.33972456 | -1.211609435 | 0.225661921 | 0.53459368 |
| EIF2AK4 | 1726.259815 | 0.026020274 | 0.361869993 | 0.071905033 | 0.942677485 | 0.978574897 |
| EIF2B1 | 1400.761664 | 0.050990231 | 0.251738647 | 0.202552257 | 0.839485009 | 0.934215701 |
| EIF2B2 | 559.9767901 | 0.180604253 | 0.288006851 | 0.627083183 | 0.53060471 | 0.781003595 |
| EIF2B3 | 655.5684015 | 0.636628892 | 0.271906433 | 2.341352809 | 0.019214001 | 0.157389967 |
| EIF2B4 | 1440.459868 | 0.159239784 | 0.291327761 | 0.546600101 | 0.5846535 | 0.814990368 |
| EIF2B5 | 1431.048461 | -0.143741022 | 0.271471969 | -0.529487528 | 0.596467294 | 0.821812117 |
| EIF2D | 1473.546556 | -0.093708882 | 0.243808896 | -0.384353826 | 0.700716219 | 0.872001128 |
| EIF2S1 | 2106.567264 | 0.47793246 | 0.276730458 | 1.727068513 | 0.08415536 | 0.332133444 |
| EIF2S2 | 5413.80732 | 0.884080868 | 0.391580758 | 2.257722962 | 0.023962937 | 0.177207433 |
| EIF2S3 | 8457.826299 | -0.0846321 | 0.330387319 | -0.256160256 | 0.797827096 | 0.915916373 |
| EIF3A | 16397.9789 | -0.185475887 | 0.249898093 | -0.742206093 | 0.457962478 | 0.736518606 |
| EIF3B | 6228.519526 | 0.035579823 | 0.24555071 | 0.144898066 | 0.884791353 | 0.95440284 |
| EIF3C | 12296.2235 | -0.561899178 | 0.365047703 | -1.539248633 | 0.123743609 | 0.405045568 |
| EIF3CL | 1380.315855 | 0.252447594 | 0.767286204 | 0.329013597 | 0.742145411 | |

| | | | | | | |
|---|---|---|---|---|---|---|
| | | | | | | 0.892150449 |
| EIF3D | 7364.616264 | -0.272829783 | 0.223490296 | -1.220767918 | 0.222173904 | 0.531066116 |
| EIF3E | 18428.78579 | 0.462171194 | 0.249192748 | 1.854673533 | 0.06364286 | 0.290852073 |
| EIF3EP1 | 20.15850087 | 0.589667103 | 0.679363438 | 0.867970028 | 0.385410741 | 0.686194853 |
| EIF3F | 4474.482455 | 0.333090174 | 0.489490968 | 0.680482778 | 0.496198823 | 0.761056737 |
| EIF3FP3 | 228.7594471 | -0.822907231 | 1.60025426 | -0.514235301 | 0.60708749 | 0.82683583 |
| EIF3G | 3887.479167 | -0.033924667 | 0.297367944 | -0.114083135 | 0.909171891 | 0.966018007 |
| EIF3H | 9259.194387 | 0.058734945 | 0.254996525 | 0.230336258 | 0.817830488 | 0.924112493 |
| EIF3I | 5129.292881 | 0.083949835 | 0.268547802 | 0.312606672 | 0.754579508 | 0.897438517 |
| EIF3J | 2216.118327 | 0.623164144 | 0.278963129 | 2.233858454 | 0.025492388 | 0.182111117 |
| EIF3K | 2574.107076 | 0.376855467 | 0.409760095 | 0.919697822 | 0.357730691 | 0.665264843 |
| EIF3L | 14947.59563 | -0.287732753 | 0.217672866 | -1.321858615 | 0.186215236 | 0.490817669 |
| EIF3M | 7089.740342 | 0.515095881 | 0.321654214 | 1.601396338 | 0.109289163 | 0.379497802 |
| EIF4A1 | 28313.14335 | 0.199593298 | 0.284999564 | 0.700328433 | 0.483722219 | 0.754278091 |
| EIF4A2 | 22346.17569 | 0.658760611 | 0.258193679 | 2.551420361 | 0.010728484 | 0.117687513 |
| EIF4A3 | 3254.477335 | 0.342442241 | 0.338357772 | 1.012071452 | 0.311503891 | 0.621492048 |
| EIF4B | 29758.7013 | -0.110886808 | 0.225920614 | -0.490822003 | 0.623552346 | 0.834324864 |
| EIF4E | 3221.909662 | 0.669632553 | 0.238524886 | 2.807390731 | 0.004994462 | 0.07561376 |
| EIF4E1B | 15.62119742 | 0.695421816 | 1.023747487 | 0.679290376 | 0.496953893 | 0.761325016 |
| EIF4E2 | 2584.30066 | 0.158774158 | 0.285200729 | 0.55671021 | 0.57772544 | 0.809936484 |
| EIF4E3 | 204.0424692 | 0.459039545 | 0.332085384 | 1.382293735 | 0.166881529 | 0.469104998 |
| EIF4EBP1 | 705.6012063 | 1.217558759 | 0.395308401 | 3.080022473 | 0.00206985 | 0.043016015 |
| EIF4EBP2 | 8649.687772 | 0.009552987 | 0.260405617 | 0.036685027 | 0.970736147 | 0.98957539 |
| EIF4EBP3 | 53.40438117 | 2.003198368 | 0.887878808 | 2.256161934 | 0.024060492 | 0.177207433 |
| EIF4ENIF1 | 1004.744296 | -0.158819727 | 0.325040804 | -0.488614736 | 0.625114479 | 0.835033444 |
| EIF4G1 | 8094.80872 | -0.612385822 | 0.353729577 | -1.731225946 | 0.083411473 | 0.330659907 |
| EIF4G2 | 26472.76187 | -0.27440545 | 0.305792241 | -0.897359102 | 0.369527329 | 0.67521115 |
| EIF4G3 | 2720.234025 | -0.237041763 | 0.415162977 | -0.570960747 | 0.568026252 | 0.805301024 |
| EIF4H | 9823.442348 | 0.054645897 | 0.284538513 | 0.192050969 | 0.847702281 | 0.938207222 |
| EIF5 | 5636.416486 | 0.214678793 | 0.256278907 | 0.837676403 | 0.40221247 | 0.699365659 |
| EIF5A | 9611.687776 | 0.437544686 | 0.24677738 | 1.773034004 | 0.076223073 | 0.316642999 |
| EIF5A2 | 145.3979158 | 0.468198866 | 0.373641135 | 1.253070987 | 0.210179874 | 0.51816316 |
| EIF5B | 5291.165124 | 0.567756396 | 0.451612896 | 1.257174898 | 0.208690311 | 0.515809824 |
| EIF6 | 2396.215372 | 0.45206261 | 0.359483404 | 1.257534019 | 0.208560329 | 0.515581062 |
| EIPR1 | 877.8962178 | 0.258721132 | 0.269090462 | 0.961465265 | 0.336318282 | 0.64425371 |
| ELAC1 | 207.8922105 | 0.219879793 | 0.378124528 | 0.581501005 | 0.560902841 | 0.800965456 |
| ELAC2 | 2036.80646 | -0.459339861 | 0.292665261 | -1.569505924 | 0.1165301 | 0.393649109 |
| ELAVL1 | 3020.60561 | 0.187671576 | 0.218585635 | 0.858572322 | 0.390576514 | 0.690466555 |
| ELAVL2 | 106.2086913 | 1.34685577 | 0.550595614 | 2.446179621 | 0.014437905 | 0.136559111 |
| ELAVL3 | 28.30332033 | 0.62144447 | 0.646969057 | 0.960547438 | 0.336779768 | 0.644459616 |

| | | | | | | |
|---|---|---|---|---|---|---|
| ELAVL4 | 12.28335477 | 2.429166081 | 1.204798137 | 2.016243225 | 0.043774548 | 0.240941849 |
| ELF1 | 1342.737101 | -1.13692633 | 0.316449851 | -3.592753564 | 0.000327202 | 0.013380004 |
| ELF2 | 1283.835464 | -0.835534312 | 0.308819086 | -2.705578602 | 0.006818551 | 0.090502394 |
| ELF3 | 1625.740386 | -1.563212983 | 0.658796207 | -2.372832395 | 0.017652272 | 0.150007193 |
| ELF4 | 106.7488392 | -0.815239981 | 0.525765972 | -1.550575777 | 0.121003381 | 0.401219509 |
| ELF5 | 81.38446527 | -1.150486149 | 0.933606782 | -1.232302691 | 0.217836035 | 0.526142769 |
| ELFN1 | 51.94704535 | 0.753403062 | 0.541448396 | 1.391458663 | 0.164086386 | 0.466164099 |
| ELFN2 | 19.64145667 | 0.243788297 | 0.863955314 | 0.282176975 | 0.777807812 | 0.908018171 |
| ELK1 | 906.1719177 | 0.111166614 | 0.270199426 | 0.411424315 | 0.680761427 | 0.862305386 |
| ELK3 | 338.5855286 | -0.191825255 | 0.342874125 | -0.559462614 | 0.575846039 | 0.808969969 |
| ELK4 | 1018.895733 | 0.163932853 | 0.430406062 | 0.380879516 | 0.703292653 | 0.872844636 |
| ELL | 302.5559259 | -0.268349582 | 0.44700559 | -0.600327128 | 0.548288243 | 0.793331515 |
| ELL2 | 959.1367018 | -0.442798578 | 0.370462541 | -1.195258708 | 0.231985967 | 0.541288697 |
| ELL3 | 117.0914234 | -0.296543479 | 0.469431092 | -0.631708218 | 0.527577557 | 0.778966897 |
| ELMO1 | 344.7130518 | -1.19071774 | 0.515311363 | -2.310676274 | 0.020850743 | 0.163645431 |
| ELMO2 | 1094.754412 | -0.450387002 | 0.41449551 | -1.086590785 | 0.2772177 | 0.591007989 |
| ELMO3 | 160.8240401 | -0.462855472 | 0.526645912 | -0.878874139 | 0.379469522 | 0.681253679 |
| ELMOD1 | 163.41165 | -1.796646162 | 0.827325174 | -2.17163241 | 0.0298834 | 0.197226575 |
| ELMOD2 | 1908.057771 | 0.294542604 | 0.241390515 | 1.22019129 | 0.222392368 | 0.531241476 |
| ELMOD3 | 387.4191211 | -0.338280117 | 0.376325467 | -0.898903067 | 0.368704294 | 0.67460916 |
| ELMSAN1 | 721.7122232 | -1.089427156 | 0.322450444 | -3.378587861 | 0.000728591 | 0.022330224 |
| ELN | 99.93586061 | -0.961048346 | 0.578646185 | -1.660856618 | 0.096742247 | 0.357177519 |
| ELOA | 1250.266124 | -0.197280969 | 0.252196559 | -0.782250837 | 0.434067173 | 0.719296316 |
| ELOB | 2715.593458 | 1.094571642 | 0.486181509 | 2.251364198 | 0.02436248 | 0.178492903 |
| ELOC | 1708.143117 | 0.852948585 | 0.416567429 | 2.047564275 | 0.040602713 | 0.231842053 |
| ELOCP19 | 10.52232961 | -1.049632363 | 1.058308526 | -0.991801858 | 0.321294191 | 0.63106605 |
| ELOF1 | 1088.057052 | 0.129921845 | 0.329750127 | 0.394000894 | 0.693580379 | 0.868790368 |
| ELOVL1 | 944.2545912 | -0.768252951 | 0.321155354 | -2.392153648 | 0.016749829 | 0.146293409 |
| ELOVL4 | 328.6672692 | 0.464717582 | 0.351158375 | 1.323384589 | 0.185707514 | 0.490232938 |
| ELOVL5 | 4865.735314 | -0.060966025 | 0.241590996 | -0.252352225 | 0.800768824 | 0.916680548 |
| ELOVL6 | 4195.760136 | -0.695254497 | 0.296312229 | -2.346357758 | 0.018957901 | 0.156140235 |
| ELOVL7 | 109.4170126 | -0.733029728 | 0.590305271 | -1.24178076 | 0.214317463 | 0.521309963 |
| ELP1 | 1996.594819 | 0.003281424 | 0.345991542 | 0.009484115 | 0.992432884 | 0.997035603 |
| ELP2 | 2838.578661 | 0.067984949 | 0.246762539 | 0.275507578 | 0.782926304 | 0.909909835 |
| ELP3 | 976.715079 | -0.256541064 | 0.241493969 | -1.062308365 | 0.288095718 | 0.601673803 |
| ELP4 | 575.1397661 | 0.671274061 | 0.276434905 | 2.428325979 | 0.015168703 | 0.139723368 |
| ELP5 | 995.6369411 | 0.550893 | 0.245371376 | 2.245139631 | 0.02475917 | 0.17998149 |
| ELP6 | 1010.103302 | 0.340877041 | 0.285696594 | 1.193143527 | 0.232813167 | 0.542441738 |

| | | | | | | |
|---|---|---|---|---|---|---|
| EMB | 963.2734035 | -0.128591448 | 0.248098735 | -0.518307554 | 0.604243703 | 0.82577977 |
| EMBP1 | 183.2951731 | 0.424339287 | 0.443100563 | 0.957659102 | 0.338234686 | 0.64545389 |
| EMC1 | 2071.317042 | -0.427891913 | 0.255047563 | -1.677694572 | 0.09340674 | 0.350724392 |
| EMC2 | 1264.833934 | 0.608363934 | 0.253142013 | 2.403251548 | 0.016250005 | 0.144168905 |
| EMC3 | 883.9844279 | 0.004895695 | 0.321858295 | 0.015210714 | 0.987864074 | 0.995541774 |
| EMC3-AS1 | 184.8150481 | 0.158563619 | 0.423460489 | 0.374447257 | 0.70807159 | 0.875585221 |
| EMC4 | 1528.675261 | 0.364085519 | 0.347945587 | 1.046386368 | 0.295382684 | 0.60699582 |
| EMC6 | 589.3849331 | 1.086228119 | 0.44257129 | 2.454357397 | 0.014113663 | 0.134922886 |
| EMC7 | 743.583123 | 0.479957646 | 0.383407095 | 1.251822546 | 0.210634533 | 0.518294063 |
| EMC8 | 788.7424461 | 0.69910164 | 0.346181118 | 2.019467858 | 0.043438615 | 0.239726164 |
| EMC9 | 103.7396427 | 1.220922995 | 0.443730792 | 2.751494861 | 0.005932395 | 0.083256582 |
| EMD | 1230.009818 | 0.46822372 | 0.271485157 | 1.724675207 | 0.084586021 | 0.332854332 |
| EME1 | 486.4044729 | 0.342492254 | 0.373150893 | 0.917838494 | 0.358703426 | 0.666432275 |
| EME2 | 351.9448802 | -0.542792295 | 0.516649518 | -1.050600603 | 0.293442064 | 0.606314063 |
| EMG1 | 1213.209971 | 0.613189325 | 0.316407333 | 1.937974446 | 0.05262634 | 0.264895225 |
| EMID1 | 509.7715676 | 0.478697683 | 0.491078441 | 0.974788635 | 0.329665116 | 0.638548751 |
| EMILIN1 | 45.81010185 | 1.047551172 | 0.905380427 | 1.157028738 | 0.247260619 | 0.559988249 |
| EMILIN2 | 653.7346891 | -1.630696454 | 0.676365866 | -2.410967992 | 0.015910245 | 0.142656219 |
| EMILIN3 | 49.79868951 | 1.099468324 | 0.606532621 | 1.812710951 | 0.069876415 | 0.303603614 |
| EML1 | 1291.809858 | 0.018914274 | 0.394476376 | 0.047947798 | 0.961757846 | 0.9870991 |
| EML2 | 626.7633694 | 0.024162745 | 0.262558833 | 0.092027926 | 0.926675852 | 0.972261514 |
| EML3 | 611.9655035 | -0.200807659 | 0.372472058 | -0.539121406 | 0.589803086 | 0.81828099 |
| EML4 | 3914.530168 | -0.433964399 | 0.348003686 | -1.247010926 | 0.212393491 | 0.519629359 |
| EML6 | 27.36226299 | -0.520835913 | 0.603516817 | -0.863001491 | 0.388136647 | 0.688370656 |
| EMP1 | 25.68705718 | -0.602109456 | 0.757335488 | -0.795036633 | 0.426592193 | 0.714361085 |
| EMP2 | 2631.713538 | -1.251673754 | 0.483337908 | -2.589645323 | 0.009607486 | 0.111098245 |
| EMP3 | 233.5722897 | 1.485388056 | 0.475661708 | 3.122782496 | 0.001791501 | 0.039332648 |
| EMSY | 1016.407668 | -0.220868391 | 0.387126106 | -0.570533446 | 0.568315944 | 0.805450586 |
| EMX2 | 15.67705713 | 0.750278438 | 1.136728845 | 0.660032901 | 0.509232716 | 0.769555686 |
| ENAH | 9396.870232 | -0.101821746 | 0.214001076 | -0.475800159 | 0.634216757 | 0.840101252 |
| ENC1 | 663.338675 | 0.274228255 | 0.361142814 | 0.759334658 | 0.44765239 | 0.729075085 |
| ENDOD1 | 133.691882 | -0.549244012 | 0.404598122 | -1.357505093 | 0.174620776 | 0.47862011 |
| ENDOG | 141.3571709 | 0.553575803 | 0.423625329 | 1.306758036 | 0.191294901 | 0.496770788 |
| ENDOV | 345.8977892 | 0.111389015 | 0.35071431 | 0.317606131 | 0.750783723 | 0.896207755 |
| ENG | 30.52519283 | -2.453470863 | 0.779731177 | -3.146559913 | 0.001652034 | 0.037682927 |
| ENGASE | 540.8391618 | 0.18815976 | 0.506798381 | 0.371271431 | 0.710435374 | 0.876831684 |
| ENHO | 265.8698805 | 1.24047792 | 0.417151299 | 2.97368826 | 0.002942438 | 0.05358664 |
| ENKD1 | 127.6917873 | 0.643120648 | 0.461713885 | 1.392898654 | 0.16365044 | 0.465194082 |
| ENKUR | 11.95180548 | -0.045266532 | 0.880525715 | -0.05140853 | 0.958999988 | 0.985770636 |
| ENO1 | 49363.00483 | -0.201681061 | 0.262343442 | -0.768767306 | 0.442031461 | 0.725453225 |
| ENO2 | 5298.159509 | 0.120277058 | 0.424195453 | 0.283541602 | 0.776761696 | 0.907767854 |
| ENO3 | 566.0926516 | 1.327560678 | 0.376098237 | 3.529824248 | 0.000415836 | 0.015721465 |
| ENO4 | 36.07449444 | 0.244453424 | 0.627036949 | 0.389854896 | 0.696643848 | 0.870023227 |
| ENOPH1 | 1556.300532 | 0.749848502 | 0.261233981 | 2.870409501 | 0.004099405 | 0.066091474 |

| | | | | | | |
|---|---|---|---|---|---|---|
| ENOSF1 | 797.1739194 | -0.014115499 | 0.322785151 | -0.043730323 | 0.965119368 | 0.988700292 |
| ENOX1 | 18.01884167 | 1.279928537 | 0.890594286 | 1.437162305 | 0.150671881 | 0.445603028 |
| ENOX2 | 356.6831413 | -0.402543724 | 0.407647241 | -0.987480555 | 0.323407111 | 0.633248487 |
| ENPEP | 143.4012254 | -1.733357743 | 0.645572091 | -2.684994857 | 0.007253092 | 0.094209471 |
| ENPP1 | 583.5228077 | -0.60599548 | 0.481101976 | -1.259598817 | 0.207814122 | 0.514566354 |
| ENPP2 | 217.1363745 | 0.637512105 | 0.386606481 | 1.648994874 | 0.099148685 | 0.361452832 |
| ENPP4 | 268.4601449 | 0.439989371 | 0.414180121 | 1.062314071 | 0.288093129 | 0.601673803 |
| ENPP5 | 329.276818 | -0.369160053 | 0.355269891 | -1.039097493 | 0.298759397 | 0.610024066 |
| ENPP7P8 | 16.97031837 | -1.985606368 | 0.9385983 | -2.115501774 | 0.034387204 | 0.211724136 |
| ENSA | 4330.796464 | 0.39707401 | 0.297395513 | 1.33517149 | 0.181820226 | 0.48591777 |
| ENTPD1 | 90.29064861 | -0.321904394 | 0.48632026 | -0.661918536 | 0.508023432 | 0.768706122 |
| ENTPD2 | 10.61395757 | -0.286076713 | 1.112104122 | -0.257239145 | 0.796994167 | 0.915732089 |
| ENTPD4 | 2782.871906 | -0.946985124 | 0.409014679 | -2.315283954 | 0.020597397 | 0.162413261 |
| ENTPD5 | 370.0964487 | -0.291091701 | 0.39033005 | -0.745757854 | 0.455813704 | 0.735306344 |
| ENTPD6 | 1325.828697 | -0.05238478 | 0.29171639 | -0.179574345 | 0.857486747 | 0.942343063 |
| ENTPD7 | 464.9289508 | -0.352588723 | 0.328779663 | -1.072416461 | 0.283533017 | 0.597427696 |
| ENTPD8 | 14.40008843 | -1.847295145 | 1.119057132 | -1.650760352 | 0.098787519 | 0.360959887 |
| ENTR1 | 1071.337481 | 0.232937275 | 0.273300314 | 0.852312504 | 0.394040668 | 0.692865578 |
| ENY2 | 2441.909384 | 1.027196106 | 0.341513412 | 3.00777677 | 0.002631664 | 0.050156997 |
| EOGT | 125.814701 | -0.69520571 | 0.495127247 | -1.404095036 | 0.160290552 | 0.460580192 |
| EP300 | 3962.405632 | -1.068259925 | 0.421292532 | -2.535672586 | 0.011223161 | 0.120297968 |
| EP400 | 2334.600167 | -0.215540733 | 0.40072903 | -0.53787152 | 0.590665751 | 0.81857953 |
| EP400P1 | 413.8749746 | -0.272910602 | 0.307173343 | -0.888457963 | 0.374294458 | 0.678668725 |
| EPAS1 | 5265.914647 | -4.742849469 | 1.399028037 | -3.390103231 | 0.000698663 | 0.021675701 |
| EPB41 | 3427.960598 | -0.302365491 | 0.39704086 | -0.761547542 | 0.446330094 | 0.728589711 |
| EPB41L1 | 1098.43238 | -0.082527737 | 0.452705599 | -0.182298909 | 0.855348155 | 0.94189496 |
| EPB41L2 | 2300.876937 | -0.295211476 | 0.332222847 | -0.888594744 | 0.374220917 | 0.67864822 |
| EPB41L3 | 5546.905811 | -0.587626288 | 0.318326863 | -1.845983979 | 0.064894532 | 0.293855407 |
| EPB41L4A | 160.5524966 | -1.387265692 | 0.58596014 | -2.367508636 | 0.017908302 | 0.150937139 |
| EPB41L4B | 306.2781651 | -0.59265376 | 0.515410611 | -1.149867208 | 0.250198569 | 0.562905438 |
| EPB41L5 | 1625.030243 | 0.093195045 | 0.339831545 | 0.274238945 | 0.783901001 | 0.910666418 |
| EPC1 | 950.8041428 | -0.190738849 | 0.289268538 | -0.659383321 | 0.50964965 | 0.769725718 |
| EPC2 | 670.5204652 | -0.489980188 | 0.316899696 | -1.546168061 | 0.122063985 | 0.402659864 |
| EPCAM | 5435.75316 | -0.134765307 | 0.751470366 | -0.179335491 | 0.857674281 | 0.942343063 |
| EPDR1 | 205.5327273 | 0.950336397 | 0.467318537 | 2.03359448 | 0.041992499 | 0.235820549 |
| EPG5 | 697.5080992 | -0.6382096 | 0.337053828 | -1.89349459 | 0.058292127 | 0.27980731 |

| | | | | | | |
|---|---|---|---|---|---|---|
| EPHA1 | 670.0116611 | -1.598893573 | 0.55234579 | -2.894732979 | 0.003794812 | 0.063596264 |
| EPHA2 | 1102.425199 | 1.420195561 | 0.493817812 | 2.875950454 | 0.004028129 | 0.065655774 |
| EPHA4 | 1192.707007 | 0.850188516 | 0.587714663 | 1.446600824 | 0.148008755 | 0.441755032 |
| EPHA6 | 28.65589895 | -2.222495889 | 0.8680015 | -2.56047471 | 0.010452927 | 0.116176818 |
| EPHA7 | 582.6279794 | 2.030749418 | 0.908367982 | 2.235602155 | 0.025377843 | 0.18194777 |
| EPHB1 | 305.7578383 | 1.251003462 | 0.496544829 | 2.519416957 | 0.011754936 | 0.122989063 |
| EPHB2 | 459.9266057 | 0.304301789 | 0.476875051 | 0.638116396 | 0.523397915 | 0.777662198 |
| EPHB3 | 1307.73296 | -0.787789697 | 0.381027375 | -2.067540942 | 0.03868321 | 0.226576733 |
| EPHB4 | 2290.644056 | 0.161385087 | 0.291581868 | 0.553481217 | 0.579933939 | 0.811340629 |
| EPHB6 | 320.5174428 | -1.572810979 | 0.690957182 | -2.276278502 | 0.022829346 | 0.172396221 |
| EPHX1 | 409.7673073 | -0.418792749 | 0.390178955 | -1.073335052 | 0.283120814 | 0.596961028 |
| EPHX2 | 676.1456519 | -0.034455566 | 0.252198688 | -0.136620718 | 0.891330599 | 0.957017706 |
| EPHX3 | 380.9842189 | -0.229753387 | 0.472617396 | -0.486129772 | 0.626875159 | 0.836041947 |
| EPHX4 | 20.10980769 | 0.527693561 | 0.993532605 | 0.531128579 | 0.595329681 | 0.821441254 |
| EPM2A | 170.2322876 | 0.181522386 | 0.345255138 | 0.525763026 | 0.599052867 | 0.823802482 |
| EPM2AIP1 | 945.4888039 | -0.025270552 | 0.350695652 | -0.072058357 | 0.942555466 | 0.978574897 |
| EPN1 | 1742.854575 | -0.279114219 | 0.360792276 | -0.773614729 | 0.439158659 | 0.722689665 |
| EPN2 | 1012.959197 | -0.067532093 | 0.387313987 | -0.174360067 | 0.861582495 | 0.94403207 |
| EPN3 | 407.8977216 | -1.293349483 | 0.760588753 | -1.700458332 | 0.089044748 | 0.342496199 |
| EPOP | 999.5074885 | 0.21173484 | 0.325842529 | 0.649807257 | 0.515816731 | 0.773832307 |
| EPOR | 248.8876053 | 0.201583755 | 0.377969777 | 0.533333 | 0.593803088 | 0.820230051 |
| EPPK1 | 364.254775 | -0.001845418 | 0.29799853 | -0.006192709 | 0.995058965 | 0.998094459 |
| EPRS | 5068.781283 | -0.139276401 | 0.289225641 | -0.481549287 | 0.630126159 | 0.838075099 |
| EPS15 | 1165.858877 | -0.207697057 | 0.283302749 | -0.733127571 | 0.463480626 | 0.740758218 |
| EPS15L1 | 1086.232436 | -0.565714189 | 0.350932806 | -1.612029939 | 0.106955425 | 0.37492307 |
| EPS8 | 187.2817867 | -0.850865337 | 0.463979401 | -1.833842915 | 0.066677315 | 0.297817842 |
| EPS8L1 | 169.2173329 | -0.459899043 | 0.553527278 | -0.830851633 | 0.40605745 | 0.701474651 |
| EPS8L2 | 871.3435531 | -2.76170973 | 0.423516943 | -6.520895511 | 6.99E-11 | 1.77E-07 |
| EPSTI1 | 107.3794015 | -0.512407334 | 0.837641601 | -0.611726224 | 0.540718909 | 0.787864037 |
| ERAL1 | 1640.385205 | 0.114735278 | 0.283553914 | 0.404633024 | 0.68574728 | 0.86493781 |
| ERAP1 | 461.5667041 | -0.057161266 | 0.347148804 | -0.164659262 | 0.869212183 | 0.947227711 |
| ERBB2 | 3219.861268 | -0.532654228 | 0.22594429 | -2.357458245 | 0.018400525 | 0.153593808 |
| ERBB3 | 2065.889802 | -1.254141202 | 0.517847234 | -2.421836247 | 0.015442307 | 0.140677297 |
| ERBB4 | 68.49132504 | -3.328944625 | 0.877237952 | -3.794802332 | 0.000147761 | 0.007793292 |
| ERBIN | 1590.006354 | -0.528530324 | 0.288984124 | -1.828925124 | 0.067410826 | 0.298244514 |
| ERC1 | 691.3813407 | -0.891179313 | 0.379103392 | -2.350755311 | 0.01873535 | 0.1554129 |
| ERC2 | 487.0380296 | -0.462922413 | 0.271708696 | -1.703745296 | 0.088428682 | 0.341796568 |
| ERCC1 | 1507.773704 | 0.451192541 | 0.307974789 | 1.465030768 | 0.142912524 | 0.433233718 |

| | | | | | | |
|---|---|---|---|---|---|---|
| ERCC2 | 2338.164954 | 0.23928577 | 0.303759951 | 0.787746274 | 0.430845132 | 0.7175707 |
| ERCC3 | 1787.109984 | -0.387461274 | 0.368176128 | -1.052380217 | 0.292625141 | 0.605732845 |
| ERCC4 | 363.0737125 | 0.042039544 | 0.298802116 | 0.140693597 | 0.888112004 | 0.955943827 |
| ERCC5 | 904.7310115 | -0.559069222 | 0.271975778 | -2.055584601 | 0.039822573 | 0.229888601 |
| ERCC6 | 330.2639375 | -0.738710076 | 0.388303284 | -1.902404914 | 0.057118239 | 0.276585 |
| ERCC6L | 563.6380947 | 0.388425675 | 0.362785723 | 1.070675196 | 0.284315499 | 0.598246396 |
| ERCC6L2 | 622.6395706 | -0.421041379 | 0.49210094 | -0.855599623 | 0.392219279 | 0.691514794 |
| ERCC8 | 668.0523059 | 0.474386192 | 0.266719529 | 1.778595642 | 0.075306079 | 0.314600339 |
| ERF | 1139.792651 | 0.26612877 | 0.474324961 | 0.561068449 | 0.574750876 | 0.808359708 |
| ERG | 117.0048379 | -1.9253626 | 0.634455065 | -3.034671338 | 0.002407979 | 0.047209009 |
| ERG28 | 1901.488345 | 0.367996137 | 0.331667807 | 1.109532276 | 0.267200629 | 0.580473201 |
| ERGIC1 | 2348.408578 | -0.097023147 | 0.266716992 | -0.363768149 | 0.716031149 | 0.879658965 |
| ERGIC2 | 985.723857 | 0.354690619 | 0.230512996 | 1.538701178 | 0.123877265 | 0.405045568 |
| ERGIC3 | 5299.858847 | 0.002224532 | 0.251392274 | 0.008848848 | 0.992939733 | 0.997243081 |
| ERH | 5933.586562 | 0.826176221 | 0.325163855 | 2.540799686 | 0.011059926 | 0.11934855 |
| ERI1 | 1061.429201 | -0.001279864 | 0.302635856 | -0.004229055 | 0.996625713 | 0.998431718 |
| ERI2 | 643.4435005 | 0.175042154 | 0.334750657 | 0.522903094 | 0.601041691 | 0.824625774 |
| ERI3 | 1181.135282 | 0.204190004 | 0.23804035 | 0.857795764 | 0.391005249 | 0.690530747 |
| ERICH1 | 895.5254149 | -0.288408421 | 0.42594919 | -0.677095832 | 0.498345153 | 0.762744065 |
| ERICH2 | 90.80925059 | 0.557987149 | 0.633737727 | 0.880470146 | 0.378604671 | 0.681253679 |
| ERICH5 | 43.92459837 | -1.597064071 | 0.746709271 | -2.138803058 | 0.032451622 | 0.205554752 |
| ERLEC1 | 1204.858015 | 0.301489524 | 0.283919549 | 1.061883639 | 0.288288513 | 0.601673803 |
| ERLEC1P1 | 51.60221667 | 1.661601501 | 0.578154902 | 2.873972867 | 0.004053438 | 0.065699472 |
| ERLIN1 | 1416.635585 | 0.412208978 | 0.246429114 | 1.672728401 | 0.094380776 | 0.352325479 |
| ERLIN2 | 2332.692514 | 0.004535974 | 0.253248802 | 0.017911136 | 0.985709745 | 0.994758368 |
| ERMAP | 222.7979658 | -0.539346919 | 0.359753767 | -1.499211316 | 0.13381882 | 0.420559542 |
| ERMARD | 290.4815154 | -0.027698849 | 0.296232563 | -0.093503729 | 0.925503387 | 0.972124891 |
| ERMN | 65.1181263 | 0.541403299 | 0.554254946 | 0.976812751 | 0.328661862 | 0.638020359 |
| ERMP1 | 3405.957119 | -0.720261454 | 0.37625044 | -1.914313919 | 0.055580057 | 0.272730921 |
| ERN1 | 176.544162 | -0.184994892 | 0.352807709 | -0.524350482 | 0.600034791 | 0.824066673 |
| ERN2 | 57.70957582 | -0.543282501 | 0.490184728 | -1.108321964 | 0.267722793 | 0.580978758 |
| ERO1A | 835.2254147 | 0.407446956 | 0.347259988 | 1.173319618 | 0.240667658 | 0.552353982 |
| ERO1B | 120.3806969 | 1.089166755 | 0.369545549 | 2.947313956 | 0.003205476 | 0.056744775 |
| ERP27 | 553.8112663 | -1.04441994 | 0.780982099 | -1.337316108 | 0.181119475 | 0.485749663 |
| ERP29 | 5857.187896 | 0.216343546 | 0.392126435 | 0.55171885 | 0.581140993 | 0.812579724 |
| ERP44 | 1397.817075 | -0.006853201 | 0.264895707 | -0.025871317 | 0.979359978 | 0.992858175 |
| ERRFI1 | 2220.123833 | -1.71525474 | 0.517990274 | -3.311364763 | 0.000928421 | 0.02597297 |
| ERV3-1 | 691.2383267 | -1.380476324 | 0.581821442 | -2.372680385 | 0.017659538 | 0.150007193 |

| | | | | | | |
|---|---|---|---|---|---|---|
| ERVK3-1 | 813.5128384 | -0.162580667 | 0.272728829 | -0.596125709 | 0.551091251 | 0.794583289 |
| ERVMER34-1 | 305.0833217 | 0.701383141 | 0.450671667 | 1.55630627 | 0.119635281 | 0.398898209 |
| ERVW-1 | 124.4858545 | -1.067692169 | 0.531776409 | -2.007784006 | 0.044666249 | 0.242965818 |
| ESCO1 | 1021.924256 | 0.181300528 | 0.29978664 | 0.604765203 | 0.545335018 | 0.79117123 |
| ESCO2 | 313.5501823 | 0.516685498 | 0.377554969 | 1.368504034 | 0.17115436 | 0.474474162 |
| ESD | 2488.242937 | 0.894396881 | 0.241582581 | 3.702240778 | 0.000213704 | 0.009905308 |
| ESF1 | 2069.053564 | 1.066998227 | 0.348406068 | 3.062513323 | 0.002194867 | 0.044516481 |
| ESM1 | 17.6978828 | -6.761316902 | 4.409436568 | -1.533374343 | 0.125183647 | 0.407021239 |
| ESPL1 | 1249.038023 | 0.043444783 | 0.472211225 | 0.09200286 | 0.926695768 | 0.972261514 |
| ESPN | 159.4492553 | 0.158976996 | 0.647427237 | 0.245551912 | 0.806029118 | 0.919089559 |
| ESPNL | 25.05424152 | 0.752562029 | 0.675394276 | 1.114255859 | 0.265169438 | 0.579082415 |
| ESPNP | 17.11116992 | -1.986190526 | 0.860593356 | -2.307931512 | 0.021002946 | 0.164636113 |
| ESR2 | 18.77192182 | -0.005716649 | 0.700611279 | -0.008159516 | 0.99348972 | 0.99730871 |
| ESRP1 | 4040.391978 | -1.54659455 | 0.677289545 | -2.28350572 | 0.022400596 | 0.170396445 |
| ESRP2 | 1021.688028 | -1.416131325 | 0.533617902 | -2.653830239 | 0.007958385 | 0.100279622 |
| ESRRA | 512.5695011 | 0.658643946 | 0.34922555 | 1.886013055 | 0.059293195 | 0.282340574 |
| ESRRAP2 | 11.0215043 | -0.839568222 | 0.946652561 | -0.886881055 | 0.375142943 | 0.679214334 |
| ESRRG | 141.8819204 | -0.073119306 | 0.399403746 | -0.183071157 | 0.854742187 | 0.941593381 |
| ESS2 | 438.5660905 | -0.437743055 | 0.408237065 | -1.072276608 | 0.28359581 | 0.597477022 |
| ESYT1 | 3463.244847 | -0.479214483 | 0.283944132 | -1.687706943 | 0.091467498 | 0.347000104 |
| ESYT2 | 4482.085411 | -1.206737365 | 0.312090381 | -3.866627865 | 0.000110351 | 0.00659106 |
| ESYT3 | 31.849479 | -0.807230894 | 0.625751871 | -1.290017547 | 0.197044566 | 0.50258876 |
| ETAA1 | 1161.576905 | 0.12060208 | 0.247675652 | 0.486935549 | 0.626304005 | 0.835767952 |
| ETF1 | 4552.716563 | -0.155327442 | 0.217486958 | -0.714191986 | 0.475108472 | 0.748791879 |
| ETFA | 2513.89724 | 0.755111788 | 0.307543382 | 2.455301696 | 0.014076639 | 0.134821147 |
| ETFB | 845.8806282 | 1.095569688 | 0.364668027 | 3.00429324 | 0.002661986 | 0.050317756 |
| ETFBKMT | 48.84250344 | 0.163456946 | 0.523500518 | 0.312238366 | 0.754859377 | 0.897503925 |
| ETFDH | 323.6352984 | 0.352926325 | 0.370446708 | 0.952704714 | 0.340739704 | 0.647546292 |
| ETFRF1 | 283.326115 | 0.982693898 | 0.297729889 | 3.300622261 | 0.000964707 | 0.026561825 |
| ETHE1 | 296.2649082 | 0.593478886 | 0.439595112 | 1.350057974 | 0.176997387 | 0.481361196 |
| ETNK1 | 1885.078091 | 0.326221938 | 0.264961317 | 1.231205905 | 0.218245863 | 0.526623336 |
| ETNK2 | 425.5492091 | -0.08575835 | 0.329355427 | -0.26038238 | 0.794568833 | 0.914875817 |
| ETNPPL | 21.5769345 | 0.264945718 | 0.757292658 | 0.349859087 | 0.726444453 | 0.884630646 |
| ETS1 | 306.5236634 | -0.235550013 | 0.385553512 | -0.610939873 | 0.541239387 | 0.788244479 |
| ETS2 | 342.7841728 | -0.558549725 | 0.29658232 | -1.883287331 | 0.059661439 | 0.282669434 |
| ETV1 | 73.52820032 | 0.405974519 | 0.566501056 | 0.716635061 | 0.473599305 | 0.747811725 |
| ETV2 | 14.76307718 | 0.692455517 | 0.902064542 | 0.767634116 | 0.442704588 | 0.726083385 |
| ETV3 | 1379.300861 | -0.920675264 | 0.258569658 | -3.560646944 | 0.000369942 | 0.014464931 |
| ETV4 | 170.194502 | 0.27330269 | 0.388561715 | 0.703370093 | 0.481825139 | 0.753118606 |
| ETV5 | 96.4134442 | 0.767234829 | 0.590449589 | 1.299407847 | 0.193804 | 0.499845061 |
| ETV6 | 395.2047152 | -0.022725057 | 0.410556728 | -0.055351808 | 0.955858189 | 0.98434188 |
| ETV7 | 37.48986525 | 0.51830474 | 0.670755667 | 0.772717646 | 0.439689501 | 0.723249422 |
| EVA1A | 42.48988424 | 0.945493739 | 0.634199471 | 1.490845993 | 0.136001933 | 0.423778383 |

| | | | | | | |
|---|---|---|---|---|---|---|
| EVA1B | 50.03013838 | 0.349671577 | 0.800501042 | 0.436815892 | 0.662244873 | 0.853821447 |
| EVA1C | 12.97901544 | 0.112418266 | 0.799657753 | 0.140582975 | 0.888199399 | 0.955943827 |
| EVC | 253.2493086 | -0.50848561 | 0.694800246 | -0.731844314 | 0.464263598 | 0.741170477 |
| EVC2 | 242.5403926 | -0.921205558 | 0.505590582 | -1.822038605 | 0.068449128 | 0.301521987 |
| EVI5 | 325.1860502 | 0.037897845 | 0.306973532 | 0.123456395 | 0.901745702 | 0.962206864 |
| EVI5L | 260.4917857 | -0.195717835 | 0.510459563 | -0.383414965 | 0.701412109 | 0.872388416 |
| EVL | 1334.242232 | -0.067146949 | 0.28952735 | -0.231919191 | 0.816600778 | 0.923554512 |
| EWSR1 | 10520.4719 | -0.147224491 | 0.225806551 | -0.651993889 | 0.514405114 | 0.77285333 |
| EXD2 | 1065.12997 | -0.688917233 | 0.28818176 | -2.390565016 | 0.01682247 | 0.146590292 |
| EXD3 | 182.9283152 | 0.441679126 | 0.522065518 | 0.846022407 | 0.397540244 | 0.696188739 |
| EXO1 | 481.0818117 | 0.158391781 | 0.389396838 | 0.406761858 | 0.684182903 | 0.863826466 |
| EXO5 | 286.1227825 | -0.234996894 | 0.279064072 | -0.842089388 | 0.399737921 | 0.697621534 |
| EXOC1 | 1292.020004 | -0.137224412 | 0.244974085 | -0.560158893 | 0.575371063 | 0.808535976 |
| EXOC2 | 1610.666901 | -0.209976461 | 0.383997916 | -0.546816669 | 0.58450469 | 0.814879678 |
| EXOC3 | 1106.073594 | -0.618925694 | 0.339722146 | -1.821858543 | 0.068476452 | 0.301554791 |
| EXOC4 | 2766.781312 | -0.356007779 | 0.226449902 | -1.572126002 | 0.115921332 | 0.392378968 |
| EXOC5 | 3772.734507 | -0.139335673 | 0.265645345 | -0.524517654 | 0.599918544 | 0.824047262 |
| EXOC6 | 509.4048479 | 0.928673132 | 0.429159406 | 2.163935169 | 0.030469325 | 0.199054492 |
| EXOC6B | 1111.395946 | -0.747159058 | 0.290990487 | -2.567640841 | 0.010239318 | 0.115067186 |
| EXOC7 | 2971.57785 | -0.617084392 | 0.417194882 | -1.479127426 | 0.139106261 | 0.427760708 |
| EXOC8 | 432.0743129 | -0.092353685 | 0.380153475 | -0.242937896 | 0.808053508 | 0.919580875 |
| EXOG | 206.173528 | 0.529232042 | 0.404391503 | 1.308712072 | 0.190631901 | 0.496058205 |
| EXOSC1 | 1076.958218 | 0.381844698 | 0.334109794 | 1.142871909 | 0.253091777 | 0.565800563 |
| EXOSC10 | 2756.399203 | -0.561071118 | 0.369717947 | -1.517565276 | 0.129124026 | 0.413453061 |
| EXOSC2 | 1198.016459 | 0.378372653 | 0.27745684 | 1.363717157 | 0.172656602 | 0.476336299 |
| EXOSC3 | 687.2959145 | 0.784269056 | 0.325428992 | 2.409954477 | 0.015954511 | 0.142746012 |
| EXOSC4 | 260.4803772 | 1.085460605 | 0.453693786 | 2.392496083 | 0.016734207 | 0.146249211 |
| EXOSC5 | 315.0965936 | 0.504529871 | 0.40817258 | 1.236069976 | 0.216432554 | 0.524101879 |
| EXOSC6 | 830.5827445 | 0.688079712 | 0.437253934 | 1.573638699 | 0.115571001 | 0.391542577 |
| EXOSC7 | 1352.583088 | 1.008103044 | 0.432515089 | 2.330792774 | 0.019764289 | 0.159152884 |
| EXOSC8 | 1801.731712 | 0.809837401 | 0.303037356 | 2.672401227 | 0.007531054 | 0.096389041 |
| EXOSC9 | 1700.510425 | 0.521803822 | 0.230294428 | 2.265811759 | 0.023462911 | 0.174992377 |
| EXPH5 | 174.1924965 | -2.576756355 | 0.851291596 | -3.026878648 | 0.002470931 | 0.047998079 |
| EXT1 | 544.1356388 | 0.072416818 | 0.350720497 | 0.206480143 | 0.836415874 | 0.932828437 |
| EXT2 | 1974.261614 | -0.125121583 | 0.274803853 | -0.455312332 | 0.648884556 | 0.847033866 |
| EXTL1 | 55.22151216 | -0.21014553 | 0.668188966 | -0.31450015 | 0.753141203 | 0.8966417 |
| EXTL2 | 1928.294594 | 0.372898991 | 0.264474481 | 1.409962086 | 0.158550878 | 0.458254025 |

| | | | | | | |
|---|---|---|---|---|---|---|
| EXTL3 | 2069.067504 | 0.465225521 | 0.517547341 | 0.898904282 | 0.368703647 | 0.67460916 |
| EYA1 | 374.9839397 | -1.767280564 | 0.609025966 | -2.901814806 | 0.003710078 | 0.062818742 |
| EYA2 | 423.8492498 | -2.285317923 | 0.622658871 | -3.670256747 | 0.000242307 | 0.010655185 |
| EYA3 | 1042.499225 | -0.054001802 | 0.283489476 | -0.190489617 | 0.848925479 | 0.938569494 |
| EYA4 | 124.1625413 | 0.202380402 | 0.592115403 | 0.34179216 | 0.73250731 | 0.888105842 |
| EZH1 | 905.9569625 | -0.399302747 | 0.441721943 | -0.903968555 | 0.366012076 | 0.672491566 |
| EZH2 | 2052.129147 | 0.298407001 | 0.301168483 | 0.990830774 | 0.32176822 | 0.631424869 |
| EZR | 6082.892395 | -1.759290559 | 0.490748448 | -3.58491314 | 0.00033719 | 0.013547304 |
| F10 | 72.19285776 | 0.37466709 | 0.567067167 | 0.66071025 | 0.508798149 | 0.769282113 |
| F11R | 6762.38803 | -0.292868158 | 0.374862525 | -0.781268169 | 0.434644788 | 0.719554351 |
| F2R | 1366.903592 | 0.550888591 | 0.402906611 | 1.367286058 | 0.17153566 | 0.474879827 |
| F2RL1 | 2217.895961 | -0.281072977 | 0.342118939 | -0.821565093 | 0.411324463 | 0.704686727 |
| F2RL3 | 23.4864523 | -0.12046858 | 0.648443643 | -0.185781111 | 0.852616418 | 0.940251776 |
| F3 | 106.7533957 | -0.009768722 | 0.450492239 | -0.02168455 | 0.982699588 | 0.993518428 |
| F5 | 50.096632 | -0.588687623 | 0.64024327 | -0.91947491 | 0.357847223 | 0.665387943 |
| F8 | 107.7514277 | 0.062924809 | 0.39096467 | 0.160947559 | 0.872134704 | 0.949014172 |
| F8A1 | 403.0787759 | 0.46210835 | 0.447282392 | 1.033146752 | 0.301535227 | 0.612395038 |
| F8A3 | 67.29642409 | -0.451415874 | 0.464661241 | -0.971494573 | 0.331302054 | 0.639138284 |
| FA2H | 36.13456571 | -1.500575473 | 0.608031107 | -2.4679255 | 0.01358986 | 0.132501135 |
| FAAH | 123.0943875 | -0.533724173 | 0.402066886 | -1.327451206 | 0.184359471 | 0.488866012 |
| FAAH2 | 142.8591289 | -0.514181768 | 0.513102527 | -1.002103364 | 0.316293673 | 0.626189654 |
| FAAP100 | 462.6764558 | -0.025968481 | 0.509794847 | -0.05093908 | 0.959374064 | 0.985887959 |
| FAAP20 | 495.1934423 | 0.415199234 | 0.426035243 | 0.974565462 | 0.329775853 | 0.638548751 |
| FAAP24 | 147.8216701 | -0.373078887 | 0.4419918 | -0.844085541 | 0.398621607 | 0.697197187 |
| FABP3 | 72.23541947 | 0.206845981 | 0.525476883 | 0.393634786 | 0.693850695 | 0.868802318 |
| FABP5 | 1145.707469 | 0.932314839 | 0.419386035 | 2.223046933 | 0.026212635 | 0.184791771 |
| FABP5P7 | 34.31537768 | -1.80174634 | 1.082129106 | -1.665001274 | 0.095912502 | 0.35559349 |
| FABP6 | 18.36093123 | 1.021004678 | 0.728866079 | 1.400812451 | 0.161270164 | 0.46167973 |
| FABP7 | 421.7645567 | 0.195349211 | 0.397495924 | 0.491449595 | 0.623108494 | 0.833907812 |
| FADD | 607.4247776 | 0.425973822 | 0.278250506 | 1.530900442 | 0.125794 | 0.407956557 |
| FADS1 | 5402.867782 | 0.377500849 | 0.440659593 | 0.856672258 | 0.391626039 | 0.691177133 |
| FADS2 | 9313.578476 | -0.34221792 | 0.384972178 | -0.888941953 | 0.374034277 | 0.678520659 |
| FADS3 | 697.4748832 | -0.235871028 | 0.259082613 | -0.910408559 | 0.362607085 | 0.669723897 |
| FAF1 | 1502.240143 | -0.177246936 | 0.249151376 | -0.711402596 | 0.476834791 | 0.749746108 |
| FAF2 | 1503.474427 | -0.493226728 | 0.250640647 | -1.967864089 | 0.049083678 | 0.255021315 |
| FAH | 487.7059573 | 0.207831085 | 0.25132655 | 0.826936449 | 0.408273083 | 0.702713988 |
| FAHD1 | 417.9266175 | 0.441000599 | 0.392188602 | 1.124460519 | 0.260817711 | 0.57415519 |
| FAHD2A | 842.5908096 | 0.068571434 | 0.26270354 | 0.261022117 | 0.794075451 | 0.914585384 |
| FAHD2B | 558.1209275 | 0.216430794 | 0.288164276 | 0.751067401 | 0.452612093 | 0.733138807 |
| FAHD2CP | 192.7008486 | 0.679190248 | 0.39849754 | 1.704377519 | 0.088310581 | |

0.341701126
FAIM 157.2996388 -0.201132192 0.353383814 -0.569160736 0.569247064
0.805601418
FAIM2 90.04948737 1.422662088 0.751070154 1.894180031 0.058201118 0.279509071
FAM102A 789.1624445 0.019533381 0.327180199 0.059702211 0.952392811
0.982774543
FAM102B 547.796785 -1.028685337 0.313275804 -3.283641202
0.001024753 0.027467362
FAM104A 842.9784731 0.020709251 0.28694166 0.07217234 0.942464757
0.978574897
FAM104B 311.9732499 0.719737908 0.33198374 2.16799144 0.030159338
0.198185614
FAM107A 25.71753664 1.298854181 0.907115842 1.4318504 0.15218664
0.448119169
FAM107B 1330.888418 -0.910462015 0.451651065 -2.015852692
0.043815382 0.240941849
FAM110A 507.1881112 -0.336820787 0.741734855 -0.454098637
0.649757834 0.84729896
FAM110B 412.0676756 -0.824905262 0.511488449 -1.61275443 0.10679787
0.374619766
FAM110C 54.69490194 -2.261165962 0.715055547 -3.16222421 0.00156569
0.036655984
FAM110D 65.87958122 1.35585193 0.74207422 1.827110946 0.06768309
0.299102871
FAM111A 691.632448 -0.546073792 0.312911795 -1.745136491
0.080961113 0.325493577
FAM111B 553.5213446 -0.069568359 0.372693725 -0.186663619
0.851924383 0.939761856
FAM114A1 236.1910533 -0.508092102 0.380271599 -1.336129502
0.181506949 0.485768051
FAM114A2 508.3608553 0.053819402 0.251128641 0.21431009 0.83030525
0.930253375
FAM117A 728.300115 0.267373681 0.272399247 0.981550733 0.326321229
0.636162664
FAM117B 805.6727735 0.068237063 0.354226284 0.192636929 0.847243321
0.938145276
FAM118A 568.7423404 -0.36657076 0.327612673 -1.118915081 0.263176369
0.576857184
FAM118B 959.9248028 0.126661876 0.244670935 0.51768256 0.604679769
0.825894731
FAM120A 1532.229536 -0.054609418 0.240582786 -0.226988054
0.820433031 0.92511751
FAM120AOS 905.4565478 0.017412903 0.269165147 0.064692264 0.948419022
0.98134156
FAM120B 553.7265496 -0.524784827 0.363882784 -1.442180971
0.149251327 0.443978801
FAM120C 147.4941566 0.075736409 0.34457619 0.219795828 0.826030169
0.927518591
FAM122A 362.640989 0.132174095 0.355507151 0.371790258 0.710049018
0.876599663
FAM122B 1056.566188 0.222472453 0.261982846 0.849187101 0.39577719
0.69442187
FAM122C 157.3653824 -0.295570047 0.335281954 -0.881556683
0.378016589 0.681253679
FAM124A 117.634416 0.635335524 0.457228615 1.389535788 0.164669886
0.46700822
FAM126A 1443.656646 0.464870332 0.3688134 1.260448597 0.20750758
0.514335219
FAM126B 342.3383811 -0.277297046 0.341902511 -0.811041267
0.417341972 0.708794572
FAM129A 32.79153278 -0.245277067 0.661207046 -0.370953499
0.710672167 0.876838602
FAM129B 1967.746021 -0.195903308 0.350255553 -0.559315354
0.575946519 0.808969969

| | | | | | | |
|---|---|---|---|---|---|---|
| FAM129C | 31.87760634 | -0.441697622 | 0.647376141 | -0.682289003 | 0.495056225 | 0.760514211 |
| FAM131A | 371.2167865 | 0.628183082 | 0.332264088 | 1.890613836 | 0.058675912 | 0.280503663 |
| FAM131B | 568.1247204 | 0.425823452 | 0.560288837 | 0.760007024 | 0.447250386 | 0.728949036 |
| FAM131C | 44.95371526 | 0.595351815 | 0.793086758 | 0.750676781 | 0.4528472 | 0.733363031 |
| FAM133A | 25.96195047 | 0.763935992 | 0.597430765 | 1.278702129 | 0.20100197 | 0.506881795 |
| FAM133B | 1584.706041 | 0.659363923 | 0.390935377 | 1.686631505 | 0.09167423 | 0.347610534 |
| FAM135A | 973.0787413 | 0.013649931 | 0.329795813 | 0.041389037 | 0.966985752 | 0.988749915 |
| FAM136A | 1764.133044 | 0.185859426 | 0.335073417 | 0.554682696 | 0.579111714 | 0.810713649 |
| FAM13A | 712.9514943 | -0.039705446 | 0.333514042 | -0.119051797 | 0.905234319 | 0.964417827 |
| FAM13B | 743.8440643 | -0.657533586 | 0.400115535 | -1.643359299 | 0.100308616 | 0.36243601 |
| FAM13C | 82.64725353 | 0.42180917 | 0.473475514 | 0.89087853 | 0.372994339 | 0.67811865 |
| FAM149A | 25.2483743 | 0.434175046 | 0.614843788 | 0.706155051 | 0.48009172 | 0.752036291 |
| FAM149B1 | 875.408629 | -0.079440354 | 0.267585107 | -0.296878832 | 0.766559017 | 0.903470079 |
| FAM151B | 35.82283542 | 0.156063781 | 0.599937219 | 0.260133521 | 0.794760781 | 0.914877916 |
| FAM153C | 22.76132765 | -0.112727661 | 0.68036899 | -0.165686066 | 0.868404012 | 0.947046732 |
| FAM155A | 65.35741315 | -0.031037275 | 0.416904514 | -0.074446963 | 0.940654742 | 0.97807441 |
| FAM155B | 100.5458844 | 0.424974824 | 0.458199603 | 0.927488416 | 0.353673004 | 0.660866257 |
| FAM156A | 546.3487989 | -0.857798764 | 0.529119808 | -1.621180593 | 0.104978914 | 0.37133017 |
| FAM157A | 301.6918254 | -0.759862114 | 0.324207904 | -2.343749504 | 0.019090988 | 0.156852604 |
| FAM157B | 51.92031809 | -0.785603275 | 0.692377062 | -1.134646594 | 0.256523423 | 0.568798135 |
| FAM160A1 | 584.6466757 | -1.078807889 | 0.355449081 | -3.035056059 | 0.00240491 | 0.047209009 |
| FAM160A2 | 915.0404579 | -0.747378648 | 0.275599836 | -2.711825445 | 0.006691382 | 0.089444885 |
| FAM160B1 | 695.6442475 | -0.238591359 | 0.330982621 | -0.720857663 | 0.470997093 | 0.745954369 |
| FAM160B2 | 514.6119167 | -0.773126206 | 0.427090134 | -1.810217902 | 0.070262003 | 0.304642709 |
| FAM161A | 761.7269437 | 0.089426745 | 0.317521808 | 0.281639695 | 0.778219799 | 0.908108663 |
| FAM161B | 70.21677804 | -0.200665816 | 0.514892254 | -0.389723898 | 0.696740723 | 0.870026319 |
| FAM162A | 1029.703564 | 1.09003123 | 0.425772 | 2.560128964 | 0.010463333 | 0.116207335 |
| FAM162B | 14.52251029 | 1.874724496 | 0.825096649 | 2.272127149 | 0.023078832 | 0.173590826 |
| FAM166B | 35.25232768 | -0.243901618 | 0.600467575 | -0.40618616 | 0.684605823 | 0.863857185 |
| FAM167A | 197.5260721 | 0.808384133 | 0.53515075 | 1.510572737 | 0.130897344 | 0.416533681 |
| FAM167B | 50.82467851 | 0.519160189 | 0.487726341 | 1.064449764 | 0.287125 | 0.601404581 |
| FAM168A | 1201.216068 | -0.643362106 | 0.405841266 | -1.585255517 | 0.112908291 | 0.38661645 |

| | | | | | | |
|---|---|---|---|---|---|---|
| FAM168B | 3713.229886 | -0.133618179 | 0.377246639 | -0.354193159 | 0.72319413 | 0.883273431 |
| FAM169A | 1471.800412 | -1.121314158 | 0.362946066 | -3.089478747 | 0.002005081 | 0.042131685 |
| FAM171A1 | 1468.264977 | -0.03646818 | 0.231678722 | -0.157408411 | 0.874922985 | 0.949740014 |
| FAM171A2 | 373.4320544 | 0.282722099 | 0.564855952 | 0.500520704 | 0.616708482 | 0.831694047 |
| FAM171B | 395.0006758 | 0.181924353 | 0.380002204 | 0.478745521 | 0.632119678 | 0.838768903 |
| FAM172A | 492.6882765 | 0.129132711 | 0.256528027 | 0.503386362 | 0.614692658 | 0.830705711 |
| FAM173A | 68.3337655 | 1.217541472 | 0.793265481 | 1.534847413 | 0.124821313 | 0.406452918 |
| FAM174A | 106.2181884 | 0.948694416 | 0.546630874 | 1.735530248 | 0.082646929 | 0.328745819 |
| FAM174B | 205.6981749 | -0.34288587 | 0.540516826 | -0.63436669 | 0.525841543 | 0.778448677 |
| FAM177A1 | 733.650765 | 0.507807051 | 0.394242488 | 1.288057648 | 0.197725897 | 0.50338976 |
| FAM178B | 14.70026118 | 1.165674688 | 0.816213923 | 1.428148498 | 0.153249121 | 0.44961176 |
| FAM183A | 64.79864251 | 0.508319889 | 0.529975918 | 0.959137713 | 0.337489374 | 0.645014619 |
| FAM184A | 742.3363991 | -1.254233908 | 0.397262986 | -3.157187939 | 0.001592986 | 0.03712319 |
| FAM184B | 10.91465143 | 0.897268664 | 1.015165602 | 0.883864329 | 0.376769455 | 0.68029735 |
| FAM185A | 120.3224374 | -0.180334574 | 0.358932761 | -0.502418818 | 0.615372945 | 0.830958873 |
| FAM185BP | 18.06468848 | -0.362016085 | 0.727318776 | -0.497740602 | 0.618666887 | 0.83229541 |
| FAM187A | 73.04923359 | -0.653188176 | 0.919301876 | -0.710526317 | 0.477377817 | 0.749955252 |
| FAM189A1 | 12.25226187 | 0.246972033 | 1.369533305 | 0.180332988 | 0.85689116 | 0.942343063 |
| FAM189A2 | 14.34231033 | -1.942546631 | 0.900963413 | -2.156077155 | 0.031077641 | 0.200942211 |
| FAM189B | 652.0529233 | 0.419109607 | 0.70426558 | 0.595101647 | 0.551775527 | 0.795324338 |
| FAM192A | 1991.133682 | -0.11647145 | 0.223524338 | -0.521068315 | 0.602319183 | 0.825021776 |
| FAM193A | 836.4402056 | -0.510926249 | 0.480918557 | -1.062396619 | 0.288055668 | 0.601673803 |
| FAM193B | 1477.577541 | -0.220021509 | 0.331527281 | -0.663660345 | 0.506907725 | 0.767724578 |
| FAM199X | 1007.421676 | -0.469632347 | 0.238419212 | -1.969775602 | 0.048864094 | 0.254713129 |
| FAM19A2 | 57.52221187 | 0.34827691 | 0.433920886 | 0.80262767 | 0.42218997 | 0.712133049 |
| FAM19A4 | 68.98429563 | 0.202422581 | 0.54772378 | 0.369570554 | 0.711702494 | 0.877287654 |
| FAM200A | 390.3673939 | 0.174412719 | 0.256225181 | 0.68070093 | 0.496060748 | 0.761056737 |
| FAM200B | 464.762289 | 0.3489449 | 0.327171927 | 1.066549027 | 0.286175528 | 0.60016159 |
| FAM204A | 732.8265938 | 0.21814986 | 0.303298639 | 0.719257631 | 0.471982196 | 0.746399031 |
| FAM206A | 640.2767556 | 0.14158787 | 0.265398431 | 0.533491737 | 0.593693229 | 0.820227665 |
| FAM207A | 276.5203278 | 0.341503149 | 0.361488284 | 0.944714293 | 0.344804758 | 0.651598528 |
| FAM20A | 192.8309184 | -4.930180248 | 0.588367335 | -8.379425494 | 5.32E-17 | 8.07E-13 |

| | | | | | | |
|---|---|---|---|---|---|---|
| FAM20B | 1627.190243 | 0.122291648 | 0.370142753 | 0.330390496 | 0.741104922 | 0.891943909 |
| FAM20C | 83.01687461 | -0.343034363 | 0.410815136 | -0.835009065 | 0.403712604 | 0.70028858 |
| FAM210A | 366.2402377 | 0.303052101 | 0.311670169 | 0.972348756 | 0.330877076 | 0.639084337 |
| FAM210B | 890.9620668 | 0.110266519 | 0.26979987 | 0.408697449 | 0.682761706 | 0.863109561 |
| FAM214A | 287.0397615 | -0.649813992 | 0.299339541 | -2.170825778 | 0.029944345 | 0.197343897 |
| FAM214B | 484.0234391 | 0.106374202 | 0.271020753 | 0.392494674 | 0.694692746 | 0.86892666 |
| FAM216A | 399.8902633 | 1.751573242 | 0.504881422 | 3.469276477 | 0.000521862 | 0.017923239 |
| FAM217B | 574.855635 | -0.211177584 | 0.254670868 | -0.829217671 | 0.406981248 | 0.701626421 |
| FAM219A | 565.4110758 | -0.505012712 | 0.253018469 | -1.995951969 | 0.045939151 | 0.246880221 |
| FAM219B | 1825.851551 | -0.008572463 | 0.270671127 | -0.031671141 | 0.97473431 | 0.991261106 |
| FAM220A | 253.5024328 | 2.014353224 | 1.809557581 | 1.113174428 | 0.265633523 | 0.579572403 |
| FAM221A | 446.379324 | 0.313483049 | 0.321454656 | 0.975201458 | 0.329460339 | 0.638506956 |
| FAM222A | 79.17968571 | 0.804093061 | 0.532365225 | 1.510416202 | 0.130937257 | 0.416533681 |
| FAM222B | 1349.75449 | -0.467782405 | 0.314103572 | -1.489261652 | 0.136418482 | 0.424447249 |
| FAM227A | 295.3795284 | 0.126702835 | 0.412782921 | 0.306947862 | 0.758883058 | 0.899440455 |
| FAM227B | 227.2612218 | 0.095969387 | 0.321163606 | 0.298817753 | 0.765079104 | 0.902991439 |
| FAM228B | 250.0699726 | 0.304903667 | 0.379164388 | 0.804146374 | 0.421312445 | 0.711694812 |
| FAM229A | 86.10634769 | -0.761586587 | 0.50636526 | -1.50402614 | 0.132574636 | 0.418641974 |
| FAM229B | 220.0959407 | 1.017325011 | 0.377409033 | 2.695550247 | 0.007027248 | 0.0921438 |
| FAM234A | 2061.853998 | -0.013813939 | 0.323168991 | -0.042745249 | 0.965904609 | 0.988700292 |
| FAM234B | 968.524656 | 0.522918524 | 0.371174746 | 1.408820319 | 0.158888305 | 0.458846746 |
| FAM241A | 120.7889134 | -0.430188813 | 0.456281709 | -0.942814066 | 0.345776016 | 0.652517297 |
| FAM241B | 237.7572142 | 0.555528522 | 0.43713052 | 1.270852748 | 0.203781039 | 0.509986087 |
| FAM32A | 2061.03252 | -0.184096898 | 0.277277448 | -0.663944723 | 0.506725691 | 0.767639613 |
| FAM3A | 524.6858544 | 0.117693412 | 0.353611624 | 0.332832418 | 0.739260786 | 0.891550525 |
| FAM3C | 1870.808502 | 0.564167443 | 0.276559434 | 2.039950094 | 0.041355297 | 0.233581982 |
| FAM3C2 | 167.0133829 | -0.338949282 | 2.205419735 | -0.153689239 | 0.87785477 | 0.951077249 |
| FAM3D | 20.81383799 | -0.436169674 | 0.735278567 | -0.593203302 | 0.553045103 | 0.795736652 |
| FAM43A | 107.2882336 | 1.449252241 | 0.610408795 | 2.37423224 | 0.017585486 | 0.14971347 |
| FAM45A | 2228.883477 | 0.174974302 | 0.305026185 | 0.573636987 | 0.566213477 | 0.804380669 |
| FAM47E | 174.3158814 | 0.568454516 | 0.433939982 | 1.309984191 | 0.190201184 | 0.495717092 |
| FAM47E-STBD1 | 150.336825 | -0.079455106 | 0.338457255 | -0.234756694 | 0.814397582 | 0.92237594 |
| FAM49A | 142.694812 | -0.795238454 | 0.748026338 | -1.06311558 | 0.28772954 | 0.601673803 |
| FAM49B | 2933.633009 | 0.584328481 | 0.242832814 | 2.406299507 | 0.016115047 | |

0.143654707
FAM50A 1547.122896 0.151772658 0.269122192 0.56395445 0.572785132
0.807670159
FAM50B 36.59596041 0.677580884 0.682658746 0.99256164 0.320923627
0.630501469
FAM53A 20.28712523 0.445911899 0.858431315 0.519449712 0.603447173
0.825211561
FAM53B 655.1704025 0.366723191 0.570853872 0.642411674 0.520605922
0.775597169
FAM53C 2521.731081 -0.261830973 0.317172978 -0.825514755
0.409079409 0.703484892
FAM57A 444.7864352 0.356310607 0.288140715 1.23658542 0.216241035
0.523980406
FAM57B 223.0179317 1.147455413 0.526346071 2.180039858 0.029254507
0.195774208
FAM71F1 31.4581176 0.830526518 0.59709182 1.390952766 0.164239751
0.466312149
FAM71F2 12.83498396 -1.548401894 0.970444358 -1.595559685
0.110587134 0.382690751
FAM72A 506.1145335 1.488170289 0.439597595 3.385301256 0.000711002
0.021968651
FAM72B 476.9893365 1.551144026 0.352467993 4.400808177 1.08E-05
0.001152231
FAM72C 242.5716058 1.548986108 0.392156829 3.949914916 7.82E-05
0.00513443
FAM72D 264.2213266 1.64767802 0.331343199 4.972723215 6.60E-07
0.000139266
FAM76A 329.1443265 0.403545536 0.271167522 1.488177985 0.136703961
0.424728769
FAM76B 905.9720564 -0.138204558 0.284263717 -0.486184307
0.626836497 0.836041947
FAM78A 99.69645894 0.440997713 0.486540789 0.906394126 0.364727281
0.671614642
FAM78B 49.14790316 0.293730998 0.717050115 0.409638032 0.682071494
0.862735748
FAM81A 10.70643871 0.307641078 0.975524435 0.315359684 0.752488574
0.896586461
FAM83B 433.9287098 -1.34295492 0.664508925 -2.020973489 0.04328251
0.23938715
FAM83D 865.7308929 0.886419717 0.308565252 2.87271399 0.004069624
0.065751076
FAM83E 29.11091684 -1.260457949 0.712731319 -1.768489634
0.076979084 0.31844659
FAM83F 57.63022616 -1.349584378 0.699663843 -1.92890399 0.053742782
0.267936823
FAM83G 724.3331776 -0.066303181 0.312991092 -0.211837277
0.832233985 0.931245375
FAM83H 1711.732743 -0.345976821 0.328195959 -1.054177578
0.291801623 0.604936107
FAM84A 88.30763853 -1.142704383 0.915193074 -1.248593784
0.211813687 0.519047883
FAM84B 3156.853498 -0.701099421 0.358768714 -1.954182163
0.050679678 0.259050335
FAM86B1 126.7827683 0.156008316 0.471759329 0.330694713 0.740875096
0.891943909
FAM86B3P 57.08205583 0.340579822 0.520409247 0.654446138 0.512824418
0.771819984
FAM86C1 196.0683 -0.116052736 0.470701259 -0.246552847
0.805254301 0.918811146
FAM86C2P 109.9195185 0.446607419 0.364916763 1.223861069 0.221004642
0.529761641
FAM86DP 198.2674915 0.055354359 0.30578576 0.181023339 0.856349257
0.942245037
FAM86EP 76.7482764 0.655717075 0.411948682 1.591744561 0.111442111

| | | | | | | |
|---|---|---|---|---|---|---|
| | | | | | | 0.383985527 |
| FAM86HP | 12.85666281 | 0.348917428 | 0.871691336 | 0.400276352 | 0.688952984 | 0.867388971 |
| FAM86JP | 41.27853589 | 1.057584102 | 0.718364737 | 1.47221049 | 0.140964045 | 0.430713136 |
| FAM89A | 973.4963593 | -1.528557085 | 0.814208126 | -1.877354248 | 0.060469558 | 0.284105089 |
| FAM89B | 437.7612832 | 0.67405212 | 0.412382333 | 1.634532001 | 0.102147193 | 0.366611924 |
| FAM8A1 | 617.2011468 | 0.078296465 | 0.273319707 | 0.286464763 | 0.774522178 | 0.90635568 |
| FAM91A1 | 819.4358228 | -0.207718101 | 0.308486851 | -0.673345073 | 0.500727784 | 0.763638219 |
| FAM92A | 1060.841159 | 0.481791249 | 0.38064691 | 1.265716957 | 0.205614435 | 0.511932925 |
| FAM95C | 51.92676381 | -0.232857843 | 0.509542842 | -0.456993649 | 0.647675611 | 0.846597949 |
| FAM98A | 1681.511146 | -0.748892757 | 0.261700828 | -2.861636941 | 0.004214594 | 0.067092973 |
| FAM98B | 2531.335012 | 0.203667915 | 0.302365517 | 0.673581818 | 0.500577216 | 0.763638219 |
| FAM98C | 137.2821857 | 0.54844454 | 0.42306723 | 1.296353159 | 0.194853842 | 0.500501471 |
| FAN1 | 563.6105117 | -0.074455789 | 0.385653414 | -0.193063996 | 0.846908849 | 0.938145276 |
| FANCA | 638.745807 | 0.05208396 | 0.420090422 | 0.123982736 | 0.901328944 | 0.961878265 |
| FANCB | 140.6352274 | 0.103945721 | 0.388676285 | 0.267435203 | 0.789134098 | 0.912566004 |
| FANCC | 693.1578573 | -0.017791227 | 0.283607268 | -0.062731913 | 0.949979984 | 0.981636196 |
| FANCD2 | 1511.049458 | 0.008207971 | 0.346974583 | 0.023655829 | 0.98112714 | 0.993241851 |
| FANCE | 656.6863127 | 0.212119596 | 0.300431029 | 0.706050893 | 0.480156488 | 0.752060096 |
| FANCF | 663.0119719 | 0.560396052 | 0.357964988 | 1.565505206 | 0.117464498 | 0.395641036 |
| FANCG | 1336.287758 | 0.169126242 | 0.236924979 | 0.713838796 | 0.475326867 | 0.748902679 |
| FANCI | 2337.512839 | 0.278014632 | 0.381280117 | 0.729161106 | 0.46590312 | 0.742755984 |
| FANCL | 1011.314203 | 0.939224096 | 0.3082028 | 3.04742233 | 0.002308132 | 0.045893402 |
| FANCM | 337.4395676 | 0.286596624 | 0.461494387 | 0.621018657 | 0.534587348 | 0.783975317 |
| FANK1 | 201.5851304 | -0.46514896 | 0.600489639 | -0.77461613 | 0.438566523 | 0.72234206 |
| FAR1 | 2228.608605 | 0.169994081 | 0.247184282 | 0.687722048 | 0.491627833 | 0.758965432 |
| FAR2 | 114.8136604 | 0.454319432 | 0.358486242 | 1.267327387 | 0.205038252 | 0.51158559 |
| FAR2P2 | 216.5901084 | 0.419961375 | 0.409147157 | 1.026431122 | 0.304688415 | 0.614996026 |
| FAR2P3 | 144.3749826 | -0.384209291 | 0.73999666 | -0.519204087 | 0.603618429 | 0.825371355 |
| FARP1 | 2841.487631 | -1.180348379 | 0.381044875 | -3.097662389 | 0.001950535 | 0.041386806 |
| FARP2 | 571.6626614 | -0.068569397 | 0.361742982 | -0.189552804 | 0.849659573 | 0.938580023 |
| FARS2 | 296.3988555 | -0.183895418 | 0.274211687 | -0.670633042 | 0.502454328 | 0.764874032 |
| FARSA | 1477.315723 | 0.344832204 | 0.256184895 | 1.34602863 | 0.178293283 | 0.4825693 |
| FARSB | 1473.036114 | 0.244172021 | 0.298446788 | 0.818142568 | 0.413275785 | 0.705652445 |
| FAS | 204.2310186 | 1.027912903 | 0.472602435 | 2.175005517 | 0.029629703 | 0.196723074 |
| FASN | 7320.604205 | -0.178459868 | 0.618163531 | -0.288693621 | 0.772815845 | 0.905717252 |
| FASTK | 896.1312954 | 0.023537889 | 0.433510868 | 0.05429596 | 0.956699368 | 0.984539826 |
| FASTKD1 | 1560.711514 | 0.415492359 | 0.283172628 | 1.467275851 | 0.14230103 | 0.432548373 |
| FASTKD2 | 1213.803491 | 0.135094359 | 0.238749974 | 0.565840309 | 0.571502342 | 0.807274555 |
| FASTKD3 | 303.5422223 | 0.107261303 | 0.320063331 | 0.335125248 | 0.737530608 | 0.890991945 |
| FASTKD5 | 503.2661063 | -0.292411895 | 0.249607189 | -1.171488272 | 0.241402569 | 0.552719345 |

| | | | | | | |
|---|---|---|---|---|---|---|
| FAT1 | 4368.145718 | -0.748283289 | 0.464458038 | -1.611089116 | 0.1071603 | 0.375111686 |
| FAT2 | 48.81145875 | -1.748126845 | 0.707095472 | -2.472264235 | 0.013426024 | 0.131761226 |
| FAT3 | 133.5450356 | -0.035488638 | 0.836733979 | -0.042413286 | 0.966169237 | 0.988700292 |
| FAT4 | 171.0528617 | 0.598815638 | 0.70938524 | 0.844133208 | 0.398594973 | 0.697197187 |
| FAU | 9536.75548 | 1.053163619 | 0.479227808 | 2.197626266 | 0.02797575 | 0.19118966 |
| FAXC | 288.8164095 | 0.415173212 | 0.331830906 | 1.251158962 | 0.210876487 | 0.518294063 |
| FAXDC2 | 500.619079 | -0.679717376 | 0.316391246 | -2.148344447 | 0.031686402 | 0.202662062 |
| FBF1 | 331.3963122 | -0.110813453 | 0.437096556 | -0.25352168 | 0.79986511 | 0.916311991 |
| FBH1 | 1304.322544 | -0.392688507 | 0.401671715 | -0.97763545 | 0.328254658 | 0.637801154 |
| FBL | 4316.806096 | 0.513853597 | 0.325882148 | 1.576808055 | 0.114839698 | 0.389985713 |
| FBLIM1 | 1032.258361 | -0.563391578 | 0.394341921 | -1.428688018 | 0.153093923 | 0.449450244 |
| FBLN1 | 5671.826041 | 0.194522162 | 0.264483569 | 0.735479193 | 0.462047707 | 0.739734673 |
| FBLN2 | 676.3986307 | 0.698557825 | 0.432716256 | 1.614355403 | 0.106450359 | 0.374388239 |
| FBLN7 | 107.0782554 | 1.38918398 | 0.418882579 | 3.316404282 | 0.000911838 | 0.025808763 |
| FBN1 | 408.6695775 | 0.922463339 | 0.710020735 | 1.299206197 | 0.193873175 | 0.499845061 |
| FBN2 | 9094.368658 | -1.223669101 | 0.393084576 | -3.112991898 | 0.001852011 | 0.040024016 |
| FBN3 | 1919.941647 | -1.669173537 | 0.44158297 | -3.779977156 | 0.000156843 | 0.008176843 |
| FBP1 | 26.80208859 | -2.228367444 | 0.928587261 | -2.399739408 | 0.016406747 | 0.144574776 |
| FBRS | 894.0149168 | -0.661198715 | 0.409586205 | -1.614309047 | 0.106460409 | 0.374388239 |
| FBRSL1 | 508.6783732 | -0.287930049 | 0.3925256 | -0.7335319 | 0.463234079 | 0.740745005 |
| FBXL12 | 572.6426517 | -0.052466631 | 0.322534768 | -0.162669689 | 0.870778513 | 0.94835469 |
| FBXL13 | 28.33077742 | 0.094109412 | 0.804072547 | 0.117040946 | 0.906827606 | 0.965234099 |
| FBXL14 | 212.2424845 | 0.757348442 | 0.914063933 | 0.828550843 | 0.407358614 | 0.701878198 |
| FBXL15 | 68.93001622 | 0.574867474 | 0.59673286 | 0.963358167 | 0.335367812 | 0.643789078 |
| FBXL16 | 26.08496904 | 0.242714885 | 0.833689351 | 0.291133484 | 0.770949232 | 0.905067213 |
| FBXL17 | 157.7507682 | -0.78621277 | 0.361164932 | -2.176880148 | 0.029489511 | 0.196221654 |
| FBXL18 | 390.8184466 | -0.049066871 | 0.335678874 | -0.146172057 | 0.883785564 | 0.953960213 |
| FBXL19 | 1450.259958 | -0.031757574 | 0.376705052 | -0.084303553 | 0.932815088 | 0.974570463 |
| FBXL2 | 247.6110977 | 0.115620993 | 0.294210155 | 0.392987769 | 0.694328515 | 0.86892666 |
| FBXL20 | 507.1907025 | -0.534774879 | 0.2907619 | -1.839219234 | 0.065882947 | 0.296326767 |
| FBXL21P | 587.4609041 | -3.722853194 | 0.742557846 | -5.013553107 | 5.34E-07 | 0.000126664 |
| FBXL3 | 599.6690126 | -0.083140257 | 0.284148415 | -0.292594477 | 0.769832135 | 0.904694332 |
| FBXL4 | 484.4576749 | -0.067390617 | 0.323112938 | -0.208566754 | 0.834786466 | 0.932107564 |
| FBXL5 | 1590.557659 | -0.035072634 | 0.278527232 | -0.12592174 | 0.899793873 | 0.961118979 |
| FBXL6 | 175.6873518 | 0.236398287 | 0.356134381 | 0.66378957 | 0.506825002 | 0.767675929 |
| FBXL7 | 469.5056494 | -0.318862319 | 0.472969275 | -0.674171316 | 0.500202404 | 0.763360896 |
| FBXO10 | 1146.462921 | -0.189715766 | 0.347118441 | -0.546544762 | 0.584691528 | 0.814990368 |

| | | | | | | |
|---|---|---|---|---|---|---|
| FBXO11 | 1955.630995 | -0.150898729 | 0.249854582 | -0.603946214 | 0.545879403 | 0.791658199 |
| FBXO15 | 70.09548577 | -0.084969002 | 0.443611674 | -0.191539148 | 0.848103213 | 0.938207222 |
| FBXO16 | 172.8837239 | 0.116272696 | 0.345421215 | 0.336611333 | 0.736409912 | 0.890400696 |
| FBXO17 | 279.5160931 | 0.517728807 | 0.382310573 | 1.354210016 | 0.175669385 | 0.479848799 |
| FBXO2 | 943.9144572 | 1.044934587 | 0.668104634 | 1.564028349 | 0.117810911 | 0.395839103 |
| FBXO21 | 1694.431453 | 0.098486097 | 0.282213666 | 0.348977063 | 0.727106529 | 0.8850111 |
| FBXO22 | 1663.314786 | 0.414973878 | 0.245396874 | 1.691031639 | 0.09083076 | 0.345388595 |
| FBXO24 | 33.75445991 | 0.065199397 | 0.606609582 | 0.107481647 | 0.914406884 | 0.968409553 |
| FBXO25 | 158.4252273 | -0.81268833 | 0.371013054 | -2.190457509 | 0.028491073 | 0.192869196 |
| FBXO27 | 62.1246241 | -0.538906288 | 0.444814231 | -1.211531131 | 0.225691911 | 0.53459368 |
| FBXO28 | 1215.958321 | -0.115046805 | 0.242217057 | -0.474974003 | 0.634805503 | 0.840313679 |
| FBXO3 | 890.6102831 | 0.246330849 | 0.237279316 | 1.038147165 | 0.299201546 | 0.610246278 |
| FBXO30 | 1273.19377 | -0.102848879 | 0.323963211 | -0.31747086 | 0.750886347 | 0.896207755 |
| FBXO31 | 344.6682575 | 0.192273033 | 0.427367636 | 0.449900781 | 0.652781985 | 0.848628373 |
| FBXO32 | 30.0057474 | -1.314825285 | 0.653170788 | -2.012988501 | 0.044115839 | 0.241443505 |
| FBXO33 | 530.9227705 | 0.249687775 | 0.272358677 | 0.91676086 | 0.359267966 | 0.666966999 |
| FBXO34 | 506.9871359 | -0.173382539 | 0.26923496 | -0.643982262 | 0.519586937 | 0.774983751 |
| FBXO36 | 153.7379948 | 0.1868125 | 0.399695107 | 0.467387508 | 0.640222658 | 0.843615231 |
| FBXO38 | 1107.026399 | -0.517078852 | 0.321722519 | -1.607219954 | 0.108006126 | 0.376853943 |
| FBXO4 | 55.05361704 | 0.47616161 | 0.466294618 | 1.021160424 | 0.307178439 | 0.617205881 |
| FBXO41 | 344.2824436 | -1.327519131 | 0.270377707 | -4.909869036 | 9.11E-07 | 0.000174186 |
| FBXO42 | 611.061314 | -0.15342755 | 0.290170451 | -0.528749736 | 0.59697907 | 0.821812117 |
| FBXO43 | 24.38550303 | 2.187493385 | 0.930824436 | 2.35006012 | 0.018770379 | 0.1554129 |
| FBXO44 | 724.7881335 | 0.148764753 | 0.353371615 | 0.420986708 | 0.673764788 | 0.85954302 |
| FBXO45 | 662.6394566 | 0.484771802 | 0.284842048 | 1.701896912 | 0.088774695 | 0.342301733 |
| FBXO46 | 492.1896419 | -0.017007044 | 0.441581763 | -0.038513919 | 0.969277934 | 0.989496332 |
| FBXO48 | 107.0781697 | 0.290629624 | 0.376877127 | 0.771152196 | 0.440616725 | 0.7241465 |
| FBXO5 | 865.6532712 | 0.543444594 | 0.249710371 | 2.176299654 | 0.029532861 | 0.196251879 |
| FBXO6 | 27.94931774 | 0.032488611 | 0.581877513 | 0.055834106 | 0.955473964 | 0.984323736 |
| FBXO7 | 2183.968166 | -0.147471556 | 0.301853523 | -0.488553371 | 0.625157932 | 0.835033444 |
| FBXO8 | 275.3967352 | 0.393020687 | 0.337208151 | 1.165513603 | 0.243811157 | 0.555134184 |
| FBXO9 | 948.8977954 | -0.522683648 | 0.287910488 | -1.81543803 | 0.069456621 | 0.30336211 |
| FBXW11 | 1658.419562 | 0.091927461 | 0.298962893 | 0.307487862 | 0.758472061 | 0.899440455 |
| FBXW12 | 16.86565878 | -0.674614353 | 0.827059252 | -0.815678382 | 0.414684106 | 0.706952756 |
| FBXW2 | 1650.110172 | -0.303194217 | 0.259295359 | -1.169300593 | 0.242282544 | 0.553231259 |
| FBXW4 | 822.856557 | -0.096342234 | 0.27215266 | -0.354000707 | 0.723338353 | 0.883273431 |
| FBXW4P1 | 24.76349226 | 0.552659842 | 0.650467543 | 0.84963477 | 0.395528175 | |

0.69426198
FBXW5 730.07824 -0.264396254 0.525631563 -0.503006808 0.614959486
0.830872437
FBXW7 727.6721764 -0.316737642 0.255445371 -1.239942775 0.214996561
0.522208266
FBXW8 836.6228963 -0.378911523 0.329083926 -1.151413038 0.249562353
0.562413758
FBXW9 325.7655144 0.425666214 0.478694484 0.889223145 0.373883167 0.678520659
FCER1G 10.39587339 0.524785571 1.007567365 0.520844154 0.602475342
0.825021776
FCF1 1961.56005 0.037905915 0.223601185 0.169524663 0.865383975 0.945250148
FCGBP 18.04508382 0.811881038 0.858396286 0.945811452 0.344244763 0.65118919
FCGRT 370.1819279 0.000915046 0.424658102 0.002154782 0.998280734 0.998939187
FCHO1 199.7953889 0.874214074 0.569553093 1.534912346 0.124805361 0.406452918
FCHO2 603.6264689 -0.844931386 0.308098385 -2.742407704 0.006099059
0.084270328
FCHSD1 606.4268861 -0.856855435 0.3176931 -2.697116921
0.006994271 0.091949811
FCHSD2 407.9443918 0.496581734 0.472893432 1.050092261 0.293675696
0.606359511
FCRL5 17.13754189 -0.338046429 0.942907195 -0.35851506 0.7199579
0.881769702
FDFT1 3398.391862 1.049030757 0.33396534 3.14113661 0.001682935 0.038017939
FDPS 4139.418957 0.41748506 0.223505966 1.86789224 0.061777093 0.286348999
FDX1 571.230183 0.656699322 0.486444286 1.349999048 0.177016288 0.481361196
FDX2 225.0427333 0.56854554 0.403455622 1.409189781 0.158779059 0.45865139
FDXACB1 97.95640058 0.815042305 0.362408827 2.248958206 0.024515155
0.178807413
FDXR 1326.576749 0.781655837 0.311159055 2.512078062 0.012002254 0.124375812
FECH 843.1668333 -0.050859714 0.307414594 -0.165443396 0.868595
0.947046732
FEM1A 849.5353311 -0.347760018 0.327512275 -1.061822852 0.288316113
0.601673803
FEM1B 2151.576212 0.115962864 0.254904271 0.454927115 0.649161675 0.847198189
FEM1C 614.0320054 -0.16137296 0.32633117 -0.49450673 0.620948354 0.833349572
FEN1 1720.172148 0.674814453 0.301357884 2.239246059 0.02513991 0.181445091
FER 765.2522561 -0.705203529 0.402601225 -1.751617942 0.079839525
0.323453279
FER1L4 193.5609636 0.04066464 0.54887011 0.074087911 0.940940435
0.978142205
FER1L5 41.00629243 0.063441797 0.618386762 0.102592424 0.918286456
0.969677217
FERMT1 330.8657746 -1.546549394 0.405693688 -3.812111059
0.000137785 0.007519192
FERMT2 2693.056316 0.010033415 0.272043872 0.036881606 0.970579406
0.98957539
FERMT3 19.44246968 -0.127798614 0.709425409 -0.180143835
0.85703965 0.942343063
FES 139.2656307 -1.297261892 0.498460186 -2.602538632 0.009253639
0.108827097
FEZ1 1327.306512 0.30433818 0.636303472 0.478290931 0.632443151 0.838768903
FEZ2 1304.063339 -0.366468142 0.375340774 -0.976361129 0.328885537
0.638209578
FEZF1 282.0939057 1.607824322 0.654937285 2.454928676 0.014091254 0.134875975
FFAR4 11.50142135 -0.46691757 1.107743653 -0.421503269 0.673387625
0.859423207
FGD1 855.1528618 -0.510208416 0.343423047 -1.485655724 0.137370208
0.425712285
FGD3 108.9285077 -1.950426777 0.69792912 -2.794591487 0.005196533
0.076636622
FGD4 424.1336595 -0.03042053 0.422721586 -0.071963512 0.942630946
0.978574897
FGD5 64.4707019 -0.564091048 0.524942825 -1.074576165 0.282564528
0.596463956

| | | | | | | |
|---|---|---|---|---|---|---|
| FGD6 | 805.2358669 | -0.426757157 | 0.426561632 | -1.000458374 | 0.317088733 | 0.627198876 |
| FGF11 | 199.5954544 | 0.224848585 | 0.402124824 | 0.559151218 | 0.576058522 | 0.808993188 |
| FGF12 | 158.7632883 | -0.322772977 | 0.462106281 | -0.698482125 | 0.484875733 | 0.755350561 |
| FGF13 | 579.3471541 | -0.144189345 | 0.244466845 | -0.589811452 | 0.555317064 | 0.796360966 |
| FGF14 | 17.36535577 | 1.727793521 | 0.775219006 | 2.228781169 | 0.025828468 | 0.183472841 |
| FGF18 | 17.53839548 | 1.298775661 | 0.945118951 | 1.374192804 | 0.169381834 | 0.472052919 |
| FGF19 | 117.203519 | 0.281165836 | 0.590449567 | 0.476189418 | 0.633939438 | 0.839809222 |
| FGF2 | 1242.602975 | 0.332110251 | 0.28462074 | 1.166851901 | 0.243270183 | 0.554511158 |
| FGF23 | 77.25820423 | 1.764004992 | 0.505420778 | 3.490171098 | 0.000482711 | 0.017231093 |
| FGF3 | 57.4518718 | 1.978780227 | 0.862355907 | 2.294621295 | 0.021754837 | 0.16744933 |
| FGF4 | 18.62443528 | 0.024366718 | 0.824154677 | 0.029565709 | 0.976413413 | 0.99184251 |
| FGF7P3 | 48.93448028 | 1.235299095 | 0.497048929 | 2.485266585 | 0.012945445 | 0.128476822 |
| FGF7P6 | 67.1712216 | 0.261101702 | 0.468376913 | 0.557460659 | 0.577212731 | 0.809901794 |
| FGF9 | 25.05621378 | 1.992820014 | 0.876765859 | 2.272921549 | 0.023030908 | 0.173530066 |
| FGFBP1 | 29.63524506 | 0.009961354 | 0.722593639 | 0.013785554 | 0.989001067 | 0.996025969 |
| FGFBP3 | 2184.126503 | 2.347617897 | 0.594316569 | 3.950113492 | 7.81E-05 | 0.00513443 |
| FGFR1 | 15208.95574 | 0.157847972 | 0.373297879 | 0.422847225 | 0.67240673 | 0.859011927 |
| FGFR1OP | 978.2474353 | 0.272017464 | 0.259069153 | 1.049980136 | 0.293727246 | 0.606359511 |
| FGFR1OP2 | 1408.349353 | 0.336950556 | 0.270732184 | 1.244589951 | 0.213282516 | 0.520461485 |
| FGFR2 | 2269.537462 | -1.016474704 | 0.435041601 | -2.336500006 | 0.019465197 | 0.158211103 |
| FGFR3 | 2649.548335 | 0.452448845 | 0.364509373 | 1.241254353 | 0.214511802 | 0.521483923 |
| FGFR4 | 681.8297565 | 0.487726107 | 0.307354979 | 1.586849542 | 0.112546725 | 0.385776405 |
| FGFRL1 | 73.91070687 | 1.609828982 | 0.62332286 | 2.582656734 | 0.009804282 | 0.112404636 |
| FGGY | 323.4304802 | 0.227877987 | 0.307280494 | 0.741596006 | 0.458332144 | 0.73670627 |
| FH | 2051.099248 | 0.364760057 | 0.24185753 | 1.50816085 | 0.131513365 | 0.41719445 |
| FHAD1 | 43.73888619 | 0.623266079 | 0.803797318 | 0.775402038 | 0.438102131 | 0.721949744 |
| FHDC1 | 254.2831743 | -1.067931386 | 0.429225778 | -2.488041121 | 0.012844888 | 0.128086828 |
| FHIT | 93.20817788 | 0.025028304 | 0.426809089 | 0.058640513 | 0.953238441 | 0.983112195 |
| FHL1 | 1307.869123 | 1.23222081 | 0.526328431 | 2.341163306 | 0.019223756 | 0.157389967 |
| FHL2 | 461.0076847 | 0.445132093 | 0.505434802 | 0.880691418 | 0.378484863 | 0.681253679 |
| FHL3 | 506.6112615 | 0.088534086 | 0.298388603 | 0.296707331 | 0.76668996 | 0.903470997 |
| FHOD1 | 426.8512879 | 0.223752911 | 0.25986281 | 0.861042449 | 0.389214664 | 0.689097897 |
| FHOD3 | 741.4834108 | 1.266360299 | 0.758348229 | 1.669892868 | 0.094940562 | 0.353777379 |
| FIBCD1 | 51.13258412 | 0.254947124 | 0.532952693 | 0.478367269 | 0.632388826 | 0.838768903 |
| FIBIN | 15.61457328 | -0.086604739 | 0.747056591 | -0.11592795 | 0.907709644 | 0.965427861 |
| FIBP | 1375.520757 | 0.239879631 | 0.275049261 | 0.872133343 | 0.383135638 | 0.68439312 |
| FICD | 121.307732 | -0.129340058 | 0.368593619 | -0.350901512 | 0.725662237 | 0.884258779 |
| FIG4 | 1365.31918 | -0.077744339 | 0.27673242 | -0.280936867 | 0.778758823 | 0.908252622 |
| FIGN | 1532.737478 | -0.782027278 | 0.324927596 | -2.406773963 | 0.016094128 | 0.143654707 |
| FIGNL1 | 473.3278793 | -0.105774543 | 0.392959215 | -0.269174354 | 0.787795514 | 0.912200103 |
| FIGNL2 | 2691.163821 | 0.050622795 | 0.352155637 | 0.143751198 | 0.885696941 | 0.954813919 |
| FILIP1 | 588.0443397 | 1.165882785 | 0.545926732 | 2.135603034 | 0.032711784 | 0.206178012 |
| FILIP1L | 15.12129818 | -2.054658474 | 1.078399043 | -1.905285884 | 0.056742919 | 0.27582404 |

| | | | | | | |
|---|---|---|---|---|---|---|
| FIP1L1 | 1579.942708 | -0.079618847 | 0.252265657 | -0.315615088 | 0.752294684 | 0.896586461 |
| FIS1 | 1475.59529 | 0.716214702 | 0.416797925 | 1.718373965 | 0.085728426 | 0.335745178 |
| FITM2 | 117.4320578 | 0.604398695 | 0.406899016 | 1.485377629 | 0.137443818 | 0.425712285 |
| FIZ1 | 395.1119984 | 0.31798286 | 0.408750189 | 0.777939358 | 0.436604764 | 0.720784523 |
| FJX1 | 253.930669 | 1.17634415 | 0.454226593 | 2.589773841 | 0.0096039 | 0.111098245 |
| FKBP10 | 5886.143065 | 0.619822104 | 0.316506125 | 1.958325776 | 0.050191798 | 0.257598027 |
| FKBP11 | 384.0744642 | 0.363559945 | 0.408453076 | 0.890089869 | 0.373417633 | 0.678294888 |
| FKBP14 | 338.5155647 | 0.168401368 | 0.303140987 | 0.555521608 | 0.578537935 | 0.810209453 |
| FKBP15 | 719.9928842 | -0.770177424 | 0.356982333 | -2.1574665 | 0.030969335 | 0.20078452 |
| FKBP1A | 5113.119773 | 0.271547047 | 0.330532074 | 0.821545224 | 0.411335775 | 0.704686727 |
| FKBP1B | 796.1879231 | 0.517801885 | 0.406164899 | 1.274856312 | 0.202360105 | 0.509139909 |
| FKBP2 | 593.4854123 | 0.872642744 | 0.554994149 | 1.572345845 | 0.115870366 | 0.392293981 |
| FKBP3 | 5025.366207 | 1.007105488 | 0.366253874 | 2.749746994 | 0.00596413 | 0.083399911 |
| FKBP4 | 3825.952162 | -0.094778532 | 0.278974176 | -0.339739444 | 0.734052754 | 0.88891398 |
| FKBP5 | 1010.703828 | 1.019516573 | 0.488195831 | 2.088335272 | 0.036767602 | 0.219399876 |
| FKBP7 | 261.803061 | 1.333272365 | 0.35699185 | 3.734741739 | 0.000187908 | 0.009050003 |
| FKBP8 | 5776.855709 | 0.291908614 | 0.491507507 | 0.593904691 | 0.552575862 | 0.795662343 |
| FKBP9 | 824.2921006 | -0.345333092 | 0.29988507 | -1.151551466 | 0.249505435 | 0.562413758 |
| FKBPL | 114.8308918 | 0.808583031 | 0.47213052 | 1.712626056 | 0.086781357 | 0.33809963 |
| FKRP | 464.2874242 | -0.247865492 | 0.385539178 | -0.642906107 | 0.520285027 | 0.775367794 |
| FKTN | 564.4982259 | -0.379326257 | 0.388122891 | -0.977335442 | 0.328403112 | 0.637925754 |
| FLAD1 | 1384.664343 | 0.200267433 | 0.251932863 | 0.794923815 | 0.42665782 | 0.714361085 |
| FLCN | 870.9267672 | -0.30044574 | 0.353064492 | -0.850965609 | 0.394788457 | 0.693708137 |
| FLII | 1444.845113 | -0.632049182 | 0.418604381 | -1.509896245 | 0.131069901 | 0.416813082 |
| FLNA | 16699.66489 | -1.044452042 | 0.560029365 | -1.864995136 | 0.062182086 | 0.287461403 |
| FLNB | 6310.956947 | -1.207706547 | 0.294844804 | -4.096075389 | 4.20E-05 | 0.003252578 |
| FLNC | 1257.63714 | -0.094738005 | 0.557958143 | -0.169794108 | 0.865172061 | 0.945248223 |
| FLOT1 | 580.8667944 | -0.659336874 | 0.787796648 | -0.836937902 | 0.402627474 | 0.699577622 |
| FLOT2 | 3865.25328 | -0.137156045 | 0.242506069 | -0.565577783 | 0.571680834 | 0.807313593 |
| FLRT2 | 129.7870741 | 0.681355998 | 0.521888631 | 1.305558232 | 0.191702832 | 0.497319369 |
| FLRT3 | 2118.469879 | -2.890925051 | 0.657694586 | -4.395543331 | 1.10E-05 | 0.001172262 |
| FLT1 | 549.4892764 | -1.971287367 | 0.425242345 | -4.635679841 | 3.56E-06 | 0.000504423 |
| FLT3 | 12.82005742 | 0.469109982 | 1.169128314 | 0.401247645 | 0.688237806 | 0.866925917 |
| FLT3LG | 18.19546025 | 0.623201825 | 0.762231321 | 0.817601964 | 0.413584505 | 0.705778726 |
| FLT4 | 117.9618684 | 0.443876999 | 0.504260447 | 0.880253451 | 0.378722023 | 0.681253679 |
| FLVCR1 | 848.5533836 | 0.171040459 | 0.246167443 | 0.694813485 | 0.487172196 | 0.756439734 |
| FLVCR2 | 229.0037654 | -0.27355122 | 0.28743408 | -0.951700718 | 0.341248785 | 0.648152255 |
| FLYWCH1 | 471.7123939 | -0.113965074 | 0.559698228 | -0.203618787 | 0.838651411 | 0.933879959 |
| FLYWCH2 | 258.1329604 | 0.973017736 | 0.480864582 | 2.023475572 | 0.04302414 | 0.238392703 |

| | | | | | | |
|---|---|---|---|---|---|---|
| FMC1 | 223.2787451 | 1.497293145 | 0.520064621 | 2.879052111 | 0.003988724 | 0.065559297 |
| FMC1-LUC7L2 | 508.061907 | -1.120830526 | 2.402261634 | -0.466573045 | 0.640805376 | 0.843724522 |
| FMN1 | 93.60747449 | -1.148958346 | 0.511506551 | -2.246224108 | 0.024689657 | 0.179648337 |
| FMN2 | 15.75375006 | -1.401477623 | 0.913474157 | -1.534227993 | 0.124973573 | 0.4067741 |
| FMNL1 | 53.08824954 | -0.353475755 | 0.550106146 | -0.642559182 | 0.520510176 | 0.775597169 |
| FMNL2 | 2553.395396 | -0.250175216 | 0.35899276 | -0.696880951 | 0.4858773 | 0.75593662 |
| FMNL3 | 435.8200099 | -0.094675071 | 0.586694842 | -0.161370212 | 0.871801826 | 0.948856124 |
| FMO4 | 110.3301342 | -0.367523307 | 0.394279582 | -0.932138826 | 0.351264785 | 0.658556359 |
| FMO5 | 18.08079607 | -0.826787138 | 0.714645075 | -1.156919942 | 0.24730507 | 0.559988249 |
| FMR1 | 4539.345102 | -1.301710405 | 0.376902808 | -3.453703121 | 0.000552946 | 0.018600309 |
| FN1 | 2927.159627 | -1.449609029 | 0.625580594 | -2.317221862 | 0.020491649 | 0.16183176 |
| FN3K | 332.1996393 | -0.110473828 | 0.335564113 | -0.329218245 | 0.741990734 | 0.892150449 |
| FN3KRP | 856.6647956 | -0.028484924 | 0.249462191 | -0.114185337 | 0.909090875 | 0.966016507 |
| FNBP1 | 1129.611899 | -0.299485351 | 0.227818769 | -1.314577163 | 0.188652049 | 0.493962743 |
| FNBP1L | 5513.14926 | -0.867295708 | 0.281656068 | -3.079272225 | 0.00207507 | 0.043065504 |
| FNBP1P1 | 38.04681527 | -0.806561677 | 0.60941647 | -1.323498324 | 0.185669713 | 0.490218451 |
| FNBP4 | 1222.701911 | -0.042508897 | 0.336199854 | -0.126439369 | 0.89938414 | 0.960836323 |
| FNDC1 | 12.3634963 | -1.337462145 | 1.019744646 | -1.311565743 | 0.189666698 | 0.495097939 |
| FNDC10 | 56.62295829 | 0.870277945 | 0.456824641 | 1.905059114 | 0.056772387 | 0.275878888 |
| FNDC11 | 40.25131063 | 0.865547941 | 0.5847871 | 1.480107787 | 0.138844484 | 0.427350307 |
| FNDC3A | 1490.841282 | -0.183819142 | 0.321682911 | -0.571429615 | 0.567708459 | 0.805151447 |
| FNDC3B | 844.0761048 | -0.142022591 | 0.288365078 | -0.492509677 | 0.622359083 | 0.833640266 |
| FNDC4 | 214.2302069 | -0.060474632 | 0.335527525 | -0.180237469 | 0.856966144 | 0.942343063 |
| FNDC5 | 263.7761474 | -0.054694263 | 0.41574688 | -0.131556641 | 0.895334983 | 0.958651071 |
| FNIP1 | 1613.12918 | -0.038026973 | 0.340131319 | -0.111800856 | 0.910981309 | 0.966866396 |
| FNIP2 | 606.8689939 | -1.105924526 | 0.283163091 | -3.905609731 | 9.40E-05 | 0.005843825 |
| FNTA | 2305.691182 | 0.007347855 | 0.225575808 | 0.03257377 | 0.974014487 | 0.99099127 |
| FNTB | 632.7638406 | -0.472352626 | 0.29858866 | -1.581950991 | 0.113660759 | 0.388022186 |
| FO393400.1 | 12.73884727 | -0.194271356 | 0.970245972 | -0.200228975 | 0.841301507 | 0.935153081 |
| FOCAD | 2122.709162 | -0.45617337 | 0.360185235 | -1.266496584 | 0.205335352 | 0.511820585 |
| FOLH1B | 16.66966032 | -3.99166463 | 1.644935796 | -2.426638559 | 0.01523943 | 0.139780768 |
| FOLR1 | 989.5263206 | -0.969851408 | 0.534984844 | -1.812857727 | 0.069853768 | 0.303603614 |
| FOPNL | 752.4487774 | 0.685695935 | 0.316474777 | 2.166668511 | 0.030260139 | 0.198390913 |
| FOS | 5157.576986 | -1.978872076 | 0.705100777 | -2.806509568 | 0.005008143 | |

| | | | | | | |
|---|---|---|---|---|---|---|
| | | | | | | 0.07561376 |
| FOSB | 652.381408 | -2.884759654 | 1.418045343 | -2.034321164 | 0.041919225 | 0.235801467 |
| FOSL1 | 76.44353936 | -0.28968177 | 0.638887068 | -0.453416236 | 0.650249047 | 0.847430646 |
| FOSL2 | 202.3507087 | -1.265937349 | 0.487245365 | -2.59815165 | 0.009372709 | 0.109379514 |
| FOXA3 | 173.7562894 | 0.544351502 | 0.385097439 | 1.413542254 | 0.157496347 | 0.456731315 |
| FOXB1 | 370.9366925 | 2.53359034 | 0.628082769 | 4.033847867 | 5.49E-05 | 0.004040998 |
| FOXC1 | 279.7983351 | -0.503523978 | 0.729935318 | -0.689819996 | 0.490307392 | 0.758593654 |
| FOXD1 | 13.92052009 | 3.502540092 | 1.193080378 | 2.935711758 | 0.003327834 | 0.058298577 |
| FOXD3 | 11.05793083 | 1.102381429 | 1.138690904 | 0.968112966 | 0.332987955 | 0.640761068 |
| FOXG1 | 19.57069875 | 2.203773997 | 1.636212448 | 1.346875217 | 0.178020425 | 0.48228793 |
| FOXH1 | 395.1689861 | 0.167316022 | 0.405125772 | 0.412997726 | 0.67960828 | 0.861682789 |
| FOXI2 | 17.74427482 | -1.80515984 | 0.919765764 | -1.962629956 | 0.049689187 | 0.256930696 |
| FOXI3 | 1014.018757 | -1.168711469 | 0.344156305 | -3.395874062 | 0.000684098 | 0.021514856 |
| FOXJ1 | 32.48699309 | 1.305687377 | 0.643287505 | 2.029710458 | 0.042385979 | 0.236937315 |
| FOXJ2 | 498.8627793 | -0.588653197 | 0.321568912 | -1.830566247 | 0.06716531 | 0.297817842 |
| FOXJ3 | 1503.382328 | -0.354999479 | 0.271452704 | -1.307776544 | 0.190949112 | 0.496058205 |
| FOXK1 | 460.6696267 | -0.651841495 | 0.345609626 | -1.886062904 | 0.059286479 | 0.282340574 |
| FOXK2 | 1593.64593 | -0.032001587 | 0.304456835 | -0.105110423 | 0.916288189 | 0.969184139 |
| FOXL2NB | 11.25356604 | -0.428411696 | 0.920607154 | -0.465357774 | 0.641675268 | 0.844038972 |
| FOXM1 | 2102.034399 | 0.071036672 | 0.354905094 | 0.20015681 | 0.841357944 | 0.935153081 |
| FOXN2 | 393.6410482 | 0.610620627 | 0.424309677 | 1.439091918 | 0.150124481 | 0.444740017 |
| FOXN3 | 1882.328195 | -0.869744989 | 0.268645107 | -3.23752403 | 0.001205718 | 0.030537472 |
| FOXN4 | 32.82077248 | -0.018431716 | 0.532770103 | -0.034596003 | 0.972401889 | 0.989954357 |
| FOXO1 | 384.0320861 | -1.199968726 | 0.514185441 | -2.333727545 | 0.019609993 | 0.158583796 |
| FOXO3 | 1839.623353 | -0.501951728 | 0.336880167 | -1.490000829 | 0.136224018 | 0.424171131 |
| FOXO3B | 101.4388944 | -0.375547796 | 0.942007637 | -0.398667464 | 0.690138244 | 0.867878589 |
| FOXO4 | 842.8190447 | -1.458558508 | 0.415645258 | -3.509142669 | 0.000449554 | 0.016498831 |
| FOXO6 | 79.63749274 | 0.208395314 | 0.584415999 | 0.356587284 | 0.721400798 | 0.882498649 |
| FOXP1 | 872.0330021 | -0.607793266 | 0.336950878 | -1.803803777 | 0.071262074 | 0.306992555 |
| FOXP2 | 21.4341806 | -1.729324793 | 1.052145272 | -1.643617891 | 0.100255156 | 0.36243601 |
| FOXP3 | 11.01479134 | 0.045724442 | 1.33547261 | 0.034238398 | 0.972687048 | 0.990112399 |
| FOXP4 | 626.2284609 | -1.312786067 | 0.453581736 | -2.894265716 | 0.003800464 | 0.063596264 |
| FOXRED1 | 1184.991812 | -0.105275795 | 0.295029617 | -0.356831278 | 0.721218119 | 0.882387104 |
| FOXRED2 | 818.8639919 | -0.503255863 | 0.308723227 | -1.630119857 | 0.103076168 | 0.368123388 |
| FP236241.2 | 10.49504699 | -0.962148663 | 0.983974079 | -0.977819115 | 0.328163795 | 0.637787615 |
| FP565260.1 | 111.3123209 | -0.529296598 | 0.661494475 | -0.800152712 | 0.423622324 | 0.712818797 |
| FP565260.2 | 14.21810668 | -0.414570537 | 1.321784405 | -0.313644597 | 0.753790984 | 0.897063305 |
| FP565260.3 | 32.82597936 | -1.944247459 | 0.619511878 | -3.138353802 | 0.001698997 | 0.038242549 |

| | | | | | | |
|---|---|---|---|---|---|---|
| FP565260.6 | 307.5383933 | 1.000040393 | 0.587321162 | 1.702714727 | 0.088621467 | 0.342280113 |
| FPGS | 1087.879492 | 0.277788842 | 0.29028389 | 0.956955764 | 0.338589583 | 0.645678097 |
| FPGT | 490.3624084 | -0.45671932 | 0.334195813 | -1.36662191 | 0.171743846 | 0.474879827 |
| FPGT-TNNI3K | 70.20250834 | -0.127030166 | 0.531302197 | -0.239092115 | 0.811034156 | 0.920897407 |
| FRA10AC1 | 924.3346638 | 0.175888006 | 0.34094238 | 0.515887775 | 0.605932793 | 0.826524582 |
| FRAS1 | 5537.061746 | -1.197439859 | 0.353162396 | -3.390621068 | 0.000697344 | 0.021675701 |
| FRAT1 | 163.3062324 | 0.196949958 | 0.559713861 | 0.351876148 | 0.724931147 | 0.883935898 |
| FRAT2 | 2175.715362 | 0.448254686 | 0.681967055 | 0.657296686 | 0.510990167 | 0.770599585 |
| FREM1 | 1385.71818 | -1.316198398 | 0.268297131 | -4.90574907 | 9.31E-07 | 0.000174319 |
| FREM2 | 8574.853635 | -2.186578137 | 0.495165999 | -4.415848708 | 1.01E-05 | 0.001114176 |
| FRG1 | 289.2408708 | 0.577697327 | 0.380843794 | 1.516887858 | 0.129295 | 0.413476905 |
| FRG1BP | 114.2802623 | -0.067324456 | 0.357535389 | -0.188301518 | 0.850640295 | 0.939164829 |
| FRG1CP | 423.2154264 | 0.646116132 | 0.406444226 | 1.589679694 | 0.111907023 | 0.384888109 |
| FRG1HP | 161.080437 | 0.053410899 | 0.362909237 | 0.147174262 | 0.882994475 | 0.953581235 |
| FRK | 70.40178666 | -0.466433937 | 0.506064864 | -0.921688049 | 0.356691316 | 0.664215534 |
| FRMD1 | 10.28813256 | 0.209460319 | 0.957977268 | 0.218648528 | 0.826923847 | 0.928082319 |
| FRMD3 | 200.0266664 | 1.502863923 | 0.736550251 | 2.040409218 | 0.041309585 | 0.233497655 |
| FRMD4A | 533.9509453 | 0.287338125 | 0.346030015 | 0.830384975 | 0.406321158 | 0.701474651 |
| FRMD4B | 231.9873234 | -0.834790997 | 0.397325359 | -2.101026217 | 0.035638665 | 0.216083122 |
| FRMD5 | 247.3417307 | 0.041292495 | 0.441864418 | 0.093450601 | 0.925545593 | 0.972124891 |
| FRMD6 | 1016.836832 | -0.364484649 | 0.295950624 | -1.231572499 | 0.218108819 | 0.526480334 |
| FRMD8 | 510.0079535 | -0.171032877 | 0.322937875 | -0.529615416 | 0.596378604 | 0.821812117 |
| FRMPD2 | 35.67441026 | 0.197628061 | 0.747962452 | 0.264221901 | 0.791608936 | 0.913478296 |
| FRMPD3 | 136.9572505 | -1.213371573 | 0.473541535 | -2.562333994 | 0.010397128 | 0.115896273 |
| FRRS1 | 201.1968857 | -1.469087117 | 0.519899081 | -2.82571593 | 0.004717508 | 0.072780519 |
| FRRS1L | 43.98299562 | -0.008309479 | 0.58745619 | -0.014144848 | 0.98871442 | 0.995803391 |
| FRS2 | 1213.709464 | -0.814500367 | 0.358114765 | -2.274411577 | 0.022941252 | 0.173068988 |
| FRS3 | 158.0791954 | 0.170250589 | 0.413140341 | 0.412088998 | 0.680274192 | 0.862002279 |
| FRY | 498.9331976 | 0.537989476 | 0.700170961 | 0.768368735 | 0.442268149 | 0.725681386 |
| FRYL | 1232.126398 | -0.314116023 | 0.372664649 | -0.842891923 | 0.399288891 | 0.697434725 |
| FRZB | 311.3890482 | 1.725031961 | 0.527783948 | 3.268443398 | 0.001081408 | 0.028433348 |
| FSBP | 762.7344389 | 0.402039673 | 0.326308498 | 1.232084591 | 0.217917487 | 0.526185929 |
| FSCN1 | 3758.475476 | 0.396531448 | 0.357489697 | 1.109210845 | 0.267339235 | 0.580644744 |
| FSD1 | 856.9818391 | 0.331625644 | 0.375162611 | 0.883951742 | 0.376722264 | 0.68029735 |
| FSD1L | 280.7341139 | 0.549483843 | 0.447920303 | 1.226744666 | 0.219918574 | 0.528494327 |
| FSIP1 | 43.64735242 | 0.719657209 | 0.524333492 | 1.372518103 | 0.169902207 | 0.472495848 |
| FSIP2 | 358.760257 | -0.635017607 | 0.389022089 | -1.632343318 | 0.102607183 | 0.367636293 |
| FST | 2375.880256 | 0.585090057 | 0.584908403 | 1.000310569 | 0.317160234 | 0.627198876 |
| FSTL1 | 23473.52561 | -0.385993243 | 0.273078513 | -1.413488151 | 0.157512243 | 0.456731315 |
| FSTL3 | 227.073926 | 0.83737532 | 0.298603684 | 2.804303381 | 0.005042543 | 0.075668075 |
| FSTL4 | 194.7087432 | -0.582646115 | 0.32852424 | -1.773525494 | 0.076141671 | 0.316564893 |

| | | | | | | |
|---|---|---|---|---|---|---|
| FTCDNL1 | 19.63592683 | -0.415634164 | 0.941415191 | -0.441499318 | 0.658851555 | 0.851400081 |
| FTH1 | 37131.72919 | 0.355285755 | 0.516721353 | 0.687577072 | 0.491719151 | 0.758965432 |
| FTL | 36344.21262 | 0.769577411 | 0.442952012 | 1.737383262 | 0.082319544 | 0.328390691 |
| FTLP12 | 12.17597684 | -1.005716344 | 0.886686009 | -1.13424181 | 0.25669313 | 0.569091258 |
| FTLP14 | 15.25639006 | -0.972321687 | 1.156851903 | -0.840489335 | 0.400634079 | 0.698141466 |
| FTO | 1269.375355 | -0.346786507 | 0.282675645 | -1.226800091 | 0.219897737 | 0.528494327 |
| FTSJ1 | 769.0453544 | 0.276162124 | 0.265195852 | 1.041351597 | 0.297712398 | 0.60928029 |
| FTSJ3 | 2498.243409 | -0.407409042 | 0.263818396 | -1.544278369 | 0.12252091 | 0.403241325 |
| FUBP1 | 5202.595377 | -0.034570811 | 0.361318517 | -0.095679598 | 0.923775045 | 0.971413208 |
| FUBP3 | 1912.348169 | -0.160672114 | 0.237774992 | -0.67573176 | 0.499210968 | 0.76281557 |
| FUCA1 | 252.7683983 | 0.062224853 | 0.281927805 | 0.220712012 | 0.825316678 | 0.927129161 |
| FUCA2 | 1710.616095 | 0.12562206 | 0.279589602 | 0.449308768 | 0.653208934 | 0.848827707 |
| FUK | 352.5759594 | -0.170206148 | 0.376481209 | -0.452097326 | 0.65119887 | 0.847876685 |
| FUNDC1 | 523.4679741 | 0.662607232 | 0.343444573 | 1.929298889 | 0.053693768 | 0.267861241 |
| FUNDC2 | 1374.148448 | 0.390528161 | 0.268955788 | 1.452016202 | 0.1464971 | 0.439317555 |
| FUOM | 289.8529327 | 0.524459418 | 0.467021087 | 1.122988731 | 0.261442281 | 0.574937735 |
| FURIN | 1888.524661 | -1.189657357 | 0.446725227 | -2.663062853 | 0.007743295 | 0.098551617 |
| FUS | 12836.86312 | -0.305522008 | 0.255890418 | -1.193956424 | 0.232495014 | 0.542005591 |
| FUT1 | 164.3193911 | -0.142787286 | 0.382469702 | -0.373329667 | 0.708903099 | 0.8756529 |
| FUT10 | 424.6716961 | -0.151179108 | 0.305720886 | -0.494500426 | 0.620952806 | 0.833349572 |
| FUT11 | 276.8805918 | 0.259271837 | 0.321045356 | 0.80758632 | 0.419328764 | 0.709924862 |
| FUT2 | 48.71295004 | -0.367770124 | 0.687181361 | -0.535186408 | 0.592520964 | 0.819859199 |
| FUT4 | 837.9672203 | -1.599014277 | 0.510085973 | -3.134793668 | 0.00171975 | 0.038595164 |
| FUT8 | 4541.160708 | -1.148698528 | 0.335049941 | -3.428439726 | 0.000607061 | 0.020018041 |
| FUT9 | 155.9649029 | 0.207557451 | 0.525517818 | 0.39495797 | 0.692873907 | 0.86854217 |
| FUZ | 442.8222827 | 0.340232536 | 0.366827056 | 0.927501204 | 0.353666367 | 0.660866257 |
| FXN | 380.9410279 | 0.684592782 | 0.366153459 | 1.869688147 | 0.061527137 | 0.285889185 |
| FXR1 | 10325.43675 | 0.045021483 | 0.239676124 | 0.187843005 | 0.850999723 | 0.939477675 |
| FXR2 | 1374.307486 | 0.090148074 | 0.30643144 | 0.294186766 | 0.768615193 | 0.904372313 |
| FXYD1 | 19.66207403 | 1.491731228 | 0.82642651 | 1.805037967 | 0.071068741 | 0.306737943 |
| FXYD2 | 15.07596285 | 0.945656114 | 0.84634328 | 1.117343444 | 0.263847507 | 0.577444856 |
| FXYD3 | 223.4053752 | -0.891777967 | 0.940783595 | -0.947909776 | 0.343175388 | 0.65005791 |
| FXYD5 | 674.8813906 | 0.244248595 | 0.302053609 | 0.808626641 | 0.418729936 | 0.709782331 |
| FXYD6 | 3215.789052 | 0.176630979 | 0.271103377 | 0.651526296 | 0.514706808 | 0.77297733 |
| FXYD7 | 11.37674267 | 2.106590087 | 1.031184276 | 2.042884223 | 0.0410639 | 0.232934829 |
| FYB2 | 18.60032676 | -1.083815483 | 0.990554978 | -1.094149752 | 0.273889305 | 0.586847211 |
| FYCO1 | 481.982535 | -0.887140146 | 0.487192196 | -1.820924377 | 0.068618353 | 0.301741749 |
| FYN | 3147.451102 | 0.349138537 | 0.361134207 | 0.966783345 | 0.333652352 | 0.641713975 |
| FYTTD1 | 1239.418336 | 0.007122562 | 0.225455302 | 0.031591902 | 0.974797501 | 0.991261106 |
| FZD1 | 401.047204 | 0.951101301 | 0.354595266 | 2.682216574 | 0.00731361 | 0.094825773 |
| FZD2 | 1028.972993 | 1.468338212 | 0.64035236 | 2.293016007 | 0.021847083 | 0.167819496 |
| FZD3 | 2575.453843 | 0.806169549 | 0.320408331 | 2.516069252 | 0.011867186 | 0.123589668 |
| FZD4 | 1506.142734 | -2.247640423 | 0.438073504 | -5.130738118 | 2.89E-07 | |

| | | | | | | |
|---|---|---|---|---|---|---|
| | | | | | | 8.32E-05 |
| FZD5 | 533.4742951 | -0.241948218 | 0.346947818 | -0.697361983 | 0.485576287 | 0.755840687 |
| FZD6 | 1116.951962 | -0.507940874 | 0.278858749 | -1.821498786 | 0.068531071 | 0.301620214 |
| FZD7 | 7054.092727 | 1.073771526 | 0.386054607 | 2.781398035 | 0.005412533 | 0.07827792 |
| FZD8 | 258.0523679 | 1.570645234 | 0.412378547 | 3.808746228 | 0.000139673 | 0.007590614 |
| FZD9 | 12.28356684 | -1.106159087 | 1.194195032 | -0.926280095 | 0.354300438 | 0.661631209 |
| FZR1 | 697.2995465 | 0.191858779 | 0.412960567 | 0.464593461 | 0.642222616 | 0.844148268 |
| G2E3 | 1916.359855 | 0.890754294 | 0.416121457 | 2.140611301 | 0.032305397 | 0.204807845 |
| G3BP1 | 7697.644953 | 0.239254684 | 0.288104223 | 0.830444903 | 0.406287287 | 0.701474651 |
| G3BP2 | 4489.314089 | 0.055268865 | 0.302733181 | 0.182566261 | 0.855138359 | 0.941800657 |
| G6PC3 | 2300.293256 | 0.277120076 | 0.261581305 | 1.059403216 | 0.289416186 | 0.60229533 |
| G6PD | 1790.76647 | -0.438056999 | 0.246140211 | -1.779705139 | 0.075124229 | 0.314600339 |
| GAA | 2202.193174 | 0.091736046 | 0.35792349 | 0.256300713 | 0.797718646 | 0.915916373 |
| GAB1 | 800.5671873 | -0.50722521 | 0.322151915 | -1.574490747 | 0.115374039 | 0.391137329 |
| GAB2 | 2287.327749 | -1.782942343 | 0.325209096 | -5.482449183 | 4.19E-08 | 2.27E-05 |
| GABARAP | 4824.496683 | 0.544337769 | 0.38665146 | 1.407825456 | 0.159182761 | 0.459030918 |
| GABARAPL1 | 1596.066384 | -0.574442853 | 0.23912165 | -2.402303819 | 0.016292171 | 0.144292038 |
| GABARAPL2 | 1357.809731 | 1.146335573 | 0.376301789 | 3.046319753 | 0.002316613 | 0.04595021 |
| GABBR1 | 572.4875428 | -1.124280125 | 0.866843306 | -1.296981955 | 0.194637396 | 0.500397209 |
| GABPA | 1333.043552 | 0.192492425 | 0.280490381 | 0.686271039 | 0.492542208 | 0.759174768 |
| GABPB1 | 856.3952793 | 0.309994691 | 0.254427415 | 1.218401293 | 0.223071514 | 0.532026087 |
| GABPB2 | 154.0962071 | -0.197462719 | 0.338039416 | -0.584141107 | 0.55912538 | 0.799405442 |
| GABRA2 | 61.66552131 | -0.537941408 | 0.782698202 | -0.687290972 | 0.491899387 | 0.759089166 |
| GABRA3 | 134.6415779 | 0.515871113 | 0.522510376 | 0.987293529 | 0.323498762 | 0.633346202 |
| GABRA5 | 13.61271199 | 0.253885057 | 0.960118668 | 0.264430914 | 0.791447893 | 0.913361935 |
| GABRB2 | 36.05545669 | -0.95698712 | 0.913293791 | -1.047841483 | 0.294711646 | 0.6068228 |
| GABRB3 | 441.7340613 | -0.490484225 | 0.29444417 | -1.665797031 | 0.095753848 | 0.355265743 |
| GABRE | 45.99338699 | 0.367298579 | 0.736409697 | 0.498769341 | 0.617941889 | 0.832176095 |
| GABRG2 | 99.22557143 | -0.178596574 | 0.404025985 | -0.442042297 | 0.658458599 | 0.851158424 |
| GABRP | 4760.841695 | -5.883707958 | 1.354863369 | -4.342657786 | 1.41E-05 | 0.00142006 |
| GABRQ | 78.89762109 | 0.201013256 | 0.601734511 | 0.334056386 | 0.738337011 | 0.891242475 |
| GABRR1 | 18.49859167 | -1.449121982 | 1.100253914 | -1.317079597 | 0.187811945 | 0.492689771 |
| GAD1 | 742.7793697 | 0.699499013 | 0.658918984 | 1.061585764 | 0.288423779 | 0.601673803 |
| GADD45A | 2865.923412 | 0.92737812 | 0.672931313 | 1.378117057 | 0.16816715 | 0.470384431 |
| GADD45B | 210.385277 | -1.090592488 | 0.729676004 | -1.494625671 | 0.135012163 | 0.421801795 |
| GADD45G | 74.09295069 | -2.043942684 | 1.046178746 | -1.953722241 | 0.050734075 | 0.259171491 |
| GADD45GIP1 | 976.9419938 | 1.102475837 | 0.505550141 | 2.180744792 | 0.029202298 | 0.195546191 |
| GADL1 | 21.68748919 | -0.245014635 | 0.937248615 | -0.261419042 | 0.793769374 | 0.914441125 |
| GAK | 1139.787892 | -0.594848773 | 0.499165197 | -1.191687195 | 0.233383921 | |

| | | | | | | |
|---|---|---|---|---|---|---|
| | | | | | | 0.543263647 |
| GAL | 32.4049676 | 0.5837647 | 0.988158258 | 0.590760331 | 0.554681018 | 0.796360966 |
| GAL3ST1 | 40.18015569 | 1.428957804 | 0.704798758 | 2.027469243 | 0.042614447 | 0.237346523 |
| GAL3ST3 | 159.1803087 | 1.615441961 | 0.439365103 | 3.676764378 | 0.000236211 | 0.010544512 |
| GAL3ST4 | 118.4827712 | 0.638809522 | 0.4843431 | 1.318919423 | 0.187196052 | 0.492107313 |
| GALC | 831.2564475 | 0.316947346 | 0.318578007 | 0.994881437 | 0.319793932 | 0.629465081 |
| GALE | 670.2264212 | 0.073380052 | 0.249999568 | 0.293520715 | 0.769124168 | 0.904597192 |
| GALK1 | 238.5578216 | 0.738915467 | 0.525079618 | 1.407244618 | 0.159354865 | 0.459178093 |
| GALK2 | 343.3638687 | 0.325968165 | 0.279852596 | 1.164785212 | 0.244105946 | 0.555457814 |
| GALM | 203.9990857 | -0.027182204 | 0.334680115 | -0.081218462 | 0.935268217 | 0.975231691 |
| GALNS | 444.5908273 | -0.36643852 | 0.319088874 | -1.148390149 | 0.250807539 | 0.563811922 |
| GALNT1 | 2778.423358 | 0.031835962 | 0.281221811 | 0.113205877 | 0.909867337 | 0.96630013 |
| GALNT10 | 510.0059332 | -0.260492239 | 0.411054151 | -0.633717573 | 0.526265154 | 0.778525486 |
| GALNT11 | 1611.236363 | -0.394259056 | 0.371916079 | -1.060075318 | 0.289110336 | 0.602227073 |
| GALNT12 | 1238.34714 | -0.116775413 | 0.458475605 | -0.254703656 | 0.798951992 | 0.916056735 |
| GALNT13 | 98.36212875 | -0.178391725 | 0.395119443 | -0.451488097 | 0.651637802 | 0.8478779 |
| GALNT14 | 26.1746678 | 0.36640726 | 0.67241693 | 0.544910819 | 0.585814855 | 0.815639728 |
| GALNT16 | 239.9938322 | 1.347644895 | 0.649656314 | 2.074396669 | 0.038042486 | 0.224307253 |
| GALNT17 | 875.221519 | 0.124715514 | 0.395354997 | 0.315451974 | 0.75241851 | 0.896586461 |
| GALNT18 | 82.76980184 | -0.136365069 | 0.601894481 | -0.22655976 | 0.820766085 | 0.925132406 |
| GALNT2 | 2079.22628 | 0.587310868 | 0.385185835 | 1.524746797 | 0.127322264 | 0.410892592 |
| GALNT3 | 117.7535198 | -1.032124841 | 0.716528856 | -1.440451187 | 0.149739791 | 0.444386222 |
| GALNT4 | 154.0269794 | -0.774927721 | 0.449303738 | -1.724730191 | 0.084576108 | 0.332854332 |
| GALNT5 | 34.62586817 | -2.221886459 | 0.775131462 | -2.866464036 | 0.004150853 | 0.066435955 |
| GALNT6 | 136.935032 | -1.681547586 | 0.650844869 | -2.583638078 | 0.009776432 | 0.112404636 |
| GALNT7 | 1227.826899 | -0.129377544 | 0.29899297 | -0.432710991 | 0.665224744 | 0.855087509 |
| GALNT8 | 87.24397969 | -0.426679542 | 0.505945519 | -0.843331003 | 0.399043348 | 0.697352242 |
| GALP | 81.59796355 | 2.476509184 | 0.913231369 | 2.711809152 | 0.006691711 | 0.089444885 |
| GALT | 698.0056976 | -0.732496475 | 0.390800907 | -1.874346915 | 0.060882625 | 0.284375094 |
| GAMT | 466.2380032 | 0.899328063 | 0.408262389 | 2.202818795 | 0.027607523 | 0.190206054 |
| GAN | 552.2486892 | -0.095210816 | 0.495527375 | -0.192140375 | 0.847632249 | 0.938207222 |
| GANAB | 13431.74969 | -0.361146366 | 0.325118961 | -1.110812993 | 0.266648853 | 0.579921585 |
| GANC | 450.7366066 | -0.344285751 | 0.269884956 | -1.275675962 | 0.202070089 | 0.508729725 |
| GAP43 | 385.7554606 | 0.979263912 | 0.303458956 | 3.227006132 | 0.001250928 | 0.031368307 |
| GAPDH | 65488.65297 | 0.093145734 | 0.22291714 | 0.417849133 | 0.676057425 | 0.860110127 |
| GAPDHP21 | 13.66783002 | -1.356131101 | 0.983520425 | -1.378854029 | 0.167939764 | 0.470384431 |
| GAPDHP35 | 13.60786498 | 0.915785879 | 0.81330338 | 1.126007713 | 0.260162255 | 0.57337005 |

| | | | | | | |
|---|---|---|---|---|---|---|
| GAPVD1 | 1039.122531 | -0.288185145 | 0.285815387 | -1.008291218 | 0.313314678 | 0.623018103 |
| GAR1 | 680.4133257 | 0.601384422 | 0.432880642 | 1.389261527 | 0.164753238 | 0.467103603 |
| GAREM1 | 304.4409233 | -0.349027429 | 0.338460767 | -1.031219754 | 0.302437779 | 0.613293571 |
| GAREM2 | 291.3136102 | 0.247323838 | 0.38892522 | 0.635916173 | 0.52483106 | 0.777787635 |
| GARNL3 | 83.78580106 | -0.116731716 | 0.723339635 | -0.161378847 | 0.871795026 | 0.948856124 |
| GARS | 3596.169153 | -0.213055527 | 0.294103833 | -0.724422817 | 0.468806196 | 0.744337782 |
| GART | 2222.478173 | -0.124032471 | 0.426008032 | -0.291150547 | 0.770936183 | 0.905067213 |
| GAS1 | 1253.043032 | -0.459520198 | 0.498210629 | -0.922341218 | 0.35635062 | 0.663825548 |
| GAS2 | 31.59220398 | 0.128178505 | 0.672222753 | 0.190678617 | 0.848777393 | 0.938569494 |
| GAS2L1 | 82.55670707 | -0.116844174 | 0.680043467 | -0.171818684 | 0.863580076 | 0.945060473 |
| GAS2L2 | 10.26934043 | 0.633628239 | 1.275783983 | 0.496657935 | 0.61943029 | 0.832691898 |
| GAS2L3 | 308.6376933 | 0.419153598 | 0.348977139 | 1.201091852 | 0.229715573 | 0.538642189 |
| GAS6 | 1297.387994 | -1.116169414 | 0.245738874 | -4.542095415 | 5.57E-06 | 0.000693377 |
| GAS7 | 937.0708803 | -0.86993612 | 0.409329838 | -2.125269257 | 0.033564152 | 0.208860439 |
| GAS8 | 426.0490395 | -0.552807858 | 0.333990782 | -1.655159027 | 0.097892244 | 0.359375073 |
| GASK1A | 89.22229201 | 0.769490273 | 0.430654769 | 1.786791481 | 0.073971197 | 0.312595272 |
| GATA2 | 392.0834917 | -1.917630272 | 0.756276661 | -2.535620059 | 0.011224844 | 0.120297968 |
| GATA3 | 2121.691976 | -1.961701645 | 0.84657352 | -2.317225378 | 0.020491458 | 0.16183176 |
| GATA5 | 107.6063138 | -0.798465308 | 0.736684426 | -1.083863429 | 0.278425357 | 0.592005173 |
| GATA6 | 233.5699513 | -3.513938867 | 1.009487723 | -3.480912931 | 0.000499708 | 0.017561181 |
| GATAD1 | 1626.386982 | 0.546836835 | 0.311998005 | 1.752693369 | 0.079654655 | 0.32319892 |
| GATAD2A | 2131.777828 | -0.093622361 | 0.246471699 | -0.379850348 | 0.704056506 | 0.873421952 |
| GATAD2B | 403.8919558 | -0.589641823 | 1.604439602 | -0.3675064 | 0.713241319 | 0.877795412 |
| GATB | 666.1133331 | -0.122428734 | 0.296097299 | -0.413474674 | 0.679258874 | 0.861625115 |
| GATC | 737.3133961 | 0.359332721 | 0.259049679 | 1.387118959 | 0.165405492 | 0.467560858 |
| GATD1 | 906.4989271 | 0.590932834 | 0.382177097 | 1.546227752 | 0.122049573 | 0.402659864 |
| GATD3A | 2560.062551 | 0.328213178 | 0.293877453 | 1.116836882 | 0.264064076 | 0.577448862 |
| GATD3B | 123.0001864 | -0.448740492 | 0.77122111 | -0.581857117 | 0.560662928 | 0.800698228 |
| GATM | 119.7052079 | 0.773213521 | 0.394966133 | 1.957670435 | 0.050268697 | 0.257653791 |
| GBA | 796.8915331 | -1.164951615 | 0.352864414 | -3.301414281 | 0.000961987 | 0.026535106 |
| GBA2 | 2382.488206 | -0.357487645 | 0.373166554 | -0.957984154 | 0.338070749 | 0.645353712 |
| GBAP1 | 227.1551421 | -0.855380114 | 0.510742508 | -1.674777603 | 0.093977875 | 0.351340153 |
| GBE1 | 399.8851739 | -0.193427477 | 0.294277254 | -0.657296729 | 0.510990139 | 0.770599585 |
| GBF1 | 1470.063259 | -0.75546163 | 0.568809106 | -1.32814616 | 0.184129827 | 0.488572217 |
| GBGT1 | 195.8065429 | -1.824710493 | 0.546304608 | -3.340097203 | 0.000837491 | 0.024340176 |

| | | | | | | |
|---|---|---|---|---|---|---|
| GBP1P1 | 16.16370803 | 0.566166109 | 0.865676974 | 0.654015443 | 0.513101857 | 0.771865966 |
| GBP4 | 15.29359273 | -0.124694367 | 0.923426714 | -0.135034394 | 0.892584681 | 0.957550733 |
| GBX2 | 87.53798957 | 1.491382472 | 0.448265385 | 3.327007886 | 0.000877839 | 0.025175225 |
| GC | 54.975419 | -3.639003977 | 1.014471304 | -3.587094049 | 0.000334384 | 0.013527829 |
| GCA | 1313.075015 | 0.564564584 | 0.391498816 | 1.442059494 | 0.149285591 | 0.443993668 |
| GCAT | 349.0978049 | 0.584446352 | 0.399318185 | 1.463610659 | 0.143300358 | 0.433760921 |
| GCC1 | 622.1776682 | -0.499569335 | 0.286380195 | -1.744426968 | 0.081084666 | 0.325604941 |
| GCC2 | 1475.296339 | -0.2977895 | 0.267802901 | -1.111972644 | 0.266149913 | 0.57970088 |
| GCDH | 950.91558 | 0.154505139 | 0.279401346 | 0.552986381 | 0.580272736 | 0.811670691 |
| GCFC2 | 776.2088795 | 0.447688376 | 0.263061341 | 1.70184024 | 0.088785321 | 0.342301733 |
| GCH1 | 416.6178613 | -0.235575883 | 0.599643928 | -0.392859616 | 0.69442317 | 0.86892666 |
| GCHFR | 114.1671186 | 0.918964887 | 0.633536494 | 1.450531888 | 0.146910254 | 0.439947781 |
| GCK | 10.21539838 | 0.575250215 | 1.231109309 | 0.467261689 | 0.640312663 | 0.843615231 |
| GCLC | 1047.838679 | -0.135110332 | 0.264646761 | -0.510530836 | 0.609679617 | 0.828409093 |
| GCLM | 272.0807602 | 0.426522015 | 0.407428085 | 1.04686454 | 0.295162059 | 0.60699582 |
| GCN1 | 4299.656015 | -0.519381698 | 0.451905694 | -1.149314346 | 0.250426385 | 0.563180949 |
| GCNA | 30.35948551 | -0.652559925 | 0.608385869 | -1.072608615 | 0.283446758 | 0.597328901 |
| GCNT1 | 726.5753634 | -0.25661505 | 0.416006469 | -0.616853509 | 0.537331354 | 0.785675856 |
| GCNT2 | 329.5692053 | -1.844731771 | 0.451019574 | -4.090136831 | 4.31E-05 | 0.003303284 |
| GCNT4 | 11.45872438 | 0.614516361 | 1.050361047 | 0.585052504 | 0.558512414 | 0.798855459 |
| GCOM1 | 66.18758119 | -1.426953471 | 0.601311197 | -2.373069847 | 0.017640928 | 0.150007193 |
| GCSAML-AS1 | 109.0861353 | -1.051940057 | 0.560475158 | -1.876871869 | 0.060535657 | 0.284105089 |
| GCSH | 2597.182722 | 1.281169352 | 0.401015218 | 3.194814799 | 0.001399206 | 0.033855429 |
| GCSHP5 | 758.5174871 | 1.264964886 | 0.362832532 | 3.486360166 | 0.000489641 | 0.0173966 |
| GDAP1 | 348.1611087 | -0.440260753 | 0.360241913 | -1.222125293 | 0.22166025 | 0.53065007 |
| GDAP1L1 | 29.64243151 | 1.183362466 | 0.724436031 | 1.633494767 | 0.10236498 | 0.367134543 |
| GDAP2 | 468.5472347 | -0.377435538 | 0.280691592 | -1.344662784 | 0.178734158 | 0.483088529 |
| GDE1 | 1841.143786 | -0.085414241 | 0.293492733 | -0.291026766 | 0.771030849 | 0.905067213 |
| GDF10 | 99.66302087 | 1.497682212 | 0.703819483 | 2.127935143 | 0.033342461 | 0.208274771 |
| GDF11 | 1974.661836 | -0.063066313 | 0.332774219 | -0.189516825 | 0.84968777 | 0.938580023 |
| GDF15 | 111.8766098 | 1.923806389 | 0.381663113 | 5.040587686 | 4.64E-07 | 0.000115425 |
| GDF6 | 459.4127874 | -5.981881061 | 0.974273951 | -6.139834753 | 8.26E-10 | 1.25E-06 |
| GDF9 | 15.81525728 | 0.25917625 | 0.810840835 | 0.319638872 | 0.749242104 | 0.895160809 |
| GDI1 | 4710.473126 | 0.249587711 | 0.24015632 | 1.039271886 | 0.298678306 | 0.610022695 |
| GDI2 | 13654.35738 | -0.071278361 | 0.216273371 | -0.329575297 | 0.74172089 | 0.892150449 |
| GDNF | 25.9078904 | 0.007947205 | 1.005393428 | 0.007904572 | 0.993693129 | 0.99730871 |
| GDPD1 | 310.6149028 | 0.373167015 | 0.357539899 | 1.043707334 | 0.296620816 | 0.607866324 |
| GDPD2 | 335.237641 | 0.058472351 | 0.371888838 | 0.157230724 | 0.875063015 | 0.949740014 |
| GDPD3 | 76.92342672 | 0.289343121 | 0.413020019 | 0.700554714 | 0.483580949 | 0.754231168 |
| GDPD5 | 315.5662892 | -0.362157435 | 0.391414686 | -0.925252547 | 0.354834556 | 0.662221067 |
| GDPGP1 | 87.22681117 | 0.093473214 | 0.480306073 | 0.194611768 | 0.845696889 | 0.937897187 |
| GEM | 116.6571489 | 0.32232959 | 0.72798973 | 0.442766671 | 0.657934513 | 0.850760404 |

| | | | | | | |
|---|---|---|---|---|---|---|
| GEMIN2 | 688.8054164 | 0.940045742 | 0.30706505 | 3.061389575 | 0.002203122 | 0.04456476 |
| GEMIN4 | 1024.829326 | -0.126336398 | 0.34121621 | -0.370253213 | 0.711193831 | 0.876982981 |
| GEMIN5 | 905.6102825 | 0.094559481 | 0.379758147 | 0.248999216 | 0.803361388 | 0.917969611 |
| GEMIN6 | 539.0385314 | 1.051208168 | 0.31370287 | 3.350967642 | 0.000805297 | 0.023815132 |
| GEMIN7 | 672.7477291 | 0.44712136 | 0.503992346 | 0.887159028 | 0.374993288 | 0.67920525 |
| GEMIN8 | 466.4033505 | 0.451789358 | 0.304711135 | 1.48268083 | 0.13815923 | 0.426306273 |
| GEMIN8P4 | 41.31166444 | 0.395144837 | 0.489677563 | 0.806949034 | 0.419695847 | 0.710229302 |
| GEN1 | 717.2877804 | -0.095422931 | 0.35655759 | -0.267622774 | 0.788989699 | 0.912533261 |
| GET4 | 1311.188281 | -0.064151942 | 0.353137311 | -0.181662883 | 0.855847296 | 0.942239429 |
| GFAP | 10.58513456 | -0.139738713 | 1.043476559 | -0.133916485 | 0.893468615 | 0.95780189 |
| GFER | 803.2192455 | 0.621157801 | 0.327902305 | 1.894338013 | 0.058180158 | 0.279509071 |
| GFI1 | 17.79473466 | -0.600168521 | 0.865429301 | -0.693492259 | 0.488000682 | 0.756726796 |
| GFM1 | 1159.181206 | 0.021139552 | 0.230322796 | 0.091782283 | 0.926871021 | 0.972378138 |
| GFM2 | 1258.240742 | 0.114364366 | 0.29299693 | 0.390326157 | 0.696295384 | 0.870023227 |
| GFOD1 | 101.4752171 | 0.935489463 | 0.417260525 | 2.241979307 | 0.02496271 | 0.180688185 |
| GFOD2 | 661.8894005 | 0.348679825 | 0.264197109 | 1.319771538 | 0.186911307 | 0.491529111 |
| GFPT1 | 2430.304756 | 0.032683989 | 0.318967073 | 0.102468222 | 0.918385035 | 0.969677217 |
| GFPT2 | 545.392273 | 0.704131332 | 0.535770806 | 1.314239828 | 0.188765509 | 0.493962743 |
| GFRA2 | 22.01329095 | 1.235840971 | 0.760059785 | 1.625978635 | 0.103954201 | 0.369392362 |
| GFRA3 | 12.04117932 | -0.77628142 | 0.915803032 | -0.847651071 | 0.396632323 | 0.695326667 |
| GFY | 26.01571094 | -0.879861638 | 0.894479592 | -0.983657588 | 0.3252839 | 0.635201705 |
| GGA1 | 1627.553047 | -0.159487963 | 0.314457568 | -0.507184368 | 0.612025462 | 0.829098987 |
| GGA2 | 2445.952732 | -0.035491494 | 0.328299556 | -0.108107042 | 0.913910782 | 0.968086893 |
| GGA3 | 1702.906739 | -0.411270267 | 0.32668981 | -1.258901423 | 0.20806594 | 0.514937745 |
| GGACT | 85.20833872 | -0.379749958 | 0.50990591 | -0.74474516 | 0.456425793 | 0.735468477 |
| GGCT | 3494.033975 | 0.754920857 | 0.430781749 | 1.752443918 | 0.079697505 | 0.323199907 |
| GGCX | 1385.641411 | -0.354551823 | 0.309477696 | -1.1456458 | 0.251941739 | 0.564747062 |
| GGH | 2277.711944 | 0.425896684 | 0.448521138 | 0.949557665 | 0.342337058 | 0.649037179 |
| GGNBP2 | 1558.833559 | -0.000939181 | 0.276289417 | -0.003399264 | 0.997287785 | 0.998674292 |
| GGPS1 | 1426.472023 | 0.321383205 | 0.345891731 | 0.929143938 | 0.352814497 | 0.659993678 |
| GGT1 | 1622.053299 | 0.509826071 | 0.401226582 | 1.270668729 | 0.203846525 | 0.509986087 |
| GGT2 | 18.53935949 | -3.51294023 | 1.34382226 | -2.614140526 | 0.008945224 | 0.10677261 |
| GGT5 | 11.52006093 | -1.131447074 | 0.934053657 | -1.211329847 | 0.225769014 | 0.53459368 |
| GGT7 | 809.9058918 | -0.224806787 | 0.486313858 | -0.462266875 | 0.643889951 | 0.845466381 |
| GHDC | 201.3725518 | -0.170550513 | 0.419062594 | -0.406980998 | 0.684021945 | 0.863826466 |
| GHITM | 4041.788357 | 0.142316609 | 0.224977403 | 0.632581794 | 0.527006779 | 0.778755887 |
| GHR | 31.03887065 | -0.594325202 | 0.67474049 | -0.880820421 | 0.378415025 | 0.681253679 |
| GID4 | 477.1649457 | 0.025955266 | 0.34380014 | 0.075495217 | 0.939820703 | 0.978003717 |
| GID8 | 3071.946309 | 0.039392626 | 0.328381875 | 0.119959806 | 0.904514989 | 0.963993604 |
| GIGYF1 | 2776.521367 | 0.113344741 | 0.296831405 | 0.381848884 | 0.702573457 | |

0.872703476
GIGYF2 2225.597822 -0.258838751 0.277249351 -0.933595516
0.350512579 0.65747111
GIN1 309.0606898 0.802259306 0.30053361 2.669449541 0.007597569 0.097104226
GINM1 577.73752 0.357880288 0.350539124 1.020942496 0.307281684 0.617289516
GINS1 1258.094502 0.559506346 0.248935311 2.247597353 0.024601877 0.179095522
GINS2 993.8783439 0.849166619 0.240601609 3.529347214 0.000416586 0.015721465
GINS3 289.7083659 0.066375237 0.354274357 0.187355466 0.851381937 0.939477675
GINS4 1044.72504 0.046378731 0.299267461 0.154974185 0.876841668 0.950454769
GIP 15.41739768 1.018201294 0.889434538 1.14477373 0.2523029 0.565139126
GIPC1 1464.870491 0.247782619 0.433426929 0.571682566 0.567537047 0.805151447
GIPC3 11.97243637 -0.841348472 1.045621543 -0.804639572 0.421027699
0.711530715
GIT1 1437.509511 0.339127912 0.356422722 0.951476691 0.341362446 0.648152255
GIT2 637.3238399 -0.344171215 0.322271434 -1.067954457 0.285541056
0.599411008
GJA1 22431.12689 -0.111048378 0.269433407 -0.412155193 0.680225677
0.862002279
GJA10 10.1072769 -0.724520923 1.057063181 -0.685409289 0.493085684
0.759389046
GJA3 11.49527989 0.172444977 0.980052984 0.175954749 0.860329493 0.943476849
GJA5 20.74723569 -0.421747137 0.715330717 -0.58958343 0.555469964
0.796360966
GJB2 14.06696458 0.138857379 1.013327227 0.137031134 0.891006186 0.957017706
GJB4 16.32108218 0.825013082 0.94043264 0.877269724 0.380340153 0.6820497
GJB5 21.42551712 -1.024122479 0.863135897 -1.186513598 0.235419535
0.546108526
GJB7 50.53002677 -0.171688578 0.514776039 -0.33352092 0.738741101
0.89135397
GJC1 1793.566478 -0.17519 0.391874924 -0.447055908 0.654834693
0.849258129
GJC2 14.00049548 0.976514012 0.958777697 1.01849888 0.308440933 0.618063319
GJC3 20.1429925 -0.130879579 0.793750966 -0.164887458 0.869032565
0.947201167
GJD2 10.8202127 -4.452382022 2.230519118 -1.996119193 0.04592095
0.246869858
GK 124.6128588 0.779965452 0.426341915 1.829436479 0.067334247 0.298083457
GK5 572.1681724 0.09037893 0.351528793 0.257102496 0.797099651 0.915732089
GKAP1 304.9353039 1.059082598 0.322194337 3.287092525 0.001012276 0.027467362
GLA 710.9125795 0.199841266 0.325454548 0.614037405 0.53919061 0.786771255
GLB1 3785.743909 -2.015766878 0.672001293 -2.99964732 0.002702924
0.05075007
GLB1L 202.9706472 0.151047905 0.322053408 0.469015082 0.639058856 0.842984254
GLB1L2 61.82468456 0.428807337 0.711603852 0.602592771 0.546779631
0.792249935
GLB1L3 89.91513889 0.32329878 0.499915376 0.646707015 0.517821582
0.773950969
GLCCI1 491.6476 0.093289051 0.288926628 0.322881457 0.74678502
0.894109554
GLCE 418.4647309 -0.198514064 0.292990604 -0.67754413 0.498060781
0.762537616
GLDC 1773.619809 -0.237692388 0.42345069 -0.561322473 0.574577724
0.808359708
GLDN 76.5826634 0.115562377 0.397641925 0.290619197 0.771342577 0.905067213
GLE1 913.9869755 -0.2583068 0.234463203 -1.101694411 0.270594548
0.583257419
GLG1 4744.289378 -0.618886111 0.331701416 -1.865792792 0.062070361
0.287269509
GLI1 210.070875 0.26244962 0.613707602 0.427646031 0.668908848 0.857385614
GLI2 389.3287976 0.350047447 0.553336727 0.632611989 0.526987056 0.778755887
GLI3 1766.905673 -0.345473421 0.355947707 -0.970573526 0.331760695
0.639616406
GLI4 158.5682958 -0.763017425 0.539716673 -1.413736989 0.15743914
0.456731315

| | | | | | | |
|---|---|---|---|---|---|---|
| GLIPR1 | 10.22469777 | 0.358219397 | 0.985226997 | 0.363590724 | 0.716163655 | 0.879750511 |
| GLIPR1L2 | 15.93801706 | 0.777007351 | 0.819441017 | 0.948216326 | 0.343019338 | 0.649843453 |
| GLIPR2 | 406.4892476 | 0.986609496 | 0.492583251 | 2.002929443 | 0.045184862 | 0.244908734 |
| GLIS2 | 112.5380767 | 0.093438304 | 0.601362937 | 0.155377556 | 0.876523677 | 0.950324661 |
| GLMN | 706.9134383 | 0.285677339 | 0.32129538 | 0.889142379 | 0.373926566 | 0.678520659 |
| GLMP | 1835.466825 | -0.668634117 | 0.311421584 | -2.147038459 | 0.03179022 | 0.203069232 |
| GLO1 | 6051.677995 | 0.466615461 | 0.230746442 | 2.022200024 | 0.043155692 | 0.238947081 |
| GLOD4 | 2223.431786 | 0.053474527 | 0.254505359 | 0.210111594 | 0.833580577 | 0.931858443 |
| GLRA3 | 15.50296658 | -0.165668632 | 0.873340875 | -0.189695269 | 0.849547929 | 0.938580023 |
| GLRA4 | 13.3588412 | -0.172003476 | 1.017822653 | -0.168991597 | 0.865803251 | 0.945447428 |
| GLRB | 38.04656225 | -0.698791507 | 0.637819753 | -1.09559402 | 0.27325648 | 0.586278328 |
| GLRX | 277.9010836 | 0.767924532 | 0.305403425 | 2.514459465 | 0.011921501 | 0.123877456 |
| GLRX2 | 135.3292788 | 0.539729516 | 0.393607227 | 1.371238837 | 0.170300515 | 0.473278827 |
| GLRX3 | 2919.760394 | 0.725926467 | 0.313571801 | 2.315024705 | 0.02061158 | 0.162440666 |
| GLRX5 | 1587.315526 | 0.900435633 | 0.371714287 | 2.422386397 | 0.015418946 | 0.140677297 |
| GLS | 1360.586329 | 0.556303417 | 0.272429278 | 2.04201039 | 0.041150501 | 0.232946328 |
| GLS2 | 91.94193504 | -0.265600429 | 0.534558348 | -0.496859566 | 0.619288087 | 0.832614282 |
| GLT8D1 | 1285.267462 | 0.365089946 | 0.252338292 | 1.446827364 | 0.147945281 | 0.441755032 |
| GLT8D2 | 172.5767684 | -0.208935023 | 0.391239638 | -0.534033371 | 0.593318446 | 0.819859199 |
| GLTP | 728.6788609 | 0.297818061 | 0.364121714 | 0.817907995 | 0.413409724 | 0.705652445 |
| GLUD1 | 2674.631644 | 0.007917349 | 0.25893505 | 0.030576585 | 0.975607216 | 0.991554705 |
| GLUD1P2 | 26.84546605 | 0.352650219 | 0.8017191 | 0.439867554 | 0.660033037 | 0.852207761 |
| GLUD1P3 | 20.27761212 | -0.81395564 | 0.699424964 | -1.163749769 | 0.244525434 | 0.555842876 |
| GLUD2 | 447.0085771 | -2.941951118 | 0.606983439 | -4.846839185 | 1.25E-06 | 0.000216263 |
| GLUL | 12767.1195 | -0.039483695 | 0.250794003 | -0.157434765 | 0.874902217 | 0.949740014 |
| GLYAT | 45.57692796 | 1.122026971 | 0.966270973 | 1.161192877 | 0.245563471 | 0.557291155 |
| GLYATL1 | 138.2920764 | -0.003139679 | 0.340009916 | -0.009234081 | 0.992632374 | 0.997035603 |
| GLYATL2 | 16.02839224 | 0.802791272 | 0.853925379 | 0.940118764 | 0.347156645 | 0.653844005 |
| GLYCTK | 131.8597037 | -0.38875785 | 0.378684906 | -1.026599804 | 0.304608946 | 0.614996026 |
| GLYR1 | 2868.678044 | -0.331921672 | 0.367679931 | -0.902746232 | 0.366660594 | 0.673276188 |
| GM2A | 2038.829393 | -0.739677472 | 0.359299682 | -2.058664421 | 0.039526397 | 0.229225905 |
| GMCL1 | 867.10939 | 0.09236295 | 0.244896552 | 0.377150882 | 0.70606148 | 0.87442112 |
| GMDS | 448.6039812 | 0.180065397 | 0.395465302 | 0.4553254 | 0.648875156 | 0.847033866 |
| GMEB1 | 735.5557861 | -0.344248642 | 0.262612658 | -1.31086081 | 0.189904793 | 0.495280824 |
| GMEB2 | 412.375636 | 0.029994194 | 0.375536941 | 0.079870155 | 0.936340527 | 0.976035601 |
| GMFB | 3851.133342 | 0.240791003 | 0.263623194 | 0.913390812 | 0.361037036 | 0.667991431 |
| GMIP | 215.880805 | -0.432109011 | 0.385639596 | -1.120499596 | 0.262500925 | 0.576074285 |
| GMNN | 1569.322572 | 1.110960996 | 0.270121786 | 4.112815239 | 3.91E-05 | 0.00308843 |
| GMPPA | 610.1836116 | -0.290934252 | 0.274717014 | -1.059032519 | 0.28958497 | 0.602398681 |
| GMPPB | 570.8392595 | -0.029819053 | 0.287674118 | -0.103655669 | 0.917442608 | 0.969661544 |
| GMPR | 36.27748103 | 0.425459572 | 0.588087795 | 0.72346268 | 0.469395674 | 0.744895583 |

| | | | | | | |
|---|---|---|---|---|---|---|
| GMPR2 | 1086.197221 | -0.227117617 | 0.253014888 | -0.897645268 | 0.369374697 | 0.675154642 |
| GMPS | 3653.415659 | 0.259771321 | 0.273932633 | 0.948303666 | 0.342974885 | 0.649840387 |
| GNA11 | 392.3213589 | -0.07939311 | 0.43427024 | -0.182819597 | 0.854939571 | 0.941701337 |
| GNA12 | 1651.513042 | -0.33136217 | 0.369770965 | -0.896128147 | 0.370184325 | 0.675901601 |
| GNA13 | 2244.229586 | -0.007176392 | 0.263996817 | -0.027183631 | 0.978313271 | 0.992395125 |
| GNA14 | 10.6870984 | -1.732614604 | 1.155432008 | -1.499538348 | 0.133734028 | 0.420405914 |
| GNAI1 | 1236.469131 | 0.598196713 | 0.244176626 | 2.449852479 | 0.014291475 | 0.135764542 |
| GNAI2 | 6414.010243 | 0.301706363 | 0.331649855 | 0.909713537 | 0.362973603 | 0.669987492 |
| GNAI3 | 3514.183453 | 0.125074239 | 0.237670637 | 0.526250281 | 0.598714323 | 0.823490027 |
| GNAL | 358.1217016 | 0.506371839 | 0.436850755 | 1.1591415 | 0.246398512 | 0.558551201 |
| GNAO1 | 186.0501986 | -0.830819395 | 0.388689594 | -2.137488135 | 0.03255831 | 0.205776324 |
| GNAQ | 2336.153328 | -0.074095082 | 0.244040507 | -0.303617965 | 0.761418963 | 0.90056031 |
| GNAQP1 | 18.51934638 | 0.381170975 | 1.132316282 | 0.336629422 | 0.736396274 | 0.890400696 |
| GNAS | 33289.95339 | -0.271954363 | 0.249974723 | -1.087927448 | 0.276627138 | 0.590171608 |
| GNAZ | 388.0813725 | 1.013919475 | 0.501579638 | 2.021452624 | 0.043232932 | 0.239200149 |
| GNB1 | 10864.69168 | 0.067509242 | 0.249217898 | 0.270884405 | 0.78647994 | 0.911315996 |
| GNB1L | 106.4559003 | 0.814182096 | 0.660122891 | 1.233379584 | 0.217434178 | 0.525520777 |
| GNB2 | 2513.958182 | 0.267912253 | 0.310091539 | 0.86397795 | 0.387600003 | 0.687752006 |
| GNB3 | 163.6597624 | 0.7870484 | 0.440845248 | 1.785316737 | 0.074209956 | 0.313236304 |
| GNB4 | 811.458277 | -0.066581264 | 0.36694648 | -0.181446798 | 0.856016889 | 0.942245037 |
| GNB5 | 627.1610699 | -0.013045373 | 0.252253224 | -0.051715385 | 0.958755478 | 0.985652867 |
| GNE | 427.2307053 | 0.818568095 | 0.414084502 | 1.976814128 | 0.048062644 | 0.252967897 |
| GNG10 | 1604.268786 | 0.663455415 | 0.50590495 | 1.311423054 | 0.189714874 | 0.49512547 |
| GNG11 | 81.52965071 | 1.763869859 | 0.665918718 | 2.648776513 | 0.008078373 | 0.100970518 |
| GNG12 | 652.1498084 | -0.048266726 | 0.27104194 | -0.178078442 | 0.858661372 | 0.942495112 |
| GNG2 | 590.6027101 | -0.132029874 | 0.426815444 | -0.309337153 | 0.757065072 | 0.898563152 |
| GNG3 | 37.12175686 | 0.433417402 | 0.668892191 | 0.647963016 | 0.517008871 | 0.773950969 |
| GNG4 | 66.18123793 | 0.172791301 | 0.500129879 | 0.345492857 | 0.729723879 | 0.886372257 |
| GNG5 | 2278.604827 | 0.918662619 | 0.431161489 | 2.130669468 | 0.033116382 | 0.207838253 |
| GNG7 | 386.269901 | 0.387734955 | 0.348266523 | 1.113328239 | 0.265567483 | 0.579572403 |
| GNG8 | 11.49058667 | 3.680286746 | 1.095804531 | 3.358524847 | 0.000783597 | 0.023439632 |
| GNGT1 | 22.52128163 | 0.920023233 | 0.666266392 | 1.380863936 | 0.1673208 | 0.469643638 |
| GNL1 | 512.9825539 | -0.176470057 | 0.345256419 | -0.511127521 | 0.609261766 | 0.8283007 |
| GNL2 | 2250.043494 | 0.128109347 | 0.32902822 | 0.389356715 | 0.697012287 | 0.870026319 |
| GNL3 | 4511.61777 | 0.530966147 | 0.274291478 | 1.935773399 | 0.05289546 | 0.265985093 |
| GNL3L | 1144.427739 | -0.171297464 | 0.345258892 | -0.496142078 | 0.619794173 | 0.832851852 |
| GNL3LP1 | 67.43645636 | -0.306867827 | 0.637954389 | -0.481018444 | 0.630503391 | 0.838177265 |
| GNPAT | 2155.998643 | 0.555506802 | 0.254998207 | 2.178473365 | 0.029370813 | 0.195861365 |
| GNPDA1 | 2233.802438 | -0.114326588 | 0.269037186 | -0.424947161 | 0.670875194 | 0.858238816 |
| GNPDA2 | 311.9708629 | 0.541900046 | 0.318019338 | 1.703984574 | 0.088383969 | 0.341710805 |
| GNPNAT1 | 1232.259571 | 0.629855051 | 0.340316848 | 1.850790092 | 0.064199757 | 0.292572699 |
| GNPTAB | 991.7780814 | -0.658207964 | 0.314945143 | -2.089913048 | 0.036625611 | 0.219135375 |
| GNPTG | 817.5420423 | 0.252106446 | 0.433261674 | 0.581880331 | 0.56064729 | 0.800698228 |
| GNRH1 | 39.12272565 | 0.394256937 | 0.607198399 | 0.649304968 | 0.516141275 | 0.773896629 |

| | | | | | | |
|---|---|---|---|---|---|---|
| GNRHR2 | 34.19806086 | 1.209635888 | 0.650829311 | 1.85860696 | 0.06308287 | 0.289778852 |
| GNS | 1592.830354 | -0.317155609 | 0.253744169 | -1.249903043 | 0.211334968 | 0.518628728 |
| GOLGA1 | 483.5400021 | -0.322012916 | 0.320291765 | -1.005373698 | 0.31471694 | 0.624045314 |
| GOLGA2 | 1816.408341 | -0.527827901 | 0.230834458 | -2.286607922 | 0.022218718 | 0.169472186 |
| GOLGA2P10 | 189.3946875 | -0.004388349 | 1.046688642 | -0.004192602 | 0.996654797 | 0.998431718 |
| GOLGA2P11 | 16.63596013 | -1.152090941 | 0.782943085 | -1.471487472 | 0.141159333 | 0.430963282 |
| GOLGA2P5 | 695.3476769 | -0.914989332 | 0.275008892 | -3.327126358 | 0.000877466 | 0.025175225 |
| GOLGA2P7 | 3771.767636 | 0.051181007 | 0.446467438 | 0.114635476 | 0.90873406 | 0.965857971 |
| GOLGA3 | 1056.715909 | -0.37874209 | 0.261685079 | -1.447320161 | 0.147807275 | 0.44174616 |
| GOLGA4 | 3619.03799 | -0.335089876 | 0.237303651 | -1.412072149 | 0.157928718 | 0.457277356 |
| GOLGA5 | 1316.358782 | -0.403734759 | 0.236378906 | -1.707998252 | 0.087636667 | 0.34024836 |
| GOLGA6L10 | 100.5610174 | -0.059527546 | 0.441430784 | -0.13485137 | 0.892729389 | 0.957550733 |
| GOLGA6L4 | 32.95697568 | -0.504077491 | 0.74782735 | -0.674055972 | 0.50027573 | 0.763396006 |
| GOLGA6L5P | 28.0356882 | -2.270100256 | 0.758603177 | -2.992473964 | 0.002767263 | 0.051511831 |
| GOLGA6L9 | 117.9978887 | -0.318266262 | 0.485793559 | -0.655147143 | 0.512373022 | 0.771687791 |
| GOLGA7 | 1280.506706 | 0.336298468 | 0.256274476 | 1.312258922 | 0.189432787 | 0.494900088 |
| GOLGA8A | 639.5360767 | 0.046307431 | 0.425423542 | 0.108850183 | 0.91332132 | 0.967876517 |
| GOLGA8B | 2320.52181 | -0.475764544 | 0.3549319 | -1.340438953 | 0.180102679 | 0.484754911 |
| GOLGA8N | 10.84243439 | 0.312005769 | 1.079489784 | 0.289030775 | 0.772557827 | 0.905717252 |
| GOLGB1 | 3026.302646 | -0.589778945 | 0.324483776 | -1.817591476 | 0.069126595 | 0.303010569 |
| GOLIM4 | 1407.324821 | -0.169186857 | 0.385766805 | -0.438572875 | 0.660971055 | 0.852848642 |
| GOLM1 | 3320.107107 | -0.052365272 | 0.261009234 | -0.20062613 | 0.840990925 | 0.935043849 |
| GOLPH3 | 2643.793503 | -0.346603734 | 0.286101283 | -1.211472138 | 0.225714507 | 0.53459368 |
| GOLPH3L | 913.6697785 | -0.077175502 | 0.250413688 | -0.308192027 | 0.757936219 | 0.899104729 |
| GOLT1A | 69.19105016 | -0.289417457 | 0.808960856 | -0.357764476 | 0.720519577 | 0.88214152 |
| GOLT1B | 1281.463862 | 0.548791189 | 0.281419648 | 1.950081283 | 0.051166432 | 0.260310508 |
| GON4L | 1770.683318 | -0.766333862 | 0.373673215 | -2.05081293 | 0.040285168 | 0.231327132 |
| GON7 | 142.0182958 | 0.642435021 | 0.412651599 | 1.556846072 | 0.119507036 | 0.398645831 |
| GOPC | 1527.375639 | -0.377694511 | 0.28490429 | -1.325689096 | 0.184942701 | 0.489406194 |
| GORAB | 456.0360596 | -0.03769668 | 0.274849224 | -0.137154036 | 0.890909041 | 0.957017706 |
| GORASP1 | 975.097314 | -0.4612636 | 0.332934331 | -1.385449195 | 0.165915157 | 0.467940842 |
| GORASP2 | 2540.46139 | -0.005731112 | 0.238860757 | -0.023993528 | 0.980857771 | 0.993241851 |
| GOSR1 | 1259.702377 | -0.082741409 | 0.229877079 | -0.359937623 | 0.718893781 | |

| | | | | | | |
|---|---|---|---|---|---|---|
| | | | | | | 0.881106605 |
| GOSR2 | 999.803364 | -0.237095424 | 0.239245608 | -0.991012648 | 0.321679404 | 0.631332243 |
| GOT1 | 1380.33551 | 0.270241649 | 0.254714199 | 1.060960283 | 0.288707951 | 0.601894782 |
| GOT2 | 2982.637047 | 0.125950696 | 0.248085348 | 0.507690988 | 0.611670069 | 0.829058038 |
| GPA33 | 29.27176675 | -0.331305331 | 0.702531687 | -0.471587741 | 0.637221079 | 0.841584486 |
| GPAA1 | 1022.958185 | 0.122119487 | 0.464880465 | 0.262690082 | 0.792789462 | 0.913936848 |
| GPALPP1 | 852.921942 | 0.30743216 | 0.247060721 | 1.244358706 | 0.213367574 | 0.52058531 |
| GPAM | 784.8213094 | -0.145130861 | 0.296758812 | -0.489053248 | 0.624804 | 0.835033444 |
| GPANK1 | 56.60582045 | 0.087709399 | 0.500713718 | 0.175168756 | 0.860947034 | 0.943676573 |
| GPAT4 | 2097.906933 | -0.227297883 | 0.304421333 | -0.746655566 | 0.455271498 | 0.735091942 |
| GPATCH1 | 501.5248225 | -0.327282057 | 0.286044579 | -1.144164515 | 0.252555416 | 0.565423035 |
| GPATCH11 | 277.1801818 | 0.338418931 | 0.358081297 | 0.945089662 | 0.344613103 | 0.651368096 |
| GPATCH2 | 621.6392244 | -0.013276189 | 0.254976804 | -0.052068221 | 0.958474334 | 0.985438602 |
| GPATCH2L | 1796.997852 | -0.161199612 | 0.367736172 | -0.438356693 | 0.661127735 | 0.852961039 |
| GPATCH3 | 553.470338 | -0.077948917 | 0.317547491 | -0.245471684 | 0.806091231 | 0.919089559 |
| GPATCH4 | 905.7112988 | 0.394805081 | 0.308694177 | 1.278952146 | 0.200913908 | 0.506881795 |
| GPATCH8 | 1985.822416 | -0.436537162 | 0.368228576 | -1.185505934 | 0.23581747 | 0.546697254 |
| GPBP1 | 4694.33302 | 0.287363194 | 0.287186317 | 1.000615897 | 0.317012542 | 0.627198876 |
| GPBP1L1 | 3012.068089 | -0.137775956 | 0.233199547 | -0.590807133 | 0.554649655 | 0.796360966 |
| GPC1 | 1006.495978 | -0.20798931 | 0.409426603 | -0.508001456 | 0.611452322 | 0.829058038 |
| GPC2 | 825.3973426 | 0.94005124 | 0.463489513 | 2.028203903 | 0.042539442 | 0.237179666 |
| GPC3 | 1796.943024 | -0.035370512 | 0.453576478 | -0.077981363 | 0.937842878 | 0.97699748 |
| GPC4 | 1886.637285 | 0.43435503 | 0.390373515 | 1.112665213 | 0.265852241 | 0.579572403 |
| GPC5 | 13.33350376 | -1.764025139 | 1.045486287 | -1.687277165 | 0.09155007 | 0.347226527 |
| GPC6 | 898.4943443 | -0.575683887 | 0.35411797 | -1.625683913 | 0.104016914 | 0.369478015 |
| GPCPD1 | 846.5023133 | -0.047159897 | 0.275018237 | -0.171479163 | 0.863847013 | 0.945079904 |
| GPD1 | 11.55011488 | 0.482675252 | 0.891174018 | 0.541617285 | 0.588082182 | 0.817253188 |
| GPD1L | 149.0057375 | 0.584927565 | 0.462755063 | 1.26401116 | 0.206226021 | 0.512653902 |
| GPD2 | 1048.184809 | -0.262567892 | 0.268180121 | -0.979072912 | 0.327543955 | 0.637399863 |
| GPER1 | 60.52421151 | -0.32113004 | 0.576662763 | -0.556876671 | 0.577611695 | 0.809936484 |
| GPHA2 | 21.1791154 | 1.01593766 | 0.759114676 | 1.338319087 | 0.180792442 | 0.485502272 |
| GPHN | 503.2294248 | 0.052668136 | 0.322288134 | 0.163419407 | 0.870188223 | 0.947915957 |
| GPI | 4685.675116 | -0.094747162 | 0.244525642 | -0.387473317 | 0.698405835 | 0.87055418 |
| GPKOW | 579.5593331 | 0.135784167 | 0.301626216 | 0.450173624 | 0.652585253 | 0.848578973 |
| GPLD1 | 118.5251629 | -0.269707916 | 0.404253439 | -0.667175316 | 0.504660156 | 0.766309601 |
| GPM6A | 189.9928359 | 0.839120736 | 0.607323851 | 1.381669326 | 0.167073257 | 0.469215449 |
| GPM6B | 727.6856364 | 1.477256548 | 0.508489656 | 2.905185051 | 0.00367036 | 0.062311776 |
| GPN1 | 1509.30856 | 0.115360135 | 0.252347657 | 0.457147636 | 0.647564933 | 0.846597949 |
| GPN2 | 647.5391275 | -0.056391741 | 0.317011747 | -0.177885337 | 0.858813027 | 0.942495112 |
| GPN3 | 1162.836149 | 0.985445705 | 0.360514561 | 2.733442174 | 0.006267613 | 0.086050637 |

| | | | | | | |
|---|---|---|---|---|---|---|
| GPNMB | 24.84725267 | -1.139689927 | 0.761113326 | -1.497398465 | 0.134289607 | 0.420619758 |
| GPR1 | 10.77656978 | -0.321620345 | 0.921060137 | -0.349184958 | 0.726950458 | 0.8850111 |
| GPR107 | 2791.773806 | -0.550139738 | 0.361611412 | -1.521356128 | 0.128170492 | 0.412317905 |
| GPR108 | 1494.290246 | -0.461094499 | 0.328597419 | -1.403220089 | 0.160551219 | 0.460961874 |
| GPR135 | 25.46895227 | 0.568225393 | 0.672811257 | 0.844553933 | 0.398359939 | 0.696980583 |
| GPR137 | 559.8325335 | -0.033840556 | 0.384409239 | -0.088032629 | 0.929850742 | 0.97351106 |
| GPR137B | 299.3574557 | 0.269290103 | 0.508915638 | 0.529144877 | 0.596704952 | 0.821812117 |
| GPR137C | 271.6462411 | 0.130696243 | 0.33481328 | 0.390355612 | 0.696273606 | 0.870023227 |
| GPR143 | 179.4354709 | 0.445610357 | 0.371394747 | 1.199829458 | 0.230205581 | 0.539041344 |
| GPR146 | 15.47507412 | 0.041797765 | 0.840327061 | 0.049739877 | 0.960329678 | 0.98646906 |
| GPR153 | 379.6617797 | 1.511754965 | 0.58537027 | 2.582561916 | 0.009806976 | 0.112404636 |
| GPR155 | 54.236683 | -1.475319279 | 0.872232079 | -1.69142974 | 0.090754756 | 0.345245839 |
| GPR156 | 25.51750863 | -0.565023615 | 0.845447184 | -0.66831332 | 0.503933613 | 0.765742872 |
| GPR157 | 445.7701605 | -0.876314908 | 0.438977699 | -1.996262933 | 0.04590531 | 0.246869858 |
| GPR160 | 166.6651462 | -0.177973296 | 0.73475252 | -0.242222097 | 0.808608074 | 0.919641737 |
| GPR161 | 793.9471741 | 0.107759945 | 0.362922422 | 0.29692281 | 0.766525441 | 0.903470079 |
| GPR162 | 1015.980632 | 0.167626151 | 0.347852461 | 0.481888645 | 0.629885053 | 0.83795038 |
| GPR173 | 507.5248537 | -0.133033982 | 0.34830399 | -0.381947912 | 0.702500001 | 0.872703476 |
| GPR176 | 332.5034243 | 1.014545513 | 0.579636333 | 1.750313869 | 0.080064169 | 0.323994001 |
| GPR180 | 396.9089923 | 0.400440536 | 0.275920096 | 1.451291668 | 0.14669866 | 0.439661276 |
| GPR19 | 463.0705208 | 0.522087216 | 0.27972876 | 1.866405213 | 0.061984694 | 0.286960573 |
| GPR26 | 11.29024177 | -0.358928804 | 0.941414622 | -0.38126538 | 0.70300634 | 0.872703476 |
| GPR27 | 192.55883 | 1.504034868 | 0.420252594 | 3.578883013 | 0.000345066 | 0.013776301 |
| GPR3 | 56.79953689 | 0.174371246 | 0.440865589 | 0.3955202 | 0.692459017 | 0.868421571 |
| GPR35 | 21.75959057 | 0.516965189 | 0.707592128 | 0.730597711 | 0.46502491 | 0.742073516 |
| GPR37 | 279.5924934 | -1.622607294 | 0.439493557 | -3.691993358 | 0.000222503 | 0.010167462 |
| GPR39 | 29.2401075 | -1.787717453 | 0.726370358 | -2.461165208 | 0.013848658 | 0.133991065 |
| GPR61 | 18.23912548 | 0.641231424 | 0.860733888 | 0.744982198 | 0.456282482 | 0.73543231 |
| GPR63 | 349.9075932 | -0.160933647 | 0.359776227 | -0.447315957 | 0.654646947 | 0.849149183 |
| GPR75 | 21.32431149 | -1.816987992 | 0.777341901 | -2.337437348 | 0.019416455 | 0.158029528 |
| GPR78 | 133.3113175 | -2.313470637 | 0.542211659 | -4.266729787 | 1.98E-05 | 0.001857856 |
| GPR83 | 33.58880156 | -2.331582036 | 0.81295347 | -2.868038727 | 0.00413025 | 0.066306898 |
| GPR85 | 34.39782024 | -0.617714335 | 0.726834453 | -0.849869365 | 0.39539772 | 0.69426198 |
| GPR87 | 35.80143987 | 0.417472743 | 0.857656719 | 0.486759719 | 0.626428619 | 0.835767952 |
| GPR89A | 498.6939646 | 0.125824351 | 0.265289921 | 0.474289979 | 0.635293137 | 0.840574932 |

| | | | | | | |
|---|---|---|---|---|---|---|
| GPR89B | 645.7802865 | -0.147020621 | 0.322150083 | -0.456373066 | 0.648121731 | 0.846620869 |
| GPRASP1 | 133.5619772 | 0.285476527 | 0.571444094 | 0.499570352 | 0.617377638 | 0.831835425 |
| GPRASP2 | 475.4682621 | 0.423066002 | 0.367158454 | 1.152270898 | 0.249209772 | 0.562027865 |
| GPRC5A | 25.33915605 | -0.986676407 | 0.851245311 | -1.159097612 | 0.246416399 | 0.558551201 |
| GPRC5B | 2027.385932 | 0.310296283 | 0.356363138 | 0.870730584 | 0.383901275 | 0.684699412 |
| GPRC5C | 592.8104841 | 0.055638914 | 0.426198436 | 0.130546968 | 0.896133697 | 0.958964824 |
| GPRIN1 | 160.6087466 | -0.287233186 | 0.36518513 | -0.786541298 | 0.431550435 | 0.718193467 |
| GPRIN2 | 65.08422898 | 0.169623763 | 0.502185338 | 0.337771238 | 0.735535586 | 0.889855747 |
| GPRIN3 | 29.60291937 | -2.575212203 | 0.898057666 | -2.867535461 | 0.004136824 | 0.066342245 |
| GPS1 | 2188.374894 | -0.284034107 | 0.382240544 | -0.743076868 | 0.457435143 | 0.73623473 |
| GPS2 | 562.269263 | -0.296060627 | 0.302762196 | -0.977865238 | 0.32814098 | 0.637787615 |
| GPSM1 | 309.4157762 | 0.303657646 | 0.525327978 | 0.578034406 | 0.563240891 | 0.801890319 |
| GPSM2 | 544.1248343 | 0.846324401 | 0.428491236 | 1.975126512 | 0.048253793 | 0.253279062 |
| GPSM3 | 142.2766541 | 2.263481517 | 0.857728513 | 2.638925351 | 0.008316929 | 0.103338357 |
| GPT2 | 1698.483494 | -0.098615804 | 0.269226711 | -0.366292793 | 0.714146607 | 0.878410749 |
| GPX1 | 2032.082492 | 1.107200486 | 0.441225338 | 2.509376484 | 0.01209445 | 0.124947219 |
| GPX2 | 14.11926117 | -0.517253089 | 1.053471588 | -0.490998613 | 0.623427428 | 0.834261049 |
| GPX3 | 770.6436928 | -0.204087309 | 0.28276444 | -0.721757335 | 0.470443684 | 0.7454951 |
| GPX4 | 4005.291419 | 0.930832402 | 0.433013767 | 2.149660063 | 0.031582114 | 0.202592072 |
| GPX7 | 251.0559092 | 0.971155291 | 0.408302369 | 2.378519852 | 0.017382301 | 0.148986942 |
| GPX8 | 1597.740316 | 0.19173855 | 0.274571326 | 0.698319644 | 0.484977317 | 0.755353112 |
| GRAMD1A | 1677.095504 | 0.028017383 | 0.366101716 | 0.076528958 | 0.938998276 | 0.977796709 |
| GRAMD1B | 390.6346906 | 0.30473969 | 0.550310061 | 0.553759982 | 0.579743119 | 0.811340629 |
| GRAMD1C | 284.4171468 | -1.096154237 | 0.426440844 | -2.570471974 | 0.010156004 | 0.11438511 |
| GRAMD2A | 149.2183495 | -1.861678944 | 0.597559796 | -3.115468874 | 0.001836527 | 0.039859736 |
| GRAMD2B | 564.6404046 | -0.86944832 | 0.466246729 | -1.864781598 | 0.062212024 | 0.287486632 |
| GRAMD4 | 790.4194168 | -0.343438237 | 0.365522638 | -0.939581305 | 0.34743237 | 0.654038526 |
| GRAMD4P3 | 10.66199929 | 1.848047629 | 1.121481583 | 1.647862664 | 0.099380856 | 0.361473741 |
| GRASP | 109.8824484 | 1.573710177 | 0.514563107 | 3.05834242 | 0.002225651 | 0.044607685 |
| GRB10 | 1131.513237 | 0.425192703 | 0.408562882 | 1.04070321 | 0.298013313 | 0.609375237 |
| GRB14 | 305.9636497 | 0.226406383 | 0.472975677 | 0.478685046 | 0.632162706 | 0.838768903 |
| GRB2 | 2820.282799 | 0.017892792 | 0.236882611 | 0.075534424 | 0.939789509 | 0.978003717 |
| GRB7 | 585.2415091 | -1.291743887 | 0.50331279 | -2.566483334 | 0.010273556 | 0.11507205 |
| GREB1 | 945.044102 | 0.461103018 | 0.532235591 | 0.866351341 | 0.38629752 | 0.686679504 |
| GREB1L | 8454.615011 | -0.461382929 | 0.376542062 | -1.225315776 | 0.220456267 | 0.529032271 |
| GREM1 | 362.7515035 | -0.447881653 | 0.585216062 | -0.765327 | 0.44407685 | 0.727127105 |
| GRHL1 | 1186.996099 | -2.821923126 | 0.752274681 | -3.75118716 | 0.000175999 | 0.008728094 |
| GRHL2 | 2287.702988 | -2.428368447 | 0.567290095 | -4.280646655 | 1.86E-05 | 0.001766957 |

| | | | | | | |
|---|---|---|---|---|---|---|
| GRHL3 | 3389.943047 | -0.951761788 | 0.435600801 | -2.1849404 | 0.028893217 | 0.194385362 |
| GRHPR | 2152.527649 | 0.36931921 | 0.281271059 | 1.313036655 | 0.189170597 | 0.49447056 |
| GRIA3 | 151.6574892 | -0.050908832 | 0.51492992 | -0.098865555 | 0.921245018 | 0.970205128 |
| GRID2 | 68.73435662 | -1.130109833 | 0.495797083 | -2.279379753 | 0.022644501 | 0.171427007 |
| GRID2IP | 32.93538621 | -0.36696621 | 0.540546536 | -0.678879958 | 0.497213925 | 0.761481169 |
| GRIK3 | 14.6000296 | -0.040425862 | 0.963777115 | -0.041945239 | 0.966542352 | 0.988700292 |
| GRIK4 | 15.52158283 | 0.143904429 | 0.903373094 | 0.159296785 | 0.873435054 | 0.949679868 |
| GRIK5 | 629.7825705 | 0.139533867 | 0.419803964 | 0.332378631 | 0.739603372 | 0.89168179 |
| GRIN2A | 12.5155324 | -0.875548501 | 1.583687955 | -0.552854177 | 0.580363267 | 0.811716708 |
| GRIN2B | 12.21186691 | 1.173786461 | 0.945675666 | 1.241214619 | 0.214526476 | 0.521483923 |
| GRIN2D | 93.74801019 | 0.171908564 | 0.610713559 | 0.281488042 | 0.778336099 | 0.908108663 |
| GRIN3B | 12.26903134 | 0.150226799 | 1.08094998 | 0.138976643 | 0.889468606 | 0.956341292 |
| GRINA | 1701.316876 | -0.194516451 | 0.272994659 | -0.712528411 | 0.476137624 | 0.749324056 |
| GRIP1 | 362.3472136 | -0.790791863 | 0.469583572 | -1.684027958 | 0.092176266 | 0.348555864 |
| GRIP2 | 67.50720939 | -0.301438254 | 0.582206116 | -0.517751782 | 0.604631465 | 0.825894731 |
| GRIPAP1 | 1378.427596 | -0.39526561 | 0.284861677 | -1.387570329 | 0.165267922 | 0.467517155 |
| GRK2 | 905.6203292 | 0.047828038 | 0.469459919 | 0.101878853 | 0.918852836 | 0.969677217 |
| GRK3 | 1187.038948 | -0.340533955 | 0.344249199 | -0.98920769 | 0.322561537 | 0.632245616 |
| GRK4 | 73.25521428 | 0.182500826 | 0.462206637 | 0.394846831 | 0.692955931 | 0.86854217 |
| GRK5 | 295.9927959 | -0.04121739 | 0.39191989 | -0.105167896 | 0.916242584 | 0.969184139 |
| GRK6 | 959.1789224 | 0.133077376 | 0.332855466 | 0.399805289 | 0.689299935 | 0.867579108 |
| GRM1 | 35.89783697 | -0.310799133 | 0.575176171 | -0.540354675 | 0.588952459 | 0.818050255 |
| GRM2 | 85.95255297 | -1.053771839 | 0.413663079 | -2.547415742 | 0.010852406 | 0.118362219 |
| GRM4 | 43.1386757 | -2.058565329 | 0.577243959 | -3.566196402 | 0.0003622 | 0.01427256 |
| GRM8 | 36.99942511 | 0.255996062 | 0.600449248 | 0.426340882 | 0.669859473 | 0.857762448 |
| GRN | 2710.07181 | -0.722185184 | 0.365946694 | -1.973470992 | 0.048441926 | 0.253705068 |
| GRPEL1 | 625.3070732 | 0.584429177 | 0.271976923 | 2.148818988 | 0.031648752 | 0.202592072 |
| GRPEL2 | 685.1812794 | 0.447417395 | 0.313682119 | 1.426340131 | 0.153770186 | 0.450608457 |
| GRPR | 27.72675281 | -6.010536308 | 1.879074035 | -3.198669237 | 0.001380635 | 0.033674613 |
| GRSF1 | 3749.275641 | 0.216432954 | 0.315379682 | 0.686261563 | 0.492548182 | 0.759174768 |
| GRTP1 | 365.9421964 | 0.166317949 | 0.360833315 | 0.460927364 | 0.644850719 | 0.845794147 |
| GRWD1 | 762.795887 | 0.254296807 | 0.286610443 | 0.887255901 | 0.374941142 | 0.67920525 |
| GSAP | 27.40511471 | -1.57485042 | 0.752064542 | -2.09403626 | 0.03625675 | 0.218014728 |
| GSDMB | 571.1040468 | -0.672234495 | 0.324218921 | -2.073396866 | 0.03813536 | 0.224762556 |
| GSDMD | 94.87924897 | 0.400648662 | 0.537457238 | 0.745452166 | 0.455998418 | 0.735399427 |
| GSDME | 631.7030165 | -1.111956733 | 0.39932561 | -2.784586579 | 0.005359601 | 0.077809098 |
| GSE1 | 2762.87478 | -0.875351957 | 0.237470325 | -3.686153031 | 0.000227669 | 0.010319181 |
| GSG1 | 12.04215205 | 0.705896947 | 0.899672537 | 0.784615422 | 0.432679085 | 0.718639059 |
| GSK3A | 1949.290352 | -0.034851552 | 0.260131179 | -0.133976836 | 0.893420892 | |

0.95780189
GSK3B 2192.467257 -0.385753178 0.253402828 -1.522292317 0.127935852
0.412171335
GSKIP 1211.295385 0.196246471 0.25770324 0.761521165 0.446345842 0.728589711
GSN 2912.860481 -0.188643802 0.33867884 -0.55699908 0.577528058
0.809916636
GSPT1 4299.176055 0.161817723 0.26458036 0.611601418 0.5408015 0.787864037
GSPT2 886.4103851 0.085618307 0.238126013 0.359550417 0.719183368 0.881260702
GSR 1317.593719 0.840971139 0.347435713 2.420508624 0.015498812 0.140797887
GSS 1202.507243 0.168796411 0.245113182 0.688646809 0.491045556 0.758593654
GSTA4 1515.667541 0.039853484 0.284377388 0.140142944 0.888547051 0.955997805
GSTCD 965.5246967 -0.33583841 0.325930669 -1.030398308 0.302823067
0.613538084
GSTK1 1716.760664 -0.563519663 0.460943826 -1.222534355 0.22150562
0.530599081
GSTM1 363.9458027 0.298684514 0.285894871 1.044735476 0.296145244 0.607412916
GSTM2 701.6345294 0.060651569 0.257325379 0.235699912 0.813665538 0.921959809
GSTM3 144.7034238 0.880180504 0.432794871 2.033712881 0.041980553 0.235820549
GSTM4 423.3741538 -0.04773798 0.350677098 -0.136130873 0.891717821
0.957210151
GSTM5 65.27867492 1.00508425 0.742731633 1.353226663 0.17598323 0.480071547
GSTO1 1845.560363 0.056733026 0.639381097 0.088731159 0.929295569 0.973104851
GSTO2 192.7572876 0.108675447 0.47026976 0.231091718 0.817243549 0.923820833
GSTP1 15388.00284 0.382355241 0.352888918 1.083500278 0.278586428 0.592103488
GSTT2B 169.6730437 -1.070628337 0.656958995 -1.629673002
0.103170626 0.368199757
GSTZ1 1420.897959 0.438005103 0.429612884 1.019534375 0.307949343 0.61773099
GTDC1 566.7640744 0.387089219 0.330919006 1.169740063 0.242105591 0.553231259
GTF2A1 1478.517173 0.118669795 0.269837329 0.439782722 0.660094483
0.852207761
GTF2A2 1415.869887 1.083428752 0.356291752 3.040847129 0.002359136
0.046602147
GTF2B 636.0210013 0.252865917 0.276160527 0.915648298 0.359851391 0.667315175
GTF2E1 934.9763266 -0.039788453 0.247478514 -0.160775385
0.872270313 0.949093661
GTF2E2 700.8402494 0.537968286 0.277136716 1.941165693 0.052238183
0.263465915
GTF2F1 2602.163426 -0.459400478 0.29760646 -1.543650897
0.122672928 0.403440492
GTF2F2 1517.924782 0.766204703 0.366482292 2.090700476 0.036554922
0.218932905
GTF2H1 719.6251817 -0.213505336 0.292501532 -0.729928949
0.465433616 0.742335301
GTF2H2 522.4600969 0.303296296 0.348379942 0.870590581 0.383977741
0.684699412
GTF2H2B 482.2186435 0.029327159 0.317828105 0.092273649 0.926480624
0.972261514
GTF2H2C 205.5980181 -0.47070086 0.519572301 -0.905939094 0.364968091
0.671631599
GTF2H3 1250.000789 0.129453187 0.264199325 0.489983035 0.624145904
0.834707114
GTF2H4 544.5717658 0.029227126 0.321626335 0.090872925 0.927593564
0.972633911
GTF2H5 1117.660818 0.986666575 0.402562766 2.450963326 0.014247447
0.135535502
GTF2I 5777.413969 -0.320180619 0.619009692 -0.517246537 0.60498407
0.82602102
GTF2IP1 4773.204227 0.011559908 0.399142105 0.028961885 0.976894989
0.992055261
GTF2IP12 291.4405114 -1.208811214 0.519156195 -2.328415274
0.019890063 0.159572793
GTF2IP13 110.6708297 -1.552366237 0.467346748 -3.321658365
0.000894842 0.025375039
GTF2IP20 212.1401106 -0.790670468 0.369146347 -2.141888914

| | | | | | | |
|---|---|---|---|---|---|---|
| | | | | | 0.032202422 | 0.204582474 |
| GTF2IP4 | 4989.193207 | -0.833667013 | 0.302087239 | -2.759689604 | 0.00578563 | 0.081954993 |
| GTF2IP7 | 20.90015559 | -1.095288257 | 0.889707932 | -1.231064958 | 0.21829857 | 0.526623336 |
| GTF2IRD1 | 1026.51995 | 0.078385737 | 0.272852267 | 0.287282705 | 0.773895865 | 0.906248714 |
| GTF2IRD2 | 161.0063974 | -0.274830129 | 0.403877619 | -0.68047873 | 0.496201385 | 0.761056737 |
| GTF2IRD2B | 155.3244654 | -0.288583523 | 0.438572112 | -0.658007007 | 0.510533629 | 0.770330936 |
| GTF3A | 2883.019554 | 0.205465764 | 0.291822843 | 0.704077044 | 0.481384796 | 0.752733727 |
| GTF3C1 | 3055.935474 | -0.383404983 | 0.514145686 | -0.745712729 | 0.455840968 | 0.735306344 |
| GTF3C2 | 2644.822895 | -0.348693739 | 0.285629981 | -1.220788299 | 0.222166185 | 0.531066116 |
| GTF3C3 | 2410.256936 | -0.140677248 | 0.341260772 | -0.412228008 | 0.68017231 | 0.862002279 |
| GTF3C4 | 2214.870666 | -0.085797413 | 0.283423816 | -0.302717725 | 0.762104988 | 0.900880067 |
| GTF3C5 | 2354.488621 | 0.347127085 | 0.266969939 | 1.300247833 | 0.193516041 | 0.499461017 |
| GTF3C6 | 1332.932271 | 0.772713048 | 0.501501292 | 1.540799716 | 0.123365539 | 0.404753154 |
| GTPBP1 | 1731.539043 | 0.072001936 | 0.339223148 | 0.212255374 | 0.831907809 | 0.931142584 |
| GTPBP10 | 630.1037868 | 0.257954132 | 0.262576489 | 0.982396152 | 0.325904721 | 0.635827259 |
| GTPBP2 | 1561.346843 | -0.1029636 | 0.2592661 | -0.397134835 | 0.691268033 | 0.868004248 |
| GTPBP3 | 624.2952041 | 0.416720058 | 0.283094317 | 1.47201845 | 0.141015895 | 0.430713136 |
| GTPBP4 | 2604.090755 | 0.355436808 | 0.302008043 | 1.176911728 | 0.23923074 | 0.55048833 |
| GTPBP6 | 745.9959408 | 0.249390421 | 0.498175084 | 0.500607977 | 0.616647048 | 0.831694047 |
| GTPBP8 | 495.8287464 | 0.441332579 | 0.315954112 | 1.396824926 | 0.162466225 | 0.463216519 |
| GTSE1 | 896.3290649 | 0.764466564 | 0.319665533 | 2.391457583 | 0.016781623 | 0.146402531 |
| GUCA1A | 299.5942807 | -0.717413408 | 0.576152916 | -1.245178821 | 0.213066026 | 0.520189768 |
| GUCA1B | 62.30222097 | 1.400464935 | 0.728131168 | 1.923369026 | 0.054433721 | 0.269716333 |
| GUCD1 | 1009.119449 | -0.333817259 | 0.356258995 | -0.937007243 | 0.348754832 | 0.655415363 |
| GUCY1A1 | 394.7332031 | -2.327954074 | 0.475658187 | -4.894174299 | 9.87E-07 | 0.000180442 |
| GUCY1B1 | 394.5741445 | -0.47649808 | 0.417778445 | -1.140552095 | 0.254056362 | 0.566474638 |
| GUCY1B2 | 10.88011209 | 0.196488748 | 0.88597467 | 0.221776936 | 0.824487535 | 0.92681808 |
| GUF1 | 1806.890106 | -0.136125298 | 0.295743404 | -0.46028177 | 0.645313984 | 0.845794147 |
| GUK1 | 1180.85685 | 0.534625348 | 0.479019189 | 1.116083364 | 0.264386452 | 0.577921357 |
| GULP1 | 2294.956593 | -0.272883286 | 0.290939284 | -0.937938949 | 0.348275786 | 0.654732583 |
| GUSB | 1591.123519 | 0.009877476 | 0.243415451 | 0.040578673 | 0.967631786 | 0.989014473 |
| GUSBP1 | 104.6756787 | -0.353223438 | 0.463810213 | -0.761568908 | 0.446317337 | 0.728589711 |
| GUSBP2 | 48.01090337 | -0.12437418 | 0.596077641 | -0.208654329 | 0.834718096 | 0.932107564 |
| GUSBP5 | 28.76551651 | -1.009021226 | 0.613331792 | -1.645147437 | 0.09993941 | 0.362399544 |
| GUSBP9 | 64.56492922 | -0.488744382 | 0.703316036 | -0.694914316 | | |

| | | | | | | |
|---|---|---|---|---|---|---|
| | | | | | 0.487109001 | 0.756439734 |
| GVINP1 | 15.87283027 | -1.763533962 | 0.920290392 | -1.916279881 | 0.05532948 | 0.272191885 |
| GVQW2 | 25.60403138 | 0.063399945 | 0.667549741 | 0.094974113 | 0.924335389 | 0.971733919 |
| GVQW3 | 217.5419975 | 0.0293092 | 0.439759752 | 0.066648209 | 0.946861766 | 0.980602079 |
| GXYLT1 | 958.7835124 | -0.571064213 | 0.342016938 | -1.669695707 | 0.094979584 | 0.353777379 |
| GXYLT2 | 129.5707765 | 0.69671514 | 0.580586904 | 1.200018697 | 0.230132079 | 0.53903563 |
| GYG1 | 765.1127479 | 0.037007688 | 0.271084309 | 0.136517263 | 0.891412379 | 0.957017706 |
| GYG2 | 1052.164389 | -0.210101927 | 0.315366171 | -0.666215802 | 0.505273174 | 0.766779966 |
| GYG2P1 | 61.4086395 | 0.302130514 | 0.600700388 | 0.50296374 | 0.614989766 | 0.830872437 |
| GYPC | 726.6098265 | 1.007603293 | 0.38846865 | 2.593782775 | 0.009492644 | 0.110354716 |
| GYS1 | 1302.781812 | -0.579229444 | 0.289733767 | -1.999178243 | 0.045589072 | 0.245771986 |
| GZF1 | 402.4425113 | 0.065463762 | 0.276142409 | 0.237065222 | 0.812606192 | 0.921654347 |
| H1F0 | 10481.29139 | -0.019473587 | 0.281054952 | -0.06928747 | 0.944760799 | 0.979553348 |
| H1FX | 2394.100757 | 0.358741277 | 0.433268481 | 0.827988401 | 0.407677068 | 0.702187648 |
| H2AFJ | 464.9656095 | 0.150876869 | 0.592030755 | 0.254846336 | 0.798841786 | 0.916056735 |
| H2AFV | 3854.016273 | 0.926665502 | 0.358866323 | 2.582202458 | 0.009817197 | 0.112405058 |
| H2AFX | 1086.570992 | 1.412435025 | 0.366025526 | 3.858842961 | 0.000113925 | 0.006725128 |
| H2AFY | 3289.665526 | 0.312973326 | 0.250395369 | 1.249916588 | 0.211330019 | 0.518628728 |
| H2AFY2 | 3713.813101 | 0.154584879 | 0.325911076 | 0.474316125 | 0.635274494 | 0.840574932 |
| H2AFZ | 9653.135786 | 1.357685657 | 0.258805667 | 5.245965709 | 1.55E-07 | 5.13E-05 |
| H2BFM | 65.26515306 | 0.864324553 | 0.470130277 | 1.83847881 | 0.065991881 | 0.296552972 |
| H3F3A | 26110.24562 | 0.71182773 | 0.408139675 | 1.74407874 | 0.081145361 | 0.325762441 |
| H3F3AP4 | 87.25625635 | 1.053479834 | 3.829868701 | 0.275069439 | 0.783262891 | 0.910216876 |
| H3F3B | 15061.618 | 0.290600394 | 0.371288954 | 0.78267988 | 0.43381512 | 0.719273746 |
| H6PD | 495.5869711 | -0.079334745 | 0.410696413 | -0.193171265 | 0.846824842 | 0.938145276 |
| HAAO | 210.3168074 | -1.375402918 | 0.522521072 | -2.632243927 | 0.008482294 | 0.104029814 |
| HABP4 | 658.0570016 | 0.379984241 | 0.277682154 | 1.368414339 | 0.171182418 | 0.474474162 |
| HACD1 | 214.2442386 | 1.15819244 | 0.49843734 | 2.323647019 | 0.02014442 | 0.160509978 |
| HACD2 | 1270.776152 | -0.183723694 | 0.256465334 | -0.716368529 | 0.473763822 | 0.747915811 |
| HACD3 | 6082.920712 | 0.479978479 | 0.285084522 | 1.6836357 | 0.092252095 | 0.348755677 |
| HACD4 | 210.563349 | -0.80591318 | 0.357111738 | -2.256753542 | 0.024023479 | 0.177207433 |
| HACE1 | 1034.436242 | -0.229558624 | 0.330660858 | -0.694241906 | 0.487530516 | 0.756439734 |
| HACL1 | 939.3207618 | 0.059529055 | 0.267710365 | 0.222363653 | 0.824030804 | 0.926576588 |
| HADH | 1889.16784 | 0.11722888 | 0.256582499 | 0.456885721 | 0.647753188 | 0.846597949 |
| HADHA | 4089.294815 | -0.189571241 | 0.314206478 | -0.603333331 | 0.546286964 | 0.791946443 |
| HADHB | 1401.707959 | 0.143721822 | 0.340105545 | 0.422580063 | 0.672601675 | 0.859011927 |
| HAGH | 558.1462961 | 0.610766794 | 0.37879337 | 1.61240096 | 0.106874716 | 0.374802662 |
| HAGHL | 41.82181536 | 1.631289184 | 0.570536819 | 2.859218072 | 0.004246867 | 0.067465155 |
| HAND1 | 1330.739523 | -4.754860331 | 0.884398162 | -5.376379708 | 7.60E-08 | 3.20E-05 |
| HAP1 | 88.8821714 | 0.570055936 | 0.523979839 | 1.087934866 | 0.276623863 | 0.590171608 |
| HAPLN1 | 10249.25095 | -1.132617858 | 0.702609601 | -1.612015915 | 0.106958477 | 0.37492307 |
| HAPLN3 | 112.3406063 | 0.452163535 | 0.422113874 | 1.071188517 | 0.284084674 | 0.598230674 |
| HAPLN4 | 120.3335344 | -0.173697415 | 0.424982238 | -0.408716882 | 0.682747443 | 0.863109561 |
| HARBI1 | 107.4858396 | -0.475142117 | 0.381946781 | -1.24400084 | 0.213499254 | 0.520678417 |

| | | | | | | |
|---|---|---|---|---|---|---|
| HARS | 1478.98483 | -0.295349355 | 0.272842365 | -1.082490822 | 0.279034492 | 0.592557709 |
| HARS2 | 1228.818801 | -0.189552286 | 0.297721802 | -0.636675866 | 0.524335997 | 0.777662198 |
| HAS1 | 21.32690375 | -1.771867868 | 1.159464413 | -1.528177879 | 0.126468379 | 0.409268725 |
| HAS2 | 312.7284294 | -1.312455957 | 0.517177373 | -2.537728883 | 0.011157438 | 0.120212934 |
| HAS3 | 1019.650534 | -1.991638625 | 0.611173733 | -3.258711097 | 0.001119196 | 0.028974947 |
| HASPIN | 258.9890933 | -0.078581551 | 0.425949321 | -0.184485683 | 0.853632458 | 0.941030229 |
| HAT1 | 1954.284407 | 0.590292446 | 0.269709497 | 2.188623144 | 0.028624242 | 0.193363702 |
| HAUS1 | 1905.746177 | 0.754731587 | 0.261463459 | 2.886566214 | 0.003894708 | 0.064617764 |
| HAUS2 | 1047.38001 | -0.020415773 | 0.233088087 | -0.08758823 | 0.930203957 | 0.973652838 |
| HAUS3 | 544.4083848 | 0.296270898 | 0.256245579 | 1.156199064 | 0.247599743 | 0.560334446 |
| HAUS4 | 1658.049189 | 0.140732314 | 0.238459564 | 0.590172653 | 0.555074904 | 0.796360966 |
| HAUS5 | 758.3457123 | 0.027356206 | 0.327534511 | 0.083521597 | 0.933436805 | 0.974750122 |
| HAUS6 | 1927.80388 | 0.299660047 | 0.307183206 | 0.975509213 | 0.329307734 | 0.638448972 |
| HAUS6P3 | 17.92726194 | 0.487833595 | 0.77732725 | 0.627578146 | 0.530280328 | 0.780679069 |
| HAUS7 | 276.8948148 | 0.713226689 | 0.304537246 | 2.342001506 | 0.019180638 | 0.157365202 |
| HAUS8 | 384.9942802 | 0.250497519 | 0.257838365 | 0.971529272 | 0.331284784 | 0.639138284 |
| HAX1 | 1639.988827 | 0.474662453 | 0.279731174 | 1.69685218 | 0.089724611 | 0.343025354 |
| HBEGF | 30.78658748 | 0.875452562 | 0.59072965 | 1.481985138 | 0.13834425 | 0.426606546 |
| HBP1 | 493.5439687 | -0.441596173 | 0.290953408 | -1.517755629 | 0.129076015 | 0.413453061 |
| HBS1L | 2169.430727 | 0.139895878 | 0.233598546 | 0.598873069 | 0.549257531 | 0.79348329 |
| HCAR1 | 15.02137172 | -2.337968827 | 1.258435024 | -1.857838332 | 0.063191976 | 0.289778852 |
| HCCS | 618.1880878 | 0.173518844 | 0.338321919 | 0.512880882 | 0.608034647 | 0.827607804 |
| HCFC1 | 2499.165051 | 0.046155206 | 0.276724143 | 0.166791396 | 0.867534189 | 0.946451976 |
| HCFC1R1 | 699.8785648 | 0.917573087 | 0.424665269 | 2.160697271 | 0.030718732 | 0.199843002 |
| HCFC2 | 120.4083633 | 0.212352142 | 0.490475715 | 0.43295139 | 0.665050085 | 0.855040241 |
| HCK | 16.49407274 | 0.198620583 | 0.820531212 | 0.24206341 | 0.80873103 | 0.919641737 |
| HCN1 | 16.5987467 | -2.905169401 | 1.177914727 | -2.466366481 | 0.01364916 | 0.132885181 |
| HCN2 | 12.57108763 | -0.775405517 | 0.869799137 | -0.891476531 | 0.372673577 | 0.678079975 |
| HCN3 | 14.3638489 | -0.872906626 | 1.194955518 | -0.730492987 | 0.465088898 | 0.742097567 |
| HCN4 | 103.6403295 | -0.842244393 | 0.399173069 | -2.109972989 | 0.034860683 | 0.213176891 |
| HDAC1 | 4591.188338 | -0.109535402 | 0.248356753 | -0.441040563 | 0.659183631 | 0.851684114 |
| HDAC10 | 282.6456817 | -0.364028195 | 0.391171653 | -0.930609856 | 0.352055414 | 0.659305356 |
| HDAC11 | 278.1199892 | 0.203923741 | 0.411566165 | 0.495482278 | 0.620259729 | 0.833216571 |
| HDAC2 | 10392.91635 | 0.471276148 | 0.24812693 | 1.89933494 | 0.057520452 | 0.277823232 |
| HDAC3 | 1983.951786 | 0.311360015 | 0.24983538 | 1.246260698 | 0.212668702 | 0.519735803 |
| HDAC4 | 358.9019816 | 0.544899105 | 0.390942551 | 1.393808639 | 0.163375399 | 0.464631632 |
| HDAC5 | 1841.050339 | -0.186990855 | 0.321076046 | -0.582388058 | 0.560305323 | 0.800514963 |
| HDAC6 | 2375.493904 | -0.25835083 | 0.442385508 | -0.583994787 | 0.559223819 | 0.799470841 |
| HDAC7 | 1206.695686 | -0.037549133 | 0.305961941 | -0.12272485 | 0.902324986 | 0.962670348 |
| HDAC8 | 694.1387495 | -0.381887939 | 0.254151542 | -1.502599341 | 0.132942393 | 0.419138571 |
| HDAC9 | 340.0332033 | 1.028227056 | 0.55264252 | 1.860564504 | 0.062805701 | 0.289348708 |
| HDC | 10.81665073 | -1.190197667 | 0.958292273 | -1.241998606 | 0.214237076 | |

| | | | | | | |
|---|---|---|---|---|---|---|
| | | | | | | 0.521197993 |
| HDDC2 | 1587.983729 | 0.997487222 | 0.299219716 | 3.333627995 | 0.000857212 | 0.024770984 |
| HDDC3 | 443.2864833 | 0.454475412 | 0.394983406 | 1.150619001 | 0.249889013 | 0.562742997 |
| HDGF | 12904.39403 | 0.324848202 | 0.235436331 | 1.379770916 | 0.16765719 | 0.46989234 |
| HDGFL2 | 2490.276776 | 0.132861983 | 0.374750848 | 0.354534175 | 0.722938597 | 0.883138856 |
| HDGFL3 | 2951.395804 | 0.520863501 | 0.241502586 | 2.156761591 | 0.031024245 | 0.200870165 |
| HDHD2 | 1424.739418 | 0.403111821 | 0.276405693 | 1.458406362 | 0.144728568 | 0.435928729 |
| HDHD3 | 602.0513065 | -0.099666255 | 0.465424267 | -0.214140651 | 0.830437376 | 0.930264006 |
| HDHD5 | 854.6736723 | 0.78521535 | 0.33576248 | 2.338603618 | 0.019355957 | 0.157622736 |
| HDLBP | 12170.99165 | -0.876495196 | 0.417622372 | -2.098774527 | 0.035836781 | 0.216490931 |
| HDX | 207.3327379 | -0.196294468 | 0.298076764 | -0.658536629 | 0.510193368 | 0.769949736 |
| HEATR1 | 2284.673538 | -0.455440672 | 0.35257775 | -1.291745358 | 0.196445344 | 0.502167241 |
| HEATR3 | 646.2971835 | 0.625890285 | 0.312075369 | 2.005574126 | 0.044901705 | 0.2438968 |
| HEATR5A | 831.4826781 | -0.538179686 | 0.265893441 | -2.024042726 | 0.042965756 | 0.238392703 |
| HEATR5B | 961.3785969 | -0.836627758 | 0.470403703 | -1.778531402 | 0.075316619 | 0.314600339 |
| HEATR6 | 697.9393815 | -0.334063769 | 0.407093181 | -0.820607625 | 0.411869803 | 0.704847916 |
| HEBP1 | 1034.95202 | 0.693509753 | 0.508355573 | 1.364221797 | 0.17249777 | 0.476244526 |
| HEBP2 | 1767.794684 | 0.082945343 | 0.526410363 | 0.157567838 | 0.874797349 | 0.949740014 |
| HECA | 413.2000051 | -0.112002535 | 0.468948 | -0.238837855 | 0.811231315 | 0.920897407 |
| HECTD1 | 2919.180704 | -0.21186662 | 0.371188514 | -0.57077903 | 0.56814944 | 0.805325156 |
| HECTD2 | 655.3530007 | 0.604549181 | 0.522948705 | 1.156039159 | 0.24766514 | 0.560334446 |
| HECTD3 | 1478.986823 | -0.706288451 | 0.274570951 | -2.572334939 | 0.010101511 | 0.114259456 |
| HECTD4 | 1162.517786 | -0.693499337 | 0.442145491 | -1.568486734 | 0.116767584 | 0.39401268 |
| HECW1 | 18.13864464 | 0.032042274 | 0.843170787 | 0.038002117 | 0.969685995 | 0.989496332 |
| HECW2 | 501.0508041 | -1.634722666 | 0.38101806 | -4.290407302 | 1.78E-05 | 0.001712458 |
| HEG1 | 394.1440456 | -0.198198396 | 0.57929612 | -0.342136585 | 0.732248106 | 0.888075467 |
| HELB | 190.3289662 | -0.071142715 | 0.392353173 | -0.181323154 | 0.856113933 | 0.942245037 |
| HELLS | 1966.666708 | 0.158904368 | 0.311261249 | 0.510517672 | 0.609688837 | 0.828409093 |
| HELQ | 369.2292042 | 0.139751211 | 0.289731762 | 0.48234688 | 0.62955955 | 0.837590803 |
| HELZ | 1014.44422 | -0.709509749 | 0.32781854 | -2.164336855 | 0.030438506 | 0.199044214 |
| HELZ2 | 231.1333799 | 0.449688852 | 0.513943029 | 0.874978017 | 0.381585867 | 0.683026643 |
| HEMK1 | 691.2528001 | -0.358975431 | 0.274419799 | -1.308125117 | 0.190830875 | 0.496058205 |
| HENMT1 | 132.1716435 | -0.058152769 | 0.384893215 | -0.15108806 | 0.879906251 | 0.952211837 |
| HEPH | 3957.087228 | -0.863900036 | 0.227368359 | -3.799561377 | 0.000144952 | 0.007733542 |
| HERC1 | 1263.35324 | -0.311992499 | 0.549154338 | -0.568132632 | 0.569944911 | 0.806205765 |
| HERC2 | 1212.821751 | -0.242502916 | 0.454659432 | -0.533372673 | 0.593775631 | 0.820230051 |
| HERC2P2 | 74.87971234 | -0.902388248 | 0.459158257 | -1.965309858 | 0.049378385 | 0.25614346 |
| HERC2P3 | 299.3885994 | -0.100521223 | 2.352091259 | -0.042736957 | 0.965911219 | 0.988700292 |
| HERC2P9 | 146.8816999 | -0.673012104 | 0.703241248 | -0.957014547 | | |

0.338559912 0.645678097
HERC3 90.50082884 -1.321443709 0.482978656 -2.736029206 0.006218551
0.085454377
HERC4 1557.637755 -0.304667156 0.228905014 -1.330976333 0.183196799
0.487250812
HERC5 205.7394143 0.074017586 0.337904626 0.219048751 0.826612072 0.92789728
HERC6 32.84489626 -2.703167912 0.804884828 -3.358453055 0.0007838
0.023439632
HERPUD1 824.0002723 -1.631975102 0.649855109 -2.511290718
0.012029059 0.124561748
HERPUD2 875.8682706 -0.173337513 0.282049118 -0.614564989
0.538842042 0.786598616
HES1 1044.122042 -0.974911452 0.801476265 -1.216394664 0.223834616
0.533008155
HES2 219.2116837 0.24394918 0.676706716 0.360494694 0.718477224 0.880880715
HES3 73.74698945 3.035617124 1.257334832 2.414326754 0.015764318 0.142219415
HES4 35.40998902 1.564051012 0.770121268 2.030915229 0.042263595 0.236528876
HES6 210.6925064 1.856717924 0.461308109 4.02489765 5.70E-05 0.00411785
HESX1 29.15895539 0.18778739 0.63717115 0.294720484 0.768207414 0.904321493
HEXA 1443.850446 -0.279087141 0.275479313 -1.013096547 0.311014047
0.621086496
HEXB 1335.150656 0.437237501 0.328281331 1.33189877 0.182893455 0.486816961
HEXD 414.6455566 -0.23610014 0.446421214 -0.528873028 0.596893533
0.821812117
HEXIM1 1961.849294 -0.409060196 0.264198752 -1.548304799
0.12154893 0.402062246
HEXIM2 185.4074061 0.308608171 0.48400063 0.637619358 0.523721492
0.777662198
HEY1 370.6868572 -3.63783932 1.423032346 -2.5563996 0.010576159 0.116946722
HEY2 286.6981769 -1.050808944 0.492243336 -2.134734727 0.032782685
0.206453345
HEYL 90.05520317 0.540525476 0.690819026 0.782441501 0.433955152 0.719296316
HFE 49.46357705 -0.253300709 0.510738321 -0.495950075 0.619929635
0.832886335
HFM1 159.9899841 0.129285951 0.367783238 0.351527577 0.725192587 0.884183617
HGF 43.85925127 -1.432869674 0.675074363 -2.122536052 0.033792749
0.209476397
HGH1 360.0780301 0.214176042 0.295463317 0.724882007 0.46852442 0.744337782
HGS 1541.089178 -0.379040372 0.409876333 -0.924767645 0.355086784
0.662528791
HGSNAT 740.7917057 -1.309514826 0.386953518 -3.384165707
0.000713949 0.022014879
HHAT 135.4170752 -0.240981826 0.37877838 -0.636207974 0.524640876
0.777662198
HHIP 236.3006369 -4.161477124 0.924714481 -4.500283285 6.79E-06
0.000817102
HHIPL1 126.8153213 -0.476563801 0.357513103 -1.33299674 0.182532874
0.486683209
HHIPL2 99.92974891 -0.867897186 0.510938218 -1.698634307
0.089388108 0.342726775
HHLA1 995.7592972 -0.607906796 0.66018603 -0.92081136 0.357148923
0.664659754
HHLA2 14.40332697 2.018557061 1.098169801 1.838110152 0.066046175 0.29670907
HHLA3 271.9133528 0.60506881 0.538856536 1.122875515 0.261490368 0.574937735
HIBADH 1605.469672 0.318022067 0.285766523 1.11287377 0.265762647
0.579572403
HIBCH 896.9939726 0.410448691 0.309980623 1.324110802 0.185466249 0.490031772
HIC2 1910.3166 -0.987474996 0.316785306 -3.117174242 0.001825936
0.039686644
HID1 703.1000425 -2.17023764 0.576857467 -3.762173093 0.000168443
0.008574622
HIF1A 4969.266082 -0.233176691 0.292862845 -0.796197589 0.425917199
0.71395509
HIF1AN 2326.263451 -0.540069361 0.311817377 -1.732005335

0.083272613 0.330368412
HIF3A 421.5890735 0.174696802 0.341802018 0.511105237 0.609277369 0.8283007
HIGD1A 2510.615783 0.631670309 0.30502463 2.07088296 0.038369735
0.225585886
HIGD2A 1350.801515 0.631790059 0.465506813 1.357209051 0.174714796
0.47862011
HIKESHI 1283.170744 0.870522835 0.341024426 2.552670043 0.010690071
0.117606289
HILPDA 751.027298 0.522913238 0.373096663 1.401548953 0.16104998
0.46167973
HINFP 857.4417894 -0.14499262 0.239948787 -0.60426486 0.545667565 0.791502307
HINT1 6928.303861 1.069255931 0.422655816 2.529850275 0.011411121 0.121146338
HINT2 574.4918851 1.056293075 0.425036209 2.48518374 0.012948458 0.128476822
HINT3 772.3504317 -0.117400854 0.273829277 -0.428737406 0.668114335
0.856874003
HIP1 2454.777873 -0.154516008 0.465770705 -0.331742649 0.740083593
0.891917214
HIP1R 508.5612586 -0.074427688 0.343964326 -0.216381999 0.828689997
0.929129846
HIPK1 1816.428284 -0.508665551 0.33117125 -1.53595927 0.12454837
0.405999855
HIPK2 1399.910469 -0.295879639 0.269301133 -1.098694372 0.271901398
0.584197156
HIPK3 534.6479569 -0.717869187 0.401270564 -1.788990402 0.073616361
0.311651561
HIRA 1406.64545 -0.319019206 0.354047164 -0.901064148 0.367554214
0.673938238
HIRIP3 293.3203839 0.25002236 0.372265393 0.671623966 0.501823117
0.764473972
HIST1H1A 18.51170556 2.244188483 0.850538029 2.638551606 0.008326103
0.103367679
HIST1H1C 107.5841351 1.102629712 0.503411513 2.190314852 0.02850141
0.192869196
HIST1H1D 12.15324415 1.351757939 0.857820204 1.575805667 0.115070596
0.390283034
HIST1H2AC 79.97391702 -0.025441089 0.677829929 -0.037533145
0.970059913 0.989496332
HIST1H2AG 28.59025361 0.977985825 0.591214756 1.654197251 0.098087441
0.359755003
HIST1H2BD 33.67005312 0.70201826 0.699554902 1.003521321 0.315609395
0.625406234
HIST1H2BE 12.93470882 0.554926073 0.87594926 0.633513948 0.526398076
0.778525486
HIST1H2BG 25.18839325 0.704106002 0.834842553 0.843399752 0.399004911
0.697352242
HIST1H2BH 72.18521714 0.961471806 0.463196446 2.075732262 0.03791872
0.223664427
HIST1H2BK 38.29620532 0.395286858 0.695562483 0.56829813 0.569832549
0.806129205
HIST1H2BN 22.32918825 1.569044958 0.672277041 2.333926138 0.01959959
0.158583796
HIST1H3E 28.09052692 -0.438719288 0.703350823 -0.623755989
0.532787847 0.782698212
HIST1H3G 13.44440366 0.792138683 0.874808934 0.905499078 0.365201048
0.671700076
HIST1H3H 17.49012985 0.893171932 0.831438862 1.074248477 0.28271133
0.596532022
HIST1H4C 23.90186829 1.510763265 0.872474894 1.731583654 0.083347719
0.330579932
HIST1H4E 13.0154383 1.272948594 0.864186971 1.473001373 0.140750666
0.43068341
HIST1H4H 14.59724395 0.907969532 0.890898901 1.019161131 0.308126476
0.617841234
HIST1H4J 77.50760527 1.122904064 0.594002739 1.890402167 0.058704194

0.280504354
HIST2H2AA3 13.73371445 1.165815219 1.598428706 0.729350777 0.465787119
0.742664886
HIST2H2AA4 15.26876405 0.520178775 1.909176713 0.272462351 0.785266536
0.910931322
HIST2H2BD 83.96632563 0.198439317 0.457954209 0.43331694 0.664784533
0.855037526
HIST2H2BE 65.43504291 -1.291998656 0.714112752 -1.80923622 0.070414313
0.304780468
HIST2H4A 48.85353519 -1.33557215 0.740720707 -1.803071167 0.071377039
0.307350563
HIST2H4B 87.8349783 0.929841443 0.457257811 2.033516803 0.042000338
0.235820549
HIST3H2A 59.24768383 0.972798675 0.546005912 1.781663262 0.074804161
0.313842348
HIST4H4 18.32571463 -1.902045755 1.099134789 -1.730493634
0.083542118 0.330745685
HIVEP1 401.2960057 -0.442173238 0.384525682 -1.149918612
0.250177394 0.562905438
HIVEP2 567.6096153 -1.503932523 0.331476943 -4.537065255 5.70E-06
0.00070357
HIVEP3 27.41887477 0.10631567 0.598182331 0.177731211 0.858934073
0.94255977
HJURP 1185.962488 0.867486215 0.40368147 2.148937417 0.031639361 0.202592072
HK1 2822.110464 -0.74059125 0.23530345 -3.147387977 0.001647362
0.037682927
HK2 1546.878914 0.287174775 0.294275299 0.975871154 0.329128318 0.638424836
HKR1 1208.676399 -0.421937574 0.252995827 -1.667764957 0.095362395
0.354701001
HLA-A 13.17803036 0.979216459 1.005657123 0.973708073 0.330201508 0.63872078
HLA-B 51.02279117 -0.141825678 0.590147203 -0.240322546 0.810080218
0.920303055
HLA-C 26.55427856 1.225396555 3.490567585 0.351059398 0.725543788 0.88422336
HLA-DOA 309.1586431 -0.19323123 0.301857192 -0.640141217 0.522080795
0.777127918
HLA-DPB2 18.12790831 0.111218563 0.963720085 0.115405463 0.908123749
0.965583536
HLA-DQB1 218.2616176 -0.560458988 0.503233088 -1.11371649 0.265400833
0.579420929
HLA-E 882.4250507 -1.533890129 0.487723057 -3.145002287 0.001660855
0.037719814
HLA-L 12.67954653 -3.397660833 1.749137215 -1.942478156 0.052079242
0.262949685
HLCS 448.8821892 -0.209631899 0.418864744 -0.500476353 0.616739703
0.831694047
HLTF 1160.143164 0.795915555 0.415654749 1.914847739 0.055511924 0.272635611
HM13 1976.532258 0.371995975 0.247625568 1.502251883 0.13303207 0.419328805
HMBOX1 468.691166 0.184854542 0.389490394 0.474606166 0.635067711
0.840423259
HMBS 619.8241036 0.691651639 0.306584487 2.255990333 0.024071237 0.177207433
HMCES 1334.758388 0.354568218 0.325629699 1.088869408 0.276211478 0.589533531
HMCN1 434.180849 -0.698705874 0.527573451 -1.324376487 0.18537804
0.490031772
HMCN2 149.2300584 -1.085138593 0.47818433 -2.26928932 0.023250738
0.174130017
HMG20A 2021.889792 -0.121218622 0.335623658 -0.361174247
0.717969192 0.880671367
HMG20B 1362.485278 0.344907495 0.421306999 0.818660728 0.412980009
0.705607651
HMGA1 36123.70467 0.145610542 0.226368765 0.64324485 0.520065235 0.775216335
HMGA2 13653.42367 0.266412498 0.3198578 0.83290918 0.404895952 0.70065889
HMGB1 17829.77326 0.712082973 0.293990699 2.422127554 0.015429933 0.140677297
HMGB1P5 2268.071328 0.558395868 0.252622336 2.210397853 0.027077563
0.188125165

| | | | | | | |
|---|---|---|---|---|---|---|
| HMGB1P6 | 775.7460457 | 0.573148958 | 0.371054092 | 1.544650687 | 0.122430779 | 0.403241325 |
| HMGB2 | 11131.86769 | 1.346113506 | 0.25037971 | 5.376288307 | 7.60E-08 | 3.20E-05 |
| HMGB3 | 4858.306784 | 0.047734663 | 0.35633938 | 0.133958428 | 0.893435448 | 0.95780189 |
| HMGCL | 943.6064594 | 0.307822746 | 0.349937049 | 0.879651774 | 0.379047982 | 0.681253679 |
| HMGCLL1 | 22.84161884 | 0.890030504 | 0.791127102 | 1.125015819 | 0.260582331 | 0.573939394 |
| HMGCR | 5463.868555 | 0.257659784 | 0.404049754 | 0.637693208 | 0.523673407 | 0.777662198 |
| HMGCS1 | 4383.494601 | 0.416980187 | 0.379622114 | 1.098408578 | 0.272026118 | 0.584230595 |
| HMGN1 | 14679.15725 | 0.556305989 | 0.388984039 | 1.430151199 | 0.152673628 | 0.448798819 |
| HMGN2 | 9657.53385 | 0.895106725 | 0.29462921 | 3.038078687 | 0.002380918 | 0.046910264 |
| HMGN3 | 3679.051484 | 0.596954781 | 0.403083759 | 1.480969569 | 0.138614683 | 0.427045019 |
| HMGN4 | 2107.10489 | 0.52035498 | 0.261215862 | 1.9920497 | 0.046365605 | 0.247615641 |
| HMGN5 | 515.9918009 | 0.944007894 | 0.405636716 | 2.327224968 | 0.019953295 | 0.159698109 |
| HMGXB3 | 1580.626673 | -0.265616507 | 0.363999899 | -0.729715882 | 0.465563871 | 0.742464994 |
| HMGXB4 | 1597.020001 | 0.176992116 | 0.247097516 | 0.716284481 | 0.473815707 | 0.747919894 |
| HMMR | 1891.709669 | 0.898764884 | 0.439293845 | 2.045930975 | 0.040763162 | 0.231983403 |
| HMOX1 | 400.0262077 | -0.923917536 | 0.605838281 | -1.525023368 | 0.127253269 | 0.410838472 |
| HMOX2 | 374.6835579 | 0.378393524 | 0.278434447 | 1.359003986 | 0.174145326 | 0.478455902 |
| HMSD | 11.68516424 | -0.724589865 | 1.060513611 | -0.683244286 | 0.494452493 | 0.760405536 |
| HNF1B | 14.28695504 | 0.882753647 | 1.235172812 | 0.714680276 | 0.474806628 | 0.748737095 |
| HNF4G | 10.61999124 | -0.048376761 | 1.201482636 | -0.04026422 | 0.967882479 | 0.989137426 |
| HNMT | 158.85732 | -1.024894285 | 0.580540026 | -1.76541537 | 0.077493982 | 0.319212925 |
| HNRNPA0 | 8247.039174 | 0.431614233 | 0.323180699 | 1.335519832 | 0.181706269 | 0.48591777 |
| HNRNPA1 | 97847.24423 | -0.019761227 | 0.271698085 | -0.072732301 | 0.942019144 | 0.978574897 |
| HNRNPA1L2 | 250.2270581 | -0.1673483 | 0.286052645 | -0.585026229 | 0.558530081 | 0.798855459 |
| HNRNPA1P10 | 25.65988975 | -1.378332816 | 0.685298022 | -2.011289647 | 0.044294871 | 0.2418127 |
| HNRNPA1P16 | 40.16996229 | 0.194443834 | 0.498245251 | 0.390257275 | 0.696346313 | 0.870023227 |
| HNRNPA1P48 | 224.9752219 | 0.948954159 | 0.570124508 | 1.664468279 | 0.096018885 | 0.355648968 |
| HNRNPA1P54 | 24.22641648 | -0.376112907 | 0.746869456 | -0.503585874 | 0.614552422 | 0.830593641 |
| HNRNPA1P7 | 12.19051283 | -0.452180541 | 0.954828525 | -0.47357251 | 0.635804782 | 0.841016203 |
| HNRNPA2B1 | 39970.25985 | 0.216217323 | 0.225951259 | 0.956920196 | 0.338607537 | 0.645678097 |
| HNRNPA3 | 17751.23171 | 0.078093117 | 0.259542026 | 0.300888138 | 0.763499796 | 0.902107118 |
| HNRNPA3P6 | 63.62426048 | -1.511323844 | 0.641711385 | -2.355145754 | 0.018515442 | 0.154302741 |
| HNRNPAB | 6366.399968 | 0.356481133 | 0.284472903 | 1.25312861 | 0.210158906 | 0.51816316 |
| HNRNPC | 30773.92372 | 0.425442827 | 0.248815364 | 1.709873619 | 0.087289246 | 0.33929417 |
| HNRNPD | 13760.25778 | 0.357585323 | 0.31612976 | 1.131134644 | 0.257998425 | 0.570734048 |
| HNRNPDL | 13267.09718 | 0.250297537 | 0.221206616 | 1.131510178 | 0.257840422 | 0.570550911 |
| HNRNPF | 7997.647271 | -0.177927075 | 0.240077597 | -0.741123193 | 0.458618748 | 0.736888903 |
| HNRNPH1 | 14038.03138 | -0.175510926 | 0.260751555 | -0.673096374 | 0.500885981 | 0.763789447 |

| | | | | | | |
|---|---|---|---|---|---|---|
| HNRNPH2 | 3340.078012 | -0.05914814 | 0.282211831 | -0.209587741 | 0.833989448 | 0.932107564 |
| HNRNPH3 | 8502.686525 | 0.337471533 | 0.249329402 | 1.353516796 | 0.175890588 | 0.480020886 |
| HNRNPK | 21011.62922 | -0.131440705 | 0.221495337 | -0.593424252 | 0.552897263 | 0.795672359 |
| HNRNPL | 6492.676709 | 0.113619005 | 0.286539352 | 0.396521471 | 0.691720372 | 0.86812349 |
| HNRNPLL | 1695.029622 | 0.389981963 | 0.244373806 | 1.595841919 | 0.110524092 | 0.38264742 |
| HNRNPM | 9532.355118 | -0.008750088 | 0.316436809 | -0.02765193 | 0.977939764 | 0.99226352 |
| HNRNPR | 4536.38124 | 0.172831902 | 0.243392154 | 0.710096439 | 0.477644335 | 0.750061298 |
| HNRNPU | 19531.93484 | -0.106862285 | 0.229662418 | -0.465301573 | 0.641715508 | 0.844038972 |
| HNRNPUL1 | 6625.390143 | -0.325435077 | 0.286396121 | -1.136311052 | 0.255826407 | 0.567999769 |
| HNRNPUL2 | 3027.23303 | -0.19985047 | 0.260167932 | -0.768159506 | 0.442392428 | 0.725806805 |
| HNRNPUL2-BSCL2 | 1662.196071 | -0.994303649 | 2.577931572 | -0.385698232 | 0.699720173 | 0.871449506 |
| HOGA1 | 15.58650681 | 1.299573783 | 0.770013038 | 1.687729582 | 0.09146315 | 0.347000104 |
| HOMER1 | 1350.629308 | 0.350833449 | 0.336310383 | 1.043183518 | 0.296863305 | 0.608198948 |
| HOMER2 | 212.6231403 | -0.757266483 | 0.413768784 | -1.83016823 | 0.067224787 | 0.297874489 |
| HOMER3 | 265.4221207 | 0.728051078 | 0.365862921 | 1.989955898 | 0.046595794 | 0.248472685 |
| HOMEZ | 639.0747727 | 0.359146019 | 0.29192339 | 1.230274899 | 0.218594181 | 0.527064894 |
| HOOK1 | 775.7820433 | -0.416535654 | 0.33765216 | -1.233623542 | 0.217343216 | 0.525468356 |
| HOOK2 | 640.178124 | -0.750728499 | 0.335044498 | -2.240682967 | 0.025046618 | 0.181116418 |
| HOOK3 | 1120.704702 | 0.001679241 | 0.3002898 | 0.005592069 | 0.995538197 | 0.998116797 |
| HOPX | 130.2584339 | 1.891369345 | 1.073934315 | 1.761159242 | 0.078211454 | 0.320774797 |
| HOXA1 | 696.1031553 | -1.570787075 | 0.724739702 | -2.16738102 | 0.030205813 | 0.198239795 |
| HOXA2 | 177.900813 | 2.236316941 | 0.841471308 | 2.657627088 | 0.007869291 | 0.099653606 |
| HOXA3 | 104.9009724 | 0.32865064 | 0.808662931 | 0.406412397 | 0.684439613 | 0.863857185 |
| HOXB1 | 263.8941052 | 0.001708085 | 1.556863659 | 0.001097132 | 0.999124616 | 0.999324484 |
| HOXB2 | 355.4934269 | 0.293901638 | 1.446358754 | 0.203201064 | 0.838977881 | 0.933915137 |
| HOXB3 | 56.695145 | -3.460251787 | 1.470143129 | -2.353683609 | 0.018588425 | 0.154438663 |
| HOXB4 | 10.37968718 | -4.583004406 | 2.085893467 | -2.197142126 | 0.028010297 | 0.191330131 |
| HOXB5 | 16.45367916 | -6.656197561 | 4.40959158 | -1.509481647 | 0.131175742 | 0.416813082 |
| HOXD1 | 42.07992275 | 2.108271784 | 0.960819819 | 2.194242605 | 0.028217973 | 0.192229399 |
| HOXD3 | 32.52499616 | 1.701331509 | 1.450105446 | 1.173246755 | 0.240696867 | 0.552353982 |
| HP1BP3 | 3907.375061 | -0.115178818 | 0.358106975 | -0.321632435 | 0.747731169 | 0.894202292 |
| HPCA | 15.8684362 | 0.332684419 | 0.749876413 | 0.443652331 | 0.657293964 | 0.850401393 |
| HPCAL1 | 183.8059807 | 0.182125444 | 0.429472773 | 0.424067497 | 0.671516589 | 0.858479663 |
| HPD | 37.18196943 | -2.126651549 | 0.602843114 | -3.527703145 | 0.000419182 | 0.015778519 |
| HPDL | 466.5530069 | 0.267633081 | 0.387224028 | 0.691158249 | 0.489466098 | 0.757800814 |
| HPF1 | 1092.454247 | 0.807746723 | 0.320690054 | 2.518776967 | 0.011776323 | 0.123127904 |
| HPGD | 597.7790038 | -2.551093454 | 0.886739019 | -2.876938307 | 0.004015541 | 0.065575642 |
| HPN | 280.8419646 | -0.215489043 | 0.795663883 | -0.270829238 | 0.786522371 | 0.911315996 |
| HPRT1 | 537.1361879 | 1.295223637 | 0.396550414 | 3.266226921 | 0.001089909 | 0.028511711 |

| | | | | | | |
|---|---|---|---|---|---|---|
| HPS1 | 1505.242521 | -0.291485637 | 0.317475529 | -0.918135764 | 0.358547793 | 0.666363907 |
| HPS3 | 1151.94249 | -0.205257197 | 0.299340602 | -0.685697814 | 0.492903685 | 0.759389046 |
| HPS4 | 1382.311908 | -0.228269847 | 0.346548292 | -0.658695636 | 0.510091236 | 0.769949736 |
| HPS5 | 435.5708182 | -0.147050649 | 0.364417324 | -0.403522664 | 0.686563765 | 0.865679761 |
| HPS6 | 409.5384446 | 0.208549619 | 0.415283634 | 0.502185982 | 0.615536703 | 0.830968556 |
| HPSE | 38.05720297 | -0.233762855 | 0.543220104 | -0.430328062 | 0.666957016 | 0.856258348 |
| HPSE2 | 47.2173019 | -0.414567946 | 0.771319725 | -0.537478729 | 0.590936974 | 0.818805811 |
| HPX | 26.43107635 | -0.1280816 | 0.594482931 | -0.215450425 | 0.82941616 | 0.929737887 |
| HRAS | 585.7387758 | 0.540188273 | 0.428508526 | 1.260624331 | 0.207444228 | 0.514322011 |
| HRASLS | 43.62630058 | 0.557836867 | 0.474681153 | 1.175182252 | 0.239921809 | 0.551576567 |
| HRASLS5 | 102.362956 | -2.848687516 | 0.562252507 | -5.066562585 | 4.05E-07 | 0.000107811 |
| HRC | 44.88021525 | 0.196109742 | 0.867057184 | 0.226178556 | 0.821062548 | 0.925158936 |
| HRH1 | 14.3090812 | -3.27775696 | 1.30329217 | -2.51498247 | 0.01190383 | 0.123863518 |
| HRH2 | 21.13222217 | -2.050581554 | 0.953977916 | -2.149506314 | 0.031594286 | 0.202592072 |
| HRH3 | 14.03843154 | 1.775564851 | 0.907603842 | 1.956321437 | 0.0504273 | 0.258046297 |
| HRK | 71.47077956 | -0.915229262 | 0.452425454 | -2.022939367 | 0.043079399 | 0.238611743 |
| HS1BP3 | 748.4401087 | -0.206481638 | 0.300790543 | -0.686463198 | 0.492421064 | 0.759174768 |
| HS2ST1 | 1200.031741 | 0.139665971 | 0.309194357 | 0.451709314 | 0.651478407 | 0.847876685 |
| HS3ST1 | 22.25218321 | 3.044961135 | 0.907710403 | 3.354551325 | 0.000794938 | 0.023647069 |
| HS3ST3A1 | 14.74232082 | -0.358425959 | 0.97452232 | -0.367796563 | 0.713024932 | 0.877743193 |
| HS3ST3B1 | 141.5057787 | -0.145466551 | 0.573643429 | -0.253583573 | 0.799817288 | 0.916311991 |
| HS3ST5 | 55.79445739 | -1.969597928 | 0.609174064 | -3.233226829 | 0.001224003 | 0.030831576 |
| HS6ST1 | 1241.716641 | 0.194291898 | 0.301153316 | 0.645159419 | 0.518823881 | 0.774467046 |
| HS6ST2 | 781.1043424 | 0.317436286 | 0.360858928 | 0.87966865 | 0.379038837 | 0.681253679 |
| HS6ST3 | 56.31725165 | -2.443843622 | 0.683656396 | -3.574666507 | 0.000350675 | 0.013963476 |
| HSBP1 | 5633.714257 | 0.919771708 | 0.315581854 | 2.914526598 | 0.003562285 | 0.061273717 |
| HSBP1L1 | 149.5533393 | -0.345895558 | 0.681604561 | -0.507472481 | 0.61182334 | 0.829098987 |
| HSCB | 323.845555 | 0.845934047 | 0.450333182 | 1.878462617 | 0.060317906 | 0.283945763 |
| HSD11B1L | 282.7510501 | 1.167005611 | 0.44625161 | 2.615129187 | 0.008919371 | 0.10668882 |
| HSD11B2 | 31.80998253 | 0.952777999 | 0.642070716 | 1.483914427 | 0.137831624 | 0.425968529 |
| HSD17B1 | 39.51730523 | 1.007331368 | 0.588917692 | 1.71047904 | 0.087177326 | 0.339206775 |
| HSD17B10 | 2061.775701 | 0.593347507 | 0.38491229 | 1.541513541 | 0.12319185 | 0.404708435 |
| HSD17B11 | 920.1736753 | 0.24958922 | 0.283771052 | 0.879544332 | 0.379106207 | 0.681253679 |
| HSD17B12 | 2538.755443 | 0.116613921 | 0.299558681 | 0.389285733 | 0.697064789 | 0.870026319 |
| HSD17B14 | 325.4205632 | 0.325901255 | 0.433687083 | 0.751466363 | 0.452372037 | 0.732985143 |
| HSD17B2 | 17.75255416 | 0.276877287 | 0.904236229 | 0.30620017 | 0.759452243 | |

0.899496446
HSD17B4 3737.986515 0.232710118 0.296166343 0.785741271 0.432019079
0.71850132
HSD17B6 66.06123076 0.917586582 0.493859365 1.85799166 0.063170199
0.289778852
HSD17B7 438.1893034 0.273553262 0.283721265 0.964161998 0.334964713
0.643096642
HSD17B7P2 73.77357621 -0.020875671 0.498927263 -0.041841111
0.966625362 0.988718539
HSD17B8 128.6527314 1.098758823 0.357374349 3.074531863 0.002108332 0.0433918
HSD3B7 97.91858401 -0.913410751 0.532962498 -1.713836816 0.0865587
0.337492171
HSDL1 1835.7816 0.340534827 0.336377518 1.012359058 0.311366407 0.621452782
HSDL2 1002.702798 0.114066379 0.262384239 0.434730299 0.663758203 0.854667747
HSF1 625.9627328 -0.359071329 0.376775745 -0.953010733 0.340584632
0.647332681
HSF2 1066.91275 0.602508311 0.294077603 2.048807198 0.040480972 0.231601821
HSF2BP 27.60957141 -0.140613584 0.697450439 -0.201610861
0.840220953 0.934461703
HSFX1 12.59006814 0.366633564 0.872287848 0.42031259 0.674257114 0.859672768
HSP90AA1 46441.02911 0.466644391 0.282591855 1.651301634 0.098676999
0.360796043
HSP90AB1 70096.15602 0.158471124 0.252864481 0.626703775 0.530853429
0.781218098
HSP90B1 24115.65219 0.006117699 0.252544709 0.024224221 0.980673758
0.993241851
HSPA12A 214.7706176 0.005479017 0.380070807 0.014415778 0.988498271
0.995803391
HSPA12B 29.22595765 0.183014439 0.691762474 0.264562542 0.791346479
0.913361935
HSPA13 1457.86718 0.515251304 0.367606816 1.401636971 0.161023681
0.46167973
HSPA14 1576.90479 0.243119675 0.236750778 1.026901274 0.304466954
0.614891928
HSPA1B 19.10871218 -5.101758882 4.365163644 -1.168744015
0.242506783 0.553576648
HSPA2 186.926407 -0.155044887 0.389776793 -0.397778652 0.690793355
0.868004248
HSPA4 8239.324577 -0.104540631 0.263578336 -0.396620728 0.691647165
0.86812349
HSPA4L 184.5926519 -0.117993165 0.328255683 -0.359455055
0.719254695 0.881264172
HSPA5 9360.390799 -0.30960917 0.317840602 -0.974102013 0.33000589
0.638681113
HSPA8 22993.60773 -0.014587921 0.347532305 -0.041975727 0.966518048
0.988700292
HSPA9 7176.349496 0.068775562 0.238298644 0.2886108 0.77287923 0.905717252
HSPB1 469.0681786 0.957151508 0.436287528 2.193854846 0.028245846 0.192252052
HSPB11 537.3308198 0.77597659 0.317220508 2.446174098 0.014438126
0.136559111
HSPB2 24.07541879 0.785171118 0.681925197 1.151403586 0.249566239 0.562413758
HSPB6 29.32158428 0.034436022 0.681260036 0.050547545 0.959686062 0.986008211
HSPB8 25.04978235 -1.709112321 1.21694589 -1.404427539 0.160191575
0.460500386
HSPBAP1 425.4133408 -0.05366514 0.274984565 -0.195156918 0.845270105
0.937778401
HSPBP1 757.6670567 0.723882437 0.402686643 1.7976321 0.072235328
0.30937269
HSPD1 18663.71436 0.388853075 0.252733592 1.538588801 0.123904714 0.405045568
HSPD1P6 61.59820086 -0.960043894 0.700183262 -1.37113231 0.170333714
0.473284392
HSPE1 4706.151366 1.389293473 0.371023876 3.744485361 0.000180764 0.008851388
HSPE1-MOB4 187.7970982 -0.666812642 0.536359251 -1.243220174
0.21378671 0.520770421

| | | | | | | |
|---|---|---|---|---|---|---|
| HSPG2 | 4062.004018 | -1.095832821 | 0.511090143 | -2.144108697 | 0.032024178 | 0.204048219 |
| HSPH1 | 2819.757807 | 0.143362556 | 0.288767315 | 0.49646393 | 0.61956713 | 0.832768045 |
| HTATIP2 | 1006.494829 | 0.347280957 | 0.365477358 | 0.950211962 | 0.342004562 | 0.648704346 |
| HTATSF1 | 3614.970166 | 0.103233778 | 0.260641965 | 0.396075047 | 0.692049666 | 0.86812349 |
| HTD2 | 204.477009 | -0.520804267 | 0.510393042 | -1.020398447 | 0.30753953 | 0.617319594 |
| HTR1B | 18.63946559 | 1.25149907 | 1.396785113 | 0.895985401 | 0.37026056 | 0.675958117 |
| HTR1D | 306.1697659 | 0.938692849 | 0.43468651 | 2.159470853 | 0.030813656 | 0.200374616 |
| HTR3A | 35.58842438 | 0.235470234 | 0.604494844 | 0.389532245 | 0.696882462 | 0.870026319 |
| HTR7 | 30.8613384 | -0.449667819 | 0.559958911 | -0.803037169 | 0.421953251 | 0.712063713 |
| HTR7P1 | 90.78618448 | -0.11442345 | 0.430997359 | -0.265485268 | 0.790635657 | 0.91311918 |
| HTRA1 | 237.6662988 | -1.503507877 | 0.630966877 | -2.382863397 | 0.017178568 | 0.147741524 |
| HTRA2 | 819.6953529 | 0.289387667 | 0.387740125 | 0.74634439 | 0.455459403 | 0.735238838 |
| HTRA3 | 47.20972892 | -0.302928816 | 0.606484198 | -0.499483445 | 0.617438847 | 0.831835425 |
| HTRA4 | 14.32036483 | -1.565518615 | 0.867892274 | -1.803816744 | 0.07126004 | 0.306992555 |
| HTT | 2724.861286 | -0.50169168 | 0.422707183 | -1.186853929 | 0.235285243 | 0.545881517 |
| HUNK | 840.8036313 | -1.557327426 | 0.591439871 | -2.633112005 | 0.008460644 | 0.103848246 |
| HUS1 | 373.5954054 | -0.128420021 | 0.278657501 | -0.460852554 | 0.644904394 | 0.845794147 |
| HUWE1 | 9780.072042 | -0.912047872 | 0.411850199 | -2.21451361 | 0.026793467 | 0.186982725 |
| HVCN1 | 95.75081202 | 1.017397426 | 0.57547489 | 1.767926704 | 0.077073158 | 0.31844659 |
| HYAL1 | 63.16064453 | -0.131323256 | 0.567458033 | -0.231423733 | 0.816985629 | 0.923735281 |
| HYAL2 | 1398.101579 | 0.679286 | 0.270712106 | 2.509256084 | 0.012098574 | 0.124947219 |
| HYAL3 | 54.66423494 | -0.102355152 | 0.503962192 | -0.203100855 | 0.839056203 | 0.933915137 |
| HYDIN | 36.80968971 | 1.109109939 | 0.687231248 | 1.613881706 | 0.106553087 | 0.374540521 |
| HYDIN2 | 38.67544582 | -0.970127443 | 0.568095354 | -1.707684171 | 0.087694961 | 0.340260934 |
| HYI | 535.5972034 | -0.381797606 | 0.399222601 | -0.956352684 | 0.338894081 | 0.645980916 |
| HYKK | 59.702762 | 0.055211827 | 0.565081684 | 0.097705921 | 0.922165814 | 0.970640219 |
| HYLS1 | 193.1000356 | 0.601411003 | 0.297132976 | 2.024046648 | 0.042965352 | 0.238392703 |
| HYOU1 | 3593.842311 | -0.175014823 | 0.345489667 | -0.506570354 | 0.612456314 | 0.829234693 |
| HYPK | 200.80431 | 0.224138089 | 0.568520883 | 0.394247767 | 0.693398122 | 0.868737544 |
| IAH1 | 224.0322872 | 0.384036464 | 0.412761454 | 0.93040777 | 0.352159997 | 0.659419811 |
| IARS | 5836.00098 | -0.284473531 | 0.314433386 | -0.904717958 | 0.365614825 | 0.672153785 |
| IARS2 | 4182.602418 | 0.023475237 | 0.254688025 | 0.092172521 | 0.92656097 | 0.972261514 |
| IBA57 | 334.2494349 | 0.068444452 | 0.29587475 | 0.231329142 | 0.817059109 | 0.923735281 |
| IBTK | 430.8719484 | -0.021326486 | 0.346446691 | -0.061557771 | 0.950915007 | 0.981826318 |
| ICA1 | 319.8285211 | -0.606254479 | 0.591238528 | -1.025397451 | 0.305175691 | 0.615135352 |
| ICA1L | 130.9176251 | -0.086705711 | 0.375395853 | -0.230971414 | 0.81733701 | 0.923820833 |
| ICAM1 | 44.28587249 | 0.567669557 | 0.616896342 | 0.920202501 | 0.357466948 | 0.665006875 |
| ICAM3 | 56.39361095 | -0.731833006 | 0.507115819 | -1.443127937 | 0.148984435 | 0.443544075 |
| ICE1 | 2191.550953 | -0.415815104 | 0.386250469 | -1.076542653 | 0.281684636 | 0.595435086 |
| ICE2 | 2238.719672 | 0.149139613 | 0.289582226 | 0.515016458 | 0.606541519 | 0.826611695 |

| | | | | | | |
|---|---|---|---|---|---|---|
| ICK | 657.434479 | -0.478829147 | 0.369495196 | -1.295900871 | 0.195009639 | 0.500590454 |
| ICMT | 3257.319386 | 0.162139812 | 0.331874331 | 0.488557857 | 0.625154755 | 0.835033444 |
| ICOSLG | 211.286455 | -1.027082737 | 0.562823815 | -1.824874339 | 0.068019994 | 0.300067268 |
| ID1 | 2651.313662 | 0.115268736 | 0.636688012 | 0.181044301 | 0.856332803 | 0.942245037 |
| ID2 | 6258.772605 | 0.129085812 | 0.390568964 | 0.330507091 | 0.741016835 | 0.891943909 |
| ID3 | 1008.715003 | 1.499310689 | 0.465407857 | 3.221498448 | 0.001275222 | 0.031663479 |
| ID4 | 2799.433625 | -0.476001412 | 0.503557646 | -0.945276902 | 0.344517528 | 0.651368096 |
| IDE | 1661.456732 | -0.301604321 | 0.286924344 | -1.05116323 | 0.293183628 | 0.60622718 |
| IDH1 | 4460.896445 | 0.330819009 | 0.233540511 | 1.416537998 | 0.156618047 | 0.4554442 |
| IDH2 | 2340.341088 | 0.79285961 | 0.321786094 | 2.463933726 | 0.013742152 | 0.13330063 |
| IDH3A | 308.6208891 | 0.11694829 | 0.350475001 | 0.333685111 | 0.738617186 | 0.89135397 |
| IDH3B | 2193.050731 | -0.067680778 | 0.255575293 | -0.264817372 | 0.791150153 | 0.913361935 |
| IDH3G | 974.4126056 | -0.10398568 | 0.234828646 | -0.442815143 | 0.657899449 | 0.850760404 |
| IDI1 | 1819.55625 | 0.833362978 | 0.349995971 | 2.381064487 | 0.01726269 | 0.14821294 |
| IDNK | 48.95808779 | -0.066983934 | 0.668650195 | -0.100177842 | 0.920203137 | 0.969770994 |
| IDS | 1566.453236 | -0.168840672 | 0.316957923 | -0.532691121 | 0.594247413 | 0.820642877 |
| IDUA | 15.13811184 | 0.61995431 | 0.923863555 | 0.671045315 | 0.502191662 | 0.764661936 |
| IER2 | 2682.448094 | -0.429613231 | 0.613393432 | -0.700387726 | 0.483685199 | 0.754278091 |
| IER3IP1 | 918.8395716 | 0.855177016 | 0.323380194 | 2.644494101 | 0.008181313 | 0.101820101 |
| IER5 | 2048.925851 | 0.487536828 | 0.451939589 | 1.078765481 | 0.280692287 | 0.594538239 |
| IER5L | 413.6605281 | 1.123962985 | 0.483771636 | 2.323333785 | 0.020161228 | 0.160559576 |
| IFFO1 | 314.1850789 | 0.561806226 | 0.448377108 | 1.252977051 | 0.210214059 | 0.51816316 |
| IFFO2 | 195.1720149 | -0.204169202 | 0.379239804 | -0.538364381 | 0.590325512 | 0.818321084 |
| IFI16 | 2074.902884 | -0.806441664 | 0.661432524 | -1.219234971 | 0.222755022 | 0.531772847 |
| IFI27 | 33.52011029 | -0.025674322 | 0.554298409 | -0.046318592 | 0.963056321 | 0.987798488 |
| IFI27L1 | 272.4470791 | 1.283593386 | 0.511703567 | 2.508470662 | 0.012125504 | 0.125055076 |
| IFI27L2 | 420.7852121 | 1.190555274 | 0.46268076 | 2.573167888 | 0.010077231 | 0.114259456 |
| IFI30 | 201.6623315 | 0.194006884 | 0.692094488 | 0.280318493 | 0.779233164 | 0.908367621 |
| IFI35 | 36.12035281 | 0.237772276 | 0.545573151 | 0.435821072 | 0.662966554 | 0.854171175 |
| IFI6 | 191.5711965 | 2.129191313 | 0.570934591 | 3.729308656 | 0.000192006 | 0.009160128 |
| IFIT1 | 146.2503858 | 2.47455831 | 0.676830415 | 3.656097973 | 0.000256084 | 0.011068499 |
| IFIT5 | 284.4346828 | 0.954394096 | 0.337501589 | 2.827821041 | 0.0046866 | 0.072477477 |
| IFITM1 | 1625.571614 | 0.67354469 | 0.531843403 | 1.266434229 | 0.205357663 | 0.511820585 |
| IFITM10 | 12.59227258 | 4.885910048 | 1.164909228 | 4.194241004 | 2.74E-05 | 0.002307566 |
| IFITM2 | 293.5547799 | 0.86782185 | 0.412966155 | 2.101435768 | 0.035602731 | 0.215965226 |
| IFITM3 | 3269.91624 | 0.363477185 | 0.403431562 | 0.900963681 | 0.367607631 | 0.673954727 |
| IFNAR1 | 651.2337226 | -0.062605757 | 0.286136604 | -0.218796742 | 0.826808385 | 0.928048979 |
| IFNAR2 | 415.8594916 | 0.39530254 | 0.28266095 | 1.398504251 | 0.161961697 | 0.462586269 |
| IFNGR1 | 758.6375793 | 0.137721108 | 0.26444427 | 0.520794451 | 0.602509969 | 0.825021776 |
| IFNGR2 | 43.56529564 | -0.960523598 | 0.813877843 | -1.18018153 | 0.237928023 | 0.54863845 |
| IFNLR1 | 221.9251337 | -1.129076856 | 0.345685545 | -3.266196321 | | |

0.001090027 0.028511711
IFRD1 1435.35975 0.441452846 0.273944174 1.6114701 0.107077299 0.375093196
IFRD2 764.0938439 0.507411653 0.304982928 1.663737891 0.096164821 0.355920101
IFT122 623.0492519 -0.155591955 0.48979784 -0.317665662
0.750738561 0.896207755
IFT140 970.6169868 -0.134776887 0.542654178 -0.248366072
0.803851182 0.917969611
IFT172 592.1729105 -0.533875237 0.503450109 -1.060433253
0.288947541 0.602063335
IFT20 377.0119771 0.131600609 0.407636079 0.322838472 0.746817575 0.894109554
IFT22 493.3041567 0.278617049 0.330818819 0.84220435 0.39967358 0.697589493
IFT27 966.7380193 0.355495562 0.239253245 1.485854714 0.137317554 0.425712285
IFT43 484.475578 0.556067962 0.40370079 1.377425992 0.168380581 0.470634525
IFT46 670.3707167 -0.273192002 0.282981282 -0.965406618 0.334341188
0.642468672
IFT52 1443.01552 0.112310319 0.258923762 0.433758256 0.664463998 0.85500751
IFT57 999.16072 0.390198986 0.367279591 1.062403126 0.288052715 0.601673803
IFT74 766.7091942 0.755028297 0.328135808 2.300962831 0.021393732 0.16601755
IFT80 1445.306796 -0.03334117 0.328249166 -0.10157275 0.91909581 0.969677217
IFT81 980.7245708 0.514754079 0.359568897 1.431586779 0.152262116 0.448119169
IFT88 303.8099079 0.236995094 0.287916576 0.823138068 0.410429483 0.704150362
IGBP1 2010.653566 -0.112655669 0.35776881 -0.314883986 0.752849741
0.896586461
IGDCC3 3526.077164 -0.123224015 0.376355634 -0.327413765
0.743354957 0.892534058
IGDCC4 689.7632519 0.145725198 0.418343306 0.348338783 0.727585771
0.88517603
IGF1R 2701.831396 -0.572283106 0.256430795 -2.231725349 0.025633121
0.182572806
IGF2 3477.09481 -2.670340798 0.857273866 -3.114921501 0.001839939
0.039876728
IGF2BP1 10322.13184 -0.256338293 0.349717034 -0.732987724
0.463565917 0.740758218
IGF2BP2 5963.261258 -0.429639024 0.220387599 -1.949470053
0.051239316 0.260426481
IGF2BP3 5570.31358 -0.267201146 0.257020537 -1.039610097
0.298521084 0.609920825
IGF2R 4401.611873 -0.874164867 0.412674713 -2.118290363 0.034150484
0.210824952
IGFBP2 8527.172996 0.875053114 0.483763145 1.808846174 0.070474905
0.304955726
IGFBP3 1613.739269 -5.338806379 1.740397902 -3.067578036
0.002158011 0.044096562
IGFBP4 1642.209986 1.359269915 0.549814198 2.472235021 0.013427121
0.131761226
IGFBP5 2823.498589 0.93650287 0.721800439 1.297454004 0.194475021 0.5003189
IGFBP6 21.49587849 1.312989232 0.880500351 1.491185358 0.135912837
0.423730301
IGFBP7 56.89398655 -1.442945213 0.943713 -1.529008516
0.126262333 0.409038191
IGFBPL1 1475.398863 0.319631244 0.304793679 1.048680686 0.294325106
0.606538766
IGFL4 22.69899636 -0.618803758 0.774946309 -0.798511782 0.424573559
0.713390792
IGFLR1 128.349627 0.46432749 0.478310313 0.97076621 0.331664712
0.639512627
IGFN1 19.35088432 -1.977017291 0.815158525 -2.425316339 0.015295053
0.140195951
IGHMBP2 349.5222654 0.176740569 0.388863324 0.454505628 0.649464942
0.847281162
IGIP 56.28076867 -0.016290668 0.676036192 -0.024097331 0.980774972
0.993241851
IGLON5 1096.39381 -0.516468821 0.354011885 -1.458902491
0.144591948 0.435928729

| | | | | | | |
|---|---|---|---|---|---|---|
| IGLV5-52 | 35.4646926 | -0.939877426 | 0.738065975 | -1.273432805 | 0.202864503 | 0.509377254 |
| IGSF1 | 989.3084808 | -0.379128684 | 0.320010298 | -1.184739012 | 0.236120653 | 0.547065733 |
| IGSF10 | 26.15523798 | -1.448381006 | 0.78923445 | -1.835172053 | 0.066480201 | 0.297513606 |
| IGSF11 | 182.9531616 | 0.022751605 | 0.311639977 | 0.073006054 | 0.9418013 | 0.978567737 |
| IGSF21 | 40.71294713 | -0.507893499 | 0.696892563 | -0.728797417 | 0.466125592 | 0.742814218 |
| IGSF3 | 3826.590096 | -1.249751345 | 0.312069431 | -4.004722087 | 6.21E-05 | 0.004422413 |
| IGSF5 | 19.26953742 | -0.759499894 | 0.855286013 | -0.888006916 | 0.374537031 | 0.678946266 |
| IGSF8 | 527.1628865 | 0.760722567 | 0.456335172 | 1.667025934 | 0.095509248 | 0.354819129 |
| IGSF9 | 1097.458238 | -0.556317308 | 0.257145907 | -2.163430539 | 0.030508081 | 0.199123233 |
| IGSF9B | 78.12435362 | 0.145225663 | 0.614082892 | 0.236491954 | 0.81305095 | 0.921802034 |
| IK | 2909.330125 | -0.209963528 | 0.258099272 | -0.81349911 | 0.415931957 | 0.707886887 |
| IKBIP | 528.1078345 | 1.084704629 | 0.250806159 | 4.324872373 | 1.53E-05 | 0.001503507 |
| IKBKB | 1076.217035 | -0.511943968 | 0.335952754 | -1.523857038 | 0.127544428 | 0.411434512 |
| IKBKE | 114.3998691 | -1.061081034 | 0.452339835 | -2.345760759 | 0.018988291 | 0.156305679 |
| IKBKG | 349.7896704 | 0.110963722 | 0.406959987 | 0.272664944 | 0.785110785 | 0.910931322 |
| IKBKGP1 | 46.56249219 | -0.981388557 | 1.149021539 | -0.85410806 | 0.393045119 | 0.692071436 |
| IKZF2 | 685.6040903 | -1.59514494 | 0.518686331 | -3.075355652 | 0.002102517 | 0.043389195 |
| IKZF3 | 15.06020684 | 0.324499383 | 0.829204713 | 0.391338084 | 0.695547352 | 0.869563155 |
| IKZF4 | 483.1261236 | -0.04472416 | 0.318641421 | -0.140358901 | 0.888376428 | 0.955943827 |
| IKZF5 | 381.7478817 | 0.12989243 | 0.260534393 | 0.498561545 | 0.618088301 | 0.832176095 |
| IL10RB | 193.719723 | -0.512713908 | 0.530296595 | -0.966843674 | 0.333622188 | 0.641713975 |
| IL11 | 20.71675001 | 0.519070255 | 0.70177084 | 0.739657771 | 0.459507678 | 0.737476588 |
| IL11RA | 218.1663285 | -0.166134625 | 0.355220447 | -0.467694433 | 0.640003123 | 0.843538624 |
| IL12A | 13.51094169 | 2.167080215 | 1.088212627 | 1.991412487 | 0.046435558 | 0.247792418 |
| IL12RB2 | 30.76780309 | -1.372725271 | 0.676739952 | -2.02843835 | 0.042515529 | 0.237179666 |
| IL13RA1 | 175.5790657 | 0.196090918 | 0.377198704 | 0.519861059 | 0.603160418 | 0.825116926 |
| IL17D | 27.42414825 | 0.376053207 | 0.817124681 | 0.46021521 | 0.645361755 | 0.845794147 |
| IL17RA | 222.8606551 | -0.874249752 | 0.374886921 | -2.332035885 | 0.019698803 | 0.158893482 |
| IL17RB | 62.64914192 | 0.597921674 | 0.48084404 | 1.243483591 | 0.213689684 | 0.520735532 |
| IL17RC | 164.3319994 | -1.374520391 | 0.419822823 | -3.274048751 | 0.001060183 | 0.028069876 |
| IL17RD | 336.4899051 | -0.458344985 | 0.520556126 | -0.880491003 | 0.378593377 | 0.681253679 |
| IL17RE | 191.0231451 | -1.58433168 | 0.715083156 | -2.215590826 | 0.026719538 | 0.186802814 |
| IL18 | 111.9822239 | -0.60411294 | 0.736940634 | -0.819757946 | 0.412354109 | 0.70515068 |
| IL18BP | 279.5578507 | -0.777104126 | 0.49121185 | -1.582014209 | 0.113646327 | 0.388022186 |
| IL1A | 25.88625441 | 0.4989746 | 0.835189493 | 0.59743879 | 0.550214462 | 0.794073783 |
| IL1R1 | 162.2196583 | -2.689617368 | 0.787704113 | -3.414502126 | 0.000638987 | 0.020581892 |
| IL1RAP | 35.43275764 | -0.456958337 | 0.57686227 | -0.792144607 | | |

0.428276363 0.715316008
IL1RAPL1 13.26682606 1.102396602 0.955285952 1.153996456 0.248501622 0.561181618
IL1RL1 15.52801629 -1.523588103 1.128999843 -1.349502493 0.17717562 0.481566699
IL20RB 164.1336026 -0.370812576 0.312738086 -1.185696892 0.235742022 0.546605872
IL21R 12.32937761 0.009573776 0.965038168 0.009920619 0.992084621 0.99688215
IL22RA1 17.40693101 -1.673701082 0.999909009 -1.673853388 0.094159417 0.351758806
IL23A 16.55432519 -0.027724942 0.879099823 -0.031537877 0.974840586 0.991261106
IL27RA 102.6388576 0.507626711 0.514576574 0.986494017 0.32389075 0.633704742
IL2RB 12.71832672 -0.607969036 1.104003665 -0.550694762 0.581842936 0.813002261
IL32 19.60013452 0.505833176 0.802348319 0.630440874 0.528406175 0.779582815
IL36RN 88.08177671 -0.254594672 0.932156758 -0.27312431 0.78475766 0.910931322
IL4R 202.0689116 -0.036041026 0.309430684 -0.11647528 0.907275876 0.965296417
IL6 10.09280235 -0.561969492 1.207008672 -0.465588612 0.641509996 0.843948481
IL6ST 763.8966298 -0.231685844 0.336948113 -0.687600955 0.491704107 0.758965432
ILDR1 47.19307324 -2.034702479 0.815021157 -2.496502652 0.012542473 0.12669427
ILDR2 843.5514076 2.329133979 1.332260779 1.74825681 0.080419568 0.324566447
ILF2 11744.09793 0.643804633 0.290794137 2.213953282 0.026831993 0.187071767
ILF3 16473.05976 -0.155822453 0.341580808 -0.456180353 0.648260292 0.846656069
ILK 2456.805124 -0.424257806 0.257106198 -1.650126711 0.098917023 0.361172118
ILKAP 554.8116871 0.114222857 0.273221602 0.418059393 0.675903693 0.860110127
ILVBL 1233.101086 0.551370864 0.406415519 1.356667838 0.174886776 0.478831849
IMMP1L 363.2350444 1.122076142 0.354110952 3.168713465 0.001531153 0.036126154
IMMP2L 251.4117507 0.474291606 0.480189349 0.987717881 0.323290835 0.63322621
IMMT 3323.810318 0.066162141 0.239625303 0.276106655 0.78246615 0.909909835
IMP3 1159.628072 1.109639534 0.465745481 2.382501986 0.017195439 0.147802838
IMP4 528.0876207 0.247059654 0.274282843 0.900747751 0.367722453 0.674002337
IMPA1 965.6550174 0.990000916 0.317356497 3.119523079 0.001811441 0.039484722
IMPA2 1298.303675 0.651458643 0.265539171 2.453342912 0.014153534 0.135106002
IMPACT 1764.552286 0.083867465 0.229175232 0.365953444 0.714399816 0.878464326
IMPAD1 1150.47978 0.145724555 0.304720205 0.47822413 0.63249069 0.838768903
IMPDH1 1012.40121 -0.480244986 0.340936771 -1.408604254 0.15895222 0.458846746
IMPDH2 10898.26371 0.204573253 0.238999191 0.855957928 0.392021051 0.691496659
IMPG1 44.47119311 0.835960081 0.746769061 1.11943588 0.262954231 0.576568672
INA 863.7445598 -0.800090673 0.481257986 -1.662498487 0.096412867 0.35626747
INAFM1 79.9854224 1.14013857 0.594146324 1.918952493 0.054990346 0.271656967
INAFM2 330.2187219 0.760293374 0.264816667 2.871017833 0.004091524 0.066034593
INAVA 1130.644819 -0.460597496 0.309659656 -1.487431401 0.136900908 0.424728769
INCENP 1055.998335 0.338799763 0.541502152 0.625666512 0.531533703 0.781915815
INF2 1355.74423 -0.649170462 0.480642295 -1.350631163 0.176813613

0.481149663
ING1 727.5971718 0.553668478 0.300011066 1.845493521 0.06496578 0.293856843
ING2 436.0702696 0.632334185 0.388693539 1.62681939 0.103775462 0.369190637
ING3 434.2724149 -0.121355157 0.286054688 -0.424237607 0.671392536 0.858479663
ING4 667.0991629 0.201802341 0.278438431 0.724764683 0.468596406 0.744337782
ING5 1213.117279 0.391826522 0.233012898 1.681565809 0.092653065 0.3491405
INHBA 15.98427473 -3.292949547 1.410694402 -2.334275618 0.019581295 0.158520715
INHBE 20.82227742 1.253042498 0.788661513 1.588821664 0.112100661 0.384920166
INIP 815.253307 0.382524226 0.298665929 1.280776241 0.200272272 0.506121566
INKA2 451.2071211 1.040386853 0.55235915 1.883533299 0.059628132 0.282669434
INMT 155.7541101 -0.124121189 0.461630803 -0.268875449 0.788025531 0.912309638
INMT-MINDY4 11.15321056 -2.162614801 1.97695138 -1.093914005 0.273992696 0.586847211
INO80 836.1653305 -0.652322218 0.312497306 -2.087449096 0.036847558 0.219736143
INO80B 265.4047579 -0.498314065 0.48239858 -1.032992398 0.301607456 0.61241443
INO80B-WBP1 255.3414831 -0.334359446 0.451427208 -0.740671897 0.458892403 0.737172453
INO80C 718.7361055 0.678882847 0.312701003 2.17102868 0.029929004 0.197328521
INO80D 174.1265197 -0.45011165 0.406447919 -1.107427617 0.268109091 0.581152025
INO80E 1083.794306 0.15320501 0.341339634 0.448834518 0.653551036 0.84883826
INPP1 47.1529104 0.010223877 0.504977264 0.020246213 0.983846963 0.994061643
INPP4A 1047.367235 -0.214867397 0.39406132 -0.545263862 0.585572055 0.815618219
INPP4B 14.93629958 -0.180157619 0.847428226 -0.212593366 0.831644146 0.930997148
INPP5A 687.2929653 -0.026442179 0.256604111 -0.103046593 0.917925992 0.969677217
INPP5B 584.8304017 -0.047299226 0.255903947 -0.184831952 0.853360846 0.940799171
INPP5D 678.3866377 0.108715498 0.408940181 0.265846945 0.790357087 0.913087091
INPP5E 533.9369122 0.145905932 0.377502458 0.386503263 0.699123987 0.871039767
INPP5F 4188.673765 -0.412219753 0.294618467 -1.399164679 0.161763606 0.462342815
INPP5J 129.620124 -1.310247892 0.47641709 -2.750211777 0.005955676 0.083399911
INPP5K 459.0344388 -0.205109282 0.295313725 -0.694547068 0.487339194 0.756439734
INPPL1 2704.871486 -0.99342394 0.474922384 -2.091760616 0.036459935 0.21871636
INSIG1 786.5678081 0.90575307 0.260085331 3.4825227 0.000496713 0.017561181
INSIG2 355.0762595 0.43244419 0.302732659 1.428468906 0.153156938 0.449515169
INSM1 60.15638566 -2.447426581 1.099528612 -2.225887124 0.026021743 0.184130533
INSR 781.6678777 -0.501860036 0.287282363 -1.746922539 0.080650776 0.325104967
INSYN1 34.49715037 0.219242763 0.597976095 0.36664135 0.71388656 0.878304517
INTS1 1706.714618 -0.270343995 0.504521695 -0.535842161 0.592067641 0.819765854
INTS10 1118.914123 0.341488284 0.228958067 1.491488327 0.135833334 0.423669306
INTS11 2026.353476 -0.162058033 0.371467453 -0.43626442 0.662644893

0.853902351
INTS12 526.4476343 -0.021015831 0.328614187 -0.063952902
0.949007729 0.98134156
INTS13 1344.733318 0.176624616 0.253719388 0.696141579 0.48634017
0.756340124
INTS14 1110.335139 -0.014572114 0.234065722 -0.062256508
0.950358563 0.981676977
INTS2 712.8813995 -0.029421722 0.354577199 -0.082976916 0.933869895
0.974933951
INTS3 1669.178886 -0.964119642 0.566334447 -1.70238566 0.088683096
0.342301733
INTS4 931.589865 -0.268744484 0.32922033 -0.81630586 0.414325224
0.706658569
INTS4P1 41.47104129 -0.454537071 0.626999351 -0.724940256
0.468488683 0.744337782
INTS5 598.5524349 -0.303108151 0.323874215 -0.935882316 0.349333783
0.656038342
INTS6 1355.872263 -0.32275732 0.270845514 -1.191665741 0.233392336
0.543263647
INTS6L 389.909634 0.209309025 0.510180714 0.410264479 0.681611946
0.862590493
INTS7 617.7810749 -0.227596066 0.339725474 -0.669941126 0.502895322
0.765161461
INTS8 2097.752617 0.117175096 0.34307779 0.341540897 0.732696423 0.888122198
INTS9 572.6755181 -0.552656483 0.31803676 -1.737712594 0.082261469
0.328245329
INTU 817.9984426 -0.28293369 0.277173059 -1.020783518 0.307357015
0.617319594
INVS 488.0841394 -0.545756821 0.366416471 -1.489444019 0.136370484
0.424384947
IP6K1 1574.530921 -0.455740949 0.286070183 -1.593108878 0.111135768
0.383475916
IP6K2 3426.903031 0.172155119 0.329947694 0.521764881 0.601834044 0.825021776
IP6K3 202.1670974 1.507346886 0.525512257 2.868338207 0.004126342 0.066306898
IPCEF1 17.75697739 0.052204957 0.756561984 0.069002882 0.944987325
0.979597042
IPMK 490.2338725 -0.509991895 0.296671661 -1.719044864 0.085606203
0.335589588
IPO11 1380.294226 0.206662496 0.286617648 0.721038978 0.470885534 0.745855888
IPO13 774.2104473 -0.464807016 0.303340848 -1.53229286 0.125450182
0.407445382
IPO4 653.743931 -0.119409646 0.39270783 -0.304067394 0.761076548
0.9003659
IPO5 16139.22574 0.171738762 0.35512107 0.483606229 0.628665353 0.836988334
IPO5P1 335.0005819 -0.180823503 0.287515904 -0.628916524
0.529403707 0.780195218
IPO7 7072.23328 0.184560458 0.363037359 0.508378692 0.611187793 0.829058038
IPO8 1898.949041 -0.138152173 0.330518317 -0.417986435 0.675957035
0.860110127
IPO9 4022.991061 -0.224560115 0.3990732 -0.562704074 0.57363641
0.808163661
IPP 239.1952652 -0.46080469 0.349337934 -1.319080025 0.18714236
0.492051429
IPPK 414.9008465 0.158419713 0.332446111 0.476527497 0.633698622 0.839689885
IQANK1 229.3480404 -1.250845676 0.56631415 -2.208748761
0.027192121 0.188542808
IQCA1 833.7126907 -1.161673281 0.568406729 -2.043735978 0.040979637
0.232847218
IQCB1 1631.838089 0.15040441 0.251536802 0.597941968 0.549878656 0.793878561
IQCC 164.1918045 0.475397649 0.349078608 1.36186417 0.173240756 0.4771935
IQCD 68.46729504 0.346667156 0.520823951 0.66561293 0.50565854 0.767134571
IQCE 591.7543423 -0.497793624 0.372403088 -1.336706488 0.181318463
0.485749663
IQCG 265.5447349 -0.192437782 0.332619826 -0.578551751 0.562891669

0.801767863
IQCH 75.39998059 -0.58567332 0.455326077 -1.2862723 0.198348046 0.50414535
IQCJ-SCHIP1 258.8412811 1.271872735 0.451981043 2.813995752 0.004892987
0.074679591
IQCK 253.2210727 0.739776188 0.369460103 2.002316846 0.045250665 0.244986785
IQCN 42.35422562 -1.551317684 0.765971811 -2.025293439 0.042837242
0.238139903
IQGAP1 4594.535116 -1.111568574 0.302687163 -3.67233471 0.000240345
0.010630519
IQGAP2 630.5126985 0.123907781 0.550103481 0.225244495 0.821789076
0.92563193
IQGAP3 811.3077795 -0.067944771 0.510507165 -0.133092688
0.894120078 0.958161736
IQSEC1 880.4034327 -0.691587173 0.366851333 -1.885197384
0.059403194 0.282486927
IQSEC2 248.8416573 -0.241970044 0.302869718 -0.79892452 0.42433418
0.713303793
IQSEC3 40.07913045 0.962106937 0.723540087 1.329721675 0.183609991
0.488007563
IQUB 41.35242361 0.033719096 0.504067226 0.066894046 0.946666053 0.980533263
IRAK1 861.2233215 -0.195090878 0.466631723 -0.418083187 0.675886297
0.860110127
IRAK1BP1 393.6038406 0.346092954 0.302559663 1.143883325 0.252672027
0.565423035
IRAK2 28.15265257 -0.175064402 0.99904949 -0.175230961 0.860898157
0.943676573
IRAK3 11.98463353 -0.112925809 1.014364007 -0.111326712 0.911357274
0.966866396
IRAK4 210.4101401 -0.340929456 0.297451254 -1.146169165 0.251725164
0.56451182
IREB2 3670.879916 0.111564709 0.281656367 0.39610221 0.692029628 0.86812349
IRF1 108.1579035 -0.797502031 0.452940392 -1.760721819 0.078285498
0.320989249
IRF2 165.5033571 0.900261909 0.449656599 2.002109856 0.045272918 0.244986785
IRF2BP1 377.9068597 0.688871576 0.423128139 1.628044822 0.103515382
0.368992449
IRF2BP2 1859.479544 0.980075221 0.259037649 3.78352423 0.000154623
0.008116919
IRF2BPL 674.8493249 -0.498088704 0.558467496 -0.891884858
0.372454651 0.677762926
IRF3 1731.865429 0.128127942 0.359615141 0.356291846 0.721622015 0.882498649
IRF4 13.27176902 -0.273843463 0.892702281 -0.306757884 0.759027667
0.899440455
IRF6 480.119859 -1.704086577 0.650156488 -2.621040639 0.008766181
0.105800902
IRF7 31.50035423 2.013704575 0.905717818 2.223324456 0.02619393 0.184791771
IRF9 114.7098477 0.101215947 0.352587494 0.287066186 0.774061643 0.90627308
IRGQ 632.3857896 -0.020360308 0.38374681 -0.053056618 0.957686797
0.98520931
IRS1 590.1822011 -0.209306157 0.525973234 -0.397940699 0.690673898
0.867976616
IRS2 514.9842006 0.587137384 0.434474619 1.351373265 0.176575893 0.480766848
IRS4 149.7620221 -1.562106763 0.961252956 -1.625073559 0.104146886
0.369853092
IRX3 2136.867883 -0.927291858 0.629013932 -1.474199235 0.140427956
0.430123922
IRX5 250.640818 -0.403458371 0.551221042 -0.731935722 0.464207801
0.741159393
IRX6 10.6334555 -2.617287363 1.589869395 -1.646227905 0.099716846
0.362001501
ISCA1 1237.122492 0.282202797 0.34774357 0.811525565 0.417063918 0.708661116
ISCA2 764.2440415 1.023221437 0.36615555 2.794499327 0.005198015 0.076636622
ISCU 1641.154669 0.617558694 0.388450343 1.589800873 0.111879697 0.384888109
ISG15 127.7069963 0.497801922 0.719133037 0.692225078 0.488795992 0.757260184

| | | | | | | |
|---|---|---|---|---|---|---|
| ISG20 | 18.73640511 | 0.118205858 | 0.776869713 | 0.152156605 | 0.87906342 | 0.951774989 |
| ISG20L2 | 1063.798523 | -0.172975517 | 0.239530076 | -0.722145293 | 0.470205154 | 0.7454951 |
| ISL1 | 214.07599 | -4.1045104 | 0.922781101 | -4.447978393 | 8.67E-06 | 0.000981385 |
| ISLR | 14.97558547 | 0.98555046 | 1.235294689 | 0.797826193 | 0.424971359 | 0.71350603 |
| ISM2 | 12.27491606 | -2.125862382 | 1.162670513 | -1.828430633 | 0.067484947 | 0.298401089 |
| ISOC1 | 819.8896065 | 0.274896703 | 0.33873673 | 0.811534972 | 0.417058518 | 0.708661116 |
| ISOC2 | 599.5348941 | 1.039176109 | 0.420641035 | 2.470458234 | 0.013494008 | 0.132246504 |
| ISPD | 96.98822111 | -1.252735897 | 0.451406279 | -2.775184918 | 0.005517033 | 0.079368821 |
| IST1 | 5714.044209 | 0.002821409 | 0.219270929 | 0.012867229 | 0.98973372 | 0.996285567 |
| ISY1 | 522.0684215 | 0.453803108 | 0.411471042 | 1.10287982 | 0.270079359 | 0.583007107 |
| ISY1-RAB43 | 830.0891715 | -0.231398684 | 0.291346087 | -0.794239892 | 0.427055789 | 0.714556109 |
| ISYNA1 | 5373.053645 | -0.279379755 | 0.407124559 | -0.686226731 | 0.492570144 | 0.759174768 |
| ITCH | 1403.843087 | 0.059878298 | 0.264128238 | 0.226701614 | 0.820655772 | 0.925132406 |
| ITFG1 | 1170.95081 | -0.140534152 | 0.232761207 | -0.603769647 | 0.545996802 | 0.791677099 |
| ITFG2 | 407.2815952 | -0.45470641 | 0.352679919 | -1.289289198 | 0.197297565 | 0.502721088 |
| ITGA1 | 125.5462635 | -0.161805919 | 0.38526933 | -0.419981313 | 0.674499105 | 0.859684611 |
| ITGA10 | 14.43484396 | -1.290696403 | 0.913921816 | -1.41226129 | 0.15787304 | 0.457277356 |
| ITGA2 | 114.1256207 | -1.299837564 | 0.589320859 | -2.205653413 | 0.027408276 | 0.189349252 |
| ITGA2B | 183.8996849 | 0.575365825 | 0.469110269 | 1.226504433 | 0.220008908 | 0.528543966 |
| ITGA3 | 318.3454912 | -0.627769827 | 0.310712194 | -2.020422238 | 0.043339609 | 0.239441079 |
| ITGA4 | 22.66000735 | 1.510392152 | 1.017459271 | 1.484474313 | 0.137683133 | 0.425849298 |
| ITGA5 | 1152.069266 | -0.064618793 | 0.374459641 | -0.172565442 | 0.862993018 | 0.944730333 |
| ITGA6 | 2937.042528 | 0.475870521 | 0.431654495 | 1.102433836 | 0.270273108 | 0.583170438 |
| ITGA7 | 1266.409366 | 0.479532056 | 0.440157793 | 1.089454881 | 0.275953341 | 0.589314209 |
| ITGA8 | 14.11229563 | 0.913595112 | 1.217561254 | 0.750348378 | 0.453044913 | 0.733436323 |
| ITGA9 | 235.564972 | -0.999859485 | 0.368368152 | -2.714294054 | 0.006641718 | 0.089169476 |
| ITGAE | 410.0319753 | 0.990942818 | 0.361653359 | 2.740034877 | 0.006143267 | 0.084669666 |
| ITGAV | 1154.015555 | -0.313907999 | 0.304751027 | -1.030047387 | 0.302987761 | 0.613538084 |
| ITGB1 | 8800.413128 | -0.115436231 | 0.293945658 | -0.392712829 | 0.694531594 | 0.86892666 |
| ITGB1BP1 | 1034.392676 | 0.712749022 | 0.337012789 | 2.114902003 | 0.0344383 | 0.211851485 |
| ITGB1BP2 | 40.02853139 | 0.24944917 | 0.535401603 | 0.46591039 | 0.641279644 | 0.843932467 |
| ITGB2 | 10.84048748 | -2.44530392 | 1.090892052 | -2.241563605 | 0.02498959 | 0.180790213 |
| ITGB3BP | 1171.683424 | 0.809191584 | 0.251270735 | 3.220397245 | 0.001280131 | 0.03168167 |
| ITGB4 | 180.0908243 | -2.780900586 | 0.603783407 | -4.60579167 | 4.11E-06 | 0.0005616 |
| ITGB5 | 2068.875588 | -1.103265766 | 0.475295146 | -2.321222456 | 0.020274841 | 0.161005801 |
| ITGB6 | 48.91267736 | -2.959809107 | 1.224155831 | -2.417836874 | 0.015613074 | 0.141376414 |
| ITGB8 | 76.85810864 | 1.183477962 | 0.65858355 | 1.797005045 | 0.072334819 | 0.309560377 |
| ITIH4 | 71.40413072 | -0.322546768 | 0.668670643 | -0.482370164 | 0.629543012 | 0.837590803 |
| ITIH5 | 120.5588345 | -1.08491916 | 0.426489866 | -2.54383339 | 0.010964337 | 0.119221957 |
| ITK | 12.30760431 | -1.034014023 | 0.996478026 | -1.037668665 | 0.299424338 | |

| | | | | | | |
|---|---|---|---|---|---|---|
| | | | | | | 0.610246278 |
| ITLN2 | 24.86603781 | -1.480339082 | 0.923258529 | -1.603385222 | 0.108849632 | 0.378925599 |
| ITM2A | 58.48004545 | -0.291071002 | 0.436522628 | -0.666794762 | 0.504903239 | 0.766569683 |
| ITM2B | 4737.905726 | 0.226421593 | 0.40201014 | 0.563223587 | 0.573282644 | 0.80799619 |
| ITM2C | 5422.674199 | -0.498751642 | 0.356499033 | -1.399026633 | 0.161804997 | 0.462374007 |
| ITPA | 890.6415114 | 0.499799078 | 0.416699105 | 1.19942441 | 0.230362961 | 0.539112606 |
| ITPK1 | 605.4452903 | 0.067019182 | 0.286382037 | 0.234020202 | 0.814969295 | 0.922747905 |
| ITPKB | 331.091325 | -0.085723093 | 0.355471969 | -0.241152891 | 0.809436621 | 0.919916322 |
| ITPKC | 610.2237945 | -0.867334193 | 0.359670415 | -2.411469382 | 0.015888386 | 0.142656219 |
| ITPR1 | 96.4310867 | 1.158262006 | 0.657321362 | 1.762093967 | 0.078053422 | 0.320406654 |
| ITPR2 | 290.7081159 | -1.502532506 | 0.449213143 | -3.34480976 | 0.00082339 | 0.024115162 |
| ITPR3 | 1786.108501 | -1.781896275 | 0.288164237 | -6.183613534 | 6.27E-10 | 1.06E-06 |
| ITPRID2 | 735.5479956 | -1.036807596 | 0.329897418 | -3.142818161 | 0.001673298 | 0.037945586 |
| ITPRIP | 216.6073344 | -0.084936882 | 0.430644416 | -0.197232052 | 0.843645949 | 0.936554419 |
| ITPRIPL1 | 113.6387287 | 0.709236439 | 0.550444771 | 1.288478838 | 0.197579331 | 0.503183597 |
| ITPRIPL2 | 200.2839082 | -1.207382357 | 0.488228666 | -2.472985389 | 0.013398962 | 0.131655216 |
| ITSN1 | 3479.042852 | -0.236631788 | 0.360059537 | -0.657201834 | 0.511051147 | 0.770614943 |
| ITSN2 | 453.4950544 | -0.862064657 | 0.345788196 | -2.493042463 | 0.01266537 | 0.127358401 |
| IVD | 1576.812299 | -0.046439676 | 0.355551014 | -0.130613257 | 0.896081255 | 0.958964824 |
| IVNS1ABP | 4589.390479 | 0.42938813 | 0.323126405 | 1.328854972 | 0.183895822 | 0.488339491 |
| IWS1 | 2330.984019 | 0.071587553 | 0.240007855 | 0.298271707 | 0.765495796 | 0.903026935 |
| IZUMO1 | 23.31075439 | -0.324745223 | 0.662371333 | -0.490276687 | 0.623938121 | 0.834649964 |
| IZUMO4 | 44.24911337 | 0.405463576 | 0.49698352 | 0.815849138 | 0.414586425 | 0.706865662 |
| JADE1 | 2149.245867 | 0.020614179 | 0.286299068 | 0.072002258 | 0.942600111 | 0.978574897 |
| JADE2 | 299.7271206 | 0.080376559 | 0.41925566 | 0.19171252 | 0.847967398 | 0.938207222 |
| JADE3 | 1042.144222 | -0.770056949 | 0.419206898 | -1.836937684 | 0.066219095 | 0.297133953 |
| JAG1 | 1554.389635 | 0.530150515 | 0.425673052 | 1.245440633 | 0.212969825 | 0.520189768 |
| JAG2 | 94.7190066 | 0.146284737 | 0.50557641 | 0.289342489 | 0.772319301 | 0.905685717 |
| JAGN1 | 1077.595612 | 0.315613844 | 0.406167963 | 0.777052533 | 0.437127777 | 0.721225178 |
| JAK1 | 1063.130674 | 0.175837713 | 0.370873591 | 0.474117643 | 0.635416018 | 0.840590898 |
| JAK2 | 109.0993618 | -0.642546733 | 0.40816618 | -1.574228254 | 0.11543469 | 0.391255513 |
| JAK3 | 146.482 | 0.606361542 | 0.499863382 | 1.213054535 | 0.225108971 | 0.534448858 |
| JAKMIP1 | 13.78203736 | 1.269039203 | 0.839682173 | 1.511332792 | 0.130703683 | 0.416052367 |
| JAKMIP2 | 642.5415975 | 0.534821694 | 0.53716961 | 0.995629096 | 0.319430392 | 0.629115732 |
| JAM2 | 60.25211374 | 0.786525507 | 0.647854414 | 1.21404669 | 0.224729896 | 0.53405032 |
| JAM3 | 1767.819417 | 0.079050316 | 0.279824126 | 0.282500001 | 0.777560146 | 0.908018171 |
| JARID2 | 2844.042413 | -0.582833243 | 0.29083336 | -2.004010969 | 0.045068885 | 0.244454795 |
| JAZF1 | 56.08825884 | -0.553545958 | 0.513048401 | -1.07893516 | 0.280616634 | 0.594538239 |
| JCAD | 821.0669613 | -2.267245157 | 0.650036052 | -3.48787602 | 0.000486874 | 0.017338878 |
| JDP2 | 712.3587085 | -0.629321568 | 0.315372113 | -1.995488955 | 0.045989578 | |

| | | | | | | |
|---|---|---|---|---|---|---|
| | | | | | | 0.246950947 |
| JHY | 117.4678691 | -0.685348551 | 0.455515292 | -1.504556626 | 0.132438105 | 0.418524946 |
| JKAMP | 1562.694371 | 0.549693407 | 0.238499497 | 2.304799022 | 0.021177832 | 0.165272062 |
| JMJD1C | 2167.467921 | -0.491574778 | 0.352743764 | -1.393574681 | 0.163446079 | 0.464700236 |
| JMJD4 | 383.4790483 | 0.289904759 | 0.403850619 | 0.717851466 | 0.472848876 | 0.747093345 |
| JMJD6 | 306.161433 | 0.063592619 | 0.306254572 | 0.207646267 | 0.835505175 | 0.932531417 |
| JMJD7 | 239.6208755 | 0.456985684 | 0.337950513 | 1.352226634 | 0.176302826 | 0.480368206 |
| JMJD7-PLA2G4B | 567.8448635 | -0.181247757 | 0.861817131 | -0.210308836 | 0.83342664 | 0.931803619 |
| JMJD8 | 878.6407083 | -0.037659363 | 0.291751313 | -0.129080356 | 0.897294065 | 0.959456082 |
| JMY | 432.2833094 | -0.569337942 | 0.28069392 | -2.028323027 | 0.04252729 | 0.237179666 |
| JOSD1 | 849.3790046 | -0.349608217 | 0.361834137 | -0.966211257 | 0.333938481 | 0.64185743 |
| JPH1 | 149.2571276 | 0.238096915 | 0.534928135 | 0.445100751 | 0.656246946 | 0.84995394 |
| JPH2 | 61.1225069 | -3.218392097 | 0.895129671 | -3.595447901 | 0.000323834 | 0.013341539 |
| JPH3 | 329.8779249 | -0.611221209 | 0.440050373 | -1.388980097 | 0.164838802 | 0.467115865 |
| JPH4 | 265.9058056 | 1.074331485 | 0.618774432 | 1.736224753 | 0.082524102 | 0.328688147 |
| JPT1 | 3176.960842 | 1.03004228 | 0.304801161 | 3.379390936 | 0.000726466 | 0.022310161 |
| JPT2 | 5043.853018 | 0.183004713 | 0.247227828 | 0.740226998 | 0.459162268 | 0.737472333 |
| JRK | 980.8171215 | -0.000901373 | 0.390952342 | -0.002305582 | 0.998160413 | 0.998939187 |
| JRKL | 410.5100831 | 0.176026308 | 0.276085178 | 0.637579712 | 0.523747306 | 0.777662198 |
| JTB | 1258.096406 | 0.782853867 | 0.427755345 | 1.830143975 | 0.067228413 | 0.297874489 |
| JUN | 205.8138514 | -2.559276938 | 1.049643198 | -2.438235148 | 0.01475917 | 0.138131624 |
| JUNB | 205.9986772 | -0.727252551 | 0.876396608 | -0.829821276 | 0.40663984 | 0.70159593 |
| JUND | 1324.373687 | 0.849578003 | 0.618810293 | 1.372921578 | 0.169776728 | 0.472495848 |
| JUP | 6122.351935 | -0.907275053 | 0.44272228 | -2.049309676 | 0.040431843 | 0.231601821 |
| KALRN | 653.4697204 | -2.163430748 | 0.580564738 | -3.726424647 | 0.000194215 | 0.00923648 |
| KANK1 | 1744.783583 | -0.816781925 | 0.25606232 | -3.18977788 | 0.001423822 | 0.034286985 |
| KANK2 | 819.6938918 | -0.14956397 | 0.23976293 | -0.623799391 | 0.53275934 | 0.782698212 |
| KANK3 | 20.67482243 | 1.136333088 | 0.726638873 | 1.563820943 | 0.117859625 | 0.395839103 |
| KANK4 | 467.1435298 | -6.347342039 | 1.779500187 | -3.566924064 | 0.000361196 | 0.014270067 |
| KANSL1 | 645.53274 | -1.557596212 | 0.737325036 | -2.112496031 | 0.034643926 | 0.21269873 |
| KANSL1L | 375.8007162 | -0.522077888 | 0.456113826 | -1.14462193 | 0.252365804 | 0.565196577 |
| KANSL2 | 978.9431259 | 0.330552547 | 0.326856145 | 1.011308958 | 0.31186858 | 0.621974264 |
| KANSL3 | 1288.790003 | -0.209039035 | 0.412224565 | -0.507099898 | 0.612084727 | 0.829098987 |
| KANTR | 260.1518074 | -0.08519835 | 0.448532678 | -0.189949036 | 0.849349067 | 0.938580023 |
| KARS | 4775.06371 | -0.154938561 | 0.273896853 | -0.565682151 | 0.571609871 | 0.807288527 |
| KAT14 | 1115.928523 | -0.122591817 | 0.2994296 | -0.409417829 | 0.682233058 | 0.862735748 |
| KAT2A | 943.787153 | 0.194310711 | 0.415502422 | 0.467652415 | 0.640033175 | 0.843538624 |
| KAT2B | 173.9339861 | -1.552380094 | 0.540048352 | -2.874520568 | 0.004046414 | 0.065655774 |
| KAT5 | 776.8867706 | -0.1906896 | 0.33967821 | -0.56138308 | 0.574536416 | 0.808359708 |
| KAT6A | 1336.211429 | 0.242066107 | 0.462598202 | 0.52327507 | 0.600782847 | 0.824475293 |

| | | | | | | |
|---|---|---|---|---|---|---|
| KAT6B | 380.6127431 | -0.509632239 | 0.300106817 | -1.698169485 | 0.089475778 | 0.342777615 |
| KAT7 | 2406.480563 | -0.639900851 | 0.346191976 | -1.848398853 | 0.064544664 | 0.293351439 |
| KAT8 | 898.8608878 | 0.176259814 | 0.258291966 | 0.682405331 | 0.494982686 | 0.760514211 |
| KATNA1 | 593.2822305 | 0.295678036 | 0.373706744 | 0.791203372 | 0.428825324 | 0.715685016 |
| KATNAL1 | 363.4973219 | 0.310547001 | 0.377691374 | 0.822224236 | 0.410949288 | 0.704426187 |
| KATNAL2 | 340.197697 | 0.244458178 | 0.308651699 | 0.792019545 | 0.42834928 | 0.715316008 |
| KATNB1 | 704.1635468 | 0.390343784 | 0.455414094 | 0.857118366 | 0.391379472 | 0.69090272 |
| KATNBL1 | 1779.765397 | 0.153476801 | 0.274651885 | 0.558804833 | 0.576294924 | 0.809009927 |
| KAZALD1 | 41.96459465 | -0.052739494 | 0.524586893 | -0.100535287 | 0.919919369 | 0.969770994 |
| KAZN | 512.0133119 | 0.000884503 | 0.34139422 | 0.002590854 | 0.9979328 | 0.998859851 |
| KBTBD2 | 734.9497108 | -0.033218078 | 0.298403382 | -0.111319377 | 0.911363091 | 0.966866396 |
| KBTBD3 | 134.9243 | 1.008685376 | 0.418287353 | 2.411465153 | 0.01588857 | 0.142656219 |
| KBTBD4 | 242.8946924 | 0.20037147 | 0.286652203 | 0.69900551 | 0.484548587 | 0.755041765 |
| KBTBD6 | 340.7727804 | 0.739486208 | 0.358112476 | 2.064955165 | 0.038927243 | 0.227316695 |
| KBTBD7 | 606.1599043 | 0.515111898 | 0.243241416 | 2.11769816 | 0.034200639 | 0.211003616 |
| KBTBD8 | 239.5323887 | 0.293260993 | 0.614435379 | 0.47728533 | 0.633158956 | 0.839434984 |
| KCMF1 | 1996.036888 | 0.320874983 | 0.344649506 | 0.93101826 | 0.351844119 | 0.65929447 |
| KCNA4 | 14.93121944 | -2.123224654 | 1.101107742 | -1.928262397 | 0.053822496 | 0.268155271 |
| KCNA5 | 10.91198487 | -2.86016642 | 1.228723851 | -2.327753643 | 0.019925189 | 0.159698109 |
| KCNAB1 | 104.5834231 | 0.433250298 | 0.640229498 | 0.676710928 | 0.498589381 | 0.762775559 |
| KCNAB2 | 251.9046757 | 0.176707841 | 0.508742811 | 0.347342188 | 0.72833426 | 0.885679973 |
| KCNAB3 | 76.29285337 | 1.439892609 | 0.60894933 | 2.364552413 | 0.018051873 | 0.15155781 |
| KCNB1 | 52.11643011 | -0.55245587 | 0.503768093 | -1.096647202 | 0.272795645 | 0.585538021 |
| KCNC4 | 240.5496896 | 0.32366712 | 0.443977334 | 0.729017216 | 0.465991132 | 0.742755984 |
| KCND1 | 214.2848494 | -0.926543357 | 0.408577262 | -2.267731082 | 0.023345602 | 0.174385097 |
| KCND2 | 24.09668373 | -0.217174723 | 0.70168544 | -0.309504389 | 0.756937874 | 0.898563152 |
| KCNE2 | 12.00496807 | 1.099439244 | 0.972108734 | 1.130983814 | 0.258061904 | 0.570791245 |
| KCNE3 | 40.56247064 | -0.495674507 | 0.578664743 | -0.856583218 | 0.391675264 | 0.691183602 |
| KCNE5 | 40.28409649 | 0.165288209 | 0.645632126 | 0.256009889 | 0.797943201 | 0.91598035 |
| KCNG1 | 395.9481745 | 0.246272744 | 0.536462224 | 0.459068194 | 0.646185196 | 0.846058135 |
| KCNG3 | 76.64581605 | 0.062629195 | 0.832804117 | 0.075202792 | 0.940053363 | 0.978024247 |
| KCNH2 | 354.8368543 | 0.727655938 | 0.532459657 | 1.366593561 | 0.171752737 | 0.474879827 |
| KCNH8 | 49.01065563 | -1.165299859 | 0.791282152 | -1.472672997 | 0.140839231 | 0.430693807 |
| KCNIP1 | 11.58780759 | 1.306057056 | 0.842515352 | 1.550187843 | 0.121096437 | 0.401219509 |
| KCNIP2 | 90.47387639 | 0.1241406 | 0.671613504 | 0.184839345 | 0.853355047 | 0.940799171 |
| KCNIP3 | 21.4778508 | 0.392315479 | 0.70331169 | 0.557811685 | 0.576972983 | 0.809737014 |
| KCNJ11 | 67.1599795 | 0.882351742 | 0.559767501 | 1.576282546 | 0.114960702 | |

| | | | | | | |
|---|---|---|---|---|---|---|
| | | | | | | 0.390084727 |
| KCNJ12 | 61.82701975 | 1.24347248 | 0.465334173 | 2.672213977 | 0.007535258 | 0.096389041 |
| KCNJ13 | 322.8819766 | -1.943101469 | 0.798789814 | -2.432556644 | 0.014992646 | 0.139113039 |
| KCNJ14 | 32.66860719 | 0.300692384 | 0.602209537 | 0.499315214 | 0.617557339 | 0.831835425 |
| KCNJ15 | 21.0036396 | 0.29316258 | 0.716196045 | 0.409332866 | 0.682295399 | 0.862735748 |
| KCNJ2 | 331.9350709 | -4.696398456 | 0.886296423 | -5.298902643 | 1.17E-07 | 4.21E-05 |
| KCNJ4 | 23.77146808 | 2.197449715 | 0.947356587 | 2.319559229 | 0.020364734 | 0.161249151 |
| KCNJ5 | 16.14350554 | -1.093272901 | 0.789724286 | -1.384372901 | 0.166244302 | 0.468047055 |
| KCNJ6 | 15.26853969 | 1.28631771 | 0.946664506 | 1.358789415 | 0.174213328 | 0.478455902 |
| KCNK1 | 320.7411778 | -0.253420088 | 0.753327252 | -0.336401062 | 0.736568449 | 0.890467761 |
| KCNK10 | 129.3931156 | 1.546765063 | 0.778692341 | 1.986362239 | 0.046993116 | 0.249626249 |
| KCNK12 | 72.68944089 | -0.56011303 | 0.590412159 | -0.948681393 | 0.34278268 | 0.649638482 |
| KCNK13 | 20.93244144 | -2.90003408 | 1.143586289 | -2.5359119 | 0.011215495 | 0.120297968 |
| KCNK5 | 268.3286916 | 0.358172276 | 0.525746513 | 0.681264198 | 0.495704333 | 0.760859008 |
| KCNK6 | 328.4123742 | -0.045687881 | 0.611504339 | -0.074713912 | 0.940442339 | 0.97807441 |
| KCNMA1 | 75.70800266 | -1.835478471 | 0.961019405 | -1.909928627 | 0.056142404 | 0.274488046 |
| KCNMB1 | 22.42464465 | -0.206968002 | 0.746358586 | -0.277303706 | 0.781546914 | 0.909755868 |
| KCNMB3 | 102.4855083 | -0.578811303 | 0.457305685 | -1.265698901 | 0.205620902 | 0.511932925 |
| KCNMB4 | 492.8331039 | 0.644927915 | 0.390407075 | 1.651937059 | 0.098547382 | 0.360516599 |
| KCNN1 | 89.67095168 | 0.55780552 | 0.410353016 | 1.359330865 | 0.174041768 | 0.478455902 |
| KCNN2 | 148.0682626 | -0.355079749 | 0.589911862 | -0.601920002 | 0.547227389 | 0.792404079 |
| KCNN3 | 177.1825122 | 1.337954178 | 0.721469648 | 1.85448436 | 0.063669896 | 0.290852073 |
| KCNN4 | 48.25847981 | -0.990147652 | 0.915945703 | -1.081011297 | 0.279692089 | 0.593293088 |
| KCNQ1 | 24.51823574 | -0.806850979 | 0.758530639 | -1.063702555 | 0.287463466 | 0.601673803 |
| KCNQ2 | 144.3296452 | 0.950417166 | 0.46331701 | 2.05133234 | 0.040234593 | 0.231327132 |
| KCNQ4 | 43.98504681 | -0.234129196 | 0.569586055 | -0.41105149 | 0.68103478 | 0.862490397 |
| KCNRG | 21.05992004 | -0.062786129 | 0.677608335 | -0.092658437 | 0.926174917 | 0.972261514 |
| KCNS1 | 40.14728249 | -3.922057763 | 1.364866309 | -2.873583836 | 0.004058433 | 0.06571024 |
| KCNS3 | 205.1896584 | 0.080943963 | 0.390311984 | 0.207382726 | 0.835710972 | 0.932590743 |
| KCNT2 | 366.5522957 | 0.324379821 | 0.461810264 | 0.702409293 | 0.482423952 | 0.753310328 |
| KCNV1 | 421.300469 | -3.577814536 | 0.765594465 | -4.673250267 | 2.96E-06 | 0.000429914 |
| KCP | 129.07106 | 0.281896921 | 0.53951392 | 0.522501665 | 0.601321088 | 0.824832027 |
| KCTD1 | 981.234488 | -0.509393905 | 0.312849207 | -1.628240996 | 0.103473795 | 0.368930892 |
| KCTD10 | 1251.54272 | -0.515544928 | 0.248304339 | -2.076262264 | 0.037869701 | 0.22354912 |
| KCTD11 | 253.1664913 | -0.019769864 | 0.383828285 | -0.051507054 | 0.958921481 | 0.985756727 |
| KCTD12 | 180.0817906 | -0.188558621 | 0.584094322 | -0.322822211 | 0.746829891 | 0.894109554 |
| KCTD13 | 270.5030074 | 0.029544741 | 0.287120582 | 0.102900116 | 0.918042246 | 0.969677217 |

| | | | | | | |
|---|---|---|---|---|---|---|
| KCTD14 | 231.3391423 | -0.342220162 | 0.417927833 | -0.818849896 | 0.412872059 | 0.705528498 |
| KCTD15 | 1678.210589 | -0.697615824 | 0.339138779 | -2.057021688 | 0.03968414 | 0.229526528 |
| KCTD16 | 25.83710219 | 0.091042456 | 0.596574478 | 0.1526087 | 0.878706863 | 0.951592678 |
| KCTD17 | 434.7827445 | 1.053677941 | 0.371025833 | 2.83990452 | 0.004512704 | 0.070434393 |
| KCTD18 | 254.0256651 | -0.055791848 | 0.292439327 | -0.190780935 | 0.848697226 | 0.938569494 |
| KCTD19 | 23.16486285 | 0.963855281 | 0.821148537 | 1.173789197 | 0.240479472 | 0.552189204 |
| KCTD2 | 1205.47087 | -0.486137671 | 0.336740421 | -1.443657015 | 0.148835478 | 0.443484094 |
| KCTD20 | 1703.147317 | -0.19573201 | 0.346162825 | -0.565433363 | 0.571779037 | 0.807351345 |
| KCTD21 | 202.8422096 | -0.709084985 | 0.449399747 | -1.577849096 | 0.114600282 | 0.389558789 |
| KCTD3 | 1601.672447 | 0.091651251 | 0.28053412 | 0.326702688 | 0.743892767 | 0.892535157 |
| KCTD5 | 763.5419037 | 0.198362769 | 0.3003638 | 0.660408375 | 0.508991799 | 0.769344882 |
| KCTD6 | 577.098449 | 0.860667546 | 0.4116938 | 2.090552606 | 0.036568188 | 0.218932905 |
| KCTD7 | 384.8051461 | -0.076415589 | 0.48933049 | -0.156163555 | 0.875904103 | 0.95018528 |
| KCTD8 | 17.88422907 | -1.22388321 | 1.073161759 | -1.140446162 | 0.25410047 | 0.566489087 |
| KCTD9 | 549.7070671 | 0.087732084 | 0.373046552 | 0.235177309 | 0.814071117 | 0.9222816 |
| KDELC1 | 579.8346992 | 0.623284269 | 0.279790736 | 2.227680152 | 0.025901851 | 0.183710607 |
| KDELC2 | 1856.061618 | -0.919055573 | 0.474278703 | -1.93779642 | 0.052648065 | 0.26491668 |
| KDELR1 | 6314.558068 | 0.401470469 | 0.461083929 | 0.870710176 | 0.383912421 | 0.684699412 |
| KDELR2 | 2977.818061 | 0.351208651 | 0.35658793 | 0.984914581 | 0.324666031 | 0.634653009 |
| KDELR3 | 141.0783079 | 0.292370586 | 0.417654822 | 0.700029236 | 0.483909046 | 0.754355132 |
| KDF1 | 306.8129738 | -0.972664974 | 0.667722968 | -1.456689407 | 0.145202135 | 0.436733637 |
| KDM1A | 9324.763437 | 0.14868523 | 0.297442443 | 0.499878997 | 0.617160282 | 0.831835425 |
| KDM1B | 351.6263162 | 0.101621461 | 0.445550999 | 0.228080426 | 0.819583717 | 0.924316427 |
| KDM2A | 2353.544211 | -0.54776457 | 0.312747282 | -1.751460692 | 0.079866587 | 0.323453279 |
| KDM2B | 2636.975049 | -0.317688763 | 0.21739283 | -1.461358057 | 0.143917205 | 0.434995877 |
| KDM3A | 1243.259761 | -0.227539045 | 0.445462014 | -0.510793375 | 0.609495749 | 0.828395867 |
| KDM3B | 4222.319117 | -0.61726131 | 0.323700075 | -1.906892701 | 0.056534484 | 0.275428597 |
| KDM4A | 2234.287133 | -0.582652739 | 0.448345298 | -1.299562506 | 0.193750957 | 0.499812238 |
| KDM4B | 1174.938053 | -0.508511607 | 0.373272636 | -1.362306147 | 0.173101289 | 0.477042625 |
| KDM4C | 461.2321374 | -1.083363095 | 0.271824619 | -3.985522352 | 6.73E-05 | 0.004643142 |
| KDM4D | 103.3645967 | 0.21113993 | 0.437504132 | 0.482600996 | 0.629379072 | 0.837497579 |
| KDM5A | 1444.651125 | -0.356818865 | 0.264754323 | -1.347735746 | 0.177743391 | 0.482042363 |
| KDM5B | 7501.768978 | -0.539797592 | 0.235631169 | -2.290858182 | 0.021971617 | 0.168393647 |
| KDM5C | 2732.39012 | -0.521074936 | 0.42098216 | -1.237760138 | 0.215805013 | 0.523378873 |
| KDM5D | 1528.798678 | -0.895278872 | 0.359292441 | -2.491783212 | 0.012710359 | 0.127363842 |
| KDM6A | 1485.336705 | -0.841527508 | 0.331123786 | -2.54142875 | 0.011040045 | |

| | | | | | | |
|---|---|---|---|---|---|---|
| | | | | | | 0.11934855 |
| KDM6B | 1281.751864 | -1.023394899 | 0.421659048 | -2.427067327 | 0.015221431 | 0.139780768 |
| KDM7A | 587.3553975 | -0.701120395 | 0.463004114 | -1.514285453 | 0.129953456 | 0.415175781 |
| KDM8 | 150.3985048 | 0.578640918 | 0.323528305 | 1.788532592 | 0.073690122 | 0.311841798 |
| KDR | 67.89522302 | -1.109568113 | 0.737933609 | -1.503615095 | 0.132680502 | 0.418742646 |
| KDSR | 925.0487164 | 0.20609446 | 0.353961289 | 0.582251412 | 0.560397348 | 0.800545025 |
| KEAP1 | 856.7839667 | 0.494349695 | 0.254431255 | 1.942959778 | 0.052021018 | 0.262949685 |
| KEL | 70.97310535 | -0.955696722 | 0.556927499 | -1.716016401 | 0.086159041 | 0.336626012 |
| KHDC1 | 143.3402054 | 0.56349571 | 0.382504265 | 1.473174974 | 0.140703861 | 0.430627049 |
| KHDC4 | 1783.500821 | 0.1326528 | 0.276854129 | 0.479143298 | 0.631836689 | 0.838768903 |
| KHDRBS1 | 11599.99054 | 0.177751418 | 0.282824878 | 0.628485796 | 0.529685749 | 0.780385606 |
| KHDRBS2 | 594.7534609 | 0.177814754 | 0.458702819 | 0.387646962 | 0.698277311 | 0.870504139 |
| KHDRBS3 | 814.0297942 | -0.403583204 | 0.508091509 | -0.794312042 | 0.427013795 | 0.714556109 |
| KHK | 147.983223 | 0.86989692 | 0.357256974 | 2.434933347 | 0.014894533 | 0.138715007 |
| KHNYN | 1198.3554 | -0.472017283 | 0.400504444 | -1.178556917 | 0.238574656 | 0.549478686 |
| KHSRP | 6963.846855 | -0.155708582 | 0.334002387 | -0.466190027 | 0.641079487 | 0.843888668 |
| KIAA0040 | 160.7016218 | -2.106334781 | 0.581666466 | -3.621207179 | 0.000293232 | 0.012323036 |
| KIAA0100 | 3916.175615 | -0.615900401 | 0.486042234 | -1.267174657 | 0.205092845 | 0.51158559 |
| KIAA0232 | 1607.106211 | -0.135783886 | 0.372980135 | -0.364051256 | 0.715819735 | 0.879470457 |
| KIAA0319L | 1582.154513 | -0.215289024 | 0.297147797 | -0.724518326 | 0.46874758 | 0.744337782 |
| KIAA0355 | 719.7058952 | -1.108482894 | 1.553588721 | -0.713498289 | 0.475537471 | 0.74892861 |
| KIAA0391 | 1189.750581 | -0.110673856 | 0.296177545 | -0.373674027 | 0.708646851 | 0.875585221 |
| KIAA0408 | 76.68534476 | 1.863277853 | 0.638844444 | 2.916637798 | 0.003538264 | 0.061068261 |
| KIAA0513 | 357.0162076 | 0.298181607 | 0.543280859 | 0.54885351 | 0.583105985 | 0.81387283 |
| KIAA0556 | 745.6471341 | -0.655738368 | 0.446727613 | -1.467870685 | 0.142139352 | 0.432308365 |
| KIAA0586 | 871.7656839 | 0.224600792 | 0.442365692 | 0.507726518 | 0.611645149 | 0.829058038 |
| KIAA0753 | 635.242455 | -0.422932058 | 0.360196241 | -1.174171213 | 0.240326454 | 0.551937498 |
| KIAA0825 | 81.13561463 | -0.416233386 | 0.407296544 | -1.021941856 | 0.30680842 | 0.616910609 |
| KIAA0895 | 175.9470465 | -0.542763175 | 0.33446645 | -1.622773152 | 0.104637907 | 0.370855378 |
| KIAA0895L | 788.6752339 | 0.039248726 | 0.437293578 | 0.089753723 | 0.928482924 | 0.972633911 |
| KIAA0930 | 1384.138661 | 0.430922871 | 0.254547381 | 1.692898464 | 0.090474796 | 0.344786015 |
| KIAA1109 | 2331.017474 | -0.330738483 | 0.406808468 | -0.813007863 | 0.41621355 | 0.708127819 |
| KIAA1143 | 1138.418669 | 0.259512457 | 0.264590071 | 0.980809509 | 0.326686688 | 0.636466385 |
| KIAA1147 | 57.08349438 | 0.083275991 | 0.739165161 | 0.112662225 | 0.910298351 | 0.966517087 |
| KIAA1191 | 3786.439078 | -0.01024584 | 0.22472244 | -0.045593309 | 0.963634403 | 0.988130579 |
| KIAA1211 | 783.1132824 | -1.142594685 | 0.75774966 | -1.507878848 | | |

| | | | | | | |
|---|---|---|---|---|---|---|
| | | | | | 0.131585537 | 0.41719445 |
| KIAA1217 | 3321.499064 | -1.543475952 | 0.387156758 | -3.986695105 | 6.70E-05 | 0.004643142 |
| KIAA1257 | 299.3566259 | 0.988594184 | 0.511476799 | 1.932823124 | 0.053257991 | 0.267056023 |
| KIAA1324 | 36.60717571 | -0.968921793 | 0.625744274 | -1.548430938 | 0.121518577 | 0.402062246 |
| KIAA1324L | 1884.117495 | -0.243652449 | 0.272246667 | -0.894969447 | 0.370803421 | 0.676136863 |
| KIAA1328 | 467.8517764 | -0.046887495 | 0.263019427 | -0.178266282 | 0.858513857 | 0.94248999 |
| KIAA1522 | 3174.09713 | -0.214502949 | 0.32120016 | -0.667817068 | 0.50425037 | 0.765917338 |
| KIAA1549 | 663.9158922 | -0.236612422 | 0.341115583 | -0.693642959 | 0.487906146 | 0.75669844 |
| KIAA1549L | 22.60765684 | 0.024136152 | 0.824249756 | 0.02928257 | 0.976639228 | 0.992005472 |
| KIAA1586 | 888.874915 | 0.276862624 | 0.238576874 | 1.160475527 | 0.245855252 | 0.557612503 |
| KIAA1614 | 37.03387984 | 0.10845266 | 0.605812519 | 0.17902017 | 0.857921864 | 0.942343063 |
| KIAA1671 | 1033.425183 | -0.640199066 | 0.386816403 | -1.655046324 | 0.097915102 | 0.359375073 |
| KIAA1755 | 16.6477138 | -0.005307134 | 0.751370713 | -0.00706327 | 0.994364373 | 0.997816146 |
| KIAA1841 | 500.4054127 | 0.002130329 | 0.268036043 | 0.007947921 | 0.993658543 | 0.99730871 |
| KIAA1958 | 1268.274735 | -0.168141123 | 0.355708032 | -0.472694198 | 0.636431367 | 0.841268221 |
| KIAA2012 | 23.54031276 | -0.559873933 | 0.824103057 | -0.679373689 | 0.496901117 | 0.761325016 |
| KIAA2013 | 1069.429233 | 0.331121555 | 0.331622825 | 0.998488434 | 0.31804257 | 0.628145491 |
| KIAA2026 | 740.4618506 | -0.300840569 | 0.333209826 | -0.902856235 | 0.366602201 | 0.673250453 |
| KIDINS220 | 2047.988111 | -0.683628528 | 0.408245261 | -1.674553492 | 0.094021871 | 0.351418034 |
| KIF11 | 2059.342801 | 0.602499228 | 0.386657044 | 1.558226438 | 0.11917958 | 0.398078689 |
| KIF12 | 37.49273265 | -2.11811514 | 0.818478847 | -2.587867906 | 0.009657201 | 0.111583694 |
| KIF13A | 682.3685151 | 0.061693709 | 0.403302808 | 0.152971187 | 0.878420997 | 0.95141893 |
| KIF13B | 633.468097 | -1.200718092 | 0.271479013 | -4.422876303 | 9.74E-06 | 0.001086461 |
| KIF14 | 1141.01146 | 0.762463056 | 0.453152272 | 1.682575819 | 0.092457236 | 0.34894241 |
| KIF15 | 547.5342489 | 1.118079533 | 0.533528031 | 2.095634097 | 0.036114662 | 0.217505176 |
| KIF16B | 541.6349743 | -1.031433099 | 0.373345038 | -2.762680616 | 0.005732883 | 0.081513644 |
| KIF17 | 135.9883732 | 0.408238605 | 0.536594689 | 0.760795091 | 0.446779467 | 0.728709125 |
| KIF18A | 650.2008636 | 0.744383633 | 0.448639232 | 1.659203161 | 0.097074861 | 0.357891303 |
| KIF18B | 655.2114189 | 0.805726853 | 0.422489336 | 1.907093942 | 0.056508424 | 0.275406162 |
| KIF19 | 14.69709294 | 1.03812066 | 0.798263723 | 1.300473303 | 0.193438801 | 0.499431594 |
| KIF1A | 2332.562119 | 0.015357045 | 0.51482327 | 0.029829742 | 0.976202838 | 0.991761418 |
| KIF1B | 3865.616484 | 0.052270368 | 0.423469332 | 0.123433657 | 0.901763707 | 0.962206864 |
| KIF1BP | 601.9257486 | 0.124859982 | 0.320224937 | 0.389913363 | 0.696600612 | 0.870023227 |
| KIF1C | 2316.818794 | -0.554777022 | 0.340900445 | -1.627387203 | 0.103654888 | 0.369064484 |
| KIF20A | 2596.834451 | 0.708808572 | 0.405610277 | 1.747511371 | 0.080548674 | 0.325001047 |
| KIF20B | 1818.405947 | 0.529681733 | 0.345792671 | 1.531789936 | 0.12557428 | 0.407592514 |

| | | | | | | |
|---|---|---|---|---|---|---|
| KIF21A | 1871.989203 | -0.581967976 | 0.335871331 | -1.732711079 | 0.083147034 | 0.330129196 |
| KIF21B | 321.2696256 | 0.571841975 | 0.666033342 | 0.858578601 | 0.390573049 | 0.690466555 |
| KIF22 | 3199.313903 | 0.552028939 | 0.225046299 | 2.452957192 | 0.01416872 | 0.135106002 |
| KIF23 | 1398.17423 | 0.607051839 | 0.367941318 | 1.649860478 | 0.098971476 | 0.36119708 |
| KIF24 | 224.9482612 | -0.293996838 | 0.568443458 | -0.517196273 | 0.605019153 | 0.82602102 |
| KIF25 | 28.81411413 | 0.213253224 | 0.728645164 | 0.292670884 | 0.769773727 | 0.904694332 |
| KIF26A | 423.1008415 | -0.126278824 | 0.538919625 | -0.234318474 | 0.814737745 | 0.922623448 |
| KIF26B | 338.4759714 | -4.255831496 | 0.915038442 | -4.650986561 | 3.30E-06 | 0.000472807 |
| KIF27 | 289.1177095 | -0.057449446 | 0.276795986 | -0.207551585 | 0.835579111 | 0.932531417 |
| KIF2A | 1611.04388 | -0.116449986 | 0.295078349 | -0.394640901 | 0.693107924 | 0.868643566 |
| KIF2C | 1663.45913 | 0.256995785 | 0.331851425 | 0.774430258 | 0.438676396 | 0.722366178 |
| KIF3A | 769.9825658 | 0.098101271 | 0.24754409 | 0.396298175 | 0.691885074 | 0.86812349 |
| KIF3B | 1470.656313 | -0.000977311 | 0.262046411 | -0.003729535 | 0.997024268 | 0.998614074 |
| KIF3C | 1292.23614 | -0.269115511 | 0.464922256 | -0.578839811 | 0.562697266 | 0.801716775 |
| KIF4A | 1647.910665 | 0.720633685 | 0.379490331 | 1.898951373 | 0.05757087 | 0.277978251 |
| KIF5A | 496.97014 | 0.95873666 | 0.749659822 | 1.278895615 | 0.200933817 | 0.506881795 |
| KIF5B | 4296.629429 | 0.311650687 | 0.228356273 | 1.364756405 | 0.172329625 | 0.475866898 |
| KIF5C | 1650.04328 | 0.556378463 | 0.442101436 | 1.258485989 | 0.208216051 | 0.51514118 |
| KIF6 | 46.31323016 | -0.466302616 | 0.474450174 | -0.982827368 | 0.32569241 | 0.635672141 |
| KIF7 | 448.7660535 | -0.42588415 | 0.389434129 | -1.093597397 | 0.274131591 | 0.586993701 |
| KIF9 | 80.89634766 | 0.54523556 | 0.471327245 | 1.156808918 | 0.247350438 | 0.559999028 |
| KIFAP3 | 546.625501 | -0.548741457 | 0.453338144 | -1.210446252 | 0.226107703 | 0.534727976 |
| KIFC1 | 1676.736536 | 0.592283662 | 0.425830839 | 1.390889546 | 0.164258923 | 0.466312149 |
| KIFC2 | 716.4200224 | -0.409886749 | 0.465357691 | -0.880799344 | 0.378426435 | 0.681253679 |
| KIFC3 | 193.9416905 | -1.068894056 | 0.470091876 | -2.27379819 | 0.022978123 | 0.173260987 |
| KIN | 764.5794261 | 0.430869331 | 0.298527093 | 1.443317341 | 0.148931097 | 0.443544075 |
| KIRREL1 | 732.1129345 | 0.553722944 | 0.542538269 | 1.020615458 | 0.307436663 | 0.617319594 |
| KIRREL2 | 132.5258893 | 1.090226422 | 0.474089337 | 2.299622324 | 0.021469626 | 0.166436229 |
| KIRREL3 | 18.6878035 | -0.034117832 | 0.712842088 | -0.047861697 | 0.961826466 | 0.9870991 |
| KIT | 1519.994026 | -0.87658218 | 0.333660059 | -2.627171452 | 0.008609794 | 0.104830807 |
| KITLG | 177.1943193 | 0.822143003 | 0.514823077 | 1.596942794 | 0.11027846 | 0.382145845 |
| KIZ | 653.5793732 | 0.182601094 | 0.376268917 | 0.485294122 | 0.627467724 | 0.836054176 |
| KLC1 | 2509.588156 | 0.382970605 | 0.271728848 | 1.409385156 | 0.158721311 | 0.458571893 |
| KLC2 | 1050.113286 | -0.105620891 | 0.430899016 | -0.245117503 | 0.806365451 | 0.919090941 |
| KLC3 | 38.27276367 | -0.43341299 | 0.756519036 | -0.572904275 | 0.56670951 | 0.804513376 |
| KLC4 | 525.1099663 | -0.049313877 | 0.316038552 | -0.156037535 | 0.876003435 | 0.95022509 |
| KLF10 | 610.9679586 | 0.66088802 | 0.335077801 | 1.972341998 | 0.048570578 | 0.25374113 |
| KLF11 | 202.2737554 | -0.190462455 | 0.416935357 | -0.456815312 | 0.647803799 | 0.846597949 |
| KLF12 | 412.9423382 | -0.471525425 | 0.38181422 | -1.234960357 | 0.216845258 | 0.524598854 |
| KLF13 | 672.9158824 | 0.099271208 | 0.334715193 | 0.29658411 | 0.766784043 | 0.903470997 |
| KLF15 | 328.003016 | -0.063744346 | 0.336654796 | -0.189346317 | 0.849821396 | |

| | | | | | | |
|---|---|---|---|---|---|---|
| | | | | | | 0.938580023 |
| KLF16 | 421.4910898 | 0.631387589 | 0.330237366 | 1.911920498 | 0.055886392 | 0.273589046 |
| KLF3 | 1139.965484 | -0.395321502 | 0.282317681 | -1.400271849 | 0.161431927 | 0.46167973 |
| KLF4 | 136.813425 | -1.413502407 | 0.641805813 | -2.202383305 | 0.027638244 | 0.190244919 |
| KLF5 | 1375.576743 | -0.749453728 | 0.728664232 | -1.028530969 | 0.303700131 | 0.614002998 |
| KLF6 | 2154.112846 | -1.023236916 | 0.558064405 | -1.833546282 | 0.066721372 | 0.297817842 |
| KLF7 | 346.6563604 | -0.321132684 | 0.280945936 | -1.143040857 | 0.253021628 | 0.565800563 |
| KLF8 | 1018.052432 | -0.971751365 | 0.259507938 | -3.744592065 | 0.000180687 | 0.008851388 |
| KLF9 | 50.95224109 | 0.917560277 | 0.586241197 | 1.565158303 | 0.117545796 | 0.395641036 |
| KLHDC1 | 14.73085891 | -0.301728984 | 0.835559067 | -0.361110299 | 0.718016995 | 0.880671367 |
| KLHDC10 | 871.2053599 | -0.018005538 | 0.282030983 | -0.063842411 | 0.949095708 | 0.98134156 |
| KLHDC2 | 1016.97298 | 0.227154746 | 0.300450892 | 0.75604617 | 0.449621514 | 0.730869816 |
| KLHDC3 | 3531.05376 | 0.088392749 | 0.266020403 | 0.33227808 | 0.73967929 | 0.89168179 |
| KLHDC4 | 726.1833677 | -0.199737505 | 0.295520554 | -0.675883633 | 0.49911453 | 0.76281557 |
| KLHDC7A | 83.40689988 | -0.623015543 | 0.505437272 | -1.23262683 | 0.217715022 | 0.526030161 |
| KLHDC8A | 58.80344634 | 1.491927759 | 0.833325252 | 1.790330673 | 0.073400768 | 0.311344591 |
| KLHDC8B | 677.3208064 | 0.828180817 | 0.404228991 | 2.048791245 | 0.040482532 | 0.231601821 |
| KLHDC9 | 243.9868668 | 0.490555466 | 0.371274814 | 1.321273214 | 0.186410282 | 0.491062752 |
| KLHL11 | 390.1224116 | 0.456093018 | 0.341608372 | 1.335134194 | 0.18183243 | 0.48591777 |
| KLHL12 | 1452.552388 | 0.069825365 | 0.240206066 | 0.290689432 | 0.771288855 | 0.905067213 |
| KLHL13 | 614.691087 | -0.065932453 | 0.374456821 | -0.176074916 | 0.860235087 | 0.943475231 |
| KLHL14 | 34.01731577 | -0.40057464 | 0.904594452 | -0.442822349 | 0.657894237 | 0.850760404 |
| KLHL15 | 604.0456747 | 0.002875454 | 0.308698241 | 0.009314773 | 0.992567994 | 0.997035603 |
| KLHL17 | 316.5326096 | -0.112558315 | 0.529437409 | -0.212599853 | 0.831639086 | 0.930997148 |
| KLHL18 | 512.4347084 | -0.414802582 | 0.391649422 | -1.059117054 | 0.289546474 | 0.602398681 |
| KLHL2 | 396.7148414 | 0.03251408 | 0.320940218 | 0.101308836 | 0.919305302 | 0.969677217 |
| KLHL20 | 302.7574656 | 0.057303491 | 0.387722546 | 0.147795097 | 0.882504479 | 0.953255639 |
| KLHL21 | 485.2585942 | -0.400686988 | 0.386221853 | -1.037452917 | 0.299524828 | 0.610246278 |
| KLHL22 | 341.2464638 | 0.136551478 | 0.465964565 | 0.293051207 | 0.769483011 | 0.904597192 |
| KLHL23 | 1974.675124 | 0.400905303 | 0.295687323 | 1.355842037 | 0.175149433 | 0.479186245 |
| KLHL24 | 630.2610013 | -0.359199829 | 0.273759962 | -1.312097746 | 0.189487156 | 0.494956896 |
| KLHL25 | 368.9587297 | 0.525775299 | 0.541049926 | 0.971768543 | 0.331165708 | 0.639084337 |
| KLHL26 | 214.3568795 | 0.255149314 | 0.375698996 | 0.679132275 | 0.497054054 | 0.761325016 |
| KLHL28 | 471.1276257 | 0.142021885 | 0.371359003 | 0.382438244 | 0.702136328 | 0.872649539 |

| | | | | | | |
|---|---|---|---|---|---|---|
| KLHL29 | 48.43483169 | -0.139459304 | 0.775368002 | -0.179862083 | 0.857260843 | 0.942343063 |
| KLHL3 | 325.5613746 | -0.231216691 | 0.427337497 | -0.541063428 | 0.588463865 | 0.81754444 |
| KLHL30 | 15.57429096 | -0.207142479 | 0.934111101 | -0.221753578 | 0.824505719 | 0.92681808 |
| KLHL31 | 15.63421036 | -2.547886101 | 1.174909586 | -2.16858057 | 0.030114542 | 0.198185614 |
| KLHL32 | 17.57181648 | -0.051104368 | 0.697451409 | -0.073273015 | 0.941588865 | 0.978517574 |
| KLHL34 | 29.4731834 | -0.152555435 | 0.747982469 | -0.203955897 | 0.838387963 | 0.933879959 |
| KLHL35 | 11.82996673 | 0.748770031 | 0.850379369 | 0.880512932 | 0.378581503 | 0.681253679 |
| KLHL36 | 742.4113098 | -0.472220447 | 0.287241827 | -1.643982186 | 0.100179882 | 0.36243601 |
| KLHL4 | 567.4729815 | 0.933001093 | 0.52271847 | 1.784901713 | 0.074277261 | 0.313236304 |
| KLHL42 | 650.2526414 | 0.037279529 | 0.318740562 | 0.11695885 | 0.906892663 | 0.965235624 |
| KLHL5 | 1184.220791 | 0.060614305 | 0.247230756 | 0.245172997 | 0.806322484 | 0.919090941 |
| KLHL7 | 1627.645281 | 0.523744425 | 0.255790731 | 2.047550445 | 0.040604069 | 0.231842053 |
| KLHL8 | 360.5544347 | 0.302738608 | 0.409142337 | 0.73993469 | 0.459339624 | 0.737476588 |
| KLHL9 | 1714.129265 | 0.01051699 | 0.279993107 | 0.037561603 | 0.970037223 | 0.989496332 |
| KLK10 | 75.0311191 | 0.078444657 | 0.865480305 | 0.090637137 | 0.927780922 | 0.972633911 |
| KLK4 | 20.32548361 | -1.813531995 | 0.818463057 | -2.215777462 | 0.026706747 | 0.186799472 |
| KLK6 | 100.9249303 | -1.240581498 | 0.668182764 | -1.856649953 | 0.063360972 | 0.290232279 |
| KLK7 | 51.02190717 | 0.034382917 | 0.811088312 | 0.042391089 | 0.966186932 | 0.988700292 |
| KLK8 | 166.2255119 | 0.122983588 | 0.701226341 | 0.175383583 | 0.860778239 | 0.943659349 |
| KLKB1 | 183.7874708 | -0.431254965 | 0.901420733 | -0.478416958 | 0.632353466 | 0.838768903 |
| KLLN | 22.89365868 | -0.080745008 | 0.824896624 | -0.097885002 | 0.92202361 | 0.970640219 |
| KLRA1P | 135.2862421 | -0.218853889 | 0.385561661 | -0.567623575 | 0.570290595 | 0.80632606 |
| KLRD1 | 54.05273994 | -0.216870781 | 0.540877596 | -0.400960925 | 0.688448893 | 0.867059891 |
| KLRG1 | 41.96310137 | 0.903926528 | 0.53962505 | 1.675101126 | 0.093914392 | 0.351340153 |
| KLRG2 | 391.2823355 | -1.826818688 | 0.367897674 | -4.965561933 | 6.85E-07 | 0.000142363 |
| KMT2A | 1527.172785 | -0.17206384 | 0.378030065 | -0.455159144 | 0.648994751 | 0.847104824 |
| KMT2B | 1284.677972 | -0.539023701 | 0.434176594 | -1.241484935 | 0.21442666 | 0.521483923 |
| KMT2C | 1651.855896 | -0.601418577 | 0.327321358 | -1.837394849 | 0.066151627 | 0.296938744 |
| KMT2D | 2124.869701 | -0.74934254 | 0.427378352 | -1.753346973 | 0.079542467 | 0.323002882 |
| KMT2E | 1909.117208 | -0.07351082 | 0.242026985 | -0.303729853 | 0.761333713 | 0.90056031 |
| KMT5A | 553.0971777 | -0.138478547 | 0.31108548 | -0.445146289 | 0.656214039 | 0.84995394 |
| KMT5B | 1525.753622 | -0.016834462 | 0.251348444 | -0.06697659 | 0.94660034 | 0.980533263 |
| KMT5C | 623.0798945 | 0.559562835 | 0.340124418 | 1.645171021 | 0.099934548 | 0.362399544 |
| KNDC1 | 102.4874425 | 0.353627052 | 0.56760898 | 0.623011729 | 0.533276814 | 0.783143103 |
| KNL1 | 952.3836778 | 0.626510439 | 0.468005904 | 1.33868063 | 0.180674665 | 0.485393189 |
| KNOP1 | 2332.909328 | 0.834159856 | 0.361764896 | 2.305806521 | 0.021121446 | 0.165086787 |
| KNSTRN | 1156.341691 | 1.065931252 | 0.270962498 | 3.933870033 | 8.36E-05 | 0.005396292 |
| KNTC1 | 2417.577739 | 0.17372255 | 0.437175658 | 0.397374709 | 0.691091163 | 0.868004248 |
| KPNA1 | 2303.527213 | -0.048651017 | 0.253579248 | -0.191857249 | 0.847854025 | 0.938207222 |

| | | | | | | |
|---|---|---|---|---|---|---|
| KPNA2 | 8916.951813 | 0.763146695 | 0.321842103 | 2.371183535 | 0.017731224 | 0.150195643 |
| KPNA2P3 | 55.7808287 | -0.399067973 | 0.497014544 | -0.802930172 | 0.422015096 | 0.712080472 |
| KPNA3 | 2242.643167 | 0.153639254 | 0.28495402 | 0.539172087 | 0.589768119 | 0.81828099 |
| KPNA4 | 2495.736756 | 0.033248551 | 0.229654942 | 0.144776119 | 0.884887637 | 0.95440284 |
| KPNA5 | 960.6993845 | 0.442139889 | 0.254278712 | 1.738800251 | 0.082069904 | 0.327912173 |
| KPNA6 | 2534.677971 | 0.085858315 | 0.27084392 | 0.317002926 | 0.751241382 | 0.896349431 |
| KPNB1 | 12852.82206 | -0.168020864 | 0.344270083 | -0.488049563 | 0.625514735 | 0.835359511 |
| KPTN | 375.637831 | 0.724053588 | 0.305983897 | 2.366312724 | 0.017966261 | 0.151238239 |
| KRAS | 2529.319542 | -0.330060785 | 0.283083831 | -1.165947145 | 0.243635816 | 0.554901511 |
| KRBA1 | 295.0348654 | -0.175677777 | 0.463153846 | -0.379307606 | 0.704459452 | 0.873435301 |
| KRBA2 | 126.2568939 | -1.181905673 | 0.520176785 | -2.272123068 | 0.023079078 | 0.173590826 |
| KRBOX4 | 285.7354873 | -0.308442317 | 0.335378061 | -0.91968543 | 0.357737169 | 0.665264843 |
| KRCC1 | 570.9770279 | 0.269108554 | 0.363317745 | 0.740697525 | 0.45887686 | 0.737172453 |
| KREMEN1 | 190.1248913 | -0.394445641 | 0.324779616 | -1.214502455 | 0.224555913 | 0.533887753 |
| KREMEN2 | 162.0053026 | 0.536488858 | 0.596532554 | 0.899345483 | 0.368468668 | 0.674555106 |
| KRI1 | 1061.652127 | 0.208696013 | 0.326600577 | 0.638994624 | 0.522826429 | 0.777662198 |
| KRIT1 | 1043.75405 | -0.514401754 | 0.340299209 | -1.511616076 | 0.13063156 | 0.416016191 |
| KRR1 | 1632.550023 | 0.253909661 | 0.235659042 | 1.07744502 | 0.2812815 | 0.595171527 |
| KRT10 | 458.5861197 | 0.770714374 | 0.476596889 | 1.617120028 | 0.105852374 | 0.373028193 |
| KRT15 | 33.79765444 | -2.759671411 | 1.035594358 | -2.664818893 | 0.007702979 | 0.098203278 |
| KRT16 | 19.41719146 | -4.136194612 | 4.271912807 | -0.968230111 | 0.33292946 | 0.640729778 |
| KRT17 | 27.20767085 | -1.684342886 | 0.918647314 | -1.833503304 | 0.066727758 | 0.297817842 |
| KRT18 | 16339.47726 | -1.273327696 | 0.71709264 | -1.775680887 | 0.075785529 | 0.315776508 |
| KRT19 | 19752.41673 | -0.98552855 | 0.661215091 | -1.490481031 | 0.136097801 | 0.423884158 |
| KRT23 | 186.0066902 | -2.509331823 | 0.985824665 | -2.545413919 | 0.010914827 | 0.118957502 |
| KRT7 | 1530.273044 | -3.220489697 | 1.501143936 | -2.145357031 | 0.031924312 | 0.203582907 |
| KRT8 | 31213.87599 | -1.482903178 | 0.607737634 | -2.440038422 | 0.0146857 | 0.137954646 |
| KRT80 | 79.42723395 | -4.163625576 | 1.200058332 | -3.469519326 | 0.000521391 | 0.017923239 |
| KRT8P12 | 76.07779346 | 0.156178281 | 0.453727241 | 0.344211823 | 0.730686991 | 0.886891139 |
| KRTAP5-11 | 18.44448269 | -0.358460713 | 1.072203345 | -0.334321577 | 0.73813691 | 0.891242475 |
| KRTCAP2 | 1061.340445 | 0.359942853 | 0.374825121 | 0.960295436 | 0.336906546 | 0.644459616 |
| KRTCAP3 | 230.0321159 | 0.793047112 | 0.720540702 | 1.100627778 | 0.271058691 | 0.583394591 |
| KSR1 | 1567.377209 | -1.106861272 | 0.25126701 | -4.405119765 | 1.06E-05 | 0.001149871 |
| KSR2 | 67.76531148 | -0.443514461 | 0.504576361 | -0.878983828 | 0.379410044 | 0.681253679 |
| KTI12 | 206.8945506 | 0.305716503 | 0.567931789 | 0.538297924 | 0.590371384 | 0.818321084 |
| KTN1 | 6947.728682 | -0.374432282 | 0.240985192 | -1.553756386 | 0.120242538 | 0.400307118 |
| KXD1 | 2654.248569 | -0.039880765 | 0.256897599 | -0.15523993 | 0.87663217 | 0.950324661 |
| KYAT1 | 389.1425314 | 1.245005229 | 0.401512851 | 3.100785502 | 0.00193008 | 0.041067673 |

KYAT3 2217.143854 0.218388784 0.261942103 0.833729216 0.40443359 0.700367583
KYNU 86.27128916 -0.191184459 0.418788196 -0.456518261 0.648017343 0.846597949
L1CAM 207.717203 0.586276296 0.582520096 1.00644819 0.314200022 0.623858474
L1TD1 3540.293872 -1.484654367 0.622112241 -2.386473482 0.017010834 0.14772259
L2HGDH 795.3592353 0.124570326 0.266643263 0.467179721 0.640371301 0.843615231
L3HYPDH 101.7072737 0.525899527 0.356489839 1.475216033 0.140154472 0.429639018
L3MBTL1 581.9802173 -0.420502754 0.392733819 -1.070706757 0.284301304 0.598246396
L3MBTL2 846.4824718 -0.241743424 0.267547617 -0.903552894 0.366232529 0.672734435
L3MBTL3 417.5911422 -0.490823833 0.265677595 -1.847441568 0.064683169 0.29355313
L3MBTL4 19.22470869 -2.724334779 1.362799841 -1.999071835 0.045600582 0.245771986
LACC1 209.1956469 -0.695719955 0.329348102 -2.112415257 0.034650848 0.21269873
LACTB 241.3125377 0.228690657 0.33860421 0.675392242 0.499426593 0.76281557
LACTB2 410.0542045 0.63368404 0.388809831 1.629804572 0.103142807 0.368199757
LAD1 354.6264178 -1.052461311 0.62765599 -1.676812342 0.093579183 0.351044935
LAG3 62.94116638 0.817773618 0.598178452 1.367106445 0.171591943 0.474879827
LAGE3 370.9647572 1.226186712 0.465469056 2.634303391 0.008431012 0.103784183
LAMA1 3002.942679 -0.554934007 0.487406909 -1.138543581 0.254893572 0.567256913
LAMA2 124.9220798 -0.420622533 0.531827329 -0.79090056 0.429002022 0.715829569
LAMA3 17.23064594 0.107251143 1.1244028 0.095384984 0.924009043 0.971458156
LAMA5 5861.506018 -0.995798907 0.452947286 -2.198487412 0.027914391 0.191019044
LAMB1 10978.32552 -1.266435385 0.352979995 -3.587838975 0.00033343 0.013525316
LAMB2 3120.9992 -0.783609831 0.397161013 -1.97302808 0.048492363 0.25374113
LAMB3 38.41866059 -1.415953793 0.57219442 -2.474602589 0.013338451 0.131315792
LAMC1 6698.489175 -0.95520514 0.404197289 -2.363215104 0.01811715 0.151937695
LAMC2 588.7624583 -2.048929571 0.405839028 -5.048626258 4.45E-07 0.000113977
LAMC3 21.25149183 -0.475318316 0.767705524 -0.619141456 0.535823175 0.784725687
LAMP1 4707.045644 -0.518670309 0.294145801 -1.763310262 0.077848176 0.320063597
LAMP2 1852.185714 -0.718337337 0.337913149 -2.125804632 0.03351953 0.208789229
LAMP3 15.37988837 -0.93488587 0.889549469 -1.050965576 0.293274401 0.606273373
LAMP5 13.56369296 0.361993158 0.795756434 0.454904469 0.649177967 0.847198189
LAMTOR1 2108.106147 0.413896095 0.478700062 0.864625113 0.387244584 0.68748084
LAMTOR2 946.4186413 0.853244573 0.455727961 1.872267332 0.061169628 0.285364214
LAMTOR3 1106.200099 0.376371385 0.257847247 1.459668037 0.14438133 0.43572064
LAMTOR4 1840.430681 1.237237931 0.474661455 2.606569203 0.009145434 0.108141373
LAMTOR5 1758.755369 0.85935086 0.432316021 1.987783978 0.046835587 0.249225774
LANCL1 2464.728525 0.101501955 0.274880662 0.369258261 0.711935233

0.877461202
LANCL2 431.8517182 -0.351957097 0.283151585 -1.242998859
0.213868254 0.520856954
LAP3 1330.093396 -0.084536566 0.256510864 -0.329563296 0.74172996
0.892150449
LAPTM4A 6700.975095 0.265876205 0.45919814 0.57900105 0.562588465
0.801706045
LAPTM4B 12765.19308 0.714875432 0.319564684 2.237028899 0.025284451
0.181849597
LAPTM5 12.64590192 1.687883859 1.031968886 1.635595687 0.101924234
0.366163454
LARGE1 676.9339612 0.075314348 0.345117126 0.218228369 0.827251182
0.928082319
LARGE2 765.4166558 0.25630431 0.265370304 0.965836442 0.33412603
0.642136559
LARP1 5765.069405 -0.302488374 0.271215385 -1.11530684 0.264718955
0.578431696
LARP1B 560.8712586 0.229449926 0.339311987 0.676221101 0.498900277
0.76281557
LARP4 1399.477376 0.128896894 0.307906015 0.418624151 0.675490836 0.860110127
LARP4B 1343.445659 -0.385055225 0.382379014 -1.006998845
0.313935329 0.623554973
LARP6 389.3866378 0.522169887 0.387209323 1.348546781 0.177482585 0.481837595
LARP7 3549.549993 0.889874831 0.386284049 2.303679983 0.021240614 0.165290607
LARS 6140.478826 0.058357177 0.258258208 0.225964461 0.821229061 0.925277838
LARS2 1057.073405 -0.206238582 0.473303858 -0.435742447 0.663023605
0.854172139
LAS1L 1711.066592 -0.136100263 0.368225253 -0.369611431 0.711672033
0.877287654
LASP1 5460.02996 -0.830496838 0.328131255 -2.530989738 0.011374118
0.120838052
LAT 88.02501191 0.411791728 0.518466629 0.794249245 0.427050345 0.714556109
LAT2 12.96095596 0.417764155 0.905847334 0.461186051 0.644665129 0.845794147
LATS1 804.1372549 -0.309511705 0.331412154 -0.933917786 0.350346303
0.657240481
LATS2 1071.931615 -0.453195965 0.302326476 -1.499028371 0.133866272
0.420559542
LAYN 110.1391289 0.177190498 0.604587343 0.293076758 0.769463481 0.904597192
LBH 83.74399895 -0.682869413 0.585784563 -1.165734735 0.243721712
0.555013823
LBHD1 78.63554041 -1.780788272 0.558890684 -3.18629085 0.001441096
0.034538505
LBR 5362.709008 -0.492252201 0.288708574 -1.705014143 0.088191787
0.341577124
LCA5 300.7284344 -0.059486048 0.376127633 -0.158153889 0.874335538
0.949740014
LCA5L 24.85392172 0.394093391 0.612033029 0.643908698 0.519634641 0.774983751
LCAT 113.6568451 -0.372375994 0.466968146 -0.797433396 0.42519937
0.71357297
LCK 12.24935623 -0.471521087 0.915056638 -0.515291696 0.606349201
0.826611695
LCLAT1 564.5290864 0.178223391 0.283874853 0.627823808 0.530119368
0.780667922
LCMT1 1833.181838 0.017320905 0.4582768 0.037795728 0.96985055 0.989496332
LCMT2 410.4084656 0.310697992 0.325086626 0.955739078 0.339204074 0.646247019
LCOR 862.1660258 -0.307429723 0.332528323 -0.924521917 0.355214646
0.662655738
LCORL 871.8643863 0.095446363 0.279236778 0.341811575 0.732492698 0.888105842
LCP1 913.4656758 -5.411026637 1.318353872 -4.10438104 4.05E-05
0.003154003
LCTL 176.9799558 1.078814683 0.472339681 2.283980634 0.022372669 0.170396445
LDAH 1189.868948 0.065623962 0.292301526 0.224507762 0.822362228 0.925840841
LDB1 2789.774442 -0.662818933 0.5410296 -1.225106599 0.220535059
0.529137654

| | | | | | | |
|---|---|---|---|---|---|---|
| LDB2 | 329.7078249 | 0.618147843 | 0.352351356 | 1.754350687 | 0.079370435 | 0.322931692 |
| LDHA | 4299.710239 | 0.891614164 | 0.28010992 | 3.183086707 | 0.00145714 | 0.034758281 |
| LDHB | 38461.40327 | 0.773554494 | 0.258585244 | 2.991487377 | 0.00277622 | 0.051552064 |
| LDHC | 20.0979091 | 1.160785597 | 0.76445712 | 1.518444354 | 0.128902417 | 0.413453061 |
| LDHD | 89.18560667 | 0.059157506 | 0.530617135 | 0.111488119 | 0.911229286 | 0.966866396 |
| LDLR | 1493.791448 | 0.526537869 | 0.452488048 | 1.163650336 | 0.244565743 | 0.55585122 |
| LDLRAD2 | 19.00388051 | 0.711027227 | 0.891861758 | 0.797239281 | 0.425312076 | 0.713604237 |
| LDLRAD3 | 561.3589841 | 0.155489132 | 0.298104421 | 0.521592842 | 0.601953848 | 0.825021776 |
| LDLRAD4 | 10.29561652 | -0.863105983 | 1.228185025 | -0.70274915 | 0.482212092 | 0.753310328 |
| LDLRAP1 | 352.9022269 | -0.405509622 | 0.313239273 | -1.294568264 | 0.195469206 | 0.50126155 |
| LEAP2 | 33.17691571 | 0.482444795 | 0.593513984 | 0.812861715 | 0.416297347 | 0.708190968 |
| LEF1 | 793.052203 | -0.187011892 | 0.288977642 | -0.647150037 | 0.517534844 | 0.773950969 |
| LEMD1 | 64.69388636 | 1.062159308 | 0.716444745 | 1.482541836 | 0.13819618 | 0.426306273 |
| LEMD2 | 1014.39643 | -0.169693332 | 0.279566059 | -0.60698832 | 0.543858664 | 0.790238463 |
| LEMD3 | 484.5293585 | 0.046510589 | 0.281375879 | 0.165297002 | 0.868710218 | 0.947046732 |
| LENG8 | 1700.525604 | -0.395025008 | 0.412200419 | -0.958332379 | 0.337895181 | 0.645164305 |
| LEO1 | 918.481468 | 0.014690736 | 0.259535566 | 0.056603943 | 0.954860693 | 0.984323736 |
| LEPR | 506.5379964 | -0.430152406 | 0.311358422 | -1.381534515 | 0.167114673 | 0.469215449 |
| LEPROT | 1703.004278 | -0.494596977 | 0.356695434 | -1.386608656 | 0.165561127 | 0.467560858 |
| LEPROTL1 | 1592.42064 | 0.565678525 | 0.265522681 | 2.130433915 | 0.033135806 | 0.207838253 |
| LETM1 | 849.2633583 | -0.106658811 | 0.36673538 | -0.290833164 | 0.77117892 | 0.905067213 |
| LETM2 | 18.25907455 | 0.984331183 | 0.806454654 | 1.220566065 | 0.222250362 | 0.531069497 |
| LETMD1 | 1610.985221 | -0.103549054 | 0.26639193 | -0.388709426 | 0.697491109 | 0.87033849 |
| LFNG | 516.2899321 | 1.474966886 | 0.431624643 | 3.417244384 | 0.000632585 | 0.020419026 |
| LGALS1 | 174.739727 | 1.623772694 | 0.504711609 | 3.217228742 | 0.001294354 | 0.031929493 |
| LGALS3 | 291.5007076 | 0.285928783 | 0.825950473 | 0.346181511 | 0.729206309 | 0.886086417 |
| LGALS3BP | 740.3956517 | 0.583924561 | 0.434550941 | 1.343742484 | 0.179031674 | 0.483466568 |
| LGALS8 | 520.0242499 | -0.380364499 | 0.276318553 | -1.376543465 | 0.16865344 | 0.471031175 |
| LGALS9 | 33.62539597 | -0.436490029 | 0.796501451 | -0.548009081 | 0.583685667 | 0.8142179 |
| LGALSL | 864.6554533 | 0.581271003 | 0.316836939 | 1.834606168 | 0.066564064 | 0.297701125 |
| LGI1 | 299.9884167 | 1.252269127 | 0.486071539 | 2.576306216 | 0.009986216 | 0.113910443 |
| LGI2 | 44.75907797 | 1.120445576 | 0.818261207 | 1.369300616 | 0.170905325 | 0.474177888 |
| LGI4 | 60.40120425 | 0.317065566 | 0.546286106 | 0.580402033 | 0.561643534 | 0.801570466 |
| LGMN | 1794.236804 | 0.51006785 | 0.358362736 | 1.423328373 | 0.154640986 | 0.452208634 |
| LGR4 | 5687.126851 | 0.643046741 | 0.370168599 | 1.73717258 | 0.082356713 | 0.328452602 |
| LGR5 | 157.2388076 | -8.547487613 | 2.854581347 | -2.994305144 | 0.002750707 | 0.051369596 |
| LGR6 | 15.90489387 | 0.014017167 | 0.851956748 | 0.016452909 | 0.98687307 | 0.995270315 |
| LHFPL1 | 53.13604556 | -0.514976553 | 0.475019567 | -1.084116506 | 0.278313145 | 0.592005173 |
| LHFPL2 | 1381.731732 | -1.361741539 | 0.496248393 | -2.744072439 | 0.006068215 | 0.084227706 |
| LHFPL4 | 92.9910617 | -0.152781107 | 0.525925057 | -0.290499767 | 0.771433929 | 0.905067213 |
| LHFPL5 | 30.19035037 | -0.196759179 | 0.585237087 | -0.336204221 | 0.73671687 | 0.890505269 |

| | | | | | | |
|---|---|---|---|---|---|---|
| LHFPL6 | 748.1096156 | -1.418092255 | 0.47887377 | -2.961307014 | 0.003063364 | 0.054934156 |
| LHPP | 437.5416711 | 0.295915649 | 0.276648331 | 1.069645526 | 0.284778895 | 0.598639409 |
| LHX2 | 19.59399521 | 1.134354516 | 1.044052447 | 1.086491889 | 0.277261427 | 0.591007989 |
| LHX4 | 16.42326476 | -0.427615852 | 0.8333266 | -0.513143169 | 0.607851176 | 0.827607804 |
| LHX5 | 90.77804895 | 1.435940907 | 1.060742117 | 1.353713484 | 0.175827804 | 0.480020214 |
| LHX8 | 45.82164373 | 0.192321484 | 0.788367318 | 0.243949083 | 0.807270254 | 0.919244692 |
| LIAS | 619.2335002 | 0.122441293 | 0.290774198 | 0.421087199 | 0.673691409 | 0.85954302 |
| LIF | 57.83029541 | -0.239497652 | 0.575090574 | -0.416452056 | 0.677079243 | 0.86029961 |
| LIFR | 306.3110174 | -1.415867179 | 0.657835151 | -2.152313048 | 0.031372707 | 0.202023044 |
| LIG1 | 1746.383755 | 0.01951469 | 0.414148892 | 0.047119987 | 0.962417598 | 0.987343615 |
| LIG3 | 2248.142547 | -0.162809118 | 0.340194871 | -0.478576052 | 0.632240259 | 0.838768903 |
| LIG4 | 501.9852844 | -0.125422907 | 0.253571273 | -0.494625853 | 0.620864249 | 0.833349572 |
| LIMA1 | 1499.178256 | -0.64256172 | 0.365711595 | -1.757017633 | 0.078914804 | 0.322507823 |
| LIMCH1 | 265.9987296 | -0.230509209 | 0.341103611 | -0.675774755 | 0.499183665 | 0.76281557 |
| LIMD1 | 918.0705751 | -0.159869151 | 0.334193373 | -0.478373193 | 0.63238461 | 0.838768903 |
| LIMD2 | 583.7825232 | 0.629766253 | 0.372556651 | 1.69039058 | 0.090953256 | 0.34565427 |
| LIME1 | 49.98614097 | 0.112497508 | 0.719258315 | 0.156407658 | 0.875711701 | 0.950112438 |
| LIMK1 | 1440.589765 | -0.103880306 | 0.410559637 | -0.253021233 | 0.800251805 | 0.916366922 |
| LIMK2 | 1860.333034 | -0.816409003 | 0.273199117 | -2.988329576 | 0.002805069 | 0.052024084 |
| LIMS1 | 1340.736848 | -0.192340285 | 0.273148646 | -0.704159761 | 0.481333287 | 0.752733727 |
| LIMS2 | 23.00839933 | -0.685553612 | 0.737478499 | -0.929591321 | 0.352582723 | 0.659729494 |
| LIMS3 | 16.78621417 | -0.969403331 | 0.819730186 | -1.182588305 | 0.23697235 | 0.547700719 |
| LIN28A | 49555.40954 | -0.06116344 | 0.341932205 | -0.178875926 | 0.858035126 | 0.942343063 |
| LIN28B | 3811.295073 | 0.064265867 | 0.283194941 | 0.226931549 | 0.820476969 | 0.92511751 |
| LIN37 | 224.2487558 | -1.222754144 | 0.40966478 | -2.984767553 | 0.002837939 | 0.052464911 |
| LIN52 | 379.9014595 | 0.383988994 | 0.314222225 | 1.222030028 | 0.221696272 | 0.53065007 |
| LIN54 | 414.6963748 | -0.084297896 | 0.366517779 | -0.229996745 | 0.818094299 | 0.924148072 |
| LIN7A | 180.8288995 | 0.960534185 | 0.526006113 | 1.826089397 | 0.067836797 | 0.299433241 |
| LIN7B | 69.47500407 | 0.386098361 | 0.576477503 | 0.66975443 | 0.503014348 | 0.765194965 |
| LIN7C | 791.0903108 | 0.435007871 | 0.369188564 | 1.178281002 | 0.238684599 | 0.54964846 |
| LIN9 | 728.1016251 | -0.402989036 | 0.428341444 | -0.940812621 | 0.346800892 | 0.65343598 |
| LINC00475 | 18.38391063 | -0.666855667 | 1.016983663 | -0.655719154 | 0.512004843 | 0.771669528 |
| LINC00634 | 17.13514252 | 1.797568035 | 0.830108042 | 2.16546274 | 0.030352266 | 0.198737258 |
| LINC00672 | 84.09875092 | 1.292117232 | 0.461653562 | 2.798889337 | 0.005127871 | 0.076062906 |
| LINC00674 | 10.30679155 | -0.158824298 | 0.97548001 | -0.162816559 | 0.87066287 | 0.94829682 |
| LINC00680 | 174.0013331 | -0.823063816 | 0.36465261 | -2.257117577 | 0.024000729 | 0.177207433 |
| LINC00869 | 338.389841 | 1.079251613 | 0.433841298 | 2.487664538 | 0.012858496 | 0.128086828 |
| LINC00888 | 342.0850058 | 0.694269594 | 0.319963536 | 2.169839732 | 0.030018989 | 0.197749929 |

| | | | | | | |
|---|---|---|---|---|---|---|
| LINC01145 | 141.9042375 | -0.272262378 | 0.412601119 | -0.659868249 | 0.509338381 | 0.769562053 |
| LINC01347 | 44.46155261 | -0.789176556 | 1.037572771 | -0.760598753 | 0.446896765 | 0.728783276 |
| LINC01881 | 27.341229 | -1.422489036 | 0.837000782 | -1.699507416 | 0.089223619 | 0.342726775 |
| LINC02210 | 448.3787882 | 0.125705524 | 0.27993348 | 0.44905498 | 0.653391995 | 0.848827707 |
| LINC02495 | 15.19547808 | 0.872415862 | 0.874421876 | 0.997705897 | 0.318421993 | 0.628475744 |
| LINGO1 | 1464.443776 | -0.139416294 | 0.328662648 | -0.424192692 | 0.67142529 | 0.858479663 |
| LINGO2 | 26.29882548 | -1.055993754 | 0.732547998 | -1.44153524 | 0.149433529 | 0.444259469 |
| LINS1 | 379.3867467 | 0.014236043 | 0.300696911 | 0.047343497 | 0.962239461 | 0.987343615 |
| LIPA | 1128.654379 | 0.457976508 | 0.354243474 | 1.292829766 | 0.196069943 | 0.501797297 |
| LIPE | 28.21019219 | 0.661479001 | 0.708851508 | 0.933170056 | 0.350732173 | 0.65780168 |
| LIPG | 111.1570078 | 1.282326148 | 0.651482832 | 1.968319171 | 0.049031325 | 0.255021315 |
| LIPH | 10.57073503 | -1.391784331 | 1.004638762 | -1.385357985 | 0.165943031 | 0.467940842 |
| LIPT1 | 207.0664111 | 0.427240574 | 0.375596337 | 1.13749931 | 0.255329613 | 0.567810841 |
| LIPT2 | 147.8579479 | 0.632548207 | 0.447911279 | 1.412217635 | 0.15788589 | 0.457277356 |
| LITAF | 3445.461902 | 0.017392908 | 0.412353811 | 0.042179575 | 0.966355545 | 0.988700292 |
| LIX1 | 1987.610288 | 0.66555537 | 0.449210786 | 1.481610396 | 0.138443991 | 0.426723647 |
| LIX1L | 178.7762891 | 1.950956262 | 0.518107862 | 3.765540739 | 0.000166189 | 0.00848907 |
| LLGL1 | 1403.938275 | 0.379672618 | 0.457315339 | 0.830220607 | 0.406414067 | 0.701474651 |
| LLGL2 | 1474.97058 | -0.907285102 | 0.539415187 | -1.681979159 | 0.092572881 | 0.3491405 |
| LLPH | 1042.263698 | 0.507663105 | 0.450001901 | 1.128135467 | 0.259262712 | 0.572529055 |
| LMAN1 | 3446.293519 | 0.131611565 | 0.275800214 | 0.477198924 | 0.633220477 | 0.839443189 |
| LMAN2 | 2614.337024 | 0.062449647 | 0.391025595 | 0.159707313 | 0.87311164 | 0.94953238 |
| LMAN2L | 686.9416741 | -0.025328767 | 0.301300413 | -0.084064828 | 0.933004889 | 0.97462713 |
| LMBR1 | 1132.997441 | -0.339459266 | 0.293490649 | -1.156627195 | 0.247424707 | 0.56008359 |
| LMBR1L | 1298.988686 | -0.428622839 | 0.25582839 | -1.675431093 | 0.09384968 | 0.351293733 |
| LMBRD1 | 1078.526177 | 0.092948188 | 0.420654326 | 0.22096097 | 0.825122823 | 0.926980032 |
| LMBRD2 | 376.8932659 | -0.121715593 | 0.499966255 | -0.243447616 | 0.807658661 | 0.919453224 |
| LMCD1 | 59.41285536 | -0.984460195 | 0.867046814 | -1.13541758 | 0.256200397 | 0.56841419 |
| LMF1 | 198.7859793 | -0.157606316 | 0.325705491 | -0.483892106 | 0.628462443 | 0.836791621 |
| LMF2 | 601.3468819 | -0.194913433 | 0.528407677 | -0.36886942 | 0.712225058 | 0.877474733 |
| LMLN | 282.0333964 | -0.319270333 | 0.430543981 | -0.741551032 | 0.458359402 | 0.73670627 |
| LMLN2 | 54.43141659 | -2.05309097 | 0.615746181 | -3.334313772 | 0.000855101 | 0.024757143 |
| LMNA | 1096.395497 | -0.642703886 | 0.444990781 | -1.444308316 | 0.148652268 | 0.443065531 |
| LMNB1 | 4547.936288 | 0.258254154 | 0.24504006 | 1.053926261 | 0.291916679 | 0.60500928 |
| LMNB2 | 4304.688942 | 0.213812183 | 0.483152018 | 0.442536044 | 0.658101354 | 0.850865488 |
| LMNTD2 | 10.12450893 | -4.53109704 | 1.732673927 | -2.615089296 | 0.008920413 | 0.10668882 |
| LMO1 | 20.63539768 | 0.629426472 | 0.971763549 | 0.647715663 | 0.517168872 | 0.773950969 |
| LMO4 | 2253.315226 | -0.243708005 | 0.43447624 | -0.560923665 | 0.574849577 | 0.808359708 |
| LMO7 | 135.3461659 | -0.976478013 | 0.415871536 | -2.348028007 | 0.018873102 | 0.155780105 |
| LMOD1 | 121.2013488 | -0.883081673 | 0.507752082 | -1.739198526 | 0.081999849 | 0.327718573 |

| | | | | | | |
|---|---|---|---|---|---|---|
| LMOD2 | 44.40365403 | -1.194095242 | 0.994724627 | -1.200427948 | 0.22997318 | 0.538913055 |
| LMTK2 | 782.1573494 | -0.729225991 | 0.348464341 | -2.092684688 | 0.036377311 | 0.218393425 |
| LMX1A | 183.7155469 | -0.060112056 | 0.456264924 | -0.131748143 | 0.895183505 | 0.958594142 |
| LMX1B | 1121.780434 | -1.947478006 | 0.471677648 | -4.128832511 | 3.65E-05 | 0.002942285 |
| LNP1 | 248.4244528 | 0.772749385 | 0.360395771 | 2.144168846 | 0.03201936 | 0.204048219 |
| LNPEP | 329.7460103 | -0.203614471 | 0.40062219 | -0.508245615 | 0.611281105 | 0.829058038 |
| LNPK | 532.2038718 | 0.404997868 | 0.282180377 | 1.435244619 | 0.151217403 | 0.446500432 |
| LNX1 | 362.842798 | -1.715768642 | 0.64628527 | -2.654816259 | 0.007935162 | 0.100236752 |
| LNX2 | 537.3164129 | -0.35826159 | 0.277178922 | -1.292528263 | 0.196174265 | 0.501797297 |
| LONP1 | 1854.662583 | -0.120255484 | 0.375030496 | -0.320655215 | 0.74847169 | 0.894522137 |
| LONP2 | 1017.366601 | -0.261744268 | 0.241324309 | -1.084616252 | 0.278091654 | 0.592005173 |
| LONRF1 | 429.1888026 | -0.465025946 | 0.652282773 | -0.712920784 | 0.475894777 | 0.749091526 |
| LOXL1 | 266.8691296 | 0.678031163 | 0.438064315 | 1.547789082 | 0.121673086 | 0.402231976 |
| LOXL2 | 1621.009382 | -0.150812031 | 0.342894393 | -0.439820639 | 0.660067018 | 0.852207761 |
| LOXL3 | 408.7998537 | 1.2093919 | 0.55677048 | 2.172155211 | 0.029843958 | 0.197226575 |
| LOXL4 | 58.7401341 | 0.67233427 | 0.721914023 | 0.931321804 | 0.351687127 | 0.659103817 |
| LPAL2 | 33.52455043 | -1.466095013 | 0.623829533 | -2.350153264 | 0.018765682 | 0.1554129 |
| LPAR1 | 181.9240544 | -0.298198824 | 0.386777904 | -0.770982056 | 0.440717568 | 0.724218535 |
| LPAR2 | 869.3967379 | 0.286736995 | 0.327642878 | 0.87515101 | 0.381491745 | 0.682984572 |
| LPAR3 | 115.0772371 | 0.666385614 | 0.516764671 | 1.289534002 | 0.197212503 | 0.50258876 |
| LPAR4 | 537.290979 | 1.47276133 | 0.496223308 | 2.967940657 | 0.002998022 | 0.05414642 |
| LPAR5 | 82.47515193 | -1.531328499 | 0.823855026 | -1.858735397 | 0.063064653 | 0.289778852 |
| LPAR6 | 90.17950918 | 0.375923081 | 0.760880229 | 0.49406341 | 0.621261398 | 0.833349572 |
| LPCAT1 | 1563.234228 | 0.878397978 | 0.707007755 | 1.242416327 | 0.214082995 | 0.520920862 |
| LPCAT2 | 143.7090059 | -1.288675696 | 0.529245374 | -2.434930488 | 0.014894651 | 0.138715007 |
| LPCAT3 | 912.563064 | -0.420866278 | 0.233549815 | -1.802040726 | 0.071538998 | 0.307638342 |
| LPCAT4 | 1528.712896 | -0.009003276 | 0.26787551 | -0.033609927 | 0.973188206 | 0.990423175 |
| LPGAT1 | 269.6147325 | -1.271543111 | 0.522481131 | -2.433663216 | 0.014946895 | 0.139030867 |
| LPIN1 | 1345.185988 | -0.548781006 | 0.24033761 | -2.283375479 | 0.02240826 | 0.170396445 |
| LPIN2 | 1639.584404 | -1.547534186 | 0.442293124 | -3.498888188 | 0.000467202 | 0.016786538 |
| LPIN3 | 418.167894 | -0.838360872 | 0.343765807 | -2.438755844 | 0.014737922 | 0.138131624 |
| LPL | 10.36151542 | 1.406635898 | 1.256319223 | 1.119648472 | 0.262863591 | 0.576453244 |
| LPP | 1340.878278 | -0.91245514 | 0.462180664 | -1.974239106 | 0.048354561 | 0.253485504 |
| LPXN | 84.56548541 | -0.429287281 | 0.469263739 | -0.914810255 | 0.36029125 | 0.667522123 |
| LRAT | 117.5544058 | 0.749215484 | 0.619207328 | 1.209959008 | 0.226294623 | 0.534919869 |
| LRBA | 2883.125147 | -0.651556474 | 0.304066169 | -2.142811468 | 0.03212824 | 0.204453662 |
| LRCH1 | 312.334709 | -1.435076639 | 0.37496101 | -3.827268975 | 0.000129573 | 0.007243505 |
| LRCH2 | 177.5374076 | 0.702658255 | 0.523776329 | 1.3415235 | 0.179750545 | 0.484454702 |

| | | | | | | |
|---|---|---|---|---|---|---|
| LRCH3 | 1159.245808 | -0.35198551 | 0.249287346 | -1.411967016 | 0.157959673 | 0.457277356 |
| LRCH4 | 647.1554213 | -0.384596934 | 0.490616341 | -0.783905676 | 0.433095459 | 0.718657207 |
| LRFN1 | 51.0703565 | 0.73504698 | 0.473645888 | 1.551891399 | 0.120688211 | 0.400973879 |
| LRFN3 | 172.6376912 | -0.484724899 | 0.485484479 | -0.998435419 | 0.318068266 | 0.628145491 |
| LRFN4 | 199.7294963 | 1.126716191 | 0.554029748 | 2.033674537 | 0.041984421 | 0.235820549 |
| LRG1 | 46.32530096 | -1.198084914 | 0.765672568 | -1.564748386 | 0.117641918 | 0.395641036 |
| LRGUK | 26.33633003 | -0.348989213 | 0.639196285 | -0.545981292 | 0.585078797 | 0.81530545 |
| LRIF1 | 581.137244 | -0.122050716 | 0.279035622 | -0.437401917 | 0.661819894 | 0.853708836 |
| LRIG1 | 1435.259631 | 1.116629599 | 0.602572228 | 1.853104986 | 0.063867312 | 0.291583207 |
| LRIG2 | 831.7953161 | -0.437701517 | 0.297207636 | -1.472712891 | 0.140828469 | 0.430693807 |
| LRIG3 | 990.2010032 | 1.611638471 | 0.695132245 | 2.318463116 | 0.020424165 | 0.161635374 |
| LRP1 | 4949.47485 | -0.296111533 | 0.392268315 | -0.754869872 | 0.450327064 | 0.73178148 |
| LRP10 | 730.789566 | -0.932805774 | 0.397527218 | -2.346520523 | 0.018949623 | 0.156140235 |
| LRP11 | 83.07273826 | 1.00600244 | 0.537846285 | 1.870427423 | 0.061424487 | 0.285640507 |
| LRP12 | 853.8937764 | 0.260847014 | 0.33642861 | 0.775341353 | 0.43813798 | 0.721949744 |
| LRP1B | 89.79870416 | -5.545245756 | 1.212036351 | -4.575148056 | 4.76E-06 | 0.000633301 |
| LRP2 | 3348.455574 | -0.267967899 | 0.698941751 | -0.383390888 | 0.701429958 | 0.872388416 |
| LRP3 | 167.9744533 | 1.501029709 | 0.704593001 | 2.130350012 | 0.033142727 | 0.207838253 |
| LRP4 | 1482.612624 | 0.831615844 | 0.703809739 | 1.181591838 | 0.237367694 | 0.547879744 |
| LRP5 | 839.8216248 | -0.897450649 | 0.489211024 | -1.834485744 | 0.066581922 | 0.297701125 |
| LRP5L | 63.7929046 | -1.046677253 | 0.434106035 | -2.411109659 | 0.015904066 | 0.142656219 |
| LRP6 | 805.0758054 | 0.055919784 | 0.377673288 | 0.14806391 | 0.882292331 | 0.953230092 |
| LRP8 | 748.7399303 | 0.903415568 | 0.566695193 | 1.594182514 | 0.110895162 | 0.383132604 |
| LRPAP1 | 1458.268776 | -0.307191775 | 0.304460942 | -1.008969404 | 0.312989308 | 0.622994042 |
| LRPPRC | 11774.06904 | 0.147293711 | 0.256720275 | 0.573751767 | 0.566135792 | 0.804380669 |
| LRR1 | 482.5459408 | 0.865610129 | 0.261465179 | 3.310613414 | 0.000930917 | 0.02597297 |
| LRRC1 | 549.2335241 | -0.500228583 | 0.473944337 | -1.055458509 | 0.291215676 | 0.604290391 |
| LRRC10B | 27.23375442 | 0.657943038 | 0.719575881 | 0.914348376 | 0.360533819 | 0.667663818 |
| LRRC14 | 564.8266197 | 0.045138433 | 0.388584143 | 0.116161285 | 0.907524719 | 0.965304913 |
| LRRC15 | 11.04531058 | -0.269479147 | 1.400263235 | -0.19244892 | 0.847390576 | 0.938171381 |
| LRRC17 | 49.40286355 | 1.790859701 | 0.652417282 | 2.74496055 | 0.006051817 | 0.084137259 |
| LRRC2 | 15.15985777 | -0.493277011 | 1.099314426 | -0.448713306 | 0.653638485 | 0.84883826 |
| LRRC20 | 388.3467929 | 1.631826825 | 0.657046372 | 2.483579386 | 0.013006934 | 0.128720283 |
| LRRC23 | 128.1629923 | 0.00371691 | 0.377836254 | 0.009837358 | 0.992151051 | 0.99688215 |
| LRRC24 | 10.5869294 | -0.353711842 | 1.360328257 | -0.260019477 | 0.79484875 | 0.914877916 |
| LRRC27 | 254.0347067 | -0.223388419 | 0.329712172 | -0.677525546 | 0.498072568 | 0.762537616 |
| LRRC28 | 479.0568057 | 0.204337004 | 0.254410281 | 0.803179036 | 0.421871261 | 0.712063713 |
| LRRC29 | 18.49458973 | 0.302177612 | 0.777489022 | 0.388658365 | 0.697528886 | |

0.87033849
LRRC3 158.8861784 0.249241552 0.540199787 0.461387727 0.644520455 0.845794147
LRRC32 88.75429321 -1.872678579 1.014554971 -1.845812828
0.064919387 0.293855407
LRRC34 145.7869115 0.676071491 0.492550221 1.372594027 0.16987859
0.472495848
LRRC36 10.37081084 -1.682346974 1.023319239 -1.644009914
0.100174155 0.36243601
LRRC37A 32.22772514 -0.344831307 0.69656619 -0.495044566
0.620568662 0.833297063
LRRC37A17P 17.92803807 -0.036337286 1.264122789 -0.028745061
0.977067918 0.992059029
LRRC37A2 114.1134172 -0.492155677 0.668033285 -0.736723286
0.461290643 0.738942035
LRRC37A3 77.29150916 -1.567390972 0.631783679 -2.480898167
0.013105181 0.129608016
LRRC37A4P 300.3658643 -0.304599787 0.841776512 -0.361853512
0.717461501 0.880516207
LRRC37A6P 27.13382727 -0.029483839 1.057371636 -0.027884083
0.977754603 0.99226352
LRRC37B 356.5808991 0.037496195 0.358768077 0.104513744 0.916761662
0.969211928
LRRC37BP1 906.2323071 0.034463441 0.24317164 0.141724756 0.88729742
0.955846706
LRRC39 19.46607718 -0.612202468 0.796346494 -0.76876394 0.442033459
0.725453225
LRRC3B 27.10237876 -2.557303464 1.22142325 -2.093707864
0.036286012 0.218104234
LRRC4 500.530227 0.329373445 0.546778215 0.60238948 0.546914911 0.792249935
LRRC40 1631.565538 0.179585431 0.322859804 0.556233476 0.578051257 0.810201
LRRC41 1768.150088 -0.270619017 0.261051898 -1.036648341
0.299899776 0.610555175
LRRC42 1639.202477 0.203971921 0.302048493 0.675295279 0.499488183
0.76281557
LRRC45 351.7630002 0.210836255 0.53017736 0.397671178 0.690872586
0.868004248
LRRC47 1066.156074 -0.142161564 0.339845058 -0.418312878
0.675718375 0.860110127
LRRC49 559.7326229 0.578082632 0.351666732 1.643836562 0.100209967
0.36243601
LRRC4C 40.81271056 0.948023603 0.749639034 1.264640128 0.206000362
0.51233303
LRRC55 76.58614473 2.318807201 1.15695038 2.004240839 0.045044268
0.244454795
LRRC57 494.3989298 0.012124809 0.341296613 0.035525724 0.971660535
0.989665121
LRRC58 1178.982658 0.326925883 0.285532427 1.144969368 0.252221847
0.565040998
LRRC59 2526.827754 -0.209754059 0.248861135 -0.842855832
0.399309078 0.697434725
LRRC6 71.3208808 -0.033110248 0.492117893 -0.06728113 0.946357899
0.980414893
LRRC63 23.97452099 -4.562082411 1.523883054 -2.993722122
0.002755968 0.051369596
LRRC69 45.44448403 0.489590529 0.630031787 0.777088616 0.43710649
0.721225178
LRRC70 11.56936918 0.693579046 1.068619813 0.649041911 0.516311287
0.773931284
LRRC74B 12.90356032 0.771792831 0.788377538 0.978963497 0.327598016
0.637423304
LRRC75A 68.22331296 0.830454826 0.518269269 1.602361699 0.109075649
0.379241044
LRRC75B 277.3638799 0.010644649 0.299266716 0.035569104 0.971625944
0.989665121

| | | | | | | |
|---|---|---|---|---|---|---|
| LRRC8A | 1255.886284 | -0.28967034 | 0.369532079 | -0.78388415 | 0.433108091 | 0.718657207 |
| LRRC8B | 742.8244649 | -0.144901225 | 0.384981385 | -0.376385018 | 0.706630683 | 0.874911784 |
| LRRC8C | 146.1024473 | -0.459500002 | 0.416382928 | -1.103551493 | 0.269787743 | 0.582543388 |
| LRRC8D | 1322.248537 | -0.18077301 | 0.27186475 | -0.664937293 | 0.506090602 | 0.767209743 |
| LRRC8E | 87.0622551 | -1.086343339 | 0.518950468 | -2.093346872 | 0.036318201 | 0.218181832 |
| LRRC9 | 67.68025305 | 0.408616895 | 0.486634734 | 0.839678854 | 0.401088474 | 0.698568129 |
| LRRCC1 | 717.2025239 | -0.371777361 | 0.288501807 | -1.288648291 | 0.197520387 | 0.503119844 |
| LRRFIP1 | 1258.896387 | -0.480991676 | 0.302476658 | -1.590177829 | 0.111794727 | 0.384888109 |
| LRRFIP1P1 | 70.26415618 | -1.048828467 | 0.639358502 | -1.640438757 | 0.100913972 | 0.363826635 |
| LRRFIP2 | 935.5772224 | 0.029319074 | 0.292772513 | 0.100142851 | 0.920230916 | 0.969770994 |
| LRRIQ1 | 119.8488699 | 0.463489322 | 0.395034314 | 1.173288766 | 0.240680026 | 0.552353982 |
| LRRIQ3 | 77.96851238 | 0.224784478 | 0.389209873 | 0.577540533 | 0.563574366 | 0.801976485 |
| LRRK1 | 8930.235558 | -1.934737127 | 0.459225945 | -4.213039676 | 2.52E-05 | 0.002159569 |
| LRRN1 | 5016.559959 | 0.183308445 | 0.450005468 | 0.40734715 | 0.683753037 | 0.863715015 |
| LRRN2 | 80.14882779 | 0.142775636 | 0.479227742 | 0.29792857 | 0.765757682 | 0.903086893 |
| LRRN3 | 262.488346 | 0.794409712 | 0.422177423 | 1.881696341 | 0.059877257 | 0.283057446 |
| LRRN4 | 178.9180247 | -5.854469698 | 1.010183892 | -5.795449465 | 6.81E-09 | 6.46E-06 |
| LRRTM1 | 176.2576639 | 0.481770512 | 1.035561667 | 0.465226289 | 0.641769414 | 0.844038972 |
| LRRTM4 | 373.773869 | -0.645559 | 0.475018374 | -1.359019009 | 0.174140565 | 0.478455902 |
| LRSAM1 | 522.9251747 | -0.877715963 | 0.393819342 | -2.228727413 | 0.025832047 | 0.183472841 |
| LRTOMT | 115.9023591 | 0.518864824 | 0.445813143 | 1.163861658 | 0.24448008 | 0.555823062 |
| LRWD1 | 447.0400174 | 0.254217516 | 0.398734304 | 0.637561187 | 0.523759368 | 0.777662198 |
| LSAMP | 180.5742114 | 0.808397622 | 0.531582082 | 1.520739034 | 0.128325339 | 0.412550057 |
| LSG1 | 1085.072686 | -0.066930274 | 0.242193744 | -0.276350136 | 0.782279152 | 0.909909835 |
| LSM1 | 834.4401438 | 0.455171598 | 0.357308692 | 1.27388896 | 0.202702772 | 0.509290789 |
| LSM10 | 507.8492861 | 0.702843927 | 0.435234627 | 1.614862155 | 0.106340549 | 0.374140185 |
| LSM11 | 422.1327179 | 0.23500196 | 0.272038381 | 0.863855897 | 0.387667057 | 0.687775519 |
| LSM12 | 2241.17976 | 0.526667286 | 0.337079686 | 1.562441488 | 0.11818402 | 0.396061358 |
| LSM12P1 | 142.2521435 | -0.511116571 | 2.149374939 | -0.237797772 | 0.812037947 | 0.921390083 |
| LSM14A | 1302.539868 | 0.853515182 | 0.708238622 | 1.205123747 | 0.228155534 | 0.537150955 |
| LSM14B | 1893.255354 | -0.204748501 | 0.247661387 | -0.826727586 | 0.408391482 | 0.702771243 |
| LSM2 | 1098.706352 | 0.511811246 | 0.372727374 | 1.373151749 | 0.169705176 | 0.472403161 |
| LSM3 | 2156.616256 | 0.826071812 | 0.423079729 | 1.952520423 | 0.050876449 | 0.259366654 |
| LSM4 | 4249.667457 | 0.816208219 | 0.365158127 | 2.235218548 | 0.025403004 | 0.18194777 |
| LSM5 | 2079.147409 | 1.153481604 | 0.397578564 | 2.901267094 | 0.003716569 | 0.062858501 |
| LSM6 | 1111.829104 | 1.020416379 | 0.424212752 | 2.405435419 | 0.016153207 | 0.143781043 |
| LSM7 | 1287.131041 | 0.911810137 | 0.423713765 | 2.151948351 | 0.031401423 | 0.202023044 |
| LSM8 | 2292.123558 | 0.675766336 | 0.320704017 | 2.107133996 | 0.035105966 | 0.213997725 |
| LSMEM1 | 15.68258401 | 0.395553386 | 0.906481901 | 0.43636104 | 0.662574801 | 0.853902351 |
| LSR | 1758.507928 | -0.187937587 | 0.412086749 | -0.456063165 | 0.648344557 | 0.846693232 |
| LSS | 1323.656024 | -0.801530294 | 0.786330887 | -1.01932953 | 0.308046549 | |

0.617841234
LTA4H 8069.370412 0.234077278 0.300381609 0.779266343 0.435822837 0.720784523
LTB4R 132.0061798 -0.761074122 0.338013417 -2.25160921 0.024346979
0.178492903
LTB4R2 41.31181155 -0.658514972 0.634424289 -1.03797251 0.299282853
0.610246278
LTBP1 3477.59437 -1.187950395 0.346554175 -3.427892324 0.000608287
0.020018041
LTBP2 24.35264107 0.113193886 0.714511699 0.15842132 0.874124815 0.949740014
LTBP3 349.7179024 -1.679827083 0.448867169 -3.742370121 0.000182293
0.008863985
LTBP4 925.4840243 0.502591954 0.571630109 0.879225824 0.379278845 0.681253679
LTBR 146.4756162 -1.477055423 0.602540075 -2.451381218 0.014230915
0.135535502
LTK 16.22933011 -2.513701118 0.995919062 -2.524001411 0.011602746
0.121816792
LTN1 1163.202831 0.032193331 0.404503994 0.079587177 0.936565594 0.97613607
LTO1 519.2509484 -0.013009269 0.331219651 -0.039276864 0.968669652
0.989475309
LTV1 1186.586763 0.407875021 0.277173895 1.47154919 0.141142655 0.430963282
LUC7L 2260.963949 0.415802452 0.219746665 1.892190046 0.058465664 0.280248566
LUC7L2 1869.566442 0.101977421 0.255611222 0.398955179 0.68992623
0.867683896
LUC7L3 8014.291556 0.638419178 0.291610567 2.189286844 0.028575998
0.19323307
LUM 15.77638331 -3.170219825 1.92562849 -1.646329934 0.09969585
0.362001501
LURAP1 34.44350633 -0.559152994 0.605832348 -0.922950047
0.356033238 0.663487122
LURAP1L 23.59178229 -0.522571842 0.842460747 -0.620292213
0.53506542 0.784173142
LUZP1 2527.452524 -0.344567127 0.362124207 -0.951516414 0.34134229
0.648152255
LUZP2 13.92253823 -2.260099765 0.958576454 -2.357766828 0.018385237
0.153593808
LXN 237.1747091 0.188271329 0.354353233 0.531309756 0.595204146 0.821417457
LY6E 3869.919073 -0.517385911 0.750955461 -0.688970169 0.49084204
0.758593654
LY6G5B 126.3880776 -1.579584587 1.992695819 -0.792687259
0.427960055 0.715278396
LY6K 23.13972275 -2.09339639 1.273582936 -1.643706374 0.100236869
0.36243601
LY6L 14.68360674 -1.978815482 1.249109709 -1.584180691 0.113152606
0.387066107
LY75 13.37390103 -1.304096199 0.944601939 -1.380577517 0.1674089
0.469714288
LYAR 773.6344489 0.657459848 0.339664635 1.93561466 0.052914914 0.265994749
LYG1 11.80184141 1.641509367 1.007555457 1.629200017 0.103270683 0.368466495
LYN 1423.991741 -0.8787912 0.468561015 -1.875510707 0.060722498
0.284105089
LYNX1 926.583255 -0.325005413 0.342937726 -0.94770971 0.343277256
0.650169695
LYPD1 111.7703148 -0.633439823 0.46037038 -1.375935227 0.168841688
0.471383372
LYPD3 83.84695244 0.044582923 0.900865807 0.049488972 0.960529625 0.986479967
LYPD5 10.69816578 -1.744710533 1.352839617 -1.289665464 0.197166835
0.50258876
LYPD6 256.4360112 -0.734155048 0.507797817 -1.445762514 0.148243824
0.442282605
LYPD6B 76.08882124 -0.254575623 0.671133934 -0.379321637
0.704449034 0.873435301
LYPLA1 3183.974613 0.319494053 0.271053855 1.178710604 0.238513432
0.549478686
LYPLA2 2595.148253 0.083617329 0.384251286 0.217611058 0.827732168

| | | | | | | |
|---|---|---|---|---|---|---|
| | | | | | | 0.928330355 |
| LYPLAL1 | 428.8462949 | 0.51604657 | 0.419952209 | 1.228822137 | 0.2191385 | 0.52778634 |
| LYRM1 | 571.1495411 | 0.697014455 | 0.266998783 | 2.610552931 | 0.009039598 | 0.1074956 |
| LYRM2 | 730.1187356 | 0.457750407 | 0.308592016 | 1.48335143 | 0.137981065 | 0.426233866 |
| LYRM4 | 1279.963799 | 0.688471626 | 0.30217286 | 2.278403252 | 0.022702563 | 0.171650943 |
| LYRM7 | 466.5814356 | 0.566833461 | 0.325669673 | 1.740516569 | 0.081768353 | 0.327017878 |
| LYRM9 | 150.4827016 | -0.590015562 | 0.437692702 | -1.348013252 | 0.17765412 | 0.482042363 |
| LYSMD1 | 291.612021 | 0.148326402 | 0.318447941 | 0.46577912 | 0.641373612 | 0.843948481 |
| LYSMD3 | 511.5655064 | -0.020973149 | 0.332767015 | -0.063026527 | 0.949745381 | 0.981628635 |
| LYSMD4 | 430.969528 | -0.470487708 | 0.270749762 | -1.737721595 | 0.082259882 | 0.328245329 |
| LYST | 125.7450971 | -0.333298259 | 0.366929207 | -0.908344858 | 0.363696049 | 0.670353372 |
| LZIC | 870.4371153 | 0.233816681 | 0.280159238 | 0.834584944 | 0.403951442 | 0.700367583 |
| LZTFL1 | 438.6208243 | 0.193588922 | 0.328160947 | 0.589920658 | 0.555243844 | 0.796360966 |
| LZTR1 | 1080.825781 | -0.473280063 | 0.430837534 | -1.09851168 | 0.27198112 | 0.584230595 |
| LZTS2 | 1658.233552 | 0.116921224 | 0.403915081 | 0.289469815 | 0.772221876 | 0.905685717 |
| LZTS3 | 423.4797664 | -0.446126906 | 0.274135673 | -1.62739457 | 0.103653324 | 0.369064484 |
| M1AP | 110.1402394 | 0.290784611 | 0.508806422 | 0.571503421 | 0.567658442 | 0.805151447 |
| M6PR | 3333.031272 | -0.193924087 | 0.22973728 | -0.844112403 | 0.398606598 | 0.697197187 |
| MAATS1 | 326.8085672 | 0.093278749 | 0.338159836 | 0.275842187 | 0.782669279 | 0.909909835 |
| MAB21L3 | 112.077798 | 0.317690269 | 0.862245727 | 0.368445165 | 0.712541325 | 0.877574643 |
| MAB21L4 | 75.73754762 | -1.968923998 | 1.295887102 | -1.519363836 | 0.12867094 | 0.413310782 |
| MACC1 | 169.3125279 | -1.070962009 | 0.478396058 | -2.238651407 | 0.025178607 | 0.18153038 |
| MACF1 | 5325.501536 | -0.213881664 | 0.510747975 | -0.418761648 | 0.675390336 | 0.860097943 |
| MACO1 | 608.0978491 | -0.496908328 | 0.258448036 | -1.92266243 | 0.054522458 | 0.269872826 |
| MACROD1 | 253.284441 | 0.206478777 | 0.549202914 | 0.375960817 | 0.706946026 | 0.87508797 |
| MACROD2 | 94.46530824 | -0.104603858 | 0.37487031 | -0.279040125 | 0.780214032 | 0.909103544 |
| MAD1L1 | 564.4993926 | -0.127422185 | 0.35618297 | -0.35774362 | 0.720535186 | 0.88214152 |
| MAD2L1 | 2504.979498 | 1.514034883 | 0.295111346 | 5.130385201 | 2.89E-07 | 8.32E-05 |
| MAD2L1BP | 635.6393177 | 0.597046555 | 0.307369334 | 1.942440212 | 0.052083831 | 0.262949685 |
| MAD2L2 | 2701.623533 | 0.546090901 | 0.327647186 | 1.666704075 | 0.095573262 | 0.354819129 |
| MADD | 848.204102 | -0.415628163 | 0.482455051 | -0.861485774 | 0.388970553 | 0.688870316 |
| MAEA | 1544.173034 | 0.032620196 | 0.328612999 | 0.099266299 | 0.920926836 | 0.970032703 |
| MAF | 173.2737834 | -0.956139472 | 0.720281871 | -1.327451808 | 0.184359272 | 0.488866012 |
| MAF1 | 1597.679774 | 0.124956186 | 0.311809311 | 0.400745525 | 0.688607489 | 0.867149602 |
| MAFB | 1027.966902 | -0.02000648 | 0.690271078 | -0.028983512 | 0.97687774 | 0.992055261 |
| MAFF | 83.51985515 | -0.569719119 | 0.693842615 | -0.821107131 | 0.411585249 | 0.704686727 |
| MAFG | 1190.159596 | 0.338864533 | 0.305535492 | 1.109084023 | 0.267393936 | 0.580680419 |
| MAFK | 160.329695 | -0.456359994 | 0.517860919 | -0.88124046 | 0.378187685 | 0.681253679 |

| | | | | | | |
|---|---|---|---|---|---|---|
| MAGED1 | 7574.211247 | -0.031029707 | 0.302186986 | -0.102683795 | 0.918213935 | 0.969677217 |
| MAGED2 | 4908.02372 | 0.175853993 | 0.262355369 | 0.670289286 | 0.502673396 | 0.764968568 |
| MAGED4 | 1205.19618 | -0.575830573 | 0.469385382 | -1.226775684 | 0.219906912 | 0.528494327 |
| MAGED4B | 2275.298229 | -0.158063885 | 0.304692741 | -0.518764852 | 0.603924731 | 0.825641353 |
| MAGEE1 | 194.3895089 | 0.378340527 | 0.33566126 | 1.127149812 | 0.259679146 | 0.572948272 |
| MAGEF1 | 1877.957479 | 0.745072976 | 0.411741295 | 1.809565823 | 0.070363144 | 0.304732874 |
| MAGEH1 | 518.5407044 | 0.891142297 | 0.404734588 | 2.201794272 | 0.027679844 | 0.190295153 |
| MAGEL2 | 129.8822947 | -0.000557704 | 0.408517331 | -0.001365191 | 0.998910736 | 0.999305953 |
| MAGI1 | 707.1965053 | -0.154391933 | 0.269653965 | -0.572555769 | 0.566945515 | 0.804595923 |
| MAGI2 | 128.5885329 | 0.243054037 | 0.410114164 | 0.592649701 | 0.55341561 | 0.795892332 |
| MAGI3 | 1263.507706 | -1.76067433 | 0.50156377 | -3.510369838 | 0.000447484 | 0.016477615 |
| MAGIX | 13.51400266 | 0.112099762 | 0.858105131 | 0.130636397 | 0.896062949 | 0.958964824 |
| MAGOH | 1257.00826 | 0.792542361 | 0.413532913 | 1.9165158 | 0.055299474 | 0.272191885 |
| MAGOHB | 691.7312831 | 0.768851971 | 0.373273081 | 2.059757348 | 0.039421744 | 0.228786879 |
| MAGT1 | 1912.521589 | 0.119505181 | 0.274484965 | 0.435379699 | 0.663286843 | 0.854409382 |
| MAIP1 | 418.6463114 | 0.322936492 | 0.381063621 | 0.847460828 | 0.396738312 | 0.695422297 |
| MAK16 | 831.1252692 | 0.196801255 | 0.26410249 | 0.74517001 | 0.456168951 | 0.735399427 |
| MAL2 | 1398.816032 | -0.52765992 | 0.589156275 | -0.895619621 | 0.370455953 | 0.676072087 |
| MALL | 97.45332609 | 0.310801098 | 0.372567431 | 0.834214353 | 0.404160205 | 0.700367583 |
| MALRD1 | 12.71852101 | 0.272004765 | 0.829475031 | 0.327923994 | 0.742969131 | 0.892445344 |
| MALSU1 | 525.9354355 | 1.158560268 | 0.465888875 | 2.486773846 | 0.012890732 | 0.128239534 |
| MALT1 | 1872.4504 | -0.958294685 | 0.298490219 | -3.210472651 | 0.001325169 | 0.032530964 |
| MAMDC2 | 189.915987 | 1.247865493 | 0.617603198 | 2.020497137 | 0.043331847 | 0.239441079 |
| MAMDC4 | 155.8472527 | 0.651495727 | 0.525370338 | 1.240069489 | 0.214949693 | 0.522178029 |
| MAML1 | 1749.633743 | -0.139381513 | 0.308424489 | -0.451914546 | 0.651330545 | 0.847876685 |
| MAML2 | 101.6710364 | -1.189131481 | 0.691925093 | -1.718584126 | 0.085690124 | 0.33574506 |
| MAML3 | 609.7838808 | -0.604238957 | 0.362432012 | -1.667178775 | 0.095478862 | 0.354819129 |
| MAMSTR | 36.94819172 | -0.013619922 | 0.647605997 | -0.021031186 | 0.983220779 | 0.993766984 |
| MAN1A1 | 384.5130031 | -1.256526277 | 0.480653212 | -2.614205511 | 0.008943522 | 0.10677261 |
| MAN1A2 | 1404.032668 | -0.148009187 | 0.344812071 | -0.429245955 | 0.667744243 | 0.856696248 |
| MAN1B1 | 1208.246789 | -0.324949289 | 0.362963733 | -0.895266549 | 0.370644617 | 0.676128642 |
| MAN1C1 | 100.3010343 | 0.731697141 | 0.475627885 | 1.538381505 | 0.123955362 | 0.405066642 |
| MAN2A1 | 2124.749685 | -0.538020245 | 0.222272373 | -2.420544838 | 0.015497268 | 0.140797887 |
| MAN2A2 | 856.2221489 | -0.533331995 | 0.392767491 | -1.357882228 | 0.174501057 | 0.478578744 |
| MAN2B1 | 2567.387708 | -0.755979471 | 0.315734885 | -2.394348889 | 0.016649903 | 0.146056278 |
| MAN2B2 | 511.8504628 | -0.77723719 | 0.327229143 | -2.375207731 | 0.017539076 | |

0.149594125
MAN2C1 1534.841423 -0.201686495 0.354634801 -0.568716025
0.569548871 0.805878187
MANBA 1189.98567 -0.581239021 0.246871608 -2.354418255 0.018551724
0.154302741
MANBAL 901.9403384 -0.02389482 0.252463462 -0.094646645 0.924595499
0.971780852
MANEA 631.4737107 -0.952214127 0.592214598 -1.607886955 0.107859939
0.37651706
MANEAL 39.96839741 -0.915712532 0.804135003 -1.138754721
0.254805472 0.567256913
MANF 1139.16946 0.642513686 0.464995639 1.381762821 0.167044538 0.469215449
MANSC1 171.8114498 -0.548783877 0.482302429 -1.137841828
0.255186535 0.567631254
MAOA 249.5501335 0.282954655 0.359699501 0.786641778 0.431491596 0.718177757
MAOB 14.43337215 0.300690692 0.791833282 0.379739901 0.704138498 0.873421952
MAP10 221.1890036 -0.055810875 0.329832126 -0.169209943 0.865631509
0.945327931
MAP1A 858.5243751 0.384826772 0.478867505 0.803618471 0.421617352 0.711972045
MAP1B 9437.014442 0.502281703 0.429025591 1.170749982 0.241699288 0.552977821
MAP1LC3A 76.83307648 0.366233376 0.768984108 0.476256104 0.633891934
0.839809222
MAP1LC3B 1616.906483 0.197936696 0.377938849 0.523726778 0.600468588
0.824475293
MAP1LC3B2 48.11638491 0.416250506 0.641495796 0.648874878 0.516419254
0.773950426
MAP1LC3C 14.13948404 -0.896962828 0.998155652 -0.898620197
0.368854997 0.674773805
MAP1S 410.5112069 -0.38176556 0.578274162 -0.660180906 0.509137744
0.769488814
MAP2 1394.55143 0.680770516 0.555865224 1.224704274 0.220686663 0.529377876
MAP2K1 1634.780605 0.192188659 0.254805262 0.754257024 0.450694902
0.732054295
MAP2K2 3155.46271 -0.077650865 0.299979628 -0.258853794
0.795748051 0.915261082
MAP2K3 393.0136483 -0.091476436 0.310975581 -0.294159547
0.76863599 0.904372313
MAP2K4 1282.433858 -0.217150352 0.272238929 -0.797646217
0.425075823 0.71357297
MAP2K5 707.1700001 -0.423692026 0.24172086 -1.752815317
0.079633713 0.32319892
MAP2K6 951.677721 0.446884806 0.382550682 1.168171506 0.242737592
0.553922067
MAP2K7 822.1023633 -0.044858025 0.390267725 -0.114941673
0.908491354 0.965717652
MAP3K1 296.6688475 0.259139105 0.368613423 0.703010494 0.48204921
0.753236025
MAP3K10 251.2051994 -0.597519035 0.634353691 -0.941933566
0.346226654 0.652902991
MAP3K11 745.2323745 -0.137633845 0.411915569 -0.334131203
0.738280555 0.891242475
MAP3K12 650.5145136 0.318730034 0.416115882 0.765964597 0.443697368
0.726848446
MAP3K13 590.1970994 0.203300456 0.39290124 0.517433989 0.604853238
0.826017506
MAP3K14 210.0450583 -0.449440934 0.467017014 -0.962365226
0.335866175 0.644086928
MAP3K15 104.545666 -1.767415047 0.640926248 -2.757595045
0.005822828 0.082336465
MAP3K2 755.2672939 0.15215346 0.281807078 0.539920648 0.589251758
0.818256545
MAP3K20 557.8048855 -0.193528333 0.24651722 -0.785049958
0.432424278 0.718639059
MAP3K21 167.471261 0.090878425 0.462202748 0.196620261 0.844124714

| | | | | | | |
|---|---|---|---|---|---|---|
| | | | | | | 0.936837827 |
| MAP3K3 | 692.672426 | 0.010528657 | 0.475601486 | 0.022137561 | 0.982338225 | 0.993518428 |
| MAP3K4 | 1502.558617 | -0.235965315 | 0.383807587 | -0.614801068 | 0.538686104 | 0.786598616 |
| MAP3K5 | 41.67078634 | -1.822245162 | 0.838332031 | -2.173655656 | 0.029731005 | 0.197143498 |
| MAP3K6 | 294.5321466 | -0.263348846 | 0.335669201 | -0.784548732 | 0.432718199 | 0.718639059 |
| MAP3K7 | 2153.863573 | -0.013618252 | 0.248635245 | -0.054772009 | 0.9563201 | 0.984447503 |
| MAP3K8 | 252.0963983 | 0.10278752 | 0.361678618 | 0.284195732 | 0.776260386 | 0.907435692 |
| MAP3K9 | 735.4304528 | -0.435345014 | 0.463809589 | -0.938628748 | 0.347921387 | 0.654471837 |
| MAP4 | 5529.028635 | -0.653673264 | 0.317990621 | -2.055636931 | 0.039817525 | 0.229888601 |
| MAP4K1 | 70.78673553 | 0.623884641 | 0.484675041 | 1.287222549 | 0.19801673 | 0.503707547 |
| MAP4K2 | 815.2769251 | -0.272001296 | 0.287358547 | -0.946557178 | 0.343864473 | 0.650875599 |
| MAP4K3 | 1253.587951 | -0.493448511 | 0.245098075 | -2.013269631 | 0.044086271 | 0.241443505 |
| MAP4K4 | 6188.621245 | -0.277506085 | 0.265521589 | -1.045135673 | 0.295960269 | 0.607250911 |
| MAP4K5 | 1467.843843 | 0.016737042 | 0.292165878 | 0.057286093 | 0.954317299 | 0.984024178 |
| MAP6 | 23.85109351 | 1.607966948 | 0.785489296 | 2.047089572 | 0.040649291 | 0.23195346 |
| MAP6D1 | 51.85263815 | 1.114050982 | 0.47571876 | 2.341826887 | 0.019189613 | 0.157365202 |
| MAP7 | 821.9390311 | -1.602492924 | 0.62432116 | -2.566776567 | 0.010264873 | 0.11507205 |
| MAP7D1 | 2485.483127 | 0.33761283 | 0.345533366 | 0.977077361 | 0.328530855 | 0.637981939 |
| MAP7D2 | 39.56517896 | 0.358364826 | 0.57729971 | 0.620760447 | 0.534757251 | 0.784009474 |
| MAP7D3 | 668.1380651 | 0.33587763 | 0.257569444 | 1.304027469 | 0.192224217 | 0.498075763 |
| MAP9 | 557.4552369 | 0.333340649 | 0.343328935 | 0.970907532 | 0.331594327 | 0.639505603 |
| MAPK1 | 2275.992057 | -0.011500796 | 0.277829881 | -0.0413951 | 0.966980919 | 0.988749915 |
| MAPK10 | 915.0530672 | 1.16017046 | 0.613322922 | 1.891614381 | 0.058542378 | 0.280288923 |
| MAPK11 | 19.03717527 | 1.172830304 | 0.738031915 | 1.589132232 | 0.112030542 | 0.384920166 |
| MAPK12 | 197.8431609 | 1.29395279 | 0.345647996 | 3.743556461 | 0.000181434 | 0.008851388 |
| MAPK13 | 246.5322184 | -0.924767511 | 0.494347764 | -1.870682097 | 0.061389158 | 0.285640507 |
| MAPK14 | 1453.785362 | -0.182762678 | 0.287971892 | -0.634654571 | 0.525653729 | 0.778399822 |
| MAPK15 | 109.6817201 | 1.326306795 | 0.743099838 | 1.78482988 | 0.074288915 | 0.313236304 |
| MAPK1IP1L | 3097.436045 | 0.115156723 | 0.408243003 | 0.282078866 | 0.777883038 | 0.908018171 |
| MAPK3 | 1796.853384 | -0.15509886 | 0.286131126 | -0.542055182 | 0.587780492 | 0.817043966 |
| MAPK4 | 403.505889 | 0.18587513 | 0.383446012 | 0.484749155 | 0.627854294 | 0.836495784 |
| MAPK6 | 3832.931851 | -0.033355528 | 0.321486733 | -0.103753979 | 0.917364589 | 0.969661544 |
| MAPK7 | 464.2668782 | -0.456234889 | 0.291282602 | -1.566296391 | 0.117279245 | 0.395279984 |
| MAPK8 | 1276.801428 | 0.254664582 | 0.336011943 | 0.757903362 | 0.448508838 | 0.729796208 |
| MAPK8IP1 | 445.5974894 | 0.473016886 | 0.326556839 | 1.448497867 | 0.147477862 | |

0.441125127

| | | | | | | |
|---|---|---|---|---|---|---|
| MAPK8IP2 | 34.93993358 | -0.408154101 | 0.577840455 | -0.706343936 | 0.479974277 | 0.752036291 |
| MAPK8IP3 | 528.0507984 | 0.073789798 | 0.715923221 | 0.10306943 | 0.917907867 | 0.969677217 |
| MAPK9 | 1172.368751 | -0.146785277 | 0.244871006 | -0.59943919 | 0.54888005 | 0.793468239 |
| MAPKAP1 | 1943.197568 | -0.224744832 | 0.287314177 | -0.782226741 | 0.434081332 | 0.719296316 |
| MAPKAPK2 | 2780.439362 | 0.227656302 | 0.234400237 | 0.971228978 | 0.331434267 | 0.639312048 |
| MAPKAPK3 | 1617.449292 | 0.006966603 | 0.263330096 | 0.026455778 | 0.978893805 | 0.992823177 |
| MAPKAPK5 | 1443.343406 | 0.19033655 | 0.284933601 | 0.668003174 | 0.504131566 | 0.765890246 |
| MAPKBP1 | 514.3120208 | -0.383116679 | 0.602145189 | -0.636252994 | 0.524611538 | 0.777662198 |
| MAPRE1 | 5951.08887 | 0.280339215 | 0.229436176 | 1.221861434 | 0.221760032 | 0.53065007 |
| MAPRE2 | 980.1085294 | 0.014531362 | 0.35928166 | 0.040445599 | 0.967737877 | 0.989056274 |
| MAPRE3 | 167.5136784 | 0.734275294 | 0.440875166 | 1.665494794 | 0.095814081 | 0.355402305 |
| MAPT | 29.16373662 | 0.32787111 | 0.710190897 | 0.461666168 | 0.644320736 | 0.845794147 |
| MARCH1 | 743.0233824 | -0.013101207 | 0.430248409 | -0.030450332 | 0.975707904 | 0.991590609 |
| MARCH2 | 413.3905147 | 0.05210551 | 0.300721915 | 0.173268085 | 0.862440709 | 0.944563095 |
| MARCH3 | 21.8844053 | 0.591059652 | 0.629157397 | 0.9394464 | 0.3475016 | 0.654087688 |
| MARCH5 | 915.2661631 | 0.143076552 | 0.30594408 | 0.467655893 | 0.640030688 | 0.843538624 |
| MARCH6 | 3676.725967 | 0.16459283 | 0.305175103 | 0.539338986 | 0.589652973 | 0.81828099 |
| MARCH7 | 3746.50428 | 0.160488993 | 0.23425706 | 0.685097786 | 0.493282217 | 0.759446369 |
| MARCH8 | 503.0423923 | -1.660317131 | 2.290833389 | -0.724765555 | 0.468595871 | 0.744337782 |
| MARCH9 | 184.0606356 | 0.442937975 | 0.34343642 | 1.289723366 | 0.197146724 | 0.50258876 |
| MARCKS | 5938.394945 | 0.941408546 | 0.398832917 | 2.360408346 | 0.018254829 | 0.152838855 |
| MARCKSL1 | 11499.28734 | 0.553570011 | 0.399733764 | 1.384846768 | 0.166099327 | 0.468031848 |
| MARF1 | 313.4103978 | 1.342381072 | 0.529399373 | 2.53566804 | 0.011223307 | 0.120297968 |
| MARK1 | 618.5797092 | -0.665906501 | 0.318380505 | -2.091542952 | 0.03647942 | 0.218746754 |
| MARK2 | 1331.088105 | -0.737624739 | 0.308521379 | -2.390838329 | 0.016809953 | 0.146565402 |
| MARK3 | 2365.688274 | 0.107692835 | 0.264443856 | 0.407242719 | 0.683829729 | 0.863739973 |
| MARK4 | 1405.067771 | -0.236377372 | 0.263521863 | -0.896993402 | 0.369722438 | 0.675302084 |
| MARS | 3919.347227 | -0.279570439 | 0.300774052 | -0.929503181 | 0.352628377 | 0.659729494 |
| MARS2 | 299.9536473 | 0.463512408 | 0.316440184 | 1.464771008 | 0.142983404 | 0.433233718 |
| MARVELD1 | 1214.676292 | 0.177113788 | 0.255096459 | 0.69430124 | 0.487493314 | 0.756439734 |
| MARVELD2 | 164.1705608 | -0.722729215 | 0.519694438 | -1.390681065 | 0.164322162 | 0.466395596 |
| MARVELD3 | 388.0661061 | -1.10592998 | 0.749904954 | -1.474760201 | 0.140277025 | 0.429883841 |
| MAS1 | 25.93959394 | -0.712987477 | 0.679675681 | -1.04901131 | 0.294172913 | 0.606454309 |
| MASP1 | 86.81319977 | -1.273574672 | 0.440101118 | -2.893822847 | 0.003805828 | |

0.063596264
MAST1 154.2108847 0.35977856 0.584588816 0.615438665 0.538265064 0.786555507
MAST2 1381.272452 -1.095432973 0.378546936 -2.893783754 0.003806302
0.063596264
MAST3 201.5410423 0.069178446 0.434596335 0.159178622 0.873528147 0.949713022
MAST4 131.3194116 -0.31007628 0.370013552 -0.838013305 0.402023231
0.699116854
MASTL 744.0235449 0.325757152 0.338771797 0.961582856 0.336259187 0.64425371
MAT2A 4673.448399 0.27377417 0.33219449 0.824138203 0.409861035 0.703966274
MAT2B 1768.491203 0.273244893 0.263609684 1.036551042 0.29994514 0.610555175
MATK 16.41538084 0.215331104 0.866296839 0.248565035 0.803697258 0.917969611
MATN2 122.7114599 -0.386201572 0.404184988 -0.955506967 0.339321384
0.646260212
MATN3 141.7406271 0.375628045 0.431835701 0.869840183 0.384387749 0.685097102
MATR3 18922.73092 0.257417232 0.366859748 0.701677505 0.482880305 0.753835883
MAU2 1282.721961 -0.563624316 0.347740921 -1.62081677 0.105056941
0.371346423
MAVS 2657.38565 -0.523647471 0.260703051 -2.008597404 0.044579847
0.242669845
MAX 876.3134906 -0.046002078 0.347270595 -0.13246753 0.894614504
0.958237013
MAZ 6568.384106 0.465763685 0.352852627 1.319994946 0.186836705 0.491503322
MB 65.42129664 -0.042650696 0.947110567 -0.045032436 0.964081455
0.98831541
MB21D2 344.7544524 -0.455718007 0.386064784 -1.180418485
0.237833814 0.548605259
MBD1 1440.151845 -0.350022609 0.254122133 -1.377379472 0.168394956
0.470634525
MBD2 740.2127761 0.047684468 0.440504947 0.108249563 0.91379773 0.968086893
MBD3 1079.142602 0.576282551 0.531744506 1.083758356 0.278471954 0.592005173
MBD4 1119.060886 0.225502432 0.230684273 0.977537087 0.328303326 0.63781403
MBD5 530.3065551 -0.307362823 0.296181484 -1.03775165 0.299385692
0.610246278
MBD6 1044.508548 -0.834712724 0.375076146 -2.225448705 0.026051131
0.184166686
MBIP 903.3111894 0.265585574 0.300505123 0.883797161 0.376805719 0.68029735
MBL1P 11.71133435 -1.898716874 1.414660524 -1.342171384 0.179540432
0.484443207
MBLAC1 12.32798802 1.599402951 0.857609092 1.864955684 0.062187616
0.287461403
MBLAC2 156.2254253 0.323378761 0.37859001 0.854166123 0.393012951
0.692071436
MBNL1 248.1372001 -0.140161179 0.320260663 -0.43764719 0.661642057
0.853552011
MBNL2 371.6519906 -0.913376039 0.485129918 -1.882745231 0.059734903
0.282669434
MBNL3 1354.919979 -0.701999153 0.279709411 -2.509744491 0.012081855
0.12494466
MBOAT1 57.13435989 -2.926269766 0.610371272 -4.794245565 1.63E-06
0.000260763
MBOAT2 1390.54815 -0.632738687 0.331188561 -1.91050888 0.056067724
0.274279592
MBOAT7 459.5088188 -1.322093596 0.523603553 -2.524989735
0.011570167 0.121704879
MBP 109.9760712 -1.499449763 0.579537989 -2.587319194 0.009672594
0.111651769
MBTD1 1920.137468 -0.231037334 0.254594058 -0.907473394 0.364156513
0.670870487
MBTPS1 2265.944036 -0.328573006 0.329568099 -0.996980616
0.318773919 0.628804869
MBTPS2 436.7358773 -0.198525931 0.324117303 -0.612512597
0.540198666 0.787481586
MC1R 90.71852688 0.383530618 0.403445815 0.950637244 0.341788556 0.648645757
MC2R 10.68917977 -2.304654807 1.1964545 -1.9262369 0.0540748

| | | | | | | |
|---|---|---|---|---|---|---|
| | | | | | | 0.268797115 |
| MCAT | 242.2495553 | 0.478651108 | 0.314644115 | 1.521246021 | 0.128198111 | 0.412317905 |
| MCC | 852.6149155 | -0.765585438 | 0.251008087 | -3.050042921 | 0.002288087 | 0.045554544 |
| MCCC1 | 1236.843937 | -0.655156774 | 0.234376394 | -2.795318943 | 0.005184854 | 0.076591444 |
| MCCC2 | 2109.512496 | -0.513338997 | 0.308930456 | -1.661665226 | 0.096579917 | 0.356760147 |
| MCEE | 198.0139304 | 0.217449071 | 0.435962427 | 0.498779385 | 0.617934812 | 0.832176095 |
| MCF2L | 525.1913479 | 0.331045269 | 0.561051872 | 0.590043961 | 0.555161177 | 0.796360966 |
| MCF2L2 | 179.8215122 | 0.096702648 | 0.391571653 | 0.246960287 | 0.804938959 | 0.918727727 |
| MCFD2 | 3096.800889 | 0.250915936 | 0.246872254 | 1.016379653 | 0.309448631 | 0.61915097 |
| MCHR1 | 17.81339704 | -0.254697333 | 0.777605189 | -0.327540681 | 0.743258979 | 0.892534058 |
| MCL1 | 6953.117892 | -0.514527797 | 0.302910669 | -1.698612331 | 0.089392252 | 0.342726775 |
| MCM10 | 727.2852115 | 0.106550359 | 0.38404068 | 0.277445502 | 0.781438047 | 0.909755868 |
| MCM2 | 3078.393761 | 0.170863559 | 0.391266471 | 0.436693588 | 0.66233358 | 0.853863251 |
| MCM3 | 5070.114047 | -0.211745486 | 0.294137774 | -0.71988539 | 0.471595564 | 0.746278951 |
| MCM3AP | 2564.406099 | -0.491395473 | 0.385824427 | -1.273624579 | 0.202796497 | 0.509290789 |
| MCM4 | 5529.446742 | 0.218406583 | 0.253369588 | 0.862007888 | 0.388683177 | 0.688626939 |
| MCM5 | 2509.585134 | -0.483468378 | 0.39391719 | -1.22733506 | 0.219696683 | 0.528259775 |
| MCM6 | 3007.821755 | 0.117213898 | 0.255991728 | 0.457881583 | 0.647037519 | 0.846597949 |
| MCM7 | 6299.801065 | 0.154157855 | 0.256885944 | 0.600102334 | 0.548438037 | 0.793396916 |
| MCM8 | 1204.159697 | -0.176910124 | 0.352092218 | -0.502453946 | 0.61534824 | 0.830958873 |
| MCM9 | 377.8432139 | -0.398733167 | 0.337400769 | -1.181779071 | 0.237293374 | 0.547879744 |
| MCMBP | 2240.100449 | 0.194271513 | 0.249349311 | 0.779113895 | 0.435912626 | 0.720784523 |
| MCMDC2 | 14.10971218 | -0.473816379 | 0.960391867 | -0.493357342 | 0.621760118 | 0.833640266 |
| MCOLN1 | 336.3386349 | -0.500102752 | 0.360927156 | -1.385605773 | 0.165867314 | 0.467940842 |
| MCOLN2 | 28.49338738 | 0.066022485 | 0.596981041 | 0.11059394 | 0.911938353 | 0.967077922 |
| MCOLN3 | 92.41231851 | -1.76398178 | 0.547890426 | -3.219588619 | 0.001283747 | 0.031719417 |
| MCPH1 | 309.0609044 | -0.209575962 | 0.328603218 | -0.637778177 | 0.523618087 | 0.777662198 |
| MCRIP1 | 1550.074469 | 0.692639694 | 0.549183003 | 1.261218375 | 0.20723018 | 0.514127402 |
| MCRIP2 | 117.9026581 | 1.556272343 | 0.519850431 | 2.993692514 | 0.002756236 | 0.051369596 |
| MCRIP2P1 | 25.93927564 | 0.736302303 | 0.780807795 | 0.943000707 | 0.345680541 | 0.652517297 |
| MCRS1 | 2082.862713 | 0.138148612 | 0.229715106 | 0.601391063 | 0.54757955 | 0.792759743 |
| MCTP1 | 26.4523565 | -0.795540945 | 0.722992193 | -1.100345139 | 0.271181772 | 0.583394591 |
| MCTP2 | 155.2180573 | -3.293852799 | 0.732557349 | -4.496375342 | 6.91E-06 | 0.000825705 |
| MCTS1 | 974.8173019 | 0.575471883 | 0.339158649 | 1.696763108 | 0.089741456 | 0.343025354 |
| MCTS2P | 14.70094086 | -0.183591129 | 1.287300521 | -0.142617148 | 0.886592554 | 0.955290883 |
| MCU | 1017.212614 | -0.033731758 | 0.307697699 | -0.109626294 | 0.912705757 | 0.967485958 |
| MCUB | 230.8077196 | -0.425984364 | 0.398501294 | -1.068966074 | 0.285084957 | 0.598867887 |
| MCUR1 | 1542.429243 | 0.168000658 | 0.285282766 | 0.588891718 | 0.555933914 | 0.796719574 |
| MDC1 | 1992.931301 | 0.011374348 | 0.441406656 | 0.025768411 | 0.979442057 | 0.992858175 |
| MDFI | 1766.097217 | -0.018852066 | 0.501257199 | -0.037609567 | 0.96999898 | |

| | | | | | | |
|---|---|---|---|---|---|---|
| | | | | | | 0.989496332 |
| MDFIC | 122.2637571 | 0.194301483 | 0.366995013 | 0.529439028 | 0.59650093 | 0.821812117 |
| MDGA1 | 208.2919402 | 0.458676014 | 0.675313812 | 0.679204254 | 0.497008452 | 0.761325016 |
| MDH1 | 6317.048883 | 0.82565587 | 0.330181299 | 2.50061367 | 0.012397834 | 0.126591155 |
| MDH1B | 71.28406733 | -0.264165968 | 0.422849871 | -0.624727561 | 0.532149881 | 0.782369637 |
| MDH2 | 5386.330927 | 0.739024003 | 0.37551876 | 1.968008212 | 0.049067093 | 0.255021315 |
| MDK | 18850.67008 | 0.46333624 | 0.482034034 | 0.961210635 | 0.33644627 | 0.644310322 |
| MDM1 | 451.5083129 | 0.416015198 | 0.372865456 | 1.115724695 | 0.264539997 | 0.578123925 |
| MDM2 | 1850.735434 | 0.187642733 | 0.482929376 | 0.388551086 | 0.697608257 | 0.87033849 |
| MDM4 | 1294.247311 | -0.167466994 | 0.419705904 | -0.399010336 | 0.689885588 | 0.867683896 |
| MDN1 | 2731.856106 | -0.668316732 | 0.397643964 | -1.680691255 | 0.0928229 | 0.349440669 |
| MDP1 | 144.0014099 | 0.782047431 | 0.410063432 | 1.907137702 | 0.056502758 | 0.275406162 |
| ME1 | 724.4780583 | -0.69141093 | 0.467526664 | -1.478869512 | 0.139175193 | 0.427760708 |
| ME2 | 1563.715889 | 0.796047157 | 0.332729439 | 2.392475876 | 0.016735128 | 0.146249211 |
| ME3 | 39.88751225 | 1.269172101 | 0.683926679 | 1.85571369 | 0.063494378 | 0.290667838 |
| MEA1 | 1195.6345 | 0.288288521 | 0.291911062 | 0.987590256 | 0.32335336 | 0.63322621 |
| MEAF6 | 1064.606664 | 0.403798342 | 0.228328135 | 1.768500159 | 0.076977326 | 0.31844659 |
| MEAK7 | 939.4818942 | -0.521384811 | 0.270778089 | -1.92550591 | 0.054166097 | 0.269074611 |
| MECOM | 28.37686371 | 0.047708719 | 0.587812348 | 0.081163179 | 0.935312182 | 0.975231691 |
| MECP2 | 964.5296924 | 0.003977741 | 0.35352997 | 0.011251497 | 0.991022794 | 0.996726674 |
| MECR | 581.8457535 | 0.103752334 | 0.2751717 | 0.3770458 | 0.706139569 | 0.874446445 |
| MED1 | 2242.452756 | -0.415988388 | 0.380181385 | -1.094184 | 0.273874287 | 0.586847211 |
| MED10 | 676.6300683 | 0.632026519 | 0.471417074 | 1.340695011 | 0.180019495 | 0.484754911 |
| MED11 | 436.9107114 | 0.832811922 | 0.383280517 | 2.172852218 | 0.029791442 | 0.197226575 |
| MED12 | 1910.941601 | -1.156827233 | 0.530946839 | -2.178800488 | 0.029346493 | 0.195861365 |
| MED12L | 781.815372 | -1.516144408 | 0.394433343 | -3.843854568 | 0.000121117 | 0.006986551 |
| MED13 | 1438.033026 | -0.404950459 | 0.354214725 | -1.143234401 | 0.252941283 | 0.565733776 |
| MED13L | 3423.046472 | -0.316123497 | 0.441605957 | -0.715849711 | 0.474084154 | 0.74811012 |
| MED14 | 1812.064171 | -0.273896972 | 0.347743893 | -0.787639918 | 0.430907358 | 0.717595558 |
| MED15 | 1433.113346 | -0.882031352 | 0.35387376 | -2.492502843 | 0.012684632 | 0.127358401 |
| MED16 | 574.7683998 | -0.135986148 | 0.640067473 | -0.212455958 | 0.831751334 | 0.931048439 |
| MED17 | 925.1007744 | -0.05897904 | 0.29230498 | -0.201772272 | 0.84009476 | 0.934461703 |
| MED18 | 272.9756608 | 0.427904563 | 0.459685199 | 0.930864347 | 0.351923739 | 0.65929447 |
| MED19 | 385.183706 | 0.758664939 | 0.411030678 | 1.845762322 | 0.064926724 | 0.293855407 |
| MED20 | 855.9732969 | -0.177727765 | 0.284519119 | -0.624660185 | 0.53219411 | 0.782369637 |
| MED21 | 804.0801124 | 0.342219526 | 0.261669082 | 1.307833249 | 0.190929874 | 0.496058205 |
| MED22 | 547.5820561 | -0.162655126 | 0.298587627 | -0.544748378 | 0.585926587 | 0.815639728 |
| MED23 | 937.0619704 | -0.447674409 | 0.32601539 | -1.373169556 | 0.169699642 | 0.472403161 |
| MED24 | 2154.44683 | -0.570531208 | 0.391344391 | -1.457875011 | 0.144874998 | 0.43609099 |
| MED25 | 1314.884545 | -0.389595076 | 0.271702246 | -1.433904509 | 0.151599515 | 0.447172604 |
| MED26 | 279.5551081 | 0.078227526 | 0.358059744 | 0.218476183 | 0.827058114 | 0.928082319 |
| MED27 | 550.7596789 | 0.236651274 | 0.245985759 | 0.962052742 | 0.336023111 | 0.644231848 |
| MED28 | 4329.742023 | 0.092438014 | 0.274661513 | 0.33655248 | 0.736454283 | 0.890400696 |
| MED29 | 1159.776311 | -0.058938898 | 0.258020194 | -0.22842746 | 0.819313943 | 0.924316427 |

| | | | | | | |
|---|---|---|---|---|---|---|
| MED30 | 385.6619788 | 1.244282714 | 0.481819911 | 2.58246429 | 0.009809751 | 0.112404636 |
| MED31 | 179.7029182 | 0.880186226 | 0.309866914 | 2.840529872 | 0.004503865 | 0.070434393 |
| MED4 | 1558.31419 | 0.408560321 | 0.309328348 | 1.320798186 | 0.186568665 | 0.49118221 |
| MED6 | 1069.169651 | 0.285505308 | 0.311646434 | 0.916119286 | 0.359604333 | 0.667101656 |
| MED7 | 370.7145121 | 0.678858915 | 0.326302873 | 2.080456446 | 0.037483687 | 0.221788225 |
| MED8 | 448.5268669 | -0.237835364 | 0.33553314 | -0.708828236 | 0.478431074 | 0.750837683 |
| MED9 | 548.9990479 | 0.048834121 | 0.266314183 | 0.183370334 | 0.854507451 | 0.941507034 |
| MEDAG | 88.16719271 | -0.452149962 | 0.645681663 | -0.700267621 | 0.483760189 | 0.754278091 |
| MEF2A | 654.4980528 | -0.224927464 | 0.298089516 | -0.754563484 | 0.45051094 | 0.731870954 |
| MEF2B | 18.54527016 | 0.533014656 | 1.007468211 | 0.529063498 | 0.596761402 | 0.821812117 |
| MEF2C | 14.62335629 | 0.120706297 | 0.803777046 | 0.150173854 | 0.880627454 | 0.952579132 |
| MEF2D | 410.4307455 | -0.354378163 | 0.266880507 | -1.327853306 | 0.184226573 | 0.488704553 |
| MEGF10 | 44.2761225 | -0.138091277 | 0.687618045 | -0.200825557 | 0.840834978 | 0.934938981 |
| MEGF6 | 452.4457924 | 0.320617998 | 0.477412417 | 0.671574484 | 0.501854627 | 0.764473972 |
| MEGF8 | 1395.242411 | -0.241933517 | 0.305363655 | -0.79228 | 0.428197431 | 0.715316008 |
| MEGF9 | 181.2295999 | 0.294087219 | 0.352954171 | 0.833216442 | 0.404722671 | 0.70065889 |
| MEI4 | 13.34293904 | -0.169846895 | 1.303209635 | -0.13032968 | 0.896305599 | 0.959013488 |
| MEIOB | 50.47697418 | 1.692660115 | 0.931438109 | 1.81725452 | 0.06917815 | 0.303081608 |
| MEIOC | 121.5816934 | 1.000543898 | 0.665878289 | 1.502592763 | 0.132944091 | 0.419138571 |
| MEIS1 | 596.0541649 | -2.738766347 | 0.835119322 | -3.279491057 | 0.001039945 | 0.027754731 |
| MEIS2 | 4600.788504 | -0.281375353 | 0.464258436 | -0.606074829 | 0.544465065 | 0.790739196 |
| MEIS3 | 1054.695454 | 0.926123091 | 0.423040535 | 2.189206502 | 0.028581834 | 0.19323307 |
| MEIS3P1 | 48.33550136 | -1.034447989 | 0.884208055 | -1.169914686 | 0.242035303 | 0.553231259 |
| MEIS3P2 | 36.18800896 | -0.791372799 | 0.77222352 | -1.024797585 | 0.305458706 | 0.61550193 |
| MELK | 1340.011137 | 0.516069257 | 0.253233332 | 2.037919946 | 0.041557939 | 0.234125072 |
| MELTF | 55.05164992 | -0.218671739 | 0.487675944 | -0.448395583 | 0.653867729 | 0.848936869 |
| MEMO1 | 1250.264521 | 0.298475901 | 0.350918048 | 0.850557282 | 0.395015328 | 0.693813886 |
| MEN1 | 1410.087005 | -0.242057085 | 0.25811732 | -0.93777932 | 0.348357831 | 0.654805681 |
| MEOX1 | 126.3621298 | 1.223843542 | 0.534580927 | 2.289351304 | 0.022058949 | 0.168823142 |
| MEPCE | 945.4396631 | 0.099168084 | 0.302766186 | 0.327540155 | 0.743259377 | 0.892534058 |
| MERTK | 739.1841832 | -2.903574688 | 0.703567002 | -4.126934152 | 3.68E-05 | 0.002950973 |
| MESD | 1924.363695 | -0.110390084 | 0.366854835 | -0.300909442 | 0.76348355 | 0.902107118 |
| MEST | 8840.718694 | 0.164587313 | 0.302937277 | 0.543304922 | 0.586919876 | 0.816520994 |
| MET | 713.5366119 | -2.430768213 | 0.639798028 | -3.799274312 | 0.00014512 | 0.007733542 |
| METAP1 | 2572.752429 | 0.020012876 | 0.242941776 | 0.082377253 | 0.934346724 | 0.97507153 |
| METAP1D | 206.1007109 | 0.01085254 | 0.314430261 | 0.034514934 | 0.972466534 | 0.989954357 |
| METAP2 | 7176.911519 | 0.594588708 | 0.358412637 | 1.658950179 | 0.097125832 | 0.357954238 |
| METRN | 394.4313411 | 1.430628741 | 0.625160214 | 2.288419368 | 0.022113111 | 0.169006551 |
| METRNL | 415.7451226 | 0.574421204 | 0.709621798 | 0.809475139 | 0.418241897 | 0.709589334 |
| METTL1 | 237.4646478 | 0.676402804 | 0.437385077 | 1.54646978 | 0.121991153 | 0.402659864 |
| METTL11B | 32.80895201 | 0.489102006 | 0.613646263 | 0.797042263 | 0.425426487 | 0.713673173 |
| METTL14 | 1238.392317 | -0.124274375 | 0.232005293 | -0.535653186 | | |

0.592198263 0.819765854
METTL15 628.4499763 0.355915445 0.258760064 1.375465128 0.16898729
0.471508612
METTL16 1506.174511 -0.105339479 0.247767092 -0.425155245
0.670723507 0.858188946
METTL17 1413.905182 0.157374087 0.227591507 0.691476098 0.489266396
0.757568942
METTL18 441.0604617 0.428309941 0.424228831 1.009620067 0.312677352
0.622907489
METTL21A 510.6521863 0.987042748 0.300219295 3.287739208 0.001009953
0.027467362
METTL22 809.1000349 0.171255926 0.27773295 0.616620843 0.537484843
0.785793828
METTL23 513.6692242 0.767698784 0.334203477 2.297099929 0.02161307
0.166611729
METTL24 56.75712331 -2.171088145 0.919479988 -2.361213048
0.018215263 0.1525918
METTL25 176.7213332 0.140230661 0.375536777 0.373413922 0.708840399
0.875646746
METTL26 593.4605826 0.628275371 0.418482499 1.501318149 0.133273293
0.419566119
METTL27 90.23799394 0.962824873 0.485348872 1.983778946 0.047280488
0.250276443
METTL2A 844.8094533 0.449897883 0.33708358 1.33467754 0.181981907
0.486131134
METTL2B 857.6440175 0.203103602 0.263724164 0.77013649 0.441218932
0.724743658
METTL3 1918.534516 0.24633501 0.248859088 0.989857399 0.322243825
0.632029271
METTL4 362.4033909 0.330297693 0.351317975 0.940167362 0.34713172
0.653844005
METTL5 1655.751746 0.938671887 0.395991563 2.370434056 0.017767213
0.150416512
METTL6 413.941423 0.135288322 0.258888039 0.522574634 0.601270297
0.824832027
METTL7A 239.8705364 -1.591239557 0.587312939 -2.709355525
0.006741406 0.089871589
METTL7B 43.48803387 1.023510079 0.610129183 1.677530116 0.093438865
0.350724392
METTL8 635.0526822 0.136527578 0.270712457 0.504326915 0.614031667
0.830316752
METTL9 2284.658296 0.518049576 0.268806803 1.927218996 0.053952344
0.268452281
MEX3A 3636.236339 0.240361417 0.287419583 0.836273627 0.403000986 0.699766232
MEX3B 425.1090679 1.719809335 0.47280934 3.637426736 0.000275375 0.011709082
MEX3C 2737.350884 -0.103138229 0.232120123 -0.444331272 0.656803095
0.85024732
MEX3D 819.7365378 0.460585339 0.384313443 1.198462733 0.230736923 0.539536045
MFAP1 1728.98782 -0.048794319 0.254222865 -0.19193521 0.847792957
0.938207222
MFAP2 3194.707064 0.65700194 0.405680663 1.619505191 0.105338611 0.372168621
MFAP3 460.8582538 -0.741064342 0.656729407 -1.128416567 0.259144035
0.572516987
MFAP3L 75.33010244 -0.319119777 0.480670065 -0.663906077
0.506750427 0.767639613
MFAP4 13.03875526 0.842206875 0.935527336 0.90024828 0.367988138 0.674179083
MFAP5 22.22723028 -1.63471448 0.901543874 -1.813238964 0.069794973
0.303603614
MFF 2027.939601 0.377623595 0.324917044 1.162215408 0.245147977 0.556630702
MFGE8 2955.446269 -0.038390994 0.273567874 -0.140334438 0.888395756
0.955943827
MFHAS1 2746.671418 -0.122967357 0.253219411 -0.485615838
0.627239565 0.836041947
MFN1 1686.010375 -0.022103609 0.228475524 -0.09674388 0.922929792

0.971075713
MFN2 2378.15722 -0.212016083 0.405908481 -0.522324841 0.601444176
0.824869221
MFNG 73.34196602 2.203726835 0.766517311 2.874986388 0.004040449 0.065655774
MFSD1 401.0748106 -0.182624714 0.325950371 -0.560283805 0.575285872
0.80849115
MFSD10 967.6687447 -0.054286107 0.510331751 -0.106374151
0.915285502 0.969069464
MFSD11 641.4854122 -0.358282995 0.3372665 -1.062314208
0.288093066 0.601673803
MFSD12 409.8715001 0.422336955 0.354754572 1.190504613 0.233848118
0.54387702
MFSD13A 81.73774497 -0.232643228 0.436944113 -0.532432458
0.59442651 0.820642877
MFSD14A 1415.865288 0.265702294 0.251825283 1.055105706 0.291376982
0.604386136
MFSD14B 1487.257653 0.109913373 0.260713148 0.42158738 0.67332622
0.859417136
MFSD14C 889.7606382 -0.130181451 0.312819316 -0.416155411
0.677296285 0.860358531
MFSD2A 53.5784579 0.673438519 0.762128357 0.883628739 0.37689666
0.68029735
MFSD3 502.7040801 1.04885949 0.539924869 1.942602666 0.052064185 0.262949685
MFSD4B 523.95021 -0.708178141 0.417135038 -1.697719148
0.089560782 0.342777615
MFSD5 451.525232 0.407634971 0.39634836 1.028476494 0.303725743 0.614002998
MFSD6 73.24255803 -2.057352595 0.69903914 -2.943115023 0.003249277
0.057339258
MFSD6L 18.08951305 -2.16142096 1.088601207 -1.985502998 0.047088537
0.25004557
MFSD8 533.3378094 -0.109798118 0.254411921 -0.431576154 0.666049493
0.855527631
MFSD9 308.6124764 -0.042986374 0.305279887 -0.140809716 0.888020267
0.955943827
MGA 1515.227526 -0.40364432 0.361360882 -1.117011665 0.263989338
0.577448862
MGARP 74.07106285 1.659441082 0.671962344 2.469544754 0.013528509 0.132400677
MGAT1 1995.12766 0.454490301 0.277130879 1.639984338 0.101008423 0.36405744
MGAT2 560.741178 0.867973575 0.26761333 3.243386924 0.001181177 0.030218607
MGAT3 677.9228386 -0.354368263 0.269605564 -1.314395211 0.188713241
0.493962743
MGAT4A 173.590576 -0.036596194 0.418827483 -0.087377729
0.930371271 0.97369593
MGAT4B 808.6440075 0.136163196 0.306426373 0.444358606 0.656783336
0.85024732
MGAT5 2369.72861 -0.030147881 0.483237176 -0.062387338 0.950254378
0.981636196
MGAT5B 109.9751218 0.005142206 0.50970114 0.010088669 0.991950543
0.99688215
MGLL 33.6943933 -0.950158675 0.570892472 -1.66433912 0.096044679
0.355648968
MGME1 1096.78722 -0.025790486 0.288924528 -0.089263748 0.928872304
0.972795922
MGMT 559.6852945 0.640290778 0.454685796 1.408204928 0.159070398 0.459030918
MGRN1 851.7102091 -0.682757483 0.449999715 -1.517239813 0.129206148
0.413472911
MGST1 4091.775014 0.956228872 0.443760529 2.154830836 0.031175074 0.201233103
MGST2 153.5432153 0.928583008 0.377436842 2.460234149 0.01388464 0.134161613
MGST3 2572.953673 0.526651718 0.528882284 0.995782492 0.319355839 0.629115732
MIA 15.06171221 0.084118984 1.104467757 0.076162463 0.939289846 0.977796709
MIA2 381.2073277 -0.012918242 0.294158828 -0.043915875 0.964971461
0.988700292
MIA3 1911.403779 -0.267051596 0.334931348 -0.797332342 0.425258042
0.713592495

| | | | | | | |
|---|---|---|---|---|---|---|
| MIB1 | 3319.158664 | 0.08097051 | 0.351898087 | 0.230096478 | 0.818016801 | 0.924129339 |
| MIB2 | 353.1568492 | -0.052941497 | 0.616645865 | -0.085853973 | 0.931582501 | 0.974341807 |
| MICAL1 | 2217.170205 | -0.207409806 | 0.320750714 | -0.64663864 | 0.517865844 | 0.773950969 |
| MICAL2 | 159.1345411 | 0.925455928 | 1.835165525 | 0.504290166 | 0.614057487 | 0.830316752 |
| MICAL3 | 597.3080064 | -0.033266274 | 0.551976522 | -0.060267552 | 0.951942545 | 0.982510399 |
| MICALL1 | 2007.47388 | -0.781279024 | 0.417617363 | -1.870801105 | 0.061372654 | 0.285640507 |
| MICALL2 | 247.79002 | 0.359669029 | 0.516222217 | 0.696732952 | 0.485969933 | 0.75593662 |
| MICB | 13.29604188 | 0.070780231 | 1.51944969 | 0.046582806 | 0.962845736 | 0.987649267 |
| MICOS10 | 783.5276343 | 1.120485945 | 0.451756847 | 2.480285474 | 0.013127723 | 0.129746377 |
| MICOS13 | 319.7706189 | 0.979571054 | 0.458269073 | 2.137545631 | 0.032553638 | 0.205776324 |
| MICU1 | 1256.930877 | -0.211188931 | 0.30516666 | -0.692044575 | 0.488909337 | 0.757325255 |
| MICU2 | 1927.501761 | 0.552033632 | 0.329281538 | 1.676479146 | 0.093644378 | 0.351044935 |
| MICU3 | 223.5165114 | 0.081038475 | 0.400790759 | 0.202196465 | 0.839763137 | 0.934363517 |
| MID1 | 907.5001564 | -0.22049479 | 0.512117046 | -0.43055546 | 0.666791632 | 0.856258128 |
| MID1IP1 | 1198.307836 | 0.791778824 | 0.274549238 | 2.883922861 | 0.003927549 | 0.064878831 |
| MID2 | 38.3021001 | -1.401570597 | 0.563596728 | -2.486832387 | 0.012888611 | 0.128239534 |
| MIDN | 2060.105702 | 0.221952696 | 0.292094426 | 0.759866251 | 0.447334537 | 0.728949036 |
| MIEF1 | 1741.341968 | 0.028700286 | 0.246874574 | 0.116254523 | 0.907450827 | 0.965304913 |
| MIEF2 | 136.1595347 | 0.245382465 | 0.432884584 | 0.566854247 | 0.570813211 | 0.806748565 |
| MIEN1 | 801.1552583 | 1.058936218 | 0.476665305 | 2.221550859 | 0.026313674 | 0.185350959 |
| MIER1 | 1469.073405 | -0.420642735 | 0.297905423 | -1.412000932 | 0.157949687 | 0.457277356 |
| MIER2 | 479.6506411 | -0.053933297 | 0.355861875 | -0.151556827 | 0.879536487 | 0.952083342 |
| MIER3 | 1123.399491 | 0.069706707 | 0.334039103 | 0.208678283 | 0.834699394 | 0.932107564 |
| MIF | 1639.422845 | 1.024439809 | 0.679200308 | 1.508302922 | 0.131477016 | 0.41719445 |
| MIF4GD | 500.5759575 | 0.576850549 | 0.311640659 | 1.851011837 | 0.06416785 | 0.292572699 |
| MIGA1 | 375.4621651 | -0.352591647 | 0.29133513 | -1.210261347 | 0.226178625 | 0.534746233 |
| MIGA2 | 676.5907137 | -0.053223809 | 0.444228924 | -0.11981167 | 0.904632337 | 0.963993604 |
| MIIP | 830.7934399 | 0.409649034 | 0.477188186 | 0.858464326 | 0.390636121 | 0.690469214 |
| MINAR1 | 47.29954575 | -0.808296192 | 0.561273282 | -1.44011165 | 0.149835814 | 0.444497287 |
| MINDY1 | 394.7135186 | -0.947549562 | 0.380791014 | -2.488371646 | 0.012832955 | 0.128084711 |
| MINDY2 | 622.4174141 | -0.483021097 | 0.355672458 | -1.358050323 | 0.174447717 | 0.478578744 |
| MINDY3 | 529.8299678 | -0.010166853 | 0.265064402 | -0.038356162 | 0.969403713 | 0.989496332 |
| MINDY4 | 158.9141924 | -0.143802072 | 0.565541346 | -0.254273314 | 0.799284415 | 0.916202785 |
| MINK1 | 2540.867882 | -0.83760234 | 0.325709393 | -2.571624763 | 0.010122254 | 0.114259456 |
| MINPP1 | 851.5158049 | 0.70719224 | 0.272088624 | 2.599124617 | 0.009346184 | 0.109346547 |
| MIOS | 1264.899684 | -0.155724407 | 0.259940819 | -0.599076389 | 0.549121946 | 0.79348329 |
| MIPEP | 803.077597 | -0.000275532 | 0.251787616 | -0.001094303 | 0.999126872 | 0.999324484 |
| MIPEPP3 | 13.75330351 | 0.498997614 | 0.890102659 | 0.560606812 | 0.575065607 | 0.808420489 |

| | | | | | | |
|---|---|---|---|---|---|---|
| MIPOL1 | 565.0691938 | -0.029836358 | 0.272551027 | -0.109470724 | 0.912829141 | 0.967549144 |
| MIS12 | 885.327564 | 0.164566697 | 0.253120648 | 0.650151217 | 0.515594549 | 0.77377435 |
| MIS18A | 1409.36852 | 1.042956952 | 0.465876033 | 2.238700593 | 0.025175404 | 0.18153038 |
| MIS18BP1 | 1967.208961 | 0.420047463 | 0.408831381 | 1.027434494 | 0.304215916 | 0.614548557 |
| MISP | 26.80826392 | -1.126243224 | 0.836926324 | -1.345689807 | 0.178402575 | 0.48262223 |
| MISP3 | 16.80795574 | 2.048306894 | 0.805299338 | 2.543534804 | 0.010973712 | 0.119221957 |
| MITD1 | 1102.247125 | 0.505922765 | 0.36494367 | 1.386303715 | 0.165654183 | 0.467560858 |
| MITF | 20.51490383 | -2.892505122 | 1.279642252 | -2.260401387 | 0.023796351 | 0.176535175 |
| MIXL1 | 29.24812518 | -2.360150359 | 1.198628514 | -1.969042394 | 0.048948223 | 0.254849017 |
| MKI67 | 4533.629699 | 0.245603123 | 0.587823312 | 0.417817936 | 0.676080236 | 0.860110127 |
| MKKS | 887.2140585 | 0.385119034 | 0.278358196 | 1.383537615 | 0.166500082 | 0.46837989 |
| MKLN1 | 1375.534616 | -0.470639874 | 0.280020505 | -1.680733611 | 0.092814669 | 0.349440669 |
| MKNK1 | 685.5474219 | -0.007535453 | 0.254270179 | -0.029635614 | 0.976357662 | 0.99184251 |
| MKNK2 | 2236.758712 | 0.007046359 | 0.383924399 | 0.018353506 | 0.985356843 | 0.99460071 |
| MKRN1 | 2515.460162 | -0.03305918 | 0.343602079 | -0.096213563 | 0.923350959 | 0.971170092 |
| MKRN2 | 614.3785435 | -0.10974506 | 0.261175721 | -0.420196256 | 0.67434209 | 0.859672768 |
| MKRN2OS | 17.20114968 | -1.501410584 | 1.385247171 | -1.083857535 | 0.27842797 | 0.592005173 |
| MKRN7P | 15.27921336 | -0.643334672 | 0.849143207 | -0.757628003 | 0.448673711 | 0.729876568 |
| MKS1 | 614.1189213 | -0.247587828 | 0.257440384 | -0.961728786 | 0.336185859 | 0.64425371 |
| MLEC | 4984.200363 | -0.2957418 | 0.360175252 | -0.821105275 | 0.411586305 | 0.704686727 |
| MLF1 | 717.3268726 | 0.731114201 | 0.326607091 | 2.238512941 | 0.025187624 | 0.18153038 |
| MLF2 | 3807.829928 | 0.48125322 | 0.304777282 | 1.579032456 | 0.114328613 | 0.389245822 |
| MLH1 | 1948.054257 | 0.279145692 | 0.220783822 | 1.264339432 | 0.206108222 | 0.512517266 |
| MLH3 | 499.3698937 | 0.049939383 | 0.426350044 | 0.117132352 | 0.906755173 | 0.965234099 |
| MLKL | 25.84890554 | -0.070355097 | 0.656897376 | -0.107102113 | 0.914707971 | 0.9686608 |
| MLLT1 | 1181.960108 | -0.216102501 | 0.411590267 | -0.525042788 | 0.599553447 | 0.823822601 |
| MLLT10 | 1652.668243 | -0.35555889 | 0.320472668 | -1.109482728 | 0.267221992 | 0.580473201 |
| MLLT11 | 2560.484284 | 0.438943477 | 0.292839375 | 1.498922328 | 0.133893783 | 0.420559542 |
| MLLT3 | 1099.830508 | 0.040447158 | 0.337636306 | 0.119795049 | 0.904645504 | 0.963993604 |
| MLLT6 | 551.97505 | -0.339312923 | 0.425440192 | -0.797557282 | 0.425127449 | 0.71357297 |
| MLPH | 283.0891314 | -0.412534587 | 0.71802663 | -0.574539397 | 0.565602848 | 0.80396897 |
| MLST8 | 1319.539717 | 0.345181399 | 0.349489452 | 0.987673296 | 0.323312677 | 0.63322621 |
| MLX | 901.117145 | 0.140275414 | 0.250046629 | 0.560997019 | 0.574799569 | 0.808359708 |
| MLXIP | 383.6211104 | 0.141323418 | 0.308465757 | 0.458149453 | 0.646845072 | 0.846556814 |
| MLXIPL | 62.23462805 | 1.036225584 | 0.58408929 | 1.774087629 | 0.076048657 | 0.316438335 |
| MLYCD | 324.8457893 | -0.572374064 | 0.290700852 | -1.96894526 | 0.048959377 | 0.254849017 |
| MMAA | 98.51299329 | 0.436293146 | 0.429337802 | 1.016200168 | 0.309534076 | 0.61915097 |
| MMAB | 653.488276 | 0.514290059 | 0.268227755 | 1.917363323 | 0.05519179 | 0.271855406 |
| MMACHC | 251.1361797 | -0.095011369 | 0.291462647 | -0.325981287 | 0.744438514 | 0.892725315 |
| MMADHC | 4475.098515 | 0.560955225 | 0.29550479 | 1.898294867 | 0.05765725 | 0.278177478 |

| | | | | | | |
|---|---|---|---|---|---|---|
| MMD | 1591.195889 | 0.808514941 | 0.315943692 | 2.559047583 | 0.010495937 | 0.116399025 |
| MME | 103.4477706 | -2.525709746 | 1.011365281 | -2.497326923 | 0.012513353 | 0.12669427 |
| MMGT1 | 861.3233359 | 0.101554149 | 0.297537549 | 0.341315405 | 0.732866152 | 0.888256962 |
| MMP11 | 220.2375937 | 0.828321901 | 0.456600539 | 1.814106271 | 0.069661366 | 0.303425952 |
| MMP14 | 2448.642461 | -0.571993885 | 0.24492966 | -2.335339404 | 0.019525697 | 0.15832408 |
| MMP15 | 1977.387406 | -0.30068427 | 0.466421261 | -0.644662444 | 0.519145959 | 0.774631448 |
| MMP16 | 479.7394683 | -0.137506072 | 0.497746042 | -0.27625749 | 0.782350304 | 0.909909835 |
| MMP17 | 64.52097869 | -0.586640111 | 0.63647469 | -0.921702183 | 0.356683942 | 0.664215534 |
| MMP19 | 35.22828812 | -0.876695939 | 0.723361778 | -1.211974375 | 0.22552219 | 0.53459368 |
| MMP2 | 3529.632391 | 0.26104006 | 0.435969563 | 0.598757533 | 0.549334585 | 0.79348329 |
| MMP23B | 28.99236789 | 0.655467718 | 0.72348504 | 0.905986554 | 0.36494297 | 0.671631599 |
| MMP24 | 63.82731169 | -2.344327871 | 0.627390991 | -3.736629793 | 0.000186503 | 0.009010956 |
| MMP24OS | 1078.755296 | 0.205601044 | 0.463823055 | 0.443274741 | 0.657567024 | 0.850609594 |
| MMP25 | 19.53936667 | 0.365615729 | 0.893022183 | 0.409413938 | 0.682235913 | 0.862735748 |
| MMP9 | 17.9704218 | -0.53659728 | 0.929133095 | -0.577524666 | 0.563585081 | 0.801976485 |
| MMRN1 | 31.00134797 | 2.74105683 | 1.428894782 | 1.918305578 | 0.055072276 | 0.271793713 |
| MMRN2 | 30.56153837 | -0.418364032 | 0.826833601 | -0.505983346 | 0.612868341 | 0.829496441 |
| MMS19 | 3541.952801 | -0.213493842 | 0.295901065 | -0.721504135 | 0.470599397 | 0.74563587 |
| MMS22L | 938.5260063 | -0.133274868 | 0.275991832 | -0.482894247 | 0.629170827 | 0.837367367 |
| MMUT | 895.2897956 | -0.12761335 | 0.249402387 | -0.511676541 | 0.608877407 | 0.828009963 |
| MN1 | 1263.99683 | 1.324441525 | 0.488039203 | 2.713801506 | 0.006651601 | 0.089223197 |
| MNAT1 | 812.9946836 | 0.734553707 | 0.31599071 | 2.324605385 | 0.020093071 | 0.160340171 |
| MND1 | 592.3503778 | 1.468012988 | 0.321944215 | 4.559836522 | 5.12E-06 | 0.000663809 |
| MNS1 | 76.69537589 | -1.327180097 | 0.614952222 | -2.15818408 | 0.030913523 | 0.200679954 |
| MNT | 585.5022526 | -0.035056611 | 0.335380216 | -0.104527962 | 0.916750379 | 0.969211928 |
| MOAP1 | 487.6880049 | -0.041076322 | 0.250303117 | -0.164106316 | 0.86964745 | 0.947530987 |
| MOB1A | 2843.359438 | 0.091464109 | 0.247418364 | 0.369673889 | 0.711625489 | 0.877287654 |
| MOB1B | 1179.637966 | 0.055167224 | 0.333208943 | 0.165563455 | 0.868500509 | 0.947046732 |
| MOB2 | 315.2371626 | -0.158081801 | 0.407192611 | -0.388223649 | 0.697850533 | 0.870467 |
| MOB3A | 398.4840714 | -0.432355542 | 0.280889573 | -1.53923671 | 0.123746519 | 0.405045568 |
| MOB3B | 1039.83469 | -0.435432293 | 0.290754523 | -1.497594217 | 0.13423871 | 0.420619758 |
| MOB3C | 77.51177223 | -0.474197066 | 0.476047558 | -0.996112801 | 0.319195341 | 0.629115732 |
| MOB4 | 2063.375653 | 0.610685985 | 0.305167288 | 2.001151528 | 0.045376063 | 0.245418984 |
| MOCOS | 34.29191715 | 0.541496771 | 0.609407843 | 0.888562195 | 0.374238416 | 0.67864822 |
| MOCS1 | 534.3441793 | 0.265373151 | 0.31506202 | 0.842288611 | 0.399626424 | 0.697589493 |
| MOCS2 | 661.4743496 | 0.192929751 | 0.317718026 | 0.607235772 | 0.543694456 | 0.790216482 |
| MOCS3 | 228.8245268 | 0.01884898 | 0.357432984 | 0.052734305 | 0.957943606 | 0.98520931 |
| MOGS | 1471.708958 | -0.206616307 | 0.332019541 | -0.622301646 | 0.533743539 | 0.783156734 |
| MOK | 119.6359065 | -0.630642732 | 0.401236845 | -1.571746811 | 0.116009281 | 0.392480553 |
| MON1A | 122.2846158 | 0.650449885 | 0.526048035 | 1.236483823 | 0.216278775 | 0.523980406 |
| MON1B | 1147.914264 | -0.383376988 | 0.242396201 | -1.581613019 | 0.11373794 | |

0.388022186
MON2 887.1013081 -0.213785932 0.33210975 -0.643720734 0.519756542
0.775026528
MORC2 2149.153454 -0.236060567 0.407187147 -0.579734821 0.562093458
0.801706045
MORC3 926.1647227 0.214851759 0.375200328 0.572632119 0.566893808 0.804595923
MORC4 284.1301545 -0.698137801 0.294029417 -2.374380798 0.017578411
0.14971347
MORF4L1 12403.73867 0.390661389 0.254698203 1.533820753 0.125073756
0.406867909
MORF4L2 7931.719582 0.364447255 0.234021976 1.55732065 0.119394375
0.398357612
MORN1 148.0554319 0.198202099 0.426454597 0.464767176 0.642098196 0.844148268
MORN2 181.3004903 0.821374453 0.428205737 1.918177135 0.055088554 0.271793713
MORN3 176.9789879 -1.310411594 0.668703155 -1.959631241 0.050038907
0.257487003
MORN4 449.6734054 0.770328305 0.347733694 2.21528232 0.026740693 0.186818803
MOSMO 150.982266 0.29255215 0.345779922 0.846064598 0.397516709 0.696188739
MOSPD1 473.9978484 -0.256096736 0.304957838 -0.839777515
0.401033143 0.698568129
MOSPD2 410.0207824 0.046521616 0.334353274 0.139139106 0.889340226
0.956341292
MOSPD3 238.8685436 0.283634493 0.405542959 0.699394445 0.484305559
0.754852802
MOV10 1406.016533 -0.68982704 0.285843902 -2.413299827 0.015808809
0.142420095
MOXD1 51.57164555 0.401151645 0.546928248 0.733463021 0.463276074 0.740745005
MPC1 135.5555027 1.017388742 0.364773312 2.789098624 0.005285497 0.077250742
MPC2 699.395612 0.630086508 0.453107333 1.39058996 0.164349803 0.466395596
MPDU1 1095.019371 0.715209108 0.309925463 2.307681015 0.021016885 0.164636113
MPDZ 1998.912567 -0.129204583 0.458996164 -0.281493818 0.778331669
0.908108663
MPG 603.1809434 0.404087065 0.58721887 0.688137057 0.491366477 0.758779636
MPHOSPH10 1092.994573 0.067255023 0.247293722 0.271964136 0.785649596
0.910966831
MPHOSPH6 759.9018149 0.855398895 0.313132193 2.731750087 0.006299891
0.086267508
MPHOSPH8 1044.944786 -0.022123957 0.439123858 -0.050382043
0.959817946 0.986076932
MPHOSPH9 1168.264434 0.535677392 0.43773659 1.223743696 0.22104893
0.529784095
MPI 1191.87641 0.328448029 0.260501878 1.260827872 0.20737087 0.514272875
MPLKIP 743.8706266 1.206618318 0.43711898 2.760388756 0.005773262
0.081921507
MPND 935.1821065 0.211462214 0.420295689 0.50312725 0.614874809 0.830872437
MPP1 312.8597278 -3.536498301 0.837524768 -4.222559663 2.42E-05
0.002118182
MPP2 847.6395885 0.44228331 0.433137081 1.021116245 0.307199368 0.617205881
MPP3 80.52878142 0.364668754 0.476519898 0.765274977 0.444107821 0.727127105
MPP5 1274.748547 0.018272848 0.294958276 0.06195062 0.950602156 0.981826318
MPP6 990.6888932 0.243135047 0.303322219 0.801573482 0.422799722 0.712269066
MPP7 453.2622009 -0.782140162 0.300497752 -2.602815352 0.009246174
0.108823663
MPPE1 386.9682035 -0.391128991 0.369488655 -1.058568337 0.289796413
0.60258519
MPPED2 33.12667976 1.269174927 0.642200392 1.976291112 0.048121816
0.253052362
MPRIP 4316.480369 -0.496688157 0.299743815 -1.657042219 0.09751094
0.358846393
MPST 1501.432964 0.72150166 0.428588565 1.683436563 0.09229061 0.348814361
MPV17 1113.724899 0.400865103 0.292839011 1.368892423 0.171032905 0.47427165
MPV17L 43.77275988 -0.63398663 0.597667327 -1.060768426 0.288795155
0.601911102
MPV17L2 385.5276368 0.486891896 0.312006323 1.560519321 0.118637208

| | | | | | | |
|---|---|---|---|---|---|---|
| | | | | | | 0.396986757 |
| MPZ | 53.54085479 | 0.343678616 | 0.570576357 | 0.602335887 | 0.546950577 | 0.792249935 |
| MPZL1 | 2679.031636 | -0.897723153 | 0.464603559 | -1.932234776 | 0.053330535 | 0.267056023 |
| MPZL2 | 137.3241149 | 0.075407397 | 0.632543417 | 0.119212997 | 0.90510661 | 0.964349489 |
| MPZL3 | 101.4065541 | -0.603545187 | 0.591612294 | -1.020170122 | 0.307647785 | 0.617370971 |
| MR1 | 35.00688334 | -0.841775967 | 0.598686235 | -1.406038619 | 0.159712657 | 0.459772433 |
| MRAP2 | 22.94379296 | 0.518201466 | 0.831944602 | 0.622879774 | 0.53336353 | 0.783143103 |
| MRAS | 277.1495562 | -0.259828263 | 0.374269917 | -0.694226952 | 0.487539893 | 0.756439734 |
| MRC1 | 11.30434865 | -0.870534481 | 1.188397481 | -0.732528043 | 0.463846333 | 0.741050201 |
| MRE11 | 1563.086915 | -0.449732708 | 0.294222148 | -1.528548113 | 0.126376507 | 0.409145965 |
| MREG | 361.7576322 | -0.26970838 | 0.335951732 | -0.802818842 | 0.42207945 | 0.712080472 |
| MRFAP1 | 7976.270958 | 0.568482812 | 0.41065646 | 1.384326968 | 0.16625836 | 0.468047055 |
| MRFAP1L1 | 2189.638229 | 0.117388587 | 0.273538071 | 0.42914899 | 0.667814802 | 0.856696248 |
| MRGBP | 516.5446974 | 0.709839224 | 0.327439309 | 2.167849748 | 0.03017012 | 0.198185614 |
| MRGPRF | 86.72673396 | 1.208092228 | 0.40810437 | 2.960253105 | 0.003073864 | 0.055057366 |
| MRI1 | 398.4828353 | 0.415137528 | 0.308098169 | 1.347419654 | 0.177845114 | 0.482075465 |
| MRM1 | 51.95518635 | 0.457134519 | 0.575530486 | 0.794283762 | 0.427030255 | 0.714556109 |
| MRM2 | 1012.509417 | 0.056193048 | 0.299454008 | 0.18765168 | 0.851149711 | 0.939477675 |
| MRM3 | 442.693026 | 0.348380843 | 0.319250599 | 1.091245698 | 0.275164785 | 0.588252711 |
| MRNIP | 425.0651 | -0.338887741 | 0.34843652 | -0.972595353 | 0.330754453 | 0.639057031 |
| MRO | 19.27500968 | 0.373497671 | 0.754583324 | 0.494972072 | 0.620619835 | 0.833297063 |
| MROH1 | 449.6004085 | -1.912809563 | 0.393532797 | -4.860610293 | 1.17E-06 | 0.000206439 |
| MROH6 | 19.3179229 | 0.257769044 | 0.731102244 | 0.352575917 | 0.724406395 | 0.883670922 |
| MROH7 | 17.01682331 | -0.793708031 | 1.058657569 | -0.749730654 | 0.453416942 | 0.733497409 |
| MROH8 | 36.88711108 | 0.107872122 | 0.522882235 | 0.206302901 | 0.836554312 | 0.932914251 |
| MRPL1 | 1000.982334 | 0.60870001 | 0.325845278 | 1.868064544 | 0.061753075 | 0.286325152 |
| MRPL10 | 2522.286504 | -0.035292668 | 0.305444563 | -0.115545249 | 0.908012957 | 0.965583536 |
| MRPL11 | 1932.859669 | 0.742530759 | 0.384198637 | 1.932674112 | 0.053276356 | 0.267056023 |
| MRPL12 | 1272.264857 | 0.941396054 | 0.395166984 | 2.382274057 | 0.017206087 | 0.147810615 |
| MRPL13 | 1301.868923 | 0.94344297 | 0.331178604 | 2.848743721 | 0.004389222 | 0.069186323 |
| MRPL14 | 1264.153698 | 0.985217609 | 0.437800982 | 2.250377798 | 0.024424973 | 0.178492903 |
| MRPL15 | 1542.288525 | 0.639164973 | 0.297810716 | 2.146212137 | 0.031856058 | 0.203318574 |
| MRPL16 | 1123.84082 | 0.648511559 | 0.467022976 | 1.388607396 | 0.164952167 | 0.467115865 |
| MRPL17 | 799.4233891 | 0.642591854 | 0.360957369 | 1.780243069 | 0.07503619 | 0.314480446 |
| MRPL18 | 973.1955893 | 0.477159442 | 0.311109654 | 1.533733961 | 0.125095115 | 0.406867909 |
| MRPL19 | 1654.53248 | 0.289732489 | 0.267000011 | 1.085140362 | 0.277859493 | 0.591968315 |
| MRPL2 | 624.5662402 | 0.543931653 | 0.314183686 | 1.731253649 | 0.083406534 | 0.330659907 |
| MRPL20 | 1530.556522 | 0.772371185 | 0.421681505 | 1.831645866 | 0.067004199 | 0.297817842 |
| MRPL21 | 741.001266 | 0.692151982 | 0.369915595 | 1.871107871 | 0.061330131 | 0.285640507 |

| | | | | | | |
|---|---|---|---|---|---|---|
| MRPL22 | 1029.166775 | 0.755598754 | 0.381050577 | 1.9829356 | 0.047374624 | 0.250687274 |
| MRPL23 | 783.3448169 | 0.622663188 | 0.360677963 | 1.726368819 | 0.084281082 | 0.332430848 |
| MRPL24 | 537.5543799 | 0.65429891 | 0.331727695 | 1.972397604 | 0.048564235 | 0.25374113 |
| MRPL27 | 1067.459441 | 0.825933209 | 0.374887955 | 2.20314683 | 0.027584401 | 0.190183846 |
| MRPL28 | 1464.996689 | 0.33155484 | 0.36326133 | 0.912717135 | 0.36139133 | 0.668292036 |
| MRPL3 | 3430.951777 | 0.452550069 | 0.221528194 | 2.042855411 | 0.041066753 | 0.232934829 |
| MRPL30 | 2413.191692 | 0.548406662 | 0.257706544 | 2.128027689 | 0.033334788 | 0.208274771 |
| MRPL32 | 1752.578457 | 0.642479719 | 0.300775038 | 2.136080582 | 0.032672846 | 0.206178012 |
| MRPL33 | 997.2294305 | 1.313981512 | 0.429641779 | 3.058318762 | 0.002225827 | 0.044607685 |
| MRPL34 | 551.3429668 | 1.223250599 | 0.438232876 | 2.791325497 | 0.005249265 | 0.077110704 |
| MRPL35 | 1010.289572 | 0.239429963 | 0.242654267 | 0.986712353 | 0.323783672 | 0.63357695 |
| MRPL36 | 560.6256059 | 1.054188505 | 0.379881155 | 2.775048174 | 0.005519353 | 0.079368821 |
| MRPL37 | 2369.048115 | 0.288009891 | 0.28432372 | 1.012964695 | 0.311077025 | 0.6211305 |
| MRPL38 | 810.4974818 | -0.028959 | 0.305568526 | -0.094770886 | 0.924496812 | 0.971780852 |
| MRPL39 | 943.5038836 | 0.546301854 | 0.247386405 | 2.208293757 | 0.027223802 | 0.188612866 |
| MRPL4 | 825.938298 | 0.558646193 | 0.422243783 | 1.323041843 | 0.185821463 | 0.490363093 |
| MRPL40 | 475.9014746 | 0.060192573 | 0.35964882 | 0.167364857 | 0.867082975 | 0.946278179 |
| MRPL41 | 335.4914058 | 0.944065621 | 0.537570377 | 1.756171212 | 0.079059179 | 0.322602824 |
| MRPL42 | 3020.783253 | 0.967010268 | 0.284209044 | 3.402461282 | 0.000667818 | 0.021195543 |
| MRPL43 | 1000.765037 | 0.607185312 | 0.405764816 | 1.496397144 | 0.134550193 | 0.420896533 |
| MRPL44 | 1234.980158 | 0.500031092 | 0.355104636 | 1.408123245 | 0.159094579 | 0.459030918 |
| MRPL45 | 671.4431975 | 0.291420137 | 0.433056799 | 0.672937446 | 0.500987088 | 0.763796851 |
| MRPL45P2 | 85.69462773 | -0.167512715 | 0.419754601 | -0.399072969 | 0.689839439 | 0.867683896 |
| MRPL46 | 469.7990451 | 0.626038386 | 0.404000984 | 1.549596191 | 0.121238468 | 0.401420514 |
| MRPL47 | 1764.286102 | 1.039971297 | 0.34406742 | 3.022579984 | 0.002506298 | 0.048313917 |
| MRPL48 | 501.9947745 | 0.527289056 | 0.366428888 | 1.438994234 | 0.150152157 | 0.444740017 |
| MRPL49 | 1153.805777 | -0.232243508 | 0.313298348 | -0.741285453 | 0.45852038 | 0.736808886 |
| MRPL50 | 1380.40847 | 0.800549083 | 0.306719809 | 2.610033845 | 0.009053327 | 0.107495835 |
| MRPL51 | 4318.62737 | 1.203048574 | 0.45225819 | 2.660092397 | 0.007811922 | 0.099092529 |
| MRPL52 | 818.7939445 | 0.773686453 | 0.354983955 | 2.179496967 | 0.02929477 | 0.195861365 |
| MRPL53 | 353.9091226 | 0.82673159 | 0.548003199 | 1.50862548 | 0.131394519 | 0.417113674 |
| MRPL54 | 554.9601254 | 1.307093764 | 0.409949699 | 3.188424741 | 0.001430502 | 0.034393268 |
| MRPL55 | 450.3386816 | 0.660141744 | 0.391794543 | 1.684918169 | 0.092004361 | 0.348253032 |
| MRPL57 | 672.0489029 | 0.889355041 | 0.414201667 | 2.147154664 | 0.031780971 | |

0.203069232
MRPL58 522.9612805 0.46306082 0.359577719 1.287790638 0.197818852 0.503399198
MRPL9 1956.531767 0.209696684 0.337019603 0.622209161 0.533804343 0.783156734
MRPS10 936.2341037 0.30606959 0.277025031 1.10484453 0.269226962 0.582188661
MRPS11 620.9719834 0.411575086 0.289723772 1.42057755 0.155439609 0.453320705
MRPS12 272.5896309 0.723197807 0.478391463 1.511728078 0.130603053 0.416016191
MRPS14 822.0307624 0.811433246 0.337889661 2.401474029 0.016329168 0.144375457
MRPS15 1932.404604 0.416217268 0.302073678 1.377866721 0.168244441 0.470495192
MRPS16 2290.997684 0.700500922 0.362831027 1.930653306 0.053525941 0.267406937
MRPS17 491.5421252 0.723643678 0.423723016 1.707822446 0.087669293 0.34024836
MRPS18A 1125.455923 0.429214517 0.388599127 1.10451745 0.269368738 0.582188661
MRPS18B 1234.809222 0.169828573 0.344773639 0.492579924 0.622309436 0.833640266
MRPS18C 1038.334644 0.760530066 0.424222019 1.79276424 0.07301063 0.31078683
MRPS2 1265.328483 0.266550419 0.296377169 0.899362186 0.368459774 0.674555106
MRPS21 2818.552221 0.707514797 0.378163959 1.87092075 0.061356066 0.285640507
MRPS22 1014.595686 0.338915442 0.278079553 1.218771529 0.222930921 0.531941648
MRPS23 1294.86547 0.866233303 0.334183737 2.592086949 0.009539566 0.110730492
MRPS24 1235.328919 1.236469349 0.459296315 2.692095077 0.00710047 0.092709113
MRPS25 1063.103422 0.411127921 0.286374515 1.435630268 0.151107577 0.446263004
MRPS26 800.8402113 0.536990924 0.324862013 1.652981579 0.098334613 0.359998651
MRPS27 3059.359058 0.069724098 0.243607176 0.286215286 0.774713237 0.906450643
MRPS28 876.7609806 1.255876836 0.398921675 3.148178993 0.001642911 0.037650448
MRPS30 1140.507671 0.708544527 0.349112874 2.029557141 0.042401575 0.236937315
MRPS31 1033.590694 0.344621685 0.295540293 1.166073437 0.243584755 0.554868517
MRPS31P4 96.30882076 -0.506374935 0.636591145 -0.795447657 0.426353147 0.714323976
MRPS31P5 358.2311952 -0.296347662 0.286532011 -1.034256734 0.301016158 0.611750318
MRPS33 1056.751664 0.834224839 0.38483177 2.167764992 0.030176572 0.198185614
MRPS34 1261.571891 1.109501158 0.461117421 2.406114162 0.016123225 0.143654707
MRPS35 1106.239592 0.185275177 0.283196056 0.654229367 0.512964044 0.771819984
MRPS36 630.5517457 0.855798545 0.385605867 2.2193608 0.026462188 0.185860119
MRPS5 1406.730969 0.600261124 0.334512383 1.794436184 0.072743576 0.310433977
MRPS6 2792.789549 1.213887046 0.417803979 2.905398481 0.003667858 0.062311776
MRPS7 1620.352733 0.427145195 0.320101398 1.334405905 0.182070866 0.486131134
MRPS9 2454.5822 0.570920829 0.378647848 1.507788391 0.131608694 0.41719445
MRRF 884.2517148 -0.338028858 0.247326714 -1.366730071 0.171709929 0.474879827
MRS2 1825.512334 -0.459312998 0.341498382 -1.344993189 0.178627434

| | | | | | | |
|---|---|---|---|---|---|---|
| | | | | | | 0.482972162 |
| MRTFA | 731.3523461 | -0.846005866 | 0.330381005 | -2.560697658 | 0.010446222 | 0.116176818 |
| MRTFB | 1179.494691 | -0.58676212 | 0.352490597 | -1.664617796 | 0.095989033 | 0.355648968 |
| MRTO4 | 952.0676991 | 0.488281777 | 0.340544635 | 1.433826075 | 0.151621902 | 0.447172604 |
| MRVI1 | 266.1081078 | -0.952172327 | 0.414267191 | -2.298449766 | 0.021536204 | 0.166469811 |
| MS4A4E | 50.57490441 | 0.711149477 | 0.602880216 | 1.179586687 | 0.23816464 | 0.548951042 |
| MS4A6A | 30.08469922 | -1.365264451 | 0.983178064 | -1.388623792 | 0.164947179 | 0.467115865 |
| MSANTD2 | 385.5613538 | -0.184632642 | 0.344876644 | -0.535358496 | 0.592401984 | 0.819859199 |
| MSANTD3 | 953.6434186 | 0.506889418 | 0.322954009 | 1.569540568 | 0.116522035 | 0.393649109 |
| MSANTD3-TMEFF1 | 4755.888578 | 0.891841462 | 0.366605968 | 2.432697611 | 0.014986811 | 0.139113039 |
| MSANTD4 | 544.5530193 | 0.3379261 | 0.25627326 | 1.318616305 | 0.18729742 | 0.49211797 |
| MSH2 | 3417.784951 | 0.095367492 | 0.254985536 | 0.374011381 | 0.708395849 | 0.875585221 |
| MSH3 | 785.5155313 | -0.083772428 | 0.279382501 | -0.299848516 | 0.764292706 | 0.902756684 |
| MSH5 | 118.9490271 | 0.138085611 | 0.893721338 | 0.154506337 | 0.877210515 | 0.950622089 |
| MSH6 | 5077.565745 | 0.016611134 | 0.25980085 | 0.063937951 | 0.949019634 | 0.98134156 |
| MSI1 | 1191.305448 | 0.436085754 | 0.251110118 | 1.73663155 | 0.082452227 | 0.328570367 |
| MSI2 | 2400.798822 | -0.22325286 | 0.267695718 | -0.833979945 | 0.404292285 | 0.700367583 |
| MSL1 | 3082.292111 | 0.308400514 | 0.3103344 | 0.993768379 | 0.320335644 | 0.630162352 |
| MSL2 | 1227.82004 | 0.069980754 | 0.263526563 | 0.265554837 | 0.790582072 | 0.91311918 |
| MSL3 | 790.4119024 | -0.226339345 | 0.270986732 | -0.835241426 | 0.403581789 | 0.700221789 |
| MSL3P1 | 503.0742167 | 0.412061207 | 0.258396714 | 1.594684392 | 0.11078283 | 0.383106067 |
| MSLN | 12.78367736 | -0.428672731 | 1.011918201 | -0.423623896 | 0.671840126 | 0.858676204 |
| MSMO1 | 2430.566773 | 0.794297323 | 0.241835385 | 3.284454523 | 0.0010218 | 0.027467362 |
| MSN | 6959.429952 | -0.311329062 | 0.34964564 | -0.890413109 | 0.373244106 | 0.678228049 |
| MSRA | 18.48950878 | 1.592056746 | 0.955385675 | 1.666402154 | 0.095633342 | 0.354905439 |
| MSRB1 | 748.3823255 | 0.50604293 | 0.316212641 | 1.600324795 | 0.109526549 | 0.380235075 |
| MSRB2 | 474.0214379 | 0.470938912 | 0.387211038 | 1.21623318 | 0.223896107 | 0.533070911 |
| MSRB3 | 100.2711771 | 0.911565914 | 0.585686041 | 1.556407101 | 0.119611318 | 0.398898209 |
| MSS51 | 66.57952124 | -0.283492444 | 0.670542807 | -0.422780531 | 0.672455394 | 0.859011927 |
| MST1 | 228.4418418 | -0.365132494 | 0.387239184 | -0.942912053 | 0.345725889 | 0.652517297 |
| MST1L | 168.8080701 | 0.215227546 | 0.421620023 | 0.510477526 | 0.609716955 | 0.828409093 |
| MST1P2 | 69.99249992 | 0.459505157 | 0.459782628 | 0.999396516 | 0.317602647 | 0.627634461 |
| MST1R | 42.72673252 | -0.11978372 | 0.543301463 | -0.220473768 | 0.8255022 | 0.92714195 |
| MSTO1 | 954.126587 | -0.593314298 | 0.356478009 | -1.6643784 | 0.096036834 | 0.355648968 |
| MSTO2P | 118.1914723 | -0.485588295 | 0.713939388 | -0.68015339 | 0.496407342 | 0.761167958 |
| MSX1 | 1226.40383 | 0.189574821 | 0.652568717 | 0.29050553 | 0.771429521 | 0.905067213 |
| MSX2 | 4144.444775 | -0.698093584 | 0.652859037 | -1.069286853 | 0.284940433 | 0.598807847 |
| MT-ATP6 | 99265.9776 | 0.170311751 | 0.391790354 | 0.434701234 | 0.663779302 | 0.854667747 |
| MT-ATP8 | 18351.36443 | -0.294906962 | 0.288807205 | -1.021120518 | 0.307197343 | 0.617205881 |
| MT-CO1 | 287140.8912 | -0.584573117 | 0.320397762 | -1.824523096 | | |

| | | | | | | |
|---|---|---|---|---|---|---|
| | | | | | 0.068073028 | 0.300140172 |
| MT-CO2 | 107019.9731 | 0.173709741 | 0.423058166 | 0.410604864 | 0.681362297 | 0.862490397 |
| MT-CO3 | 214199.117 | 0.246413458 | 0.440538146 | 0.559346472 | 0.575925285 | 0.808969969 |
| MT-CYB | 117818.0817 | -0.579309364 | 0.357272206 | -1.621478957 | 0.104914959 | 0.37133017 |
| MT-ND1 | 50582.47353 | -0.89747688 | 0.370659032 | -2.42130045 | 0.015465089 | 0.140744373 |
| MT-ND2 | 56416.19381 | -0.191131172 | 0.371253221 | -0.514826973 | 0.606673935 | 0.826667578 |
| MT-ND3 | 26880.95326 | 1.138154683 | 0.502959013 | 2.262917361 | 0.023640784 | 0.175897172 |
| MT-ND4 | 141973.3451 | -0.553009142 | 0.384098713 | -1.439757861 | 0.149935917 | 0.444707291 |
| MT-ND4L | 16966.53545 | -1.093486555 | 0.671524276 | -1.628364892 | 0.103447537 | 0.368923975 |
| MT-ND5 | 49512.85839 | -1.107847471 | 0.538268153 | -2.058170197 | 0.039573798 | 0.229237914 |
| MT-ND6 | 1964.568854 | 0.404327223 | 0.488739513 | 0.827285726 | 0.408075132 | 0.702586667 |
| MT1E | 247.6882695 | 1.827710491 | 0.947408499 | 1.929168349 | 0.053709966 | 0.267861241 |
| MT1F | 42.85721183 | 0.572718423 | 0.810976713 | 0.706208223 | 0.480058657 | 0.752036291 |
| MT1G | 64.21406108 | 2.02650788 | 1.36895402 | 1.48033305 | 0.138784387 | 0.427314124 |
| MT1H | 25.80748186 | 1.171916037 | 1.408229134 | 0.832191302 | 0.405300973 | 0.701039911 |
| MT1X | 458.3192241 | 1.292718931 | 1.434500708 | 0.901162978 | 0.367501672 | 0.673938238 |
| MT2A | 164.5910538 | 1.796206818 | 0.861498798 | 2.084978901 | 0.037071216 | 0.220638452 |
| MTA1 | 2517.963337 | 0.129293818 | 0.351616587 | 0.367712509 | 0.713087612 | 0.877748654 |
| MTA2 | 3496.158791 | -0.098185563 | 0.249885235 | -0.392922629 | 0.694376628 | 0.86892666 |
| MTA3 | 4115.370619 | -0.467131775 | 0.345893863 | -1.350506108 | 0.176853696 | 0.481149663 |
| MTAP | 1292.430151 | 0.013375983 | 0.340341951 | 0.039301599 | 0.968649931 | 0.989475309 |
| MTATP6P1 | 12463.75389 | -0.247353276 | 0.346055158 | -0.714779913 | 0.474745049 | 0.748737095 |
| MTBP | 408.5495624 | -0.107666613 | 0.399259646 | -0.269665652 | 0.787417487 | 0.911971196 |
| MTCH1 | 6618.681492 | 0.03518876 | 0.260876284 | 0.134886772 | 0.892701399 | 0.957550733 |
| MTCH2 | 1366.136258 | 0.427291674 | 0.382199994 | 1.11797928 | 0.263575843 | 0.577444856 |
| MTCL1 | 2185.104838 | -0.462601198 | 0.382743991 | -1.208643921 | 0.226799675 | 0.535481989 |
| MTCO1P12 | 205.3559923 | 0.523821927 | 0.511799838 | 1.023489826 | 0.306076308 | 0.616173522 |
| MTCO1P40 | 123.5766423 | 1.440966532 | 0.5968597 | 2.414246652 | 0.015767784 | 0.142219415 |
| MTCP1 | 124.3804395 | 0.369638095 | 0.610796291 | 0.605174099 | 0.545063323 | 0.791079659 |
| MTDH | 1773.320345 | 0.408246356 | 0.222982544 | 1.830844465 | 0.067123761 | 0.297817842 |
| MTERF1 | 561.8573624 | 0.206387735 | 0.261802791 | 0.788332829 | 0.43050205 | 0.717078019 |
| MTERF2 | 392.8649641 | 0.46106908 | 0.343287083 | 1.343100581 | 0.179239407 | 0.483850721 |
| MTERF3 | 557.8634689 | 0.808435582 | 0.243282066 | 3.323038137 | 0.000890427 | 0.025375039 |
| MTERF4 | 737.5466804 | 0.174329895 | 0.254716749 | 0.684406877 | 0.493718274 | 0.759809286 |
| MTF1 | 495.3832343 | -0.130952716 | 0.356268248 | -0.367567744 | 0.71319557 | 0.877795412 |
| MTF2 | 2716.56609 | 0.18102145 | 0.229438201 | 0.788976942 | 0.430125483 | 0.716765565 |
| MTFMT | 346.3706589 | 0.436562544 | 0.280968515 | 1.553777452 | 0.120237511 | 0.400307118 |
| MTFP1 | 339.4793004 | 0.932978223 | 0.318047728 | 2.933453507 | 0.003352139 | 0.058568145 |
| MTFR1 | 1103.080312 | 0.461432574 | 0.299909418 | 1.538573134 | 0.123908541 | 0.405045568 |
| MTFR1L | 902.8243672 | -0.070532249 | 0.24668966 | -0.285914899 | 0.774943302 | 0.906450643 |
| MTFR2 | 192.3262361 | 0.347975111 | 0.314003893 | 1.108187251 | 0.267780956 | 0.581021864 |

MTG1 1283.44889 0.037406045 0.352492295 0.106118759 0.915488127 0.969148725
MTG2 530.3299659 -0.145329488 0.269378501 -0.539499208 0.589542444
0.81828099
MTHFD1 2907.400878 -0.180580915 0.417480859 -0.432548969
0.66534247 0.85512628
MTHFD1L 1311.281294 0.332347841 0.243235741 1.366361045 0.171825669
0.474879827
MTHFD2 2920.071608 0.931732354 0.28921716 3.221566636 0.001274918
0.031663479
MTHFD2L 229.6417982 0.238382831 0.298058395 0.799785665 0.42383499
0.712860381
MTHFR 629.610245 -0.324730146 0.288143091 -1.126975301 0.259752924
0.573027717
MTHFS 364.7387131 0.763782594 0.55268135 1.381958329 0.166984496 0.469215449
MTHFSD 427.3314188 -0.382532679 0.266107839 -1.437509997
0.150573134 0.445465098
MTIF2 1450.627659 -0.153850933 0.246240783 -0.624798751 0.532103151
0.782369637
MTIF3 1267.208332 0.286667583 0.338425442 0.847062743 0.39696015 0.695574317
MTLN 86.33840528 0.721068513 0.594098365 1.213719066 0.224855021 0.53418032
MTM1 408.1754018 0.346206181 0.384748097 0.899825583 0.368213078 0.674331313
MTMR1 584.4306582 -0.3747406 0.329138842 -1.138548698 0.254891436
0.567256913
MTMR10 597.8110426 -0.41110591 0.294621244 -1.3953709 0.162904022
0.464171436
MTMR11 232.8376591 -1.419479676 0.417711963 -3.398226055
0.000678244 0.021392167
MTMR12 858.5593975 -0.029611124 0.315706696 -0.093793144
0.925273478 0.971979223
MTMR14 1251.226795 -0.33960551 0.273228167 -1.24293741 0.213890899
0.520856954
MTMR2 2062.047222 -0.128763231 0.247052959 -0.52119688 0.602229628
0.825021776
MTMR3 4780.313817 -0.388161554 0.381821807 -1.016603941 0.309341879
0.61913267
MTMR4 1129.684666 -0.406156267 0.439988648 -0.923106242 0.355951842
0.663487122
MTMR6 678.1162711 -0.165806953 0.262098596 -0.632612898 0.526986462
0.778755887
MTMR7 258.6820054 -0.694284089 0.386002741 -1.798650672 0.072073956
0.309054266
MTMR8 26.68679181 -0.349518201 0.612254079 -0.570871167 0.568086977
0.80531186
MTMR9 822.3063306 0.023841006 0.298932791 0.079753734 0.936433122 0.976065056
MTMR9LP 409.6353239 0.364212534 0.28962659 1.257524505 0.208563771
0.515581062
MTND1P23 7611.600674 1.194616729 0.646623326 1.847469279 0.064679157
0.29355313
MTND2P28 230.2365053 -0.49134178 0.586298659 -0.838040089 0.402008189
0.699116854
MTND4P12 41.5542274 0.603799287 0.521136304 1.158620656 0.246610845
0.558824964
MTND5P11 24.54920503 -0.328877519 0.779668036 -0.421817369
0.673158327 0.85934744
MTO1 571.4835075 -0.016130317 0.265792519 -0.060687626 0.951607986
0.982350991
MTOR 2960.234091 -0.671909328 0.566361417 -1.186361409 0.235479605
0.546164361
MTPAP 889.5974761 0.215833732 0.300961813 0.717146568 0.473283664 0.747624581
MTPN 4380.813476 -0.190115181 0.247048969 -0.769544526 0.441570122
0.725085001
MTR 994.7878879 -0.420472727 0.363079213 -1.158074359 0.246833693
0.559246409
MTREX 3410.256945 -0.061477739 0.310806234 -0.197800857 0.843200876

| | | | | | | |
|---|---|---|---|---|---|---|
| | | | | | | 0.936162912 |
| MTRF1 | 474.635491 | -0.242198479 | 0.279543591 | -0.866406839 | 0.386267095 | 0.686679504 |
| MTRF1L | 670.6053399 | -0.053757979 | 0.28509454 | -0.18856194 | 0.850436165 | 0.939032396 |
| MTRNR2L1 | 14544.38581 | 1.081254231 | 0.453196394 | 2.385840323 | 0.017040148 | 0.14773715 |
| MTRNR2L10 | 51.99957415 | 1.85594788 | 0.557832316 | 3.327071287 | 0.000877639 | 0.025175225 |
| MTRNR2L11 | 515.9848331 | 1.468394107 | 0.629863085 | 2.331290948 | 0.019738023 | 0.159110282 |
| MTRNR2L12 | 90105.88211 | 0.073682381 | 0.343976791 | 0.21420742 | 0.83038531 | 0.930264006 |
| MTRNR2L4 | 283.805111 | 1.312626135 | 0.315049867 | 4.166407525 | 3.09E-05 | 0.002537553 |
| MTRNR2L6 | 619.2857139 | 0.78272462 | 0.313053434 | 2.5002908 | 0.01240914 | 0.126591155 |
| MTRNR2L8 | 13559.68509 | 0.853959979 | 0.331725284 | 2.574298731 | 0.010044351 | 0.114059016 |
| MTRR | 844.4855548 | 0.112097501 | 0.30857958 | 0.363269343 | 0.716403692 | 0.879925471 |
| MTSS1 | 191.6069279 | 0.165746008 | 0.444660449 | 0.372747359 | 0.709336485 | 0.875945015 |
| MTSS1L | 2328.562202 | -0.644933301 | 0.445958292 | -1.446174031 | 0.148128396 | 0.442025156 |
| MTTP | 14.90550062 | 1.307001849 | 0.872833331 | 1.49742431 | 0.134282887 | 0.420619758 |
| MTURN | 249.820983 | 0.288555654 | 0.283908738 | 1.016367639 | 0.30945435 | 0.61915097 |
| MTUS1 | 300.1084156 | -2.343965582 | 0.523206715 | -4.479999039 | 7.46E-06 | 0.000875358 |
| MTUS2 | 122.4769354 | -1.731037471 | 1.174304325 | -1.474096139 | 0.140455708 | 0.430123922 |
| MTX1 | 1071.295417 | 0.507079562 | 0.394889163 | 1.284106046 | 0.199104853 | 0.504867077 |
| MTX2 | 1098.534174 | 0.90513388 | 0.30635272 | 2.954548206 | 0.003131272 | 0.055756493 |
| MTX3 | 1053.272315 | -0.066469593 | 0.318987383 | -0.208376872 | 0.834934713 | 0.932204484 |
| MUC1 | 145.0737399 | -0.770411117 | 0.603110888 | -1.277395471 | 0.201462663 | 0.507874719 |
| MUC12 | 31.41081221 | 0.296740871 | 0.718862257 | 0.412792393 | 0.679758725 | 0.861682789 |
| MUC15 | 891.0343117 | -1.607222776 | 1.330691089 | -1.20781058 | 0.227120131 | 0.535952637 |
| MUC16 | 82.79831539 | -3.068809572 | 0.833674891 | -3.681062732 | 0.000232264 | 0.010434843 |
| MUC19 | 74.39045039 | -0.658196687 | 0.507792062 | -1.296193337 | 0.194908885 | 0.500501471 |
| MUC3A | 334.0868141 | -1.394135983 | 0.509546575 | -2.736032484 | 0.006218489 | 0.085454377 |
| MUL1 | 531.8127602 | -0.369125376 | 0.342501501 | -1.077733603 | 0.281152658 | 0.595171527 |
| MUS81 | 688.9809355 | 0.007624574 | 0.311655114 | 0.024464781 | 0.980481876 | 0.993221987 |
| MUSTN1 | 568.5712046 | 1.298139519 | 0.683414093 | 1.899491879 | 0.057499833 | 0.277812093 |
| MUTYH | 692.1481661 | 0.051180934 | 0.338376364 | 0.151254459 | 0.879774992 | 0.952137709 |
| MVB12A | 734.6518548 | 0.217943305 | 0.372182225 | 0.58558225 | 0.558156279 | 0.798855459 |
| MVB12B | 401.8653782 | 0.112677596 | 0.565438509 | 0.199274713 | 0.842047859 | 0.935422069 |
| MVD | 840.039994 | 0.467603307 | 0.456038845 | 1.025358501 | 0.305194062 | 0.615135352 |
| MVK | 348.1058209 | 0.345395433 | 0.390930583 | 0.883521137 | 0.376954768 | 0.680321292 |
| MVP | 260.8235132 | -0.721350072 | 0.428178719 | -1.68469389 | 0.092047646 | 0.348263686 |
| MX1 | 37.19200934 | 0.17240834 | 0.741164311 | 0.232618243 | 0.81605786 | 0.923222505 |
| MX2 | 121.4445305 | -0.741405667 | 0.350391615 | -2.115934384 | 0.034350389 | 0.211669271 |
| MXD1 | 97.60722496 | -0.374094615 | 0.383037208 | -0.976653461 | 0.328740742 | 0.638091838 |
| MXD3 | 100.8079296 | 1.45210216 | 0.584234511 | 2.485478235 | 0.01293775 | 0.128476822 |

| | | | | | | |
|---|---|---|---|---|---|---|
| MXD4 | 1112.276628 | -0.072700875 | 0.583857066 | -0.124518275 | 0.900904931 | 0.961704658 |
| MXI1 | 1216.510603 | -0.20945289 | 0.335345504 | -0.624588335 | 0.532241278 | 0.782369637 |
| MXRA5 | 312.0968321 | -3.090321396 | 0.876578977 | -3.525434077 | 0.000422789 | 0.015778519 |
| MXRA7 | 1639.587654 | -0.009889075 | 0.221030452 | -0.044740782 | 0.964313927 | 0.988486931 |
| MXRA8 | 2816.740887 | -1.426297647 | 0.694614913 | -2.053364562 | 0.040037234 | 0.230689279 |
| MYADM | 1675.877044 | -0.312749051 | 0.556513111 | -0.561979663 | 0.574129875 | 0.808359708 |
| MYB | 67.01808907 | 0.178108786 | 0.53449557 | 0.333227806 | 0.73896233 | 0.891443823 |
| MYBBP1A | 605.7105238 | -0.248339731 | 0.403970212 | -0.614747632 | 0.538721398 | 0.786598616 |
| MYBL1 | 168.1214568 | 0.613353137 | 0.536861967 | 1.142478282 | 0.253255268 | 0.565852088 |
| MYBL2 | 2956.939695 | -0.736990328 | 0.301648756 | -2.443206923 | 0.014557388 | 0.137477204 |
| MYBPC1 | 59.61367533 | 3.397542093 | 2.780593738 | 1.221876482 | 0.22175434 | 0.53065007 |
| MYBPC2 | 106.88469 | -0.777742238 | 0.553425383 | -1.405324479 | 0.159924812 | 0.460121245 |
| MYC | 89.05501212 | -1.390687206 | 0.716077591 | -1.942090109 | 0.052126193 | 0.263030768 |
| MYCBP | 1037.735723 | 0.33711804 | 0.328954861 | 1.024815501 | 0.305450251 | 0.61550193 |
| MYCBP2 | 1031.334077 | -0.421750342 | 0.391415775 | -1.077499603 | 0.281257128 | 0.595171527 |
| MYCBPAP | 55.36589889 | 0.6270402 | 0.648891738 | 0.966324833 | 0.333881664 | 0.64183994 |
| MYCL | 313.0259927 | 0.632165489 | 0.373655506 | 1.691840422 | 0.090676404 | 0.345245839 |
| MYCN | 1348.033521 | 0.931861163 | 0.274821275 | 3.390789754 | 0.000696915 | 0.021675701 |
| MYCT1 | 70.46261064 | -3.027734663 | 1.089884682 | -2.778032129 | 0.005468921 | 0.078942912 |
| MYD88 | 574.0454216 | -0.58676131 | 0.330667888 | -1.77447321 | 0.07598491 | 0.316259825 |
| MYDGF | 855.2812113 | 0.896582477 | 0.396071589 | 2.263687931 | 0.023593315 | 0.175630121 |
| MYEF2 | 2891.674843 | 0.580116293 | 0.319432553 | 1.816083822 | 0.069357515 | 0.30336211 |
| MYF6 | 18.8546634 | 0.01088619 | 0.950898963 | 0.011448314 | 0.990865767 | 0.996726674 |
| MYH10 | 10063.75086 | -0.282951296 | 0.411870249 | -0.686991345 | 0.492088182 | 0.759174768 |
| MYH14 | 41.96601538 | -1.293918982 | 0.642676714 | -2.013327935 | 0.044080141 | 0.241443505 |
| MYH15 | 24.28922939 | -1.539570452 | 0.764209143 | -2.014593082 | 0.043947303 | 0.241112355 |
| MYH3 | 41.04383453 | 0.098417731 | 0.669801013 | 0.146935775 | 0.883182714 | 0.953716631 |
| MYH6 | 162.2736362 | -4.080318217 | 0.912081921 | -4.473631286 | 7.69E-06 | 0.000883852 |
| MYH7B | 21.61603666 | -1.196966479 | 0.748924653 | -1.598246865 | 0.109988047 | 0.381488036 |
| MYH9 | 12797.27616 | -1.272153687 | 0.366327184 | -3.472725317 | 0.000515202 | 0.017892193 |
| MYL12A | 7092.921106 | 0.009898555 | 0.457520715 | 0.021635207 | 0.982738949 | 0.993518428 |
| MYL12B | 4802.003172 | -0.200006043 | 0.486884933 | -0.410787086 | 0.681228663 | 0.862490397 |
| MYL3 | 88.58731746 | -0.893246979 | 0.785780019 | -1.136764689 | 0.255636668 | 0.567910953 |
| MYL4 | 51.81077132 | 1.144963468 | 0.691846431 | 1.654938751 | 0.097936923 | 0.359375073 |
| MYL5 | 150.320828 | -0.256206523 | 0.492735331 | -0.519967835 | 0.603085994 | 0.825089513 |
| MYL6 | 16105.52937 | 0.590684274 | 0.463156838 | 1.275343955 | 0.202187526 | 0.508856497 |
| MYL6B | 2847.940384 | 0.381269743 | 0.372708966 | 1.022969067 | 0.306322472 | 0.616341939 |
| MYL7 | 20.92277315 | -0.981065737 | 0.756709373 | -1.296489474 | 0.194806904 | 0.500501471 |
| MYL9 | 2989.599018 | 0.225983813 | 0.2282702 | 0.989983858 | 0.322182009 | 0.632029271 |

| | | | | | | |
|---|---|---|---|---|---|---|
| MYLIP | 762.8719229 | -1.129520632 | 0.639504157 | -1.766244393 | 0.077354856 | 0.318812965 |
| MYLK | 486.3977619 | 0.101995019 | 0.506144263 | 0.201513732 | 0.840296893 | 0.934477655 |
| MYLK3 | 230.6258984 | -1.869381137 | 0.427952144 | -4.368201362 | 1.25E-05 | 0.001310711 |
| MYLK4 | 20.93609049 | -1.545952896 | 0.73222433 | -2.111310472 | 0.034745635 | 0.212894193 |
| MYLPF | 14.66544162 | -0.290639735 | 0.832852378 | -0.348969088 | 0.727112517 | 0.8850111 |
| MYNN | 744.166952 | 0.290383194 | 0.305850578 | 0.9494283 | 0.342402823 | 0.649080747 |
| MYO10 | 2727.129039 | -0.268839396 | 0.346956818 | -0.774849728 | 0.438428461 | 0.722219855 |
| MYO15A | 63.71859 | 0.26702485 | 0.48288307 | 0.552980352 | 0.580276865 | 0.811670691 |
| MYO15B | 512.6572248 | -1.036924535 | 0.3949054 | -2.625754252 | 0.008645721 | 0.104918628 |
| MYO18A | 1235.773446 | -0.227129834 | 0.350379162 | -0.648240132 | 0.516829649 | 0.773950969 |
| MYO19 | 832.8047455 | -0.151113965 | 0.33867127 | -0.446196587 | 0.655455248 | 0.849471345 |
| MYO1B | 2411.726792 | -0.375931384 | 0.376023548 | -0.999754898 | 0.317429138 | 0.627531109 |
| MYO1C | 2063.227144 | -0.87574679 | 0.300122431 | -2.917965131 | 0.003523237 | 0.060947589 |
| MYO1D | 282.8558174 | -0.73995552 | 0.413390542 | -1.789967221 | 0.073459181 | 0.311470066 |
| MYO1E | 1107.559819 | -1.316121658 | 0.258375273 | -5.093837517 | 3.51E-07 | 9.51E-05 |
| MYO1F | 19.22236537 | -0.461092939 | 0.77429361 | -0.595501413 | 0.551508355 | 0.795033566 |
| MYO3A | 24.08701328 | -0.710797327 | 0.658624976 | -1.079214049 | 0.280492318 | 0.594488539 |
| MYO3B | 483.2328355 | 0.079115016 | 1.410368017 | 0.056095299 | 0.955265889 | 0.984323736 |
| MYO5A | 894.8806381 | -0.299599969 | 0.398687349 | -0.751465955 | 0.452372282 | 0.732985143 |
| MYO5B | 1979.519732 | -2.070403393 | 0.492509256 | -4.203785754 | 2.62E-05 | 0.002237189 |
| MYO5C | 564.3900935 | -1.15914101 | 0.439735807 | -2.635994141 | 0.008389118 | 0.103556802 |
| MYO6 | 3215.669791 | -0.627641174 | 0.365058256 | -1.719290452 | 0.085561498 | 0.335501029 |
| MYO7A | 874.4800345 | -2.429842912 | 0.580989623 | -4.182248384 | 2.89E-05 | 0.002406024 |
| MYO9A | 791.6208568 | -0.509882734 | 0.305093694 | -1.671233275 | 0.09467561 | 0.3531654 |
| MYO9B | 1321.872029 | -0.438381354 | 0.430883976 | -1.017399992 | 0.308963186 | 0.618619571 |
| MYOCD | 51.65949056 | -0.849419992 | 0.654140402 | -1.298528556 | 0.19410577 | 0.500132242 |
| MYOF | 1709.092019 | -2.599497331 | 0.615946033 | -4.22033294 | 2.44E-05 | 0.00212692 |
| MYOM1 | 39.96796189 | -1.75960055 | 0.628206292 | -2.800991606 | 0.005094584 | 0.075897737 |
| MYOM3 | 16.55309549 | 0.050244388 | 0.760192787 | 0.06609427 | 0.947302773 | 0.980790989 |
| MYORG | 165.4514177 | -1.910255665 | 0.635385676 | -3.006450627 | 0.00264317 | 0.050167035 |
| MYOZ1 | 67.40697346 | 1.030095393 | 0.461665694 | 2.231258261 | 0.025664027 | 0.182645569 |
| MYOZ3 | 65.48570774 | 0.434167525 | 0.544420642 | 0.797485421 | 0.425169167 | 0.71357297 |
| MYPOP | 138.6937496 | 0.498794733 | 0.483520997 | 1.031588569 | 0.302264898 | 0.613115536 |
| MYRF | 237.5048713 | 0.323855122 | 0.539990301 | 0.599742478 | 0.548677874 | 0.793468239 |
| MYRFL | 18.57220853 | -0.468209444 | 0.863091733 | -0.542479351 | 0.587488328 | 0.816787521 |
| MYRIP | 37.64287303 | -0.700459539 | 0.98006639 | -0.714706214 | 0.474790597 | 0.748737095 |

| | | | | | | |
|---|---|---|---|---|---|---|
| MYSM1 | 1784.585236 | -0.658329589 | 0.350511332 | -1.87819773 | 0.060354121 | 0.283945763 |
| MYT1 | 20.09096121 | 0.323451078 | 0.816000409 | 0.396385927 | 0.691820346 | 0.86812349 |
| MYT1L | 30.0307059 | -0.516334325 | 0.791437362 | -0.652400745 | 0.514142684 | 0.77274209 |
| MYZAP | 95.44955162 | -0.988843275 | 0.776837285 | -1.272909134 | 0.203050288 | 0.509506436 |
| MZF1 | 397.1786853 | -0.494069368 | 0.541822827 | -0.911865176 | 0.361839695 | 0.668713609 |
| MZT1 | 1233.354472 | 1.371388739 | 0.264348737 | 5.187801363 | 2.13E-07 | 6.73E-05 |
| MZT2A | 375.427245 | 0.922866489 | 0.547504808 | 1.685586092 | 0.09187555 | 0.347939083 |
| MZT2B | 238.9425182 | 1.025448869 | 0.694380167 | 1.476783062 | 0.139733798 | 0.428675153 |
| N4BP2 | 1075.886335 | -0.470851815 | 0.349721951 | -1.346360483 | 0.178186289 | 0.482467283 |
| N4BP2L1 | 53.2469066 | -0.567044424 | 0.651564581 | -0.870281229 | 0.384146735 | 0.684816878 |
| N4BP2L2 | 2032.086144 | -0.012875276 | 0.252103884 | -0.051071313 | 0.959268695 | 0.985871727 |
| N4BP3 | 323.6995479 | 0.002446423 | 0.395783558 | 0.006181214 | 0.995068136 | 0.998094459 |
| N6AMT1 | 344.679857 | 0.396838038 | 0.443155546 | 0.895482503 | 0.370529215 | 0.676124456 |
| NAA10 | 1358.155459 | 0.461445463 | 0.351275426 | 1.313628646 | 0.188971204 | 0.494119637 |
| NAA15 | 2825.244138 | 0.366689982 | 0.282219092 | 1.299309623 | 0.193837693 | 0.499845061 |
| NAA16 | 690.8134915 | 0.098748599 | 0.331200716 | 0.298153338 | 0.765586134 | 0.903026935 |
| NAA20 | 784.3575261 | 0.506587465 | 0.302460037 | 1.674890573 | 0.093955704 | 0.351340153 |
| NAA25 | 1070.664248 | 0.372897778 | 0.385855289 | 0.96641873 | 0.333834696 | 0.64183994 |
| NAA30 | 1238.600666 | -0.056060029 | 0.310382065 | -0.1806162 | 0.856668839 | 0.942343063 |
| NAA35 | 1296.037404 | 0.060305236 | 0.261905773 | 0.230255467 | 0.817893263 | 0.924112493 |
| NAA38 | 410.9961015 | 1.083282194 | 0.50862598 | 2.129820804 | 0.03318641 | 0.207874084 |
| NAA40 | 1615.986486 | -0.746834421 | 0.396525457 | -1.883446343 | 0.059639905 | 0.282669434 |
| NAA50 | 3539.140708 | 0.338711709 | 0.260948524 | 1.298002013 | 0.194286643 | 0.500187881 |
| NAA60 | 964.9386956 | -0.276884428 | 0.325113732 | -0.851654053 | 0.39440613 | 0.693260965 |
| NAA80 | 166.3846077 | 0.35522248 | 0.340743934 | 1.042490987 | 0.297184102 | 0.608609612 |
| NAAA | 22.10970657 | -0.276055437 | 0.720567536 | -0.383108347 | 0.701639431 | 0.872479299 |
| NAALAD2 | 525.0493463 | 0.024545926 | 0.33556626 | 0.073147778 | 0.941688522 | 0.978517574 |
| NAALADL1 | 22.16838694 | -0.807630717 | 0.691279815 | -1.16831231 | 0.242680812 | 0.553890567 |
| NAALADL2 | 82.79602518 | -2.534393689 | 0.757117662 | -3.347423811 | 0.000815664 | 0.024028038 |
| NAB1 | 1215.53177 | -0.707840324 | 0.521659178 | -1.356901889 | 0.174812387 | 0.478752143 |
| NAB2 | 1208.294748 | 0.689722383 | 0.361425801 | 1.908337428 | 0.056347619 | 0.275048173 |
| NABP1 | 263.7702651 | -0.630736885 | 0.454964336 | -1.386343576 | 0.165642017 | 0.467560858 |
| NABP2 | 880.6826605 | 0.55839264 | 0.304857091 | 1.831653773 | 0.06700302 | 0.297817842 |
| NACA | 21377.75125 | 0.37258923 | 0.271140038 | 1.374157915 | 0.169392663 | 0.472052919 |
| NACA3P | 10.58778233 | 1.404638479 | 1.160362883 | 1.210516554 | 0.226080743 | 0.534727976 |
| NACAD | 48.75007987 | -0.130944571 | 0.49258039 | -0.265833909 | 0.790367127 | 0.913087091 |
| NACC1 | 3266.425077 | -0.174640968 | 0.323811518 | -0.539329078 | 0.589659808 | 0.81828099 |
| NACC2 | 521.8947768 | 0.261728041 | 0.404775791 | 0.646600036 | 0.517890835 | 0.773950969 |
| NADK | 1327.138449 | 0.126325979 | 0.290873423 | 0.434298803 | 0.664071473 | 0.854858903 |
| NADK2 | 755.764056 | 0.165537457 | 0.241673468 | 0.684963302 | 0.493367079 | 0.759455148 |
| NADSYN1 | 1454.631269 | -0.614393568 | 0.288678453 | -2.128297285 | 0.033312443 | 0.208274771 |
| NAE1 | 3402.683625 | 0.542631143 | 0.227666964 | 2.383442613 | 0.017151558 | 0.147741524 |
| NAF1 | 356.1035331 | -0.007743388 | 0.271747902 | -0.028494747 | 0.977267558 | |

| | | | | | | |
|---|---|---|---|---|---|---|
| | | | | | | 0.99216657 |
| NAGA | 595.0310569 | 0.134798694 | 0.340152599 | 0.396288886 | 0.691891925 | 0.86812349 |
| NAGK | 1466.238982 | -0.534843596 | 0.434543105 | -1.230818278 | 0.218390838 | 0.526742035 |
| NAGLU | 550.6422036 | 0.305265733 | 0.319347937 | 0.955903258 | 0.339121113 | 0.646170109 |
| NAGPA | 203.1916025 | -0.556422253 | 0.392348917 | -1.418182207 | 0.156137575 | 0.454657035 |
| NAGS | 18.21915079 | 0.064992198 | 0.68813239 | 0.09444723 | 0.924753899 | 0.971780852 |
| NAIF1 | 195.1835112 | 0.091733874 | 0.338291705 | 0.271167967 | 0.786261849 | 0.911261183 |
| NAIP | 14.89098952 | 1.226684902 | 1.000615104 | 1.225930827 | 0.220224707 | 0.528727493 |
| NAMPT | 2815.964195 | 0.28498949 | 0.250248831 | 1.138824461 | 0.254776377 | 0.567256913 |
| NAMPTP1 | 147.8797721 | -0.088170959 | 0.464388467 | -0.189864661 | 0.849415185 | 0.938580023 |
| NANOG | 49.88800337 | -0.94634973 | 0.838827327 | -1.128181808 | 0.259243145 | 0.572529055 |
| NANOS1 | 88.46113196 | 1.428709391 | 0.401942102 | 3.554515399 | 0.000378676 | 0.014655354 |
| NANP | 206.9104828 | 0.37048762 | 0.376257964 | 0.984663861 | 0.32478921 | 0.634714054 |
| NANS | 681.9797135 | 0.484054937 | 0.33712813 | 1.435818889 | 0.151053884 | 0.44619129 |
| NAP1L1 | 33079.24955 | 0.588759979 | 0.344296449 | 1.710037908 | 0.087258863 | 0.33929417 |
| NAP1L2 | 457.1274689 | 0.414319657 | 0.290243945 | 1.427487681 | 0.153439374 | 0.450006832 |
| NAP1L3 | 637.5287519 | 0.64180612 | 0.398195034 | 1.61178836 | 0.107008002 | 0.375010025 |
| NAP1L4 | 1820.954475 | 0.104385284 | 0.324692644 | 0.321489527 | 0.747839448 | 0.894202292 |
| NAP1L5 | 163.5513665 | 0.121390722 | 0.313007543 | 0.387820437 | 0.69814892 | 0.870467 |
| NAP1L6 | 255.3061667 | 2.845984692 | 0.507989626 | 5.602446476 | 2.11E-08 | 1.34E-05 |
| NAPA | 1487.922807 | -0.141121067 | 0.271108842 | -0.520532881 | 0.602692216 | 0.825021776 |
| NAPB | 523.20222 | -0.144804753 | 0.325209738 | -0.445265735 | 0.656127727 | 0.84995394 |
| NAPEPLD | 91.55472743 | 0.242681314 | 0.515822726 | 0.470474257 | 0.638016222 | 0.842182924 |
| NAPG | 826.7384192 | 0.085018223 | 0.254547275 | 0.333997772 | 0.73838124 | 0.891242475 |
| NAPRT | 397.2133493 | 0.577160976 | 0.593923134 | 0.971777227 | 0.331161387 | 0.639084337 |
| NARF | 1596.314366 | 0.540027268 | 0.337174079 | 1.601627471 | 0.109238012 | 0.379407025 |
| NARS | 5096.464838 | -0.097635267 | 0.239599594 | -0.407493456 | 0.683645599 | 0.863715015 |
| NARS2 | 823.8239575 | 0.415697651 | 0.243778863 | 1.705224345 | 0.088152591 | 0.341577124 |
| NASP | 12887.57318 | 0.093279397 | 0.267899107 | 0.348188532 | 0.727698599 | 0.88517603 |
| NAT1 | 150.5358109 | -0.374766835 | 0.434358544 | -0.862805257 | 0.388244548 | 0.688412581 |
| NAT10 | 2010.813899 | -0.26460674 | 0.29057251 | -0.910639273 | 0.362485471 | 0.669662332 |
| NAT14 | 1289.814901 | 1.156433053 | 0.544540428 | 2.123686311 | 0.033696383 | 0.209168504 |
| NAT8L | 204.962414 | 1.226310031 | 0.462357942 | 2.652295805 | 0.007994647 | 0.100486153 |
| NAT9 | 1238.555411 | 0.102572363 | 0.256141438 | 0.400452045 | 0.688823597 | 0.867303742 |
| NATD1 | 974.9595505 | -1.656781197 | 0.376171052 | -4.40432933 | 1.06E-05 | 0.001149871 |
| NAV1 | 2398.907357 | -0.475649704 | 0.388573391 | -1.22409232 | 0.220917403 | 0.529719917 |
| NAV2 | 3266.746127 | -0.687221509 | 0.313519818 | -2.191955565 | 0.028382715 | 0.192544525 |
| NAV3 | 21.83964748 | -0.853149311 | 0.788114014 | -1.082520163 | 0.279021461 | 0.592557709 |
| NAXD | 1151.230502 | -0.287671659 | 0.266614626 | -1.07897929 | 0.28059696 | 0.594538239 |
| NAXE | 1068.770147 | 0.741202018 | 0.341319617 | 2.171577552 | 0.029887542 | 0.197226575 |
| NBAS | 1600.700036 | -0.345504799 | 0.419521958 | -0.823567855 | 0.410185147 | 0.704109399 |
| NBDY | 963.3821466 | 0.526034388 | 0.421444033 | 1.248171398 | 0.211968296 | 0.519175172 |
| NBEA | 907.3079545 | -0.593080278 | 0.440426588 | -1.346604165 | 0.178107752 | |

| | | | | | | |
|---|---|---|---|---|---|---|
| | | | | | | 0.482426835 |
| NBEAL1 | 127.2676322 | -1.225198447 | 0.689120524 | -1.77791606 | 0.075417642 | 0.314738565 |
| NBEAL2 | 2009.787754 | -1.754529234 | 0.342059953 | -5.129303263 | 2.91E-07 | 8.32E-05 |
| NBL1 | 27.51677241 | 1.238829189 | 0.713292974 | 1.736774698 | 0.082426947 | 0.328570367 |
| NBN | 1465.847102 | 0.09628719 | 0.287716264 | 0.334660228 | 0.737881407 | 0.891202836 |
| NBPF1 | 978.7178414 | -0.070434232 | 0.336894806 | -0.209068917 | 0.834394439 | 0.932107564 |
| NBPF10 | 152.0742624 | -0.340657266 | 0.490961557 | -0.693857311 | 0.487771697 | 0.756677644 |
| NBPF11 | 483.6387659 | -0.622818578 | 0.330727774 | -1.883175915 | 0.059676532 | 0.282669434 |
| NBPF12 | 535.4258727 | -0.831406314 | 0.343262811 | -2.42206929 | 0.015432408 | 0.140677297 |
| NBPF14 | 804.0347293 | -0.357106179 | 0.42356092 | -0.843104646 | 0.399169921 | 0.697352242 |
| NBPF15 | 895.7124617 | -0.108153602 | 0.282605405 | -0.382701815 | 0.701940868 | 0.872523139 |
| NBPF19 | 765.1007544 | -0.384453741 | 0.435776205 | -0.882227475 | 0.377653807 | 0.680934859 |
| NBPF20 | 128.3643089 | -0.947103563 | 0.596135387 | -1.588739042 | 0.112119321 | 0.384920166 |
| NBPF25P | 59.82853113 | 1.2130123 | 0.708182929 | 1.712851653 | 0.086739835 | 0.33809963 |
| NBPF26 | 122.9925322 | 2.149356121 | 0.763059265 | 2.816761712 | 0.00485105 | 0.07433867 |
| NBPF3 | 379.4127998 | -0.581102803 | 0.492473354 | -1.17996801 | 0.238012939 | 0.548684742 |
| NBPF8 | 564.6200641 | -0.258327422 | 0.431928117 | -0.598079661 | 0.54978678 | 0.793878561 |
| NBPF9 | 863.3772995 | -0.077845038 | 0.30568374 | -0.254658748 | 0.79898668 | 0.916056735 |
| NBR1 | 2689.528152 | -0.406297982 | 0.230011752 | -1.766422709 | 0.077324958 | 0.318807331 |
| NCALD | 139.1126125 | 0.901899491 | 0.514175172 | 1.75407048 | 0.079418431 | 0.322931692 |
| NCAM1 | 490.7508102 | 0.82177254 | 0.620352957 | 1.324685455 | 0.185275499 | 0.489859637 |
| NCAM2 | 15.7750562 | 0.268931544 | 0.965043978 | 0.278672838 | 0.780495909 | 0.909230088 |
| NCAN | 38.85474893 | 0.961211991 | 0.641832602 | 1.497605431 | 0.134235795 | 0.420619758 |
| NCAPD2 | 4897.164409 | 0.348656561 | 0.431966132 | 0.807138651 | 0.419586606 | 0.710202879 |
| NCAPD3 | 1210.507999 | 0.113432978 | 0.44649248 | 0.254053501 | 0.799454226 | 0.91629415 |
| NCAPG | 2563.270983 | 0.817180657 | 0.366078067 | 2.232257902 | 0.025597923 | 0.18249346 |
| NCAPG2 | 1806.200514 | 0.094468213 | 0.408245864 | 0.231400295 | 0.817003836 | 0.923735281 |
| NCAPH | 1030.874166 | 0.531568405 | 0.393938829 | 1.349367886 | 0.177218831 | 0.481566699 |
| NCAPH2 | 1241.049951 | 0.110759143 | 0.371341562 | 0.298267564 | 0.765498959 | 0.903026935 |
| NCBP1 | 1793.776319 | -0.037812253 | 0.256678907 | -0.147313441 | 0.882884624 | 0.953581235 |
| NCBP2 | 3584.315382 | 0.153534718 | 0.297110125 | 0.516760303 | 0.605323494 | 0.826077544 |
| NCBP2-AS2 | 511.109915 | 1.097364362 | 0.480818743 | 2.282282831 | 0.022472648 | 0.170466268 |
| NCBP3 | 2118.065672 | -0.515144948 | 0.342331162 | -1.504814652 | 0.132371736 | 0.418524946 |
| NCCRP1 | 115.1538435 | -0.184803111 | 0.790909594 | -0.233658958 | 0.815249753 | 0.922920552 |
| NCDN | 535.2847632 | -0.094827194 | 0.474194061 | -0.199975499 | 0.841499743 | 0.935153081 |
| NCEH1 | 155.4274615 | -0.258094288 | 0.371763919 | -0.694242435 | 0.487530185 | 0.756439734 |
| NCF1 | 17.79403595 | -0.884437765 | 0.845199772 | -1.046424519 | 0.295365077 | 0.60699582 |

| | | | | | | |
|---|---|---|---|---|---|---|
| NCF1C | 60.25825649 | -0.633697146 | 0.552663174 | -1.146624518 | 0.251536839 | 0.564383659 |
| NCK1 | 787.8626909 | 0.324890546 | 0.242501742 | 1.339745202 | 0.180328197 | 0.484774033 |
| NCK2 | 1084.729883 | -0.581680745 | 0.296914415 | -1.959085569 | 0.050102767 | 0.257487003 |
| NCKAP1 | 6497.966454 | -0.082149709 | 0.246762361 | -0.332910207 | 0.739202064 | 0.891550525 |
| NCKAP1L | 12.67181642 | 1.504327041 | 0.813324349 | 1.849602859 | 0.06437081 | 0.293255043 |
| NCKAP5 | 60.06012514 | -0.550275574 | 0.611316242 | -0.900148787 | 0.368041076 | 0.674179083 |
| NCKAP5L | 429.1925199 | -0.076578662 | 0.484158064 | -0.158168721 | 0.87432385 | 0.949740014 |
| NCKIPSD | 864.3435109 | -0.147448733 | 0.249639453 | -0.590646757 | 0.554757129 | 0.796360966 |
| NCL | 25257.83095 | -0.04215999 | 0.236347629 | -0.178381271 | 0.858423557 | 0.942476753 |
| NCLN | 1237.568165 | 0.365832191 | 0.383378412 | 0.954232632 | 0.3399659 | 0.646642762 |
| NCOA1 | 963.7534002 | -0.526095599 | 0.341428325 | -1.540866884 | 0.123349187 | 0.404753154 |
| NCOA2 | 1749.209761 | -0.86368545 | 0.254359871 | -3.395525593 | 0.00068497 | 0.021514856 |
| NCOA3 | 1759.888694 | -0.739892292 | 0.300113796 | -2.465372476 | 0.013687089 | 0.133106937 |
| NCOA4 | 3793.258387 | -0.413060276 | 0.242392861 | -1.704094235 | 0.088363483 | 0.341710805 |
| NCOA5 | 786.0356334 | -0.542399039 | 0.360207383 | -1.505796563 | 0.132119406 | 0.418458996 |
| NCOA6 | 1469.371236 | -0.188088021 | 0.392126539 | -0.479661544 | 0.631468077 | 0.838659038 |
| NCOA7 | 445.5091473 | -1.030608462 | 0.293853382 | -3.507220012 | 0.000452815 | 0.016498831 |
| NCOR1 | 3183.595211 | -0.182649948 | 0.25663646 | -0.711706934 | 0.476646273 | 0.749735677 |
| NCOR2 | 2187.523836 | -0.216952202 | 0.374658185 | -0.579067029 | 0.562543946 | 0.801706045 |
| NCR3LG1 | 18.23368824 | -0.337548981 | 1.127842558 | -0.299287324 | 0.764720826 | 0.902893924 |
| NCS1 | 986.6001603 | 0.616646824 | 0.319622958 | 1.929294526 | 0.053694309 | 0.267861241 |
| NCSTN | 2825.721392 | -0.708728094 | 0.266017739 | -2.664213655 | 0.007716853 | 0.09829755 |
| NDC1 | 2601.508483 | 0.262332947 | 0.236875469 | 1.107471989 | 0.268089916 | 0.581152025 |
| NDC80 | 1114.307323 | 0.965997215 | 0.292229849 | 3.305607615 | 0.000947707 | 0.026284577 |
| NDE1 | 452.575027 | 0.110769793 | 0.488852101 | 0.226591627 | 0.820741304 | 0.925132406 |
| NDEL1 | 676.6669893 | -0.043089955 | 0.34531116 | -0.124785875 | 0.90069307 | 0.961673205 |
| NDFIP1 | 1620.410625 | 0.12936966 | 0.351059136 | 0.368512442 | 0.712491169 | 0.877574643 |
| NDFIP2 | 781.4310781 | 0.005699671 | 0.332688865 | 0.017132135 | 0.986331202 | 0.994988408 |
| NDN | 873.8630635 | 0.510966421 | 0.319694686 | 1.59829501 | 0.109977337 | 0.381488036 |
| NDNF | 49.03536096 | 0.599963262 | 0.635170759 | 0.944570027 | 0.344878435 | 0.651656587 |
| NDOR1 | 310.2803708 | -0.564680455 | 0.752213225 | -0.750692006 | 0.452838035 | 0.733363031 |
| NDRG1 | 327.5107629 | -1.543858693 | 0.577653647 | -2.672637316 | 0.007525757 | 0.096389041 |
| NDRG2 | 255.3105281 | -0.507123033 | 0.36638993 | -1.384107453 | 0.166325555 | 0.468062511 |
| NDRG3 | 885.8263438 | 0.043356709 | 0.302746963 | 0.143211045 | 0.886123507 | 0.955049604 |
| NDRG4 | 303.6954009 | -0.026732407 | 0.337601984 | -0.079183205 | 0.936886903 | 0.976155095 |
| NDST1 | 2724.253015 | -1.160102423 | 0.386272986 | -3.003322686 | 0.002670491 | 0.050390573 |
| NDST2 | 468.0945354 | -0.453115912 | 0.415197404 | -1.091326457 | 0.27512926 | |

| | | | | | | |
|---|---|---|---|---|---|---|
| | | | | | | 0.588252711 |
| NDST3 | 31.40535099 | -0.989882337 | 0.867674515 | -1.140845236 | 0.253934332 | 0.566445924 |
| NDUFA1 | 1848.681564 | 1.247881278 | 0.418102462 | 2.984630306 | 0.002839212 | 0.052464911 |
| NDUFA10 | 1065.409189 | 0.006875749 | 0.237971623 | 0.028893146 | 0.976949812 | 0.992055261 |
| NDUFA11 | 1994.292114 | 0.745572011 | 0.43643356 | 1.70832878 | 0.087575354 | 0.340144828 |
| NDUFA12 | 1334.457685 | 0.827198899 | 0.41526927 | 1.991957889 | 0.046375679 | 0.247615641 |
| NDUFA13 | 3998.618906 | 1.1499463 | 0.427669423 | 2.688867237 | 0.007169493 | 0.093523967 |
| NDUFA2 | 1472.95989 | 1.266870164 | 0.458989592 | 2.760128304 | 0.005777866 | 0.081921507 |
| NDUFA3 | 156.4320982 | 0.890714825 | 0.529885922 | 1.68095582 | 0.092771496 | 0.349440669 |
| NDUFA4 | 7193.428912 | 1.015471981 | 0.351403781 | 2.889758269 | 0.003855382 | 0.064133766 |
| NDUFA5 | 1782.359565 | 0.793245988 | 0.252549677 | 3.140950312 | 0.001684006 | 0.038017939 |
| NDUFA6 | 42.57336152 | 1.330762614 | 0.720334364 | 1.847423475 | 0.06468579 | 0.29355313 |
| NDUFA7 | 1153.662925 | 1.164121241 | 0.465239351 | 2.502198576 | 0.012342468 | 0.126347892 |
| NDUFA8 | 1005.847568 | 0.993853848 | 0.454693849 | 2.185764881 | 0.028832811 | 0.194237381 |
| NDUFA9 | 1130.306207 | 0.330119515 | 0.351789019 | 0.938401989 | 0.348037864 | 0.654528628 |
| NDUFAB1 | 1665.073577 | 1.151802684 | 0.450852445 | 2.554722054 | 0.010627262 | 0.117085107 |
| NDUFAF1 | 567.1911172 | 0.438744909 | 0.447860645 | 0.979646044 | 0.327260868 | 0.637339491 |
| NDUFAF2 | 424.2593245 | 1.473186069 | 0.396084742 | 3.719370916 | 0.00019972 | 0.00940977 |
| NDUFAF3 | 323.2736836 | 0.860050169 | 0.489717868 | 1.75621562 | 0.079051599 | 0.322602824 |
| NDUFAF4 | 675.7914095 | 1.33405496 | 0.346758464 | 3.847216715 | 0.000119467 | 0.006917703 |
| NDUFAF5 | 334.6896527 | 0.655247482 | 0.280063795 | 2.339636522 | 0.019302515 | 0.157609505 |
| NDUFAF6 | 343.9373604 | 0.457585749 | 0.368297144 | 1.24243632 | 0.214075622 | 0.520920862 |
| NDUFAF7 | 657.5559773 | -0.273687321 | 0.326537782 | -0.838149016 | 0.401947017 | 0.699116854 |
| NDUFAF8 | 291.4626343 | 1.329871765 | 0.441502986 | 3.012146706 | 0.002594072 | 0.049502724 |
| NDUFB1 | 1174.030143 | 1.227917193 | 0.447140143 | 2.746157358 | 0.006029783 | 0.083980132 |
| NDUFB10 | 1496.04845 | 0.812782352 | 0.433326182 | 1.875682535 | 0.060698885 | 0.284105089 |
| NDUFB11 | 2057.768481 | 0.866522261 | 0.465297312 | 1.862298015 | 0.062561095 | 0.288484612 |
| NDUFB2 | 1254.053911 | 1.171046744 | 0.387410995 | 3.022750407 | 0.002504887 | 0.048313917 |
| NDUFB3 | 1145.493786 | 1.247408168 | 0.351104199 | 3.552814728 | 0.000381133 | 0.014675547 |
| NDUFB4 | 3645.733482 | 0.869889366 | 0.455709732 | 1.908867214 | 0.056279223 | 0.27489121 |
| NDUFB5 | 2361.744464 | 0.960072255 | 0.446569903 | 2.149881235 | 0.03156461 | 0.202592072 |
| NDUFB6 | 1140.083908 | 1.390947088 | 0.37930526 | 3.667091485 | 0.000245325 | 0.010725724 |
| NDUFB7 | 804.5546074 | 0.917828212 | 0.650620866 | 1.410695937 | 0.15833429 | |

0.45780246
NDUFB8 2613.389828 0.846136271 0.439569927 1.92491847 0.054239559
0.269175122
NDUFB9 3028.923393 1.03530017 0.481038847 2.152217388 0.031380237
0.202023044
NDUFC1 866.6899322 0.983745814 0.451120095 2.180673894 0.029207545
0.195546191
NDUFC2 2356.004871 1.222586079 0.473245998 2.583405006 0.00978304
0.112404636
NDUFC2-KCTD14 35.37185032 1.100897971 0.650831031 1.691526554 0.090736281
0.345245839
NDUFS1 1152.848079 0.173680959 0.341902336 0.507984125 0.611464476
0.829058038
NDUFS2 2766.981453 0.347061661 0.264181443 1.313724602 0.188938899
0.494119637
NDUFS3 770.1673159 0.837584617 0.379743357 2.205659695 0.027407836
0.189349252
NDUFS4 1816.433531 1.220281394 0.351257121 3.474040301 0.000512684
0.017880297
NDUFS5 4863.257252 1.338932988 0.450882742 2.969581361 0.002982058
0.053986643
NDUFS6 2133.352097 1.314456682 0.514393467 2.555352596 0.010608028
0.117043195
NDUFS7 550.5827087 0.698611273 0.526322021 1.327345702 0.184394352
0.488866012
NDUFS8 1109.735089 0.943042658 0.38290365 2.46287195 0.013782913
0.13361059
NDUFV1 2419.261046 0.221332755 0.304986647 0.725712936 0.46801477
0.744224533
NDUFV2 2962.706315 1.024477475 0.459867994 2.227764245 0.02589624
0.183710607
NDUFV2P1 88.16178526 0.443765314 0.400917664 1.106873941 0.268348434
0.581480913
NDUFV3 1009.069463 0.706750138 0.324315543 2.179205265 0.029316423
0.195861365
NEB 36.53728168 -0.683619386 0.577971742 -1.182790327 0.236892255
0.547682475
NEBL 2303.326735 -0.161689691 0.361084041 -0.447789635 0.654305024
0.849068645
NECAB1 14.72125469 -2.206070066 0.954362543 -2.311563968
0.020801724 0.163514486
NECAB2 303.2727259 1.538397109 0.530471526 2.900055954 0.003730961
0.063017919
NECAB3 518.0216564 0.226907653 0.364465154 0.622577086 0.533562473
0.783156734
NECAP1 570.2861919 0.141893035 0.279452821 0.507753097 0.611626507
0.829058038
NECAP2 1520.413237 -0.010957155 0.294643402 -0.037187852
0.970335225 0.98957539
NECTIN1 2932.981847 -0.515430799 0.374895233 -1.37486624 0.169172917
0.471613806
NECTIN2 3889.918495 -0.069281606 0.30346843 -0.228299219
0.819413632 0.924316427
NECTIN3 1492.642527 -0.09958977 0.515886941 -0.193045729 0.846923155
0.938145276
NECTIN4 349.3721418 -0.773166447 0.75155614 -1.028754081
0.303595249 0.614002998
NEDD1 1081.263964 0.33640601 0.282769374 1.189683327 0.234170881 0.544210545
NEDD4 683.772832 -1.256491679 0.367224097 -3.421593758 0.000622553
0.020311279
NEDD4L 1686.086353 -0.496790387 0.25300605 -1.963551416
0.049582136 0.256639574
NEDD8 2020.970928 0.807593796 0.398373958 2.027225372 0.04263937 0.237387844
NEDD9 1001.470001 -2.0152439 0.53109682 -3.794494384 0.000147945

| | | | | | | |
|---|---|---|---|---|---|---|
| | | | | | | 0.007793292 |
| NEFH | 33.66849918 | -2.247168568 | 0.714355937 | -3.145726733 | 0.001656747 | 0.037682927 |
| NEFL | 392.309207 | 0.668863029 | 0.30863715 | 2.167150097 | 0.030223411 | 0.198239795 |
| NEFM | 375.703904 | 0.433624651 | 0.264256316 | 1.640924456 | 0.100813098 | 0.363642563 |
| NEGR1 | 25.39350634 | -0.046502658 | 0.754490582 | -0.06163451 | 0.950853894 | 0.981826318 |
| NEIL1 | 147.2954206 | -0.780782766 | 0.377242216 | -2.069712067 | 0.038479316 | 0.225742345 |
| NEIL2 | 314.599485 | 1.034514948 | 0.316504512 | 3.268563032 | 0.001080951 | 0.028433348 |
| NEIL3 | 277.8449639 | 1.005382155 | 0.340256058 | 2.954781054 | 0.00312891 | 0.055756493 |
| NEK1 | 685.363586 | -0.108739908 | 0.253786263 | -0.428470427 | 0.66830866 | 0.856934809 |
| NEK10 | 17.55058421 | -0.090023118 | 0.743511997 | -0.121078232 | 0.903629071 | 0.963499647 |
| NEK11 | 107.0857947 | -0.662648895 | 0.383114951 | -1.729634652 | 0.083695571 | 0.330936505 |
| NEK2 | 1052.563731 | 1.129236182 | 0.327288366 | 3.450279014 | 0.000560007 | 0.018754686 |
| NEK3 | 798.9477364 | -0.180104425 | 0.328460119 | -0.548329659 | 0.583465565 | 0.814179185 |
| NEK4 | 945.1231812 | 0.1045147 | 0.261384544 | 0.399850346 | 0.689266746 | 0.867579108 |
| NEK5 | 33.89015089 | -1.887446215 | 0.671369444 | -2.811337681 | 0.004933598 | 0.075072829 |
| NEK6 | 269.6551134 | 0.557100171 | 0.387698419 | 1.436942075 | 0.150734453 | 0.445603028 |
| NEK7 | 544.3998012 | -0.027625008 | 0.332918105 | -0.08297839 | 0.933868723 | 0.974933951 |
| NEK9 | 2911.936848 | -0.191102442 | 0.309817981 | -0.616821663 | 0.537352362 | 0.785675856 |
| NELFA | 965.8209578 | 0.17774569 | 0.397928259 | 0.446677728 | 0.655107765 | 0.849331122 |
| NELFB | 1499.342759 | 0.185682891 | 0.386331103 | 0.480631483 | 0.630778435 | 0.838256801 |
| NELFCD | 2101.691855 | -0.149642391 | 0.320384706 | -0.467070956 | 0.640449113 | 0.843644482 |
| NELFE | 1657.363539 | 0.61617526 | 0.446495078 | 1.380026994 | 0.167578334 | 0.469844927 |
| NELL2 | 890.2749343 | 0.371113562 | 0.40956166 | 0.906123786 | 0.364870337 | 0.671614642 |
| NEMF | 2384.809027 | 0.236865301 | 0.332613617 | 0.712133504 | 0.476382109 | 0.74962941 |
| NEMP1 | 1493.967215 | 0.062911593 | 0.390123495 | 0.161260714 | 0.871888064 | 0.94888191 |
| NEMP2 | 136.0010139 | -0.670097361 | 0.456532572 | -1.467797484 | 0.142159241 | 0.432308365 |
| NENF | 471.2882312 | 0.628323705 | 0.547808102 | 1.146977752 | 0.251390816 | 0.564262475 |
| NEO1 | 2808.330316 | -0.683134876 | 0.391329113 | -1.745678645 | 0.080866808 | 0.325332894 |
| NEPRO | 1882.907449 | -0.070533822 | 0.23961769 | -0.294359826 | 0.768482962 | 0.904332533 |
| NES | 6959.049091 | 0.521076405 | 0.390057074 | 1.335897845 | 0.181582666 | 0.485768051 |
| NET1 | 1970.106253 | -0.181567748 | 0.30979806 | -0.586084199 | 0.557818934 | 0.798666577 |
| NETO1 | 12.34187498 | 0.00297737 | 0.877219856 | 0.003394098 | 0.997291907 | 0.998674292 |
| NETO2 | 469.5526709 | -0.133729292 | 0.295930123 | -0.451894828 | 0.651344751 | 0.847876685 |
| NEU1 | 197.1278782 | -1.164747315 | 0.706424475 | -1.648792413 | 0.09919017 | 0.361452832 |
| NEU3 | 560.376648 | -0.000629722 | 0.403110958 | -0.001562155 | 0.998753581 | 0.999214625 |
| NEURL1 | 97.05484975 | -0.983565002 | 0.513723811 | -1.914579354 | 0.05554617 | 0.272715517 |
| NEURL1B | 211.9433592 | 1.477089088 | 0.60918794 | 2.424685374 | 0.015321659 | 0.140195951 |
| NEURL4 | 1064.077101 | -0.273199944 | 0.457001475 | -0.597809766 | 0.549966874 | 0.793878561 |
| NEUROG1 | 40.13601639 | -1.295000661 | 1.010383201 | -1.28169259 | 0.199950506 | 0.50598138 |
| NEXMIF | 10.67772522 | 1.824533988 | 0.986581908 | 1.849348719 | 0.064407475 | 0.293255043 |
| NEXN | 148.2530988 | -0.564317703 | 0.390605497 | -1.44472545 | 0.148535018 | |

0.442803058
NF1 1996.63651 -0.604315225 0.386170391 -1.564892698 0.117608071
0.395641036
NF2 581.8316107 -0.20207691 0.286748731 -0.704717712 0.480985927
0.75260258
NFAM1 12.33181652 -1.387800518 1.133564989 -1.224279624 0.220846761
0.529634241
NFASC 163.673719 0.060571157 0.628208353 0.096418898 0.923187884 0.971120333
NFAT5 847.0414485 -0.673338037 0.394514708 -1.70675015 0.087868502
0.340847107
NFATC1 124.1060913 0.233176401 0.448072125 0.520399257 0.602785328
0.825021776
NFATC2 20.79369327 -1.828569194 0.840197889 -2.176355377
0.029528698 0.196251879
NFATC2IP 1911.82581 -0.41399245 0.276253487 -1.498596284 0.133978397
0.420619758
NFATC3 1168.073785 0.42206668 0.28948088 1.458012287 0.144837156
0.436063603
NFATC4 1160.894125 0.538701114 0.358695855 1.50183256 0.133140357
0.419488075
NFE2 201.8479364 -0.661647522 0.575851917 -1.148989007 0.250560514
0.563399074
NFE2L1 3553.106815 -0.009883748 0.457882905 -0.021585756
0.982778396 0.993518428
NFE2L2 1244.352781 0.117332081 0.261820158 0.448139983 0.654052174
0.849043068
NFE2L3 569.2540737 -0.561988411 0.425090009 -1.322045682
0.18615294 0.490817669
NFE4 15.31439627 -4.05858523 1.612867228 -2.516379004 0.01185676
0.123589668
NFIA 22.92846631 1.048770184 0.858224443 1.222023204 0.221698852 0.53065007
NFIB 21.31660338 0.664730945 0.955781379 0.695484302 0.486751846 0.756340124
NFIC 135.5749018 0.427197068 0.352547655 1.211742758 0.225610866 0.53459368
NFIL3 540.3810075 -0.176620734 0.290180598 -0.60865797 0.542751171
0.789309626
NFKB1 227.9249597 -0.762101752 0.374719638 -2.033791864 0.041972585
0.235820549
NFKB2 682.9385709 -0.754380252 0.288374134 -2.615977524 0.008897241
0.10668882
NFKBIA 721.9673682 -0.10756224 0.523985103 -0.205277286 0.837355489
0.933338258
NFKBIB 175.6777958 0.336127781 0.41008951 0.819644915 0.412418561
0.70515068
NFKBIE 162.3005082 0.66975386 0.326461823 2.051553394 0.040213086
0.231327132
NFKBIZ 307.4788807 -0.530410235 0.461309841 -1.149791719
0.250229667 0.562905438
NFRKB 1312.589606 -0.335154636 0.343706756 -0.975117973 0.329501744
0.638506956
NFS1 584.764419 0.264470723 0.254740559 1.038196369 0.299178643 0.610246278
NFU1 746.8456127 0.654584684 0.369174255 1.773104908 0.076211325 0.316642999
NFX1 1073.940379 -0.303745708 0.272554865 -1.114438766 0.265091
0.578994467
NFXL1 709.3900922 -0.264686804 0.271353165 -0.9754329 0.329345571
0.638448972
NFYA 1404.813934 -0.34509992 0.389494786 -0.886019358 0.375607096
0.67966045
NFYB 1527.685136 0.755323502 0.243820501 3.097867071 0.001949188 0.041386806
NFYC 1249.043775 -0.52777378 0.271546642 -1.943584263 0.051945605
0.262776517
NGDN 1173.408705 0.557365839 0.340472531 1.637036145 0.101622918 0.365596699
NGEF 83.83120818 0.201007514 0.615851522 0.326389571 0.744129627 0.892691858
NGF 24.40528718 -1.347146436 0.988098036 -1.363373255 0.172764906
0.476461804

| | | | | | | |
|---|---|---|---|---|---|---|
| NGFR | 481.5074535 | 1.333164434 | 0.378190286 | 3.525115483 | 0.000423298 | 0.015778519 |
| NGLY1 | 918.788168 | 0.035024711 | 0.260994844 | 0.134196947 | 0.89324684 | 0.95780189 |
| NGRN | 3312.096175 | 0.382308632 | 0.28402852 | 1.346021985 | 0.178295427 | 0.4825693 |
| NHEJ1 | 575.9051691 | -0.006361756 | 0.27500374 | -0.023133342 | 0.98154391 | 0.993395774 |
| NHLH1 | 17.32598555 | -0.360783449 | 0.83973912 | -0.429637539 | 0.667459325 | 0.856540807 |
| NHLH2 | 78.27997109 | 1.803923276 | 0.485693386 | 3.714119497 | 0.000203912 | 0.009548011 |
| NHLRC2 | 880.1428091 | -0.072247741 | 0.329277669 | -0.219412816 | 0.826328488 | 0.927784894 |
| NHLRC3 | 675.9747079 | -0.161443913 | 0.264756333 | -0.609783007 | 0.542005559 | 0.788754565 |
| NHP2 | 2497.356255 | 0.894732628 | 0.436260454 | 2.0509139 | 0.040275332 | 0.231327132 |
| NHS | 308.0287849 | -0.051477991 | 0.365820878 | -0.140719118 | 0.888091842 | 0.955943827 |
| NHSL1 | 520.940172 | -0.523413616 | 0.35246031 | -1.485028528 | 0.137536267 | 0.425742238 |
| NHSL2 | 367.3381694 | -0.093416407 | 0.588836337 | -0.158645791 | 0.87394795 | 0.949740014 |
| NICN1 | 423.4125658 | -0.218353762 | 0.300775303 | -0.725969718 | 0.467857335 | 0.744166872 |
| NID1 | 472.4079252 | 0.758335862 | 0.461357811 | 1.643704394 | 0.100237278 | 0.36243601 |
| NID2 | 606.4398388 | -0.491328953 | 0.510855892 | -0.961776033 | 0.33616212 | 0.64425371 |
| NIF3L1 | 1072.133641 | 0.444759682 | 0.245125508 | 1.814416159 | 0.06961368 | 0.303425952 |
| NIFK | 1449.058072 | 0.63001192 | 0.228616839 | 2.755754658 | 0.00585569 | 0.082561966 |
| NIN | 904.5363127 | 0.12443471 | 0.451290956 | 0.275730563 | 0.782755019 | 0.909909835 |
| NINJ1 | 418.1264873 | 0.485103353 | 0.345423898 | 1.404371139 | 0.160208361 | 0.460500386 |
| NINL | 989.9782278 | -0.224004 | 0.346876346 | -0.64577479 | 0.518425217 | 0.774169079 |
| NIP7 | 998.5533479 | 0.120933373 | 0.32217073 | 0.375370453 | 0.707384976 | 0.87520286 |
| NIPA1 | 515.4338726 | -0.345971724 | 0.338538251 | -1.021957557 | 0.306800989 | 0.616910609 |
| NIPA2 | 1519.561539 | -0.090522749 | 0.273325466 | -0.331190323 | 0.740500729 | 0.891917214 |
| NIPAL1 | 163.8526166 | -1.911336263 | 0.509525643 | -3.751207204 | 0.000175985 | 0.008728094 |
| NIPAL2 | 36.88743075 | -1.437087226 | 0.607243911 | -2.3665733 | 0.017953619 | 0.151235063 |
| NIPAL3 | 287.7221987 | -0.242677789 | 0.356648361 | -0.68043994 | 0.496225939 | 0.761056737 |
| NIPBL | 2925.243296 | -0.380926345 | 0.300601834 | -1.267212311 | 0.205079385 | 0.51158559 |
| NIPSNAP1 | 7045.083025 | 0.196269228 | 0.248874692 | 0.788626701 | 0.43033022 | 0.717027981 |
| NIPSNAP2 | 1623.608432 | 0.537432813 | 0.27440016 | 1.958573247 | 0.050162785 | 0.257598027 |
| NIPSNAP3A | 625.2526871 | 0.530896807 | 0.312543078 | 1.698635627 | 0.089387859 | 0.342726775 |
| NIPSNAP3B | 58.20714727 | 0.34545994 | 0.542913964 | 0.636306971 | 0.524576362 | 0.777662198 |
| NISCH | 3101.726657 | -0.391090322 | 0.369107512 | -1.059556659 | 0.28934634 | 0.60229533 |
| NIT1 | 1010.388692 | -0.003981349 | 0.260136547 | -0.015304843 | 0.987788979 | 0.995541774 |
| NIT2 | 869.232525 | -0.03267239 | 0.361920457 | -0.090275058 | 0.92806864 | 0.972633911 |
| NKAIN1 | 483.2414171 | 0.470110365 | 0.470119577 | 0.999980405 | 0.317319991 | 0.627402787 |
| NKAIN2 | 15.37788 | 0.892204933 | 0.944052461 | 0.945079823 | 0.344618125 | 0.651368096 |
| NKAIN4 | 126.672669 | -0.257622045 | 0.495217197 | -0.520220312 | 0.60291003 | 0.825021776 |
| NKAP | 803.6954374 | 0.805124189 | 0.427265052 | 1.884367059 | 0.059515342 | 0.282669434 |

| | | | | | | |
|---|---|---|---|---|---|---|
| NKAPD1 | 1381.889381 | -0.092578672 | 0.260436565 | -0.355474938 | 0.722233817 | 0.882846607 |
| NKAPP1 | 28.06116764 | -0.285928291 | 0.669546093 | -0.427047956 | 0.669344399 | 0.857712777 |
| NKD1 | 730.9675413 | 0.954054512 | 0.641402433 | 1.487450721 | 0.136895809 | 0.424728769 |
| NKD2 | 58.63171065 | -0.781781457 | 0.513637298 | -1.522049627 | 0.127996646 | 0.412255816 |
| NKIRAS1 | 226.9753641 | 0.525598429 | 0.342666615 | 1.533847788 | 0.125067103 | 0.406867909 |
| NKIRAS2 | 1243.627127 | -0.193572124 | 0.254447303 | -0.76075526 | 0.446803262 | 0.728709125 |
| NKPD1 | 18.87421426 | 0.174012659 | 0.720346125 | 0.241568119 | 0.809114828 | 0.919688398 |
| NKRF | 682.123479 | 0.128730786 | 0.253515512 | 0.507782682 | 0.611605756 | 0.829058038 |
| NKTR | 3235.253096 | -0.436799089 | 0.378416679 | -1.154280752 | 0.248385085 | 0.561085486 |
| NKX1-2 | 120.5455938 | -0.55239168 | 0.601258326 | -0.918726039 | 0.358238887 | 0.665871373 |
| NKX3-1 | 18.25738402 | -0.64911866 | 1.083529748 | -0.59907784 | 0.549120979 | 0.79348329 |
| NKX6-1 | 16.57511825 | 1.776498922 | 0.880928075 | 2.016621985 | 0.043734977 | 0.240923506 |
| NLE1 | 1228.47822 | 0.535359503 | 0.246332736 | 2.173318544 | 0.029756351 | 0.197218697 |
| NLGN1 | 118.2792019 | 0.401657078 | 0.529461474 | 0.758614361 | 0.448083279 | 0.729463615 |
| NLGN2 | 1581.461538 | 0.249317694 | 0.356897623 | 0.698569219 | 0.484821286 | 0.755350561 |
| NLGN3 | 284.0603808 | 0.952595356 | 0.618461284 | 1.540266757 | 0.123495343 | 0.40487218 |
| NLGN4X | 638.9575197 | -0.424226146 | 0.409832131 | -1.035121734 | 0.300612064 | 0.611214323 |
| NLGN4Y | 700.089992 | -0.027121014 | 0.375112979 | -0.07230092 | 0.942362432 | 0.978574897 |
| NLK | 1008.99959 | -0.076741134 | 0.235624484 | -0.325692531 | 0.744656996 | 0.892846858 |
| NLN | 2554.968001 | -0.178519473 | 0.254054453 | -0.702681932 | 0.48225399 | 0.753310328 |
| NLRP1 | 64.36641869 | 1.318047222 | 0.712143817 | 1.850816073 | 0.064196018 | 0.292572699 |
| NLRP12 | 18.17478071 | -1.31818941 | 0.745675433 | -1.767779051 | 0.077097848 | 0.31844659 |
| NLRX1 | 526.8774049 | -0.020319222 | 0.303696112 | -0.066906426 | 0.946656197 | 0.980533263 |
| NMB | 83.87901501 | 1.520104394 | 0.517636668 | 2.936624256 | 0.003318059 | 0.058194529 |
| NMBR | 15.49676879 | -2.065411199 | 1.026632719 | -2.011830679 | 0.044237789 | 0.241762065 |
| NMD3 | 1775.408135 | 0.26514534 | 0.240872066 | 1.100772472 | 0.270995695 | 0.583394591 |
| NME1 | 2284.810886 | 1.037176404 | 0.252477514 | 4.107995159 | 3.99E-05 | 0.003121069 |
| NME1-NME2 | 683.6479137 | 0.386885207 | 0.524883397 | 0.737087911 | 0.46106889 | 0.73885513 |
| NME2 | 8642.508905 | 0.844467843 | 0.434429615 | 1.943854226 | 0.051913032 | 0.262699338 |
| NME3 | 239.3945145 | 1.006735053 | 0.704089937 | 1.429838717 | 0.152763314 | 0.44888093 |
| NME4 | 3075.3628 | 0.821079604 | 0.264402744 | 3.105412569 | 0.001900138 | 0.04071609 |
| NME5 | 137.7820841 | 0.878492128 | 0.331284354 | 2.651776693 | 0.008006948 | 0.100518204 |
| NME6 | 592.2287925 | -0.393187286 | 0.299578632 | -1.312467726 | 0.189362369 | 0.494879496 |
| NME7 | 620.8099845 | 0.437113536 | 0.307690241 | 1.420628535 | 0.155424778 | 0.453320705 |
| NME8 | 51.60917127 | -0.134463209 | 0.53988237 | -0.249060197 | 0.803314218 | 0.917969611 |
| NMI | 250.5448546 | -0.232573086 | 0.598580518 | -0.388541022 | 0.697615703 | 0.87033849 |
| NMNAT1 | 232.0561129 | -0.29396001 | 0.341853883 | -0.859899579 | 0.389844401 | 0.689717715 |
| NMNAT2 | 2086.385857 | -1.526091752 | 0.289642554 | -5.268879629 | 1.37E-07 | 4.73E-05 |
| NMNAT3 | 492.6189269 | 0.037950046 | 0.375506108 | 0.101063724 | 0.919499874 | 0.969677217 |
| NMRAL1 | 348.781598 | 0.107113021 | 0.437830896 | 0.24464473 | 0.806731527 | 0.919137032 |

| | | | | | | |
|---|---|---|---|---|---|---|
| NMRK1 | 306.9353963 | -0.359739235 | 0.480085769 | -0.749322846 | 0.453662641 | 0.733509104 |
| NMRK2 | 83.04538733 | 0.259372374 | 0.489865365 | 0.529476859 | 0.596474694 | 0.821812117 |
| NMT1 | 3502.079354 | 0.003385712 | 0.225871396 | 0.014989556 | 0.988040512 | 0.995653445 |
| NMT2 | 504.7935347 | 0.257636081 | 0.303943233 | 0.847645392 | 0.396635487 | 0.695326667 |
| NMU | 412.0609926 | 1.581750314 | 0.484932444 | 3.261795189 | 0.001107091 | 0.028759722 |
| NNT | 908.559536 | -0.572005814 | 0.359165658 | -1.592596065 | 0.111250837 | 0.383708973 |
| NOA1 | 1667.706866 | -0.111123825 | 0.335414414 | -0.331303069 | 0.740415574 | 0.891917214 |
| NOB1 | 2707.076661 | -0.002678449 | 0.265237046 | -0.01009832 | 0.991942843 | 0.99688215 |
| NOC2L | 2558.491452 | -0.142247761 | 0.342648321 | -0.415142152 | 0.678037844 | 0.860940084 |
| NOC3L | 869.7087169 | 0.626317677 | 0.359495596 | 1.74221238 | 0.081471289 | 0.326379963 |
| NOC4L | 304.6115212 | -0.057607496 | 0.446543633 | -0.129007542 | 0.897351681 | 0.959456082 |
| NOCT | 256.8452961 | -1.004617782 | 0.378084105 | -2.657127787 | 0.007880956 | 0.099718089 |
| NOD1 | 562.4687452 | -0.593448593 | 0.307976754 | -1.92692658 | 0.053988781 | 0.268545505 |
| NOD2 | 10.51620149 | -0.343967192 | 1.042191185 | -0.330042316 | 0.741367989 | 0.892005215 |
| NOG | 103.445784 | 2.053735808 | 0.81302724 | 2.526035668 | 0.011535777 | 0.121534216 |
| NOL10 | 979.1171073 | 0.067517873 | 0.304008425 | 0.222092111 | 0.824242179 | 0.926745579 |
| NOL11 | 2181.838319 | 0.459661677 | 0.240266495 | 1.913132654 | 0.055731072 | 0.273181292 |
| NOL12 | 369.021575 | 0.322100781 | 0.326995497 | 0.985031244 | 0.324608724 | 0.634653009 |
| NOL4L | 621.9253528 | 0.106461451 | 0.397778108 | 0.267640297 | 0.788976209 | 0.912533261 |
| NOL6 | 1082.807115 | -0.134636919 | 0.359455044 | -0.374558436 | 0.707988889 | 0.875585221 |
| NOL7 | 1352.473037 | 0.756326138 | 0.402045714 | 1.881194381 | 0.059945482 | 0.283136022 |
| NOL8 | 1138.158373 | -0.016057485 | 0.250749116 | -0.064038052 | 0.948939928 | 0.98134156 |
| NOL9 | 646.8275583 | -0.239478664 | 0.297372882 | -0.8053144 | 0.420638274 | 0.711245864 |
| NOLC1 | 4145.368894 | -0.039631284 | 0.251238373 | -0.157743753 | 0.874658722 | 0.949740014 |
| NOM1 | 751.9086121 | 0.028968233 | 0.261422674 | 0.110809948 | 0.911767056 | 0.96696386 |
| NOMO1 | 1888.791279 | -0.421732353 | 0.401195421 | -1.051189349 | 0.293171634 | 0.60622718 |
| NOMO2 | 2904.84619 | -0.187782255 | 0.253053927 | -0.742064181 | 0.458048451 | 0.736518606 |
| NOMO3 | 52.47504518 | 0.037402859 | 0.682306687 | 0.054818251 | 0.95628326 | 0.984447503 |
| NONO | 27350.92912 | -0.150032676 | 0.234482198 | -0.639846767 | 0.522272225 | 0.777260341 |
| NOP10 | 2067.795848 | 0.951037581 | 0.429074437 | 2.216486228 | 0.02665822 | 0.186715644 |
| NOP14 | 1282.847888 | -0.15229054 | 0.258022655 | -0.590221584 | 0.555042104 | 0.796360966 |
| NOP16 | 586.1525149 | 0.711379533 | 0.306459314 | 2.321285407 | 0.020271445 | 0.161005801 |
| NOP2 | 1098.823648 | -0.223014267 | 0.356359172 | -0.625813182 | 0.531437485 | 0.781850086 |
| NOP53 | 4643.042661 | 0.0190604 | 0.421137967 | 0.045259277 | 0.963900647 | 0.988196831 |
| NOP56 | 5203.143018 | 0.293297208 | 0.259489328 | 1.130286205 | 0.258355645 | 0.571274376 |
| NOP58 | 3298.823493 | 0.678626481 | 0.26383593 | 2.572153384 | 0.01010681 | 0.114259456 |
| NOP9 | 571.1370015 | -0.484696549 | 0.381151748 | -1.271662929 | 0.203492907 | 0.509858117 |
| NOS1 | 34.04587519 | -1.752552737 | 0.648770592 | -2.701344296 | 0.006905981 | 0.091263619 |
| NOS1AP | 58.61143893 | -0.331892651 | 0.627284043 | -0.529094682 | 0.59673977 | 0.821812117 |
| NOS2 | 130.0468771 | -4.315753832 | 1.700703895 | -2.537628005 | 0.011160654 | 0.120212934 |
| NOS3 | 151.2350506 | 0.123581746 | 0.364875215 | 0.338695919 | 0.734838816 | 0.889483063 |
| NOSIP | 848.0373267 | 0.339928422 | 0.380380286 | 0.893654153 | 0.371506963 | 0.67682226 |

NOTCH1 912.9430981 0.197100977 0.482883133 0.408175319 0.683144967
0.863415274
NOTCH2 2690.752402 -0.058246853 0.418235462 -0.139268089
0.889238306 0.956307814
NOTCH2NLA 109.9162399 0.059164658 0.516205701 0.114614498 0.908750687
0.965857971
NOTCH3 3854.97068 -0.49204045 0.288856259 -1.703409341 0.088491491
0.341952218
NOTUM 102.2444959 0.462488745 0.591964818 0.781277418 0.434639349 0.719554351
NOVA1 438.8886076 0.857768998 0.335616644 2.555799936 0.010594401 0.116977919
NOVA2 154.6822068 1.105615381 0.621476391 1.779014292 0.075237419 0.314600339
NOX4 330.0605168 -1.376067178 0.456283466 -3.01581644 0.002562884
0.049092812
NOXA1 37.33838008 -0.79800247 0.792242399 -1.007270592 0.313804758
0.623434514
NOXRED1 19.20477579 -0.49930335 0.876041213 -0.569954178 0.568708777
0.805450586
NPAS1 130.7234209 -0.819332015 0.603102218 -1.358529269 0.174295802
0.478542911
NPAS4 20.10548002 -4.625160706 1.573184582 -2.93999875 0.003282136
0.057789532
NPAT 551.995637 -0.468897814 0.339662842 -1.380480157 0.167438855
0.469714288
NPB 10.33068534 0.645383564 0.935972931 0.689532296 0.490488357 0.758593654
NPC1 1306.27545 -0.631175759 0.276542247 -2.282384578 0.022466645
0.170466268
NPC2 4522.852901 0.306422416 0.440706096 0.695298791 0.486868072 0.756403023
NPDC1 193.4813264 0.37208568 0.535554783 0.694766794 0.487201461 0.756439734
NPEPL1 646.6837564 -0.320213884 0.404751263 -0.791137454
0.428863785 0.715685016
NPEPPS 4466.567343 -0.321110131 0.266129986 -1.206591316
0.227589573 0.536368893
NPFF 18.57080211 0.270971203 1.244068509 0.217810515 0.827576751 0.928323608
NPFFR2 11.35939498 -2.52427315 1.1218172 -2.25016442 0.02443851
0.178505842
NPHP1 214.5572243 0.160292427 0.290990081 0.550851859 0.581735231 0.813002261
NPHP3 664.5421723 -0.487891422 0.338001923 -1.443457533 0.148891627
0.443517548
NPHP3-ACAD11 44.5279797 -0.509247769 2.412457434 -0.211090882
0.832816355 0.931484585
NPHP4 460.3436777 0.080251008 0.554224246 0.144798804 0.884869726 0.95440284
NPHS1 10.89306371 -0.272726733 1.037443891 -0.262883357 0.792640485
0.913834546
NPIPA1 1037.324063 -0.114455053 0.367735288 -0.31124305 0.755615858
0.897896779
NPIPA2 85.47208075 0.102629082 0.673523692 0.152376351 0.878890109
0.951723186
NPIPA5 33.80667134 0.217605834 1.071807284 0.203027016 0.839113917
0.933915137
NPIPA7 127.8832841 0.14336136 0.52630305 0.272393177 0.785319719
0.910931322
NPIPA8 14.77870445 0.888876967 1.25754693 0.706834032 0.479669623
0.751840877
NPIPA9 568.4583159 0.426104788 0.263383911 1.617808721 0.105703826
0.372686009
NPIPB11 110.7433486 -1.048723311 0.45057745 -2.327509537
0.019938162 0.159698109
NPIPB12 752.4619834 -0.429682358 0.388777307 -1.105214604
0.26906661 0.582096023
NPIPB13 55.35950287 0.603636438 0.475740536 1.268835409 0.204499773
0.510835602
NPIPB14P 148.2687717 -1.455571415 0.476250859 -3.056312417
0.002240776 0.044848046
NPIPB15 111.0106751 0.389764491 0.568671894 0.685394329 0.493095122

| | | | | | | |
|---|---|---|---|---|---|---|
| | | | | | | 0.759389046 |
| NPIPB2 | 23.4626266 | -0.509486652 | 0.655057473 | -0.777773971 | 0.436702275 | 0.720784523 |
| NPIPB3 | 940.6691034 | -0.375088239 | 0.422999751 | -0.886733948 | 0.375222157 | 0.679214334 |
| NPIPB4 | 236.2986723 | -0.738565106 | 0.414003269 | -1.783959599 | 0.07443023 | 0.313367219 |
| NPIPB5 | 2382.315677 | -1.294948941 | 0.35798066 | -3.617371233 | 0.00029761 | 0.012472503 |
| NPIPB6 | 10.8406305 | -0.062598375 | 1.002602967 | -0.062435856 | 0.950215741 | 0.981636196 |
| NPIPB7 | 47.25225846 | -0.281479617 | 0.610500179 | -0.461063938 | 0.644752733 | 0.845794147 |
| NPIPP1 | 324.0991127 | 0.411171905 | 0.358267634 | 1.1476669 | 0.2511061 | 0.564075361 |
| NPL | 50.16984579 | 0.568002313 | 0.656885495 | 0.864689992 | 0.387208964 | 0.68748084 |
| NPLOC4 | 2006.044627 | -0.357133037 | 0.309807343 | -1.152758464 | 0.249009537 | 0.5617433 |
| NPM1 | 69438.72895 | 0.627464013 | 0.278028281 | 2.256835208 | 0.024018374 | 0.177207433 |
| NPM1P40 | 32.7571093 | 3.114988908 | 1.118559643 | 2.78482147 | 0.00535572 | 0.077809098 |
| NPM3 | 1334.704882 | 1.177028206 | 0.465508251 | 2.528479791 | 0.011455768 | 0.121279761 |
| NPNT | 225.0459838 | -2.662662552 | 0.851439653 | -3.12724753 | 0.001764513 | 0.039027635 |
| NPPB | 28.27928981 | -1.66357967 | 0.893608195 | -1.861643256 | 0.062653392 | 0.288822427 |
| NPR2 | 289.1290851 | 0.418564343 | 0.484049124 | 0.864714596 | 0.387195456 | 0.68748084 |
| NPRL2 | 470.9094624 | 0.318913887 | 0.283610461 | 1.124478573 | 0.260810055 | 0.57415519 |
| NPRL3 | 919.1585976 | -0.420792552 | 0.263309853 | -1.598088891 | 0.110023196 | 0.38152272 |
| NPTN | 2506.590644 | 0.202702384 | 0.259151024 | 0.782178596 | 0.434109622 | 0.719296316 |
| NPTX1 | 23.91032479 | 2.57886281 | 0.944722184 | 2.729757862 | 0.006338086 | 0.086470416 |
| NPTX2 | 232.31292 | 1.73659408 | 0.508012897 | 3.418405498 | 0.000629892 | 0.020418995 |
| NPTXR | 200.7893342 | -0.503969413 | 0.493905621 | -1.020375941 | 0.3075502 | 0.617319594 |
| NPY | 11.76272735 | 0.773184196 | 1.115113202 | 0.693368345 | 0.488078422 | 0.756733546 |
| NPY1R | 69.7676797 | -0.420951983 | 0.49684828 | -0.847244523 | 0.396858842 | 0.695477127 |
| NQO1 | 872.9487974 | 0.883750433 | 0.386049537 | 2.289215108 | 0.022066857 | 0.168823142 |
| NQO2 | 1101.110668 | 0.594370783 | 0.279018237 | 2.130221988 | 0.033153291 | 0.207838253 |
| NR1D1 | 105.6777284 | 0.528044238 | 0.509052432 | 1.037308154 | 0.299592267 | 0.610246278 |
| NR1D2 | 1066.558937 | -0.012793074 | 0.28117182 | -0.045499131 | 0.963709467 | 0.988130579 |
| NR1H2 | 461.3678685 | 0.325491831 | 0.43012759 | 0.756733207 | 0.449209716 | 0.730513518 |
| NR1H3 | 427.0601953 | 0.065181214 | 0.294862825 | 0.221056057 | 0.825048785 | 0.926977119 |
| NR1I2 | 10.88822971 | -0.212901429 | 0.949503973 | -0.224223843 | 0.822583132 | 0.925840841 |
| NR2C1 | 1291.969201 | 0.21346577 | 0.23757818 | 0.898507473 | 0.368915063 | 0.674802318 |
| NR2C2 | 1394.348443 | -0.525933578 | 0.36232898 | -1.451536053 | 0.146630651 | 0.43954428 |
| NR2C2AP | 300.3904296 | 0.522031172 | 0.370916745 | 1.407407938 | 0.159306459 | 0.459178093 |
| NR2F1 | 523.0808134 | 1.305935202 | 0.833022251 | 1.567707465 | 0.116949419 | 0.39436311 |
| NR2F2 | 1403.80806 | -0.742723094 | 0.605113787 | -1.227410629 | 0.219668293 | 0.528259775 |
| NR2F6 | 1505.349128 | 0.396184011 | 0.475015353 | 0.834044644 | 0.404255827 | 0.700367583 |
| NR3C1 | 52.58638366 | -2.911134038 | 0.803590248 | -3.622659738 | 0.000291589 | 0.012288058 |
| NR3C2 | 19.17808721 | -1.360043391 | 0.821843567 | -1.654868939 | 0.097951086 | 0.359375073 |
| NR4A1 | 751.9735511 | -2.577009407 | 0.835621009 | -3.08394521 | 0.002042752 | 0.042686766 |
| NR4A3 | 50.06970592 | -0.752922236 | 0.70312927 | -1.070816233 | 0.284252066 | 0.598246396 |

| | | | | | | |
|---|---|---|---|---|---|---|
| NR6A1 | 7285.2303 | 0.217059542 | 0.259445234 | 0.836629523 | 0.402800845 | 0.699667005 |
| NRADDP | 11.91808406 | 0.557099923 | 1.146249416 | 0.48601981 | 0.62695312 | 0.836041947 |
| NRARP | 893.0011068 | 0.679034127 | 0.348895074 | 1.946241659 | 0.051625718 | 0.261823226 |
| NRAS | 4695.362349 | 0.227559402 | 0.254654796 | 0.893599515 | 0.371536206 | 0.67682226 |
| NRBF2 | 960.3891115 | 0.196517561 | 0.345118489 | 0.569420553 | 0.569070772 | 0.805450586 |
| NRBP1 | 3568.39004 | -0.242795628 | 0.289662567 | -0.838201604 | 0.401917486 | 0.699116854 |
| NRBP2 | 644.7145233 | -0.454075525 | 0.343150719 | -1.323253892 | 0.185750959 | 0.490262318 |
| NRCAM | 836.9520953 | 1.61850807 | 0.731843759 | 2.211548641 | 0.026997867 | 0.188055394 |
| NRDC | 4781.619694 | -0.468221509 | 0.339879709 | -1.37760948 | 0.168323893 | 0.470630625 |
| NRDE2 | 749.3582846 | -0.180190597 | 0.294271257 | -0.612328225 | 0.540320619 | 0.787481586 |
| NREP | 7470.623612 | -0.038013064 | 0.218374561 | -0.174072768 | 0.861808274 | 0.944211261 |
| NRF1 | 616.6791799 | 0.246386059 | 0.310346354 | 0.79390673 | 0.427249732 | 0.714721103 |
| NRG1 | 706.2106534 | 2.027659437 | 0.648581387 | 3.126299147 | 0.001770214 | 0.039034758 |
| NRG3 | 23.214705 | -0.698405156 | 0.732835324 | -0.953017866 | 0.340581019 | 0.647332681 |
| NRGN | 29.72413064 | 1.552412389 | 0.61284931 | 2.533106206 | 0.011305669 | 0.120702539 |
| NRIP1 | 3369.6927 | -0.728625466 | 0.390653673 | -1.865144288 | 0.062161182 | 0.287461403 |
| NRIP2 | 14.94932693 | 1.577958973 | 0.881966216 | 1.789137662 | 0.073592648 | 0.311651561 |
| NRIP3 | 126.1491278 | 0.353682216 | 0.558261718 | 0.633541948 | 0.526379797 | 0.778525486 |
| NRK | 47.91667966 | -4.319177962 | 1.454436687 | -2.969656912 | 0.002981325 | 0.053986643 |
| NRM | 119.0316888 | 0.568116624 | 0.359900425 | 1.578538353 | 0.114441984 | 0.389282364 |
| NRN1 | 10.288985 | 0.058686947 | 0.911446706 | 0.064388786 | 0.948660659 | 0.98134156 |
| NRN1L | 39.16451075 | -0.659397993 | 0.885321041 | -0.744812291 | 0.456385204 | 0.735468477 |
| NRP1 | 339.0000709 | -4.052969362 | 1.69960695 | -2.384650969 | 0.017095332 | 0.14773715 |
| NRP2 | 3428.266733 | 0.281040044 | 0.525830046 | 0.534469352 | 0.593016848 | 0.819859199 |
| NRROS | 31.71066339 | -0.401164451 | 0.649761084 | -0.617403012 | 0.536968936 | 0.785493754 |
| NRSN2 | 633.9155798 | -0.015951282 | 0.242612022 | -0.065748109 | 0.94757837 | 0.980942439 |
| NRTN | 34.3106866 | 0.801545857 | 0.560882771 | 1.429079119 | 0.152981494 | 0.449260986 |
| NRXN1 | 265.7807621 | -0.055289481 | 0.430025195 | -0.128572656 | 0.897695804 | 0.959688749 |
| NRXN2 | 126.9796661 | 0.488637826 | 0.477997836 | 1.022259494 | 0.306658099 | 0.616853623 |
| NRXN3 | 998.4468945 | 0.02442204 | 0.288729385 | 0.084584532 | 0.932591697 | 0.97453841 |
| NSA2 | 4001.452378 | 0.669300805 | 0.332977746 | 2.01004666 | 0.044426251 | 0.242233633 |
| NSD1 | 3882.846678 | -0.608522447 | 0.488802086 | -1.244926043 | 0.213158937 | 0.52032731 |
| NSD2 | 5222.593526 | 0.06101695 | 0.418634486 | 0.145752325 | 0.884116913 | 0.954046354 |
| NSD3 | 2274.643878 | -0.120162112 | 0.324223057 | -0.370615566 | 0.710923886 | 0.876976263 |
| NSDHL | 765.4557564 | 0.170175679 | 0.249719223 | 0.681468078 | 0.495575359 | 0.760859008 |
| NSF | 568.8468712 | 0.087608187 | 0.332851738 | 0.263204834 | 0.792392705 | 0.91361831 |
| NSFL1C | 2046.250798 | -0.019108266 | 0.232167803 | -0.082303686 | 0.934405223 | 0.97507153 |
| NSFP1 | 43.6692116 | 0.362328956 | 1.02564626 | 0.353268929 | 0.723886837 | 0.883585743 |
| NSG1 | 655.9047112 | -0.360771425 | 0.44394666 | -0.812645883 | 0.416421117 | 0.708268589 |
| NSL1 | 1257.572776 | 0.445366779 | 0.251236139 | 1.772701893 | 0.076278117 | 0.316784921 |
| NSMAF | 1014.757589 | -0.488841235 | 0.274032464 | -1.783880738 | 0.074443046 | 0.313367219 |
| NSMCE1 | 999.0142263 | 0.873206913 | 0.406348397 | 2.148911921 | 0.031641383 | 0.202592072 |
| NSMCE2 | 769.998502 | 0.477386365 | 0.282310977 | 1.690994698 | 0.090837815 | 0.345388595 |

| | | | | | | |
|---|---|---|---|---|---|---|
| NSMCE4A | 2012.348215 | 0.776529051 | 0.350142008 | 2.21775461 | 0.026571569 | 0.186369517 |
| NSMF | 1080.202457 | 0.243855957 | 0.452255145 | 0.539199963 | 0.589748886 | 0.81828099 |
| NSRP1 | 1081.32249 | 0.356265923 | 0.414035824 | 0.860471249 | 0.389529326 | 0.689321054 |
| NSUN2 | 2274.167211 | -0.349783575 | 0.246420984 | -1.41945531 | 0.155766317 | 0.454098923 |
| NSUN3 | 378.5665196 | -0.042550741 | 0.275158015 | -0.15464111 | 0.877104259 | 0.950622089 |
| NSUN4 | 740.9359312 | -0.075596089 | 0.246826816 | -0.306271782 | 0.759397722 | 0.899496446 |
| NSUN5 | 858.2657436 | 0.21450816 | 0.301481959 | 0.711512427 | 0.476766753 | 0.749746108 |
| NSUN5P1 | 1864.747669 | -0.14197049 | 0.388506298 | -0.365426484 | 0.714793075 | 0.87855355 |
| NSUN5P2 | 294.3827909 | 0.727482975 | 0.448469686 | 1.62214526 | 0.104772249 | 0.371204996 |
| NSUN6 | 334.2327676 | 0.216440823 | 0.291797285 | 0.741750639 | 0.458238432 | 0.73666793 |
| NSUN7 | 642.3327026 | 0.023200924 | 0.325767743 | 0.071219219 | 0.943223286 | 0.978607681 |
| NT5C | 681.2520911 | 0.683600083 | 0.390770001 | 1.749366842 | 0.080227629 | 0.324292679 |
| NT5C2 | 2217.578926 | -0.892020692 | 0.297162879 | -3.001790444 | 0.002683969 | 0.050458917 |
| NT5C3A | 595.0639363 | 0.423635067 | 0.340981078 | 1.242400515 | 0.214088826 | 0.520920862 |
| NT5C3B | 1408.45855 | 0.242049362 | 0.314158338 | 0.770469323 | 0.441021545 | 0.724576333 |
| NT5DC1 | 1595.029357 | 0.107081942 | 0.252428998 | 0.424206184 | 0.671415451 | 0.858479663 |
| NT5DC2 | 7299.965871 | 0.04621493 | 0.238448981 | 0.193814751 | 0.846320936 | 0.938145276 |
| NT5DC3 | 266.2175952 | 0.355069747 | 0.303622711 | 1.169443966 | 0.242224805 | 0.553231259 |
| NT5M | 210.824504 | 1.301042473 | 0.36852433 | 3.530411337 | 0.000414914 | 0.015721465 |
| NTAN1 | 209.0212928 | 0.734060139 | 0.588370628 | 1.2476152 | 0.212172009 | 0.519367057 |
| NTF3 | 164.3063058 | 0.649628058 | 0.454396647 | 1.429649761 | 0.152817566 | 0.448899925 |
| NTF4 | 64.57020636 | -0.482150781 | 0.694918622 | -0.693823371 | 0.487792984 | 0.756677644 |
| NTHL1 | 480.9021288 | 1.167730213 | 0.474896173 | 2.458916873 | 0.013935688 | 0.134161613 |
| NTM | 71.49282469 | -0.128556332 | 0.438362019 | -0.293265216 | 0.769319439 | 0.904597192 |
| NTMT1 | 610.6452161 | 0.73726071 | 0.317740492 | 2.320323434 | 0.020323387 | 0.161005801 |
| NTN1 | 152.1154869 | 1.038692864 | 0.488651552 | 2.125630953 | 0.033534 | 0.208789229 |
| NTN4 | 25.40159738 | 1.372564973 | 0.728566423 | 1.883925652 | 0.059575033 | 0.282669434 |
| NTN5 | 13.64553294 | 0.244478767 | 1.033934061 | 0.236454892 | 0.813079706 | 0.921802034 |
| NTNG2 | 20.69211258 | 1.540301779 | 0.714430402 | 2.155985767 | 0.031084777 | 0.200942211 |
| NTPCR | 1529.123676 | 0.822286562 | 0.274200349 | 2.998853088 | 0.002709979 | 0.050819652 |
| NTRK1 | 11.32110693 | 1.834777947 | 0.992537104 | 1.84857366 | 0.064519399 | 0.293324484 |
| NTRK2 | 69.15049011 | 4.080137014 | 1.063564626 | 3.836284994 | 0.000124909 | 0.007078495 |
| NTRK3 | 61.90750476 | 1.217440595 | 0.572136003 | 2.127886706 | 0.033346478 | 0.208274771 |
| NTS | 23.08290884 | 0.300779963 | 0.858679803 | 0.350281866 | 0.726127173 | 0.8844924 |
| NUAK1 | 1066.945104 | 0.305963135 | 0.572849234 | 0.534107609 | 0.593267086 | 0.819859199 |
| NUAK2 | 720.6543952 | -0.509421633 | 0.531870468 | -0.957792666 | 0.338167318 | 0.645406515 |
| NUB1 | 1312.775631 | -0.021898992 | 0.240349714 | -0.091113036 | 0.927402774 | 0.972633911 |
| NUBP1 | 587.2853024 | 0.148883161 | 0.256896697 | 0.579544865 | 0.562221583 | 0.801706045 |
| NUBP2 | 568.2798492 | 0.263950635 | 0.521522462 | 0.506115564 | 0.612775525 | 0.829444816 |
| NUBPL | 441.771069 | -0.321388503 | 0.278525639 | -1.153891988 | 0.248544454 | 0.561194808 |
| NUCB1 | 3408.626089 | -0.438235373 | 0.271243153 | -1.615655058 | 0.106168913 | 0.373883142 |
| NUCB2 | 1090.713651 | 0.546887008 | 0.257669598 | 2.122435137 | 0.033801214 | 0.209476397 |
| NUCKS1 | 18010.55567 | 0.673719112 | 0.269630597 | 2.498674553 | 0.012465873 | 0.126670974 |
| NUDC | 2721.68733 | 0.191245451 | 0.311216513 | 0.614509331 | 0.538878809 | 0.786598616 |
| NUDCD1 | 1363.575714 | 0.622726633 | 0.301572394 | 2.064932486 | 0.038929389 | |

0.227316695

| | | | | | | |
|---|---|---|---|---|---|---|
| NUDCD2 | 1243.577799 | 1.201194589 | 0.345099106 | 3.480723565 | 0.000500061 | 0.017561181 |
| NUDCD3 | 1498.74011 | -0.287584711 | 0.340259371 | -0.845192624 | 0.3980033 | 0.696517252 |
| NUDT1 | 832.1462649 | 0.893087077 | 0.41384909 | 2.158001788 | 0.030927693 | 0.200686068 |
| NUDT10 | 481.0782755 | 0.724031419 | 0.327546111 | 2.210471735 | 0.02707244 | 0.188125165 |
| NUDT11 | 1374.615216 | 0.62933846 | 0.290400185 | 2.167142075 | 0.030224023 | 0.198239795 |
| NUDT12 | 889.2384301 | 0.479937242 | 0.256499292 | 1.871105526 | 0.061330456 | 0.285640507 |
| NUDT13 | 25.51018314 | -0.391612061 | 0.737724241 | -0.530838 | 0.595531045 | 0.821568613 |
| NUDT14 | 285.9309189 | 0.730895503 | 0.411140065 | 1.777728725 | 0.07544842 | 0.314738565 |
| NUDT15 | 1470.907618 | 0.571008053 | 0.301136883 | 1.896174417 | 0.057936985 | 0.278947 |
| NUDT16 | 1077.089228 | -0.094725642 | 0.357370198 | -0.26506307 | 0.790960876 | 0.91335572 |
| NUDT16L1 | 659.1178949 | 0.63658437 | 0.475175277 | 1.339683271 | 0.180348338 | 0.484774033 |
| NUDT16P1 | 191.6248173 | 0.967826009 | 0.365449312 | 2.648318053 | 0.008089338 | 0.100970518 |
| NUDT17 | 91.82396852 | -0.72986911 | 0.442958995 | -1.647712586 | 0.099411664 | 0.361499127 |
| NUDT18 | 69.60496597 | -0.063410455 | 0.451803329 | -0.140349685 | 0.88838371 | 0.955943827 |
| NUDT19 | 173.5301108 | -0.225061052 | 0.578851375 | -0.388806284 | 0.697419452 | 0.87033849 |
| NUDT2 | 407.8718968 | 0.732157959 | 0.379450114 | 1.929523622 | 0.05366589 | 0.267861241 |
| NUDT21 | 5860.178142 | 0.141264743 | 0.222365978 | 0.635280381 | 0.525245567 | 0.778249877 |
| NUDT22 | 339.5151929 | 0.527855047 | 0.411560204 | 1.282570669 | 0.199642531 | 0.505774645 |
| NUDT3 | 1550.927641 | 0.215965154 | 0.270300826 | 0.798980742 | 0.424301579 | 0.713303793 |
| NUDT4 | 4105.8141 | -0.312857634 | 0.475024511 | -0.658613665 | 0.510143885 | 0.769949736 |
| NUDT4B | 45.4259321 | -0.407336077 | 1.265934378 | -0.321767134 | 0.747629114 | 0.894202292 |
| NUDT4P2 | 171.2519727 | -1.604987895 | 0.698310674 | -2.298386598 | 0.021539796 | 0.166469811 |
| NUDT5 | 2502.242761 | 0.599958666 | 0.280926656 | 2.135641644 | 0.032708634 | 0.206178012 |
| NUDT6 | 216.0534372 | 0.278362278 | 0.34764446 | 0.800709661 | 0.423299748 | 0.712671233 |
| NUDT7 | 157.4217973 | 1.03256473 | 0.380360121 | 2.714702918 | 0.006633525 | 0.089169476 |
| NUDT8 | 57.91986739 | 0.827223614 | 0.619363081 | 1.335603685 | 0.181678845 | 0.48591777 |
| NUDT9 | 401.9333123 | 0.156592284 | 0.336247142 | 0.465705917 | 0.641426016 | 0.843948481 |
| NUF2 | 1159.521515 | 1.366512384 | 0.282195851 | 4.842425503 | 1.28E-06 | 0.000218639 |
| NUFIP1 | 600.5050123 | 0.237644325 | 0.315684992 | 0.752789429 | 0.45157646 | 0.732712992 |
| NUFIP2 | 3567.583531 | -0.111306218 | 0.365281738 | -0.304713338 | 0.760584491 | 0.899924139 |
| NUMA1 | 4827.442234 | -0.437398303 | 0.441936614 | -0.989730856 | 0.322305689 | 0.632029271 |
| NUMB | 1818.083228 | -0.647910556 | 0.232278141 | -2.789373777 | 0.005281008 | 0.077250742 |
| NUMBL | 312.9894182 | 0.344952122 | 0.360734719 | 0.956248745 | 0.338946578 | 0.645999817 |
| NUP107 | 3384.186424 | 0.202299253 | 0.251159143 | 0.805462428 | 0.420552879 | 0.711203626 |
| NUP133 | 2852.571813 | -0.092796859 | 0.232172391 | -0.399689466 | 0.689385252 | 0.867579108 |
| NUP153 | 2596.330887 | -0.219185518 | 0.34042706 | -0.643854568 | 0.519669745 | 0.774983751 |
| NUP155 | 1782.145846 | 0.038273491 | 0.381675594 | 0.100277543 | 0.920123986 | 0.969770994 |

| | | | | | | |
|---|---|---|---|---|---|---|
| NUP160 | 1746.216391 | -0.137014043 | 0.29690474 | -0.461474759 | 0.644458027 | 0.845794147 |
| NUP188 | 2681.784378 | -0.255481595 | 0.506635495 | -0.504271014 | 0.614070944 | 0.830316752 |
| NUP205 | 4257.065361 | -0.130554111 | 0.368888286 | -0.353912325 | 0.72340459 | 0.883279173 |
| NUP210 | 895.3168354 | 0.082946955 | 0.692251719 | 0.119821957 | 0.904624188 | 0.963993604 |
| NUP214 | 1355.214888 | -0.757894484 | 0.487754627 | -1.553843762 | 0.120221689 | 0.400307118 |
| NUP35 | 519.4729833 | 0.782219598 | 0.269261831 | 2.905051914 | 0.003671922 | 0.062311776 |
| NUP37 | 1539.167015 | 0.609536425 | 0.275648009 | 2.211285424 | 0.027016078 | 0.188095879 |
| NUP43 | 2108.039509 | 0.162299018 | 0.247987208 | 0.654465281 | 0.512812088 | 0.771819984 |
| NUP50 | 3183.019682 | -0.301467002 | 0.325146347 | -0.927173271 | 0.353836579 | 0.660911171 |
| NUP54 | 2344.651382 | 0.517062739 | 0.251290339 | 2.057630795 | 0.039625588 | 0.229275285 |
| NUP58 | 2132.584618 | 0.177596442 | 0.331574422 | 0.535615625 | 0.592224228 | 0.819765854 |
| NUP62 | 2872.926526 | 0.21678115 | 0.249639375 | 0.868377237 | 0.385187852 | 0.685935162 |
| NUP62CL | 109.9509131 | 0.264988912 | 0.375004502 | 0.706628615 | 0.479797302 | 0.751963313 |
| NUP85 | 1502.515289 | 0.083707873 | 0.228471861 | 0.366381545 | 0.714080389 | 0.878400517 |
| NUP88 | 1432.61636 | 0.118514823 | 0.297534308 | 0.398323217 | 0.690391949 | 0.867938158 |
| NUP93 | 2603.53577 | -0.290001792 | 0.30888126 | -0.938877911 | 0.347793432 | 0.654455642 |
| NUP98 | 2861.102838 | -0.227144032 | 0.361237565 | -0.628794051 | 0.529483895 | 0.780195218 |
| NUPL2 | 885.8801833 | 0.165655429 | 0.250381706 | 0.661611549 | 0.508220204 | 0.768714727 |
| NUPR1 | 27.34005976 | 0.296454946 | 0.811118078 | 0.36548926 | 0.714746222 | 0.87855355 |
| NUPR2 | 22.61937863 | 1.941361351 | 0.738049867 | 2.6303932 | 0.008528616 | 0.10430915 |
| NUS1 | 3192.154984 | 0.173222339 | 0.299211569 | 0.578929283 | 0.56263689 | 0.801706045 |
| NUS1P2 | 48.1699014 | -0.568391041 | 0.557218527 | -1.020050508 | 0.307704507 | 0.617403132 |
| NUSAP1 | 2440.619654 | 0.950284587 | 0.378136441 | 2.51307328 | 0.011968447 | 0.124280158 |
| NUTF2 | 6093.536479 | 0.54961217 | 0.430621426 | 1.276323324 | 0.201841248 | 0.50842771 |
| NUTM2A | 29.19160178 | -0.303872911 | 0.653934146 | -0.464684269 | 0.642157575 | 0.844148268 |
| NUTM2D | 158.7125063 | -0.589293557 | 0.343236926 | -1.716871098 | 0.086002727 | 0.33625288 |
| NVL | 1010.858866 | -0.192028706 | 0.280230918 | -0.685251675 | 0.493185121 | 0.759389046 |
| NWD2 | 15.70493213 | 0.743631663 | 0.905791495 | 0.820974437 | 0.411660829 | 0.704686727 |
| NXF1 | 1565.788576 | -0.616511042 | 0.285869487 | -2.15661716 | 0.031035506 | 0.200870165 |
| NXF2 | 10.24427837 | 0.688844224 | 0.933351987 | 0.738032632 | 0.46049462 | 0.738395128 |
| NXF2B | 11.83873469 | 0.518993863 | 0.937612201 | 0.553527207 | 0.579902456 | 0.811340629 |
| NXN | 2325.385453 | 0.060736797 | 0.233576003 | 0.260030124 | 0.794840537 | 0.914877916 |
| NXNL2 | 23.39670413 | 0.445818611 | 1.064407731 | 0.418841951 | 0.675331643 | 0.860095396 |
| NXPE3 | 759.625543 | 0.576711333 | 0.355209825 | 1.62357934 | 0.104465618 | 0.370464676 |
| NXPH2 | 94.21140879 | 0.197239965 | 0.778576531 | 0.253334075 | 0.800010067 | 0.916311991 |
| NXPH3 | 542.3664194 | -1.329718804 | 0.415858319 | -3.197528448 | 0.001386107 | 0.033753829 |
| NXT1 | 341.9880654 | 1.083223048 | 0.399039029 | 2.714579197 | 0.006636003 | 0.089169476 |
| NXT2 | 461.4718914 | 0.183197389 | 0.261641168 | 0.700185639 | 0.483811379 | 0.754280386 |
| NYAP1 | 109.2424101 | 0.544612045 | 0.622036985 | 0.875530006 | 0.381285591 | 0.682967936 |
| NYNRIN | 2061.063125 | -0.890148726 | 0.269836547 | -3.298844193 | 0.000970838 | 0.026682211 |
| OAF | 26.64758915 | 0.936462603 | 0.578460888 | 1.618886641 | 0.105471656 | 0.372436233 |
| OARD1 | 1419.564436 | 0.40755203 | 0.298051589 | 1.367387542 | 0.171503865 | 0.474879827 |
| OAS3 | 30.32188652 | -0.906415277 | 0.610299976 | -1.485196318 | 0.137491827 | 0.425712285 |
| OAT | 1976.896123 | 0.319935418 | 0.312024033 | 1.025355051 | 0.305195689 | 0.615135352 |
| OAZ1 | 9771.741374 | 0.759216644 | 0.423845237 | 1.791259115 | 0.073251724 | 0.311262322 |
| OAZ2 | 5030.526706 | -0.035185664 | 0.30562427 | -0.115127192 | 0.908344307 | |

| | | | | | | |
|---|---|---|---|---|---|---|
| | | | | | | 0.965629002 |
| OAZ3 | 11.49458479 | -1.066326625 | 1.005696562 | -1.060286637 | 0.289014217 | 0.602119567 |
| OBSCN | 1431.019261 | -0.579755066 | 0.501786917 | -1.155380992 | 0.247934442 | 0.560705687 |
| OBSL1 | 2437.119122 | -0.72988408 | 0.474465774 | -1.538328199 | 0.123968388 | 0.405066642 |
| OCEL1 | 310.3452421 | 0.790649283 | 0.573664743 | 1.378242767 | 0.168128347 | 0.470384431 |
| OCIAD1 | 3645.844224 | 0.226316459 | 0.256507891 | 0.882298231 | 0.377615553 | 0.680934859 |
| OCIAD2 | 568.548353 | 0.564858714 | 0.530512878 | 1.064740816 | 0.286993234 | 0.601236469 |
| OCLM | 10.77155637 | -1.510328641 | 0.9964976 | -1.515637008 | 0.129611164 | 0.414313309 |
| OCLN | 796.944143 | -0.224002299 | 0.520721589 | -0.430176707 | 0.667067104 | 0.856327216 |
| OCRL | 1127.68424 | -0.419681569 | 0.315256228 | -1.331239581 | 0.183110191 | 0.487105859 |
| ODC1 | 3727.400551 | 1.313817109 | 0.34140714 | 3.848241453 | 0.000118969 | 0.00691523 |
| ODF2 | 1398.277617 | -0.008235126 | 0.410703614 | -0.020051262 | 0.984002479 | 0.994061643 |
| ODF2L | 1208.353658 | -0.473207462 | 0.269864116 | -1.753502726 | 0.079515752 | 0.323002882 |
| ODR4 | 820.9880833 | -0.140680519 | 0.278292238 | -0.505513629 | 0.613198129 | 0.829794738 |
| OFCC1 | 17.48117077 | -0.426428218 | 0.842351654 | -0.50623539 | 0.612691413 | 0.829404964 |
| OFD1 | 1151.278604 | -0.429197594 | 0.249428732 | -1.720722351 | 0.085301219 | 0.334653423 |
| OGA | 5623.278811 | -0.305822994 | 0.240358063 | -1.272364197 | 0.203243749 | 0.509654697 |
| OGDH | 2503.302257 | 0.045393616 | 0.413643765 | 0.109740844 | 0.912614907 | 0.967485958 |
| OGDHL | 224.2633626 | 0.229726379 | 0.426364076 | 0.538803318 | 0.590022574 | 0.818321084 |
| OGFOD1 | 1139.13777 | -0.100975495 | 0.326023056 | -0.309718876 | 0.756774747 | 0.898563152 |
| OGFOD2 | 348.5589216 | 0.007432366 | 0.388411442 | 0.019135292 | 0.984733178 | 0.994433294 |
| OGFOD3 | 657.7351433 | 0.134869983 | 0.270035108 | 0.499453513 | 0.617459929 | 0.831835425 |
| OGFR | 398.6377009 | -0.358592564 | 0.463141776 | -0.774260891 | 0.438776526 | 0.722407379 |
| OGFRL1 | 274.6150096 | 0.445245731 | 0.292123561 | 1.524169188 | 0.127466453 | 0.411270428 |
| OGG1 | 607.6222079 | 0.119458255 | 0.284218562 | 0.420304199 | 0.674263243 | 0.859672768 |
| OGT | 5814.792842 | -0.30507047 | 0.343215648 | -0.888859446 | 0.374078623 | 0.678520659 |
| OIP5 | 546.3162704 | 1.235028411 | 0.353723777 | 3.491505212 | 0.000480307 | 0.017185705 |
| OLA1 | 5160.490863 | 0.275059289 | 0.21931331 | 1.254184202 | 0.209775061 | 0.517783588 |
| OLA1P3 | 14.91639854 | -1.107652383 | 0.879306719 | -1.259688296 | 0.207781829 | 0.514566354 |
| OLFM1 | 139.230211 | 0.519008983 | 0.396819586 | 1.307921789 | 0.190899838 | 0.496058205 |
| OLFM2 | 658.6006114 | 0.062962993 | 0.36085956 | 0.174480601 | 0.861487775 | 0.94403207 |
| OLFM3 | 558.7589515 | 0.577389388 | 0.444486665 | 1.299002723 | 0.193942995 | 0.499882632 |
| OLFML1 | 77.82149248 | -1.821913799 | 0.851943863 | -2.138537382 | 0.032473154 | 0.205554752 |
| OLFML2A | 301.9784818 | 0.985525187 | 0.548637432 | 1.796314157 | 0.072444568 | 0.309775814 |
| OLFML2B | 212.2520744 | 0.213466605 | 0.500547924 | 0.426465867 | 0.669768414 | 0.857762448 |
| OLFML3 | 3575.049378 | -1.567971268 | 0.449929212 | -3.484928801 | 0.000492268 | 0.017449061 |
| OLIG3 | 242.9670846 | 2.198392434 | 0.515410305 | 4.265324951 | 2.00E-05 | 0.001857856 |
| OMA1 | 324.1925639 | 0.055080537 | 0.27639075 | 0.199285024 | 0.842039793 | 0.935422069 |
| ONECUT2 | 63.59783292 | 1.263036046 | 0.591220386 | 2.136320188 | 0.032653324 | |

| | | | | | | |
|---|---|---|---|---|---|---|
| | | | | | | 0.206152137 |
| OPA1 | 3085.183615 | 0.052011516 | 0.274881809 | 0.189214107 | 0.849925012 | 0.938580023 |
| OPA3 | 605.2943998 | -0.216316562 | 0.263806838 | -0.819980875 | 0.41222701 | 0.70515068 |
| OPCML | 41.58401865 | -1.922251656 | 0.593368284 | -3.239559154 | 0.001197146 | 0.030472998 |
| OPHN1 | 717.5437104 | -1.976564697 | 0.433384982 | -4.560759556 | 5.10E-06 | 0.000663809 |
| OPLAH | 183.6232151 | 0.079594621 | 0.65629786 | 0.121278196 | 0.90347069 | 0.963418418 |
| OPN3 | 904.9805461 | 0.42842324 | 0.281170652 | 1.523712509 | 0.127580543 | 0.411463525 |
| OPRD1 | 65.78612977 | -0.190258549 | 0.591350615 | -0.321735606 | 0.747653001 | 0.894202292 |
| OPRK1 | 49.16552551 | 0.070926072 | 0.584097768 | 0.121428425 | 0.903351704 | 0.963418418 |
| OPRL1 | 29.37908052 | -1.686198021 | 1.067622979 | -1.57939465 | 0.114245564 | 0.389050381 |
| OPRM1 | 55.38864165 | 0.227620922 | 0.499425887 | 0.455765165 | 0.648558856 | 0.846827301 |
| OPTC | 28.50719988 | 1.389024533 | 0.963014302 | 1.442371655 | 0.149197556 | 0.443905887 |
| OPTN | 1347.44874 | -0.868487163 | 0.310455825 | -2.797458104 | 0.005150645 | 0.076234566 |
| OR10G3 | 22.15352075 | 0.226251567 | 0.823176308 | 0.274851894 | 0.783430027 | 0.910341371 |
| OR10H5 | 14.70438599 | 0.252408713 | 0.821642766 | 0.307200068 | 0.758691094 | 0.899440455 |
| OR10K1 | 13.71137812 | -0.470884786 | 0.979424439 | -0.480777043 | 0.630674968 | 0.838192723 |
| OR13A1 | 14.45366193 | 0.534465623 | 0.976644641 | 0.547246768 | 0.58420921 | 0.814692336 |
| OR14J1 | 14.27189131 | -1.324661551 | 1.051899238 | -1.259304602 | 0.207920332 | 0.514730716 |
| OR1A1 | 24.24842494 | -0.166300239 | 0.656602548 | -0.253273825 | 0.800056622 | 0.916311991 |
| OR1D2 | 72.83948428 | 0.143042562 | 0.708087717 | 0.202012489 | 0.839906961 | 0.934455009 |
| OR1I1 | 15.8563667 | -0.26501159 | 0.802231113 | -0.330343196 | 0.741140657 | 0.891943909 |
| OR2A20P | 18.09337815 | -2.464847524 | 0.97384455 | -2.531048231 | 0.011372221 | 0.120838052 |
| OR2A7 | 12.24701206 | -2.237292189 | 1.013514939 | -2.207458522 | 0.027282042 | 0.188904612 |
| OR2A9P | 43.73460811 | 0.983198677 | 0.666149442 | 1.475943108 | 0.139959166 | 0.429213767 |
| OR2AT4 | 27.29538614 | -0.085867776 | 0.723616332 | -0.118664785 | 0.905540937 | 0.964473572 |
| OR2G6 | 12.05251356 | -0.71371164 | 0.905154901 | -0.788496685 | 0.430406237 | 0.717075886 |
| OR2L2 | 11.24859465 | -1.015475068 | 0.908647372 | -1.117567827 | 0.263751616 | 0.577444856 |
| OR2M3 | 69.51107664 | 0.101759712 | 0.562071818 | 0.181043967 | 0.856333065 | 0.942245037 |
| OR2T12 | 15.99413311 | 0.352771741 | 0.723801008 | 0.487387745 | 0.625983576 | 0.835544329 |
| OR4D1 | 24.09335335 | 0.998524243 | 0.833428555 | 1.19809219 | 0.230881129 | 0.539623726 |
| OR4D9 | 25.26967127 | 0.100221923 | 0.697266289 | 0.143735506 | 0.885709332 | 0.954813919 |
| OR4F17 | 83.6138091 | 0.260666089 | 0.452860331 | 0.575599298 | 0.564886055 | 0.803326429 |
| OR4K17 | 16.08778505 | -0.964409161 | 0.918582156 | -1.049888848 | 0.29376922 | 0.60636365 |
| OR52A1 | 11.13487019 | -0.126863446 | 0.861813366 | -0.147205243 | 0.882970022 | 0.953581235 |
| OR52I1 | 13.90993469 | 1.036181332 | 0.887362481 | 1.1677092 | 0.242924085 | 0.554112357 |
| OR52K1 | 13.96214983 | 0.44814368 | 0.825222451 | 0.543058032 | 0.587089847 | 0.81660769 |
| OR55B1P | 11.58019288 | -0.155516468 | 0.88283678 | -0.176155402 | 0.860171857 | 0.943475231 |
| OR56A1 | 14.1566944 | -1.498728621 | 0.875593539 | -1.71167163 | 0.0869572 | |

0.338697734
OR5A1 13.0640896 -0.060609304 1.228934598 -0.049318576 0.960665416
0.986479967
OR5A2 62.10300968 0.255107723 0.66875719 0.38146539 0.702857948 0.872703476
OR5AS1 124.98887 -0.280206999 0.468567271 -0.598008047
0.549834564 0.793878561
OR5AU1 10.68809627 -1.538015239 1.113325279 -1.381460808
0.16713732 0.469215449
OR6C75 13.95037802 0.473911879 0.967890388 0.489633831 0.624393033
0.834816841
OR6D1P 58.55975328 0.461223002 0.576621034 0.799871968 0.42378498
0.712860381
OR6Y1 13.94621064 -1.41589693 0.915745628 -1.546168375 0.122063909
0.402659864
OR7C1 18.76349554 -2.602025443 0.944122253 -2.756025966 0.005850835
0.082561966
OR7E104P 101.80501 -0.410443093 0.422763191 -0.970858159
0.331618916 0.639505603
OR7E122P 122.8176502 -1.698218229 0.422076892 -4.023480699 5.73E-05
0.004123081
OR7E14P 43.53598809 -0.111128576 0.716007418 -0.155205901
0.876658996 0.950324661
OR7E156P 64.81128795 -0.690938967 0.445707707 -1.550206459
0.12109197 0.401219509
OR7E22P 20.72708805 -0.237328867 1.063740669 -0.223107825
0.823451588 0.926199885
OR7E25P 13.65332809 1.007690966 0.966201083 1.042941251 0.296975503
0.608346659
OR7E38P 84.58055422 0.405236666 0.582702934 0.695442982 0.486777732
0.756340124
OR7E91P 29.00568207 -1.261666438 0.973348425 -1.296212544
0.194902269 0.500501471
OR9M1P 14.17289253 -0.47404331 0.908834964 -0.521594491 0.6019527
0.825021776
ORAI1 398.2127675 -0.301422988 0.455553198 -0.661663641 0.508186811
0.768714727
ORAI2 527.7283041 0.553235996 0.311625258 1.77532463 0.075844301 0.31593462
ORAI3 267.307176 1.073816841 0.42826598 2.507359657 0.012163687 0.125278548
ORC1 951.3072406 -0.206949032 0.399792721 -0.517640819 0.604708897
0.825894731
ORC2 1560.988989 0.09376095 0.297604998 0.315051665 0.752722427 0.896586461
ORC3 1781.443166 0.265119671 0.300290088 0.882878529 0.377301905 0.680704865
ORC4 1633.377807 0.465803793 0.224965817 2.070553644 0.038400528 0.225590715
ORC5 598.8021446 -0.027555371 0.253848201 -0.108550585 0.913558957
0.967939236
ORC6 1899.996409 0.23671303 0.292354607 0.809677783 0.41812539 0.709589334
ORMDL1 1378.445717 0.299417146 0.279298464 1.072032913 0.283705249
0.597624595
ORMDL2 560.7419833 0.191781601 0.42487195 0.451386826 0.651710777 0.8478779
ORMDL3 2671.521949 -0.271101214 0.410029618 -0.661174711
0.508500277 0.768908373
OS9 4359.193269 -0.536226138 0.28162245 -1.90406034 0.056902326 0.2761211
OSBP 1558.72459 -0.574995613 0.251837463 -2.283201261 0.022418516
0.170396445
OSBP2 28.95335704 0.338091742 0.608061825 0.556015406 0.578200323 0.810209453
OSBPL10 312.9970321 -0.821342464 0.293234874 -2.800971297
0.005094905 0.075897737
OSBPL11 680.7763592 -0.097802277 0.353815546 -0.276421649
0.782224231 0.909909835
OSBPL1A 1740.083977 0.184178791 0.285229688 0.64572097 0.518460078
0.774169079
OSBPL2 848.8064083 -0.528272956 0.297166168 -1.777702219
0.075452775 0.314738565
OSBPL3 538.3757602 -0.789005752 0.326159573 -2.419078937

0.015559863 0.141183424
OSBPL5 793.1962293 -0.959512768 0.543209233 -1.766377871
0.077332475 0.318807331
OSBPL6 304.8709881 1.224563913 0.515573596 2.375148615 0.017541886
0.149594125
OSBPL8 2402.82484 -0.058978436 0.379971277 -0.155218143
0.876649346 0.950324661
OSBPL9 3621.039554 -0.788806596 0.323768039 -2.4363325 0.014837039
0.138518598
OSCP1 124.5455693 0.130801842 0.372258613 0.351373581 0.7253081 0.884193635
OSER1 838.0356189 0.085573388 0.262426924 0.326084636 0.744360321 0.892725315
OSGEP 1044.874029 -0.249539945 0.236684441 -1.054314953 0.291738745
0.60488841
OSGEPL1 474.4172343 0.215957396 0.302254 0.714489787 0.474924369
0.748737095
OSGIN1 18.42815818 0.203129525 0.785807535 0.258497807 0.796022741
0.915299455
OSGIN2 625.9977853 -0.08915251 0.317932122 -0.28041366 0.779160158
0.908367621
OSMR 14.19082659 -1.991592854 0.910526359 -2.187298407 0.028720747
0.193826714
OST4 2503.988648 0.876212572 0.446169447 1.96385606 0.049546787 0.25654413
OSTC 4361.427324 0.684186378 0.371932192 1.839546003 0.065834918 0.29619856
OSTF1 395.6390807 -0.407399644 0.64609342 -0.630558417 0.528329296
0.7795452
OSTM1 572.6285978 0.714881841 0.264731035 2.700408142 0.006925446 0.091361686
OTOF 52.34199233 0.570908664 0.805383382 0.708865711 0.478407815 0.750837683
OTOG 16.6970267 -0.510157746 0.863014012 -0.591134951 0.554430003
0.796360966
OTOGL 69.10075461 -0.498851563 0.595838994 -0.837225439 0.402465861
0.699485574
OTUB1 2490.531182 0.115752886 0.288121096 0.401750819 0.68786742 0.866689498
OTUB2 114.1766977 -0.228900189 0.663933939 -0.344763501 0.730272175
0.886742369
OTUD1 123.3784626 0.123598131 0.378350506 0.326676266 0.743912754 0.892535157
OTUD3 922.8524568 -0.053584728 0.391928492 -0.136720676 0.891251586
0.957017706
OTUD4 1608.132567 -0.166805274 0.322989745 -0.516441394 0.605546161
0.826145757
OTUD5 1791.427685 -0.200287328 0.266619292 -0.751210935 0.45252572
0.733138807
OTUD6B 602.8158909 0.460067287 0.301489456 1.525981347 0.12701451
0.410423244
OTUD7A 51.91434295 0.372547427 0.481880828 0.773111121 0.43945662
0.72310156
OTUD7B 807.0409305 -0.59505203 0.276411825 -2.152773422 0.03133649
0.202023044
OTULIN 609.9075075 0.112346668 0.284486043 0.394911002 0.69290857
0.86854217
OTULINL 165.4596685 0.330118762 0.413562415 0.798232021 0.424735858
0.713505449
OTX1 116.6908015 -1.909476694 1.009284795 -1.891910691 0.058502881
0.280248566
OTX2 1844.577558 -0.233878571 0.734007353 -0.318632463 0.750005234
0.895752516
OVCA2 349.2923392 0.261183039 0.548009511 0.476603113 0.633644766 0.839689885
OVCH2 12.59612254 -1.225246964 1.08509146 -1.129164692 0.258828365
0.57198618
OVGP1 80.47886614 -0.105031358 0.483199906 -0.217366263 0.827922921
0.928475653
OVOL1 159.9294221 -1.28808915 0.645208318 -1.996392658 0.045891199
0.246869858
OVOL2 277.728237 -1.09092374 0.857715282 -1.271894955 0.203410445
0.509735688

| | | | | | | |
|---|---|---|---|---|---|---|
| OXA1L | 3445.156934 | -0.291282333 | 0.286938135 | -1.015139842 | 0.310039172 | 0.619783075 |
| OXCT1 | 1041.239866 | 0.035154297 | 0.27156296 | 0.129451738 | 0.897000211 | 0.95928311 |
| OXCT2 | 16.89302613 | 1.685843945 | 0.967720261 | 1.74207776 | 0.081494839 | 0.326388119 |
| OXCT2P1 | 12.09813328 | -0.784216945 | 1.119342938 | -0.700604719 | 0.483549732 | 0.754231168 |
| OXER1 | 97.33851759 | -1.003790792 | 0.554156516 | -1.811384984 | 0.070081279 | 0.304116772 |
| OXGR1 | 13.57423619 | 1.064664538 | 0.85428362 | 1.246265892 | 0.212666796 | 0.519735803 |
| OXLD1 | 348.5189801 | 0.57951097 | 0.406774363 | 1.424649692 | 0.154258488 | 0.451476933 |
| OXNAD1 | 454.4442763 | 0.361134232 | 0.278987175 | 1.294447429 | 0.195510917 | 0.501283779 |
| OXR1 | 933.4513796 | 0.215941587 | 0.322730217 | 0.669108673 | 0.503426158 | 0.76527838 |
| OXSM | 171.3279039 | 0.362665399 | 0.327667069 | 1.106810642 | 0.268375806 | 0.581480913 |
| OXSR1 | 2236.440583 | -0.047278239 | 0.271072397 | -0.174411853 | 0.8615418 | 0.94403207 |
| OXTR | 60.37476266 | 0.211875817 | 0.563985058 | 0.375676294 | 0.707157563 | 0.875135606 |
| P2RX3 | 22.26547269 | 0.127733603 | 1.26431887 | 0.101029579 | 0.91952698 | 0.969677217 |
| P2RX4 | 412.7026356 | -0.886562839 | 0.364856257 | -2.429896217 | 0.015103147 | 0.139261407 |
| P2RX5 | 11.3084159 | -0.089812489 | 0.950288153 | -0.094510795 | 0.924703407 | 0.971780852 |
| P2RX5-TAX1BP3 | 23.86269294 | 1.197045419 | 0.795856904 | 1.504096292 | 0.132556575 | 0.418641974 |
| P2RX6 | 43.55664513 | 0.177757596 | 0.593207194 | 0.299655158 | 0.764440206 | 0.902796386 |
| P2RX7 | 11.37103601 | 0.802899008 | 0.907629327 | 0.884611133 | 0.3763664 | 0.680079109 |
| P2RY1 | 151.2378462 | 0.474926026 | 0.526574808 | 0.901915586 | 0.367101711 | 0.673678488 |
| P2RY11 | 66.39097574 | 0.351568676 | 0.49466928 | 0.710714592 | 0.477261116 | 0.74992526 |
| P2RY2 | 60.52439688 | -2.286519871 | 0.684321983 | -3.341292443 | 0.000833893 | 0.02432884 |
| P2RY6 | 159.4538797 | -3.927811923 | 1.092817606 | -3.594206299 | 0.000325382 | 0.013341539 |
| P3H1 | 1262.217901 | 0.438774137 | 0.302001015 | 1.45288961 | 0.146254406 | 0.438763218 |
| P3H2 | 438.4776558 | -1.89207706 | 0.562152439 | -3.365772214 | 0.000763297 | 0.02306769 |
| P3H3 | 1106.349974 | 0.835642051 | 0.395944219 | 2.110504485 | 0.034814925 | 0.213146583 |
| P3H4 | 808.9336787 | 0.78046894 | 0.263528629 | 2.961609691 | 0.003060355 | 0.054934156 |
| P4HA1 | 1071.598932 | 0.834071393 | 0.243766684 | 3.421597156 | 0.000622545 | 0.020311279 |
| P4HA2 | 168.6312818 | -0.791859243 | 0.474876528 | -1.667505544 | 0.095413923 | 0.354701001 |
| P4HB | 8807.641666 | -0.512620139 | 0.322642984 | -1.588815391 | 0.112102077 | 0.384920166 |
| P4HTM | 511.9212098 | 0.155633506 | 0.275667969 | 0.564568697 | 0.572367162 | 0.807456036 |
| PA2G4 | 6632.059971 | 0.335465623 | 0.217812593 | 1.540157156 | 0.123522051 | 0.40487218 |
| PAAF1 | 690.3037298 | 0.086516805 | 0.300623187 | 0.287791522 | 0.773506328 | 0.906158747 |
| PABPC1 | 47616.30013 | -0.19685786 | 0.32037058 | -0.614469219 | 0.538905308 | 0.786598616 |
| PABPC1L | 765.5526933 | -0.42558183 | 0.298597301 | -1.425270184 | 0.154079115 | 0.451087275 |
| PABPC1L2A | 11.93470884 | 0.15473409 | 0.905275237 | 0.170924912 | 0.864282808 | 0.945136254 |
| PABPC1P4 | 13.92966881 | -1.000408687 | 0.883679253 | -1.1320948 | 0.257594581 | 0.57017324 |
| PABPC4 | 5158.34001 | -0.064902515 | 0.24359922 | -0.266431536 | 0.78990688 | 0.912949942 |
| PABPC4L | 180.9169283 | -0.306870802 | 0.367008425 | -0.836141027 | 0.40307557 | 0.699766232 |
| PABPC5 | 84.94119107 | 0.024022283 | 0.455791427 | 0.052704553 | 0.957967312 | 0.98520931 |
| PABPN1 | 10061.24713 | 0.476202184 | 0.330841859 | 1.439364976 | 0.150047142 | 0.444740017 |
| PACRG | 29.90815636 | -0.126034827 | 0.606992078 | -0.207638339 | 0.835511367 | 0.932531417 |

| | | | | | | |
|---|---|---|---|---|---|---|
| PACRGL | 474.9645117 | 0.645415821 | 0.268761246 | 2.401446746 | 0.016330386 | 0.144375457 |
| PACS1 | 827.990435 | -1.205550258 | 0.385570115 | -3.126669338 | 0.001767987 | 0.039034758 |
| PACS2 | 860.0435761 | -0.175622182 | 0.406348849 | -0.432195593 | 0.665599262 | 0.855161526 |
| PACSIN1 | 220.3551984 | -1.695126328 | 0.367602398 | -4.611303782 | 4.00E-06 | 0.000552393 |
| PACSIN2 | 1788.823862 | -0.737121068 | 0.386518919 | -1.907076296 | 0.056510708 | 0.275406162 |
| PACSIN3 | 663.9458606 | 0.497822054 | 0.460816441 | 1.080304456 | 0.280006626 | 0.593790959 |
| PADI2 | 294.9373788 | -1.030137752 | 0.367400731 | -2.803853299 | 0.005049588 | 0.075698905 |
| PADI3 | 30.14535703 | 0.620988058 | 0.740546066 | 0.83855426 | 0.401719487 | 0.699116854 |
| PAF1 | 997.0004268 | -0.364894301 | 0.360471452 | -1.012269625 | 0.311409154 | 0.621452782 |
| PAFAH1B1 | 4337.462206 | -0.015390888 | 0.225919405 | -0.068125567 | 0.945685678 | 0.979919228 |
| PAFAH1B2 | 3378.121718 | 0.564971068 | 0.390435699 | 1.447027179 | 0.147889312 | 0.44174616 |
| PAFAH1B3 | 1112.153041 | 0.731864959 | 0.372390788 | 1.965314347 | 0.049377866 | 0.25614346 |
| PAFAH2 | 499.7764182 | -0.108170762 | 0.35054366 | -0.308579999 | 0.757641037 | 0.898965445 |
| PAGR1 | 1264.17711 | 0.898421434 | 0.321149674 | 2.797516256 | 0.005149718 | 0.076234566 |
| PAH | 76.3613151 | -1.138955651 | 0.632942649 | -1.799461061 | 0.071945778 | 0.308679129 |
| PAICS | 14216.04769 | 0.454792727 | 0.256251936 | 1.774787475 | 0.075932985 | 0.316130438 |
| PAIP1 | 3692.928209 | 0.137357098 | 0.329282104 | 0.417141095 | 0.676575208 | 0.860199564 |
| PAIP2 | 3233.078636 | 0.727354681 | 0.303107491 | 2.399659203 | 0.016410342 | 0.144574776 |
| PAIP2B | 688.8952684 | -0.1216179 | 0.267103202 | -0.45532176 | 0.648877774 | 0.847033866 |
| PAK1 | 2806.492123 | -0.494538063 | 0.2779659 | -1.779132127 | 0.075218103 | 0.314600339 |
| PAK1IP1 | 440.2578985 | 0.677508975 | 0.300420203 | 2.255204441 | 0.024120499 | 0.177464644 |
| PAK2 | 2567.521407 | -0.205851207 | 0.294794296 | -0.698287618 | 0.484997341 | 0.755353112 |
| PAK3 | 115.011641 | -0.021840957 | 0.504165302 | -0.043321023 | 0.965445633 | 0.988700292 |
| PAK4 | 1381.648637 | 0.190796672 | 0.342368169 | 0.557285078 | 0.577332669 | 0.809901794 |
| PAK5 | 31.54879712 | 0.873853019 | 0.826748824 | 1.056975218 | 0.290522898 | 0.603522235 |
| PAK6 | 417.1039015 | -4.371688177 | 0.873909754 | -5.002448088 | 5.66E-07 | 0.000128176 |
| PALB2 | 689.043178 | -0.114379713 | 0.305162464 | -0.374815801 | 0.707797462 | 0.875570393 |
| PALD1 | 378.635569 | 0.859423687 | 0.427261323 | 2.011470827 | 0.044275749 | 0.2418127 |
| PALLD | 3504.651451 | -1.019419287 | 0.395968205 | -2.574497838 | 0.010038571 | 0.114059016 |
| PALM | 795.9202517 | -0.156188405 | 0.441563233 | -0.353716961 | 0.723551011 | 0.883317967 |
| PALM2 | 44.89518123 | 0.432838089 | 0.640895458 | 0.675364576 | 0.499444166 | 0.76281557 |
| PALM2-AKAP2 | 131.8137805 | 0.054537142 | 2.313814045 | 0.023570236 | 0.981195414 | 0.993241851 |
| PALM3 | 388.3976906 | 0.075180981 | 0.432729922 | 0.173736497 | 0.862072553 | 0.944228049 |
| PAM | 2628.659432 | 0.061803383 | 0.298998154 | 0.206701553 | 0.836242944 | 0.93270414 |
| PAM16 | 202.3251491 | 0.906581259 | 0.456919173 | 1.984117353 | 0.047242759 | 0.250276443 |
| PAMR1 | 881.1668959 | 0.553504328 | 0.592204151 | 0.934651214 | 0.349968076 | 0.656612131 |
| PAN2 | 1521.979917 | -0.289767118 | 0.402165883 | -0.720516409 | 0.4712071 | 0.74605332 |
| PAN3 | 1303.231292 | -0.356967793 | 0.230946778 | -1.545671242 | 0.122183986 | 0.40285432 |
| PANK1 | 887.292587 | 0.368321014 | 0.280256334 | 1.314229043 | 0.188769138 | 0.493962743 |

PANK2 979.0533854 0.326597527 0.358846202 0.910132324 0.36275273 0.66991134
PANK3 3798.928779 0.102256141 0.377098829 0.271165364 0.786263851 0.911261183
PANK4 114.6015045 0.526300613 1.164172534 0.452081283 0.651210427 0.847876685
PANX1 535.0218435 -0.061321687 0.357222953 -0.171662227 0.863703084 0.94507782
PANX2 16.94317423 -0.647971254 0.820970487 -0.789274723 0.429951457 0.716554273
PAOX 98.82885199 0.119462226 0.415643313 0.287415249 0.773794386 0.90622469
PAPLN 729.8471431 2.839294833 1.72434608 1.646592217 0.099641891 0.362001501
PAPOLA 8664.482524 -0.136745846 0.223697331 -0.611298514 0.541001974 0.788008477
PAPOLG 921.4480505 -0.027966532 0.265713471 -0.105250712 0.916176872 0.969184139
PAPPA 89.14927078 -0.122113034 0.821302259 -0.148682209 0.8818044 0.953217539
PAPPA2 82.36350335 -2.904521643 1.007555993 -2.882739681 0.003942331 0.064934636
PAPSS1 3488.48074 -0.008584154 0.314922908 -0.027257954 0.978253992 0.992395125
PAPSS2 142.2107807 -0.406218272 0.359456921 -1.130088889 0.258438771 0.57129165
PAQR3 1800.70056 -0.005517816 0.257003404 -0.021469817 0.982870881 0.993545718
PAQR4 350.8855882 0.365421001 0.331520414 1.102257917 0.270349559 0.583170438
PAQR5 68.50624548 -0.529480075 0.630666063 -0.839556947 0.401156847 0.698568129
PAQR6 474.9254544 0.421183306 0.379844598 1.108830582 0.267503275 0.580717148
PAQR7 366.0494651 -0.731498856 0.411911172 -1.77586554 0.075755082 0.315769427
PAQR8 525.9500382 -0.236228927 0.546300676 -0.432415587 0.665439392 0.855161526
PAQR9 56.22643491 0.563303542 0.493381443 1.141720163 0.253570357 0.565942797
PARD3 1481.711636 -0.07914507 0.336921871 -0.234906299 0.81428146 0.92237594
PARD3B 192.007136 -0.891541526 0.484087341 -1.8416956 0.065519688 0.295305761
PARD6A 51.89609312 1.027021992 0.484661908 2.119048298 0.034086385 0.210632641
PARD6B 538.9085788 0.19833979 0.431382495 0.459777094 0.645676226 0.845794147
PARD6G 597.3641538 0.806444008 0.316495615 2.54804165 0.010832954 0.118235065
PARG 1266.095142 -0.137457074 0.286310868 -0.480097296 0.631158211 0.838320891
PARGP1 144.2526962 -0.425032674 0.76442262 -0.556017919 0.578198605 0.810209453
PARK7 5350.161688 0.646703392 0.320414497 2.01833375 0.043556514 0.240202063
PARL 1445.562811 0.329538335 0.323487296 1.018705644 0.308342732 0.617978528
PARM1 75.31482555 2.331292579 0.931893742 2.501672106 0.012360835 0.126450589
PARN 1401.786953 -0.592948285 0.253847798 -2.33584175 0.019499491 0.158280778
PARP1 15776.07008 -0.241333473 0.210721554 -1.145271894 0.252096545 0.564998953
PARP10 24.81286957 0.033313067 0.781492883 0.042627474 0.965998494 0.988700292
PARP11 514.1496965 0.203070745 0.248645272 0.816708652 0.414094946 0.706345224
PARP12 75.80574796 0.013955093 0.459711833 0.030356175 0.975782996 0.991600498
PARP14 396.343158 -0.684225043 0.357787767 -1.912376851 0.055827875 0.273390798
PARP15 24.25691009 -0.727752284 0.661445811 -1.100244754 0.271225496 0.583405926
PARP16 187.1624517 0.710561038 0.317489777 2.238059586 0.02521717

0.181657019
PARP2 1221.477606 0.087544333 0.233966534 0.374174593 0.708274425 0.875585221
PARP3 32.4380562 -0.832586953 0.631813036 -1.317774255 0.187579229
0.492618714
PARP4 1665.998526 -0.676323976 0.340846276 -1.984249276 0.047228058
0.250276443
PARP6 1019.398633 -0.850527276 0.382172406 -2.225506768 0.026047237
0.184166686
PARP8 677.7372434 -0.734279166 0.304518203 -2.411281689 0.015896565
0.142656219
PARP9 116.2099068 -0.74621068 0.350941262 -2.126312178 0.033477275
0.208789229
PARPBP 844.9398337 0.32923023 0.304371474 1.081672423 0.279398111
0.593066767
PARS2 281.8995078 0.105191354 0.302862923 0.347323313 0.728348439 0.885679973
PARVA 752.4710484 -0.803715439 0.425998071 -1.886664506 0.059205465
0.282188534
PARVB 142.1908404 0.8126808 0.388708246 2.090721788 0.036553011 0.218932905
PASK 1156.283329 -0.208355475 0.313155817 -0.665341223 0.50583227
0.767164369
PATJ 2293.60473 -1.983872142 0.46501374 -4.266265641 1.99E-05
0.001857856
PATL1 1210.883075 -0.516769085 0.338795479 -1.525312813 0.127181093
0.410771662
PATZ1 2679.317221 0.160245516 0.244444728 0.655549077 0.512114299 0.771681172
PAWR 3382.167869 0.185417006 0.274954229 0.674355899 0.500085074 0.763360896
PAX2 20.89637142 0.83710244 0.841238588 0.995083264 0.31969577 0.629465081
PAX3 1062.537726 0.369757736 0.525804204 0.70322324 0.481916638 0.753184024
PAX6 158.7027857 3.076251933 0.93350391 3.295381946 0.00098288 0.026915652
PAX7 367.5241034 -0.467854815 0.579045495 -0.807975916 0.419104446
0.709924862
PAXBP1 2558.409186 0.07291267 0.29596169 0.246358473 0.805404749
0.918913699
PAXIP1 908.9622048 -0.098315896 0.299321188 -0.328462868
0.742561716 0.892270716
PAXX 335.817965 0.475983501 0.554314651 0.858688292 0.390512511 0.690466555
PBDC1 933.9357122 0.573016659 0.432426459 1.325119327 0.185131577 0.489735162
PBK 425.5380288 1.440503715 0.292332166 4.92762645 8.32E-07 0.000165643
PBLD 80.49946273 -0.641053038 0.415898926 -1.54136738 0.123227398
0.404737576
PBRM1 2411.425269 -0.337333463 0.315926661 -1.067758771 0.28562934
0.599513381
PBX1 4634.073431 -0.600909889 0.330365381 -1.818925115 0.068922856
0.302794854
PBX2 3494.064893 -0.300873086 0.30160358 -0.997577967 0.31848405
0.628475744
PBX3 2453.890845 0.105719078 0.373809495 0.282815391 0.777318356 0.907898743
PBX4 47.27713463 0.478394991 0.563003135 0.849719942 0.395480809 0.69426198
PBXIP1 1662.104983 -0.434422515 0.324617111 -1.338261296
0.180811274 0.485502272
PC 539.55722 0.003352055 0.477257212 0.007023581 0.994396039 0.997816146
PCBD1 690.9399434 0.292732319 0.394264057 0.742477824 0.457797883 0.736506011
PCBD2 92.25464272 -0.164833685 0.453781363 -0.363244722 0.716422082
0.879925471
PCBP1 8537.835041 0.630676336 0.341667231 1.845878913 0.064909789 0.293855407
PCBP2 24828.68985 0.059595877 0.300584331 0.198266745 0.842836369 0.935861125
PCBP3 67.64423655 0.166483205 0.467533861 0.356088016 0.721774652 0.882498649
PCBP4 1995.571244 0.751740225 0.498774725 1.507173856 0.1317661 0.417506996
PCCA 734.4248677 -0.202897086 0.315494139 -0.643108891 0.520153446
0.775247856
PCCB 1573.372202 0.323934862 0.259026202 1.250587234 0.211085111 0.518518817
PCDH1 686.4279692 -1.366369796 0.27310972 -5.003006828 5.64E-07
0.000128176
PCDH10 154.5520265 -1.560683807 0.599464931 -2.603461397

0.009228766 0.108795292
PCDH11X 331.8876728 -0.982376372 0.536963987 -1.82950141 0.067324528
0.298083457
PCDH11Y 73.1482771 -0.372809329 0.784642466 -0.475132746
0.63469236 0.840313679
PCDH17 37.48479625 -0.534961854 0.791643465 -0.675761093
0.499192341 0.76281557
PCDH18 4653.925066 1.451569798 0.612146186 2.371279658 0.017726613
0.150195643
PCDH19 40.70076978 -0.875582591 0.682085064 -1.28368533 0.199252079
0.505071561
PCDH20 30.29784693 -0.491032339 1.067799167 -0.459854582
0.645620602 0.845794147
PCDH7 41.10272819 -1.125023149 0.650781488 -1.728726417 0.083858071
0.331304895
PCDH8 681.5160631 1.336070416 0.76395223 1.748892617 0.080309583 0.324292679
PCDH9 91.86602923 -1.544878265 0.799840452 -1.931483037 0.053423345
0.26722241
PCDHA1 69.35014074 1.713373789 0.911912224 1.878880164 0.060260858
0.283904551
PCDHA10 967.410263 -0.526578647 0.310447348 -1.696193091
0.089849318 0.34317825
PCDHA11 245.6409691 -0.553645894 0.691156615 -0.801042604
0.423106982 0.712504832
PCDHA12 238.6591149 -0.891006063 0.485416722 -1.83554876 0.066424421
0.297462837
PCDHA13 12.36584452 -1.614237457 1.181472236 -1.366293179
0.171846961 0.474879827
PCDHA2 96.70971846 -0.392981653 0.499570989 -0.786638259
0.431493656 0.718177757
PCDHA3 137.6097292 0.847169386 0.443733145 1.909186624 0.056238021
0.274778428
PCDHA4 366.6550758 -0.115387375 0.348447596 -0.331146996
0.740533454 0.891917214
PCDHA5 19.9861032 -0.707960525 0.794694436 -0.890858792
0.37300493 0.67811865
PCDHA6 11.49807909 0.987126591 1.021538107 0.966314017 0.333887074
0.64183994
PCDHA7 52.64011036 1.197716729 0.861993213 1.389473503 0.164688813
0.46700822
PCDHA9 30.98703963 0.732915003 0.686873918 1.067029892 0.285958338
0.599871951
PCDHAC1 149.043331 -1.629082363 0.640909541 -2.541828851
0.011027416 0.119327334
PCDHAC2 109.2462412 -0.492899734 2.08959412 -0.235883002
0.813523459 0.921959809
PCDHB1 11.62734346 1.129671435 0.848857023 1.330814736 0.183249978
0.487306821
PCDHB10 210.9014934 0.007220316 0.317768762 0.022721918 0.981872093
0.993518428
PCDHB11 68.93884179 -0.895375595 0.439198601 -2.038657666
0.041484206 0.233961666
PCDHB12 148.1460753 0.526741057 0.365146249 1.442548181 0.14914779
0.443844865
PCDHB13 211.9331383 -1.243018637 0.358104627 -3.471104655
0.000518322 0.017912215
PCDHB14 188.1878937 -0.507530952 0.373488498 -1.358893124
0.174180458 0.478455902
PCDHB15 592.5424443 -0.551835209 0.275870245 -2.000343345
0.045463202 0.245452751
PCDHB16 488.7855386 -0.19827659 0.296454507 -0.668826362 0.503606248
0.765398757
PCDHB17P 124.6693656 0.814136826 0.412745848 1.972489437 0.048553761
0.25374113

| | | | | | | |
|---|---|---|---|---|---|---|
| PCDHB18P | 16.78297942 | -0.209292496 | 0.79474514 | -0.263345424 | 0.792284351 | 0.913570955 |
| PCDHB2 | 1116.893687 | -0.147243835 | 0.29363313 | -0.501455117 | 0.616050858 | 0.831265761 |
| PCDHB3 | 85.69226033 | -0.066963811 | 0.4313106 | -0.155256586 | 0.87661904 | 0.950324661 |
| PCDHB4 | 32.39168734 | 0.19511336 | 0.615826744 | 0.31683158 | 0.751371401 | 0.89636357 |
| PCDHB5 | 1611.047972 | 0.009752739 | 0.289783724 | 0.033655235 | 0.973152076 | 0.990423175 |
| PCDHB6 | 31.26931473 | 0.373221652 | 0.657861209 | 0.567325822 | 0.570492836 | 0.806522093 |
| PCDHB7 | 34.45550911 | -0.602044032 | 0.560897443 | -1.073358489 | 0.283110302 | 0.596961028 |
| PCDHB8 | 233.5035139 | -2.625169388 | 0.379342205 | -6.920319836 | 4.51E-12 | 1.37E-08 |
| PCDHB9 | 129.0567814 | 0.135010419 | 0.41020275 | 0.329130945 | 0.742056715 | 0.892150449 |
| PCDHGA1 | 101.6042259 | -0.49789141 | 0.412531645 | -1.206916889 | 0.227464153 | 0.536263973 |
| PCDHGA10 | 33.42986859 | -0.765892307 | 0.670909693 | -1.141572874 | 0.253631605 | 0.565942797 |
| PCDHGA11 | 648.4986406 | 0.044745828 | 1.445939393 | 0.030945853 | 0.975312722 | 0.991429622 |
| PCDHGA12 | 13.32279193 | -2.265232611 | 1.03520859 | -2.188189542 | 0.028655799 | 0.193474464 |
| PCDHGA2 | 32.53092701 | -0.53690079 | 0.627932645 | -0.85502927 | 0.392534944 | 0.691717122 |
| PCDHGA4 | 39.60237118 | -0.699214138 | 0.690654548 | -1.012393447 | 0.31134997 | 0.621452782 |
| PCDHGA5 | 40.11574069 | -0.509120539 | 0.598297391 | -0.850948953 | 0.39479771 | 0.693708137 |
| PCDHGA6 | 102.3700575 | -0.386465671 | 0.49620254 | -0.77884662 | 0.436070071 | 0.720784523 |
| PCDHGA7 | 18.81442138 | 0.260856591 | 0.901967494 | 0.289208417 | 0.772421891 | 0.905717252 |
| PCDHGA8 | 56.65044974 | 0.185292679 | 0.477804272 | 0.387800381 | 0.698163764 | 0.870467 |
| PCDHGA9 | 16.88178832 | -1.270272229 | 0.964428034 | -1.317124953 | 0.187796744 | 0.492689771 |
| PCDHGB1 | 94.97815934 | -1.224289277 | 0.890258657 | -1.375206259 | 0.169067508 | 0.471579918 |
| PCDHGB2 | 45.48320623 | -1.090874241 | 0.663657601 | -1.643730503 | 0.100231882 | 0.36243601 |
| PCDHGB3 | 37.11727823 | -1.281999517 | 0.984846962 | -1.301724598 | 0.193010548 | 0.498908723 |
| PCDHGB4 | 506.3770856 | -0.639318134 | 0.520595189 | -1.228052329 | 0.219427325 | 0.527925634 |
| PCDHGB5 | 292.1665553 | -1.273096502 | 0.883569913 | -1.440855425 | 0.149625531 | 0.444386222 |
| PCDHGB6 | 300.2349543 | -0.724274226 | 0.590793852 | -1.225933925 | 0.220223541 | 0.528727493 |
| PCDHGB7 | 604.3842014 | -0.693636414 | 1.630435089 | -0.425430254 | 0.670523055 | 0.858101085 |
| PCDHGB8P | 11.76729486 | -1.379965429 | 1.029432423 | -1.340510944 | 0.180079289 | 0.484754911 |
| PCDHGC3 | 1348.335579 | 0.261572109 | 0.325561309 | 0.803449618 | 0.421714906 | 0.712057523 |
| PCDHGC4 | 194.5284682 | 0.800761292 | 1.597117991 | 0.501378919 | 0.616104473 | 0.831265761 |
| PCDHGC5 | 38.33520982 | 0.229144691 | 1.305507886 | 0.175521492 | 0.860669885 | 0.943645503 |
| PCED1A | 1048.188613 | 0.40922841 | 0.271995339 | 1.504541998 | 0.132441868 | 0.418524946 |
| PCED1B | 17.3591624 | 0.748254343 | 0.871949314 | 0.858139724 | 0.390815315 | |

| | | | | | | |
|---|---|---|---|---|---|---|
| | | | | | | 0.690469214 |
| PCF11 | 1693.165089 | -0.238264223 | 0.375424277 | -0.634653211 | 0.525654616 | 0.778399822 |
| PCGF1 | 271.7705188 | 0.457678201 | 0.342413606 | 1.336623878 | 0.181345441 | 0.485749663 |
| PCGF2 | 365.0237206 | -0.496149342 | 0.639753435 | -0.775532126 | 0.43802529 | 0.721949744 |
| PCGF3 | 1901.693833 | -0.281166117 | 0.358357387 | -0.78459696 | 0.432689913 | 0.718639059 |
| PCGF5 | 144.8333461 | 0.235624783 | 0.362059938 | 0.650789434 | 0.515182422 | 0.773232343 |
| PCGF6 | 646.5215522 | 0.455247914 | 0.35118754 | 1.296309983 | 0.194868711 | 0.500501471 |
| PCID2 | 1583.55189 | 0.465460108 | 0.240906353 | 1.932120515 | 0.053344633 | 0.267056023 |
| PCIF1 | 843.7764371 | -0.202012271 | 0.266275417 | -0.758659111 | 0.448056502 | 0.729463615 |
| PCK2 | 496.6674618 | 0.409640703 | 0.276736125 | 1.480257422 | 0.138804561 | 0.427314124 |
| PCLAF | 2269.31411 | 1.377450118 | 0.352448213 | 3.908234079 | 9.30E-05 | 0.005828498 |
| PCLO | 83.29902655 | -0.262565105 | 0.610020724 | -0.430419976 | 0.666890167 | 0.856258128 |
| PCM1 | 4440.873717 | -0.043041243 | 0.252212404 | -0.170654742 | 0.864495251 | 0.94517566 |
| PCMT1 | 2439.270078 | 0.443405367 | 0.342051783 | 1.296310644 | 0.194868483 | 0.500501471 |
| PCMTD1 | 1242.502306 | -0.135097977 | 0.259485663 | -0.520637537 | 0.602619295 | 0.825021776 |
| PCMTD2 | 1647.799203 | -0.145810004 | 0.316971249 | -0.46001019 | 0.645508906 | 0.845794147 |
| PCNA | 4117.186386 | 0.972235244 | 0.32785228 | 2.965467392 | 0.003022234 | 0.054402443 |
| PCNP | 5087.106686 | 0.33592756 | 0.318708988 | 1.054026001 | 0.291871013 | 0.604997287 |
| PCNT | 1048.161435 | -0.230581136 | 0.402552082 | -0.57279827 | 0.566781291 | 0.804513376 |
| PCNX1 | 1764.567478 | -0.180531096 | 0.387097149 | -0.466371546 | 0.640949575 | 0.843790871 |
| PCNX2 | 715.2728738 | -0.508939726 | 0.337170281 | -1.5094442 | 0.131185305 | 0.416813082 |
| PCNX3 | 910.4678716 | -0.51223509 | 0.36204032 | -1.414856471 | 0.157110583 | 0.456439037 |
| PCNX4 | 1695.5268 | 0.009293684 | 0.358490231 | 0.025924512 | 0.979317549 | 0.992858175 |
| PCOLCE | 420.8258546 | 0.597858029 | 0.342684221 | 1.744632503 | 0.081048859 | 0.325604941 |
| PCOLCE2 | 735.0160768 | -0.191713783 | 0.45947804 | -0.417242536 | 0.676501015 | 0.860199564 |
| PCP4L1 | 120.8903498 | -0.964753178 | 0.800608045 | -1.205025586 | 0.228193425 | 0.537150955 |
| PCSK1 | 15.36325925 | 0.783680457 | 0.91605316 | 0.855496702 | 0.39227623 | 0.691514794 |
| PCSK2 | 63.43546896 | 0.169741343 | 0.583479707 | 0.290912162 | 0.7711185 | 0.905067213 |
| PCSK4 | 63.48513631 | -0.408421451 | 0.712726818 | -0.573040667 | 0.566617159 | 0.804513376 |
| PCSK5 | 378.5970611 | 0.235759861 | 0.474324549 | 0.497043347 | 0.619158484 | 0.832513814 |
| PCSK6 | 170.970832 | 0.196579025 | 0.463204836 | 0.424388975 | 0.67128216 | 0.858479663 |
| PCSK7 | 1445.309228 | -0.449188618 | 0.240526698 | -1.867520825 | 0.061828891 | 0.286447296 |
| PCSK9 | 145.8559895 | 1.378710409 | 0.491897234 | 2.80284237 | 0.005065442 | 0.07586162 |
| PCTP | 771.7277985 | 0.223496378 | 0.257949943 | 0.866433137 | 0.386252679 | 0.686679504 |
| PCYOX1 | 2055.955254 | -0.38641835 | 0.252772625 | -1.528719142 | 0.126334085 | 0.409095923 |
| PCYOX1L | 848.6297306 | 0.135926108 | 0.302133081 | 0.4498882 | 0.652791056 | 0.848628373 |
| PCYT1A | 872.8963286 | -0.220863466 | 0.28874449 | -0.764909717 | 0.444325307 | 0.727151225 |
| PCYT1B | 821.396476 | 0.041876622 | 0.311300256 | 0.134521644 | 0.892990097 | 0.957627254 |
| PCYT2 | 658.81944 | -0.596907507 | 0.338472871 | -1.763531314 | 0.077810921 | 0.319997148 |
| PDAP1 | 3381.504998 | -0.064930351 | 0.237506621 | -0.273383331 | 0.784558565 | 0.910931322 |
| PDCD10 | 1486.441623 | 0.377597029 | 0.25907536 | 1.457479512 | 0.144984063 | |

0.436332715

| | | | | | | |
|---|---|---|---|---|---|---|
| PDCD11 | 2802.632056 | -0.26966106 | 0.366884673 | -0.73500225 | 0.462338124 | 0.73986733 |
| PDCD2 | 3950.246438 | 0.480090606 | 0.550768009 | 0.871674821 | 0.3833858 | 0.68451387 |
| PDCD2L | 363.0591519 | 0.885447767 | 0.427570054 | 2.070883495 | 0.038369685 | 0.225585886 |
| PDCD4 | 1200.542553 | -0.22126182 | 0.365277719 | -0.605735879 | 0.544690155 | 0.790739196 |
| PDCD5 | 1227.889454 | 1.085105782 | 0.413325951 | 2.625302811 | 0.008657194 | 0.104918628 |
| PDCD6 | 1523.631752 | 0.377248623 | 0.381301043 | 0.989372124 | 0.322481108 | 0.632222913 |
| PDCD6IP | 3201.882284 | -0.432366841 | 0.252272273 | -1.713889665 | 0.086548992 | 0.337492171 |
| PDCD6IPP2 | 49.5640753 | -0.090292912 | 0.484616787 | -0.186318168 | 0.852195261 | 0.939923977 |
| PDCD7 | 848.2246097 | 0.311650239 | 0.283168292 | 1.100583109 | 0.27107814 | 0.583394591 |
| PDCL | 930.3290728 | 0.110373578 | 0.265338475 | 0.41597276 | 0.677429936 | 0.860419848 |
| PDCL3 | 561.1920264 | 0.76286842 | 0.401932509 | 1.898001288 | 0.057695912 | 0.278227809 |
| PDE10A | 85.43995204 | 0.897025452 | 0.674873954 | 1.3291748 | 0.183790307 | 0.488175154 |
| PDE11A | 180.3269396 | -0.208555881 | 0.799661474 | -0.260805212 | 0.794242725 | 0.914628224 |
| PDE12 | 401.7330294 | 0.132302823 | 0.319098366 | 0.414614543 | 0.6784241 | 0.861138002 |
| PDE1B | 55.97001396 | 1.195711427 | 0.531585291 | 2.249331288 | 0.024491427 | 0.178720266 |
| PDE1C | 16.22909768 | -1.050730054 | 0.889018095 | -1.181899513 | 0.237245575 | 0.547879744 |
| PDE2A | 29.33160405 | 0.247159207 | 0.718214575 | 0.344130036 | 0.730748496 | 0.886894834 |
| PDE3A | 132.2883203 | -1.226537801 | 0.749558259 | -1.636347525 | 0.101766875 | 0.365921972 |
| PDE3B | 604.2255085 | 0.377392107 | 0.405490543 | 0.930705075 | 0.352006144 | 0.65929447 |
| PDE4B | 25.93895666 | -0.597124432 | 0.76762321 | -0.777887412 | 0.43663539 | 0.720784523 |
| PDE4C | 22.28990165 | -1.350642359 | 0.855924058 | -1.577993219 | 0.114567168 | 0.389533505 |
| PDE4D | 150.4783589 | -1.004926019 | 0.611356828 | -1.643763468 | 0.10022507 | 0.36243601 |
| PDE4DIP | 1400.638038 | -0.774908153 | 0.225117925 | -3.442232129 | 0.000576935 | 0.019194474 |
| PDE5A | 648.4936924 | -0.722013337 | 0.265850754 | -2.715859659 | 0.006610393 | 0.089064189 |
| PDE6A | 11.464407 | -1.075023424 | 1.03320666 | -1.040472797 | 0.298120296 | 0.609375237 |
| PDE6B | 204.9443986 | -0.197139026 | 0.416876769 | -0.472895206 | 0.636287945 | 0.84124553 |
| PDE6D | 496.7334475 | 0.635499545 | 0.318364123 | 1.996140581 | 0.045918623 | 0.246869858 |
| PDE7A | 3624.771289 | -0.083730479 | 0.220524167 | -0.379688449 | 0.704176696 | 0.873421952 |
| PDE8A | 206.7897263 | -0.695972014 | 0.369112007 | -1.885530682 | 0.059358226 | 0.282486927 |
| PDE9A | 493.9525291 | 0.559604028 | 0.36041161 | 1.552680358 | 0.120499517 | 0.400810825 |
| PDF | 161.5340174 | 0.40131233 | 0.584187573 | 0.686958006 | 0.492109192 | 0.759174768 |
| PDGFA | 841.3408319 | -0.250973052 | 0.540181041 | -0.464609147 | 0.642211381 | 0.844148268 |
| PDGFB | 53.97473885 | -1.777560875 | 0.87617012 | -2.02878509 | 0.042480185 | 0.237110699 |
| PDGFC | 286.3748028 | -1.372692579 | 0.555979038 | -2.468964628 | 0.013550461 | 0.132400677 |
| PDGFD | 93.50521906 | -1.270995767 | 0.41048683 | -3.096313139 | 0.001959433 | 0.041483956 |
| PDGFRA | 167.9022061 | -1.037578358 | 0.426894999 | -2.430523575 | 0.015077025 | 0.139217011 |
| PDGFRB | 245.8446181 | 0.717139518 | 0.591981892 | 1.211421375 | 0.225733951 | 0.53459368 |
| PDGFRL | 142.0063869 | -0.817915893 | 0.418158239 | -1.955996119 | 0.050465611 | 0.258046297 |

| | | | | | | |
|---|---|---|---|---|---|---|
| PDHA1 | 2691.011615 | 0.219927248 | 0.283842208 | 0.774822214 | 0.438444721 | 0.722219855 |
| PDHB | 3851.513283 | 0.566166474 | 0.332966478 | 1.70037079 | 0.089061203 | 0.342496199 |
| PDHX | 1004.210932 | 0.267741792 | 0.271876788 | 0.98479092 | 0.324726782 | 0.634685649 |
| PDIA3 | 13995.03871 | 0.088421675 | 0.299178363 | 0.29554836 | 0.767575015 | 0.903824942 |
| PDIA3P1 | 120.6850759 | -1.193561227 | 0.425280313 | -2.806528284 | 0.005007852 | 0.07561376 |
| PDIA4 | 5783.641349 | -0.100511401 | 0.268017367 | -0.375018238 | 0.707646903 | 0.87545553 |
| PDIA5 | 714.152762 | -0.07292103 | 0.370929145 | -0.196590188 | 0.844148249 | 0.936837827 |
| PDIA6 | 9587.569967 | 0.256729369 | 0.271019215 | 0.947273678 | 0.34349934 | 0.650427919 |
| PDIK1L | 311.8856744 | -0.00957894 | 0.392328697 | -0.024415599 | 0.980521106 | 0.993221987 |
| PDK1 | 279.6258125 | -0.247850338 | 0.411362616 | -0.602510603 | 0.546834308 | 0.792249935 |
| PDK2 | 492.3164639 | -0.17676576 | 0.321472047 | -0.549863546 | 0.582412969 | 0.813383702 |
| PDK3 | 450.6358895 | 0.152858266 | 0.358625821 | 0.426233296 | 0.669937858 | 0.857762448 |
| PDLIM1 | 7469.360462 | -0.598930684 | 0.386278499 | -1.550515199 | 0.121017908 | 0.401219509 |
| PDLIM2 | 409.8621845 | 0.251473771 | 0.553240008 | 0.454547335 | 0.649434931 | 0.847281162 |
| PDLIM3 | 92.46560414 | 0.566467292 | 0.525746336 | 1.077453619 | 0.281277661 | 0.595171527 |
| PDLIM4 | 193.8739343 | 0.745467267 | 0.490855294 | 1.518710864 | 0.128835291 | 0.413453061 |
| PDLIM5 | 683.5585552 | -0.352158849 | 0.332000065 | -1.060719218 | 0.288817524 | 0.601911102 |
| PDLIM7 | 1403.171125 | 0.370352646 | 0.343990581 | 1.076636008 | 0.281642911 | 0.59542985 |
| PDP1 | 591.5721058 | -0.265879086 | 0.309781993 | -0.858278054 | 0.390738944 | 0.690469214 |
| PDP2 | 562.3361493 | 0.444906412 | 0.355567235 | 1.251258181 | 0.210840297 | 0.518294063 |
| PDPK1 | 1225.785818 | -0.527966648 | 0.341391611 | -1.546513242 | 0.121980664 | 0.402659864 |
| PDPN | 1725.212071 | -0.011905524 | 0.234776901 | -0.050709945 | 0.959556652 | 0.985942023 |
| PDPR | 2565.634711 | -0.52504865 | 0.338040021 | -1.553214466 | 0.120371907 | 0.400562008 |
| PDRG1 | 904.8640743 | 0.920419136 | 0.367664384 | 2.503422075 | 0.012299876 | 0.126252654 |
| PDS5A | 3334.018542 | -0.004301451 | 0.293642354 | -0.014648607 | 0.988312521 | 0.995803391 |
| PDS5B | 1730.110176 | 0.016441595 | 0.253380475 | 0.064888959 | 0.948262412 | 0.98134156 |
| PDSS1 | 316.3690737 | 0.813476533 | 0.295315058 | 2.754605673 | 0.005876291 | 0.082775498 |
| PDSS2 | 415.8592815 | -0.202053421 | 0.285013111 | -0.708926759 | 0.478369928 | 0.750837683 |
| PDXDC1 | 2318.717455 | -0.791559904 | 0.357331389 | -2.21519835 | 0.026746453 | 0.186818803 |
| PDXDC2P | 118.7037833 | -1.205720196 | 0.524688055 | -2.297975309 | 0.021563194 | 0.166565796 |
| PDXK | 1111.297068 | 0.226224405 | 0.282241479 | 0.801527848 | 0.422826129 | 0.712269066 |
| PDXP | 600.777582 | 1.854105424 | 0.37641691 | 4.925669852 | 8.41E-07 | 0.000165643 |
| PDYN | 145.9680473 | 0.704964911 | 0.526948598 | 1.337824816 | 0.18095355 | 0.485626449 |
| PDZD11 | 618.6238746 | 0.188011353 | 0.275781881 | 0.681739321 | 0.495403799 | 0.760859008 |
| PDZD2 | 171.9782055 | -2.302683634 | 0.613194122 | -3.755227832 | 0.000173184 | 0.008728094 |
| PDZD4 | 172.6368311 | 2.265743405 | 0.615553326 | 3.68082392 | 0.000232482 | 0.010434843 |
| PDZD7 | 78.08196768 | -1.121469777 | 0.614032164 | -1.826402332 | 0.067789681 | 0.299425477 |
| PDZD8 | 771.430366 | 0.173826419 | 0.324009424 | 0.536485689 | 0.591622924 | 0.819382087 |
| PDZRN3 | 161.5522717 | -0.470581446 | 0.598506298 | -0.786259805 | 0.431715295 | 0.718310237 |
| PDZRN4 | 20.48265277 | -1.601720915 | 0.969138906 | -1.652725843 | | |

| | | | | | | |
|---|---|---|---|---|---|---|
| | | | | | 0.098386672 | 0.360015486 |
| PEA15 | 1134.905261 | 0.12160932 | 0.408965479 | 0.297358398 | 0.766192901 | 0.903373663 |
| PEAK1 | 1535.710606 | -0.60051482 | 0.411431697 | -1.459573543 | 0.144407315 | 0.43572064 |
| PEAR1 | 71.437934 | -2.984056574 | 0.885284817 | -3.370730544 | 0.000749691 | 0.022792724 |
| PEBP1 | 6793.961499 | 0.657748515 | 0.38651946 | 1.701721604 | 0.088807568 | 0.342301733 |
| PECAM1 | 13.61804195 | -0.510825493 | 0.836314429 | -0.610805548 | 0.54132832 | 0.788298325 |
| PECR | 383.4461391 | 0.498353252 | 0.326362819 | 1.526991507 | 0.126763123 | 0.410047621 |
| PEF1 | 1480.850417 | 0.181988705 | 0.294102999 | 0.618792415 | 0.53605312 | 0.784910904 |
| PEG10 | 4710.2031 | 0.380530003 | 0.392147822 | 0.970373879 | 0.331860164 | 0.63972688 |
| PELI1 | 1314.267321 | 0.342572159 | 0.342683394 | 0.999675402 | 0.31746762 | 0.627531109 |
| PELI2 | 478.7059185 | 0.442787457 | 0.274051946 | 1.615706305 | 0.106157827 | 0.373883142 |
| PELI3 | 117.4883062 | -0.436881878 | 0.520421008 | -0.839477792 | 0.401201246 | 0.698568129 |
| PELO | 573.4517858 | 0.266833881 | 0.266693391 | 1.000526785 | 0.317055642 | 0.627198876 |
| PELP1 | 2842.829467 | 0.098762837 | 0.289118938 | 0.341599335 | 0.732652437 | 0.888122198 |
| PEMT | 372.9840537 | 0.503228483 | 0.308683411 | 1.630241422 | 0.103050483 | 0.368123388 |
| PEPD | 1370.272035 | -0.072817326 | 0.342475497 | -0.212620542 | 0.831622947 | 0.930997148 |
| PER2 | 151.8909551 | -0.206540734 | 0.469062235 | -0.440326931 | 0.659700338 | 0.852112718 |
| PER3 | 282.0238392 | 0.586054687 | 0.576875387 | 1.015912102 | 0.309671245 | 0.61915097 |
| PERP | 7321.857714 | -0.341212921 | 0.671992244 | -0.507763184 | 0.611619432 | 0.829058038 |
| PES1 | 3132.749559 | 0.081964159 | 0.268271095 | 0.305527359 | 0.759964536 | 0.899725004 |
| PET100 | 1633.994671 | 1.202604127 | 0.5343055 | 2.250779987 | 0.024399476 | 0.178492903 |
| PET117 | 405.4960886 | 0.772020366 | 0.468398866 | 1.648211433 | 0.099309292 | 0.361473431 |
| PEX1 | 1186.114344 | -0.253505632 | 0.28315609 | -0.895285821 | 0.370634318 | 0.676128642 |
| PEX10 | 550.5682438 | 0.123109479 | 0.37529006 | 0.32803821 | 0.742882772 | 0.892412269 |
| PEX11A | 159.0361733 | -0.124740395 | 0.312463145 | -0.399216345 | 0.689733801 | 0.867683896 |
| PEX11B | 513.0154916 | -0.188860317 | 0.268495351 | -0.70340256 | 0.481804912 | 0.753118606 |
| PEX11G | 35.0556364 | -0.060303086 | 0.835425176 | -0.07218251 | 0.942456664 | 0.978574897 |
| PEX12 | 180.8930876 | 0.292766095 | 0.301246752 | 0.97184814 | 0.331126103 | 0.639084337 |
| PEX13 | 864.0136691 | 0.288259611 | 0.306787755 | 0.939605984 | 0.347419706 | 0.654038526 |
| PEX14 | 557.6135527 | 0.085092676 | 0.322574772 | 0.263792098 | 0.791940124 | 0.913570955 |
| PEX16 | 647.6268101 | 0.231448416 | 0.412978335 | 0.560437186 | 0.575181273 | 0.808460804 |
| PEX19 | 2552.494058 | 0.165996439 | 0.227924588 | 0.728295445 | 0.46643275 | 0.743017201 |
| PEX2 | 910.2920412 | 0.221241436 | 0.257690559 | 0.858554685 | 0.390586248 | 0.690466555 |
| PEX26 | 1389.239105 | -0.554165605 | 0.300753469 | -1.842590899 | 0.065388764 | 0.294978571 |
| PEX3 | 286.9767833 | 0.030307657 | 0.281784621 | 0.107556108 | 0.914347816 | 0.968409553 |
| PEX5 | 1028.423945 | 0.466301524 | 0.25346303 | 1.839722045 | 0.065809055 | 0.29619856 |
| PEX6 | 563.7562867 | -0.136116578 | 0.309838573 | -0.439314501 | 0.660433668 | 0.852355524 |
| PEX7 | 346.2681851 | 0.931883416 | 0.420902751 | 2.214011226 | 0.026828007 | 0.187071767 |
| PFAS | 1823.30075 | -0.229281879 | 0.501665994 | -0.457040904 | 0.647641645 | 0.846597949 |
| PFDN1 | 1371.075985 | 0.0023339 | 0.249373779 | 0.009359042 | 0.992532674 | 0.997035603 |
| PFDN2 | 1201.91575 | 1.199942848 | 0.398341862 | 3.012344326 | 0.002592384 | 0.049502724 |
| PFDN4 | 2461.999962 | 1.224510601 | 0.451740768 | 2.710648866 | 0.00671517 | 0.089679436 |
| PFDN5 | 5573.994018 | 0.733456855 | 0.384868028 | 1.905735996 | 0.056684467 | 0.27571659 |
| PFDN6 | 189.4080566 | 0.736498209 | 0.462958801 | 1.590850434 | 0.11164324 | 0.384591187 |
| PFKFB1 | 14.05686078 | 0.017999943 | 0.805442875 | 0.022347883 | 0.982170453 | 0.993518428 |
| PFKFB2 | 465.6036253 | -0.364247421 | 0.30589134 | -1.190773888 | 0.233742363 | 0.543788993 |

| | | | | | | |
|---|---|---|---|---|---|---|
| PFKFB3 | 411.8188964 | 0.350239755 | 0.323091019 | 1.084028135 | 0.278352325 | 0.592005173 |
| PFKFB4 | 523.2522781 | 0.166554891 | 0.323297898 | 0.515174679 | 0.606430961 | 0.826611695 |
| PFKL | 2866.570232 | -0.614951137 | 0.39326572 | -1.563703893 | 0.117887123 | 0.395839103 |
| PFKM | 3908.368638 | -0.254464981 | 0.235322938 | -1.081343719 | 0.279544247 | 0.593225034 |
| PFKP | 1911.946767 | -0.689407727 | 0.240801941 | -2.862965827 | 0.004196958 | 0.066981369 |
| PFN1 | 16118.58594 | 0.789884313 | 0.458899252 | 1.721258665 | 0.085203897 | 0.334358075 |
| PFN2 | 11725.8478 | 0.825223754 | 0.283731062 | 2.908471667 | 0.003632001 | 0.062081109 |
| PGA5 | 12.06746183 | 0.123325231 | 0.856350427 | 0.144012575 | 0.885490539 | 0.95478159 |
| PGAM1 | 14002.16566 | 0.447007304 | 0.258412313 | 1.729822 | 0.083662083 | 0.330936505 |
| PGAM2 | 140.1121023 | 1.908745891 | 0.596123905 | 3.201928111 | 0.001365111 | 0.033349586 |
| PGAM5 | 1572.623514 | 0.235129754 | 0.312640613 | 0.752076808 | 0.45200487 | 0.732888936 |
| PGAP1 | 1673.116739 | 0.174020387 | 0.371598199 | 0.468302557 | 0.63956824 | 0.843436176 |
| PGAP2 | 926.0188068 | -0.101078853 | 0.406343398 | -0.248752297 | 0.803552393 | 0.917969611 |
| PGAP3 | 619.9868804 | 0.042554706 | 0.311443904 | 0.136636826 | 0.891317867 | 0.957017706 |
| PGBD1 | 389.3448629 | -0.153315734 | 0.269899185 | -0.568048153 | 0.570002271 | 0.806205765 |
| PGBD2 | 236.1084997 | -0.732173993 | 0.327446716 | -2.23600958 | 0.025351144 | 0.181931031 |
| PGBD4 | 99.61076829 | -0.278473119 | 0.527441362 | -0.527969815 | 0.597520286 | 0.822220432 |
| PGBD5 | 131.3212457 | 1.295459689 | 0.606095551 | 2.137385248 | 0.03256667 | 0.205776324 |
| PGD | 10138.34203 | -0.270999188 | 0.299815753 | -0.903885752 | 0.366055985 | 0.672491566 |
| PGF | 152.5375811 | 0.291059829 | 0.395305044 | 0.736291714 | 0.461553187 | 0.73902094 |
| PGGHG | 685.2147137 | -0.112861795 | 0.361149483 | -0.312507147 | 0.754655132 | 0.897458102 |
| PGGT1B | 724.0207663 | 0.117046652 | 0.318106694 | 0.367947781 | 0.712912172 | 0.877743193 |
| PGK1 | 8658.081674 | 0.408422832 | 0.248593062 | 1.642937371 | 0.100395892 | 0.362557981 |
| PGM1 | 903.0236225 | 0.079449139 | 0.300409916 | 0.264469096 | 0.791418475 | 0.913361935 |
| PGM2 | 1209.544844 | -0.065619568 | 0.254922927 | -0.257409443 | 0.796862714 | 0.915710299 |
| PGM2L1 | 615.3525026 | -0.224372321 | 0.401407758 | -0.558963589 | 0.57618657 | 0.808993188 |
| PGM3 | 1877.580643 | -0.04706018 | 0.241390415 | -0.194954636 | 0.845428462 | 0.937778401 |
| PGM5 | 14.5978842 | -1.095269192 | 1.254191174 | -0.873287274 | 0.382506515 | 0.683824871 |
| PGM5P2 | 41.57240823 | 0.89371742 | 0.848324914 | 1.053508397 | 0.292108048 | 0.605157886 |
| PGP | 596.9721615 | 0.633820632 | 0.355192062 | 1.784444814 | 0.074351415 | 0.313285577 |
| PGPEP1 | 155.0486656 | 0.072473805 | 0.37383236 | 0.193867126 | 0.846279925 | 0.938145276 |
| PGRMC1 | 4077.774699 | 0.40977143 | 0.274812511 | 1.491094523 | 0.13593668 | 0.423730301 |
| PGRMC2 | 1711.10053 | 0.237181476 | 0.323442193 | 0.733304068 | 0.463372995 | 0.740758218 |
| PGS1 | 1240.424213 | -0.325178644 | 0.258427135 | -1.258299149 | 0.208283589 | 0.515167134 |
| PHACTR1 | 256.0306739 | -0.805246094 | 0.506515415 | -1.589776085 | 0.111885286 | 0.384888109 |
| PHACTR2 | 955.1604754 | -0.225504468 | 0.266251723 | -0.846959656 | 0.397017609 | 0.695594658 |
| PHACTR3 | 11.09192014 | 0.137520498 | 0.945213783 | 0.145491423 | 0.884322887 | 0.954200749 |
| PHACTR4 | 1434.085452 | -0.012453302 | 0.253776473 | -0.049071932 | 0.960861971 | 0.986559996 |
| PHAX | 1310.100704 | 0.091370177 | 0.25364156 | 0.360233459 | 0.718672555 | 0.880993556 |

| | | | | | | |
|---|---|---|---|---|---|---|
| PHB | 4230.804222 | 0.358108943 | 0.238429229 | 1.501950683 | 0.133109846 | 0.419486804 |
| PHB2 | 7408.104185 | 0.24283246 | 0.352329924 | 0.689218951 | 0.490685493 | 0.758593654 |
| PHC1 | 2515.798538 | -0.522128104 | 0.317845791 | -1.64270888 | 0.100443181 | 0.36264243 |
| PHC1P1 | 283.9211832 | -0.850708103 | 1.776892335 | -0.478761761 | 0.632108123 | 0.838768903 |
| PHC2 | 2484.069126 | 0.225810964 | 0.224596358 | 1.005407949 | 0.314700454 | 0.624045314 |
| PHC3 | 661.5810643 | -0.391351205 | 0.345608373 | -1.132354526 | 0.257485415 | 0.570028755 |
| PHETA1 | 233.4853968 | 0.049430663 | 0.385928549 | 0.12808242 | 0.898083748 | 0.959968192 |
| PHETA2 | 222.0104199 | -0.155527485 | 0.447080875 | -0.347873269 | 0.727935361 | 0.885393038 |
| PHF1 | 106.7376557 | 1.534124715 | 0.867227122 | 1.768999926 | 0.076893887 | 0.31844659 |
| PHF10 | 1816.640837 | -0.011563136 | 0.270134443 | -0.042805114 | 0.965856888 | 0.988700292 |
| PHF11 | 168.5339439 | -0.27141099 | 0.400668429 | -0.6773955 | 0.498155053 | 0.762537616 |
| PHF12 | 1265.982931 | -0.335735156 | 0.408346513 | -0.82218201 | 0.410973316 | 0.704426187 |
| PHF13 | 1431.038929 | 0.077729788 | 0.242009987 | 0.321184218 | 0.748070791 | 0.894254352 |
| PHF14 | 2053.977819 | 0.137500733 | 0.264325399 | 0.520194929 | 0.602927719 | 0.825021776 |
| PHF19 | 384.8770394 | 0.373522671 | 0.40185313 | 0.929500464 | 0.352629785 | 0.659729494 |
| PHF2 | 476.8548773 | -0.197840437 | 0.469169278 | -0.42168242 | 0.673256838 | 0.859400882 |
| PHF20 | 2804.377028 | 0.239818531 | 0.240032055 | 0.999110434 | 0.317741197 | 0.627826479 |
| PHF20L1 | 1117.694815 | 0.041196855 | 0.277761287 | 0.148317485 | 0.882092217 | 0.953217539 |
| PHF21A | 1469.193967 | -0.345922193 | 0.401249035 | -0.862113457 | 0.388625087 | 0.688604437 |
| PHF21B | 396.6759683 | 0.491085536 | 0.372916078 | 1.316879495 | 0.187879021 | 0.492689771 |
| PHF23 | 1865.615738 | 0.436385355 | 0.236647277 | 1.84403286 | 0.065178351 | 0.294467174 |
| PHF24 | 12.42636088 | 0.451078658 | 0.84039108 | 0.536748507 | 0.591441345 | 0.81920539 |
| PHF3 | 2600.312801 | -0.421386911 | 0.356560057 | -1.181811878 | 0.237280354 | 0.547879744 |
| PHF5A | 1427.98487 | 0.415413544 | 0.244724747 | 1.69747256 | 0.089607355 | 0.342777615 |
| PHF6 | 2462.782067 | 0.485781313 | 0.387280072 | 1.254341104 | 0.20971805 | 0.517759566 |
| PHF7 | 181.0308485 | 0.417614125 | 0.405738243 | 1.029269813 | 0.303352904 | 0.613785931 |
| PHF8 | 1168.132565 | -0.363807332 | 0.334767055 | -1.086747714 | 0.277148321 | 0.590951114 |
| PHGDH | 8426.524174 | 0.178978185 | 0.227494243 | 0.786737206 | 0.431435719 | 0.718177757 |
| PHIP | 3627.735481 | 0.110458029 | 0.300258214 | 0.367876795 | 0.712965104 | 0.877743193 |
| PHKA1 | 306.5597585 | -0.72979323 | 0.347108301 | -2.102494315 | 0.035509997 | 0.215661396 |
| PHKA2 | 694.8775407 | -0.36466008 | 0.386270986 | -0.944052475 | 0.345142833 | 0.651993765 |
| PHKB | 2162.226669 | -0.499767152 | 0.235939088 | -2.118204133 | 0.034157783 | 0.210824952 |
| PHKG2 | 1046.652687 | -0.184055871 | 0.3501232 | -0.525688874 | 0.599104395 | 0.823802482 |
| PHLDA1 | 858.1235029 | -1.960836112 | 0.795491013 | -2.464938109 | 0.013703692 | 0.133173911 |
| PHLDA2 | 18.58861964 | 1.63655513 | 0.770784096 | 2.123234169 | 0.033734234 | 0.209317818 |
| PHLDA3 | 680.7795265 | 0.633427229 | 0.362204332 | 1.748811855 | 0.080323547 | 0.324292679 |
| PHLDB1 | 1563.106478 | -0.078065299 | 0.386606278 | -0.201924551 | 0.839975709 | 0.934461703 |
| PHLDB2 | 634.8986283 | 0.225423113 | 0.250140265 | 0.901186832 | 0.367488991 | 0.673938238 |
| PHLDB3 | 155.3424365 | -0.130189153 | 0.519548239 | -0.250581453 | 0.802137723 | 0.917532337 |
| PHLPP1 | 724.9175043 | -0.678570354 | 0.288869564 | -2.349054517 | 0.018821151 | 0.155449937 |

| | | | | | | |
|---|---|---|---|---|---|---|
| PHLPP2 | 645.2993652 | -0.076595324 | 0.473251881 | -0.161848958 | 0.871424798 | 0.948581876 |
| PHOSPHO2 | 110.8279772 | 0.220672626 | 0.359910975 | 0.613131139 | 0.539789632 | 0.787188413 |
| PHOX2A | 25.97871268 | -1.65013195 | 0.872835558 | -1.89054162 | 0.05868556 | 0.280503663 |
| PHPT1 | 1385.825733 | 1.14245575 | 0.326564476 | 3.498407928 | 0.000468045 | 0.016786538 |
| PHRF1 | 18.02045869 | 7.351112819 | 4.332064285 | 1.696907602 | 0.089714131 | 0.343025354 |
| PHTF1 | 1138.455559 | 0.3312075 | 0.29799275 | 1.111461605 | 0.266369708 | 0.57970088 |
| PHTF2 | 669.3180693 | 0.212755576 | 0.299524091 | 0.710312066 | 0.477510639 | 0.750016059 |
| PHYH | 341.3661115 | 0.713987146 | 0.387583032 | 1.842152742 | 0.065452811 | 0.295179725 |
| PHYHD1 | 149.6172766 | -0.619649413 | 0.463778881 | -1.336088033 | 0.181520501 | 0.485768051 |
| PHYHIP | 79.16205898 | 1.351699422 | 0.611537454 | 2.210329741 | 0.027082286 | 0.188125165 |
| PHYKPL | 591.3536438 | -0.158738715 | 0.284151481 | -0.558641165 | 0.57640664 | 0.809091889 |
| PI15 | 36.49364149 | 0.121689927 | 0.53275237 | 0.22841743 | 0.81932174 | 0.924316427 |
| PI4K2A | 754.2822294 | -0.295173226 | 0.386497198 | -0.763713754 | 0.445037843 | 0.727707384 |
| PI4K2B | 773.0223657 | -0.407879462 | 0.415218248 | -0.982325473 | 0.325939529 | 0.635827259 |
| PI4KA | 1706.230906 | -1.041026771 | 0.462441263 | -2.251154588 | 0.024375748 | 0.178492903 |
| PI4KAP1 | 777.7739182 | -0.714797533 | 0.394444255 | -1.812163633 | 0.069960917 | 0.303858307 |
| PI4KAP2 | 1797.145954 | -1.469191304 | 0.419592201 | -3.501474291 | 0.000462692 | 0.016673383 |
| PI4KB | 1058.799277 | -0.31540689 | 0.315957895 | -0.998256081 | 0.318155198 | 0.628235392 |
| PIANP | 145.3632089 | 0.17783133 | 0.564051781 | 0.315274831 | 0.752552993 | 0.896586461 |
| PIAS1 | 1482.994517 | -0.294228982 | 0.271359836 | -1.084276093 | 0.278242402 | 0.592005173 |
| PIAS2 | 739.4484286 | -0.027508618 | 0.275883618 | -0.099710953 | 0.920573805 | 0.969843271 |
| PIAS3 | 1159.622529 | 0.340604143 | 0.311615426 | 1.09302722 | 0.274381849 | 0.587352209 |
| PIAS4 | 600.4037175 | -0.383045219 | 0.472299351 | -0.811022116 | 0.41735297 | 0.708794572 |
| PIBF1 | 654.1902476 | 0.395179773 | 0.386676301 | 1.021991189 | 0.30678507 | 0.616910609 |
| PICALM | 2694.515712 | -0.35919585 | 0.314460051 | -1.142262267 | 0.253345021 | 0.565885941 |
| PICK1 | 527.2053046 | -0.463837835 | 0.344843356 | -1.345068209 | 0.178603209 | 0.482972162 |
| PID1 | 38.8614689 | 1.396211397 | 0.95370351 | 1.463988946 | 0.143196969 | 0.433589745 |
| PIDD1 | 590.5711214 | 0.080766348 | 0.542388476 | 0.148908673 | 0.881625697 | 0.953217539 |
| PIEZO1 | 1074.447931 | -2.764089363 | 0.446864625 | -6.185518409 | 6.19E-10 | 1.06E-06 |
| PIEZO2 | 173.5485613 | -1.226275317 | 0.438658581 | -2.795511975 | 0.005181758 | 0.076591444 |
| PIF1 | 923.384177 | 1.030515091 | 0.302201721 | 3.410023903 | 0.000649572 | 0.020790411 |
| PIFO | 183.5603815 | 0.220612487 | 0.3573118 | 0.617422896 | 0.536955824 | 0.785493754 |
| PIGA | 335.3661741 | -0.14635039 | 0.275736922 | -0.530760949 | 0.595584444 | 0.821568613 |
| PIGB | 365.4231471 | 0.085741303 | 0.298698984 | 0.287049193 | 0.774074654 | 0.90627308 |
| PIGBOS1 | 256.5058905 | 0.696435494 | 0.519233034 | 1.341277324 | 0.179830429 | 0.484583915 |
| PIGC | 766.0295949 | -0.108094598 | 0.306365946 | -0.35282837 | 0.724217113 | 0.883667763 |
| PIGF | 833.8378043 | 1.052332648 | 0.350085313 | 3.005932008 | 0.002647682 | 0.050167035 |
| PIGG | 1089.356713 | -0.512461607 | 0.307322902 | -1.667502174 | 0.095414593 | 0.354701001 |
| PIGH | 191.6016461 | 0.627468082 | 0.314577652 | 1.994636547 | 0.046082535 | 0.246950947 |
| PIGK | 739.6972021 | 0.292539581 | 0.247083995 | 1.183968151 | 0.236425671 | 0.547336452 |
| PIGL | 424.9266608 | 0.110453515 | 0.39078205 | 0.282647362 | 0.777447171 | 0.907979294 |

| | | | | | | |
|---|---|---|---|---|---|---|
| PIGM | 455.5528316 | 0.125513446 | 0.34689074 | 0.361824148 | 0.717483446 | 0.880516207 |
| PIGN | 1463.302804 | -0.558348716 | 0.313814658 | -1.779230834 | 0.075201925 | 0.314600339 |
| PIGO | 829.8270491 | -0.472685877 | 0.334480098 | -1.413195825 | 0.157598154 | 0.4568931 |
| PIGP | 375.9732818 | 0.88677615 | 0.44450402 | 1.994978918 | 0.046045179 | 0.246950947 |
| PIGQ | 754.6044579 | -0.271583126 | 0.493237312 | -0.550613506 | 0.581898648 | 0.813002261 |
| PIGS | 1191.160231 | -0.42901846 | 0.278993918 | -1.537734094 | 0.124113644 | 0.40536665 |
| PIGT | 1942.776514 | -0.527856572 | 0.238805929 | -2.210399779 | 0.027077429 | 0.188125165 |
| PIGU | 1009.299736 | 0.670988947 | 0.267593168 | 2.507496552 | 0.012158977 | 0.125278548 |
| PIGV | 389.8730252 | -0.240950684 | 0.264887075 | -0.909635487 | 0.363014777 | 0.669987492 |
| PIGW | 158.9720943 | -0.016387181 | 0.456940069 | -0.035862868 | 0.971391704 | 0.989658654 |
| PIGX | 390.7285841 | 0.386858962 | 0.397550137 | 0.973107355 | 0.330499949 | 0.638890765 |
| PIH1D1 | 1927.916292 | 0.476969154 | 0.331081206 | 1.440640983 | 0.149686136 | 0.444386222 |
| PIH1D2 | 49.59513063 | 0.434076061 | 0.50873766 | 0.853241454 | 0.39352542 | 0.692506234 |
| PIK3AP1 | 19.63311052 | -0.250482358 | 0.902144541 | -0.277652135 | 0.781279407 | 0.909732666 |
| PIK3C2A | 2647.92146 | -0.405294413 | 0.365222595 | -1.109718892 | 0.26712018 | 0.580460198 |
| PIK3C2B | 1212.833279 | 0.229370367 | 0.460517228 | 0.498071197 | 0.618433861 | 0.832277132 |
| PIK3C2G | 44.93503381 | -3.27675265 | 0.848938061 | -3.859825351 | 0.000113468 | 0.006724314 |
| PIK3C3 | 1203.684287 | -0.567237432 | 0.298457206 | -1.90056538 | 0.057358964 | 0.277308107 |
| PIK3CA | 571.6190283 | -0.295259432 | 0.363699709 | -0.81182202 | 0.416893765 | 0.708661116 |
| PIK3CB | 994.5681477 | -0.226592574 | 0.289939187 | -0.781517588 | 0.434498137 | 0.719554351 |
| PIK3CD | 275.4201808 | -0.914564447 | 0.291261871 | -3.140007457 | 0.001689435 | 0.038083839 |
| PIK3IP1 | 138.2185878 | -1.026136865 | 0.447071889 | -2.295239066 | 0.021719428 | 0.167346589 |
| PIK3R1 | 698.519226 | -0.448466824 | 0.481472913 | -0.931447672 | 0.351622041 | 0.659063255 |
| PIK3R2 | 2085.352047 | -0.117626717 | 0.236991253 | -0.496333582 | 0.619659077 | 0.832817847 |
| PIK3R3 | 1539.991638 | 0.67978797 | 0.548334647 | 1.23973193 | 0.215074563 | 0.522230505 |
| PIK3R4 | 1092.041386 | -0.332645479 | 0.319636982 | -1.040697722 | 0.298015861 | 0.609375237 |
| PIK3R5 | 79.70866335 | 0.442193836 | 0.514352789 | 0.859709221 | 0.38994935 | 0.689813912 |
| PIKFYVE | 1730.091063 | -0.184448428 | 0.353167573 | -0.522268868 | 0.601483142 | 0.824869221 |
| PILRA | 17.28764118 | 0.372310636 | 0.807712183 | 0.460944682 | 0.644838293 | 0.845794147 |
| PILRB | 929.2688373 | -0.472477322 | 0.336515236 | -1.404029509 | 0.160310063 | 0.460580192 |
| PIM1 | 1382.486525 | 0.013987402 | 0.368395674 | 0.03796842 | 0.969712861 | 0.989496332 |
| PIM2 | 431.7648334 | 0.610282017 | 0.335217069 | 1.820557701 | 0.068674118 | 0.301812004 |
| PIM3 | 574.1435214 | 0.709649732 | 0.391493508 | 1.812673051 | 0.069882263 | 0.303603614 |
| PIMREG | 1520.867325 | 1.551187423 | 0.376623151 | 4.118672524 | 3.81E-05 | 0.003042674 |
| PIN1 | 979.766931 | 0.633689978 | 0.345347318 | 1.83493528 | 0.066515279 | 0.297582809 |
| PIN4 | 888.2921122 | 0.436732271 | 0.420363128 | 1.038940481 | 0.298832418 | 0.610091053 |
| PINK1 | 999.2923698 | -0.384284548 | 0.38371225 | -1.001491476 | 0.316589259 | 0.626693064 |

| | | | | | | |
|---|---|---|---|---|---|---|
| PINLYP | 169.2540241 | 1.061061936 | 0.367949284 | 2.883717897 | 0.003930106 | 0.064878831 |
| PINX1 | 476.2032439 | 0.297725147 | 0.280370849 | 1.061897656 | 0.288282149 | 0.601673803 |
| PIP4K2A | 942.7649686 | -0.638403585 | 0.532607199 | -1.198638671 | 0.230668475 | 0.539536045 |
| PIP4K2B | 1165.419469 | -0.129381326 | 0.405099065 | -0.319381941 | 0.749436903 | 0.895323038 |
| PIP4K2C | 1098.744827 | -1.046542231 | 0.25851049 | -4.048354987 | 5.16E-05 | 0.0038721 |
| PIP4P1 | 1152.346409 | 0.151871306 | 0.334933306 | 0.453437456 | 0.650233771 | 0.847430646 |
| PIP4P2 | 579.8118134 | 0.476303747 | 0.32923899 | 1.446680867 | 0.147986326 | 0.441755032 |
| PIP5K1A | 2430.304468 | -0.005926313 | 0.271280061 | -0.021845736 | 0.982571011 | 0.993518428 |
| PIP5K1B | 60.6323169 | -0.995727227 | 0.5714269 | -1.742527744 | 0.081416141 | 0.326289185 |
| PIP5K1C | 930.9169733 | -0.586046967 | 0.355514283 | -1.648448442 | 0.099260682 | 0.361452832 |
| PIP5KL1 | 72.68494595 | 0.587375583 | 0.446061938 | 1.316802742 | 0.187904754 | 0.492689771 |
| PIPOX | 205.9985962 | 1.823502367 | 0.687029422 | 2.654183806 | 0.007950051 | 0.100258644 |
| PIPSL | 14.31366522 | -0.738311274 | 0.890159801 | -0.829414306 | 0.406870009 | 0.701626421 |
| PIR | 526.8475977 | 0.931348902 | 0.372568424 | 2.499806321 | 0.012426122 | 0.126606244 |
| PISD | 889.8052999 | -0.237877311 | 0.296633369 | -0.80192364 | 0.42259713 | 0.712269066 |
| PITHD1 | 1514.76032 | 0.598799853 | 0.439970946 | 1.360998627 | 0.173514123 | 0.477540818 |
| PITPNA | 3896.426297 | -0.56118331 | 0.24148429 | -2.323891585 | 0.020131305 | 0.160489771 |
| PITPNB | 1950.551125 | -0.270077107 | 0.320566374 | -0.842499805 | 0.399508249 | 0.697589493 |
| PITPNC1 | 596.5543319 | 0.329027792 | 0.374836817 | 0.877789421 | 0.380058007 | 0.68184242 |
| PITPNM1 | 488.5795925 | 0.086825953 | 0.592997267 | 0.146418809 | 0.88359078 | 0.953885699 |
| PITPNM2 | 14.58087023 | 0.512758613 | 0.858957015 | 0.596954917 | 0.550537481 | 0.794328924 |
| PITPNM3 | 188.3129824 | -0.030529535 | 0.331332967 | -0.092141555 | 0.926585573 | 0.972261514 |
| PITRM1 | 1020.153627 | -0.307861905 | 0.383356093 | -0.803070332 | 0.421934084 | 0.712063713 |
| PITX2 | 38.45914236 | 0.820890436 | 0.809290345 | 1.014333658 | 0.310423569 | 0.62031559 |
| PIWIL4 | 157.9473683 | -2.138758875 | 0.734481447 | -2.911930431 | 0.003592026 | 0.061575854 |
| PJA1 | 1554.410331 | -0.200720835 | 0.335810878 | -0.597719871 | 0.550026864 | 0.793878561 |
| PJA2 | 2629.65919 | -0.058659489 | 0.343204562 | -0.170916986 | 0.86428904 | 0.945136254 |
| PJVK | 153.1381905 | -0.145572001 | 0.34797553 | -0.418339765 | 0.67569872 | 0.860110127 |
| PKD1 | 612.7087751 | 0.250692957 | 0.573463746 | 0.43715572 | 0.66199842 | 0.853756491 |
| PKD1L1 | 14.38969711 | -1.379789341 | 0.967561315 | -1.426048479 | 0.153854349 | 0.450690158 |
| PKD1L2 | 87.35194135 | 1.327476392 | 0.57156988 | 2.322509352 | 0.020205525 | 0.160763185 |
| PKD1P1 | 271.1678345 | -0.292372626 | 1.793456556 | -0.163021861 | 0.870501222 | 0.948188831 |
| PKD1P5 | 54.01387762 | 0.112985789 | 1.264060183 | 0.089383236 | 0.928777346 | 0.972763626 |
| PKD1P6 | 97.27036249 | -0.271179539 | 0.96049294 | -0.282333715 | 0.777687636 | 0.908018171 |
| PKD2 | 1285.464068 | -1.028806691 | 0.462442454 | -2.224723709 | 0.026099792 | |

0.184424751
PKDCC 6842.59061 -1.159454122 0.369991911 -3.133728296 0.001726006
0.03867834
PKHD1 19.04674856 -1.309722399 0.770522543 -1.69978466 0.089171438
0.342726775
PKIA 355.7199513 0.188961111 0.285262753 0.662410738 0.507708024 0.768476348
PKIB 105.8993857 -0.461917863 0.677731471 -0.681564724 0.495514227
0.760859008
PKIG 713.1116839 0.269133629 0.323793353 0.831189481 0.405866596 0.701371854
PKM 40178.38805 0.195732085 0.323605975 0.604846944 0.545280699 0.79117123
PKMYT1 440.6721425 0.078575304 0.381710973 0.205850262 0.836907878
0.933183529
PKN1 2085.259156 0.546351792 0.307317044 1.777811556 0.07543481 0.314738565
PKN2 1643.840514 -0.257740694 0.350843645 -0.734631219 0.46256412
0.740092835
PKN3 718.8179489 0.221500784 0.369760303 0.599038842 0.549146983 0.79348329
PKNOX1 440.3001999 -0.708188871 0.444486932 -1.593272647
0.11109904 0.383475916
PKNOX2 378.3253123 0.887915528 0.672330939 1.320652489 0.186617263
0.49118221
PKP1 52.43696623 -0.881583132 0.782779552 -1.126221462 0.260071791
0.57337005
PKP2 3093.797283 -1.556749473 0.497576295 -3.128664867 0.001756025
0.039027635
PKP3 515.0680793 -1.302767875 0.730442028 -1.783533564 0.074499489
0.313517823
PKP4 2832.126855 -0.48409381 0.284031613 -1.704365953 0.08831274
0.341701126
PLA2G12A 577.5508175 0.426151446 0.398974185 1.068117844 0.285467358
0.599411008
PLA2G12B 32.44147899 0.133441333 0.987930273 0.13507161 0.892555256
0.957550733
PLA2G15 246.7914248 0.263172611 0.326325558 0.806472569 0.419970418
0.710456202
PLA2G16 401.3289796 -1.796166501 0.573768476 -3.130472615
0.001745253 0.038937101
PLA2G2A 126.644338 -0.764496333 0.92922508 -0.8227246 0.410664623
0.704374561
PLA2G3 125.9760705 0.419161108 0.409097012 1.024600758 0.305551607
0.615607361
PLA2G4A 99.03996299 0.071313339 0.445896321 0.159932558 0.872934201
0.949475535
PLA2G4B 91.73472439 0.673143376 0.385738046 1.74507903 0.080971113
0.325493577
PLA2G4C 26.54289324 0.186800819 0.76882572 0.242969003 0.80802941
0.919580875
PLA2G4F 130.9698052 -2.072545798 0.810000229 -2.558697792
0.010506503 0.116405896
PLA2G6 616.0773799 -0.590189086 0.343184094 -1.719744873
0.085478827 0.335263518
PLA2G7 110.4928577 -0.817147166 0.54349451 -1.503505834
0.132708654 0.418744381
PLA2R1 95.26096891 -1.297806894 0.534751867 -2.426932893
0.015227072 0.139780768
PLAA 1663.982388 -0.158550718 0.24466498 -0.648031926 0.516964301
0.773950969
PLAC1 30.17974474 0.327842483 0.697852003 0.46978798 0.638506503 0.842491082
PLAC8 20.07997506 -1.0349188 0.896181826 -1.154808957 0.248168668
0.56079644
PLAC8L1 12.95643627 -0.114252647 0.91416391 -0.124980483
0.900539001 0.961576379
PLAC9 15.08991717 -0.381046994 1.128489968 -0.337660949 0.735618706
0.889855747
PLAG1 776.4075368 0.02199808 0.347785623 0.063251839 0.949565966 0.981593436

| | | | | | | |
|---|---|---|---|---|---|---|
| PLAGL1 | 164.5449081 | -0.024337007 | 0.667705217 | -0.036448729 | 0.97092456 | 0.989604013 |
| PLAGL2 | 1066.996137 | -0.238757397 | 0.273450664 | -0.873127875 | 0.382593382 | 0.683824871 |
| PLAT | 230.5882754 | 1.267811842 | 0.620799856 | 2.042223159 | 0.0411294 | 0.232946328 |
| PLAU | 629.0288723 | 0.712475943 | 0.469037721 | 1.519016298 | 0.128758394 | 0.413329156 |
| PLAUR | 331.3222823 | 0.786876646 | 0.31404303 | 2.505633209 | 0.012223235 | 0.125721147 |
| PLB1 | 60.07608578 | -0.750894012 | 0.656520017 | -1.143748847 | 0.252727809 | 0.565423035 |
| PLBD1 | 292.2268438 | -0.298521325 | 0.33783265 | -0.88363669 | 0.376892366 | 0.68029735 |
| PLBD2 | 1238.728411 | -0.438626554 | 0.241311699 | -1.817676294 | 0.069113623 | 0.303010569 |
| PLCB1 | 917.645797 | -0.24043547 | 0.236242395 | -1.017749039 | 0.308797236 | 0.618441002 |
| PLCB2 | 98.80701758 | -0.442558779 | 0.499003023 | -0.886885968 | 0.375140297 | 0.679214334 |
| PLCB3 | 1537.987662 | -0.35902636 | 0.361149865 | -0.994120154 | 0.320164374 | 0.62990711 |
| PLCB4 | 316.8105981 | -0.286475193 | 0.343925346 | -0.832957491 | 0.404868704 | 0.70065889 |
| PLCD1 | 582.7631478 | 0.694503168 | 0.381250361 | 1.821645929 | 0.068508727 | 0.301609373 |
| PLCD3 | 619.0916615 | -0.308264536 | 0.32006368 | -0.963135012 | 0.335479773 | 0.643919384 |
| PLCD4 | 157.0481413 | -0.875938831 | 0.450087319 | -1.9461531 | 0.051636352 | 0.261823226 |
| PLCE1 | 538.4682069 | -2.5044148 | 0.327209103 | -7.653866521 | 1.95E-14 | 9.86E-11 |
| PLCG1 | 4982.793117 | -0.302685822 | 0.50872561 | -0.59498837 | 0.551851245 | 0.795324338 |
| PLCG2 | 216.8541703 | -0.042207849 | 0.322664636 | -0.130810273 | 0.895925397 | 0.958944842 |
| PLCH1 | 666.5028445 | 0.040320318 | 0.463250366 | 0.087037855 | 0.930641422 | 0.973909155 |
| PLCL2 | 183.8058877 | 0.220678616 | 0.422571077 | 0.522228397 | 0.601511317 | 0.824869221 |
| PLCXD1 | 2133.989037 | 0.126362556 | 0.236315596 | 0.534719493 | 0.59284384 | 0.819859199 |
| PLCXD2 | 189.963833 | 0.379576915 | 0.447940242 | 0.847382931 | 0.396781716 | 0.695422297 |
| PLCXD3 | 47.27106511 | -1.84888835 | 0.598389755 | -3.089772737 | 0.002003097 | 0.042131685 |
| PLD1 | 85.54985267 | -1.998851137 | 0.535975243 | -3.729372136 | 0.000191957 | 0.009160128 |
| PLD2 | 2077.56439 | -0.886652533 | 0.242932648 | -3.649787461 | 0.000262457 | 0.011279719 |
| PLD3 | 4943.714846 | -0.593301049 | 0.342216353 | -1.73370163 | 0.082971037 | 0.329644853 |
| PLD5 | 71.42222864 | -1.765572393 | 0.760215328 | -2.322463556 | 0.020207989 | 0.160763185 |
| PLD6 | 57.50129901 | 0.819897856 | 0.48917868 | 1.676070299 | 0.093724424 | 0.351226741 |
| PLEC | 1494.081948 | -0.6945635 | 0.44570804 | -1.558337381 | 0.119153292 | 0.398078527 |
| PLEK2 | 36.30914293 | -2.101895487 | 0.841905286 | -2.496593765 | 0.012539251 | 0.12669427 |
| PLEKHA1 | 1270.971233 | -0.404500877 | 0.273565539 | -1.478625115 | 0.139240537 | 0.427816061 |
| PLEKHA2 | 37.09919273 | -0.975624982 | 0.668558013 | -1.459297418 | 0.144483266 | 0.435776467 |
| PLEKHA3 | 389.89882 | 0.52532631 | 0.345345893 | 1.52115986 | 0.128219726 | 0.412317905 |
| PLEKHA4 | 512.3107019 | 0.370265644 | 0.460327385 | 0.80435285 | 0.421193223 | 0.711651898 |
| PLEKHA5 | 1487.574253 | -0.101938818 | 0.377964858 | -0.269704486 | 0.787387609 | 0.911971196 |
| PLEKHA6 | 343.2905576 | -1.244697852 | 0.602442837 | -2.066084575 | 0.038820494 | 0.226954035 |

| | | | | | | |
|---|---|---|---|---|---|---|
| PLEKHA7 | 1098.322691 | -0.989989589 | 0.384935486 | -2.57183249 | 0.010116183 | 0.114259456 |
| PLEKHA8 | 2064.931446 | -0.134486414 | 0.34295488 | -0.392140256 | 0.694954585 | 0.869142545 |
| PLEKHA8P1 | 152.6134908 | -0.509971427 | 0.391613272 | -1.302232235 | 0.192837009 | 0.498908723 |
| PLEKHB1 | 638.531933 | 1.14610666 | 0.445979962 | 2.569861335 | 0.010173923 | 0.114501918 |
| PLEKHB2 | 4063.043925 | 0.044585039 | 0.229157969 | 0.194560283 | 0.845737199 | 0.937897187 |
| PLEKHD1 | 16.52943687 | -0.962251246 | 1.068200595 | -0.900815119 | 0.367686628 | 0.674002337 |
| PLEKHF1 | 63.46781212 | -0.137175423 | 0.652429874 | -0.210253129 | 0.833470115 | 0.931803619 |
| PLEKHF2 | 494.4306713 | -0.324917258 | 0.483025339 | -0.672671249 | 0.501156463 | 0.763971533 |
| PLEKHG1 | 148.9115295 | -0.884773964 | 0.402607928 | -2.19760691 | 0.02797713 | 0.19118966 |
| PLEKHG2 | 1408.202014 | -0.698601275 | 0.40229817 | -1.736526108 | 0.082470852 | 0.328570367 |
| PLEKHG3 | 1056.646945 | 0.461589875 | 0.427422391 | 1.079938453 | 0.280169589 | 0.594053505 |
| PLEKHG4 | 186.7275661 | -0.742854712 | 0.306948177 | -2.420130717 | 0.015514929 | 0.140859955 |
| PLEKHG4B | 2307.458971 | -0.799456667 | 0.357437156 | -2.236635596 | 0.025310166 | 0.181849597 |
| PLEKHG5 | 458.2656334 | -0.278299214 | 0.466206533 | -0.596944046 | 0.550544739 | 0.794328924 |
| PLEKHG6 | 505.7609267 | -0.699468091 | 0.416649142 | -1.678794027 | 0.093192193 | 0.350041783 |
| PLEKHH1 | 1262.094016 | -1.241634559 | 0.249700167 | -4.972501918 | 6.61E-07 | 0.000139266 |
| PLEKHH2 | 201.5940961 | 0.192453085 | 0.5955858 | 0.323132427 | 0.746594953 | 0.894109554 |
| PLEKHH3 | 344.6992877 | 0.377998471 | 0.521099737 | 0.725386034 | 0.468215239 | 0.744267958 |
| PLEKHJ1 | 1853.631305 | 0.460639818 | 0.392855566 | 1.172542425 | 0.24097935 | 0.552437022 |
| PLEKHM1 | 196.6731156 | -0.334492111 | 0.607535958 | -0.55057171 | 0.581927306 | 0.813002261 |
| PLEKHM1P1 | 236.2540703 | -1.063004335 | 0.473607508 | -2.244483708 | 0.024801296 | 0.180028929 |
| PLEKHM2 | 1280.938524 | -0.490994412 | 0.492020561 | -0.99791442 | 0.318320859 | 0.628398926 |
| PLEKHM3 | 107.9682896 | -0.779245307 | 0.493800355 | -1.578057404 | 0.114552423 | 0.389533505 |
| PLEKHN1 | 50.53246541 | -2.527836867 | 0.815079822 | -3.101336577 | 0.001926492 | 0.041048886 |
| PLEKHO1 | 1039.790274 | 0.538696705 | 0.312113108 | 1.725966293 | 0.084353477 | 0.332479605 |
| PLEKHO2 | 559.2624266 | -0.196037308 | 0.360809741 | -0.543325985 | 0.586905376 | 0.816520994 |
| PLGLB1 | 30.46373446 | 0.121802174 | 0.631961451 | 0.192736714 | 0.847165169 | 0.938145276 |
| PLGRKT | 627.3979216 | 0.586898463 | 0.370234497 | 1.585207397 | 0.11291922 | 0.38661645 |
| PLIN2 | 2588.313828 | -0.248589747 | 0.326366976 | -0.76168781 | 0.446246352 | 0.728589711 |
| PLIN3 | 1708.729833 | 0.33147397 | 0.362275177 | 0.914978423 | 0.360202958 | 0.667522123 |
| PLIN4 | 22.38243318 | 0.142758354 | 0.724978085 | 0.196914027 | 0.843894816 | 0.93669361 |
| PLIN5 | 77.09577165 | 0.865525538 | 0.406089984 | 2.131363915 | 0.033059173 | 0.207677316 |
| PLK1 | 2190.591544 | 0.644908517 | 0.427307818 | 1.509236408 | 0.131238379 | 0.416813082 |
| PLK2 | 744.4311423 | -0.282574559 | 0.371996579 | -0.759616014 | 0.447484144 | 0.729035862 |

| | | | | | | |
|---|---|---|---|---|---|---|
| PLK3 | 140.2895776 | 0.606073262 | 0.361678141 | 1.67572544 | 0.093791984 | 0.351226741 |
| PLK4 | 981.1235402 | 0.277104055 | 0.380165923 | 0.72890293 | 0.466061043 | 0.742789377 |
| PLK5 | 12.63380405 | 0.058120829 | 0.835799039 | 0.069539239 | 0.9445604 | 0.979422174 |
| PLLP | 53.45458138 | -0.057895942 | 0.678345455 | -0.085348758 | 0.93198413 | 0.974430344 |
| PLOD1 | 1984.278359 | -0.420786104 | 0.234090181 | -1.797538465 | 0.072250177 | 0.30937269 |
| PLOD2 | 807.3201 | -0.779826086 | 0.354595168 | -2.199201105 | 0.027863626 | 0.190853383 |
| PLOD3 | 667.0344035 | -0.10781168 | 0.323613287 | -0.333149732 | 0.73902126 | 0.891444028 |
| PLP1 | 1368.387454 | 0.89108885 | 0.474887772 | 1.876419867 | 0.060597649 | 0.284105089 |
| PLP2 | 202.906013 | 0.225331698 | 0.461229066 | 0.488546179 | 0.625163025 | 0.835033444 |
| PLPBP | 820.1835033 | 0.199804163 | 0.234395447 | 0.852423396 | 0.39397914 | 0.692865578 |
| PLPP1 | 447.2287733 | 0.069829406 | 0.500225648 | 0.139595814 | 0.888979348 | 0.956217735 |
| PLPP2 | 237.4990997 | 0.628055222 | 0.734026201 | 0.855630523 | 0.392202181 | 0.691514794 |
| PLPP3 | 1273.528281 | 0.024195371 | 0.266107196 | 0.090923401 | 0.927553456 | 0.972633911 |
| PLPP4 | 28.27045572 | -3.320746483 | 0.90369326 | -3.674638983 | 0.000238186 | 0.010565846 |
| PLPP5 | 839.2810851 | 0.691606869 | 0.334967125 | 2.064700736 | 0.038951325 | 0.227316695 |
| PLPP6 | 452.1735793 | -0.307433085 | 0.352137895 | -0.873047433 | 0.382637224 | 0.683824871 |
| PLPP7 | 61.15726793 | 1.055606074 | 0.57663652 | 1.830626465 | 0.067156316 | 0.297817842 |
| PLPPR1 | 16.14084684 | 0.612917955 | 0.853761759 | 0.717902797 | 0.472817223 | 0.747093345 |
| PLPPR2 | 109.2440572 | 0.422975219 | 0.429594826 | 0.984591047 | 0.324824989 | 0.634714054 |
| PLPPR3 | 688.3612825 | -0.036200601 | 0.422760819 | -0.085629035 | 0.931761317 | 0.974341807 |
| PLRG1 | 2273.358582 | -0.02499952 | 0.247437744 | -0.101033575 | 0.919523807 | 0.969677217 |
| PLS1 | 853.4587848 | -0.014677164 | 0.297378292 | -0.049355197 | 0.960636232 | 0.986479967 |
| PLS3 | 5755.248946 | -0.467293561 | 0.229537706 | -2.035803044 | 0.041770137 | 0.235049979 |
| PLSCR1 | 467.3215413 | -0.324654939 | 0.431696368 | -0.752044638 | 0.452024215 | 0.732888936 |
| PLSCR3 | 352.5531965 | -0.240004152 | 0.414061124 | -0.579634596 | 0.562161058 | 0.801706045 |
| PLSCR4 | 23.24081964 | -1.34964889 | 0.774252676 | -1.743163352 | 0.081305085 | 0.326151133 |
| PLSCR5 | 20.19582572 | -6.951742905 | 4.409182959 | -1.576651042 | 0.114875841 | 0.389985713 |
| PLTP | 2779.656292 | -0.093302075 | 0.252486047 | -0.36953359 | 0.711730041 | 0.877287654 |
| PLXDC1 | 26.31558116 | -0.147863655 | 0.712494566 | -0.20752952 | 0.835596341 | 0.932531417 |
| PLXDC2 | 997.2438924 | -1.135806034 | 0.336361249 | -3.376744608 | 0.000733491 | 0.022388677 |
| PLXNA1 | 2328.068868 | -0.572276757 | 0.497382162 | -1.150577566 | 0.249906067 | 0.562742997 |
| PLXNA2 | 405.9048382 | -0.575475943 | 0.472151399 | -1.218837738 | 0.222905785 | 0.531941648 |
| PLXNA3 | 868.4258607 | -0.480803593 | 0.493734416 | -0.973810164 | 0.330150805 | 0.638717861 |
| PLXNB1 | 3114.9756 | -0.458532481 | 0.339523523 | -1.350517563 | 0.176850024 | 0.481149663 |
| PLXNB2 | 2302.119603 | 0.089899378 | 0.570967785 | 0.157450876 | 0.87488952 | 0.949740014 |
| PLXNB3 | 24.23589399 | 1.491227338 | 0.832942293 | 1.790312907 | 0.073403623 | 0.311344591 |
| PLXNC1 | 21.68988219 | 0.75313417 | 0.802568448 | 0.938404907 | 0.348036365 | 0.654528628 |
| PLXND1 | 677.5336143 | -0.468943134 | 0.343852208 | -1.363792707 | | |

0.172632816 0.476336299
PM20D2 544.7560268 -0.396337248 0.263938366 -1.501628025
0.133193201 0.419488075
PMAIP1 144.3954318 0.232833723 0.493209685 0.47207857 0.636870708
0.841268221
PMEL 4092.751025 0.763667596 0.451003686 1.693262426 0.090405527 0.344608605
PMEPA1 42.87403188 0.88325356 0.637481045 1.385536977 0.165888334
0.467940842
PMF1 1572.159904 0.41643396 0.355376976 1.171809062 0.241273724 0.552673201
PMF1-BGLAP 71.66036153 -0.130367569 0.55109872 -0.236559375
0.81299864 0.921802034
PMFBP1 55.14174569 -0.512407012 0.773446231 -0.662498557
0.507651759 0.768467855
PML 681.7051527 -0.062497429 0.250304556 -0.249685544 0.802830539
0.917969611
PMM1 603.0974364 0.457770482 0.437748455 1.045738658 0.295681709 0.60699582
PMM2 811.5582223 -0.488638846 0.277136059 -1.763173106 0.077871298
0.320071922
PMP22 294.55357 -0.844518185 0.739072204 -1.142673451 0.253174197
0.565800563
PMPCA 1714.734305 -0.053640373 0.281728967 -0.190397082 0.848997984
0.938580023
PMPCB 3196.952461 0.122634585 0.253220124 0.484300311 0.628172756 0.836626185
PMS1 1549.415982 0.388365625 0.273723005 1.418827131 0.155949421 0.454370782
PMS2 942.559597 -0.201180782 0.240821998 -0.83539205 0.403497004
0.700154758
PMS2CL 308.7959305 -0.078879988 0.316027312 -0.249598643
0.802897748 0.917969611
PMS2P1 178.8637167 0.649722535 0.355942367 1.825358809 0.067946901
0.299832005
PMS2P2 15.89602951 0.325014727 0.735343604 0.441990282 0.658496238
0.851158424
PMS2P3 97.11622058 -0.02763146 0.366515328 -0.075389643 0.939904699
0.978003717
PMS2P4 31.76064697 -0.144167053 0.594482186 -0.242508617
0.808386081 0.91963689
PMS2P5 18.4457058 0.710255461 0.80364558 0.883791908 0.376808555
0.68029735
PMS2P6 110.2416879 0.122967278 0.481592668 0.255334614 0.798464668
0.916056735
PMS2P7 21.04712959 0.24067014 0.677963339 0.354989903 0.722597155
0.882935039
PMVK 626.3685472 0.962024058 0.446781865 2.153229872 0.031300617 0.201896965
PNCK 326.0970048 2.014011986 0.743511275 2.708784727 0.006753014 0.089947301
PNISR 7153.875295 0.330702572 0.278977466 1.185409618 0.235855531 0.546701949
PNKD 846.9196424 0.649698956 0.374683623 1.73399347 0.082919242 0.329609655
PNKP 772.322546 0.377447929 0.302013795 1.249770492 0.211383398 0.518651612
PNMA1 1800.88625 0.218798884 0.246138943 0.888924287 0.374043772 0.678520659
PNMA2 130.4565668 1.927097594 0.690554464 2.790652578 0.00526019 0.077178282
PNMA3 52.23163546 0.363788807 0.573848558 0.633945667 0.526116281 0.778525486
PNMA6A 23.13506239 1.582361842 0.818728255 1.932707016 0.0532723
0.267056023
PNMA8A 1401.027593 0.411671983 0.336334974 1.223993978 0.220954499
0.529725144
PNMA8B 12.36573375 0.402926679 0.80819112 0.498553707 0.618093824
0.832176095
PNMA8C 14.10830181 1.450920731 0.90195887 1.60863292 0.107696631
0.376206676
PNN 6262.456289 0.026948681 0.276189907 0.09757301 0.922271357 0.970640219
PNO1 738.6932178 0.268954416 0.262260958 1.025522131 0.305116888 0.615135352
PNP 2094.947985 -0.268033945 0.305845536 -0.876370302 0.380828758
0.682522514
PNPLA2 629.2529554 -0.583716785 0.376930367 -1.548606415
0.121476363 0.402032701

| | | | | | | |
|---|---|---|---|---|---|---|
| PNPLA3 | 254.0406752 | 0.54522711 | 0.51339543 | 1.062002265 | 0.288234657 | 0.601673803 |
| PNPLA4 | 513.8816004 | 0.220272861 | 0.268730887 | 0.819678242 | 0.412399557 | 0.70515068 |
| PNPLA5 | 15.47525015 | 0.138256961 | 0.862670729 | 0.160266202 | 0.872671382 | 0.949330774 |
| PNPLA6 | 1788.477906 | 0.067132756 | 0.365877781 | 0.183484101 | 0.854418193 | 0.941485939 |
| PNPLA7 | 38.59643023 | -0.725681165 | 0.541517984 | -1.340086916 | 0.180217089 | 0.484754911 |
| PNPLA8 | 845.9378118 | -0.19022119 | 0.31998744 | -0.594464553 | 0.552201444 | 0.795643979 |
| PNPO | 358.0619391 | -0.091236261 | 0.382082912 | -0.238786551 | 0.8112711 | 0.920897407 |
| PNPT1 | 1205.22633 | 0.526045941 | 0.285250763 | 1.844152616 | 0.065160902 | 0.294467174 |
| PNRC1 | 1301.089833 | 0.101147347 | 0.456331311 | 0.221653314 | 0.824583776 | 0.92681808 |
| PNRC2 | 6335.013673 | 0.240002823 | 0.23676427 | 1.013678387 | 0.310736241 | 0.620719059 |
| POC1A | 623.9638267 | 0.238929799 | 0.287620604 | 0.830711692 | 0.40613652 | 0.701474651 |
| POC1B | 765.5344797 | -0.261876174 | 0.286267579 | -0.914795085 | 0.360299215 | 0.667522123 |
| POC1B-GALNT4 | 27.58742724 | -4.052199508 | 1.057056279 | -3.833475652 | 0.000126345 | 0.007125594 |
| POC5 | 336.3825343 | 0.179720227 | 0.364486443 | 0.493077946 | 0.621957513 | 0.833640266 |
| PODN | 10.69508245 | -1.016974297 | 1.032032167 | -0.985409496 | 0.324422967 | 0.634500559 |
| PODNL1 | 27.43234082 | 1.342884435 | 0.735293863 | 1.826323464 | 0.067801553 | 0.299425477 |
| PODXL | 9423.759861 | -0.124642992 | 0.625823714 | -0.199166297 | 0.842132662 | 0.935422069 |
| PODXL2 | 238.8136495 | 0.349588774 | 0.518499174 | 0.674232075 | 0.50016378 | 0.763360896 |
| POF1B | 44.54213182 | -3.571737114 | 0.810681411 | -4.40584558 | 1.05E-05 | 0.001149871 |
| POFUT1 | 1444.120156 | -0.033997469 | 0.345415563 | -0.098424833 | 0.921594957 | 0.970399576 |
| POFUT2 | 1056.927592 | -0.044643845 | 0.330030705 | -0.135271792 | 0.892396987 | 0.957550733 |
| POGK | 2020.97929 | -0.249404289 | 0.31329472 | -0.79606924 | 0.425991792 | 0.71395509 |
| POGLUT1 | 489.6447317 | 0.149664995 | 0.291972315 | 0.512599954 | 0.608231185 | 0.82765049 |
| POGZ | 2629.702301 | -0.458846473 | 0.317057934 | -1.447200728 | 0.147840713 | 0.44174616 |
| POLA1 | 1035.588365 | -0.234241839 | 0.456676407 | -0.512927393 | 0.608002111 | 0.827607804 |
| POLA2 | 640.0346121 | 0.012602969 | 0.331717669 | 0.037993058 | 0.969693217 | 0.989496332 |
| POLB | 1081.942899 | 0.478130964 | 0.27759144 | 1.722426904 | 0.084992214 | 0.334045822 |
| POLD1 | 1124.269051 | 0.003238527 | 0.575831438 | 0.005624089 | 0.99551265 | 0.998116797 |
| POLD2 | 2332.045331 | 0.355679124 | 0.274190637 | 1.297196463 | 0.194563598 | 0.500370345 |
| POLD3 | 1079.122763 | 0.28096055 | 0.285861255 | 0.982856351 | 0.325678143 | 0.635672141 |
| POLD4 | 231.4726095 | 0.010514534 | 0.46696942 | 0.022516537 | 0.982035921 | 0.993518428 |
| POLDIP2 | 2109.112394 | -0.174514948 | 0.223689914 | -0.78016458 | 0.435294008 | 0.720471896 |
| POLDIP3 | 3615.579657 | -0.282796568 | 0.375190994 | -0.753740287 | 0.451005186 | 0.732208864 |
| POLE | 2077.641442 | -0.330610924 | 0.443396139 | -0.745633295 | 0.455888964 | 0.735306344 |
| POLE2 | 466.7384523 | 0.716150807 | 0.254398156 | 2.815078609 | 0.00487653 | 0.074653726 |
| POLE3 | 1486.59675 | 0.393017066 | 0.277915325 | 1.414161186 | 0.157314581 | 0.456681882 |
| POLE4 | 343.7538331 | 0.9523772 | 0.468852995 | 2.031291706 | 0.042225412 | 0.236528876 |
| POLG | 1447.593471 | -0.276231453 | 0.302279073 | -0.913829232 | 0.360806584 | 0.667943464 |
| POLG2 | 486.8564958 | 0.418865454 | 0.276705973 | 1.513756458 | 0.13008762 | 0.415181944 |
| POLH | 349.4423104 | 0.100585132 | 0.639383396 | 0.157315834 | 0.874995942 | 0.949740014 |

| | | | | | | |
|---|---|---|---|---|---|---|
| POLI | 1729.061449 | -0.575304004 | 0.249931978 | -2.301842323 | 0.021344066 | 0.165801753 |
| POLK | 399.6950528 | 0.099466552 | 0.419896839 | 0.236883308 | 0.81274732 | 0.921745503 |
| POLL | 1026.856794 | -0.278605245 | 0.295421187 | -0.943078077 | 0.345640968 | 0.652517297 |
| POLM | 585.8706671 | -0.43282959 | 0.276571314 | -1.564983673 | 0.117586738 | 0.395641036 |
| POLN | 27.06349554 | -0.629888152 | 0.832257622 | -0.756842755 | 0.449144075 | 0.730485072 |
| POLQ | 409.6831738 | 0.084903354 | 0.452612377 | 0.187585135 | 0.85120188 | 0.939477675 |
| POLR1A | 1403.126576 | -0.451557879 | 0.460277165 | -0.981056443 | 0.326564908 | 0.636392578 |
| POLR1B | 753.6524644 | 0.081162351 | 0.395625109 | 0.205149646 | 0.83745521 | 0.93337004 |
| POLR1C | 786.5665558 | 0.398635808 | 0.357190882 | 1.116030192 | 0.264409211 | 0.577921357 |
| POLR1D | 3808.736173 | 0.453633025 | 0.410068479 | 1.106237247 | 0.268623845 | 0.581542504 |
| POLR1E | 980.4743061 | 0.022243516 | 0.239503125 | 0.092873595 | 0.926003983 | 0.972261514 |
| POLR2A | 4114.649533 | -0.078787863 | 0.2659643 | -0.296234731 | 0.767050827 | 0.903470997 |
| POLR2B | 4547.14243 | -0.135149144 | 0.302718071 | -0.446452183 | 0.655270646 | 0.849449796 |
| POLR2C | 2000.622661 | -0.064351335 | 0.224429713 | -0.286732687 | 0.774317007 | 0.906279513 |
| POLR2D | 1138.90972 | 0.373396473 | 0.286615273 | 1.302779399 | 0.192650086 | 0.498838446 |
| POLR2E | 2219.395138 | 0.108983579 | 0.347984921 | 0.313184776 | 0.754140284 | 0.897293999 |
| POLR2F | 1212.025607 | 0.902551635 | 0.422084884 | 2.138317834 | 0.032490956 | 0.205554752 |
| POLR2G | 1597.925493 | 0.843889876 | 0.402485933 | 2.096694088 | 0.036020664 | 0.217061721 |
| POLR2H | 1845.558005 | 1.080581191 | 0.315382953 | 3.426251102 | 0.000611974 | 0.020095807 |
| POLR2I | 932.6176506 | 0.859984802 | 0.434175048 | 1.980732901 | 0.047621235 | 0.251378482 |
| POLR2J | 935.9454302 | 0.45922712 | 0.386697714 | 1.187560989 | 0.235006415 | 0.545567302 |
| POLR2J2 | 59.39355646 | -0.779088655 | 0.540787804 | -1.440655004 | 0.149682173 | 0.444386222 |
| POLR2J3 | 1544.492714 | -0.221736581 | 0.34193832 | -0.64846953 | 0.516681312 | 0.773950651 |
| POLR2J4 | 98.27231427 | -0.247574984 | 0.416897114 | -0.593851517 | 0.552611428 | 0.795662343 |
| POLR2K | 1355.980315 | 1.100392652 | 0.33097104 | 3.324739989 | 0.00088501 | 0.025285294 |
| POLR2L | 939.0221181 | 1.10204612 | 0.464524791 | 2.37241616 | 0.017672173 | 0.150008791 |
| POLR2M | 1624.257504 | 0.042733101 | 0.291268691 | 0.146713679 | 0.883358021 | 0.953770161 |
| POLR3A | 1406.532119 | -0.289490139 | 0.352572512 | -0.821079718 | 0.411600862 | 0.704686727 |
| POLR3B | 531.5373312 | 0.019178545 | 0.326121745 | 0.058807929 | 0.953105093 | 0.983108339 |
| POLR3C | 1054.010505 | -0.550068124 | 0.255023264 | -2.156933121 | 0.031010876 | 0.200870165 |
| POLR3D | 1477.593474 | 0.217506813 | 0.30750628 | 0.707324786 | 0.479364666 | 0.751673525 |
| POLR3E | 542.5989622 | 0.033251157 | 0.294410916 | 0.112941318 | 0.910077079 | 0.966444934 |
| POLR3F | 502.6734289 | -0.011081428 | 0.248410015 | -0.044609425 | 0.96441863 | 0.988527467 |

| | | | | | | |
|---|---|---|---|---|---|---|
| POLR3G | 349.7083387 | 0.488244805 | 0.277449125 | 1.759763364 | 0.078447937 | 0.321397152 |
| POLR3GL | 359.8364383 | 1.277604941 | 0.401029682 | 3.185811423 | 0.001443487 | 0.034541221 |
| POLR3H | 844.4423809 | 0.193173009 | 0.26531934 | 0.728077377 | 0.466566221 | 0.743017201 |
| POLR3K | 699.8354169 | 1.55783139 | 0.470317312 | 3.312298631 | 0.000925327 | 0.02597297 |
| POLRMT | 993.4759109 | 0.388078696 | 0.423384136 | 0.916611331 | 0.359346345 | 0.667030882 |
| POLRMTP1 | 11.12077363 | 0.047599862 | 1.154485791 | 0.041230357 | 0.967112253 | 0.988749915 |
| POM121 | 1760.956951 | -0.493306382 | 0.372567297 | -1.324073224 | 0.185478727 | 0.490031772 |
| POM121B | 15.02695042 | -0.33253643 | 1.28598385 | -0.25858523 | 0.79595528 | 0.915299455 |
| POM121C | 1818.709474 | -0.162961153 | 0.252544563 | -0.645276822 | 0.51874781 | 0.774446273 |
| POMGNT1 | 1852.851682 | -0.005242233 | 0.250567689 | -0.020921426 | 0.983308335 | 0.993789271 |
| POMGNT2 | 334.9888937 | 0.358673431 | 0.324404489 | 1.105636461 | 0.268883902 | 0.581986006 |
| POMK | 47.4391465 | -0.784477559 | 0.621508716 | -1.262214895 | 0.20687147 | 0.513642439 |
| POMP | 2736.47563 | 0.931116296 | 0.367316209 | 2.534917526 | 0.01124738 | 0.120339982 |
| POMT1 | 1275.457637 | -0.227275767 | 0.255999414 | -0.887797997 | 0.37464942 | 0.678987738 |
| POMT2 | 935.8419352 | -0.624251874 | 0.357413759 | -1.746580422 | 0.080710146 | 0.325104967 |
| POMZP3 | 197.6477266 | 0.38235426 | 0.37991357 | 1.006424328 | 0.314211496 | 0.623858474 |
| PON1 | 34.5069138 | 0.609115715 | 0.543747894 | 1.120217147 | 0.262621238 | 0.57612824 |
| PON2 | 1203.817831 | 0.46505285 | 0.231582801 | 2.008149347 | 0.044627424 | 0.242841695 |
| POP1 | 349.2168899 | 0.117956785 | 0.376931724 | 0.312939394 | 0.754326708 | 0.897419266 |
| POP4 | 652.1595595 | 0.275244102 | 0.265485718 | 1.036756719 | 0.299849251 | 0.610555175 |
| POP5 | 655.622033 | 0.951986497 | 0.310699194 | 3.064013412 | 0.002183891 | 0.044412618 |
| POP7 | 743.3523581 | 1.084546157 | 0.469154115 | 2.311705521 | 0.020793917 | 0.163514486 |
| POPDC2 | 23.22149544 | -1.211881162 | 0.885656101 | -1.368342815 | 0.171204795 | 0.474474162 |
| POPDC3 | 29.16340448 | -1.05381154 | 0.614593349 | -1.71464846 | 0.086409698 | 0.337171176 |
| POR | 1495.404383 | 0.032839506 | 0.340822361 | 0.096353731 | 0.923239639 | 0.971120333 |
| PORCN | 229.6093193 | 0.451369702 | 0.438426428 | 1.02952211 | 0.303234395 | 0.613656879 |
| POSTN | 644.3005627 | -3.266408044 | 0.649710609 | -5.027481466 | 4.97E-07 | 0.000120267 |
| POT1 | 1281.348258 | 0.193442764 | 0.278404097 | 0.694827289 | 0.487163544 | 0.756439734 |
| POTEC | 13.42981439 | -0.59549545 | 1.129277936 | -0.527324081 | 0.597968554 | 0.822688033 |
| POU2AF1 | 21.66402149 | -0.268055325 | 0.649091532 | -0.412969993 | 0.679628599 | 0.861682789 |
| POU2F1 | 2238.36022 | -0.184981618 | 0.319360274 | -0.579225512 | 0.562437019 | 0.801706045 |
| POU2F2 | 144.2517547 | 1.028842397 | 0.472411633 | 2.177851528 | 0.029417092 | 0.195997675 |
| POU2F3 | 45.62939099 | -2.737457982 | 0.930156369 | -2.943008372 | 0.003250396 | 0.057339258 |
| POU3F1 | 161.4289513 | 1.978452299 | 0.780684286 | 2.534254032 | 0.011268701 | 0.12047742 |
| POU3F2 | 194.8210945 | 3.042902789 | 1.583952081 | 1.921082604 | 0.054721294 | 0.270592163 |
| POU4F1 | 19.00560816 | -2.533701172 | 0.861842324 | -2.939866261 | 0.003283539 | 0.057789532 |
| POU5F1 | 668.7012605 | -0.799824046 | 0.408562464 | -1.957654252 | 0.050270597 | 0.257653791 |

| | | | | | | |
|---|---|---|---|---|---|---|
| POU5F1B | 37.51778176 | -0.2425078 | 0.583919888 | -0.415310055 | 0.677914942 | 0.860856079 |
| POU5F2 | 15.75017718 | 0.453084523 | 0.903788724 | 0.501316858 | 0.616148143 | 0.831265761 |
| POU6F2 | 323.0818577 | -0.273905879 | 0.352331255 | -0.777410109 | 0.43691685 | 0.721033997 |
| PPA1 | 4587.957812 | 0.581976612 | 0.317074569 | 1.835456603 | 0.066438064 | 0.297462837 |
| PPA2 | 2413.432427 | 0.878194784 | 0.403558702 | 2.176126496 | 0.029545803 | 0.196251918 |
| PPAN | 618.3651175 | 0.470985555 | 0.321409018 | 1.465377537 | 0.142817943 | 0.433233718 |
| PPAN-P2RY11 | 24.26995499 | -1.12726526 | 0.821125271 | -1.372829823 | 0.169805257 | 0.472495848 |
| PPARA | 226.2791645 | -0.536158185 | 0.372896775 | -1.437819314 | 0.150485328 | 0.445342388 |
| PPARD | 583.1167709 | -0.706408227 | 0.385672864 | -1.831625434 | 0.067007245 | 0.297817842 |
| PPARG | 124.5443954 | -2.083231536 | 0.880658724 | -2.365537839 | 0.018003903 | 0.151322559 |
| PPARGC1A | 33.30565271 | 0.023332214 | 0.616927361 | 0.037820034 | 0.969831171 | 0.989496332 |
| PPARGC1B | 102.0395927 | 0.741273368 | 0.563851239 | 1.314661239 | 0.188623779 | 0.493962743 |
| PPAT | 2205.080378 | 0.417503742 | 0.290758455 | 1.435912644 | 0.1510272 | 0.44619129 |
| PPCDC | 475.4816282 | 0.300309926 | 0.385954287 | 0.778097139 | 0.43651175 | 0.720784523 |
| PPCS | 898.4270517 | 0.249969869 | 0.443730003 | 0.563337767 | 0.573204906 | 0.807961686 |
| PPDPF | 1278.801102 | 0.105662299 | 0.443333876 | 0.238335722 | 0.811620717 | 0.921156423 |
| PPEF1 | 12.31729071 | 0.596001804 | 1.125005275 | 0.529776897 | 0.596266625 | 0.821812117 |
| PPFIA1 | 1404.022769 | -0.257767324 | 0.459187202 | -0.561355637 | 0.57455512 | 0.808359708 |
| PPFIA2 | 10.10584075 | 0.543769439 | 1.104726425 | 0.492220904 | 0.622563188 | 0.833766431 |
| PPFIA3 | 347.489297 | 0.085236974 | 0.43960559 | 0.1938942 | 0.846258725 | 0.938145276 |
| PPFIA4 | 72.75557794 | 0.654220973 | 0.74610071 | 0.876853438 | 0.380566249 | 0.682213231 |
| PPFIBP1 | 1006.765183 | -0.952923807 | 0.342211053 | -2.784608499 | 0.005359239 | 0.077809098 |
| PPFIBP2 | 247.3666346 | -1.604679445 | 0.51417673 | -3.120871391 | 0.001803168 | 0.039360945 |
| PPHLN1 | 1811.941238 | 0.055340443 | 0.219589645 | 0.252017546 | 0.801027502 | 0.916680548 |
| PPIA | 29731.66358 | 0.814584717 | 0.365240912 | 2.2302669 | 0.025729729 | 0.183003152 |
| PPIAP14 | 12.13104298 | 1.147870596 | 0.945026555 | 1.214643747 | 0.224501996 | 0.533874112 |
| PPIAP46 | 35.23081128 | 0.153218771 | 0.589719762 | 0.25981624 | 0.795005523 | 0.914961978 |
| PPIAP53 | 14.50855273 | 0.483705066 | 1.056936737 | 0.457648078 | 0.647205296 | 0.846597949 |
| PPIB | 6652.43903 | 0.211641001 | 0.411392395 | 0.514450446 | 0.606937098 | 0.826788682 |
| PPIC | 47.93571993 | -0.896213048 | 0.915119918 | -0.979339461 | 0.327412279 | 0.637370783 |
| PPID | 1422.355073 | 0.16769124 | 0.290689956 | 0.576873183 | 0.564025127 | 0.802433524 |
| PPIE | 1204.49058 | 0.366715119 | 0.266589603 | 1.375579222 | 0.168951943 | 0.471508612 |
| PPIEL | 17.04230558 | 0.183818434 | 0.728303758 | 0.252392538 | 0.800737668 | 0.916680548 |
| PPIF | 1174.423136 | 0.544271869 | 0.320844954 | 1.696370359 | 0.089815763 | 0.343136476 |
| PPIG | 3979.044881 | 0.628303292 | 0.345514214 | 1.818458596 | 0.068994069 | 0.302794854 |
| PPIH | 1106.192223 | 0.644127034 | 0.323120236 | 1.993459284 | 0.046211178 | 0.247290927 |
| PPIL1 | 1731.104414 | 0.498300789 | 0.26200506 | 1.901874681 | 0.05718754 | 0.276743914 |
| PPIL2 | 1221.824524 | -0.579299786 | 0.254902141 | -2.272636013 | 0.023048124 | 0.173530066 |
| PPIL3 | 1100.049941 | 0.388968768 | 0.285220906 | 1.363745644 | 0.172647633 | 0.476336299 |
| PPIL4 | 1099.422801 | 0.002209149 | 0.260271389 | 0.008487866 | 0.993227744 | 0.997303469 |
| PPIL6 | 37.79622453 | 1.442026247 | 0.551195138 | 2.616181001 | 0.008891941 | 0.10668882 |
| PPIP5K1 | 785.6775277 | -0.712054297 | 0.436168893 | -1.632519672 | 0.102570058 | 0.367636293 |

| | | | | | | |
|---|---|---|---|---|---|---|
| PPIP5K2 | 1915.996875 | -0.093804651 | 0.3532153 | -0.26557358 | 0.790567636 | 0.91311918 |
| PPL | 3364.647419 | -1.587863868 | 0.324936889 | -4.886683909 | 1.03E-06 | 0.00018521 |
| PPM1A | 1153.895705 | 0.07950774 | 0.234887434 | 0.338492949 | 0.73499174 | 0.889483064 |
| PPM1B | 1741.561114 | 0.148384395 | 0.256581662 | 0.578312549 | 0.563053123 | 0.801847267 |
| PPM1D | 561.8454581 | 0.208369249 | 0.349475735 | 0.596233811 | 0.551019042 | 0.794554689 |
| PPM1E | 220.1780799 | 0.967169022 | 0.613120181 | 1.577454229 | 0.114691046 | 0.389779986 |
| PPM1F | 671.9774664 | 0.22513432 | 0.361674153 | 0.622478323 | 0.533627393 | 0.783156734 |
| PPM1G | 4645.676213 | 0.122525906 | 0.241705806 | 0.506921652 | 0.612209793 | 0.829098987 |
| PPM1H | 1524.35534 | -1.638817666 | 0.448977901 | -3.650107637 | 0.00026213 | 0.011279719 |
| PPM1J | 315.8983224 | -0.218607674 | 0.469199896 | -0.465915863 | 0.641275726 | 0.843932467 |
| PPM1K | 368.91543 | 0.319771725 | 0.298536979 | 1.071129367 | 0.284111265 | 0.598230674 |
| PPM1L | 947.9243558 | -0.509283618 | 0.311073786 | -1.637179477 | 0.101592974 | 0.365596699 |
| PPM1M | 140.3182083 | -0.551026852 | 0.391360617 | -1.407977269 | 0.159137801 | 0.459030918 |
| PPM1N | 272.4393228 | 0.346878685 | 0.642382386 | 0.539987853 | 0.589205409 | 0.818256545 |
| PPME1 | 1290.679896 | -0.44920883 | 0.317377893 | -1.4153753 | 0.156958488 | 0.456084508 |
| PPOX | 554.4478322 | -0.001871044 | 0.285446117 | -0.006554805 | 0.994770059 | 0.997993425 |
| PPP1CA | 3462.861278 | -0.198239213 | 0.299519446 | -0.66185757 | 0.508062506 | 0.768706122 |
| PPP1CB | 7351.568709 | 0.19681647 | 0.308992923 | 0.636961094 | 0.524150186 | 0.777662198 |
| PPP1CC | 13806.36576 | 0.389821257 | 0.266773679 | 1.461243319 | 0.143948679 | 0.434995877 |
| PPP1R10 | 422.3572013 | -0.700718841 | 1.953522671 | -0.358695014 | 0.719823259 | 0.881675978 |
| PPP1R12A | 2734.207774 | 0.124083821 | 0.257690099 | 0.481523433 | 0.630144529 | 0.838075099 |
| PPP1R12B | 423.2224951 | -0.282297931 | 0.420386342 | -0.671520224 | 0.501889181 | 0.764473972 |
| PPP1R12C | 1113.964685 | 0.126667249 | 0.44024048 | 0.287722859 | 0.773558891 | 0.906158747 |
| PPP1R13B | 206.910358 | -1.708885555 | 0.476724516 | -3.584639549 | 0.000337544 | 0.013547304 |
| PPP1R13L | 603.3458674 | -1.191449919 | 0.596507185 | -1.997377315 | 0.04578421 | 0.246484121 |
| PPP1R14A | 79.95758576 | 1.285880385 | 0.595753292 | 2.158410878 | 0.030895901 | 0.200651417 |
| PPP1R14B | 1669.962977 | 1.143026265 | 0.329977475 | 3.463952394 | 0.000532301 | 0.018147267 |
| PPP1R14C | 38.32730025 | 1.91219024 | 0.88009315 | 2.172713468 | 0.029801889 | 0.197226575 |
| PPP1R15A | 1158.977292 | -0.753603615 | 0.343429926 | -2.194344634 | 0.028210643 | 0.192229399 |
| PPP1R15B | 1155.901463 | 0.219785463 | 0.282390945 | 0.778302092 | 0.436390942 | 0.720784523 |
| PPP1R16A | 308.3310115 | -0.422440317 | 0.594547322 | -0.710524296 | 0.47737907 | 0.749955252 |
| PPP1R16B | 22.42177446 | 2.355934357 | 1.01301296 | 2.325670499 | 0.020036135 | 0.160028741 |
| PPP1R18 | 780.405026 | -0.208635937 | 0.619446309 | -0.336810364 | 0.73625986 | 0.890378504 |
| PPP1R1A | 255.4659272 | 0.899417269 | 0.454179449 | 1.980312562 | 0.047668418 | 0.251480612 |
| PPP1R1B | 38.35829489 | 1.911727071 | 1.073802122 | 1.7803346 | 0.075021218 | 0.314480446 |
| PPP1R1C | 186.2920961 | -0.341385805 | 0.805856324 | -0.423631105 | 0.671834867 | 0.858676204 |
| PPP1R2 | 1504.068877 | -0.177930816 | 0.348976059 | -0.509865395 | | |

| | | | | | | |
|---|---|---|---|---|---|---|
| | | | | | 0.610145767 | 0.828605673 |
| PPP1R21 | 779.3319057 | -0.721961795 | 0.23938899 | -3.015852135 | 0.002562582 | 0.049092812 |
| PPP1R26 | 870.0929488 | -0.429722002 | 0.334883696 | -1.283197741 | 0.199422807 | 0.505419881 |
| PPP1R32 | 118.2859356 | 0.43022925 | 0.401982917 | 1.070267497 | 0.28449892 | 0.598465491 |
| PPP1R35 | 399.2405916 | 1.244394358 | 0.502557873 | 2.476121506 | 0.013281837 | 0.13092836 |
| PPP1R37 | 500.6280564 | -0.229315566 | 0.482904077 | -0.474867737 | 0.634881248 | 0.840313679 |
| PPP1R3B | 1163.197696 | -1.467199123 | 0.508445575 | -2.885656195 | 0.003905986 | 0.064691825 |
| PPP1R3C | 66.19745173 | 1.98864898 | 0.782695185 | 2.540770683 | 0.011060844 | 0.11934855 |
| PPP1R3D | 75.06969571 | -0.28312678 | 0.519169143 | -0.545345932 | 0.585515619 | 0.815614494 |
| PPP1R3E | 119.1361637 | 0.628736458 | 0.497587175 | 1.263570465 | 0.20638424 | 0.512832712 |
| PPP1R3F | 206.5720995 | 0.984639408 | 0.356200976 | 2.764280485 | 0.005704848 | 0.081265958 |
| PPP1R7 | 1159.356446 | 0.115549073 | 0.271083765 | 0.426248592 | 0.669926713 | 0.857762448 |
| PPP1R8 | 1753.518807 | 0.107078849 | 0.233944546 | 0.457710389 | 0.647160523 | 0.846597949 |
| PPP1R9A | 534.8701135 | -0.513700665 | 0.39950186 | -1.285853 | 0.198494369 | 0.504246161 |
| PPP1R9B | 1136.566745 | -0.248181959 | 0.403710841 | -0.614751781 | 0.538718657 | 0.786598616 |
| PPP2CA | 5263.58337 | 0.223358779 | 0.239981666 | 0.93073268 | 0.351991861 | 0.65929447 |
| PPP2CB | 3084.206213 | 0.520911194 | 0.315960828 | 1.648657519 | 0.099217818 | 0.361452832 |
| PPP2R1A | 10861.87411 | 0.167362877 | 0.291736496 | 0.57367823 | 0.566185562 | 0.804380669 |
| PPP2R1B | 1944.710614 | -0.096053081 | 0.236383531 | -0.406344218 | 0.6844897 | 0.863857185 |
| PPP2R2A | 2065.121687 | 0.049387443 | 0.24844571 | 0.198785654 | 0.842430419 | 0.935547316 |
| PPP2R2B | 59.93992699 | 1.201911335 | 0.650746566 | 1.846972995 | 0.064751054 | 0.293666076 |
| PPP2R2C | 54.38002189 | 0.595543638 | 0.576544301 | 1.03295382 | 0.301625511 | 0.61241443 |
| PPP2R2D | 705.0093313 | -0.02155122 | 0.335343152 | -0.064266171 | 0.948758289 | 0.98134156 |
| PPP2R3A | 986.5261226 | -0.688900106 | 0.269998963 | -2.55149167 | 0.010726289 | 0.117687513 |
| PPP2R3B | 97.262179 | -0.159494101 | 0.552772648 | -0.28853472 | 0.772937457 | 0.905717252 |
| PPP2R3C | 723.3300996 | 0.5520543 | 0.255390547 | 2.161608202 | 0.030648389 | 0.199470916 |
| PPP2R5A | 1284.01292 | -0.211206589 | 0.304046007 | -0.694653389 | 0.487272545 | 0.756439734 |
| PPP2R5B | 157.2238174 | -0.368664515 | 0.546243065 | -0.674909282 | 0.499733403 | 0.762952144 |
| PPP2R5C | 3251.52457 | 0.088663799 | 0.226123469 | 0.392103481 | 0.694981756 | 0.869142545 |
| PPP2R5D | 2122.775604 | 0.008245795 | 0.312506093 | 0.026386029 | 0.978949437 | 0.992823177 |
| PPP2R5E | 2193.629987 | -0.085218799 | 0.247495687 | -0.344324381 | 0.730602351 | 0.886859359 |
| PPP3CA | 1050.483887 | 0.042583995 | 0.305188044 | 0.139533627 | 0.889028485 | 0.956217735 |
| PPP3CB | 1619.466908 | 0.036726071 | 0.233013571 | 0.157613441 | 0.874761411 | |

0.949740014

| | | | | | | |
|---|---|---|---|---|---|---|
| PPP3CC | 199.390483 | -0.223054792 | 0.463692415 | -0.481040415 | 0.630487775 | 0.838177265 |
| PPP3R1 | 1506.908057 | 0.125120221 | 0.246803766 | 0.506962365 | 0.612181226 | 0.829098987 |
| PPP4C | 3164.650505 | 0.380840747 | 0.371046391 | 1.026396578 | 0.30470469 | 0.614996026 |
| PPP4R1 | 2294.427385 | -0.311727158 | 0.260678571 | -1.195829624 | 0.231763053 | 0.540934965 |
| PPP4R1L | 326.3753287 | -0.097806572 | 0.399109125 | -0.24506223 | 0.806408247 | 0.919090941 |
| PPP4R2 | 3330.760977 | 0.238483919 | 0.229397406 | 1.039610352 | 0.298520965 | 0.609920825 |
| PPP4R3A | 2440.479223 | -0.213113485 | 0.25132413 | -0.847962687 | 0.396458751 | 0.695257856 |
| PPP4R3B | 1997.730952 | -0.05980225 | 0.261093491 | -0.229045349 | 0.818833669 | 0.924316427 |
| PPP4R4 | 87.37829042 | -2.114465566 | 1.183366598 | -1.786822079 | 0.07396625 | 0.312595272 |
| PPP5C | 1811.236755 | -0.123777078 | 0.393097571 | -0.314876222 | 0.752855636 | 0.896586461 |
| PPP5D1 | 13.35405377 | 0.409478798 | 0.804433159 | 0.509027747 | 0.610732775 | 0.829046791 |
| PPP6C | 1756.857631 | 0.235582238 | 0.338618744 | 0.695715291 | 0.486607147 | 0.756340124 |
| PPP6R1 | 1437.11835 | -0.292395835 | 0.476948472 | -0.613055398 | 0.539839711 | 0.787188413 |
| PPP6R2 | 1386.274886 | -0.254001829 | 0.394713212 | -0.643509821 | 0.519893344 | 0.775088632 |
| PPP6R3 | 3038.697772 | -0.304820146 | 0.323729718 | -0.941588396 | 0.346403413 | 0.653093035 |
| PPRC1 | 1165.363746 | -0.307227002 | 0.397833629 | -0.772249956 | 0.439966399 | 0.7234571 |
| PPT1 | 4292.021009 | -0.042979416 | 0.253444544 | -0.169581145 | 0.865339552 | 0.945250148 |
| PPT2 | 142.2831284 | -1.478502235 | 0.653847247 | -2.26123493 | 0.023744713 | 0.176324544 |
| PPT2-EGFL8 | 27.16778898 | -2.585258223 | 0.818019856 | -3.160385661 | 0.001575604 | 0.036831268 |
| PPTC7 | 507.2699793 | -0.001433258 | 0.325135876 | -0.004408182 | 0.996482791 | 0.998431718 |
| PPWD1 | 1341.740405 | -0.180267742 | 0.30536549 | -0.590334363 | 0.554966506 | 0.796360966 |
| PQBP1 | 1378.221345 | 0.038582211 | 0.268863638 | 0.143501038 | 0.885894491 | 0.954877821 |
| PQLC1 | 502.9879545 | 0.445806917 | 0.425274128 | 1.048281303 | 0.294509021 | 0.606702791 |
| PQLC2 | 309.578286 | -0.073972577 | 0.360376177 | -0.205264892 | 0.837365172 | 0.933338258 |
| PQLC2L | 13.42457893 | 0.463959826 | 0.889647778 | 0.521509565 | 0.602011844 | 0.825021776 |
| PQLC3 | 471.3121668 | -0.442608779 | 0.344580118 | -1.284487279 | 0.198971512 | 0.504782074 |
| PRADC1 | 172.3717127 | 0.999287223 | 0.316275604 | 3.159545698 | 0.001580153 | 0.036880775 |
| PRAF2 | 379.203998 | 0.620216589 | 0.379045817 | 1.636257574 | 0.101785691 | 0.365921972 |
| PRAG1 | 850.7701032 | -0.738712698 | 0.29547485 | -2.500086551 | 0.012416297 | 0.126591155 |
| PRAM1 | 13.54257382 | 0.794002471 | 0.795906264 | 0.997608019 | 0.318469471 | 0.628475744 |
| PRAP1 | 32.63587473 | 0.7218797 | 0.690177925 | 1.045932756 | 0.29559208 | 0.60699582 |
| PRB3 | 12.4416904 | -1.366317535 | 1.569407456 | -0.870594522 | 0.383975588 | 0.684699412 |
| PRC1 | 1854.713618 | 0.701009794 | 0.364475448 | 1.923338866 | 0.054437507 | 0.269716333 |
| PRCC | 1889.699154 | -0.049430196 | 0.300375282 | -0.164561465 | 0.869289164 | 0.947227711 |
| PRCP | 2801.915971 | -0.254442526 | 0.303625921 | -0.838013188 | 0.402023297 | 0.699116854 |
| PRDM1 | 628.9173249 | -0.532260809 | 0.555635347 | -0.957931873 | 0.338097113 | |

0.645353712
PRDM10 573.036841 -0.116139845 0.295773327 -0.392665039
0.694566896 0.86892666
PRDM11 190.9916778 -0.764533102 0.457092298 -1.672601148
0.094405841 0.352332352
PRDM12 130.7252285 1.90684839 0.910513379 2.09425631 0.036237154
0.217983293
PRDM14 84.18672829 -1.268920029 0.687377609 -1.846030497
0.064887777 0.293855407
PRDM15 432.8565282 -0.06813192 0.412355682 -0.165226097 0.868766025
0.947046732
PRDM2 1126.554304 -0.40145376 0.328822107 -1.220884336 0.222129816
0.531066116
PRDM4 1033.871545 -0.591337348 0.302858873 -1.952517824 0.050876757
0.259366654
PRDM5 502.7886391 0.033066365 0.345095123 0.095818118 0.923665027 0.971365737
PRDX1 17421.90664 0.724713999 0.332331547 2.180695772 0.029205926 0.195546191
PRDX2 6622.294449 0.559380388 0.33173128 1.68624553 0.091748518 0.347684361
PRDX3 5896.882744 0.480114279 0.228669112 2.099602675 0.035763807 0.21633681
PRDX4 1992.486309 0.657800326 0.341966423 1.923581623 0.054407046 0.269716333
PRDX5 4572.133384 0.709020527 0.45873351 1.545604391 0.12220014 0.40285432
PRDX6 5802.265995 0.442586721 0.42286385 1.046641185 0.295265099 0.60699582
PREB 1284.300556 0.18651805 0.231372571 0.806137262 0.420163709 0.710703939
PRELID1 2049.138684 0.644547229 0.292743524 2.201747181 0.027683172
0.190295153
PRELID2 83.7261666 0.881441899 0.463270746 1.9026496 0.057086282
0.276551437
PRELID3A 100.7994314 0.416528486 0.4759342 0.875180827 0.381475524
0.682984572
PRELID3B 1884.825242 0.531930754 0.297342821 1.788947692 0.07362324
0.311651561
PRELP 37.82393882 0.107575634 0.700456036 0.153579423 0.877941362 0.951103142
PREP 1533.249256 -0.186940099 0.242118489 -0.772101708 0.44005419
0.7234571
PREPL 1563.972112 0.279459779 0.344939327 0.810170825 0.417842003 0.709307489
PREX1 522.8916198 0.168362602 0.417748241 0.403024084 0.686930507 0.865854331
PREX2 822.9239297 0.268223882 0.531909359 0.504266145 0.614074364 0.830316752
PRH1 19.71485699 0.691041091 0.72073615 0.958798986 0.33766002 0.645098755
PRICKLE1 511.3634257 -0.578474233 0.340701336 -1.697892473
0.089528058 0.342777615
PRICKLE2 378.4967391 -0.189520561 0.327871595 -0.578032874
0.563241925 0.801890319
PRICKLE3 290.5823291 -0.414858304 0.271498658 -1.52803077 0.126504898
0.409299598
PRICKLE4 18.01112401 -5.371130823 1.692826434 -3.172877453
0.001509362 0.035778959
PRIM1 1108.324803 0.593238291 0.274270883 2.162964895 0.03054388 0.199123233
PRIM2 970.8688828 0.099388274 0.237499312 0.418478158 0.675597553 0.860110127
PRIMA1 37.8206362 -0.453701305 0.646928903 -0.701315559
0.483106106 0.753955635
PRIMPOL 267.1385567 0.354036362 0.333613373 1.061217537 0.288591051
0.601816473
PRKAA1 965.6647108 -0.605653566 0.277313495 -2.184003219
0.028962012 0.194675532
PRKAA2 683.1485348 -0.60396122 0.329249335 -1.834358207 0.066600839
0.297701125
PRKAB1 788.5739453 -0.297444293 0.260521547 -1.141726266
0.253567819 0.565942797
PRKAB2 1460.019233 0.159200238 0.322832256 0.493136094 0.621916429
0.833640266
PRKACA 1291.622213 -0.367120189 0.37882748 -0.969095984
0.332497301 0.640222942
PRKACB 799.2344442 -0.007784306 0.287464061 -0.027079232
0.978396539 0.992395125

| | | | | | | |
|---|---|---|---|---|---|---|
| PRKAG1 | 1672.178732 | 0.067641811 | 0.236758958 | 0.285699055 | 0.775108628 | 0.906557532 |
| PRKAG2 | 518.7175711 | 0.147685929 | 0.285861115 | 0.516635253 | 0.605410802 | 0.826077544 |
| PRKAR1A | 2954.458599 | 0.094944572 | 0.291255366 | 0.325983941 | 0.744436505 | 0.892725315 |
| PRKAR1B | 235.6768167 | -0.298899875 | 0.408225527 | -0.732193004 | 0.464050774 | 0.741119432 |
| PRKAR2A | 1113.005601 | 0.16234523 | 0.283046956 | 0.573562889 | 0.566263631 | 0.804380669 |
| PRKAR2B | 328.1595666 | 0.329096449 | 0.405082527 | 0.812418277 | 0.416551662 | 0.708268589 |
| PRKCA | 382.7734984 | 0.291454191 | 0.54980919 | 0.530100616 | 0.596042172 | 0.821812117 |
| PRKCB | 268.5303754 | 1.073800882 | 0.596638605 | 1.799750926 | 0.071899976 | 0.308679129 |
| PRKCD | 1395.702199 | 0.17510001 | 0.39282278 | 0.445748105 | 0.655779212 | 0.849746021 |
| PRKCE | 103.2670468 | -1.264246165 | 0.944013463 | -1.339224719 | 0.180497528 | 0.485089104 |
| PRKCG | 202.3800989 | -0.459605041 | 0.342962236 | -1.340103934 | 0.180211557 | 0.484754911 |
| PRKCH | 872.5729174 | -1.302584979 | 0.716799182 | -1.817224421 | 0.069182757 | 0.303081608 |
| PRKCI | 2626.299505 | -0.041185726 | 0.238224685 | -0.172886057 | 0.862740992 | 0.944730333 |
| PRKCQ | 166.9071201 | -1.227112178 | 0.378400367 | -3.242893727 | 0.001183223 | 0.030220003 |
| PRKCSH | 8605.467229 | -0.300366198 | 0.282415648 | -1.063560747 | 0.287527733 | 0.601673803 |
| PRKCZ | 1383.269627 | -0.722590148 | 0.323381242 | -2.234483802 | 0.025451257 | 0.182046687 |
| PRKD1 | 383.9386768 | -1.505058441 | 0.435229639 | -3.45807892 | 0.000544042 | 0.018382319 |
| PRKD2 | 754.7876647 | -0.714354369 | 0.347638271 | -2.054878387 | 0.039890752 | 0.230107455 |
| PRKD3 | 1580.474232 | 0.001568244 | 0.298134473 | 0.005260191 | 0.995802994 | 0.998171604 |
| PRKDC | 10515.74293 | -0.314898222 | 0.435029538 | -0.723854807 | 0.469154878 | 0.744591344 |
| PRKG1 | 1005.645959 | -1.002064291 | 0.351668801 | -2.849454626 | 0.004379425 | 0.069136586 |
| PRKG2 | 82.79293294 | -0.10883108 | 0.386131291 | -0.281849938 | 0.778058578 | 0.908058754 |
| PRKN | 147.8226257 | -0.483764703 | 0.374923459 | -1.290302571 | 0.196945624 | 0.50258876 |
| PRKRA | 1758.436626 | 0.505900337 | 0.296679805 | 1.705206518 | 0.088155915 | 0.341577124 |
| PRKRIP1 | 951.1590302 | 0.144562857 | 0.309725229 | 0.466745501 | 0.640681971 | 0.843724522 |
| PRKX | 1261.057401 | 0.190288732 | 0.338294357 | 0.562494551 | 0.573779115 | 0.808170361 |
| PRKY | 416.3855138 | -0.184619402 | 0.298800002 | -0.617869482 | 0.536661378 | 0.785348759 |
| PRLR | 150.0200343 | -2.860029411 | 0.855908972 | -3.341511195 | 0.000833237 | 0.02432884 |
| PRMT1 | 6890.814987 | 0.615196054 | 0.256967381 | 2.394062823 | 0.016662894 | 0.146056278 |
| PRMT2 | 2011.892685 | -0.245769922 | 0.269519661 | -0.911881237 | 0.36183124 | 0.668713609 |
| PRMT3 | 726.9220693 | 0.545767797 | 0.290824751 | 1.876620871 | 0.060570075 | 0.284105089 |
| PRMT5 | 3447.269524 | 0.1849802 | 0.274006417 | 0.67509441 | 0.499615785 | 0.762849343 |
| PRMT6 | 951.6703703 | 0.19543203 | 0.271808864 | 0.71900536 | 0.472137617 | 0.746435993 |
| PRMT7 | 874.7082553 | -0.318087992 | 0.43257481 | -0.735336374 | 0.462134661 | 0.739795815 |
| PRMT8 | 128.2922516 | 1.782289673 | 0.581420143 | 3.06540751 | 0.002173736 | 0.044324931 |
| PRMT9 | 395.3527497 | -0.079181524 | 0.378078094 | -0.209431663 | 0.834111278 | 0.932107564 |
| PRNP | 292.0565033 | 0.493348394 | 0.410167661 | 1.202796907 | 0.229054922 | 0.537758004 |
| PROB1 | 42.95491896 | 1.02991055 | 0.537354604 | 1.916631109 | 0.055284813 | 0.272191885 |
| PROC | 190.4995082 | 1.67419382 | 1.002881355 | 1.669383733 | 0.095041355 | 0.353833719 |

| | | | | | | |
|---|---|---|---|---|---|---|
| PROCA1 | 426.9416112 | 0.214801718 | 0.273741903 | 0.784687019 | 0.432637096 | 0.718639059 |
| PROCR | 207.9211121 | -0.162654023 | 0.378931114 | -0.429244307 | 0.667745442 | 0.856696248 |
| PRODH | 590.6800888 | -2.016225663 | 0.497212231 | -4.055060473 | 5.01E-05 | 0.003783039 |
| PROKR1 | 15.58753068 | 0.29042689 | 0.946782908 | 0.306751302 | 0.759032678 | 0.899440455 |
| PROM1 | 1042.810609 | -0.461435457 | 0.351116726 | -1.314193895 | 0.188780963 | 0.493962743 |
| PROM2 | 309.1727912 | -2.332263986 | 0.487649006 | -4.782669415 | 1.73E-06 | 0.00027224 |
| PRORSD1P | 18.58356884 | -0.698052061 | 0.873860916 | -0.798813688 | 0.424398452 | 0.713303793 |
| PROS1 | 566.0503819 | -0.215573563 | 0.305274219 | -0.706163669 | 0.480086361 | 0.752036291 |
| PROSER1 | 3073.754324 | -0.486758927 | 0.280724704 | -1.733936918 | 0.082929277 | 0.329609655 |
| PROSER2 | 112.7775147 | -0.396003185 | 0.490957951 | -0.806592875 | 0.41990108 | 0.710456202 |
| PROSER3 | 304.5341079 | -0.204914589 | 0.305742681 | -0.670219113 | 0.502718122 | 0.764968568 |
| PROX1 | 130.0157383 | -1.17945597 | 0.627199632 | -1.880511259 | 0.060038433 | 0.283398591 |
| PRPF18 | 432.765396 | 0.427405955 | 0.300267034 | 1.423419516 | 0.154614578 | 0.452208634 |
| PRPF19 | 3152.129152 | -0.207037619 | 0.316724229 | -0.653684184 | 0.513315295 | 0.772068737 |
| PRPF3 | 1512.533942 | -0.2222363 | 0.322775799 | -0.688515994 | 0.491127902 | 0.758593654 |
| PRPF31 | 1083.524922 | 0.045810689 | 0.353742996 | 0.129502746 | 0.896959853 | 0.95928311 |
| PRPF38A | 2390.305613 | 0.137809864 | 0.25700644 | 0.536211717 | 0.591812237 | 0.819494655 |
| PRPF38B | 2154.441862 | 0.326341209 | 0.285305246 | 1.143831785 | 0.252693405 | 0.565423035 |
| PRPF39 | 1452.795076 | 0.146971666 | 0.319984564 | 0.459308613 | 0.646012563 | 0.84590511 |
| PRPF4 | 1232.56141 | -0.183078964 | 0.269582611 | -0.679120079 | 0.497061781 | 0.761325016 |
| PRPF40A | 6836.746857 | 0.323050639 | 0.235506545 | 1.371726795 | 0.170148504 | 0.47294301 |
| PRPF40B | 982.8689135 | 0.106263757 | 0.324829327 | 0.327137202 | 0.743564115 | 0.892534058 |
| PRPF4B | 3502.217606 | -0.001963398 | 0.303650977 | -0.00646597 | 0.994840939 | 0.997998537 |
| PRPF6 | 3160.181695 | -0.15234467 | 0.256056824 | -0.594964303 | 0.551867333 | 0.795324338 |
| PRPF8 | 9314.000501 | -0.611469301 | 0.484995413 | -1.260773368 | 0.207390511 | 0.514272875 |
| PRPH | 17.8426878 | 0.014565026 | 0.896222235 | 0.016251578 | 0.987033687 | 0.995287969 |
| PRPH2 | 31.019513 | 0.316390712 | 0.640378475 | 0.494068312 | 0.621257936 | 0.833349572 |
| PRPS1 | 1760.550251 | 0.783262165 | 0.357579804 | 2.190454147 | 0.028491317 | 0.192869196 |
| PRPS2 | 499.90518 | 0.615067298 | 0.310028153 | 1.983907887 | 0.04726611 | 0.250276443 |
| PRPSAP1 | 1081.149513 | 0.29253457 | 0.332898637 | 0.878749676 | 0.379537018 | 0.681253679 |
| PRPSAP2 | 1554.582375 | 0.492434514 | 0.294246081 | 1.67354655 | 0.094219751 | 0.351897547 |
| PRR11 | 1388.894499 | 0.110181307 | 0.359198719 | 0.306741926 | 0.759039815 | 0.899440455 |
| PRR12 | 1048.359665 | -0.186527095 | 0.285389172 | -0.653588549 | 0.513376924 | 0.772068737 |
| PRR13 | 2883.128066 | 0.30403856 | 0.444859222 | 0.683448933 | 0.494323209 | 0.760354598 |
| PRR14 | 430.683419 | 0.367946351 | 0.331956511 | 1.108417335 | 0.267681622 | 0.580972515 |
| PRR14L | 1261.222201 | -0.314887219 | 0.314674296 | -1.000676647 | | |

0.316983161 0.627198876
PRR15 229.4781992 -1.303882636 0.694902712 -1.87635278 0.060606854
0.284105089
PRR15L 18.22423281 -1.835745117 1.199024212 -1.531032567
0.125761344 0.407937855
PRR19 24.76621466 -0.151995691 0.650916221 -0.233510374 0.815365115
0.922920552
PRR22 18.47416001 -0.008850757 0.788123869 -0.01123016 0.991039817
0.996726674
PRR29 23.1073857 -1.335757195 0.727678519 -1.835641921 0.066410633
0.297462837
PRR36 956.4216974 -0.482868188 0.391485955 -1.233424038 0.217417601
0.525520777
PRR5 222.1470357 0.075034732 0.396510611 0.18923764 0.849906569 0.938580023
PRR5-ARHGAP8 32.29206754 -1.984346695 0.836562915 -2.372023261
0.017690977 0.150022253
PRR5L 144.4800328 1.171718745 0.487555263 2.403253198 0.016249932 0.144168905
PRR7 28.89748186 1.637181675 1.052133777 1.556058469 0.11969419 0.399006932
PRR9 11.62466991 -1.921796196 1.210902632 -1.587077396 0.112495117
0.385686647
PRRC1 1605.93698 -0.086035276 0.302737427 -0.284191079 0.776263951
0.907435692
PRRC2A 277.9281365 0.930230332 0.730979323 1.272580909 0.203166797
0.509545953
PRRC2B 7710.551436 -0.418590461 0.376898058 -1.110619838
0.26673202 0.579921585
PRRC2C 9627.725133 -0.240595262 0.27132328 -0.886747581
0.375214815 0.679214334
PRRG1 332.5141181 0.048406227 0.278210353 0.173991466 0.861872169 0.944213076
PRRG2 129.9672473 -1.067891578 0.741154437 -1.440848931 0.149627366
0.444386222
PRRG3 84.06539315 0.289921449 0.456247572 0.635447653 0.525136497 0.77816427
PRRG4 145.3529279 -3.060577612 1.066679462 -2.869257093 0.004114372
0.066262358
PRRT2 1084.855692 1.122728862 0.608334365 1.84557856 0.064953422 0.293856843
PRRT3 165.5792684 -0.200210328 0.338504937 -0.591454676 0.554215815
0.796360966
PRRT4 69.39185003 -2.309832597 0.914941784 -2.524567831 0.011584065
0.121704879
PRRX1 50.08964827 -1.49316237 0.864981632 -1.726235927 0.084304977
0.332430848
PRRX2 109.379097 1.815264952 0.614833075 2.952451693 0.003152614 0.055939539
PRSS12 1769.210215 -4.945974666 0.799596145 -6.185590932 6.19E-10
1.06E-06
PRSS16 440.04172 -0.97606863 0.503433663 -1.93882273 0.052522927
0.264550665
PRSS22 148.9082555 -2.031260676 0.91656134 -2.21617538 0.026679493
0.186715644
PRSS23 1933.090014 -0.432866627 0.366622079 -1.18068892 0.237726325
0.548440704
PRSS27 64.65583127 -0.208019559 0.636650608 -0.326740533
0.743864141 0.892535157
PRSS35 285.6526659 -1.402820112 0.666502013 -2.104750001
0.035313076 0.21486064
PRSS36 24.46273233 0.870429225 0.764248265 1.138935167 0.254730196
0.567256913
PRSS42 42.03787682 1.177305758 0.637365219 1.847144656 0.064726178
0.293648578
PRSS44 23.11199059 -0.792949303 0.681249456 -1.163963208
0.244438922 0.555812783
PRSS46P 12.67631208 -0.474465299 0.949188715 -0.499864033
0.617170819 0.831835425
PRSS50 68.66944008 0.08086581 0.454573135 0.177893948 0.858806264
0.942495112

| | | | | | | |
|---|---|---|---|---|---|---|
| PRSS51 | 17.20067415 | -2.188306993 | 1.307202957 | -1.67403767 | 0.094123196 | 0.351710101 |
| PRSS53 | 107.3231249 | 0.188330189 | 0.431688773 | 0.436263811 | 0.662645335 | 0.853902351 |
| PRSS8 | 4194.861703 | -0.665725705 | 0.727805351 | -0.914702955 | 0.360347592 | 0.667522123 |
| PRTFDC1 | 385.6862529 | -0.187978625 | 0.332002361 | -0.566196651 | 0.571260106 | 0.807019934 |
| PRTG | 25842.50602 | -0.127730098 | 0.318217239 | -0.401392766 | 0.688130974 | 0.866925917 |
| PRUNE1 | 858.5223936 | -0.443199699 | 0.240475673 | -1.843012614 | 0.065327169 | 0.294876072 |
| PRUNE2 | 153.9851493 | 0.471961577 | 0.607885026 | 0.776399412 | 0.437513192 | 0.721468765 |
| PRX | 289.3981449 | 0.138262814 | 0.467553996 | 0.295715179 | 0.767447604 | 0.903747194 |
| PRXL2A | 3428.226844 | -0.426449527 | 0.422931721 | -1.008317668 | 0.313301984 | 0.623018103 |
| PRXL2B | 260.6922566 | 0.433686937 | 0.417801691 | 1.038021019 | 0.29926027 | 0.610246278 |
| PRXL2C | 208.9726199 | 0.34554889 | 0.340953337 | 1.013478541 | 0.310831641 | 0.620803953 |
| PSAP | 19541.71948 | -0.003682197 | 0.236196709 | -0.015589537 | 0.987561853 | 0.995541774 |
| PSAT1 | 5405.551063 | 0.817305764 | 0.273883542 | 2.984136095 | 0.002843803 | 0.052472041 |
| PSCA | 13.49928911 | -0.449630063 | 1.392687183 | -0.322850723 | 0.746808297 | 0.894109554 |
| PSD | 42.45433862 | -0.027481134 | 0.549275235 | -0.050031627 | 0.960097185 | 0.98629702 |
| PSD3 | 818.4477532 | 0.925957857 | 0.5770526 | 1.604633368 | 0.108574514 | 0.3783151 |
| PSD4 | 201.1601315 | -2.479897311 | 0.687266344 | -3.608349706 | 0.000308151 | 0.012860767 |
| PSEN1 | 1877.564196 | -0.409325225 | 0.246218374 | -1.662447927 | 0.096422996 | 0.35626747 |
| PSEN2 | 380.0531109 | 0.507084677 | 0.295388931 | 1.716667834 | 0.08603988 | 0.33625288 |
| PSENEN | 1121.250496 | 0.516166965 | 0.433501578 | 1.19069224 | 0.233774426 | 0.543788993 |
| PSIP1 | 8975.789061 | 0.686739949 | 0.287845963 | 2.38578975 | 0.017042491 | 0.14773715 |
| PSKH1 | 1135.057979 | -0.136675782 | 0.272019128 | -0.50244916 | 0.615351606 | 0.830958873 |
| PSMA1 | 4104.589047 | 0.604171182 | 0.341871113 | 1.767248412 | 0.077186635 | 0.31844659 |
| PSMA2 | 6224.622364 | 0.972284568 | 0.366856346 | 2.650314157 | 0.008041696 | 0.100743653 |
| PSMA3 | 4328.175384 | 0.867174086 | 0.323510195 | 2.680515484 | 0.007350886 | 0.095072717 |
| PSMA4 | 6426.222295 | 0.914095302 | 0.272684428 | 3.352209398 | 0.000801694 | 0.023754869 |
| PSMA5 | 3220.917663 | 0.729300058 | 0.317473865 | 2.297197154 | 0.021607525 | 0.166611729 |
| PSMA6 | 4304.917912 | 0.64251968 | 0.307685034 | 2.088238323 | 0.036776342 | 0.219399876 |
| PSMA7 | 4303.770791 | 0.744883531 | 0.366757341 | 2.030998286 | 0.042255169 | 0.236528876 |
| PSMB1 | 5663.856463 | 0.7495486 | 0.415540641 | 1.803791315 | 0.071264028 | 0.306992555 |
| PSMB10 | 76.12243649 | 1.037652275 | 0.504263681 | 2.05775731 | 0.039613436 | 0.229275285 |
| PSMB2 | 3059.927675 | 0.488884105 | 0.385538648 | 1.268054726 | 0.204778408 | 0.511170593 |
| PSMB3 | 2457.828411 | 0.512778273 | 0.348891098 | 1.469737336 | 0.141632906 | 0.431641789 |
| PSMB4 | 6410.692148 | 0.579775957 | 0.417900179 | 1.38735513 | 0.1653335 | 0.467525541 |
| PSMB5 | 5042.662998 | 0.58704109 | 0.364493292 | 1.610567608 | 0.107273999 | 0.375248751 |
| PSMB6 | 3433.746741 | 0.6830836 | 0.453846085 | 1.505099688 | 0.132298449 | 0.418521131 |
| PSMB7 | 4485.594278 | 0.248212937 | 0.298262941 | 0.832195028 | 0.40529887 | 0.701039911 |
| PSMC1 | 2739.966237 | 0.340894294 | 0.332081679 | 1.026537493 | 0.3046383 | 0.614996026 |
| PSMC1P1 | 253.9282826 | -0.699414766 | 2.264422163 | -0.308871189 | 0.757419514 | 0.898913514 |
| PSMC2 | 2090.260637 | 0.402821095 | 0.234515434 | 1.717674132 | 0.08585607 | 0.335996519 |
| PSMC3 | 3428.171868 | 0.002058167 | 0.260192351 | 0.007910176 | 0.993688658 | 0.99730871 |
| PSMC3IP | 363.6300503 | 0.335080814 | 0.293551927 | 1.141470326 | 0.253674254 | 0.565954722 |
| PSMC4 | 996.1451142 | -0.072556457 | 0.2594481 | -0.27965692 | 0.779740733 | 0.908699236 |

| | | | | | | |
|---|---|---|---|---|---|---|
| PSMC5 | 3297.13773 | -0.118134963 | 0.266497467 | -0.443287376 | 0.657557886 | 0.850609594 |
| PSMC6 | 3455.571078 | 0.638129509 | 0.259705255 | 2.457129756 | 0.014005209 | 0.134477513 |
| PSMD1 | 5191.698976 | 0.21196992 | 0.22279743 | 0.951401996 | 0.341400347 | 0.648152255 |
| PSMD10 | 1762.405842 | 0.791007736 | 0.300118439 | 2.635651904 | 0.008397583 | 0.103577017 |
| PSMD11 | 2721.014716 | 0.13766754 | 0.23300745 | 0.590828921 | 0.554635055 | 0.796360966 |
| PSMD12 | 1756.45285 | 0.20230091 | 0.270259446 | 0.748543348 | 0.454132488 | 0.734034091 |
| PSMD13 | 3404.500293 | 0.583764976 | 0.307210354 | 1.900212566 | 0.05740523 | 0.277443371 |
| PSMD14 | 2799.674456 | 0.502503249 | 0.26109143 | 1.924625593 | 0.054276215 | 0.269268954 |
| PSMD2 | 6747.770546 | -0.134493822 | 0.22312532 | -0.602772567 | 0.546659999 | 0.792249935 |
| PSMD3 | 3667.135632 | -0.226866148 | 0.280017578 | -0.810185378 | 0.41783364 | 0.709307489 |
| PSMD4 | 5492.785657 | 0.330933416 | 0.344610676 | 0.96031098 | 0.336898725 | 0.644459616 |
| PSMD5 | 1274.760683 | -0.260329921 | 0.238662596 | -1.090786432 | 0.275366868 | 0.588475948 |
| PSMD6 | 2255.170927 | 0.369204058 | 0.269677454 | 1.369057934 | 0.170981166 | 0.474214884 |
| PSMD7 | 3192.967816 | 0.393205506 | 0.304197129 | 1.292600977 | 0.196149101 | 0.501797297 |
| PSMD8 | 3360.234442 | 0.243906651 | 0.266655733 | 0.914687444 | 0.360355738 | 0.667522123 |
| PSMD9 | 1016.896603 | 0.730513571 | 0.394491234 | 1.851786577 | 0.064056476 | 0.292206722 |
| PSME1 | 613.5045595 | 0.753184087 | 0.397787234 | 1.893434539 | 0.058300106 | 0.27980731 |
| PSME2 | 579.9447141 | 0.627487543 | 0.336050054 | 1.867244286 | 0.061867482 | 0.286505362 |
| PSME3 | 3904.012313 | -0.175761273 | 0.248562147 | -0.707111986 | 0.479496888 | 0.751803151 |
| PSME4 | 2705.378464 | -0.144343276 | 0.324493304 | -0.444826672 | 0.656445018 | 0.849955395 |
| PSMF1 | 1289.138822 | -0.403642103 | 0.251580901 | -1.60442268 | 0.108620915 | 0.378356736 |
| PSMG1 | 1197.804449 | 0.797506312 | 0.330689571 | 2.411646393 | 0.015880675 | 0.142656219 |
| PSMG2 | 1774.447083 | 0.620169667 | 0.395968424 | 1.566209904 | 0.117299485 | 0.395279984 |
| PSMG3 | 829.7356794 | 1.176537591 | 0.444102109 | 2.649250179 | 0.008067059 | 0.100970518 |
| PSMG4 | 237.5627587 | 0.751732715 | 0.331056095 | 2.270710995 | 0.023164479 | 0.173867082 |
| PSPC1 | 4322.499838 | 0.193554776 | 0.304211678 | 0.636250317 | 0.524613282 | 0.777662198 |
| PSPH | 895.4786263 | 0.552717256 | 0.336622425 | 1.641950196 | 0.100600326 | 0.363123375 |
| PSPHP1 | 61.6749868 | 0.740298018 | 0.550889428 | 1.343823242 | 0.179005552 | 0.483466568 |
| PSPN | 83.01398777 | 0.417964221 | 0.462752648 | 0.903213029 | 0.366412844 | 0.672984172 |
| PSRC1 | 1433.258821 | 1.023636553 | 0.265744212 | 3.851961799 | 0.000117175 | 0.006837179 |
| PSTK | 166.949955 | 0.371314936 | 0.353114222 | 1.051543417 | 0.29300908 | 0.606114092 |
| PSTPIP1 | 15.57894581 | 1.474695056 | 0.952832885 | 1.547695383 | 0.121695654 | 0.402231976 |
| PSTPIP2 | 157.3038308 | -1.999622244 | 0.665132524 | -3.006351624 | 0.002644031 | 0.050167035 |
| PTAFR | 23.26935037 | 0.298645234 | 0.846847858 | 0.352655121 | 0.724347009 | 0.883670922 |
| PTAR1 | 1396.936077 | -0.343524327 | 0.475089575 | -0.723072753 | 0.469635188 | 0.744930517 |
| PTBP1 | 9960.273868 | -0.059287471 | 0.263200293 | -0.225256099 | 0.82178005 | 0.92563193 |
| PTBP2 | 1799.666186 | 0.228609077 | 0.253696249 | 0.901113349 | 0.367528056 | 0.673938238 |
| PTBP3 | 2790.059414 | -0.406353426 | 0.402877544 | -1.008627639 | 0.313153247 | 0.623010823 |
| PTCD1 | 485.9757846 | -0.214009072 | 0.333163817 | -0.642353882 | 0.520643436 | 0.775597169 |
| PTCD2 | 478.6096633 | 0.141952587 | 0.271212566 | 0.523399743 | 0.600696103 | 0.824475293 |
| PTCD3 | 3405.965967 | 0.109823803 | 0.31284685 | 0.351046537 | 0.725553436 | 0.88422336 |
| PTCH1 | 2926.92863 | -0.385060861 | 0.440701399 | -0.873745493 | 0.38225687 | 0.683628312 |
| PTCH2 | 27.07193063 | 0.50156936 | 0.674043419 | 0.744120254 | 0.456803727 | 0.735735031 |
| PTCHD1 | 20.83979409 | -3.085703109 | 1.064645443 | -2.898338719 | | |

| | | | | | | |
|---|---|---|---|---|---|---|
| | | | | | 0.003751452 | 0.06316679 |
| PTCHD4 | 257.9682473 | 0.767802516 | 0.399753532 | 1.920689763 | 0.054770831 | 0.270748869 |
| PTDSS1 | 1644.69568 | -0.155769953 | 0.237608248 | -0.655574687 | 0.512097817 | 0.771681172 |
| PTDSS2 | 562.4432379 | -0.810345452 | 0.495793335 | -1.634442005 | 0.102166074 | 0.366611924 |
| PTEN | 880.2446118 | 0.224312076 | 0.374142577 | 0.599536353 | 0.548815276 | 0.793468239 |
| PTENP1 | 17.39001855 | 0.938757628 | 0.709896871 | 1.322385923 | 0.186039674 | 0.490682875 |
| PTER | 248.6825167 | -0.334599255 | 0.316179956 | -1.058255743 | 0.289938864 | 0.602639061 |
| PTGDS | 20.63887793 | -0.433154581 | 0.956216876 | -0.452987802 | 0.650557525 | 0.847540422 |
| PTGER3 | 17.9724771 | -0.432689144 | 0.904063863 | -0.478604623 | 0.632219929 | 0.838768903 |
| PTGER4 | 10.78249839 | -1.221464385 | 1.273734218 | -0.958963312 | 0.337577228 | 0.64509184 |
| PTGES | 121.035445 | 0.292912672 | 0.368422551 | 0.795045449 | 0.426587065 | 0.714361085 |
| PTGES2 | 944.3753312 | 0.248793908 | 0.36991636 | 0.672568003 | 0.501222164 | 0.763991735 |
| PTGES3 | 10884.2171 | 0.750808271 | 0.311871421 | 2.407428898 | 0.01606529 | 0.143566101 |
| PTGES3L | 71.82331798 | 0.142384037 | 0.533586314 | 0.266843494 | 0.789589664 | 0.91286231 |
| PTGES3L-AARSD1 | 76.94478986 | -0.511359021 | 0.57909929 | -0.883024777 | 0.377222885 | 0.680674553 |
| PTGFRN | 813.9220772 | 0.144828918 | 0.317237012 | 0.456532221 | 0.648007306 | 0.846597949 |
| PTGIS | 1283.653689 | -0.713779429 | 0.282560235 | -2.526114226 | 0.011533198 | 0.121534216 |
| PTGR1 | 2480.338152 | 0.378667256 | 0.31825725 | 1.189815017 | 0.234119106 | 0.544210545 |
| PTGR2 | 357.7509492 | 0.198175693 | 0.280134132 | 0.707431442 | 0.479298403 | 0.751673525 |
| PTGS1 | 68.47924014 | 1.419405528 | 0.809925503 | 1.752513686 | 0.079685519 | 0.323199907 |
| PTH1R | 497.9329911 | -2.865310337 | 0.578503697 | -4.952968061 | 7.31E-07 | 0.000149844 |
| PTK2 | 4465.850936 | -0.061537202 | 0.280786076 | -0.219160446 | 0.826525066 | 0.927868268 |
| PTK2B | 283.6249464 | -0.799398933 | 0.293453655 | -2.724106246 | 0.006447576 | 0.087257964 |
| PTK7 | 11699.18282 | -0.579894271 | 0.255968299 | -2.265492536 | 0.023482472 | 0.174992377 |
| PTMA | 56724.65299 | 0.946434046 | 0.4767657 | 1.98511354 | 0.047131841 | 0.250100439 |
| PTMS | 14519.64074 | 1.144286405 | 0.47068633 | 2.431101845 | 0.015052983 | 0.139165886 |
| PTN | 1319.729355 | -0.266069194 | 0.586573804 | -0.453598834 | 0.650117593 | 0.847430646 |
| PTOV1 | 4099.026246 | 0.202249611 | 0.323823312 | 0.624567792 | 0.532254764 | 0.782369637 |
| PTP4A1 | 2461.683 | 0.322842785 | 0.389324596 | 0.829238092 | 0.406969694 | 0.701626421 |
| PTP4A2 | 3671.674173 | 0.417534438 | 0.320755874 | 1.301720315 | 0.193012012 | 0.498908723 |
| PTP4A3 | 85.36213053 | 0.474371588 | 0.470186819 | 1.008900226 | 0.313022487 | 0.622994042 |
| PTPA | 4741.88381 | 0.181646853 | 0.252527423 | 0.719315358 | 0.471946635 | 0.746399031 |
| PTPDC1 | 335.3126998 | -0.306099618 | 0.310598326 | -0.985515993 | 0.32437068 | 0.63448009 |
| PTPMT1 | 398.2057406 | 0.741954493 | 0.514970151 | 1.440771841 | 0.149649151 | 0.444386222 |
| PTPN1 | 1349.663235 | -0.267320237 | 0.334655089 | -0.798793281 | 0.424410287 | 0.713303793 |
| PTPN11 | 3197.092584 | -0.070398125 | 0.270684797 | -0.260074174 | 0.794806559 | 0.914877916 |
| PTPN12 | 3867.252071 | -0.346544063 | 0.228901315 | -1.513945271 | 0.130039721 | 0.415175781 |

| | | | | | | |
|---|---|---|---|---|---|---|
| PTPN13 | 5422.967445 | -0.433882771 | 0.55633038 | -0.779901272 | 0.43544899 | 0.720542613 |
| PTPN14 | 3187.848064 | -0.154318057 | 0.301443015 | -0.511931111 | 0.608699223 | 0.827916077 |
| PTPN18 | 205.6293796 | 0.378330883 | 0.290174733 | 1.303803672 | 0.19230053 | 0.498188413 |
| PTPN2 | 1611.128058 | 0.044042522 | 0.257119878 | 0.171291781 | 0.863994343 | 0.945104779 |
| PTPN23 | 980.8635861 | -0.649934093 | 0.393791572 | -1.650452016 | 0.09885052 | 0.361016186 |
| PTPN3 | 86.65044221 | -0.512026956 | 0.431670198 | -1.186153128 | 0.235561833 | 0.546271563 |
| PTPN4 | 1041.158877 | -0.073345256 | 0.344885146 | -0.212665742 | 0.831587689 | 0.930997148 |
| PTPN5 | 84.64938725 | 0.055943516 | 0.599970624 | 0.093243759 | 0.925709911 | 0.972124891 |
| PTPN6 | 533.3761073 | -1.419591721 | 0.476742129 | -2.977693043 | 0.002904267 | 0.053021219 |
| PTPN9 | 881.8395069 | -0.549215001 | 0.250536886 | -2.192152262 | 0.028368514 | 0.192544525 |
| PTPRA | 2387.989651 | -0.192389461 | 0.313309332 | -0.614055955 | 0.539178353 | 0.786771255 |
| PTPRD | 408.0274022 | -0.045616624 | 0.406881793 | -0.11211272 | 0.910734032 | 0.966746851 |
| PTPRE | 134.639831 | -1.625241171 | 0.572968695 | -2.836526997 | 0.004560712 | 0.071037541 |
| PTPRF | 13424.23439 | -1.1853194 | 0.245374333 | -4.830657648 | 1.36E-06 | 0.000226869 |
| PTPRG | 1235.634774 | 0.102333533 | 0.341658437 | 0.299519994 | 0.764543319 | 0.902847878 |
| PTPRJ | 467.1928148 | -0.020080123 | 0.35549289 | -0.056485302 | 0.954955204 | 0.984323736 |
| PTPRK | 765.2214359 | -0.984904287 | 0.30819277 | -3.19574105 | 0.001394722 | 0.03385493 |
| PTPRM | 411.9280045 | -0.162567983 | 0.435087384 | -0.373644442 | 0.708668865 | 0.875585221 |
| PTPRN2 | 20.22405615 | 0.343150467 | 0.918829818 | 0.373464661 | 0.708802642 | 0.875646746 |
| PTPRO | 24.68645552 | 1.251436251 | 0.778950862 | 1.60656636 | 0.108149527 | 0.377180798 |
| PTPRS | 2059.700518 | 0.032931344 | 0.752245114 | 0.043777411 | 0.965081833 | 0.988700292 |
| PTPRU | 1360.404116 | -0.167084616 | 0.276849461 | -0.603521549 | 0.546161784 | 0.791840638 |
| PTPRZ1 | 1002.18015 | 1.833182159 | 0.774877313 | 2.36577085 | 0.017992577 | 0.15131119 |
| PTRH1 | 69.45249891 | 0.40762946 | 0.600194978 | 0.679161731 | 0.497035392 | 0.761325016 |
| PTRH2 | 604.4607719 | 0.574207907 | 0.383954569 | 1.495510026 | 0.134781385 | 0.421290623 |
| PTRHD1 | 596.2121052 | 1.036910689 | 0.404233159 | 2.565130213 | 0.010313709 | 0.115220384 |
| PTS | 869.5345944 | 0.624563136 | 0.420886017 | 1.483924651 | 0.137828911 | 0.425968529 |
| PTTG1 | 3326.318553 | 1.866993457 | 0.439309711 | 4.249834249 | 2.14E-05 | 0.001966978 |
| PTTG1IP | 5134.217329 | -0.106880863 | 0.282572524 | -0.378242234 | 0.705250653 | 0.873844958 |
| PTX3 | 22.51437465 | 2.69630187 | 0.904587525 | 2.980697605 | 0.002875926 | 0.052821643 |
| PUDP | 842.5948451 | 0.186756659 | 0.286496469 | 0.651863737 | 0.51448908 | 0.77285333 |
| PUF60 | 390.894106 | -1.039537762 | 1.901048788 | -0.546823295 | 0.584500137 | 0.814879678 |
| PUM1 | 4141.214491 | -0.207881133 | 0.234876607 | -0.885065292 | 0.376121418 | 0.680079109 |
| PUM2 | 3442.492138 | -0.084599803 | 0.258453518 | -0.327330823 | 0.743417682 | 0.892534058 |
| PUM3 | 1240.367417 | 0.290342517 | 0.23854737 | 1.217127303 | 0.223555784 | 0.53268495 |
| PURA | 406.2271044 | 0.134503118 | 0.385823043 | 0.34861349 | 0.727379499 | 0.885092921 |
| PURB | 1301.748789 | -0.313146094 | 0.319451916 | -0.980260497 | 0.326957549 | 0.636830527 |
| PURG | 165.03332 | 1.012132519 | 0.463794993 | 2.182284275 | 0.029088559 | 0.195093955 |
| PUS1 | 519.3194536 | 0.349435194 | 0.306962595 | 1.138364087 | 0.254968484 | 0.567340403 |
| PUS10 | 228.0023095 | 0.16916933 | 0.350428345 | 0.482750134 | 0.629273162 | 0.837430099 |

| | | | | | | |
|---|---|---|---|---|---|---|
| PUS3 | 481.6494468 | 0.244966973 | 0.393276351 | 0.622887627 | 0.533358369 | 0.783143103 |
| PUS7 | 796.1848847 | 0.186320573 | 0.296888953 | 0.627576643 | 0.530281314 | 0.780679069 |
| PUS7L | 445.2509684 | -0.043587041 | 0.354658432 | -0.122898647 | 0.902187357 | 0.962591209 |
| PUSL1 | 224.5085437 | 0.699106905 | 0.502584888 | 1.391022536 | 0.164218593 | 0.466312149 |
| PVR | 308.9534244 | -0.185841786 | 0.382938968 | -0.485303929 | 0.627460769 | 0.836054176 |
| PWP1 | 1672.970986 | 0.169582482 | 0.236951735 | 0.715683648 | 0.47418671 | 0.748145033 |
| PWP2 | 1168.115755 | -0.058409792 | 0.285902947 | -0.204299371 | 0.838119561 | 0.933836065 |
| PWWP2A | 708.2226102 | 0.007405927 | 0.286788738 | 0.025823632 | 0.979398013 | 0.992858175 |
| PWWP2B | 115.9268841 | 0.690200854 | 0.706552225 | 0.976857519 | 0.328639695 | 0.638020359 |
| PWWP3A | 1173.354493 | -0.031191232 | 0.369488445 | -0.08441734 | 0.932724621 | 0.974543059 |
| PWWP3B | 173.6615772 | 0.661503784 | 0.463408403 | 1.427474727 | 0.153443105 | 0.450006832 |
| PXDC1 | 566.2766897 | -1.681834926 | 0.699753828 | -2.403466559 | 0.016240452 | 0.144168905 |
| PXDN | 5561.71579 | -0.318217927 | 0.357758548 | -0.889476797 | 0.373746888 | 0.678520659 |
| PXDNL | 18.61585831 | -1.714127511 | 0.806431319 | -2.125571603 | 0.033538946 | 0.208789229 |
| PXK | 280.2644187 | -0.596763664 | 0.321465657 | -1.85638388 | 0.063398861 | 0.290318176 |
| PXMP2 | 189.0161887 | 0.562886519 | 0.471459128 | 1.193924321 | 0.232507573 | 0.542005591 |
| PXMP4 | 224.3358741 | 0.152481158 | 0.295667287 | 0.515718729 | 0.606050871 | 0.826611325 |
| PXN | 3729.131527 | -0.989325352 | 0.37620667 | -2.629738998 | 0.008545044 | 0.104377512 |
| PXYLP1 | 673.2282693 | -0.048766722 | 0.284986759 | -0.171119257 | 0.864129994 | 0.9451245 |
| PYCARD | 20.05994517 | 1.075700842 | 0.974374919 | 1.103990693 | 0.269597175 | 0.582376584 |
| PYCR1 | 1377.376706 | 0.947868137 | 0.28335786 | 3.34512739 | 0.000822448 | 0.024115162 |
| PYCR2 | 998.2315431 | 0.760311749 | 0.241059993 | 3.154035392 | 0.001610296 | 0.037175747 |
| PYCR3 | 49.01871571 | -0.033310602 | 0.884018794 | -0.037680876 | 0.969942124 | 0.989496332 |
| PYGB | 1603.093839 | 0.115209557 | 0.308310168 | 0.373680693 | 0.708641891 | 0.875585221 |
| PYGL | 1040.205302 | 0.452340125 | 0.349293763 | 1.295013462 | 0.195315586 | 0.501121723 |
| PYGM | 335.6827326 | -0.508014022 | 0.371805834 | -1.366342254 | 0.171831565 | 0.474879827 |
| PYGO1 | 71.71247876 | 0.180216621 | 0.64294104 | 0.280300385 | 0.779247055 | 0.908367621 |
| PYGO2 | 1469.813382 | -0.056412477 | 0.224719937 | -0.25103459 | 0.801787367 | 0.917339075 |
| PYM1 | 1675.33018 | 0.567535929 | 0.456292979 | 1.2437972 | 0.213574211 | 0.520678417 |
| PYROXD1 | 192.2633527 | 0.170924864 | 0.321768606 | 0.531204291 | 0.59527722 | 0.821441254 |
| PYROXD2 | 182.1456708 | 0.542783019 | 0.472991936 | 1.147552375 | 0.2511534 | 0.564075361 |
| PYURF | 1449.016203 | 0.735186611 | 0.368447918 | 1.995361011 | 0.04600352 | 0.246950947 |
| PYY | 10.26256378 | 0.776041394 | 0.997286927 | 0.778152579 | 0.436479069 | 0.720784523 |
| QARS | 10115.0744 | -0.20098866 | 0.232713015 | -0.863676062 | 0.387765867 | 0.687804977 |
| QDPR | 752.9704132 | -0.441607434 | 0.407804869 | -1.082889068 | 0.278857665 | 0.592513955 |
| QKI | 5709.349942 | 0.006515138 | 0.2734515 | 0.023825571 | 0.980991743 | 0.993241851 |
| QPCT | 48.04404229 | -0.451788501 | 0.848503924 | -0.532453049 | 0.594412252 | 0.820642877 |
| QPCTL | 542.8921071 | 0.961409729 | 0.400489345 | 2.400587531 | 0.016368775 | 0.144378309 |
| QPRT | 5388.123058 | 0.458430471 | 0.303922782 | 1.508378108 | 0.131457783 | 0.41719445 |
| QRFPR | 14.1290719 | 1.015914264 | 0.882962629 | 1.150574476 | 0.249907339 | 0.562742997 |
| QRICH1 | 2485.817298 | -0.141567337 | 0.233601022 | -0.606021909 | 0.544500205 | 0.790739196 |

| | | | | | | |
|---|---|---|---|---|---|---|
| QRICH2 | 51.5702883 | -1.094048399 | 0.6411269 | -1.706445947 | 0.087925082 | 0.340967522 |
| QRSL1 | 726.1677442 | -0.190182189 | 0.245666982 | -0.774146319 | 0.438844269 | 0.722407379 |
| QSER1 | 1851.62148 | -0.378609115 | 0.41100798 | -0.921172177 | 0.356960542 | 0.664415851 |
| QSOX1 | 1680.702894 | 0.18310237 | 0.34514472 | 0.53050897 | 0.595759091 | 0.82173481 |
| QSOX2 | 816.8908251 | 0.63718765 | 0.407706367 | 1.562859209 | 0.118085714 | 0.395958766 |
| QTRT1 | 605.3900929 | 0.4330351 | 0.393318445 | 1.10097837 | 0.270906071 | 0.583394591 |
| QTRT2 | 778.4065048 | -0.299980049 | 0.323520413 | -0.92723685 | 0.353803574 | 0.660911171 |
| R3HCC1 | 670.4919168 | 0.222815059 | 0.324717468 | 0.686181316 | 0.492598778 | 0.759174768 |
| R3HCC1L | 638.9129333 | -0.567125795 | 0.316231628 | -1.793387332 | 0.072911012 | 0.310624251 |
| R3HDM1 | 1334.21551 | -0.259887138 | 0.422957662 | -0.614451898 | 0.53891675 | 0.786598616 |
| R3HDM2 | 1711.326602 | -0.580992944 | 0.288913094 | -2.010960926 | 0.044329584 | 0.241915151 |
| R3HDM4 | 1473.204685 | 0.410165044 | 0.3959624 | 1.035868669 | 0.300263416 | 0.610801113 |
| RAB10 | 4383.324539 | 0.238578091 | 0.231169237 | 1.032049482 | 0.302048938 | 0.612946018 |
| RAB11A | 4887.558597 | 0.292764645 | 0.346204663 | 0.845640387 | 0.397753388 | 0.696366306 |
| RAB11B | 1174.577361 | 0.145101332 | 0.356659475 | 0.406834369 | 0.684129642 | 0.863826466 |
| RAB11FIP1 | 2806.355414 | -0.889189025 | 0.305259036 | -2.912899934 | 0.003580893 | 0.061524046 |
| RAB11FIP2 | 728.0967715 | 0.011576772 | 0.299669174 | 0.03863184 | 0.969183917 | 0.989496332 |
| RAB11FIP3 | 465.0547851 | 0.112270573 | 0.37450778 | 0.299781685 | 0.764343687 | 0.902756684 |
| RAB11FIP4 | 3214.960395 | -1.33485743 | 0.298332312 | -4.474397771 | 7.66E-06 | 0.000883852 |
| RAB11FIP5 | 269.4219054 | -0.201534312 | 0.302371747 | -0.666511715 | 0.505084078 | 0.766569683 |
| RAB12 | 705.7908882 | 0.525121179 | 0.417944135 | 1.256438684 | 0.208956964 | 0.516300667 |
| RAB13 | 2536.191406 | 0.030863571 | 0.432999961 | 0.071278462 | 0.943176137 | 0.978607681 |
| RAB14 | 2235.765425 | 0.203655928 | 0.241374716 | 0.843733476 | 0.398818357 | 0.697300137 |
| RAB15 | 1438.20029 | -1.593933449 | 0.279620675 | -5.700341889 | 1.20E-08 | 1.01E-05 |
| RAB17 | 494.41256 | -1.733464627 | 0.793357909 | -2.184971759 | 0.028890917 | 0.194385362 |
| RAB18 | 2040.673702 | 0.159531365 | 0.252505223 | 0.631794319 | 0.527521287 | 0.778966897 |
| RAB1A | 4638.348816 | 0.248596865 | 0.34912574 | 0.71205539 | 0.476430477 | 0.74962941 |
| RAB1B | 3631.30354 | -0.105786632 | 0.28464652 | -0.371642105 | 0.710159335 | 0.876599663 |
| RAB20 | 302.1268231 | -0.259808703 | 0.626800319 | -0.414499953 | 0.678508001 | 0.861138002 |
| RAB21 | 1048.360791 | 0.013060784 | 0.307455606 | 0.042480227 | 0.966115874 | 0.988700292 |
| RAB22A | 475.1475442 | 0.139643324 | 0.274875132 | 0.50802458 | 0.611436106 | 0.829058038 |
| RAB23 | 805.7822656 | 0.219981274 | 0.310044818 | 0.709514433 | 0.478005298 | 0.750550442 |
| RAB24 | 516.0568344 | 0.834516048 | 0.433695485 | 1.924198147 | 0.054329751 | 0.269446439 |
| RAB25 | 1986.697182 | -0.873238144 | 0.770806485 | -1.132888943 | 0.257260894 | 0.569767157 |
| RAB26 | 22.70432283 | 1.144550933 | 0.7087556 | 1.614873919 | 0.106338001 | 0.374140185 |
| RAB27A | 154.2201374 | -0.091774523 | 0.359428434 | -0.25533462 | 0.798464663 | 0.916056735 |
| RAB27B | 79.23882412 | -1.099151178 | 0.511509364 | -2.148838818 | 0.031647179 | 0.202592072 |
| RAB28 | 764.4865082 | 1.05928669 | 0.315556801 | 3.356881186 | 0.00078827 | 0.023494776 |
| RAB29 | 314.1413758 | 0.000749846 | 0.271771998 | 0.002759098 | 0.997798561 | 0.998851994 |
| RAB2A | 1765.827502 | 0.247734808 | 0.333890848 | 0.741963457 | 0.458109476 | 0.736538667 |

| | | | | | | |
|---|---|---|---|---|---|---|
| RAB2B | 817.5176056 | 0.174795028 | 0.233986801 | 0.747029434 | 0.455045794 | 0.734911996 |
| RAB30 | 308.7298016 | -0.272910939 | 0.297948775 | -0.91596597 | 0.359684744 | 0.667169244 |
| RAB31 | 1785.965399 | -1.517455789 | 0.553469874 | -2.741713437 | 0.006111964 | 0.084371795 |
| RAB32 | 85.48425195 | 0.678532219 | 0.418489781 | 1.621383008 | 0.104935523 | 0.37133017 |
| RAB33A | 13.54080439 | 2.797420076 | 1.288124781 | 2.171699603 | 0.029878329 | 0.197226575 |
| RAB33B | 407.2061543 | -0.126652898 | 0.282627165 | -0.448127122 | 0.654061455 | 0.849043068 |
| RAB34 | 2381.175752 | 0.626760951 | 0.395575984 | 1.584426195 | 0.113096765 | 0.386997793 |
| RAB35 | 1034.111295 | -0.56909929 | 0.376489803 | -1.511592837 | 0.130637475 | 0.416016191 |
| RAB36 | 106.5744606 | 0.158544007 | 0.51052788 | 0.310549166 | 0.756143377 | 0.898288428 |
| RAB37 | 87.0108608 | -0.733215609 | 0.562770042 | -1.302868942 | 0.192619509 | 0.498838446 |
| RAB38 | 145.8850036 | 0.170117404 | 0.367853507 | 0.462459649 | 0.643751732 | 0.845466381 |
| RAB39A | 66.79375648 | -0.360861822 | 0.703633922 | -0.512854498 | 0.608053105 | 0.827607804 |
| RAB39B | 223.3566476 | 0.444367089 | 0.303154609 | 1.465810104 | 0.142700028 | 0.433233718 |
| RAB3A | 62.48321533 | 0.915064024 | 0.629311691 | 1.454071228 | 0.14592656 | 0.438251053 |
| RAB3B | 2910.733076 | -1.09765102 | 0.274826571 | -3.993977062 | 6.50E-05 | 0.004569989 |
| RAB3C | 220.8884643 | 0.18482075 | 0.339247602 | 0.544796041 | 0.585893802 | 0.815639728 |
| RAB3D | 505.9407182 | -0.189001857 | 0.43664411 | -0.432851039 | 0.665122991 | 0.855061512 |
| RAB3GAP1 | 1549.386917 | -0.38906348 | 0.30135837 | -1.291032599 | 0.196692374 | 0.50258876 |
| RAB3GAP2 | 1802.470217 | 0.003411069 | 0.252184158 | 0.013526103 | 0.989208061 | 0.996168303 |
| RAB3IL1 | 108.8962281 | -0.173287873 | 0.527629087 | -0.328427444 | 0.742588495 | 0.892270716 |
| RAB3IP | 1388.508969 | 0.022047813 | 0.236947632 | 0.093049306 | 0.92586439 | 0.972196059 |
| RAB40A | 26.27640386 | -0.713996322 | 0.679707203 | -1.050446897 | 0.293512694 | 0.606329123 |
| RAB40B | 97.98659373 | 0.727660365 | 0.519477344 | 1.400754765 | 0.16128742 | 0.46167973 |
| RAB40C | 382.9952558 | -0.186424303 | 0.508816596 | -0.366388016 | 0.714075561 | 0.878400517 |
| RAB42 | 22.61165167 | 1.269150136 | 0.808187327 | 1.570366293 | 0.11632992 | 0.393147964 |
| RAB43 | 193.340184 | -0.739092874 | 0.669264972 | -1.104335211 | 0.269447754 | 0.582188661 |
| RAB43P1 | 11.885499 | -0.036117102 | 1.948327161 | -0.018537493 | 0.985210067 | 0.99460071 |
| RAB4A | 1865.756269 | 0.229546697 | 0.299629631 | 0.766101457 | 0.443615936 | 0.726848446 |
| RAB4B | 404.2517189 | 0.467462808 | 0.435589843 | 1.073171965 | 0.283193967 | 0.596962021 |
| RAB4B-EGLN2 | 36.9035022 | -0.182366655 | 0.692297286 | -0.263422461 | 0.792224981 | 0.913570955 |
| RAB5A | 1771.832104 | -0.194849717 | 0.316698078 | -0.615253867 | 0.538387079 | 0.786560766 |
| RAB5B | 3041.02657 | -0.534998065 | 0.245560669 | -2.178679783 | 0.029355465 | 0.195861365 |
| RAB5C | 2243.380254 | -0.223462141 | 0.283836286 | -0.787292366 | 0.431110738 | 0.71785545 |
| RAB5IF | 1686.220037 | 0.842823881 | 0.408548198 | 2.062972951 | 0.039115198 | 0.227886587 |
| RAB6A | 2914.706111 | -0.380136786 | 0.289373625 | -1.313653882 | 0.188962707 | 0.494119637 |
| RAB6B | 400.3597241 | 0.022350263 | 0.3246099 | 0.068852685 | 0.945106881 | 0.979654037 |
| RAB7A | 3838.22634 | -0.008956015 | 0.298555386 | -0.029997833 | 0.976068781 | 0.991758053 |
| RAB8A | 1237.313859 | -0.309278295 | 0.325325833 | -0.950672413 | 0.341770697 | |

0.648645757
RAB8B 973.2999832 0.470315279 0.294885886 1.594906031 0.110733251 0.383021922
RAB9A 519.2304632 0.564517735 0.460241698 1.226567989 0.219985007 0.528543966
RAB9B 23.29481136 0.430867405 0.708155908 0.608435798 0.542898475 0.789372509
RABAC1 995.3502973 0.902943311 0.542610059 1.664074035 0.096097635
0.355758229
RABEP1 1927.200891 0.090885563 0.410289636 0.221515619 0.824690977
0.92681808
RABEP2 266.7392293 -0.554978747 0.533251872 -1.040744113
0.297994324 0.609375237
RABEPK 360.1916909 0.835303954 0.358148397 2.332284499 0.019685729
0.158893482
RABGAP1 1626.255789 -0.424577937 0.35763173 -1.187193143
0.235151445 0.54573697
RABGAP1L 1142.933803 -0.705512483 0.282421694 -2.498081764
0.012486739 0.12669427
RABGEF1 603.0042498 0.090302838 0.249441336 0.362020342 0.71733683
0.880516207
RABGGTA 371.5102882 -0.333872453 0.329375395 -1.013653289
0.31074822 0.620719059
RABGGTB 3143.122292 0.27889305 0.231038794 1.207126499 0.227383431
0.536186965
RABIF 287.8258446 -0.051241253 0.4445189 -0.115273509 0.908228335
0.965583536
RABL2A 432.5031686 -0.16417784 0.297658491 -0.551564445 0.581246802
0.812652772
RABL2B 748.4644237 -0.340896813 0.297091672 -1.147446543
0.251197114 0.564075361
RABL3 509.2461203 -0.118734922 0.257021588 -0.461964781 0.644106577
0.845598899
RABL6 2859.75344 0.05800768 0.41299451 0.140456299 0.888299478 0.955943827
RAC1 6865.315548 0.336959449 0.432081558 0.779851494 0.435478292 0.720542613
RAC2 41.72480549 -0.587651784 0.621829263 -0.945037197 0.344639886
0.651368096
RAC3 1603.183735 0.965528343 0.420053806 2.298582535 0.021528656 0.166469811
RACGAP1 1627.553077 0.866612687 0.355526134 2.437549883 0.014787174
0.138223176
RACK1 40074.39292 -0.013861032 0.283016953 -0.048975979 0.960938439
0.986559996
RAD1 837.8458511 0.152348136 0.273431634 0.557170848 0.577410705 0.809901794
RAD17 1370.091764 0.000694317 0.286226939 0.002425757 0.998064528 0.998920501
RAD18 576.7841688 -0.024408864 0.299935769 -0.081380302 0.935139514
0.975231691
RAD21 8578.655794 0.380338179 0.225998591 1.682922791 0.092390039 0.348900321
RAD23A 1768.92665 0.294417953 0.255916072 1.150447296 0.24995969
0.562742997
RAD23B 8474.666284 -0.350068176 0.279676246 -1.251690771
0.210682564 0.518294063
RAD50 1875.074549 -0.255986666 0.271298956 -0.943559345 0.345394876
0.652226246
RAD51 594.8984288 0.182392475 0.333969611 0.546134943 0.584973182 0.815233156
RAD51AP1 1131.808962 0.700849515 0.294800063 2.377372335 0.017436478
0.149104584
RAD51B 122.1991875 -0.144245507 0.342627391 -0.420998177
0.673756413 0.85954302
RAD51C 522.6735373 0.717069835 0.288069468 2.489225397 0.012802177
0.128036515
RAD51D 383.9075274 -0.300198517 0.30328841 -0.989812032
0.322266003 0.632029271
RAD52 473.5561012 -0.106009968 0.450605578 -0.235261109 0.814006078
0.92227679
RAD54B 536.6653813 0.121920284 0.532275649 0.229054784 0.818826337
0.924316427
RAD54L 952.9147796 0.050675891 0.300302389 0.168749542 0.865993648

0.945587278
RAD54L2 1370.88756 -0.134730549 0.256963148 -0.524318565
0.600056986 0.824066673
RAD9A 412.3593822 0.293600618 0.30595376 0.959624153 0.337244409 0.64481093
RADIL 112.6281721 -0.223556328 0.618939076 -0.361192784 0.717955336
0.880671367
RADX 181.3392033 0.105358919 0.402677894 0.261645648 0.793594647 0.914378704
RAE1 1207.076625 0.070673487 0.265467777 0.26622247 0.790067879 0.912949942
RAF1 2958.816685 -0.07170573 0.283429146 -0.252993493 0.800273242
0.916366922
RAG1 83.69618973 -0.609017454 0.402486658 -1.513136998 0.130244863
0.41537625
RAI1 242.5610985 0.591277979 0.665015236 0.889119448 0.373938888 0.678520659
RAI14 2040.558766 0.32829172 0.401310732 0.818048695 0.413329383 0.705652445
RALA 2304.972356 0.173186901 0.308039397 0.562223217 0.573963945 0.808355645
RALB 1386.561031 0.312118863 0.250947378 1.243762199 0.213587097 0.520678417
RALBP1 3358.938965 -0.235864846 0.261941412 -0.900448862
0.367881428 0.674179083
RALGAPA1 641.7841397 -0.47644902 0.465119913 -1.024357388 0.305666501
0.615675317
RALGAPA2 742.1805064 -0.626538643 0.516060457 -1.214079929
0.224717204 0.53405032
RALGAPB 1924.633976 -0.480360485 0.371708775 -1.292303323
0.196252122 0.501911824
RALGDS 1220.08344 0.051017099 0.429595931 0.118756011 0.90546866
0.964473572
RALGPS1 561.491441 -0.583313439 0.343816811 -1.696582077
0.089775701 0.343069812
RALGPS2 1905.10578 -0.25526507 0.313934892 -0.813114682 0.41615231
0.708127819
RALY 3300.922289 0.524556932 0.376338939 1.393841768 0.163365392 0.464631632
RAMAC 1143.691754 0.104014281 0.382149008 0.272182523 0.785481679 0.910931322
RAMP1 26.26277755 -0.45129367 1.058721291 -0.426262959 0.669916246
0.857762448
RAMP2 34.76517075 1.23876969 0.613602745 2.018846396 0.043503187 0.239995219
RAN 17388.44509 0.599810759 0.306001282 1.960157669 0.049977364 0.257487003
RANBP1 5350.598884 1.218429657 0.331538008 3.675082878 0.000237772
0.010565846
RANBP10 738.9783181 -0.588445002 0.436269857 -1.34880967 0.177398108
0.481794969
RANBP17 367.9167159 -0.81135459 0.448838876 -1.807674498 0.070657176
0.30557013
RANBP2 4890.966651 -0.550534769 0.348030326 -1.581858614
0.113681851 0.388022186
RANBP3 1354.012699 -0.477783012 0.3692408 -1.293960504
0.195679064 0.501375962
RANBP6 1336.554136 0.196915112 0.278383622 0.707351642 0.479347981
0.751673525
RANBP9 1734.542773 -0.309104879 0.258194047 -1.197180505
0.231236207 0.540284656
RANGAP1 3547.834887 0.517385893 0.260662728 1.984886356 0.047157117
0.250147069
RANGRF 579.519602 1.108717431 0.383415884 2.891683615 0.003831836
0.063812059
RAP1A 892.8639973 0.453874681 0.30656084 1.480537048 0.138729981 0.427257927
RAP1B 2797.18944 0.274110163 0.256016256 1.070674835 0.284315662 0.598246396
RAP1GAP 139.655553 0.284981643 0.505622228 0.563625621 0.573008946
0.807760521
RAP1GAP2 451.9076264 -0.064963153 0.360428985 -0.180238425
0.856965393 0.942343063
RAP1GDS1 1043.477624 0.040679257 0.243840041 0.166827635 0.867505674
0.946451976
RAP2A 1269.765196 0.274753763 0.39902816 0.688557327 0.491101883 0.758593654
RAP2B 1334.049207 0.377032724 0.369746196 1.019706835 0.307867519 0.617648523

| | | | | | | |
|---|---|---|---|---|---|---|
| RAP2C | 1997.816368 | -0.242650647 | 0.398937977 | -0.608241534 | 0.543027292 | 0.78943022 |
| RAPGEF1 | 1766.807231 | -1.327656416 | 0.487312553 | -2.724445345 | 0.006440959 | 0.08724624 |
| RAPGEF2 | 1114.783326 | -0.751789482 | 0.32795684 | -2.292342742 | 0.021885873 | 0.167946678 |
| RAPGEF3 | 18.53865165 | 0.503513003 | 0.826295136 | 0.609362177 | 0.542284402 | 0.78908466 |
| RAPGEF4 | 15.70176922 | 1.64200591 | 0.883230159 | 1.859091759 | 0.063014133 | 0.289778852 |
| RAPGEF5 | 76.83818479 | 0.357233692 | 0.603834659 | 0.591608459 | 0.554112808 | 0.796360966 |
| RAPGEF6 | 559.0555762 | -0.115108518 | 0.375643255 | -0.306430415 | 0.759276953 | 0.899449351 |
| RAPGEFL1 | 443.186038 | -0.015318659 | 0.367759966 | -0.04165396 | 0.966774557 | 0.988749915 |
| RAPH1 | 819.0216076 | -1.010977804 | 0.35922368 | -2.814340645 | 0.00488774 | 0.074674632 |
| RARA | 241.5125189 | -0.786523701 | 0.509574444 | -1.543491261 | 0.122711626 | 0.403480295 |
| RARB | 204.4659171 | -0.833507928 | 0.797009385 | -1.045794371 | 0.295655981 | 0.60699582 |
| RARG | 1579.369185 | 0.282820067 | 0.311809802 | 0.907027503 | 0.364392253 | 0.671176869 |
| RARRES2 | 45.76521606 | 0.371071175 | 0.727803714 | 0.509850621 | 0.610156119 | 0.828605673 |
| RARS | 3608.748791 | 0.079880114 | 0.301115431 | 0.265280705 | 0.790793227 | 0.91323164 |
| RARS2 | 1335.035117 | 0.027379478 | 0.249630766 | 0.109679901 | 0.912663241 | 0.967485958 |
| RASA1 | 1806.783148 | -0.201267311 | 0.231428914 | -0.869672281 | 0.384479525 | 0.68518018 |
| RASA2 | 684.9230017 | 0.208775004 | 0.327311828 | 0.637847415 | 0.523573011 | 0.777662198 |
| RASA3 | 107.3835462 | 0.785771129 | 0.584673363 | 1.343948911 | 0.178964908 | 0.483466568 |
| RASA4 | 1274.260144 | 0.762005108 | 0.36570957 | 2.083634582 | 0.037193419 | 0.220938673 |
| RASA4B | 295.8453352 | 0.190949212 | 0.739335083 | 0.258271542 | 0.796197348 | 0.91536147 |
| RASA4CP | 25.37440004 | -0.177218881 | 0.598247481 | -0.296230049 | 0.767054402 | 0.903470997 |
| RASAL1 | 18.75606788 | -0.44584338 | 0.838102441 | -0.531967643 | 0.594748406 | 0.820937865 |
| RASAL2 | 760.7798448 | -0.632700561 | 0.311783407 | -2.029295169 | 0.042428235 | 0.236995123 |
| RASD1 | 18.8299924 | 0.260357306 | 0.783796526 | 0.332174611 | 0.739757414 | 0.891693924 |
| RASEF | 261.9218697 | -0.946930032 | 0.480386757 | -1.971182632 | 0.04870299 | 0.254345287 |
| RASGEF1A | 256.1428567 | -0.036913135 | 0.304109304 | -0.121381144 | 0.903389152 | 0.963418418 |
| RASGEF1B | 696.4531432 | -1.40811527 | 0.43553266 | -3.233087663 | 0.0012246 | 0.030831576 |
| RASGEF1C | 18.27850424 | -0.486224347 | 0.719186473 | -0.676075491 | 0.498992717 | 0.76281557 |
| RASGRF1 | 178.3262453 | -2.91236727 | 0.534247115 | -5.451348618 | 5.00E-08 | 2.53E-05 |
| RASGRF2 | 20.17993121 | -1.957491653 | 1.003453611 | -1.950754507 | 0.051086256 | 0.259989798 |
| RASGRP1 | 97.84547151 | -0.805476906 | 0.580326938 | -1.387970907 | 0.165145904 | 0.467343502 |
| RASGRP2 | 80.26195126 | -0.334137995 | 0.714509312 | -0.467646802 | 0.64003719 | 0.843538624 |
| RASGRP4 | 26.08878521 | -1.458805165 | 0.657503311 | -2.218703908 | 0.026506875 | 0.18608783 |
| RASIP1 | 62.66534753 | 0.46006424 | 0.541795126 | 0.849147985 | 0.395798952 | 0.69442187 |
| RASL10B | 1242.048612 | 1.143044795 | 0.611153331 | 1.870307722 | 0.061441098 | 0.285640507 |
| RASL11A | 16.89377952 | 1.146147863 | 0.847578865 | 1.352261022 | 0.176291829 | |

| | | | | | | |
|---|---|---|---|---|---|---|
| | | | | | | 0.480368206 |
| RASL11B | 667.9651992 | 2.322440679 | 0.741255504 | 3.133117618 | 0.001729601 | 0.03870174 |
| RASL12 | 509.8417007 | 0.001035676 | 0.36881863 | 0.00280809 | 0.997759472 | 0.998851994 |
| RASSF1 | 236.7852723 | 0.316498564 | 0.365350212 | 0.866288162 | 0.386332156 | 0.686679504 |
| RASSF10 | 14.48030858 | -1.144180061 | 1.040711216 | -1.099421284 | 0.271584351 | 0.583820065 |
| RASSF2 | 418.4818508 | -0.223338206 | 0.441161937 | -0.506249943 | 0.612681198 | 0.829404964 |
| RASSF3 | 1155.084756 | 0.158186517 | 0.364504017 | 0.433977432 | 0.66430483 | 0.85500751 |
| RASSF4 | 323.1665314 | 0.627903441 | 0.491401272 | 1.277781472 | 0.201326489 | 0.507615783 |
| RASSF5 | 11.57592187 | 0.726401306 | 1.263631587 | 0.574852127 | 0.565391309 | 0.803818907 |
| RASSF6 | 52.65662539 | -1.17331207 | 0.890500038 | -1.317587894 | 0.18764164 | 0.492618714 |
| RASSF7 | 449.034011 | 0.045366444 | 0.65039103 | 0.069752568 | 0.9443906 | 0.979313042 |
| RASSF8 | 1154.411926 | -0.042877498 | 0.321188627 | -0.133496315 | 0.893800878 | 0.958022688 |
| RASSF9 | 26.57560534 | -2.949765888 | 1.006298188 | -2.931303985 | 0.003375423 | 0.058860395 |
| RAVER1 | 416.6109908 | -0.439959561 | 0.919580264 | -0.478435193 | 0.63234049 | 0.838768903 |
| RAVER2 | 781.3504014 | -0.124417307 | 0.329728451 | -0.377332641 | 0.705926418 | 0.874325226 |
| RB1 | 782.1753451 | 0.18786584 | 0.418713438 | 0.448674017 | 0.653666831 | 0.84883826 |
| RB1CC1 | 2306.498798 | 0.082048729 | 0.313285683 | 0.261897473 | 0.793400488 | 0.914254706 |
| RBAK | 608.0151448 | -0.545932745 | 0.305017052 | -1.789843362 | 0.073479096 | 0.311470066 |
| RBBP4 | 8499.350994 | 0.284360266 | 0.254396014 | 1.117785857 | 0.263658463 | 0.577444856 |
| RBBP5 | 747.4650706 | -0.10493752 | 0.307859936 | -0.340861243 | 0.733208043 | 0.888457729 |
| RBBP6 | 3165.708698 | 0.237435904 | 0.286077013 | 0.829971976 | 0.406554628 | 0.701528693 |
| RBBP7 | 4051.005794 | 0.255216699 | 0.268699867 | 0.949820709 | 0.342203361 | 0.648945899 |
| RBBP8 | 2071.978283 | 0.177658078 | 0.362804437 | 0.489680005 | 0.624360353 | 0.834816841 |
| RBBP8NL | 273.5668614 | -1.656901021 | 0.749524768 | -2.210602093 | 0.027063404 | 0.188125165 |
| RBBP9 | 759.0729378 | -0.152730177 | 0.246211964 | -0.620319886 | 0.535047205 | 0.784173142 |
| RBCK1 | 823.0491126 | 0.006774842 | 0.347226805 | 0.019511288 | 0.984433232 | 0.994286144 |
| RBFA | 493.5499381 | 0.250072967 | 0.253339343 | 0.987106714 | 0.323590327 | 0.633361998 |
| RBFOX2 | 5335.296372 | -0.40331613 | 0.311663608 | -1.29407515 | 0.195639464 | 0.501359174 |
| RBFOX3 | 72.12130941 | 0.668830012 | 0.623175701 | 1.073260736 | 0.283154147 | 0.596961028 |
| RBKS | 110.5982652 | -0.452869157 | 0.569794617 | -0.794793674 | 0.426733532 | 0.714409006 |
| RBL1 | 331.5137773 | -0.305929631 | 0.389574786 | -0.785291149 | 0.432282884 | 0.718639059 |
| RBL2 | 1194.41547 | -0.264403641 | 0.294603262 | -0.897490542 | 0.369457218 | 0.67521115 |
| RBM10 | 1894.804835 | -0.063464809 | 0.438723004 | -0.144658037 | 0.884980872 | 0.954416194 |
| RBM11 | 92.84711828 | -0.277749289 | 0.434486463 | -0.639258787 | 0.522654594 | 0.777662198 |
| RBM12 | 4832.894772 | 0.102226869 | 0.285721754 | 0.357784689 | 0.720504449 | 0.88214152 |
| RBM12B | 1387.015713 | 0.276134971 | 0.43396937 | 0.6363006 | 0.524580514 | 0.777662198 |
| RBM14 | 1725.396814 | 0.105086609 | 0.260636329 | 0.403192483 | 0.686806629 | 0.865835142 |

| | | | | | | |
|---|---|---|---|---|---|---|
| RBM14-RBM4 | 56.22256049 | -0.753913466 | 0.637469027 | -1.182666818 | 0.23694122 | 0.547700719 |
| RBM15 | 466.164048 | 0.517142793 | 0.358648283 | 1.441921842 | 0.149324424 | 0.444022115 |
| RBM15B | 3189.127164 | -0.236194952 | 0.249901077 | -0.945153799 | 0.344580362 | 0.651368096 |
| RBM17 | 2549.759233 | 0.157491897 | 0.267126557 | 0.589577835 | 0.555473716 | 0.796360966 |
| RBM18 | 631.1850568 | 0.509914156 | 0.274320872 | 1.858823765 | 0.063052123 | 0.289778852 |
| RBM19 | 872.6128493 | -0.470262425 | 0.364056797 | -1.291728183 | 0.196451294 | 0.502167241 |
| RBM20 | 201.5170989 | -1.290587202 | 0.409174471 | -3.154124445 | 0.001609804 | 0.037175747 |
| RBM22 | 1389.310855 | -0.101710217 | 0.270926781 | -0.375415883 | 0.707351194 | 0.87520286 |
| RBM23 | 1514.502928 | 0.20850015 | 0.262056037 | 0.795631929 | 0.426246002 | 0.714238498 |
| RBM24 | 78.72223616 | 2.174181849 | 0.719972944 | 3.019810489 | 0.002529329 | 0.048634283 |
| RBM25 | 5516.948284 | 0.167546902 | 0.304119089 | 0.550925306 | 0.581684879 | 0.813002261 |
| RBM26 | 2812.727213 | 0.210413554 | 0.271051846 | 0.776285265 | 0.437580572 | 0.721501451 |
| RBM27 | 1486.563914 | 0.180603258 | 0.238753194 | 0.756443318 | 0.449383445 | 0.730561106 |
| RBM28 | 1553.485666 | -0.128139825 | 0.254168717 | -0.504152621 | 0.614154132 | 0.830316752 |
| RBM3 | 5189.722395 | -0.019794907 | 0.275617807 | -0.071820131 | 0.942745052 | 0.978574897 |
| RBM33 | 1724.170933 | -0.274511575 | 0.336894899 | -0.814828529 | 0.415170468 | 0.707398889 |
| RBM34 | 1896.187094 | 0.441774287 | 0.339300825 | 1.302013598 | 0.192911737 | 0.498908723 |
| RBM38 | 787.0541266 | 0.295719675 | 0.430853326 | 0.686358111 | 0.492487313 | 0.759174768 |
| RBM39 | 7650.15294 | -0.051787884 | 0.305823945 | -0.16933888 | 0.865530096 | 0.945285227 |
| RBM4 | 3820.300175 | 0.059320637 | 0.286805067 | 0.206832597 | 0.836140598 | 0.93270414 |
| RBM41 | 483.1336283 | -0.610866206 | 0.310246474 | -1.968970666 | 0.04895646 | 0.254849017 |
| RBM42 | 1210.839921 | 0.301970434 | 0.402645417 | 0.749966152 | 0.453275091 | 0.733497409 |
| RBM43 | 197.0988849 | 0.007891683 | 0.360353889 | 0.021899812 | 0.982527874 | 0.993518428 |
| RBM45 | 375.9931456 | 0.283054686 | 0.344673814 | 0.821224807 | 0.411518229 | 0.704686727 |
| RBM47 | 1354.155536 | -1.703735415 | 1.27403895 | -1.337271059 | 0.181134174 | 0.485749663 |
| RBM48 | 362.0271001 | -0.145290596 | 0.328893046 | -0.441756362 | 0.658665519 | 0.851304702 |
| RBM4B | 822.9464308 | -0.050912658 | 0.265765481 | -0.191569868 | 0.848079147 | 0.938207222 |
| RBM5 | 4494.346127 | -0.666683827 | 0.263705118 | -2.528141403 | 0.011466816 | 0.121279761 |
| RBM6 | 3120.709255 | -0.175802729 | 0.319156489 | -0.550835514 | 0.581746436 | 0.813002261 |
| RBM7 | 1499.432596 | -0.012660082 | 0.291756886 | -0.043392572 | 0.965388599 | 0.988700292 |
| RBM8A | 3895.743326 | 0.343693245 | 0.301339549 | 1.140551404 | 0.25405665 | 0.566474638 |
| RBMS1 | 4521.657499 | -0.107347203 | 0.368178971 | -0.291562559 | 0.770621106 | 0.905067213 |
| RBMS2 | 739.0325859 | -0.53850807 | 0.271335736 | -1.984655903 | 0.047182768 | 0.250195657 |
| RBMS3 | 54.5715312 | -0.389157973 | 0.578794861 | -0.672359068 | 0.501355134 | 0.763991735 |
| RBMX | 16748.7367 | 0.194230111 | 0.218724966 | 0.888010703 | 0.374534994 | 0.678946266 |
| RBMX2 | 1083.077725 | 0.517057269 | 0.404275642 | 1.278972103 | 0.20090688 | 0.506881795 |
| RBMXL1 | 776.4856695 | -0.274996834 | 0.262014534 | -1.049548015 | 0.293925969 | 0.606430922 |
| RBP1 | 422.5493449 | 0.171753856 | 0.612833336 | 0.280261934 | 0.779276553 | 0.908367621 |
| RBP5 | 20.13713198 | 0.249648656 | 0.720752387 | 0.346372291 | 0.729062946 | 0.886086417 |
| RBP7 | 18.05406543 | 0.716598849 | 0.963961284 | 0.743389657 | 0.457245804 | 0.736164289 |
| RBPJ | 2375.61851 | 0.394639302 | 0.32433257 | 1.216773579 | 0.223690375 | 0.532852544 |
| RBPMS | 3569.27854 | -0.384880342 | 0.54543481 | -0.705639492 | 0.480412358 | 0.752383182 |
| RBPMS2 | 1488.002604 | -0.358415008 | 0.36960744 | -0.969718057 | | |

| | | | | | | |
|---|---|---|---|---|---|---|
| | | | | | 0.332187047 | 0.640031711 |
| RBSN | 942.6565177 | -0.265477838 | 0.286350995 | -0.927106394 | 0.353871297 | 0.660911171 |
| RBX1 | 2486.23733 | 1.180294149 | 0.459028548 | 2.571287025 | 0.010132131 | 0.114285922 |
| RC3H1 | 694.3938328 | -0.520437062 | 0.38847033 | -1.339708653 | 0.180340083 | 0.484774033 |
| RC3H2 | 1425.034873 | -0.384453388 | 0.26326837 | -1.460309828 | 0.144204942 | 0.435369787 |
| RCAN1 | 248.2322512 | 0.470911808 | 0.30339431 | 1.552144493 | 0.120627654 | 0.400973879 |
| RCAN2 | 253.1969694 | -0.902063702 | 0.354705242 | -2.543136085 | 0.010986243 | 0.119221957 |
| RCAN3 | 819.107911 | -0.585149854 | 0.455406777 | -1.284894919 | 0.198829007 | 0.504782074 |
| RCBTB1 | 1973.774212 | -0.128974839 | 0.258132554 | -0.499645773 | 0.617324521 | 0.831835425 |
| RCBTB2 | 525.9234249 | 0.517019126 | 0.387399947 | 1.334587497 | 0.182011392 | 0.486131134 |
| RCC1 | 2598.586623 | 0.194011656 | 0.345958003 | 0.560795397 | 0.574937025 | 0.808375311 |
| RCC1L | 1173.686687 | 0.022948736 | 0.294674223 | 0.07787833 | 0.937924837 | 0.977015772 |
| RCC2 | 3571.022836 | 0.118990033 | 0.224860684 | 0.529172242 | 0.596685971 | 0.821812117 |
| RCCD1 | 546.343035 | 0.706985578 | 0.250703142 | 2.820010835 | 0.004802203 | 0.073664528 |
| RCE1 | 678.7873244 | 0.320973344 | 0.371130288 | 0.86485354 | 0.38711918 | 0.68748084 |
| RCHY1 | 658.8534815 | 0.076283128 | 0.303839296 | 0.251064063 | 0.80176458 | 0.917339075 |
| RCL1 | 394.7121489 | 0.507124583 | 0.298754145 | 1.697464589 | 0.08960886 | 0.342777615 |
| RCN1 | 7456.669436 | 0.679780237 | 0.275020811 | 2.47174108 | 0.013445686 | 0.131858113 |
| RCN1P2 | 23.5124457 | 0.487752913 | 0.864644419 | 0.564108091 | 0.572680572 | 0.807670159 |
| RCN2 | 6606.618346 | 0.93655556 | 0.293809944 | 3.18762377 | 0.001434471 | 0.034434101 |
| RCN3 | 119.3601487 | 1.423014901 | 0.530466628 | 2.682571956 | 0.007305843 | 0.094813472 |
| RCOR1 | 1775.714087 | -0.553296109 | 0.270197767 | -2.047744938 | 0.040584998 | 0.231842053 |
| RCOR2 | 2312.317051 | 0.451988204 | 0.393774716 | 1.147834501 | 0.251036892 | 0.564052234 |
| RCOR3 | 864.0566748 | -0.419147579 | 0.293094082 | -1.430078616 | 0.152694457 | 0.448798819 |
| RCSD1 | 72.12604146 | -2.816736654 | 1.107419468 | -2.543513759 | 0.010974373 | 0.119221957 |
| RD3 | 56.30602014 | 0.438280414 | 0.676264371 | 0.648090352 | 0.516926514 | 0.773950969 |
| RDH10 | 210.2731055 | 0.468564338 | 0.425481436 | 1.101256833 | 0.270784893 | 0.583394591 |
| RDH11 | 2255.970108 | -0.094759676 | 0.336291983 | -0.28177798 | 0.778113757 | 0.908058754 |
| RDH12 | 39.18689316 | 0.66013334 | 0.527978736 | 1.250302892 | 0.211188923 | 0.518607329 |
| RDH13 | 148.3999077 | 0.198741713 | 0.324140777 | 0.613133944 | 0.539787778 | 0.787188413 |
| RDH14 | 513.3300815 | 0.860512008 | 0.326416029 | 2.636243108 | 0.008382965 | 0.103556802 |
| RDH16 | 42.67614486 | -0.016659803 | 0.70965603 | -0.023475885 | 0.981270674 | 0.993251761 |
| RDH5 | 59.99776711 | 0.03600288 | 0.428936093 | 0.083935301 | 0.933107873 | 0.97462713 |
| RDM1 | 65.13580556 | 1.570353228 | 0.644462287 | 2.436687545 | 0.014822481 | 0.138467893 |
| RDM1P5 | 13.70148391 | 0.699353676 | 0.848554213 | 0.824170885 | 0.409842468 | 0.703966274 |
| RDX | 4100.94962 | 0.023003476 | 0.223781917 | 0.10279417 | 0.918126332 | 0.969677217 |
| REC8 | 697.2037669 | 0.0018163 | 0.424481357 | 0.004278869 | 0.996585967 | 0.998431718 |
| RECK | 320.9784129 | 0.761950205 | 0.290867774 | 2.619575879 | 0.008803918 | 0.106171897 |
| RECQL | 1356.169829 | -0.087551292 | 0.263945118 | -0.331702637 | 0.74011381 | 0.891917214 |
| RECQL4 | 664.6513999 | 0.564357797 | 0.503250306 | 1.12142564 | 0.262106727 | 0.575601582 |
| RECQL5 | 828.1771084 | -0.732042705 | 0.35047422 | -2.08872055 | 0.036732886 | 0.219399453 |
| REEP1 | 58.67589802 | -1.428215948 | 0.481657703 | -2.9652094 | 0.00302477 | 0.054402443 |
| REEP2 | 1423.710974 | 0.859365219 | 0.36773982 | 2.336883778 | 0.019445228 | 0.15817885 |
| REEP3 | 1504.904574 | 0.149321156 | 0.251337757 | 0.594105548 | 0.55244152 | 0.795662343 |
| REEP4 | 487.7167709 | 0.60760964 | 0.25975439 | 2.339169857 | 0.019326644 | 0.157622736 |
| REEP5 | 1517.382092 | 0.079246799 | 0.39881729 | 0.198704522 | 0.842493886 | 0.935549316 |

| | | | | | | |
|---|---|---|---|---|---|---|
| REEP6 | 182.5555403 | -0.286352919 | 0.427678582 | -0.669551692 | 0.503143618 | 0.765232263 |
| REL | 548.0915229 | -0.302653624 | 0.347509866 | -0.870920953 | 0.383797315 | 0.684699412 |
| RELA | 596.7048389 | -0.5425376 | 0.322044664 | -1.684665704 | 0.092053087 | 0.348263686 |
| RELB | 97.550235 | -0.063392264 | 0.428795024 | -0.147838152 | 0.882470499 | 0.953255639 |
| RELCH | 816.3397216 | -0.346840266 | 0.297374001 | -1.166343609 | 0.243475548 | 0.554786353 |
| RELL1 | 396.29485 | 0.34864711 | 0.5710376 | 0.610550181 | 0.541497413 | 0.788468879 |
| RELL2 | 68.59727098 | -0.068216612 | 0.510402843 | -0.133652492 | 0.893677374 | 0.957957991 |
| RELN | 18.6085062 | -2.989389634 | 1.35931922 | -2.199181466 | 0.027865022 | 0.190853383 |
| RELT | 196.1470853 | 0.776044146 | 0.36624617 | 2.118914023 | 0.034097734 | 0.210632641 |
| REM2 | 30.98813345 | 0.499133766 | 0.615704343 | 0.810671179 | 0.417554529 | 0.709057506 |
| RENBP | 283.6780487 | 0.470618858 | 0.449336879 | 1.047363081 | 0.294932152 | 0.606966925 |
| REPIN1 | 1937.851749 | 0.061335483 | 0.374091621 | 0.163958451 | 0.869763852 | 0.947589759 |
| REPS1 | 2915.972043 | -0.299001123 | 0.321668279 | -0.929532511 | 0.352613185 | 0.659729494 |
| REPS2 | 35.58797065 | -0.724977128 | 0.617855094 | -1.173377277 | 0.240644545 | 0.552353982 |
| RER1 | 1630.253145 | 0.064285869 | 0.309549322 | 0.20767569 | 0.835482201 | 0.932531417 |
| RERE | 2985.662881 | -0.671038205 | 0.222666244 | -3.013650351 | 0.002581251 | 0.049382295 |
| RERG | 13.32754713 | -1.213747945 | 1.018030586 | -1.192250961 | 0.233162857 | 0.543032499 |
| RESF1 | 4093.826655 | -1.477781031 | 0.506144711 | -2.919680871 | 0.0035039 | 0.060751619 |
| REST | 2310.940532 | -0.077342734 | 0.313526554 | -0.246686391 | 0.80515094 | 0.918762309 |
| RET | 72.19843018 | 0.503479388 | 0.575131341 | 0.875416365 | 0.381347399 | 0.682967936 |
| RETREG1 | 79.29641556 | -0.944012042 | 0.702655408 | -1.343492175 | 0.179112658 | 0.483594614 |
| RETREG2 | 1724.122452 | -0.269668955 | 0.281763847 | -0.957074365 | 0.338529722 | 0.645678097 |
| RETREG3 | 949.8890272 | -0.144857604 | 0.324315472 | -0.446656471 | 0.655123116 | 0.849331122 |
| RETSAT | 714.2024739 | -0.410143081 | 0.435718238 | -0.941303451 | 0.346549375 | 0.653186802 |
| REV1 | 963.94689 | 0.068092506 | 0.32013189 | 0.212701415 | 0.831559863 | 0.930997148 |
| REV3L | 590.9212919 | -0.786574155 | 0.382670874 | -2.055484774 | 0.039832205 | 0.229888601 |
| REX1BD | 365.0937111 | 0.51249586 | 0.456810862 | 1.121899462 | 0.261905189 | 0.575432821 |
| REXO1 | 529.2815856 | -0.479604938 | 0.415692194 | -1.153750168 | 0.248602609 | 0.561242587 |
| REXO1L1P | 20.38902329 | 0.588539087 | 0.81949308 | 0.718174566 | 0.472649658 | 0.747011977 |
| REXO2 | 1195.151844 | -0.358452076 | 0.471730711 | -0.75986589 | 0.447334752 | 0.728949036 |
| REXO4 | 529.7577143 | -0.282849615 | 0.293797086 | -0.962737985 | 0.335679029 | 0.644086928 |
| REXO5 | 129.0129105 | 0.622037194 | 0.444132781 | 1.400565823 | 0.161343947 | 0.46167973 |
| RFC1 | 2345.481947 | 0.035984782 | 0.237757047 | 0.151351066 | 0.879698789 | 0.952123159 |
| RFC2 | 1141.84485 | 0.290734487 | 0.249631587 | 1.164654244 | 0.244158977 | 0.555457814 |
| RFC3 | 1275.044163 | 0.421226625 | 0.264392093 | 1.59318919 | 0.111117756 | 0.383475916 |
| RFC4 | 1765.629728 | 0.477706681 | 0.305677554 | 1.562779716 | 0.118104417 | 0.395958766 |
| RFC5 | 992.5610236 | 0.361509879 | 0.254732743 | 1.419173185 | 0.155848532 | 0.454251311 |
| RFESD | 55.76520225 | 0.809826369 | 0.466351586 | 1.736514668 | 0.082472873 | 0.328570367 |
| RFFL | 681.6505367 | -0.495704275 | 0.247575369 | -2.002235835 | 0.045259373 | 0.244986785 |

| | | | | | | |
|---|---|---|---|---|---|---|
| RFK | 546.4987585 | 0.706763517 | 0.291078214 | 2.428087997 | 0.01517866 | 0.139730249 |
| RFLNA | 107.196003 | -0.998164807 | 0.764147142 | -1.306246863 | 0.191468621 | 0.496966715 |
| RFLNB | 39.07520169 | 0.73340114 | 0.526840164 | 1.392075225 | 0.16389962 | 0.465769928 |
| RFNG | 657.7004152 | -0.035334103 | 0.403243055 | -0.087624828 | 0.930174869 | 0.973652838 |
| RFT1 | 703.4204832 | -0.065212325 | 0.239345467 | -0.272461083 | 0.785267511 | 0.910931322 |
| RFTN2 | 202.6830663 | 0.048802339 | 0.600423245 | 0.081279896 | 0.935219362 | 0.975231691 |
| RFWD3 | 2387.445042 | -0.171637124 | 0.331092715 | -0.518395955 | 0.604182036 | 0.82576988 |
| RFX1 | 286.8858294 | 0.037054955 | 0.378634027 | 0.097864831 | 0.922039626 | 0.970640219 |
| RFX2 | 149.3355371 | 0.653181221 | 0.422231627 | 1.546973696 | 0.121869589 | 0.40263143 |
| RFX3 | 745.8859323 | -0.115154405 | 0.284097091 | -0.405334686 | 0.685231514 | 0.864502893 |
| RFX4 | 124.7140523 | 2.419466931 | 1.126050911 | 2.148630144 | 0.03166373 | 0.202602468 |
| RFX5 | 1926.213489 | -0.069548784 | 0.383581858 | -0.181314061 | 0.85612107 | 0.942245037 |
| RFX7 | 1364.073973 | -0.182003308 | 0.398275192 | -0.456978772 | 0.647686304 | 0.846597949 |
| RFXANK | 1105.906085 | 0.301136443 | 0.393016477 | 0.766218367 | 0.443546381 | 0.726848446 |
| RFXAP | 322.2988455 | 0.539228867 | 0.329870848 | 1.63466663 | 0.102118952 | 0.366611924 |
| RGCC | 23.97659661 | 0.286375054 | 1.036763207 | 0.27622031 | 0.782378859 | 0.909909835 |
| RGL1 | 414.7014887 | 0.214359208 | 0.292772405 | 0.732170125 | 0.464064736 | 0.741119432 |
| RGL2 | 495.0986419 | -1.10791384 | 0.416801363 | -2.658133919 | 0.007857466 | 0.099586986 |
| RGL3 | 489.3345361 | -1.53912853 | 0.592971181 | -2.595621135 | 0.009442011 | 0.109934578 |
| RGMA | 1856.495416 | 0.817059325 | 0.423211992 | 1.930614774 | 0.053530709 | 0.267406937 |
| RGMB | 891.3219788 | 1.248508373 | 0.535900191 | 2.329740488 | 0.019819871 | 0.159346723 |
| RGN | 13.77763746 | 0.723402862 | 0.884996573 | 0.817407529 | 0.413695574 | 0.705778726 |
| RGP1 | 1259.801154 | -0.550222409 | 0.410613758 | -1.339999934 | 0.180245366 | 0.484754911 |
| RGPD1 | 42.75302485 | -0.361063927 | 0.855163745 | -0.42221613 | 0.672867269 | 0.859151631 |
| RGPD3 | 10.37834577 | -1.352087906 | 1.046726792 | -1.291729529 | 0.196450828 | 0.502167241 |
| RGPD5 | 231.6405361 | -0.613339898 | 0.700569849 | -0.875487146 | 0.381308901 | 0.682967936 |
| RGPD6 | 1742.491783 | -0.439093289 | 0.265587817 | -1.653288519 | 0.098272159 | 0.359965304 |
| RGPD8 | 217.0622137 | -0.819709248 | 0.84168918 | -0.973885928 | 0.330113181 | 0.638717861 |
| RGS10 | 278.8738838 | 0.342523192 | 0.508699259 | 0.673331415 | 0.500736471 | 0.763638219 |
| RGS12 | 692.1609529 | 0.088259712 | 0.470746868 | 0.187488686 | 0.851277494 | 0.939477675 |
| RGS14 | 188.7272142 | 0.721389602 | 0.412874743 | 1.747235971 | 0.080596415 | 0.325022513 |
| RGS16 | 257.568587 | -0.432112821 | 0.776251727 | -0.556665841 | 0.577755759 | 0.809936484 |
| RGS17 | 58.61544175 | 0.847884112 | 0.507040067 | 1.672223099 | 0.094480337 | 0.352523659 |
| RGS19 | 83.48690811 | 0.790990751 | 0.476460442 | 1.660139396 | 0.096886413 | 0.357434422 |
| RGS2 | 316.059404 | -0.218168923 | 0.326157169 | -0.668907336 | 0.503554591 | 0.765396923 |
| RGS20 | 21.61379145 | 0.592923418 | 0.652815678 | 0.908255481 | 0.363743257 | 0.670353372 |
| RGS3 | 529.4360155 | 0.454166843 | 0.315317689 | 1.44034686 | 0.14976929 | 0.444386837 |
| RGS4 | 330.2399426 | -5.742710674 | 1.610911035 | -3.564883813 | 0.000364017 | 0.014277637 |
| RGS5 | 1282.515718 | 0.110461952 | 0.446590575 | 0.247345014 | 0.804641225 | 0.918595231 |
| RGS6 | 87.1966786 | 0.120828496 | 0.438587498 | 0.275494619 | 0.78293626 | 0.909909835 |
| RGS8 | 23.17614872 | 0.838716829 | 0.965828111 | 0.868391404 | 0.385180099 | 0.685935162 |
| RGS9 | 189.0718244 | 0.338571184 | 0.380052556 | 0.89085359 | 0.373007721 | 0.67811865 |
| RHBDD1 | 574.3940229 | -0.409424394 | 0.312542734 | -1.309978923 | 0.190202966 | 0.495717092 |
| RHBDD2 | 861.8230429 | 0.131912149 | 0.292980414 | 0.45024221 | 0.652535804 | |

0.848578973
RHBDD3	632.4727291 0.205625695 0.427541695 0.480948869 0.630552839
0.838177265
RHBDF1	973.4259386 -0.601670548	0.352867051 -1.705091323
0.088177394 0.341577124
RHBDF2	212.1329277 0.074235914 0.324604017 0.228696842 0.819104549
0.924316427
RHBDL1	43.08799963 0.778326679 0.705122307 1.103817978 0.269672105
0.582376584
RHBDL3	682.7200701 -1.85053946 0.467277316 -3.960259566	7.49E-05
0.004975677
RHCE	14.64188436 1.116423741 0.863316018 1.293180848 0.195948518 0.501797297
RHEB	2426.923272 0.987386965 0.297815169 3.315435443 0.000915005 0.025850161
RHEBL1	24.22763534 1.222440744 0.812177925 1.505139091 0.132288321
0.418521131
RHEBP2	29.83744867 0.427857893 0.551041613 0.776452963 0.437481583
0.721468765
RHNO1 1002.710227 0.474704872 0.29391919 1.615086348 0.106291997 0.374140185
RHOA	13812.55173 0.197442394 0.36617139 0.539207593 0.589743621 0.81828099
RHOB	1423.947477 -0.177563701	0.509362219 -0.348600063	0.727389581
0.885092921
RHOBTB1	345.2172634 -0.14539933 0.398755262 -0.364633006	0.715385374
0.879026964
RHOBTB2	860.9004237 -2.360448909	0.60858851 -3.878563055
0.000105075 0.006358893
RHOBTB3	855.0443532 -1.729742022	0.492217597 -3.514181596
0.000441111 0.016402199
RHOC	1812.81353 -0.583405274	0.397247639 -1.468618606	0.141936265
0.432219004
RHOD	88.00664926 -0.129409233	0.635451285 -0.203649338	0.838627535
0.933879959
RHOF	568.0826723 0.29741456 0.379015928 0.784702009 0.432628305 0.718639059
RHOG	435.5255459 -0.043001487	0.294956428 -0.145789287	0.884087734
0.954046354
RHOQ	718.9477485 0.249484189 0.395408401 0.630953181 0.528071137 0.779319497
RHOQP1	10.39760875 0.275729698 1.142927818 0.241248567 0.809362471
0.919900963
RHOT1 1439.087478 0.155335805 0.353115225 0.439901181 0.660008681 0.852207761
RHOT2 1999.063727 0.197051733 0.466185469 0.422689565 0.67252177 0.859011927
RHOU	1676.053378 -0.599335717	0.383849384 -1.561382515	0.118433525
0.396722236
RHOV	157.4243907 0.913006197 0.821882076 1.110872501 0.266623234 0.579921585
RHOXF1P1	10.17807298 -4.382929214	2.067344187 -2.12007717 0.033999538
0.210275984
RHOXF1P3	24.5971547 -3.075467155	1.594604874 -1.928670359
0.053771798 0.267993414
RHPN1 477.7100812 0.097567025 0.648083285 0.150547047 0.880333036 0.952483146
RHPN2 1154.006821 0.294028873 0.314025278 0.936322309 0.349107265 0.655726918
RIBC1 46.80914867 -0.698748759	0.548584552 -1.273730287	0.202759019
0.509290789
RIBC2 18.36694087 -0.617944253	0.719426793 -0.858939727	0.39037377
0.690412738
RIC1	976.1350178 -0.323638327	0.349077481 -0.927124621	0.353861834
0.660911171
RIC3	335.8402382 -0.010507231	0.290754605 -0.036137797	0.971172484
0.989629517
RIC8A 2188.498303 -0.351911649	0.290228653 -1.212532412	0.225308643
0.53459368
RIC8B 508.1628795 0.645756625 0.439315556 1.469915229 0.141584715 0.431581616
RICTOR	805.1737544 -0.457911926	0.295103011 -1.551701977
0.120733549 0.400973879
RIDA	973.5437763 1.196024437 0.348103246 3.435832479 0.000590736 0.019567812
RIF1	2346.069359 -0.075861118	0.371761432 -0.204058602	0.838307705
0.933879959

| | | | | | | |
|---|---|---|---|---|---|---|
| RILP | 17.96927736 | 0.414033935 | 0.726803515 | 0.56966419 | 0.568905483 | 0.805450586 |
| RILPL1 | 387.9282566 | 0.313802423 | 0.27604151 | 1.136794331 | 0.255624274 | 0.567910953 |
| RILPL2 | 207.6315069 | 0.706673659 | 0.325268064 | 2.172588511 | 0.029811301 | 0.197226575 |
| RIMBP2 | 21.7900374 | -1.546287352 | 0.812910523 | -1.902161811 | 0.057150004 | 0.276650512 |
| RIMKLA | 36.80996076 | 0.289852187 | 0.549552359 | 0.527433251 | 0.597892758 | 0.822658356 |
| RIMKLB | 2324.93346 | -0.792750194 | 0.271978761 | -2.914750374 | 0.003559731 | 0.061273717 |
| RIMS1 | 59.87091257 | -1.006424677 | 0.634497976 | -1.586174762 | 0.112699671 | 0.38618767 |
| RIMS2 | 276.8218374 | -0.702260642 | 0.34414131 | -2.040617098 | 0.041288901 | 0.233467731 |
| RIMS3 | 1016.621715 | -0.233621593 | 0.459473848 | -0.508454604 | 0.611134568 | 0.829058038 |
| RIMS4 | 267.7157044 | 0.644579469 | 0.53065924 | 1.214676803 | 0.224489383 | 0.533874112 |
| RIN1 | 60.30615587 | 1.111628196 | 0.569030238 | 1.953548548 | 0.05075463 | 0.259171491 |
| RIN2 | 476.3060708 | -0.262801239 | 0.304787017 | -0.862245515 | 0.388552428 | 0.688604437 |
| RIN3 | 81.35310424 | -2.153354385 | 0.666077784 | -3.232887263 | 0.001225459 | 0.030831576 |
| RING1 | 709.7061313 | -0.159520818 | 0.449491064 | -0.354892078 | 0.722670444 | 0.882953475 |
| RINL | 72.46618664 | 1.012103674 | 0.422977779 | 2.392805783 | 0.016720089 | 0.146249211 |
| RINT1 | 705.4526346 | 0.064669233 | 0.243063044 | 0.266059502 | 0.790193384 | 0.913025425 |
| RIOK1 | 911.2798513 | 0.426071592 | 0.257295652 | 1.655961106 | 0.097729696 | 0.359375073 |
| RIOK2 | 1070.174462 | 0.342392357 | 0.242872662 | 1.409760795 | 0.158610327 | 0.458338527 |
| RIOK3 | 1504.99307 | -0.299681731 | 0.226982623 | -1.320284908 | 0.186739913 | 0.491419118 |
| RIOX1 | 461.4735468 | 0.302525371 | 0.265998129 | 1.137321424 | 0.255403942 | 0.567820048 |
| RIOX2 | 438.0359253 | -0.245048825 | 0.281088753 | -0.871784525 | 0.383325938 | 0.684494911 |
| RIPK1 | 425.1187547 | -0.65075922 | 0.308927743 | -2.106509481 | 0.03516012 | 0.214222565 |
| RIPK2 | 622.6511319 | 0.194766756 | 0.336771119 | 0.578335687 | 0.563037505 | 0.801847267 |
| RIPK4 | 394.7801522 | -0.113749433 | 0.371086898 | -0.306530448 | 0.759200801 | 0.899449351 |
| RIPOR1 | 738.0527697 | -1.263653133 | 0.342963756 | -3.684509254 | 0.000229144 | 0.010346247 |
| RIPOR2 | 439.0324043 | -0.028258382 | 0.779775067 | -0.036239145 | 0.971091673 | 0.989629517 |
| RIPOR3 | 22.71840099 | 0.061848546 | 0.859136475 | 0.071989198 | 0.942610505 | 0.978574897 |
| RIPPLY3 | 43.21134394 | 1.131716236 | 0.867839456 | 1.304061746 | 0.192212531 | 0.498075763 |
| RIT1 | 905.6833287 | 0.313427586 | 0.322412884 | 0.972131082 | 0.330985341 | 0.639084337 |
| RITA1 | 424.6875307 | 0.27288992 | 0.296250369 | 0.921146264 | 0.356974069 | 0.664415851 |
| RLF | 912.3509518 | -0.141579043 | 0.37093385 | -0.381682725 | 0.702696715 | 0.872703476 |
| RLIM | 1343.614355 | -0.232062899 | 0.308967206 | -0.751092331 | 0.452597091 | 0.733138807 |
| RLN1 | 25.52636905 | 0.439585313 | 0.822879243 | 0.534203915 | 0.593200461 | 0.819859199 |
| RLN2 | 114.5292492 | 0.976966504 | 0.577573599 | 1.691501319 | 0.090741096 | 0.345245839 |
| RMC1 | 397.9309554 | -0.143612703 | 0.279767228 | -0.513329257 | 0.607721022 | 0.827550096 |
| RMDN1 | 1115.036383 | 0.125066695 | 0.252146575 | 0.496007906 | 0.619888833 | 0.832886335 |
| RMDN2 | 91.09858495 | -0.090562252 | 0.415850779 | -0.217775839 | 0.82760377 | 0.928323608 |
| RMDN3 | 658.8671963 | -0.007318419 | 0.261392997 | -0.027997764 | 0.977663935 | 0.99226352 |
| RMI1 | 722.3018477 | 0.356781717 | 0.299578616 | 1.190945206 | 0.233675096 | 0.543724676 |
| RMI2 | 961.2276837 | 0.64137976 | 0.379808856 | 1.688690904 | 0.09127868 | 0.346543756 |

| | | | | | | |
|---|---|---|---|---|---|---|
| RMND1 | 877.5859613 | -0.571377676 | 0.25028208 | -2.28293482 | 0.022434208 | 0.170430331 |
| RMND5A | 2454.529689 | 0.086587059 | 0.251430571 | 0.344377609 | 0.730562325 | 0.886859359 |
| RMND5B | 727.2104741 | 0.097546977 | 0.255631556 | 0.381592077 | 0.702763962 | 0.872703476 |
| RNA5-8SP6 | 226.6384939 | 1.425716985 | 0.4559072 | 3.127208751 | 0.001764746 | 0.039027635 |
| RNA5SP199 | 11.46234946 | -0.001805691 | 0.884842912 | -0.002040691 | 0.998371766 | 0.998964388 |
| RNASE1 | 30.77983013 | -0.817203084 | 0.672276796 | -1.215575324 | 0.224146739 | 0.533486437 |
| RNASE10 | 42.7250763 | -0.456226489 | 0.577966628 | -0.789364761 | 0.429898846 | 0.716554273 |
| RNASE4 | 49.2111678 | -0.449342655 | 0.784289198 | -0.572929802 | 0.566692225 | 0.804513376 |
| RNASEH1 | 1495.086075 | 0.027320722 | 0.251017281 | 0.108840005 | 0.913329393 | 0.967876517 |
| RNASEH2A | 912.7050881 | 0.39087515 | 0.239535844 | 1.631802341 | 0.102721132 | 0.367636293 |
| RNASEH2B | 486.7368542 | 0.413532986 | 0.308471332 | 1.340588068 | 0.180054233 | 0.484754911 |
| RNASEH2C | 501.3420628 | 0.94589761 | 0.396606347 | 2.3849785 | 0.01708012 | 0.14773715 |
| RNASEK | 1630.318437 | 0.467537006 | 0.46314253 | 1.009488387 | 0.312740468 | 0.622907489 |
| RNASEK-C17orf49 | 24.62413009 | 0.019234858 | 0.785383531 | 0.024491038 | 0.980460932 | 0.993221987 |
| RNASEL | 136.0062791 | -0.912722417 | 0.34886956 | -2.616228305 | 0.008890709 | 0.10668882 |
| RNASET2 | 457.8531745 | 0.246666051 | 0.498661823 | 0.494655976 | 0.620842982 | 0.833349572 |
| RND1 | 405.4446861 | -0.330783928 | 0.402323398 | -0.822184168 | 0.410972088 | 0.704426187 |
| RND2 | 2848.28205 | 0.746464816 | 0.344773944 | 2.165084773 | 0.030381194 | 0.198840851 |
| RND3 | 743.1640824 | -0.476536598 | 0.632410152 | -0.753524585 | 0.451134743 | 0.732309564 |
| RNF10 | 2762.478976 | -0.379526087 | 0.232499566 | -1.632373313 | 0.102600867 | 0.367636293 |
| RNF103 | 556.0050336 | -0.069256504 | 0.282254154 | -0.2453693 | 0.806170497 | 0.919090941 |
| RNF11 | 2527.617997 | 0.287973899 | 0.342160498 | 0.841633973 | 0.399992867 | 0.697652919 |
| RNF111 | 856.2116723 | -0.241142542 | 0.377914344 | -0.638087827 | 0.52341651 | 0.777662198 |
| RNF112 | 53.43096733 | 0.927860985 | 0.686558142 | 1.351467456 | 0.176545738 | 0.480766848 |
| RNF113A | 337.9915199 | 0.266650794 | 0.411616587 | 0.647813531 | 0.517105563 | 0.773950969 |
| RNF114 | 1688.45189 | 0.175983047 | 0.305990303 | 0.575126223 | 0.565205934 | 0.803630668 |
| RNF115 | 892.2375409 | -0.212360078 | 0.256431069 | -0.828137085 | 0.407592869 | 0.702164665 |
| RNF121 | 684.7596332 | -0.529133556 | 0.3251452 | -1.627376189 | 0.103657225 | 0.369064484 |
| RNF122 | 411.7851238 | 0.389376518 | 0.281745608 | 1.382014507 | 0.166967247 | 0.469215449 |
| RNF123 | 830.7652084 | -0.222533026 | 0.451722561 | -0.492632084 | 0.622272574 | 0.833640266 |
| RNF125 | 198.3996228 | 0.761668164 | 0.409219479 | 1.861270546 | 0.06270598 | 0.288977044 |
| RNF126 | 449.7024781 | -0.088607553 | 0.432813575 | -0.204724524 | 0.837787359 | 0.933603058 |
| RNF13 | 650.9067853 | -0.435654159 | 0.319540837 | -1.36337553 | 0.172764189 | 0.476461804 |

| | | | | | | |
|---|---|---|---|---|---|---|
| RNF130 | 2864.889457 | 0.645601124 | 0.3304682 | 1.953595305 | 0.050749096 | 0.259171491 |
| RNF135 | 381.6409203 | -0.472383176 | 0.329907713 | -1.431864601 | 0.152182575 | 0.448119169 |
| RNF138 | 1330.008663 | 0.549325219 | 0.266712916 | 2.059612364 | 0.039435613 | 0.228786879 |
| RNF139 | 813.9263005 | 0.249327976 | 0.275637953 | 0.904548786 | 0.365704478 | 0.672171387 |
| RNF14 | 649.0954269 | 0.096029026 | 0.29209667 | 0.328757688 | 0.742338847 | 0.892150449 |
| RNF141 | 1034.568614 | 0.154629218 | 0.257981409 | 0.599381246 | 0.54891868 | 0.793468239 |
| RNF144A | 885.1236042 | -0.23637366 | 0.308628983 | -0.765882898 | 0.443745983 | 0.726848446 |
| RNF144B | 55.5715716 | -0.70670311 | 0.740368335 | -0.954529086 | 0.339815893 | 0.646629378 |
| RNF145 | 2476.293217 | 0.339614949 | 0.337399675 | 1.006565727 | 0.314143511 | 0.623858474 |
| RNF146 | 1158.297187 | -0.145767731 | 0.245470789 | -0.593829234 | 0.552626334 | 0.795662343 |
| RNF149 | 1504.38439 | -0.052613203 | 0.585284105 | -0.089893443 | 0.928371892 | 0.972633911 |
| RNF150 | 115.4571508 | -0.00468606 | 0.408351194 | -0.011475563 | 0.990844027 | 0.996726674 |
| RNF152 | 189.6250186 | -0.332353454 | 0.357870253 | -0.928698184 | 0.353045522 | 0.660100273 |
| RNF157 | 350.9145431 | 0.379722277 | 0.502432434 | 0.755767843 | 0.4497884 | 0.731062762 |
| RNF165 | 359.8258984 | 1.255631942 | 0.521035679 | 2.409877084 | 0.015957896 | 0.142746012 |
| RNF166 | 310.8932559 | 0.388223006 | 0.452746095 | 0.857485046 | 0.391176876 | 0.690625437 |
| RNF167 | 2564.17178 | -0.125208414 | 0.308196084 | -0.406262183 | 0.684549969 | 0.863857185 |
| RNF168 | 946.8336733 | 0.406434922 | 0.333558064 | 1.218483276 | 0.223040376 | 0.532026087 |
| RNF169 | 551.0082102 | 0.109792983 | 0.447709495 | 0.245232643 | 0.806276303 | 0.919090941 |
| RNF170 | 703.9535828 | -0.045368454 | 0.24287581 | -0.186796923 | 0.85181986 | 0.939761856 |
| RNF175 | 76.35345217 | 1.53539032 | 0.566545015 | 2.710094131 | 0.006726412 | 0.089750563 |
| RNF180 | 94.01769669 | -0.363302976 | 0.377503255 | -0.96238369 | 0.335856903 | 0.644086928 |
| RNF181 | 661.5341579 | 0.723687937 | 0.376880291 | 1.920206375 | 0.054831837 | 0.270962148 |
| RNF182 | 423.1000129 | -0.046033726 | 0.341153918 | -0.134935357 | 0.892662984 | 0.957550733 |
| RNF185 | 1182.712125 | -0.394780543 | 0.265252179 | -1.488321581 | 0.136666106 | 0.424728769 |
| RNF187 | 1015.123229 | 0.071988133 | 0.312728021 | 0.230194061 | 0.817940977 | 0.924112493 |
| RNF19A | 673.3417538 | -0.421571812 | 0.306616262 | -1.374916676 | 0.169157279 | 0.471613806 |
| RNF19B | 708.8618515 | -0.731024621 | 0.261898721 | -2.791249302 | 0.005250501 | 0.077110704 |
| RNF2 | 1178.030821 | 0.384036498 | 0.248779761 | 1.543680626 | 0.122665722 | 0.403440492 |
| RNF20 | 1776.794121 | -0.088112252 | 0.238664053 | -0.369189458 | 0.711986513 | 0.877461202 |
| RNF207 | 61.41820593 | 0.836397212 | 0.523835997 | 1.596677616 | 0.110337588 | 0.382176151 |
| RNF208 | 18.8528469 | -0.197122498 | 0.970525205 | -0.203109097 | 0.839049761 | 0.933915137 |
| RNF212 | 16.13919438 | -0.019264886 | 0.77399037 | -0.024890343 | 0.98014243 | 0.993196129 |

| | | | | | | |
|---|---|---|---|---|---|---|
| RNF214 | 990.3514056 | -0.425310002 | 0.235014025 | -1.809721787 | 0.070338942 | 0.304732874 |
| RNF215 | 587.8142514 | 0.10775453 | 0.307712012 | 0.350179799 | 0.726203766 | 0.8844924 |
| RNF216 | 988.6502712 | -0.284176921 | 0.296109307 | -0.959702767 | 0.33720483 | 0.64481093 |
| RNF216P1 | 575.1221371 | -0.418451976 | 0.272267399 | -1.536915464 | 0.124314012 | 0.405846326 |
| RNF217 | 497.6084944 | -0.342257455 | 0.32237543 | -1.061673515 | 0.288383927 | 0.601673803 |
| RNF219 | 817.7833014 | 0.022227978 | 0.335809751 | 0.066192176 | 0.947224826 | 0.980790989 |
| RNF220 | 2122.341806 | 0.291172511 | 0.22855103 | 1.273993433 | 0.202665744 | 0.509290789 |
| RNF227 | 35.32339221 | 0.512816958 | 0.572488736 | 0.895767769 | 0.370376807 | 0.676008968 |
| RNF24 | 589.3259063 | -0.485060926 | 0.377629311 | -1.284489609 | 0.198970697 | 0.504782074 |
| RNF25 | 467.3745645 | 0.284773396 | 0.326298285 | 0.87273948 | 0.382805093 | 0.683935 |
| RNF26 | 1821.157558 | 0.051040059 | 0.377724579 | 0.135125066 | 0.892512992 | 0.957550733 |
| RNF31 | 696.2384506 | -0.582405599 | 0.312649715 | -1.86280547 | 0.06248964 | 0.288330393 |
| RNF32 | 40.1715247 | 0.477384357 | 0.514006792 | 0.928751068 | 0.353018109 | 0.660100273 |
| RNF34 | 1199.517644 | -0.035554274 | 0.267099799 | -0.133112319 | 0.894104552 | 0.958161736 |
| RNF38 | 1470.062041 | -0.355474729 | 0.325420657 | -1.092354533 | 0.2746773 | 0.587581685 |
| RNF4 | 2074.828041 | -0.006670981 | 0.280265188 | -0.023802388 | 0.981010235 | 0.993241851 |
| RNF40 | 2107.275842 | -0.521568944 | 0.400429163 | -1.302524872 | 0.192737021 | 0.498908723 |
| RNF41 | 1491.734596 | 0.096019503 | 0.232606586 | 0.41279787 | 0.679754712 | 0.861682789 |
| RNF43 | 1314.039447 | -0.433988702 | 0.331556926 | -1.308941749 | 0.190554083 | 0.496058205 |
| RNF44 | 1932.914472 | 0.103354337 | 0.295364966 | 0.349920773 | 0.726398157 | 0.884630646 |
| RNF5 | 1993.528532 | 0.926387199 | 0.363516158 | 2.54840721 | 0.010821607 | 0.118235065 |
| RNF6 | 589.6833697 | -0.642316431 | 0.273661715 | -2.347118348 | 0.018919244 | 0.15607605 |
| RNF7 | 2115.896373 | 0.38491991 | 0.420560986 | 0.915253488 | 0.36005857 | 0.667522123 |
| RNF8 | 904.60188 | 0.324634412 | 0.266177099 | 1.219618115 | 0.222609676 | 0.531509505 |
| RNFT1 | 235.1324982 | -0.256665822 | 0.388107249 | -0.661327052 | 0.508402597 | 0.7688373 |
| RNFT2 | 443.5827965 | 0.862717977 | 0.353126729 | 2.443083192 | 0.01456238 | 0.137477204 |
| RNGTT | 1060.219647 | 0.169655858 | 0.343015902 | 0.494600563 | 0.620882105 | 0.833349572 |
| RNH1 | 419.375428 | 0.374118196 | 0.460422971 | 0.812553282 | 0.416474226 | 0.708268589 |
| RNLS | 149.2148877 | 0.223286825 | 0.331542937 | 0.673477851 | 0.500643335 | 0.763638219 |
| RNMT | 1499.577689 | -0.443570352 | 0.336833068 | -1.316884816 | 0.187877237 | 0.492689771 |
| RNPC3 | 767.6989831 | 0.187467952 | 0.263652521 | 0.71104176 | 0.477058358 | 0.749806646 |
| RNPEP | 2952.799446 | 0.068139961 | 0.359820865 | 0.1893719 | 0.849801346 | 0.938580023 |
| RNPEPL1 | 279.1109535 | -0.944701609 | 0.520256694 | -1.815837489 | 0.069395305 | 0.30336211 |
| RNPS1 | 4990.37714 | 0.310830187 | 0.216355868 | 1.436661692 | 0.150814145 | 0.445743501 |
| RO60 | 1098.92128 | -0.069167606 | 0.307922474 | -0.224626688 | 0.822269701 | 0.925831501 |
| ROBO1 | 2366.084627 | -0.35849154 | 0.313405377 | -1.143858932 | 0.252682145 | 0.565423035 |
| ROBO2 | 47.45465601 | 0.02699485 | 0.685620528 | 0.039372874 | 0.968593107 | 0.989475309 |
| ROBO3 | 162.1941177 | -0.332167423 | 0.414951494 | -0.800496994 | 0.423422905 | 0.712688899 |
| ROCK1 | 2567.503892 | -0.1248739 | 0.227248022 | -0.549504892 | 0.582659008 | 0.813577525 |
| ROCK1P1 | 49.76307517 | 0.799275552 | 0.597884033 | 1.336840438 | 0.181274726 | 0.485749663 |
| ROCK2 | 1124.14986 | -0.147576696 | 0.32019826 | -0.460891625 | 0.644876361 | |

| | | | | | | |
|---|---|---|---|---|---|---|
| | | | | | | 0.845794147 |
| ROGDI | 884.1276912 | -0.127035802 | 0.355536651 | -0.357307192 | 0.720861846 | 0.882227889 |
| ROM1 | 24.38830245 | 0.135954524 | 0.705209009 | 0.192786142 | 0.847126457 | 0.938145276 |
| ROMO1 | 894.6456841 | 1.292222227 | 0.543525857 | 2.377480686 | 0.017431356 | 0.149104584 |
| ROR1 | 973.7642435 | 0.044132331 | 0.346670124 | 0.127303532 | 0.898700165 | 0.960250212 |
| ROR2 | 1934.035367 | -0.349409792 | 0.277395328 | -1.259609503 | 0.207810266 | 0.514566354 |
| RORA | 312.5982073 | 0.190839686 | 0.422983342 | 0.451175416 | 0.651863126 | 0.848003386 |
| RORB | 50.61256575 | 1.42353796 | 0.599517337 | 2.374473385 | 0.017574003 | 0.14971347 |
| RP2 | 364.9156987 | -0.019645802 | 0.347367028 | -0.056556323 | 0.954898628 | 0.984323736 |
| RP9 | 434.7886367 | 0.776751672 | 0.477625288 | 1.626278366 | 0.103890452 | 0.369319502 |
| RP9P | 29.3398906 | 2.21907192 | 0.62181986 | 3.568673281 | 0.000358794 | 0.014212158 |
| RPA1 | 5356.418955 | 0.046377044 | 0.255995926 | 0.181163212 | 0.856239469 | 0.942245037 |
| RPA2 | 1329.635873 | 0.308154888 | 0.246203368 | 1.251627424 | 0.210705657 | 0.518294063 |
| RPA3 | 1543.84497 | 1.367007996 | 0.425326301 | 3.214021781 | 0.001308897 | 0.032235849 |
| RPAIN | 2544.771519 | 0.424663234 | 0.337629571 | 1.257778556 | 0.208471853 | 0.515521839 |
| RPAP1 | 1289.325768 | -0.007875121 | 0.290221179 | -0.027134895 | 0.978352142 | 0.992395125 |
| RPAP2 | 824.536577 | -0.101664528 | 0.256566437 | -0.396250304 | 0.691920385 | 0.86812349 |
| RPAP3 | 1502.804886 | 0.401501872 | 0.239594062 | 1.675758861 | 0.093785435 | 0.351226741 |
| RPE | 1546.104626 | 0.138609245 | 0.243243709 | 0.569836917 | 0.568788314 | 0.805450586 |
| RPF1 | 1183.12159 | 0.645080848 | 0.277999339 | 2.320440218 | 0.020317075 | 0.161005801 |
| RPF2 | 1255.376602 | 0.851762932 | 0.32257937 | 2.640475527 | 0.008278977 | 0.102951117 |
| RPGR | 302.7765231 | -0.0350753 | 0.289061547 | -0.121341977 | 0.903420173 | 0.963418418 |
| RPGRIP1L | 540.9909328 | 0.226727212 | 0.309144688 | 0.733401612 | 0.463313516 | 0.740745005 |
| RPH3A | 33.33376418 | -2.29170976 | 0.721295584 | -3.177213073 | 0.001486977 | 0.035303498 |
| RPH3AL | 31.49639744 | -0.749863934 | 0.576176106 | -1.301449204 | 0.193104741 | 0.498908723 |
| RPIA | 650.5584798 | 0.554438515 | 0.352876363 | 1.571197658 | 0.116136744 | 0.392757589 |
| RPL10 | 30039.34298 | 0.300594441 | 0.248047817 | 1.211840703 | 0.225573364 | 0.53459368 |
| RPL10A | 35613.31088 | 0.698210194 | 0.389134716 | 1.79426344 | 0.072771131 | 0.310464236 |
| RPL10P16 | 12.51831586 | 0.083123353 | 0.880226441 | 0.094434056 | 0.924764364 | 0.971780852 |
| RPL10P9 | 916.6469276 | 1.18549407 | 0.496579184 | 2.387321312 | 0.016971651 | 0.147615406 |
| RPL11 | 34513.19349 | 0.487511728 | 0.367621227 | 1.326125079 | 0.18479827 | 0.489365432 |
| RPL12 | 36125.87775 | 0.371030838 | 0.329651179 | 1.125525589 | 0.26036638 | 0.573630315 |
| RPL13 | 28808.15676 | 1.00068668 | 0.466158412 | 2.146666569 | 0.031819836 | 0.203172868 |
| RPL13A | 94507.87474 | 0.731703154 | 0.427561858 | 1.7113387 | 0.087018606 | 0.338762965 |
| RPL13AP7 | 17.67649591 | 1.199034572 | 0.969450147 | 1.236819218 | 0.216154206 | 0.523980406 |
| RPL13P12 | 214.7599748 | 0.612576002 | 0.632508135 | 0.968487151 | 0.332801133 | 0.640564069 |
| RPL13P5 | 79.66256896 | 0.016703456 | 0.465784272 | 0.035860927 | 0.971393252 | 0.989658654 |
| RPL14 | 23021.26642 | 0.351470631 | 0.317911065 | 1.105562751 | 0.26891582 | 0.581986006 |
| RPL14P1 | 43.35782052 | 0.199213778 | 0.599561516 | 0.332265786 | 0.739688572 | 0.89168179 |
| RPL15 | 52590.75338 | 0.595603701 | 0.344618301 | 1.728299684 | 0.08393451 | 0.331434265 |
| RPL17 | 55215.72647 | 0.959985802 | 0.411937564 | 2.330415788 | 0.019784186 | 0.159228588 |
| RPL17-C18orf32 | 720.7888995 | 0.975874917 | 0.294894453 | 3.309234571 | 0.000935514 | 0.026041626 |
| RPL18 | 22979.60357 | 0.627788913 | 0.418149235 | 1.501351337 | 0.133264713 | 0.419566119 |
| RPL18A | 28389.58945 | 0.583879815 | 0.416984384 | 1.400243841 | 0.161440312 | 0.46167973 |
| RPL19 | 35648.67599 | 0.54775362 | 0.353147162 | 1.55106335 | 0.120886503 | 0.401219509 |

| | | | | | | |
|---|---|---|---|---|---|---|
| RPL21 | 22110.3232 | 1.004897131 | 0.424367855 | 2.367985983 | 0.017885214 | 0.150854036 |
| RPL21P16 | 11543.95325 | 1.285530412 | 0.478475118 | 2.686723644 | 0.007215663 | 0.093964649 |
| RPL21P28 | 10.10114185 | 1.188289221 | 0.962895335 | 1.234079322 | 0.217173348 | 0.52522507 |
| RPL21P75 | 12.39458205 | -0.062333435 | 0.855365302 | -0.072873467 | 0.941906808 | 0.978574897 |
| RPL22 | 24811.97626 | 0.746276448 | 0.401675386 | 1.857909333 | 0.063181891 | 0.289778852 |
| RPL22L1 | 925.5618504 | 1.206586996 | 0.411354215 | 2.933206839 | 0.003354803 | 0.058568145 |
| RPL22P1 | 2684.30844 | 1.469958048 | 0.529543817 | 2.775895024 | 0.005504998 | 0.079368821 |
| RPL23 | 37152.34147 | 0.720145866 | 0.368162698 | 1.956053315 | 0.050458874 | 0.258046297 |
| RPL23A | 23668.45799 | 0.954388887 | 0.484298958 | 1.970660625 | 0.048762707 | 0.254481951 |
| RPL23AP42 | 9639.472257 | 0.13989571 | 0.251736509 | 0.555722771 | 0.578400387 | 0.810209453 |
| RPL23AP49 | 32.88637742 | 0.538005759 | 0.686024898 | 0.784236492 | 0.432901356 | 0.718657207 |
| RPL23AP7 | 76.63803903 | 0.740827275 | 0.418488725 | 1.770244288 | 0.076686454 | 0.317958513 |
| RPL23AP82 | 202.1725452 | 0.783271711 | 0.319274699 | 2.453284628 | 0.014155828 | 0.135106002 |
| RPL23AP87 | 13.95190227 | -1.136819275 | 0.848356406 | -1.340025568 | 0.180237033 | 0.484754911 |
| RPL24 | 31850.53861 | 1.020470003 | 0.455133749 | 2.242132134 | 0.024952834 | 0.180688185 |
| RPL26 | 37714.7839 | 0.717150047 | 0.360039873 | 1.991862846 | 0.046386108 | 0.247615641 |
| RPL26L1 | 915.8444551 | 0.976839951 | 0.338433108 | 2.886360486 | 0.003897255 | 0.064617764 |
| RPL27 | 25844.60975 | 0.900846304 | 0.427016011 | 2.109631212 | 0.034890134 | 0.213176891 |
| RPL27A | 32199.26665 | 0.829329197 | 0.470643952 | 1.762115915 | 0.078049715 | 0.320406654 |
| RPL28 | 18608.63805 | 0.335212184 | 0.409654724 | 0.818279795 | 0.413197441 | 0.705607651 |
| RPL29 | 23507.12614 | 0.474478506 | 0.325368786 | 1.458279117 | 0.144763624 | 0.435928729 |
| RPL3 | 68139.50538 | -0.019613634 | 0.221928979 | -0.088377976 | 0.929576266 | 0.973331598 |
| RPL30 | 24337.75154 | 0.814989552 | 0.390169745 | 2.08880766 | 0.036725041 | 0.219399453 |
| RPL31 | 35751.40339 | 1.001904873 | 0.416409512 | 2.406056646 | 0.016125764 | 0.143654707 |
| RPL32 | 27872.95366 | 0.781491623 | 0.382786433 | 2.041586523 | 0.041192563 | 0.233096746 |
| RPL32P29 | 268.9426978 | 0.792943984 | 0.422551987 | 1.876559589 | 0.06057848 | 0.284105089 |
| RPL32P3 | 363.1447688 | -0.313572353 | 0.384969975 | -0.814537167 | 0.415337289 | 0.70750977 |
| RPL34 | 14578.15606 | 1.006460456 | 0.430159848 | 2.339735937 | 0.019297378 | 0.157609505 |
| RPL35 | 15639.30617 | 0.941511509 | 0.399942742 | 2.354115751 | 0.018566828 | 0.154343756 |
| RPL35A | 21193.06094 | 1.015533287 | 0.455460479 | 2.229684756 | 0.025768379 | 0.183192161 |
| RPL36 | 9576.287052 | 1.163227208 | 0.496246157 | 2.344052829 | 0.019075469 | 0.156852604 |
| RPL36A | 23642.58689 | 1.054999533 | 0.41372731 | 2.549987655 | 0.010772673 | 0.117984572 |
| RPL36A-HNRNPH2 | 22.17760075 | 0.047433635 | 0.911160771 | 0.052058469 | 0.958482105 | 0.985438602 |
| RPL36AL | 4613.619029 | 0.914579947 | 0.499176634 | 1.832176997 | 0.066925055 | 0.297817842 |
| RPL37 | 34457.42525 | 0.821454062 | 0.419297985 | 1.959117601 | 0.050099016 | 0.257487003 |
| RPL37A | 43754.97944 | 1.085744372 | 0.514768563 | 2.109189353 | 0.034928241 | 0.213323807 |
| RPL38 | 14025.18736 | 1.00945484 | 0.527407549 | 1.913993917 | 0.055620933 | 0.272730921 |
| RPL39 | 20694.41563 | 1.149464998 | 0.439562267 | 2.615022001 | 0.008922171 | 0.10668882 |
| RPL39L | 198.575344 | 0.763969965 | 0.430075909 | 1.776360753 | 0.075673476 | 0.315569626 |
| RPL39P40 | 10.82432356 | 1.564149758 | 1.014595837 | 1.541648113 | 0.123159127 | 0.404688568 |
| RPL3P4 | 3479.198105 | 5.479456939 | 1.694713462 | 3.233264538 | 0.001223842 | |

0.030831576

| | | | | | | |
|---|---|---|---|---|---|---|
| RPL4 | 81444.23536 | 0.210617733 | 0.282598238 | 0.745290325 | 0.456096229 | 0.735399427 |
| RPL41 | 75530.46719 | 1.081522242 | 0.43964889 | 2.459968093 | 0.013894937 | 0.134161613 |
| RPL4P4 | 14.05910336 | 0.819594205 | 0.780157455 | 1.050549732 | 0.293465438 | 0.606314063 |
| RPL5 | 52569.35413 | 0.598417525 | 0.330058891 | 1.813062883 | 0.069822123 | 0.303603614 |
| RPL6 | 54857.48854 | 0.73097852 | 0.346410598 | 2.11015057 | 0.034845389 | 0.213176891 |
| RPL6P27 | 87.17689914 | 0.381449069 | 0.533937381 | 0.714407873 | 0.474975005 | 0.748737095 |
| RPL7 | 63671.02543 | 0.759317702 | 0.346229896 | 2.19310265 | 0.028299984 | 0.192356209 |
| RPL7A | 29271.32354 | 0.487741423 | 0.390108424 | 1.250271445 | 0.211200406 | 0.518607329 |
| RPL7L1 | 3717.552087 | 0.159931399 | 0.269463063 | 0.593518819 | 0.552833992 | 0.795672359 |
| RPL7P1 | 14.54952829 | 0.846031761 | 0.817332909 | 1.035112807 | 0.300616232 | 0.611214323 |
| RPL7P9 | 32.38826585 | 1.268315543 | 0.711740593 | 1.781991299 | 0.074750651 | 0.313842348 |
| RPL8 | 39360.87718 | 0.534677929 | 0.408701131 | 1.30823697 | 0.190792946 | 0.496058205 |
| RPL9 | 27194.6858 | 0.928563112 | 0.391352648 | 2.372701746 | 0.017658517 | 0.150007193 |
| RPL9P7 | 15.20218469 | 0.364701204 | 0.91016951 | 0.400695915 | 0.688644019 | 0.867149602 |
| RPL9P9 | 1480.918965 | 0.525435701 | 0.550575284 | 0.954339426 | 0.339911857 | 0.646629378 |
| RPLP0 | 92783.19568 | 0.624122616 | 0.367126429 | 1.700020935 | 0.089126988 | 0.342662324 |
| RPLP0P2 | 60.56964975 | -0.295125396 | 0.948545531 | -0.311134665 | 0.755698249 | 0.897924353 |
| RPLP0P6 | 15.86978236 | 0.433811273 | 0.811375154 | 0.534661767 | 0.592883764 | 0.819859199 |
| RPLP1 | 28059.05515 | 1.05440338 | 0.505816812 | 2.084555821 | 0.037109638 | 0.220693972 |
| RPLP1P6 | 11.40516674 | 1.674387057 | 1.05488529 | 1.58726932 | 0.112451661 | 0.385686647 |
| RPLP2 | 14265.01502 | 1.061503066 | 0.529672093 | 2.00407588 | 0.045061933 | 0.244454795 |
| RPN1 | 5410.391595 | -0.061038582 | 0.225616212 | -0.270541651 | 0.786743579 | 0.911399422 |
| RPN2 | 8235.874474 | -0.090069457 | 0.221674753 | -0.406313556 | 0.684512227 | 0.863857185 |
| RPP14 | 769.9310838 | 0.370960361 | 0.299580676 | 1.238265319 | 0.215617699 | 0.523214349 |
| RPP25 | 375.6242147 | 1.136034208 | 0.556331604 | 2.042009119 | 0.041150627 | 0.232946328 |
| RPP25L | 428.8978404 | 0.951735105 | 0.400361637 | 2.377188564 | 0.017445168 | 0.149104584 |
| RPP30 | 1218.672683 | 0.544709392 | 0.288030039 | 1.891154806 | 0.058603682 | 0.280288923 |
| RPP38 | 584.5060128 | 0.722926574 | 0.446002125 | 1.620903879 | 0.105038255 | 0.371346423 |
| RPP40 | 298.1115707 | 0.602872167 | 0.416105038 | 1.448846115 | 0.147380562 | 0.440921023 |
| RPRD1A | 3125.210888 | 0.329900433 | 0.282164223 | 1.169178818 | 0.242331593 | 0.553259986 |
| RPRD1B | 869.0966077 | 0.058661377 | 0.237301254 | 0.247202138 | 0.80475179 | 0.918652326 |
| RPRD2 | 1119.652073 | -0.40000311 | 0.305736919 | -1.308324526 | 0.19076326 | 0.496058205 |
| RPRM | 158.9862374 | 0.637369474 | 0.37729915 | 1.689294754 | 0.091162958 | 0.346217714 |
| RPS10 | 32896.29257 | 0.820154843 | 0.409981299 | 2.000468912 | 0.045449654 | 0.245452751 |
| RPS10-NUDT3 | 781.7368042 | 0.917279327 | 0.280575678 | 3.269275986 | 0.001078231 | 0.028433348 |
| RPS10P7 | 13.30958592 | -0.231412343 | 0.907981998 | -0.254864462 | 0.798827785 | 0.916056735 |
| RPS11 | 41836.31528 | 0.561112588 | 0.374171767 | 1.499612311 | 0.133714857 | 0.420405914 |
| RPS12 | 37766.89498 | 0.93527704 | 0.419882078 | 2.227475497 | 0.025915512 | 0.183721602 |
| RPS13 | 24687.10639 | 1.032854198 | 0.470637804 | 2.19458401 | 0.028193452 | 0.192229399 |
| RPS14 | 32340.85665 | 1.09594849 | 0.47445124 | 2.309928603 | 0.020892107 | 0.163885294 |
| RPS15 | 9188.042444 | 0.839610612 | 0.445925159 | 1.882850957 | 0.059720569 | 0.282669434 |
| RPS15A | 30570.43253 | 0.893995811 | 0.383360012 | 2.332000683 | 0.019700655 | 0.158893482 |
| RPS15AP10 | 10.57550374 | 0.752026289 | 1.548734387 | 0.485574734 | 0.627268713 | 0.836041947 |

| | | | | | | |
|---|---|---|---|---|---|---|
| RPS16 | 23752.99289 | 0.896574544 | 0.465762776 | 1.924959639 | 0.054234408 | 0.269175122 |
| RPS17 | 46247.89429 | 0.891155306 | 0.392096533 | 2.272795683 | 0.023038495 | 0.173530066 |
| RPS18 | 52672.27994 | 0.704741634 | 0.393380588 | 1.791500788 | 0.073212968 | 0.311211528 |
| RPS18P9 | 21.6309979 | 0.982655854 | 0.865290066 | 1.135637508 | 0.256108304 | 0.568292977 |
| RPS19 | 44609.36811 | 0.925475421 | 0.459685707 | 2.013278652 | 0.044085323 | 0.241443505 |
| RPS19BP1 | 972.7061733 | 0.902797574 | 0.440599363 | 2.049021515 | 0.040460011 | 0.231601821 |
| RPS2 | 57869.05377 | 0.765062873 | 0.417712608 | 1.831553223 | 0.067018011 | 0.297817842 |
| RPS20 | 42670.35788 | 0.908246714 | 0.472767809 | 1.921126391 | 0.054715775 | 0.270592163 |
| RPS20P22 | 90.50712144 | 0.355791275 | 0.591582361 | 0.601423063 | 0.547558242 | 0.792759743 |
| RPS21 | 12787.05957 | 1.226446868 | 0.495183157 | 2.476754008 | 0.013258325 | 0.130781562 |
| RPS23 | 44953.79575 | 1.070653924 | 0.41012222 | 2.610572826 | 0.009039072 | 0.1074956 |
| RPS24 | 35510.51661 | 0.808141166 | 0.333028607 | 2.426641886 | 0.01523929 | 0.139780768 |
| RPS25 | 16674.77029 | 1.011387673 | 0.421291988 | 2.400681006 | 0.016364595 | 0.144378309 |
| RPS26 | 4622.199257 | 0.926164744 | 0.473035677 | 1.957917315 | 0.050239716 | 0.257653791 |
| RPS27 | 29542.15641 | 0.847123969 | 0.409846637 | 2.066929171 | 0.038740829 | 0.226725106 |
| RPS27A | 41977.97713 | 1.086426161 | 0.470799776 | 2.307618264 | 0.021020378 | 0.164636113 |
| RPS27L | 3373.678621 | 1.214115637 | 0.318906612 | 3.807119672 | 0.000140595 | 0.007590614 |
| RPS28 | 9831.247263 | 1.164208907 | 0.568369545 | 2.048330909 | 0.040527586 | 0.231666922 |
| RPS28P7 | 2379.290458 | 0.620178176 | 0.370254176 | 1.675006567 | 0.093932943 | 0.351340153 |
| RPS29 | 16497.97439 | 1.348005449 | 0.501434429 | 2.688298551 | 0.007181715 | 0.093602925 |
| RPS2P46 | 22.9895331 | -0.004607291 | 0.68811531 | -0.006695522 | 0.994657786 | 0.997946781 |
| RPS2P5 | 6918.535558 | 0.408416784 | 0.258210623 | 1.581719521 | 0.113713614 | 0.388022186 |
| RPS3 | 62651.73446 | 0.657170169 | 0.442487879 | 1.485171008 | 0.13749853 | 0.425712285 |
| RPS3A | 68029.41579 | 0.812848662 | 0.354683712 | 2.291756384 | 0.021919705 | 0.168121254 |
| RPS3AP26 | 72.43118813 | 1.01439074 | 1.12027606 | 0.905482833 | 0.365209651 | 0.671700076 |
| RPS3AP5 | 13.75715653 | 1.825525386 | 0.830229968 | 2.19881895 | 0.027890798 | 0.190943729 |
| RPS3AP6 | 37.15422199 | 1.028794638 | 0.525084094 | 1.959294998 | 0.050078249 | 0.257487003 |
| RPS4X | 22875.88428 | 0.284242017 | 0.337754324 | 0.841564405 | 0.40003182 | 0.697652919 |
| RPS4XP16 | 20.43087901 | -0.172207711 | 0.754406593 | -0.228269096 | 0.819437048 | 0.924316427 |
| RPS4Y1 | 8466.552903 | 0.417548841 | 0.334047701 | 1.249967714 | 0.211311342 | 0.518628728 |
| RPS5 | 20109.9723 | 0.696386195 | 0.369426694 | 1.885045686 | 0.05942367 | 0.282486927 |
| RPS6 | 70458.11441 | 0.697583878 | 0.429447607 | 1.624374816 | 0.104295839 | 0.370122145 |
| RPS6KA1 | 519.3657968 | -0.292623753 | 0.35267001 | -0.829738126 | 0.406686861 | 0.701597267 |
| RPS6KA2 | 379.0601567 | 0.000447477 | 0.463298342 | 0.000965851 | 0.999229362 | 0.999361108 |
| RPS6KA3 | 714.728988 | -0.437553768 | 0.322832295 | -1.355359344 | 0.175303096 | 0.479279739 |
| RPS6KA4 | 355.6643739 | -0.771626083 | 0.431896628 | -1.786599 | 0.074002324 | 0.312639724 |
| RPS6KA5 | 219.8689305 | 0.217907879 | 0.367251106 | 0.593348463 | 0.552947972 | 0.795672359 |
| RPS6KA6 | 421.557966 | -0.469331595 | 0.290089062 | -1.617887941 | 0.10568675 | 0.372686009 |
| RPS6KB1 | 778.0984801 | -0.235342298 | 0.283683356 | -0.829595013 | 0.406767797 | 0.701626421 |
| RPS6KB2 | 874.3659696 | -0.031108058 | 0.307318574 | -0.101224139 | 0.919372535 | 0.969677217 |
| RPS6KC1 | 770.4145116 | -0.391091082 | 0.305250259 | -1.281214576 | 0.200118308 | 0.506083488 |
| RPS6KL1 | 111.786689 | -0.677051559 | 0.4514013 | -1.499888368 | | |

| | | | | | | |
|---|---|---|---|---|---|---|
| | | | | | 0.133643322 | 0.420294949 |
| RPS7 | 36567.60658 | 0.718747995 | 0.353886561 | 2.031012401 | 0.042253737 | 0.236528876 |
| RPS7P1 | 10.25981447 | 0.203294466 | 0.971466861 | 0.209265467 | 0.834241009 | 0.932107564 |
| RPS8 | 48926.56077 | 0.885496967 | 0.444077541 | 1.994014298 | 0.046150492 | 0.247053322 |
| RPS9 | 4919.213154 | 1.455219462 | 0.47555154 | 3.06006676 | 0.002212877 | 0.044579397 |
| RPSA | 71161.245 | 0.576433454 | 0.352431351 | 1.63559074 | 0.10192527 | 0.366163454 |
| RPSAP14 | 16.96695059 | 1.646635596 | 0.817295372 | 2.014737453 | 0.043932166 | 0.241112355 |
| RPSAP41 | 12.70524181 | -1.081405099 | 0.873272508 | -1.23833636 | 0.215591367 | 0.523214349 |
| RPSAP58 | 119.4766748 | -0.925049464 | 0.429079276 | -2.155894066 | 0.031091938 | 0.200942211 |
| RPTOR | 512.408388 | -0.594808962 | 0.534915158 | -1.111968792 | 0.266151569 | 0.57970088 |
| RPUSD1 | 375.8306293 | 0.056059574 | 0.530811056 | 0.105611166 | 0.915890865 | 0.969169304 |
| RPUSD2 | 346.6382818 | 0.182396227 | 0.324521645 | 0.562046414 | 0.574084396 | 0.808359708 |
| RPUSD3 | 1029.064121 | -0.151453512 | 0.288947852 | -0.524155175 | 0.600170615 | 0.824148118 |
| RPUSD4 | 1754.266263 | 0.091720737 | 0.258980462 | 0.354160837 | 0.723218352 | 0.883273431 |
| RRAGA | 1179.119293 | 0.347182762 | 0.341153452 | 1.017673307 | 0.308833237 | 0.618441002 |
| RRAGB | 492.2344712 | -0.24841923 | 0.338667922 | -0.733518628 | 0.46324217 | 0.740745005 |
| RRAGC | 769.7181506 | -0.052817895 | 0.340274428 | -0.155221465 | 0.876646727 | 0.950324661 |
| RRAGD | 215.7965727 | 1.064807515 | 0.480100699 | 2.2178837 | 0.026562763 | 0.186369517 |
| RRAS | 325.8489058 | 0.827584927 | 0.486703223 | 1.700389248 | 0.089057733 | 0.342496199 |
| RRAS2 | 1904.30984 | 0.395565357 | 0.288845848 | 1.36946873 | 0.170852803 | 0.474118872 |
| RRBP1 | 3484.814731 | -1.300212145 | 0.368096004 | -3.532263676 | 0.000412018 | 0.015709208 |
| RREB1 | 1653.222464 | -1.508890633 | 0.303446477 | -4.972509978 | 6.61E-07 | 0.000139266 |
| RRM1 | 3887.320997 | 0.513467171 | 0.25769664 | 1.992525668 | 0.046313412 | 0.247615641 |
| RRM2 | 1472.96138 | 0.493046434 | 0.36657686 | 1.345001522 | 0.178624743 | 0.482972162 |
| RRM2B | 1651.1803 | 0.82070846 | 0.409531615 | 2.004017346 | 0.045068202 | 0.244454795 |
| RRN3 | 574.8342941 | 0.004352909 | 0.251931963 | 0.017278115 | 0.986214745 | 0.994937085 |
| RRN3P1 | 304.5194705 | 0.069310399 | 0.388930436 | 0.178207701 | 0.858559862 | 0.94248999 |
| RRN3P3 | 44.3890732 | -0.841306196 | 0.575938342 | -1.460757402 | 0.14408203 | 0.435085286 |
| RRNAD1 | 657.8462203 | 0.231849837 | 0.271751556 | 0.853168387 | 0.393565932 | 0.692506234 |
| RRP1 | 828.3421754 | 0.145736301 | 0.251813778 | 0.578746334 | 0.562760347 | 0.801731358 |
| RRP12 | 840.9602234 | -0.06889602 | 0.327419912 | -0.210420985 | 0.833339116 | 0.931794496 |
| RRP15 | 958.9723188 | 0.536930838 | 0.308935501 | 1.738003035 | 0.082210279 | 0.328245329 |
| RRP1B | 1704.929909 | 0.139040659 | 0.306557242 | 0.45355529 | 0.650148939 | 0.847430646 |
| RRP36 | 1196.2408 | 0.12139569 | 0.337007785 | 0.360216278 | 0.718685403 | 0.880993556 |
| RRP7A | 1073.919486 | 0.099186131 | 0.289983289 | 0.342040851 | 0.732320149 | 0.888091845 |
| RRP7BP | 368.0356551 | 0.553577181 | 0.334223603 | 1.656307862 | 0.097659489 | 0.359260938 |
| RRP8 | 312.15424 | 0.239406525 | 0.317436049 | 0.754188206 | 0.450736218 | 0.732054295 |
| RRP9 | 945.5166791 | 0.468351454 | 0.297161582 | 1.576083456 | 0.115006571 | 0.390153107 |
| RRS1 | 587.2895106 | 0.630455711 | 0.417738626 | 1.509210957 | 0.131244881 | 0.416813082 |
| RSAD1 | 640.127676 | 0.25725094 | 0.29717621 | 0.865651189 | 0.386681475 | 0.687008387 |
| RSBN1 | 612.4087912 | 0.505231655 | 0.247322606 | 2.04280419 | 0.041071826 | 0.232934829 |
| RSBN1L | 608.8320307 | 0.29481977 | 0.295987642 | 0.996054322 | 0.319223752 | 0.629115732 |
| RSC1A1 | 32.81692757 | 1.71834668 | 4.171710133 | 0.411904621 | 0.680409334 | 0.86207533 |
| RSF1 | 2569.826792 | 0.081890547 | 0.243654775 | 0.33609252 | 0.736801099 | 0.890536127 |

| | | | | | | |
|---|---|---|---|---|---|---|
| RSKR | 316.2050576 | -0.333712952 | 0.446737161 | -0.747000655 | 0.455063166 | 0.734911996 |
| RSL1D1 | 5327.920005 | 0.279321823 | 0.263909566 | 1.058399765 | 0.289873226 | 0.60258519 |
| RSL24D1 | 6235.871746 | 0.775494283 | 0.331434693 | 2.339810223 | 0.019293541 | 0.157609505 |
| RSPH1 | 18.51400853 | 0.294074529 | 0.690439704 | 0.425923548 | 0.670163555 | 0.857922099 |
| RSPH14 | 12.039488 | 1.510264123 | 0.855757133 | 1.764827969 | 0.077592683 | 0.319405602 |
| RSPH3 | 151.3564594 | -0.288327392 | 0.342428911 | -0.842006568 | 0.399784277 | 0.697622184 |
| RSPH4A | 13.55346222 | -0.612726683 | 0.834744015 | -0.734029441 | 0.462930795 | 0.740523312 |
| RSPH9 | 64.5262429 | 1.062292676 | 0.566424321 | 1.8754362 | 0.060732739 | 0.284105089 |
| RSPO1 | 24.38639035 | -1.22207444 | 1.011298627 | -1.208420943 | 0.226885388 | 0.535481989 |
| RSPO3 | 27.32134633 | 0.191603809 | 0.750627816 | 0.255258072 | 0.798523781 | 0.916056735 |
| RSPO4 | 51.17197198 | 0.231081347 | 0.480261827 | 0.481157016 | 0.630404908 | 0.838177265 |
| RSPRY1 | 938.8336909 | -0.247527135 | 0.285661361 | -0.866505481 | 0.386213022 | 0.686679504 |
| RSRC1 | 723.7506521 | 0.12524283 | 0.289151767 | 0.433138735 | 0.664913984 | 0.855040241 |
| RSRC2 | 3226.989221 | 0.097818652 | 0.253509382 | 0.385858113 | 0.699601754 | 0.87140051 |
| RSRP1 | 1061.592034 | 0.092770425 | 0.286419012 | 0.323897582 | 0.746015575 | 0.89396527 |
| RSU1 | 1043.077436 | -0.067795402 | 0.233216151 | -0.290697714 | 0.77128252 | 0.905067213 |
| RTBDN | 18.65790153 | 0.432048431 | 0.745636235 | 0.579435937 | 0.562295061 | 0.801706045 |
| RTCA | 1210.559875 | 0.211193028 | 0.385016552 | 0.548529739 | 0.583328215 | 0.8140624 |
| RTCB | 4337.124936 | 0.055293279 | 0.220970121 | 0.250229665 | 0.802409745 | 0.917774295 |
| RTEL1 | 340.7504389 | -0.142752722 | 0.566519582 | -0.251981973 | 0.801054998 | 0.916680548 |
| RTEL1-TNFRSF6B | 66.90360927 | -0.404048553 | 0.857043753 | -0.471444488 | 0.637323353 | 0.84164629 |
| RTF1 | 3824.431707 | 0.289492251 | 0.346499838 | 0.835475861 | 0.403449832 | 0.700152986 |
| RTF2 | 2021.556716 | -0.048522947 | 0.224073343 | -0.216549395 | 0.828559527 | 0.929120894 |
| RTKN | 895.9837834 | -0.049719869 | 0.288283505 | -0.172468657 | 0.8630691 | 0.944730333 |
| RTKN2 | 768.5449723 | 0.804992056 | 0.332519846 | 2.420884242 | 0.015482807 | 0.140797887 |
| RTL1 | 25.86634299 | 1.658723625 | 1.270193546 | 1.30588258 | 0.191592491 | 0.49715242 |
| RTL10 | 719.2082497 | -0.386538498 | 0.288027067 | -1.342021434 | 0.179589046 | 0.484443366 |
| RTL5 | 258.472222 | -1.089450941 | 0.626013535 | -1.740299339 | 0.08180647 | 0.327031873 |
| RTL6 | 1241.953868 | -0.051639375 | 0.400072918 | -0.129074907 | 0.897298377 | 0.959456082 |
| RTL8A | 664.330649 | 0.401390018 | 0.364314929 | 1.101766594 | 0.270563157 | 0.583257419 |
| RTL8B | 269.982992 | 0.079316322 | 0.385868837 | 0.205552548 | 0.837140447 | 0.933224905 |
| RTL8C | 1082.093315 | 0.35513348 | 0.412059758 | 0.861849461 | 0.388770363 | 0.688700978 |
| RTN1 | 64.97401253 | 1.173268223 | 0.463361893 | 2.532077501 | 0.011338893 | 0.120838052 |
| RTN2 | 442.4221937 | -0.805430716 | 0.404345915 | -1.991934839 | 0.046378208 | 0.247615641 |
| RTN3 | 6942.676194 | 0.15408658 | 0.295725004 | 0.521046843 | 0.60233414 | 0.825021776 |
| RTN4 | 5057.062007 | 0.188978405 | 0.242861458 | 0.778132546 | 0.436490878 | 0.720784523 |
| RTN4IP1 | 295.0514115 | 0.402030092 | 0.334069844 | 1.203431254 | 0.22880948 | 0.537506211 |
| RTN4R | 90.88833286 | 0.943975645 | 0.69510753 | 1.358028226 | 0.174454728 | 0.478578744 |
| RTN4RL1 | 35.92820598 | 1.564727428 | 0.689711825 | 2.268668408 | 0.023288498 | 0.174130017 |
| RTN4RL2 | 136.4778773 | 0.488818298 | 0.473560955 | 1.032218331 | 0.30196985 | 0.612867503 |
| RTRAF | 5947.981831 | 0.439592244 | 0.318427112 | 1.380511355 | 0.167429256 | 0.469714288 |
| RTTN | 862.9208542 | -0.191510923 | 0.417304198 | -0.458924027 | 0.646288724 | 0.846120662 |
| RUBCN | 544.0157121 | -0.270525463 | 0.310747832 | -0.870562673 | 0.383992985 | |

0.684699412
RUBCNL 108.2683491 -1.376110627 0.640277611 -2.149240584 0.031615334 0.202592072
RUFY1 555.7867325 -0.33863882 0.320868819 -1.05538089 0.291251159 0.604290391
RUFY2 471.7968015 -0.088245322 0.283822648 -0.310917126 0.755863625 0.898050517
RUFY3 1558.61186 -0.238774423 0.243476767 -0.980686682 0.326747273 0.636502681
RUNDC1 507.9468381 -0.218057218 0.301069612 -0.724275082 0.468896872 0.744337782
RUNDC3A 25.25889884 0.184139968 0.642922276 0.286410932 0.774563403 0.90635568
RUNX1 48.52229253 -1.702152153 1.29185701 -1.317601049 0.187637234 0.492618714
RUNX1T1 248.7774993 -4.583926894 0.815483504 -5.621115412 1.90E-08 1.25E-05
RUSC1 1280.373819 0.069099673 0.272701606 0.253389314 0.799967385 0.916311991
RUSC2 246.4599963 0.29717682 0.525887171 0.565096159 0.572008361 0.807351345
RUVBL1 1078.076915 0.370647716 0.27837583 1.331465146 0.183036006 0.486993905
RUVBL2 2609.044559 0.659897688 0.320225682 2.060726935 0.039329098 0.228518481
RWDD1 1475.827347 1.026365849 0.439341733 2.33614467 0.019483703 0.158237289
RWDD2A 298.3675166 0.120475957 0.326794872 0.368659264 0.712381716 0.877520543
RWDD2B 216.9264435 -0.394060865 0.374814528 -1.051348963 0.293098347 0.606216091
RWDD3 408.0540564 0.667185537 0.463725892 1.438749804 0.150221423 0.444771508
RWDD4 881.7427321 0.622376902 0.28227622 2.20485063 0.027464578 0.189565564
RXRA 2101.28032 -0.646389829 0.336149128 -1.922925793 0.05448937 0.269872826
RXYLT1 277.9625443 0.58768022 0.323791054 1.81499832 0.069524168 0.303425952
RYBP 1803.28835 0.110285459 0.287325825 0.383834132 0.701101389 0.872239782
RYK 1919.612344 0.346417507 0.223456744 1.55026651 0.121077562 0.401219509
RYR1 693.3698999 -1.166444965 0.357936027 -3.258808495 0.001118812 0.028974947
RYR2 42.58057615 -2.091975988 1.086568658 -1.925304924 0.054191222 0.269111302
RYR3 42.00243086 -0.267896816 0.582547192 -0.459871439 0.645608502 0.845794147
S100A1 26.09350976 0.856906738 0.870856573 0.983981478 0.32512462 0.635057014
S100A10 3319.441586 0.522199035 0.643816725 0.811098896 0.41730888 0.708794572
S100A11 3038.810279 -0.080608208 0.693976471 -0.116154094 0.907530418 0.965304913
S100A13 1607.197053 0.449668775 0.423184512 1.062583252 0.287970985 0.601673803
S100A14 251.3231491 -1.928247787 0.975806958 -1.976054558 0.048148598 0.25310547
S100A16 1774.323469 -1.539927185 0.723803331 -2.127549182 0.033374479 0.20836388
S100A4 568.6827488 1.646723894 0.515409109 3.194984073 0.001398386 0.033855429
S100A6 422.9723342 1.125169135 0.516296015 2.179310128 0.029308637 0.195861365
S100B 24.24326702 1.559568105 0.893346967 1.74575855 0.080852916 0.325332894
S100PBP 1066.867538 0.046589344 0.292223288 0.159430634 0.873329605 0.949633274
S1PR1 24.09507146 2.080920863 0.87600256 2.375473496 0.017526451 0.149594125
S1PR2 295.9768156 0.483657124 0.455160955 1.062606796 0.287960304 0.601673803
S1PR3 174.2312272 0.485277886 0.475478669 1.020609162 0.307439647 0.617319594
SAAL1 1045.336177 0.426967153 0.239641418 1.781691816 0.074799502 0.313842348

| | | | | | | |
|---|---|---|---|---|---|---|
| SAC3D1 | 194.214606 | 1.129996025 | 0.496103123 | 2.27774423 | 0.022741821 | 0.171820799 |
| SACM1L | 1228.507266 | -0.015699372 | 0.292411919 | -0.053689233 | 0.957182762 | 0.984903668 |
| SACS | 2289.837968 | 0.880661354 | 0.560006183 | 1.572592196 | 0.115813276 | 0.392188216 |
| SAE1 | 7206.291417 | 0.098067041 | 0.233712357 | 0.419605718 | 0.67477351 | 0.859745438 |
| SAFB | 2656.2742 | -0.442445972 | 0.408137714 | -1.084060495 | 0.278337977 | 0.592005173 |
| SAFB2 | 1076.601395 | -0.250621647 | 0.330307974 | -0.758751427 | 0.448001267 | 0.729463615 |
| SALL1 | 4108.182341 | 0.319933259 | 0.455090291 | 0.703010513 | 0.482049198 | 0.753236025 |
| SALL2 | 4953.668143 | 0.10655311 | 0.419477684 | 0.254013775 | 0.799484916 | 0.91629415 |
| SALL3 | 189.6754025 | 0.263642004 | 0.323813857 | 0.814177645 | 0.41554319 | 0.707542731 |
| SALL4 | 4193.557448 | -0.083416791 | 0.265069215 | -0.314698149 | 0.75299085 | 0.896586461 |
| SAMD1 | 1332.320475 | 0.788035941 | 0.407706139 | 1.932852774 | 0.053254337 | 0.267056023 |
| SAMD10 | 77.05796112 | -0.07845069 | 0.521179992 | -0.150525138 | 0.88035032 | 0.952483146 |
| SAMD11 | 217.3324743 | -1.844972812 | 0.824987962 | -2.236363312 | 0.025327982 | 0.181850837 |
| SAMD12 | 313.7573385 | -0.670271934 | 0.32656708 | -2.05247857 | 0.040123176 | 0.231096696 |
| SAMD13 | 76.52779211 | 1.227422132 | 0.432176059 | 2.840097472 | 0.004509975 | 0.070434393 |
| SAMD14 | 385.1981759 | -0.850863165 | 0.332255952 | -2.560866585 | 0.010441144 | 0.116176818 |
| SAMD15 | 57.08997953 | -1.170913535 | 0.513648216 | -2.279602067 | 0.0226313 | 0.171427007 |
| SAMD4A | 100.6286336 | -0.892165997 | 0.744602916 | -1.198176877 | 0.230848165 | 0.539623726 |
| SAMD4B | 2101.073679 | -0.3787608 | 0.295036754 | -1.28377497 | 0.199220704 | 0.505071561 |
| SAMD8 | 1022.516733 | -0.090834974 | 0.31346968 | -0.28977276 | 0.771990089 | 0.905547141 |
| SAMHD1 | 217.0331594 | -0.485417206 | 0.3796118 | -1.278720012 | 0.20099567 | 0.506881795 |
| SAMM50 | 1909.956855 | -0.04814353 | 0.269179712 | -0.178852743 | 0.85805333 | 0.942343063 |
| SAP130 | 1997.871246 | -0.501069126 | 0.283974708 | -1.76448505 | 0.07765035 | 0.31942339 |
| SAP18 | 4305.253982 | 0.602392261 | 0.369112903 | 1.632000008 | 0.102679485 | 0.367636293 |
| SAP25 | 57.2663154 | 0.249882515 | 0.719773968 | 0.347168037 | 0.728465082 | 0.885682302 |
| SAP30 | 357.0666529 | 1.422029442 | 0.378963797 | 3.752415013 | 0.000175139 | 0.008728094 |
| SAP30BP | 1875.286473 | -0.006878357 | 0.278623956 | -0.024686883 | 0.980304718 | 0.993201742 |
| SAP30L | 666.7640613 | -0.253697981 | 0.282021886 | -0.899568414 | 0.368349972 | 0.674500595 |
| SAPCD2 | 323.4208502 | 0.656196344 | 0.368078832 | 1.782760342 | 0.074625324 | 0.313779586 |
| SAR1A | 3490.263268 | 0.264371243 | 0.346681659 | 0.762576377 | 0.445716078 | 0.728439722 |
| SAR1B | 627.4279445 | 0.20588167 | 0.255719886 | 0.805106216 | 0.420758389 | 0.711245864 |
| SARAF | 3362.922271 | 0.281220974 | 0.307337141 | 0.91502437 | 0.360178837 | 0.667522123 |
| SARDH | 154.7505289 | 0.921036828 | 0.451015975 | 2.042137928 | 0.041137852 | 0.232946328 |
| SARM1 | 65.33211212 | 0.220394581 | 0.610231422 | 0.361165573 | 0.717975677 | 0.880671367 |
| SARNP | 3011.592036 | 0.898386971 | 0.360215165 | 2.494028732 | 0.012630232 | 0.127233234 |
| SARS | 3356.641673 | -0.113798729 | 0.232179359 | -0.49013284 | 0.624039901 | 0.834707114 |
| SARS2 | 492.3509617 | -0.158463706 | 0.423835115 | -0.373880551 | 0.708493187 | 0.875585221 |
| SART1 | 1592.256447 | -0.361027864 | 0.272679074 | -1.324002823 | 0.185502107 | 0.490031772 |
| SART3 | 2295.448013 | -0.139064754 | 0.346086489 | -0.401820811 | 0.687815905 | 0.866689498 |
| SASH1 | 1598.226752 | 0.59179874 | 0.474269186 | 1.247811914 | 0.212099944 | 0.519329932 |

| | | | | | | |
|---|---|---|---|---|---|---|
| SASS6 | 556.0259668 | 0.12041304 | 0.269400283 | 0.446967015 | 0.654898876 | 0.849258129 |
| SAT1 | 8515.229033 | 1.270766467 | 0.765080156 | 1.660958602 | 0.096721761 | 0.357177519 |
| SAT2 | 1298.772073 | 0.929312735 | 0.429677871 | 2.162812653 | 0.030555592 | 0.199123233 |
| SATB1 | 1709.018594 | -1.365099214 | 0.412268308 | -3.311191252 | 0.000928997 | 0.02597297 |
| SATB2 | 127.3271344 | 0.707262475 | 0.507060658 | 1.39482814 | 0.163067671 | 0.464494862 |
| SATL1 | 23.91535749 | 0.139967478 | 0.688714107 | 0.203230159 | 0.838955141 | 0.933915137 |
| SAV1 | 1282.061271 | 0.040664491 | 0.288363294 | 0.141018264 | 0.887855514 | 0.955943827 |
| SAXO2 | 22.70147242 | 1.655880784 | 0.745385014 | 2.2215107 | 0.026316391 | 0.185350959 |
| SAYSD1 | 286.3277538 | -0.106896617 | 0.462919854 | -0.230918194 | 0.817378356 | 0.923820833 |
| SBDS | 1009.17725 | 0.565832068 | 0.337681066 | 1.675640495 | 0.093808632 | 0.351226741 |
| SBDSP1 | 129.320752 | -0.239709964 | 0.397982878 | -0.602312255 | 0.546966305 | 0.792249935 |
| SBF1 | 1870.494234 | 0.059600846 | 0.574560182 | 0.10373299 | 0.917381246 | 0.969661544 |
| SBF2 | 776.4186929 | -0.224088722 | 0.373807473 | -0.599476302 | 0.548855308 | 0.793468239 |
| SBK1 | 2049.109804 | -0.054803985 | 0.258372723 | -0.212112115 | 0.832019569 | 0.931142584 |
| SBK2 | 68.30179217 | -2.334518083 | 0.74420702 | -3.136920264 | 0.001707326 | 0.03837309 |
| SBK3 | 69.6173966 | -1.818478545 | 0.730613828 | -2.488973622 | 0.012811247 | 0.128036515 |
| SBNO1 | 1732.067212 | 0.091334873 | 0.26574326 | 0.343695913 | 0.731074985 | 0.886936313 |
| SBNO2 | 61.2108442 | 0.793459211 | 1.03812808 | 0.764317261 | 0.444678202 | 0.727526247 |
| SBSPON | 11.08774616 | -0.119064598 | 1.065507643 | -0.111744481 | 0.91102601 | 0.966866396 |
| SC5D | 2547.809889 | 0.411200893 | 0.305513106 | 1.345935364 | 0.178323363 | 0.4825693 |
| SCAF1 | 1143.786156 | -0.301093945 | 0.434446136 | -0.693052419 | 0.488276656 | 0.756963535 |
| SCAF11 | 3124.593263 | -0.486771662 | 0.360454271 | -1.350439434 | 0.176875069 | 0.481149663 |
| SCAF4 | 1722.229158 | -0.26205784 | 0.324317121 | -0.80802962 | 0.419073531 | 0.709924862 |
| SCAF8 | 2002.2458 | -0.535495813 | 0.288269632 | -1.857621317 | 0.063222809 | 0.289778852 |
| SCAI | 730.3006567 | -0.334802392 | 0.331673441 | -1.00943383 | 0.312766621 | 0.622907489 |
| SCAMP1 | 1857.222924 | 0.175653305 | 0.293691571 | 0.598087662 | 0.549781442 | 0.793878561 |
| SCAMP2 | 1649.959156 | -0.349598965 | 0.289047795 | -1.209484977 | 0.226476579 | 0.535186874 |
| SCAMP3 | 131.1134221 | 0.569407045 | 0.341018406 | 1.669725256 | 0.094973734 | 0.353777379 |
| SCAMP4 | 1153.816007 | -0.630094184 | 0.269781346 | -2.33557358 | 0.019513477 | 0.158309603 |
| SCAMP5 | 1032.887442 | -0.308014525 | 0.256708979 | -1.199858791 | 0.230194187 | 0.539041344 |
| SCAND1 | 294.3175816 | 0.649598107 | 0.73734401 | 0.880997334 | 0.378319263 | 0.681253679 |
| SCAND2P | 103.3530907 | -0.294483587 | 0.485043466 | -0.607128243 | 0.543765808 | 0.790216482 |
| SCAP | 1891.58951 | 0.102236862 | 0.262617659 | 0.389299264 | 0.69705478 | 0.870026319 |
| SCAPER | 320.5687569 | -0.012854217 | 0.305330014 | -0.042099422 | 0.966419441 | 0.988700292 |
| SCARA3 | 691.9338382 | -1.096895281 | 0.312902884 | -3.505545453 | 0.000455673 | 0.016498831 |
| SCARB1 | 376.3450836 | 0.591923973 | 0.488101165 | 1.21270756 | 0.225241649 | 0.53459368 |
| SCARB2 | 3574.861455 | -0.307362721 | 0.261227341 | -1.176610077 | 0.239351174 | 0.550578888 |
| SCARF2 | 16.3279384 | 0.172742305 | 0.771976202 | 0.22376636 | 0.822939107 | 0.926014081 |
| SCART1 | 14.82582475 | -0.357378946 | 0.783291721 | -0.456252679 | 0.648208288 | 0.846656069 |

| | | | | | | |
|---|---|---|---|---|---|---|
| SCCPDH | 437.2975974 | 1.634696015 | 0.338300447 | 4.832083518 | 1.35E-06 | 0.000226869 |
| SCD | 13251.95575 | -0.122692601 | 0.299551601 | -0.409587532 | 0.682108544 | 0.862735748 |
| SCD5 | 79.241846 | 1.593050519 | 0.580028381 | 2.746504432 | 0.006023407 | 0.083980132 |
| SCFD1 | 1860.993241 | 0.196660265 | 0.230149444 | 0.854489424 | 0.392833867 | 0.692011554 |
| SCFD2 | 189.6558299 | 0.080400404 | 0.344670524 | 0.233267419 | 0.815553756 | 0.922975941 |
| SCG3 | 311.7376419 | -1.363185508 | 0.541296488 | -2.518371241 | 0.011789898 | 0.123184948 |
| SCG5 | 588.343069 | 0.868302536 | 0.423822908 | 2.048739035 | 0.04048764 | 0.231601821 |
| SCGB3A2 | 513.1707632 | 1.001539947 | 0.561281299 | 1.784381465 | 0.074361701 | 0.313285577 |
| SCHIP1 | 174.6740465 | 0.169832513 | 0.710748066 | 0.238948963 | 0.811145158 | 0.920897407 |
| SCIN | 27.17343483 | -0.979142106 | 0.814225909 | -1.202543539 | 0.229153008 | 0.537806762 |
| SCLT1 | 426.0913411 | 0.458785474 | 0.334107991 | 1.373165222 | 0.169700989 | 0.472403161 |
| SCLY | 683.7890061 | 0.58282901 | 0.368917171 | 1.579837037 | 0.114144191 | 0.388887104 |
| SCMH1 | 380.0560092 | -0.098150086 | 0.473475214 | -0.2072972 | 0.835777761 | 0.932596676 |
| SCML1 | 374.9790676 | 1.159653264 | 0.476592163 | 2.433219332 | 0.014965233 | 0.139113039 |
| SCML2 | 602.302782 | 0.349383636 | 0.383100192 | 0.911990241 | 0.361773855 | 0.668713609 |
| SCML4 | 19.48467552 | -1.341866115 | 0.882147914 | -1.521135054 | 0.128225949 | 0.412317905 |
| SCN11A | 19.47058459 | -0.45449659 | 0.71206 | -0.638284119 | 0.523288748 | 0.777662198 |
| SCN1B | 120.5677302 | -0.298469086 | 0.533614002 | -0.559335183 | 0.575932988 | 0.808969969 |
| SCN2A | 17.92390351 | -0.440909268 | 0.778306434 | -0.566498295 | 0.571055091 | 0.806805438 |
| SCN2B | 13.33874434 | 1.143925352 | 1.02725928 | 1.113570229 | 0.265463604 | 0.579474581 |
| SCN4A | 233.8072923 | 0.054041834 | 0.535750426 | 0.100871286 | 0.919652637 | 0.969700456 |
| SCN4B | 307.9922509 | 1.359530168 | 0.679584239 | 2.000532223 | 0.045442824 | 0.245452751 |
| SCN5A | 110.2489076 | -0.077721053 | 0.619888545 | -0.125379077 | 0.900223452 | 0.96137477 |
| SCN7A | 24.56075198 | -0.784999438 | 0.649586695 | -1.208459847 | 0.226870431 | 0.535481989 |
| SCN8A | 382.3224523 | -0.292617654 | 0.628497884 | -0.46558256 | 0.641514329 | 0.843948481 |
| SCN9A | 117.4537425 | -2.28548197 | 0.462686135 | -4.939594683 | 7.83E-07 | 0.000158355 |
| SCNM1 | 759.3161057 | -0.016420638 | 0.388341646 | -0.042283999 | 0.966272301 | 0.988700292 |
| SCNN1A | 1563.941377 | -1.153424142 | 0.555086854 | -2.077916517 | 0.037717049 | 0.2229082 |
| SCNN1D | 75.11147528 | 0.092043804 | 0.683431938 | 0.134678816 | 0.892865823 | 0.957604542 |
| SCNN1G | 29.70866288 | -0.836000658 | 0.989003728 | -0.84529576 | 0.397945728 | 0.696517252 |
| SCO1 | 1451.37676 | 0.232437027 | 0.241456789 | 0.962644408 | 0.335726004 | 0.644086928 |
| SCO2 | 198.9024128 | 0.521109094 | 0.468842053 | 1.11148113 | 0.266361308 | 0.57970088 |
| SCOC | 2652.962047 | 0.815500377 | 0.288570954 | 2.825996051 | 0.004713385 | 0.072780519 |
| SCP2 | 1641.710343 | 0.046339789 | 0.272060016 | 0.17032929 | 0.864751178 | 0.945248223 |
| SCPEP1 | 1197.940883 | 0.073394122 | 0.234750478 | 0.312647378 | 0.754548579 | 0.897438517 |
| SCRG1 | 10.09352346 | -1.102064495 | 1.103501953 | -0.998697367 | 0.317941317 | 0.628058296 |
| SCRIB | 498.0836891 | 0.161447678 | 0.71738153 | 0.225051345 | 0.821939331 | 0.92563193 |
| SCRN1 | 3417.877859 | -0.606335841 | 0.32239149 | -1.880743937 | 0.06000676 | 0.28333724 |
| SCRN2 | 980.8560007 | 0.138352831 | 0.400465723 | 0.345479833 | 0.729733669 | 0.886372257 |
| SCRN3 | 538.5322823 | -0.240578168 | 0.285815835 | -0.841724421 | 0.399942225 | 0.697652919 |
| SCUBE1 | 19.92544234 | 0.764513804 | 0.751962012 | 1.016692057 | 0.309299945 | |

0.61913267

| | | | | | | |
|---|---|---|---|---|---|---|
| SCUBE2 | 41.94058896 | 0.066872155 | 0.532303546 | 0.125627859 | 0.900026509 | 0.961249486 |
| SCUBE3 | 1977.589509 | -1.627306293 | 0.538796684 | -3.020260411 | 0.002525574 | 0.048623717 |
| SCYL1 | 1167.141377 | -0.458771771 | 0.384235737 | -1.193985168 | 0.23248377 | 0.542005591 |
| SCYL2 | 2100.853643 | -0.570360709 | 0.307434326 | -1.855227804 | 0.063563703 | 0.290809691 |
| SCYL3 | 672.5329992 | -0.484422249 | 0.361203278 | -1.3411347 | 0.179876723 | 0.484622582 |
| SDAD1 | 1516.598834 | -0.031495641 | 0.230441142 | -0.136675425 | 0.891287356 | 0.957017706 |
| SDAD1P1 | 51.54190766 | -0.775961607 | 0.475526812 | -1.631793597 | 0.102722975 | 0.367636293 |
| SDC1 | 803.1759004 | 1.040163245 | 0.416691588 | 2.496242484 | 0.012551677 | 0.12669427 |
| SDC2 | 2352.667728 | 0.753237669 | 0.295600386 | 2.548161993 | 0.010829217 | 0.118235065 |
| SDC3 | 1625.931992 | -0.461662657 | 0.564813934 | -0.817371225 | 0.413716314 | 0.705778726 |
| SDC4 | 3949.90209 | -0.604938116 | 0.645625083 | -0.936980504 | 0.348768587 | 0.655415363 |
| SDCBP | 3959.799367 | 0.331965999 | 0.342146316 | 0.970245722 | 0.331924025 | 0.639768693 |
| SDCBP2 | 33.22658514 | 0.866245569 | 0.619267404 | 1.39882313 | 0.161866028 | 0.462461302 |
| SDCCAG8 | 105.0577656 | -0.056468922 | 0.673601673 | -0.083831327 | 0.93319054 | 0.97462713 |
| SDE2 | 527.0918513 | 0.288803727 | 0.256845118 | 1.12442755 | 0.26083169 | 0.57415519 |
| SDF2 | 897.2079214 | 0.209287251 | 0.417928565 | 0.500772783 | 0.616531042 | 0.831694047 |
| SDF2L1 | 202.8251333 | 0.667907841 | 0.565907566 | 1.180241935 | 0.237904005 | 0.54863845 |
| SDF4 | 1466.070137 | -0.158740527 | 0.336963351 | -0.47109137 | 0.637575487 | 0.841782104 |
| SDHA | 1694.137884 | 0.250246459 | 0.232672057 | 1.075532929 | 0.282136198 | 0.595962806 |
| SDHAF1 | 177.9813819 | 1.08290952 | 0.470096138 | 2.303591609 | 0.021245579 | 0.165290607 |
| SDHAF2 | 521.5775286 | 0.481016657 | 0.436515352 | 1.101946712 | 0.270484839 | 0.583257419 |
| SDHAF3 | 513.3516516 | 0.93021905 | 0.357109945 | 2.604853385 | 0.009191358 | 0.108599763 |
| SDHAF4 | 593.147972 | 1.391742349 | 0.428032762 | 3.251485566 | 0.001148036 | 0.029521884 |
| SDHAP1 | 355.2046864 | 0.084754928 | 0.288866452 | 0.293405232 | 0.769212427 | 0.904597192 |
| SDHAP2 | 32.81721876 | -0.334840571 | 0.76053094 | -0.440272122 | 0.65974003 | 0.852112718 |
| SDHAP3 | 263.4025621 | -0.572296581 | 0.346859854 | -1.649936061 | 0.098956014 | 0.36119708 |
| SDHB | 1093.839125 | 0.991212144 | 0.370732814 | 2.673656357 | 0.007502929 | 0.096381827 |
| SDHC | 3883.173412 | 0.245713945 | 0.30516126 | 0.805193768 | 0.420707872 | 0.711245864 |
| SDHD | 1851.404738 | 0.720404984 | 0.333037594 | 2.16313412 | 0.030530866 | 0.199123233 |
| SDK1 | 262.3861483 | -1.331350273 | 0.607057179 | -2.193121703 | 0.028298612 | 0.192356209 |
| SDK2 | 1841.219858 | 0.702173742 | 0.570877561 | 1.229990089 | 0.218700816 | 0.527143548 |
| SDR39U1 | 589.6250409 | 0.506360877 | 0.31785052 | 1.593078649 | 0.111142548 | 0.383475916 |
| SDR42E1 | 64.9007315 | -3.749038351 | 0.655906797 | -5.715809576 | 1.09E-08 | 9.74E-06 |
| SDSL | 173.6741356 | 0.390189845 | 0.50491983 | 0.772775838 | 0.439655056 | 0.723249422 |
| SEC11A | 7550.034037 | 0.573362078 | 0.314256611 | 1.824502835 | 0.068076088 | 0.300140172 |
| SEC11C | 557.9004659 | 1.006495109 | 0.348672527 | 2.886648735 | 0.003893687 | 0.064617764 |
| SEC13 | 2478.60963 | 0.030149815 | 0.253771418 | 0.118806977 | 0.905428281 | 0.964473572 |
| SEC14L1 | 2754.082171 | -0.189723011 | 0.296241896 | -0.640432746 | 0.5218913 | |

0.776922079
SEC14L1P1 22.03870841 -1.709101618 0.775230191 -2.204637587 0.027479536 0.189582556
SEC14L2 35.66221272 0.600541513 0.659464791 0.91064985 0.362479896 0.669662332
SEC14L4 15.49449624 -0.204562812 0.816206324 -0.25062635 0.802103007 0.917532337
SEC14L5 47.10407414 1.288191388 0.75872459 1.69783793 0.089538355 0.342777615
SEC14L6 90.38510466 -0.617919693 0.388754975 -1.589483693 0.111951233 0.384920166
SEC16A 1869.980725 -0.479222502 0.358513794 -1.336691948 0.181323211 0.485749663
SEC16B 84.87377719 0.477861852 0.543267756 0.879606505 0.379072514 0.681253679
SEC22A 384.7494586 0.323939374 0.268860466 1.204860569 0.228257133 0.537150955
SEC22B 1398.539922 -0.096523934 0.224476345 -0.429996014 0.66719854 0.856421651
SEC22B4P 23.04255282 0.292247907 0.665867749 0.438897826 0.660735572 0.852672623
SEC22C 750.5666952 -0.023029154 0.272200119 -0.084603762 0.932576408 0.97453841
SEC23A 1694.225943 0.058516866 0.285791391 0.204753776 0.837764503 0.933603058
SEC23B 1527.500722 -0.066682311 0.276831656 -0.240876755 0.809650638 0.920090624
SEC23IP 1698.438712 -0.182186701 0.267899263 -0.680056744 0.496468532 0.761184851
SEC24A 1014.338793 -0.480381928 0.297006522 -1.617412052 0.105789366 0.372892767
SEC24B 1380.915696 -0.354058719 0.259551344 -1.364118226 0.172530359 0.476247831
SEC24C 3159.552617 -0.466075205 0.288077727 -1.617880045 0.105688452 0.372686009
SEC24D 794.5185438 -0.185442529 0.320171368 -0.579197727 0.562455764 0.801706045
SEC31A 4033.225175 -0.471785128 0.310798135 -1.517979273 0.129019624 0.413453061
SEC31B 374.8013347 -0.12260031 0.527944922 -0.232221782 0.816365761 0.923501973
SEC61A1 4728.011887 -0.469326585 0.306455406 -1.531467794 0.12565382 0.407763445
SEC61A2 574.2989106 0.226495476 0.318871914 0.710302369 0.477516651 0.750016059
SEC61B 2083.411789 1.073984333 0.501029192 2.143556403 0.032068447 0.204244505
SEC61G 1562.620499 1.364336219 0.486031644 2.807093399 0.004999075 0.07561376
SEC62 2859.647579 0.467647981 0.29376497 1.591912003 0.111404478 0.383943057
SEC63 2552.080809 -0.144168169 0.273707733 -0.526723041 0.598385933 0.823112973
SECISBP2 2248.736037 0.220787212 0.276411355 0.798763177 0.424427746 0.713303793
SECISBP2L 1461.194499 -0.629025513 0.431014099 -1.459408208 0.144452788 0.435771178
SEH1L 1183.200886 0.369505804 0.280632211 1.316690636 0.187942344 0.492703179
SEL1L 1345.755004 -0.171229023 0.300214663 -0.570355296 0.568436743 0.805450586
SEL1L3 1741.63732 0.201783363 0.502279206 0.40173545 0.687878732 0.866689498
SELENBP1 1487.94129 0.014447537 0.247809506 0.05830098 0.953508887 0.983324269
SELENOF 3424.249566 0.32607585 0.267230724 1.220203444 0.222387762

0.531241476
SELENOH 2241.925532 1.268230685 0.51589083 2.458331513 0.013958426
0.134282356
SELENOI 1507.013272 0.012062287 0.379680688 0.031769556 0.974655825
0.991261106
SELENOK 1064.814953 0.79767445 0.395971921 2.014472259 0.043959975
0.241112355
SELENOM 97.72274991 0.856782408 0.710186082 1.206419599 0.227655743
0.536382245
SELENON 3374.650876 -0.553969259 0.353237529 -1.56826275 0.116819825
0.394101306
SELENOO 255.4555368 0.100994503 0.427314903 0.236346784 0.813163587
0.921802034
SELENOP 3404.076558 -0.674109262 0.618115349 -1.09058813 0.275454155
0.588496689
SELENOS 754.3796183 0.375053482 0.416624342 0.9002198 0.368003291
0.674179083
SELENOT 1479.600595 0.447455326 0.291852696 1.533154676 0.125237749
0.407109896
SELENOV 78.36880487 0.077996478 0.766962088 0.10169535 0.918998493
0.969677217
SELENOW 4724.823007 0.217103775 0.485073604 0.447568726 0.654464478 0.8491303
SELL 13.77705572 -3.750521909 1.445807168 -2.594067862 0.009484776
0.110347809
SEM1 4582.349872 1.098122215 0.507496295 2.163803413 0.03047944 0.199054492
SEMA3A 674.1235673 -0.879077494 0.252521968 -3.481192152
0.000499187 0.017561181
SEMA3B 424.1563949 0.224141282 0.458637605 0.488711087 0.625046254
0.835033444
SEMA3C 1157.55102 0.593918798 0.542393973 1.094995202 0.273518738
0.586675071
SEMA3D 1175.480116 -2.343546446 0.386572736 -6.062368675 1.34E-09
1.57E-06
SEMA3E 1160.903974 0.577542901 0.396673912 1.455963914 0.145402596
0.437158696
SEMA3F 1202.035751 -0.336807903 0.381603238 -0.882612802
0.377445509 0.680802024
SEMA3G 61.38600543 0.686890174 0.602711641 1.139666345 0.254425333
0.566963386
SEMA4A 90.65053854 -0.977917204 0.465883856 -2.099057934
0.035811794 0.216454473
SEMA4B 623.2515116 -0.291186068 0.353328473 -0.82412285 0.409869758
0.703966274
SEMA4C 2690.655127 -0.341225642 0.405676733 -0.841126972
0.400276806 0.697919713
SEMA4D 974.446759 -0.256497388 0.297712757 -0.861559952
0.388929717 0.688870316
SEMA4F 269.7558042 0.417418591 0.399720612 1.044275873 0.296357772
0.607655597
SEMA4G 349.6209034 -1.318314942 0.385438821 -3.420296223
0.00062553 0.020321012
SEMA5A 317.4296839 1.162754541 0.590873732 1.967856206 0.049084585
0.255021315
SEMA5B 565.6976777 0.111422047 0.528524866 0.210817039 0.833030042
0.931554473
SEMA6A 4397.510574 -1.428580821 0.412548898 -3.462815749
0.000534554 0.018183236
SEMA6B 343.7861017 1.093472283 0.424276726 2.577262001 0.009958644 0.1136814
SEMA6C 417.1216529 -0.234320188 0.476256348 -0.492004336
0.622716278 0.833787151
SEMA6D 549.0913047 -0.232691622 0.577223308 -0.40312236 0.686858212
0.865835142
SEMA7A 513.5298064 0.454581875 0.355132482 1.280034631 0.200532957
0.506625393

| | | | | | | |
|---|---|---|---|---|---|---|
| SENP1 | 937.0165198 | -0.450836968 | 0.238234619 | -1.892407453 | 0.058436714 | 0.280248566 |
| SENP2 | 1201.026606 | -0.30158307 | 0.241012923 | -1.251314932 | 0.2108196 | 0.518294063 |
| SENP3 | 1765.281921 | -0.020290195 | 0.313926293 | -0.064633626 | 0.948465711 | 0.98134156 |
| SENP3-EIF4A1 | 53.14863618 | -0.031224007 | 0.789068739 | -0.039570706 | 0.968435382 | 0.989435867 |
| SENP5 | 995.5402623 | 0.055328778 | 0.248596165 | 0.22256489 | 0.823874165 | 0.926469124 |
| SENP6 | 2939.463955 | -0.147574225 | 0.257035198 | -0.57414014 | 0.565872972 | 0.804126907 |
| SENP7 | 620.16627 | -0.418389892 | 0.38674062 | -1.081835913 | 0.279325446 | 0.593009563 |
| SENP8 | 189.7713333 | 0.280130334 | 0.366594984 | 0.764141208 | 0.444783098 | 0.727526247 |
| SEPHS1 | 2247.267908 | 0.098010491 | 0.25141734 | 0.389831868 | 0.696660877 | 0.870023227 |
| SEPHS2 | 1655.255974 | 0.063992277 | 0.398514536 | 0.160577021 | 0.872426554 | 0.949195586 |
| SEPSECS | 362.2598708 | 0.208009069 | 0.302149785 | 0.688430307 | 0.491181844 | 0.758599181 |
| SEPT1 | 38.15909806 | 0.116486295 | 0.631978923 | 0.184319905 | 0.8537625 | 0.941105201 |
| SEPT10 | 2505.096972 | 0.15825033 | 0.243956948 | 0.648681381 | 0.516544342 | 0.773950426 |
| SEPT11 | 4369.733013 | -0.305903583 | 0.262155567 | -1.166878075 | 0.243259611 | 0.554511158 |
| SEPT14 | 115.3615 | 0.151915889 | 0.373379433 | 0.406867321 | 0.684105439 | 0.863826466 |
| SEPT14P19 | 11.13380684 | 0.542312674 | 1.049791213 | 0.516590982 | 0.605441712 | 0.826077544 |
| SEPT2 | 13021.55998 | 0.053389757 | 0.220801126 | 0.241800203 | 0.808934982 | 0.919641737 |
| SEPT3 | 441.60992 | 0.675122629 | 0.604147505 | 1.117479792 | 0.263789235 | 0.577444856 |
| SEPT4 | 186.9759648 | 0.282622334 | 0.461589656 | 0.612280475 | 0.540352205 | 0.787481586 |
| SEPT5 | 64.58242792 | 1.094803761 | 0.557715813 | 1.963013663 | 0.049644586 | 0.256787597 |
| SEPT6 | 1903.726312 | -0.910832684 | 0.238370593 | -3.821078228 | 0.000132869 | 0.007303489 |
| SEPT7 | 5713.020338 | 0.103130126 | 0.261416849 | 0.394504509 | 0.693208599 | 0.868643566 |
| SEPT7P2 | 430.0756911 | -0.037230999 | 0.265012615 | -0.140487648 | 0.888274711 | 0.955943827 |
| SEPT8 | 2035.702422 | -0.904017346 | 0.388107035 | -2.329299047 | 0.019843229 | 0.159450013 |
| SEPT9 | 10959.34727 | -0.438961556 | 0.330039323 | -1.330028048 | 0.18350903 | 0.487824688 |
| SERAC1 | 551.2304887 | -0.187838656 | 0.333063936 | -0.563971765 | 0.572773348 | 0.807670159 |
| SERBP1 | 18587.83839 | 0.219460326 | 0.2416277 | 0.908258143 | 0.363741852 | 0.670353372 |
| SERF1A | 59.6443292 | 1.677982646 | 0.849912658 | 1.9743001 | 0.048347629 | 0.253485504 |
| SERF1B | 51.74716083 | 0.857421142 | 0.609096808 | 1.407692718 | 0.159222079 | 0.459057043 |
| SERF2 | 11234.97683 | 0.958512033 | 0.499594469 | 1.918580154 | 0.055037489 | 0.271793713 |
| SERGEF | 317.2055958 | 0.176741629 | 0.297149809 | 0.594789643 | 0.551984091 | 0.795417045 |
| SERHL | 85.90678968 | -1.161337873 | 0.606703631 | -1.914176564 | 0.055597599 | 0.272730921 |
| SERHL2 | 92.30572414 | 0.00474922 | 0.552518306 | 0.008595589 | 0.993141796 | 0.997283174 |
| SERINC1 | 3253.095806 | -0.539946844 | 0.319197151 | -1.691577892 | 0.090726485 | 0.345245839 |
| SERINC2 | 532.342699 | -0.576028288 | 0.322757742 | -1.784707886 | 0.074308711 | 0.313236304 |
| SERINC3 | 1680.829257 | -0.600823486 | 0.263221901 | -2.282574076 | 0.022455469 | 0.170466268 |
| SERINC5 | 4334.200866 | -1.20689338 | 0.540239344 | -2.233997566 | 0.025483233 | |

0.182111117
SERP1 3928.547668 0.359355751 0.279497861 1.285719143 0.198541098 0.504280428
SERP2 34.02637023 0.705519806 0.543722702 1.297572831 0.194434163 0.500298624
SERPINA1 17.43051358 0.462044106 0.726252254 0.636203335 0.524643899
0.777662198
SERPINA10 26.40035702 0.863469512 0.588445493 1.467373822 0.142274392
0.432548373
SERPINA5 186.2340815 -0.653620947 0.332919187 -1.963302125
0.049611079 0.256701799
SERPINB1 538.4266524 0.189664918 0.3699171 0.512722764 0.608145264
0.827607804
SERPINB6 260.741315 0.354377007 0.368919137 0.960581797 0.336762485
0.644459616
SERPINB8 37.41420994 -1.514233463 0.657268306 -2.30382851 0.021232272
0.165290607
SERPINB9 491.8482823 -2.603868541 0.666676679 -3.905744154 9.39E-05
0.005843825
SERPINE1 30.57345949 -1.416126349 1.116011758 -1.268917051
0.20447065 0.510835602
SERPINE2 1102.113173 0.744051418 0.490282291 1.517597989 0.129115774
0.413453061
SERPINF1 609.9115309 0.458466452 0.384607474 1.192037294 0.233246623
0.543144207
SERPINF2 18.59686853 0.852748634 1.000519125 0.85230618 0.394044177
0.692865578
SERPING1 2453.753051 -0.494995614 0.30998736 -1.596825156
0.110304687 0.382149441
SERPINH1 6677.365754 0.085888143 0.252662418 0.339932404 0.733907433
0.888879903
SERPINI1 238.7473094 -0.193845534 0.295774069 -0.655383803
0.512220677 0.771687791
SERTAD1 235.1766573 -0.292084244 0.503878054 -0.579672485
0.562135502 0.801706045
SERTAD2 1292.107377 -0.647656648 0.231244792 -2.800740468
0.005098551 0.075897737
SERTAD3 962.4015576 0.059883624 0.377891579 0.15846774 0.874088239
0.949740014
SERTAD4 114.1157088 -0.569511488 0.380258183 -1.497696863
0.134212027 0.420619758
SERTM2 17.37681293 -6.737592944 2.169277659 -3.105915426
0.00189691 0.040704407
SESN1 1246.611105 0.595972614 0.298052346 1.99955686 0.045548136 0.245736406
SESN2 421.549422 -0.160044682 0.304769937 -0.525132774 0.599490894
0.823816451
SESN3 4147.913428 -0.527622682 0.49800551 -1.059471576 0.289385067
0.60229533
SESTD1 807.1725854 0.389083462 0.386962295 1.005481586 0.314665012
0.624045314
SET 25981.51194 0.268592048 0.256670521 1.046446811 0.29535479 0.60699582
SETBP1 556.8552982 0.728934055 0.598900569 1.217120324 0.223558439
0.53268495
SETD1A 972.3964427 -0.414846669 0.388715296 -1.067224969
0.28587026 0.599853073
SETD1B 693.7297578 -0.326847506 0.316823358 -1.03163955 0.302241006
0.613115536
SETD2 1934.823278 -0.595250039 0.39781505 -1.496298441 0.134575901
0.420896533
SETD3 1499.160663 0.094257872 0.250218134 0.3767028 0.706394482 0.874690719
SETD4 558.4428498 -0.374053325 0.29065982 -1.286911018 0.198125304
0.503899244
SETD5 7081.42848 -0.297683915 0.287595079 -1.035079999 0.300631552
0.611214323
SETD6 524.1696787 -0.183104575 0.266850146 -0.686170039 0.492605889
0.759174768

| | | | | | | |
|---|---|---|---|---|---|---|
| SETD7 | 920.1808345 | 0.190555474 | 0.374244707 | 0.509173464 | 0.610630642 | 0.828982323 |
| SETD9 | 255.1135002 | 0.32058514 | 0.316868727 | 1.011728557 | 0.311667858 | 0.621655675 |
| SETDB1 | 1948.894705 | -0.458039667 | 0.272576768 | -1.680406113 | 0.092878327 | 0.349511008 |
| SETDB2 | 343.54914 | -0.137165324 | 0.359147605 | -0.381919083 | 0.702521385 | 0.872703476 |
| SETMAR | 679.0168393 | 0.274175522 | 0.316443383 | 0.866428363 | 0.386255295 | 0.686679504 |
| SETX | 1547.737631 | -0.521385972 | 0.434808973 | -1.199115023 | 0.230483223 | 0.53927529 |
| SEZ6L2 | 1159.480577 | -0.105583308 | 0.309760636 | -0.340854506 | 0.733213115 | 0.888457729 |
| SF1 | 5820.832931 | -0.369778185 | 0.230767706 | -1.602382721 | 0.109071003 | 0.379241044 |
| SF3A1 | 3351.037884 | -0.427918428 | 0.361034204 | -1.185257307 | 0.235915728 | 0.546757945 |
| SF3A2 | 1401.655806 | -0.463745941 | 0.394558109 | -1.17535524 | 0.239852623 | 0.551501083 |
| SF3A3 | 3538.734827 | 0.199966573 | 0.231833604 | 0.862543524 | 0.388388493 | 0.688506874 |
| SF3B1 | 14366.1782 | -0.418596867 | 0.279760518 | -1.49626856 | 0.134583685 | 0.420896533 |
| SF3B2 | 6907.547475 | -0.411192983 | 0.327896575 | -1.254032565 | 0.209830169 | 0.517783588 |
| SF3B3 | 5631.036924 | -0.273118235 | 0.306901525 | -0.889921402 | 0.373508091 | 0.678377978 |
| SF3B4 | 1868.887461 | -0.493926635 | 0.26420212 | -1.869502923 | 0.061552877 | 0.285921219 |
| SF3B5 | 1779.426326 | 1.018131065 | 0.420170799 | 2.423136182 | 0.015387157 | 0.14062564 |
| SF3B6 | 2699.767078 | 0.969173093 | 0.394879696 | 2.45435028 | 0.014113942 | 0.134922886 |
| SFI1 | 547.5303597 | -0.199635553 | 0.505321272 | -0.395066593 | 0.692793743 | 0.86854217 |
| SFMBT1 | 282.1731119 | 0.051779694 | 0.442158014 | 0.117106763 | 0.906775451 | 0.965234099 |
| SFMBT2 | 440.112426 | -1.657988563 | 0.48787664 | -3.39837661 | 0.00067787 | 0.021392167 |
| SFN | 451.63229 | 1.134485302 | 0.707911226 | 1.602581313 | 0.109027122 | 0.379241044 |
| SFPQ | 12631.11104 | -0.086821174 | 0.237265586 | -0.365924007 | 0.714421782 | 0.878464326 |
| SFR1 | 305.1839081 | 0.540311647 | 0.348547777 | 1.550179578 | 0.12109842 | 0.401219509 |
| SFRP1 | 2320.576308 | 1.71557353 | 0.625509588 | 2.742681427 | 0.006093978 | 0.084270328 |
| SFRP2 | 764.4318735 | 1.600189439 | 0.424258893 | 3.771728687 | 0.00016212 | 0.008377957 |
| SFSWAP | 1490.639243 | -0.201966432 | 0.280104871 | -0.721038628 | 0.470885749 | 0.745855888 |
| SFT2D1 | 889.3325043 | 0.821746663 | 0.449515935 | 1.828070149 | 0.067539024 | 0.298553187 |
| SFT2D2 | 792.349583 | -0.08640998 | 0.522708877 | -0.165311866 | 0.86869852 | 0.947046732 |
| SFT2D3 | 617.0110538 | 0.641603459 | 0.307032325 | 2.089693519 | 0.03664534 | 0.219135375 |
| SFTPB | 12.84106836 | 1.411873585 | 1.025174349 | 1.377203386 | 0.168449375 | 0.470634525 |
| SFTPD | 23.69129265 | -0.683144218 | 0.964147597 | -0.708547343 | 0.478605423 | 0.751026363 |
| SFXN1 | 2056.696023 | 0.176446392 | 0.262987457 | 0.670930828 | 0.502264596 | 0.764661936 |
| SFXN2 | 610.7905962 | -0.282832682 | 0.336712869 | -0.839981802 | 0.40091859 | 0.698476795 |
| SFXN3 | 84.9572859 | -2.686674251 | 0.855141539 | -3.141788966 | 0.00167919 | 0.038017939 |
| SFXN4 | 526.6353215 | 0.726002124 | 0.261588131 | 2.775363399 | 0.005514006 | 0.079368821 |
| SFXN5 | 348.6574892 | -0.026445624 | 0.370378566 | -0.071401604 | 0.943078133 | 0.978607681 |
| SGCA | 15.32367514 | -0.272097404 | 0.941861314 | -0.288893279 | 0.772663047 | 0.905717252 |
| SGCB | 986.4638322 | 0.309715085 | 0.299862829 | 1.032855876 | 0.30167135 | 0.612425539 |
| SGCE | 693.1803019 | 0.055598493 | 0.248293924 | 0.223922082 | 0.822817933 | 0.926014081 |

| | | | | | | |
|---|---|---|---|---|---|---|
| SGCG | 171.3639252 | -1.762862495 | 1.057168429 | -1.667532293 | 0.095408609 | 0.354701001 |
| SGF29 | 276.7979561 | 0.447659416 | 0.309136476 | 1.448096391 | 0.147590095 | 0.441286821 |
| SGK1 | 367.3822893 | -1.342508261 | 0.582786398 | -2.303602599 | 0.021244962 | 0.165290607 |
| SGK2 | 32.8960185 | -0.222509663 | 0.577652929 | -0.385196113 | 0.700092123 | 0.871796568 |
| SGK3 | 304.5009926 | -1.285486185 | 0.575658833 | -2.233069506 | 0.025544361 | 0.182282927 |
| SGMS1 | 273.5858081 | -0.439648017 | 0.342875306 | -1.282238789 | 0.199758893 | 0.505887637 |
| SGMS2 | 15.33058761 | -0.871117701 | 0.782724261 | -1.112930498 | 0.26573828 | 0.579572403 |
| SGO1 | 583.1243835 | 0.724961928 | 0.25991014 | 2.789279123 | 0.005282552 | 0.077250742 |
| SGO2 | 1539.04159 | 1.105962345 | 0.386635259 | 2.860479792 | 0.004230005 | 0.067267723 |
| SGPL1 | 2056.240366 | -0.334370102 | 0.351362335 | -0.951639004 | 0.341280093 | 0.648152255 |
| SGPP1 | 391.3736917 | -1.123720915 | 0.655236666 | -1.714984788 | 0.086348016 | 0.337103899 |
| SGPP2 | 411.8406082 | -0.312836027 | 0.907853981 | -0.344588484 | 0.730403764 | 0.886760204 |
| SGSH | 446.6646537 | -0.186442059 | 0.311367527 | -0.598784534 | 0.549316577 | 0.79348329 |
| SGSM1 | 516.1448937 | 0.25061948 | 0.35764628 | 0.700746783 | 0.483461054 | 0.754199245 |
| SGSM2 | 2456.708446 | -1.055568718 | 0.370056963 | -2.852449279 | 0.004338374 | 0.068559859 |
| SGSM3 | 1954.056597 | 0.016854706 | 0.329235509 | 0.051193463 | 0.959171361 | 0.985871727 |
| SGTA | 1814.260848 | 0.021690169 | 0.387238638 | 0.056012409 | 0.955331922 | 0.984323736 |
| SGTB | 250.0165941 | 0.198763097 | 0.430127577 | 0.462102659 | 0.644007703 | 0.845542265 |
| SH2B1 | 1148.536012 | -0.306993424 | 0.457959979 | -0.670349894 | 0.502634768 | 0.764968568 |
| SH2B2 | 152.369821 | -0.350435006 | 0.465400627 | -0.752974934 | 0.451464977 | 0.732638351 |
| SH2B3 | 137.0180049 | 0.499734292 | 0.500320456 | 0.998828423 | 0.317877815 | 0.628014628 |
| SH2D2A | 57.00016554 | 1.471737965 | 0.654744968 | 2.247803399 | 0.024588729 | 0.179085746 |
| SH2D3A | 205.5762879 | -1.151467866 | 0.601071234 | -1.915692851 | 0.055404203 | 0.272371083 |
| SH2D3C | 10.71787274 | 0.997871126 | 0.922588468 | 1.081599392 | 0.279430575 | 0.593066767 |
| SH2D4A | 701.9942781 | -1.741728109 | 0.695378501 | -2.504719527 | 0.012254853 | 0.125960958 |
| SH2D5 | 124.9878083 | 0.873000234 | 0.449326714 | 1.942907483 | 0.052027338 | 0.262949685 |
| SH2D6 | 21.14263903 | 0.50193771 | 0.655326707 | 0.765935074 | 0.443714935 | 0.726848446 |
| SH3BGR | 398.465304 | 0.95688103 | 0.519500613 | 1.841924739 | 0.065486159 | 0.295242355 |
| SH3BGRL | 1646.505489 | -0.088469244 | 0.301041816 | -0.293876928 | 0.768851949 | 0.904486113 |
| SH3BGRL2 | 1425.909904 | -0.624536685 | 0.406746072 | -1.535446137 | 0.124674278 | 0.406173955 |
| SH3BGRL3 | 2713.165084 | 0.56197457 | 0.445322583 | 1.261949408 | 0.206966992 | 0.513642439 |
| SH3BP1 | 550.8857095 | -0.269255061 | 0.472614019 | -0.569714503 | 0.568871352 | 0.805450586 |
| SH3BP2 | 505.1413165 | -0.276539975 | 0.302006567 | -0.915675371 | 0.359837187 | 0.667315175 |
| SH3BP4 | 1419.115958 | 0.035590728 | 0.400793995 | 0.088800553 | 0.929240418 | 0.973104851 |
| SH3BP5 | 1293.285828 | -1.001524659 | 0.494946869 | -2.02349933 | 0.043021693 | 0.238392703 |
| SH3BP5L | 1048.015515 | -0.267589911 | 0.278535615 | -0.960702677 | 0.336701685 | 0.644459616 |
| SH3D19 | 1530.22995 | -0.575145634 | 0.242887401 | -2.367951695 | 0.017886871 | 0.150854036 |

SH3D21 462.0704651 -1.027898671 0.305175573 -3.368220667 0.00075655 0.022909426
SH3GL1 1089.119627 0.252045062 0.44615925 0.564921746 0.572126992 0.807358601
SH3GL1P1 18.09087859 -1.578024306 0.795358018 -1.984042745 0.047251075 0.250276443
SH3GL2 25.05140796 -0.857789055 0.683609402 -1.254794115 0.209553509 0.517437548
SH3GL3 130.9654757 0.004904421 0.41978841 0.011683078 0.990678465 0.996726674
SH3GLB1 1652.246908 -0.159020848 0.229697699 -0.692304925 0.488745858 0.757260184
SH3GLB2 2635.948827 -0.612672385 0.359473626 -1.704359765 0.088313896 0.341701126
SH3KBP1 1569.901179 -0.611368945 0.325554777 -1.877929576 0.0603908 0.284001495
SH3PXD2A 855.5427393 -0.310137382 0.3483092 -0.890408239 0.37324672 0.678228049
SH3PXD2B 1919.600698 -0.168586578 0.484014145 -0.348309197 0.727607988 0.88517603
SH3RF1 684.5236837 -0.290157696 0.340863498 -0.851243086 0.394634333 0.693581727
SH3RF2 55.26912228 -1.445118853 0.594437135 -2.431070954 0.015054266 0.139165886
SH3TC1 15.29390806 -2.76688806 1.088325888 -2.542334139 0.011011485 0.119239998
SH3TC2 16.46527066 0.464533314 0.906831103 0.512260015 0.608469046 0.827858453
SH3YL1 873.4394158 -0.957307618 0.47217298 -2.027451078 0.042616303 0.237346523
SHANK2 658.0940737 -1.327237866 0.328966593 -4.034567318 5.47E-05 0.004040998
SHANK3 81.69183956 -0.370846143 0.884076688 -0.419472822 0.674870613 0.859796949
SHARPIN 650.747374 0.100656216 0.43907348 0.229246859 0.818677054 0.924316427
SHB 444.1206785 -0.477708132 0.465146199 -1.027006418 0.304417442 0.614873787
SHBG 10.59340171 -0.66723575 1.029952217 -0.647831752 0.517093776 0.773950969
SHC1 4234.139093 0.173947235 0.351307826 0.495141931 0.620499937 0.833283575
SHC2 260.6603218 -1.112469126 0.64346607 -1.728869908 0.083832381 0.331289671
SHC3 18.92466076 0.935323063 0.804805937 1.162172171 0.245165536 0.556630702
SHC4 20.90569722 0.939207127 0.681951765 1.377233955 0.168439927 0.470634525
SHCBP1 273.5807935 0.490309323 0.349394974 1.403309605 0.160524535 0.460961874
SHD 21.40808421 -0.416477423 0.881528152 -0.472449374 0.63660607 0.841268221
SHF 891.4093849 0.831157728 0.319184088 2.604007403 0.009214077 0.108783468
SHISA2 345.183697 0.265869258 0.861215913 0.308713824 0.757539227 0.898914948
SHISA4 287.3982903 0.162339474 0.322033519 0.504107382 0.61418592 0.830316752
SHISA5 4393.809866 -0.066238783 0.384068891 -0.172465889 0.863071276 0.944730333
SHISA9 319.5724389 -0.594005881 0.341030923 -1.74179478 0.081544361 0.326496916
SHISAL1 69.30360265 0.942669248 0.755727598 1.247366446 0.212263164 0.519478054
SHISAL2A 13.54999815 -0.879376702 1.031199415 -0.852770754 0.393786446 0.692653237
SHISAL2B 26.04408115 1.023814343 0.638283726 1.604011354 0.10871155 0.37853177

| | | | | | | |
|---|---|---|---|---|---|---|
| SHKBP1 | 1596.656736 | -0.338877022 | 0.34659411 | -0.977734509 | 0.32820565 | 0.637787615 |
| SHLD1 | 99.77257033 | 0.891450912 | 0.423347819 | 2.105717505 | 0.035228898 | 0.214555446 |
| SHLD2 | 571.2717939 | 0.445496434 | 0.417873787 | 1.066102846 | 0.286377153 | 0.60033547 |
| SHLD2P1 | 32.5553273 | 0.566866657 | 0.613180812 | 0.924469008 | 0.35524218 | 0.662655738 |
| SHLD2P3 | 18.22946956 | -0.382705947 | 0.786987118 | -0.486292518 | 0.626759783 | 0.836041947 |
| SHLD3 | 111.1016125 | 0.322955778 | 0.361735926 | 0.892794313 | 0.371967332 | 0.677193964 |
| SHMT1 | 822.7367778 | 0.42741534 | 0.237132448 | 1.80243296 | 0.071477313 | 0.307538946 |
| SHMT2 | 3199.052382 | 0.412707858 | 0.262737609 | 1.570798563 | 0.116229446 | 0.392895928 |
| SHOC2 | 982.7315079 | -0.030783417 | 0.259260895 | -0.118735286 | 0.90548508 | 0.964473572 |
| SHPK | 924.468672 | -0.172085847 | 0.342734188 | -0.502097116 | 0.615599209 | 0.830968556 |
| SHPRH | 921.7411639 | -0.235931541 | 0.391966933 | -0.601916951 | 0.54722942 | 0.792404079 |
| SHQ1 | 661.0833391 | -0.030796976 | 0.254779795 | -0.120876838 | 0.903788588 | 0.963500625 |
| SHROOM1 | 29.85869193 | -0.536336934 | 0.660941081 | -0.811474652 | 0.417093144 | 0.708661116 |
| SHROOM2 | 565.0901078 | -0.344643407 | 0.428573163 | -0.804164694 | 0.421301866 | 0.711694812 |
| SHROOM3 | 881.3564084 | 0.38604739 | 0.525275515 | 0.734942672 | 0.462374409 | 0.73986733 |
| SHROOM4 | 94.02763257 | -0.59589167 | 0.419340279 | -1.42102178 | 0.155310428 | 0.453268877 |
| SHTN1 | 1768.183674 | -1.068963468 | 0.44858732 | -2.382955158 | 0.017174286 | 0.147741524 |
| SIAE | 229.7508985 | -0.121328213 | 0.31118972 | -0.389885028 | 0.696621566 | 0.870023227 |
| SIAH1 | 1142.287886 | 0.397488975 | 0.329399349 | 1.206708443 | 0.227544447 | 0.536368893 |
| SIAH2 | 666.9179861 | -0.506919532 | 0.415250959 | -1.220754632 | 0.222178936 | 0.531066116 |
| SIDT2 | 1238.354871 | -0.455252772 | 0.323496223 | -1.407289297 | 0.159341622 | 0.459178093 |
| SIGIRR | 377.1831237 | -0.782828882 | 0.637134642 | -1.22867104 | 0.219195169 | 0.52778634 |
| SIGLEC6 | 14.2852173 | -2.594041088 | 1.628973019 | -1.592439566 | 0.111285972 | 0.383708973 |
| SIGMAR1 | 1270.744198 | 0.158303352 | 0.356645311 | 0.443867751 | 0.657138202 | 0.850335056 |
| SIK1 | 111.6139297 | 0.421039873 | 0.418762215 | 1.005439026 | 0.314685496 | 0.624045314 |
| SIK1B | 144.270488 | 0.556121259 | 0.531663643 | 1.046002048 | 0.295560087 | 0.60699582 |
| SIK2 | 900.3321626 | -0.504069728 | 0.374536598 | -1.345849058 | 0.178351201 | 0.4825693 |
| SIK3 | 1004.424256 | -0.519309486 | 0.417553567 | -1.243695485 | 0.213611658 | 0.520678417 |
| SIKE1 | 1714.812629 | -0.068717229 | 0.230287039 | -0.298398162 | 0.765399291 | 0.903026935 |
| SIL1 | 345.9333673 | 0.414065281 | 0.312519868 | 1.324924666 | 0.185196139 | 0.489784764 |
| SIMC1 | 326.0186018 | -0.196730568 | 0.331104868 | -0.594163927 | 0.552402477 | 0.795662343 |
| SIN3A | 2497.498886 | -0.230855907 | 0.372500294 | -0.619746912 | 0.535424425 | 0.784368864 |
| SIN3B | 1559.968819 | -0.971340801 | 0.365966271 | -2.654181211 | 0.007950112 | 0.100258644 |
| SINHCAF | 7095.564775 | -0.047106169 | 0.334925337 | -0.140646778 | 0.888148992 | 0.955943827 |
| SIPA1 | 265.8475161 | 0.798319394 | 0.367708087 | 2.171068365 | 0.029926005 | 0.197328521 |
| SIPA1L1 | 1739.50478 | -0.716209963 | 0.42751544 | -1.675284438 | 0.093878437 | 0.351314695 |
| SIPA1L2 | 1936.397758 | -0.701922368 | 0.322374054 | -2.177353793 | 0.02945418 | 0.196075451 |

| | | | | | | |
|---|---|---|---|---|---|---|
| SIPA1L3 | 552.0694972 | -0.378913929 | 0.422081436 | -0.897727066 | 0.369331076 | 0.675154642 |
| SIRPA | 451.6071809 | -0.088193427 | 0.339992175 | -0.259398403 | 0.795327862 | 0.915124686 |
| SIRT1 | 1129.508462 | 0.104469191 | 0.272725855 | 0.383055692 | 0.701678471 | 0.872479299 |
| SIRT2 | 205.4067584 | -0.077145508 | 0.337966862 | -0.228263526 | 0.819441378 | 0.924316427 |
| SIRT3 | 478.1062487 | 0.058524701 | 0.270427828 | 0.216415233 | 0.828664094 | 0.929129846 |
| SIRT4 | 91.43994001 | 0.383129764 | 0.415144223 | 0.922883525 | 0.356067907 | 0.663487122 |
| SIRT5 | 580.5320838 | 0.254565583 | 0.301874174 | 0.843283741 | 0.399069774 | 0.697352242 |
| SIRT6 | 307.4580688 | 0.247435643 | 0.492212067 | 0.502701295 | 0.615174299 | 0.830958873 |
| SIRT7 | 549.6740059 | -0.055804368 | 0.373159095 | -0.149545782 | 0.881122987 | 0.952848873 |
| SIT1 | 20.89727223 | 0.936936266 | 0.689498187 | 1.358866903 | 0.174188768 | 0.478455902 |
| SIVA1 | 612.9421442 | 0.623477448 | 0.498074573 | 1.251775301 | 0.210651753 | 0.518294063 |
| SIX1 | 50.34108103 | 0.02795764 | 0.874826608 | 0.031957921 | 0.974505608 | 0.991231953 |
| SIX4 | 689.6256243 | -0.238243478 | 0.243662256 | -0.977761107 | 0.328192491 | 0.637787615 |
| SIX5 | 590.6826868 | 0.569363005 | 0.434312401 | 1.310952677 | 0.189873752 | 0.495280824 |
| SKA1 | 394.2045252 | 0.617662696 | 0.37581875 | 1.64351219 | 0.100277005 | 0.36243601 |
| SKA2 | 1814.127699 | 0.780588016 | 0.266788652 | 2.925866641 | 0.003434982 | 0.059693132 |
| SKA3 | 630.7272772 | 0.601494271 | 0.318021215 | 1.891365237 | 0.058575606 | 0.280288923 |
| SKAP2 | 633.1596344 | -0.78622535 | 0.554116606 | -1.418880688 | 0.155933804 | 0.454370782 |
| SKI | 1169.518885 | -0.087510086 | 0.306014142 | -0.285967456 | 0.774903048 | 0.906450643 |
| SKIDA1 | 1746.853551 | 1.087704203 | 0.571230775 | 1.904141462 | 0.056891763 | 0.2761211 |
| SKIL | 861.8257226 | -0.845194375 | 0.425751111 | -1.98518419 | 0.047123983 | 0.250100439 |
| SKIV2L | 82.83550415 | -0.104612879 | 0.687279697 | -0.152212964 | 0.87901897 | 0.951774989 |
| SKP1 | 7744.820446 | 0.588364568 | 0.364647692 | 1.613515128 | 0.106632639 | 0.374599763 |
| SKP2 | 1879.568227 | 0.293514803 | 0.248232921 | 1.182416908 | 0.237040318 | 0.547774358 |
| SLA2 | 13.38950189 | -1.445190637 | 0.93779414 | -1.541053175 | 0.123303845 | 0.404753154 |
| SLAIN1 | 526.4911963 | 0.335366321 | 0.394753602 | 0.849558608 | 0.395570533 | 0.69426198 |
| SLAIN2 | 942.0132737 | -0.041387328 | 0.31570033 | -0.131096878 | 0.895698671 | 0.958837464 |
| SLBP | 1693.877084 | 0.3943129 | 0.361124925 | 1.091901647 | 0.274876334 | 0.587924553 |
| SLC10A3 | 211.1533483 | -0.899503585 | 0.489542765 | -1.837436173 | 0.066145531 | 0.296938744 |
| SLC10A5 | 10.40732229 | 1.251878232 | 0.90722438 | 1.379899239 | 0.167617671 | 0.469868383 |
| SLC10A7 | 209.0465487 | -0.38876418 | 0.305236832 | -1.273647672 | 0.202788309 | 0.509290789 |
| SLC11A1 | 13.73148797 | 1.780596402 | 1.037196241 | 1.716740122 | 0.086026666 | 0.33625288 |
| SLC11A2 | 1926.378994 | -0.335171032 | 0.285855039 | -1.172520986 | 0.240987952 | 0.552437022 |
| SLC12A2 | 424.1190955 | 0.031900216 | 0.497402935 | 0.064133551 | 0.948863887 | 0.98134156 |
| SLC12A3 | 73.56549099 | -4.54904954 | 1.008405678 | -4.511130432 | 6.45E-06 | 0.000782618 |
| SLC12A4 | 1004.714945 | -0.433980102 | 0.307218212 | -1.4126119 | 0.157769868 | 0.457216174 |
| SLC12A5 | 45.82802277 | 0.121074469 | 0.491794932 | 0.246188933 | 0.805535982 | 0.918930547 |
| SLC12A6 | 1023.9438 | -0.965839771 | 0.246671007 | -3.915497739 | 9.02E-05 | 0.00572676 |
| SLC12A7 | 425.4269157 | 0.181291184 | 0.442408487 | 0.409782338 | 0.681965624 | 0.862735748 |
| SLC12A8 | 87.65051879 | -0.23967912 | 0.435268725 | -0.550646316 | 0.581876153 | 0.813002261 |

| | | | | | | |
|---|---|---|---|---|---|---|
| SLC12A9 | 887.0967709 | -0.623942178 | 0.399050404 | -1.563567336 | 0.117919211 | 0.395839103 |
| SLC13A3 | 265.0276362 | 0.331241766 | 0.382651451 | 0.86564879 | 0.386682791 | 0.687008387 |
| SLC13A4 | 253.6229374 | -2.677754104 | 0.552085269 | -4.85025458 | 1.23E-06 | 0.000215015 |
| SLC15A1 | 426.4826222 | -6.854204346 | 2.046088455 | -3.349906172 | 0.000808389 | 0.023860069 |
| SLC15A2 | 59.57198628 | -0.266998082 | 0.538072898 | -0.496211726 | 0.619745039 | 0.832851852 |
| SLC15A3 | 171.3752955 | 2.349361734 | 0.808024124 | 2.9075391 | 0.003642848 | 0.062135365 |
| SLC16A1 | 10220.27743 | -0.384558026 | 0.338129393 | -1.137310256 | 0.255408609 | 0.567820048 |
| SLC16A10 | 147.7734052 | -0.105340922 | 0.529811828 | -0.198827049 | 0.842398036 | 0.935547316 |
| SLC16A11 | 18.58413576 | 2.227780468 | 1.166554119 | 1.909710345 | 0.056170518 | 0.274537027 |
| SLC16A12 | 82.28377173 | 0.084517004 | 0.568949566 | 0.148549203 | 0.881909359 | 0.953217539 |
| SLC16A13 | 131.4945396 | -0.601316588 | 0.420255955 | -1.430834188 | 0.152477742 | 0.448453866 |
| SLC16A14 | 51.69956628 | 1.151666078 | 0.752419062 | 1.530617891 | 0.125863857 | 0.408095871 |
| SLC16A2 | 951.8753047 | 0.309673636 | 0.457403955 | 0.677024396 | 0.498390476 | 0.762744065 |
| SLC16A3 | 140.7100395 | -0.37516097 | 0.596644263 | -0.628785012 | 0.529489814 | 0.780195218 |
| SLC16A4 | 337.2002639 | -1.696834255 | 0.551963029 | -3.074181 | 0.002110813 | 0.0433918 |
| SLC16A5 | 264.8304351 | -0.651699597 | 0.480300263 | -1.356858715 | 0.174826107 | 0.478752143 |
| SLC16A6 | 34.01969706 | -0.598628534 | 0.873609125 | -0.685236128 | 0.493194929 | 0.759389046 |
| SLC16A7 | 41.6610226 | 0.067355921 | 0.568123548 | 0.118558579 | 0.905625083 | 0.964495481 |
| SLC16A9 | 815.2556822 | -1.544132456 | 0.437808745 | -3.526956632 | 0.000420366 | 0.015778519 |
| SLC17A5 | 444.3659816 | 0.668926441 | 0.321087486 | 2.083315204 | 0.037222502 | 0.220973863 |
| SLC17A7 | 128.8388622 | 0.661505559 | 0.406667117 | 1.626651214 | 0.103811196 | 0.369190637 |
| SLC17A9 | 49.83656877 | 0.598669649 | 0.692178524 | 0.86490642 | 0.387090154 | 0.68748084 |
| SLC18A2 | 22.13455439 | 0.184030849 | 0.630254104 | 0.291994685 | 0.770290688 | 0.904985675 |
| SLC18B1 | 701.9257225 | 0.112447961 | 0.393073983 | 0.286073272 | 0.774822002 | 0.906450643 |
| SLC19A1 | 274.2443439 | 0.442944143 | 0.337130413 | 1.31386587 | 0.188891346 | 0.494119637 |
| SLC19A2 | 1011.00114 | -0.459620158 | 0.36605984 | -1.255587494 | 0.209265568 | 0.516894811 |
| SLC19A3 | 148.409989 | -0.579622144 | 0.568910598 | -1.018828171 | 0.308284549 | 0.617978528 |
| SLC1A1 | 375.5942655 | -0.203890165 | 0.47771124 | -0.426806297 | 0.66952042 | 0.857712777 |
| SLC1A2 | 74.6503582 | 1.096620285 | 0.513595981 | 2.135180814 | 0.032746243 | 0.206309493 |
| SLC1A3 | 288.0282864 | -0.706709049 | 0.635630284 | -1.11182407 | 0.266213801 | 0.57970088 |
| SLC1A4 | 47.18420484 | -0.210385011 | 0.67545643 | -0.311470884 | 0.755442674 | 0.897831634 |
| SLC1A5 | 3298.459495 | 0.583825178 | 0.259368182 | 2.250951426 | 0.024388614 | 0.178492903 |

| | | | | | | |
|---|---|---|---|---|---|---|
| SLC1A6 | 196.3042088 | 0.592283446 | 0.512696282 | 1.155232574 | 0.247995199 | 0.560705687 |
| SLC20A1 | 1140.735481 | 0.333739511 | 0.302211757 | 1.104323384 | 0.269452883 | 0.582188661 |
| SLC20A2 | 1075.423828 | -0.556163091 | 0.230769185 | -2.410040543 | 0.015950748 | 0.142746012 |
| SLC22A15 | 59.71816541 | -1.292322903 | 0.532381528 | -2.427437532 | 0.015205905 | 0.139780768 |
| SLC22A16 | 64.40471874 | 1.816864853 | 1.007706952 | 1.802969455 | 0.071393012 | 0.307350563 |
| SLC22A17 | 725.1472317 | 0.406164311 | 0.36541437 | 1.111517072 | 0.266345846 | 0.57970088 |
| SLC22A18 | 68.40513621 | -0.076180347 | 0.698977984 | -0.108988193 | 0.913211855 | 0.967876517 |
| SLC22A20P | 36.20002445 | -0.930307032 | 0.590670711 | -1.575001121 | 0.115256186 | 0.390825122 |
| SLC22A23 | 527.555391 | -1.735596942 | 0.694036879 | -2.500727259 | 0.012393859 | 0.126591155 |
| SLC22A25 | 15.29462966 | -0.991640338 | 0.909202532 | -1.090670454 | 0.275417916 | 0.588496689 |
| SLC22A31 | 25.97355152 | -0.856113808 | 0.850894218 | -1.00613424 | 0.314351 | 0.623972134 |
| SLC22A5 | 199.4314305 | -0.773295887 | 0.37570861 | -2.058233071 | 0.039567765 | 0.229237914 |
| SLC23A2 | 428.3089197 | -0.759259155 | 0.563302189 | -1.347871834 | 0.177699609 | 0.482042363 |
| SLC23A3 | 10.44703369 | -0.726167064 | 1.152365413 | -0.63015347 | 0.528594179 | 0.779660133 |
| SLC24A1 | 441.7791576 | -0.968985571 | 0.331580666 | -2.922322291 | 0.003474318 | 0.060307645 |
| SLC24A3 | 26.65886051 | -1.081308244 | 0.841741223 | -1.284608872 | 0.198928997 | 0.504782074 |
| SLC24A4 | 28.03967197 | -0.5657921 | 0.752145313 | -0.752237754 | 0.451908093 | 0.732888936 |
| SLC25A1 | 2081.753628 | 0.091648354 | 0.270405308 | 0.338929568 | 0.734662789 | 0.889483064 |
| SLC25A10 | 344.7580099 | 0.367740327 | 0.383743439 | 0.958297366 | 0.337912831 | 0.645164305 |
| SLC25A11 | 1451.351106 | 0.458519287 | 0.252150383 | 1.818435815 | 0.068997549 | 0.302794854 |
| SLC25A12 | 118.9373981 | -0.337106995 | 0.450757464 | -0.747867805 | 0.454539899 | 0.734536089 |
| SLC25A13 | 4118.2243 | -0.415080818 | 0.262506007 | -1.581224075 | 0.113826813 | 0.388234391 |
| SLC25A14 | 258.6162744 | 0.059480257 | 0.29494579 | 0.201665048 | 0.840178589 | 0.934461703 |
| SLC25A15 | 902.3388492 | 0.315263166 | 0.262390678 | 1.201502923 | 0.229556173 | 0.538434864 |
| SLC25A16 | 290.3825265 | -0.1026719 | 0.392076273 | -0.261867159 | 0.793423859 | 0.914254706 |
| SLC25A17 | 848.7576677 | 0.385627831 | 0.27130769 | 1.421367125 | 0.155210059 | 0.453173941 |
| SLC25A18 | 14.72593741 | -0.452845816 | 0.773559885 | -0.585404989 | 0.558275435 | 0.798855459 |
| SLC25A19 | 168.2181214 | 0.621558937 | 0.326657457 | 1.902785085 | 0.057068594 | 0.276551437 |
| SLC25A20 | 140.8967185 | 0.141776946 | 0.402234647 | 0.352473231 | 0.72448339 | 0.883674024 |
| SLC25A21 | 154.3343835 | 0.857149819 | 0.390706865 | 2.193843766 | 0.028246643 | 0.192252052 |
| SLC25A22 | 518.7298612 | 0.704107207 | 0.460712757 | 1.528299782 | 0.126438124 | 0.409258114 |
| SLC25A23 | 1200.671233 | -0.263665457 | 0.325897208 | -0.809044848 | 0.41848935 | 0.709738117 |

| | | | | | | |
|---|---|---|---|---|---|---|
| SLC25A24 | 1191.822704 | -0.038483097 | 0.231412465 | -0.166296562 | 0.867923571 | 0.9466285 |
| SLC25A25 | 430.893363 | 0.160592613 | 0.467503388 | 0.343511122 | 0.731213975 | 0.887005928 |
| SLC25A26 | 330.525628 | 0.386399829 | 0.482012755 | 0.801638183 | 0.422762284 | 0.712269066 |
| SLC25A27 | 232.4950665 | -0.093242593 | 0.316122634 | -0.29495703 | 0.768026705 | 0.904216447 |
| SLC25A28 | 420.0753896 | 0.095216692 | 0.295476038 | 0.32224844 | 0.74726449 | 0.894202292 |
| SLC25A29 | 3532.96985 | -0.176848838 | 0.307003532 | -0.576048219 | 0.56458259 | 0.803120719 |
| SLC25A3 | 19513.67057 | 0.364524363 | 0.4006393 | 0.909856729 | 0.362898073 | 0.669987492 |
| SLC25A30 | 296.9294099 | -0.083152603 | 0.369548446 | -0.225011372 | 0.821970428 | 0.92563193 |
| SLC25A32 | 442.5934131 | 0.341700428 | 0.254291102 | 1.343737258 | 0.179033365 | 0.483466568 |
| SLC25A33 | 853.1262492 | 0.742595573 | 0.409338191 | 1.814137035 | 0.069656631 | 0.303425952 |
| SLC25A34 | 31.82363411 | -0.716061087 | 0.614884367 | -1.164545931 | 0.244202841 | 0.555457814 |
| SLC25A35 | 101.6736792 | -0.28033146 | 0.388416777 | -0.721728506 | 0.470461413 | 0.7454951 |
| SLC25A36 | 7370.895123 | -0.449301607 | 0.25301115 | -1.775817418 | 0.075763016 | 0.315769427 |
| SLC25A37 | 1878.138979 | -0.278951485 | 0.270960399 | -1.02949171 | 0.303248673 | 0.613656879 |
| SLC25A38 | 1209.838153 | -0.088675808 | 0.342885591 | -0.258616314 | 0.795931294 | 0.915299455 |
| SLC25A39 | 3205.032559 | 0.252911806 | 0.308810588 | 0.818986834 | 0.412793926 | 0.705474445 |
| SLC25A4 | 985.448072 | 0.554795358 | 0.316613949 | 1.752277052 | 0.07972618 | 0.323229791 |
| SLC25A40 | 407.1833425 | 0.106821603 | 0.336818673 | 0.317148697 | 0.751130776 | 0.896349431 |
| SLC25A41 | 11.63949073 | 1.469500382 | 0.879357003 | 1.671107839 | 0.094700379 | 0.353170957 |
| SLC25A42 | 133.9046399 | 0.96648543 | 0.39899117 | 2.422322858 | 0.015421642 | 0.140677297 |
| SLC25A43 | 167.8180434 | -0.188791068 | 0.312022827 | -0.60505531 | 0.545142247 | 0.791118522 |
| SLC25A44 | 647.7584602 | -0.601316099 | 0.292682168 | -2.054501998 | 0.03992713 | 0.230229757 |
| SLC25A45 | 37.61178606 | -0.923879882 | 0.609679919 | -1.515352324 | 0.129683204 | 0.414456265 |
| SLC25A46 | 851.5469601 | -0.177752513 | 0.287638465 | -0.617971984 | 0.536593807 | 0.785348759 |
| SLC25A47 | 69.58885474 | 0.590632546 | 0.585598926 | 1.008595679 | 0.313168581 | 0.623010823 |
| SLC25A5 | 9634.543151 | 0.50594508 | 0.367133561 | 1.378095423 | 0.168173829 | 0.470384431 |
| SLC25A51 | 154.3428806 | -0.303559162 | 0.333896622 | -0.909141159 | 0.363275619 | 0.670061327 |
| SLC25A53 | 126.4449159 | 0.479969843 | 0.450425518 | 1.065592033 | 0.286608102 | 0.600653615 |
| SLC25A6 | 18996.04275 | 0.372458107 | 0.381128418 | 0.977250945 | 0.328444932 | 0.637925754 |
| SLC26A10 | 53.98381336 | 0.452084043 | 0.47265719 | 0.956473427 | 0.338833103 | 0.645945842 |
| SLC26A11 | 686.0899111 | -0.453684618 | 0.31051831 | -1.461055928 | 0.144000094 | 0.434995877 |
| SLC26A2 | 448.1629457 | 0.802781747 | 0.472503056 | 1.698997998 | 0.089319561 | 0.342726775 |

| | | | | | | |
|---|---|---|---|---|---|---|
| SLC26A6 | 1350.123352 | -0.192896483 | 0.31439186 | -0.613554316 | 0.53950988 | 0.78700361 |
| SLC26A7 | 13.35784001 | -0.010043676 | 0.940516473 | -0.010678894 | 0.991479637 | 0.996814923 |
| SLC26A8 | 22.01330527 | -0.230504552 | 0.689810193 | -0.334156489 | 0.738261475 | 0.891242475 |
| SLC27A1 | 294.9429307 | 0.391259558 | 0.360624227 | 1.084950838 | 0.27794343 | 0.591980875 |
| SLC27A2 | 63.20119038 | -0.232865776 | 0.683635364 | -0.340628628 | 0.733383175 | 0.888457729 |
| SLC27A3 | 163.1472903 | 0.014302638 | 0.563127902 | 0.02539856 | 0.97973706 | 0.993024514 |
| SLC27A4 | 740.7412176 | -0.140509237 | 0.347018434 | -0.404904245 | 0.685547898 | 0.864830131 |
| SLC27A5 | 197.9460881 | 1.005409962 | 0.427720625 | 2.350623053 | 0.018742009 | 0.1554129 |
| SLC27A6 | 202.4356669 | -1.536922094 | 0.833772609 | -1.843334835 | 0.065280138 | 0.294751481 |
| SLC28A2 | 28.58976617 | -1.234761397 | 0.977256678 | -1.263497528 | 0.206410434 | 0.512832712 |
| SLC29A1 | 2654.812135 | 0.151665173 | 0.227495686 | 0.666672744 | 0.504981192 | 0.766569683 |
| SLC29A2 | 809.0036257 | 0.118166345 | 0.32601777 | 0.362453693 | 0.717013022 | 0.880371898 |
| SLC29A3 | 362.8584256 | -0.459002534 | 0.296859371 | -1.546195199 | 0.122057433 | 0.402659864 |
| SLC29A4 | 348.3621935 | 0.57603194 | 0.406021789 | 1.418721742 | 0.155980156 | 0.45437307 |
| SLC2A1 | 4428.993469 | -0.71776971 | 0.521417164 | -1.376574765 | 0.168643757 | 0.471031175 |
| SLC2A10 | 324.7069976 | -0.610929916 | 0.311570475 | -1.96080812 | 0.04990141 | 0.257391965 |
| SLC2A11 | 194.6108164 | -0.325335667 | 0.386287973 | -0.84221019 | 0.399670311 | 0.697589493 |
| SLC2A12 | 540.0642826 | -0.378332296 | 0.416142136 | -0.909142004 | 0.363275173 | 0.670061327 |
| SLC2A13 | 566.4707953 | -0.933156645 | 0.350680663 | -2.660986887 | 0.007791199 | 0.09891237 |
| SLC2A14 | 385.1540086 | 0.582190938 | 0.584005657 | 0.996892635 | 0.318816627 | 0.628804869 |
| SLC2A3 | 27839.4847 | -0.507796853 | 0.391227147 | -1.297959145 | 0.194301374 | 0.500187881 |
| SLC2A4 | 233.2870397 | 0.657335194 | 0.365488961 | 1.798509025 | 0.07209638 | 0.309063062 |
| SLC2A4RG | 546.2486473 | 0.554875505 | 0.434753291 | 1.276299724 | 0.201849588 | 0.50842771 |
| SLC2A6 | 35.3275092 | 0.518927658 | 0.531891027 | 0.975627773 | 0.329248956 | 0.638424836 |
| SLC2A8 | 323.9580251 | -0.133992033 | 0.490368349 | -0.273247719 | 0.784662801 | 0.910931322 |
| SLC30A1 | 758.1593031 | -0.117634368 | 0.371922955 | -0.316286926 | 0.751784734 | 0.896586461 |
| SLC30A2 | 10.18531317 | -1.430910171 | 1.216453123 | -1.17629701 | 0.23947621 | 0.550719052 |
| SLC30A3 | 14.49079803 | 1.341735973 | 0.826843446 | 1.622720697 | 0.104649125 | 0.370855378 |
| SLC30A4 | 36.95068907 | -0.729774942 | 0.660858024 | -1.104284 | 0.269469961 | 0.582188661 |
| SLC30A5 | 1208.780165 | 0.252422087 | 0.250142416 | 1.009113495 | 0.312920207 | 0.622994042 |
| SLC30A6 | 1307.495593 | -0.160455048 | 0.252741237 | -0.634858999 | 0.52552038 | 0.778371208 |
| SLC30A7 | 593.6081867 | -0.243940866 | 0.377129929 | -0.646835074 | 0.51773869 | 0.773950969 |
| SLC30A8 | 171.7410716 | -0.745532143 | 0.338739886 | -2.200898604 | | |

0.027743203 0.190621435
SLC30A9 3379.094345 0.090669241 0.268556287 0.337617273 0.735651624
0.889855747
SLC31A1 1530.88935 -0.282403867 0.300809334 -0.938813511
0.347826501 0.654455642
SLC31A2 90.69549125 -0.576161822 0.580367773 -0.992752956
0.320830361 0.63048153
SLC33A1 612.9093782 -0.053587443 0.287393498 -0.186460179
0.852083904 0.939869485
SLC34A2 145.6189525 -0.598144958 0.332311915 -1.799950384
0.071868473 0.308679129
SLC35A1 912.7195362 0.058269656 0.303681086 0.191877792 0.847837934
0.938207222
SLC35A2 706.341528 -0.038989895 0.457905946 -0.085148261
0.932143523 0.97444726
SLC35A3 592.1973629 -0.135787671 0.314159855 -0.43222477 0.665578058
0.855161526
SLC35A4 1549.494498 -0.241531585 0.247755191 -0.974880019
0.329619779 0.638548751
SLC35A5 1242.072395 0.015191407 0.270491621 0.056162207 0.955212588
0.984323736
SLC35B1 1124.614194 0.205522291 0.325077093 0.632226311 0.527239007
0.778906646
SLC35B2 1545.956398 0.003199896 0.323961247 0.009877402 0.992119101
0.99688215
SLC35B3 541.2643154 -0.437397446 0.394776696 -1.107961667
0.267878371 0.581031837
SLC35B4 1272.662648 0.036857998 0.334031092 0.110343015 0.912137345
0.967221336
SLC35C1 437.5224855 -0.45511806 0.366054992 -1.243305157 0.213755405
0.520770421
SLC35C2 1093.096361 -0.344178131 0.329477381 -1.044618388
0.296199377 0.607412916
SLC35D1 408.696978 -0.48186343 0.402690345 -1.196610338 0.231458469
0.540473516
SLC35D2 1396.866614 -0.139517756 0.548310585 -0.254450232
0.799147748 0.916115346
SLC35E1 1000.78685 -0.47754605 0.295129461 -1.618090069 0.105643189
0.372686009
SLC35E2A 1588.834086 -0.326677527 0.311018635 -1.050347121
0.293558548 0.606341284
SLC35E2B 1954.165034 0.020290233 0.232220282 0.087374939 0.930373489
0.97369593
SLC35E3 388.7177646 -0.039215992 0.336484854 -0.116546084
0.907219765 0.965296417
SLC35E4 61.00483656 1.232170971 0.53160447 2.317834104 0.020458339
0.161737081
SLC35F1 904.0518461 0.833303196 0.363767463 2.290757916 0.021977419
0.168393647
SLC35F2 1120.146598 -1.073711175 0.380463391 -2.822114299
0.004770817 0.073402552
SLC35F3 29.33873448 0.241989404 0.751387655 0.322056667 0.747409765
0.894202292
SLC35F5 1028.493199 0.056649781 0.311505788 0.181857876 0.855694263
0.942139318
SLC35F6 658.8000827 -0.530879076 0.392767357 -1.351637469
0.176491318 0.480709117
SLC35G1 367.5931587 0.127609476 0.329821437 0.386904736 0.698826735
0.870873025
SLC35G2 23.60173966 0.82025318 0.676915102 1.211751929 0.225607355
0.53459368
SLC36A1 784.9271175 -1.145318024 0.471154193 -2.43087728 0.015062315
0.139165886
SLC36A2 17.08610201 -0.274918424 0.921020408 -0.298493303

| | | | | | | |
|---|---|---|---|---|---|---|
| | | | | | 0.765326687 | 0.903026935 |
| SLC36A4 | 1800.106764 | -0.320647947 | 0.273493123 | -1.172416853 | 0.241029736 | 0.552449332 |
| SLC37A1 | 309.2399711 | -0.755951203 | 0.401330272 | -1.883613712 | 0.059617246 | 0.282669434 |
| SLC37A2 | 28.14013506 | 0.159286636 | 0.808056307 | 0.197123189 | 0.843731137 | 0.936580455 |
| SLC37A3 | 1550.816862 | -0.057735508 | 0.334814945 | -0.172440056 | 0.863091584 | 0.944730333 |
| SLC37A4 | 1726.282046 | 0.235432671 | 0.309194297 | 0.761439244 | 0.446394755 | 0.72859116 |
| SLC38A1 | 9834.808522 | -1.233088803 | 0.439606913 | -2.804980465 | 0.005031963 | 0.075658979 |
| SLC38A10 | 965.3252825 | -0.59361049 | 0.467042322 | -1.270999357 | 0.203728878 | 0.509986087 |
| SLC38A2 | 7852.148205 | -0.48475862 | 0.382134744 | -1.268554164 | 0.204600121 | 0.510860506 |
| SLC38A3 | 52.46637006 | 0.200447318 | 0.567868821 | 0.352981729 | 0.724102138 | 0.883667763 |
| SLC38A4 | 70.31927287 | 0.706944059 | 0.57332262 | 1.233065005 | 0.217551512 | 0.525720611 |
| SLC38A5 | 60.29206565 | 0.20928624 | 0.466718668 | 0.448420547 | 0.653849716 | 0.848936869 |
| SLC38A6 | 231.3996703 | 0.256582177 | 0.343916413 | 0.746059704 | 0.45563135 | 0.735306344 |
| SLC38A7 | 1191.459271 | -0.850360597 | 0.299022143 | -2.843804771 | 0.004457835 | 0.069937766 |
| SLC38A9 | 843.4529057 | -0.372741219 | 0.280437874 | -1.329140083 | 0.183801758 | 0.488175154 |
| SLC39A1 | 3908.000229 | 0.364709019 | 0.25379608 | 1.437015966 | 0.150713457 | 0.445603028 |
| SLC39A10 | 3054.082872 | 0.316154842 | 0.34273068 | 0.922458538 | 0.356289448 | 0.663793099 |
| SLC39A11 | 275.0397486 | -0.482219007 | 0.3494105 | -1.380093063 | 0.167557993 | 0.469844927 |
| SLC39A13 | 477.0291786 | 0.424595365 | 0.409276468 | 1.037429215 | 0.299535869 | 0.610246278 |
| SLC39A14 | 2108.362068 | -0.247806945 | 0.287945541 | -0.860603516 | 0.389456449 | 0.68927249 |
| SLC39A2 | 41.22205636 | 0.431024115 | 1.040149908 | 0.414386534 | 0.678591049 | 0.861138002 |
| SLC39A3 | 326.14646 | 0.329495225 | 0.289810333 | 1.136934013 | 0.255565872 | 0.567910953 |
| SLC39A4 | 42.40610843 | -1.04868169 | 0.819884569 | -1.279060163 | 0.200875871 | 0.506881795 |
| SLC39A6 | 1954.374408 | 0.661181301 | 0.323021569 | 2.046864249 | 0.040671415 | 0.23195346 |
| SLC39A7 | 312.3161053 | 0.565836277 | 0.428434368 | 1.320707019 | 0.186599073 | 0.49118221 |
| SLC39A8 | 489.8292807 | 0.466431568 | 0.251400394 | 1.855333483 | 0.06354862 | 0.290809691 |
| SLC39A9 | 1388.870336 | -0.015116068 | 0.239586288 | -0.063092375 | 0.949692946 | 0.981628635 |
| SLC3A2 | 5777.613229 | -0.377990372 | 0.393852163 | -0.959726536 | 0.337192864 | 0.64481093 |
| SLC40A1 | 293.3880945 | -2.563791521 | 0.956042464 | -2.681671179 | 0.007325543 | 0.094825773 |
| SLC41A1 | 755.4531713 | 0.392503217 | 0.395071246 | 0.993499834 | 0.320466432 | 0.63024811 |
| SLC41A2 | 17.47747017 | -0.435078036 | 0.883258656 | -0.492582816 | 0.622307392 | 0.833640266 |
| SLC41A3 | 1297.531004 | 0.280874447 | 0.227360255 | 1.235371797 | 0.216692164 | 0.524312093 |
| SLC43A1 | 67.63447511 | 1.064390576 | 0.529039689 | 2.011929536 | 0.044227365 | |

| | | | | | | |
|---|---|---|---|---|---|---|
| | | | | | | 0.241762065 |
| SLC43A2 | 92.47223729 | 0.412844885 | 0.444754834 | 0.928252721 | 0.353276492 | 0.660450729 |
| SLC43A3 | 262.2605564 | 0.200415471 | 0.37015337 | 0.541439002 | 0.588205031 | 0.817334541 |
| SLC44A1 | 2012.389909 | -0.270542941 | 0.265711423 | -1.018183327 | 0.308590842 | 0.618282047 |
| SLC44A2 | 10219.31136 | -0.90699824 | 0.348420562 | -2.603170821 | 0.009236592 | 0.108795292 |
| SLC44A3 | 134.3401407 | -2.018448876 | 0.766262876 | -2.634146765 | 0.008434902 | 0.103784183 |
| SLC44A3-AS1 | 185.0796502 | 0.807628071 | 0.842525133 | 0.958580391 | 0.337770175 | 0.645135506 |
| SLC44A5 | 334.7293605 | -1.967723483 | 0.322604328 | -6.099494985 | 1.06E-09 | 1.47E-06 |
| SLC45A1 | 21.42873156 | 1.759457596 | 0.804850487 | 2.18606763 | 0.028810658 | 0.194174362 |
| SLC45A3 | 227.8204375 | 1.419163442 | 0.581905163 | 2.438822564 | 0.014735201 | 0.138131624 |
| SLC45A4 | 338.6528249 | -0.265597755 | 0.450983631 | -0.588929922 | 0.555908284 | 0.796719574 |
| SLC46A1 | 135.9103361 | 0.066410397 | 0.447533492 | 0.148392015 | 0.882033402 | 0.953217539 |
| SLC46A3 | 29.60730143 | -0.705029969 | 0.595659368 | -1.18361266 | 0.236566427 | 0.547336452 |
| SLC47A1 | 45.75570326 | 0.56816118 | 0.697668894 | 0.814370807 | 0.415432558 | 0.707513171 |
| SLC48A1 | 315.9853614 | -0.075863846 | 0.295112013 | -0.25706797 | 0.797126303 | 0.915732089 |
| SLC49A3 | 72.32083461 | -2.01716458 | 0.646253337 | -3.121321726 | 0.001800412 | 0.039360945 |
| SLC49A4 | 621.8482782 | -0.170577053 | 0.376467903 | -0.453098528 | 0.650477795 | 0.847509329 |
| SLC4A11 | 174.4238291 | -0.197122579 | 0.37052653 | -0.532006655 | 0.594721387 | 0.820937865 |
| SLC4A1AP | 875.4537142 | -0.119787105 | 0.272130925 | -0.440181889 | 0.659805376 | 0.852124573 |
| SLC4A2 | 1892.833353 | -0.45406625 | 0.378291213 | -1.200308742 | 0.230019456 | 0.53893445 |
| SLC4A3 | 284.3705794 | -1.416734604 | 0.449235736 | -3.153655174 | 0.001612395 | 0.037175747 |
| SLC4A5 | 49.88767905 | -1.222703809 | 0.495208487 | -2.469068767 | 0.013546518 | 0.132400677 |
| SLC4A7 | 1255.584618 | 0.517305082 | 0.441924809 | 1.170572622 | 0.241770607 | 0.552977821 |
| SLC4A8 | 498.2114962 | 0.481130795 | 0.613090627 | 0.784762926 | 0.432592581 | 0.718639059 |
| SLC50A1 | 706.4510819 | 0.223310616 | 0.323564325 | 0.690158337 | 0.49009462 | 0.758505823 |
| SLC52A1 | 16.70467368 | -0.514771152 | 0.910718536 | -0.565236274 | 0.571913067 | 0.807351345 |
| SLC52A2 | 303.6978571 | 0.159908509 | 0.416658686 | 0.383787773 | 0.701135751 | 0.872239782 |
| SLC52A3 | 49.01283292 | -0.70319212 | 0.531582861 | -1.322826922 | 0.185892942 | 0.490427539 |
| SLC5A12 | 202.0806101 | -0.192149925 | 0.422914491 | -0.454346988 | 0.649579101 | 0.847284373 |
| SLC5A3 | 2103.66485 | 0.445734897 | 0.407925557 | 1.092686861 | 0.27453131 | 0.587352209 |
| SLC5A5 | 42.87000629 | 0.014424534 | 0.757482585 | 0.019042727 | 0.984807021 | 0.994433294 |
| SLC5A6 | 3042.371542 | -1.06674918 | 0.398118341 | -2.67947761 | 0.007373713 | 0.095205623 |
| SLC5A7 | 79.80058784 | -5.019302758 | 1.191095802 | -4.214021029 | 2.51E-05 | |

| | | | | | | |
|---|---|---|---|---|---|---|
| | | | | | | 0.002159569 |
| SLC6A1 | 36.01650086 | 1.539050888 | 0.644600059 | 2.387605874 | 0.016958517 | 0.147606234 |
| SLC6A10P | 17.86072974 | 1.495536598 | 1.409196487 | 1.061269036 | 0.288567653 | 0.601816473 |
| SLC6A11 | 13.70102506 | -0.077351261 | 0.795381754 | -0.097250484 | 0.922527477 | 0.970775082 |
| SLC6A12 | 41.29356598 | 0.75771764 | 0.622519354 | 1.217179248 | 0.223536024 | 0.53268495 |
| SLC6A15 | 172.2794105 | 0.873808551 | 0.59918876 | 1.458319329 | 0.144752545 | 0.435928729 |
| SLC6A16 | 81.23489617 | 0.296484083 | 0.47901706 | 0.618942639 | 0.535954147 | 0.784841735 |
| SLC6A20 | 23.03621655 | -1.584954058 | 0.840877091 | -1.884881958 | 0.059445776 | 0.282486927 |
| SLC6A4 | 67.42517877 | -1.098900823 | 0.743386397 | -1.478236389 | 0.139344517 | 0.42802099 |
| SLC6A6 | 2780.33954 | -0.812368055 | 0.329596755 | -2.46473317 | 0.013711532 | 0.133173911 |
| SLC6A8 | 3129.625487 | 1.193501494 | 0.532704373 | 2.240457473 | 0.025061239 | 0.181135804 |
| SLC6A9 | 304.4024133 | 0.638054428 | 0.57823501 | 1.103451741 | 0.269831038 | 0.58255396 |
| SLC7A1 | 1183.901747 | 0.501524959 | 0.472371451 | 1.061717337 | 0.288364026 | 0.601673803 |
| SLC7A10 | 17.18440366 | 0.566379409 | 0.888307324 | 0.637593988 | 0.52373801 | 0.777662198 |
| SLC7A11 | 3097.593979 | -0.662071783 | 0.49923679 | -1.326167855 | 0.184784104 | 0.489365432 |
| SLC7A14 | 12.79434639 | -0.504692699 | 0.854794902 | -0.59042549 | 0.554905426 | 0.796360966 |
| SLC7A2 | 2383.729714 | -2.17897568 | 0.627563172 | -3.472121654 | 0.000516362 | 0.017892193 |
| SLC7A3 | 914.8810962 | 0.123378851 | 0.368338769 | 0.334960263 | 0.737655062 | 0.891065064 |
| SLC7A4 | 20.03987953 | -1.944623777 | 1.020124549 | -1.906261132 | 0.056616334 | 0.275490291 |
| SLC7A5 | 4686.103549 | -1.762831239 | 0.42272486 | -4.17016222 | 3.04E-05 | 0.00250967 |
| SLC7A6 | 737.5935163 | 0.068530244 | 0.411491505 | 0.16654109 | 0.86773115 | 0.946530722 |
| SLC7A6OS | 738.1456376 | -0.001161799 | 0.266589916 | -0.004358 | 0.99652283 | 0.998431718 |
| SLC7A7 | 70.585175 | -0.057796262 | 0.580423665 | -0.099575992 | 0.920680955 | 0.969843271 |
| SLC7A8 | 3972.820155 | -0.988617977 | 0.304988448 | -3.241493194 | 0.001189053 | 0.030317847 |
| SLC8A1 | 38.48386691 | -3.643778339 | 1.039103442 | -3.506656017 | 0.000453775 | 0.016498831 |
| SLC8A2 | 184.9417198 | 0.55491659 | 0.417686022 | 1.328549581 | 0.183996616 | 0.488493979 |
| SLC8B1 | 242.4506712 | -0.135382999 | 0.312434305 | -0.433316689 | 0.664784715 | 0.855037526 |
| SLC9A1 | 919.9432206 | -1.015687493 | 0.293609154 | -3.459318213 | 0.000541545 | 0.018350186 |
| SLC9A3 | 199.3342787 | 0.607324671 | 0.309694732 | 1.961042953 | 0.049874012 | 0.257360083 |
| SLC9A3R1 | 2706.425263 | -0.481127477 | 0.450720996 | -1.06746187 | 0.285763323 | 0.59971163 |
| SLC9A3R2 | 162.6602367 | 0.790657292 | 0.505906215 | 1.562853485 | 0.118087061 | 0.395958766 |
| SLC9A5 | 84.49276482 | 0.546258984 | 0.464024498 | 1.177220139 | 0.239107652 | 0.550288561 |
| SLC9A6 | 577.6031289 | -0.138322507 | 0.248320549 | -0.557032058 | | |

| | | | | | | |
|---|---|---|---|---|---|---|
| | | | | | 0.577505526 | 0.809916636 |
| SLC9A7 | 331.2238329 | -1.231331418 | 0.54581063 | -2.255968189 | 0.024072623 | 0.177207433 |
| SLC9A7P1 | 31.16682779 | -1.616363388 | 0.8642103 | -1.870335713 | 0.061437213 | 0.285640507 |
| SLC9A8 | 383.8284536 | -0.842372562 | 0.318311635 | -2.646376913 | 0.008135911 | 0.10133818 |
| SLC9A9 | 13.84092662 | -1.68175358 | 0.887514358 | -1.894902955 | 0.05810526 | 0.279402502 |
| SLC9B1 | 21.30627972 | 0.474922699 | 0.631994964 | 0.751465956 | 0.452372282 | 0.732985143 |
| SLC9B2 | 204.8830824 | 0.494263866 | 0.363833142 | 1.358490497 | 0.174308096 | 0.478542911 |
| SLCO1A2 | 707.8581931 | 0.457509244 | 0.36524572 | 1.252606721 | 0.210348868 | 0.51816316 |
| SLCO1B1 | 24.40563931 | 1.201360839 | 0.669491333 | 1.794438225 | 0.07274325 | 0.310433977 |
| SLCO2A1 | 59.27750675 | -3.414194058 | 1.030902902 | -3.311848334 | 0.000926818 | 0.02597297 |
| SLCO3A1 | 72.49027788 | 0.846433984 | 0.622913958 | 1.358829696 | 0.174200561 | 0.478455902 |
| SLCO4A1 | 11.5291751 | 0.793178466 | 0.901855451 | 0.879496227 | 0.379132278 | 0.681253679 |
| SLCO4C1 | 235.6076303 | -2.074742034 | 0.402209231 | -5.158365033 | 2.49E-07 | 7.56E-05 |
| SLCO5A1 | 55.65499842 | 1.675169422 | 0.673168428 | 2.488484833 | 0.012828871 | 0.128084711 |
| SLCO6A1 | 20.66396649 | -0.344761898 | 0.725441534 | -0.475244223 | 0.63461291 | 0.840313679 |
| SLF1 | 606.5305706 | 0.652566703 | 0.316145977 | 2.064130973 | 0.0390053 | 0.227508422 |
| SLF2 | 761.9570179 | -0.301634428 | 0.440930544 | -0.684086036 | 0.493920839 | 0.759966841 |
| SLFN11 | 11.97337965 | -1.072801017 | 1.10978087 | -0.966678239 | 0.333704909 | 0.641733702 |
| SLFN12 | 356.1473275 | -0.311336831 | 0.364062609 | -0.855173873 | 0.392454898 | 0.691674403 |
| SLFN12L | 15.51269493 | -0.359853187 | 0.829607122 | -0.433763377 | 0.664460279 | 0.85500751 |
| SLFN13 | 355.4553225 | -1.005823124 | 0.373623856 | -2.692074152 | 0.007100916 | 0.092709113 |
| SLFN5 | 700.2669202 | -1.191557852 | 0.306429447 | -3.888522674 | 0.000100856 | 0.006194697 |
| SLIRP | 1826.008744 | 1.272223959 | 0.41205703 | 3.08749485 | 0.002018513 | 0.042355264 |
| SLIT1 | 728.9773203 | -1.336217103 | 0.595784806 | -2.242784793 | 0.024910696 | 0.180563862 |
| SLIT2 | 388.1465467 | -1.035779513 | 0.298322114 | -3.472017211 | 0.000516563 | 0.017892193 |
| SLIT3 | 300.440679 | -1.020058124 | 0.405307317 | -2.516752305 | 0.011844206 | 0.123582156 |
| SLITRK1 | 12.77296847 | 1.775565975 | 0.893174486 | 1.987927335 | 0.046819727 | 0.249225774 |
| SLITRK3 | 46.3741316 | 1.769537348 | 0.896075335 | 1.974764038 | 0.048294932 | 0.253347998 |
| SLITRK4 | 76.65369427 | 5.014751164 | 0.617409392 | 8.122246322 | 4.58E-16 | 3.47E-12 |
| SLITRK5 | 578.8954199 | 0.978673483 | 0.553795089 | 1.767212283 | 0.077192683 | 0.31844659 |
| SLITRK6 | 246.8924543 | -2.055242518 | 0.888577931 | -2.31295697 | 0.020725005 | 0.163165051 |
| SLK | 1578.591356 | -0.01362164 | 0.305995807 | -0.044515772 | 0.96449328 | 0.988532608 |
| SLMAP | 959.1183664 | -0.119455798 | 0.312376598 | -0.382409563 | 0.702157598 | 0.872649539 |
| SLTM | 3506.87246 | 0.03855956 | 0.250841749 | 0.153720665 | 0.87782999 | 0.951077249 |
| SLU7 | 1018.592819 | -0.124186359 | 0.364680479 | -0.340534703 | 0.733453894 | |

| | | | | | | |
|---|---|---|---|---|---|---|
| | | | | | | 0.888472455 |
| SLX1A | 479.3976093 | 0.697544526 | 0.519889084 | 1.34171797 | 0.179687458 | 0.484443366 |
| SLX1B | 567.8725519 | -0.27994939 | 0.396881509 | -0.705372722 | 0.480578315 | 0.752509463 |
| SLX4 | 405.224222 | -0.210576231 | 0.666326743 | -0.316025484 | 0.751983165 | 0.896586461 |
| SLX4IP | 403.8371081 | -0.487846693 | 0.403534199 | -1.20893519 | 0.226687745 | 0.535481989 |
| SMAD1 | 664.9302766 | -0.23224495 | 0.246311499 | -0.942891225 | 0.345736544 | 0.652517297 |
| SMAD2 | 2186.908943 | -0.160327216 | 0.299270926 | -0.535725999 | 0.592147932 | 0.819765854 |
| SMAD3 | 874.4570374 | -0.196198048 | 0.470823232 | -0.41671276 | 0.67688852 | 0.86029961 |
| SMAD4 | 1945.124568 | 0.260124108 | 0.253528673 | 1.026014553 | 0.304884724 | 0.615135352 |
| SMAD5 | 2806.497721 | 0.117351768 | 0.273109313 | 0.429687904 | 0.667422683 | 0.856540807 |
| SMAD6 | 386.0894221 | -0.835109511 | 0.360497019 | -2.316550394 | 0.020528236 | 0.161952092 |
| SMAD7 | 251.7605125 | -1.587752754 | 0.523301558 | -3.034106681 | 0.002412491 | 0.047225679 |
| SMAD9 | 44.51734839 | -0.499092549 | 0.645667608 | -0.772986817 | 0.439530183 | 0.723144171 |
| SMAGP | 2015.310448 | 0.16904761 | 0.662496893 | 0.255167401 | 0.798593808 | 0.916056735 |
| SMAP1 | 1449.115844 | -0.207293691 | 0.353121169 | -0.587032749 | 0.557181712 | 0.798206209 |
| SMAP2 | 208.0145578 | 0.210677544 | 0.423778521 | 0.497140686 | 0.619089845 | 0.832495306 |
| SMARCA1 | 4086.874607 | 0.058105752 | 0.242029748 | 0.240076903 | 0.810270639 | 0.920450458 |
| SMARCA2 | 1220.406757 | -3.859356398 | 0.725493235 | -5.319631134 | 1.04E-07 | 3.96E-05 |
| SMARCA4 | 8225.239358 | -0.095918553 | 0.258122111 | -0.371601459 | 0.710189602 | 0.876599663 |
| SMARCA5 | 8332.142418 | 0.003286992 | 0.261974299 | 0.012547 | 0.989989205 | 0.996425809 |
| SMARCAD1 | 2443.307703 | -0.307476963 | 0.349446185 | -0.879897897 | 0.378914624 | 0.681253679 |
| SMARCAL1 | 872.2238438 | -0.450297354 | 0.306723533 | -1.468088705 | 0.142080129 | 0.432308365 |
| SMARCB1 | 359.2856704 | 1.553197855 | 0.621193717 | 2.500343794 | 0.012407284 | 0.126591155 |
| SMARCC1 | 9120.863298 | -0.08075252 | 0.303034448 | -0.266479672 | 0.789869813 | 0.912949942 |
| SMARCC2 | 2859.849528 | -0.374624597 | 0.325709132 | -1.150181436 | 0.250069151 | 0.562794702 |
| SMARCD1 | 5054.148596 | -0.200220109 | 0.256852395 | -0.779514276 | 0.435676831 | 0.720633799 |
| SMARCD2 | 3293.044404 | -0.610870705 | 0.325679179 | -1.875682402 | 0.060698904 | 0.284105089 |
| SMARCD3 | 211.0429776 | 0.810515271 | 0.378395649 | 2.141978307 | 0.032195228 | 0.204582474 |
| SMARCE1 | 6060.407751 | -0.176778373 | 0.225492623 | -0.783965218 | 0.43306052 | 0.718657207 |
| SMBD1P | 10.64648466 | 0.324752184 | 0.922515961 | 0.352028797 | 0.724816666 | 0.883935898 |
| SMC1A | 4346.317117 | -0.108579813 | 0.384889382 | -0.282106543 | 0.777861817 | 0.908018171 |
| SMC2 | 1668.952355 | 0.155684011 | 0.336527983 | 0.462618322 | 0.643637973 | 0.845466381 |
| SMC3 | 3994.991957 | 0.473118847 | 0.297999226 | 1.587651259 | 0.11236522 | 0.385502659 |
| SMC4 | 5899.276462 | 0.686369514 | 0.271788029 | 2.525385376 | 0.011557148 | 0.121674869 |
| SMC5 | 1303.96498 | -0.088008512 | 0.294525506 | -0.298814568 | 0.765081535 | 0.902991439 |
| SMC6 | 2050.703813 | 0.208230226 | 0.277445369 | 0.750526948 | 0.4529374 | 0.733430813 |
| SMCHD1 | 1836.362406 | -0.179647783 | 0.417353464 | -0.43044517 | 0.666871843 | 0.856258128 |

SMCO4 236.2678951 1.17935396 0.384075832 3.070627883 0.002136092 0.04379277
SMCR8 537.3584154 -0.752086842 0.422067499 -1.781911289 0.074763699 0.313842348
SMDT1 125.3962671 0.729899278 0.521667119 1.39916673 0.161762991 0.462342815
SMG1 3678.041131 -0.561520239 0.446264034 -1.258269088 0.208294457 0.515167134
SMG1P1 205.2138458 -0.756834014 0.709364553 -1.066918287 0.286008736 0.599894724
SMG1P2 183.563117 -0.181448652 0.368856358 -0.491922256 0.622774304 0.833787151
SMG1P3 440.4194757 -0.667224868 0.519499655 -1.28436056 0.199015826 0.504810082
SMG1P4 105.2897901 -1.039331408 0.664237795 -1.564697786 0.117653788 0.395641036
SMG1P5 560.1475717 0.205671266 0.291890807 0.704617141 0.481048528 0.75260258
SMG1P7 120.8810457 0.401552791 0.412926048 0.972456915 0.330823289 0.639084337
SMG5 3298.374414 -0.563437305 0.407072866 -1.38411904 0.166322008 0.468062511
SMG6 1546.224533 -0.501447921 0.362055866 -1.385001511 0.166052005 0.468031848
SMG7 3329.910746 -0.503817089 0.254888554 -1.976617155 0.048084922 0.252967897
SMG8 801.2941353 -0.333847162 0.302015802 -1.10539634 0.268987889 0.582058945
SMG9 602.5866387 0.083506731 0.293459647 0.284559502 0.775981642 0.907315413
SMIM1 25.50214854 0.315902133 0.901255252 0.3505135 0.72595336 0.884471445
SMIM10 22.80362589 0.233928523 0.675190764 0.34646286 0.728994891 0.886086417
SMIM10L1 793.5984369 0.228810328 0.388003338 0.589712267 0.55538357 0.796360966
SMIM10L2A 392.8706245 -1.971421608 0.373824466 -5.273655917 1.34E-07 4.72E-05
SMIM10L2B 100.0465508 -0.769918431 0.390171334 -1.973282926 0.048463337 0.253705068
SMIM11A 159.4385224 0.470022578 0.561184416 0.837554579 0.402280912 0.699404505
SMIM11B 171.3763589 1.365775186 1.979052585 0.690115663 0.490121454 0.758505823
SMIM12 714.9069836 0.267461354 0.387828535 0.689638152 0.490421769 0.758593654
SMIM13 397.4002338 0.008337416 0.298762478 0.027906503 0.977736722 0.99226352
SMIM14 795.9693381 0.300823916 0.432751678 0.695142113 0.486966245 0.756439734
SMIM15 1093.727844 0.457590218 0.312275025 1.465343624 0.14282719 0.433233718
SMIM17 27.05889938 0.547287166 0.676961251 0.808446813 0.418833413 0.709799118
SMIM19 871.9176324 0.979880242 0.379757928 2.580275934 0.00987214 0.112863778
SMIM20 959.7056498 1.235118516 0.394901889 3.127659171 0.001762044 0.039027635
SMIM24 18.0564795 1.285237321 0.939711854 1.367692996 0.171408193 0.474791619
SMIM26 1211.749631 1.295531224 0.492553308 2.630235556 0.008532572 0.10430915
SMIM27 100.9638575 1.118027924 0.582985416 1.91776311 0.055141055 0.271793713
SMIM29 101.2067134 0.841172882 0.45940072 1.831022122 0.067097241 0.297817842
SMIM3 511.8183045 0.486865709 0.392474363 1.240503215 0.214789326 0.522039228
SMIM30 1200.832627 0.853623755 0.384621451 2.219386755 0.026460424

0.185860119
SMIM32 18.63598577 0.590809893 0.763145447 0.774177314 0.438825942 0.722407379
SMIM4 210.2876871 1.341755292 0.454977428 2.949059029 0.003187431 0.056491255
SMIM7 1389.743864 0.328483837 0.386717126 0.849416319 0.395649676 0.694320559
SMIM8 392.4036402 1.013474136 0.303517265 3.339098798 0.000840507 0.024381121
SMN1 57.33435134 -0.143036627 0.449004833 -0.31856367 0.750057407 0.895752516
SMNDC1 1499.160553 0.343508577 0.305879437 1.123019516 0.261429206 0.574937735
SMO 2436.460217 0.07371837 0.426723995 0.172754217 0.862844625 0.944730333
SMOC1 1112.405813 1.139061245 1.48437965 0.767365172 0.44286443 0.726188529
SMOC2 31.42804204 0.607062166 0.786217201 0.772130354 0.440037226 0.7234571
SMOX 259.2164127 -0.146431184 0.342954289 -0.426969975 0.669401197 0.857712777
SMPD1 302.0515787 -0.08946434 0.285050886 -0.313853927 0.753631984 0.896944444
SMPD2 237.8305446 0.409778235 0.354806107 1.154935689 0.248116763 0.56079644
SMPD3 12.49664344 0.426627057 0.932154623 0.457678422 0.647183492 0.846597949
SMPD4 1131.673958 -0.356085373 0.280480091 -1.269556677 0.20424259 0.510640133
SMPDL3A 110.5578297 -1.531035267 0.818105875 -1.871439032 0.061284253 0.285640507
SMPDL3B 310.4560042 -0.66076056 0.311033817 -2.124401022 0.033636625 0.209054171
SMPX 46.22351951 -3.262052045 1.196166514 -2.727088585 0.006389588 0.086705227
SMS 4051.022813 0.014195705 0.509884021 0.027841047 0.977788928 0.99226352
SMTN 249.1381082 0.461406212 0.380414693 1.212903235 0.225166819 0.534502552
SMTNL2 1237.706785 -0.616306446 0.687916228 -0.895903339 0.370304391 0.675958117
SMU1 2077.272413 0.270305377 0.272252053 0.992849731 0.32078319 0.630470499
SMUG1 667.7194167 -0.227373413 0.357268225 -0.636422153 0.524501306 0.777662198
SMURF1 659.1062076 -0.507330998 0.405562556 -1.250931551 0.210959451 0.518294063
SMURF2 785.0008556 -0.06257866 0.283637401 -0.220629084 0.825381253 0.927133052
SMURF2P1 27.55301359 -0.791150806 0.87525614 -0.903907747 0.366044321 0.672491566
SMYD2 677.5254433 -0.470613547 0.30326874 -1.551803677 0.120709206 0.400973879
SMYD3 559.376212 0.384375342 0.313650734 1.225488419 0.220391251 0.528959922
SMYD4 736.9240756 -0.211750128 0.280327695 -0.755366423 0.450029154 0.731375715
SMYD5 842.0984712 0.122316315 0.388247221 0.315047496 0.752725592 0.896586461
SNAI1 226.9509026 -0.29472755 0.467116179 -0.630951279 0.528072381 0.779319497
SNAI2 82.2143038 0.235442069 0.631717341 0.372701608 0.70937054 0.875945015
SNAI3 28.70918809 -0.016901403 0.707710895 -0.02388179 0.9809469 0.993241851
SNAP23 2241.801682 -0.183321892 0.295017546 -0.621393182 0.534340957 0.783689739
SNAP29 635.3618094 -0.101975564 0.29253701 -0.3485903 0.727396911 0.885092921
SNAP47 781.2602653 0.084000195 0.250471493 0.335368286 0.737347288 0.89093937
SNAP91 33.16298473 0.715264648 0.727241208 0.983531516 0.325345912 0.635241033
SNAPC1 428.476911 1.236955055 0.284186087 4.352623551 1.35E-05 0.001389524
SNAPC2 412.8073722 0.440931069 0.333338168 1.322774023 0.185910538 0.490427539
SNAPC3 1241.975805 0.319722895 0.281530925 1.13565817 0.256099654

| | | | | | | |
|---|---|---|---|---|---|---|
| | | | | | | 0.568292977 |
| SNAPC4 | 487.5390921 | -0.134855574 | 0.515665236 | -0.261517676 | 0.79369332 | 0.914422947 |
| SNAPC5 | 506.1259125 | 0.542983697 | 0.365017857 | 1.48755379 | 0.136868608 | 0.424728769 |
| SNAPIN | 474.6874782 | 0.276365553 | 0.319022463 | 0.866288697 | 0.386331863 | 0.686679504 |
| SNCA | 364.6456729 | 0.645683313 | 0.38835858 | 1.662595719 | 0.096393389 | 0.35626747 |
| SNCAIP | 125.4377838 | 0.656715827 | 0.606163251 | 1.083397625 | 0.27863197 | 0.592117329 |
| SNCB | 12.353901 | 1.451915195 | 1.001381781 | 1.449911735 | 0.147083135 | 0.44029168 |
| SNCG | 38.202373 | 1.493773028 | 0.570594529 | 2.617923852 | 0.008846654 | 0.106517932 |
| SND1 | 7980.891395 | -0.532607623 | 0.261059216 | -2.040179359 | 0.041332465 | 0.233539972 |
| SNED1 | 11.00760378 | -1.343387102 | 0.97143174 | -1.382893978 | 0.166697377 | 0.468674186 |
| SNF8 | 1617.372884 | 0.228485889 | 0.414054607 | 0.551825497 | 0.581067917 | 0.812552435 |
| SNIP1 | 562.1041583 | -0.10496786 | 0.287359899 | -0.365283606 | 0.714899715 | 0.87855355 |
| SNN | 2395.688437 | -0.091097015 | 0.226996788 | -0.401314114 | 0.688188873 | 0.866925917 |
| SNORC | 15.05424126 | 0.146785906 | 0.774588476 | 0.189501795 | 0.849699549 | 0.938580023 |
| SNPH | 31.46054888 | -0.219562605 | 0.817796115 | -0.268480861 | 0.788329205 | 0.912309638 |
| SNRK | 605.1289281 | -0.41665439 | 0.283018368 | -1.472181444 | 0.140971887 | 0.430713136 |
| SNRNP200 | 11037.49729 | -0.536181178 | 0.451186149 | -1.188381289 | 0.234683225 | 0.545067238 |
| SNRNP25 | 1054.616082 | 0.953716038 | 0.307496424 | 3.101551639 | 0.001925093 | 0.041048886 |
| SNRNP27 | 1111.344607 | 0.577048868 | 0.374433722 | 1.541124193 | 0.123286563 | 0.404753154 |
| SNRNP35 | 427.6258849 | 0.214559302 | 0.402873144 | 0.532572858 | 0.594329295 | 0.820642877 |
| SNRNP40 | 2295.879198 | 0.45050989 | 0.222824245 | 2.021817194 | 0.043195241 | 0.239078803 |
| SNRNP48 | 1048.158152 | 0.354583459 | 0.305697614 | 1.159915689 | 0.246083135 | 0.557962522 |
| SNRNP70 | 5473.04756 | 0.197470159 | 0.291094794 | 0.678370629 | 0.497536727 | 0.761898625 |
| SNRPA | 3416.946617 | 0.430776292 | 0.329757798 | 1.306341487 | 0.191436455 | 0.496966715 |
| SNRPA1 | 2047.649338 | 0.662159926 | 0.34296179 | 1.930710492 | 0.053518865 | 0.267406937 |
| SNRPB | 6026.441839 | 0.865958713 | 0.372145993 | 2.326932789 | 0.019968843 | 0.159698109 |
| SNRPB2 | 2761.127834 | 0.843567904 | 0.415983615 | 2.027887333 | 0.042571748 | 0.237272591 |
| SNRPC | 2841.431387 | 0.792679658 | 0.357127475 | 2.219598642 | 0.026446025 | 0.185860119 |
| SNRPD1 | 3212.666366 | 1.129416425 | 0.370754666 | 3.046263554 | 0.002317046 | 0.04595021 |
| SNRPD2 | 7965.548654 | 1.072716742 | 0.425385184 | 2.521753891 | 0.011677137 | 0.122513037 |
| SNRPD3 | 2426.623604 | 0.320842465 | 0.342894645 | 0.935688176 | 0.34943376 | 0.656038342 |
| SNRPE | 4456.457142 | 1.368484018 | 0.419394741 | 3.262997563 | 0.001102405 | 0.028736394 |
| SNRPF | 2919.113259 | 1.289472471 | 0.458432519 | 2.812785783 | 0.004911436 | 0.074885822 |
| SNRPG | 2831.584311 | 1.244113916 | 0.346740756 | 3.588023309 | 0.000333195 | 0.013525316 |
| SNRPN | 6885.415071 | 0.459963844 | 0.323408696 | 1.422237096 | 0.154957433 | 0.452697712 |
| SNTA1 | 90.63689763 | 0.828349897 | 0.467612265 | 1.771446045 | 0.076486557 | 0.317498731 |
| SNTB1 | 50.88480211 | -0.196497918 | 0.58596858 | -0.335338659 | 0.737369634 | 0.89093937 |
| SNTB2 | 522.3772191 | 0.164924181 | 0.3354218 | 0.491691897 | 0.622937167 | 0.833825637 |
| SNTG2 | 12.51356121 | 2.109648821 | 1.046968168 | 2.015007606 | 0.043903852 | 0.241112355 |
| SNU13 | 2697.80791 | 0.514967956 | 0.330536145 | 1.557977741 | 0.119238525 | 0.398100277 |
| SNUPN | 473.1078013 | 0.296413171 | 0.322584847 | 0.918868861 | 0.358164169 | 0.665814068 |

| | | | | | | |
|---|---|---|---|---|---|---|
| SNURF | 65.71053627 | -0.542515196 | 0.58599433 | -0.925802808 | 0.354548468 | 0.661914443 |
| SNW1 | 2293.720973 | 0.06099631 | 0.224425686 | 0.271788453 | 0.785784684 | 0.910984215 |
| SNX1 | 1676.578441 | -0.14700972 | 0.285565649 | -0.514801833 | 0.606691505 | 0.826667578 |
| SNX10 | 465.4164012 | -0.315805039 | 0.334010826 | -0.945493421 | 0.344407028 | 0.651368096 |
| SNX11 | 601.5881204 | 0.285187562 | 0.258693639 | 1.102414279 | 0.270281607 | 0.583170438 |
| SNX12 | 2431.231884 | -0.195478811 | 0.233816296 | -0.836035873 | 0.403134722 | 0.699766232 |
| SNX13 | 1152.726849 | -0.28115441 | 0.276265607 | -1.017696023 | 0.308822438 | 0.618441002 |
| SNX14 | 1179.840636 | -0.467110492 | 0.339566362 | -1.375608848 | 0.168942766 | 0.471508612 |
| SNX15 | 586.7571264 | -0.15931382 | 0.336545803 | -0.47337931 | 0.635942588 | 0.841016203 |
| SNX16 | 334.5387753 | 0.425493503 | 0.278771334 | 1.526317275 | 0.126930868 | 0.41024035 |
| SNX17 | 1907.246895 | 0.245533029 | 0.292670427 | 0.838940343 | 0.401502787 | 0.698932735 |
| SNX18 | 902.9750037 | -0.403034637 | 0.314354984 | -1.282100359 | 0.199807443 | 0.505887637 |
| SNX18P3 | 50.13904305 | -0.169890603 | 0.594935134 | -0.285561556 | 0.775213952 | 0.906557532 |
| SNX19 | 1032.593915 | -0.420586361 | 0.325293502 | -1.292944242 | 0.196030344 | 0.501797297 |
| SNX2 | 3717.603597 | -0.367714383 | 0.237766769 | -1.546533961 | 0.121975665 | 0.402659864 |
| SNX21 | 291.8223094 | -0.114105896 | 0.305719072 | -0.37323774 | 0.70897151 | 0.875666106 |
| SNX22 | 99.51731961 | 0.426309775 | 0.452765476 | 0.941568644 | 0.34641353 | 0.653093035 |
| SNX24 | 781.3103385 | -0.747063619 | 0.395525654 | -1.888786764 | 0.058920407 | 0.281360244 |
| SNX25 | 522.0273118 | 0.286393175 | 0.32858479 | 0.871595959 | 0.383428836 | 0.68451387 |
| SNX25P1 | 32.1588618 | 0.883166025 | 0.630632114 | 1.400445688 | 0.161379897 | 0.46167973 |
| SNX27 | 1084.752493 | -0.251268226 | 0.328249646 | -0.765479046 | 0.443986339 | 0.727127105 |
| SNX29 | 380.1760599 | -0.51408046 | 0.302752565 | -1.698021815 | 0.089503645 | 0.342777615 |
| SNX3 | 2573.734983 | 0.311893612 | 0.372998854 | 0.836178472 | 0.403054508 | 0.699766232 |
| SNX30 | 1070.477858 | 0.130158873 | 0.293282224 | 0.443800757 | 0.657186641 | 0.850335056 |
| SNX32 | 81.46543794 | 0.807896509 | 0.558947157 | 1.445389781 | 0.148348432 | 0.442420693 |
| SNX33 | 1206.705948 | -0.666281774 | 0.349461166 | -1.906597471 | 0.056572733 | 0.275490291 |
| SNX4 | 1407.020878 | 0.380378299 | 0.294257338 | 1.292672262 | 0.196124435 | 0.501797297 |
| SNX5 | 6016.47486 | 0.29036611 | 0.278876256 | 1.041200545 | 0.297782483 | 0.609341523 |
| SNX6 | 3744.82641 | 0.122661628 | 0.251112902 | 0.488472025 | 0.625215536 | 0.835033444 |
| SNX7 | 719.625849 | 0.344140851 | 0.365038052 | 0.942753364 | 0.345807071 | 0.652517297 |
| SNX8 | 952.6492046 | -0.092201403 | 0.268088709 | -0.343921245 | 0.730905514 | 0.886911603 |
| SNX9 | 755.6975559 | -0.468536091 | 0.247158418 | -1.895691411 | 0.058000862 | 0.278988926 |
| SOAT1 | 622.4802268 | 0.202627059 | 0.287329255 | 0.705208589 | 0.480680437 | 0.752563699 |
| SOBP | 92.80874446 | 0.818039686 | 0.533649894 | 1.532914546 | 0.125296911 | 0.407127745 |
| SOCS1 | 83.65407271 | 0.274072629 | 0.595839977 | 0.459976907 | 0.645532797 | 0.845794147 |
| SOCS2 | 617.0042316 | -0.493806032 | 0.658161965 | -0.750280414 | 0.453085836 | 0.733436323 |
| SOCS3 | 878.2720509 | -1.804805901 | 0.644617546 | -2.799808834 | 0.005113288 | 0.075978148 |
| SOCS4 | 1194.272043 | 0.013515715 | 0.371116814 | 0.036419032 | 0.970948239 | 0.989604013 |
| SOCS5 | 798.2709428 | 0.2352315 | 0.272152281 | 0.864337784 | 0.38740236 | 0.687545052 |
| SOCS6 | 1695.027725 | -0.870768204 | 0.293296973 | -2.968895984 | 0.002988717 | 0.054042709 |
| SOCS7 | 96.98809267 | -0.53926609 | 0.585837545 | -0.920504489 | 0.357309189 | 0.66487645 |
| SOD1 | 4276.667607 | 1.09229438 | 0.389212796 | 2.806419502 | 0.005009543 | 0.07561376 |

| | | | | | | |
|---|---|---|---|---|---|---|
| SOD2 | 1685.287187 | 0.527795566 | 0.248644223 | 2.122693859 | 0.033779514 | 0.209476397 |
| SOD3 | 19.83321818 | -0.493895305 | 0.754919173 | -0.654236006 | 0.512959768 | 0.771819984 |
| SOGA1 | 1206.65456 | -0.043004563 | 0.514640683 | -0.083562308 | 0.933404436 | 0.974750122 |
| SOGA3 | 64.64501396 | 1.561968066 | 0.682750387 | 2.287758595 | 0.022151583 | 0.169130181 |
| SOHLH2 | 115.3245634 | 0.813163554 | 0.495456021 | 1.641242652 | 0.100747055 | 0.363566502 |
| SON | 8270.891313 | -0.127345657 | 0.25846258 | -0.492704426 | 0.62222145 | 0.833640266 |
| SORBS1 | 471.0284531 | 0.583419309 | 0.4804392 | 1.214345769 | 0.224615715 | 0.533946257 |
| SORBS2 | 29.06662742 | 0.959625133 | 1.047259092 | 0.916320651 | 0.35949874 | 0.66703422 |
| SORBS3 | 1221.363857 | -0.91110758 | 0.402667452 | -2.262679972 | 0.023655424 | 0.175919824 |
| SORCS1 | 77.94246328 | -0.151830012 | 0.658432491 | -0.230593135 | 0.817630901 | 0.92398615 |
| SORCS2 | 176.6268523 | -1.178361632 | 0.389461509 | -3.025617693 | 0.002481258 | 0.048014245 |
| SORD | 948.6031742 | 0.531855352 | 0.344378034 | 1.544393948 | 0.122492925 | 0.403241325 |
| SORD2P | 454.4859386 | 0.773959651 | 0.318526172 | 2.429814941 | 0.015106534 | 0.139261407 |
| SORL1 | 1925.917866 | -0.349519265 | 0.391489067 | -0.892794447 | 0.37196726 | 0.677193964 |
| SORT1 | 2114.073737 | -0.736699021 | 0.283297319 | -2.600444735 | 0.009310301 | 0.10932398 |
| SOS1 | 1592.688061 | -1.077354959 | 0.444287695 | -2.424903884 | 0.01531244 | 0.140195951 |
| SOS2 | 591.7721392 | -0.207317572 | 0.345714767 | -0.599678091 | 0.548720792 | 0.793468239 |
| SOST | 264.4797113 | 2.180118055 | 0.920122059 | 2.369379187 | 0.017817975 | 0.150762132 |
| SOSTDC1 | 89.18544514 | -1.910945366 | 0.525482077 | -3.636556693 | 0.000276307 | 0.011709082 |
| SOWAHA | 27.79200895 | -7.414328165 | 2.020076833 | -3.670319882 | 0.000242247 | 0.010655185 |
| SOWAHB | 36.23460735 | -3.593487724 | 0.855636007 | -4.199785533 | 2.67E-05 | 0.00226436 |
| SOWAHC | 452.4002876 | -0.465023995 | 0.446904466 | -1.040544523 | 0.29808699 | 0.609375237 |
| SOWAHD | 10.40119035 | 0.417641704 | 1.172692491 | 0.356139147 | 0.721736361 | 0.882498649 |
| SOX1 | 282.4663903 | 3.636008922 | 2.072931275 | 1.754042194 | 0.079423277 | 0.322931692 |
| SOX10 | 54.75303328 | 0.38517959 | 0.679014227 | 0.567262916 | 0.570535568 | 0.806522093 |
| SOX11 | 5142.464801 | 0.74131672 | 0.407840541 | 1.817663144 | 0.069115634 | 0.303010569 |
| SOX12 | 2016.402016 | 0.382556981 | 0.394752707 | 0.969105403 | 0.332492602 | 0.640222942 |
| SOX13 | 3037.407213 | -0.489749097 | 0.232290091 | -2.108351218 | 0.035000621 | 0.213593893 |
| SOX14 | 35.51180646 | 0.412802407 | 0.929631361 | 0.444049571 | 0.657006745 | 0.85024732 |
| SOX15 | 1115.66598 | -1.078303295 | 0.552706186 | -1.9509521 | 0.051062744 | 0.259957346 |
| SOX2 | 4592.654498 | 2.181902658 | 0.55633273 | 3.921938331 | 8.78E-05 | 0.005599213 |
| SOX21 | 571.5849696 | 2.71039303 | 0.789117777 | 3.434712927 | 0.000593182 | 0.019606017 |
| SOX3 | 202.7485905 | 0.880042489 | 0.453847253 | 1.939071976 | 0.052492575 | 0.264550665 |
| SOX4 | 4983.605417 | -0.519211523 | 0.462642622 | -1.122273433 | 0.261746199 | 0.57525012 |
| SOX5 | 319.1413494 | 0.06799474 | 0.512626528 | 0.132639917 | 0.894478162 | 0.958237013 |
| SOX6 | 16.44000268 | -0.288979322 | 0.780028486 | -0.370472781 | 0.711030254 | 0.876976263 |
| SOX7 | 50.86550867 | -2.186337336 | 1.123825443 | -1.945442106 | 0.051721791 | 0.262081261 |
| SOX8 | 41.11436154 | 1.717783049 | 0.526070079 | 3.265312204 | 0.001093435 | 0.028551635 |
| SOX9 | 486.5779761 | -0.347545904 | 0.587382159 | -0.591686176 | 0.554060755 | 0.796360966 |

| | | | | | | |
|---|---|---|---|---|---|---|
| SP1 | 3273.051906 | -0.54259883 | 0.266941217 | -2.032652865 | 0.042087607 | 0.236135754 |
| SP100 | 30.18652977 | -1.105333522 | 0.867158001 | -1.274662197 | 0.202428833 | 0.509210383 |
| SP110 | 26.17218334 | -0.644163981 | 0.717281495 | -0.898063014 | 0.369151956 | 0.674991482 |
| SP140L | 30.62928894 | -0.427336928 | 0.612666035 | -0.697503867 | 0.48548752 | 0.755840687 |
| SP2 | 495.7597288 | -0.361161209 | 0.266901835 | -1.353161209 | 0.176004135 | 0.480071547 |
| SP3 | 3039.729812 | 0.119959355 | 0.304068472 | 0.394514282 | 0.693201385 | 0.868643566 |
| SP4 | 759.4401761 | 0.288290012 | 0.331287677 | 0.870210492 | 0.384185383 | 0.684816878 |
| SP5 | 588.494467 | 1.850728874 | 0.505351698 | 3.662259138 | 0.000250001 | 0.010898741 |
| SP6 | 1284.679912 | -1.732295948 | 0.659204908 | -2.62785657 | 0.008592474 | 0.104788117 |
| SP8 | 629.3533715 | 0.328648179 | 0.460836622 | 0.713155516 | 0.47574953 | 0.749091526 |
| SPA17 | 415.2614618 | 0.789237368 | 0.28547877 | 2.764609668 | 0.005699095 | 0.081260305 |
| SPACA6 | 127.1109557 | -0.223570407 | 0.365358871 | -0.611920019 | 0.540590676 | 0.787753448 |
| SPACA7 | 10.27786549 | 0.946086016 | 1.134743301 | 0.833744527 | 0.404424961 | 0.700367583 |
| SPACA9 | 131.2276949 | 0.572232528 | 0.478058752 | 1.196992056 | 0.231309651 | 0.540292336 |
| SPAG1 | 78.85691981 | 0.087477806 | 0.47631072 | 0.183657017 | 0.85428253 | 0.941473214 |
| SPAG16 | 716.0164749 | 0.430369361 | 0.442150799 | 0.973354254 | 0.330377267 | 0.638816405 |
| SPAG17 | 46.19250082 | -0.7277792 | 0.497008416 | -1.464319672 | 0.143106625 | 0.433433939 |
| SPAG4 | 35.64478416 | 0.317654007 | 0.631645399 | 0.502899265 | 0.615035098 | 0.830872437 |
| SPAG5 | 2291.064769 | 0.503187397 | 0.386613976 | 1.301524073 | 0.19307913 | 0.498908723 |
| SPAG6 | 22.04568289 | 0.655395581 | 0.74487497 | 0.879873276 | 0.378927963 | 0.681253679 |
| SPAG7 | 1895.677542 | 0.699503689 | 0.319106956 | 2.192066567 | 0.0283747 | 0.192544525 |
| SPAG8 | 14.60172328 | 0.079523186 | 0.758246225 | 0.104877786 | 0.916472786 | 0.969211928 |
| SPAG9 | 1912.064282 | 0.632610899 | 0.459018794 | 1.378180823 | 0.168147466 | 0.470384431 |
| SPAM1 | 24.12411125 | -1.917061446 | 1.212739389 | -1.580769506 | 0.113930749 | 0.388326982 |
| SPARC | 5369.309837 | 0.611931293 | 0.395364819 | 1.547763645 | 0.121679212 | 0.402231976 |
| SPART | 2120.783853 | 0.242496687 | 0.235395627 | 1.030166489 | 0.302931858 | 0.613538084 |
| SPAST | 983.1016911 | 0.190141679 | 0.282226945 | 0.673719086 | 0.500489925 | 0.763638219 |
| SPATA13 | 665.9009581 | 0.81888852 | 0.590590181 | 1.386559659 | 0.165576077 | 0.467560858 |
| SPATA17 | 130.9290391 | 0.063831288 | 0.54304249 | 0.117543819 | 0.906429122 | 0.965055357 |
| SPATA18 | 311.8320773 | -0.236133005 | 0.319861256 | -0.738235721 | 0.46037122 | 0.738395128 |
| SPATA2 | 390.936388 | 0.062454245 | 0.309730493 | 0.201640608 | 0.840197696 | 0.934461703 |
| SPATA20 | 630.3596591 | -0.079427709 | 0.396148692 | -0.200499739 | 0.841089762 | 0.93508521 |
| SPATA24 | 132.0284481 | 0.837271751 | 0.427373569 | 1.959109808 | 0.050099928 | 0.257487003 |
| SPATA2L | 127.6466682 | 0.263826715 | 0.553694205 | 0.476484517 | 0.633729236 | 0.839689885 |
| SPATA33 | 219.8471897 | 0.327435132 | 0.286227745 | 1.143967129 | 0.25263727 | 0.565423035 |
| SPATA5 | 207.6983999 | 0.262399004 | 0.352240548 | 0.74494264 | 0.456306396 | 0.73543231 |
| SPATA5L1 | 279.3955275 | 0.583873639 | 0.398168117 | 1.466399778 | 0.142539407 | 0.432925994 |
| SPATA6 | 485.0341709 | -0.456189198 | 0.275557572 | -1.655513199 | 0.097820441 | 0.359375073 |
| SPATA6L | 62.24638292 | -0.949721405 | 0.629883262 | -1.507773681 | 0.13161246 | 0.41719445 |
| SPATA7 | 583.1252408 | 0.333445279 | 0.279168086 | 1.194424777 | 0.232311849 | |

| | | | | | | |
|---|---|---|---|---|---|---|
| | | | | | | 0.54188239 |
| SPATC1L | 59.26592227 | 0.020783705 | 0.567244504 | 0.036639764 | 0.970772238 | 0.98957539 |
| SPATS2 | 1494.288011 | 0.057786417 | 0.273493829 | 0.211289656 | 0.832661254 | 0.931432899 |
| SPATS2L | 346.4846019 | -1.002828751 | 0.524931207 | -1.910400331 | 0.056081688 | 0.274279592 |
| SPC24 | 380.2561265 | 1.408934414 | 0.309202244 | 4.556675905 | 5.20E-06 | 0.000668161 |
| SPC25 | 242.9404425 | 1.148469433 | 0.337850928 | 3.399337805 | 0.000675492 | 0.021392167 |
| SPCS1 | 2162.749649 | 0.805139759 | 0.45545241 | 1.767780216 | 0.077097654 | 0.31844659 |
| SPCS2 | 2364.394679 | 0.310382357 | 0.407881687 | 0.760961736 | 0.446679922 | 0.728709125 |
| SPCS2P4 | 24.44386638 | -0.001422865 | 3.476760705 | -0.00040925 | 0.999673466 | 0.999739364 |
| SPCS3 | 2896.944293 | 0.020182521 | 0.278668768 | 0.072424768 | 0.942263874 | 0.978574897 |
| SPDL1 | 1494.708953 | 0.519947094 | 0.318834426 | 1.630774632 | 0.102937882 | 0.367884714 |
| SPDYC | 15.0192533 | 0.742093739 | 0.858633686 | 0.864272798 | 0.38743805 | 0.687545052 |
| SPECC1 | 2045.587772 | -0.321822057 | 0.232100302 | -1.386564572 | 0.165574578 | 0.467560858 |
| SPECC1L | 1822.853751 | -0.461190479 | 0.444094273 | -1.038496796 | 0.299038826 | 0.610246278 |
| SPECC1L-ADORA2A | 12.10645171 | 1.632050659 | 1.965337095 | 0.830417674 | 0.406302676 | 0.701474651 |
| SPEF2 | 87.81418589 | -0.381265378 | 0.437823763 | -0.870819289 | 0.383852831 | 0.684699412 |
| SPEG | 253.4341877 | -0.09153039 | 0.290674483 | -0.314889664 | 0.75284543 | 0.896586461 |
| SPEN | 1799.795635 | -0.21398673 | 0.509949192 | -0.419623628 | 0.674760425 | 0.859745438 |
| SPG11 | 1761.005201 | -0.400949117 | 0.354427385 | -1.131258853 | 0.257946157 | 0.570701641 |
| SPG21 | 2430.199813 | 0.424250727 | 0.378282436 | 1.121518439 | 0.262067247 | 0.575601582 |
| SPG7 | 2164.405608 | -0.386139697 | 0.25453147 | -1.517060726 | 0.129251353 | 0.413472911 |
| SPHK1 | 159.0036739 | 0.802179761 | 0.621459597 | 1.290799539 | 0.196773198 | 0.50258876 |
| SPHK2 | 1106.661906 | -0.74550353 | 0.469483329 | -1.587923328 | 0.112303678 | 0.385402109 |
| SPIB | 13.42555296 | -0.012466261 | 0.957964928 | -0.013013275 | 0.989617202 | 0.996285567 |
| SPICE1 | 696.2021535 | 0.007993863 | 0.278294112 | 0.028724513 | 0.977084306 | 0.992059029 |
| SPIDR | 981.0081689 | -0.066584912 | 0.291190784 | -0.228664215 | 0.81912991 | 0.924316427 |
| SPIN1 | 3317.738159 | -0.561527318 | 0.263444497 | -2.131482436 | 0.033049418 | 0.207677316 |
| SPIN2A | 47.85067207 | 0.217868901 | 0.473669887 | 0.459959367 | 0.645545386 | 0.845794147 |
| SPIN2B | 143.0404271 | 0.816908181 | 0.495707385 | 1.647964516 | 0.099359953 | 0.361473741 |
| SPIN3 | 463.456678 | -0.107456079 | 0.328353437 | -0.327257359 | 0.743473241 | 0.892534058 |
| SPIN4 | 1667.879026 | -1.489419933 | 0.529585262 | -2.812427079 | 0.004916917 | 0.074894126 |
| SPINDOC | 1811.891007 | 0.142079536 | 0.316143947 | 0.449414065 | 0.653132987 | 0.848827707 |
| SPINK1 | 90.37964114 | 1.248542295 | 0.649908988 | 1.92110329 | 0.054718687 | 0.270592163 |
| SPINK2 | 28.73635329 | 1.007494447 | 0.944950208 | 1.066187867 | 0.286338726 | 0.60033547 |
| SPINK5 | 592.4023582 | -0.005365253 | 0.413727825 | -0.012968074 | 0.989653264 | 0.996285567 |
| SPINK9 | 61.60312091 | 1.241843285 | 0.694443454 | 1.788256883 | 0.073734573 | 0.311905115 |
| SPINT1 | 3151.873822 | -0.960101858 | 0.632555996 | -1.517813229 | 0.129061489 | 0.413453061 |

| | | | | | | |
|---|---|---|---|---|---|---|
| SPINT2 | 6907.883075 | 0.297559462 | 0.602411665 | 0.493947046 | 0.621343578 | 0.833354021 |
| SPIRE1 | 352.1736357 | 0.140356095 | 0.420468305 | 0.333808977 | 0.738523709 | 0.891339952 |
| SPIRE2 | 269.3543989 | -0.07768209 | 0.281477389 | -0.275979859 | 0.782563536 | 0.909909835 |
| SPN | 28.70751304 | 0.026378143 | 0.613300488 | 0.043010145 | 0.965693447 | 0.988700292 |
| SPNS1 | 1055.227591 | -0.305171652 | 0.423064398 | -0.72133617 | 0.470702708 | 0.745721678 |
| SPOCD1 | 74.33351901 | -0.418346371 | 0.480088025 | -0.871395138 | 0.383538439 | 0.684548431 |
| SPOCK1 | 198.3788746 | 0.381621424 | 0.447189069 | 0.853378247 | 0.393449581 | 0.69249409 |
| SPOCK2 | 47.51945794 | 1.543002874 | 0.72570021 | 2.126226302 | 0.033484421 | 0.208789229 |
| SPON1 | 704.7933222 | 0.76089933 | 0.560440985 | 1.357679669 | 0.174565351 | 0.478578744 |
| SPON2 | 18.23778045 | 1.037066165 | 0.856752588 | 1.210461667 | 0.226101791 | 0.534727976 |
| SPOP | 640.5870278 | 0.426335457 | 0.331487977 | 1.286126455 | 0.198398933 | 0.504172564 |
| SPOPL | 685.9846864 | -0.855185273 | 0.40148543 | -2.130053072 | 0.033167232 | 0.207839767 |
| SPOUT1 | 1363.695032 | -0.025289242 | 0.252250117 | -0.100254631 | 0.920142175 | 0.969770994 |
| SPP1 | 273.5303462 | -0.397929711 | 0.782889312 | -0.508283489 | 0.611254548 | 0.829058038 |
| SPPL2A | 1066.165912 | -0.6928332 | 0.447773302 | -1.547285638 | 0.121794383 | 0.402470614 |
| SPPL2B | 1114.302353 | -0.291952862 | 0.494438899 | -0.590473086 | 0.554873524 | 0.796360966 |
| SPPL3 | 1509.073823 | -0.078598339 | 0.356838672 | -0.220262952 | 0.825666372 | 0.927178721 |
| SPR | 109.0448273 | -1.114115351 | 0.86363513 | -1.290030144 | 0.197040192 | 0.50258876 |
| SPRED1 | 954.6249044 | -0.108852645 | 0.328224571 | -0.331640758 | 0.740160539 | 0.891917214 |
| SPRED2 | 603.8490661 | -0.630197556 | 0.292339065 | -2.15570764 | 0.031106501 | 0.200942211 |
| SPRED3 | 32.99513521 | -0.868976195 | 0.653655182 | -1.329410702 | 0.183712509 | 0.488132305 |
| SPRTN | 408.5325907 | -0.170940667 | 0.262534863 | -0.651116067 | 0.514971565 | 0.773145336 |
| SPRY1 | 74.09046105 | 0.147000733 | 0.530277678 | 0.277214634 | 0.781615303 | 0.909765672 |
| SPRY2 | 388.6936565 | -0.444201312 | 0.532700221 | -0.833867332 | 0.404355748 | 0.700367583 |
| SPRY3 | 50.70977795 | 0.26767589 | 0.775255507 | 0.345274413 | 0.729888081 | 0.886488838 |
| SPRY4 | 59.51672094 | -0.36203421 | 0.735755509 | -0.492057763 | 0.622678509 | 0.833787151 |
| SPRYD3 | 215.016858 | 0.08700307 | 0.338541729 | 0.256993636 | 0.797183686 | 0.915732089 |
| SPRYD4 | 339.002504 | -0.185826916 | 0.326164479 | -0.569733763 | 0.568858286 | 0.805450586 |
| SPRYD7 | 280.4173693 | 0.551332217 | 0.348284213 | 1.582995141 | 0.113422572 | 0.387639973 |
| SPSB1 | 193.8699199 | -0.446437113 | 0.296622755 | -1.505066977 | 0.132306858 | 0.418521131 |
| SPSB2 | 344.1963811 | 0.072205486 | 0.487167742 | 0.148214834 | 0.882173225 | 0.953230092 |
| SPSB3 | 394.753588 | -0.194969454 | 0.51375824 | -0.379496499 | 0.704319204 | 0.873435301 |
| SPSB4 | 2484.968657 | -0.545573969 | 0.35445515 | -1.539190414 | 0.123757817 | 0.405045568 |
| SPTAN1 | 7738.482902 | -0.776298865 | 0.438715828 | -1.769479956 | 0.076813813 | 0.31831258 |
| SPTB | 154.3688812 | -0.857126251 | 0.532634356 | -1.609220738 | 0.107568083 | 0.376005887 |
| SPTBN1 | 5642.60346 | -0.455332783 | 0.384758346 | -1.183425357 | | |

| | | | | | | |
|---|---|---|---|---|---|---|
| | | | | | 0.236640613 | 0.547336452 |
| SPTBN2 | 1626.799219 | -0.690964695 | 0.323837606 | -2.133676515 | 0.03286927 | 0.206912735 |
| SPTBN4 | 68.06427774 | 0.640330443 | 0.513749676 | 1.246386078 | 0.21262269 | 0.519735803 |
| SPTLC1 | 2876.725978 | -0.072241291 | 0.236998202 | -0.304817889 | 0.760504856 | 0.899902262 |
| SPTLC2 | 894.3341675 | -0.399105909 | 0.262944974 | -1.517830526 | 0.129057128 | 0.413453061 |
| SPTLC3 | 327.4381049 | -0.866125245 | 0.562110726 | -1.540844544 | 0.123354625 | 0.404753154 |
| SPTSSA | 1039.175731 | 0.921033856 | 0.38005634 | 2.423414001 | 0.015375393 | 0.140602827 |
| SPTSSB | 137.3923514 | -0.222832809 | 0.728737439 | -0.30577928 | 0.759772706 | 0.899725004 |
| SPTY2D1 | 771.9524549 | -0.279674946 | 0.286420615 | -0.976448383 | 0.328842315 | 0.638207339 |
| SPTY2D1OS | 35.81923089 | 1.04658166 | 0.637805593 | 1.640910132 | 0.100816072 | 0.363642563 |
| SPX | 17.87148772 | -0.51404314 | 0.742478816 | -0.692333745 | 0.488727763 | 0.757260184 |
| SQLE | 2247.161338 | 0.541232309 | 0.305140437 | 1.773715457 | 0.076110228 | 0.316520909 |
| SQOR | 26.16169402 | -0.332200295 | 0.916547442 | -0.362447463 | 0.717017677 | 0.880371898 |
| SQSTM1 | 1790.38503 | 0.316561384 | 0.302709368 | 1.045760115 | 0.2956718 | 0.60699582 |
| SRA1 | 734.8229418 | 0.413365627 | 0.359008557 | 1.151408841 | 0.249564078 | 0.562413758 |
| SRBD1 | 753.6258449 | -0.014774699 | 0.26993464 | -0.054734359 | 0.956350096 | 0.984447503 |
| SRC | 1407.776788 | -0.444682082 | 0.379856449 | -1.17065824 | 0.241736177 | 0.552977821 |
| SRCAP | 1566.735052 | -0.995712048 | 0.780421143 | -1.275865033 | 0.202003234 | 0.508645818 |
| SRCIN1 | 118.2674239 | -1.15656296 | 0.51177597 | -2.259900871 | 0.023827404 | 0.176679151 |
| SRD5A1 | 925.9832327 | 0.330300328 | 0.34733489 | 0.950956373 | 0.341626523 | 0.648500498 |
| SRD5A3 | 708.9317125 | 0.679757563 | 0.345105524 | 1.96970931 | 0.048871695 | 0.254713129 |
| SREBF1 | 547.1093833 | -0.072351182 | 0.257255418 | -0.281242598 | 0.778524334 | 0.908160087 |
| SREBF2 | 6460.157525 | -0.14163464 | 0.342502566 | -0.413528696 | 0.679219303 | 0.861625115 |
| SREK1 | 2373.542993 | 0.15172854 | 0.238433658 | 0.636355374 | 0.52454482 | 0.777662198 |
| SREK1IP1 | 1322.802236 | 0.916136605 | 0.314559321 | 2.912444633 | 0.003586118 | 0.061544108 |
| SRF | 1789.951292 | 0.018212832 | 0.300552972 | 0.060597743 | 0.951679571 | 0.982350991 |
| SRFBP1 | 426.2072098 | 0.064138378 | 0.33649147 | 0.190609224 | 0.848831763 | 0.938569494 |
| SRGAP1 | 246.2109907 | -0.296041684 | 0.423304418 | -0.699358833 | 0.484327808 | 0.754852802 |
| SRGAP2 | 1513.434294 | -0.097257812 | 0.390670019 | -0.248951308 | 0.803398446 | 0.917969611 |
| SRGAP2B | 691.8773161 | 0.276797103 | 0.342353979 | 0.808511424 | 0.418796232 | 0.709799118 |
| SRGAP2C | 824.8020633 | 0.355338068 | 0.316652456 | 1.122170574 | 0.261789922 | 0.575262877 |
| SRGAP2D | 56.87696757 | -0.580052329 | 1.220986274 | -0.475068673 | 0.634738027 | 0.840313679 |
| SRGAP3 | 632.0708859 | 0.331984877 | 0.525886411 | 0.631286283 | 0.527853354 | 0.779221878 |
| SRI | 1812.308289 | 0.198592317 | 0.381384692 | 0.520713917 | 0.602566078 | 0.825021776 |
| SRM | 1006.718309 | 0.110820876 | 0.242652291 | 0.456706489 | 0.647882026 | 0.846597949 |
| SRP14 | 7654.951653 | 0.542541025 | 0.353183372 | 1.536145437 | 0.124502714 | 0.405999855 |

| | | | | | | |
|---|---|---|---|---|---|---|
| SRP19 | 1249.135068 | 0.751867878 | 0.3803538 | 1.97675921 | 0.048068854 | 0.252967897 |
| SRP54 | 1745.303842 | -0.075648705 | 0.260349386 | -0.290566096 | 0.771383193 | 0.905067213 |
| SRP54-AS1 | 34.83235237 | 1.156831785 | 0.565206346 | 2.046742386 | 0.040683385 | 0.23195346 |
| SRP68 | 2338.660189 | -0.226941349 | 0.269173147 | -0.843105455 | 0.399169468 | 0.697352242 |
| SRP72 | 4226.068717 | 0.068513745 | 0.223294341 | 0.306831535 | 0.758971604 | 0.899440455 |
| SRP9 | 11583.96229 | 0.806866024 | 0.303842941 | 2.655536516 | 0.007918236 | 0.100106299 |
| SRPK1 | 2877.695718 | 0.239724353 | 0.247134852 | 0.970014351 | 0.332039339 | 0.639828376 |
| SRPK2 | 1637.836226 | 0.042801438 | 0.251891395 | 0.169920209 | 0.865072888 | 0.945248223 |
| SRPRA | 1746.843009 | -0.457448923 | 0.26728213 | -1.71148338 | 0.086991917 | 0.338745988 |
| SRPRB | 709.4655146 | 0.243113557 | 0.29523437 | 0.823459537 | 0.410246718 | 0.704135419 |
| SRPX | 179.3136142 | 0.269785525 | 0.431420668 | 0.625342142 | 0.531746526 | 0.78215305 |
| SRPX2 | 10.9077571 | 0.129106745 | 0.933937075 | 0.138239233 | 0.890051349 | 0.956707455 |
| SRR | 365.8078899 | 0.297442173 | 0.272147641 | 1.092944155 | 0.27441832 | 0.587352209 |
| SRRD | 672.3846353 | 0.285913553 | 0.279096044 | 1.024427106 | 0.305633585 | 0.615675317 |
| SRRM1 | 4885.964962 | 0.024588463 | 0.298663591 | 0.082328291 | 0.934385658 | 0.97507153 |
| SRRM2 | 14948.13442 | -0.547088506 | 0.438876164 | -1.246566915 | 0.212556339 | 0.519735803 |
| SRRM3 | 169.0461644 | 1.255540157 | 0.345668248 | 3.632211417 | 0.000281003 | 0.01187491 |
| SRRM4 | 33.88791205 | -0.056876246 | 0.594939525 | -0.095600046 | 0.923838228 | 0.971413208 |
| SRRT | 3685.214096 | -0.084631903 | 0.342815727 | -0.246872872 | 0.805006612 | 0.918735824 |
| SRSF1 | 8149.471564 | 0.016169461 | 0.214718239 | 0.075305487 | 0.939971656 | 0.978006309 |
| SRSF10 | 4602.078487 | 0.186456777 | 0.245083456 | 0.760788915 | 0.446783156 | 0.728709125 |
| SRSF11 | 7685.113232 | 0.270715158 | 0.274790181 | 0.98517042 | 0.324540367 | 0.634648352 |
| SRSF12 | 560.9322018 | 0.447084161 | 0.268245193 | 1.666699618 | 0.095574149 | 0.354819129 |
| SRSF2 | 6872.23114 | 0.465280911 | 0.372603594 | 1.248728992 | 0.211764213 | 0.519010481 |
| SRSF3 | 15166.79999 | 0.533915701 | 0.241982852 | 2.206419569 | 0.027354636 | 0.189237202 |
| SRSF4 | 3260.099805 | -0.168263937 | 0.221270151 | -0.76044571 | 0.446988209 | 0.728783454 |
| SRSF5 | 7545.893239 | 0.208898791 | 0.344599779 | 0.606206979 | 0.544377319 | 0.790739196 |
| SRSF6 | 5807.493358 | 0.056268184 | 0.295387428 | 0.190489435 | 0.848925621 | 0.938569494 |
| SRSF7 | 7667.479 | 0.577978732 | 0.220827556 | 2.617330654 | 0.008862045 | 0.106618623 |
| SRSF8 | 761.7773704 | 0.071116357 | 0.387180512 | 0.183677523 | 0.854266443 | 0.941473214 |
| SRSF9 | 5234.374818 | 0.581552422 | 0.3887895 | 1.495802798 | 0.134705052 | 0.421189271 |
| SRXN1 | 487.4420193 | 0.6141571 | 0.346741603 | 1.771224145 | 0.076523436 | 0.317542955 |
| SRY | 14.78854119 | 1.089927 | 0.788897724 | 1.381582132 | 0.167100043 | 0.469215449 |
| SS18 | 2529.562919 | -0.321893569 | 0.25624458 | -1.256196595 | 0.209044701 | 0.516433343 |
| SS18L1 | 782.3605899 | 0.162928782 | 0.249663093 | 0.652594583 | 0.514017678 | 0.772630754 |
| SS18L2 | 966.7250206 | 1.080376333 | 0.444396558 | 2.431108689 | 0.015052698 | 0.139165886 |
| SSB | 6922.25554 | 0.620529571 | 0.284234393 | 2.183161459 | 0.029023922 | 0.194746536 |
| SSBP1 | 3082.859551 | 0.691073393 | 0.369431212 | 1.870641598 | 0.061394775 | 0.285640507 |
| SSBP2 | 2930.336608 | 0.081001293 | 0.276388949 | 0.29306994 | 0.769468693 | 0.904597192 |
| SSBP3 | 4485.040667 | -0.130983172 | 0.286505501 | -0.457175068 | 0.647545217 | 0.846597949 |
| SSBP4 | 332.5479394 | 1.009552025 | 0.543259472 | 1.858323833 | 0.063123041 | 0.289778852 |
| SSC4D | 340.3302371 | -0.542545639 | 0.316503729 | -1.714184035 | 0.086494932 | 0.337416975 |
| SSC5D | 77.66993141 | -0.823101678 | 0.486305265 | -1.692561725 | 0.090538922 | 0.344933707 |
| SSH1 | 898.8676585 | -0.568605862 | 0.391931886 | -1.450777245 | 0.146841898 | 0.439926258 |
| SSH2 | 353.6003441 | -0.400936939 | 0.41698399 | -0.961516384 | 0.336292592 | 0.64425371 |

| | | | | | | |
|---|---|---|---|---|---|---|
| SSH3 | 569.394535 | -1.289499501 | 0.304465694 | -4.235286684 | 2.28E-05 | 0.002051038 |
| SSNA1 | 789.7893494 | 0.734058686 | 0.405983255 | 1.808100893 | 0.070590799 | 0.305370121 |
| SSPN | 32.46841085 | 0.148041163 | 0.731380349 | 0.202413372 | 0.839593576 | 0.934243372 |
| SSPO | 175.2176716 | 0.117886851 | 0.513554947 | 0.229550609 | 0.81844099 | 0.924316427 |
| SSR1 | 4305.43509 | 0.166188356 | 0.325634314 | 0.510352715 | 0.609804377 | 0.828453677 |
| SSR2 | 6634.736728 | 0.572751166 | 0.387263531 | 1.478970053 | 0.139148319 | 0.427760708 |
| SSR3 | 3610.75885 | 0.419766743 | 0.325090816 | 1.291229166 | 0.196624225 | 0.502524614 |
| SSR4 | 2160.482118 | 0.601350253 | 0.417843808 | 1.439174741 | 0.15010102 | 0.444740017 |
| SSR4P1 | 26.8944866 | 0.287058614 | 0.593154113 | 0.483952834 | 0.628419343 | 0.836791621 |
| SSRP1 | 9265.165226 | 0.126510345 | 0.236698106 | 0.534479751 | 0.593009655 | 0.819859199 |
| SSSCA1 | 261.8908934 | 0.461854597 | 0.410724366 | 1.124487942 | 0.260806083 | 0.57415519 |
| SSTR2 | 40.21815022 | 0.805350713 | 0.577491206 | 1.394567926 | 0.163146173 | 0.464573649 |
| SSTR3 | 18.91975795 | 1.679934478 | 0.85606528 | 1.962390624 | 0.049717024 | 0.256987041 |
| SSU72 | 2912.300483 | 0.409846945 | 0.360215643 | 1.137782194 | 0.255211442 | 0.567631254 |
| SSUH2 | 98.61111948 | 0.332710372 | 0.516139963 | 0.644612694 | 0.519178206 | 0.774631448 |
| SSX2IP | 691.0434811 | 0.048711352 | 0.320922347 | 0.151785478 | 0.879356137 | 0.95195604 |
| ST13 | 13007.63628 | -0.037304202 | 0.258310268 | -0.144416257 | 0.88517178 | 0.954505727 |
| ST14 | 920.7682234 | -1.64723366 | 0.667392758 | -2.468162322 | 0.013580872 | 0.132501135 |
| ST20 | 192.112251 | 0.789210141 | 0.499554593 | 1.579827614 | 0.11414635 | 0.388887104 |
| ST20-MTHFS | 48.78296114 | -0.171585555 | 0.668941036 | -0.256503258 | 0.797562265 | 0.915916373 |
| ST3GAL1 | 518.2401103 | -2.691509271 | 0.904894182 | -2.974391176 | 0.002935706 | 0.053530758 |
| ST3GAL2 | 531.1191386 | 0.551192717 | 0.389231626 | 1.416104656 | 0.156744865 | 0.455611392 |
| ST3GAL3 | 318.1065358 | 0.922292472 | 0.425284779 | 2.168646796 | 0.03010951 | 0.198185614 |
| ST3GAL4 | 368.0771512 | 0.243515176 | 0.284364122 | 0.856349862 | 0.391804289 | 0.691330875 |
| ST3GAL5 | 201.0183168 | 0.384264831 | 0.386053107 | 0.995367797 | 0.319557415 | 0.629284208 |
| ST3GAL6 | 595.4699592 | -0.267155253 | 0.389064313 | -0.686660903 | 0.49229644 | 0.759174768 |
| ST5 | 933.835259 | -0.219664194 | 0.470857289 | -0.466519684 | 0.640843561 | 0.843724522 |
| ST6GAL1 | 1577.104931 | -0.418902543 | 0.384361934 | -1.089864801 | 0.275772704 | 0.58901136 |
| ST6GAL2 | 168.0070473 | 0.131593269 | 0.645582128 | 0.203836605 | 0.838481187 | 0.933879959 |
| ST6GALNAC2 | 157.6505614 | -0.844976417 | 0.651430948 | -1.297108189 | 0.194593964 | 0.500370345 |
| ST6GALNAC3 | 570.5106007 | 0.582858897 | 0.361373758 | 1.612897682 | 0.106766739 | 0.374599763 |
| ST6GALNAC4 | 111.2985209 | 1.704567094 | 0.5402868 | 3.154930111 | 0.001605366 | 0.037175747 |
| ST6GALNAC5 | 288.4090706 | -0.188656161 | 0.30078034 | -0.627222383 | 0.530513473 | 0.780945071 |
| ST6GALNAC6 | 944.8686837 | 0.579941669 | 0.300392265 | 1.930614521 | 0.053530741 | 0.267406937 |
| ST7 | 184.2573182 | -0.215943129 | 0.385205538 | -0.560591964 | 0.575075731 | 0.808420489 |
| ST7L | 357.3432889 | 0.229800279 | 0.331989713 | 0.692190962 | 0.488817414 | 0.757260184 |
| ST8SIA1 | 71.25316597 | 1.983054549 | 0.639110258 | 3.102836362 | 0.001916756 | 0.040960574 |
| ST8SIA2 | 120.6585356 | 0.734284097 | 0.601076723 | 1.221614595 | 0.221853407 | 0.530789786 |
| ST8SIA3 | 16.95856209 | 1.012326097 | 0.962249849 | 1.052040796 | 0.292780833 | 0.605827098 |

| | | | | | | |
|---|---|---|---|---|---|---|
| ST8SIA4 | 26.71677586 | 0.461816566 | 0.59885082 | 0.771171301 | 0.440605402 | 0.7241465 |
| ST8SIA5 | 14.89756806 | -0.699692254 | 0.785961159 | -0.890237699 | 0.373338266 | 0.678294888 |
| ST8SIA6 | 98.84114576 | -0.262640317 | 0.627534364 | -0.418527387 | 0.675561568 | 0.860110127 |
| STAC | 101.1996048 | 0.650137564 | 0.620955114 | 1.046996069 | 0.295101391 | 0.60699582 |
| STAC2 | 39.57366714 | 0.439001617 | 0.694616464 | 0.632005775 | 0.527383103 | 0.778906646 |
| STAC3 | 78.2094649 | 0.260205225 | 0.411907676 | 0.631707638 | 0.527577936 | 0.778966897 |
| STAG1 | 1060.908216 | -0.169891376 | 0.366924767 | -0.463014197 | 0.643354191 | 0.845415888 |
| STAG2 | 4046.232431 | 0.085813403 | 0.325310894 | 0.263788902 | 0.791942587 | 0.913570955 |
| STAG3 | 69.41184158 | -0.924505754 | 0.653642844 | -1.414389772 | 0.157247491 | 0.456574487 |
| STAG3L1 | 78.53209921 | -0.670415946 | 0.535309497 | -1.252389412 | 0.210428003 | 0.51816316 |
| STAG3L2 | 321.8984825 | -0.249193778 | 0.308292762 | -0.808302393 | 0.418916525 | 0.709860673 |
| STAG3L3 | 550.9848701 | -0.449419668 | 0.283865035 | -1.583216006 | 0.113372239 | 0.38755526 |
| STAG3L4 | 425.906434 | 0.458607912 | 0.344114253 | 1.332719898 | 0.182623741 | 0.486683209 |
| STAG3L5P | 296.213315 | -0.712152622 | 0.679487161 | -1.048073699 | 0.294604653 | 0.606767199 |
| STAM | 893.1342304 | 0.110120057 | 0.275514187 | 0.399689243 | 0.689385416 | 0.867579108 |
| STAM2 | 633.7866701 | -0.300662828 | 0.265381887 | -1.132944041 | 0.257237753 | 0.569767157 |
| STAMBP | 1318.485885 | -0.12281233 | 0.2299384 | -0.534109703 | 0.593265637 | 0.819859199 |
| STAMBPL1 | 43.17895776 | 0.256735371 | 0.55788558 | 0.460193596 | 0.645377267 | 0.845794147 |
| STAP2 | 706.7376536 | -0.267165414 | 0.711117648 | -0.375697909 | 0.707141492 | 0.875135606 |
| STAR | 17.29534927 | 1.023630447 | 0.917917105 | 1.115166545 | 0.264779059 | 0.578479711 |
| STARD10 | 888.1113581 | -0.564331522 | 0.505153278 | -1.117149085 | 0.263930586 | 0.577448862 |
| STARD13 | 69.49969712 | 0.225676472 | 0.511266928 | 0.441406356 | 0.658918842 | 0.851414509 |
| STARD3 | 1049.066925 | 0.021168158 | 0.283954523 | 0.074547706 | 0.940574583 | 0.97807441 |
| STARD3NL | 1453.702996 | 0.37815796 | 0.24402076 | 1.549695853 | 0.121214534 | 0.401420514 |
| STARD4 | 954.0465242 | 0.670552837 | 0.358897024 | 1.868371127 | 0.061710359 | 0.28627362 |
| STARD5 | 211.6106487 | -0.975034714 | 0.399484113 | -2.440734645 | 0.01465742 | 0.137859718 |
| STARD7 | 3725.202981 | 0.232120269 | 0.22143469 | 1.048256121 | 0.29452062 | 0.606702791 |
| STARD8 | 147.7690303 | 0.79579571 | 0.611216395 | 1.301986853 | 0.19292088 | 0.498908723 |
| STARD9 | 601.3743088 | 0.279174792 | 0.662497831 | 0.421397292 | 0.673464996 | 0.859449651 |
| STAT1 | 1796.481259 | 0.011600846 | 0.262770053 | 0.04414828 | 0.964786208 | 0.988700292 |
| STAT2 | 1693.410571 | -0.771113779 | 0.318865594 | -2.418303488 | 0.015593065 | 0.141315649 |
| STAT3 | 1894.970935 | 0.231467708 | 0.332737653 | 0.695646271 | 0.486650381 | 0.756340124 |
| STAT5A | 132.0507142 | -0.193145524 | 0.454864947 | -0.424621694 | 0.671112476 | 0.858469424 |
| STAT5B | 1504.550379 | 0.435305852 | 0.368411758 | 1.181574264 | 0.23737467 | 0.547879744 |
| STAT6 | 1744.124415 | -1.05089293 | 0.314591648 | -3.340498505 | 0.000836281 | 0.024340176 |
| STAU1 | 4258.817475 | -0.095349573 | 0.214678311 | -0.444150936 | 0.656933462 | 0.85024732 |
| STAU2 | 1921.232634 | -0.08230536 | 0.256147304 | -0.321320425 | 0.74796758 | |

0.894202292
STBD1 28.39984884 -2.273412127 0.986285832 -2.305023608 0.021165251
0.165258892
STC1 394.2375886 0.798609265 0.441299182 1.809677646 0.070345791 0.304732874
STC2 1514.593701 -0.827826027 0.299056387 -2.768126889 0.00563795
0.080615777
STEAP1 130.2043326 0.48912602 0.69005597 0.708820794 0.478435692
0.750837683
STEAP1B 34.38503539 1.624651015 0.675444446 2.405306645 0.0161589
0.143781043
STEAP2 211.9061609 -0.555251333 0.460206667 -1.20652605 0.227614721
0.536368893
STEAP3 585.6688305 -0.571981382 0.514167081 -1.112442634
0.265947882 0.579614326
STEAP4 20.81557457 -0.208957566 0.796027944 -0.26250029 0.792935762
0.913966604
STIL 712.3358055 0.00336019 0.418303561 0.008032899 0.993590743 0.99730871
STIM1 793.3115295 0.066964905 0.295723833 0.226444061 0.820856062 0.925132406
STIM2 570.3697642 0.124047774 0.30371634 0.408432993 0.682955815 0.863283009
STIMATE 558.8747249 -0.238157426 0.445616218 -0.534445148
0.59303359 0.819859199
STIP1 6199.493211 -0.073774 0.304118549 -0.242583034 0.808328425
0.91963689
STK10 313.0681495 -0.549000034 0.304656741 -1.802028186 0.071540971
0.307638342
STK11 961.1803637 -0.072341712 0.464122545 -0.155867696 0.876137309
0.950296086
STK11IP 857.0898179 -0.197914782 0.319632712 -0.619194388
0.535788308 0.784725687
STK16 868.5375338 -0.07615692 0.300680692 -0.253281711 0.800050528
0.916311991
STK17A 447.1865941 0.427244421 0.315318184 1.354962836 0.175429398
0.479452242
STK17B 250.0099147 0.409257644 0.356696209 1.147356305 0.251234392
0.564075361
STK19 101.8825219 0.196274056 0.820018548 0.239353191 0.810831723 0.920880976
STK24 4412.658669 0.212261673 0.34660918 0.612394841 0.540276555 0.787481586
STK25 3417.958326 0.093280326 0.312052545 0.298925062 0.764997224 0.902991439
STK26 4200.176172 0.084204176 0.222233967 0.378898765 0.704763041 0.873723655
STK3 615.3347234 0.229826685 0.293941011 0.781880298 0.434284925 0.719429635
STK31 14.74626554 -0.251652696 0.780576539 -0.322393364 0.747154711
0.89418504
STK32A 59.34428133 -0.68557112 0.500384045 -1.370089886 0.170658844
0.473840653
STK32B 28.550688 1.022517508 0.751541137 1.36056093 0.173652485 0.4776939
STK32C 288.7635268 0.973066174 0.435361053 2.235078605 0.025412189
0.18194777
STK33 343.0934352 0.72342176 0.455019599 1.589869449 0.111864235 0.384888109
STK35 1414.241991 -0.261795197 0.222861509 -1.174699025 0.24011515
0.551686648
STK36 1256.419771 -0.57819235 0.362629941 -1.594441839 0.110837108
0.383119107
STK38 1066.679963 -0.417999708 0.289312108 -1.444805439 0.148512543
0.442803058
STK38L 1327.201918 0.144770744 0.279514472 0.51793649 0.604502582
0.825894731
STK39 373.167126 0.531432856 0.414891847 1.280894911 0.200230581 0.506121566
STK4 1382.98257 -0.135883124 0.314373114 -0.432235194 0.665570483
0.855161526
STK40 608.6257396 -0.309877555 0.477801548 -0.648548663 0.516630147
0.773950426
STMN1 19360.21344 0.843086025 0.399255374 2.111646026 0.034716822 0.212806795
STMN2 1695.451923 -0.068808183 0.752662924 -0.091419653 0.927159146
0.972482583

| | | | | | | |
|---|---|---|---|---|---|---|
| STMN3 | 199.6428674 | -0.066885994 | 0.388399244 | -0.172209383 | 0.863272922 | 0.94479249 |
| STMP1 | 2311.279757 | 0.612169567 | 0.422709861 | 1.448202713 | 0.147560366 | 0.441284903 |
| STN1 | 379.7899187 | 0.320476171 | 0.318111125 | 1.007434652 | 0.313725946 | 0.62342819 |
| STOM | 10079.53078 | -1.622583564 | 0.707240182 | -2.294246856 | 0.021776323 | 0.167529716 |
| STOML1 | 80.10493923 | 0.344971485 | 0.502044342 | 0.687133498 | 0.491998607 | 0.75916506 |
| STOML2 | 3564.852393 | 0.464779355 | 0.293991401 | 1.580928399 | 0.11389441 | 0.388324469 |
| STOML3 | 20.10848449 | 2.28312331 | 1.12670241 | 2.026376521 | 0.042726215 | 0.237784082 |
| STON1 | 1183.549143 | -0.087279063 | 0.368074791 | -0.237123176 | 0.812561234 | 0.921654347 |
| STON2 | 1076.578763 | 0.218880109 | 0.612148858 | 0.357560267 | 0.720672417 | 0.882172735 |
| STOX1 | 435.8529005 | 0.509290572 | 0.374239028 | 1.360869748 | 0.173554855 | 0.477540818 |
| STOX2 | 602.5111271 | 0.426471598 | 0.380315689 | 1.121362096 | 0.262133764 | 0.575601582 |
| STPG1 | 337.705713 | 0.024513126 | 0.30079077 | 0.081495607 | 0.935047818 | 0.975224148 |
| STPG2 | 10.57585791 | -0.004348632 | 0.875191046 | -0.00496878 | 0.996035504 | 0.998272751 |
| STRA6 | 1145.264102 | -1.630249896 | 0.468731549 | -3.478003348 | 0.000505164 | 0.017658611 |
| STRADA | 1172.84134 | -0.181333755 | 0.276979943 | -0.654681897 | 0.512672583 | 0.771819984 |
| STRADB | 1415.653357 | -0.178050177 | 0.321662997 | -0.553530183 | 0.579900419 | 0.811340629 |
| STRAP | 7922.381488 | 0.312695599 | 0.318443198 | 0.981950945 | 0.326124016 | 0.635960913 |
| STRBP | 2488.961554 | -0.058564949 | 0.245631332 | -0.238426215 | 0.811550537 | 0.921145683 |
| STRC | 29.00510271 | -1.146037758 | 0.719805735 | -1.592148691 | 0.111351299 | 0.383846979 |
| STRCP1 | 23.14250485 | -0.062681083 | 0.789130699 | -0.079430547 | 0.936690172 | 0.976155095 |
| STRIP1 | 1239.965027 | -0.495874461 | 0.357892623 | -1.385539764 | 0.165887483 | 0.467940842 |
| STRIP2 | 249.2301432 | 0.464139522 | 0.478782426 | 0.969416372 | 0.332337487 | 0.640217269 |
| STRN | 1133.275448 | -0.720192704 | 0.342923871 | -2.100153318 | 0.035715356 | 0.216129905 |
| STRN3 | 1164.651749 | 0.206155768 | 0.244045903 | 0.844741771 | 0.398255032 | 0.696877404 |
| STRN4 | 2798.464148 | 0.042576787 | 0.419069638 | 0.101598357 | 0.919075484 | 0.969677217 |
| STS | 198.9009386 | -2.436987706 | 0.943183185 | -2.58379045 | 0.009772115 | 0.112404636 |
| STT3A | 4590.218531 | -0.455193204 | 0.294832595 | -1.543903935 | 0.122611607 | 0.403413725 |
| STT3B | 9320.95384 | -0.186228877 | 0.27576529 | -0.675316597 | 0.499474642 | 0.76281557 |
| STUB1 | 1289.402123 | 0.639280269 | 0.441763166 | 1.447110844 | 0.147865881 | 0.44174616 |
| STUM | 36.31751738 | 0.516877011 | 0.870162204 | 0.59400076 | 0.552511605 | 0.795662343 |
| STX10 | 542.0067777 | 0.745203697 | 0.32631738 | 2.283677614 | 0.022390485 | 0.170396445 |
| STX12 | 542.7052272 | 0.096181168 | 0.254683682 | 0.37764951 | 0.70569098 | 0.874104985 |
| STX16 | 3541.550755 | -0.091927332 | 0.242041749 | -0.379799486 | 0.704094264 | 0.873421952 |
| STX16-NPEPL1 | 141.2875964 | -0.950675458 | 0.811419286 | -1.171620486 | 0.241349459 | 0.552681155 |
| STX17 | 1279.11053 | -0.55675253 | 0.307651317 | -1.80968681 | 0.070344369 | 0.304732874 |
| STX18 | 732.3439211 | -0.38224624 | 0.276405483 | -1.382918444 | 0.166689874 | 0.468674186 |
| STX1A | 228.2933778 | 0.308597513 | 0.349908216 | 0.88193846 | 0.377810088 | 0.681135691 |
| STX1B | 88.2787837 | 0.924704339 | 0.710276059 | 1.30189428 | 0.192952528 | 0.498908723 |
| STX2 | 492.997878 | 0.63682895 | 0.369921048 | 1.721526669 | 0.085155297 | 0.334340324 |
| STX3 | 2010.93991 | -0.283934622 | 0.263602803 | -1.077130511 | 0.281421964 | 0.595211573 |
| STX4 | 423.9822853 | -0.316101393 | 0.29728942 | -1.063278312 | 0.287655757 | |

| | | | | | | |
|---|---|---|---|---|---|---|
| | | | | | | 0.601673803 |
| STX5 | 996.4354977 | -0.135040383 | 0.272674352 | -0.49524417 | 0.620427775 | 0.833260426 |
| STX6 | 1396.335918 | 0.012192799 | 0.25120487 | 0.048537272 | 0.961288061 | 0.986785383 |
| STX7 | 1842.677579 | -0.083931225 | 0.226563468 | -0.370453477 | 0.711044634 | 0.876976263 |
| STX8 | 987.0699023 | 0.338376512 | 0.412773584 | 0.819763 | 0.412351228 | 0.70515068 |
| STXBP1 | 726.2743129 | 0.225336285 | 0.428743352 | 0.525573827 | 0.599184346 | 0.823816451 |
| STXBP2 | 1391.412076 | -0.596516267 | 0.31279227 | -1.907068442 | 0.056511725 | 0.275406162 |
| STXBP3 | 1463.331819 | 0.319804067 | 0.28175983 | 1.135023638 | 0.256365415 | 0.568552367 |
| STXBP4 | 394.7608982 | 0.162205058 | 0.31976879 | 0.507257316 | 0.611974284 | 0.829098987 |
| STXBP5 | 285.0936059 | -0.702244586 | 0.294622324 | -2.383541667 | 0.017146943 | 0.147741524 |
| STXBP5L | 15.01871301 | 0.057184972 | 0.809378777 | 0.070652917 | 0.943673994 | 0.97883756 |
| STXBP6 | 66.33297056 | 0.777766282 | 0.415125423 | 1.873569382 | 0.060989801 | 0.28478802 |
| STYK1 | 98.58742513 | -1.609775729 | 0.720119689 | -2.235428018 | 0.025389262 | 0.18194777 |
| STYX | 1009.57073 | 0.174384278 | 0.323242168 | 0.53948493 | 0.589552293 | 0.81828099 |
| STYXL1 | 661.7562151 | 0.116297169 | 0.35302419 | 0.329431162 | 0.741829817 | 0.892150449 |
| SUB1 | 6017.701395 | 1.004103784 | 0.327625975 | 3.064786868 | 0.002178252 | 0.04435739 |
| SUCLA2 | 1282.225415 | 0.537722341 | 0.247056799 | 2.176513026 | 0.029516921 | 0.196251879 |
| SUCLG1 | 1398.498864 | 0.409067475 | 0.311852881 | 1.31173223 | 0.189610498 | 0.495097939 |
| SUCLG2 | 1320.37955 | 0.613051987 | 0.231274078 | 2.650759622 | 0.008031098 | 0.100694036 |
| SUCO | 2656.186713 | -0.909836899 | 0.405821029 | -2.241965879 | 0.024963578 | 0.180688185 |
| SUDS3 | 2049.602599 | -0.31937675 | 0.235600868 | -1.35558393 | 0.175231588 | 0.479195921 |
| SUFU | 1121.610548 | -0.286177261 | 0.319748041 | -0.895008646 | 0.370782466 | 0.676136863 |
| SUGCT | 63.86827216 | 0.45799729 | 0.504189235 | 0.908383714 | 0.363675527 | 0.670353372 |
| SUGP2 | 3711.060483 | 0.116529139 | 0.472264619 | 0.246745434 | 0.805105242 | 0.918762309 |
| SUGT1 | 2425.26829 | 0.528843949 | 0.294570026 | 1.795308088 | 0.072604627 | 0.310190031 |
| SUGT1P3 | 19.22416671 | -0.01782175 | 0.86607527 | -0.020577599 | 0.98358261 | 0.994000251 |
| SULF1 | 741.9241123 | -1.022923352 | 0.32915504 | -3.107725015 | 0.001885334 | 0.040513318 |
| SULF2 | 2553.154333 | -0.631378236 | 0.313651057 | -2.012995725 | 0.044115079 | 0.241443505 |
| SULT1A1 | 167.4211654 | -0.672165001 | 0.413408073 | -1.625911647 | 0.103968452 | 0.369392362 |
| SULT1A2 | 32.07075749 | -1.226111372 | 0.62009334 | -1.977301307 | 0.048007582 | 0.252967897 |
| SULT1A3 | 672.6875754 | -0.164818727 | 0.399575756 | -0.412484302 | 0.679984484 | 0.861896952 |
| SULT1A4 | 123.2789735 | -1.298233108 | 0.411508171 | -3.154817328 | 0.001605986 | 0.037175747 |
| SULT1B1 | 47.17104808 | -0.035743363 | 0.6752499 | -0.052933534 | 0.957784866 | 0.98520931 |
| SULT1C2 | 140.8606133 | 0.463567866 | 0.459777423 | 1.008244082 | 0.3133373 | 0.623018103 |
| SULT1C4 | 137.9607608 | -0.867816211 | 0.383064317 | -2.265458231 | 0.023484574 | 0.174992377 |
| SULT2B1 | 15.45623851 | 0.880181224 | 0.849077931 | 1.036631848 | 0.299907465 | 0.610555175 |
| SULT4A1 | 188.2715562 | 0.144329036 | 0.365231906 | 0.395170942 | 0.692716736 | |

| | | | | | | |
|---|---|---|---|---|---|---|
| | | | | | | 0.86854217 |
| SUMF1 | 621.983375 | -0.218437006 | 0.490802623 | -0.445060796 | 0.656275819 | 0.84995394 |
| SUMF2 | 2000.445819 | 0.036976097 | 0.233283495 | 0.158502842 | 0.874060581 | 0.949740014 |
| SUMO1 | 4667.739971 | 0.601951822 | 0.279832661 | 2.151113524 | 0.03146724 | 0.202283687 |
| SUMO2 | 14848.33735 | 0.695230275 | 0.354866055 | 1.959134341 | 0.050097056 | 0.257487003 |
| SUMO3 | 2443.056518 | 0.34638171 | 0.257738353 | 1.343927694 | 0.17897177 | 0.483466568 |
| SUN1 | 2452.549086 | 0.056566638 | 0.303007842 | 0.186683743 | 0.851908603 | 0.939761856 |
| SUN2 | 560.8078084 | 0.480326879 | 0.28112253 | 1.708603287 | 0.087524459 | 0.340034206 |
| SUOX | 876.8197664 | -0.281526625 | 0.245572376 | -1.146409988 | 0.251625552 | 0.564383659 |
| SUPT16H | 8398.225381 | -0.137881078 | 0.33149715 | -0.41593443 | 0.677457984 | 0.860419848 |
| SUPT20H | 1678.970186 | -0.426471899 | 0.376629006 | -1.132339496 | 0.257491731 | 0.570028755 |
| SUPT3H | 671.7274288 | 0.04713051 | 0.251305577 | 0.187542634 | 0.851235199 | 0.939477675 |
| SUPT4H1 | 2028.179474 | 0.251105027 | 0.395907963 | 0.634251016 | 0.525917019 | 0.778458782 |
| SUPT5H | 3612.148854 | -0.49760807 | 0.321952593 | -1.545594229 | 0.122202596 | 0.40285432 |
| SUPT6H | 3083.981375 | -0.925361566 | 0.451619111 | -2.04898673 | 0.040463413 | 0.231601821 |
| SUPT7L | 1348.586538 | -0.327205161 | 0.294054602 | -1.112736067 | 0.265821801 | 0.579572403 |
| SUPV3L1 | 799.4657035 | 0.070495289 | 0.281884384 | 0.250085827 | 0.802520976 | 0.917815781 |
| SURF1 | 332.6291464 | 0.354502882 | 0.414840468 | 0.854552315 | 0.392799036 | 0.692011554 |
| SURF2 | 171.3291677 | 0.751488092 | 0.462558327 | 1.62463423 | 0.104240519 | 0.37001238 |
| SURF4 | 2318.676567 | -0.03391619 | 0.227916793 | -0.148809527 | 0.881703933 | 0.953217539 |
| SURF6 | 474.5216886 | 0.197982926 | 0.299209042 | 0.661687641 | 0.508171427 | 0.768714727 |
| SUSD1 | 206.0776749 | -1.102083317 | 0.40404948 | -2.727594937 | 0.00637979 | 0.086649766 |
| SUSD2 | 37.75970365 | -1.902796114 | 0.69369744 | -2.742976988 | 0.006088495 | 0.084270328 |
| SUSD3 | 39.25301271 | 1.290680062 | 0.542617519 | 2.378618486 | 0.017377651 | 0.148986942 |
| SUSD4 | 158.5249874 | -1.595069578 | 0.464238101 | -3.435886828 | 0.000590618 | 0.019567812 |
| SUSD5 | 78.57730553 | -0.24028674 | 0.399927035 | -0.600826449 | 0.547955586 | 0.792941769 |
| SUSD6 | 312.5879556 | -0.013674225 | 0.38324802 | -0.035679831 | 0.971537653 | 0.989665121 |
| SUV39H1 | 642.0992288 | -0.120652557 | 0.304746214 | -0.395911587 | 0.692170253 | 0.868131192 |
| SUV39H2 | 878.782556 | 0.625713977 | 0.289239195 | 2.163309772 | 0.030517362 | 0.199123233 |
| SUZ12 | 1474.84452 | 0.201649462 | 0.24931873 | 0.808801897 | 0.418629105 | 0.709770021 |
| SUZ12P1 | 578.5829836 | 0.722450348 | 0.323020111 | 2.236549129 | 0.025315823 | 0.181849597 |
| SV2A | 2184.548542 | -0.025880735 | 0.286154599 | -0.090443191 | 0.927935036 | 0.972633911 |
| SVBP | 767.0137071 | 0.903513413 | 0.422770052 | 2.137127283 | 0.03258764 | 0.205823103 |
| SVEP1 | 151.8606314 | -1.140753041 | 0.836922744 | -1.363032669 | 0.172872215 | 0.476584476 |
| SVIL | 1434.394524 | -0.66485261 | 0.353865828 | -1.878826826 | 0.060268143 | 0.283904551 |
| SVIL-AS1 | 51.14618104 | 0.001803939 | 0.580546743 | 0.00310731 | 0.997520729 | 0.99877158 |
| SVIL2P | 12.22799263 | -0.81899012 | 1.2940458 | -0.632891139 | 0.526804735 | 0.778755887 |
| SVIP | 388.8821477 | 0.517711251 | 0.341272906 | 1.517000743 | 0.129266496 | 0.413472911 |
| SVOP | 12.7158551 | 0.318010317 | 0.91580093 | 0.347248301 | 0.728404787 | 0.885679973 |
| SVOPL | 77.73960337 | -1.253626652 | 0.750417884 | -1.670571395 | 0.094806366 | |

| | | | | | | |
|---|---|---|---|---|---|---|
| | | | | | | 0.353479328 |
| SWAP70 | 1371.477342 | -0.174529763 | 0.302055036 | -0.577807823 | 0.563393873 | 0.801924557 |
| SWI5 | 438.063257 | 0.415320034 | 0.415152737 | 1.000402976 | 0.31711553 | 0.627198876 |
| SWSAP1 | 87.48564564 | 0.394737268 | 0.638954964 | 0.617785745 | 0.536716581 | 0.785348759 |
| SWT1 | 351.6318529 | -0.271765471 | 0.287483299 | -0.945326121 | 0.344492407 | 0.651368096 |
| SYAP1 | 1046.417184 | 0.24315309 | 0.341989443 | 0.710995893 | 0.477086781 | 0.749806646 |
| SYBU | 158.6809511 | -1.851358423 | 0.639722095 | -2.894004191 | 0.003803631 | 0.063596264 |
| SYCE1L | 17.13842341 | 1.10410668 | 1.156161596 | 0.95497609 | 0.339589789 | 0.646575889 |
| SYCE2 | 23.22513052 | 1.219936766 | 0.695257012 | 1.754655824 | 0.079318197 | 0.322931692 |
| SYCE3 | 24.56335237 | 1.440441816 | 0.777762526 | 1.852032938 | 0.064021093 | 0.292197354 |
| SYCP2 | 31.77149262 | -0.482699786 | 0.563258579 | -0.856977246 | 0.39145746 | 0.690959992 |
| SYCP2L | 120.0186685 | -0.576900575 | 0.367803915 | -1.568500363 | 0.116764405 | 0.39401268 |
| SYDE1 | 266.9282668 | 0.576124183 | 0.394624161 | 1.45993135 | 0.144308942 | 0.435597088 |
| SYDE2 | 775.1733601 | -2.0264336 | 0.466002066 | -4.348550677 | 1.37E-05 | 0.001395326 |
| SYF2 | 1565.596893 | 0.353734291 | 0.415277361 | 0.851802494 | 0.394323722 | 0.693260965 |
| SYK | 226.4031909 | -2.599512236 | 0.616523915 | -4.216401298 | 2.48E-05 | 0.002151957 |
| SYMPK | 2220.212682 | -0.179630494 | 0.402423129 | -0.446372192 | 0.655328417 | 0.849452103 |
| SYN1 | 54.04753865 | -0.707814419 | 0.65895113 | -1.07415313 | 0.282754055 | 0.596532022 |
| SYN2 | 104.1816988 | 0.346265975 | 0.504693607 | 0.686091463 | 0.492655434 | 0.759174768 |
| SYN3 | 68.48003333 | -0.255225948 | 0.600303442 | -0.425161561 | 0.670718903 | 0.858188946 |
| SYNCRIP | 6020.493386 | -0.024822367 | 0.241666021 | -0.102713518 | 0.918190345 | 0.969677217 |
| SYNDIG1 | 159.2594519 | -0.417851701 | 0.411020432 | -1.016620265 | 0.30933411 | 0.61913267 |
| SYNDIG1L | 20.62588553 | 0.578833527 | 0.70143299 | 0.825215715 | 0.409249133 | 0.703508181 |
| SYNE1 | 242.2276003 | -1.43208316 | 0.377343961 | -3.795166504 | 0.000147544 | 0.007793292 |
| SYNE2 | 3895.435953 | -1.146475953 | 0.33952749 | -3.376680786 | 0.000733661 | 0.022388677 |
| SYNE3 | 48.71617247 | -3.932571464 | 1.079241319 | -3.6438296 | 0.000268611 | 0.01147916 |
| SYNE4 | 266.0352782 | -0.098433687 | 0.630240557 | -0.156184311 | 0.875887743 | 0.95018528 |
| SYNGAP1 | 2296.221183 | 0.303620692 | 0.46183173 | 0.657427093 | 0.510906336 | 0.770599585 |
| SYNGR1 | 742.0902356 | 0.188226975 | 0.307931177 | 0.611263128 | 0.541025397 | 0.788008477 |
| SYNGR2 | 790.2456371 | -0.036509701 | 0.343848038 | -0.106179755 | 0.915439733 | 0.969148725 |
| SYNGR3 | 170.6937525 | 0.888531756 | 0.540504053 | 1.643894715 | 0.100197952 | 0.36243601 |
| SYNJ1 | 254.333813 | -0.459074617 | 0.389870886 | -1.177504227 | 0.238994311 | 0.550165913 |
| SYNJ2 | 195.1029628 | -1.34849183 | 0.401053021 | -3.362377938 | 0.000772743 | 0.023268821 |
| SYNJ2BP | 1123.557664 | -0.04499758 | 0.306469514 | -0.14682563 | 0.883269654 | 0.953742628 |
| SYNJ2BP-COX16 | 253.5085967 | 1.071813848 | 0.361225011 | 2.967163995 | 0.003005606 | 0.054218845 |
| SYNM | 28.31760824 | 0.744376321 | 0.687811739 | 1.082238467 | 0.27914658 | 0.592712773 |
| SYNPO | 173.2957433 | -1.807357929 | 0.425267725 | -4.249929687 | 2.14E-05 | |

0.001966978
SYNPO2 32.6293336 -2.665594972 0.881829454 -3.022801019
0.002504469 0.048313917
SYNPR 69.22890332 0.295603184 0.823999151 0.358742098 0.719788032 0.881675978
SYNRG 1243.949637 -0.69535825 0.390900668 -1.778861761 0.075262429
0.314600339
SYP 44.07363788 1.256851533 0.657163502 1.912540075 0.055806958 0.273376608
SYPL1 2152.048602 0.100288241 0.338332193 0.296419446 0.766909776 0.903470997
SYS1 678.4510323 0.078033242 0.260699776 0.299322244 0.764694184 0.902893924
SYS1-DBNDD2 23.12232444 -0.046139558 0.848507527 -0.054377311
0.956634555 0.984539826
SYT1 382.0795913 0.341997295 0.395444582 0.864842535 0.387125221 0.68748084
SYT10 18.05054239 1.41872258 1.28250369 1.106213254 0.268634228 0.581542504
SYT11 61.16049419 1.446711642 0.843424098 1.71528374 0.086293218 0.336976683
SYT12 48.76873927 0.235637411 0.512512748 0.459768877 0.645682125 0.845794147
SYT13 215.8539789 -0.21874868 0.361696994 -0.604784346 0.545322296
0.79117123
SYT14 212.9318685 0.097091615 0.431089932 0.225223574 0.821805351 0.92563193
SYT15 90.3546545 -0.106456418 0.433710477 -0.24545503 0.806104124
0.919089559
SYT16 11.22956474 -0.787100854 1.067189253 -0.737545709 0.460790557
0.73857935
SYT17 139.2050171 -1.637641282 0.651284696 -2.514478372 0.011920861
0.123877456
SYT2 33.41783882 -0.974287538 0.846258501 -1.151288333 0.249613635
0.562437021
SYT6 322.7431542 -0.893725786 0.410231288 -2.178590008 0.029362139
0.195861365
SYT7 2407.827142 -2.004366515 0.456618564 -4.389586131 1.14E-05
0.001196471
SYT9 12.46138386 0.015734215 1.02217953 0.015392809 0.9877188 0.995541774
SYTL1 453.8705954 -0.068251534 0.406680331 -0.167826001 0.866720167
0.946040122
SYTL2 95.66853864 0.934415279 0.493277872 1.894297985 0.058185468 0.279509071
SYTL3 15.97803657 -0.184369911 0.75348738 -0.244688785 0.806697413
0.919137032
SYTL4 19.96925249 -0.752090337 0.7726138 -0.973436323 0.330336493
0.638816405
SYTL5 26.99329941 -0.882425111 1.157253971 -0.762516382 0.445751869
0.728439722
SYVN1 685.6173297 -0.333301982 0.281681337 -1.183259016 0.23670651
0.547336452
SZRD1 2551.778028 0.01013116 0.264131325 0.038356526 0.969403423 0.989496332
SZT2 839.5234797 -0.939299103 0.482794745 -1.945545418 0.051709369
0.262081261
TAB1 1006.156695 -0.246645921 0.364529407 -0.676614607 0.498650509
0.762775559
TAB2 2454.129099 -0.273870901 0.290637892 -0.942309687 0.34603411
0.652702162
TAB3 930.0998393 -0.86541729 0.334185458 -2.589631802 0.009607864
0.111098245
TAC3 431.5520222 0.504367787 0.499279338 1.010191587 0.312403509 0.622795484
TAC4 12.95836434 -0.535492901 0.928319142 -0.576841387 0.564046608
0.802433524
TACC1 1721.806709 -2.179662186 0.611449794 -3.564744329 0.000364211
0.014277637
TACC2 897.0417018 -0.46976315 0.369033247 -1.272956174 0.203033594
0.509506436
TACC3 3143.409385 0.797648666 0.28183929 2.830154252 0.004652557 0.072024426
TACO1 456.3003591 0.540000324 0.333579166 1.618807105 0.105488773 0.372436233
TACR1 38.01226306 -1.326083633 1.081931313 -1.225663421 0.220325361
0.528885451
TACR3 18.96639135 -0.695391659 0.780876866 -0.890526649 0.373183166
0.678228049

| | | | | | | |
|---|---|---|---|---|---|---|
| TACSTD2 | 1131.387446 | 0.015797775 | 1.462790777 | 0.01079975 | 0.991383214 | 0.996814923 |
| TADA1 | 880.4491424 | 0.67075135 | 0.25694447 | 2.610491478 | 0.009041222 | 0.1074956 |
| TADA2A | 519.0260338 | 0.132104863 | 0.272074125 | 0.485547324 | 0.627288152 | 0.836041947 |
| TADA2B | 471.4212155 | -0.267628683 | 0.293188409 | -0.912821499 | 0.361336429 | 0.668292036 |
| TADA3 | 2184.364342 | 0.119451319 | 0.303418055 | 0.3936856 | 0.693813174 | 0.868802318 |
| TAF1 | 1391.98458 | -0.538077586 | 0.326948121 | -1.645758307 | 0.099813529 | 0.362265802 |
| TAF10 | 1730.701533 | 0.813557411 | 0.454658759 | 1.789380268 | 0.073553595 | 0.311651561 |
| TAF11 | 1809.220687 | 0.506294675 | 0.450891175 | 1.122875547 | 0.261490354 | 0.574937735 |
| TAF12 | 946.0741719 | 0.248806709 | 0.309040534 | 0.805094095 | 0.420765383 | 0.711245864 |
| TAF13 | 553.852343 | -0.204329515 | 0.419839443 | -0.486684894 | 0.626481652 | 0.835767952 |
| TAF15 | 8319.287591 | -0.023471943 | 0.287343113 | -0.081686118 | 0.934896318 | 0.975224148 |
| TAF1A | 335.3444539 | 0.6604348 | 0.292716678 | 2.256225385 | 0.02405652 | 0.177207433 |
| TAF1B | 888.5155272 | -0.145891779 | 0.23643715 | -0.617042536 | 0.53720667 | 0.785675856 |
| TAF1C | 1759.647887 | -0.489870312 | 0.400755671 | -1.222366514 | 0.221569057 | 0.530611548 |
| TAF1D | 5021.574172 | 0.442721521 | 0.282217397 | 1.568725121 | 0.116712002 | 0.394000398 |
| TAF2 | 1170.477818 | -0.15586409 | 0.359394933 | -0.433684719 | 0.664517405 | 0.85500751 |
| TAF3 | 441.3434452 | 0.184161634 | 0.265161436 | 0.694526463 | 0.487352111 | 0.756439734 |
| TAF4 | 190.3778822 | -0.999392568 | 0.842052678 | -1.186852788 | 0.235285693 | 0.545881517 |
| TAF4B | 229.569428 | 0.056011409 | 0.294865922 | 0.189955518 | 0.849343987 | 0.938580023 |
| TAF5 | 348.5068362 | 0.094193035 | 0.276439613 | 0.340736386 | 0.733302044 | 0.888457729 |
| TAF5L | 715.5126616 | -0.200476668 | 0.383054667 | -0.523363075 | 0.600721615 | 0.824475293 |
| TAF6 | 2048.222754 | -0.472227024 | 0.347012311 | -1.360836515 | 0.17356536 | 0.477540818 |
| TAF6L | 488.4017763 | -0.294882269 | 0.355557839 | -0.829351056 | 0.406905788 | 0.701626421 |
| TAF7 | 5515.477487 | 0.193135201 | 0.298456483 | 0.64711344 | 0.517558527 | 0.773950969 |
| TAF8 | 732.9245927 | -0.170041676 | 0.235961902 | -0.72063191 | 0.471136015 | 0.746018629 |
| TAF9 | 1600.796702 | 0.291414793 | 0.261413221 | 1.114766852 | 0.264950345 | 0.57877058 |
| TAF9B | 1795.968262 | 0.19813298 | 0.29802171 | 0.664827336 | 0.506160936 | 0.767209743 |
| TAGLN | 1150.223696 | 0.931141428 | 0.44815458 | 2.077723781 | 0.037734807 | 0.22292631 |
| TAGLN2 | 2326.467017 | -0.31511679 | 0.440144834 | -0.715938859 | 0.474029102 | 0.748101062 |
| TAGLN3 | 288.2192969 | 1.580790113 | 0.589459993 | 2.6817598 | 0.007323602 | 0.094825773 |
| TALDO1 | 4042.935366 | 0.348362807 | 0.335558873 | 1.038157042 | 0.299196948 | 0.610246278 |
| TAMM41 | 399.0104243 | 0.370392124 | 0.39704318 | 0.932876179 | 0.350883903 | 0.657923581 |
| TANC1 | 2358.42598 | -1.049211398 | 0.295663339 | -3.548669238 | 0.000387183 | 0.014833219 |
| TANC2 | 1019.820794 | -1.783267591 | 0.38165125 | -4.672505574 | 2.98E-06 | 0.000429914 |
| TANGO2 | 249.6084351 | -0.513981291 | 0.356186767 | -1.443010628 | 0.149017477 | 0.443544075 |
| TANGO6 | 395.7683804 | -0.200673965 | 0.302856548 | -0.662604017 | 0.507584197 | 0.768444894 |
| TANK | 471.8954601 | -0.117736897 | 0.269742634 | -0.436478637 | 0.662489495 | 0.853902351 |
| TAOK1 | 3120.874605 | -0.238461207 | 0.403203337 | -0.59141675 | 0.55424122 | 0.796360966 |
| TAOK2 | 1260.596435 | -0.108907991 | 0.519991817 | -0.20944174 | 0.834103412 | 0.932107564 |

TAOK3 266.9139586 -0.990462078 0.414687564 -2.388453774 0.016919437 0.147350616
TAP1 109.8565397 -0.25298823 0.405316605 -0.624174355 0.532513087 0.782597698
TAP2 42.81840944 0.53693022 0.965934207 0.555866245 0.578302295 0.810209453
TAPBPL 10.99731029 -0.901274889 1.248594788 -0.721831373 0.470398159 0.7454951
TAPT1 236.1574017 0.282880685 0.346625472 0.816098952 0.414443543 0.706780912
TARBP1 1745.616243 0.290973968 0.391044876 0.744093545 0.456819884 0.735735031
TARBP2 1387.367995 0.23082536 0.277601888 0.831497801 0.40569247 0.701371854
TARDBP 5554.095121 0.155982694 0.247029833 0.631432619 0.527757694 0.779156479
TARS 2836.535627 -0.250463368 0.256611643 -0.976040546 0.329044372 0.638424836
TARS2 1148.641884 -0.020939537 0.241051386 -0.086867525 0.930776813 0.973983655
TARSL2 380.4763683 0.201343092 0.268666984 0.74941509 0.453607059 0.733497409
TAS2R5 18.63921303 -1.283612074 0.959385682 -1.337952086 0.180912056 0.485600992
TASOR 1312.749127 -0.076442283 0.426623865 -0.179179576 0.8577967 0.942343063
TASOR2 1167.326002 -0.354955911 0.476330987 -0.745187529 0.456158361 0.735399427
TASP1 245.6289947 -0.309827574 0.376288583 -0.823377558 0.410293321 0.704135744
TATDN1 1403.916507 0.818174162 0.308471861 2.652346182 0.007993454 0.100486153
TATDN2 662.26548 -0.212107351 0.308390129 -0.687789042 0.491585638 0.758965432
TATDN2P3 13.1482282 -3.84355773 1.335162301 -2.878719485 0.003992933 0.065559297
TATDN3 531.2905014 0.486697573 0.282514267 1.722736262 0.08493623 0.333912294
TAX1BP1 3673.091183 -0.132785062 0.226915775 -0.58517334 0.558431169 0.798855459
TAX1BP3 1584.491221 -0.029728742 0.415563283 -0.071538423 0.942969246 0.978574897
TAZ 665.1173608 0.128304705 0.267084649 0.480389665 0.630950341 0.838320891
TBC1D1 5324.174644 -1.179542632 0.222091726 -5.311060678 1.09E-07 4.03E-05
TBC1D10A 232.3647191 -0.227228234 0.476734191 -0.476635068 0.633622007 0.839689885
TBC1D10B 1035.361969 -0.241798985 0.302625251 -0.799004658 0.424287711 0.713303793
TBC1D12 179.4540812 -0.034738376 0.351823951 -0.09873795 0.921346337 0.970205128
TBC1D13 1019.638946 -0.604929503 0.256902904 -2.354700919 0.018537619 0.154302741
TBC1D14 2339.920563 -0.424814332 0.387939127 -1.095054101 0.273492935 0.586675071
TBC1D15 928.9596384 0.169169758 0.266622885 0.634490764 0.525760592 0.778438656
TBC1D16 1276.758974 0.026059792 0.486630286 0.05355152 0.957292482 0.984949762
TBC1D17 820.4766097 -0.462640569 0.301872749 -1.532568179 0.125382287 0.407317918
TBC1D19 166.9304116 0.362906841 0.395611711 0.917330886 0.358969277 0.666738847
TBC1D2 159.3353052 -1.619585025 0.494244647 -3.27688936 0.001049575 0.027896179
TBC1D20 917.0132132 -0.37216248 0.335978665 -1.107696765 0.267992797

0.581149046

| | | | | | | |
|---|---|---|---|---|---|---|
| TBC1D22A | 603.1380405 | -0.212015996 | 0.310959383 | -0.681812504 | 0.495357517 | 0.760859008 |
| TBC1D22B | 555.2553089 | -0.474786857 | 0.282594735 | -1.680098027 | 0.092938245 | 0.349595587 |
| TBC1D23 | 917.0897601 | -0.142977471 | 0.255137583 | -0.56039361 | 0.575210989 | 0.808460804 |
| TBC1D24 | 240.1113231 | 0.038270563 | 0.33421009 | 0.114510495 | 0.908833127 | 0.965877925 |
| TBC1D25 | 301.0128721 | -0.372287298 | 0.271929549 | -1.369057904 | 0.170981176 | 0.474214884 |
| TBC1D26 | 96.07154321 | 0.776378239 | 0.541813567 | 1.432925062 | 0.151879254 | 0.44767052 |
| TBC1D29P | 16.32080415 | -0.831394121 | 1.04110914 | -0.79856577 | 0.424542243 | 0.713390792 |
| TBC1D2B | 814.2557415 | -0.512751187 | 0.359437336 | -1.426538472 | 0.15371297 | 0.450536992 |
| TBC1D3 | 15.56523681 | -0.442849641 | 4.183070758 | -0.105867117 | 0.915687784 | 0.969158739 |
| TBC1D30 | 24.70828514 | 0.087160646 | 0.613199324 | 0.142140806 | 0.886968787 | 0.955628398 |
| TBC1D31 | 563.8948684 | 0.684166833 | 0.351138639 | 1.948423663 | 0.051364291 | 0.260792387 |
| TBC1D32 | 278.1749539 | 0.227255408 | 0.357972049 | 0.634841209 | 0.525531984 | 0.778371208 |
| TBC1D3B | 22.85645674 | -1.308795849 | 0.881720666 | -1.484365626 | 0.137711949 | 0.425851605 |
| TBC1D3C | 212.174494 | -1.811208055 | 0.833456845 | -2.173127579 | 0.029770717 | 0.197226575 |
| TBC1D3F | 32.21924001 | -2.401024899 | 1.196946597 | -2.005958249 | 0.044860703 | 0.2438487 |
| TBC1D3G | 27.37484209 | 8.917609319 | 1.571685075 | 5.673916142 | 1.40E-08 | 1.03E-05 |
| TBC1D3I | 11.62968131 | -6.156163218 | 1.706255154 | -3.607996849 | 0.00030857 | 0.012860767 |
| TBC1D3L | 323.0135188 | -0.674720384 | 0.438915417 | -1.537244667 | 0.124233406 | 0.40567047 |
| TBC1D3P5 | 22.66906401 | -2.028440448 | 1.052970036 | -1.926399022 | 0.054054569 | 0.268784619 |
| TBC1D4 | 680.8646004 | 0.159814977 | 0.272995391 | 0.585412731 | 0.558270231 | 0.798855459 |
| TBC1D5 | 1296.780948 | -0.521031938 | 0.357897325 | -1.455814006 | 0.145444043 | 0.437196666 |
| TBC1D7 | 837.4511517 | 0.986388678 | 0.422867282 | 2.332619997 | 0.019668098 | 0.158893482 |
| TBC1D8 | 312.0295952 | -0.73932363 | 0.465004136 | -1.589929147 | 0.111850777 | 0.384888109 |
| TBC1D8B | 350.2242995 | -0.102321948 | 0.350687783 | -0.291775057 | 0.770458617 | 0.905042794 |
| TBC1D9 | 373.1482533 | -1.427594444 | 0.366621237 | -3.89392185 | 9.86E-05 | 0.006082981 |
| TBC1D9B | 961.6718309 | -0.817188756 | 0.433031706 | -1.887133771 | 0.059142335 | 0.281976232 |
| TBCA | 4493.354763 | 0.5461517 | 0.501499033 | 1.089038392 | 0.276136956 | 0.589457402 |
| TBCB | 2462.764798 | 0.857426429 | 0.388140812 | 2.209060223 | 0.027170452 | 0.188528766 |
| TBCC | 481.0418277 | 0.224945699 | 0.340994099 | 0.659676221 | 0.509461629 | 0.769564758 |
| TBCCD1 | 319.9004851 | -0.158715885 | 0.403415192 | -0.39343061 | 0.694001465 | 0.868919389 |
| TBCD | 2404.767963 | 0.05103792 | 0.541745658 | 0.094210113 | 0.924942251 | 0.97180348 |
| TBCE | 986.3909891 | 0.425694651 | 0.272072 | 1.564639699 | 0.117667415 | 0.395641036 |
| TBCEL | 455.2632832 | -0.594588726 | 0.287096391 | -2.071042145 | 0.038354858 | 0.225585886 |
| TBCK | 786.9725649 | -0.888518691 | 0.394680419 | -2.251235804 | 0.024370606 | 0.178492903 |
| TBK1 | 1005.733429 | 0.12385404 | 0.324342091 | 0.381862373 | 0.702563451 | 0.872703476 |

| | | | | | | |
|---|---|---|---|---|---|---|
| TBKBP1 | 376.2826575 | 0.28405772 | 0.43918575 | 0.646782643 | 0.517772627 | 0.773950969 |
| TBL1X | 2642.088571 | -1.022704148 | 0.452186281 | -2.261687697 | 0.023716705 | 0.17620281 |
| TBL1XR1 | 2251.129224 | 0.202656742 | 0.332868245 | 0.60881969 | 0.542643961 | 0.789277749 |
| TBL1Y | 122.1812668 | -0.93078046 | 0.450417853 | -2.066482165 | 0.038782975 | 0.226822093 |
| TBL2 | 886.3295107 | 0.288133698 | 0.24327752 | 1.184382751 | 0.236261585 | 0.547141583 |
| TBL3 | 763.0820104 | 0.216517407 | 0.428718506 | 0.505033965 | 0.61353498 | 0.830176524 |
| TBP | 714.5354869 | 0.100676479 | 0.368790987 | 0.272990618 | 0.784860428 | 0.910931322 |
| TBPL1 | 1117.897841 | 0.732877646 | 0.275264737 | 2.662446535 | 0.007757489 | 0.098649513 |
| TBRG1 | 888.0043246 | -0.376624787 | 0.263963861 | -1.426804356 | 0.153636295 | 0.450399273 |
| TBRG4 | 1489.502771 | 0.22305017 | 0.310103028 | 0.719277627 | 0.471969878 | 0.746399031 |
| TBX15 | 217.5042456 | 0.463267464 | 0.446755571 | 1.036959569 | 0.2997547 | 0.610495175 |
| TBX19 | 67.26240632 | 0.211322249 | 0.406105632 | 0.520362764 | 0.602810758 | 0.825021776 |
| TBX3 | 516.7901034 | -5.048093501 | 0.934746671 | -5.400493695 | 6.65E-08 | 3.12E-05 |
| TBX6 | 74.98151231 | -0.37123835 | 0.520206941 | -0.713635903 | 0.475452351 | 0.74892861 |
| TBXAS1 | 14.28802221 | 1.501244878 | 0.882510785 | 1.701106551 | 0.088922981 | 0.342496199 |
| TC2N | 147.863115 | -2.689266127 | 0.532932879 | -5.046162909 | 4.51E-07 | 0.000113977 |
| TCAF1 | 5053.169035 | 0.112164784 | 0.220701583 | 0.508219212 | 0.611299619 | 0.829058038 |
| TCAF1P1 | 242.7158011 | -0.561290095 | 2.259743642 | -0.248386624 | 0.803835282 | 0.917969611 |
| TCAF2 | 37.95371484 | -2.078706956 | 0.696653959 | -2.983844314 | 0.002846516 | 0.052472041 |
| TCAIM | 960.4877853 | -0.021653793 | 0.31553214 | -0.068626268 | 0.945287109 | 0.979754792 |
| TCAP | 34.81833157 | 0.49956617 | 0.537795781 | 0.92891426 | 0.352933522 | 0.660053559 |
| TCEA1 | 5358.16201 | 0.146441874 | 0.237072071 | 0.617710358 | 0.536766283 | 0.785348759 |
| TCEA2 | 789.8153176 | 0.739093607 | 0.354957797 | 2.082201359 | 0.037324081 | 0.22118892 |
| TCEA3 | 15.30231795 | 0.253718886 | 0.793474264 | 0.319756919 | 0.749152608 | 0.895124377 |
| TCEAL1 | 159.7477435 | 0.787580188 | 0.407639847 | 1.932049071 | 0.05335345 | 0.267056023 |
| TCEAL2 | 89.70446138 | 1.021352056 | 0.387688489 | 2.634465775 | 0.00842698 | 0.103784183 |
| TCEAL3 | 163.5110408 | 0.762061507 | 0.431092934 | 1.76774298 | 0.077103881 | 0.31844659 |
| TCEAL4 | 1290.034476 | 0.063272083 | 0.353656573 | 0.178908262 | 0.858009735 | 0.942343063 |
| TCEAL5 | 108.8912811 | 1.878308663 | 0.377671254 | 4.973395888 | 6.58E-07 | 0.000139266 |
| TCEAL7 | 253.7287826 | 1.291030373 | 0.339429962 | 3.803525073 | 0.000142652 | 0.00767435 |
| TCEAL8 | 1740.141955 | 0.38773979 | 0.346632316 | 1.118591003 | 0.263314664 | 0.577025388 |
| TCEAL9 | 1990.3758 | 0.228995351 | 0.550262818 | 0.416156323 | 0.677295618 | 0.860358531 |
| TCEANC | 74.65877591 | 0.069688253 | 0.401054573 | 0.173762519 | 0.862052102 | 0.944228049 |
| TCEANC2 | 432.3123466 | 0.075473513 | 0.323665135 | 0.233183946 | 0.815618572 | 0.922975941 |
| TCERG1 | 3371.491399 | 0.161727583 | 0.286291693 | 0.564904911 | 0.572138443 | 0.807358601 |
| TCF12 | 2411.444609 | -0.221906184 | 0.35623671 | -0.622917789 | 0.533338547 | 0.783143103 |
| TCF15 | 13.27827163 | 2.092738739 | 0.945894518 | 2.212444093 | 0.026935995 | 0.187710602 |
| TCF20 | 187.6971683 | -0.645427354 | 0.539459259 | -1.196433916 | 0.231527273 | 0.540550978 |
| TCF21 | 16.05046215 | -3.072835211 | 1.112268061 | -2.762675042 | 0.005732981 | |

| | | | | | | |
|---|---|---|---|---|---|---|
| | | | | | | 0.081513644 |
| TCF25 | 3056.367322 | 0.136997101 | 0.315873878 | 0.433708231 | 0.664500328 | 0.85500751 |
| TCF3 | 4394.185382 | 0.308187603 | 0.333473307 | 0.924174729 | 0.355395351 | 0.662859954 |
| TCF4 | 2438.733406 | 0.123420973 | 0.355359326 | 0.347313167 | 0.72835606 | 0.885679973 |
| TCF7 | 198.020538 | -0.264813377 | 0.410501145 | -0.645097779 | 0.518863823 | 0.774467046 |
| TCF7L1 | 1923.348825 | -0.751626178 | 0.283458262 | -2.651629106 | 0.008010448 | 0.100518204 |
| TCF7L2 | 1888.897732 | -0.417848418 | 0.357330154 | -1.169362321 | 0.242257684 | 0.553231259 |
| TCFL5 | 484.7223664 | 0.672249045 | 0.349073186 | 1.925811183 | 0.054127955 | 0.268973206 |
| TCHH | 32.10558883 | -1.571486608 | 0.744564282 | -2.110612404 | 0.034805641 | 0.213146583 |
| TCHP | 654.5553416 | -0.459295743 | 0.340078462 | -1.350558162 | 0.176837011 | 0.481149663 |
| TCIM | 15.45573216 | -0.165441676 | 1.529270924 | -0.108183366 | 0.91385024 | 0.968086893 |
| TCIRG1 | 427.6571601 | -0.418936955 | 0.456343284 | -0.918030285 | 0.358603011 | 0.666384894 |
| TCL1B | 14.68663496 | 1.463997585 | 0.978957799 | 1.495465469 | 0.134793005 | 0.421290623 |
| TCN2 | 1716.715673 | -0.040449449 | 0.304781357 | -0.132716283 | 0.894417764 | 0.958237013 |
| TCOF1 | 3850.516311 | -0.020463977 | 0.248812287 | -0.082246648 | 0.934450579 | 0.97507153 |
| TCP1 | 8655.242506 | 0.348239077 | 0.266146004 | 1.308451271 | 0.190720292 | 0.496058205 |
| TCP10L | 26.81817904 | 0.661064402 | 0.591973227 | 1.116713344 | 0.264116911 | 0.577448862 |
| TCP11L1 | 331.5463771 | -0.460220495 | 0.305279461 | -1.507538349 | 0.131672722 | 0.417298279 |
| TCP11L2 | 95.0309069 | -1.831341823 | 0.540536721 | -3.388006316 | 0.000704026 | 0.02179752 |
| TCTA | 257.0089432 | -0.698279715 | 0.393673403 | -1.773753854 | 0.076103874 | 0.316520909 |
| TCTE3 | 26.11283739 | 0.54525445 | 0.732441985 | 0.744433636 | 0.456614176 | 0.735622968 |
| TCTEX1D2 | 388.1830942 | 1.760261085 | 0.435342024 | 4.043398037 | 5.27E-05 | 0.003917842 |
| TCTN1 | 432.8170094 | 0.199980015 | 0.2639237 | 0.757719046 | 0.448619194 | 0.729866144 |
| TCTN2 | 314.8244353 | -0.490646295 | 0.27973857 | -1.75394582 | 0.079439791 | 0.322931692 |
| TCTN3 | 1612.236837 | 0.07240776 | 0.231219577 | 0.313155838 | 0.754162269 | 0.897293999 |
| TDG | 2163.295591 | 0.376653955 | 0.292457843 | 1.287891449 | 0.197783753 | 0.503399198 |
| TDGF1 | 128.7805034 | -1.370338023 | 0.903896049 | -1.516034974 | 0.129510509 | 0.414078806 |
| TDH | 24.81781923 | -0.420367248 | 0.807898338 | -0.520321961 | 0.602839192 | 0.825021776 |
| TDP1 | 1195.368063 | 0.153189635 | 0.27955508 | 0.547976575 | 0.583707988 | 0.8142179 |
| TDP2 | 1384.56146 | 0.110046141 | 0.254343116 | 0.432668056 | 0.66525594 | 0.855087509 |
| TDRD3 | 811.2575447 | 0.216102371 | 0.239432118 | 0.902562167 | 0.366758313 | 0.673335968 |
| TDRD7 | 193.9875548 | -0.819469047 | 0.525134582 | -1.560493395 | 0.11864333 | 0.396986757 |
| TDRKH | 506.6533286 | -0.05945574 | 0.381626643 | -0.15579557 | 0.876194163 | 0.950296086 |
| TDRP | 2507.19194 | -0.760728596 | 0.58255715 | -1.305843721 | 0.191605708 | 0.49715242 |
| TEAD1 | 3381.517058 | -0.687994359 | 0.372973159 | -1.844621635 | 0.065092598 | 0.294255005 |
| TEAD2 | 4333.968868 | 0.120229517 | 0.220324576 | 0.545692717 | 0.585277179 | 0.815432095 |
| TEAD3 | 2596.153767 | -1.341271342 | 0.39366054 | -3.407177514 | 0.000656384 | 0.02093243 |
| TEAD4 | 632.169432 | -0.508829216 | 0.391307286 | -1.300331567 | 0.193487353 | 0.499461017 |
| TEC | 15.64934499 | -1.560844722 | 0.870605787 | -1.792826036 | 0.073000745 | 0.31078683 |
| TECPR1 | 683.2440617 | 0.046259048 | 0.3565371 | 0.129745398 | 0.896767864 | 0.95928311 |

| | | | | | | |
|---|---|---|---|---|---|---|
| TECPR2 | 534.8852374 | -0.501893352 | 0.522560561 | -0.960450116 | 0.336828726 | 0.644459616 |
| TECR | 3369.324298 | -0.058520185 | 0.250239403 | -0.233856795 | 0.815096156 | 0.92282267 |
| TECRP1 | 123.7715853 | 10.13870192 | 2.85074294 | 3.556512157 | 0.000375811 | 0.014581663 |
| TECTA | 69.33142285 | -0.842332642 | 0.502415528 | -1.676565702 | 0.093627438 | 0.351044935 |
| TEDC1 | 94.11557987 | 0.769840423 | 0.547989112 | 1.404846203 | 0.160067017 | 0.460355774 |
| TEDC2 | 405.0480833 | 0.767782256 | 0.33822854 | 2.270010261 | 0.02320696 | 0.174034996 |
| TEF | 793.3273118 | 0.253491014 | 0.434842558 | 0.582948953 | 0.559927665 | 0.800250834 |
| TEFM | 303.481659 | 0.405733816 | 0.316623466 | 1.281439502 | 0.200039337 | 0.506052489 |
| TEK | 748.4427822 | 0.673587584 | 0.680861522 | 0.989316568 | 0.32250828 | 0.632222913 |
| TEKT2 | 99.4447945 | 0.396220095 | 0.596258886 | 0.664510172 | 0.506363841 | 0.767347299 |
| TEKT3 | 42.49914222 | -0.857029419 | 0.953144213 | -0.899160282 | 0.368567293 | 0.67460916 |
| TEKT5 | 11.76903663 | -0.393229565 | 0.907463808 | -0.433328096 | 0.664776429 | 0.855037526 |
| TELO2 | 688.9950909 | 0.167536026 | 0.510445857 | 0.328215077 | 0.742749049 | 0.892392953 |
| TEN1 | 387.2924282 | -0.232576396 | 0.520334288 | -0.446974957 | 0.654893142 | 0.849258129 |
| TEN1-CDK3 | 129.4195258 | -1.038231234 | 0.599828546 | -1.730879999 | 0.08347317 | 0.330745685 |
| TENM1 | 26.83763778 | -0.413841887 | 0.726714201 | -0.569469933 | 0.56903727 | 0.805450586 |
| TENM2 | 217.4410169 | 0.710502959 | 0.86807397 | 0.818482046 | 0.41308199 | 0.705607651 |
| TENM3 | 2005.111402 | -0.656251363 | 0.431771506 | -1.519904288 | 0.128535033 | 0.413049138 |
| TENM4 | 509.4011518 | 0.335499506 | 0.758820988 | 0.442132612 | 0.658393246 | 0.851158424 |
| TENT2 | 1388.301386 | -0.275886583 | 0.25369739 | -1.087463231 | 0.27683214 | 0.590525927 |
| TENT4A | 748.3895979 | 0.103552989 | 0.382593527 | 0.270660587 | 0.786652093 | 0.911363031 |
| TENT4B | 459.7283424 | 0.366504437 | 0.273176478 | 1.341639806 | 0.179712813 | 0.484443366 |
| TENT5A | 171.4993893 | -0.008769972 | 0.389459508 | -0.022518316 | 0.982034501 | 0.993518428 |
| TENT5B | 1067.328426 | -0.675469966 | 0.568673048 | -1.187800211 | 0.234912131 | 0.545431886 |
| TENT5C | 40.21063242 | -1.077181028 | 0.968053759 | -1.112728522 | 0.265825042 | 0.579572403 |
| TEP1 | 835.8834711 | -0.843498824 | 0.256185144 | -3.292536051 | 0.000992882 | 0.027091741 |
| TEPP | 30.71230718 | 0.819150386 | 0.972081088 | 0.842677011 | 0.399409108 | 0.697529133 |
| TEPSIN | 359.2777271 | -0.578968036 | 0.433363473 | -1.335987161 | 0.18155347 | 0.485768051 |
| TERF1 | 1832.278613 | 0.310367382 | 0.245930955 | 1.262010231 | 0.206945105 | 0.513642439 |
| TERF2 | 1046.912552 | 0.156428641 | 0.246856416 | 0.633682705 | 0.526287914 | 0.778525486 |
| TERF2IP | 775.0913234 | 0.444175108 | 0.261003676 | 1.701796371 | 0.088793547 | 0.342301733 |
| TERT | 104.0073815 | 0.951187368 | 0.418920349 | 2.27056855 | 0.023173109 | 0.173867082 |
| TES | 857.5622195 | -1.704980654 | 0.595717275 | -2.862063477 | 0.004208926 | 0.067073127 |
| TESC | 26.86529308 | 1.106776307 | 0.80256368 | 1.379051077 | 0.167879006 | 0.470340241 |
| TESK1 | 261.1307519 | -0.393896778 | 0.523372403 | -0.752612815 | 0.451682615 | 0.73280686 |
| TESK2 | 573.407801 | -0.766192495 | 0.321017545 | -2.386762054 | 0.016997489 | 0.147691237 |
| TESMIN | 465.7388154 | -0.288672584 | 0.428644666 | -0.673454278 | 0.500658328 | 0.763638219 |
| TESPA1 | 24.46842997 | 0.59372643 | 0.79621103 | 0.745689782 | 0.455854833 | 0.735306344 |
| TET1 | 2298.244205 | -0.105837267 | 0.377242585 | -0.280554929 | 0.77905179 | 0.908367621 |

| | | | | | | |
|---|---|---|---|---|---|---|
| TET2 | 452.625437 | -1.202297805 | 0.444501936 | -2.704820174 | 0.006834137 | 0.090629982 |
| TET3 | 995.1045932 | -0.309314239 | 0.317809688 | -0.973268753 | 0.330419748 | 0.638817127 |
| TEX10 | 1622.96517 | -0.215325698 | 0.2399608 | -0.897336974 | 0.369539133 | 0.67521115 |
| TEX11 | 11.08192948 | 0.575349946 | 1.060535127 | 0.542509089 | 0.587467848 | 0.816787521 |
| TEX14 | 12.96924688 | -0.338906575 | 0.882743825 | -0.38392404 | 0.701034748 | 0.872239782 |
| TEX15 | 92.30380992 | 0.917534873 | 0.6563633 | 1.397907033 | 0.162140987 | 0.462724023 |
| TEX19 | 10.34579615 | 0.645201303 | 0.99138699 | 0.650806708 | 0.51517127 | 0.773232343 |
| TEX2 | 368.1298614 | -0.269827808 | 0.572144364 | -0.471607911 | 0.637206679 | 0.841584486 |
| TEX21P | 25.64445213 | 0.220592038 | 0.617627256 | 0.357160465 | 0.720971681 | 0.882227889 |
| TEX22 | 18.72118304 | 0.229158993 | 0.709021033 | 0.323204788 | 0.746540155 | 0.894109554 |
| TEX261 | 823.2299174 | -0.585903869 | 0.355719558 | -1.647094896 | 0.099538542 | 0.361873765 |
| TEX264 | 174.3492743 | -0.583157222 | 0.461393665 | -1.263903835 | 0.206264545 | 0.512653902 |
| TEX30 | 315.1696329 | 0.538980161 | 0.335669962 | 1.605684816 | 0.108343179 | 0.377595766 |
| TEX35 | 17.5462477 | 0.281161354 | 0.757672103 | 0.371085794 | 0.710573632 | 0.876838602 |
| TEX37 | 26.52211941 | -0.260680918 | 0.805880064 | -0.32347359 | 0.746336605 | 0.894109554 |
| TEX9 | 183.8468515 | 0.019938496 | 0.33612427 | 0.059318822 | 0.95269817 | 0.982955926 |
| TF | 345.4267219 | -0.447045658 | 0.319164176 | -1.400676175 | 0.16131093 | 0.46167973 |
| TFAM | 2367.734074 | 0.050264779 | 0.223487683 | 0.224910736 | 0.822048717 | 0.925651384 |
| TFAP2A | 9788.597628 | -0.82584806 | 0.595177021 | -1.387567113 | 0.165268901 | 0.467517155 |
| TFAP2B | 892.2451445 | -1.335791501 | 0.766957245 | -1.741676619 | 0.081565046 | 0.326496916 |
| TFAP2C | 2862.448398 | -0.944642958 | 0.586421262 | -1.610860689 | 0.10721009 | 0.375111686 |
| TFAP2D | 11.96751624 | -2.706202037 | 1.23488968 | -2.191452468 | 0.028419066 | 0.192647743 |
| TFAP2E | 26.33226724 | -0.477513516 | 0.880191838 | -0.54251073 | 0.587466718 | 0.816787521 |
| TFAP4 | 347.9572207 | 0.223467874 | 0.384576077 | 0.581075859 | 0.561189328 | 0.801299134 |
| TFB1M | 510.9572103 | 0.376949134 | 0.306541518 | 1.229683784 | 0.218815542 | 0.527263434 |
| TFB2M | 447.1095706 | 0.509636812 | 0.36394795 | 1.400301367 | 0.161423092 | 0.46167973 |
| TFCP2 | 1587.893441 | -0.087017565 | 0.274584189 | -0.316906685 | 0.751314409 | 0.89636357 |
| TFCP2L1 | 47.78144943 | -2.966577591 | 0.774183892 | -3.831877185 | 0.000127169 | 0.007145497 |
| TFDP1 | 2258.925195 | 0.16795318 | 0.240832352 | 0.697386286 | 0.485561082 | 0.755840687 |
| TFDP2 | 2167.167586 | 0.606874134 | 0.291842186 | 2.079459939 | 0.037575096 | 0.222242412 |
| TFE3 | 1188.295962 | -0.384427557 | 0.263097059 | -1.461162502 | 0.143970851 | 0.434995877 |
| TFEB | 47.7636256 | -1.084653495 | 0.802404289 | -1.351754358 | 0.176453909 | 0.480693528 |
| TFG | 2714.417998 | 0.104031294 | 0.329751456 | 0.315483955 | 0.752394232 | 0.896586461 |
| TFIP11 | 856.3552391 | -0.47857167 | 0.290904546 | -1.645115818 | 0.099945929 | 0.362399544 |
| TFPI | 730.0547714 | 0.568864938 | 0.448625461 | 1.268017506 | 0.204791699 | 0.511170593 |
| TFPT | 72.14311787 | 0.458149964 | 0.70642056 | 0.648551289 | 0.516628449 | 0.773950426 |
| TFR2 | 30.54628255 | 0.206383666 | 0.560614389 | 0.368138367 | 0.712770064 | 0.87771385 |
| TFRC | 1632.374315 | 0.163571394 | 0.409498603 | 0.399443107 | 0.689566736 | 0.867663346 |
| TGDS | 468.231261 | 0.425820937 | 0.281510875 | 1.512626956 | 0.130374443 | 0.415496314 |
| TGFA | 17.97291725 | -0.761151655 | 0.945843268 | -0.804733386 | 0.42097355 | 0.711518463 |
| TGFB1I1 | 10.27213062 | -0.444572778 | 1.041466242 | -0.426872 | 0.66947256 | 0.857712777 |
| TGFB2 | 18.8651489 | -1.434078423 | 1.118039532 | -1.282672376 | 0.199606882 | |

| | | | | | | |
|---|---|---|---|---|---|---|
| | | | | | | 0.505774645 |
| TGFB3 | 14.27914349 | -0.801085953 | 0.861842011 | -0.929504413 | 0.352627739 | 0.659729494 |
| TGFBI | 218.8276443 | -1.327724543 | 0.468945426 | -2.831298631 | 0.004635941 | 0.071913976 |
| TGFBR1 | 3229.19056 | 0.135748048 | 0.228688754 | 0.59359302 | 0.552784351 | 0.795672359 |
| TGFBR2 | 116.6910052 | -0.616508845 | 0.480990385 | -1.281748791 | 0.199930783 | 0.50598138 |
| TGFBR3 | 214.8454987 | -0.137123019 | 0.347759061 | -0.394304661 | 0.693356122 | 0.868737544 |
| TGFBRAP1 | 579.2095359 | -0.346973406 | 0.352622955 | -0.9839785 | 0.325126084 | 0.635057014 |
| TGIF1 | 2311.421092 | -0.244439079 | 0.290340362 | -0.841905265 | 0.399840983 | 0.697640892 |
| TGIF2 | 1957.24995 | 0.16277685 | 0.34549561 | 0.47114014 | 0.637540662 | 0.841782104 |
| TGIF2-RAB5IF | 88.28729428 | 0.100471848 | 0.686938543 | 0.146260315 | 0.883715893 | 0.953952882 |
| TGM1 | 157.3915347 | -2.006595204 | 0.455814023 | -4.402223498 | 1.07E-05 | 0.001152231 |
| TGM2 | 65.37619786 | -0.9527211 | 0.620115216 | -1.536361429 | 0.124449761 | 0.405999855 |
| TGOLN2 | 2575.894093 | -0.361850908 | 0.221845668 | -1.631092964 | 0.102870705 | 0.367817927 |
| TGS1 | 1123.173668 | -0.197807636 | 0.240910805 | -0.821082459 | 0.411599301 | 0.704686727 |
| THADA | 837.7313264 | -0.15685647 | 0.280627257 | -0.558949519 | 0.576196173 | 0.808993188 |
| THAP1 | 348.3352914 | 0.125100551 | 0.269535177 | 0.464134412 | 0.642551448 | 0.844507322 |
| THAP10 | 118.9441685 | 0.724579845 | 0.347814232 | 2.083238058 | 0.03722953 | 0.220973863 |
| THAP11 | 572.0670567 | 1.235653506 | 0.322903809 | 3.826692258 | 0.000129877 | 0.007243505 |
| THAP12 | 2059.796538 | 0.236734029 | 0.220380985 | 1.074203518 | 0.282731476 | 0.596532022 |
| THAP2 | 228.1861097 | 1.214643761 | 0.312631131 | 3.885229717 | 0.000102233 | 0.00625395 |
| THAP3 | 315.9105121 | 0.952187337 | 0.325237816 | 2.927664897 | 0.003415179 | 0.059485284 |
| THAP4 | 1030.75546 | 0.037085445 | 0.308536351 | 0.120197977 | 0.904326321 | 0.963992033 |
| THAP5 | 599.7895999 | 0.426248731 | 0.283544717 | 1.503285746 | 0.132765374 | 0.418836243 |
| THAP6 | 296.5090709 | -0.113168733 | 0.306605012 | -0.369102684 | 0.712051189 | 0.877469627 |
| THAP7 | 268.0700754 | 0.362068312 | 0.477724431 | 0.757902021 | 0.448509641 | 0.729796208 |
| THAP8 | 153.0786094 | 0.447999019 | 0.406719546 | 1.101493705 | 0.270681843 | 0.583311682 |
| THAP9 | 472.1806569 | 0.238795989 | 0.327955455 | 0.728135438 | 0.466530682 | 0.743017201 |
| THBD | 11.18351591 | -2.171835323 | 1.181092046 | -1.838836634 | 0.065939218 | 0.296404112 |
| THBS1 | 453.3299548 | -2.452620618 | 0.682362627 | -3.594306781 | 0.000325256 | 0.013341539 |
| THBS2 | 33.01281249 | 1.727640896 | 0.672989228 | 2.567115231 | 0.010254853 | 0.11507205 |
| THBS3 | 1503.613005 | -0.736487158 | 0.375809424 | -1.959735737 | 0.050026686 | 0.257487003 |
| THBS4 | 245.1359193 | 0.310430346 | 0.37375782 | 0.830565489 | 0.406219138 | 0.701474651 |
| THEM4 | 212.0211065 | 0.342218423 | 0.305502809 | 1.12018094 | 0.262636664 | 0.57612824 |
| THEM6 | 348.5141095 | -0.782943447 | 0.539385531 | -1.451546997 | 0.146627606 | 0.43954428 |
| THEMIS2 | 152.0667009 | 0.555410097 | 0.438993456 | 1.265189923 | 0.205803254 | 0.512178669 |
| THG1L | 749.7833301 | -0.046294679 | 0.235614676 | -0.196484702 | 0.844230805 | 0.936860913 |
| THNSL1 | 315.911969 | 0.14066981 | 0.342360469 | 0.410882162 | 0.681158943 | 0.862490397 |
| THNSL2 | 264.8196673 | -1.040669426 | 0.41170012 | -2.527736511 | 0.011480047 | 0.121279761 |
| THOC1 | 1584.734525 | 0.220927684 | 0.30563544 | 0.722847076 | 0.469773841 | 0.74495024 |

| | | | | | | |
|---|---|---|---|---|---|---|
| THOC2 | 2864.498451 | 0.375641307 | 0.25789722 | 1.456554312 | 0.145239447 | 0.436754738 |
| THOC3 | 1135.350471 | 0.447557798 | 0.351382374 | 1.273705887 | 0.202767669 | 0.509290789 |
| THOC5 | 1655.325845 | -1.05537539 | 0.222012215 | -4.75368164 | 2.00E-06 | 0.000306095 |
| THOC6 | 1082.859286 | 0.369309737 | 0.308795853 | 1.195967283 | 0.231709327 | 0.540892783 |
| THOC7 | 1713.739275 | 0.603232188 | 0.447700364 | 1.347401603 | 0.177850924 | 0.482075465 |
| THOP1 | 1524.758 | 0.136594523 | 0.46118413 | 0.296182183 | 0.767090954 | 0.903470997 |
| THPO | 35.44035935 | 0.2849469 | 0.552622565 | 0.51562661 | 0.606115221 | 0.826611695 |
| THRA | 582.9922587 | 0.154215554 | 0.440880595 | 0.349789843 | 0.726496422 | 0.884630646 |
| THRAP3 | 5261.946394 | -0.257215512 | 0.315209102 | -0.816015498 | 0.414491272 | 0.706782858 |
| THRB | 121.1304559 | -1.98575443 | 0.61581971 | -3.224571085 | 0.001261615 | 0.031428515 |
| THSD4 | 304.3418998 | -0.096685515 | 0.463521996 | -0.20858884 | 0.834769223 | 0.932107564 |
| THSD7A | 25.96505896 | -2.269284352 | 1.064486101 | -2.131812101 | 0.033022296 | 0.207677316 |
| THSD8 | 1048.671497 | 1.294342273 | 0.561755131 | 2.304104053 | 0.021216803 | 0.165290607 |
| THTPA | 317.2577308 | 0.415931525 | 0.320418209 | 1.298089537 | 0.194256569 | 0.500187881 |
| THUMPD1 | 1941.166423 | 0.202493235 | 0.270638573 | 0.748205376 | 0.454336289 | 0.734285271 |
| THUMPD2 | 455.6124317 | 0.594267728 | 0.30683108 | 1.936791173 | 0.052770875 | 0.265446601 |
| THUMPD3 | 1335.867447 | 0.068223358 | 0.247642382 | 0.275491444 | 0.782938698 | 0.909909835 |
| THY1 | 723.1674633 | 0.581748951 | 0.344197058 | 1.690162471 | 0.090996876 | 0.345733435 |
| THYN1 | 1133.514593 | 0.578314479 | 0.328032155 | 1.762981067 | 0.077903683 | 0.320085498 |
| TIA1 | 5167.08989 | -0.092120716 | 0.324931615 | -0.28350801 | 0.776787442 | 0.907767854 |
| TIAF1 | 39.69151577 | -0.977000949 | 0.625943663 | -1.560844861 | 0.11856036 | 0.396971798 |
| TIAL1 | 3588.833307 | 0.188855366 | 0.238703334 | 0.791171882 | 0.428843697 | 0.715685016 |
| TIAM1 | 1629.797797 | 0.112335195 | 0.41885058 | 0.268198733 | 0.78854635 | 0.912309638 |
| TIAM2 | 888.4993921 | 0.549387754 | 0.501446716 | 1.095605448 | 0.273251476 | 0.586278328 |
| TICAM1 | 22.74664276 | -0.920806735 | 0.970300234 | -0.948991562 | 0.342624904 | 0.649420591 |
| TICAM2 | 12.09073406 | 6.752477956 | 1.751909298 | 3.854353625 | 0.000116036 | 0.006823174 |
| TICRR | 517.7014176 | 0.197364123 | 0.505070747 | 0.390765302 | 0.695970725 | 0.869877399 |
| TIE1 | 19.71509838 | 0.402496858 | 0.839910006 | 0.479214267 | 0.631786205 | 0.838768903 |
| TIFA | 191.156285 | 0.506812465 | 0.379623068 | 1.335041278 | 0.181862837 | 0.48591777 |
| TIGAR | 767.9911755 | 0.111849438 | 0.342058461 | 0.326989246 | 0.743676019 | 0.892535157 |
| TIGD1 | 251.8026888 | -0.217304582 | 0.379159489 | -0.573121834 | 0.566562204 | 0.804513376 |
| TIGD2 | 313.788977 | -0.277809381 | 0.363595055 | -0.764062595 | 0.444829941 | 0.727526247 |
| TIGD3 | 15.06035686 | 1.606926189 | 0.786113741 | 2.044139551 | 0.040939763 | 0.232707809 |
| TIGD4 | 46.62522873 | 0.353081248 | 0.553668384 | 0.637712497 | 0.523660849 | 0.777662198 |
| TIGD5 | 61.26295765 | 0.57253958 | 0.478400751 | 1.19677818 | 0.231393025 | 0.540403877 |
| TIGD6 | 142.0830146 | -0.33914954 | 0.355379448 | -0.954330763 | 0.339916241 | 0.646629378 |
| TIGD7 | 243.721024 | 0.319233763 | 0.286825409 | 1.112989831 | 0.265712797 | 0.579572403 |
| TIMELESS | 3895.593376 | -0.005516579 | 0.430849491 | -0.01280396 | 0.989784197 | 0.996285567 |
| TIMM10 | 1035.493856 | 1.089944435 | 0.432421381 | 2.52056092 | 0.011716795 | 0.122759326 |
| TIMM10B | 736.680546 | -0.017475764 | 0.252588107 | -0.069186804 | 0.944840927 | 0.979553348 |
| TIMM13 | 1377.414628 | 0.890535972 | 0.477592194 | 1.864636782 | 0.062232334 | 0.287492915 |
| TIMM17A | 1499.587924 | 0.549444778 | 0.302798277 | 1.814557144 | 0.069591993 | 0.303425952 |
| TIMM17B | 1415.116716 | 0.645189 | 0.343879711 | 1.876205487 | 0.060627069 | 0.284105089 |
| TIMM21 | 674.8737504 | 0.444255916 | 0.253445366 | 1.752866598 | 0.079624908 | |

0.32319892
TIMM22 602.6578972 0.224324955 0.294203721 0.762481704 0.445772559
0.728439722
TIMM23 1760.326391 0.634058205 0.342915655 1.849020879 0.064454798
0.293286926
TIMM23B 355.8014511 0.15416897 0.316609642 0.486937064 0.626302932
0.835767952
TIMM29 320.5992166 0.628891431 0.467102867 1.346366028 0.178184501
0.482467283
TIMM44 970.4460012 -0.036421697 0.327637093 -0.111164753
0.911485701 0.966866396
TIMM50 1843.224783 0.221421503 0.320416875 0.691041951 0.489539178
0.757836619
TIMM8A 435.7338396 0.837004445 0.360550169 2.321464575 0.020261784
0.161005801
TIMM8B 1082.966727 1.025282795 0.377857736 2.713409565 0.006659474
0.089249897
TIMM9 860.4353329 0.608215263 0.389132539 1.563002837 0.118051928 0.395958766
TIMMDC1 1646.886813 0.390065671 0.369834297 1.054703888 0.291560773
0.604602035
TIMP1 1077.823653 0.546205692 0.404920449 1.348920988 0.177362346 0.481794969
TIMP2 1738.283565 0.022788997 0.284205666 0.08018488 0.936090215 0.975841727
TIMP3 104.9385374 0.957754178 0.732003353 1.308401354 0.190737214 0.496058205
TINAGL1 111.7551395 -3.512547322 1.122875735 -3.128171008
0.001758978 0.039027635
TINCR 521.8455914 -1.508824256 0.656320152 -2.29891502 0.021509765
0.166469811
TINF2 440.6352508 0.209236684 0.31386382 0.666647988 0.504997009 0.766569683
TIPARP 224.3827812 0.130915986 0.438556525 0.29851565 0.765309633
0.903026935
TIPIN 570.6566851 0.435265849 0.42073857 1.034528041 0.300889375 0.611656533
TIPRL 1293.634517 0.63119809 0.234379054 2.693065265 0.007079841 0.092634974
TIRAP 94.50951476 -1.040469048 0.458415771 -2.269706047 0.023225424
0.174087407
TJAP1 974.8426667 -0.311833198 0.363552927 -0.857738104 0.391037094
0.690530747
TJP1 1193.308675 -0.309555516 0.317564122 -0.974781137 0.329668836
0.638548751
TJP2 3411.710111 -1.125572789 0.392357235 -2.868744826 0.004121041
0.066299381
TJP3 1024.113258 -1.122576256 0.49881987 -2.250464192 0.024419494
0.178492903
TK1 649.0676182 0.328039667 0.283985563 1.155127971 0.248038026 0.560718952
TK2 233.3588179 -0.721502343 0.315513349 -2.286756946 0.022210014
0.169472186
TKFC 1395.998684 0.199904215 0.299616646 0.667199962 0.504644415 0.766309601
TKT 14084.55807 0.289382288 0.257639565 1.123205933 0.261350043 0.574937735
TKTL1 14.16735611 1.010798421 0.807873562 1.251183933 0.210867379 0.518294063
TLCD1 273.8659371 0.392607265 0.411780728 0.953437686 0.340368355 0.647245965
TLCD2 91.74153624 -0.748667179 0.506079272 -1.479347644 0.139047425
0.427760708
TLDC2 34.50638304 -0.155641175 0.541359953 -0.287500349 0.773729235
0.906218345
TLE1 1577.928776 -0.820530277 0.503495317 -1.62966814 0.103171654
0.368199757
TLE3 1942.826585 -0.615095103 0.259761991 -2.367918033 0.017888499
0.150854036
TLE4 2347.15437 -0.525740763 0.31302388 -1.67955481 0.093043966
0.349658165
TLK1 1419.748092 -0.058369795 0.279821086 -0.208596842 0.834762976
0.932107564
TLK2 1525.284197 -0.225896989 0.269064413 -0.83956472 0.401152487
0.698568129
TLL2 24.85910784 1.01586787 0.719020525 1.412849613 0.157699947 0.457100859

| | | | | | | |
|---|---|---|---|---|---|---|
| TLN1 | 3581.488175 | -0.538996161 | 0.546233265 | -0.986750891 | 0.323764775 | 0.63357695 |
| TLN2 | 1624.52583 | -1.814867178 | 0.52177494 | -3.478256692 | 0.000504686 | 0.017658611 |
| TLNRD1 | 336.2533728 | 0.694811613 | 0.365191058 | 1.90259755 | 0.057093079 | 0.276551437 |
| TLR3 | 45.51454849 | 0.344402592 | 0.61981321 | 0.555655456 | 0.578446413 | 0.810209453 |
| TLR5 | 42.66183857 | -0.695758797 | 0.528762863 | -1.315823871 | 0.188233165 | 0.49338033 |
| TM2D1 | 496.7900514 | 0.737443544 | 0.390565761 | 1.888141813 | 0.059006915 | 0.281596069 |
| TM2D2 | 1310.246276 | 0.313698485 | 0.412898265 | 0.759747645 | 0.447405443 | 0.728985928 |
| TM2D3 | 664.4050682 | 0.63135128 | 0.371269126 | 1.7005219 | 0.089032801 | 0.342496199 |
| TM6SF2 | 122.5105691 | -0.277396445 | 0.361485968 | -0.767378183 | 0.442856696 | 0.726188529 |
| TM7SF2 | 1623.274363 | -0.092970182 | 0.377893209 | -0.246022367 | 0.805664918 | 0.918930547 |
| TM7SF3 | 2566.940609 | 0.293563445 | 0.257964478 | 1.13799949 | 0.255120695 | 0.567595846 |
| TM9SF1 | 398.2733905 | 0.099802415 | 0.412786124 | 0.241777544 | 0.80895254 | 0.919641737 |
| TM9SF2 | 4453.997956 | -0.125142025 | 0.254144192 | -0.492405606 | 0.622432637 | 0.833665183 |
| TM9SF3 | 5385.591142 | -0.282896884 | 0.271646952 | -1.041413798 | 0.297683541 | 0.60928029 |
| TM9SF4 | 1945.982393 | -0.573516132 | 0.322272916 | -1.779597672 | 0.075141828 | 0.314600339 |
| TMA16 | 989.4752723 | 0.874216792 | 0.387946649 | 2.253445916 | 0.024231047 | 0.178105241 |
| TMA7 | 5129.242727 | 1.22613555 | 0.464899186 | 2.637422448 | 0.008353872 | 0.103542973 |
| TMBIM1 | 1751.008018 | -0.753707532 | 0.369907324 | -2.037557741 | 0.041594182 | 0.234233606 |
| TMBIM4 | 1084.942023 | 0.640231285 | 0.412257562 | 1.552988577 | 0.120425864 | 0.400653679 |
| TMBIM6 | 12356.61055 | -0.461463072 | 0.311609469 | -1.480901956 | 0.138632702 | 0.427045019 |
| TMC3 | 22.86161865 | 0.12850502 | 0.819841799 | 0.156743679 | 0.875446862 | 0.94989303 |
| TMC4 | 21.80028 | -3.549340311 | 1.042057206 | -3.406089693 | 0.000659005 | 0.020959683 |
| TMC5 | 715.0227958 | -3.302061914 | 0.715605687 | -4.61435952 | 3.94E-06 | 0.000552393 |
| TMC6 | 980.2164017 | -0.513267685 | 0.480572482 | -1.068033864 | 0.285505237 | 0.599411008 |
| TMC7 | 269.5301715 | -0.960403875 | 0.350053127 | -2.74359462 | 0.006077053 | 0.084270328 |
| TMCC1 | 991.1221471 | -0.311247492 | 0.2430023 | -1.280841757 | 0.200249254 | 0.506121566 |
| TMCC2 | 205.3458966 | 0.363065719 | 0.376257096 | 0.964940522 | 0.334574604 | 0.64263567 |
| TMCC3 | 629.7262993 | -1.961239292 | 0.519229554 | -3.777210438 | 0.000158595 | 0.008239865 |
| TMCO1 | 1897.14527 | 0.330674332 | 0.310795436 | 1.063961353 | 0.287346207 | 0.601618729 |
| TMCO3 | 1256.87163 | 0.042600311 | 0.236959924 | 0.179778548 | 0.857326426 | 0.942343063 |
| TMCO4 | 238.3494923 | -1.533820943 | 0.584256786 | -2.625251396 | 0.008658501 | 0.104918628 |
| TMCO6 | 527.4908197 | -0.051175889 | 0.284452793 | -0.179909956 | 0.857223259 | 0.942343063 |
| TMED1 | 695.2134791 | 0.950764949 | 0.380967826 | 2.495656808 | 0.012572417 | 0.126734979 |
| TMED10 | 4672.88325 | 0.105790963 | 0.285665892 | 0.370331097 | 0.711135806 | 0.876982711 |
| TMED2 | 4424.696301 | -0.0909248 | 0.232540399 | -0.391006467 | 0.695792456 | 0.869726238 |
| TMED3 | 1621.004274 | 0.434666923 | 0.38327962 | 1.134072622 | 0.256764087 | 0.569149579 |
| TMED4 | 1992.409401 | 0.291761894 | 0.237572201 | 1.228097783 | 0.219410263 | 0.527925634 |
| TMED5 | 895.4421944 | 0.167576595 | 0.268141954 | 0.624954778 | 0.532000739 | 0.782369637 |
| TMED7 | 1854.86262 | 0.065811996 | 0.227868352 | 0.288815867 | 0.772722289 | 0.905717252 |
| TMED7-TICAM2 | 137.5681208 | 0.118670475 | 0.43413835 | 0.273347138 | 0.784586384 | |

0.910931322
TMED8 933.0292821 -0.662292219 0.290248271 -2.281812793 0.022500395
0.170591452
TMED9 3276.384905 0.324252487 0.393559662 0.823896649 0.409998285 0.70410731
TMEFF1 546.4447466 -0.093669253 0.59527487 -0.157354624
0.874965373 0.949740014
TMEFF2 34.7894217 0.845964252 0.95860884 0.882491603 0.377511019
0.680839237
TMEM100 35.04141243 0.296912448 0.535258949 0.554708051 0.579094369
0.810713649
TMEM101 1217.77882 0.491430561 0.408016685 1.204437416 0.228420558 0.5372664
TMEM102 62.21507621 -0.586320301 0.551740524 -1.062673985
0.287929822 0.601673803
TMEM104 497.7581538 -0.11024567 0.370825501 -0.297297974 0.766239028
0.903373663
TMEM106A 69.06373391 -0.583724757 0.468073318 -1.247079751
0.212368257 0.519629359
TMEM106B 1602.756095 0.169741689 0.237594727 0.714416903 0.474969423
0.748737095
TMEM106C 898.6441861 -0.491810221 0.500319747 -0.982991823
0.325611463 0.635672141
TMEM107 155.3164953 0.519549719 0.323837604 1.60435265 0.108636342
0.378356736
TMEM108 249.3836617 0.072605775 0.427215119 0.169951324 0.865048417
0.945248223
TMEM109 1181.673625 0.419373823 0.314113116 1.33510446 0.18184216
0.48591777
TMEM11 551.9488273 0.535108863 0.346401937 1.544762909 0.122403622
0.403241325
TMEM110-MUSTN1 166.9876381 0.108457335 0.331545898 0.327126154 0.743572471
0.892534058
TMEM114 10.1641242 1.149470848 1.290145939 0.890961877 0.372949623
0.67811865
TMEM115 694.8558598 -0.224643623 0.338982187 -0.662700377
0.507522468 0.768444894
TMEM116 322.1646288 0.494764709 0.290912976 1.700730973 0.088993516
0.342496199
TMEM117 10.41136883 -1.781954684 1.124701127 -1.584380634
0.113107126 0.386997793
TMEM119 42.25608681 1.003527383 0.675914864 1.484694946 0.137624651
0.425842052
TMEM120A 693.2214834 0.412515683 0.322363825 1.27965873 0.200665185
0.506835677
TMEM120B 568.0348998 -2.108323308 0.350173213 -6.020801221 1.74E-09
1.76E-06
TMEM121B 160.3302948 -0.674792216 0.384028243 -1.757142158
0.078893582 0.322507823
TMEM123 6596.10159 0.285324935 0.270160317 1.056131922 0.290907948
0.604006016
TMEM125 334.3018835 -1.308241801 0.648252453 -2.018105437
0.043580282 0.240245804
TMEM126A 569.5223529 0.99016185 0.355393788 2.786097795 0.005334677
0.077809098
TMEM126B 1314.728647 0.694228943 0.308217275 2.252401145 0.024296933
0.178492903
TMEM127 949.6875836 -0.594261118 0.284491456 -2.08885401 0.036720867
0.219399453
TMEM128 996.5104487 0.510658517 0.342055849 1.492909765 0.135460809
0.422942154
TMEM129 704.7894523 0.591209442 0.363455179 1.626636447 0.103814334
0.369190637
TMEM130 12.41943369 2.011126953 1.128783027 1.781677173 0.074801891
0.313842348
TMEM131 1821.205479 -0.772631275 0.249713767 -3.094067597

0.001974325 0.041716552
TMEM131L 430.4949155 0.052154009 0.543383829 0.095980053 0.923536415
0.971297812
TMEM132A 2721.41812 0.357475414 0.412666901 0.866256569 0.386349478
0.686679504
TMEM132B 1146.664373 0.800516314 0.646712819 1.237823484 0.215781519
0.523378873
TMEM132E 54.0621766 -0.405869381 0.796337637 -0.509669971
0.610282694 0.828658261
TMEM134 391.3820166 0.685343963 0.396030648 1.730532641 0.083535155
0.330745685
TMEM135 1753.166914 -0.612797738 0.28297324 -2.165567802
0.030344229 0.198737258
TMEM136 413.4052469 0.289779993 0.369904235 0.783391932 0.433396992
0.71890019
TMEM138 495.7026148 0.323529332 0.271177401 1.193054181 0.232848155
0.542441738
TMEM139 150.3518659 -1.241245644 0.53657025 -2.313295684
0.020706388 0.16310312
TMEM141 909.3800394 0.573029741 0.431732622 1.327279226 0.184416333
0.488866012
TMEM143 162.4371087 -0.076123107 0.317449626 -0.239795862
0.810488515 0.920629025
TMEM144 373.7541413 -0.819476512 0.560857341 -1.461113999
0.143984159 0.434995877
TMEM145 15.22593096 2.377256476 0.827430943 2.873057258 0.004065204
0.065749696
TMEM147 1874.72411 0.748108003 0.487787887 1.533674828 0.125109669
0.406867909
TMEM14A 398.1359612 0.741651862 0.337108977 2.200035931 0.027804346
0.190782329
TMEM14B 1748.570388 0.897793726 0.386913431 2.320399485 0.020319277
0.161005801
TMEM14C 1499.015072 0.739035518 0.354910109 2.082317464 0.037313482
0.22118892
TMEM150A 459.8377673 -0.1308377 0.269987342 -0.484606793 0.627955294
0.836556882
TMEM150C 104.2684292 -0.116742999 0.359472422 -0.324762045
0.745361173 0.893479326
TMEM154 452.1096468 -3.243754048 0.72679392 -4.46310014 8.08E-06
0.000921465
TMEM158 60.60367939 1.709279133 0.433694684 3.941203788 8.11E-05
0.005278838
TMEM159 343.6019742 0.130122304 0.366346115 0.355189528 0.722447609
0.882935039
TMEM160 140.3081591 0.706232053 0.492919552 1.432753176 0.151928387
0.447727291
TMEM161A 611.7058587 0.452336219 0.431013822 1.049470331 0.293961703
0.606430922
TMEM161B 692.1203877 0.135754982 0.255954477 0.530387215 0.595843488
0.821776506
TMEM163 16.10689508 0.436060425 0.822588122 0.530107856 0.596037153
0.821812117
TMEM164 969.0073882 -0.389665605 0.258392738 -1.508036211
0.13154526 0.41719445
TMEM165 1387.474655 0.027017967 0.335854122 0.080445544 0.935882906
0.975692658
TMEM167A 4941.349663 0.103213686 0.248708769 0.414998177 0.678143238
0.861001846
TMEM167B 802.0502619 -0.08146237 0.256892837 -0.317106427 0.751162848
0.896349431
TMEM168 958.7211576 -0.232230139 0.285010138 -0.814813609
0.41517901 0.707398889
TMEM169 286.3854315 0.80113088 0.476670344 1.680681188 0.092824856

0.349440669
TMEM17 63.95292015 0.729922846 0.44100692 1.655127876 0.097898561
0.359375073
TMEM170A 1060.55708 0.181386227 0.32181227 0.563639873 0.572999245
0.807760521
TMEM170B 490.5429886 0.775450663 0.397215536 1.952216351 0.050912524
0.259366654
TMEM171 19.45863893 -2.645619013 1.076827073 -2.456865246
0.014015525 0.134490529
TMEM175 117.6648272 -0.619168667 0.550857778 -1.124008214
0.261009541 0.574380005
TMEM176B 10.39544629 1.40158156 0.996134269 1.407020723 0.159421244
0.459194929
TMEM177 440.9692611 0.636120923 0.308103169 2.064636092 0.038957446
0.227316695
TMEM178A 47.74671471 0.791886737 0.576900526 1.372657333 0.169858899
0.472495848
TMEM178B 204.0316128 0.246305554 0.581674688 0.423442105 0.671972731
0.858752635
TMEM179 63.16187474 0.568420029 0.462000963 1.230343819 0.218568382
0.527064894
TMEM179B 629.7195144 0.310785097 0.510513572 0.608769509 0.542677226
0.789277749
TMEM18 829.1454593 0.522775488 0.296270454 1.764521171 0.077644274
0.31942339
TMEM181 584.4092267 0.146983571 0.321662798 0.456949239 0.647707531
0.846597949
TMEM182 125.9312213 -0.295037523 0.364036726 -0.810460873
0.417675344 0.709183284
TMEM183A 3202.283309 0.17233719 0.221947619 0.776476859 0.437467479
0.721468765
TMEM184A 25.06374858 -0.67832898 0.719735109 -0.942470323 0.345951898
0.652683033
TMEM184B 1260.688559 -0.342738415 0.259270817 -1.321932095
0.186190764 0.490817669
TMEM184C 995.7570398 -0.656673719 0.302107538 -2.173642288
0.02973201 0.197143498
TMEM185A 582.5036815 -0.294884737 0.252076227 -1.169823669
0.242071937 0.553231259
TMEM185B 525.1769471 0.063561667 0.298860151 0.212680303 0.831576331
0.930997148
TMEM186 311.0265301 0.471482478 0.35379256 1.332652327 0.182645925
0.486683209
TMEM187 110.2226657 0.801083011 0.542122254 1.477679629 0.139493549
0.428218664
TMEM189 1399.53182 0.026492522 0.430730031 0.061506095 0.950956161
0.981826318
TMEM189-UBE2V1 183.8562417 -1.404765087 2.107527573 -0.666546481
0.505061864 0.766569683
TMEM19 339.5931001 -0.251895311 0.318951502 -0.789760541
0.429667627 0.716389544
TMEM190 13.36276851 0.272188086 1.443711963 0.188533512 0.850458448
0.939032396
TMEM191A 50.73576532 0.019427435 0.779313404 0.024928912 0.980111666
0.993196129
TMEM191B 20.66073678 -0.009901775 1.747728384 -0.005665511 0.9954796
0.998116797
TMEM191C 54.12280054 -0.624509918 0.594434119 -1.050595682
0.293444325 0.606314063
TMEM192 479.6970372 0.324944316 0.325588216 0.998022347 0.318268522
0.628377376
TMEM198 55.52318038 -0.15460533 0.67877922 -0.227769686 0.819825295
0.924520147
TMEM198B 589.5581853 -0.84269997 0.280356815 -3.005812326 0.002648724

0.050167035
TMEM199 717.9106665 -0.035173106 0.290634811 -0.121021656
0.903673883 0.963499647
TMEM200A 384.3041589 -1.296159533 0.364207808 -3.558846093
0.000372488 0.014489772
TMEM200B 40.65256671 -0.584525381 0.904472699 -0.646260945
0.518110376 0.773958483
TMEM200C 259.5616076 0.415375294 0.377002955 1.101782594 0.270556199
0.583257419
TMEM201 249.251679 0.448571397 0.335527209 1.336915115 0.181250347
0.485749663
TMEM203 895.7440826 0.750436631 0.355173292 2.112874613 0.034611501
0.21269873
TMEM205 956.214879 0.505297822 0.434343308 1.163360441 0.244683293
0.556035085
TMEM206 633.06624 0.527330281 0.262385427 2.009754455 0.044457183
0.242262905
TMEM208 683.395173 0.838675861 0.465485774 1.801721788 0.071589188
0.307758452
TMEM209 860.9505888 0.159247685 0.350679873 0.454111278 0.649748736
0.84729896
TMEM212 40.6502341 -0.541821877 0.551499513 -0.982452141
0.32587715 0.635827259
TMEM214 1121.313515 -0.585307758 0.36153248 -1.618963136
0.105455195 0.372436233
TMEM216 427.658119 0.315753211 0.288639618 1.093935799 0.273983137
0.586847211
TMEM217 11.48582382 -0.694805767 0.909748271 -0.763734089
0.445025722 0.727707384
TMEM218 380.8009213 0.754201834 0.338831777 2.225888732 0.026021635
0.184130533
TMEM219 1643.752307 0.759781065 0.466986811 1.626986133 0.103740043
0.369190637
TMEM220 64.94207925 0.324648188 0.574942713 0.5646618 0.572303821
0.807456036
TMEM221 13.6951961 0.189241002 0.774811344 0.244241393 0.807043871
0.919137032
TMEM222 738.250656 -0.221397783 0.266117642 -0.831954553
0.405434598 0.701111169
TMEM223 323.0888266 0.645110091 0.368411445 1.751058769 0.079935787
0.323627252
TMEM225B 28.3474166 1.834150642 0.715275965 2.564255939 0.010339727
0.115426046
TMEM229A 10.41880086 0.781250225 0.997938156 0.782864369 0.433706763
0.71918874
TMEM229B 11.92791884 0.266225131 0.913993554 0.291276815 0.770839618
0.905067213
TMEM230 2363.572709 0.266960585 0.407987961 0.654334467 0.512896345
0.771819984
TMEM231 384.2708457 0.150838004 0.33267085 0.453415151 0.650249828
0.847430646
TMEM231P1 22.10534315 -1.152712757 0.842943166 -1.367485737
0.171473105 0.474879827
TMEM232 38.91713257 -0.510008658 0.567256844 -0.899078898
0.368610637 0.67460916
TMEM234 253.4066626 -0.318768937 0.379111414 -0.840831811
0.400442164 0.698047577
TMEM237 1395.010274 0.264357652 0.224681675 1.176587505 0.239360188
0.550578888
TMEM240 23.57743771 0.694936865 0.849221397 0.818322368 0.413173137
0.705607651
TMEM241 237.5756225 0.163237555 0.331198293 0.492869554 0.622104761
0.833640266
TMEM242 430.1560135 0.469558797 0.35527141 1.321690359 0.18627128

| | | | | | | |
|---|---|---|---|---|---|---|
| | | | | | | 0.490817669 |
| TMEM243 | 221.5103482 | 0.478360586 | 0.386488326 | 1.237710312 | 0.215823494 | 0.523378873 |
| TMEM245 | 2579.471624 | -0.259781633 | 0.314281591 | -0.826588768 | 0.408470186 | 0.702771243 |
| TMEM246 | 665.828721 | 0.849324396 | 0.476475015 | 1.78251612 | 0.074665105 | 0.313779586 |
| TMEM248 | 1854.752553 | -0.134520615 | 0.249853233 | -0.538398538 | 0.590301936 | 0.818321084 |
| TMEM25 | 400.7844893 | -0.432883405 | 0.294445984 | -1.47016237 | 0.141517784 | 0.431464288 |
| TMEM250 | 960.4690372 | 0.654779494 | 0.357561635 | 1.831235319 | 0.067065427 | 0.297817842 |
| TMEM251 | 196.9316856 | 0.490771873 | 0.412800906 | 1.188882743 | 0.234485812 | 0.544775536 |
| TMEM253 | 27.75833755 | -0.081634715 | 0.638549153 | -0.127844058 | 0.898272383 | 0.960048173 |
| TMEM254 | 893.2853214 | -0.53683777 | 0.33370041 | -1.608741716 | 0.10767283 | 0.376206676 |
| TMEM255A | 119.947941 | 1.789813232 | 0.768161926 | 2.32999472 | 0.01980643 | 0.159323092 |
| TMEM255B | 44.56367141 | 1.231242994 | 0.476887527 | 2.581830985 | 0.00982777 | 0.112441253 |
| TMEM256 | 934.4348177 | 1.468615038 | 0.509507567 | 2.882420462 | 0.003946328 | 0.064934636 |
| TMEM258 | 2831.33798 | 1.084687957 | 0.431724009 | 2.512456883 | 0.011989376 | 0.124327283 |
| TMEM259 | 1589.950846 | -0.149594052 | 0.390708743 | -0.38287869 | 0.701809712 | 0.872479299 |
| TMEM260 | 746.1728007 | -0.531138332 | 0.269877342 | -1.968073082 | 0.04905963 | 0.255021315 |
| TMEM263 | 786.6978856 | 0.252124873 | 0.23486812 | 1.073474224 | 0.283058398 | 0.596961028 |
| TMEM265 | 301.3366767 | 0.71853572 | 0.91405501 | 0.786096802 | 0.431810778 | 0.718311547 |
| TMEM266 | 17.12595204 | 0.092853403 | 0.724910694 | 0.128089438 | 0.898078195 | 0.959968192 |
| TMEM267 | 206.0621425 | 0.438565686 | 0.316192662 | 1.387020444 | 0.165435529 | 0.467560858 |
| TMEM268 | 55.19196168 | -0.89444889 | 0.619847266 | -1.443014979 | 0.149016252 | 0.443544075 |
| TMEM269 | 24.24167565 | 1.036217837 | 0.645197981 | 1.606046311 | 0.108263735 | 0.377405589 |
| TMEM30A | 3286.442465 | -0.130200226 | 0.303493184 | -0.429005436 | 0.667919268 | 0.856696248 |
| TMEM30B | 299.6921136 | -0.926066706 | 0.78247131 | -1.183515221 | 0.236605018 | 0.547336452 |
| TMEM31 | 17.66995364 | 0.879299318 | 0.770193761 | 1.141659883 | 0.253595422 | 0.565942797 |
| TMEM33 | 1435.405396 | 0.25430298 | 0.457014864 | 0.556443564 | 0.577907665 | 0.81007458 |
| TMEM35A | 205.8975728 | 0.679211762 | 0.363527163 | 1.868393427 | 0.061707253 | 0.28627362 |
| TMEM35B | 72.14257518 | 0.902186598 | 0.634233961 | 1.422482322 | 0.15488628 | 0.452576994 |
| TMEM37 | 152.5676457 | -0.827745054 | 0.449162282 | -1.842864119 | 0.065348852 | 0.294886209 |
| TMEM38A | 51.89538279 | 0.12723991 | 0.460528884 | 0.276290835 | 0.782324695 | 0.909909835 |
| TMEM38B | 538.3533082 | 0.869947585 | 0.266086887 | 3.269411709 | 0.001077714 | 0.028433348 |
| TMEM39A | 1094.152077 | -0.044617223 | 0.284038724 | -0.15708148 | 0.875180633 | 0.949740014 |
| TMEM39B | 485.3110242 | 0.00593851 | 0.25408426 | 0.023372206 | 0.981353375 | |

0.993269201
TMEM40 44.51246644 -1.108760803 0.872578837 -1.270671206
0.203845643 0.509986087
TMEM41A 472.965287 0.186913482 0.319806797 0.584457503 0.558912549
0.799176463
TMEM41B 1006.88523 0.463074059 0.436898836 1.059911404 0.289184907
0.602227073
TMEM42 208.4589404 1.068171473 0.385332237 2.772079182 0.005569949
0.079919047
TMEM43 790.1385089 -0.03564846 0.26071112 -0.136735481 0.891239883
0.957017706
TMEM44 376.7813721 -0.57035124 0.305980277 -1.864013081 0.062319867
0.287743217
TMEM45A 458.7167332 0.523867277 0.280390427 1.868349366 0.06171339
0.28627362
TMEM47 904.6360148 0.497919745 0.411329526 1.210513018 0.226082099
0.534727976
TMEM50A 2684.048777 0.441178023 0.353029858 1.249690397 0.211412667
0.518651612
TMEM50B 342.0852672 -0.292185304 0.305612398 -0.956064955
0.339039419 0.646095594
TMEM51 262.831965 0.01124711 0.494569645 0.022741205 0.981856707
0.993518428
TMEM52 63.57123071 1.273049532 0.48472665 2.626324615 0.008631246
0.104839578
TMEM53 305.7059827 0.30182786 0.415243987 0.726868706 0.467306389
0.743758417
TMEM54 2666.048074 0.077560068 1.296038082 0.059843973 0.952279903
0.982774543
TMEM56 102.3878377 -0.769436105 0.543374146 -1.416033706
0.156765636 0.455611392
TMEM56-RWDD3 30.81383481 -1.055625376 1.047270814 -1.007977461
0.313465283 0.623029189
TMEM59 2295.209142 0.420471333 0.346988383 1.211773519 0.225599088
0.53459368
TMEM59L 14.04102755 2.104505995 0.807069679 2.607588972 0.009118237
0.107988114
TMEM60 635.2584979 1.023264036 0.427086898 2.395915305 0.016578921
0.145808003
TMEM62 248.368057 -0.326876236 0.43478825 -0.751805588
0.452167981 0.732888936
TMEM63A 503.3790074 -1.436596016 0.46299886 -3.102806812
0.001916947 0.040960574
TMEM63B 1641.28801 -0.248768057 0.239767081 -1.037540499
0.299484032 0.610246278
TMEM63C 78.23027699 0.544017384 0.57987733 0.938159428 0.348162485
0.654681837
TMEM64 202.9949818 -0.541320535 0.43430719 -1.246400125
0.212617535 0.519735803
TMEM65 572.8941425 0.37792259 0.272962601 1.384521499 0.16619883
0.468047055
TMEM67 589.0397124 0.180765109 0.312863439 0.577776391 0.563415096
0.801924557
TMEM68 1012.968758 0.645172202 0.255076614 2.529327134 0.011428145
0.121242234
TMEM69 1506.602328 0.135866856 0.330663742 0.410891303 0.68115224
0.862490397
TMEM70 539.6318131 0.819561322 0.406384101 2.016716008 0.043725158
0.240923506
TMEM74 125.2325815 1.070092456 0.366597631 2.918983553 0.003511748
0.060818175
TMEM74B 10.31392727 0.120435873 0.905802152 0.132960462 0.89422465
0.958206114
TMEM79 332.0576186 -0.958650345 0.591287767 -1.621292369

| | | | | | | |
|---|---|---|---|---|---|---|
| | | | | | 0.104954951 | 0.37133017 |
| TMEM80 | 381.6187333 | 0.715331851 | 0.340387829 | 2.101520059 | 0.035595339 | 0.215965226 |
| TMEM81 | 94.18150696 | 0.929274576 | 0.554489276 | 1.675910818 | 0.093755663 | 0.351226741 |
| TMEM82 | 21.91747608 | -0.888600788 | 0.964317069 | -0.921481966 | 0.356798852 | 0.664252717 |
| TMEM86A | 18.03917169 | -1.355177969 | 0.908567308 | -1.491554844 | 0.135815884 | 0.423669306 |
| TMEM86B | 69.31540275 | 0.180582285 | 0.498952087 | 0.361923099 | 0.717409498 | 0.880516207 |
| TMEM87A | 1281.054982 | -0.112053363 | 0.250172995 | -0.447903512 | 0.654222834 | 0.849068645 |
| TMEM87B | 644.4779993 | -0.527864977 | 0.329999666 | -1.599592455 | 0.109689023 | 0.380712005 |
| TMEM88 | 312.5321002 | 1.2472796 | 0.380698122 | 3.276295647 | 0.001051784 | 0.027896179 |
| TMEM8A | 565.1478741 | 0.466085002 | 0.42787461 | 1.089302779 | 0.276020387 | 0.589374426 |
| TMEM8B | 412.2103529 | 0.268797431 | 0.411834416 | 0.652683264 | 0.513960494 | 0.772630754 |
| TMEM9 | 2189.585945 | 0.564871721 | 0.328665606 | 1.718682183 | 0.085672258 | 0.33574506 |
| TMEM91 | 100.3646144 | 0.688695899 | 0.459142077 | 1.499962502 | 0.133624116 | 0.420294949 |
| TMEM92 | 344.1198902 | 0.054973356 | 0.505504333 | 0.108749525 | 0.91340116 | 0.967884962 |
| TMEM94 | 1772.179851 | -0.630292622 | 0.404412699 | -1.558538155 | 0.119105731 | 0.398078527 |
| TMEM97 | 2555.752673 | 0.151413636 | 0.227382116 | 0.665899497 | 0.505475342 | 0.767010044 |
| TMEM98 | 2152.089723 | 0.363311057 | 0.245653965 | 1.478954581 | 0.139152454 | 0.427760708 |
| TMEM99 | 260.8889305 | 0.09664896 | 0.347114149 | 0.278435668 | 0.780677942 | 0.909271055 |
| TMEM9B | 483.739025 | -0.02878165 | 0.341852014 | -0.084193301 | 0.932902744 | 0.974594927 |
| TMF1 | 1075.413864 | -0.282094744 | 0.267105729 | -1.056116413 | 0.290915033 | 0.604006016 |
| TMLHE | 354.5705181 | -0.176607429 | 0.313851588 | -0.562710005 | 0.57363237 | 0.808163661 |
| TMOD1 | 174.1509708 | -1.513692078 | 0.630849113 | -2.399451859 | 0.016419639 | 0.144574776 |
| TMOD2 | 94.57023745 | -0.056801968 | 0.549098924 | -0.103445782 | 0.917609178 | 0.969677217 |
| TMOD3 | 1088.711685 | -0.357794773 | 0.33282314 | -1.075029739 | 0.282361414 | 0.596201116 |
| TMPO | 4901.502359 | 0.524920262 | 0.244153779 | 2.149957554 | 0.031558572 | 0.202592072 |
| TMPOP2 | 23.63863114 | 0.439973711 | 0.631001764 | 0.697262251 | 0.485638687 | 0.755840687 |
| TMPPE | 52.70658786 | -2.369332286 | 0.92918592 | -2.549901193 | 0.010775345 | 0.117984572 |
| TMPRSS11E | 33.2572981 | -0.277898731 | 0.772936067 | -0.359536504 | 0.719193775 | 0.881260702 |
| TMPRSS11F | 15.35810446 | -3.402511603 | 1.144342117 | -2.973334244 | 0.002945835 | 0.05358664 |
| TMPRSS13 | 1035.582131 | -1.708131062 | 0.601466958 | -2.839941647 | 0.004512179 | 0.070434393 |
| TMPRSS2 | 496.9707338 | -0.89414443 | 0.612021188 | -1.46096973 | 0.144023749 | 0.434995877 |
| TMPRSS4 | 614.0332932 | -1.541909279 | 0.734044902 | -2.10056534 | 0.035679139 | 0.216083122 |
| TMPRSS5 | 30.05631648 | 0.539011792 | 0.605800369 | 0.889751507 | 0.37359933 | 0.678462465 |
| TMPRSS9 | 56.98243235 | -1.400842169 | 0.680754491 | -2.057778812 | | |

0.039611371 0.229275285
TMSB10 20797.24484 0.889313408 0.540530938 1.645259033 0.099916404
0.362399544
TMSB15A 2341.937691 1.81347432 0.395219843 4.588520419 4.46E-06
0.00059932
TMSB15B 386.0077314 0.677576791 0.293020951 2.31238343 0.020756563
0.163328741
TMSB4X 17113.84441 0.545717621 0.466167803 1.170646317 0.241740972
0.552977821
TMSB4Y 60.57910524 0.237745837 0.554130085 0.429043365 0.667891666
0.856696248
TMTC1 430.7527764 -0.622655544 0.376098838 -1.655563592 0.097810228
0.359375073
TMTC2 756.3999843 -0.803985566 0.463179494 -1.735796975 0.082599739
0.328745819
TMTC3 1117.334508 -0.062863011 0.29217571 -0.215154817 0.829646617
0.929858808
TMTC4 1278.090201 0.325509493 0.259904374 1.252420217 0.210416783 0.51816316
TMUB1 267.5248329 0.020094407 0.546698202 0.036755941 0.970679605 0.98957539
TMUB2 1067.132988 -0.622607581 0.259198774 -2.402046777 0.016303623
0.144307043
TMX1 1685.490698 0.784286503 0.265532732 2.953633987 0.003140562 0.055790951
TMX2 2137.01882 0.074693958 0.299236388 0.249615226 0.802884923 0.917969611
TMX3 1533.283027 -0.067334814 0.261381453 -0.257611293 0.796706913
0.915710299
TMX4 600.559053 -0.115742477 0.263323331 -0.439545089 0.660266619
0.852284938
TNC 2096.769613 -1.699941092 0.391532087 -4.341766992 1.41E-05
0.00142006
TNFAIP1 785.0309216 -0.592956153 0.268302515 -2.210028309
0.027103198 0.188184262
TNFAIP2 132.6289928 0.30418987 0.377150233 0.806548277 0.419926784
0.710456202
TNFAIP3 263.5594043 -2.183898476 0.705361832 -3.09613928 0.001960582
0.041483956
TNFAIP8 449.3564138 0.139702398 0.28038237 0.498256714 0.618303113
0.832248827
TNFAIP8L1 398.6073569 0.815706136 0.327590735 2.490015888 0.012773738
0.127914447
TNFRSF10A 83.08034732 0.389122179 0.601814049 0.646582079 0.51790246
0.773950969
TNFRSF10B 3365.037271 0.492664358 0.442950552 1.112233308 0.26603785
0.579643865
TNFRSF10C 251.7891954 0.185065654 0.440322808 0.420295408 0.674269664
0.859672768
TNFRSF10D 631.5975931 0.192980611 0.316179201 0.610352012 0.54162865
0.788584285
TNFRSF11A 33.6519052 0.419388181 0.626681633 0.669220477 0.503354846
0.765246655
TNFRSF11B 12.27353537 -0.394204684 1.140853361 -0.345534928
0.729692257 0.886372257
TNFRSF12A 344.1604171 0.730840712 0.48110401 1.519090875 0.128739623
0.413329156
TNFRSF13C 42.1309692 0.636072018 0.53449083 1.190052256 0.234025854
0.544206964
TNFRSF19 2179.389726 -0.064729691 0.512036909 -0.126416065
0.899402586 0.960836323
TNFRSF1A 677.2713343 -0.09582097 0.248466214 -0.385649897 0.699755974
0.871449506
TNFRSF1B 31.43056778 0.564060211 0.698637651 0.807371617 0.419452414
0.710054962
TNFRSF21 897.4381083 -0.066443245 0.3941766 -0.168562125
0.866141074 0.945612136
TNFRSF25 178.7671851 -0.039407701 0.436580866 -0.090264379

| | | | | | | |
|---|---|---|---|---|---|---|
| | | | | | 0.928077126 | 0.972633911 |
| TNFRSF8 | 10.31218036 | -0.014599051 | 0.954766233 | -0.015290707 | 0.987800256 | 0.995541774 |
| TNFRSF9 | 10.38829189 | -0.393231761 | 0.972681232 | -0.404276086 | 0.686009709 | 0.865124962 |
| TNFSF12 | 97.01406603 | -0.876805411 | 0.679474649 | -1.290416667 | 0.196906028 | 0.50258876 |
| TNFSF12-TNFSF13 | 20.32612787 | 8.495076819 | 4.331579942 | 1.961195899 | 0.049856175 | 0.257355574 |
| TNFSF13 | 100.1554346 | -2.046427902 | 0.514460393 | -3.977814291 | 6.96E-05 | 0.004731696 |
| TNFSF13B | 15.17485233 | 0.897318095 | 0.80074341 | 1.120606281 | 0.26245549 | 0.576074285 |
| TNFSF15 | 84.00703161 | -4.127820938 | 0.957102784 | -4.312829307 | 1.61E-05 | 0.001557478 |
| TNFSF4 | 33.79752207 | 0.297314306 | 0.634827752 | 0.468338546 | 0.639542507 | 0.843436176 |
| TNFSF8 | 21.61407134 | -7.053006509 | 1.766074779 | -3.993605815 | 6.51E-05 | 0.004569989 |
| TNFSF9 | 268.0922579 | 1.36909269 | 0.404701155 | 3.382972036 | 0.000717059 | 0.022065937 |
| TNIK | 579.1666889 | -1.557508632 | 0.402094979 | -3.87348441 | 0.00010729 | 0.0064336 |
| TNIP1 | 1175.383787 | -0.662404574 | 0.303330953 | -2.18376848 | 0.028979265 | 0.194705238 |
| TNIP2 | 716.8838544 | -0.078317848 | 0.331990013 | -0.23590423 | 0.813506986 | 0.921959809 |
| TNK1 | 93.80246313 | -1.386430754 | 0.51814235 | -2.675771926 | 0.007455736 | 0.096019504 |
| TNK2 | 465.4675705 | -0.698224244 | 0.416355118 | -1.676992102 | 0.093544026 | 0.351015687 |
| TNKS | 2298.033699 | -0.309352373 | 0.275175297 | -1.124201106 | 0.26092772 | 0.574283251 |
| TNKS1BP1 | 2800.171465 | -0.00273592 | 0.376899027 | -0.007259027 | 0.994208185 | 0.997759633 |
| TNKS2 | 3005.589404 | -0.252618917 | 0.347846035 | -0.726237738 | 0.467693041 | 0.744139604 |
| TNN | 16.57309278 | -0.008017332 | 1.010873919 | -0.00793109 | 0.993671972 | 0.99730871 |
| TNNC1 | 384.4937528 | 0.845222616 | 0.485636725 | 1.740442127 | 0.081781414 | 0.327017878 |
| TNNC2 | 27.45605069 | 1.157887624 | 0.77307313 | 1.497772434 | 0.134192385 | 0.420619758 |
| TNNI3 | 452.3829486 | 1.619897748 | 0.541038505 | 2.994052609 | 0.002752985 | 0.051369596 |
| TNNI3K | 12.43859926 | 0.344731638 | 0.892660712 | 0.386184396 | 0.69936011 | 0.87117105 |
| TNNT1 | 3283.277354 | 0.917383281 | 0.446189156 | 2.056041185 | 0.039778547 | 0.229888601 |
| TNNT2 | 148.1318848 | 0.312558068 | 0.526675695 | 0.593454511 | 0.552877017 | 0.795672359 |
| TNNT3 | 21.39547732 | -1.299353029 | 0.788249903 | -1.648402395 | 0.099270125 | 0.361452832 |
| TNPO1 | 7375.830691 | -0.071562021 | 0.292883145 | -0.244336428 | 0.806970273 | 0.919137032 |
| TNPO2 | 2295.19216 | -0.105277106 | 0.319691217 | -0.329308721 | 0.741922353 | 0.892150449 |
| TNPO3 | 2718.786173 | -0.39745548 | 0.357854488 | -1.110662278 | 0.266713745 | 0.579921585 |
| TNR | 19.31402801 | -0.321543025 | 0.869695347 | -0.36971915 | 0.711591762 | 0.877287654 |
| TNRC18 | 1088.719006 | -0.905898161 | 0.395001313 | -2.293405442 | 0.021824674 | 0.167816586 |
| TNRC6A | 2412.087997 | 0.157235417 | 0.317435582 | 0.495330159 | 0.620367086 | 0.833252683 |
| TNRC6B | 1281.614221 | -0.544598148 | 0.294740889 | -1.847718349 | 0.064643098 | 0.29355313 |
| TNRC6C | 771.822668 | 0.229428014 | 0.351903394 | 0.651963061 | 0.514425002 | 0.77285333 |
| TNS1 | 75.11137075 | -0.575130512 | 0.656475532 | -0.876088267 | 0.380982051 | |

| | | | | | | |
|---|---|---|---|---|---|---|
| | | | | | | 0.682555348 |
| TNS2 | 20.05388092 | -0.514930185 | 0.684817484 | -0.751923245 | 0.452097218 | 0.732888936 |
| TNS3 | 3122.628273 | -1.038200397 | 0.428708722 | -2.421691805 | 0.015448446 | 0.140677297 |
| TNXB | 74.86363918 | 0.363721413 | 0.68817061 | 0.528533779 | 0.597128908 | 0.821867951 |
| TOB1 | 1154.068764 | 0.094621859 | 0.308809865 | 0.306408148 | 0.759293905 | 0.899449351 |
| TOB2 | 1646.142281 | -0.174265739 | 0.338719576 | -0.514483814 | 0.606913774 | 0.826788682 |
| TOB2P1 | 18.50588009 | -0.383039717 | 0.837165003 | -0.457543872 | 0.647280176 | 0.846597949 |
| TOE1 | 380.5757637 | -0.095599709 | 0.369008507 | -0.25907183 | 0.79557982 | 0.915261082 |
| TOGARAM1 | 424.0355821 | -0.056709345 | 0.412904708 | -0.137342453 | 0.890760116 | 0.957017706 |
| TOGARAM2 | 35.2354623 | 0.429913884 | 0.524678068 | 0.81938604 | 0.412566197 | 0.705244144 |
| TOLLIP | 575.1390942 | 0.233689818 | 0.390929834 | 0.597779442 | 0.54998711 | 0.793878561 |
| TOM1 | 706.0666911 | 0.184805211 | 0.369989671 | 0.499487489 | 0.617435999 | 0.831835425 |
| TOM1L1 | 1747.361444 | -0.270900592 | 0.288595144 | -0.938687284 | 0.347891324 | 0.654471837 |
| TOM1L2 | 321.250504 | -1.220582988 | 0.548706987 | -2.224471377 | 0.026116746 | 0.184458639 |
| TOMM20 | 10703.21227 | 0.306924869 | 0.276109853 | 1.11160419 | 0.26630837 | 0.57970088 |
| TOMM22 | 2451.320267 | 0.45223185 | 0.353499703 | 1.279299094 | 0.200791751 | 0.506881795 |
| TOMM34 | 1216.042779 | 0.1823981 | 0.303587045 | 0.600809893 | 0.547966614 | 0.792941769 |
| TOMM40L | 365.364239 | -0.012474314 | 0.270420581 | -0.046129308 | 0.963207187 | 0.987886441 |
| TOMM5 | 1578.315358 | 1.175846604 | 0.318398231 | 3.693006086 | 0.000221619 | 0.010157636 |
| TOMM6 | 2870.794035 | 0.914499891 | 0.398819951 | 2.293014402 | 0.021847176 | 0.167819496 |
| TOMM7 | 2583.721482 | 1.034516721 | 0.432241822 | 2.393374886 | 0.016694174 | 0.146184286 |
| TOMM70 | 2306.072666 | -0.076835828 | 0.2319081 | -0.331320158 | 0.740402667 | 0.891917214 |
| TONSL | 487.8745812 | 0.179921392 | 0.485728299 | 0.370415708 | 0.711072772 | 0.876976263 |
| TOP1 | 4400.110831 | 0.074653856 | 0.251639217 | 0.296670195 | 0.766718314 | 0.903470997 |
| TOP1MT | 1500.84522 | -0.355190836 | 0.259960432 | -1.366326532 | 0.171836497 | 0.474879827 |
| TOP2A | 9565.156522 | 1.028942018 | 0.353792716 | 2.908318828 | 0.003633777 | 0.062081109 |
| TOP2B | 7999.400953 | -0.206473388 | 0.253626155 | -0.814085548 | 0.415595944 | 0.707553144 |
| TOP3A | 498.9645502 | -0.265793545 | 0.317665286 | -0.836709445 | 0.402755909 | 0.699667005 |
| TOP3B | 770.3992996 | -0.267174559 | 0.425269054 | -0.628248298 | 0.529841295 | 0.780409931 |
| TOPBP1 | 1820.106798 | -0.50432652 | 0.336633829 | -1.498145688 | 0.134095403 | 0.420619758 |
| TOPORS | 1083.604261 | 0.113853581 | 0.250953605 | 0.453683784 | 0.65005644 | 0.847430646 |
| TOR1A | 948.2179829 | -0.196760722 | 0.285707814 | -0.688678128 | 0.491025843 | 0.758593654 |
| TOR1AIP1 | 815.2088959 | 0.165401356 | 0.240021665 | 0.68911011 | 0.490753978 | 0.758593654 |
| TOR1AIP2 | 1347.952711 | 0.06910362 | 0.364997929 | 0.189326061 | 0.849837271 | 0.938580023 |
| TOR1B | 563.8489947 | -0.403710619 | 0.275569424 | -1.465005125 | 0.14291952 | 0.433233718 |
| TOR2A | 239.7968794 | 0.007204321 | 0.330739155 | 0.021782486 | 0.982621465 | 0.993518428 |
| TOR3A | 565.3965497 | 0.092504288 | 0.338142884 | 0.27356568 | 0.784418411 | 0.910931322 |
| TOR4A | 58.77981768 | -0.982861382 | 0.840245914 | -1.16973063 | 0.242109388 | 0.553231259 |

| | | | | | | |
|---|---|---|---|---|---|---|
| TOX | 414.2201725 | -0.443904756 | 0.519630104 | -0.854270668 | 0.392955036 | 0.692071436 |
| TOX2 | 28.52912357 | 1.728392168 | 0.697383831 | 2.478394382 | 0.013197518 | 0.130266456 |
| TOX3 | 396.4245677 | 0.756162931 | 0.314766461 | 2.402298288 | 0.016292417 | 0.144292038 |
| TOX4 | 1710.825442 | -0.475577772 | 0.25601491 | -1.857617482 | 0.063223354 | 0.289778852 |
| TP53 | 2098.796458 | -0.07098969 | 0.286163568 | -0.248073824 | 0.804077288 | 0.918089602 |
| TP53BP1 | 2847.673128 | -0.214789144 | 0.444396058 | -0.483328195 | 0.628862723 | 0.837135782 |
| TP53BP2 | 3178.093487 | -0.662141105 | 0.242717006 | -2.728037545 | 0.006371236 | 0.086611125 |
| TP53I11 | 2750.933372 | 0.050865924 | 0.351767234 | 0.144601086 | 0.88502584 | 0.954416194 |
| TP53I13 | 270.5657844 | 0.600530914 | 0.471448886 | 1.273798564 | 0.202734814 | 0.509290789 |
| TP53I3 | 1097.381404 | 0.12219981 | 0.35895683 | 0.34043038 | 0.733532443 | 0.888496663 |
| TP53INP1 | 1582.55916 | -0.436394248 | 0.265067601 | -1.646350766 | 0.099691563 | 0.362001501 |
| TP53INP2 | 294.9285702 | -1.172536665 | 0.355009958 | -3.302827532 | 0.000957152 | 0.026449833 |
| TP53RK | 445.0229857 | 0.818499069 | 0.368639971 | 2.220320997 | 0.026396986 | 0.185660023 |
| TP63 | 74.9850666 | -2.834825374 | 0.968650052 | -2.926573293 | 0.003427187 | 0.059625986 |
| TP73 | 297.1333642 | -0.559577841 | 0.775176917 | -0.721871135 | 0.470373709 | 0.7454951 |
| TP73-AS1 | 1226.641406 | -0.206601844 | 0.31874648 | -0.648169804 | 0.51687513 | 0.773950969 |
| TPBG | 831.1932457 | 0.43244616 | 0.496781137 | 0.870496337 | 0.38402922 | 0.684699412 |
| TPCN1 | 446.3978855 | -0.451402242 | 0.36042687 | -1.252410071 | 0.210420478 | 0.51816316 |
| TPCN2 | 517.731764 | -0.674714403 | 0.331827058 | -2.033331481 | 0.042019045 | 0.235838301 |
| TPD52 | 1368.817654 | -0.18393539 | 0.404980015 | -0.454183868 | 0.649696493 | 0.84729896 |
| TPD52L1 | 374.9597235 | -0.389415266 | 0.57868574 | -0.672930468 | 0.500991528 | 0.763796851 |
| TPD52L2 | 2380.418648 | -0.354644584 | 0.253116693 | -1.401111002 | 0.161180882 | 0.46167973 |
| TPGS1 | 30.56833636 | 0.370262616 | 0.795962972 | 0.465175679 | 0.641805654 | 0.844038972 |
| TPGS2 | 3706.356108 | -0.166999315 | 0.233240606 | -0.71599589 | 0.473993887 | 0.748101062 |
| TPH1 | 15.42203587 | -0.757115012 | 0.800049956 | -0.946334671 | 0.343977915 | 0.6510091 |
| TPH2 | 33.55881901 | -0.179302521 | 0.587630136 | -0.305128192 | 0.76026852 | 0.899760802 |
| TPI1 | 15515.93615 | 0.675178028 | 0.272895313 | 2.474128336 | 0.013356171 | 0.131319811 |
| TPI1P2 | 25.43155878 | -2.24272946 | 0.735187397 | -3.050554824 | 0.00228419 | 0.045536718 |
| TPK1 | 38.1605714 | 0.673595536 | 0.574877642 | 1.171719836 | 0.241309557 | 0.552673201 |
| TPM1 | 14334.50203 | -1.074362022 | 0.473480871 | -2.26907165 | 0.023263969 | 0.174130017 |
| TPM2 | 3746.534559 | 1.380276373 | 0.281917296 | 4.896032951 | 9.78E-07 | 0.000180442 |
| TPM3 | 11701.4191 | -0.220598188 | 0.288361912 | -0.765004596 | 0.444268808 | 0.727151225 |
| TPM3P9 | 274.7564626 | -0.472720113 | 0.307100978 | -1.539298626 | 0.123731409 | 0.405045568 |
| TPM4 | 14549.19191 | -0.327977443 | 0.224372585 | -1.461753641 | 0.143808732 | 0.434973978 |
| TPMT | 736.7915919 | 0.462375249 | 0.303900394 | 1.521469723 | 0.128142004 | 0.412317905 |
| TPP1 | 2409.283199 | -0.845176794 | 0.281337224 | -3.004141369 | 0.002663315 | 0.050317756 |

| | | | | | | |
|---|---|---|---|---|---|---|
| TPP2 | 2195.365737 | -0.231559419 | 0.350598032 | -0.660469818 | 0.508952381 | 0.769344882 |
| TPPP | 12.03662916 | -0.236959268 | 1.056236432 | -0.224343017 | 0.822490406 | 0.925840841 |
| TPPP3 | 16.14116135 | 1.651731617 | 0.93766919 | 1.761529157 | 0.078148883 | 0.32060484 |
| TPR | 6420.754801 | -0.590221231 | 0.394204299 | -1.497247068 | 0.134328982 | 0.420620225 |
| TPRA1 | 731.1620771 | 0.123696935 | 0.294606588 | 0.419871586 | 0.674579266 | 0.859714553 |
| TPRG1 | 39.1425207 | 0.819164196 | 0.526514428 | 1.555824785 | 0.119749763 | 0.399101344 |
| TPRG1L | 959.4906475 | -0.400963252 | 0.28804286 | -1.39202635 | 0.163914419 | 0.465769928 |
| TPRKB | 1026.487188 | 0.866140672 | 0.366269432 | 2.364763737 | 0.018041576 | 0.151555235 |
| TPRN | 302.4535444 | 0.344543998 | 0.478767789 | 0.719647407 | 0.471742115 | 0.746399031 |
| TPRXL | 11.27961744 | -0.415353496 | 1.149829262 | -0.361230584 | 0.717927081 | 0.880671367 |
| TPST1 | 462.0123449 | 0.402955538 | 0.271912113 | 1.48193302 | 0.138358118 | 0.426606546 |
| TPST2 | 660.6799103 | 1.143067981 | 0.338565724 | 3.37620704 | 0.000734926 | 0.022388677 |
| TPT1 | 55737.21753 | 0.285809174 | 0.339910978 | 0.840835371 | 0.400440169 | 0.698047577 |
| TPTE2P1 | 32.19023316 | -1.579573996 | 0.617560791 | -2.557762763 | 0.010534793 | 0.11657429 |
| TPTE2P2 | 27.1573027 | -0.447384206 | 0.592794403 | -0.754703831 | 0.450426708 | 0.731865008 |
| TPTEP1 | 172.8864362 | -0.914348505 | 0.366164599 | -2.497096957 | 0.012521472 | 0.12669427 |
| TPTEP2 | 150.6841057 | -0.059848832 | 0.388174973 | -0.154180038 | 0.877467782 | 0.95079378 |
| TPTEP2-CSNK1E | 48.69999628 | -0.093577792 | 0.673104944 | -0.139024075 | 0.889431125 | 0.956341292 |
| TPX2 | 5158.27393 | 1.045331167 | 0.320433615 | 3.262239413 | 0.001105357 | 0.028759722 |
| TRA2A | 3465.775131 | 0.038847851 | 0.229246402 | 0.169458935 | 0.86543567 | 0.945250148 |
| TRA2B | 5951.967358 | 0.296151584 | 0.232959416 | 1.27125827 | 0.203636783 | 0.509986087 |
| TRABD | 707.7686753 | 0.028764145 | 0.490204466 | 0.058677851 | 0.953208701 | 0.983112195 |
| TRABD2A | 76.94357079 | 0.543324203 | 0.407926281 | 1.331917624 | 0.182887259 | 0.486816961 |
| TRABD2B | 80.77878097 | 0.935476632 | 0.583879901 | 1.602173034 | 0.109117351 | 0.379241044 |
| TRADD | 44.04212602 | -0.973424193 | 0.769578755 | -1.264879243 | 0.205914619 | 0.512218582 |
| TRAF1 | 89.20307023 | -0.29199735 | 0.518961754 | -0.562656781 | 0.573668619 | 0.808163661 |
| TRAF2 | 699.6559955 | 0.177534856 | 0.311454618 | 0.570018377 | 0.568665234 | 0.805450586 |
| TRAF3 | 227.6937931 | 0.987605188 | 0.415562318 | 2.376551351 | 0.017475329 | 0.149278273 |
| TRAF3IP1 | 395.415449 | -0.413934268 | 0.296086853 | -1.398016372 | 0.162108151 | 0.462717357 |
| TRAF3IP2 | 433.2824498 | -1.515023316 | 0.405068347 | -3.740167126 | 0.000183898 | 0.008913469 |
| TRAF4 | 4898.574968 | -0.018019411 | 0.436575448 | -0.041274449 | 0.967077102 | 0.988749915 |
| TRAF5 | 926.59447 | 0.075576209 | 0.521037342 | 0.145049505 | 0.884671786 | 0.95440284 |
| TRAF6 | 447.4781986 | -0.43705536 | 0.333997318 | -1.30855949 | 0.190683611 | 0.496058205 |
| TRAF7 | 2662.637566 | -0.218248287 | 0.345999384 | -0.630776518 | 0.528186659 | 0.779410544 |
| TRAFD1 | 1391.309114 | 0.196902558 | 0.246174344 | 0.799850035 | 0.42379769 | 0.712860381 |
| TRAIP | 509.2315573 | 0.429892656 | 0.281188681 | 1.528840543 | 0.126303979 | 0.409085752 |
| TRAK1 | 1790.686132 | -0.420155529 | 0.323722021 | -1.297889862 | 0.194325184 | 0.500187881 |
| TRAK2 | 678.9984346 | -0.825729096 | 0.26919916 | -3.067353914 | 0.00215963 | 0.044096562 |
| TRAM1 | 3481.462375 | -0.008534971 | 0.233082178 | -0.036617864 | 0.9707897 | 0.98957539 |
| TRAM1L1 | 205.7430713 | 0.278358792 | 0.395244479 | 0.704269905 | 0.481264705 | 0.752722364 |
| TRAM2 | 692.4320157 | -0.385286819 | 0.354098216 | -1.088078962 | 0.27656025 | |

0.590171608
TRANK1 332.7307364 -0.136554518 0.530191245 -0.257557098
0.796748744 0.915710299
TRAP1 3483.742152 0.056068869 0.312728808 0.179289109 0.857710698 0.942343063
TRAPPC1 1996.243541 0.196718611 0.373876631 0.526159152 0.598777632
0.823502444
TRAPPC10 1054.863476 -0.965134497 0.583936125 -1.652808339
0.098369876 0.360015486
TRAPPC11 1408.937473 -0.197359006 0.345304176 -0.57155117 0.567626084
0.805151447
TRAPPC12 520.9988381 -0.082042612 0.320971645 -0.25560704 0.798254282
0.916056735
TRAPPC13 564.7225087 0.215758888 0.257329353 0.838454245 0.401775635
0.699116854
TRAPPC2 431.01651 0.080622703 0.362113946 0.222644568 0.823812147
0.926468056
TRAPPC2B 171.502579 0.639761328 0.413218748 1.54823887 0.121564796
0.402062246
TRAPPC2L 1017.881682 0.384578923 0.26368997 1.458451087 0.144716248
0.435928729
TRAPPC3 1352.466276 0.236885726 0.32566442 0.727392098 0.466985794
0.743404143
TRAPPC3L 50.75330264 -0.046905779 0.592545889 -0.079159741
0.936905566 0.976155095
TRAPPC4 544.2721728 0.884826385 0.429015824 2.062456292 0.039164315
0.228085151
TRAPPC5 422.9892388 0.385405795 0.500737075 0.769676972 0.441491533
0.725034429
TRAPPC6A 628.6118016 0.532139987 0.512096055 1.03914096 0.298739184
0.610024066
TRAPPC6B 691.6366377 0.058169285 0.262635266 0.221483148 0.824716257
0.92681808
TRAPPC8 1375.882189 -0.307627386 0.274465965 -1.120821612
0.262363803 0.576023337
TRAPPC9 523.8280955 -0.66262545 0.350222448 -1.892013075 0.058489239
0.280248566
TRDMT1 505.1553878 0.295656948 0.285275319 1.036391614 0.300019481
0.610624571
TRDN 15.16293799 1.045955758 0.869840419 1.202468562 0.229182039 0.537806762
TREH 11.86605673 0.043906868 0.929133779 0.047255701 0.962309434 0.987343615
TRERF1 812.2220433 -0.097005373 0.384670001 -0.25217816 0.800903358
0.916680548
TREX1 37.9582029 2.286431578 0.668407029 3.420717437 0.000624562 0.020321012
TRH 32.69574223 -0.837900221 0.870481037 -0.962571482 0.335762615
0.644086928
TRIAP1 1504.473325 0.855852241 0.302224097 2.831846466 0.004628006
0.071864367
TRIB1 344.2130172 0.352498193 0.305726622 1.152984945 0.248916564 0.56168903
TRIB2 2094.667394 0.698369757 0.39188704 1.782069031 0.074737975 0.313842348
TRIB3 89.23373157 0.253269374 0.437331859 0.579123997 0.562505509 0.801706045
TRIL 611.0231208 0.192551092 0.312517182 0.616129618 0.537808976 0.786158891
TRIM11 796.781432 -0.085850228 0.308378098 -0.278392755
0.780710881 0.909271055
TRIM13 1718.509116 0.352354526 0.256185353 1.37538904 0.169010865
0.471508612
TRIM14 65.58598578 0.192841908 0.546984975 0.352554306 0.724422599
0.883670922
TRIM16 652.1556712 -0.329644768 0.24740404 -1.332414656
0.182723969 0.486683209
TRIM16L 332.1935011 -0.057052053 0.270401384 -0.210990241
0.832894887 0.931503747
TRIM17 90.01459406 -2.002767671 0.504896847 -3.966686824 7.29E-05
0.00492431
TRIM2 1673.157422 0.670618002 0.467988311 1.432980238 0.151863485 0.44767052

| | | | | | | |
|---|---|---|---|---|---|---|
| TRIM21 | 546.7101116 | -0.645967585 | 0.335150204 | -1.927397262 | 0.053930141 | 0.268429843 |
| TRIM22 | 602.4213734 | 1.296880344 | 0.614655174 | 2.10993155 | 0.034864252 | 0.213176891 |
| TRIM23 | 524.6148309 | 0.100931814 | 0.282363302 | 0.357453727 | 0.720752162 | 0.882172735 |
| TRIM24 | 2677.566538 | -0.431183618 | 0.371513338 | -1.160614099 | 0.245798869 | 0.557567979 |
| TRIM25 | 1018.21214 | -1.25638148 | 0.363689872 | -3.454540738 | 0.000551231 | 0.018583832 |
| TRIM26 | 248.5333486 | -1.138060759 | 0.569025194 | -2.000018224 | 0.045498296 | 0.245554838 |
| TRIM27 | 219.4625609 | 1.383181389 | 1.724007377 | 0.802305958 | 0.422375997 | 0.712222546 |
| TRIM28 | 14313.49477 | -0.088397391 | 0.255277742 | -0.346279274 | 0.729132843 | 0.886086417 |
| TRIM29 | 132.7681351 | -1.775815916 | 0.781818822 | -2.271390591 | 0.023123343 | 0.173699516 |
| TRIM3 | 307.5824944 | 0.189520522 | 0.412236591 | 0.459737264 | 0.645704819 | 0.845794147 |
| TRIM32 | 367.7002397 | 0.174448861 | 0.409671959 | 0.42582573 | 0.670234836 | 0.857925473 |
| TRIM33 | 2715.17466 | -0.090864634 | 0.333118071 | -0.272770054 | 0.785029981 | 0.910931322 |
| TRIM34 | 122.259398 | -0.87637412 | 0.489408879 | -1.790678832 | 0.073344848 | 0.311344591 |
| TRIM35 | 381.4357486 | -0.287414573 | 0.318818436 | -0.901499225 | 0.367322946 | 0.673921442 |
| TRIM36 | 312.1097587 | 0.395772251 | 0.375629348 | 1.053624411 | 0.292054908 | 0.605130431 |
| TRIM37 | 2171.54355 | -0.158398953 | 0.282197667 | -0.561304971 | 0.574589654 | 0.808359708 |
| TRIM38 | 1089.918306 | -0.727967594 | 0.731801694 | -0.994760739 | 0.319852646 | 0.629465081 |
| TRIM39 | 64.05828132 | -0.492060052 | 0.477841933 | -1.029754859 | 0.303125096 | 0.61357768 |
| TRIM4 | 592.6862012 | -0.338144807 | 0.276900516 | -1.221177959 | 0.222018648 | 0.531017642 |
| TRIM41 | 1225.547066 | -0.212924827 | 0.296977007 | -0.716974117 | 0.473390067 | 0.747714806 |
| TRIM44 | 1677.276785 | -0.135202413 | 0.316817165 | -0.426752172 | 0.669559846 | 0.857712777 |
| TRIM45 | 1013.855337 | -0.49308302 | 0.316466242 | -1.558090423 | 0.119211815 | 0.39809871 |
| TRIM46 | 311.7557332 | 0.217944059 | 0.421807936 | 0.516690277 | 0.605372384 | 0.826077544 |
| TRIM5 | 1155.100762 | -0.323962772 | 0.317514081 | -1.020309938 | 0.307581492 | 0.617319594 |
| TRIM52 | 186.7994527 | -0.521187667 | 0.32663329 | -1.595635482 | 0.110570201 | 0.382690751 |
| TRIM55 | 124.9659731 | -3.341740118 | 0.816455232 | -4.092986348 | 4.26E-05 | 0.003279499 |
| TRIM56 | 476.6901943 | 0.050538544 | 0.447746206 | 0.112873192 | 0.91013109 | 0.966444934 |
| TRIM58 | 28.40130792 | -2.515896961 | 1.071038376 | -2.349025971 | 0.018822594 | 0.155449937 |
| TRIM59 | 1227.155162 | 0.575804455 | 0.261104463 | 2.205264705 | 0.027435525 | 0.189451229 |
| TRIM6 | 925.3166772 | -1.217459848 | 0.324558998 | -3.751120305 | 0.000176046 | 0.008728094 |
| TRIM6-TRIM34 | 16.5901202 | -0.755952498 | 1.918445614 | -0.394044268 | 0.693548357 | 0.868790368 |
| TRIM61 | 15.41937182 | -0.097449459 | 0.751423009 | -0.129686552 | 0.896814423 | 0.95928311 |
| TRIM62 | 395.4577707 | -0.832344907 | 0.267543131 | -3.111068123 | | |

| | | | | | | |
|---|---|---|---|---|---|---|
| | | | | | 0.001864119 | 0.040171241 |
| TRIM65 | 688.9102221 | 1.000973083 | 0.313195608 | 3.19599974 | 0.001393472 | 0.03385493 |
| TRIM66 | 297.9280691 | -0.789458922 | 0.459662698 | -1.71747441 | 0.085892526 | 0.336017409 |
| TRIM68 | 384.6199622 | -0.839047092 | 0.288280418 | -2.910524061 | 0.003608232 | 0.061783843 |
| TRIM69 | 18.74109555 | 0.39031197 | 0.758711225 | 0.514440748 | 0.606943876 | 0.826788682 |
| TRIM71 | 5475.206395 | -0.007891543 | 0.394407264 | -0.020008614 | 0.984036501 | 0.994061643 |
| TRIM72 | 23.70784809 | 0.748563797 | 0.846228048 | 0.884588734 | 0.376378485 | 0.680079109 |
| TRIM73 | 60.25600191 | 0.23264706 | 0.497961208 | 0.467199164 | 0.640357392 | 0.843615231 |
| TRIM8 | 535.7668444 | 0.677852876 | 0.414362074 | 1.635895072 | 0.10186155 | 0.366107929 |
| TRIM9 | 26.92461386 | 0.540296398 | 0.784726647 | 0.688515422 | 0.491128262 | 0.758593654 |
| TRIML1 | 44.88535127 | -0.431555255 | 0.993188032 | -0.434515158 | 0.66391439 | 0.854739071 |
| TRIML2 | 2788.07048 | -5.163383393 | 1.311460809 | -3.937123669 | 8.25E-05 | 0.005346423 |
| TRIO | 1621.783855 | -0.59851227 | 0.455130271 | -1.315035077 | 0.188498114 | 0.493818838 |
| TRIOBP | 1319.254563 | -0.518858033 | 0.295730445 | -1.754496507 | 0.079345468 | 0.322931692 |
| TRIP10 | 2852.291735 | -0.149762545 | 0.286421922 | -0.522873893 | 0.601062014 | 0.824625774 |
| TRIP11 | 949.0754003 | -0.325715941 | 0.380737355 | -0.855487219 | 0.392281477 | 0.691514794 |
| TRIP12 | 4074.581498 | -0.439487036 | 0.329694235 | -1.333014015 | 0.182527205 | 0.486683209 |
| TRIP13 | 604.521845 | 0.35922866 | 0.334051331 | 1.075369643 | 0.282209267 | 0.595962806 |
| TRIP4 | 754.2578931 | -0.05072379 | 0.277113448 | -0.183043407 | 0.85476396 | 0.941593381 |
| TRIP6 | 2137.540307 | 0.014049539 | 0.432288597 | 0.032500369 | 0.974073022 | 0.99099127 |
| TRIQK | 1139.539295 | 0.111198716 | 0.271521575 | 0.409539154 | 0.682144039 | 0.862735748 |
| TRIR | 2749.024143 | 0.762657381 | 0.455835979 | 1.673096061 | 0.094308387 | 0.352141899 |
| TRIT1 | 822.8808099 | 0.398008649 | 0.331326024 | 1.201259847 | 0.229650421 | 0.538572659 |
| TRMO | 108.9174826 | -0.355241975 | 0.363106155 | -0.978341926 | 0.32790524 | 0.637775693 |
| TRMT1 | 1058.312394 | 0.225195122 | 0.329727913 | 0.682972576 | 0.494624172 | 0.760514211 |
| TRMT10A | 279.7234826 | 0.202570037 | 0.368040908 | 0.550400872 | 0.582044451 | 0.813018724 |
| TRMT10B | 224.8567605 | -0.121974832 | 0.341730833 | -0.356932474 | 0.721142358 | 0.882365571 |
| TRMT10C | 1197.187574 | 0.813470039 | 0.295582642 | 2.752090019 | 0.005921624 | 0.08318237 |
| TRMT11 | 792.6152184 | 0.849979623 | 0.302679312 | 2.808185392 | 0.004982154 | 0.07561376 |
| TRMT112 | 4712.368596 | 1.14947588 | 0.469516615 | 2.448211296 | 0.014356743 | 0.13604382 |
| TRMT12 | 456.4787032 | -0.085792002 | 0.281043293 | -0.305262585 | 0.760166169 | 0.899760802 |
| TRMT13 | 478.7055485 | 0.1614746 | 0.341183241 | 0.473278229 | 0.636014692 | 0.841016203 |
| TRMT1L | 775.3817175 | 0.122384262 | 0.311813238 | 0.392492193 | 0.694694579 | 0.86892666 |
| TRMT2A | 645.0526651 | 0.112233698 | 0.416953109 | 0.269175827 | 0.787794381 | 0.912200103 |
| TRMT2B | 421.8228131 | -0.33383765 | 0.2739871 | -1.218442949 | 0.223055692 | 0.532026087 |
| TRMT44 | 318.7133835 | -0.383582098 | 0.391653272 | -0.979392041 | 0.327386308 | 0.637370783 |

| | | | | | | |
|---|---|---|---|---|---|---|
| TRMT5 | 787.5089984 | 0.383062544 | 0.26033885 | 1.471399848 | 0.141183015 | 0.430963282 |
| TRMT6 | 900.1573579 | -0.025244972 | 0.296656927 | -0.085098206 | 0.932183316 | 0.97444726 |
| TRMT61A | 135.9379117 | 0.32379414 | 0.457979854 | 0.707005202 | 0.479563245 | 0.751829491 |
| TRMT61B | 575.5057795 | 0.31826826 | 0.321072789 | 0.991265128 | 0.321556136 | 0.631253641 |
| TRMT9B | 299.3564507 | -2.61316668 | 0.616631297 | -4.237810653 | 2.26E-05 | 0.002050447 |
| TRMU | 850.0916804 | 0.263491887 | 0.246295633 | 1.069819567 | 0.284700533 | 0.59855762 |
| TRNAU1AP | 741.3133932 | -0.088367871 | 0.331671886 | -0.266431599 | 0.789906832 | 0.912949942 |
| TRNP1 | 89.89768722 | -0.020599777 | 0.470959672 | -0.043740002 | 0.965111653 | 0.988700292 |
| TRNT1 | 691.3951573 | 0.153667495 | 0.242931488 | 0.632554867 | 0.527024368 | 0.778755887 |
| TRO | 2380.868216 | -0.314339366 | 0.387402254 | -0.81140304 | 0.417134254 | 0.708661116 |
| TROAP | 990.9195481 | 0.422278279 | 0.316087813 | 1.335952422 | 0.181564825 | 0.485768051 |
| TRPC1 | 475.9566483 | 0.076311323 | 0.277951044 | 0.274549509 | 0.783662362 | 0.910512111 |
| TRPC3 | 12.49414641 | -0.388898418 | 0.948354107 | -0.410077222 | 0.681749301 | 0.862620404 |
| TRPC4 | 84.02379046 | 1.307695733 | 0.813058195 | 1.608366709 | 0.107754888 | 0.376323529 |
| TRPC4AP | 2794.489143 | -0.430421686 | 0.286938702 | -1.500047508 | 0.133602097 | 0.420294949 |
| TRPC6 | 39.26632198 | -1.079221961 | 0.922207682 | -1.170259131 | 0.241896703 | 0.553182829 |
| TRPM3 | 63.11937759 | -0.448046138 | 0.686351001 | -0.65279447 | 0.513888789 | 0.772630754 |
| TRPM4 | 527.0117082 | -1.70392698 | 0.347149554 | -4.908336935 | 9.19E-07 | 0.000174186 |
| TRPM6 | 14.48962538 | -1.712339578 | 1.173849874 | -1.458738138 | 0.144637195 | 0.435928729 |
| TRPM7 | 1940.249248 | -0.737302417 | 0.419900155 | -1.755899369 | 0.079105593 | 0.322697216 |
| TRPM8 | 13.75248096 | 1.417532953 | 0.828983553 | 1.709965112 | 0.087272325 | 0.33929417 |
| TRPS1 | 110.0720402 | -0.330192962 | 0.421132632 | -0.784059313 | 0.433005308 | 0.718657207 |
| TRPT1 | 528.9716611 | 0.326344401 | 0.425400737 | 0.767145828 | 0.442994817 | 0.726245339 |
| TRPV1 | 125.0658001 | -0.303602123 | 0.345105016 | -0.879738366 | 0.37900106 | 0.681253679 |
| TRPV2 | 44.95991322 | 0.046344919 | 0.68861262 | 0.067301873 | 0.946341386 | 0.980414893 |
| TRPV3 | 113.6299183 | 0.160120518 | 0.504354071 | 0.317476406 | 0.75088214 | 0.896207755 |
| TRPV4 | 244.5566044 | -1.028533139 | 0.324306444 | -3.171485355 | 0.001516615 | 0.035838892 |
| TRPV6 | 386.4207779 | -4.759400395 | 0.79048725 | -6.020843971 | 1.74E-09 | 1.76E-06 |
| TRRAP | 2487.613402 | -0.597114649 | 0.449965998 | -1.327021712 | 0.184501501 | 0.48892092 |
| TRUB1 | 791.5636547 | 0.27506345 | 0.277426375 | 0.991482696 | 0.321449938 | 0.631253641 |
| TRUB2 | 1011.983613 | 0.027323633 | 0.335049712 | 0.081550983 | 0.935003781 | 0.975224148 |
| TSACC | 13.88885976 | -0.058797313 | 0.928865869 | -0.063300111 | 0.949527527 | 0.981593436 |
| TSC1 | 1759.539647 | -0.772326033 | 0.37039785 | -2.085125583 | 0.037057903 | 0.220638452 |
| TSC2 | 1839.96336 | -0.757507347 | 0.435897608 | -1.737810286 | 0.082244248 | 0.328245329 |
| TSC22D1 | 4617.750305 | 0.025862171 | 0.252799527 | 0.102303081 | 0.91851611 | 0.969677217 |
| TSC22D2 | 998.0303762 | -0.478268649 | 0.287689456 | -1.662447611 | 0.096423059 | 0.35626747 |
| TSC22D3 | 223.7335169 | 0.281250653 | 0.387562909 | 0.725690323 | 0.468028636 | 0.744224533 |
| TSC22D4 | 594.1954728 | -0.407900215 | 0.513130978 | -0.794924167 | 0.426657615 | 0.714361085 |

| | | | | | | |
|---|---|---|---|---|---|---|
| TSEN15 | 884.2591274 | 0.555285301 | 0.269377031 | 2.061368411 | 0.039267906 | 0.228431056 |
| TSEN2 | 343.7701175 | 0.183081463 | 0.332684317 | 0.55031588 | 0.582102735 | 0.813025279 |
| TSEN34 | 59.34038617 | -0.245035076 | 0.637009918 | -0.38466446 | 0.70048603 | 0.872001128 |
| TSEN54 | 672.4931288 | 0.119961023 | 0.376149954 | 0.318918085 | 0.74978863 | 0.895672701 |
| TSFM | 826.146664 | 0.557408828 | 0.3893697 | 1.431567038 | 0.152267769 | 0.448119169 |
| TSG101 | 1865.005348 | 0.239926338 | 0.330699991 | 0.725510567 | 0.468138865 | 0.744224533 |
| TSGA10 | 112.3771603 | -0.383641381 | 0.381484834 | -1.005653034 | 0.314582503 | 0.624045314 |
| TSGA10IP | 86.86152159 | 1.57009535 | 0.766355674 | 2.048781529 | 0.040483483 | 0.231601821 |
| TSHR | 26.93564187 | 0.727919064 | 0.57788576 | 1.259624504 | 0.207804851 | 0.514566354 |
| TSHZ1 | 272.593723 | -0.039745421 | 0.707945857 | -0.056141893 | 0.955228771 | 0.984323736 |
| TSHZ2 | 75.73137662 | 0.328847595 | 0.840550438 | 0.391228867 | 0.695628073 | 0.86959241 |
| TSHZ3 | 86.66032022 | 1.94976678 | 0.941088291 | 2.071821314 | 0.038282108 | 0.225369757 |
| TSKU | 560.1152803 | -0.151730128 | 0.261764543 | -0.579643547 | 0.56215502 | 0.801706045 |
| TSN | 2835.248946 | 0.060577919 | 0.27720505 | 0.218531082 | 0.827015344 | 0.928082319 |
| TSNARE1 | 212.5198824 | 0.305184226 | 0.428120289 | 0.712846912 | 0.475940493 | 0.749091526 |
| TSNAX | 1289.637522 | 0.289208589 | 0.258791065 | 1.117536996 | 0.263764791 | 0.577444856 |
| TSNAX-DISC1 | 13.65620912 | 1.000175002 | 1.083517681 | 0.923081386 | 0.355964794 | 0.663487122 |
| TSNAXIP1 | 40.34427251 | -0.067720241 | 0.597936596 | -0.113256559 | 0.909827157 | 0.96630013 |
| TSPAN1 | 31.49586364 | -3.547134061 | 1.099463516 | -3.226240806 | 0.001254278 | 0.03140041 |
| TSPAN10 | 55.60481914 | -0.169667078 | 0.561638714 | -0.302092918 | 0.762581231 | 0.901372797 |
| TSPAN11 | 371.8149438 | 0.750859136 | 0.533314539 | 1.407910494 | 0.159157575 | 0.459030918 |
| TSPAN12 | 704.1731354 | 0.252112231 | 0.35758993 | 0.705031686 | 0.480790518 | 0.752563699 |
| TSPAN13 | 979.3242176 | 0.695020725 | 0.305059569 | 2.278311506 | 0.022708025 | 0.171650943 |
| TSPAN14 | 2999.370646 | -0.319169683 | 0.262021458 | -1.218105133 | 0.223184024 | 0.532210755 |
| TSPAN15 | 587.0861602 | -0.906120143 | 0.442014667 | -2.049977547 | 0.040366622 | 0.231601821 |
| TSPAN17 | 1006.875515 | 0.352983874 | 0.240310874 | 1.468863512 | 0.141869813 | 0.43212631 |
| TSPAN18 | 1629.479267 | -0.041654186 | 0.426556987 | -0.097652102 | 0.922208551 | 0.970640219 |
| TSPAN19 | 12.72054554 | 0.709862155 | 0.791211372 | 0.897183963 | 0.369620761 | 0.675245088 |
| TSPAN2 | 249.7889064 | -0.263731613 | 0.380315511 | -0.693454789 | 0.488024189 | 0.756726796 |
| TSPAN3 | 4380.918616 | 0.508925837 | 0.287912589 | 1.767640095 | 0.077121091 | 0.31844659 |
| TSPAN31 | 666.221045 | -0.060187202 | 0.280145743 | -0.214842464 | 0.829890144 | 0.93006304 |
| TSPAN33 | 989.0997875 | 0.462797652 | 0.32519915 | 1.423120729 | 0.15470116 | 0.452297418 |
| TSPAN4 | 495.4328036 | 0.705896769 | 0.433630483 | 1.627876259 | 0.103551126 | 0.369033154 |
| TSPAN5 | 2054.435807 | -0.279342088 | 0.23603097 | -1.183497607 | 0.236611994 | 0.547336452 |
| TSPAN6 | 5136.747703 | 0.472714084 | 0.314050558 | 1.505216507 | 0.132268423 | 0.418521131 |
| TSPAN7 | 231.4195522 | 0.791281196 | 0.377070374 | 2.098497391 | 0.03586123 | |

| | | | | | | |
|---|---|---|---|---|---|---|
| | | | | | | 0.216490931 |
| TSPAN9 | 756.4923812 | -0.09225731 | 0.292565849 | -0.315338616 | 0.752504569 | 0.896586461 |
| TSPEAR | 35.78281754 | -2.751982118 | 0.746539055 | -3.686320359 | 0.00022752 | 0.010319181 |
| TSPO | 67.55834432 | 1.111978748 | 0.728328374 | 1.526754673 | 0.126822027 | 0.410063292 |
| TSPOAP1 | 212.9635547 | 0.252538655 | 0.500878504 | 0.504191442 | 0.614126854 | 0.830316752 |
| TSPY26P | 432.7000039 | -0.115621141 | 0.280764215 | -0.41180868 | 0.680479659 | 0.862092435 |
| TSPYL1 | 2174.055213 | 0.12967004 | 0.282307181 | 0.4593225 | 0.646002593 | 0.84590511 |
| TSPYL2 | 1057.447096 | -0.601157586 | 0.444364492 | -1.352847937 | 0.176104215 | 0.480085722 |
| TSPYL4 | 1142.170608 | 0.055728767 | 0.328553268 | 0.169618667 | 0.865310041 | 0.945250148 |
| TSPYL5 | 1646.742031 | 0.035195646 | 0.278850726 | 0.1262168 | 0.899560313 | 0.960937157 |
| TSR1 | 2324.6536 | -0.301557638 | 0.324603879 | -0.929001954 | 0.352888074 | 0.660049928 |
| TSR2 | 1208.73274 | 0.187267379 | 0.344444326 | 0.543679673 | 0.586661923 | 0.816311844 |
| TSR3 | 904.1012962 | 0.447431021 | 0.487843228 | 0.917161488 | 0.359058024 | 0.666802166 |
| TSSC2 | 1179.923212 | -0.084311888 | 0.367875898 | -0.229185681 | 0.818724601 | 0.924316427 |
| TSSC4 | 486.0849011 | 0.071425418 | 0.515240619 | 0.138625362 | 0.889746201 | 0.956447219 |
| TSSK6 | 46.4902 | 0.23891748 | 0.601337274 | 0.397310278 | 0.691138669 | 0.868004248 |
| TST | 816.2389765 | 0.734007192 | 0.476074597 | 1.541790293 | 0.123124562 | 0.404662636 |
| TSTA3 | 662.5284023 | -0.438645556 | 0.459124534 | -0.955395592 | 0.339377682 | 0.646260212 |
| TSTD1 | 664.2411335 | 0.784327961 | 0.603421901 | 1.299800288 | 0.193669427 | 0.499686884 |
| TSTD2 | 847.4459709 | -0.108139516 | 0.404184153 | -0.267550114 | 0.789045634 | 0.912533261 |
| TSTD3 | 119.7484049 | 0.562093869 | 0.333354774 | 1.686173152 | 0.091762454 | 0.347684361 |
| TTBK1 | 18.98603282 | 1.14767163 | 0.826364833 | 1.388819543 | 0.164887631 | 0.467115865 |
| TTBK2 | 578.2009915 | 0.191872218 | 0.335971245 | 0.571097144 | 0.567933795 | 0.805288664 |
| TTC1 | 1686.285623 | 0.485833223 | 0.325791956 | 1.491237626 | 0.135899118 | 0.423730301 |
| TTC12 | 472.6895325 | 0.224157141 | 0.268445814 | 0.835018202 | 0.40370746 | 0.70028858 |
| TTC13 | 431.8087475 | 0.395212724 | 0.281870787 | 1.402106012 | 0.160883592 | 0.461479481 |
| TTC14 | 1182.627249 | -0.002074216 | 0.302094515 | -0.006866115 | 0.994521675 | 0.997876213 |
| TTC17 | 2723.997693 | -0.406786931 | 0.387069401 | -1.050940554 | 0.293285893 | 0.606273373 |
| TTC19 | 2136.908532 | 0.140951412 | 0.293067813 | 0.480951524 | 0.630550952 | 0.838177265 |
| TTC21A | 117.273784 | 0.262326136 | 0.478567379 | 0.548148802 | 0.583589733 | 0.8142179 |
| TTC21B | 479.89769 | 0.158421298 | 0.362422146 | 0.43711815 | 0.662025665 | 0.853756491 |
| TTC23 | 776.7470071 | -0.169457625 | 0.294450774 | -0.575504093 | 0.564950423 | 0.803342663 |
| TTC23L | 28.53021611 | 0.324929877 | 0.611081756 | 0.531728978 | 0.594913719 | 0.821091342 |
| TTC25 | 67.21913131 | 0.081143691 | 0.469739609 | 0.172741853 | 0.862854344 | 0.944730333 |
| TTC26 | 184.0702318 | 0.003405262 | 0.414934229 | 0.008206751 | 0.993452034 | 0.99730871 |
| TTC27 | 714.5830315 | 0.013202797 | 0.250007204 | 0.052809665 | 0.957883561 | 0.98520931 |
| TTC28 | 622.0036678 | 0.024609536 | 0.417755616 | 0.058908929 | 0.953024646 | 0.983108339 |
| TTC29 | 341.9685994 | -0.145509843 | 0.470014353 | -0.309585957 | 0.756875837 | 0.898563152 |
| TTC3 | 4165.225026 | 0.143757469 | 0.304004327 | 0.472879682 | 0.636299021 | 0.84124553 |
| TTC30A | 366.9428131 | 0.236139039 | 0.268798649 | 0.878497865 | 0.379673596 | 0.681418209 |
| TTC30B | 295.2708929 | -0.077821548 | 0.283923169 | -0.274093685 | 0.784012627 | 0.910669593 |
| TTC31 | 835.9331945 | -0.305380939 | 0.289797881 | -1.053772162 | 0.291987241 | 0.605072864 |
| TTC32 | 272.9685387 | 0.907640127 | 0.458776108 | 1.978394496 | 0.04788422 | 0.252503129 |

| | | | | | | |
|---|---|---|---|---|---|---|
| TTC33 | 652.0147502 | 0.645717263 | 0.364520768 | 1.771414192 | 0.07649185 | 0.317498731 |
| TTC34 | 20.06413514 | 0.122374195 | 0.929162644 | 0.13170374 | 0.895218627 | 0.958594142 |
| TTC37 | 2891.912329 | -0.12642485 | 0.384278866 | -0.328992462 | 0.742161386 | 0.892150449 |
| TTC38 | 614.4021085 | -0.029424984 | 0.291395425 | -0.100979566 | 0.919566681 | 0.969677217 |
| TTC39A | 64.45794255 | -0.642001828 | 0.437689106 | -1.466798737 | 0.142430813 | 0.432682794 |
| TTC39B | 175.2670912 | 0.413190046 | 0.46921958 | 0.880589949 | 0.378539801 | 0.681253679 |
| TTC39C | 311.5013016 | 0.350890645 | 0.335705849 | 1.045232444 | 0.295915552 | 0.607241288 |
| TTC3P1 | 133.4952517 | -0.635938456 | 0.572605779 | -1.110604326 | 0.2667387 | 0.579921585 |
| TTC4 | 788.5863007 | -0.005847335 | 0.243067677 | -0.024056408 | 0.980807615 | 0.993241851 |
| TTC5 | 501.5842684 | 0.140431771 | 0.277057364 | 0.506868936 | 0.612246783 | 0.829098987 |
| TTC7A | 565.7696454 | -0.38997471 | 0.420235699 | -0.927990437 | 0.35341253 | 0.660553718 |
| TTC7B | 702.5998348 | -0.400410657 | 0.23834031 | -1.679995541 | 0.092958183 | 0.349595587 |
| TTC8 | 405.7547416 | 0.156120065 | 0.279334306 | 0.558900432 | 0.576229675 | 0.808993188 |
| TTC9 | 403.5969385 | -0.558735119 | 0.286351028 | -1.951224422 | 0.051030355 | 0.259879663 |
| TTC9C | 862.3284205 | 0.159270811 | 0.244195741 | 0.652225997 | 0.514255392 | 0.772834923 |
| TTF1 | 1164.327904 | 0.279610243 | 0.386619487 | 0.723218183 | 0.469545849 | 0.744930517 |
| TTF2 | 1080.428078 | -0.277308551 | 0.477719561 | -0.580483977 | 0.561588289 | 0.801570466 |
| TTI1 | 1232.895059 | -0.326314189 | 0.442271089 | -0.737814876 | 0.460626952 | 0.738395128 |
| TTI2 | 427.4362422 | 0.414258066 | 0.268903186 | 1.540547259 | 0.123427013 | 0.404779768 |
| TTK | 2120.257543 | 0.594247843 | 0.350067846 | 1.697521921 | 0.08959803 | 0.342777615 |
| TTL | 1449.294315 | 0.029754852 | 0.364978545 | 0.081524935 | 0.935024495 | 0.975224148 |
| TTLL1 | 331.0677284 | 0.348817102 | 0.29320102 | 1.189685842 | 0.234169892 | 0.544210545 |
| TTLL11 | 11.32675272 | 0.312550223 | 1.104990706 | 0.28285326 | 0.777289325 | 0.907898743 |
| TTLL12 | 1234.717405 | 0.365311498 | 0.348204751 | 1.049128415 | 0.29411902 | 0.606454309 |
| TTLL3 | 659.1164921 | -0.370905606 | 0.350102287 | -1.059420688 | 0.289408232 | 0.60229533 |
| TTLL4 | 1816.737365 | -0.622791632 | 0.419144344 | -1.485864335 | 0.137315009 | 0.425712285 |
| TTLL5 | 1010.633439 | -0.850960593 | 0.414925643 | -2.050874916 | 0.040279129 | 0.231327132 |
| TTLL6 | 126.0596113 | 0.497397924 | 0.697208822 | 0.713413125 | 0.475590154 | 0.74892861 |
| TTLL7 | 597.5168592 | -0.830826486 | 0.469644177 | -1.769055227 | 0.076884659 | 0.31844659 |
| TTPA | 26.19248765 | -0.877260224 | 0.828226488 | -1.059203293 | 0.289507206 | 0.602398681 |
| TTPAL | 526.9077319 | -0.507920895 | 0.276738661 | -1.835381054 | 0.066449249 | 0.297462837 |
| TTYH1 | 52.21628772 | 1.930085696 | 0.836668024 | 2.306871592 | 0.02106198 | 0.16470868 |
| TTYH2 | 133.6146968 | -0.677650004 | 0.420662515 | -1.610911312 | 0.107199054 | 0.375111686 |
| TTYH3 | 3141.975583 | 0.245588852 | 0.344505529 | 0.712873473 | 0.475924056 | 0.749091526 |
| TUB | 430.031312 | 0.685068662 | 0.694859748 | 0.985909262 | 0.324177641 | 0.634184268 |
| TUBA1A | 14387.91527 | 0.598346877 | 0.255305575 | 2.343649869 | 0.019096089 | 0.156852604 |
| TUBA1B | 49253.50495 | 0.76059395 | 0.23882433 | 3.184742314 | 0.00144883 | 0.034614478 |
| TUBA1C | 10898.80749 | 0.543222162 | 0.390749982 | 1.390203933 | 0.16446696 | 0.466640778 |
| TUBA4A | 76.77291842 | 0.269751887 | 0.506564982 | 0.532511912 | 0.594371494 | 0.820642877 |

| | | | | | | |
|---|---|---|---|---|---|---|
| TUBB | 49315.0991 | -0.155828238 | 0.299856766 | -0.519675577 | 0.603289712 | 0.825211561 |
| TUBB2A | 1303.556925 | 0.395575153 | 0.350013946 | 1.130169692 | 0.258404728 | 0.57129165 |
| TUBB2B | 11239.24627 | 0.165605827 | 0.285910878 | 0.579221848 | 0.562439491 | 0.801706045 |
| TUBB3 | 3984.317368 | 0.226634961 | 0.342410833 | 0.661880232 | 0.508047982 | 0.768706122 |
| TUBB4A | 430.8047714 | -0.033203853 | 0.462748624 | -0.071753542 | 0.942798045 | 0.978574897 |
| TUBB4B | 6106.053695 | 0.669452863 | 0.250251835 | 2.675116698 | 0.007470324 | 0.096125775 |
| TUBB6 | 1940.950444 | 0.816277511 | 0.274064674 | 2.978411992 | 0.002897462 | 0.052996691 |
| TUBB8 | 19.87357472 | 1.186062837 | 0.675085858 | 1.756906655 | 0.078933721 | 0.322507823 |
| TUBBP5 | 55.57327627 | 0.105687949 | 0.478068826 | 0.221072664 | 0.825035855 | 0.926977119 |
| TUBD1 | 352.4299488 | 0.129582242 | 0.310391457 | 0.417480054 | 0.676327311 | 0.860199564 |
| TUBE1 | 318.7969784 | 0.367565354 | 0.332214017 | 1.106411334 | 0.268548522 | 0.581542504 |
| TUBG1 | 1803.155881 | 0.259128187 | 0.385415915 | 0.672333905 | 0.50137115 | 0.763991735 |
| TUBG2 | 484.6381583 | -0.308116149 | 0.299104913 | -1.030127343 | 0.302950231 | 0.613538084 |
| TUBGCP2 | 1400.811499 | -0.333073046 | 0.415496708 | -0.801626197 | 0.422769219 | 0.712269066 |
| TUBGCP3 | 693.4807581 | -0.356611756 | 0.796750998 | -0.447582441 | 0.654454578 | 0.8491303 |
| TUBGCP4 | 1004.400619 | -0.244280885 | 0.277044031 | -0.881740292 | 0.377917267 | 0.681247962 |
| TUBGCP5 | 601.4787897 | 0.017796005 | 0.319459389 | 0.055706627 | 0.95557552 | 0.984323736 |
| TUBGCP6 | 981.9193019 | -0.671899804 | 0.634599343 | -1.05877797 | 0.289700909 | 0.602474638 |
| TUFM | 6200.460664 | 0.216952204 | 0.294500271 | 0.73667913 | 0.461317501 | 0.738942035 |
| TUFT1 | 689.2547011 | -1.175358492 | 0.381189649 | -3.083395613 | 0.002046529 | 0.042697375 |
| TULP1 | 19.27814876 | -0.85624325 | 0.696184015 | -1.229909379 | 0.218731042 | 0.527143548 |
| TULP3 | 1091.777089 | -0.810660768 | 0.336081025 | -2.412099191 | 0.015860966 | 0.142656219 |
| TULP4 | 775.3807359 | -0.456825477 | 0.275697336 | -1.656981833 | 0.097523148 | 0.358846393 |
| TUNAR | 388.9329761 | 1.015244787 | 0.553900292 | 1.8329017 | 0.066817191 | 0.297817842 |
| TUSC1 | 201.9858456 | 0.76101999 | 0.648143423 | 1.174153688 | 0.240333472 | 0.551937498 |
| TUSC2 | 850.7141493 | 0.535702003 | 0.37655413 | 1.422642752 | 0.154839744 | 0.452575411 |
| TUSC3 | 2851.589291 | 0.208939712 | 0.342572904 | 0.609913128 | 0.541919355 | 0.788704772 |
| TUT1 | 606.3826477 | 0.012562467 | 0.309574909 | 0.040579731 | 0.967630943 | 0.989014473 |
| TUT4 | 2265.048531 | -0.008546729 | 0.485780513 | -0.017593807 | 0.985962897 | 0.994881484 |
| TUT7 | 749.2080934 | -0.742244423 | 0.389186192 | -1.907170497 | 0.056498513 | 0.275406162 |
| TVP23A | 56.68407122 | 0.734795347 | 0.48838995 | 1.504525937 | 0.132446 | 0.418524946 |
| TVP23B | 720.3231601 | 0.356141094 | 0.285916153 | 1.245613756 | 0.21290623 | 0.52012889 |
| TVP23C | 129.5541855 | -1.005687893 | 0.432122862 | -2.327319337 | 0.019948275 | 0.159698109 |
| TVP23C-CDRT4 | 501.3531327 | -1.462005655 | 0.756385076 | -1.932885381 | 0.053250319 | 0.267056023 |
| TWF1 | 1406.414179 | 0.014768273 | 0.297893274 | 0.049575718 | 0.960460497 | 0.986479967 |
| TWF2 | 760.7067931 | 0.280154108 | 0.3756011 | 0.745882024 | 0.455738685 | 0.735306344 |
| TWIST1 | 10.75372981 | -0.326669785 | 0.90481805 | -0.361033674 | 0.718074274 | 0.880671367 |
| TWISTNB | 1437.191597 | 0.021668058 | 0.345440927 | 0.062725797 | 0.949984855 | 0.981636196 |
| TWNK | 588.1988752 | -0.095914949 | 0.307626607 | -0.31179016 | 0.755200003 | 0.897683871 |

| | | | | | | |
|---|---|---|---|---|---|---|
| TWSG1 | 1305.652716 | 0.180099671 | 0.269085204 | 0.669303509 | 0.503301889 | 0.765246655 |
| TXLNA | 1557.307856 | -0.191235805 | 0.392352769 | -0.487407813 | 0.625969358 | 0.835544329 |
| TXLNG | 815.8907119 | 0.145694475 | 0.281043147 | 0.518406077 | 0.604174975 | 0.82576988 |
| TXLNGY | 1722.097634 | 0.011084106 | 0.319692251 | 0.034671177 | 0.972341945 | 0.989954357 |
| TXN | 4729.002332 | 1.288997827 | 0.420881713 | 3.062613054 | 0.002194136 | 0.044516481 |
| TXN2 | 2027.067854 | 0.445103629 | 0.296711595 | 1.500122126 | 0.133582771 | 0.420294949 |
| TXNDC11 | 217.6000766 | -0.397998476 | 0.314289055 | -1.266345329 | 0.205389475 | 0.511820585 |
| TXNDC12 | 1388.815672 | 0.035061775 | 0.264419349 | 0.132599129 | 0.894510422 | 0.958237013 |
| TXNDC15 | 709.1732104 | -0.063554306 | 0.243746499 | -0.26073936 | 0.794293511 | 0.914628224 |
| TXNDC16 | 533.3410172 | 0.167047224 | 0.397729041 | 0.420002582 | 0.674483567 | 0.859684611 |
| TXNDC17 | 1486.038659 | 1.217471359 | 0.433743089 | 2.806895121 | 0.005002152 | 0.07561376 |
| TXNDC5 | 3147.308852 | 0.410974036 | 0.289896997 | 1.417655373 | 0.156291405 | 0.454843067 |
| TXNDC9 | 1104.52471 | 0.615913794 | 0.321156998 | 1.917796586 | 0.055136809 | 0.271793713 |
| TXNIP | 3934.602465 | 1.144921194 | 0.483359511 | 2.368674181 | 0.017851972 | 0.150854036 |
| TXNL1 | 2407.631167 | 0.36507832 | 0.273699028 | 1.333867801 | 0.182247186 | 0.486415681 |
| TXNL4A | 1510.094484 | 1.007074449 | 0.366246008 | 2.749721301 | 0.005964597 | 0.083399911 |
| TXNL4B | 298.4446498 | -0.114789894 | 0.295787405 | -0.388082425 | 0.697955037 | 0.870467 |
| TXNRD1 | 3847.731777 | 0.130471668 | 0.379869821 | 0.343464158 | 0.7312493 | 0.887005928 |
| TXNRD2 | 157.65756 | -0.23526936 | 0.378122863 | -0.622203478 | 0.533808079 | 0.783156734 |
| TXNRD3 | 146.4711212 | -0.368518579 | 0.49534957 | -0.743956594 | 0.456902735 | 0.735735031 |
| TYK2 | 671.442087 | -0.128946712 | 0.385042808 | -0.334889288 | 0.737708603 | 0.891065064 |
| TYMS | 3197.112572 | 1.100320757 | 0.278700575 | 3.948039064 | 7.88E-05 | 0.005152513 |
| TYRO3 | 5844.220427 | -0.577980885 | 0.236833783 | -2.440449488 | 0.014668997 | 0.137883121 |
| TYSND1 | 416.0927668 | 0.715469776 | 0.307447516 | 2.327128174 | 0.019958444 | 0.159698109 |
| TYW1 | 688.9456634 | -0.254587099 | 0.287335661 | -0.886026808 | 0.375603081 | 0.67966045 |
| TYW1B | 60.20720875 | 0.125771021 | 0.422680941 | 0.297555459 | 0.766042474 | 0.903282323 |
| TYW3 | 1268.521898 | 0.588228728 | 0.24545454 | 2.396487462 | 0.01655306 | 0.145665006 |
| TYW5 | 558.3463971 | 0.126835771 | 0.248784881 | 0.509821056 | 0.610176833 | 0.828605673 |
| U2AF1 | 1327.047782 | 0.746645219 | 0.621918383 | 1.200551775 | 0.229925117 | 0.538883664 |
| U2AF1L4 | 221.4663148 | 0.050757297 | 0.348110461 | 0.145808019 | 0.884072945 | 0.954046354 |
| U2AF1L5 | 4175.481485 | 0.333560026 | 0.294222994 | 1.13369802 | 0.25692124 | 0.569347376 |
| U2AF2 | 5012.559848 | 0.05185814 | 0.375823181 | 0.137985476 | 0.890251896 | 0.956787441 |
| U2SURP | 4169.245206 | -0.042892732 | 0.264806527 | -0.161977623 | 0.871323475 | 0.948581876 |
| UACA | 1503.263715 | 0.267380451 | 0.320433844 | 0.834432616 | 0.404037244 | 0.700367583 |
| UAP1 | 802.9975828 | 0.542185719 | 0.287087855 | 1.888570727 | 0.058949373 | 0.281409985 |
| UAP1L1 | 66.67677709 | -0.502005063 | 0.539696524 | -0.930161749 | 0.352287343 | 0.659576859 |
| UBA1 | 9867.221783 | -0.182076437 | 0.335807369 | -0.542205008 | 0.587677286 | 0.81697536 |
| UBA2 | 6500.900537 | 0.453456711 | 0.273627257 | 1.657205922 | 0.09747785 | 0.358846393 |
| UBA3 | 2629.651673 | 0.252314213 | 0.227376858 | 1.109674109 | 0.267139484 | 0.580460198 |
| UBA5 | 1359.292847 | 0.168380767 | 0.291267177 | 0.578097296 | 0.563198432 | 0.801890319 |
| UBA52 | 15223.87486 | 0.664837496 | 0.452920071 | 1.467891443 | 0.142133713 | 0.432308365 |

| | | | | | | |
|---|---|---|---|---|---|---|
| UBA6 | 1778.000064 | 0.013929133 | 0.236858109 | 0.058807922 | 0.953105099 | 0.983108339 |
| UBA7 | 12.47607134 | 0.946501955 | 1.068486912 | 0.885833925 | 0.375707026 | 0.679686536 |
| UBAC1 | 1211.43032 | 0.473567254 | 0.301028477 | 1.5731643 | 0.115680779 | 0.391826994 |
| UBAC2 | 2238.388431 | 0.084072121 | 0.260396172 | 0.322862355 | 0.746799487 | 0.894109554 |
| UBALD1 | 254.3676744 | -1.114618615 | 0.683767389 | -1.630113739 | 0.10307746 | 0.368123388 |
| UBALD2 | 316.2618382 | 0.639859543 | 0.51591482 | 1.24024261 | 0.214885672 | 0.522125669 |
| UBAP1 | 1019.259576 | -0.297417462 | 0.324534724 | -0.916442649 | 0.359434775 | 0.66703422 |
| UBAP1L | 50.62850318 | -0.25210027 | 0.515937112 | -0.488625966 | 0.625106527 | 0.835033444 |
| UBAP2 | 2170.098692 | -0.76348634 | 0.227074064 | -3.362278932 | 0.00077302 | 0.023268821 |
| UBAP2L | 4325.804022 | -0.46073627 | 0.430613596 | -1.06995291 | 0.284640505 | 0.59851436 |
| UBASH3B | 261.7076772 | -1.744718496 | 0.663916943 | -2.627916809 | 0.008590952 | 0.104788117 |
| UBB | 21334.72136 | 0.513021559 | 0.357906696 | 1.433394693 | 0.151745074 | 0.447448886 |
| UBBP4 | 39.42321279 | 1.387158468 | 0.814986196 | 1.702063759 | 0.088743417 | 0.342301733 |
| UBC | 33705.66348 | -0.102632231 | 0.264625636 | -0.387839337 | 0.698134933 | 0.870467 |
| UBE2A | 1396.525439 | 0.403494356 | 0.372332875 | 1.083692532 | 0.278501148 | 0.592005173 |
| UBE2B | 1134.523002 | 0.403118366 | 0.397135957 | 1.015063883 | 0.310075377 | 0.619783075 |
| UBE2C | 2285.537266 | 1.529129263 | 0.396085667 | 3.86060237 | 0.000113108 | 0.006724314 |
| UBE2D1 | 1390.063992 | 0.196336786 | 0.369767105 | 0.530974181 | 0.595436671 | 0.821514163 |
| UBE2D2 | 3372.135281 | 0.627293972 | 0.381043043 | 1.646254888 | 0.099711293 | 0.362001501 |
| UBE2D3 | 9211.364506 | 0.203292521 | 0.250042099 | 0.813033173 | 0.416199039 | 0.708127819 |
| UBE2D4 | 179.8279195 | 4.14E-05 | 0.397890411 | 0.0001041 | 0.99991694 | 0.99991694 |
| UBE2E1 | 3445.602321 | 0.294637141 | 0.376482438 | 0.782605272 | 0.433858945 | 0.719273746 |
| UBE2E2 | 692.9090566 | 0.160574246 | 0.391012567 | 0.410662622 | 0.681319939 | 0.862490397 |
| UBE2E3 | 3010.827038 | 0.656775638 | 0.376258895 | 1.745541826 | 0.080890598 | 0.32534233 |
| UBE2F | 1172.864189 | 0.127753178 | 0.380506549 | 0.335745019 | 0.737063154 | 0.890781894 |
| UBE2F-SCLY | 29.25609026 | -7.486983651 | 1.592462863 | -4.701512246 | 2.58E-06 | 0.000391779 |
| UBE2G1 | 3228.942582 | 0.142511574 | 0.301815385 | 0.472181278 | 0.636797402 | 0.841268221 |
| UBE2G2 | 2399.455573 | 0.113687205 | 0.295670515 | 0.3845064 | 0.700603153 | 0.872001128 |
| UBE2H | 5066.87254 | -0.456564219 | 0.232461938 | -1.9640386 | 0.049525616 | 0.25654413 |
| UBE2I | 5774.256839 | 0.500958364 | 0.325497647 | 1.539053715 | 0.123791183 | 0.405045568 |
| UBE2J1 | 1869.31017 | -0.541014195 | 0.405837822 | -1.333079782 | 0.182505624 | 0.486683209 |
| UBE2J2 | 940.8110934 | 0.05540407 | 0.310034547 | 0.178702892 | 0.858170998 | 0.942404069 |
| UBE2K | 3858.677955 | 0.200541235 | 0.258707061 | 0.775167226 | 0.438240852 | 0.72204083 |
| UBE2L3 | 3257.27535 | 0.279534042 | 0.315237818 | 0.88674019 | 0.375218795 | 0.679214334 |
| UBE2L6 | 1642.761997 | 0.304484968 | 0.323515106 | 0.941176972 | 0.346614176 | 0.653227784 |
| UBE2M | 1586.269024 | 0.582476844 | 0.368000348 | 1.582816015 | 0.113463405 | 0.38769219 |
| UBE2N | 3269.716601 | 0.459418735 | 0.283373885 | 1.621245852 | 0.104964923 | 0.37133017 |
| UBE2O | 1626.672636 | -0.015222788 | 0.402005239 | -0.037867137 | 0.969793615 | 0.989496332 |
| UBE2Q1 | 2332.361541 | 0.231300489 | 0.236209469 | 0.979217683 | 0.327472433 | 0.637370783 |

| | | | | | | |
|---|---|---|---|---|---|---|
| UBE2Q2 | 1222.128172 | -0.107872687 | 0.376055537 | -0.286853073 | 0.774224823 | 0.906279513 |
| UBE2Q2P1 | 17.3736093 | -0.804417734 | 0.74202715 | -1.084081268 | 0.278328767 | 0.592005173 |
| UBE2Q2P2 | 29.6448304 | -0.265483444 | 0.908062907 | -0.292362393 | 0.770009559 | 0.904795525 |
| UBE2R2 | 4144.22089 | 0.277832256 | 0.289869922 | 0.95847218 | 0.337824713 | 0.64515845 |
| UBE2S | 2684.054947 | 1.751039363 | 0.329220141 | 5.318749209 | 1.04E-07 | 3.96E-05 |
| UBE2T | 1055.939948 | 1.385535645 | 0.268254522 | 5.165003874 | 2.40E-07 | 7.44E-05 |
| UBE2V1 | 5841.029206 | -0.084011449 | 0.261863309 | -0.320821763 | 0.748345466 | 0.894441745 |
| UBE2V2 | 2742.671382 | 0.755672399 | 0.304224619 | 2.483929153 | 0.012994166 | 0.128720283 |
| UBE2W | 735.2768217 | 0.443157357 | 0.237200752 | 1.868279727 | 0.061723091 | 0.28627362 |
| UBE2Z | 2416.69453 | -0.210670288 | 0.229761298 | -0.916909377 | 0.35919013 | 0.666904107 |
| UBE3A | 2450.975074 | 0.429850355 | 0.220606824 | 1.948490747 | 0.051356271 | 0.260792387 |
| UBE3B | 1172.20829 | -0.677048944 | 0.282896612 | -2.393273424 | 0.016698792 | 0.146184286 |
| UBE3C | 1790.531478 | -0.393441402 | 0.370557605 | -1.061755034 | 0.288346908 | 0.601673803 |
| UBE3D | 499.176666 | 0.019889481 | 0.345522226 | 0.057563537 | 0.954096295 | 0.983863165 |
| UBE4A | 1677.096754 | -0.322623117 | 0.305759008 | -1.055154904 | 0.291354485 | 0.604386136 |
| UBE4B | 3275.393316 | -0.89087146 | 0.349037175 | -2.552368414 | 0.010699332 | 0.117622869 |
| UBFD1 | 2679.746993 | -0.355857709 | 0.248402201 | -1.432586776 | 0.151975963 | 0.447727291 |
| UBIAD1 | 472.8353372 | 0.030173679 | 0.280102947 | 0.107723534 | 0.914215001 | 0.968341533 |
| UBL3 | 1091.440801 | -0.358830202 | 0.43629615 | -0.82244641 | 0.410822875 | 0.704426187 |
| UBL4A | 811.3041558 | 0.124515194 | 0.274717902 | 0.453247467 | 0.650370556 | 0.847442386 |
| UBL5 | 4183.038343 | 1.416190672 | 0.49901894 | 2.837949739 | 0.004540433 | 0.070794359 |
| UBL7 | 995.367067 | -0.158395409 | 0.268100917 | -0.590805173 | 0.554650969 | 0.796360966 |
| UBLCP1 | 1776.500086 | 0.263885963 | 0.300801002 | 0.87727754 | 0.380335908 | 0.6820497 |
| UBN1 | 1671.590884 | -0.137205853 | 0.32882663 | -0.417258946 | 0.676489014 | 0.860199564 |
| UBN2 | 691.4083319 | -0.577967345 | 0.356078642 | -1.623145218 | 0.104558365 | 0.370706931 |
| UBOX5 | 256.9680006 | 0.047561495 | 0.298186165 | 0.159502687 | 0.873272841 | 0.949633274 |
| UBP1 | 2564.21866 | -0.355349123 | 0.243926671 | -1.456786674 | 0.145175276 | 0.436733637 |
| UBQLN1 | 4651.380747 | 0.124672862 | 0.267516774 | 0.466037551 | 0.641188623 | 0.843932467 |
| UBQLN2 | 1404.249209 | 0.136065715 | 0.268267501 | 0.507201634 | 0.612013349 | 0.829098987 |
| UBQLN4 | 2220.540362 | -0.078490899 | 0.243269412 | -0.322650093 | 0.746960252 | 0.89418504 |
| UBR1 | 932.1085876 | -0.320295875 | 0.36517799 | -0.877095237 | 0.380434911 | 0.682138996 |
| UBR2 | 1089.121311 | -0.727299117 | 0.298829341 | -2.433827668 | 0.014940106 | 0.139030867 |
| UBR3 | 586.7057568 | -0.079351189 | 0.363647727 | -0.218208952 | 0.82726631 | 0.928082319 |
| UBR4 | 4827.81088 | -0.736067604 | 0.545856191 | -1.348464333 | 0.177509084 | 0.481837595 |
| UBR5 | 4114.426241 | -0.487886505 | 0.373877474 | -1.304936879 | 0.191914342 | 0.497612799 |
| UBR7 | 1018.866798 | -0.005334585 | 0.299568899 | -0.017807538 | 0.985792391 | 0.9947756 |
| UBTD1 | 287.2574772 | 0.967669135 | 0.542025739 | 1.785282626 | 0.074215486 | 0.313236304 |

| | | | | | | |
|---|---|---|---|---|---|---|
| UBTD2 | 1617.296238 | 0.332321747 | 0.262733472 | 1.264862616 | 0.205920581 | 0.512218582 |
| UBTF | 2352.147422 | -0.203566545 | 0.261043903 | -0.779817275 | 0.435498436 | 0.720542613 |
| UBXN1 | 3345.760468 | 0.350529683 | 0.439852501 | 0.79692552 | 0.425494288 | 0.713673173 |
| UBXN10 | 54.24736342 | -0.37053603 | 0.447911663 | -0.827252472 | 0.408093976 | 0.702586667 |
| UBXN11 | 443.1580483 | 0.364005309 | 0.379022407 | 0.960379392 | 0.336864306 | 0.644459616 |
| UBXN2A | 700.3330888 | 0.388137134 | 0.321758271 | 1.20630041 | 0.22770168 | 0.536407171 |
| UBXN2B | 1327.384426 | -0.467769611 | 0.259945332 | -1.799492253 | 0.071940848 | 0.308679129 |
| UBXN4 | 3242.815836 | -0.115982854 | 0.25900571 | -0.447800373 | 0.654297274 | 0.849068645 |
| UBXN6 | 485.854924 | -0.398638758 | 0.456575091 | -0.873106673 | 0.382604937 | 0.683824871 |
| UBXN7 | 2244.110341 | -0.097621371 | 0.30985935 | -0.315050589 | 0.752723244 | 0.896586461 |
| UBXN8 | 423.096772 | 0.14058293 | 0.309391208 | 0.454385666 | 0.649551267 | 0.847284373 |
| UCHL1 | 7085.06037 | 1.056404263 | 0.394304509 | 2.679158464 | 0.007380746 | 0.095215383 |
| UCHL3 | 873.080327 | 0.75074996 | 0.400906845 | 1.872629437 | 0.061119574 | 0.28521841 |
| UCHL5 | 871.3882203 | 0.431258363 | 0.242302335 | 1.779835771 | 0.075102842 | 0.314600339 |
| UCK1 | 653.0142145 | 0.107169076 | 0.317072991 | 0.33799497 | 0.735366979 | 0.889724255 |
| UCK2 | 1302.708233 | 0.426085017 | 0.266878619 | 1.596549841 | 0.110366087 | 0.382187608 |
| UCKL1 | 497.0197887 | -0.021217344 | 0.400313883 | -0.053001769 | 0.957730498 | 0.98520931 |
| UCP2 | 843.4556903 | -0.228758074 | 0.287150161 | -0.796649643 | 0.425654537 | 0.713784126 |
| UCP3 | 17.3485139 | -0.251607664 | 0.911499423 | -0.276037108 | 0.782519565 | 0.909909835 |
| UEVLD | 365.5201087 | -0.302593216 | 0.319893269 | -0.945919298 | 0.34418975 | 0.65118919 |
| UFC1 | 2533.99905 | 0.117093838 | 0.350586806 | 0.333993852 | 0.738384198 | 0.891242475 |
| UFD1 | 1304.894237 | 0.045504543 | 0.288862016 | 0.157530381 | 0.874826866 | 0.949740014 |
| UFL1 | 1346.708439 | -0.181452476 | 0.242709889 | -0.747610561 | 0.454695093 | 0.734645643 |
| UFM1 | 1868.056323 | -0.095701597 | 0.310357458 | -0.308359262 | 0.757808977 | 0.899024084 |
| UFSP1 | 11.87952042 | 0.512737818 | 0.816932133 | 0.627638206 | 0.530240974 | 0.780679069 |
| UFSP2 | 763.0923201 | 0.255131587 | 0.303512877 | 0.840595594 | 0.400574528 | 0.698117882 |
| UGCG | 904.2557031 | 0.610979314 | 0.254185286 | 2.403676952 | 0.01623111 | 0.144168905 |
| UGDH | 1380.747015 | 0.069842477 | 0.237798296 | 0.2937047 | 0.768983562 | 0.904570801 |
| UGGT1 | 2698.450334 | -0.282545896 | 0.339584754 | -0.832033515 | 0.405390027 | 0.701111169 |
| UGGT2 | 621.3127364 | -0.005981389 | 0.275481793 | -0.021712466 | 0.98267732 | 0.993518428 |
| UGP2 | 4776.37878 | -0.663736097 | 0.357247873 | -1.857914763 | 0.06318112 | 0.289778852 |
| UGT1A1 | 24.90093514 | -0.60529272 | 0.621393979 | -0.974088486 | 0.330012606 | 0.638681113 |
| UGT3A1 | 419.6494979 | -1.371190531 | 0.482394616 | -2.842466492 | 0.004476594 | 0.070159507 |
| UGT3A2 | 990.5280207 | -1.80386819 | 0.438170495 | -4.116818017 | 3.84E-05 | 0.003051191 |
| UGT8 | 153.0139675 | 0.631649004 | 0.427196508 | 1.478591215 | 0.139249602 | 0.427816061 |
| UHMK1 | 1010.46594 | 0.006232116 | 0.306818386 | 0.020312068 | 0.983794429 | 0.994061643 |
| UHRF1 | 1735.850686 | 0.517837425 | 0.285084337 | 1.816435906 | 0.069303532 | 0.30336211 |
| UHRF1BP1 | 681.14773 | -0.521778434 | 0.436935696 | -1.194176715 | 0.232408849 | 0.542005591 |
| UHRF1BP1L | 529.3695545 | -0.261483714 | 0.333408922 | -0.784273293 | 0.432879767 | 0.718657207 |
| UHRF2 | 1714.51125 | -0.071307527 | 0.341858076 | -0.208588102 | 0.8347698 | 0.932107564 |
| UIMC1 | 1170.652678 | 0.041231901 | 0.272342145 | 0.151397431 | 0.879662216 | 0.952123159 |

| | | | | | | |
|---|---|---|---|---|---|---|
| ULBP1 | 16.6998371 | -0.191619101 | 0.771154611 | -0.248483375 | 0.803760431 | 0.917969611 |
| ULBP3 | 123.8954733 | 0.277618665 | 0.37990779 | 0.730752756 | 0.464930185 | 0.742000404 |
| ULK1 | 1170.598354 | -0.380404837 | 0.382652987 | -0.994124833 | 0.320162097 | 0.62990711 |
| ULK2 | 604.6429056 | 0.033010087 | 0.363335174 | 0.09085299 | 0.927609404 | 0.972633911 |
| ULK3 | 925.7931966 | -0.251920723 | 0.277040253 | -0.909328951 | 0.363176513 | 0.670041454 |
| ULK4 | 115.5931446 | -0.460579139 | 0.36190336 | -1.272657813 | 0.203139494 | 0.509545953 |
| UMAD1 | 403.5711633 | 0.112562672 | 0.295609658 | 0.380781443 | 0.70336543 | 0.872863553 |
| UMPS | 1269.115499 | 0.032500124 | 0.245839453 | 0.132200603 | 0.894825624 | 0.95833177 |
| UNC119 | 942.2946699 | 0.245061745 | 0.286309583 | 0.855932738 | 0.392034985 | 0.691496659 |
| UNC119B | 2429.924869 | 0.030786295 | 0.3263674 | 0.094330178 | 0.924846878 | 0.971800249 |
| UNC13A | 51.95845601 | -0.19923468 | 0.656947733 | -0.30327326 | 0.761681623 | 0.900715944 |
| UNC13B | 2075.422928 | -0.486446202 | 0.409493381 | -1.187922015 | 0.234864135 | 0.545403918 |
| UNC13C | 10.95521123 | 0.448840273 | 0.995172467 | 0.451017575 | 0.651976882 | 0.848078649 |
| UNC13D | 63.9926504 | 0.394020043 | 0.51504198 | 0.765025102 | 0.444256597 | 0.727151225 |
| UNC45A | 2311.668242 | -0.332313222 | 0.297511226 | -1.116977086 | 0.264004123 | 0.577448862 |
| UNC50 | 997.1345744 | 0.395429054 | 0.380384051 | 1.03955214 | 0.298548022 | 0.609920825 |
| UNC5A | 33.11886114 | 0.977769609 | 0.664806258 | 1.470758732 | 0.141356377 | 0.431058812 |
| UNC5B | 332.3764791 | 0.282692803 | 0.487431833 | 0.579963769 | 0.561939051 | 0.801706045 |
| UNC5C | 396.0771586 | -5.211305221 | 0.96546626 | -5.397708276 | 6.75E-08 | 3.12E-05 |
| UNC5CL | 109.551699 | -0.220822866 | 0.372158041 | -0.593357772 | 0.552941743 | 0.795672359 |
| UNC5D | 457.1823839 | 0.401453631 | 0.448479069 | 0.895144631 | 0.370709778 | 0.676128642 |
| UNC79 | 37.24162317 | -0.607080039 | 0.559362137 | -1.085307708 | 0.277785393 | 0.591968315 |
| UNC80 | 103.4671093 | -0.361670892 | 0.425361668 | -0.85026677 | 0.395176787 | 0.693972339 |
| UNC93B1 | 166.9285246 | -0.692931124 | 0.382150493 | -1.813241477 | 0.069794585 | 0.303603614 |
| UNG | 2314.677656 | 0.120205954 | 0.332082872 | 0.361975772 | 0.717370135 | 0.880516207 |
| UNK | 818.7614034 | -0.238386787 | 0.317000529 | -0.752007537 | 0.452046526 | 0.732888936 |
| UNKL | 423.2140017 | 0.24503001 | 0.396367293 | 0.618189276 | 0.536450578 | 0.785265508 |
| UPF1 | 1711.239396 | -0.374707852 | 0.427110356 | -0.877309218 | 0.380318707 | 0.6820497 |
| UPF2 | 1517.643509 | -0.382595992 | 0.223589787 | -1.711151469 | 0.087053155 | 0.338810522 |
| UPF3A | 2779.084628 | 0.235805888 | 0.346386513 | 0.680759437 | 0.49602372 | 0.761056737 |
| UPF3B | 2018.904758 | 0.442573857 | 0.410919268 | 1.077033596 | 0.281465257 | 0.595220158 |
| UPK1A | 14.37930982 | -1.285266879 | 0.8891239 | -1.445543055 | 0.148305409 | 0.442379346 |
| UPK2 | 70.33941608 | -1.32986873 | 0.915176638 | -1.453127927 | 0.146188239 | 0.438651457 |
| UPK3BL1 | 598.9356829 | 0.765576245 | 0.878685437 | 0.871274534 | 0.383604272 | 0.684585391 |
| UPP1 | 293.0631419 | -0.166783744 | 0.430003805 | -0.38786574 | 0.698115393 | 0.870467 |
| UPRT | 379.6366003 | -0.409981301 | 0.335111542 | -1.223417427 | 0.221172074 | 0.529995505 |
| UQCC1 | 1253.854438 | 0.086312505 | 0.255308227 | 0.338071774 | 0.735309101 | 0.889724255 |
| UQCC2 | 1755.238905 | 1.23751787 | 0.330382548 | 3.745711983 | 0.000179883 | 0.008851388 |
| UQCC3 | 253.3558472 | 1.397522901 | 0.562700443 | 2.483600143 | 0.013006176 | 0.128720283 |
| UQCR10 | 1975.030866 | 1.004359814 | 0.471668714 | 2.12937552 | 0.033223204 | 0.207932851 |

| | | | | | | |
|---|---|---|---|---|---|---|
| UQCR11 | 1494.230552 | 1.033809221 | 0.409063262 | 2.527260006 | 0.011495636 | 0.121279761 |
| UQCRB | 7926.171224 | 1.052158333 | 0.427613417 | 2.460536297 | 0.013872954 | 0.134140588 |
| UQCRC1 | 5340.886084 | 0.527735806 | 0.317769441 | 1.660750652 | 0.096763536 | 0.357177519 |
| UQCRC2 | 2499.398807 | 0.122631239 | 0.237491617 | 0.516360283 | 0.6056028 | 0.826148734 |
| UQCRFS1 | 4178.69838 | 1.009284955 | 0.434919199 | 2.320626351 | 0.020307019 | 0.161005801 |
| UQCRH | 9206.97393 | 0.995636295 | 0.433091607 | 2.298904617 | 0.021510356 | 0.166469811 |
| UQCRQ | 1062.914804 | 1.427816318 | 0.411886777 | 3.46652623 | 0.00052723 | 0.018055555 |
| URAHP | 33.28894299 | 1.660620146 | 0.684764447 | 2.425096912 | 0.015304301 | 0.140195951 |
| URB1 | 845.3301708 | -0.115645418 | 0.42378681 | -0.272885835 | 0.784940975 | 0.910931322 |
| URB2 | 516.2493193 | -0.035781391 | 0.349925885 | -0.1022542 | 0.918554908 | 0.969677217 |
| URGCP | 222.9010413 | -0.165769127 | 0.466128847 | -0.35562941 | 0.722118115 | 0.882776304 |
| URI1 | 1815.910331 | 0.411615503 | 0.251801657 | 1.634681471 | 0.102115839 | 0.366611924 |
| URM1 | 1572.237697 | 0.260665706 | 0.32831684 | 0.793945587 | 0.427227109 | 0.714721103 |
| UROD | 1420.56065 | 0.467625895 | 0.263449361 | 1.775012449 | 0.075895832 | 0.316062493 |
| UROS | 1273.670686 | 0.055496436 | 0.268330228 | 0.20682141 | 0.836149335 | 0.93270414 |
| USB1 | 1800.427113 | -0.492879158 | 0.232852568 | -2.116700544 | 0.034285271 | 0.211353857 |
| USE1 | 250.8405332 | 0.6263741 | 0.437244755 | 1.432548002 | 0.151987051 | 0.447727291 |
| USF1 | 1240.979219 | -0.153473715 | 0.261576824 | -0.586725203 | 0.557388279 | 0.798275991 |
| USF2 | 2118.913032 | 0.121588182 | 0.309681192 | 0.392623722 | 0.694597416 | 0.86892666 |
| USF3 | 498.8811899 | -0.269687578 | 0.364855841 | -0.739162014 | 0.459808624 | 0.73770692 |
| USH1C | 86.27747248 | 0.317033884 | 0.512736415 | 0.618317474 | 0.536366086 | 0.78521759 |
| USH1G | 116.7829141 | -0.523621267 | 0.473266746 | -1.106397758 | 0.268554395 | 0.581542504 |
| USHBP1 | 79.12249393 | -0.093263742 | 0.522719853 | -0.178420125 | 0.858393045 | 0.942476753 |
| USO1 | 4519.030398 | -0.045605931 | 0.312768893 | -0.145813512 | 0.884068609 | 0.954046354 |
| USP1 | 2175.527371 | 0.303097122 | 0.22651834 | 1.338068792 | 0.180874013 | 0.48558479 |
| USP10 | 3286.421274 | -0.088025289 | 0.224923452 | -0.391356653 | 0.695533628 | 0.869563155 |
| USP11 | 2498.780118 | -0.207586184 | 0.386579606 | -0.536981726 | 0.591280237 | 0.819070234 |
| USP12 | 823.2056189 | -0.341347645 | 0.353639818 | -0.965240981 | 0.334424125 | 0.642472578 |
| USP13 | 686.2203823 | 0.36115155 | 0.355458365 | 1.016016463 | 0.309621547 | 0.61915097 |
| USP14 | 2689.615468 | 0.087707439 | 0.241922321 | 0.362543806 | 0.716945694 | 0.880371898 |
| USP15 | 1007.951621 | 0.227194893 | 0.309924381 | 0.733065569 | 0.463518439 | 0.740758218 |
| USP16 | 1167.38457 | 0.173843184 | 0.294800577 | 0.58969757 | 0.555393425 | 0.796360966 |
| USP18 | 30.92683932 | -1.537893744 | 0.728293545 | -2.111639948 | 0.034717343 | 0.212806795 |
| USP19 | 1835.743306 | -0.435448335 | 0.354468176 | -1.228455371 | 0.219276073 | 0.52778634 |
| USP2 | 94.79574651 | 0.849481659 | 0.513651586 | 1.653809083 | 0.09816631 | 0.359903597 |
| USP20 | 643.5818893 | -0.264569551 | 0.397689947 | -0.665265875 | 0.505880453 | 0.767164369 |
| USP21 | 870.8520227 | -0.069043177 | 0.251821237 | -0.274175355 | 0.783949866 | 0.910666418 |
| USP22 | 8299.865076 | -0.365798216 | 0.330985304 | -1.105179629 | 0.269081762 | 0.582096023 |
| USP24 | 1917.862436 | -0.475068587 | 0.375852882 | -1.263974843 | 0.206239056 | 0.512653902 |
| USP25 | 1879.117509 | -0.808495831 | 0.299068513 | -2.703379977 | 0.006863823 | 0.090803834 |
| USP27X | 209.9626106 | -0.929724625 | 0.517090163 | -1.797993255 | | |

| | | | | | | |
|---|---|---|---|---|---|---|
| | | | | | 0.072178077 | 0.30923852 |
| USP28 | 1906.287804 | -1.252730725 | 0.25136216 | -4.983768146 | 6.24E-07 | 0.000139122 |
| USP30 | 469.9229586 | 0.041165431 | 0.32773117 | 0.125607311 | 0.900042775 | 0.961249486 |
| USP31 | 422.602451 | -0.148565111 | 0.355966262 | -0.417357279 | 0.676417098 | 0.860199564 |
| USP32 | 1313.686974 | -0.4316657 | 0.392488379 | -1.099817787 | 0.271411521 | 0.583632536 |
| USP32P1 | 116.3027722 | 0.683403199 | 0.565411521 | 1.208682833 | 0.226784719 | 0.535481989 |
| USP32P2 | 38.46926475 | 0.52596233 | 0.527593462 | 0.996908353 | 0.318808997 | 0.628804869 |
| USP32P3 | 14.54656786 | -0.817609545 | 0.996704233 | -0.820313106 | 0.412037637 | 0.705055604 |
| USP33 | 2161.703864 | -0.179346999 | 0.247031548 | -0.72600848 | 0.467833572 | 0.744166872 |
| USP34 | 3610.755852 | -0.448481787 | 0.332982728 | -1.346862009 | 0.178024679 | 0.48228793 |
| USP35 | 249.4550407 | -0.268809391 | 0.370431316 | -0.725665946 | 0.468043583 | 0.744224533 |
| USP36 | 1145.295775 | -0.110336303 | 0.291421649 | -0.378613955 | 0.704974558 | 0.873744266 |
| USP37 | 624.5640503 | -0.145436492 | 0.357913728 | -0.406345108 | 0.684489046 | 0.863857185 |
| USP38 | 403.9836106 | -0.386543501 | 0.32854246 | -1.176540472 | 0.239378969 | 0.550578888 |
| USP39 | 2178.834552 | -0.042130596 | 0.230523792 | -0.182760292 | 0.854986106 | 0.941701337 |
| USP4 | 1915.314172 | -0.367881332 | 0.250457772 | -1.468835761 | 0.141877342 | 0.43212631 |
| USP40 | 608.4861845 | -0.586583361 | 0.410310031 | -1.429610092 | 0.152828957 | 0.448899925 |
| USP42 | 583.0389601 | 0.29680803 | 0.267184307 | 1.110873737 | 0.266622701 | 0.579921585 |
| USP43 | 211.0810346 | -1.735496977 | 0.585322043 | -2.965029247 | 0.003026542 | 0.054402443 |
| USP44 | 769.1193719 | 1.555852971 | 0.565656018 | 2.750528451 | 0.005949922 | 0.083399911 |
| USP45 | 656.7418198 | 0.39273017 | 0.366469905 | 1.071657357 | 0.283873961 | 0.597896968 |
| USP46 | 2066.377637 | 0.158748683 | 0.278405718 | 0.570206258 | 0.568537812 | 0.805450586 |
| USP47 | 2126.8031 | -0.511176464 | 0.242879177 | -2.104653312 | 0.035321497 | 0.21486064 |
| USP48 | 1732.465191 | -0.306306328 | 0.284453135 | -1.076825284 | 0.281558327 | 0.595333991 |
| USP49 | 875.3072178 | -0.11362846 | 0.433922664 | -0.26186339 | 0.793426765 | 0.914254706 |
| USP5 | 3274.948766 | 0.133486404 | 0.276200811 | 0.483294758 | 0.628886462 | 0.837135782 |
| USP51 | 304.7338493 | 0.103188994 | 0.294640417 | 0.350220091 | 0.72617353 | 0.8844924 |
| USP53 | 1121.991844 | -0.652023145 | 0.267379507 | -2.438568134 | 0.014745579 | 0.138131624 |
| USP54 | 1270.213241 | -0.419415829 | 0.595859902 | -0.703883292 | 0.481505457 | 0.75277404 |
| USP6 | 66.40972101 | -1.180955972 | 0.673240625 | -1.754136526 | 0.079407116 | 0.322931692 |
| USP6NL | 983.330775 | -0.350548133 | 0.305782422 | -1.146397266 | 0.251630813 | 0.564383659 |
| USP7 | 3952.031045 | -0.292111828 | 0.264381113 | -1.10488917 | 0.269207616 | 0.582188661 |
| USP8 | 1849.541864 | -0.167102344 | 0.235938506 | -0.708245324 | 0.478792924 | 0.751165197 |
| USP9X | 4324.7582 | -0.371539883 | 0.350680292 | -1.059483214 | 0.28937977 | 0.60229533 |
| USP9Y | 2008.897536 | -0.557813488 | 0.310450738 | -1.79678583 | 0.072369627 | 0.309621999 |
| USPL1 | 855.7147368 | -0.037091666 | 0.361152468 | -0.102703619 | 0.918198201 | 0.969677217 |
| UST | 944.6791262 | 0.546686675 | 0.331444682 | 1.64940548 | 0.099064593 | 0.361363053 |

| | | | | | | |
|---|---|---|---|---|---|---|
| UTP11 | 960.1829112 | 0.606055888 | 0.326138017 | 1.85828041 | 0.063129204 | 0.289778852 |
| UTP14A | 874.1535613 | -0.544135763 | 0.298883556 | -1.820561055 | 0.068673607 | 0.301812004 |
| UTP14C | 261.0842274 | 0.138033887 | 2.017860785 | 0.068406051 | 0.945462405 | 0.979754792 |
| UTP15 | 557.4076344 | 0.260077472 | 0.329309961 | 0.789764972 | 0.429665039 | 0.716389544 |
| UTP18 | 1735.439487 | 0.785005883 | 0.267178356 | 2.938134269 | 0.00330194 | 0.058046037 |
| UTP20 | 1002.977129 | -0.284990306 | 0.424737354 | -0.670980087 | 0.502233215 | 0.764661936 |
| UTP23 | 406.0024572 | 0.426634522 | 0.304556606 | 1.400838181 | 0.161262468 | 0.46167973 |
| UTP25 | 386.5344429 | -0.271943458 | 0.268063946 | -1.01447234 | 0.310357421 | 0.620265109 |
| UTP3 | 1134.705091 | 0.074604765 | 0.40700575 | 0.183301501 | 0.854561457 | 0.941507034 |
| UTP4 | 1060.454098 | -0.741989545 | 1.330807656 | -0.557548299 | 0.577152869 | 0.809901794 |
| UTP6 | 1862.422432 | 0.089628778 | 0.242985754 | 0.368864335 | 0.712228848 | 0.877474733 |
| UTRN | 3706.871988 | -1.195219757 | 0.288120325 | -4.148335441 | 3.35E-05 | 0.002731607 |
| UTS2B | 11.74841983 | 0.364865286 | 0.880368916 | 0.414445899 | 0.678547581 | 0.861138002 |
| UTY | 793.6397551 | -0.725718372 | 0.362196798 | -2.003657615 | 0.04510675 | 0.24457273 |
| UVRAG | 624.3288159 | -0.554759448 | 0.34084438 | -1.627603331 | 0.103609022 | 0.369064484 |
| UVSSA | 780.1301463 | -0.420535528 | 0.383516359 | -1.096525658 | 0.272848801 | 0.585569269 |
| UXS1 | 1652.121811 | 0.750177024 | 0.301397292 | 2.488997222 | 0.012810397 | 0.128036515 |
| UXT | 907.1333458 | 0.559343628 | 0.367192262 | 1.523299061 | 0.127683902 | 0.411621861 |
| VAC14 | 1093.085694 | -0.439047744 | 0.368442211 | -1.191632585 | 0.233405342 | 0.543263647 |
| VAMP1 | 319.1988734 | 0.248155932 | 0.387783367 | 0.63993444 | 0.522215222 | 0.777251755 |
| VAMP2 | 882.3233258 | 0.389455537 | 0.395702165 | 0.984213812 | 0.325010396 | 0.634994554 |
| VAMP3 | 1679.225345 | -0.015590032 | 0.276713467 | -0.056339983 | 0.955070967 | 0.984323736 |
| VAMP4 | 349.4846504 | 0.354131207 | 0.26711353 | 1.325770383 | 0.184915766 | 0.489406194 |
| VAMP5 | 261.7873951 | 0.600939842 | 0.508533366 | 1.181711726 | 0.237320104 | 0.547879744 |
| VAMP7 | 1222.079562 | -0.154564911 | 0.249192181 | -0.620263888 | 0.535084066 | 0.784173142 |
| VAMP8 | 1188.888332 | -0.502773739 | 0.765231649 | -0.657021621 | 0.511167014 | 0.770636467 |
| VANGL1 | 747.2202784 | -0.809702784 | 0.280655098 | -2.885045707 | 0.003913568 | 0.064746724 |
| VANGL2 | 3231.737709 | 0.095556375 | 0.308902177 | 0.309341864 | 0.757061489 | 0.898563152 |
| VAPA | 2913.880339 | 0.560315982 | 0.270260173 | 2.073246592 | 0.038149336 | 0.224762556 |
| VAPB | 1748.276457 | 0.114879417 | 0.228063403 | 0.503717014 | 0.614460251 | 0.830593641 |
| VARS | 600.1187698 | 0.002856826 | 0.90652346 | 0.003151409 | 0.997485544 | 0.99877158 |
| VARS2 | 336.3583979 | 0.347172605 | 0.708861067 | 0.489761141 | 0.624302931 | 0.834816841 |
| VASH1 | 342.8729048 | 0.682510065 | 0.596024218 | 1.145104584 | 0.252165837 | 0.564998953 |
| VASH2 | 674.7772588 | 0.169773153 | 0.397164942 | 0.427462586 | 0.66904243 | 0.857484389 |
| VASN | 32.37027154 | -4.63453867 | 1.930081135 | -2.401214429 | 0.016340758 | 0.144378309 |
| VASP | 2385.872412 | 0.217339339 | 0.510900239 | 0.425404652 | 0.670541715 | 0.858101085 |
| VAT1 | 8729.949646 | -0.5564077 | 0.219273935 | -2.53750041 | 0.011164724 | 0.120212934 |
| VAT1L | 1798.903202 | 0.457837851 | 0.472311514 | 0.969355684 | 0.332367755 | 0.640217269 |
| VAV2 | 1649.076526 | -0.541545252 | 0.295785564 | -1.830871137 | 0.067119779 | 0.297817842 |
| VAV3 | 90.47814885 | -1.183523594 | 0.508256815 | -2.328593655 | 0.019880603 | 0.159572793 |
| VAX2 | 49.00277397 | -0.516628895 | 0.66072629 | -0.781910609 | 0.43426711 | 0.719429635 |
| VBP1 | 3014.5169 | 0.944278536 | 0.26894907 | 3.510993865 | 0.000446435 | 0.016477615 |
| VCAM1 | 35.48808807 | -0.30881187 | 0.934671484 | -0.330396161 | 0.741100642 | 0.891943909 |
| VCAN | 3803.613059 | 0.234550754 | 0.50163566 | 0.467571931 | 0.640090742 | 0.843538624 |

| | | | | | | |
|---|---|---|---|---|---|---|
| VCL | 5332.572258 | -0.294664802 | 0.302721002 | -0.973387379 | 0.330360809 | 0.638816405 |
| VCP | 10720.6571 | -0.169215143 | 0.261741177 | -0.646497982 | 0.517956904 | 0.773950969 |
| VCPIP1 | 462.1397901 | -0.073624429 | 0.296416652 | -0.248381556 | 0.803839203 | 0.917969611 |
| VCPKMT | 274.7484317 | 0.369369148 | 0.395912605 | 0.932956273 | 0.350842546 | 0.657923581 |
| VCX | 24.96851399 | 0.777231444 | 0.742603268 | 1.04663079 | 0.295269896 | 0.60699582 |
| VDAC1 | 6381.776426 | 0.470951832 | 0.310255569 | 1.517948037 | 0.129027499 | 0.413453061 |
| VDAC2 | 6161.885843 | 0.43286119 | 0.39610702 | 1.092788484 | 0.274486679 | 0.587352209 |
| VDAC3 | 3065.153638 | 0.601445967 | 0.227083261 | 2.648570236 | 0.008083305 | 0.100970518 |
| VEGFA | 504.3525412 | -0.176263236 | 0.346963168 | -0.50801714 | 0.611441323 | 0.829058038 |
| VEGFB | 276.660338 | 0.508810257 | 0.443108883 | 1.148273656 | 0.250855611 | 0.563811922 |
| VEGFC | 12.67978103 | 0.663058768 | 1.15749984 | 0.572837027 | 0.566755046 | 0.804513376 |
| VENTX | 65.96486795 | -0.346173009 | 0.816290224 | -0.424080798 | 0.671506889 | 0.858479663 |
| VEPH1 | 22.74320917 | -1.058852302 | 0.782239398 | -1.353616686 | 0.175858701 | 0.480020214 |
| VEZF1 | 1685.657236 | 0.326318624 | 0.28287335 | 1.1535856 | 0.248670105 | 0.561311435 |
| VEZT | 1925.848268 | 0.218577949 | 0.286916044 | 0.761818497 | 0.446168339 | 0.728589711 |
| VGF | 52.00042994 | 0.622160746 | 0.665297433 | 0.935161801 | 0.349704921 | 0.656280722 |
| VGLL1 | 198.4939601 | -1.381459245 | 1.014754585 | -1.361372754 | 0.173395922 | 0.477420967 |
| VGLL3 | 475.7481791 | 1.392875416 | 0.705017792 | 1.97565995 | 0.048193304 | 0.253165032 |
| VGLL4 | 1602.3342 | -0.085487191 | 0.259932051 | -0.32888284 | 0.742244245 | 0.892150449 |
| VHL | 1421.153799 | -0.263916317 | 0.270793012 | -0.974605343 | 0.329756062 | 0.638548751 |
| VIL1 | 42.22869202 | -1.39200533 | 0.716781839 | -1.942020924 | 0.052134567 | 0.263030768 |
| VILL | 208.4436629 | -1.207598112 | 0.495250524 | -2.438358071 | 0.014754151 | 0.138131624 |
| VIM | 24064.96299 | 1.169812792 | 0.454817391 | 2.572049386 | 0.010109847 | 0.114259456 |
| VIPAS39 | 1270.592732 | -0.443799089 | 0.254143884 | -1.7462513 | 0.080767294 | 0.325104967 |
| VIPR1 | 84.50685202 | -0.87199774 | 0.426141796 | -2.046261946 | 0.040730606 | 0.23195346 |
| VIPR2 | 21.55217954 | -2.969253455 | 1.277406423 | -2.324439114 | 0.020101971 | 0.160340171 |
| VIRMA | 1964.674318 | -0.316882602 | 0.335824654 | -0.94359541 | 0.345376439 | 0.652226246 |
| VKORC1 | 1078.445151 | 0.757775382 | 0.337074703 | 2.248093301 | 0.024570241 | 0.179037045 |
| VKORC1L1 | 1193.457048 | 0.428695788 | 0.282352796 | 1.518298364 | 0.1289392 | 0.413453061 |
| VLDLR | 230.2672624 | 0.542921312 | 0.504840132 | 1.075432156 | 0.282181292 | 0.595962806 |
| VMA21 | 1440.443277 | 0.780606575 | 0.385601966 | 2.024384325 | 0.042930623 | 0.238392703 |
| VMAC | 119.7093575 | -0.788944544 | 0.392441042 | -2.01035177 | 0.044393971 | 0.24217941 |
| VMP1 | 2343.609177 | -0.005710362 | 0.287781021 | -0.019842734 | 0.984168828 | 0.994129122 |
| VN1R1 | 40.33589015 | -1.032813296 | 0.662341554 | -1.559336401 | 0.11891678 | 0.397638633 |
| VN1R81P | 24.36969749 | -0.34497672 | 0.867785007 | -0.397537083 | 0.690971447 | 0.868004248 |
| VN1R83P | 16.45552282 | -1.306234309 | 1.148732215 | -1.137109495 | 0.255492515 | 0.567840162 |
| VOPP1 | 1400.012348 | 0.496240503 | 0.264901283 | 1.873303504 | 0.061026486 | 0.284871638 |
| VPS11 | 488.0594418 | -0.861125125 | 0.308394496 | -2.792284349 | 0.005233734 | 0.07701356 |
| VPS13A | 1358.094947 | -0.546319326 | 0.356560783 | -1.53219129 | 0.125475237 | 0.407445382 |

| | | | | | | |
|---|---|---|---|---|---|---|
| VPS13B | 897.7974116 | -0.598541865 | 0.531005968 | -1.127184817 | 0.259664348 | 0.572948272 |
| VPS13C | 1011.870403 | -0.368879343 | 0.378203508 | -0.975346171 | 0.329388575 | 0.638450757 |
| VPS13D | 879.3400735 | -0.7513491 | 0.484673211 | -1.550217926 | 0.121089219 | 0.401219509 |
| VPS16 | 1164.046383 | -0.318729519 | 0.320867232 | -0.993337702 | 0.32054541 | 0.63024811 |
| VPS18 | 925.8132492 | -0.444416318 | 0.279710806 | -1.588842146 | 0.112096035 | 0.384920166 |
| VPS25 | 2285.898705 | 0.373088233 | 0.291564456 | 1.279608077 | 0.200683008 | 0.506835677 |
| VPS26A | 1606.236639 | -0.074548017 | 0.255396905 | -0.291890838 | 0.770370089 | 0.905008875 |
| VPS26B | 1046.39383 | -0.318539469 | 0.304359654 | -1.04658901 | 0.295289173 | 0.60699582 |
| VPS26C | 1434.847885 | -0.015516037 | 0.251218172 | -0.061763194 | 0.950751414 | 0.981826318 |
| VPS28 | 211.1009621 | 0.237920123 | 0.505292938 | 0.470855825 | 0.637743696 | 0.84190825 |
| VPS29 | 1286.105323 | 0.460078676 | 0.348061096 | 1.321833097 | 0.186223735 | 0.490817669 |
| VPS33A | 1422.870769 | -0.275111019 | 0.232333726 | -1.184120035 | 0.23636555 | 0.547298804 |
| VPS33B | 734.9704533 | -0.18607402 | 0.361045145 | -0.515376048 | 0.606290267 | 0.826611695 |
| VPS35 | 3386.726709 | -0.359932894 | 0.265429574 | -1.356039151 | 0.175086712 | 0.479137654 |
| VPS35L | 1616.142278 | -0.62088223 | 0.333094817 | -1.863980458 | 0.062324449 | 0.287743217 |
| VPS36 | 1566.138956 | -0.090013869 | 0.225592043 | -0.399011718 | 0.68988457 | 0.867683896 |
| VPS37A | 817.44623 | 0.132499784 | 0.243245381 | 0.54471655 | 0.585948481 | 0.815639728 |
| VPS37C | 883.5563641 | -0.663167016 | 0.297123896 | -2.231954498 | 0.02561797 | 0.182550601 |
| VPS37D | 115.9141322 | 1.031736479 | 0.609608968 | 1.692456202 | 0.090559024 | 0.344933707 |
| VPS39 | 2298.009252 | -0.629719851 | 0.297191767 | -2.118900728 | 0.034098857 | 0.210632641 |
| VPS41 | 1817.504505 | -0.596830584 | 0.273138829 | -2.185081435 | 0.028882876 | 0.194385362 |
| VPS45 | 799.8493696 | -0.177556907 | 0.253107614 | -0.701507566 | 0.482986314 | 0.753923796 |
| VPS4A | 1275.522526 | 0.502829513 | 0.305398336 | 1.646471031 | 0.099666819 | 0.362001501 |
| VPS4B | 1398.382475 | -0.233421508 | 0.272123836 | -0.857776781 | 0.391015733 | 0.690530747 |
| VPS50 | 524.6707401 | -0.128296194 | 0.308869562 | -0.41537338 | 0.677868591 | 0.860856079 |
| VPS51 | 1534.25917 | -0.147743796 | 0.41795018 | -0.35349619 | 0.723716484 | 0.883448888 |
| VPS52 | 331.416995 | -2.195576694 | 0.485419715 | -4.523048049 | 6.10E-06 | 0.00074577 |
| VPS53 | 524.7814744 | -0.443241288 | 0.268103404 | -1.653247522 | 0.098280498 | 0.359965304 |
| VPS54 | 956.95788 | -0.15582801 | 0.277305832 | -0.561935565 | 0.574159921 | 0.808359708 |
| VPS72 | 2137.129131 | 0.335690297 | 0.298875944 | 1.123176035 | 0.261362739 | 0.574937735 |
| VPS8 | 1183.452525 | -0.474092729 | 0.461207899 | -1.027937141 | 0.303979397 | 0.614152542 |
| VPS9D1 | 167.6487728 | -0.026603553 | 0.483003183 | -0.055079457 | 0.956075162 | 0.984426596 |
| VRK1 | 1342.056177 | 0.97585952 | 0.263186935 | 3.707856995 | 0.000209021 | 0.009727153 |
| VRK2 | 675.4542593 | -0.120975857 | 0.263356279 | -0.459361962 | 0.645974259 | 0.84590511 |
| VRK3 | 878.1422546 | -0.24465949 | 0.238866111 | -1.024253665 | 0.305715477 | 0.615683213 |

| | | | | | | |
|---|---|---|---|---|---|---|
| VRTN | 597.8242258 | 0.934455486 | 0.495779945 | 1.884819051 | 0.059454272 | 0.282486927 |
| VSIG10 | 503.9728601 | -1.063632959 | 0.316738943 | -3.358074477 | 0.000784875 | 0.023439632 |
| VSIG10L | 240.2775293 | 0.411341903 | 0.306484987 | 1.342127414 | 0.179554687 | 0.484443207 |
| VSIR | 10.36288231 | -0.393319808 | 1.100202976 | -0.357497495 | 0.720719401 | 0.882172735 |
| VSNL1 | 332.9836271 | -0.145870921 | 0.756923564 | -0.192715524 | 0.847181765 | 0.938145276 |
| VSTM2L | 100.7508235 | 1.489325667 | 0.558492746 | 2.666687576 | 0.007660285 | 0.097741108 |
| VSTM4 | 24.91155709 | 0.548292997 | 0.901084117 | 0.60848148 | 0.542868186 | 0.789372509 |
| VSTM5 | 160.2719786 | -4.91400409 | 1.06983561 | -4.59323287 | 4.36E-06 | 0.00059117 |
| VTA1 | 2032.025447 | 0.293372822 | 0.234883073 | 1.249016446 | 0.211659059 | 0.518836577 |
| VTCN1 | 687.5977723 | -5.044865778 | 1.514254084 | -3.331584726 | 0.00086353 | 0.024906112 |
| VTI1A | 448.8253781 | -0.302577716 | 0.286153289 | -1.057397302 | 0.290330302 | 0.60328736 |
| VTI1B | 2810.449632 | 0.14103358 | 0.261027968 | 0.540300646 | 0.588989713 | 0.818050255 |
| VWA1 | 447.7416709 | -0.62397553 | 0.533038238 | -1.170601816 | 0.241758867 | 0.552977821 |
| VWA2 | 154.0378939 | -2.83647779 | 0.822733208 | -3.447627691 | 0.000565533 | 0.018856486 |
| VWA5A | 876.8101355 | -0.581438435 | 0.31399395 | -1.851750439 | 0.064061667 | 0.292206722 |
| VWA8 | 542.5207155 | 0.031768287 | 0.340805571 | 0.09321528 | 0.925732535 | 0.972124891 |
| VWCE | 292.4414166 | 0.714744161 | 0.397176707 | 1.799562132 | 0.071929805 | 0.308679129 |
| VWDE | 120.5109121 | -1.24869656 | 0.40501953 | -3.083052709 | 0.002048889 | 0.042697375 |
| VWF | 315.821664 | -1.324616565 | 0.509798184 | -2.598315583 | 0.009368235 | 0.109379514 |
| WAC | 4711.839523 | -0.082849434 | 0.233381993 | -0.354994972 | 0.722593358 | 0.882935039 |
| WAPL | 2350.723422 | -0.386258871 | 0.257002691 | -1.502937071 | 0.132855272 | 0.41903271 |
| WARS | 2242.586998 | -0.482106134 | 0.270063307 | -1.785159707 | 0.074235416 | 0.313236304 |
| WARS2 | 577.5904928 | 0.034880762 | 0.299355685 | 0.116519458 | 0.907240866 | 0.965296417 |
| WAS | 15.74663817 | -0.972232158 | 0.786256851 | -1.236532511 | 0.216260689 | 0.523980406 |
| WASF1 | 2181.997689 | -0.112162641 | 0.271459939 | -0.413183033 | 0.679472518 | 0.861682789 |
| WASF2 | 4408.09171 | -0.267866409 | 0.223506662 | -1.198471703 | 0.230733433 | 0.539536045 |
| WASF3 | 1797.861742 | 0.521060944 | 0.505312144 | 1.031166478 | 0.302462758 | 0.613293571 |
| WASH2P | 280.0166937 | -0.414971293 | 0.363485221 | -1.141645571 | 0.253601374 | 0.565942797 |
| WASH3P | 541.6606068 | 0.413993601 | 0.406719034 | 1.017885976 | 0.308732147 | 0.618441002 |
| WASH4P | 28.52517274 | -0.970691451 | 0.807618651 | -1.201918072 | 0.229395272 | 0.538167075 |
| WASH5P | 1215.006926 | -0.865471159 | 0.808774962 | -1.070101326 | 0.284573703 | 0.598479579 |
| WASH6P | 759.3219093 | -0.211913588 | 0.309917581 | -0.683774014 | 0.494117878 | 0.760192914 |
| WASH7P | 214.3417542 | -0.92976701 | 1.503351816 | -0.618462691 | 0.536270384 | 0.785153251 |
| WASH8P | 240.5923201 | 0.315360836 | 0.394232514 | 0.79993614 | 0.423747798 | 0.712860381 |
| WASH9P | 236.79312 | -0.190324343 | 0.419410644 | -0.453789969 | 0.649980004 | 0.847430646 |
| WASHC1 | 378.4891775 | -0.012489588 | 0.372988523 | -0.033485182 | 0.973287683 | 0.990457971 |
| WASHC2A | 1534.597746 | -0.622728068 | 0.357242092 | -1.74315424 | 0.081306676 | |

0.326151133

| | | | | | | |
|---|---|---|---|---|---|---|
| WASHC2C | 1447.872036 | -0.838127092 | 0.355490223 | -2.35766566 | 0.018390248 | 0.153593808 |
| WASHC3 | 298.8080135 | 0.118429017 | 0.453238496 | 0.261295141 | 0.793864913 | 0.914481746 |
| WASHC4 | 1771.695646 | -0.251643245 | 0.348108057 | -0.722888311 | 0.469748505 | 0.74495024 |
| WASHC5 | 1303.384395 | -0.391648819 | 0.274600776 | -1.426248043 | 0.153796756 | 0.450608457 |
| WASL | 1662.853974 | -0.784299417 | 0.347388242 | -2.25770283 | 0.023964193 | 0.177207433 |
| WBP1 | 1995.667413 | 0.393031612 | 0.396884104 | 0.990293154 | 0.322030852 | 0.631858517 |
| WBP11 | 3528.378731 | -0.084498002 | 0.252685703 | -0.334399616 | 0.738078029 | 0.891242475 |
| WBP1L | 1377.833936 | -0.79032838 | 0.277619253 | -2.846806814 | 0.004416015 | 0.069497268 |
| WBP2 | 1413.358229 | 0.059707939 | 0.246777636 | 0.241950365 | 0.808818624 | 0.919641737 |
| WBP4 | 1025.125836 | 0.498231286 | 0.398873631 | 1.24909557 | 0.211630121 | 0.518836577 |
| WDCP | 495.9935517 | 0.129069681 | 0.475434513 | 0.271477306 | 0.786023952 | 0.911122354 |
| WDFY1 | 2390.745384 | 0.207696181 | 0.250179161 | 0.830189775 | 0.406431495 | 0.701474651 |
| WDFY2 | 2515.911579 | -0.716442343 | 0.289409072 | -2.475535192 | 0.013303665 | 0.131058378 |
| WDFY3 | 1160.175099 | -0.839672366 | 0.359656474 | -2.334651052 | 0.019561657 | 0.158520715 |
| WDHD1 | 2180.889172 | 0.04363926 | 0.363481162 | 0.120059207 | 0.904436247 | 0.963993604 |
| WDPCP | 255.496859 | -0.253380136 | 0.296757575 | -0.853828703 | 0.393199908 | 0.692183315 |
| WDR1 | 8142.51912 | -0.59904325 | 0.244810303 | -2.446969112 | 0.014406318 | 0.136428375 |
| WDR11 | 2398.662357 | -0.446470157 | 0.223895413 | -1.994101397 | 0.046140975 | 0.247053322 |
| WDR12 | 1859.582525 | 0.352175179 | 0.250905945 | 1.403614324 | 0.160433727 | 0.460798954 |
| WDR13 | 734.3699718 | 0.249954587 | 0.364036207 | 0.686620127 | 0.492322142 | 0.759174768 |
| WDR17 | 125.3511554 | 0.301497432 | 0.712920985 | 0.422904414 | 0.672365002 | 0.859011927 |
| WDR18 | 801.2529999 | 0.155643078 | 0.437012867 | 0.356152164 | 0.721726614 | 0.882498649 |
| WDR19 | 1225.550088 | -0.345846613 | 0.333317734 | -1.037588394 | 0.299461723 | 0.610246278 |
| WDR20 | 756.7575255 | -0.592010185 | 0.273032184 | -2.168279856 | 0.0301374 | 0.198185614 |
| WDR24 | 289.5817328 | -0.515168362 | 0.460440427 | -1.118859969 | 0.263199883 | 0.576857184 |
| WDR25 | 155.3598017 | 0.106276691 | 0.401827897 | 0.264483109 | 0.791407679 | 0.913361935 |
| WDR26 | 2413.447537 | -0.180878898 | 0.245614806 | -0.7364332 | 0.461467106 | 0.73902094 |
| WDR27 | 1073.27667 | -0.302040141 | 0.360139606 | -0.838675159 | 0.401651622 | 0.699111606 |
| WDR3 | 1369.280744 | 0.306540213 | 0.34866772 | 0.879175775 | 0.379305977 | 0.681253679 |
| WDR31 | 82.91073638 | -0.486274145 | 0.447329178 | -1.087061093 | 0.277009811 | 0.590738803 |
| WDR33 | 2049.929383 | -0.234551223 | 0.250660835 | -0.935731435 | 0.349411481 | 0.656038342 |
| WDR34 | 957.4468686 | 0.487143979 | 0.411262739 | 1.184507939 | 0.236212056 | 0.547141583 |
| WDR35 | 704.3136948 | 0.468017827 | 0.507524588 | 0.922157937 | 0.3564462 | 0.663922075 |
| WDR36 | 849.5899874 | -0.076861739 | 0.336797597 | -0.228213441 | 0.819480312 | 0.924316427 |
| WDR37 | 548.7678804 | -0.275872446 | 0.311752858 | -0.884907511 | 0.376206517 | 0.680079109 |
| WDR4 | 462.2907873 | 0.570829213 | 0.32436455 | 1.759838467 | 0.078435199 | 0.321397152 |
| WDR41 | 1882.508476 | 0.323632065 | 0.241222159 | 1.341634893 | 0.179714407 | 0.484443366 |
| WDR43 | 1989.486618 | 0.247339693 | 0.325684208 | 0.759446382 | 0.447585577 | 0.729044534 |
| WDR44 | 434.2139867 | -0.450758736 | 0.275011067 | -1.639056711 | 0.101201449 | 0.36459919 |
| WDR45 | 445.7019082 | -0.844897998 | 0.433940207 | -1.947037828 | 0.0515302 | 0.261459752 |

| | | | | | | |
|---|---|---|---|---|---|---|
| WDR45B | 2975.933332 | 0.056352243 | 0.217217596 | 0.259427617 | 0.795305324 | 0.915124686 |
| WDR46 | 260.6103032 | 0.872776167 | 0.765081924 | 1.140761713 | 0.253969097 | 0.566445924 |
| WDR47 | 233.3803771 | -0.210972909 | 0.531197765 | -0.397164527 | 0.691246139 | 0.868004248 |
| WDR48 | 1550.563765 | -0.325592013 | 0.363957543 | -0.894587897 | 0.371007424 | 0.676265004 |
| WDR5 | 1616.091116 | -0.121320395 | 0.232712178 | -0.521332386 | 0.602135244 | 0.825021776 |
| WDR53 | 354.8858385 | 0.150135268 | 0.343659551 | 0.436872095 | 0.66220411 | 0.853821447 |
| WDR54 | 964.5894862 | 0.785481277 | 0.285865551 | 2.747729749 | 0.006000945 | 0.08383088 |
| WDR55 | 985.0115878 | -0.534671119 | 0.233631672 | -2.28852156 | 0.022107166 | 0.169006551 |
| WDR59 | 1601.822798 | -0.219093934 | 0.302505524 | -0.724264244 | 0.468903524 | 0.744337782 |
| WDR5B | 280.188724 | -0.013232101 | 0.320532041 | -0.041281679 | 0.967071339 | 0.988749915 |
| WDR6 | 5427.766147 | -0.135576362 | 0.292803698 | -0.463028173 | 0.643344173 | 0.845415888 |
| WDR60 | 825.4646403 | -0.071705525 | 0.314303696 | -0.228140889 | 0.819536714 | 0.924316427 |
| WDR61 | 1226.860067 | 0.505580581 | 0.24901297 | 2.030338343 | 0.042322159 | 0.236751283 |
| WDR62 | 527.6335408 | 0.193791025 | 0.60806334 | 0.318702038 | 0.749952469 | 0.895752516 |
| WDR63 | 32.28369343 | -0.202939093 | 0.627983729 | -0.323159794 | 0.746574228 | 0.894109554 |
| WDR66 | 43.53829764 | -0.05760656 | 0.517499144 | -0.111317208 | 0.91136481 | 0.966866396 |
| WDR7 | 429.1546948 | -0.08305799 | 0.368183751 | -0.225588417 | 0.82152155 | 0.925538648 |
| WDR70 | 1113.39318 | -0.298308294 | 0.242671441 | -1.229268237 | 0.218971252 | 0.527527654 |
| WDR72 | 620.7917793 | -0.302738389 | 0.321559372 | -0.941469646 | 0.346464237 | 0.653107473 |
| WDR73 | 1115.124493 | -0.248203274 | 0.240887078 | -1.030371892 | 0.302835463 | 0.613538084 |
| WDR74 | 916.8407519 | 0.56082786 | 0.325079249 | 1.725203504 | 0.084490804 | 0.332764796 |
| WDR75 | 2408.894042 | 0.243584909 | 0.308216072 | 0.790305668 | 0.429349277 | 0.71613003 |
| WDR76 | 414.7242633 | 0.119428868 | 0.371682631 | 0.321319474 | 0.7479683 | 0.894202292 |
| WDR77 | 1559.483162 | 0.169484198 | 0.267371572 | 0.633890122 | 0.526152533 | 0.778525486 |
| WDR78 | 69.70130289 | -0.449829479 | 0.53474767 | -0.841199513 | 0.400236174 | 0.697919713 |
| WDR81 | 26.22326843 | -1.18327977 | 1.114217928 | -1.061982347 | 0.288243699 | 0.601673803 |
| WDR82 | 2722.816393 | -0.205341195 | 0.344197855 | -0.59657895 | 0.550788528 | 0.794461475 |
| WDR83 | 251.4355043 | 0.422673893 | 0.349562931 | 1.209149643 | 0.22660536 | 0.535404129 |
| WDR83OS | 1594.080948 | 0.729075173 | 0.54476938 | 1.338318928 | 0.180792494 | 0.485502272 |
| WDR86 | 236.757837 | -1.605305286 | 0.713018392 | -2.251421987 | 0.024358823 | 0.178492903 |
| WDR89 | 774.7295386 | 0.244233403 | 0.331008559 | 0.737846187 | 0.460607923 | 0.738395128 |
| WDR90 | 409.1383657 | -0.117762407 | 0.478639066 | -0.246035928 | 0.805654421 | 0.918930547 |
| WDR91 | 582.668326 | -0.656165174 | 0.26418547 | -2.483729232 | 0.013001462 | 0.128720283 |
| WDR92 | 564.2483473 | 0.094912445 | 0.307742001 | 0.308415636 | 0.757766086 | 0.899024084 |
| WDR93 | 11.60552541 | -0.94542688 | 1.057838527 | -0.893734588 | 0.371463915 | 0.67682226 |
| WDR97 | 20.98205931 | -0.726782101 | 0.838460832 | -0.86680507 | 0.386048824 | 0.686679504 |
| WDSUB1 | 327.9580666 | -0.426697743 | 0.411925908 | -1.035860417 | 0.300267266 | 0.610801113 |
| WDTC1 | 1446.947431 | -0.623140493 | 0.230817681 | -2.699708661 | 0.006940022 | 0.091474435 |

| | | | | | | |
|---|---|---|---|---|---|---|
| WDYHV1 | 1042.665309 | 0.805241001 | 0.390961051 | 2.05964507 | 0.039432484 | 0.228786879 |
| WEE1 | 2377.897663 | 0.256695724 | 0.391795914 | 0.655177133 | 0.512353715 | 0.771687791 |
| WFDC1 | 13.1429448 | -0.884564763 | 0.920123942 | -0.961353925 | 0.336374243 | 0.64425371 |
| WFDC2 | 503.7684844 | 0.431011623 | 0.791001974 | 0.544893233 | 0.585826951 | 0.815639728 |
| WFIKKN1 | 164.0803953 | 0.394363903 | 0.655087469 | 0.602001904 | 0.54717287 | 0.792404079 |
| WFS1 | 621.1042169 | -0.148654488 | 0.39244124 | -0.378794257 | 0.704840653 | 0.873723655 |
| WHAMM | 178.5973686 | -0.229593239 | 0.336302583 | -0.682698411 | 0.494797434 | 0.760514211 |
| WHAMMP3 | 12.55502994 | 0.116149417 | 1.002974842 | 0.115804916 | 0.907807154 | 0.965463886 |
| WHRN | 489.149126 | -0.573297571 | 0.385421725 | -1.487455255 | 0.136894612 | 0.424728769 |
| WIPF1 | 440.9948283 | -0.413926446 | 0.375982984 | -1.100918031 | 0.270932334 | 0.583394591 |
| WIPF2 | 1376.564866 | -0.09579743 | 0.271529132 | -0.352807188 | 0.724232995 | 0.883667763 |
| WIPF3 | 211.6659091 | 0.099891593 | 0.387722253 | 0.257636987 | 0.796687082 | 0.915710299 |
| WIPI1 | 425.1841604 | -0.401611869 | 0.401286402 | -1.000811058 | 0.316918163 | 0.627198876 |
| WIPI2 | 2195.444262 | -0.273205765 | 0.278392301 | -0.981369689 | 0.326410468 | 0.636173332 |
| WIZ | 939.9808155 | -0.278287723 | 0.407774908 | -0.68245426 | 0.494951756 | 0.760514211 |
| WLS | 11392.46808 | -0.09090325 | 0.238334072 | -0.381411055 | 0.702898259 | 0.872703476 |
| WNK1 | 4690.613702 | -0.516832086 | 0.397987577 | -1.298613614 | 0.194076563 | 0.500132242 |
| WNK2 | 303.464835 | 0.218331829 | 0.489548744 | 0.445985883 | 0.655607443 | 0.849596013 |
| WNK3 | 556.8596265 | 0.326435614 | 0.328858636 | 0.992632025 | 0.320889312 | 0.630501469 |
| WNK4 | 20.93996553 | 0.143124371 | 0.741574262 | 0.193000726 | 0.8469584 | 0.938145276 |
| WNT1 | 31.79919865 | -0.079351526 | 1.052443208 | -0.075397442 | 0.939898494 | 0.978003717 |
| WNT10B | 141.1088961 | 0.752927569 | 0.604618922 | 1.245292765 | 0.213024154 | 0.520189768 |
| WNT11 | 44.22880547 | -2.702357368 | 1.226647941 | -2.203042354 | 0.027591764 | 0.190183846 |
| WNT2B | 34.91426185 | 0.356237868 | 0.788735477 | 0.45165696 | 0.651516129 | 0.847876685 |
| WNT3 | 11.93709763 | 1.468839866 | 1.794861992 | 0.818358109 | 0.413152734 | 0.705607651 |
| WNT4 | 110.3752104 | -0.697888571 | 0.897174221 | -0.77787408 | 0.43664325 | 0.720784523 |
| WNT5A | 180.0966675 | 1.097266809 | 0.487809373 | 2.249376232 | 0.024488569 | 0.178720266 |
| WNT5B | 75.81533883 | -0.938174873 | 0.661774273 | -1.417665981 | 0.156288307 | 0.454843067 |
| WNT6 | 199.3643918 | -1.136726196 | 0.925198067 | -1.228630103 | 0.219210524 | 0.52778634 |
| WNT7B | 30.17311012 | -0.365972736 | 0.692029903 | -0.528839483 | 0.596916806 | 0.821812117 |
| WNT8A | 135.0553364 | 2.007748666 | 0.762425301 | 2.633370986 | 0.008454195 | 0.103848246 |
| WNT8B | 21.89900397 | -0.500084181 | 0.845706077 | -0.591321494 | 0.554305031 | 0.796360966 |
| WNT9A | 71.21806145 | -0.94421429 | 0.444389228 | -2.124746123 | 0.033607803 | 0.208960645 |
| WRAP53 | 880.7959807 | 0.657414725 | 0.260735326 | 2.521387245 | 0.011689313 | 0.122556026 |
| WRAP73 | 497.9909068 | 0.19172379 | 0.256982923 | 0.746056539 | 0.455633262 | 0.735306344 |
| WRB | 1237.586923 | 0.81027335 | 0.309085308 | 2.621520104 | 0.00875386 | 0.105800902 |
| WRB-SH3BGR | 21.9829069 | 0.265054672 | 1.040321959 | 0.254781387 | 0.798891952 | 0.916056735 |
| WRN | 1026.957924 | 0.095237837 | 0.392528128 | 0.242626782 | 0.808294532 | 0.91963689 |

| | | | | | | |
|---|---|---|---|---|---|---|
| WRNIP1 | 888.5019569 | 0.299908452 | 0.245828956 | 1.219988309 | 0.222469307 | 0.531341602 |
| WSB1 | 7538.063747 | -0.209996113 | 0.323319196 | -0.649500914 | 0.516014656 | 0.773896629 |
| WSB2 | 719.9877815 | 0.220119339 | 0.27589001 | 0.797851792 | 0.424956502 | 0.71350603 |
| WSCD1 | 701.3703419 | 0.945984513 | 0.540660974 | 1.749681516 | 0.080173285 | 0.324202401 |
| WSCD2 | 118.8407344 | -1.10373292 | 0.800756182 | -1.378363283 | 0.168091154 | 0.470384431 |
| WTAP | 3393.830168 | 0.391613776 | 0.271784451 | 1.440898385 | 0.149613392 | 0.444386222 |
| WTAPP1 | 46.50018341 | -0.422779433 | 0.663674715 | -0.637028086 | 0.524106549 | 0.777662198 |
| WTIP | 508.0341154 | -0.094605104 | 0.312010247 | -0.303211529 | 0.761728664 | 0.900715944 |
| WWC1 | 946.8215737 | -0.821676637 | 0.396896851 | -2.070252349 | 0.038428719 | 0.225620008 |
| WWC2 | 1499.122678 | -0.341535757 | 0.386793427 | -0.882992659 | 0.377240238 | 0.680674553 |
| WWC3 | 1408.365764 | -1.05219158 | 0.452445794 | -2.325563844 | 0.02004183 | 0.160028741 |
| WWOX | 480.3512697 | -0.147697219 | 0.315537223 | -0.46808176 | 0.639726123 | 0.843538624 |
| WWP1 | 1487.819801 | -0.23897973 | 0.289623478 | -0.825139355 | 0.409292479 | 0.703508181 |
| WWP2 | 1108.833761 | -0.536826466 | 0.377359315 | -1.422587028 | 0.154855907 | 0.452575411 |
| WWTR1 | 836.6255716 | -1.032057119 | 0.604309604 | -1.707828425 | 0.087668183 | 0.34024836 |
| XAB2 | 1422.675926 | -0.371695644 | 0.406802156 | -0.913701263 | 0.36087384 | 0.667986461 |
| XAF1 | 55.60474218 | -0.352105942 | 0.560687172 | -0.627990009 | 0.530010485 | 0.780583348 |
| XAGE2 | 10.10229142 | -1.540839947 | 1.086209208 | -1.418548044 | 0.156030822 | 0.454433405 |
| XBP1 | 1852.816345 | 0.208722111 | 0.475932022 | 0.438554461 | 0.6609844 | 0.852848642 |
| XCL1 | 22.38495128 | 0.051038888 | 0.85484521 | 0.059705415 | 0.952390259 | 0.982774543 |
| XDH | 174.8437027 | -0.877074231 | 0.313373195 | -2.79881702 | 0.005129019 | 0.076062906 |
| XG | 10.85300404 | 1.655747899 | 1.37529303 | 1.203923718 | 0.228619063 | 0.53738497 |
| XIAP | 1007.894356 | -0.129093377 | 0.303787316 | -0.424946569 | 0.670875626 | 0.858238816 |
| XK | 20.1149942 | 0.826846389 | 0.746266658 | 1.107977129 | 0.267871693 | 0.581031837 |
| XKR4 | 28.89619586 | -1.495900446 | 0.692961878 | -2.158705252 | 0.030873041 | 0.200588825 |
| XKR5 | 117.19534 | 0.742875254 | 0.53017849 | 1.401179541 | 0.161160391 | 0.46167973 |
| XKR6 | 92.19564709 | 0.072267758 | 0.424652513 | 0.170180926 | 0.864867852 | 0.945248223 |
| XKR7 | 35.93026009 | -1.695486459 | 0.765630318 | -2.214497543 | 0.026794572 | 0.186982725 |
| XKR8 | 470.1778674 | 0.290823515 | 0.251379354 | 1.156910901 | 0.247308765 | 0.559988249 |
| XKR9 | 17.21311735 | 1.00404066 | 0.767556157 | 1.308100588 | 0.190839194 | 0.496058205 |
| XKRX | 29.00468545 | -0.475533203 | 0.703042344 | -0.676393403 | 0.498790905 | 0.76281557 |
| XPA | 175.2809512 | 0.639851053 | 0.341030214 | 1.876229806 | 0.060623731 | 0.284105089 |
| XPC | 1853.599663 | -0.660180055 | 0.312505452 | -2.11253932 | 0.034640217 | 0.21269873 |
| XPNPEP1 | 1886.357412 | 0.330652014 | 0.367851926 | 0.89887259 | 0.36872053 | 0.67460916 |
| XPNPEP2 | 14.00727922 | 0.806377924 | 0.891734996 | 0.904279778 | 0.365847067 | 0.672352011 |
| XPNPEP3 | 727.5181376 | 0.009053903 | 0.364971498 | 0.024807151 | 0.980208787 | 0.993196129 |
| XPO1 | 10860.76749 | 0.203120323 | 0.320475097 | 0.633810005 | 0.526204823 | 0.778525486 |
| XPO4 | 1164.494939 | -0.142526249 | 0.344092025 | -0.414209684 | 0.67872055 | 0.8612303 |
| XPO5 | 2634.931895 | -0.095338544 | 0.426284298 | -0.223650143 | 0.823029543 | |

0.926014081

| | | | | | | |
|---|---|---|---|---|---|---|
| XPO6 | 2253.94389 | -0.148728716 | 0.421248375 | -0.353066562 | 0.72403854 | 0.883667763 |
| XPO7 | 3281.148714 | -0.266251285 | 0.34875722 | -0.76342874 | 0.445207746 | 0.727906747 |
| XPOT | 4456.898014 | 0.070331392 | 0.260358988 | 0.270132377 | 0.787058414 | 0.911694526 |
| XPR1 | 2143.254867 | -0.044481529 | 0.260584901 | -0.170698798 | 0.864460608 | 0.94517566 |
| XRCC1 | 1510.243782 | -0.434752056 | 0.346397701 | -1.25506623 | 0.209454718 | 0.517277801 |
| XRCC2 | 612.9205422 | 0.217470434 | 0.435507221 | 0.499349777 | 0.617532993 | 0.831835425 |
| XRCC3 | 529.5754238 | 0.315775616 | 0.382380536 | 0.825815089 | 0.408908993 | 0.703351285 |
| XRCC4 | 476.7905278 | 0.538093963 | 0.27598632 | 1.949712447 | 0.051210402 | 0.260426481 |
| XRCC5 | 20242.54857 | 0.115495245 | 0.243574644 | 0.474167768 | 0.635380276 | 0.840590898 |
| XRCC6 | 11739.20245 | 0.064665553 | 0.218916771 | 0.295388758 | 0.76769692 | 0.90389833 |
| XRN1 | 661.9764866 | -0.732599573 | 0.408249447 | -1.794490056 | 0.072734984 | 0.310433977 |
| XRN2 | 5235.427252 | -0.096361926 | 0.251711458 | -0.382826934 | 0.70184809 | 0.872479299 |
| XRRA1 | 119.7330355 | -0.349210072 | 0.446934906 | -0.78134437 | 0.43459998 | 0.719554351 |
| XXYLT1 | 850.7377496 | 0.246066118 | 0.260148882 | 0.945866521 | 0.344216671 | 0.65118919 |
| XYLB | 82.22046482 | -0.757012473 | 0.658032868 | -1.150417417 | 0.24997199 | 0.562742997 |
| XYLT1 | 216.0755031 | -1.395489386 | 0.481856139 | -2.896070573 | 0.003778674 | 0.063484236 |
| XYLT2 | 642.3355516 | 0.074717762 | 0.470546662 | 0.158789273 | 0.873834901 | 0.949740014 |
| YAE1 | 499.0704559 | 0.497332244 | 0.308311319 | 1.61308461 | 0.106726127 | 0.374599763 |
| YAF2 | 311.6925401 | 0.231530381 | 0.38395946 | 0.603007363 | 0.546503791 | 0.792185077 |
| YAP1 | 9890.002792 | -0.996057342 | 0.413092532 | -2.411220889 | 0.015899216 | 0.142656219 |
| YARS | 2336.58221 | -0.121237728 | 0.294918039 | -0.411089564 | 0.681006862 | 0.862490397 |
| YARS2 | 856.3280213 | 0.189163388 | 0.273540428 | 0.691537224 | 0.489227996 | 0.757568942 |
| YBEY | 544.5770914 | 0.462727529 | 0.370796464 | 1.247928645 | 0.212057189 | 0.519309059 |
| YBX1 | 36237.37658 | 0.622414359 | 0.24391509 | 2.551766515 | 0.010717832 | 0.117687513 |
| YBX2 | 609.0858558 | -0.251133405 | 0.394172852 | -0.637114919 | 0.524049991 | 0.777662198 |
| YBX3 | 4842.567144 | 0.537829177 | 0.35349755 | 1.52145093 | 0.128146716 | 0.412317905 |
| YDJC | 211.161759 | 0.503027757 | 0.453599827 | 1.108968139 | 0.267443927 | 0.580705856 |
| YEATS2 | 2811.550744 | -0.360928747 | 0.469278722 | -0.769113812 | 0.441825749 | 0.725269254 |
| YEATS4 | 1128.297293 | 0.736446884 | 0.336231268 | 2.190298628 | 0.028502586 | 0.192869196 |
| YES1 | 2986.110082 | -0.03784128 | 0.236240823 | -0.160180952 | 0.872738534 | 0.949330774 |
| YIF1A | 692.7551215 | 0.394031761 | 0.378029001 | 1.042332096 | 0.297257737 | 0.608678246 |
| YIF1B | 1050.478831 | 0.364473954 | 0.377754075 | 0.964844534 | 0.334622686 | 0.64263567 |
| YIPF1 | 867.4693459 | 0.025658351 | 0.365813949 | 0.070140438 | 0.944081881 | 0.979059828 |
| YIPF2 | 841.1239345 | 0.018490624 | 0.438829767 | 0.042136212 | 0.966390113 | 0.988700292 |
| YIPF3 | 2256.180176 | -0.141403263 | 0.31192466 | -0.453325051 | 0.650314697 | 0.847442386 |
| YIPF4 | 881.6359065 | 0.513600108 | 0.305740009 | 1.679859005 | 0.092984752 | 0.349608839 |
| YIPF5 | 1090.781826 | 0.003864903 | 0.267093323 | 0.014470235 | 0.988454826 | 0.995803391 |
| YIPF6 | 1543.800318 | 0.073048686 | 0.268593595 | 0.271967342 | 0.78564713 | 0.910966831 |
| YJEFN3 | 220.5255421 | 0.185174848 | 0.576110344 | 0.321422536 | 0.747890207 | 0.894202292 |
| YJU2 | 336.4328386 | -0.282818168 | 0.343887733 | -0.822414236 | 0.41084118 | 0.704426187 |
| YKT6 | 2769.104323 | 0.106594367 | 0.224067409 | 0.475724547 | 0.634270631 | 0.840101252 |
| YLPM1 | 3111.600199 | -0.624661121 | 0.332255765 | -1.880061048 | 0.060099758 | 0.28351164 |
| YME1L1 | 3908.936757 | -0.204465112 | 0.230884993 | -0.885571253 | | |

| | | | | | | |
|---|---|---|---|---|---|---|
| | | | | | 0.375848608 | 0.679780547 |
| YOD1 | 752.3705854 | 0.153683775 | 0.315038424 | 0.487825496 | 0.625673451 | 0.83542439 |
| YPEL1 | 562.9319353 | 0.587615681 | 0.248693731 | 2.362808584 | 0.018137035 | 0.152020417 |
| YPEL2 | 496.8862676 | -1.230781522 | 0.365229686 | -3.369883582 | 0.000751999 | 0.022817168 |
| YPEL3 | 556.0887834 | -0.419561497 | 0.598582178 | -0.700925475 | 0.483349525 | 0.754102802 |
| YPEL5 | 998.0793605 | 0.090064189 | 0.312871447 | 0.287863242 | 0.773451425 | 0.906158747 |
| YRDC | 434.706511 | 0.559790974 | 0.339024965 | 1.65117921 | 0.098701987 | 0.360796043 |
| YTHDC1 | 1500.003874 | -0.145112907 | 0.253696101 | -0.571995021 | 0.567325348 | 0.804984367 |
| YTHDC2 | 867.9597681 | -0.254640432 | 0.305579437 | -0.833303557 | 0.404673551 | 0.70065889 |
| YTHDF1 | 1952.29642 | -0.273185131 | 0.369578496 | -0.739180266 | 0.459797542 | 0.73770692 |
| YTHDF2 | 4313.540637 | -0.081273267 | 0.33180857 | -0.24494023 | 0.806502712 | 0.919129555 |
| YTHDF3 | 2834.359363 | -0.347445724 | 0.218045337 | -1.593456337 | 0.111057856 | 0.383475916 |
| YWHAB | 8426.158432 | 0.316248827 | 0.403733038 | 0.783311736 | 0.433444073 | 0.71890019 |
| YWHAE | 20850.24182 | 0.10854276 | 0.232495777 | 0.466859061 | 0.640600717 | 0.843724522 |
| YWHAG | 4868.180912 | 0.076913116 | 0.361679088 | 0.21265569 | 0.831595531 | 0.930997148 |
| YWHAH | 3735.417633 | 0.365422842 | 0.301385458 | 1.212476688 | 0.225329961 | 0.53459368 |
| YWHAQ | 13429.33582 | 0.371348661 | 0.249813621 | 1.486502858 | 0.137146161 | 0.425402661 |
| YWHAZ | 26377.18689 | -0.027382229 | 0.270510683 | -0.101224206 | 0.919372482 | 0.969677217 |
| YY1 | 2666.386115 | 0.197595975 | 0.238478446 | 0.828569533 | 0.407348034 | 0.701878198 |
| YY1AP1 | 1425.676984 | -0.53626677 | 0.250488045 | -2.140887684 | 0.032283097 | 0.204752031 |
| YY2 | 29.34329123 | -0.725926775 | 0.579527659 | -1.252617996 | 0.210344762 | 0.51816316 |
| Z83820.1 | 13.45586153 | -4.867977473 | 1.590999124 | -3.0596984 | 0.0022156 | 0.044579397 |
| Z83844.3 | 312.9116414 | -1.899602228 | 2.192534949 | -0.866395415 | 0.386273357 | 0.686679504 |
| Z84492.2 | 14.42511278 | 0.708501281 | 0.815001836 | 0.869324768 | 0.38466952 | 0.685277276 |
| ZACN | 12.25373931 | -0.407211839 | 0.832535229 | -0.489122652 | 0.624754865 | 0.835033444 |
| ZADH2 | 432.153767 | 0.394485223 | 0.31087557 | 1.268948932 | 0.204459279 | 0.510835602 |
| ZBBX | 120.2796297 | 0.317720783 | 0.385754012 | 0.82363572 | 0.410146574 | 0.704109399 |
| ZBED1 | 968.4198789 | -0.285294891 | 0.338111293 | -0.843789889 | 0.398786826 | 0.697300137 |
| ZBED3 | 135.1422062 | -0.762402644 | 0.42053141 | -1.812950536 | 0.069839451 | 0.303603614 |
| ZBED4 | 1501.022796 | -0.027060999 | 0.30780573 | -0.087915839 | 0.929943568 | 0.97351462 |
| ZBED5 | 1526.22939 | -0.086721205 | 0.262416722 | -0.330471337 | 0.741043847 | 0.891943909 |
| ZBED6 | 343.738513 | -1.977741074 | 2.176351181 | -0.908741701 | 0.363486487 | 0.670353372 |
| ZBED8 | 238.9045771 | 1.152027864 | 0.402040284 | 2.865453815 | 0.00416412 | 0.066568879 |
| ZBED9 | 264.8940236 | -0.289429961 | 0.28408076 | -1.018829862 | 0.308283746 | 0.617978528 |
| ZBTB1 | 910.517051 | -0.242356136 | 0.243941505 | -0.993501027 | 0.32046585 | 0.63024811 |
| ZBTB10 | 1615.986146 | -0.032532871 | 0.332347006 | -0.097888263 | 0.92202102 | 0.970640219 |
| ZBTB11 | 540.1845438 | 0.080991267 | 0.327739799 | 0.247120634 | 0.804814865 | 0.918655204 |
| ZBTB12 | 160.314858 | 0.31947869 | 0.642338854 | 0.497367842 | 0.618929678 | 0.832427495 |
| ZBTB14 | 344.7230242 | -0.13216127 | 0.297086615 | -0.444857706 | 0.656422589 | 0.849955395 |

ZBTB16 97.15370038 1.391488407 0.931176523 1.494333644 0.135088436
0.421953193
ZBTB17 561.8855727 -0.296474727 0.491866031 -0.602755036
0.546671663 0.792249935
ZBTB18 798.3751707 0.337824608 0.293867496 1.149581403 0.250316322
0.563016889
ZBTB2 420.4230788 -0.420957012 0.254606863 -1.653360821 0.098257451
0.359965304
ZBTB20 117.5289224 0.414197664 0.508325639 0.814827411 0.415171108
0.707398889
ZBTB21 420.965846 0.267262912 0.411470868 0.649530581 0.515995487
0.773896629
ZBTB22 354.8267696 0.106526274 0.294995953 0.361110969 0.718016494
0.880671367
ZBTB24 530.0350378 -0.106872278 0.345353512 -0.309457627
0.75697344 0.898563152
ZBTB25 321.6746431 -0.269534349 0.359369199 -0.750020729
0.45324222 0.733497409
ZBTB26 406.9600247 0.392546108 0.331739631 1.183295788 0.236691941
0.547336452
ZBTB3 238.2672852 -0.077419298 0.333853542 -0.231895991 0.816618799
0.923554512
ZBTB33 1057.454471 0.138812102 0.285856958 0.485599871 0.627250888
0.836041947
ZBTB34 699.3450546 -0.243031698 0.278864283 -0.871505291
0.383478318 0.684521658
ZBTB37 486.7541681 -0.589456247 0.471091487 -1.251256417
0.210840941 0.518294063
ZBTB38 568.6504746 -0.936865933 0.290887731 -3.220713119
0.001278721 0.03168167
ZBTB39 1212.199681 -0.31416158 0.357266752 -0.879347376 0.379212956
0.681253679
ZBTB4 369.711561 -0.589205407 0.289287308 -2.036748208 0.041675281
0.234603223
ZBTB40 931.9476494 -0.7858816 0.49141248 -1.59923004 0.109769498
0.380904175
ZBTB41 444.5128846 0.208144779 0.326761504 0.636992965 0.524129426
0.777662198
ZBTB43 437.0666606 -0.340584168 0.333468849 -1.021337281
0.307094669 0.617205881
ZBTB44 1181.466831 0.083191469 0.245701755 0.338587201 0.734920726
0.889483064
ZBTB45 226.8645758 -0.289772093 0.431867233 -0.670974945
0.502236491 0.764661936
ZBTB46 463.675831 -0.610507892 0.42023974 -1.452760968
0.146290133 0.43878363
ZBTB47 122.822391 0.22298073 0.436518392 0.510816346 0.609479662
0.828395867
ZBTB48 470.9207964 -0.353532192 0.406998184 -0.86863334 0.385047713
0.685870477
ZBTB49 166.2577111 -0.451583651 0.419215081 -1.077212323
0.281385421 0.595211573
ZBTB5 735.3128785 0.137713826 0.345528337 0.398560151 0.690217329 0.8679061
ZBTB6 506.907785 0.170029888 0.281357008 0.604320782 0.545630392 0.791502307
ZBTB7A 551.6486565 0.307669204 0.523666928 0.587528423 0.556848867
0.797880068
ZBTB7B 1012.399199 -0.151102386 0.303716614 -0.497511097
0.618828679 0.832365448
ZBTB7C 115.0118594 -0.80612756 0.439968337 -1.832239941 0.066915681
0.297817842
ZBTB8A 610.775075 0.280084517 0.389480596 0.71912316 0.472065039
0.746399031
ZBTB8B 205.1137413 0.386969125 0.345351574 1.120507777 0.26249744
0.576074285

ZBTB8OS 593.8617369 0.760679285 0.347002816 2.192141529 0.028369289 0.192544525
ZBTB9 104.6858101 7.202127126 2.198163192 3.27642968 0.001051285 0.027896179
ZC2HC1A 611.1482828 -0.641268337 0.369763133 -1.734267912 0.082870558 0.329548948
ZC2HC1C 21.67708596 0.022014321 0.774396513 0.028427712 0.977321022 0.99216657
ZC3H10 331.2996519 0.482171062 0.32721756 1.473548856 0.1406031 0.430405495
ZC3H11A 3522.356145 -0.134661554 0.270331107 -0.498135622 0.618388455 0.832277132
ZC3H12A 94.42697639 -0.616714763 0.485919202 -1.269171419 0.204379932 0.510835602
ZC3H12B 215.4473293 -0.348742907 0.354497268 -0.983767543 0.325229821 0.63517786
ZC3H12C 326.2258875 -0.022793024 0.305578206 -0.074589823 0.940541072 0.97807441
ZC3H13 3708.096236 -0.09743902 0.287648809 -0.338742998 0.734803346 0.889483064
ZC3H14 2449.329604 -0.136097516 0.294417054 -0.462260981 0.643894177 0.845466381
ZC3H15 2919.736503 0.563624414 0.284620671 1.980265213 0.047673735 0.251480612
ZC3H18 1614.319482 -0.282472792 0.440307275 -0.641535601 0.521174758 0.776211464
ZC3H3 298.062276 -0.673266738 0.588808284 -1.143439649 0.252856098 0.565626639
ZC3H6 479.4368676 0.582315449 0.390103875 1.492718956 0.135510769 0.423011087
ZC3H7A 1501.128931 -0.072370034 0.255491011 -0.283258633 0.776978585 0.907794497
ZC3H7B 2703.072124 -0.444936383 0.410015459 -1.085169774 0.277846469 0.591968315
ZC3H8 474.5074899 0.105214497 0.305910763 0.343938525 0.730892519 0.886911603
ZC3HAV1 948.6799505 -0.277440108 0.316051739 -0.877831297 0.380035277 0.68184242
ZC3HAV1L 40.65206912 -1.171463051 0.71435264 -1.639894621 0.10102708 0.36405744
ZC3HC1 755.0986477 0.255600262 0.250914745 1.018673739 0.308357884 0.617978528
ZC4H2 746.552598 -0.096552504 0.30346706 -0.318164695 0.750360012 0.896002498
ZCCHC10 746.4473004 0.459855527 0.241598978 1.903383574 0.056990513 0.276407631
ZCCHC14 1031.90474 -0.13366996 0.396506369 -0.337119326 0.736026949 0.890167797
ZCCHC17 1394.888508 0.52655265 0.271049175 1.9426462 0.052058921 0.262949685
ZCCHC18 92.92031893 0.752674503 0.487410886 1.544229979 0.122532629 0.403241325
ZCCHC2 444.0255226 -0.632116118 0.302681469 -2.08838724 0.036762918 0.219399876
ZCCHC24 458.1527954 0.968934511 0.415088227 2.334285699 0.019580767 0.158520715
ZCCHC3 1816.575603 0.572696631 0.256380408 2.233776894 0.025497757 0.182111117
ZCCHC4 406.1066483 -0.020294573 0.277152822 -0.073225207 0.941626908 0.978517574
ZCCHC7 821.3185766 0.095443721 0.30746629 0.310420114 0.7562415 0.898288428
ZCCHC8 985.3137723 0.006236368 0.356468869 0.017494846 0.986041844 0.994894974
ZCCHC9 1056.031772 0.390899113 0.332824832 1.174489028 0.240199204 0.551796203
ZCRB1 1657.531758 1.393445164 0.442911463 3.146103185 0.001654616 0.037682927

| | | | | | | |
|---|---|---|---|---|---|---|
| ZCWPW1 | 61.21996706 | -0.614858274 | 0.529902929 | -1.160322467 | 0.245917541 | 0.557670405 |
| ZCWPW2 | 42.99141174 | -0.017624369 | 0.565095064 | -0.031188326 | 0.97511935 | 0.99132451 |
| ZDBF2 | 745.0695628 | -0.135801254 | 0.370097951 | -0.366933276 | 0.713668789 | 0.878107802 |
| ZDHHC1 | 73.70067222 | 0.019140086 | 0.421320673 | 0.045428785 | 0.963765537 | 0.988130579 |
| ZDHHC11 | 271.788442 | -0.434111812 | 0.41890981 | -1.036289437 | 0.300067132 | 0.610639633 |
| ZDHHC11B | 23.5279452 | -0.439661862 | 0.843876982 | -0.521002316 | 0.602365158 | 0.825021776 |
| ZDHHC12 | 551.1955663 | 0.144345773 | 0.484682726 | 0.297814972 | 0.765844387 | 0.903118942 |
| ZDHHC13 | 817.4099461 | 0.006318182 | 0.255013189 | 0.024775903 | 0.980233712 | 0.993196129 |
| ZDHHC14 | 125.2275413 | -0.084681352 | 0.393003835 | -0.215472076 | 0.829399281 | 0.929737887 |
| ZDHHC15 | 199.635615 | -0.459303169 | 0.304252562 | -1.509611506 | 0.131142584 | 0.416813082 |
| ZDHHC16 | 1048.207688 | -0.061950288 | 0.235251644 | -0.263336259 | 0.792291415 | 0.913570955 |
| ZDHHC17 | 1362.286689 | 0.29591793 | 0.302391528 | 0.978592002 | 0.327781613 | 0.637698749 |
| ZDHHC18 | 1941.421068 | -0.098657223 | 0.235038925 | -0.419748444 | 0.674669231 | 0.859745438 |
| ZDHHC2 | 726.712044 | -0.197953109 | 0.285816551 | -0.692587986 | 0.488568152 | 0.757183311 |
| ZDHHC20 | 2638.394174 | -0.394536522 | 0.244273868 | -1.615140111 | 0.106280356 | 0.374140185 |
| ZDHHC21 | 707.0708341 | -0.336196042 | 0.320480771 | -1.049036551 | 0.294161296 | 0.606454309 |
| ZDHHC22 | 34.59674846 | -0.813664625 | 0.625902677 | -1.299985853 | 0.193605818 | 0.499607734 |
| ZDHHC23 | 90.74366271 | -1.202355332 | 0.497323885 | -2.417650484 | 0.015621073 | 0.141376414 |
| ZDHHC24 | 288.3720422 | 0.111458997 | 0.456446706 | 0.244188413 | 0.807084901 | 0.919137032 |
| ZDHHC3 | 740.7438924 | -0.406116155 | 0.292945208 | -1.386321207 | 0.165648844 | 0.467560858 |
| ZDHHC4 | 631.7709811 | -0.054244693 | 0.283150736 | -0.191575321 | 0.848074875 | 0.938207222 |
| ZDHHC5 | 2205.441918 | -0.683406108 | 0.237738578 | -2.874611743 | 0.004045246 | 0.065655774 |
| ZDHHC6 | 879.8213158 | -0.187430048 | 0.311926044 | -0.600879766 | 0.547920071 | 0.792941769 |
| ZDHHC7 | 1637.298161 | -0.356263271 | 0.258495168 | -1.378220237 | 0.168135301 | 0.470384431 |
| ZDHHC8 | 407.1194038 | -0.212942318 | 0.550771546 | -0.386625489 | 0.699033485 | 0.871039767 |
| ZDHHC8P1 | 84.19976006 | -1.455422525 | 0.630888333 | -2.306941574 | 0.021058077 | 0.16470868 |
| ZDHHC9 | 1124.904544 | -0.359034504 | 0.252833005 | -1.420046024 | 0.155594282 | 0.453684578 |
| ZEB1 | 190.9320296 | 1.601675397 | 0.760980582 | 2.104751995 | 0.035312902 | 0.21486064 |
| ZEB2 | 998.2782641 | 1.837951483 | 0.783649241 | 2.345375183 | 0.019007942 | 0.156382583 |
| ZER1 | 882.5309234 | -1.002610016 | 0.394319896 | -2.542631062 | 0.011002133 | 0.119239998 |
| ZFAND1 | 1115.662001 | 0.71570605 | 0.250517597 | 2.856909287 | 0.004277881 | 0.067815809 |
| ZFAND2A | 309.2882482 | 0.348085231 | 0.447192607 | 0.778378769 | 0.436345751 | 0.720784523 |
| ZFAND2B | 479.5267725 | 0.833311823 | 0.365719325 | 2.278555618 | 0.022693495 | 0.171650943 |

| | | | | | | |
|---|---|---|---|---|---|---|
| ZFAND3 | 2468.603492 | -0.39686118 | 0.251376093 | -1.578754669 | 0.11439234 | 0.38927175 |
| ZFAND4 | 203.4065923 | 0.274951325 | 0.388399167 | 0.707909151 | 0.479001677 | 0.751414997 |
| ZFAND5 | 4196.258944 | -0.068115981 | 0.250871109 | -0.271517837 | 0.785992783 | 0.911122354 |
| ZFAND6 | 2501.296428 | 0.378655498 | 0.289734841 | 1.306903572 | 0.191245463 | 0.496727429 |
| ZFAT | 219.0341493 | -1.049231865 | 0.477081396 | -2.199272231 | 0.027858571 | 0.190853383 |
| ZFC3H1 | 1211.415862 | 0.050288824 | 0.31999677 | 0.157154162 | 0.875123353 | 0.949740014 |
| ZFHX2 | 437.1516002 | 0.172912836 | 0.51080457 | 0.338510745 | 0.734978331 | 0.889483064 |
| ZFHX3 | 962.1208576 | -1.744943466 | 0.566848978 | -3.078321621 | 0.002081701 | 0.043144109 |
| ZFHX4 | 465.2280484 | -0.308173193 | 0.658896931 | -0.467710773 | 0.639991436 | 0.843538624 |
| ZFP1 | 195.2490646 | 0.084536486 | 0.381723549 | 0.221459971 | 0.824734301 | 0.92681808 |
| ZFP14 | 435.6982882 | 0.248213574 | 0.288625776 | 0.859984087 | 0.389797815 | 0.689715727 |
| ZFP2 | 147.0708283 | -0.085148563 | 0.328861574 | -0.258919162 | 0.795697613 | 0.915261082 |
| ZFP28 | 237.8193142 | -0.011667521 | 0.379289134 | -0.030761548 | 0.975459706 | 0.991471205 |
| ZFP3 | 39.2843744 | -2.590606375 | 0.642594389 | -4.031479919 | 5.54E-05 | 0.004042687 |
| ZFP30 | 322.4745722 | -0.024907816 | 0.308861346 | -0.080644005 | 0.935725069 | 0.97559515 |
| ZFP36 | 276.4951634 | -1.200788915 | 0.688812281 | -1.743274543 | 0.08128567 | 0.326151133 |
| ZFP36L1 | 3674.195022 | 0.127022619 | 0.50379956 | 0.252129278 | 0.800941141 | 0.916680548 |
| ZFP36L2 | 8165.857251 | -0.623522793 | 0.675581782 | -0.922941989 | 0.356037437 | 0.663487122 |
| ZFP37 | 303.4498859 | 0.612642866 | 0.302481744 | 2.025387907 | 0.042827548 | 0.238139903 |
| ZFP41 | 245.667596 | -0.728741839 | 0.383398747 | -1.900741317 | 0.057335904 | 0.277284985 |
| ZFP42 | 11.72152747 | -0.310449468 | 0.87697803 | -0.35399914 | 0.723339528 | 0.883273431 |
| ZFP62 | 935.7982828 | -0.287744518 | 0.301626917 | -0.953974931 | 0.340096332 | 0.64680976 |
| ZFP64 | 1017.029872 | -0.114636928 | 0.295089151 | -0.388482353 | 0.697659111 | 0.87033849 |
| ZFP69 | 145.0972894 | 0.471067977 | 0.314378579 | 1.498409903 | 0.134026785 | 0.420619758 |
| ZFP69B | 138.4984165 | 0.27618173 | 0.443833269 | 0.622264596 | 0.533767897 | 0.783156734 |
| ZFP82 | 445.1471234 | 0.632588985 | 0.323209374 | 1.957211132 | 0.050322651 | 0.257746432 |
| ZFP90 | 1105.33251 | -0.281913081 | 0.379367959 | -0.743112524 | 0.457413558 | 0.73623473 |
| ZFP91 | 1930.310229 | 0.096898778 | 0.263061487 | 0.368350302 | 0.71261205 | 0.877590503 |
| ZFPL1 | 652.2948918 | -0.031263567 | 0.344983224 | -0.090623441 | 0.927791806 | 0.972633911 |
| ZFPM1 | 46.68475352 | -0.111488544 | 0.560209183 | -0.199012346 | 0.842253087 | 0.93548734 |
| ZFPM2 | 12.48238507 | 0.308777454 | 0.95923448 | 0.32189987 | 0.747528552 | 0.894202292 |
| ZFR | 3090.740431 | -0.160058563 | 0.323942717 | -0.494095266 | 0.621238901 | 0.833349572 |
| ZFR2 | 143.2897933 | 0.276320381 | 0.434171336 | 0.636431652 | 0.524495116 | 0.777662198 |
| ZFX | 398.0055208 | -0.295111864 | 0.291627538 | -1.011947864 | 0.311562982 | 0.621528206 |
| ZFY | 437.6300782 | -0.492857808 | 0.280747573 | -1.755519389 | 0.079170507 | 0.322875205 |
| ZFYVE1 | 755.0043692 | -0.841625111 | 0.346077404 | -2.431898473 | 0.015019917 | 0.139113039 |
| ZFYVE16 | 1386.727538 | -0.16430358 | 0.400620589 | -0.410122655 | 0.681715975 | |

0.862620404
ZFYVE19 658.4571904 -0.067032253 0.401377285 -0.167005597
0.867365645 0.946404214
ZFYVE21 1344.067922 -0.459549611 0.403516034 -1.138863321
0.254760166 0.567256913
ZFYVE26 913.8067605 -0.739164159 0.472762521 -1.56349991 0.117935057
0.395839103
ZFYVE27 579.655777 -0.230212668 0.373178496 -0.616896929
0.537302713 0.785675856
ZFYVE9 861.2042672 -0.538229142 0.499546078 -1.077436429
0.281285337 0.595171527
ZGLP1 55.07226052 -0.14624773 0.516125556 -0.283356885 0.776903275
0.907794497
ZGPAT 468.4440352 -0.203096315 0.377853564 -0.537500064 0.590922241
0.818805811
ZGRF1 695.5977181 0.222309591 0.434619842 0.511503548 0.608998505 0.828100414
ZHX1 148.9703491 -0.123086907 0.495273867 -0.248522918 0.80372984
0.917969611
ZHX1-C8orf76 51.57297806 0.577931284 0.69531002 0.831185037 0.405869107
0.701371854
ZHX2 1890.585266 -1.527827438 0.335778232 -4.55010865 5.36E-06
0.000683565
ZHX3 275.4599672 0.050288178 0.480605423 0.104635061 0.916665393 0.969211928
ZIC1 13.69090898 -0.52551372 0.974110538 -0.539480582 0.589555292
0.81828099
ZIC2 5080.801354 1.605869259 0.528735685 3.037187207 0.002387971 0.046988207
ZIC3 2152.305757 1.627368814 0.526563608 3.090545546 0.001997892 0.042097241
ZIC5 1482.18408 1.452543933 0.518348736 2.802252293 0.005074717 0.075897737
ZIK1 747.1622701 -0.399463241 0.294306221 -1.35730478 0.174684389
0.47862011
ZKSCAN1 2692.461587 -0.236754745 0.348307287 -0.67972952 0.496675741
0.761325016
ZKSCAN2 586.1654334 -0.333042556 0.318326483 -1.046229498
0.295455087 0.60699582
ZKSCAN3 320.2166288 -0.285858905 0.293351558 -0.974458452
0.329828959 0.638570079
ZKSCAN4 328.1970737 -0.177966212 0.300107921 -0.593007378
0.553176215 0.79574357
ZKSCAN5 820.2243317 -0.312869795 0.31203943 -1.002661089
0.31602441 0.625961211
ZKSCAN7 151.7461897 -0.446674171 0.329004434 -1.357653956
0.174573513 0.478578744
ZKSCAN8 1427.199319 -0.495863792 0.429224158 -1.155256018
0.247985601 0.560705687
ZMAT1 93.2345631 -0.068083283 0.409477713 -0.166268591 0.867945582
0.9466285
ZMAT2 2781.111033 0.036299373 0.314930727 0.115261452 0.908237892 0.965583536
ZMAT3 1315.503774 0.483217055 0.330698267 1.461202259 0.143959943 0.434995877
ZMAT4 91.11320981 0.028668318 0.526163303 0.054485591 0.956548288 0.984539826
ZMAT5 243.6852161 0.729540719 0.33187796 2.19821985 0.027933443 0.191063238
ZMIZ1 2798.875831 -0.744267823 0.396748164 -1.875920018 0.060666263
0.284105089
ZMIZ2 1601.913103 -0.597359315 0.394571141 -1.513945781 0.130039592
0.415175781
ZMPSTE24 1223.122839 -0.133439781 0.247785857 -0.538528643
0.590212136 0.818321084
ZMYM1 857.4761748 0.190869766 0.304557489 0.626711779 0.530848181 0.781218098
ZMYM2 5446.387746 0.180673908 0.311527157 0.579961985 0.561940254 0.801706045
ZMYM3 4090.688057 -0.357027949 0.341984499 -1.043988688 0.296490623
0.607790454
ZMYM4 2918.280493 -0.253784481 0.415949592 -0.610132779 0.541773854
0.788644324
ZMYM5 521.29695 0.243680075 0.282879103 0.861428336 0.389002175 0.688870316
ZMYM6 371.6210832 -0.17920631 0.335296852 -0.534470601 0.593015984

0.819859199
ZMYND11 1168.031177 -0.15217373 0.39883784 -0.38154286 0.702800475
0.872703476
ZMYND12 32.82101508 0.009965401 0.595372258 0.016738101 0.986645551
0.995107011
ZMYND19 687.478437 0.635692639 0.321761587 1.975663552 0.048192896
0.253165032
ZMYND8 4036.558691 -0.080321122 0.28308831 -0.283731682
0.776616013 0.907755593
ZNF10 1161.103538 -0.451958127 0.661494585 -0.683237834 0.49445657
0.760405536
ZNF100 473.341908 -0.09502528 0.280668154 -0.338568088 0.734935127
0.889483064
ZNF101 475.2820164 0.415069138 0.294083694 1.411398004 0.158127291
0.457552763
ZNF106 3245.530017 0.465432347 0.407458521 1.142281541 0.253337011
0.565885941
ZNF107 837.0497483 -0.3747377 0.331911107 -1.129030309 0.258885048
0.572028119
ZNF112 212.9295363 -0.212787476 0.450587217 -0.472244813
0.636752056 0.841268221
ZNF114 172.5399042 -0.157410391 0.35448464 -0.444054193
0.657003403 0.85024732
ZNF117 1832.998345 -1.049437706 0.293160675 -3.579735602
0.000343942 0.013767664
ZNF12 940.2567921 -0.151548815 0.279360851 -0.542484084 0.587485069
0.816787521
ZNF121 3328.263474 0.547308483 0.411979855 1.328483604 0.184018396
0.488493979
ZNF124 259.9653977 -0.023350157 0.301757738 -0.077380476
0.938320872 0.977342821
ZNF131 1126.697841 0.213244786 0.243929791 0.874205587 0.382006304
0.683341308
ZNF132 124.6869441 -0.284404674 0.478627626 -0.594208646
0.55237257 0.795662343
ZNF133 531.6104641 -0.313024845 0.247975283 -1.262322765
0.206832668 0.513642439
ZNF134 1233.586467 -0.224711633 0.274081222 -0.819872415
0.412288843 0.70515068
ZNF135 352.013849 -0.675017443 0.352898073 -1.912783022
0.055775836 0.273312403
ZNF136 348.6807734 -0.367353027 0.27690666 -1.326631243
0.184630695 0.489177835
ZNF137P 89.49190671 -1.920770322 0.739909748 -2.595952177
0.009432919 0.109913071
ZNF138 799.6809723 0.109566355 0.275811694 0.397250579 0.691182687
0.868004248
ZNF14 275.2989947 -0.12324074 0.293162589 -0.420383586 0.674205257
0.859672768
ZNF140 491.1039116 0.060769238 0.345967492 0.175650138 0.86056881
0.943645503
ZNF141 770.4918876 0.082266313 0.397556804 0.206929707 0.836064755
0.93270414
ZNF142 500.6760334 -0.718926337 0.488635278 -1.471294378
0.141211523 0.430963592
ZNF143 644.1473056 0.302346894 0.287483061 1.051703336 0.29293568
0.606044893
ZNF146 4813.261714 0.203177249 0.256267726 0.792831982 0.42787572
0.715216235
ZNF148 765.2843079 -0.075139733 0.254983934 -0.294684185
0.768235146 0.904321493
ZNF154 578.60358 -0.529442069 0.329576312 -1.60643241 0.108178935
0.377196648
ZNF155 266.4203777 -0.370294553 0.326538351 -1.134000192

0.256794468 0.569149579
ZNF157 19.50854065 -0.262582614 0.891822424 -0.294433742
0.768426486 0.904332533
ZNF16 396.490522 -0.34085131 0.330983969 -1.029812141 0.3030982
0.61357768
ZNF160 1343.01863 -0.761364142 0.279356398 -2.725422246
0.00642193 0.08706622
ZNF165 171.6499537 -0.338780165 0.697973417 -0.485376888
0.627409024 0.836054176
ZNF169 126.5428559 0.297168867 0.412572554 0.720282684 0.471350964
0.746147696
ZNF17 161.3974503 -0.220853503 0.346705327 -0.637006373 0.524120693
0.777662198
ZNF174 283.92679 0.028800759 0.321004612 0.0897207 0.928509166
0.972633911
ZNF175 339.7650595 -0.549228248 0.341857939 -1.606597901
0.108142603 0.377180798
ZNF177 157.2080988 0.18310567 0.414712658 0.441524189 0.658833554
0.851400081
ZNF18 281.8964652 0.419367849 0.291129159 1.440487277 0.149729587 0.444386222
ZNF180 328.7859887 0.140676384 0.293595645 0.479150103 0.631831848
0.838768903
ZNF181 360.4388773 0.403729124 0.321199799 1.256940774 0.208775083
0.515935296
ZNF182 353.3024937 -0.313519323 0.409290863 -0.766006162
0.443672636 0.726848446
ZNF184 319.2004763 0.248905601 0.39449374 0.630949431 0.52807359
0.779319497
ZNF185 378.4829859 -0.742530415 0.372165878 -1.995159845
0.046025449 0.246950947
ZNF189 605.5610747 -0.289484795 0.297793327 -0.97209967 0.331000966
0.639084337
ZNF19 144.5620753 -0.436667472 0.33846716 -1.290132469 0.197004668
0.50258876
ZNF192P1 37.46081743 -0.166868501 0.746569386 -0.223513721
0.823135705 0.926014081
ZNF195 1833.36915 0.192748834 0.29600634 0.651164546 0.514940274
0.773145336
ZNF197 456.4111915 -0.195420307 0.346140848 -0.564568752
0.572367125 0.807456036
ZNF2 96.36728064 -0.167173199 0.394847532 -0.423386715 0.672013136
0.858752635
ZNF20 101.1514094 0.159638389 0.462973851 0.34481081 0.730236606 0.886742369
ZNF200 334.3973308 -0.048572195 0.29884044 -0.162535548
0.870884137 0.94840164
ZNF202 461.9541331 -0.108441573 0.385515001 -0.281290151
0.778487864 0.908160087
ZNF204P 517.2652878 -0.229933428 0.28408932 -0.809370193
0.418302242 0.709602481
ZNF205 252.5939257 -0.674196738 0.59759937 -1.128175115
0.259245971 0.572529055
ZNF207 7525.235164 -0.1629032 0.248168807 -0.656420931 0.511553327
0.771142242
ZNF208 13.20274223 -0.762487098 1.044295541 -0.730144933
0.465301599 0.742280815
ZNF211 558.8199612 -0.407578193 0.263743731 -1.545356894
0.122259961 0.402868346
ZNF212 510.9236742 -0.111604398 0.268131847 -0.416229549
0.677242039 0.860358531
ZNF213 228.1606907 -0.469588882 0.446052615 -1.052765674
0.292448401 0.605587347
ZNF214 160.250328 -0.225572424 0.429472798 -0.525230991
0.599422623 0.823816451
ZNF215 357.7117161 0.038133554 0.311548506 0.122400054 0.902582195

| | | | | | | |
|---|---|---|---|---|---|---|
| | | | | | | 0.962877047 |
| ZNF217 | 2765.710719 | -0.336620994 | 0.255935349 | -1.315257919 | 0.188423236 | 0.493707929 |
| ZNF219 | 1032.053333 | 0.651120812 | 0.394933359 | 1.648685273 | 0.099212129 | 0.361452832 |
| ZNF22 | 2056.195533 | 0.438055369 | 0.274780817 | 1.594199239 | 0.110891417 | 0.383132604 |
| ZNF222 | 148.8253569 | 0.206850461 | 0.362117643 | 0.571224477 | 0.567847488 | 0.805273345 |
| ZNF224 | 504.5761828 | -0.739218859 | 0.295235811 | -2.503825185 | 0.012285872 | 0.12619429 |
| ZNF225 | 200.1426512 | -0.452269316 | 0.344737684 | -1.311923054 | 0.189546098 | 0.495025625 |
| ZNF226 | 757.9296759 | -0.09142924 | 0.246450257 | -0.370984557 | 0.710649034 | 0.876838602 |
| ZNF227 | 655.342981 | -0.119483299 | 0.266313872 | -0.448655936 | 0.653679876 | 0.84883826 |
| ZNF229 | 355.1536119 | -0.460582223 | 0.361812729 | -1.272985127 | 0.203023319 | 0.509506436 |
| ZNF23 | 474.6058615 | -0.072517864 | 0.270371443 | -0.268215694 | 0.788533295 | 0.912309638 |
| ZNF230 | 139.1717356 | -0.416999919 | 0.357040781 | -1.167933582 | 0.242833557 | 0.553989157 |
| ZNF232 | 595.047239 | 0.070962065 | 0.30389118 | 0.233511433 | 0.815364292 | 0.922920552 |
| ZNF233 | 57.52765465 | -0.925714799 | 0.509931156 | -1.815372113 | 0.069466744 | 0.30336211 |
| ZNF234 | 244.9693945 | -0.455102727 | 0.400619124 | -1.135998507 | 0.255957189 | 0.56812385 |
| ZNF235 | 148.6987224 | -0.409887329 | 0.399608137 | -1.025723178 | 0.305022086 | 0.615135352 |
| ZNF236 | 429.1019078 | -0.308155329 | 0.345270977 | -0.892502844 | 0.372123469 | 0.677322753 |
| ZNF239 | 117.4753425 | 0.276628068 | 0.426090879 | 0.649223163 | 0.516194142 | 0.773896629 |
| ZNF24 | 2812.27481 | 0.264930345 | 0.266325887 | 0.994760021 | 0.319852995 | 0.629465081 |
| ZNF248 | 298.4605138 | -2.007897128 | 0.357227256 | -5.620783676 | 1.90E-08 | 1.25E-05 |
| ZNF25 | 75.93849258 | 0.809717497 | 0.57914371 | 1.398128795 | 0.162074394 | 0.462717357 |
| ZNF250 | 250.0993993 | 0.146072601 | 0.296324956 | 0.492947348 | 0.622049791 | 0.833640266 |
| ZNF251 | 476.2119015 | -0.249695658 | 0.274833682 | -0.908533683 | 0.363596326 | 0.670353372 |
| ZNF252P | 621.6687292 | -0.544434134 | 0.284449579 | -1.913991702 | 0.055621216 | 0.272730921 |
| ZNF253 | 852.1997764 | -0.154255119 | 0.288311939 | -0.535028553 | 0.592630113 | 0.819859199 |
| ZNF254 | 1498.685735 | 0.14749628 | 0.261944589 | 0.563081988 | 0.573379056 | 0.808057005 |
| ZNF256 | 298.5610144 | -0.13452066 | 0.372756085 | -0.360881191 | 0.718188265 | 0.88073997 |
| ZNF26 | 416.5121273 | -0.044859593 | 0.261735568 | -0.171392806 | 0.863914911 | 0.945086034 |
| ZNF260 | 1459.66217 | -0.112171697 | 0.293761687 | -0.381845902 | 0.702575669 | 0.872703476 |
| ZNF263 | 951.4505478 | -0.173380092 | 0.233880239 | -0.74131997 | 0.458499456 | 0.736808886 |
| ZNF264 | 491.7209269 | -0.099289607 | 0.415037303 | -0.239230561 | 0.810926807 | 0.920897407 |
| ZNF266 | 881.4639385 | -0.396977603 | 0.239383315 | -1.658334468 | 0.097249976 | 0.358274005 |
| ZNF267 | 467.9139875 | -0.141821807 | 0.282774494 | -0.501536772 | 0.615993405 | 0.831265761 |
| ZNF268 | 900.0980199 | -0.236813939 | 0.242383277 | -0.977022598 | 0.328557965 | 0.637981939 |

| | | | | | | |
|---|---|---|---|---|---|---|
| ZNF271P | 877.3347858 | 0.269840909 | 0.262376935 | 1.028447523 | 0.303739364 | 0.614002998 |
| ZNF273 | 501.2375953 | 0.305140077 | 0.283942798 | 1.074653343 | 0.28252996 | 0.596463956 |
| ZNF274 | 890.6954741 | -0.181462071 | 0.250104949 | -0.725543703 | 0.468118544 | 0.744224533 |
| ZNF275 | 377.9007224 | -0.35572771 | 0.443260842 | -0.802524555 | 0.42224959 | 0.712133049 |
| ZNF276 | 371.4972522 | 0.032352852 | 0.392391484 | 0.082450444 | 0.934288524 | 0.97507153 |
| ZNF277 | 960.3106835 | 0.159898058 | 0.308424634 | 0.518434783 | 0.604154952 | 0.82576988 |
| ZNF28 | 551.8782153 | -0.738399268 | 0.389387389 | -1.896310177 | 0.057919041 | 0.278947 |
| ZNF280B | 149.9144514 | 0.589561847 | 0.526312537 | 1.120174431 | 0.262639437 | 0.57612824 |
| ZNF280C | 355.611796 | 0.110903609 | 0.402464813 | 0.275561007 | 0.782885262 | 0.909909835 |
| ZNF280D | 1013.047275 | 0.587240129 | 0.298688921 | 1.966059298 | 0.049291761 | 0.25592242 |
| ZNF281 | 1526.432239 | 0.242205362 | 0.228281385 | 1.060994799 | 0.288692265 | 0.601894782 |
| ZNF282 | 708.339364 | -0.717126091 | 0.467388629 | -1.53432507 | 0.124949701 | 0.4067741 |
| ZNF283 | 20.57612664 | -0.570477807 | 0.697415682 | -0.817988212 | 0.413363918 | 0.705652445 |
| ZNF284 | 99.57094187 | 0.489950512 | 0.36774554 | 1.33230851 | 0.182758832 | 0.486683209 |
| ZNF285 | 176.0794383 | 0.063090919 | 0.383439978 | 0.164539231 | 0.869306665 | 0.947227711 |
| ZNF286A | 2458.280671 | 0.105949305 | 0.318666027 | 0.332477567 | 0.739528676 | 0.89168179 |
| ZNF286B | 259.9181742 | 0.752542042 | 0.382990781 | 1.964909027 | 0.049424768 | 0.256262184 |
| ZNF287 | 220.6789472 | 0.618815381 | 0.300086521 | 2.06212321 | 0.039196007 | 0.228182127 |
| ZNF292 | 1434.409833 | -0.19704682 | 0.289504087 | -0.680635713 | 0.496102024 | 0.761056737 |
| ZNF296 | 26.04338208 | -0.533173881 | 0.730163092 | -0.730212041 | 0.465260584 | 0.742280815 |
| ZNF3 | 1831.800115 | -0.102031361 | 0.263235355 | -0.387605079 | 0.69830831 | 0.870504139 |
| ZNF30 | 212.0511433 | 0.143944762 | 0.31966563 | 0.4502979 | 0.652495653 | 0.848578973 |
| ZNF300 | 1137.284526 | -0.019218707 | 0.307572605 | -0.062485108 | 0.950176521 | 0.981636196 |
| ZNF302 | 1451.017432 | 0.317269411 | 0.229350538 | 1.383338419 | 0.166561123 | 0.468464738 |
| ZNF304 | 415.3867305 | -0.605061835 | 0.350587911 | -1.72584911 | 0.084374562 | 0.332479605 |
| ZNF316 | 692.1106969 | -0.138158124 | 0.303084156 | -0.455840801 | 0.648504462 | 0.846827301 |
| ZNF317 | 981.4764444 | -0.760924372 | 0.344472325 | -2.208956472 | 0.027177669 | 0.188528766 |
| ZNF318 | 1318.037859 | -0.105362376 | 0.386449365 | -0.272642124 | 0.785128328 | 0.910931322 |
| ZNF319 | 214.6922845 | 0.178010903 | 0.297020378 | 0.599322189 | 0.548958054 | 0.793468239 |
| ZNF32 | 740.0249679 | 0.588839559 | 0.281060934 | 2.095060141 | 0.036165647 | 0.217725803 |
| ZNF320 | 93.82211654 | -0.576342216 | 0.521775618 | -1.104578664 | 0.2693422 | 0.582188661 |
| ZNF321P | 104.4483663 | 0.056012192 | 0.410127445 | 0.136572651 | 0.891368596 | 0.957017706 |
| ZNF322 | 1357.554833 | 0.305548842 | 0.337203784 | 0.906125189 | 0.364869594 | 0.671614642 |

| | | | | | | |
|---|---|---|---|---|---|---|
| ZNF324 | 176.6187963 | -0.267032402 | 0.330618389 | -0.807675588 | 0.41927736 | 0.709924862 |
| ZNF324B | 46.77332034 | -0.593840983 | 0.550821704 | -1.078100188 | 0.280989048 | 0.595042552 |
| ZNF326 | 1388.677398 | -0.158323404 | 0.299429924 | -0.528749439 | 0.596979276 | 0.821812117 |
| ZNF329 | 568.2681384 | -0.146471446 | 0.29051728 | -0.504174641 | 0.614138659 | 0.830316752 |
| ZNF330 | 1087.719212 | 0.424776591 | 0.291605184 | 1.456683948 | 0.145203643 | 0.436733637 |
| ZNF331 | 966.5956551 | 0.015595722 | 0.384125961 | 0.040600541 | 0.967614353 | 0.989014473 |
| ZNF333 | 340.9602443 | 0.370379031 | 0.334304092 | 1.107910551 | 0.267900448 | 0.581031837 |
| ZNF334 | 852.8327004 | -0.495396416 | 0.332981976 | -1.487757452 | 0.13681487 | 0.424728769 |
| ZNF335 | 596.0423652 | -0.826157741 | 0.44758395 | -1.845816279 | 0.064918886 | 0.293855407 |
| ZNF337 | 431.2315531 | -0.1910881 | 0.310566058 | -0.61528971 | 0.538363412 | 0.786560766 |
| ZNF33A | 1571.211617 | -0.398691941 | 0.269048543 | -1.481858762 | 0.13837788 | 0.426606546 |
| ZNF33B | 1128.697707 | -0.455192282 | 0.270426708 | -1.683237153 | 0.092329191 | 0.348873265 |
| ZNF34 | 103.9097478 | -0.167438381 | 0.358872422 | -0.466567981 | 0.640809 | 0.843724522 |
| ZNF341 | 154.5708392 | -0.079440148 | 0.386296946 | -0.205645292 | 0.837067996 | 0.93321271 |
| ZNF343 | 197.3019961 | 0.25676489 | 0.538888483 | 0.476471288 | 0.633738658 | 0.839689885 |
| ZNF345 | 168.9377679 | -0.562920988 | 0.332387119 | -1.693570408 | 0.090346945 | 0.344491755 |
| ZNF346 | 753.922468 | -0.227518972 | 0.263739782 | -0.862664594 | 0.388321904 | 0.688469278 |
| ZNF347 | 438.3939185 | -0.157561972 | 0.32918827 | -0.478637868 | 0.632196274 | 0.838768903 |
| ZNF35 | 151.6251141 | -0.380289452 | 0.346062217 | -1.098904858 | 0.271809568 | 0.584165314 |
| ZNF350 | 327.9672847 | -0.910590579 | 0.308128723 | -2.955227835 | 0.003124382 | 0.055756493 |
| ZNF354A | 781.6437865 | 0.071912561 | 0.236850854 | 0.303619599 | 0.761417718 | 0.90056031 |
| ZNF354B | 410.8458605 | -0.245795048 | 0.324882123 | -0.75656686 | 0.449309402 | 0.730519016 |
| ZNF354C | 692.6991142 | -0.392151837 | 0.290603354 | -1.349440161 | 0.177195629 | 0.481566699 |
| ZNF358 | 864.6942809 | 0.079917442 | 0.4429472 | 0.180422049 | 0.856821246 | 0.942343063 |
| ZNF362 | 804.7607729 | -0.210043981 | 0.282659864 | -0.743098004 | 0.457422348 | 0.73623473 |
| ZNF367 | 346.6717695 | 0.442448485 | 0.40725248 | 1.08642306 | 0.277291864 | 0.591007989 |
| ZNF37A | 548.5607799 | -0.261416484 | 0.368516869 | -0.70937454 | 0.478092083 | 0.750609023 |
| ZNF37BP | 335.7781859 | -0.858081933 | 0.372376882 | -2.304337286 | 0.021203717 | 0.165290607 |
| ZNF382 | 226.9088429 | 0.032179459 | 0.375656705 | 0.085661877 | 0.931735208 | 0.974341807 |
| ZNF383 | 213.0385176 | -0.087804794 | 0.295490018 | -0.29714978 | 0.766352161 | 0.903436836 |
| ZNF384 | 1785.156805 | -0.536609155 | 0.285570349 | -1.879078683 | 0.06023375 | 0.283904551 |
| ZNF385A | 1095.766268 | -0.598953407 | 0.33001823 | -1.814910063 | 0.069537732 | 0.303425952 |

| | | | | | | |
|---|---|---|---|---|---|---|
| ZNF385B | 26.83964832 | -1.581248592 | 0.795869357 | -1.986819291 | 0.046942426 | 0.249564641 |
| ZNF385C | 307.537514 | -0.42670954 | 0.349054419 | -1.22247282 | 0.221528876 | 0.530599081 |
| ZNF385D | 18.28724115 | 0.669784976 | 0.681835929 | 0.982325729 | 0.325939403 | 0.635827259 |
| ZNF391 | 354.2657834 | -0.267782537 | 0.293137366 | -0.913505296 | 0.360976849 | 0.667991431 |
| ZNF394 | 450.0129615 | -0.171564168 | 0.318920929 | -0.537952052 | 0.590610152 | 0.818577253 |
| ZNF395 | 914.6710043 | -0.414275997 | 0.283364654 | -1.461988963 | 0.143744234 | 0.434930949 |
| ZNF396 | 81.02221538 | -0.397619476 | 0.424645261 | -0.936356795 | 0.349089515 | 0.655726918 |
| ZNF397 | 2164.791845 | 0.06606041 | 0.299901186 | 0.22027392 | 0.82565783 | 0.927178721 |
| ZNF398 | 441.6913475 | -0.075852557 | 0.326756452 | -0.232137903 | 0.816430907 | 0.923506806 |
| ZNF404 | 136.98004 | 0.055073667 | 0.520619972 | 0.105784776 | 0.915753116 | 0.969158739 |
| ZNF407 | 269.6632166 | -0.234990868 | 0.480454252 | -0.489101444 | 0.624769879 | 0.835033444 |
| ZNF408 | 320.3416463 | -0.578702782 | 0.373225238 | -1.550545684 | 0.121010597 | 0.401219509 |
| ZNF41 | 131.7732515 | -0.11143387 | 0.582863451 | -0.191183493 | 0.848381836 | 0.938447017 |
| ZNF410 | 918.725116 | -0.194109214 | 0.241263503 | -0.804552747 | 0.421077819 | 0.711536154 |
| ZNF414 | 115.390797 | -0.072217644 | 0.498431866 | -0.144889702 | 0.884797957 | 0.95440284 |
| ZNF415 | 328.5434048 | -0.02175203 | 0.440656839 | -0.049362742 | 0.960630219 | 0.986479967 |
| ZNF416 | 225.279754 | -0.228699571 | 0.304165741 | -0.751891287 | 0.452116438 | 0.732888936 |
| ZNF417 | 606.1607115 | -0.534192478 | 0.379789198 | -1.406549952 | 0.159560883 | 0.459433126 |
| ZNF418 | 330.1351399 | -0.408519249 | 0.335515163 | -1.217588038 | 0.223380562 | 0.532595711 |
| ZNF419 | 464.9144822 | -0.065252178 | 0.30891707 | -0.211228789 | 0.832708747 | 0.931432899 |
| ZNF420 | 334.8512226 | 0.261795761 | 0.306043587 | 0.855419856 | 0.392318755 | 0.691514794 |
| ZNF423 | 2011.493579 | 0.171604737 | 0.394793994 | 0.434669068 | 0.663802654 | 0.854667747 |
| ZNF425 | 123.8051163 | -0.462870075 | 0.381068071 | -1.21466507 | 0.22449386 | 0.533874112 |
| ZNF426 | 977.2058607 | -0.25285974 | 0.276445743 | -0.914681257 | 0.360358987 | 0.667522123 |
| ZNF428 | 430.0356153 | 0.69917853 | 0.40029239 | 1.746669552 | 0.080694675 | 0.325104967 |
| ZNF429 | 352.9135001 | -0.273755511 | 0.296563975 | -0.923090915 | 0.355959829 | 0.663487122 |
| ZNF43 | 1020.427694 | -0.513218245 | 0.705258695 | -0.727702116 | 0.466795955 | 0.743189974 |
| ZNF430 | 418.3350625 | -0.03749458 | 0.366787706 | -0.102224201 | 0.918578719 | 0.969677217 |
| ZNF431 | 1596.230861 | -0.2848525 | 0.317327343 | -0.897661376 | 0.369366107 | 0.675154642 |
| ZNF432 | 522.3212065 | -0.06375871 | 0.312777972 | -0.20384655 | 0.838473415 | 0.933879959 |
| ZNF433 | 440.2739845 | -0.695924859 | 0.444089002 | -1.567084202 | 0.117095011 | 0.394766315 |
| ZNF436 | 179.2510872 | -0.645965744 | 0.434230524 | -1.487610171 | 0.13685373 | 0.424728769 |

| | | | | | | |
|---|---|---|---|---|---|---|
| ZNF438 | 115.9364289 | -0.190645777 | 0.449186479 | -0.424424568 | 0.671256206 | 0.858479663 |
| ZNF439 | 132.3340205 | 0.086762779 | 0.356462205 | 0.243399658 | 0.807695809 | 0.919453224 |
| ZNF44 | 278.4866802 | -1.068144307 | 0.429339015 | -2.487880832 | 0.012850678 | 0.128086828 |
| ZNF440 | 447.2161784 | -0.492870823 | 0.347788073 | -1.417158496 | 0.156436593 | 0.455090998 |
| ZNF441 | 106.7065022 | 0.033291946 | 0.444911004 | 0.074828326 | 0.940351304 | 0.97807441 |
| ZNF442 | 77.43287996 | 0.103028565 | 0.386178809 | 0.266789794 | 0.789631013 | 0.91286231 |
| ZNF443 | 221.5481254 | -0.544025186 | 0.311775173 | -1.744927864 | 0.080997426 | 0.325513101 |
| ZNF444 | 1103.336162 | 0.440082768 | 0.407966838 | 1.078721914 | 0.280711714 | 0.594538239 |
| ZNF445 | 643.2019651 | -0.350803199 | 0.426118123 | -0.823253412 | 0.410363901 | 0.704150362 |
| ZNF446 | 258.3551955 | -0.280547304 | 0.450354376 | -0.62294788 | 0.533318772 | 0.783143103 |
| ZNF449 | 291.588624 | -0.486231243 | 0.44559312 | -1.091200067 | 0.275184859 | 0.588252711 |
| ZNF45 | 328.5787938 | -0.653126957 | 0.419823443 | -1.555718167 | 0.119775125 | 0.399101344 |
| ZNF451 | 1750.931973 | 0.078920155 | 0.307340616 | 0.256784006 | 0.797345517 | 0.915848641 |
| ZNF454 | 137.4402952 | -0.315212131 | 0.385595468 | -0.817468455 | 0.413660769 | 0.705778726 |
| ZNF460 | 56.45978742 | -0.429731074 | 0.501008265 | -0.857732505 | 0.391040187 | 0.690530747 |
| ZNF461 | 163.2641271 | -0.203206822 | 0.312733856 | -0.649775577 | 0.515837197 | 0.773832307 |
| ZNF462 | 2376.266333 | -0.540752499 | 0.298069002 | -1.81418563 | 0.069649152 | 0.303425952 |
| ZNF467 | 90.79048314 | -0.471819396 | 0.55570851 | -0.849041157 | 0.395858391 | 0.694445843 |
| ZNF468 | 1553.39458 | -0.392860828 | 0.272139428 | -1.443601283 | 0.148851164 | 0.443484094 |
| ZNF469 | 13.19414323 | 0.666562496 | 0.967184586 | 0.689178163 | 0.490711157 | 0.758593654 |
| ZNF470 | 315.1056634 | -0.210409322 | 0.289557332 | -0.726658587 | 0.467435129 | 0.743885276 |
| ZNF471 | 146.2530152 | -0.781454488 | 0.357058408 | -2.188590074 | 0.028626648 | 0.193363702 |
| ZNF473 | 359.9841754 | -0.100098134 | 0.392430271 | -0.255072409 | 0.798667174 | 0.916056735 |
| ZNF48 | 1369.47637 | 0.43312318 | 0.36939022 | 1.17253559 | 0.240982092 | 0.552437022 |
| ZNF480 | 972.6866306 | -0.244036187 | 0.282799264 | -0.862930773 | 0.38817553 | 0.688370656 |
| ZNF483 | 65.8145436 | 0.408552701 | 0.60877069 | 0.67111099 | 0.502149826 | 0.764661936 |
| ZNF484 | 291.7012077 | 0.147879111 | 0.388218018 | 0.380917691 | 0.703264325 | 0.872844636 |
| ZNF485 | 136.0046864 | 0.48597797 | 0.393224547 | 1.235879025 | 0.216503535 | 0.524190093 |
| ZNF486 | 939.8508947 | -0.41126802 | 0.336706799 | -1.221442577 | 0.221918495 | 0.530861792 |
| ZNF487 | 66.12945785 | -0.127135252 | 0.487520668 | -0.260779203 | 0.794262783 | 0.914628224 |
| ZNF490 | 147.9564023 | -0.33284026 | 0.501718667 | -0.663400192 | 0.507074283 | 0.767900174 |
| ZNF491 | 67.6931884 | 0.707007636 | 0.465130865 | 1.520018749 | 0.128506264 | 0.413044179 |
| ZNF493 | 668.1261955 | -0.354715063 | 0.272617154 | -1.301147262 | | |

0.193208053 0.499090647
ZNF496 2580.809012 -0.434225818 0.297063378 -1.461727865
0.143815798 0.434973978
ZNF497 91.6481914 -0.246322704 0.447455767 -0.55049621 0.581979076
0.813002261
ZNF500 311.3530618 -0.242237938 0.279412892 -0.866953335
0.385967579 0.686679504
ZNF502 136.9001052 -0.07327189 0.385996549 -0.189825245 0.849446073
0.938580023
ZNF503 296.70311 0.98913558 0.712360452 1.388532418 0.16497498
0.467115865
ZNF506 541.3860161 -0.356031023 0.337279665 -1.055595874
0.291152887 0.604251771
ZNF507 1126.208151 -0.107114777 0.441995467 -0.242343611
0.808513924 0.91963689
ZNF510 438.8941313 -0.228945879 0.455933515 -0.502147509
0.615563763 0.830968556
ZNF511 692.1073397 0.597985996 0.344930459 1.733642191 0.082981589
0.329644853
ZNF512 1559.613374 0.049335668 0.322349841 0.153050076 0.878358786
0.95141893
ZNF512B 444.4694816 -0.720622603 0.553271979 -1.302474425
0.192754255 0.498908723
ZNF513 678.8007919 -0.02972528 0.384686985 -0.077271344 0.938407687
0.977342821
ZNF514 519.8673196 -0.352125273 0.267038019 -1.31863348 0.187291675
0.49211797
ZNF516 1537.131427 0.109420194 0.391074881 0.27979346 0.779635969
0.908654895
ZNF517 144.1222099 -0.169363598 0.355315224 -0.476657306
0.633606169 0.839689885
ZNF518A 1798.28543 -0.42643969 0.378079685 -1.127909555 0.259358118
0.572656383
ZNF518B 1462.030791 -0.003119313 0.312881332 -0.009969637
0.992045512 0.99688215
ZNF519 789.4400437 0.122152169 0.318988736 0.382935683 0.701767453
0.872479299
ZNF521 1441.670242 -0.4690673 0.379622418 -1.235615385 0.216601563
0.524289385
ZNF524 63.7863411 0.132902729 0.752921537 0.17651604 0.859888548
0.943267672
ZNF525 521.0906915 -0.640415059 0.378151056 -1.69354296 0.090352165
0.344491755
ZNF526 367.0276373 -0.347140927 0.305850901 -1.135000505
0.256375107 0.568552367
ZNF527 138.1110605 -0.093195575 0.334638606 -0.278496184
0.780631494 0.909271055
ZNF528 164.6467186 -1.292895912 0.502021936 -2.575377325
0.010013079 0.114014677
ZNF529 472.2790729 0.256554087 0.32145835 0.798094331 0.424815751
0.71350603
ZNF530 159.3323552 -0.558439666 0.358176879 -1.559117012
0.118968688 0.39772454
ZNF532 4158.552535 -0.388692437 0.355180068 -1.094353181
0.27380011 0.586847211
ZNF536 15.31974528 -0.385659443 0.842738076 -0.457626698
0.647220659 0.846597949
ZNF540 12.82257004 0.249356675 0.804526043 0.309942329 0.756604813
0.898563152
ZNF542P 856.8370249 0.06765729 0.245583264 0.275496338 0.782934939
0.909909835
ZNF543 183.608938 -0.025805667 0.403113555 -0.064015874
0.948957587 0.98134156
ZNF544 1371.163275 -0.334643746 0.261179134 -1.281280552

| | | | | | | |
|---|---|---|---|---|---|---|
| | | | | | 0.200095141 | 0.506083488 |
| ZNF546 | 39.55166798 | 0.360156494 | 0.580308703 | 0.620629144 | 0.534843659 | 0.784048039 |
| ZNF547 | 67.99193881 | 0.489821105 | 0.48622871 | 1.007388284 | 0.313748219 | 0.62342819 |
| ZNF548 | 287.1323664 | -0.30679858 | 0.292309403 | -1.049567945 | 0.293916801 | 0.606430922 |
| ZNF549 | 474.5787045 | -0.40520456 | 0.334591845 | -1.211041352 | 0.225879557 | 0.534727976 |
| ZNF550 | 361.3755775 | -0.088226653 | 0.299495451 | -0.294584284 | 0.768311469 | 0.904332533 |
| ZNF551 | 567.2127694 | -0.495926803 | 0.366119132 | -1.354550362 | 0.175560857 | 0.47969026 |
| ZNF552 | 415.2240965 | -1.259163797 | 0.481859974 | -2.613132166 | 0.00897166 | 0.106919917 |
| ZNF554 | 271.6939939 | -0.59195433 | 0.400520463 | -1.47796276 | 0.139417746 | 0.428072582 |
| ZNF555 | 178.078042 | -0.446255463 | 0.370330295 | -1.205020136 | 0.228195529 | 0.537150955 |
| ZNF557 | 214.2936762 | -0.193227559 | 0.330032377 | -0.585480614 | 0.558224598 | 0.798855459 |
| ZNF558 | 708.5695743 | -0.193251302 | 0.241467426 | -0.800320379 | 0.423525199 | 0.71273442 |
| ZNF559 | 577.9578568 | 0.10210792 | 0.268628615 | 0.380108128 | 0.703865153 | 0.873412255 |
| ZNF559-ZNF177 | 93.58214219 | -0.150705916 | 2.104054948 | -0.071626416 | 0.942899217 | 0.978574897 |
| ZNF561 | 1782.232036 | -0.344713214 | 0.223520818 | -1.542197356 | 0.123025642 | 0.40442514 |
| ZNF562 | 713.9193037 | 0.096642888 | 0.298023757 | 0.324279141 | 0.74572671 | 0.893776262 |
| ZNF563 | 88.51515685 | 0.042614657 | 0.470055211 | 0.090658833 | 0.927763683 | 0.972633911 |
| ZNF564 | 356.2867993 | 0.24961517 | 0.354116229 | 0.70489616 | 0.48087486 | 0.752563699 |
| ZNF565 | 67.44052582 | 0.279624362 | 0.528469887 | 0.529120711 | 0.596721715 | 0.821812117 |
| ZNF566 | 321.6431016 | 0.436521465 | 0.287745942 | 1.517037779 | 0.129257146 | 0.413472911 |
| ZNF567 | 553.4924904 | 0.054967929 | 0.284066036 | 0.193504052 | 0.846564233 | 0.938145276 |
| ZNF568 | 300.2412094 | -0.612523071 | 0.284648544 | -2.151857387 | 0.031408588 | 0.202023044 |
| ZNF569 | 491.511187 | 0.714168296 | 0.401408657 | 1.779155191 | 0.075214322 | 0.314600339 |
| ZNF57 | 153.5692796 | -0.127659865 | 0.390197867 | -0.327166999 | 0.74354158 | 0.892534058 |
| ZNF570 | 294.1641925 | 0.03723585 | 0.281720033 | 0.132173243 | 0.894847265 | 0.95833177 |
| ZNF571 | 472.6748695 | -0.029826678 | 0.37626529 | -0.079270341 | 0.936817596 | 0.976155095 |
| ZNF572 | 32.49555725 | -0.346964216 | 0.680584624 | -0.509803195 | 0.610189347 | 0.828605673 |
| ZNF573 | 73.71840507 | 0.389653006 | 0.545341648 | 0.714511732 | 0.474910804 | 0.748737095 |
| ZNF574 | 490.0332048 | 0.125526779 | 0.332231137 | 0.377829663 | 0.705557137 | 0.874010559 |
| ZNF576 | 248.9632667 | 0.572193317 | 0.330901065 | 1.72919757 | 0.083773741 | 0.331144197 |
| ZNF577 | 822.1193341 | -0.472712056 | 0.241422514 | -1.958028052 | 0.050226721 | 0.257653791 |
| ZNF578 | 308.2821583 | -0.999579572 | 0.433632864 | -2.305128729 | 0.021159365 | 0.165258892 |
| ZNF579 | 356.4000748 | 0.449920812 | 0.446316969 | 1.008074625 | 0.313418638 | |

0.623018103
ZNF580 388.8203834 0.705139129 0.434997039 1.621020526 0.105013237
0.371346423
ZNF581 1307.344955 0.379533886 0.469651791 0.808117616 0.419022877
0.709924862
ZNF582 152.2356148 -0.104621864 0.314442872 -0.332721372
0.739344615 0.891550525
ZNF583 123.8938114 -0.02877038 0.339683626 -0.084697578 0.932501821
0.97453841
ZNF584 157.2612405 0.230739643 0.324429053 0.71121757 0.476949423
0.749746108
ZNF585A 579.4381234 -0.23493817 0.308600976 -0.7613008 0.446477423
0.728647696
ZNF585B 319.9085 -0.262657379 0.36238165 -0.724808719
0.468569386 0.744337782
ZNF586 351.7459974 -0.18956739 0.290735704 -0.652026522 0.514384063
0.77285333
ZNF587 1194.688035 -0.444861071 0.311083145 -1.430039134
0.152705788 0.448798819
ZNF587B 490.5692324 0.135241273 0.403430333 0.335228319 0.737452861
0.89096897
ZNF589 1378.845895 -0.193440096 0.369924617 -0.52291761 0.60103159
0.824625774
ZNF592 837.4275714 -0.309275237 0.372465291 -0.830346461
0.406342927 0.701474651
ZNF593 352.7459857 1.116597437 0.553956097 2.015678576 0.043833597
0.240941849
ZNF594 290.586034 -0.471503581 0.55433679 -0.850572413
0.395006919 0.693813886
ZNF595 327.5764574 0.108625256 0.334718419 0.324527274 0.745538877
0.893621735
ZNF596 76.14348427 -0.297400736 0.414159728 -0.718082219
0.472706593 0.747024138
ZNF597 136.5467362 -0.361917011 0.358988012 -1.008159044
0.313378116 0.623018103
ZNF598 549.1437401 0.294614423 0.468904603 0.628303542 0.529805112
0.780409931
ZNF599 147.9414129 0.121499743 0.345797244 0.351361226 0.725317367
0.884193635
ZNF600 409.8466827 -0.97274411 0.300964366 -3.232090639 0.001228881
0.030866472
ZNF605 742.0723363 -0.371650046 0.354866761 -1.047294609
0.294963722 0.606966925
ZNF606 617.3509347 -0.009894705 0.284758848 -0.034747663
0.972280954 0.989954357
ZNF607 486.955131 -0.115308405 0.462797434 -0.249155238
0.803240703 0.917969611
ZNF608 3334.109523 -0.104245056 0.37259044 -0.279784571
0.77964279 0.908654895
ZNF609 2456.789003 -0.536556602 0.421833224 -1.271963827
0.203385972 0.509735688
ZNF610 313.8127953 -0.137487838 0.33509343 -0.410297026
0.681588074 0.862590493
ZNF611 415.0296542 -0.99367519 0.397653139 -2.498849101 0.012459735
0.126670974
ZNF613 402.7845072 -0.92089186 0.271373614 -3.393446569 0.00069019
0.021634041
ZNF614 1113.190743 -0.632586372 0.232982057 -2.715172057
0.006624135 0.089169476
ZNF615 490.7947664 -0.717409561 0.323604884 -2.216930573
0.026627836 0.186591641
ZNF616 730.1273223 -0.351792217 0.373288577 -0.942413561
0.345980947 0.652683033
ZNF618 1379.679558 -0.189776927 0.388424884 -0.488580767

| | | | | | | |
|---|---|---|---|---|---|---|
| | | | | | 0.625138532 | 0.835033444 |
| ZNF619 | 191.9724169 | -0.290748603 | 0.422546983 | -0.688085857 | 0.491398716 | 0.758779636 |
| ZNF620 | 407.3190843 | -0.412239561 | 0.302778973 | -1.361519784 | 0.173349486 | 0.477379751 |
| ZNF621 | 1298.890578 | -0.621124225 | 0.365804048 | -1.697969796 | 0.089513463 | 0.342777615 |
| ZNF622 | 699.2326795 | -0.211528888 | 0.37425137 | -0.565205381 | 0.571934077 | 0.807351345 |
| ZNF623 | 473.9191044 | -0.053725174 | 0.278652955 | -0.192803173 | 0.847113118 | 0.938145276 |
| ZNF624 | 124.7634268 | 0.129427425 | 0.346329342 | 0.373711982 | 0.70861861 | 0.875585221 |
| ZNF625 | 200.3891047 | 0.797534822 | 0.499096659 | 1.597956645 | 0.110052626 | 0.381537567 |
| ZNF625-ZNF20 | 285.161372 | -0.062856416 | 0.326159373 | -0.192716879 | 0.847180704 | 0.938145276 |
| ZNF626 | 665.8377 | -0.094322115 | 0.335431435 | -0.28119641 | 0.778559758 | 0.908160087 |
| ZNF627 | 762.3277397 | -0.206840222 | 0.32691262 | -0.632707975 | 0.526924361 | 0.778755887 |
| ZNF628 | 71.49637877 | 0.105866752 | 0.62905405 | 0.168295161 | 0.866351081 | 0.945705299 |
| ZNF629 | 1293.630162 | -0.546530542 | 0.255198131 | -2.141593044 | 0.032226244 | 0.204648115 |
| ZNF630 | 85.39643685 | -0.489041012 | 0.398298859 | -1.227824286 | 0.219512937 | 0.528021367 |
| ZNF638 | 4718.85527 | -0.128530844 | 0.31136301 | -0.41280062 | 0.679752697 | 0.861682789 |
| ZNF639 | 836.5274275 | 0.310365668 | 0.281593856 | 1.102174856 | 0.270385661 | 0.583170438 |
| ZNF641 | 532.0756507 | -0.92290023 | 0.338064705 | -2.729951446 | 0.006334365 | 0.086470416 |
| ZNF644 | 1724.029635 | 0.110432983 | 0.35098305 | 0.314639077 | 0.753035706 | 0.896586461 |
| ZNF646 | 684.8752035 | -0.415929845 | 0.493144075 | -0.8434246 | 0.398991018 | 0.697352242 |
| ZNF648 | 16.45808823 | -0.895789317 | 0.749181053 | -1.195691366 | 0.231817022 | 0.540977702 |
| ZNF649 | 1232.392342 | -0.63628734 | 0.252516506 | -2.519785143 | 0.011742649 | 0.122945289 |
| ZNF652 | 663.0670006 | 0.180162534 | 0.392778601 | 0.458687243 | 0.646458776 | 0.846270264 |
| ZNF653 | 142.0430922 | 0.156904628 | 0.478214085 | 0.328105409 | 0.742831964 | 0.892412269 |
| ZNF654 | 277.1272231 | -0.240255813 | 0.393833089 | -0.610044761 | 0.541832157 | 0.788653521 |
| ZNF655 | 1629.8458 | -0.250398142 | 0.304644636 | -0.821935176 | 0.411113792 | 0.704587363 |
| ZNF658 | 167.3437798 | 0.275929423 | 0.431860342 | 0.638932071 | 0.522867123 | 0.777662198 |
| ZNF658B | 104.0204045 | 0.213726961 | 0.427475774 | 0.499974441 | 0.617093074 | 0.831835425 |
| ZNF66 | 58.62337869 | -0.752462198 | 0.681642617 | -1.103895471 | 0.269638484 | 0.582376584 |
| ZNF660 | 237.5442169 | 0.0765545 | 0.313555976 | 0.244149388 | 0.807115123 | 0.919137032 |
| ZNF663P | 82.45938606 | 0.22941188 | 0.43414934 | 0.528416973 | 0.597209959 | 0.821867951 |
| ZNF664 | 5215.872813 | -0.171376312 | 0.305543626 | -0.560889828 | 0.574872644 | 0.808359708 |
| ZNF665 | 177.4406092 | 0.164929262 | 0.31711455 | 0.520093644 | 0.602998308 | 0.825043952 |
| ZNF667 | 229.6888141 | 0.044869126 | 0.403380335 | 0.111232805 | 0.911431738 | |

0.966866396
ZNF668 164.6485581 0.490384156 0.548165418 0.89459156 0.371005465
0.676265004
ZNF669 317.4867939 0.046966807 0.273770309 0.171555517 0.863786981
0.945079904
ZNF670 539.8584165 0.377437505 0.284183179 1.328148648 0.184129005
0.488572217
ZNF670-ZNF695 33.59173074 -0.624642583 0.621855615 -1.004481697
0.315146489 0.624652128
ZNF671 232.73544 -0.486389506 0.299967151 -1.6214759 0.104915614
0.37133017
ZNF672 672.8123896 -0.101055869 0.343283141 -0.294380518
0.768467152 0.904332533
ZNF674 141.1722081 -0.501991855 0.353107327 -1.421641004
0.155130495 0.453028823
ZNF675 599.0587735 -0.204523953 0.259420791 -0.788386898
0.430470432 0.717078019
ZNF677 1061.086405 -0.010160317 0.238491866 -0.042602364
0.966018511 0.988700292
ZNF678 816.0590162 0.307523438 0.296838154 1.035997001 0.300203542
0.610801113
ZNF680 741.0209373 0.373356372 0.256011596 1.45835727 0.144742092
0.435928729
ZNF681 519.2035172 0.332864625 0.261560079 1.272612498 0.203155582
0.509545953
ZNF682 454.8131055 0.183328184 0.298831969 0.613482502 0.539557349
0.78700361
ZNF684 108.9298362 0.751513359 0.392365772 1.915338725 0.05544932
0.272416332
ZNF687 1219.321762 -0.777583362 0.397539384 -1.955990761
0.050466242 0.258046297
ZNF688 67.25276379 0.950041412 0.499241342 1.902970231 0.05704443
0.276551437
ZNF689 465.6194721 0.254310623 0.365966056 0.694902216 0.487116584
0.756439734
ZNF69 172.9915965 0.160958455 0.406195878 0.396258218 0.691914547 0.86812349
ZNF691 361.3187456 0.400148697 0.341226613 1.172677283 0.240925245
0.552437022
ZNF692 1191.605931 -0.01905441 0.270748473 -0.070376795 0.94389376
0.978984717
ZNF695 458.8857234 0.455865784 0.343942948 1.32541105 0.185034854
0.489564662
ZNF696 129.6195699 -0.047008723 0.34320271 -0.136970722
0.891053937 0.957017706
ZNF697 443.8227905 -0.219596125 0.327885866 -0.669733428
0.503027738 0.765194965
ZNF699 65.8886232 -0.56071334 0.761477928 -0.736348776 0.46151847
0.73902094
ZNF7 671.4844156 -0.132669443 0.257537795 -0.515145526 0.606451332
0.826611695
ZNF70 155.0014515 -0.224917946 0.469115828 -0.479450772 0.631617981
0.838768903
ZNF700 484.9304393 -0.194904529 0.247917188 -0.786167876
0.431769143 0.718311547
ZNF701 366.5479506 -0.877924618 0.409171846 -2.145613455
0.031903831 0.203537856
ZNF702P 738.9518314 0.027942854 0.365732925 0.076402347 0.939099002
0.977796709
ZNF703 1542.323179 0.019781899 0.628669867 0.031466276 0.974897687
0.991261106
ZNF704 522.6615381 -0.071366921 0.480272156 -0.148596831
0.881871773 0.953217539
ZNF706 1360.558322 0.606686705 0.285391973 2.125801572 0.033519785
0.208789229

| | | | | | | |
|---|---|---|---|---|---|---|
| ZNF707 | 92.59518738 | -0.97962264 | 0.529933745 | -1.848575692 | 0.064519105 | 0.293324484 |
| ZNF708 | 905.2370291 | 0.200800213 | 0.248057332 | 0.809491142 | 0.418232696 | 0.709589334 |
| ZNF709 | 148.2970714 | 0.096603684 | 0.442561729 | 0.21828296 | 0.82720865 | 0.928082319 |
| ZNF71 | 381.3876267 | -0.023435629 | 0.355658559 | -0.065893619 | 0.947462521 | 0.980889444 |
| ZNF710 | 792.4637793 | 0.182633341 | 0.422641644 | 0.432123392 | 0.665651734 | 0.855161526 |
| ZNF711 | 1042.227548 | -0.1940971 | 0.372614398 | -0.520906065 | 0.60243221 | 0.825021776 |
| ZNF713 | 116.6144005 | -0.551993939 | 0.368480286 | -1.498028418 | 0.134125867 | 0.420619758 |
| ZNF714 | 1123.784941 | -0.015544605 | 0.319763936 | -0.048612751 | 0.961227908 | 0.986785383 |
| ZNF717 | 154.1819711 | 0.333042197 | 0.436946107 | 0.762204289 | 0.445938086 | 0.728501578 |
| ZNF718 | 432.0047437 | -0.345430253 | 0.347835873 | -0.993084039 | 0.320669002 | 0.630327731 |
| ZNF720 | 511.7282784 | -0.093660278 | 0.29705362 | -0.315297547 | 0.752535747 | 0.896586461 |
| ZNF721 | 1231.084059 | 0.102247335 | 0.268027123 | 0.381481301 | 0.702846144 | 0.872703476 |
| ZNF724 | 176.5818662 | -0.062372431 | 0.619772767 | -0.100637579 | 0.919838164 | 0.969770994 |
| ZNF726 | 103.7020556 | 0.213846213 | 0.51320345 | 0.416688962 | 0.676905929 | 0.86029961 |
| ZNF730 | 324.435273 | 0.152031902 | 0.412297186 | 0.368743486 | 0.712318932 | 0.877514455 |
| ZNF732 | 138.9723876 | 0.068955202 | 0.487121481 | 0.141556479 | 0.887430346 | 0.955922023 |
| ZNF738 | 1324.567987 | 0.347778079 | 0.305238825 | 1.139363839 | 0.254551432 | 0.567161077 |
| ZNF74 | 990.4205955 | 0.261423563 | 0.297454401 | 0.878869373 | 0.379472106 | 0.681253679 |
| ZNF740 | 1134.455371 | -0.298111854 | 0.391148529 | -0.762144893 | 0.445973531 | 0.728501578 |
| ZNF746 | 502.9670312 | -0.073442722 | 0.343511023 | -0.21380019 | 0.830702877 | 0.930492716 |
| ZNF747 | 211.2420249 | 0.147795286 | 0.313073055 | 0.472079228 | 0.636870238 | 0.841268221 |
| ZNF749 | 186.4155288 | 0.355018816 | 0.373578543 | 0.950319079 | 0.341950147 | 0.648704346 |
| ZNF750 | 479.2967511 | -1.441307219 | 0.82168386 | -1.754089728 | 0.079415133 | 0.322931692 |
| ZNF75A | 790.7733145 | -0.517756831 | 0.24846135 | -2.083852602 | 0.037173577 | 0.220938673 |
| ZNF75D | 338.40439 | -0.932759795 | 0.359945723 | -2.591390134 | 0.009558906 | 0.110870153 |
| ZNF76 | 752.2296929 | -0.065761148 | 0.440451881 | -0.14930382 | 0.881313901 | 0.952987398 |
| ZNF761 | 797.4691401 | -0.958251862 | 0.326227155 | -2.937376145 | 0.003310024 | 0.058120796 |
| ZNF763 | 158.0855663 | -1.464721225 | 0.490606097 | -2.985534085 | 0.002830836 | 0.052437865 |
| ZNF764 | 147.0515092 | 0.260627485 | 0.338386288 | 0.770206991 | 0.441177118 | 0.724743658 |
| ZNF765 | 545.7435575 | -0.589022853 | 0.300414697 | -1.960699195 | 0.049914123 | 0.257391965 |
| ZNF766 | 1060.927944 | -0.188869853 | 0.294314298 | -0.64172843 | 0.521049526 | 0.776125907 |
| ZNF767P | 270.7820772 | -0.23055325 | 0.352451454 | -0.654141859 | 0.513020415 | 0.771819984 |
| ZNF768 | 928.2308594 | 0.233959221 | 0.330960742 | 0.706909284 | 0.479622854 | |

0.751840877

| | | | | | | |
|---|---|---|---|---|---|---|
| ZNF77 | 192.5233119 | 0.508897001 | 0.307653245 | 1.654125252 | 0.098102066 | 0.359755003 |
| ZNF770 | 1767.097952 | 0.249384599 | 0.304342874 | 0.819419874 | 0.4125469 | 0.705244144 |
| ZNF771 | 97.661259 | 0.593874445 | 0.455070831 | 1.305015405 | 0.191887602 | 0.497612799 |
| ZNF772 | 308.5519592 | -0.013509674 | 0.312476905 | -0.043234151 | 0.965514882 | 0.988700292 |
| ZNF773 | 222.3066953 | -0.508828291 | 0.337223775 | -1.508874308 | 0.131330907 | 0.416998992 |
| ZNF774 | 88.32047524 | -0.690562273 | 0.405233291 | -1.704110419 | 0.088360461 | 0.341710805 |
| ZNF775 | 167.2083496 | 1.15612654 | 0.481053434 | 2.403322496 | 0.016246852 | 0.144168905 |
| ZNF776 | 706.5831448 | -0.467626108 | 0.252176857 | -1.854357744 | 0.063687996 | 0.290852073 |
| ZNF777 | 341.3583877 | -1.111069159 | 0.525990771 | -2.112335845 | 0.034657653 | 0.21269873 |
| ZNF778 | 872.1429107 | -0.681217274 | 0.328014595 | -2.07678952 | 0.03782099 | 0.223348477 |
| ZNF780A | 244.5803482 | -0.155474145 | 0.344258154 | -0.451620806 | 0.651542179 | 0.847876685 |
| ZNF780B | 240.514028 | -0.147842022 | 0.354875272 | -0.416602772 | 0.676968982 | 0.86029961 |
| ZNF781 | 113.7580979 | 0.446535528 | 0.434224115 | 1.028352671 | 0.303783963 | 0.614002998 |
| ZNF782 | 120.3692971 | -0.617255422 | 0.509516934 | -1.211452221 | 0.225722136 | 0.53459368 |
| ZNF783 | 300.0635547 | -0.215846108 | 0.272191373 | -0.792993937 | 0.427781355 | 0.715137294 |
| ZNF784 | 250.7901571 | 0.116530139 | 0.388722448 | 0.299777231 | 0.764347084 | 0.902756684 |
| ZNF785 | 363.0475522 | 0.315217238 | 0.26187745 | 1.203682248 | 0.228712416 | 0.537452922 |
| ZNF786 | 294.9618561 | 0.131910204 | 0.268266645 | 0.491713025 | 0.622922229 | 0.833825637 |
| ZNF787 | 656.1682551 | 1.035731808 | 0.470322436 | 2.20217393 | 0.027653025 | 0.190260335 |
| ZNF788P | 164.9030265 | 0.330269794 | 0.835940664 | 0.395087604 | 0.692778237 | 0.86854217 |
| ZNF789 | 259.6922501 | -0.192525419 | 0.306359082 | -0.628430592 | 0.529721902 | 0.780385606 |
| ZNF79 | 137.7929601 | -0.412804853 | 0.449012271 | -0.919362074 | 0.35790622 | 0.665416085 |
| ZNF790 | 154.7046652 | -0.047862822 | 0.323636157 | -0.147890838 | 0.882428919 | 0.953255639 |
| ZNF791 | 972.8982323 | -0.157675079 | 0.36790613 | -0.428574209 | 0.668233118 | 0.856934809 |
| ZNF792 | 146.1335682 | -0.467639356 | 0.326847398 | -1.430757471 | 0.152499736 | 0.448453866 |
| ZNF793 | 519.5889547 | -0.329845526 | 0.300213846 | -1.098701911 | 0.271898109 | 0.584197156 |
| ZNF799 | 191.9601996 | -0.838934871 | 0.375383043 | -2.234876848 | 0.025425435 | 0.18194777 |
| ZNF8 | 591.7244485 | -0.087639318 | 0.392199512 | -0.223455958 | 0.823180657 | 0.926014081 |
| ZNF800 | 237.715071 | -0.165100272 | 0.35142778 | -0.469798581 | 0.638498928 | 0.842491082 |
| ZNF804A | 10.11777582 | -2.242415505 | 1.374763127 | -1.631128637 | 0.102863179 | 0.367817927 |
| ZNF805 | 228.5954808 | -0.030464981 | 0.548765359 | -0.055515495 | 0.955727786 | 0.984323736 |
| ZNF808 | 315.3137418 | -0.374884532 | 0.371595353 | -1.008851506 | 0.313045855 | 0.622994042 |

ZNF81 189.3824473 -0.611880788 0.504386574 -1.213118705 0.225084439
0.534448858
ZNF813 731.2208298 -0.136449243 0.284180279 -0.480150287
0.631120533 0.838320891
ZNF814 474.7009718 0.22674567 0.395459008 0.573373385 0.566391907
0.80448756
ZNF815P 80.82913859 0.038349423 0.447279566 0.08573927 0.931673684
0.974341807
ZNF816 295.6236425 -0.660665372 0.412382451 -1.602069562
0.109140228 0.379241044
ZNF816-ZNF321P 10.45897165 -5.999265365 1.738510868 -3.450806938
0.000558913 0.018754686
ZNF821 336.32328 -0.008713952 0.351704022 -0.024776379
0.980233332 0.993196129
ZNF823 340.4969844 -0.505032807 0.348140018 -1.450660025
0.146874552 0.439927706
ZNF826P 920.0691602 0.187542515 0.307960057 0.608983247 0.542535544
0.789223006
ZNF827 765.6553227 -0.18990142 0.279306054 -0.679904418 0.496564984
0.761255797
ZNF829 252.2088045 -0.284017684 0.312603076 -0.908556906
0.363584063 0.670353372
ZNF83 1218.751943 -0.421362498 0.287628948 -1.464951635 0.142934114
0.433233718
ZNF830 572.2435724 0.337990017 0.395024241 0.85561842 0.392208878
0.691514794
ZNF833P 33.33377144 0.326914104 0.570604322 0.572926091 0.566694738
0.804513376
ZNF836 948.7860812 -1.105090065 0.538789815 -2.051059679
0.040261135 0.231327132
ZNF837 22.71312864 0.130639358 0.853179044 0.153120683 0.878303106
0.95141893
ZNF839 568.9781731 -0.356193122 0.243208422 -1.464559158
0.143041232 0.43332239
ZNF84 1269.785476 -0.049990871 0.2594616 -0.192671557 0.8472162
0.938145276
ZNF841 388.4482235 -1.178709828 0.372293612 -3.166075884
0.001545105 0.036390779
ZNF843 18.56901393 -0.065651457 0.729979789 -0.089935993
0.928338079 0.972633911
ZNF844 274.3424912 0.107310705 0.28345263 0.378584264 0.704996609
0.873744266
ZNF845 722.1412402 -0.531745667 0.280641712 -1.89474923 0.058125632
0.279411903
ZNF846 160.0793996 0.071038628 0.326436616 0.217618442 0.827726414
0.928330355
ZNF85 527.3997411 0.130648592 0.25367074 0.515032171 0.60653054 0.826611695
ZNF850 290.0754569 -0.035773711 0.399172162 -0.089619754
0.928589386 0.972633911
ZNF852 85.9868617 -0.621426094 0.469425939 -1.323800077
0.185569451 0.490113028
ZNF853 145.841769 0.324383643 0.368519008 0.880235851 0.378731556
0.681253679
ZNF860 14.26518608 -1.624561734 0.995412733 -1.632048375
0.102669296 0.367636293
ZNF862 364.644232 -0.390316677 0.53292412 -0.732405728
0.463920964 0.741091391
ZNF865 266.7534779 0.33442903 0.440350839 0.759460413 0.447577186
0.729044534
ZNF876P 344.9525102 0.487988757 0.307879143 1.585001022 0.112966102
0.386689697
ZNF878 53.21063792 -0.611246324 0.521183539 -1.172804355
0.240874271 0.552437022
ZNF879 211.7806503 -0.272932878 0.342738949 -0.796328749

0.42584098 0.713938944
ZNF880 460.6588617 -0.195655892 0.348785723 -0.560963017
0.574822749 0.808359708
ZNF883 377.2997492 0.190408203 0.305717682 0.622823652 0.533400414
0.783143103
ZNF887P 59.24058928 -0.449204005 0.559341519 -0.803094335
0.421920212 0.712063713
ZNF888 263.483656 -0.279955433 0.302560042 -0.925288847
0.354815679 0.662221067
ZNF891 237.7954021 0.217184222 0.410618711 0.528919448 0.59686133
0.821812117
ZNF90 1306.743432 -1.086167537 0.390049247 -2.78469333 0.005357837
0.077809098
ZNF91 1370.568836 0.027055976 0.266691263 0.101450551 0.919192809 0.969677217
ZNF92 1166.863304 0.159950552 0.301955689 0.529715311 0.596309331 0.821812117
ZNF93 956.2298553 -0.306700409 0.328067737 -0.934869158 0.349855733
0.656482539
ZNF969P 25.08648833 -0.034019245 0.645963133 -0.052664376
0.957999324 0.98520931
ZNFX1 342.0229356 -0.166698324 0.41288568 -0.403739661 0.686404171
0.865550468
ZNHIT1 807.3088808 0.305609331 0.412933464 0.740093399 0.459243324
0.737472333
ZNHIT2 44.42848283 0.562676422 0.757569462 0.74273905 0.457639683
0.736329582
ZNHIT3 1088.304379 0.604159685 0.372639825 1.621296611 0.104954042
0.37133017
ZNHIT6 711.1074526 -0.094074192 0.261235488 -0.360112604
0.718762927 0.881017401
ZNRD1 279.5926628 0.684707952 0.390351023 1.754082638 0.079416348 0.322931692
ZNRD1ASP 162.316027 -0.317075497 0.380232272 -0.833899488
0.404337626 0.700367583
ZNRF1 1148.710647 0.246256476 0.438954072 0.561007385 0.574792503 0.808359708
ZNRF2 1192.124607 -0.496778169 0.311372379 -1.595447135 0.110612283
0.382690751
ZNRF2P1 29.5503713 -0.405574489 0.647398532 -0.626468039
0.531007994 0.781369765
ZNRF3 728.5590147 0.216529707 0.328260153 0.659628362 0.509492349 0.769564758
ZP3 50.73361491 1.555681215 0.73664652 2.111842211 0.034699985 0.212806795
ZPR1 1594.021886 0.130728765 0.267466265 0.488767302 0.62500645 0.835033444
ZRANB1 966.1962586 -0.536494412 0.266298278 -2.014637186
0.043942678 0.241112355
ZRANB2 3557.818098 0.409326241 0.269396967 1.519416661 0.128657651
0.413310782
ZRANB3 505.4944361 0.106854019 0.413252537 0.258568332 0.79596832
0.915299455
ZRSR2 249.5991056 -0.341323338 0.309604533 -1.102449421 0.270266336
0.583170438
ZRSR2P1 10.12467456 -0.349793092 1.205976769 -0.290049611
0.771778284 0.905401203
ZSCAN1 23.03699237 -0.198475808 0.745283382 -0.266309182
0.790001102 0.912949942
ZSCAN10 240.452898 -1.302955777 0.429865921 -3.031074839
0.002436848 0.047457544
ZSCAN12 554.5737895 -0.188658762 0.337821622 -0.558456739
0.576532537 0.809193739
ZSCAN12P1 13.96258235 -4.049662099 1.362729946 -2.971727531
0.002961294 0.053803335
ZSCAN16 342.3275006 -0.67064639 0.35760691 -1.875373131 0.060741409
0.284105089
ZSCAN18 1155.560313 0.397032285 0.29250659 1.357344753 0.174671694
0.47862011
ZSCAN2 228.4268301 0.284835869 0.319065806 0.892718251 0.372008074
0.677193964

| | | | | | | |
|---|---|---|---|---|---|---|
| ZSCAN20 | 151.9788626 | -0.427263347 | 0.479479185 | -0.891098843 | 0.372876146 | 0.67811865 |
| ZSCAN21 | 637.79803 | -0.30828441 | 0.256766294 | -1.200642053 | 0.229890081 | 0.538883664 |
| ZSCAN22 | 108.7083408 | -0.709852603 | 0.366152746 | -1.938678902 | 0.052540449 | 0.264550665 |
| ZSCAN23 | 103.429575 | -0.15872608 | 0.889395192 | -0.178465188 | 0.858357658 | 0.942476753 |
| ZSCAN25 | 600.2535761 | -0.458587827 | 0.247280856 | -1.854522158 | 0.063664493 | 0.290852073 |
| ZSCAN26 | 425.8641367 | -0.132223166 | 0.309644387 | -0.42701619 | 0.669367536 | 0.857712777 |
| ZSCAN29 | 1020.331543 | -0.389808535 | 0.29948812 | -1.301582633 | 0.1930591 | 0.498908723 |
| ZSCAN30 | 1013.960556 | -0.724598202 | 0.304004027 | -2.38351514 | 0.017148179 | 0.147741524 |
| ZSCAN31 | 145.2600436 | -0.155628673 | 0.348756383 | -0.446238924 | 0.655424669 | 0.849471345 |
| ZSCAN32 | 491.5196445 | 0.053446625 | 0.267407011 | 0.199869946 | 0.841582296 | 0.935153081 |
| ZSCAN5A | 136.0369756 | -0.283976164 | 0.341794211 | -0.830839596 | 0.406064251 | 0.701474651 |
| ZSCAN9 | 558.1869658 | 0.12947914 | 0.269651033 | 0.48017298 | 0.631104398 | 0.838320891 |
| ZSWIM1 | 239.0449903 | -0.101947166 | 0.361516769 | -0.281998444 | 0.777944703 | 0.908018171 |
| ZSWIM3 | 85.03905842 | 0.007845488 | 0.425456862 | 0.018440149 | 0.985287724 | 0.99460071 |
| ZSWIM4 | 380.204484 | -0.248391476 | 0.296666525 | -0.837275039 | 0.402437987 | 0.699485574 |
| ZSWIM5 | 108.1379485 | 0.137186537 | 0.655756873 | 0.209203354 | 0.834289494 | 0.932107564 |
| ZSWIM6 | 1092.346783 | 0.031025257 | 0.285883625 | 0.108524077 | 0.913579983 | 0.967939236 |
| ZSWIM7 | 801.1527145 | 0.185642199 | 0.413221412 | 0.449256001 | 0.653246994 | 0.848827707 |
| ZSWIM8 | 1043.433247 | -0.564135153 | 0.470303564 | -1.199512817 | 0.230328604 | 0.539112606 |
| ZSWIM9 | 335.6959073 | 0.054315372 | 0.295567975 | 0.183766093 | 0.854196957 | 0.941473214 |
| ZUP1 | 492.6094262 | 0.368860586 | 0.322995537 | 1.141999017 | 0.253454428 | 0.565942797 |
| ZW10 | 909.2511217 | -0.118265853 | 0.250308749 | -0.472479899 | 0.636584286 | 0.841268221 |
| ZWILCH | 1689.414904 | 0.166633429 | 0.256615722 | 0.649350037 | 0.51611215 | 0.773896629 |
| ZWINT | 1790.600601 | 0.201602951 | 0.267997009 | 0.752258214 | 0.451895791 | 0.732888936 |
| ZXDA | 85.8136533 | -0.260687952 | 0.422326203 | -0.617266818 | 0.537058749 | 0.785549392 |
| ZXDB | 374.3397875 | -0.269693486 | 0.290134186 | -0.929547426 | 0.352605459 | 0.659729494 |
| ZXDC | 421.33181 | -0.490571083 | 0.465959575 | -1.052818978 | 0.292423965 | 0.605587347 |
| ZYG11A | 63.10260636 | -0.311809192 | 0.550317433 | -0.56659879 | 0.570986797 | 0.806784083 |
| ZYG11B | 847.9036683 | 0.029268044 | 0.276509653 | 0.105848182 | 0.915702808 | 0.969158739 |
| ZYX | 1117.535755 | -0.726730339 | 0.324446093 | -2.239910896 | 0.025096708 | 0.18130579 |
| ZZEF1 | 773.5588392 | -0.891987837 | 0.497349247 | -1.793483839 | 0.072895592 | 0.310624251 |

### WEB-based GEne SeT AnaLysis Toolkit

WebGestalt

*Translating gene lists into biological insights...*

#### Summary

Result Download

**Job summary**

**GO Slim summary for the user uploaded IDs**

#### Enrichment Results

Redundancy reduction: (•) None ( ) Weighted set cover

Table | **Bar chart** | Volcano plot | DAG

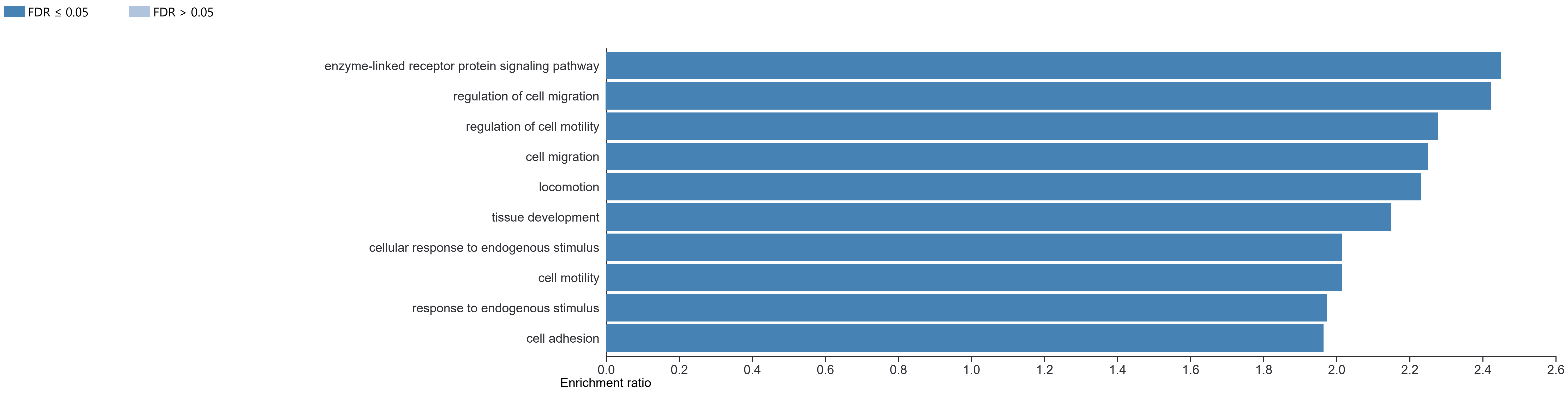

**Select an enriched analyte set...**

GO:0009888: tissue development

##### Analyte set: GO:0009888 tissue development

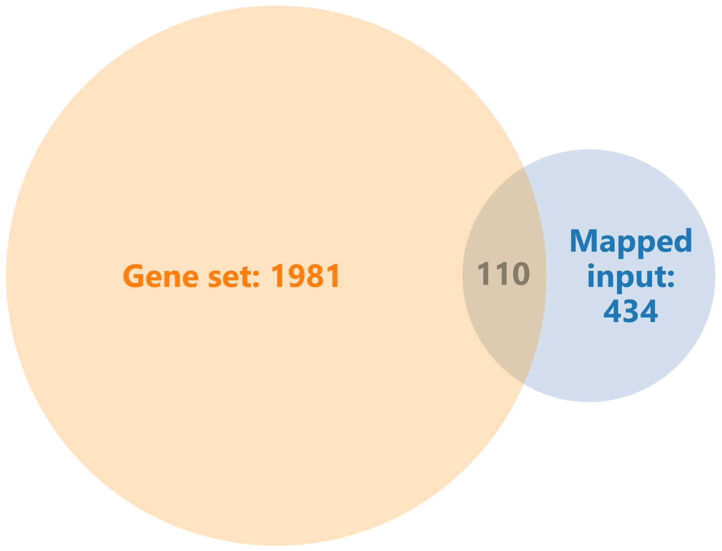

| FDR | P Value |
|---|---|
| 1.9573e-11 | 2.1658e-15 |

| Analyte Set Size | Expected Value | Overlap | Enrichment Ratio |
|---|---|---|---|
| 1981 | 51.182 | 110 | 2.1492 |

| User ID ↑ | Gene Symbol | Gene Name | Entrez Gene ID |
|---|---|---|---|
| ADAMTSL4 | ADAMTSL4 | ADAMTS like 4 | 54507 |
| ALPK2 | ALPK2 | alpha kinase 2 | 115701 |
| AMBN | AMBN | ameloblastin | 258 |
| AMER2 | AMER2 | APC membrane recruitment protein 2 | 219287 |
| APELA | APELA | apelin receptor early endogenous ligand | 100506013 |
| ATP2B1 | ATP2B1 | ATPase plasma membrane Ca2+ transporting 1 | 490 |
| ATP2C2 | ATP2C2 | ATPase secretory pathway Ca2+ transporting 2 | 9914 |
| BMPER | BMPER | BMP binding endothelial regulator | 168667 |
| BNC1 | BNC1 | basonuclin zinc finger protein 1 | 646 |
| BNC2 | BNC2 | basonuclin zinc finger protein 2 | 54796 |

10 per page

1 2 ... 11

WebGestalt is currently developed and maintained by Yuxing Liao, Suhas Vasaikar, Zhiao Shi and Bing Zhang at the Zhang Lab. Other people who have made significant contribution to the project include Jing Wang, Dexter Duncan, Stefan Kirov and Jay Snoddy.
**Funding credits:** NIH/NCI (U24 CA210954); Leidos (15X038); CPRIT (RR160027); NIH/NIAAA (U01 AA016662, U01 AA013512); NIH/NIDA (P01 DA015027); NIH/NIMH (P50 MH078028, P50 MH096972); NIH/NCI (U24 CA159988); NIH/NIGMS (R01 GM088822).

WebGestalt

### WEB-based GEne SeT AnaLysis Toolkit

*Translating gene lists into biological insights...*

#### Summary

Result Download

**Job summary**

**GO Slim summary for the user uploaded IDs**

#### Enrichment Results

Redundancy reduction: ◉ None ○ Weighted set cover

Table | Bar chart | Volcano plot | DAG

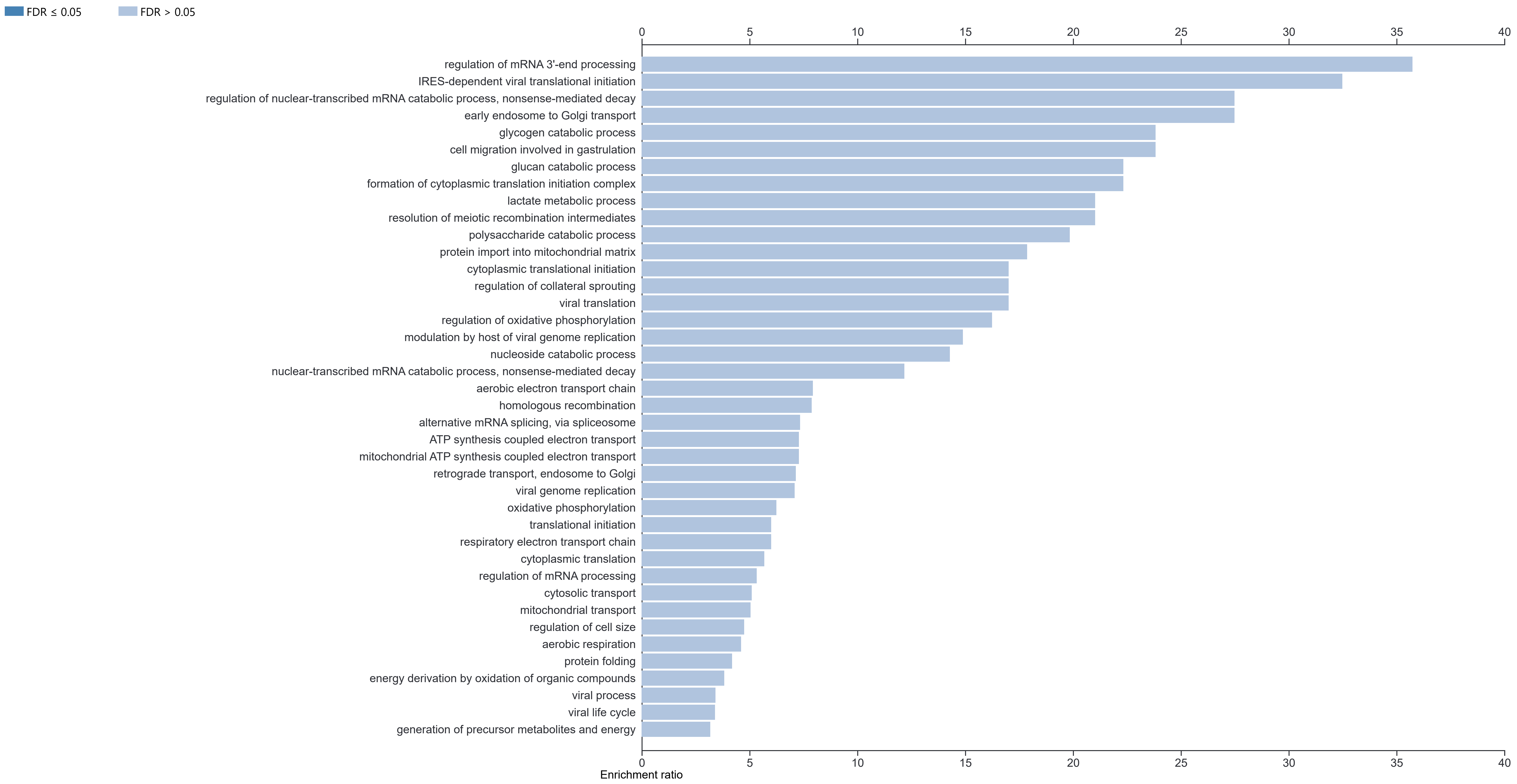

**Select an enriched analyte set...**

GO:0002183: cytoplasmic translational initiatior

##### Analyte set: GO:0002183 cytoplasmic translational initiation

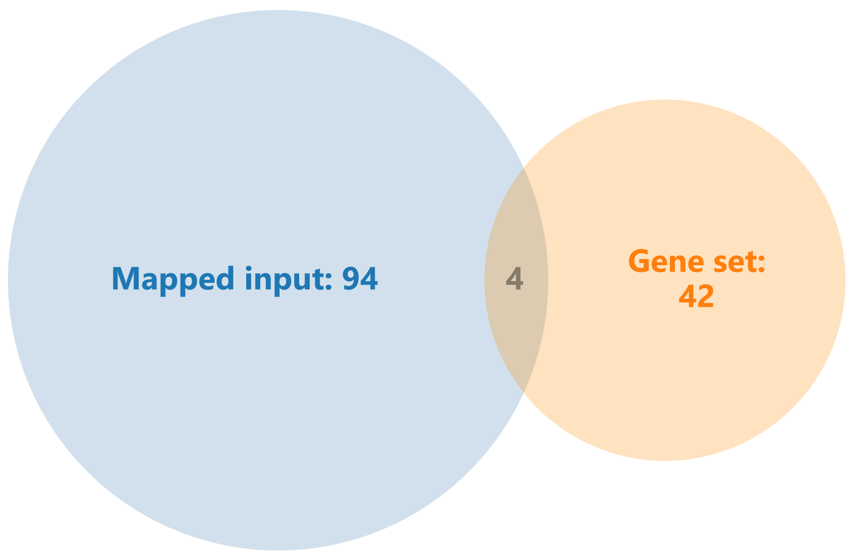

| FDR | P Value |
|---|---|
| 0.79032 | 0.000087454 |

| Analyte Set Size | Expected Value | Overlap | Enrichment Ratio |
|---|---|---|---|
| 42 | 0.23503 | 4 | 17.019 |

| User ID ↑ | Gene Symbol | Gene Name | Entrez Gene ID |
|---|---|---|---|
| DENR | DENR | density regulated re-initiation and release factor | 8562 |
| EIF3B | EIF3B | eukaryotic translation initiation factor 3 subunit B | 8662 |
| EIF3K | EIF3K | eukaryotic translation initiation factor 3 subunit K | 27335 |
| NCBP2 | NCBP2 | nuclear cap binding protein subunit 2 | 22916 |

10 per page

1

WebGestalt is currently developed and maintained by Yuxing Liao, Suhas Vasaikar, Zhiao Shi and Bing Zhang at the Zhang Lab. Other people who have made significant contribution to the project include Jing Wang, Dexter Duncan, Stefan Kirov and Jay Snoddy.
**Funding credits:** NIH/NCI (U24 CA210954); Leidos (15X038); CPRIT (RR160027); NIH/NIAAA (U01 AA016662, U01 AA013512); NIH/NIDA (P01 DA015027); NIH/NIMH (P50 MH078028, P50 MH096972); NIH/NCI (U24 CA159988); NIH/NIGMS (R01 GM088822).

WebGestalt

### WEB-based GEne SeT AnaLysis Toolkit

*Translating gene lists into biological insights...*

#### Summary

Result Download

**Job summary**

**GO Slim summary for the user uploaded IDs**

#### Enrichment Results

Redundancy reduction: (•) None ( ) Weighted set cover

Table | Bar chart | Volcano plot | DAG

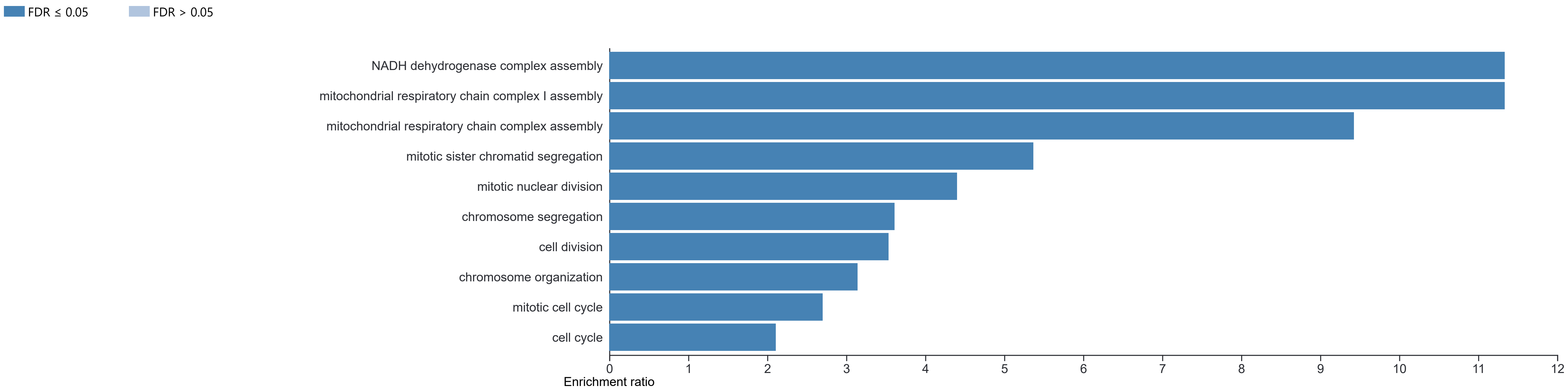

**Select an enriched analyte set...**

GO:0033108: mitochondrial respiratory chain cc

##### Analyte set: GO:0033108 mitochondrial respiratory chain complex assembly

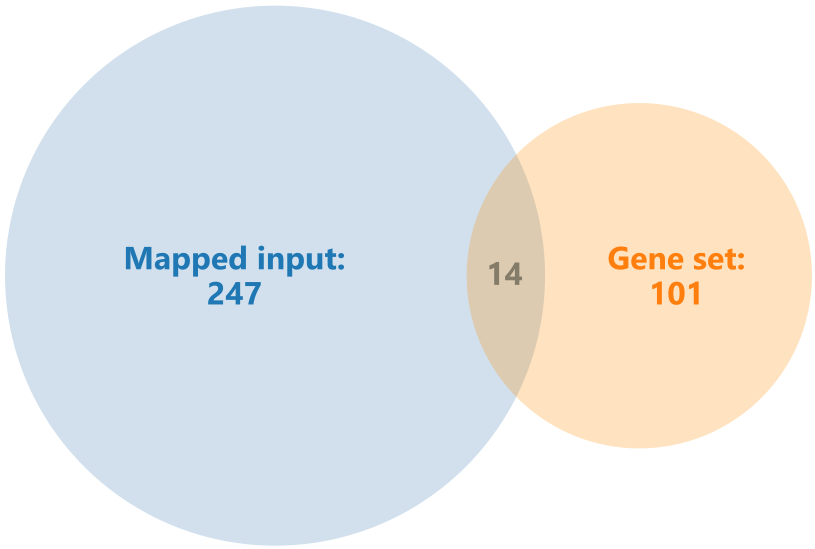

| FDR | P Value |
|---|---|
| 0.0000012680 | 2.5175e-10 |

| Analyte Set Size | Expected Value | Overlap | Enrichment Ratio |
|---|---|---|---|
| 101 | 1.4851 | 14 | 9.4269 |

| User ID ↑ | Gene Symbol | Gene Name | Entrez Gene ID |
|---|---|---|---|
| COA1 | COA1 | cytochrome c oxidase assembly factor 1 | 55744 |
| COX17 | COX17 | cytochrome c oxidase copper chaperone COX17 | 10063 |
| DMAC1 | DMAC1 | distal membrane arm assembly component 1 | 90871 |
| NDUFA5 | NDUFA5 | NADH:ubiquinone oxidoreductase subunit A5 | 4698 |
| NDUFAF2 | NDUFAF2 | NADH:ubiquinone oxidoreductase complex assembly factor 2 | 91942 |
| NDUFAF4 | NDUFAF4 | NADH:ubiquinone oxidoreductase complex assembly factor 4 | 29078 |
| NDUFAF8 | NDUFAF8 | NADH:ubiquinone oxidoreductase complex assembly factor 8 | 284184 |
| NDUFB2 | NDUFB2 | NADH:ubiquinone oxidoreductase subunit B2 | 4708 |
| NDUFB3 | NDUFB3 | NADH:ubiquinone oxidoreductase subunit B3 | 4709 |
| NDUFB6 | NDUFB6 | NADH:ubiquinone oxidoreductase subunit B6 | 4712 |

10 per page

1 2 < >

WebGestalt is currently developed and maintained by Yuxing Liao, Suhas Vasaikar, Zhiao Shi and Bing Zhang at the Zhang Lab. Other people who have made significant contribution to the project include Jing Wang, Dexter Duncan, Stefan Kirov and Jay Snoddy.
**Funding credits:** NIH/NCI (U24 CA210954); Leidos (15X038); CPRIT (RR160027); NIH/NIAAA (U01 AA016662, U01 AA013512); NIH/NIDA (P01 DA015027); NIH/NIMH (P50 MH078028, P50 MH096972); NIH/NCI (U24 CA159988); NIH/NIGMS (R01 GM088822).
